# Genome-wide analysis reveals the genetic basis of key agronomic traits and modern wheat breeding in Henan Province

**DOI:** 10.1101/2025.03.13.642997

**Authors:** Chaojun Peng, Ying Chen, Xuelian Han, Haibin Dong, Ao Zheng, Xijun Du, Xiaoyu Chang, Miaomiao Zhao, Xueli Qi, Yuanming Zhang, Lin Hu

## Abstract

Despite the continuous improvement of wheat cultivars in Henan Province, the largest Chinese wheat-producing area, there have been no reports of studies on their breeding advances in the past decades. This study aims to elucidate their genetic basis and breeding advances. A total of 351 landraces and 265 cultivars in Henan Province were employed in selective sweep studies and genome-wide association studies. All the wheat accessions were planted in five locations in Henan Province in 2016−2017, 2017−2018 and 2018−2019. Candidate genes were identified through differential expression, GO annotation, gene homology, and haplotype analyses. Breeding advances were summarized based on four breeding periods (1950s, 1960s−1970s, 1980s−1990s, and post−2000s) of all 265 cultivars. The results demonstrated that 263 known genes with selection signals were found to be associated with important traits. Around the detected quantitative trait loci, 51 genes had been previously reported, while 252 genes were identified as candidate genes. Of the 303 genes and loci identified, 80 genes and 50 loci contributed to the improvement of 12 traits like plant height, and 12 key candidate genes was verified by qRT-PCR, while superior haplotypes of 24 genes explained cultivar climate adaptation. The superior haplotypes of 164 genes are available for future breeding. Transmission of superior haplotypes in four varietal pedigrees revealed pyramiding of superior haplotypes in elite cultivars. The analysis of post−2000 cultivars suggests a broader range of germplasm resources should be considered in parental selection than only recent accessions. This study provides a genetic basis and guide for wheat improvement.

## Introduction

Wheat is the most widely cultivated and traded crop globally, providing sustenance for approximately 40% of the world’s population [1]. Henan Province is the largest wheat-producing regions in China, currently representing approximately 24% of China’s perennial sown area and 29% of China’s total wheat production. The annual genetic gain in wheat yield in Henan Province has been increased by 1.09% from the 1940s to the 2020s, as a result of genetic improvement and cultivar renewal [2]. Nevertheless, the genetic basis of wheat breeding selection and important agronomic traits in Henan Province remains unclear.

The development of genomics-related technologies has accelerated the time to high-quality wheat genome sequences. In 2018, the hexaploid bread wheat “Chinese Spring” was released with a reference genome sequence and annotation information [3]. Subsequently, the sequencing and assembly of a number of important modern wheat cultivars was conducted to obtain genome information at the chromosome level [4−8], revealing the important characteristics and the richness of variation in the current age of wheat cultivars from the genomic perspective. A high-density wheat 660K SNP microarray, designed from the reference genome with genome-specific SNPs at reliable physical locations, is a reliable and cost-effective alternative to genotyping by sequencing [9]. The utilization of the 660K SNP microarrays enabled the genotyping of more than 600 common wheats and their diploid and tetraploid progenitors, explained the mechanism of modern breeding efforts on the Chinese wheat genome, and revealed the genetic basis of selection signatures in the breeding history through genomic information [10]. However, the existing literature on Henan wheat accessions is relatively scarce.

A comprehensive understanding of the genetic architecture of important traits, such as yield, is essential for accurate selection decisions and ideal allele combinations in the breeding. Genome-wide association study (GWAS) is a valuable tool for elucidating the genetic basis of important traits in wheat and for identifying alleles that can be employed in breeding [11,12]. The application of GWAS methods to identify quantitative trait loci (QTLs) for yield, stress tolerance and resistance in wheat has been reported [13−22]. However, a comprehensive genome-wide analysis of the genetic basis of agronomic traits and breeding progress in wheat is still lacking, especially in Henan Province.

In this study, a panel of 616 wheat accessions was genotyped using high-density 660K SNP microarrays, including 351 landraces and 265 modern cultivars from different periods in Henan Province. Subsequently, the selection sweep study was conducted on the wheat landraces and cultivars. The selection regions and their known genes between the landraces and cultivars, and between the cultivars of different periods were identified. Furthermore, the number and functions of genes with selective signals across the different sub-genomes were investigated. Finally, GWAS were conducted on thirteen traits, such as yield and filling rate, to identify known and candidate genes using multi-omics analysis. These genes can be utilized for future wheat breeding in Henan Province. This study elucidated the genetic basis for genetic improvement and important traits in wheat breeding progress, and provided a wealth of genetic resources and novel approaches for future genetic improvement in Henan Province.

## Results

### Genomic and phenotypic signatures trace wheat breeding improvement from landraces to modern cultivars

To evaluate the population structure of 616 wheat accessions, all the 429,721 high quality SNPs were used to perform clustering analysis and principal component analysis (PCA). Although these landraces and cultivars were grouped on the basis of variety source, the clusters were confirmed by both types of analysis. In Figure 1A and 1B, the 616 wheat accessions were clustered into landraces (orange) and cultivars (blue), and about ten varieties were mixed between landraces and cultivars, possibly because these cultivars were bred from landraces. Based on the breeding progress made in Henan Province, 265 cultivars were further divided into four groups: 50s, 60s**−**70s, 80s**−**90s and 00s**−**, to illustrate the breeding progress over different decades. Then, linkage disequilibrium (LD) analysis was performed to calculate the decay distance (2 Mb) of the correlation coefficient (*r^2^*) from the maximum value down to the half of the maximum value, and the distance is considered as the regions of mining candidate genes (Figure S2).

**Figure 1.**
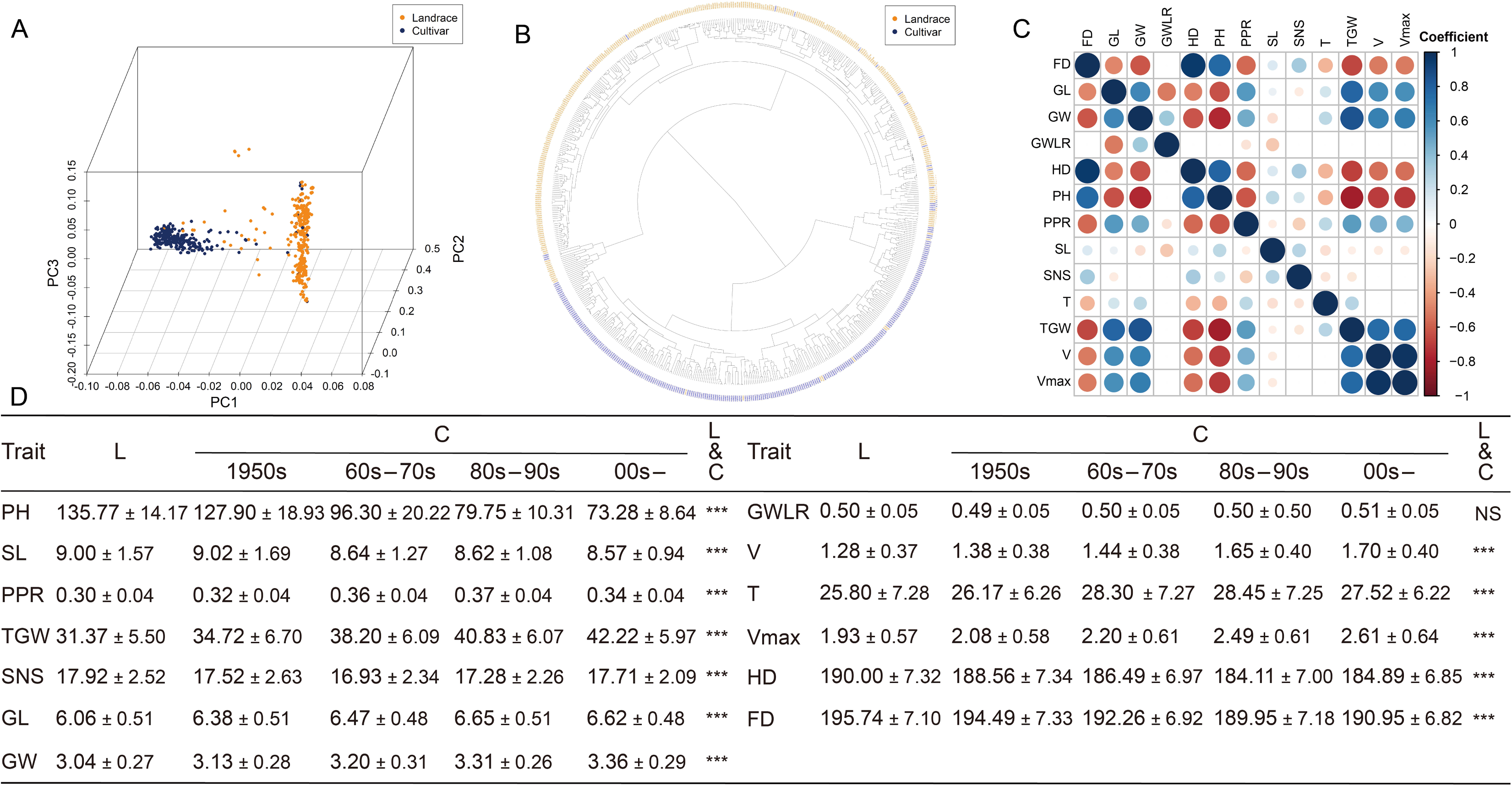
Population structure and phenotypic analyses of 616 wheat accessions. **A.** Principal component analysis of marker genotypes. **B.** Cluster analysis of marker genotypes. **C.** Correlation analysis between the 13 traits. Red represents positive correlation, while blue represents negative correlation. **D.** Significant differences of 13 traits among landraces and cultivars with different decades. L: landraces, C: cultivars, NS: no significance, ***: the 0.001 probability level of significance, PH: plant height, SL: spike length, PPR: ratio of peduncle length to plant height, SNS: spikelet number per spike, TGW: thousand grain weight, GL: grain length, GW: grain width, GWLR: ratio of grain width to length, Vmax: maximum filling rate, V: average grain filling rate, T: active filling period, HD: heading date, FD: flowering date.

A total of 13 important agronomic traits were measured, including morphological traits (plant height, PH; spike length, SL; the ratio of peduncle length to plant height, PPR), yield-related traits (spikelet number per spike, SNS; thousand grain weight, TGW; grain length, GL; grain width GW; grain width/length ratio GWLR), grain filling-related traits (maximum filling rate, Vmax; average grain filling rate, V; active filling period, T), and growth period-related traits (heading date, HD; flowering date, FD). Correlation analysis revealed that SL exhibited a moderate positive correlation with PH, while displaying a negative correlation with PPR. Furthermore, strong positive correlations were observed between GW, GL and TGW. However, the correlation between SNS and GL, as well as between SNS and GWLR, did not reach statistical significance. Conversely, a negative correlation was identified between SNS and TGW, as well as between SNS and GL. Furthermore, a positive correlation was observed between V and Vmax, as well as between HD and FD (Figure 1C).

All traits except for GWLR demonstrated notable discrepancies between landraces and cultivars. For instance, cultivars exhibited markedly reduced PH and SL in comparison to landraces, whereas cultivars displayed considerably elevated TGW, GL, and GW in relation to landraces. During the breeding process and across different periods, significant changes were observed (Figure 1D). The most notable alterations in three morphological traits occurred between the 1950s and 1970s. For instance, PH underwent a significant change, decreasing from 127.90 cm to 96.30 cm. The most notable changes in FD and HD occurred from the 60s−70s to the 80s−90s, with reductions of 2.31 and 2.39 days, respectively. With regard to yield-related traits, the most substantial changes in grain shape were observed from landraces to the 50s, with an increase of 0.31 mm in GL. HXM393 in the 50s exhibited the greatest GL (7.30 mm). The most significant changes were observed in TGW and SNS during the periods from 50s to 60s−70s and from 80s−90s to 00s−, respectively. TGW increased by 3.49 g (from 34.71 to 38.20 g), while SNS increased by 0.43 (from 17.28 to 17.71). The most substantial alterations in grain-filling related traits were observed from the 50s to the 80s−90s. For instance, the T increased by 2.13 days from the 50s to the 60s−70s, while the V and Vmax increased by 0.20 and 0.29 (g/DAP), respectively, from the 60s−70s to the 80s−90s.

Unravelling the significant differences of these traits between landraces and cultivars and between different periods is essential for a comprehensive understanding of the genetic improvement from landraces to cultivars and the trait change trends across different periods.

### Selective gene introgression drives the improvement of important traits in wheat breeding

In order to identify selective regions, five distinct indicators were employed. The top 5% of intervals identified by at least two indicators were considered as candidate regions for further investigation in selective sweep studies. The number of genes in selective regions was 12,602 from landraces to cultivars, 17,450 from landraces to 50s, 20,666 from 50s to 60s−70s, 24,335 from 60s−70s to 80s−90s and 19,843 from 80s−90s to 00s (Figure 2A). All the known genes are listed in Table S1. In Table S1, 59 known genes were found to be in selective regions from landraces to cultivars (Figure 2B), including 14 growth period-related genes (GPRGs), 12 yield-related genes (YRGs), 16 quality-related genes (QRGs), 12 resistance-related genes (RRGs) and 6 morphology-related genes (MRGs); 88 known genes from landraces to the 50s, including 17 GPRGs, 10 MRGs, 31 QRGs, 17 YRGs and 20 RRGs; 88 known genes from 50s to 60s-70s, including 17 GPRGs, 12 MRGs, 21 QRGs, 27 YRGs and 10 RRGs, and *TaDWF4-3D1* and *TaDWF4-3D2* for nitrogen use were detected simultaneously by five indicators; 102 known genes from 60s-70s to 80s-90s, including 13 GPRGs, 16 MRGs, 25 QRGs, 32 YRGs and 23 RRGs; 81 known genes from 80s-90s to 00s, including 14 GPRGs, 15 MRGs, 18 QRGs, 20 YRGs and 17 RRGs (Table S1). It should be noted that many YRGs were detected in all generational selection regions (Table S1). Thus, yield improvement has played a crucial role in breeding progress.

**Figure 2.**
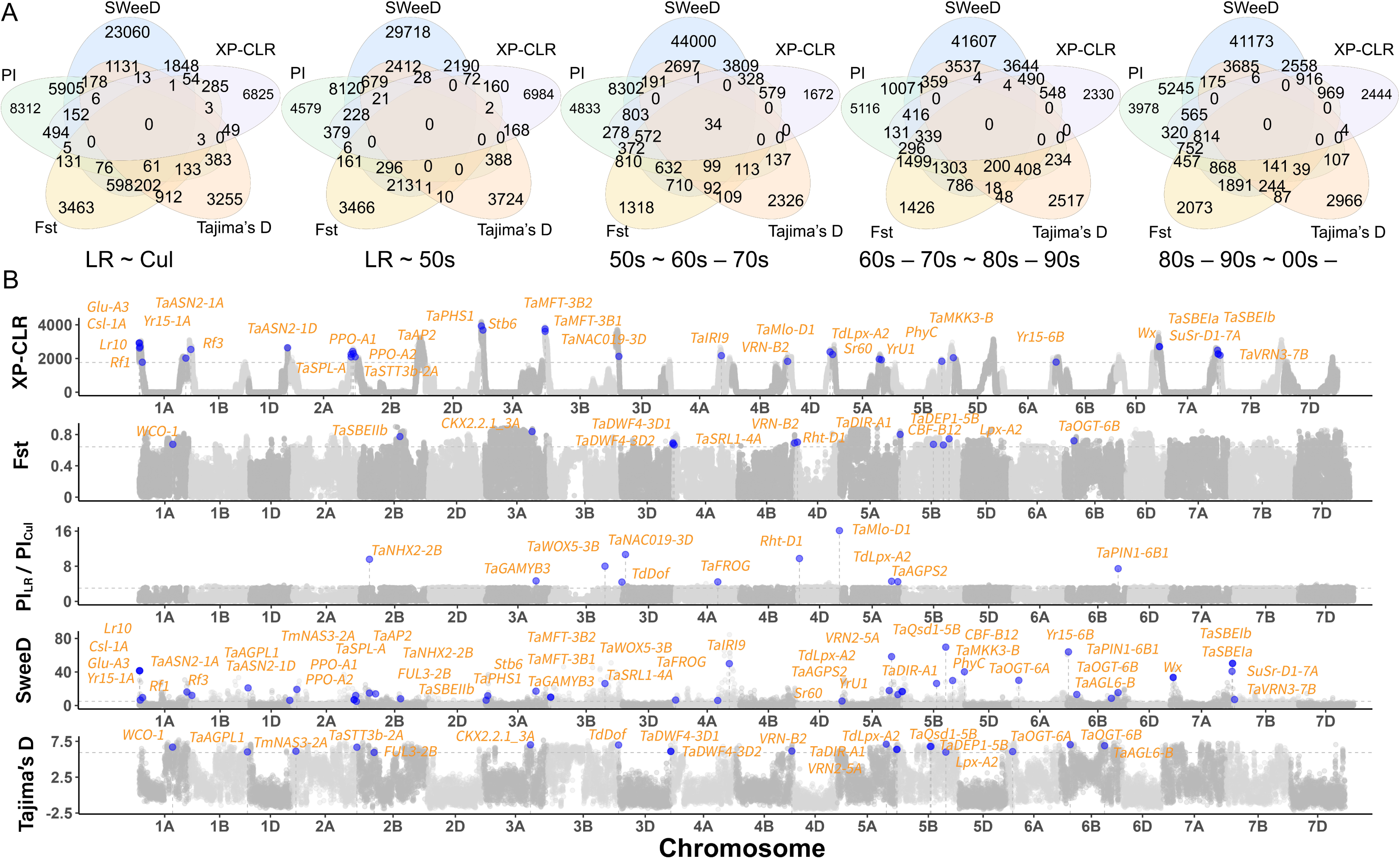
The number of genes with selection signals and the distribution of known genes in the comparison between landraces and cultivars. **A.** Number of genes with selection signals identified by the XP-CLR, SweeD, Fst, π, and Tajima’s D statistics in five comparisons: landraces and cultivars, landraces and 50s cultivars, 60s−70s and 80s−90s cultivars, and 80s−90s and 00s− cultivars. **B.** Known genes identified by XP−CLR, SweeD, Fst, π, and Tajima’s D statistics between landraces and cultivars. The grey dashed line indicates the genetic indicator threshold set at the top 5%. LR: landraces, Cul: cultivars.

For three morphological traits, the largest changes of traits between different breeding periods occurred from the 50s to the 60s−70s based on the trait values in Figure 1D, while 12 known MRGs were found in the selection intervals, including 6 genes (e.g., *SVP3*−*4A*, *TaSRL1*−*4A* and *TaARF12*−*2D*) for PH and 2 genes (*SVP3*−*4A* and *TaARF12*−*2D*) for SL. For five yield related traits, the largest grain shape changes occurred from landraces to the 50s, while grain shape-related gene *TaARF25*−*5A* was found in the selection intervals; the largest changes of TGW occurred from 50s to 60s-70s, while 8 TGW related genes (such as *GNI*−*B1* and *TaPIN1*−*6D*) were found in the selection intervals; the largest SNS changes occurred from the 80s−90s to the 00s−, while 8 spike-related genes (such as *FT*−*A2* and *TaPIN1*−*6B1*) were found in the selection intervals. For three grain-filling related traits, the largest trait changes mainly occurred from the 50s to the 80s−90s, while 8 starch-related genes were found in the selection intervals from the 50s to the 60s−70s, and 7 starch-elated genes (such as *TaSuSy4*, *BGC1* and *TaSBEIa*) were found in the selection intervals from the 60s−70s to the 80s−90s. For two resistance traits, seven genes for stripe rust resistance, such as *TaPsIPK1*−*6B* and *YrU1*, and one gene for powdery mildew resistance, *Lr67*, were found in the selection intervals from the 60s−70s to the 80s−90s, three gene for stripe rust resistance, such as *Lr67*, and four genes for powdery mildew resistance, such as *Lr67* and *TaMlo*−*D1*, were found in the selection intervals from the 80s−90s to the 00s−, while the number of varieties resistant to stripe rust and powdery mildew increased in the 80s−90s and the 00s− (Figure 1D; Table S2). In summary, it is quite possible that the above significant trait changes in different periods of wheat cultivars are caused by the introduction of many excellent genes in wheat breeding.

Zhengmai 7698 is a representative cultivar from Henan Province. The genes in the 7698 [23]. The results showed that five genes in Li et al. [23] underwent selection, three glutenin−related genes *Bx7* (*Glu-1Bx*), *By9* (*Glu*−*1By*) and *Ax1* (*Glu*−*A1*) influenced flour quality, the polyphenol oxidase gene *PPO*−*A1b* influenced flour color, and the gene *PHS1* was associated with preharvest sprouting resistance (Table S3).

### Multiple GWAS methods identifies known and candidate genes for important traits

#### Identification of genes for wheat morphological traits

For morphological traits, the 3VmrMLM, BLUPmrMLM, EMMAX, GEMMA and FarmCPU methods identified 552, 291 and 164 significant QTLs associated with PH, SL and PPR, respectively, while 557, 470 and 167 suggested QTLs associated with PH, SL and PPR were identified (Tables S4−S6). Around these QTLs, all the known genes are listed in Table S7, while all the candidate genes are listed in Table S8.

Among all the genes around PH, SL and PPR QTLs, 12, 3 and 3 genes were reported to be responsible for PH, SL and PPR in previous studies, such as *TaARF12*−*2A*, *TaARF12*−*2B* [24] and green revolution gene *Rht*−*D1* [25] for PH, *TaARF12*−*2A* for PPR and *TaARF12*−*2B* for SL [24]. 181 out of 5987 differential expression genes (DEGs), 333 out of 1909 DEGs and 202 out of 2758 DEGs were identified by GO annotation to be associated with PH, SL and PPR, respectively, while 286, 175 and 121 genes were found to be homologous to the known rice PH, SL and PPR genes, such as *d35* for PH [26] and *DLT* for PPR [27]. Among the above genes, 22, 15 and 11 genes were identified by haplotype analysis to be significantly associated with PH, SL and PPR, respectively. 4 out of 22 PH candidate genes were simultaneously identified by differential expression (|log_2_FC| > 1; adjust *P*-value < 0.05), GO annotation, rice gene homology and haplotype analysis, including *TraesCS2A01G552400*, *TraesCS4A01G460100*, *TraesCS6D01G173000* and *TraesCS7A01G362300*, while two key PPR candidate genes *TraesCS2A01G552400* and *TraesCS7A01G059000* were simultaneously identified by the above four analysis (Table S8; Figure S3).

#### Identification of genes for wheat yield-related traits

For yield-related traits, the above five GWAS methods identified 202, 328, 251, 269 and 130 significant QTLs associated with GL, GW, GWLR, TGW and SNS, respectively, while 380, 442, 334, 439 and 461 suggested QTLs associated with the five traits were identified (Tables S8−S13). Around these QTLs, all the known genes are listed in Table S14, while all the candidate genes are listed in Table S15.

Among all the genes around the GL, GW, GWLR, TGW and SNS QTLs, 4, 3, 7, 8 and 2 genes were reported to be responsible for the five traits in previous studies, including *DA1* for grain weight and shape traits [28] and *GW7* for grain shape traits [29]. 26 out of 4719 DEGs, 30 out of 5559 DEGs, 27 out of 5530 DEGs, 25 out of 5858 DEGs and 10 out of 4475 DEGs are identified by GO annotation to be associated with the above five traits, respectively, while 42, 36, 9, 64 and 3 candidate genes were found to be homologous to the known genes for the five traits in rice, respectively. Among the above genes, 24, 17, 19, 34 and 13 genes were identified by haplotype analysis to be significantly associated with the above five traits, respectively (Table S15; Figure 3A and S4).

**Figure 3.**
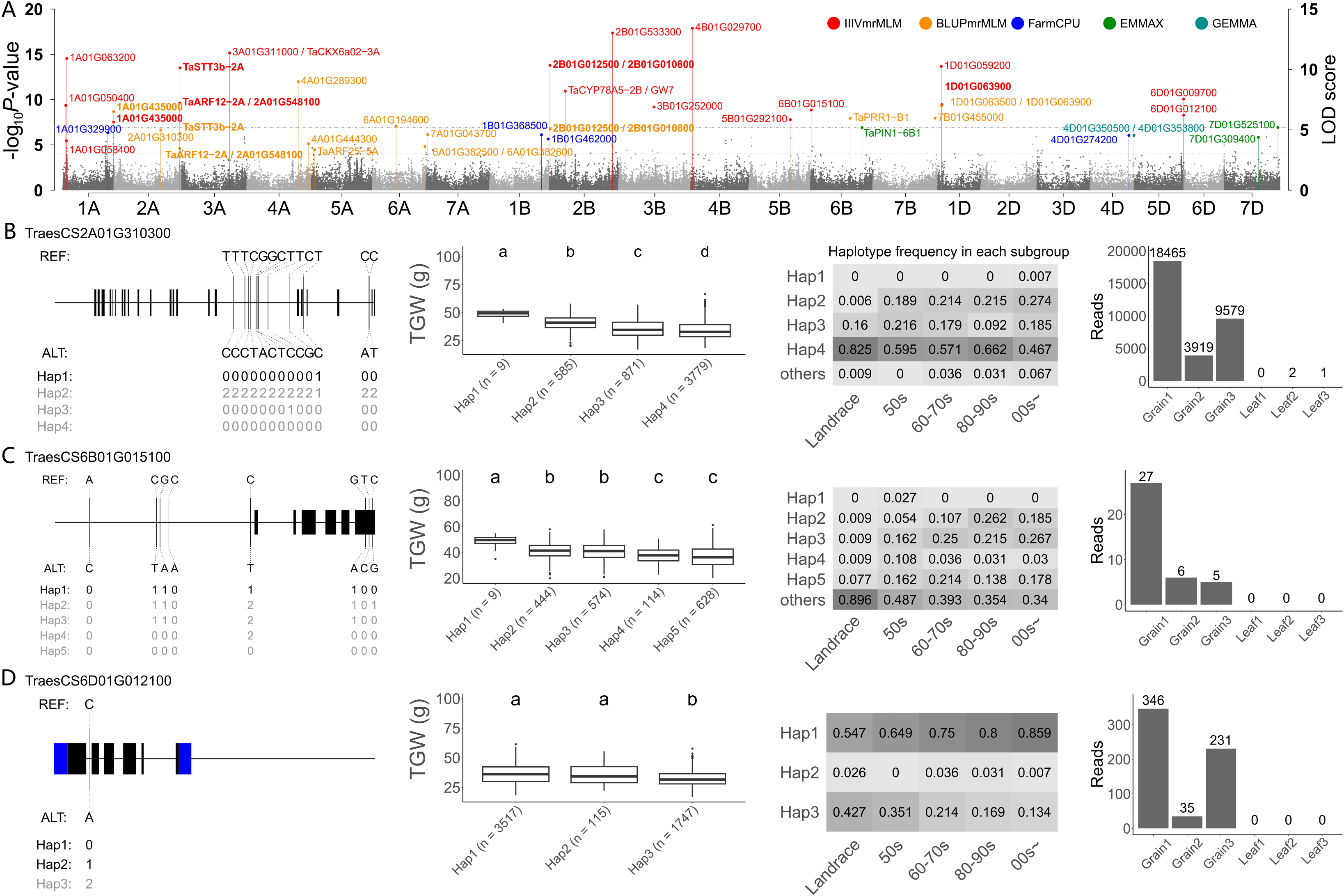
The genes identified for thousand grain weight (TGW) in wheat. **A.** Manhattan plot for TGW. IIIVmrMLM: red; BLUPmrMLM: orange, FarmCPU: blue; EMMAX: green; GEMMA: cyan. The TraesCS in gene names have been deleted. Genes in bold are detected by multiple methods. Identification of gene-trait associations for *TraesCS2A01G310300* (B), *TraesCS6D01G012100* (C) and *TraesCS6B01G015100* (D). For each gene in the subplots B to D, the superior haplotype (black), the multiple comparisons of the haplotypes over all environments, the frequency of the haplotypes in different sub-populations, and the expressional levels in leaves (*n* = 3) and grains (*n* = 3) at the dough stage are presented, respectively, in the first to fourth columns. 0: REF base, 2: ALT base, 1: heterozygous genotype.

*TraesCS2A01G310300*, *TraesCS6D01G012100* and *TraesCS6B01G015100* were simultaneously found to be associated with TGW using differential expression, GO were much higher in seeds than in leaves at dough stage (Figure 3B, 3C and 3D). The superior haplotype (Hap1) of *TraesCS6D01G012100* has a high frequency that increased with time, indicating its important role in increasing yield (Figure 3C), while the superior haplotypes of other genes were not high in all subgroups, indicating their high breeding values in the future (Figure 3B and 3D).

#### Identification of genes for wheat grain filling-related traits

For grain filling-related traits, the five GWAS methods identified 77, 178 and 225 significant QTLs associated with T, V and Vmax, respectively, while 408, 312 and 433 suggested QTLs associated with T, V and Vmax were identified (Tables S16-S18). Around these QTLs, all the candidate genes are listed in Table S19.

Among all the genes around the T, V and Vmax QTLs, 24 out of 3477 DEGs, 36 out of 4,318 DEGs and 52 out of 5053 DEGs are identified by GO annotation to be associated with T, V and Vmax, respectively; while 3, 6 and 3 genes for T, V and Vmax were found to be homologous to the known rice grain filling genes, such as *OsFl3* for V [30], and 14, 13 and 18 genes for T, V and Vmax were reported to be known genes for wheat grain-related traits, such as such as *GW7* [29]. Among the above candidate genes, 9, 23 and 25 genes were identified by haplotype analysis to be significantly associated with T, V and Vmax, respectively (Table S19; Figure S5 and S5A). Two out of 9 candidate genes for T, *TraesCS2B02G194200* and *TraesCS7D02G159800*, were simultaneously identified by differential expression analysis, GO annotation, rice homology genes, wheat grain-related genes and haplotype analysis, while there were three key V and four key Vmax candidate genes (*TraesCS1A02G247100*, *TraesCS2A02G293400*, *TraesCS7A02G549100* and *TraesCS7A02G549300*) (Table S19; Figure S4B and S6).

Among the three key V genes *TraesCS2A01G293400*, *TraesCS7A01G549100* and *TraesCS7A01G549300*, the superior haplotypes (Hap1, Hap2 and Hap3) of *TraesCS2A01G293400* have a high frequency in cultivars that increased with time, indicating their important role in wheat breeding for improving filling rate (Figure S6), whereas the superior haplotypes of *TraesCS7A01G549100* and *TraesCS7A01G549300* were not high in all subgroups, indicating their high breeding value in the future (Figure S6).

#### Identification of genes for wheat growth period-related traits

For growth period-related traits, the five GWAS methods identified 413 and 314 significant QTLs associated with FD and HD, respectively, while 493 and 675 suggested QTLs associated with FD and HD were identified (Tables S14−S15). Around these QTLs, all the known genes are listed in Table S22, while all the candidate genes are listed in Table S23.

Among all the genes around the FD and HD QTLs, 5 and 4 genes were reported to be responsible for FD and HD, respectively, in previous studies, such as *TaFT3*−*B1* and *TaTOE*−*D1* for FD [31] and *PhyC* for HD [32]. 33 out of 10,042 DEGs and 8 out of 2603 DEGs are identified by GO annotation to be associated with FD and HD, respectively, while 33 and 10 genes for FD and HD, respectively, were found to be homologous to their known genes in rice, such as *Se14* for FD [33]. Among the above genes, 31 and 9 genes were identified by haplotype analysis to be significantly associated with FD and HD, respectively (Table S23; Figure S7A and S4). One out of the 31 candidate genes for FD has relatively sufficient evidence to be associated with FD, while 7 genes for HD and 9 genes for FD were detected simultaneously by FD was detected by four methods except for FarmCPU (Table S23), its expression levels were different in floral organs and leaves, and its superior haplotype was extremely fully used in cultivars rather than in landraces (Figure S7).

## Discussion

In this study, 12 important traits except for GWLR were found to be significant between landraces and cultivars, while PH, TGW, GW and FD were significant between cultivars of different periods (Figure 1D). In the genetic diversity analysis, 11.37% ∼ 21.96% of all the genes were located on selection intervals, among which 263 genes were reported in previous studies (Figure 2; Table S1), including genes related to hormone signaling, protein synthesis, growth and development, disease resistance and so on, such as *Lr67*, which affects resistance to a wide range of pathogens [34], and *Rht-B1* and *Rht-D1*, known as green revolution genes [35]. In the GWAS for 13 traits, 51 genes were reported in previous studies, such as *TaARF12* [34] and *DA1* [28], while 252 genes were identified as candidates, of which 19 genes were very important (Tables S8, S15, S19, and S23).

To confirm the correctness and breeding value of the results in this study, the trait BLUP values and the significant and suggested loci were used to perform genomic selection using the R package rrBLUP. The results showed its high prediction accuracy (Table S24; Figure S8), with similar results observed in cassava, chrysanthemum, and maize [35–37], demonstrating its general but not random results.

### Known and candidate genes responsible for trait changes from landraces to different period cultivars

The correlation coefficient between the trait mean value and the frequency of superior haplotype for each of the 303 genes in each group were used to ascertain whether a specific gene was responsible for trait changes from landraces to different period cultivars [38]. As a result, 80 known and candidate genes for all the traits except for GWLR contributed to trait changes from landraces to different period cultivars in Henan wheat breeding (Figure 4; Table S25).

**Figure 4.**
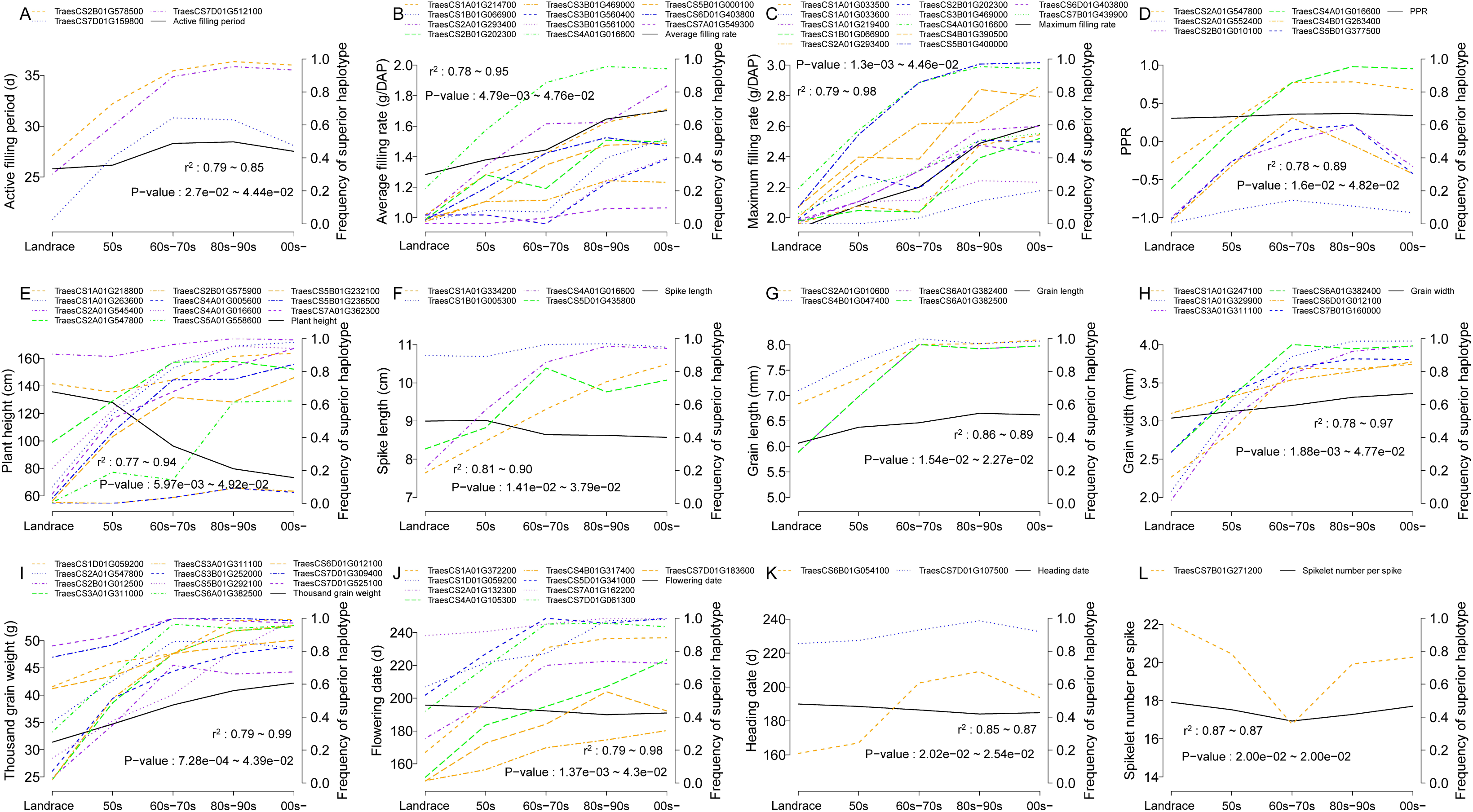
Genes responsible for the changes of 12 traits between landraces and cultivars with different periods. The black lines (phenotypes) are based on the left-side vertical coordinate axis, while the colored lines (gene frequencies) are based on the right-side vertical coordinate axis. PPR: ratio of peduncle length to plant height.

For three morphological traits, the most significant changes occurred during the period from the 1950s to the 1960s–1970s due to the changes in the frequencies and effects of the superior haplotypes of 21 genes (Figure 4; Table S25). First, the largest frequency changes (LFCs) (about 30%) of their superior haplotypes for seven morphological trait genes occurred between the two periods, including *TraesCS4A01G016600* and *TraesCS4B01G263400* for PPR, *TraesCS5B01G236500*, *TraesCS4A01G016600* and *TraesCS1A01G263600* for PH, and *TraesCS4A01G016600* and *TraesCS5D01G435800* for SL. Second, the superior haplotype genetic values (SHGVs) for these genes were changed, e.g., the superior haplotype genetic values for eight PH genes decreased by more than 20 cm from the 50s to the 60s–70s.

For yield–related traits, the largest GL and GW changes occurred mainly from the landraces to the 50s, and the largest TGW changes occurred from the 50s to the 60s– 70s, due to the changes in the frequencies and effects of the superior haplotypes of 21 genes. First, the LFCs (>10%–18%) of superior haplotypes for three GL–related genes (*TraesCS4B01G047400*, *TraesCS6A01G382400* and *TraesCS6A01G382500*), four GW–related genes (e.g. *TraesCS1A01G329900* and *TraesCS3A01G311100*), and four TGW–related genes (e.g. *TraesCS6D01G012100* and *TraesCS7D01G309400*) occurred between their corresponding change periods. Second, the SHGVs of these genes were changed, e.g. four GL genes increased by over 0.30 mm, and six TGW genes increased by more than 2.20 g.

For grain–filling traits, the largest V and Vmax changes occurred from the 60s–70s to the 80s–90s, and the largest T changes occurred from the 50s to the 60s–70s, due to the changes in the frequencies and effects of the superior haplotypes of 12 genes. First, the LFCs (>25%, >10% and >30%) of their superior haplotypes for four V genes (e.g. *TraesCS1B01G066900* and *TraesCS2B01G202300*), seven Vmax genes (e.g. *TraesCS1A01G219400* and *TraesCS1B01G066900*) and one T gene (*TraesCS7D01G512100*) occurred between their corresponding periods. Second, the SHGVs of five V genes and nine Vmax genes increased by >0.25, while the superior haplotype genetic values of *TraesCS2B01G578500* and *TraesCS7D01G512100* increased by 1.86 and 1.98, respectively.

For two growth period traits, the largest trait changes occurred from the 60s–70s to the 80s–90s. First, the LFCs (approximately 20%) of their superior haplotype of *TraesCS1D01G059200* occurred between the two periods. Second, the SHGVs of eight FD genes and two HD genes decreased by more than 2.00 d.

To determine the important loci contributing to the trait changes, as described above, the correlation coefficient for the most significant 20 loci for each of 12 traits were used to ascertain whether a specific locus was responsible for trait changes. As a result, the frequency changes of superior genotypes of 50 loci were found to be significantly correlated with the trait changes from landraces to different period cultivars. In particular, the superior genotypes of 36 out of 50 loci were fully utilized in wheat breeding because their superior genotype frequencies exceeded 0.50 (Table In summary, the increase in superior haplotype/genotype frequency of 80 genes and 50 loci was positively correlated with trait improvement, with the majority of superior haplotype/genotype frequencies exceeding 0.50, while the superior haplotype genetic values of the 80 genes were more favorable for trait improvement, indicating that the accumulation of superior haplotypes/genotypes of 80 known and candidate genes and 50 loci contributed to the improvement of the 12 traits (Tables S25–S26; Figures 4 and 5A).

**Figure 5.**
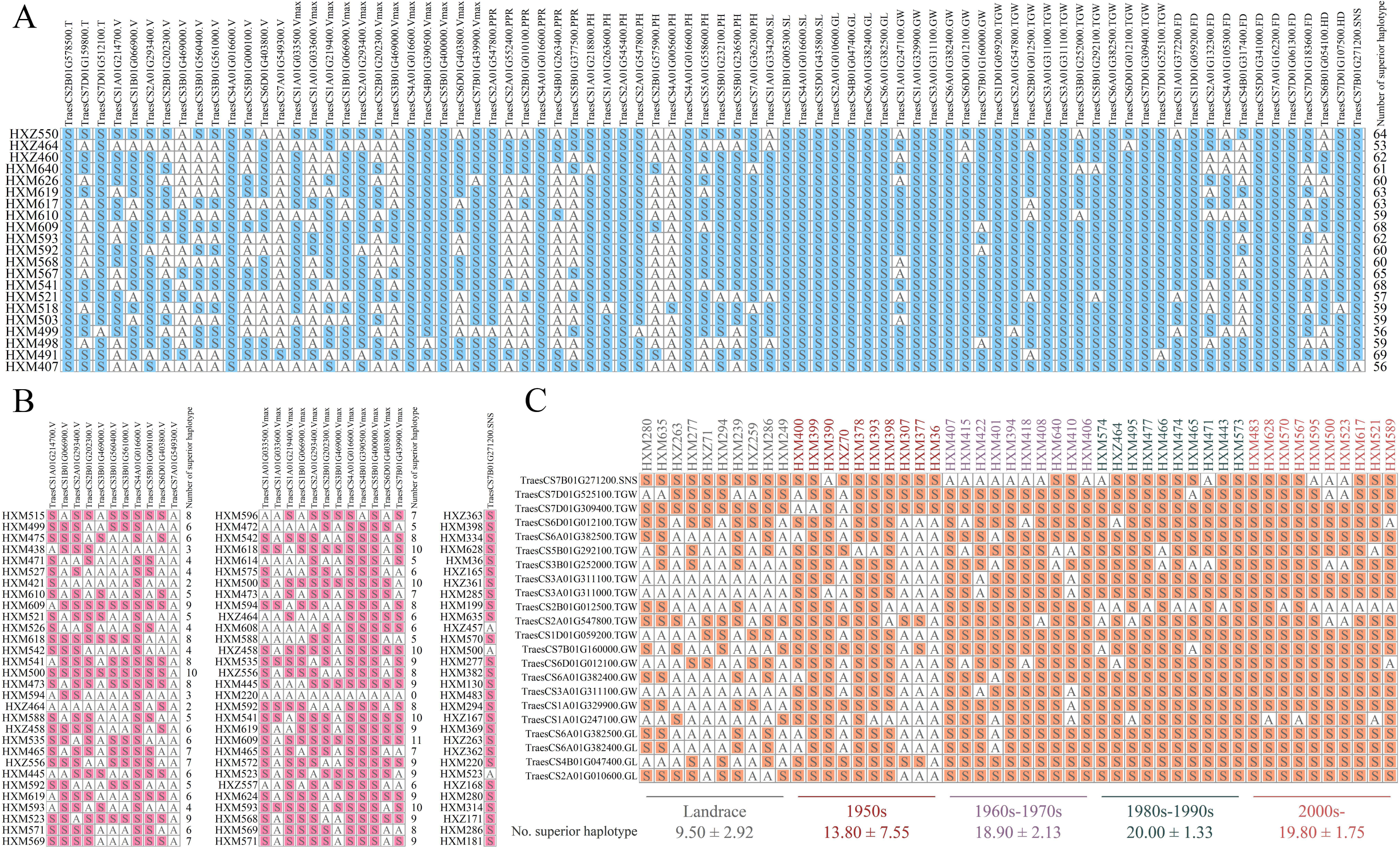
Distribution of superior haplotypes in superior accessions. **A.** Distribution of superior haplotypes of 80 genes contributing to the improvement of 12 traits in the accessions with excellent overall performance. **B.** Distribution of superior haplotypes in 30 accessions with top 5% phenotypes for grain filling rate, maximum grain filling rate, and spikelet number per spike. **C.** Distribution of superior haplotypes of the yield-related genes in the top 10 accessions of each period. The number is the average and standard deviation of superior haplotypes in each period. No.: number, PH: plant height, SL: spike length, PPR: peduncle length to plant height ratio, SNS: spikelet number per spike, TGW: thousand grain weight, GL: grain length, GW: grain width, Vmax: maximum filling rate, V: average grain filling rate, T: active filling period, HD: heading date, FD: flowering date. S filled with color represents a superior haplotype, while A without color represents a non-superior haplotype.

**Figure 6.**
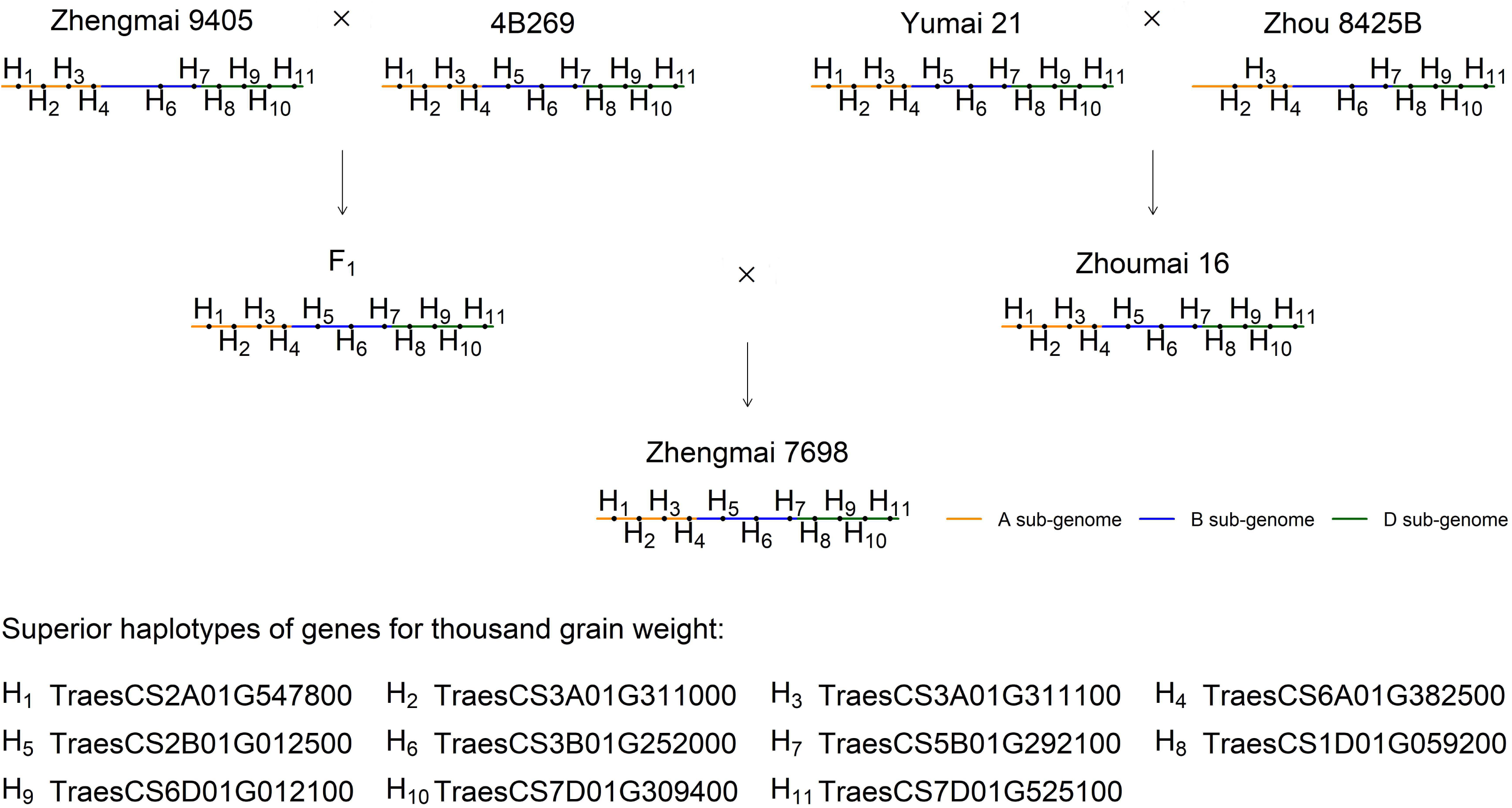
Transmission of superior haplotypes of eleven known and candidate genes for thousand grain weight in HXM518 (Zhengmai 7698) pedigree H is the gene superior haplotypes for thousand grain weight. Zhengmai 9504, 4B269, Yumai 21, Zhou 8425B and Zhoumai 16 were the parents of cultivar Zhengmai 7698.

### Genomic findings with the region characteristics

To adapt to the climatic conditions of Henan Province, wheat accessions have three unique growth characteristics compared to other eco–regions: longer tillering period, longer spike differentiation period and shorter grain filling period. In this study, SNS, V and Vmax were found to increase from the landraces to different period cultivars (Figure 1D). To explain this phenomenon, 30 accessions with top 5% phenotypes for each trait were selected to analyze the contribution of superior haplotypes of one SNS, 11 V and 12 Vmax genes in the above 80 known and candidate genes. On average, there are about 6 out of 11 superior haplotypes of the V genes and about 8 out of 12 superior haplotypes of the Vmax genes in the selected 30 accessions (Table S27; Figure 5B), while 27 of the 30 selected accessions have the superior haplotype of the SNS gene *TraesCS7B01G271200* (Table S27; Figure 5B). These genes are helpful to improve the climate adaptation of these accessions in Henan Province, demonstrating the potential of pyramiding more superior haplotypes to improve fitness.

Yield improvement in Henan Province has been going on since the 1950s. The top 10 varieties from each period were selected based on the performance of yield–related traits. The largest changes of the average number of superior haplotypes for the top 10 varieties in each period occurred from the 50s to the 60s–70s, which is consistent with the period for the largest changes in SNS and TGW between the five periods (Figure 1D and 5C), indicating the reliability of these results in this study.

### Transmission of superior haplotypes in important varietal pedigree

Four cultivars HXM497 (Zhoumai 17), HXM518 (Zhengmai 7698), HXM523 (Luomai 21), and HXM590 (Xinmai 30) with complete pedigrees, were selected for analysis of the transmission of superior haplotypes of genes driving trait variation in the pedigree. Among the above four varieties, the number of superior haplotypes in the variety is higher than in any of its parents. For example, HXM518 had 46 superior haplotypes compared with 35, 41, 29, 39 and 31 superior haplotypes from its parents HXM446 (Yumai 21), HXM496 (Zhoumai 16), HXM516 (Zhengmai 9405), HXM573 (4B269) and HXM574 (Zhou 8425B), respectively (Tables S28–S31). Some of the superior haplotypes were from one or a few parents. For instance, the superior haplotype of *TraesCS1A01G214700* in HXM518 was derived exclusively from HXM496 (Table S29). Additionally, the number of superior haplotypes in the varieties was augmented through the polymerization of superior haplotypes through crosses between parents. Nevertheless, it was found that very few superior haplotypes did not originate from the parents, which could be attributed to recombination, mutation or other causes.

Furthermore, the superior haplotypes of TGW in HXM518 were taken as an example. It was observed that HXM516 lacked H1 and HXM574 lacked H1 and H5. As a consequence of the complementation of superior haplotypes between the parents, the final HXM518 was obtained with 11 superior haplotypes (Figure 5). Consequently, the combination or complementation between the parents aggregated the superior haplotypes, thereby increasing the number of superior haplotypes and improving the performance of the cultivar.

### Known and candidate genes available for future wheat breeding in Henan province

All identified candidate genes were categorized into genes that could be used for breeding and genes that had been fully utilized according to whether the frequency of superior haplotypes was greater than 0.50 [39]. In total, 277 superior haplotypes of known and candidate genes were identified for the 13 traits. Of these, 158 superior haplotypes with frequencies lower than 0.50 were genes that could be utilized for breeding (Figure S9; Table S32).

To obtain the haplotypes of known genes in the selection intervals, all varieties with extreme phenotypes were grouped into two extreme pools. If a significant difference in the frequency of a gene haplotype was observed in the two pools, this haplotype was used to scan the frequent changes in each group (landraces, 50s, 60s–70s, 80s– 90s and 00s–) (Table S33). For disease resistance traits, three out of the six known genes related to powdery mildew were fully used in the 00s, with particular reference to *Lr67*; six out of the eleven known genes related to stripe rust were fully used, such as *Lr67* and *YrU1* (Table S33). For breeding progress in grain quality, the majority of the haplotypes with high grain protein content exhibited a decline in frequency, whereas all the haplotypes with low grain protein content demonstrated an increase, such as *Glu*–*A3* and *Glu*–*1Ax* haplotype frequency change of 52.22% and 31.29% from 50s to 80s–90s, indicating the breeding process towards lower grain protein content, which is consistent with our findings that the annual genetic gain in grain nitrogen content decreases during genetic improvement [40]. For FD, 7 out of the 17 known genes were found to be associated with FD, exhibiting superior haplotype frequencies higher than 0.50 (Table S32). Furthermore, 3 out of the 15 genes were also related to TGW with superior haplotype frequencies in excess of 0.50 (Figure 1D), which were fully used in the breeding progress. These known genes were also considered to be a powerful factor for the improvement of critical traits in the breeding progress. And superior haplotypes of 46 genes could be used for breeding in the future (Table S32). In conclusion, the distributions of superior haplotypes of 164 genes were described in Table S34, which can be employed for breeding in the future.

Among the genes available for breeding, 64 and 15 gene superior haplotypes were absent in the 00s– (Table S34), which could be attributed to the loss of some superior haplotypes during the breeding process and suggests the selection of a wider range of germplasms in parental selection than only recent germplasms, such as HXM239, HXM280, HXM232 and HXM635, which had relatively more superior haplotypes (Table S35). In addition, TGW had the most superior haplotypes available for breeding among the 13 traits, and TGW can make a significant contribution to genetic gain in yield.

### Candidate genes for future molecular biological validation

In the GWAS for 13 traits, 51 known genes identified in this study were reported in previous studies (Tables S7, S14 and S22), indicating the reliability of the methods. Meanwhile, 252 genes were identified as candidates (Tables S8, S15, S19 and S23). Furthermore, ATAC-seq data during grain development [41] was used to substantiate the candidate genes for yield and grain filling related traits. Consequently, accessible chromatin regions were found to overlap with 61 gene bodies and located less than 100 bp, 100–2000 bp away and 2000–8000 bp away from the gene bodies of 11, 20 and 8 genes, respectively (Table S36). This further demonstrates the reliability of the candidate genes in this study.

A total of 19 focus candidate genes were confirmed by all four aforementioned methods to be associated with seven traits (Tables S8, S15, S19 and S23), namely focus candidates. After removing replicated genes identified between traits, 15 key candidate genes were obtained. In ATAC-seq analysis, accessible chromatin regions overlapped with the regions of 7 key candidate genes related to wheat grain (Table S36). More importantly, qRT-PCR showed that 12 out of 15 key candidate genes were found to be expressed in the corresponding issues (Figure S10). It is possible that these genes may play an important role and further verification is required through molecular biology experiments in the future.

### Asymmetric selection among the three sub-genomes of common wheat in breeding

Although the numbers of all the genes in the sub-genomes were largely similar, but the numbers of the genes in the selection intervals were significantly different between the sub-genomes, indicating the asymmetry in the numbers of the genes with selection signals in the sub–genomes (Figure S11). The number of gene with selection signals in sub-genomes is A > B > D sub–genome (Figure S11), which is consistent with Hao et al. [42]. To investigate the functions of the genes with selection signals in the sub-genomes, all the genes with selection signals in five comparisons were annotated, and the majority of biological processes of the genes in the sub-genomes A, B and D were mainly involved in protein phosphorylation, oxidation-reduction process, regulation of transcription, DNA–templated and other related functions, demonstrating the similarity in the majority of the gene functions observed across the three sub-genomes, although the number of genes involved in pollen recognition was significantly greater in D sub–genome than in A and B sub-genomes in the selection interval from landraces to 50s, indicating a partial asymmetry in gene functions across the sub–genomes (Table S37). In addition, the carbohydrate metabolic process, with a relatively large number of genes in this study and Hao et al. [41], was a very important biological process, while yield improvement has been ongoing since the 1950s, in which is consistent with Hao et al. [42], and the yield-related genes were found to be in the selection interval of each comparison in this study (Table S37).

## Conclusion

The wheat genome has undergone a relatively large number of selections, with 11.37% ∼ 21.96% of the total genes in the whole genome falling in the selection intervals, in which 263 genes have been identified as responsible for important traits in wheat in Henan Province. A total of 51 known genes and 252 (of which 19 were identified as key) candidate genes were found to be associated with 13 traits, of which 164 genes with frequencies of their superior haplotypes less than 0.50 can be used for future breeding, 80 genes and 50 loci for 12 traits contributed to trait improvement between landraces and cultivars with different periods, except for GWLR, 12 key candidate genes was preliminarily verified by qRT-PCR, and superior haplotypes of 24 genes explained cultivar climate adaptation. The transmission of superior haplotypes in important varietal pedigree demonstrated that the combination or complementation between parents can aggregate the superior haplotypes, increase the number of superior haplotypes and improve the performance of cultivars.

## Materials and methods

### Plant materials

The 616 wheat accessions, including 351 landraces and 265 cultivars, were obtained from the Wheat Institute of Henan Academy of Agricultural Sciences and used as association mapping populations. All 265 cultivars were roughly divided into four groups: 1950s (50s, 37), 1960−1970s (60s−70s, 28), 1980−1990s (80s−90s, 65), and post−2000s (00s−, 135). The names, origins, and corresponding breeding years of the 616 accessions are provided in File S1.

### Field trials for trait evaluation

All the wheat accessions were planted in five locations, namely Xinxiang (35.01°N, 113.70°E), Anyang (36.13°N, 114.58°E), Jiyuan (35.12°N, 112.28°E), Shangqiu (34. 24°N, 116.13°E) and Zhumadian (33.01°N, 114.40°E) in Henan Province (Figure S1), with Xinxiang in 2016-2017, 2017-2018 and 2018-2019 and the others in 2016-2017 and 2017-2018. A total of 13 important agronomic traits were measured, including morphological traits (PH, SL, and PPR), yield-related traits (SNS, TGW, GL, GW, and GWLR), grain filling-related traits (Vmax, V, and T), and growth period-related traits (HD and FD). The measurement methods were shown in File S1. Best linear unbiased prediction (BLUP) was used to calculate the breeding value using the R package lme4 (version 1.1-31).

### SNP genotyping

Wheat 660K microarrays [43] were used to sequence 616 wheat accessions, yielding 552,470 original SNPs. Missing markers were imputed using Beagle (version 5.2) [44] and minor allele frequencies (MAF ≥0.05) were filtered using Plink (version 1.90) [45]. Finally, 429,721 high-quality SNPs were selected for further analysis.

### Scanning for selective sweeps

Selective sweep signals were determined using XP*−*CLR [46], SweeD [47], and VCFtools [48] that was used to calculate fixation index (*FST*), nucleotide diversity 𝜋 and Tajima’s *D* value. The comparison groups were landraces and cultivars, landraces and 50s, 50s and 60s−70s, 60s−70s and 80s−90s, and 80s−90s and 00s−. Each indicator was swept with the 500 kb windows. The P-value for XP−CLR was set at 0.95. Intervals detected in the top 5% of at least two indicators were considered as selection intervals.

### Population structure analysis

All the markers were used to perform principal component analysis (PCA), which was implemented using *plink* software (version 1.90), and cluster analysis, which was implemented using R package *amap* (version 0.8-19). These analyses were used to elucidate the population structure of 616 wheat accessions in the present study.

### Genome-wide association study

The 3VmrMLM [49,50], BLUPmrMLM [51] (https://github.com/YuanmingZhang65), EMMAX [52], GEMMA [53] and FarmCPU [54] methods were used to perform genome-wide association studies. In these studies, eleven principal components with a variance explanation rate >1% were used to control for population structure, and the kinship matrix was calculated by the corresponding software packages. The software packages were executed with the default parameters. The *P*−value threshold for identifying significant QTLs was set at 0.05/*m* for all the five methods, while the LOD score threshold for suggested QTLs was set as LOD score ≥ 3 for BLUPmrMLM and IIIVmrMLM and 1/*m* for EMMAX, GEMMA, and FarmCPU, where *m* is the number of markers.

### LD analysis

PopLDdecay [55] was used for LD decay analysis, in which the decay of the squared correlation (*r^2^*) of gene frequencies in the population of gametes to half of the maximum value was used to determine search radius in the mining of candidate genes. The candidate genes within the physical intervals of the identified QTLs were extracted from the IWGSC reference genome version 1.0 (https://www.wheatgenome.org/Resources/Annotations/IWGSC-RefSeq-v1.0-annotation).

### Identification of candidate genes

Differential expression analysis was based on transcriptome data from public databases [56,57] using the R package DESeq2 [58] with the criterion of |log_2_FC| > 1 and *P* < 0.05, and the specific grouping is shown in File S1.

Gene annotation in GWAS was performed using eggNOG-mapper (http://eggnog-mapper.embl.de/) and AgBase (https://agbase.arizona.edu/). GO annotation of genes in selection intervals was performed using GFAP [59] with corrected *P*-values <0.05.

Known rice genes were obtained from the China Rice Data Centre (https://ricedata.cn/gene/). Wheat protein sequences (https://urgi.versailles.inra.fr/, version 1.0) and rice protein sequences (http://rice.uga.edu/, version 7.0) were used to find the homologous genes using the orthofinder [60]. Candidate genes for wheat grain filling rate also included known wheat grain-related genes from WheatOmics 1.0 (http://202.194.139.32/) and reported literature.

The potential candidate genes that were significant in the haplotype analysis using one−way ANOVA were considered as candidate genes, in which the markers in the region of each candidate gene were used to construct the haplotypes of the candidate gene. The superior haplotype was determined by multiple comparisons and its genetic value was the trait average value of the accessions with the superior haplotype.

### Genomic selection

Genomic selection was conducted by R package rrBLUP [61] (version 4.6.3) using the BLUP values of each trait and the SIG and SUG markers associated with the trait and identified by all the methods. And five−fold cross-validation was used to assess the accuracy, while the training dataset and validation dataset were obtained by createFolds function of R package caret. Correlation analysis was conducted by R cor function.

### Quantitative real-time PCR for candidate genes

On the first day after anthesis, Zhengmai 7698 tissues were collected in three biological replicates. These issues included the root, stem, spike, flag leaf, leaf sheath, and tiller base. The total RNA was extracted according to the instructions of the Trizol reagent kit (Invitrogen, Carlsbad, CA, USA). The quality of the extracted RNA was verified by means of RNase-free agarose gel electrophoresis. Subsequently, cDNA was synthesized by using the MonScript RTIII All-in-One Mix with dsDNase kit (MR05101, Monad Biotech Co., LTD, Suzhou, China). The yield and purity of the extracted cDNA were then analyzed using a spectrophotometer (NanoDrop2000, Thermo Scientific, USA). The subsequent 20 µl PCR reaction was prepared according to the instructions of the TB Green® Premix Ex Taq™ II (Tli RNaseH Plus) kit (RR820A, Takara Biotech Co., LTD, Dalian, China). Each sample was subjected to three technical replicates, with TaActin serving as the reference gene. The PCR reactions were subsequently performed using the BioRad CFX Connect Real-time System fluorescence PCR quantification instrument, adhering to the following PCR conditions: Initial denaturation at 95°C for three minutes; followed by 40 cycles of 95° C for 10 seconds and 60°C for 30 seconds. The calculation of relative gene expression levels was performed by using the 2^-△△Ct^ method. Primer design for quantitative real-time PCR was conducted using Primer Premier 6.0, with the specific primers being designed based on the CDS sequence of the corresponding gene ID in IWGSC RefSeq v1.0. The specific primer information can be found in Table A3 of File S1. The expression level of each candidate gene in root tissue was then utilized as a control.

## Data availability

The variation data reported in this paper have been deposited in the Genome Variation Map (GVM) in National Genomics Data Center, Beijing Institute of Genomics, Chinese Academy of Sciences and China National Center for Bioinformation, under accession number GVM000856 (BioProject: PRJCA030529), and are publicly accessible at https://ngdc.cncb.ac.cn/bioproject/browse/PRJCA030529.

## Code availability

The code is publicly accessible at https://ngdc.cncb.ac.cn/biocode.

## CRediT author statement

**Chaojun Peng:** Investigation, Data curation, Writing − review. **Ying Chen:** Writing − original draft & editing, Data curation, Form analysis. **Xuelian Han:** Writing − original draft, Form analysis. **Haibin Dong**: Investigation. **Ao Zheng:** Writing − original draft, Form analysis. **Xijun Du:** Investigation. **Xiaoyu Chang:** Form analysis. **Miaomiao Zhao:** Writing − original draft. **Xueli Qi:** Investigation. **Yuanming Zhang:** Writing − review & editing, Funding acquisition, Project administration. **Lin Hu:** Writing − review & editing, Funding acquisition, Project administration.

## Competing interest

The authors declared that they have no conflict of interest to this work.

## Acknowledgments

This work was supported by the National Key Research and Development Program of China (2021YFD1200603), the National Natural Science Foundation of China (32270673; 32470657), the Science and Technology R&D Program Joint Fund of Henan Province (222301420025), earmarked fund for CARS-wheat (CARS–03–7).

## Supplementary material

**File S1 Supplementary information**

**Figure S1 Geographic distribution of the planting locations**

**Figure S2 Decay of linkage disequilibrium (LD) in all sub-genomes and whole genome**

A sub-genome: red line, B sub-genome: blue line, D sub-genome: black line and whole genome: green line.

PH: plant height, SL: spike length, PPR: ratio of peduncle length to plant height.

**Figure S3 Manhattan plots for plant height, spike length and ratio of peduncle length to plant height**

**Figure S4 Manhattan plots for grain length, grain width, ratio of grain width to length and spikelet number per spike**

GL: grain length, GW: grain width, GWLR: ratio of grain width to length, SNS: spikelet number per spike.

**Figure S5 Manhattan plots for active filling period, maximum filling rate and heading date**

Vmax: maximum filling rate, T: active filling period, HD: heading date.

**Figure S6 The genes identified for average filling rate in wheat**

**A.** Manhattan plot for average filling rate (V). IIIVmrMLM: red; BLUPmrMLM: orange, FarmCPU: blue; EMMAX: green; GEMMA: cyan. Genes with black boxes are detected by multiple methods. B, C and D. Identification of *TraesCS7A01G549300*, *TraesCS7A01G549100* and *TraesCS2A01G293400*, respectively. The superior haplotype is represented by the black color in the first column, the comparisons of the haplotypes are presented in the second column, the frequencies of the haplotypes are shown in the third column, and the expression levels of the genes in grains in the 2, 14, and 30 days after pollination are displayed in the fourth column.

**Figure S7 The genes identified for flowering date in wheat**

**A.** Manhattan plot for flowering date (FD). IIIVmrMLM: red; BLUPmrMLM: orange, FarmCPU: blue; EMMAX: green; GEMMA: cyan. Genes with black boxes are detected by multiple methods. **B, C and D.** Identification of *TraesCS5D01G403200*. The superior haplotype is represented by the black color in the first column, the comparisons of the haplotypes are presented in the second column, the frequencies of the haplotypes are shown in the third column, and the expression levels of the genes in grains in the 2, 14, and 30 days after pollination are displayed in the fourth column.

**Figure S8 Genomic prediction of 13 important traits in wheat using rrBLUP**

PH: plant height, SL: spike length, PPR: ratio of peduncle length to plant height, SNS: spikelet number per spike, TGW: thousand grain weight, GL: grain length, GW: grain width, GWLR: ratio of grain width to length, Vmax: maximum filling rate, V: average grain filling rate, T: active filling period, HD: heading date, FD: flowering date, blue: observation, red: prediction.

**Figure S9 Superior haplotype frequencies for 13 traits in landraces and cultivars with different periods**

PH: plant height, SL: spike length, PPR: ratio of peduncle length to plant height, SNS: spikelet number per spike, TGW: thousand grain weight, GL: grain length, GW: grain width, GWLR: ratio of grain width to length, Vmax: maximum filling rate, V: average grain filling rate, T: active filling period, HD: heading date, FD: flowering date.

**Figure S10 Relative expression levels of each candidate gene in six different issues on the first day after anthesis**

Expression in roots was used as a control

**Figure S11 The asymmetry of gene number in the selection intervals between the A, B and D sub-genomes**

**A.** the number of all genes in sub-genomes. **B**−**F.** the number of genes in the selection intervals of landraces and cultivars, landraces and 50s, 50s and 60s−70s, 60s−70s and 80s−90s, and 80s−90s and 00s−.

**Table S1 List of known genes in selection sweeps between landraces and cultivars with different periods**

**Table S2 Genes in the selection intervals associated with wheat trait changes**

**Table S3 Five selection signal genes out of 28 important genes in Zhengmai 7698**

**Table S4 All QTNs identified to be associated with wheat plant height**

**Table S5 All QTNs identified as associated with spike length in wheat**

**Table S6 All QTNs identified as associated with ratio of peduncle length to plant height in wheat**

**Table S7 All the known genes for three morphological traits in genome-wide association studies and their pieces of evidence**

**Table S8 Candidate genes for three morphological traits in genome-wide association studies and their pieces of evidence**

**Table S9 All QTNs identified as associated with grain length in wheat**

**Table S10 All QTNs identified as associated with grain width in wheat**

**Table S11 All QTNs identified as associated with ratio of grain width to length in wheat**

**Table S12 All QTNs identified as associated with thousand grain weight in wheat**

**Table S13 All QTNs identified as associated with spikelet number per spike in wheat**

**Table S14 All the known genes for five yield-related traits in genome-wide association studies and their pieces of evidence**

**Table S15 Candidate genes for five yield-related traits in genome-wide association studies and their pieces of evidence**

**Table S16 All QTNs identified as associated with active filling period in wheat**

**Table S17 All QTNs identified as associated with average grain filling rate in wheat**

**Table S18 All QTNs identified as associated with maximum filling rate in wheat**

**Table S19 Candidate genes for three grain filling-related traits in genome-wide association studies and their pieces of evidence**

**Table S20 All QTNs identified as associated with flowering date in wheat**

**Table S21 All QTNs identified as associated with heading date in wheat**

**Table S22 All the known genes for two growth period-related wheat traits in genome-wide association studies and their pieces of evidence**

**Table S23 Candidate genes for two growth period-related traits in genome-wide association studies and their pieces of evidence**

**Table S24 The genomic prediction accuracy of thirteen wheat traits using all the significant/suggested QTLs and rrBLUP with fivefold cross-validation**

**Table S25 Frequencies and genetic values of superior haplotypes for eighty genes associated with the trait changes between landraces and cultivars in different time periods**

**Table S26 Frequencies and genetic values of superior genotypes for 50 loci associated with the trait changes between landraces and cultivars in different periods**

**Table S27 Gene superior haplotypes in 30 varieties with the top 5% SNS, V and Vmax phenotypes**

**Table S28 Transmission of superior haplotypes of HXM497 (Zhoumai 17) in its varietal pedigree**

**Table S29 Transmission of superior haplotypes of HXM518 (Zhengmai 7698) in its varietal pedigree**

**Table S30 Transmission of superior haplotypes of HXM523 (Luomai 21) in its varietal pedigree**

**Table S31 Transmission of superior haplotypes of HXM590 (Xinmai 30) in its varietal pedigree**

**Table S32 The superior haplotype frequencies for 13 wheat traits between landraces and cultivars with different periods**

**Table S33 Haplotype frequencies of the genes for important traits in landraces and cultivars with different periods**

**Table S34 Distribution of the superior haplotypes of 164 genes for future breeding**

**Table S35 Four landraces with the most superior haplotypes for future breeding**

**Table S36 The results for accessible chromatin regions of candidate genes for wheat grain-related traits**

**Table S37 GO annotation analysis for the genes with selection signals between landraces and cultivars with different periods**

## Notes

https://ngdc.cncb.ac.cn/bioproject/browse/PRJCA030529

### Competing Interest Statement

The authors have declared no competing interest.

