## Supplementary material for "Genome-wide analysis reveals the genetic basis of key agronomic traits and modern wheat breeding in Henan Province": File S1

**Methods to measure the phenotypic data**

The experimental materials were planted in five environments, namely Xinxiang (35.01°N, 113.70°E), Anyang (36.13°N, 114.58°E), Jiyuan (35.12°N, 112.28°E), Shangqiu (34.24°N, 116.13°E), and Zhumadian (33.01°N, 114.40°E) in Henan Province, in which Xinxiang was in 2016-2017, 2017-2018 and 2018-2019 and others were in 2016-2017 and 2017-2018. Each accession was planted in a 2-m two-row plot with 23.33 cm between rows and 3.33 cm between plants with three replicates.

Here 13 important agronomic traits were measured, including plant height (PH), spike length (SL), ratio of peduncle length to plant height (PPR), spikelet number per spike (SNS), thousand grain weight (TKW), grain length (GL), grain width (GW), ratio of grain width to length (GWLR), heading date (HD), flowering date (FD), maximum filling rate (Vmax), average grain filling rate (V) and active filling period (T).

PH: After the milky stage, 10 individual plants were randomly sampled and the length to the top of the spike, excluding awn, was measured and averaged; SL: At the maturity stage, 10 main panicles were randomly selected from each experimental plot, and the length of the first fruiting spikelet at the base of each spike to the top of the spike (excluding awn) was measured and averaged; PPR: Ratio of length of the first internode in the lower part of the spike to plant height; SNS: At the maturity stage, ten main panicles were randomly selected, and the total number of spikelets born on each main panicle, including sterile spikelets, was investigated and averaged; TKW: The seeds were randomly sampled from the cleared samples, 200 seeds per material were counted in two portions, weighed, and the weight of 1000 seeds was calculated; GL and GW: Ten seeds were randomly sampled from the cleared seeds and placed side by side to measure the total length and width, respectively, and the average value was calculated; GWLR: The ratio of grain width (GW) to grain length (GL); HD: The number of days from sowing of the experimental material until 1cm of the flag leaf sheath was exposed at the top of the spike (excluding awn) of more than 50% of the plants was counted; FD: The number of days from sowing to more than 50% panicle flowering was calculated for the experimental material.

Calculation of parameters related to grain filling rate: 25 single panicles with the same size at flowering stage and panicle type were selected and labeled at flowering stage. The first two samples were taken once every 7 days, and the next three samples were taken once every 5 days. For each experiment, 5 primary panicles were selected from individual plants. The hulled seeds were then fixed at 105℃ for 30 minutes in an oven, followed by further fixation at 80℃ for 16-18 hours until reaching a constant weight. Logistic equation
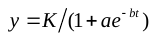
 was used to fit the grain filling process (Yin et al., 2020), where model parameters “k”, “a” and “b” have biological meanings, i.e., “K” is the maximum 1000-grain weight, “a” is the duration of grain filling and filling rate, “b” is filling rate, and *t* is the time after flowering. Based on this equation, the average filling rate (V), the maximum filling rate (Vmax) and the active filling duration (T) were estimated by the integral of the formula
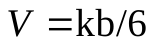
,
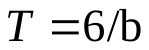
 and
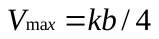
 respectively.

**Table A1**. Information of 616 wheat accessions.

| ID | Group | Name of germplasm | Origin |
| --- | --- | --- | --- |
| HXZ1 | 1950s | Fumai | Sanmenxia, Henan province |
| HXZ2 | Landrace | Qitoufumai (Shanxian1) | Sanmenxia, Henan province |
| HXZ3 | Landrace | Qisiniu | Luoyang, Henan province |
| HXZ4 | Landrace | Baixiantiao | Luoyang, Henan province |
| HXZ5 | Landrace | Baifumai | Luoyang, Henan province |
| HXZ6 | Landrace | Erjimai | Pingdingshan, Henan province |
| HXZ7 | Landrace | Baiheshangtou (Xiaxian) | Pingdingshan, Henan province |
| HXZ8 | Landrace | Huangguaxian (Yuxian) | Xuchang, Henan province |
| HXZ9 | Landrace | Yaerzui | Puyang, Henan province |
| HXZ10 | Landrace | Heshangtou (Anyangxian) | Anyang, Henan province |
| HXZ11 | 1950s | Baiyupi | Unkown, Henan province |
| HXZ12 | Landrace | Shuanghoumai | Anyang, Henan province |
| HXZ13 | Landrace | Zhuganqing (Tangxian) | Anyang, Henan province |
| HXZ14 | Landrace | Baihulutou | Anyang, Henan province |
| HXZ15 | Landrace | Xiaobaimai (Mianchixian) | Sanmenxia, Henan province |
| HXZ16 | Landrace | Huitucao | Luoyang, Henan province |
| HXM17 | Landrace | Ziganyangmai | Luoyang, Henan province |
| HXM18 | Landrace | Baitiaoyu (Yiyangxian) | Luoyang, Henan province |
| HXM19 | Landrace | Dazihong (Yanchengxian) | Luohe, Henan province |
| HXM20 | Landrace | Baiheshangtou | Nanyang, Henan province |
| HXM21 | Landrace | Zhuaziya | Nanyang, Henan province |
| HXM22 | Landrace | Dajiantou | Nanyang, Henan province |
| HXM23 | Landrace | Tiepachimai (Wuzhixian) | Jiaozuo, Henan province |
| HXM24 | Landrace | Xiaofoshou (Jiyuanxian1) | Jiyuan, Henan province |
| HXM25 | Landrace | Sanyuehuang (Wenxian1) | Jiaozuo, Henan province |
| HXM26 | Landrace | Baitutou | Shangqiu, Henan province |
| HXM27 | Landrace | Tuhulutou | Kaifeng, Henan province |
| HXM28 | 1950s | Dakoumai | Luoyang, Henan province |
| HXM29 | Landrace | Guangbaitousanmai | Zhumadian, Henan province |
| HXM30 | 1950s | Huixianhong | Jiaozuo, Henan province |
| HXM31 | Landrace | Sanyuehuang (Wenxian2) | Jiaozuo, Henan province |
| HXM32 | Landrace | Sanyuehuang (Mengxian) | Jiaozuo, Henan province |
| HXM33 | Landrace | Huoshaomai | Shangqiu, Henan province |
| HXM34 | Landrace | Zhuganqing | Jiaozuo, Henan province |
| HXM35 | Landrace | Jinguoyin (Linbaoxian) | Jiyuan, Henan province |
| HXM36 | 1950s | Mazhamai | Luoyang, Henan province |
| HXM37 | Landrace | Hongheshangtou (Baofengxian) | Pingdingshan, Henan province |
| HXM38 | Landrace | Hongheshangtou (Nanyangxian1) | Nanyang, Henan province |
| HXM39 | Landrace | Honggoudou | Jiaozuo, Henan province |
| HXM40 | Landrace | Yangmai | Kaifeng, Henan province |
| HXM41 | Landrace | Baiyintiao | Luoyang, Henan province |
| HXM42 | Landrace | Zigancao (Yinchuanxian) | Luoyang, Henan province |
| HXM43 | Landrace | Baimai (Xichuangxian) | Nanyang, Henan province |
| HXM44 | Landrace | Baimai (Xiangchengxian) | Xuchang, Henan province |
| HXM45 | Landrace | Baipimai | Xinyang, Henan province |
| HXM46 | Landrace | Xiaomanshu | Anyang, Henan province |
| HXM47 | Landrace | Zaosantian | Anyang, Henan province |
| HXM48 | Landrace | Sanyuehuang | Puyang, Henan province |
| HXM49 | Landrace | Ziganbaimangxian | Puyang, Henan province |
| HXM50 | Landrace | Liulinger | Anyang, Henan province |
| HXM51 | Landrace | Xiaobaimang (Puyangxian) | Puyang, Henan province |
| HXM52 | Landrace | Dabaimai | Xinxiang, Henan province |
| HXM53 | Landrace | Wuhuatou (Fengqiu xian) | Xinxiang, Henan province |
| HXM54 | Landrace | Baimangcao (Tongxuxian) | Kaifeng, Henan province |
| HXM55 | Landrace | Duangancao (Zhongmuxian) | Zhengzhou, Henan province |
| HXM56 | Landrace | Baimangcao (Kaifengxian) | Kaifeng, Henan province |
| HXM57 | Landrace | Baitiaoyu (Xinmixian) | Zhengzhou, Henan province |
| HXM58 | Landrace | Gedacao (Luanchuanxian) | Luoyang, Henan province |
| HXM59 | Landrace | Youzitou | Pingdingshan, Henan province |
| HXM60 | Landrace | Laolaiqiao | Xuchang, Henan province |
| HXM61 | Landrace | Xiaozibai (Lushanxian) | Pingdingshan, Henan province |
| HXM62 | Landrace | Hongboyan (Yuxian) | Xuchang, Henan province |
| HXM63 | Landrace | Yulincao (Zhengpingxian) | Nanyang, Henan province |
| HXM64 | Landrace | Gedacao (Fangchenxian) | Shangqiu, Henan province |
| HXZ65 | Landrace | Yanghuomai | Shangqiu, Henan province |
| HXZ66 | Landrace | Nvmai | Nanyang, Henan province |
| HXZ67 | Landrace | Bodijiang (Xichuanxian) | Nanyang, Henan province |
| HXZ68 | Landrace | Baimangcao (Dengxian) | Nanyang, Henan province |
| HXZ69 | Landrace | Kumai (Dengxian) | Nanyang, Henan province |
| HXZ70 | 1950s | Baihuomai | Nanyang, Henan province |
| HXZ71 | Landrace | Meiguocao | Nanyang, Henan province |
| HXZ72 | Landrace | Dachuantiaomai | Nanyang, Henan province |
| HXZ73 | Landrace | Shengsimai | Nanyang, Henan province |
| HXZ74 | Landrace | Jiyulin | Xinxiang, Henan province |
| HXZ75 | Landrace | Duanganchaoxiaomai | Xinxiang, Henan province |
| HXZ76 | Landrace | Sinenggan | Xinxiang, Henan province |
| HXZ77 | Landrace | Wugongcao (Xinxian) | Xinxiang, Henan province |
| HXZ78 | Landrace | Jinbangchui | Xinxiang, Henan province |
| HXZ79 | Landrace | Duangancao (Luoshanxian) | Xinxiang, Henan province |
| HXZ80 | Landrace | Datoucao (Guangshanxian) | Xinxiang, Henan province |
| HXM81 | Landrace | Lvmangcao | Xinxiang, Henan province |
| HXM82 | Landrace | Luanmai | Xinxiang, Henan province |
| HXM83 | Landrace | Silengmai | Xinxiang, Henan province |
| HXM84 | Landrace | Bagucha | Xinxiang, Henan province |
| HXM85 | Landrace | Shuilaomai | Xinxiang, Henan province |
| HXM86 | Landrace | Datoucao (Gushixian) | Xinxiang, Henan province |
| HXM87 | Landrace | Changshengmai | Xinxiang, Henan province |
| HXM88 | Landrace | Wugongcao (Xincaixian) | Zhumadian, Henan province |
| HXM89 | Landrace | Xiaozihong | Zhumadian, Henan province |
| HXM90 | Landrace | Datouben (Lunanxian) | Zhumadian, Henan province |
| HXM91 | Landrace | Jiuyisi | Zhumadian, Henan province |
| HXM92 | Landrace | Changsuimai | Zhumadian, Henan province |
| HXM93 | Landrace | Datouben (Biyangxian1) | Zhumadian, Henan province |
| HXM94 | Landrace | Datouben (Queshanxian) | Zhumadian, Henan province |
| HXM95 | Landrace | Yaozhouhong | Puyang, Henan province |
| HXM96 | Landrace | Baimangben | Puyang, Henan province |
| HXM97 | Landrace | Ziganbai | Shangqiu, Henan province |
| HXM98 | Landrace | Laohongpi | Zhoukou, Henan province |
| HXM99 | Landrace | Jinzihong | Kaifeng, Henan province |
| HXM100 | Landrace | Baidadu | Kaifeng, Henan province |
| HXM101 | Landrace | Wuhuatou (Zhongmuxian) | Zhengzhou, Henan province |
| HXM102 | Landrace | Helubai | Zhengzhou, Henan province |
| HXM103 | Landrace | Hongnangbai (Xixian) | Xinxiang, Henan province |
| HXM104 | Landrace | Daizimai | Xinxiang, Henan province |
| HXM105 | Landrace | Xiaohongbai (Puyangxian) | Puyang, Henan province |
| HXM106 | Landrace | Fanglumai | Puyang, Henan province |
| HXM107 | Landrace | Shizitou | Puyang, Henan province |
| HXM108 | Landrace | Huangguaxian (Kaifengxian) | Kaifeng, Henan province |
| HXM109 | Landrace | Dazipang | Kaifeng, Henan province |
| HXM110 | Landrace | Hongmanghong (Zhongmuxian) | Zhengzhou, Henan province |
| HXM111 | Landrace | Huaiqingfumai | Jiyuan, Henan province |
| HXM112 | Landrace | Hongdanvmai | Luoyang, Henan province |
| HXM113 | Landrace | Jinsihuang | Luoyang, Henan province |
| HXM114 | Landrace | Laoliangkou | Pingdingshan, Henan province |
| HXM115 | Landrace | Hongcantiao | Luoyang, Henan province |
| HXM116 | Landrace | Hongmanghongmai (Yanlin) | Xuchang, Henan province |
| HXM117 | Landrace | Zaobanyue (Zhengpingxian) | Nanyang, Henan province |
| HXM118 | Landrace | Huolibeng | Nanyang, Henan province |
| HXM119 | Landrace | Yixiaomai | Nanyang, Henan province |
| HXM120 | Landrace | Qiuzimai (Shangchenxian) | Xinxiang, Henan province |
| HXM121 | Landrace | Erxiaomai | Nanyang, Henan province |
| HXM122 | Landrace | Zaobanyue (Dengxian) | Nanyang, Henan province |
| HXM123 | Landrace | Huomai (Xichuanxian) | Nanyang, Henan province |
| HXM124 | Landrace | Hongyouzi (Xichuanxian) | Nanyang, Henan province |
| HXM125 | Landrace | Hongquanmang | Nanyang, Henan province |
| HXM126 | Landrace | Xianmai | Xinxiang, Henan province |
| HXM127 | Landrace | Huomai (Xinyangxian) | Xinxiang, Henan province |
| HXM128 | Landrace | Sanyuehuang (Gushixian) | Xinxiang, Henan province |
| HXM129 | Landrace | Loukucao | Xinxiang, Henan province |
| HXM130 | Landrace | Hongsuihong | Zhumadian, Henan province |
| HXM131 | Landrace | Sanhongmai | Zhumadian, Henan province |
| HXM132 | Landrace | Dazihong (Shangcaixian) | Zhumadian, Henan province |
| HXM133 | Landrace | Hongmanghongmai (Nanlexian) | Puyang, Henan province |
| HXM134 | Landrace | Jiandimai | Hebi, Henan province |
| HXM135 | Landrace | Hongjianmai (Fengqiuxian) | Xinxiang, Henan province |
| HXM136 | Landrace | Jianmai (Fengqiuxian) | Xinxiang, Henan province |
| HXM137 | Landrace | Hongjianmai (Yuchenxian) | Shangqiu, Henan province |
| HXM138 | Landrace | Jianmai (Shangqiuxian） | Shangqiu, Henan province |
| HXM139 | Landrace | Qiuzimai (Shangqiuxian) | Shangqiu, Henan province |
| HXM140 | Landrace | Kaosanhong (Shangqiuxian) | Shangqiu, Henan province |
| HXM141 | Landrace | Wuhuatou (Shuixian) | Shangqiu, Henan province |
| HXM142 | Landrace | Dazibai (Shuixian) | Shangqiu, Henan province |
| HXM143 | Landrace | Wuhuatou (Tuochenxian) | Shangqiu, Henan province |
| HXM144 | Landrace | Jianmai (Kaifeng) | Kaifeng, Henan province |
| HXM145 | Landrace | Hongboyan (Zhongmuxian) | Zhengzhou, Henan province |
| HXM146 | Landrace | Baihulu | Zhengzhou, Henan province |
| HXM147 | Landrace | Silengmai (Zhechenxian1) | Shangqiu, Henan province |
| HXM148 | Landrace | Laochusanbao | Luoyang, Henan province |
| HXM149 | Landrace | Huisancao | Luoyang, Henan province |
| HXM150 | Landrace | Siyuecao | Pingdingshan, Henan province |
| HXM151 | Landrace | Duangancao (Xinyangxian) | Xinxiang, Henan province |
| HXM153 | Landrace | Duanganzhuganqing | Jiaozuo, Henan province |
| HXM154 | 1950s | Hongheshangtou | Jiaozuo, Henan province |
| HXM155 | Landrace | Liekoucao | Pingdingshan, Henan province |
| HXM156 | Landrace | Shajiangbai | Luoyang, Henan province |
| HXM157 | Landrace | Yangguangtou | Pingdingshan, Henan province |
| HXM158 | Landrace | Jiacangmai | Xuchang, Henan province |
| HXM159 | Landrace | Quanmangmai (Yuxian) | Xuchang, Henan province |
| HXM160 | Landrace | Xiaoquanmang (Xianchenxian) | Xuchang, Henan province |
| HXZ161 | Landrace | Daquanmang | Luohe, Henan province |
| HXZ162 | Landrace | Liangguangtou | Xuchang, Henan province |
| HXZ163 | Landrace | Tiepachimai (Wuzhixian2) | Jiaozuo, Henan province |
| HXZ164 | Landrace | Baijiantou | Nanyang, Henan province |
| HXZ165 | Landrace | Shandatou | Nanyang, Henan province |
| HXZ166 | Landrace | Heshangtou (Zhengpingxian) | Nanyang, Henan province |
| HXZ167 | Landrace | Siyixiaomai | Nanyang, Henan province |
| HXZ168 | Landrace | Xinpeimai | Pingdingshan, Henan province |
| HXZ169 | Landrace | Baimai (Lushanxian1) | Pingdingshan, Henan province |
| HXZ170 | Landrace | Ermangmai (Biyangxian1) | Zhumadian, Henan province |
| HXZ171 | Landrace | Kumai (Mengxian) | Jiaozuo, Henan province |
| HXZ172 | Landrace | Chushanbao (Xinanxian) | Luoyang, Henan province |
| HXZ173 | Landrace | Yinghuinong | Luoyang, Henan province |
| HXZ174 | Landrace | Baimayidan | Luoyang, Henan province |
| HXZ175 | Landrace | Lengmai | Luoyang, Henan province |
| HXZ176 | Landrace | Yuercao | Luoyang, Henan province |
| HXM177 | Landrace | Quanmangmai | Xuchang, Henan province |
| HXM178 | Landrace | Daquanmai | Xuchang, Henan province |
| HXM179 | Landrace | Cantiaomai (Zhengpingxian) | Nanyang, Henan province |
| HXM180 | Landrace | Baiquanmang | Nanyang, Henan province |
| HXM181 | Landrace | Deguohong | Jiaozuo, Henan province |
| HXM182 | Landrace | Kuquanmang | Kaifeng, Henan province |
| HXM183 | Landrace | Hongyouzitou (Linluxian) | Zhumadian, Henan province |
| HXM184 | Landrace | Hongqulianmang | Nanyang, Henan province |
| HXM185 | Landrace | Chushanbao (Songxian1) | Luoyang, Henan province |
| HXM186 | Landrace | Qitoufumai (Shanxian2) | Luoyang, Henan province |
| HXM187 | Landrace | Cantiaomai (Luanchuanxian) | Luoyang, Henan province |
| HXM188 | Landrace | Huisanbao | Luoyang, Henan province |
| HXM189 | Landrace | Bodijiang (Lushixian) | Luoyang, Henan province |
| HXM190 | Landrace | Juanmaohu | Luoyang, Henan province |
| HXM191 | Landrace | Duanermang | Pingdingshan, Henan province |
| HXM192 | Landrace | Pishanba | Nanyang, Henan province |
| HXM193 | Landrace | Ermangmai (Nanzhaoxian) | Nanyang, Henan province |
| HXM194 | Landrace | Ermangmai (Zhengpingxian) | Nanyang, Henan province |
| HXM195 | Landrace | Ermangmai (Fangchenxian) | Nanyang, Henan province |
| HXM196 | Landrace | Lvweibamai | Jiaozuo, Henan province |
| HXM197 | Landrace | Hongquanmang | Nanyang, Henan province |
| HXM198 | Landrace | Zhangfeihu | Jiyuan, Henan province |
| HXM199 | Landrace | Wangshimai | Jiaozuo, Henan province |
| HXM200 | Landrace | Saogudan | Puyang, Henan province |
| HXM201 | Landrace | Youmai | Puyang, Henan province |
| HXM202 | Landrace | Baisuimai | Kaifeng, Henan province |
| HXM203 | Landrace | Lengdingzi | Kaifeng, Henan province |
| HXM204 | Landrace | Silengmai (Zhechenxian2) | Shangqiu, Henan province |
| HXM205 | Landrace | Baimangmai (Xiangchen1) | Xuchang, Henan province |
| HXM206 | Landrace | Dawangshuibai | Xinxiang, Henan province |
| HXM207 | Landrace | Kumai (Zhengpingxian) | Nanyang, Henan province |
| HXM208 | Landrace | Youzitou (Suopingxian) | Zhumadian, Henan province |
| HXM209 | Landrace | Youzitou (Lunanxian) | Zhumadian, Henan province |
| HXM220 | Landrace | Quermang | Shangqiu, Henan province |
| HXM221 | Landrace | Youmangsaogudan | Anyang, Henan province |
| HXM222 | Landrace | Hongmai | Jiaozuo, Henan province |
| HXM223 | Landrace | Gedacao (Xiangchenxian1) | Xuchang, Henan province |
| HXM224 | Landrace | Baishapi | Kaifeng, Henan province |
| HXM225 | Landrace | Hongmangbai (Zhechenxian) | Shangqiu, Henan province |
| HXM226 | Landrace | Bodanbamai | Nanyang, Henan province |
| HXM227 | Landrace | Honghulutou | Nanyang, Henan province |
| HXM228 | Landrace | Hongyouzitou (Nanlexian) | Nanyang, Henan province |
| HXM229 | Landrace | Baimangyouzi | Anyang, Henan province |
| HXM230 | Landrace | Youzimai (Qinfengxian) | Puyang, Henan province |
| HXM231 | Landrace | Xiaobaimang (Xunxian) | Puyang, Henan province |
| HXM232 | Landrace | Hanbadan | Puyang, Henan province |
| HXM233 | Landrace | Yududasuibai | Xinxiang, Henan province |
| HXM234 | Landrace | Kumai (Nanyangxian) | Nanyang, Henan province |
| HXM235 | Landrace | Laolaihong | Jiaozuo, Henan province |
| HXM236 | Landrace | Xiaofoshou (Jiyuan2) | Jiyuan, Henan province |
| HXM237 | Landrace | Hongtutou (Jiyuan) | Jiyuan, Henan province |
| HXM238 | Landrace | Baigedatou | Jiaozuo, Henan province |
| HXM239 | Landrace | Fangtoumai (Xiuwuxian) | Jiaozuo, Henan province |
| HXM240 | Landrace | Banjiemang (Xiuwuxian) | Jiaozuo, Henan province |
| HXM241 | Landrace | Youzimai (Fengqiuxian) | Xinxiang, Henan province |
| HXM242 | Landrace | Xiaohongmang (Zhengxian) | Zhengzhou, Henan province |
| HXM243 | Landrace | Hongqumang | Zhengzhou, Henan province |
| HXM244 | Landrace | Zaobanyue (Xichuanxian) | Nanyang, Henan province |
| HXM245 | Landrace | Hanzaobadou | Zhengzhou, Henan province |
| HXM246 | Landrace | Hongtutou (Lankaoxian) | Kaifeng, Henan province |
| HXM247 | Landrace | Toulongbei | Xuchang, Henan province |
| HXM248 | Landrace | Tutoumai (Qixian) | Kaifeng, Henan province |
| HXM249 | Landrace | Hongmanghong (Zhechenxian) | Shangqiu, Henan province |
| HXM250 | Landrace | Misuimai | Shangqiu, Henan province |
| HXM251 | Landrace | Sanyuehuang (Luoyang) | Luoyang, Henan province |
| HXM252 | Landrace | Laohongmai | Luoyang, Henan province |
| HXM253 | Landrace | Chushanbao (Songxian2) | Luoyang, Henan province |
| HXM254 | Landrace | Yingsizhuang (Songxian) | Luoyang, Henan province |
| HXM255 | Landrace | Sanyuehuang (Lushixian) | Luoyang, Henan province |
| HXM256 | Landrace | Zhiliaomai | Luoyang, Henan province |
| HXZ257 | Landrace | Xiaosanyuehuang | Luoyang, Henan province |
| HXZ258 | Landrace | Baigeda | Luoyang, Henan province |
| HXZ259 | Landrace | Bodijiang | Luoyang, Henan province |
| HXZ260 | Landrace | Huihuicao | Luoyang, Henan province |
| HXZ261 | Landrace | Huomai (Lushixian) | Luoyang, Henan province |
| HXZ262 | Landrace | Bensiyuehuang | Luoyang, Henan province |
| HXZ263 | Landrace | Heshangtou (Luanchuanxian) | Luoyang, Henan province |
| HXZ264 | Landrace | Honggancao | Luoyang, Henan province |
| HXZ265 | Landrace | Xiaobaimai (Linbaoxian) | Jiyuan, Henan province |
| HXZ266 | Landrace | Baigeda | Jiyuan, Henan province |
| HXZ267 | Landrace | Hongheshangtou (Linbaoxian) | Jiyuan, Henan province |
| HXZ268 | Landrace | Jinguoyin (Sanmenxia) | Sanmenxia, Henan province |
| HXZ269 | Landrace | Yingsizhuang (Yichuanxian) | Luoyang, Henan province |
| HXZ270 | Landrace | Yingbazhuang | Sanmenxia, Henan province |
| HXZ271 | Landrace | Quanyangmai | Xuchang, Henan province |
| HXZ272 | Landrace | Huangguaxian (Yanchengxian) | Luohe, Henan province |
| HXM273 | Landrace | Xiaozibai (Yuxian) | Xuchang, Henan province |
| HXM274 | Landrace | Hongmangbaimai | Xuchang, Henan province |
| HXM275 | Landrace | Hongheshangtou (Biyangxian) | Zhumadian, Henan province |
| HXM276 | Landrace | Ermangmai (Biyangxian2) | Zhumadian, Henan province |
| HXM277 | Landrace | Mazhamei | Zhumadian, Henan province |
| HXM278 | Landrace | Datouben (Biyangxian2) | Zhumadian, Henan province |
| HXM279 | Landrace | Sanbaimai | Zhumadian, Henan province |
| HXM280 | Landrace | Yumai (Pingyuxian) | Zhumadian, Henan province |
| HXM281 | Landrace | Youzimai (Pingyuxian) | Zhumadian, Henan province |
| HXM282 | Landrace | Hongmangbai (Shangcaixian) | Zhumadian, Henan province |
| HXM283 | Landrace | Youzimai (Shangcaixian) | Zhumadian, Henan province |
| HXM284 | 1950s | Wangshuibai | Xinyang, Henan province |
| HXM285 | Landrace | Wangshanbai | Xinyang, Henan province |
| HXM286 | Landrace | Baiheshangtou (Xinxian) | Xinyang, Henan province |
| HXM287 | Landrace | Ercaomai (Huanyangxian) | Zhoukou, Henan province |
| HXM288 | Landrace | Simangmai | Zhoukou, Henan province |
| HXM289 | Landrace | Youzimai (Taikangxian) | Zhoukou, Henan province |
| HXM290 | Landrace | Hongsuibai | Shangqiu, Henan province |
| HXM291 | Landrace | Zigancao (Shenqiuxian) | Zhoukou, Henan province |
| HXM292 | Landrace | Hongmangmai (Nanyangxian) | Nanyang, Henan province |
| HXM293 | Landrace | Zhuganqing (Nanyangxian) | Nanyang, Henan province |
| HXM294 | Landrace | Datouhong (Neixiangxian) | Nanyang, Henan province |
| HXM295 | Landrace | Hongyouzi (Zhengpingxian) | Nanyang, Henan province |
| HXM296 | Landrace | Hongmangmai (Fangchenxian) | Nanyang, Henan province |
| HXM297 | Landrace | Pushanbamai | Nanyang, Henan province |
| HXM298 | Landrace | Hongquanmang (Dengxian) | Nanyang, Henan province |
| HXM299 | Landrace | Datouhong (Dengxian) | Nanyang, Henan province |
| HXM300 | Landrace | Yumai (Dengxian) | Nanyang, Henan province |
| HXM303 | Landrace | Hongquanmang (Lushanxian) | Zhumadian, Henan province |
| HXM304 | Landrace | Yumai (Yexian) | Pingdingshan, Henan province |
| HXM305 | Landrace | Ercaomai (Xuchangxian1) | Xuchang, Henan province |
| HXM306 | Landrace | Ercaomai (Xuchangxian2) | Xuchang, Henan province |
| HXM307 | 1950s | Sanyuehuang (Linyinxian) | Xuchang, Henan province |
| HXM308 | Landrace | Sanyuehuang (Yexian) | Pingdingshan, Henan province |
| HXM309 | Landrace | Sanmangmai | Pingdingshan, Henan province |
| HXM310 | Landrace | Sanbaoxian | Xuchang, Henan province |
| HXM311 | Landrace | Damai | Pingdingshan, Henan province |
| HXM312 | Landrace | Dazibai (Changgexian) | Xuchang, Henan province |
| HXM313 | Landrace | Dazihong (Xuchangxian) | Xuchang, Henan province |
| HXM314 | Landrace | Dazicao (Xuchangxian) | Xuchang, Henan province |
| HXM315 | Landrace | Dazicao (Yexian) | Pingdingshan, Henan province |
| HXM316 | Landrace | Xiaoguangtou | Xuchang, Henan province |
| HXM317 | Landrace | Xiaozibai (Xuchangxian) | Xuchang, Henan province |
| HXM318 | Landrace | Xiaozicao | Xuchang, Henan province |
| HXM319 | Landrace | Xiaoquanmang (Linyinxian) | Luohe, Henan province |
| HXM320 | Landrace | Wuhuatou (Linyinxian) | Luohe, Henan province |
| HXM321 | Landrace | Wuhuatou (Xihuaxian1) | Zhoukou, Henan province |
| HXM322 | Landrace | Wuhuatou (Xihuaxian2) | Zhoukou, Henan province |
| HXM323 | Landrace | Changzicao | Xuchang, Henan province |
| HXM324 | Landrace | Wukelan | Xuchang, Henan province |
| HXM325 | Landrace | Yubianmai | Xuchang, Henan province |
| HXM326 | Landrace | Baimanghong (Yanlinxian) | Xuchang, Henan province |
| HXM327 | Landrace | Baimanghong (Xihuaxian1) | Zhoukou, Henan province |
| HXM328 | Landrace | Baimanghong (Xihuaxian2) | Zhoukou, Henan province |
| HXM329 | Landrace | Baimanghong (Xihuaxian3) | Zhoukou, Henan province |
| HXM330 | Landrace | Baimanghong (Yexian) | Pingdingshan, Henan province |
| HXM331 | Landrace | Baimangmai (Xiangchen2) | Xuchang, Henan province |
| HXM332 | Landrace | Baimai (Linyinxian) | Luohe, Henan province |
| HXM333 | Landrace | Baimai (Yexian) | Pingdingshan, Henan province |
| HXM334 | Landrace | Baimai (Lushanxian2) | Pingdingshan, Henan province |
| HXM335 | Landrace | Baikeheimai | Xuchang, Henan province |
| HXM336 | Landrace | Baizhamang | Xuchang, Henan province |
| HXM337 | Landrace | Baiquanmang | Xuchang, Henan province |
| HXM338 | Landrace | Baiyouzitou | Pingdingshan, Henan province |
| HXM339 | Landrace | Baishenmai | Xuchang, Henan province |
| HXM340 | Landrace | Youmanggedacao | Pingdingshan, Henan province |
| HXM341 | Landrace | Huitoucao | Pingdingshan, Henan province |
| HXM342 | Landrace | Guangtoumai | Xuchang, Henan province |
| HXM343 | Landrace | Hongmangbai (Xihuaxian) | Zhoukou, Henan province |
| HXM344 | Landrace | Hongmanghong (Xuchangxian) | Xuchang, Henan province |
| HXM345 | Landrace | Hongmangmai (Wuyangxian) | Pingdingshan, Henan province |
| HXM346 | Landrace | Hongmangmai (Changgexian) | Xuchang, Henan province |
| HXM347 | Landrace | Hongmangshen | Xuchang, Henan province |
| HXM348 | Landrace | Hongmangben | Pingdingshan, Henan province |
| HXM349 | Landrace | Hongmangcao (Xihuaxian) | Zhoukou, Henan province |
| HXM350 | Landrace | Hongmangcao (Yanlinxian) | Xuchang, Henan province |
| HXM351 | Landrace | Hongheshangtou (Lushanxian) | Pingdingshan, Henan province |
| HXM352 | Landrace | Hongyouzitou (Jiaxian) | Pingdingshan, Henan province |
| HXZ353 | Landrace | Hongyouzitou (Lushanxian) | Pingdingshan, Henan province |
| HXZ354 | Landrace | Hongsuimai | Pingdingshan, Henan province |
| HXZ355 | Landrace | Yuanzicao | Xuchang, Henan province |
| HXZ356 | Landrace | Tutoumai (Yexian) | Pingdingshan, Henan province |
| HXZ357 | Landrace | Yulincao (Xiangchenxian) | Xuchang, Henan province |
| HXZ358 | Landrace | Gedacao (Xiangchenxian2) | Xuchang, Henan province |
| HXZ359 | 1950s | Hulutou | , Henan province |
| HXZ360 | Landrace | Nanyanghong | Pingdingshan, Henan province |
| HXZ361 | Landrace | Quanmang | Pingdingshan, Henan province |
| HXZ362 | Landrace | Quanmangmai (Xihuaxian) | Zhoukou, Henan province |
| HXZ363 | Landrace | Quanmangcao | Pingdingshan, Henan province |
| HXZ364 | Landrace | Huangguaxian (Xihuaxian) | Zhoukou, Henan province |
| HXZ365 | Landrace | Huangguaxian (Jiaxian) | Pingdingshan, Henan province |
| HXZ366 | Landrace | Youzitou (Yanlinxian) | Xuchang, Henan province |
| HXZ367 | Landrace | Youzitou (Wuyangxian) | Pingdingshan, Henan province |
| HXZ368 | Landrace | Zigancao (Xuchangxian) | Xuchang, Henan province |
| HXM369 | Landrace | Ziboyan | Pingdingshan, Henan province |
| HXM370 | Landrace | Hunanbai | Xuchang, Henan province |
| HXM371 | Landrace | Hunanhong | Xuchang, Henan province |
| HXM372 | Landrace | Pushanba (Shanxian) | Pingdingshan, Henan province |
| HXM373 | Landrace | Pushanba (Yexian) | Pingdingshan, Henan province |
| HXM374 | 1950s | Pingyuan 50 | Jiaozuo, Henan province |
| HXM375 | 1950s | Youzimai | Puyang, Henan province |
| HXM376 | 1950s | Xuzhou 438 | Xuzhou, Jiangsu province |
| HXM377 | 1950s | Kaifeng 124 | Kaifeng, Henan province |
| HXM378 | 1950s | Biyumai | Australia |
| HXM379 | 1950s | Danmai 1 | Denmark |
| HXM380 | 1950s | Bima 1 | Yanglin, Shanxi province |
| HXM381 | 1950s | Bima 4 | Yanglin, Shanxi province |
| HXM382 | 1950s | Bima 6 | Yanglin, Shanxi province |
| HXM383 | 1950s | Xinong 6028 | Yanglin, Shanxi province |
| HXM384 | 1950s | Nanda 2419 | Nanjing, Jiangsu province |
| HXM385 | 1950s | Zaoyangmai | Beijing |
| HXM386 | 1960s-1970s | Xuzhou 8 | Xuzhou, Jiangsu province |
| HXM387 | 1960s-1970s | Zhengzhou 15 | Zhengzhou, Henan province |
| HXM388 | 1960s-1970s | Zhengzhou 24 | Zhengzhou, Henan province |
| HXM389 | 1960s-1970s | Jinan 4 | Jinan, Shangdong province |
| HXM390 | 1950s | Afu | Italy |
| HXM391 | 1950s | Abo | Italy |
| HXM392 | 1950s | Neixiang 36 | Nanyang, Henan province |
| HXM393 | 1950s | Neixiang 5 | Nanyang, Henan province |
| HXM394 | 1960s-1970s | Bonong 7422 | Jiaozuo, Henan province |
| HXM395 | 1950s | Tiegancao | Xuchang, Henan province |
| HXM396 | 1950s | Xiannong 39 | Xianyang, Shanxi province |
| HXM397 | 1950s | Mala | Russia |
| HXM398 | 1950s | Gaojiasuo | Russia |
| HXM399 | 1950s | Shanqianmai | Russia |
| HXM400 | 1950s | Luofulin 10 | Romania |
| HXM401 | 1960s-1970s | Aimengniu | Jinan, Shangdong province |
| HXM402 | 1950s | Fulanni | Russia |
| HXM403 | 1950s | Zhaomai 2 | Nanyang, Henan province |
| HXM404 | 1960s-1970s | Zhengmai 6 | Zhengzhou, Henan province |
| HXM405 | 1960s-1970s | Zhengmai 5 | Zhengzhou, Henan province |
| HXM406 | 1960s-1970s | Boai 7023 | Jiaozuo, Henan province |
| HXM407 | 1960s-1970s | Zhengzhou 683 | Zhengzhou, Henan province |
| HXM408 | 1960s-1970s | Zhengyin 1 | Zhengzhou, Henan province |
| HXM409 | 1960s-1970s | Zhengyin 4 | Zhengzhou, Henan province |
| HXM410 | 1960s-1970s | Xuzhou 14 | Xuzhou, Jiangsu province |
| HXM411 | 1960s-1970s | Puyang 5 | Puyang, Henan province |
| HXM412 | 1960s-1970s | Anxuan 5 | Anyang, Henan province |
| HXM413 | 1960s-1970s | Yanda 25 | Luoyang, Henan province |
| HXM414 | 1960s-1970s | Xiaoyan 4 | Yanglin, Shanxi province |
| HXM415 | 1960s-1970s | Zhengzhou 17 | Zhengzhou, Henan province |
| HXM416 | 1960s-1970s | Zhengzhou 761 | Zhengzhou, Henan province |
| HXM417 | 1960s-1970s | Zhengliufu | Zhengzhou, Henan province |
| HXM418 | 1960s-1970s | Aifeng 3 | Yanglin, Shanxi province |
| HXM419 | 1960s-1970s | Zhengzhou 722 | Zhengzhou, Henan province |
| HXM420 | 1950s | Fengchan 1 | Yanglin, Shanxi province |
| HXM421 | 1950s | Fengchan 3 | Yanglin, Shanxi province |
| HXM422 | 1960s-1970s | Xiaoyan 6 | Yanglin, Shanxi province |
| HXM423 | 1960s-1970s | Yanshi 4 | Luoyang, Henan province |
| HXM424 | 1960s-1970s | Xian 8 | Xian, Shanxi province |
| HXM425 | 1980s-1990s | Bainong 3217 | Jiaozuo, Henan province |
| HXM426 | 1980s-1990s | Yumai 1 | Zhengzhou, Henan province |
| HXM427 | 1980s-1990s | Yumai 2 | Pingdingshan, Henan province |
| HXM428 | 1980s-1990s | Yumai 4 | Luoyang, Henan province |
| HXM429 | 1980s-1990s | Yumai 5 | Zhengzhou, Henan province |
| HXM430 | 1980s-1990s | Yumai 6 | Luoyang, Henan province |
| HXM431 | 1980s-1990s | Yumai 7 | Luoyang, Henan province |
| HXM432 | 1980s-1990s | Yumai 8 | Luoyang, Henan province |
| HXM433 | 1980s-1990s | Yumai 10 | Xian, Shanxi province |
| HXM434 | 1980s-1990s | Yumai 12 | Anyang, Henan province |
| HXM435 | 1980s-1990s | Yumai 13 | Zhengzhou, Henan province |
| HXM436 | 1980s-1990s | Xuzhou 21 | Xuzhou, Jiangsu province |
| HXM437 | 1980s-1990s | Shannong 7859 | Yanglin, Shanxi province |
| HXM438 | 1980s-1990s | Een 1 | Enshi, Hubei province |
| HXM439 | 1980s-1990s | Zhengtaiyu 1 | Zhengzhou, Henan province |
| HXM440 | 1980s-1990s | Yumai 14 | Luoyang, Henan province |
| HXM441 | 1980s-1990s | Yumai 15 | Zhoukou, Henan province |
| HXM442 | 1980s-1990s | Yumai 16 | Zhengzhou, Henan province |
| HXM443 | 1980s-1990s | Yumai 17 | Nanyang, Henan province |
| HXM444 | 1980s-1990s | Yumai 18 | Luoyang, Henan province |
| HXM445 | 1980s-1990s | Yumai 19 | Kaifeng, Henan province |
| HXM446 | 1980s-1990s | Yumai 21 | Zhengzhou, Henan province |
| HXM447 | 1980s-1990s | Yumai 24 | Puyang, Henan province |
| HXM448 | 1980s-1990s | Yumai 25 | Jiaozuo, Henan province |
| HXZ449 | 1980s-1990s | Yumai 26 | Shangqiu, Henan province |
| HXZ450 | 1980s-1990s | Yumai 28 | Zhengzhou, Henan province |
| HXZ451 | 1980s-1990s | Yumai 29 | Zhengzhou, Henan province |
| HXZ452 | 1980s-1990s | Yumai 30 | Shangqiu, Henan province |
| HXZ453 | 1980s-1990s | Yumai 32 | Jiaozuo, Henan province |
| HXZ454 | 1980s-1990s | Yumai 33 | Zhengzhou, Henan province |
| HXZ455 | 1980s-1990s | Yumai 34 | Zhengzhou, Henan province |
| HXZ456 | 1980s-1990s | Yumai 35 | Nanyang, Henan province |
| HXZ457 | 1980s-1990s | Yumai 36 | Zhengzhou, Henan province |
| HXZ458 | 1980s-1990s | Yumai 40 | Kaifeng, Henan province |
| HXZ459 | 1980s-1990s | Yumai 41 | Jiaozuo, Henan province |
| HXZ460 | 1980s-1990s | Yumai 43 | Zhengzhou, Henan province |
| HXZ461 | 1980s-1990s | Zhongyu 3 | Anyang, Henan province |
| HXZ462 | 1980s-1990s | Yumai 45 | Zhengzhou, Henan province |
| HXZ463 | 1980s-1990s | Yumai 47 | Zhengzhou, Henan province |
| HXZ464 | 1980s-1990s | Yumai 48 | Luoyang, Henan province |
| HXM465 | 1980s-1990s | Yumai 49 | Jiaozuo, Henan province |
| HXM466 | 1980s-1990s | Yumai 50 | Zhengzhou, Henan province |
| HXM467 | 1980s-1990s | Yumai 51 | Zhoukou, Henan province |
| HXM468 | 1980s-1990s | Yumai 52 | Zhengzhou, Henan province |
| HXM469 | 1980s-1990s | Yumai 54 | Jiaozuo, Henan province |
| HXM470 | 1980s-1990s | Yumai 55 | Zhengzhou, Henan province |
| HXM471 | 1980s-1990s | Yumai 56 | Zhengzhou, Henan province |
| HXM472 | 1980s-1990s | Yumai 57 | Luohe, Henan province |
| HXM473 | 1980s-1990s | Yumai 58 | Jiaozuo, Henan province |
| HXM474 | 1980s-1990s | Yumai 59 | Zhengzhou, Henan province |
| HXM475 | 1980s-1990s | Yumai 60 | Zhengzhou, Henan province |
| HXM476 | 1980s-1990s | Yumai 61 | Zhengzhou, Henan province |
| HXM477 | 1980s-1990s | Yumai 62 | Zhoukou, Henan province |
| HXM478 | 1980s-1990s | Yumai 63 | Luoyang, Henan province |
| HXM479 | 1980s-1990s | Yumai 64 | Shangqiu, Henan province |
| HXM480 | 1980s-1990s | Yumai 67 | Jiaozuo, Henan province |
| HXM481 | 1980s-1990s | Yumai 68 | Zhengzhou, Henan province |
| HXM482 | Post-2000 | Yumai 69 | Xinxiang, Henan province |
| HXM483 | Post-2000 | Yumai 70 | Nanyang, Henan province |
| HXM484 | Post-2000 | Zhongyu 5 | Anyang, Henan province |
| HXM485 | Post-2000 | Zhongyu 6 | Anyang, Henan province |
| HXM486 | 1980s-1990s | Zhoumai 11 | Zhoukou, Henan province |
| HXM487 | Post-2000 | Luomai 1 | Luoyang, Henan province |
| HXM488 | Post-2000 | Luohan 2 | Luoyang, Henan province |
| HXM489 | Post-2000 | Luohan 3 | Luoyang, Henan province |
| HXM490 | 1980s-1990s | Zhengmai 9023 | Zhengzhou, Henan province |
| HXM491 | Post-2000 | Aikang 58 | Xinxiang, Henan province |
| HXM492 | Post-2000 | Zhengmai 366 | Zhengzhou, Henan province |
| HXM493 | Post-2000 | Zhengmai 004 | Zhengzhou, Henan province |
| HXM494 | Post-2000 | Yumai 70-36 | Zhengzhou, Henan province |
| HXM495 | 1980s-1990s | Zhoumai 13 | Zhoukou, Henan province |
| HXM496 | Post-2000 | Zhoumai 16 | Zhoukou, Henan province |
| HXM497 | Post-2000 | Zhoumai 17 | Zhoukou, Henan province |
| HXM498 | Post-2000 | Zhoumai 18 | Zhoukou, Henan province |
| HXM499 | Post-2000 | Xuke 718 | Xuchang, Henan province |
| HXM500 | Post-2000 | Xuke 316 | Xuchang, Henan province |
| HXM501 | Post-2000 | Yumai 18-99 | Luoyang, Henan province |
| HXM502 | Post-2000 | Fanmai 5 | Zhengzhou, Henan province |
| HXM503 | Post-2000 | Xinmai 12 | Xinxiang, Henan province |
| HXM504 | Post-2000 | Xinmai 13 | Xinxiang, Henan province |
| HXM505 | Post-2000 | Xinmai 16 | Xinxiang, Henan province |
| HXM506 | Post-2000 | Xinmai 18 | Xinxiang, Henan province |
| HXM507 | Post-2000 | Yanzhan 4110 | Luoyang, Henan province |
| HXM508 | Post-2000 | Jimai 1 | Jinan, Shangdong province |
| HXM509 | Post-2000 | Xinmai 11 | Xinxiang, Henan province |
| HXM510 | Post-2000 | Wan 369 | Nanyang, Henan province |
| HXM511 | Post-2000 | Taikong 6 | Zhengzhou, Henan province |
| HXM512 | Post-2000 | Pumai 9 | Puyang, Henan province |
| HXM513 | Post-2000 | Zhongyuan 98-68 | Zhengzhou, Henan province |
| HXM514 | Post-2000 | Xinmai 208 | Xinxiang, Henan province |
| HXM515 | Post-2000 | Yumai 49-198 | Jiaozuo, Henan province |
| HXM516 | Post-2000 | Zhengmai 9405 | Zhengzhou, Henan province |
| HXM517 | Post-2000 | Zhengmai 9694 | Zhengzhou, Henan province |
| HXM518 | Post-2000 | Zhengmai 7698 | Zhengzhou, Henan province |
| HXM519 | Post-2000 | Zhengmai 0856 | Zhengzhou, Henan province |
| HXM520 | Post-2000 | Zhengmai 0943 | Zhengzhou, Henan province |
| HXM521 | Post-2000 | Zhengmai 379 | Zhengzhou, Henan province |
| HXM522 | Post-2000 | Zhengmai 3596 | Zhengzhou, Henan province |
| HXM523 | Post-2000 | Luomai 21 | Luoyang, Henan province |
| HXM524 | Post-2000 | Luomai 24 | Luoyang, Henan province |
| HXM525 | Post-2000 | Luohan 6 | Luoyang, Henan province |
| HXM526 | Post-2000 | Luohan 10 | Luoyang, Henan province |
| HXM527 | Post-2000 | Luohan 12 | Luoyang, Henan province |
| HXM528 | Post-2000 | Xinmai 19 | Xinxiang, Henan province |
| HXM529 | Post-2000 | Xinmai 23 | Xinxiang, Henan province |
| HXM530 | Post-2000 | Xinmai 26 | Xinxiang, Henan province |
| HXM531 | Post-2000 | Huapei 3 | Zhengzhou, Henan province |
| HXM532 | Post-2000 | Huapei 8 | Zhengzhou, Henan province |
| HXM533 | Post-2000 | Zhoumai 19 | Zhoukou, Henan province |
| HXM534 | Post-2000 | Zhoumai 20 | Zhoukou, Henan province |
| HXM535 | Post-2000 | Zhoumai 22 | Zhoukou, Henan province |
| HXM536 | Post-2000 | Zhoumai 23 | Zhoukou, Henan province |
| HXM537 | Post-2000 | Zhoumai 24 | Zhoukou, Henan province |
| HXM538 | Post-2000 | Zhoumai 26 | Zhoukou, Henan province |
| HXM539 | Post-2000 | Zhoumai 27 | Zhoukou, Henan province |
| HXM540 | Post-2000 | Zhoumai 32 | Zhoukou, Henan province |
| HXM541 | Post-2000 | Fengdecunmai 1 | Zhengzhou, Henan province |
| HXM542 | Post-2000 | Zhongyu 9 | Anyang, Henan province |
| HXM543 | Post-2000 | 04 zhong 36 | Anyang, Henan province |
| HXM544 | Post-2000 | Zhongyu 12 | Anyang, Henan province |
| HXZ545 | Post-2000 | Kaimai 18 | Kaifeng, Henan province |
| HXZ546 | Post-2000 | Jimai 19 | Jinan, Shangdong province |
| HXZ547 | Post-2000 | Jimai 20 | Jinan, Shangdong province |
| HXZ548 | Post-2000 | Jimai 22 | Jinan, Shangdong province |
| HXZ549 | Post-2000 | Pingan 6 | Jiaozuo, Henan province |
| HXZ550 | Post-2000 | Pingan 8 | Jiaozuo, Henan province |
| HXZ551 | Post-2000 | Kaimai 21 | Kaifeng, Henan province |
| HXZ552 | Post-2000 | Bainong 207 | Xinxiang, Henan province |
| HXZ553 | Post-2000 | Zhongmai 1 | Shangqiu, Henan province |
| HXZ554 | Post-2000 | Hengguan 35 | Hengshui, Hebei province |
| HXZ555 | Post-2000 | Kaimai 20 | Kaifeng, Henan province |
| HXZ556 | Post-2000 | Yangao 21 | Luoyang, Henan province |
| HXZ557 | Post-2000 | Xinong 979 | Yanglin, Shanxi province |
| HXZ558 | Post-2000 | Luomai 9 | Luohe, Henan province |
| HXZ559 | Post-2000 | Zhongluotiegan | Luoyang, Henan province |
| HXZ560 | Post-2000 | Xuke 1 | Xuchang, Henan province |
| HXM561 | Post-2000 | Zhengyumai 9987 | Zhengzhou, Henan province |
| HXM562 | Post-2000 | Luomai 22 | Luoyang, Henan province |
| HXM563 | Post-2000 | Luomai 23 | Luoyang, Henan province |
| HXM564 | Post-2000 | Xinmai 21 | Xinxiang, Henan province |
| HXM565 | Post-2000 | Bainong 419 | Xinxiang, Henan province |
| HXM566 | Post-2000 | R 93127 | Zhengzhou, Henan province |
| HXM567 | Post-2000 | Zhengmai 1348 | Zhengzhou, Henan province |
| HXM568 | Post-2000 | Yulong 1325 | Zhengzhou, Henan province |
| HXM569 | Post-2000 | 07H508 | Zhengzhou, Henan province |
| HXM570 | Post-2000 | Zhengmai 1354 | Zhengzhou, Henan province |
| HXM571 | Post-2000 | Zhengmai 1345 | Zhengzhou, Henan province |
| HXM572 | Post-2000 | Zhengmai 1342 | Zhengzhou, Henan province |
| HXM573 | 1980s-1990s | 4B269 | Zhoukou, Henan province |
| HXM574 | 1980s-1990s | Zhou 8425B | Zhoukou, Henan province |
| HXM575 | Post-2000 | 07H508-05-1 | Zhengzhou, Henan province |
| HXM576 | Post-2000 | 07H508-05-2 | Zhengzhou, Henan province |
| HXM577 | Post-2000 | 07H508-05-3 | Zhengzhou, Henan province |
| HXM586 | Post-2000 | Zhoumai 28 | Zhoukou, Henan province |
| HXM587 | Post-2000 | Zhoumai 31 | Zhoukou, Henan province |
| HXM588 | Post-2000 | Xinmai 26-5 | Xinxiang, Henan province |
| HXM589 | Post-2000 | Xinmai 29 | Xinxiang, Henan province |
| HXM590 | Post-2000 | Xinmai 30 | Xinxiang, Henan province |
| HXM591 | Post-2000 | Chunmai 5 | Zhengzhou, Henan province |
| HXM592 | Post-2000 | Chunmai 8 | Zhengzhou, Henan province |
| HXM593 | Post-2000 | Zhongyu 1023 | Anyang, Henan province |
| HXM594 | Post-2000 | Zhongyu 9307 | Anyang, Henan province |
| HXM595 | Post-2000 | Zhongyu 1220 | Anyang, Henan province |
| HXM596 | Post-2000 | Zhongyu 1311 | Anyang, Henan province |
| HXM597 | Post-2000 | Zhongyu 1326 | Anyang, Henan province |
| HXM598 | Post-2000 | Gaoyou 8 | Shijiazhuang, Hebei province |
| HXM599 | Post-2000 | Gaoyou 9828 | Shijiazhuang, Hebei province |
| HXM600 | Post-2000 | Gaoyou 5766 | Shijiazhuang, Hebei province |
| HXM601 | Post-2000 | Gaoyou 2018 | Shijiazhuang, Hebei province |
| HXM602 | Post-2000 | Gaoyou 5218 | Shijiazhuang, Hebei province |
| HXM603 | Post-2000 | Shiluan 02-1 | Shijiazhuang, Hebei province |
| HXM604 | Post-2000 | Shiluan 08-4 | Shijiazhuang, Hebei province |
| HXM605 | Post-2000 | Jichuang 2 | Jiyuan, Henan province |
| HXM606 | Post-2000 | Zhengpinmai 7 | Zhengzhou, Henan province |
| HXM607 | Post-2000 | Luomai 28 | Luoyang, Henan province |
| HXM608 | Post-2000 | Luomai 29 | Luoyang, Henan province |
| HXM609 | Post-2000 | Luomai 31 | Luoyang, Henan province |
| HXM610 | Post-2000 | Guomai 301 | Zhengzhou, Henan province |
| HXM611 | Post-2000 | Huaimai 30 | Huaian, Jiangsu province |
| HXM612 | Post-2000 | Luomai 26 | Luoyang, Henan province |
| HXM613 | Post-2000 | Xuke 168 | Xuchang, Henan province |
| HXM614 | Post-2000 | Bonong 6 | Jiaozuo, Henan province |
| HXM615 | Post-2000 | Luomai 8 | Luohe, Henan province |
| HXM616 | Post-2000 | Xinmai 30 | Xinxiang, Henan province |
| HXM617 | Post-2000 | Guomai 0319 | Zhengzhou, Henan province |
| HXM618 | Post-2000 | Zhengyumai 043 | Zhengzhou, Henan province |
| HXM619 | Post-2000 | Kaimai 22 | Kaifeng, Henan province |
| HXM620 | Post-2000 | Wanmai 98 | Nanyang, Henan province |
| HXM621 | Post-2000 | Zhongmai 875 | Anyang, Henan province |
| HXM622 | Post-2000 | Zhengmai 119 | Zhengzhou, Henan province |
| HXM623 | Post-2000 | Zhengmai 1867 | Zhengzhou, Henan province |
| HXM624 | Post-2000 | 08H250-41-7-12-4-23-1 | Zhengzhou, Henan province |
| HXM625 | Post-2000 | 08H250-41-7-12-4-23-2 | Zhengzhou, Henan province |
| HXM626 | Post-2000 | 08H250-41-7-12-4-23-3 | Zhengzhou, Henan province |
| HXM627 | Post-2000 | 08H250-41-7-12-4-23-4 | Zhengzhou, Henan province |
| HXM628 | Post-2000 | 08H250-41-7-12-4-23-5 | Zhengzhou, Henan province |
| HXM629 | Post-2000 | Zhongmai 895 | Anyang, Henan province |
| HXM632 | 1950s | Sulianzaoshu 1 | Russia |
| HXM634 | 1980s-1990s | Jimai 30 | Shijiazhuang, Hebei province |
| HXM635 | Landrace | Fangtoumai | Jiaozuo, Henan province |
| HXM636 | 1980s-1990s | Yumai 65 | Jiaozuo, Henan province |
| HXM637 | 1960s-1970s | Baiquan 3039 | Jiaozuo, Henan province |
| HXM638 | 1960s-1970s | Baiquan 41 | Jiaozuo, Henan province |
| HXM639 | 1980s-1990s | Shiai 1 | Shijiazhuang, Hebei province |
| HXM640 | 1960s-1970s | Zhengzhou 871 | Zhengzhou, Henan province |

**Table A2**. The designs of differential expression analysis.

| Trait | Tissue | Age | Short |
| --- | --- | --- | --- |
| PH | shoot apical meristem | Seedling stage ~ Tillering stage | PH1 |
| PH | shoot axis | Tillering stage ~ milk grain stage | PH2 |
| PH | spike | Full boot ~ 30% spike | PH3 |
| SL | spike | Full boot ~ 30% spike | SL |
| PPR | shoot axis | Tillering stage ~ milk grain stage | PPR1 |
| PPR | spike | Full boot ~ 30% spike | PPR2 |
| HD | Spikelets~Spike | 30% spike~Full Boot | HD |
| FD | flag leaf blade~stigma & ovary | Anthesis | FD1 |
| FD | flag leaf blade~anther | Anthesis | FD2 |
| SNS | flag leaf blade~spike | 30% spike | SNS1 |
| SNS | flag leaf blade~spike | full boot | SNS2 |
| GL, GW, TKW and GWLR | flag leaf blade~grain | Dough | G |
| Vmax, V and T | grain | 2 ~ 14 DAP | V1 |
| Vmax, V and T | grain | 2 ~ 30 DAP | V2 |

**Table A3**. Primer for qRT-PCR.

| Gene ID | 5'—3' | Primer sequence | Product size |
| --- | --- | --- | --- |
| TraesCS1A03G0483700, TaActin | Forward | CCTGATGGTCAAGTGATCACCA | 180bp |
|  | Reverse | CCCACCACTGAGGACAATGTTA |  |
| TraesCS2A01G552400, TaMYB16 | Forward | TACAGTCTTCTTCTTTCCGGCC | 180bp |
|  | Reverse | TTTACTGCATCCTCAGGGTCAC |  |
| TraesCS6D01G173000, TaGATA | Forward | AAAGAAGGAGAAGAGGGCTGTG | 176bp |
|  | Reverse | CTTTGCTCAGTCTCTCCACCAT |  |
| TraesCS7A01G362300, TaGAD35 | Forward | ACGATGGTTGCTACAAGTGACT | 211bp |
|  | Reverse | GGAACAGCTCGTCCTTGAAAAC |  |
| TraesCS7A01G059000, TaDLT | Forward | TCGGTCCACATCCATCCATCCA | 181bp |
|  | Reverse | AGGCGTCACCAGCGGATTCA |  |
| TraesCS2A01G310300, TaBEI1 | Forward | CCACCAATTCCTCACGGCTCAC | 105bp |
|  | Reverse | TCCTGGAGTCTGCACGGAGTAC |  |
| TraesCS6B01G015100, TaSWEET14-1 | Forward | GGTTCGCCTTCGGTGTGATTCA | 206bp |
|  | Reverse | ATCCTTCGTTGGTGGTGGCATC |  |
| TraesCS6D01G012100, TaSWEET14-2 | Forward | CGCTGCTCAAGTCTGACGAGTG | 135bp |
|  | Reverse | GGAGGAGGATCTTCGCCGTGAA |  |
| TraesCS2A01G293400, TaSBEIa-1 | Forward | TGGTTTCGATGGCACTGATACA | 281bp |
|  | Reverse | GCATCAAGTAAACTACCGCGTC |  |
| TraesCS7A01G549100, TaSBEIa-2 | Forward | GCACTCCGAGATGCAACTGACA | 115bp |
|  | Reverse | GGTCGGTCTTCGCTTGCTTCTT |  |
| TraesCS7A01G549300, TaSBEIb | Forward | GCCAGTTGATGAAGGTGGAGTA | 278bp |
|  | Reverse | GTGGGCTTGTCTCTAATTGGGA |  |
| TraesCS1A01G247100, TaISA2 | Forward | CTATCTGAACCACGATGGGAGG | 135bp |
|  | Reverse | TGCTGGTTCCTCATTTGCATTG |  |
| TraesCS2B01G194200, TaSuSy2 | Forward | GCCTTACCTGGAAGCTTACACT | 84bp |
|  | Reverse | TTTCCAATGATCAGGTCAGGGG |  |
| TraesCS7D01G159800, TaSuSyl | Forward | GGATCCAAGCTACTGGGACAAA | 131bp |
|  | Reverse | TGCTCACATACTTCCAGAACCC |  |
