## Supplementary material for "Genome-wide analysis reveals the genetic basis of key agronomic traits and modern wheat breeding in Henan Province": Figure S

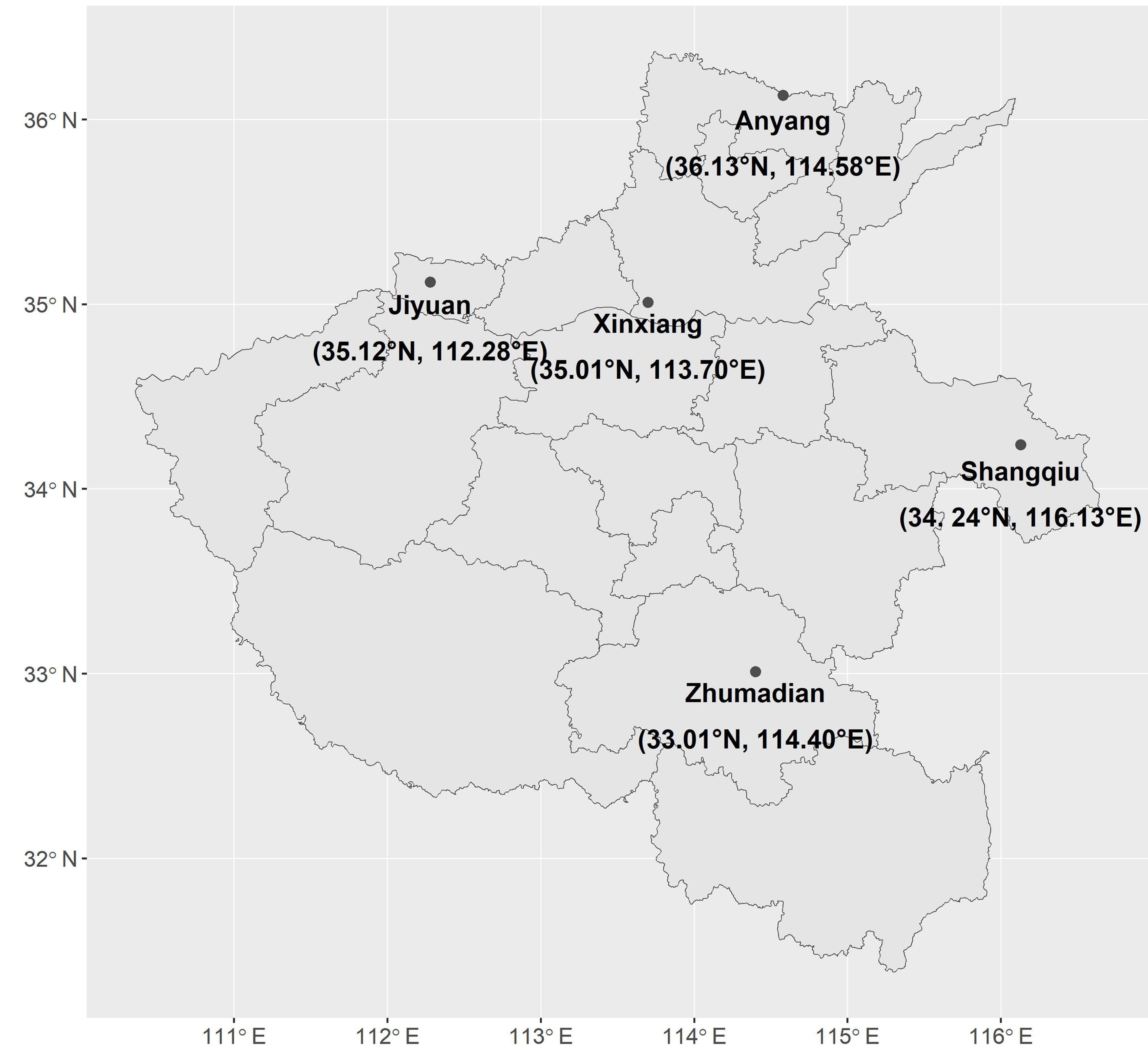

36° N
35° N
34° N
33° N
32° N
111° E
112° E
113° E
114° E
115° E
116° E
Anyang
(36.13°N, 114.58°E)
Jiyuan
(35.12°N, 112.28°E)
Xinxiang
(35.01°N, 113.70°E)
Shangqiu
(34. 24°N, 116.13°E)
Zhumadian
(33.01°N, 114.40°E)

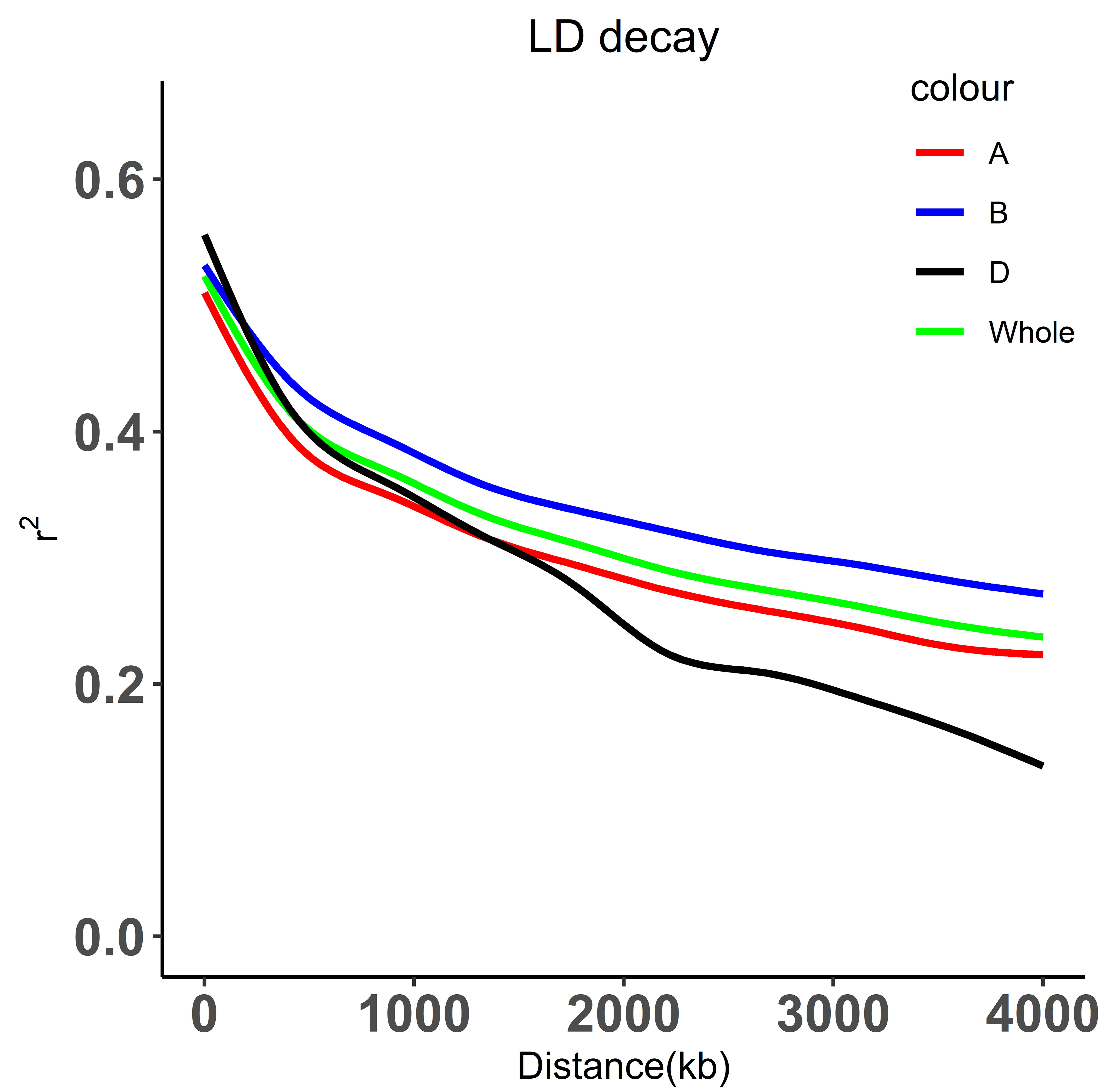

LD decay
colour
A
B
D
Whole
0.6
0.4
0.2
0.0
$r^2$
0
1000
2000
3000
4000
Distance(kb)

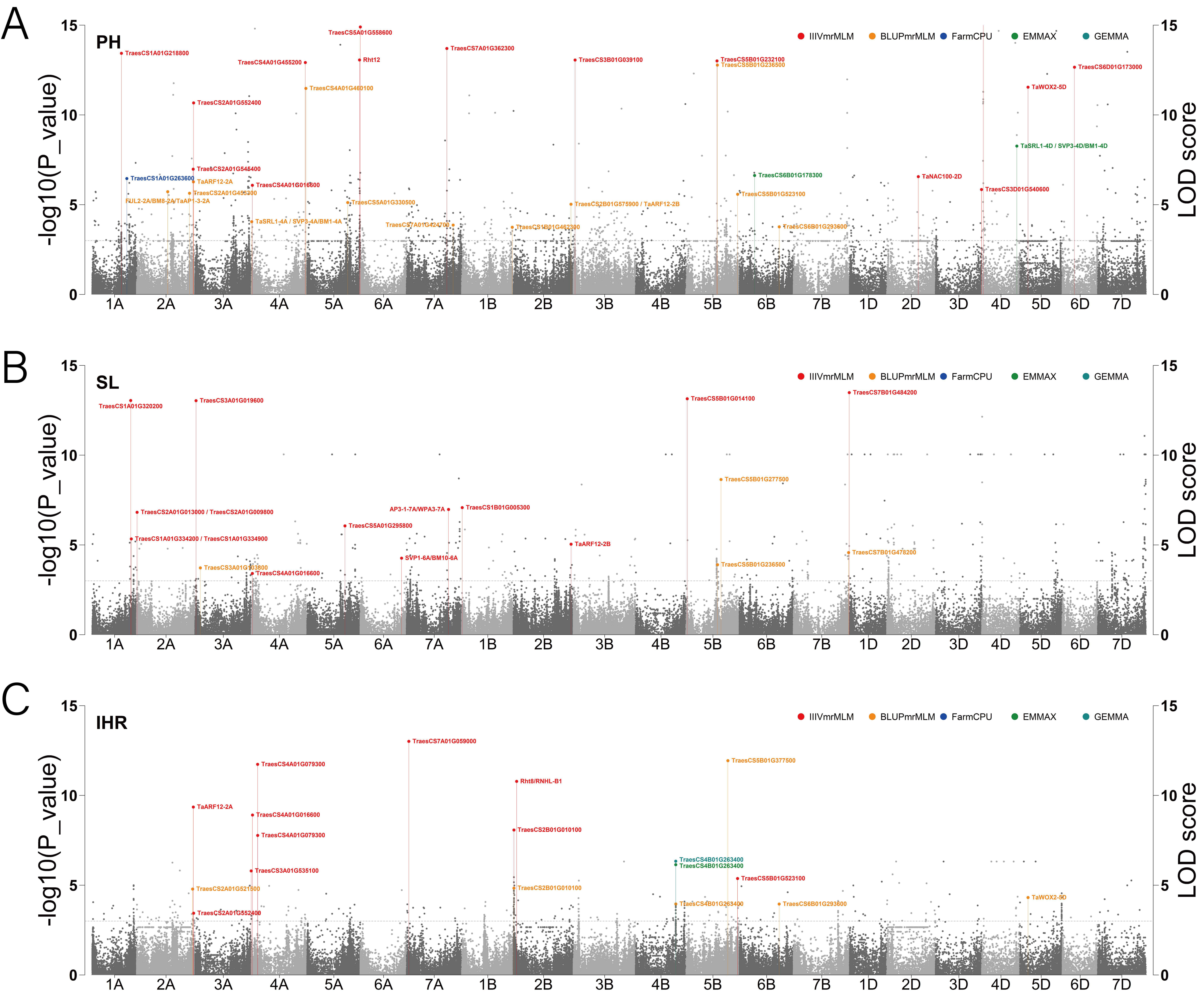
A
PH
IIIVmrMLM
BLUPmrMLM
FarmCPU
EMMAX
GEMMA
-log10(P_value)
LOD score
TraesCS5A01G558600
TraesCS1A01G218800
TraesCS7A01G362300
TraesCS4A01G455200
Rht12
TraesCS3B01G039100
TraesCS5B01G232100
TraesCS5B01G236500
TraesCS6D01G173000
TraesCS4A01G460100
TaWOX2-5D
TraesCS2A01G552400
TaSRL1-4D / SVP3-4D/BM1-4D
TraesCS2A01G545400
TraesCS1A01G263600
TraesCS6B01G178300
TaNAC100-2D
TaARF12-2A
TraesCS4A01G016600
TraesCS2A01G459300
TraesCS3D01G540600
FUL2-2A/BM8-2A/TaAP1-3-2A
TraesCS5B01G523100
TraesCS5A01G330500
TraesCS2B01G575900 / TaARF12-2B
TaSRL1-4A / SVP3-4A/BM1-4A
TraesCS7A01G424700
TraesCS1B01G462300
TraesCS6B01G293600
B
SL
TraesCS1A01G320200
TraesCS3A01G019600
TraesCS5B01G014100
TraesCS7B01G484200
TraesCS5B01G277500
TraesCS2A01G013000 / TraesCS2A01G009800
AP3-1-7A/WPA3-7A
TraesCS1B01G005300
TraesCS5A01G295800
TraesCS1A01G334200 / TraesCS1A01G334900
TaARF12-2B
TraesCS7B01G478200
SVP1-6A/BM10-6A
TraesCS5B01G236500
TraesCS3A01G103600
TraesCS4A01G016600
C
IHR
TraesCS7A01G059000
TraesCS5B01G377500
TraesCS4A01G079300
Rht8/RNHL-B1
TaARF12-2A
TraesCS4A01G016600
TraesCS2B01G010100
TraesCS4A01G079300
TraesCS4B01G263400
TraesCS4B01G263400
TraesCS3A01G535100
TraesCS5B01G523100
TraesCS2A01G521600
TraesCS2B01G010100
TaWOX2-5D
TraesCS2A01G552400
TraesCS4B01G263400
TraesCS6B01G293600
1A
2A
3A
4A
5A
6A
7A
1B
2B
3B
4B
5B
6B
7B
1D
2D
3D
4D
5D
6D
7D

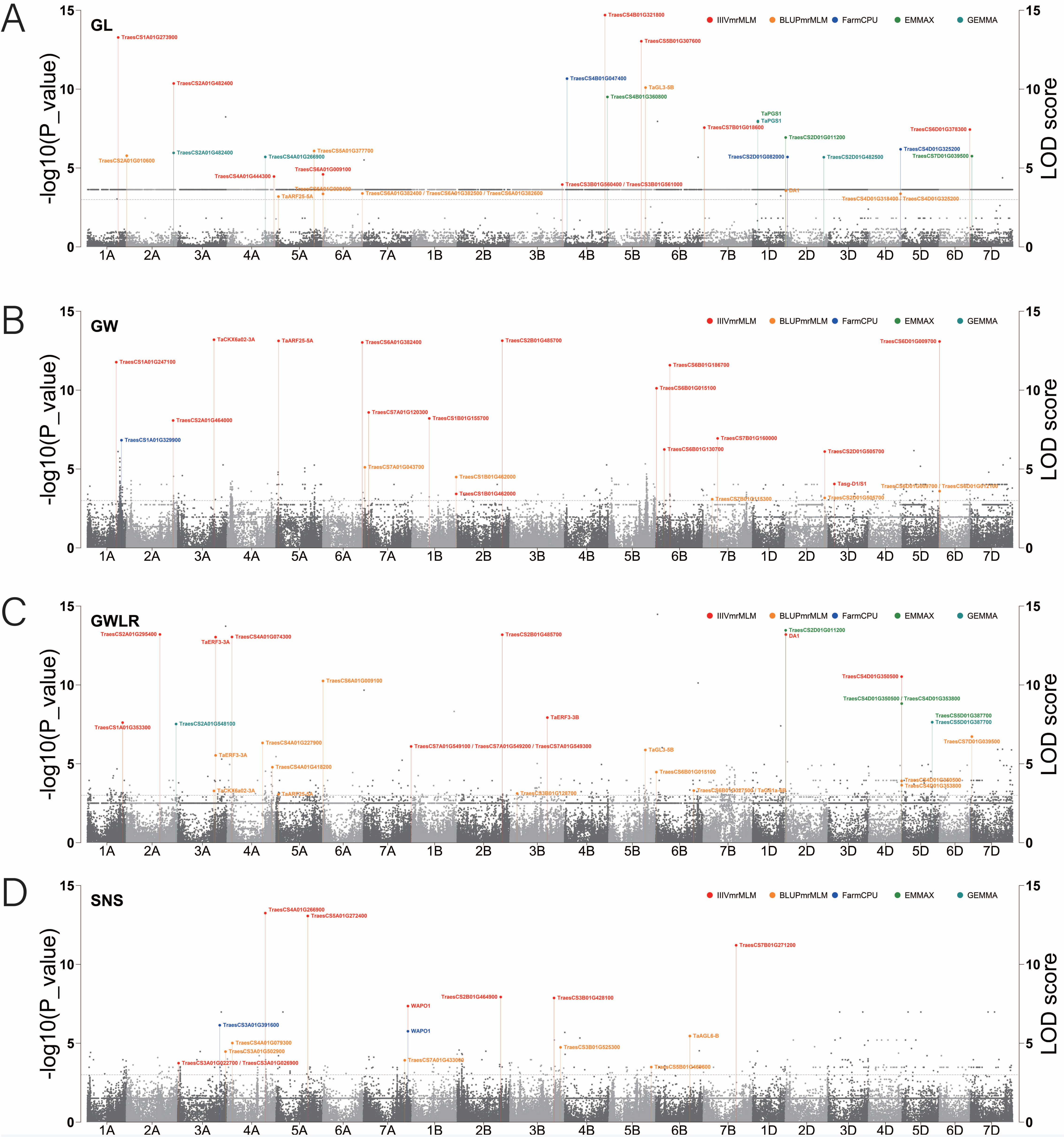

A
GL
B
GW
C
GWLR
D
SNS
IIIVmrMLM
BLUPmrMLM
FarmCPU
EMMAX
GEMMA
-log10(P_value)
LOD score
1A
2A
3A
4A
5A
6A
7A
1B
2B
3B
4B
5B
6B
7B
1D
2D
3D
4D
5D
6D
7D
TraesCS1A01G273900
TraesCS4B01G321800
TraesCS5B01G307600
TraesCS2A01G482400
TraesCS4B01G047400
TaGL3-5B
TraesCS4B01G360800
TaPGS1
TraesCS7B01G018600
TraesCS6D01G378300
TraesCS2D01G011200
TraesCS2A01G010600
TraesCS5A01G377700
TraesCS4A01G266900
TraesCS2D01G082000
TraesCS2D01G482500
TraesCS4D01G325200
TraesCS7D01G039500
TraesCS6A01G009100
TraesCS4A01G444300
TraesCS3B01G560400 / TraesCS3B01G561000
TaARF25-5A
TraesCS6A01G382400 / TraesCS6A01G382500 / TraesCS6A01G382600
DA1
TraesCS4D01G318400 / TraesCS4D01G325200
TaCKX6a02-3A
TraesCS6A01G382400
TraesCS2B01G485700
TraesCS6D01G009700
TraesCS1A01G247100
TraesCS6B01G186700
TraesCS6B01G015100
TraesCS7A01G120300
TraesCS1B01G155700
TraesCS2A01G464000
TraesCS1A01G329900
TraesCS7B01G160000
TraesCS6B01G130700
TraesCS2D01G505700
TraesCS7A01G043700
TraesCS1B01G462000
Tasg-D1/S1
TraesCS2A01G295400
TaERF3-3A
TraesCS4A01G074300
DA1
TraesCS4D01G350500
TraesCS4D01G350500 / TraesCS4D01G353800
TraesCS1A01G353300
TraesCS2A01G548100
TaERF3-3B
TraesCS5D01G387700
TraesCS4A01G227900
TraesCS7A01G549100 / TraesCS7A01G549200 / TraesCS7A01G549300
TraesCS7D01G039500
TaGL3-5B
TraesCS4A01G418200
TraesCS6B01G015100
TraesCS4D01G353800
TraesCS4A01G266900
TraesCS5A01G272400
TraesCS7B01G271200
TraesCS2B01G464900
TraesCS3B01G428100
WAPO1
TraesCS3A01G391600
TraesCS4A01G079300
TraesCS3A01G502900
TaAGL6-B
TraesCS3B01G525300
TraesCS3A01G022700 / TraesCS3A01G026900
TraesCS7A01G433000

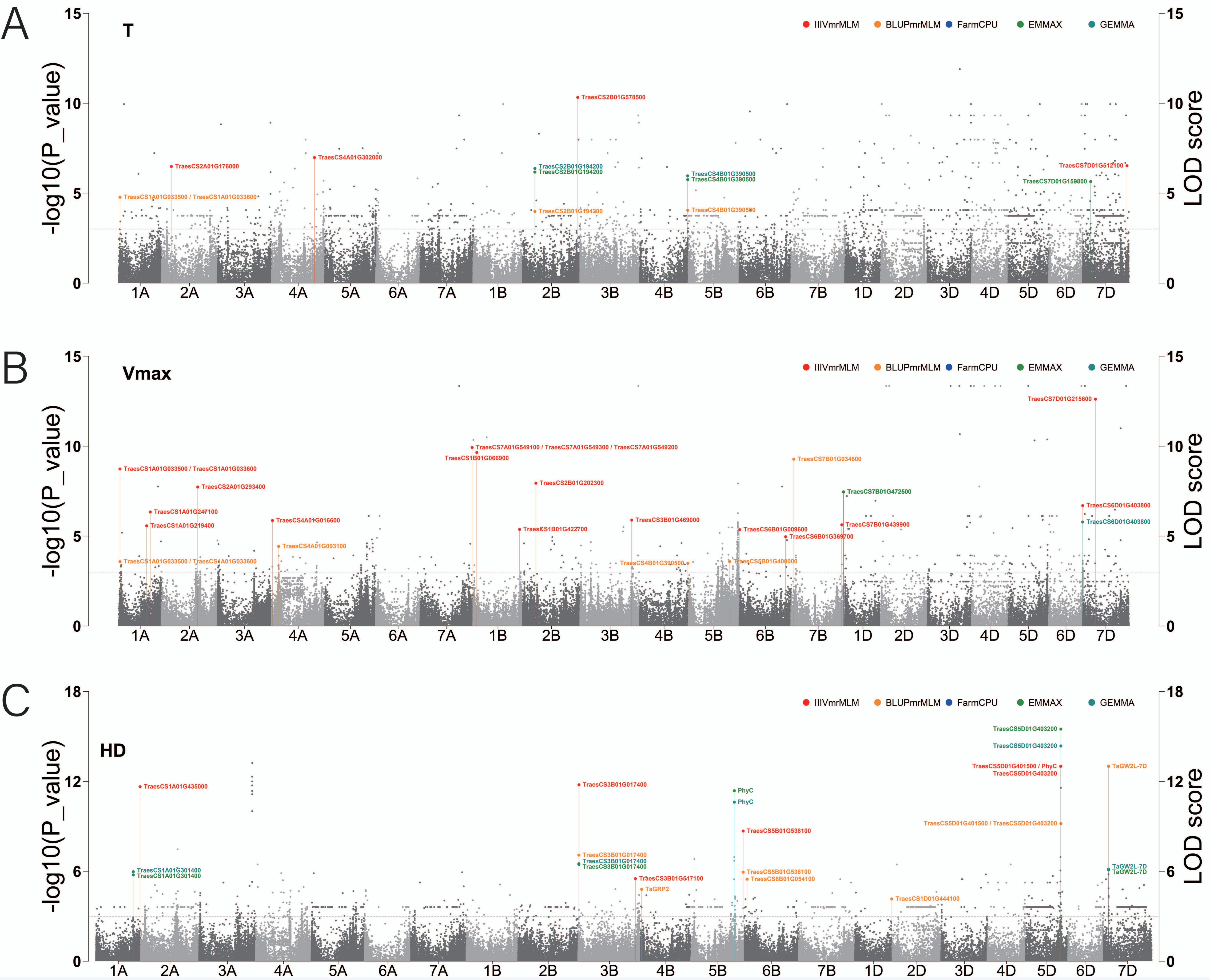
A
T
IIIVmrMLM
BLUPmrMLM
FarmCPU
EMMAX
GEMMA
-log10(P_value)
LOD score
TraesCS1A01G033500 / TraesCS1A01G033600
TraesCS2A01G176000
TraesCS4A01G302000
TraesCS2B01G194200
TraesCS2B01G194200
TraesCS2B01G194200
TraesCS2B01G578500
TraesCS4B01G390500
TraesCS4B01G390500
TraesCS4B01G390500
TraesCS7D01G512100
TraesCS7D01G159800
1A
2A
3A
4A
5A
6A
7A
1B
2B
3B
4B
5B
6B
7B
1D
2D
3D
4D
5D
6D
7D
B
Vmax
TraesCS1A01G033500 / TraesCS1A01G033600
TraesCS1A01G247100
TraesCS1A01G219400
TraesCS2A01G293400
TraesCS1A01G033500 / TraesCS1A01G033600
TraesCS4A01G016600
TraesCS4A01G093100
TraesCS7A01G549100 / TraesCS7A01G549300 / TraesCS7A01G549200
TraesCS1B01G066900
TraesCS2B01G202300
TraesCS1B01G422700
TraesCS3B01G469000
TraesCS4B01G390500
TraesCS5B01G400000
TraesCS6B01G009600
TraesCS6B01G369700
TraesCS7B01G034600
TraesCS7B01G472500
TraesCS7B01G439900
TraesCS7D01G215600
TraesCS6D01G403800
TraesCS6D01G403800
C
HD
TraesCS1A01G435000
TraesCS1A01G301400
TraesCS1A01G301400
TraesCS3B01G017400
TraesCS3B01G017400
TraesCS3B01G017400
TraesCS3B01G017400
TraesCS3B01G517100
TaGRP2
PhyC
PhyC
TraesCS5B01G538100
TraesCS5B01G538100
TraesCS6B01G054100
TraesCS1D01G444100
TraesCS5D01G403200
TraesCS5D01G403200
TraesCS5D01G401500 / PhyC
TraesCS5D01G403200
TraesCS5D01G401500 / TraesCS5D01G403200
TaGW2L-7D
TaGW2L-7D
TaGW2L-7D

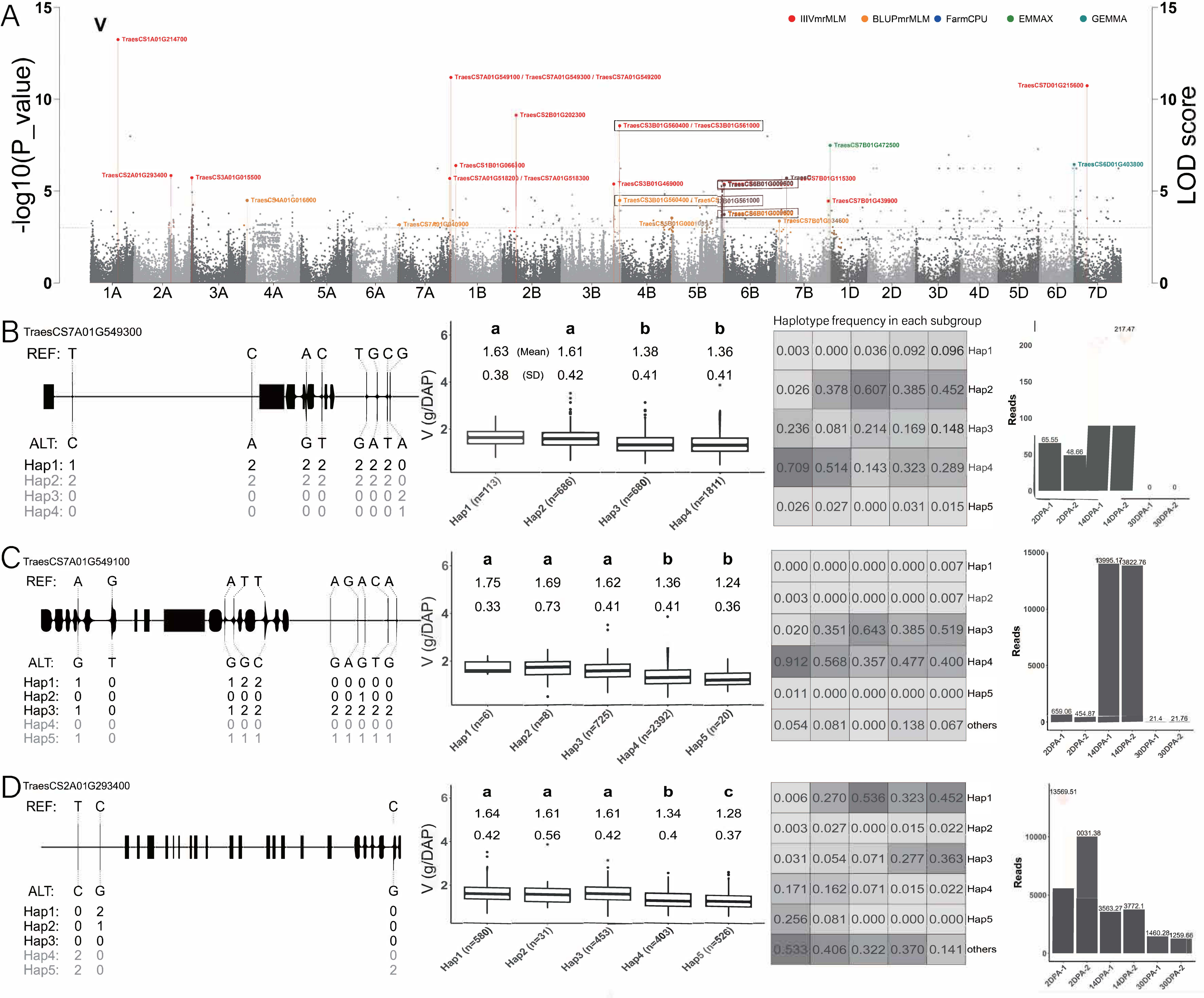
A
-log10(P_value)
LOD score
IIIVmrMLM
BLUPmrMLM
FarmCPU
EMMAX
GEMMA
TraesCS1A01G214700
TraesCS7A01G549100 / TraesCS7A01G549300 / TraesCS7A01G549200
TraesCS7D01G215600
TraesCS2B01G202300
TraesCS3B01G560400 / TraesCS3B01G561000
TraesCS7B01G472500
TraesCS6D01G403800
TraesCS2A01G293400
TraesCS3A01G015500
TraesCS1B01G066100
TraesCS7A01G518200 / TraesCS7A01G518300
TraesCS3B01G469000
TraesCS6B01G009600
TraesCS7B01G115300
TraesCS4A01G016600
TraesCS7B01G439900
TraesCS7B01G134600
1A 2A 3A 4A 5A 6A 7A 1B 2B 3B 4B 5B 6B 7B 1D 2D 3D 4D 5D 6D 7D
B TraesCS7A01G549300
REF: T C A C T G C G
ALT: C A G T G A T A
Hap1: 1 2 2 2 2 2 2 0
Hap2: 2 2 2 2 2 2 2 0
Hap3: 0 0 0 0 0 0 0 2
Hap4: 0 0 0 0 0 0 0 1
V (g/DAP)
a a b b
1.63 (Mean) 1.61 1.38 1.36
0.38 (SD) 0.42 0.41 0.41
Hap1 (n=113) Hap2 (n=686) Hap3 (n=680) Hap4 (n=1811)
Haplotype frequency in each subgroup
0.003 0.000 0.036 0.092 0.096 Hap1
0.026 0.378 0.607 0.385 0.452 Hap2
0.236 0.081 0.214 0.169 0.148 Hap3
0.709 0.514 0.143 0.323 0.289 Hap4
0.026 0.027 0.000 0.031 0.015 Hap5
Reads
217.47
65.55
48.66
0
0
2DPA-1 2DPA-2 14DPA-1 14DPA-2 30DPA-1 30DPA-2
C TraesCS7A01G549100
REF: A G A T T A G A C A
ALT: G T G G C G A G T G
Hap1: 1 0 1 2 2 0 0 0 0 0
Hap2: 0 0 0 0 0 0 0 1 0 0
Hap3: 1 0 1 2 2 2 2 2 2 2
Hap4: 0 0 0 0 0 0 0 0 0 0
Hap5: 1 0 1 1 1 1 1 1 1 1
V (g/DAP)
a a a b b
1.75 1.69 1.62 1.36 1.24
0.33 0.73 0.41 0.41 0.36
Hap1 (n=6) Hap2 (n=8) Hap3 (n=725) Hap4 (n=2392) Hap5 (n=20)
0.000 0.000 0.000 0.000 0.007 Hap1
0.003 0.000 0.000 0.000 0.007 Hap2
0.020 0.351 0.643 0.385 0.519 Hap3
0.912 0.568 0.357 0.477 0.400 Hap4
0.011 0.000 0.000 0.000 0.000 Hap5
0.054 0.081 0.000 0.138 0.067 others
Reads
13995.17
13822.76
659.06
454.87
21.4
21.76
2DPA-1 2DPA-2 14DPA-1 14DPA-2 30DPA-1 30DPA-2
D TraesCS2A01G293400
REF: T C C
ALT: C G G
Hap1: 0 2 0
Hap2: 0 1 0
Hap3: 0 0 0
Hap4: 2 0 0
Hap5: 2 0 2
V (g/DAP)
a a a b c
1.64 1.61 1.61 1.34 1.28
0.42 0.56 0.42 0.4 0.37
Hap1 (n=580) Hap2 (n=31) Hap3 (n=453) Hap4 (n=403) Hap5 (n=526)
0.006 0.270 0.536 0.323 0.452 Hap1
0.003 0.027 0.000 0.015 0.022 Hap2
0.031 0.054 0.071 0.277 0.363 Hap3
0.171 0.162 0.071 0.015 0.022 Hap4
0.256 0.081 0.000 0.000 0.000 Hap5
0.533 0.406 0.322 0.370 0.141 others
Reads
13569.51
0031.38
3563.27
3772.1
1460.28
1259.66
2DPA-1 2DPA-2 14DPA-1 14DPA-2 30DPA-1 30DPA-2

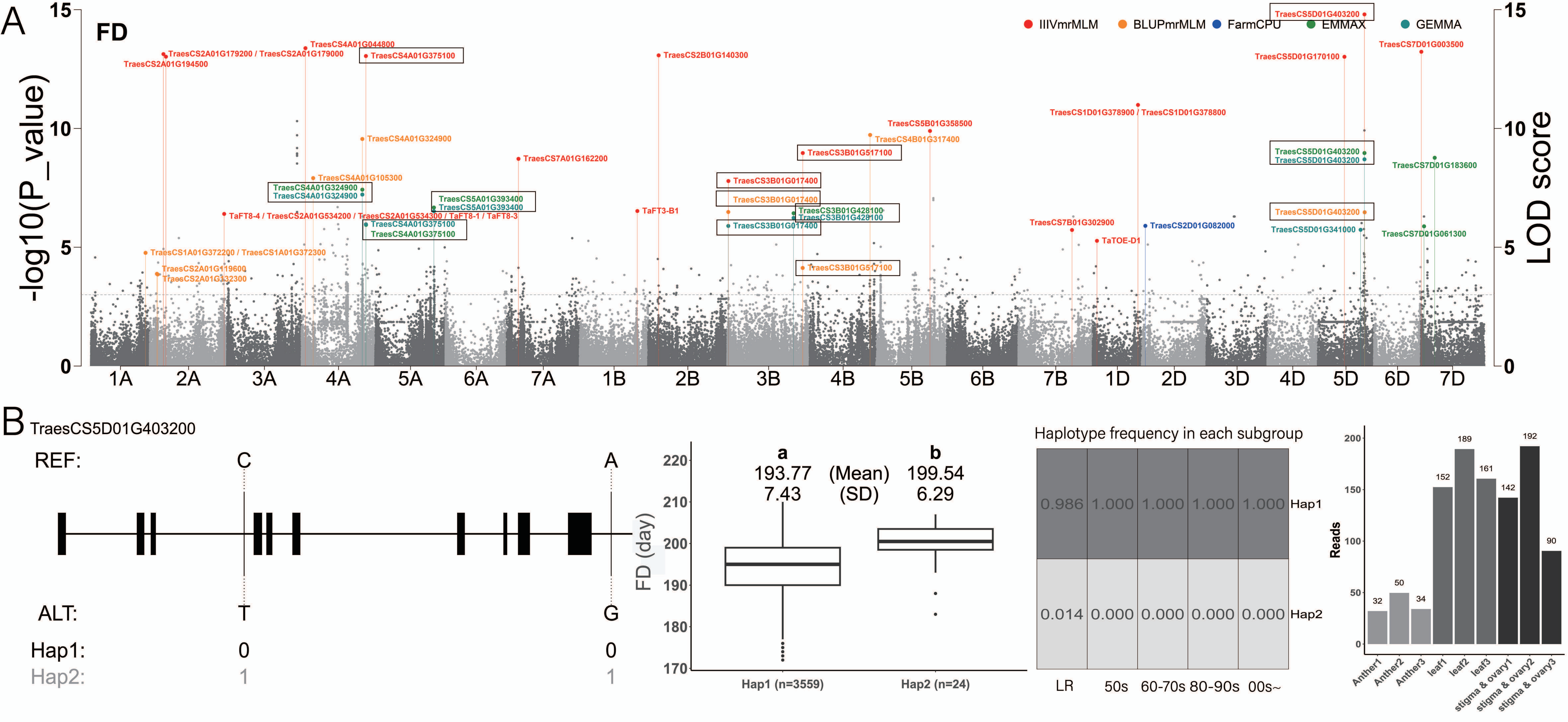

A
FD
IIIVmrMLM
BLUPmrMLM
FarmCPU
EMMAX
GEMMA
-log10(P_value)
LOD score
TraesCS5D01G403200
TraesCS4A01G044800
TraesCS2A01G179200 / TraesCS2A01G179000
TraesCS2A01G194500
TraesCS4A01G375100
TraesCS2B01G140300
TraesCS7D01G003500
TraesCS5D01G170100
TraesCS1D01G378900 / TraesCS1D01G378800
TraesCS5B01G358500
TraesCS4B01G317400
TraesCS4A01G324900
TraesCS3B01G517100
TraesCS7A01G162200
TraesCS5D01G403200
TraesCS5D01G403200
TraesCS7D01G183600
TraesCS4A01G105300
TraesCS4A01G324900
TraesCS4A01G324900
TraesCS3B01G017400
TraesCS3B01G017400
TraesCS5A01G393400
TraesCS5A01G393400
TaFT3-B1
TraesCS3B01G428100
TraesCS3B01G428100
TraesCS5D01G403200
TaFT8-4 / TraesCS2A01G534200 / TraesCS2A01G534300 / TaFT8-1 / TaFT8-3
TraesCS4A01G375100
TraesCS4A01G375100
TraesCS7B01G302900
TraesCS3B01G017400
TraesCS2D01G082000
TraesCS5D01G341000
TraesCS7D01G061300
TaTOE-D1
TraesCS1A01G372200 / TraesCS1A01G372300
TraesCS2A01G119600
TraesCS2A01G132300
TraesCS3B01G517100
1A 2A 3A 4A 5A 6A 7A 1B 2B 3B 4B 5B 6B 7B 1D 2D 3D 4D 5D 6D 7D
B
TraesCS5D01G403200
REF:
C
A
ALT:
T
G
Hap1:
0
0
Hap2:
1
1
FD (day)
a
b
193.77
(Mean)
199.54
7.43
(SD)
6.29
Hap1 (n=3559)
Hap2 (n=24)
Haplotype frequency in each subgroup
0.986 1.000 1.000 1.000 1.000 Hap1
0.014 0.000 0.000 0.000 0.000 Hap2
LR 50s 60-70s 80-90s 00s~
Reads
32 50 34 152 189 161 142 192 90
Anther1
Anther2
Anther3
leaf1
leaf2
leaf3
stigma & ovary1
stigma & ovary2
stigma & ovary3

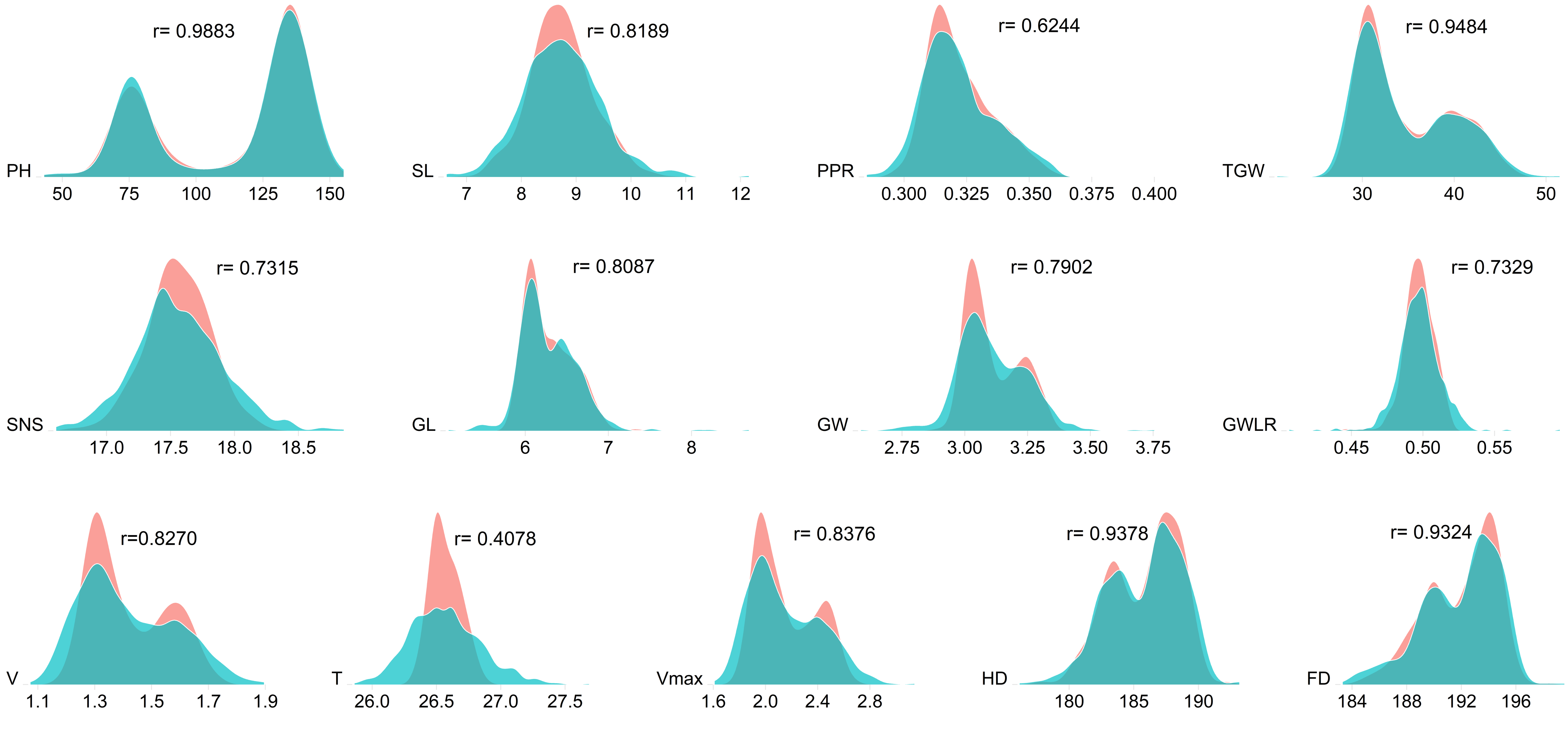

r= 0.9883
PH
50 75 100 125 150
r= 0.8189
SL
7 8 9 10 11 12
r= 0.6244
PPR
0.300 0.325 0.350 0.375 0.400
r= 0.9484
TGW
30 40 50
r= 0.7315
SNS
17.0 17.5 18.0 18.5
r= 0.8087
GL
6 7 8
r= 0.7902
GW
2.75 3.00 3.25 3.50 3.75
r= 0.7329
GWLR
0.45 0.50 0.55
r=0.8270
V
1.1 1.3 1.5 1.7 1.9
r= 0.4078
T
26.0 26.5 27.0 27.5
r= 0.8376
Vmax
1.6 2.0 2.4 2.8
r= 0.9378
HD
180 185 190
r= 0.9324
FD
184 188 192 196

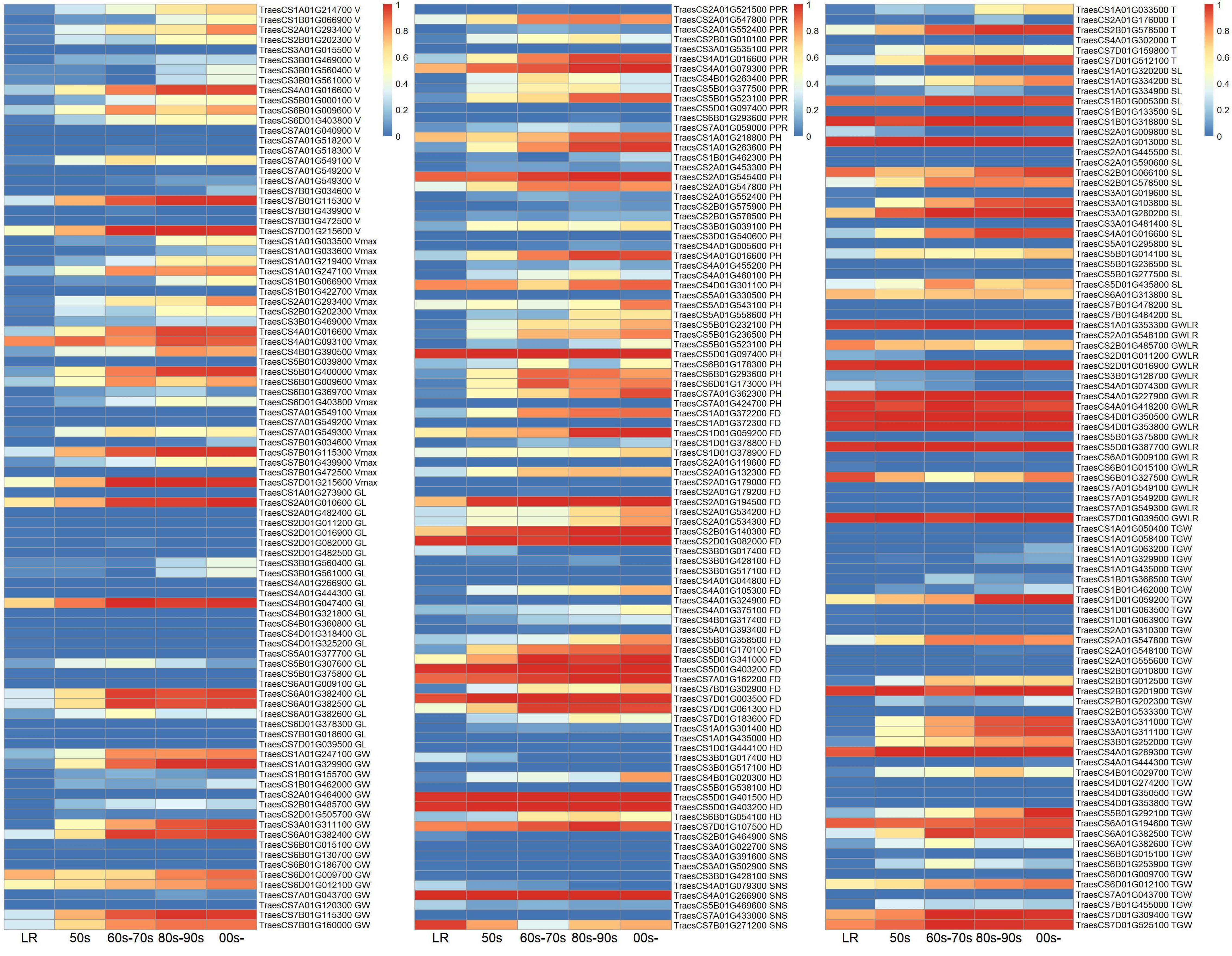
TraesCS1A01G214700 V
TraesCS1B01G066900 V
TraesCS2A01G293400 V
TraesCS2B01G202300 V
TraesCS3A01G015500 V
TraesCS3B01G469000 V
TraesCS3B01G560400 V
TraesCS3B01G561000 V
TraesCS4A01G016600 V
TraesCS5B01G000100 V
TraesCS6B01G009600 V
TraesCS6D01G403800 V
TraesCS7A01G040900 V
TraesCS7A01G518200 V
TraesCS7A01G518300 V
TraesCS7A01G549100 V
TraesCS7A01G549200 V
TraesCS7A01G549300 V
TraesCS7B01G034600 V
TraesCS7B01G115300 V
TraesCS7B01G439900 V
TraesCS7B01G472500 V
TraesCS7D01G215600 V
TraesCS1A01G033500 Vmax
TraesCS1A01G033600 Vmax
TraesCS1A01G219400 Vmax
TraesCS1A01G247100 Vmax
TraesCS1B01G066900 Vmax
TraesCS1B01G422700 Vmax
TraesCS2A01G293400 Vmax
TraesCS2B01G202300 Vmax
TraesCS3B01G469000 Vmax
TraesCS4A01G016600 Vmax
TraesCS4A01G093100 Vmax
TraesCS4B01G390500 Vmax
TraesCS5B01G039800 Vmax
TraesCS5B01G400000 Vmax
TraesCS6B01G009600 Vmax
TraesCS6B01G369700 Vmax
TraesCS6D01G403800 Vmax
TraesCS7A01G549100 Vmax
TraesCS7A01G549200 Vmax
TraesCS7A01G549300 Vmax
TraesCS7B01G034600 Vmax
TraesCS7B01G115300 Vmax
TraesCS7B01G439900 Vmax
TraesCS7B01G472500 Vmax
TraesCS7D01G215600 Vmax
TraesCS1A01G273900 GL
TraesCS2A01G010600 GL
TraesCS2A01G482400 GL
TraesCS2D01G011200 GL
TraesCS2D01G016900 GL
TraesCS2D01G082000 GL
TraesCS2D01G482500 GL
TraesCS3B01G560400 GL
TraesCS3B01G561000 GL
TraesCS4A01G266900 GL
TraesCS4A01G444300 GL
TraesCS4B01G047400 GL
TraesCS4B01G321800 GL
TraesCS4B01G360800 GL
TraesCS4D01G318400 GL
TraesCS4D01G325200 GL
TraesCS5A01G377700 GL
TraesCS5B01G307600 GL
TraesCS5B01G375800 GL
TraesCS6A01G009100 GL
TraesCS6A01G382400 GL
TraesCS6A01G382500 GL
TraesCS6A01G382600 GL
TraesCS6D01G378300 GL
TraesCS7B01G018600 GL
TraesCS7D01G039500 GL
TraesCS1A01G247100 GW
TraesCS1A01G329900 GW
TraesCS1B01G155700 GW
TraesCS1B01G462000 GW
TraesCS2A01G464000 GW
TraesCS2B01G485700 GW
TraesCS2D01G505700 GW
TraesCS3A01G311100 GW
TraesCS6A01G382400 GW
TraesCS6B01G015100 GW
TraesCS6B01G130700 GW
TraesCS6B01G186700 GW
TraesCS6D01G009700 GW
TraesCS6D01G012100 GW
TraesCS7A01G043700 GW
TraesCS7A01G120300 GW
TraesCS7B01G115300 GW
TraesCS7B01G160000 GW
LR
50s
60s-70s
80s-90s
00s-
1
0.8
0.6
0.4
0.2
0
TraesCS2A01G521500 PPR
TraesCS2A01G547800 PPR
TraesCS2A01G552400 PPR
TraesCS2B01G010100 PPR
TraesCS3A01G535100 PPR
TraesCS4A01G016600 PPR
TraesCS4A01G079300 PPR
TraesCS4B01G263400 PPR
TraesCS5B01G377500 PPR
TraesCS5B01G523100 PPR
TraesCS5D01G097400 PPR
TraesCS6B01G293600 PPR
TraesCS7A01G059000 PPR
TraesCS1A01G218800 PH
TraesCS1A01G263600 PH
TraesCS1B01G462300 PH
TraesCS2A01G453300 PH
TraesCS2A01G545400 PH
TraesCS2A01G547800 PH
TraesCS2A01G552400 PH
TraesCS2B01G575900 PH
TraesCS2B01G578500 PH
TraesCS3B01G039100 PH
TraesCS3D01G540600 PH
TraesCS4A01G005600 PH
TraesCS4A01G016600 PH
TraesCS4A01G455200 PH
TraesCS4A01G460100 PH
TraesCS4D01G301100 PH
TraesCS5A01G330500 PH
TraesCS5A01G543100 PH
TraesCS5A01G558600 PH
TraesCS5B01G232100 PH
TraesCS5B01G236500 PH
TraesCS5B01G523100 PH
TraesCS5D01G097400 PH
TraesCS6B01G178300 PH
TraesCS6B01G293600 PH
TraesCS6D01G173000 PH
TraesCS7A01G362300 PH
TraesCS7A01G424700 PH
TraesCS1A01G372200 FD
TraesCS1A01G372300 FD
TraesCS1D01G059200 FD
TraesCS1D01G378800 FD
TraesCS1D01G378900 FD
TraesCS2A01G119600 FD
TraesCS2A01G132300 FD
TraesCS2A01G179000 FD
TraesCS2A01G179200 FD
TraesCS2A01G194500 FD
TraesCS2A01G534200 FD
TraesCS2A01G534300 FD
TraesCS2B01G140300 FD
TraesCS2D01G082000 FD
TraesCS3B01G017400 FD
TraesCS3B01G428100 FD
TraesCS3B01G517100 FD
TraesCS4A01G044800 FD
TraesCS4A01G105300 FD
TraesCS4A01G324900 FD
TraesCS4A01G375100 FD
TraesCS4B01G317400 FD
TraesCS5A01G393400 FD
TraesCS5B01G358500 FD
TraesCS5D01G170100 FD
TraesCS5D01G341000 FD
TraesCS5D01G403200 FD
TraesCS7A01G162200 FD
TraesCS7B01G302900 FD
TraesCS7D01G003500 FD
TraesCS7D01G061300 FD
TraesCS7D01G183600 FD
TraesCS1A01G301400 HD
TraesCS1A01G435000 HD
TraesCS1D01G444100 HD
TraesCS3B01G017400 HD
TraesCS3B01G517100 HD
TraesCS4B01G020300 HD
TraesCS5B01G538100 HD
TraesCS5D01G401500 HD
TraesCS5D01G403200 HD
TraesCS6B01G054100 HD
TraesCS7D01G107500 HD
TraesCS2B01G464900 SNS
TraesCS3A01G022700 SNS
TraesCS3A01G391600 SNS
TraesCS3A01G502900 SNS
TraesCS3B01G428100 SNS
TraesCS4A01G079300 SNS
TraesCS4A01G266900 SNS
TraesCS5B01G469600 SNS
TraesCS7A01G433000 SNS
TraesCS7B01G271200 SNS
LR
50s
60s-70s
80s-90s
00s-
1
0.8
0.6
0.4
0.2
0
TraesCS1A01G033500 T
TraesCS2A01G176000 T
TraesCS2B01G578500 T
TraesCS4A01G302000 T
TraesCS7D01G159800 T
TraesCS7D01G512100 T
TraesCS1A01G320200 SL
TraesCS1A01G334200 SL
TraesCS1A01G334900 SL
TraesCS1B01G005300 SL
TraesCS1B01G133500 SL
TraesCS1B01G318800 SL
TraesCS2A01G009800 SL
TraesCS2A01G013000 SL
TraesCS2A01G445500 SL
TraesCS2A01G590600 SL
TraesCS2B01G066100 SL
TraesCS2B01G578500 SL
TraesCS3A01G019600 SL
TraesCS3A01G103800 SL
TraesCS3A01G280200 SL
TraesCS3A01G481400 SL
TraesCS4A01G016600 SL
TraesCS5A01G295800 SL
TraesCS5B01G014100 SL
TraesCS5B01G236500 SL
TraesCS5B01G277500 SL
TraesCS5D01G435800 SL
TraesCS6A01G313800 SL
TraesCS7B01G478200 SL
TraesCS7B01G484200 SL
TraesCS1A01G353300 GWLR
TraesCS2A01G548100 GWLR
TraesCS2B01G485700 GWLR
TraesCS2D01G011200 GWLR
TraesCS2D01G016900 GWLR
TraesCS3B01G128700 GWLR
TraesCS4A01G074300 GWLR
TraesCS4A01G227900 GWLR
TraesCS4A01G418200 GWLR
TraesCS4D01G350500 GWLR
TraesCS4D01G353800 GWLR
TraesCS5B01G375800 GWLR
TraesCS5D01G387700 GWLR
TraesCS6A01G009100 GWLR
TraesCS6B01G015100 GWLR
TraesCS6B01G327500 GWLR
TraesCS7A01G549100 GWLR
TraesCS7A01G549200 GWLR
TraesCS7A01G549300 GWLR
TraesCS7D01G039500 GWLR
TraesCS1A01G050400 TGW
TraesCS1A01G058400 TGW
TraesCS1A01G063200 TGW
TraesCS1A01G329900 TGW
TraesCS1A01G435000 TGW
TraesCS1B01G368500 TGW
TraesCS1B01G462000 TGW
TraesCS1D01G059200 TGW
TraesCS1D01G063500 TGW
TraesCS1D01G063900 TGW
TraesCS2A01G310300 TGW
TraesCS2A01G547800 TGW
TraesCS2A01G548100 TGW
TraesCS2A01G555600 TGW
TraesCS2B01G010800 TGW
TraesCS2B01G012500 TGW
TraesCS2B01G201900 TGW
TraesCS2B01G202300 TGW
TraesCS2B01G533300 TGW
TraesCS3A01G311000 TGW
TraesCS3A01G311100 TGW
TraesCS3B01G252000 TGW
TraesCS4A01G289300 TGW
TraesCS4A01G444300 TGW
TraesCS4B01G029700 TGW
TraesCS4D01G274200 TGW
TraesCS4D01G350500 TGW
TraesCS4D01G353800 TGW
TraesCS5B01G292100 TGW
TraesCS6A01G194600 TGW
TraesCS6A01G382500 TGW
TraesCS6A01G382600 TGW
TraesCS6B01G015100 TGW
TraesCS6B01G253900 TGW
TraesCS6D01G009700 TGW
TraesCS6D01G012100 TGW
TraesCS7A01G043700 TGW
TraesCS7B01G455000 TGW
TraesCS7D01G309400 TGW
TraesCS7D01G525100 TGW
LR
50s
60s-70s
80s-90s
00s-
1
0.8
0.6
0.4
0.2
0

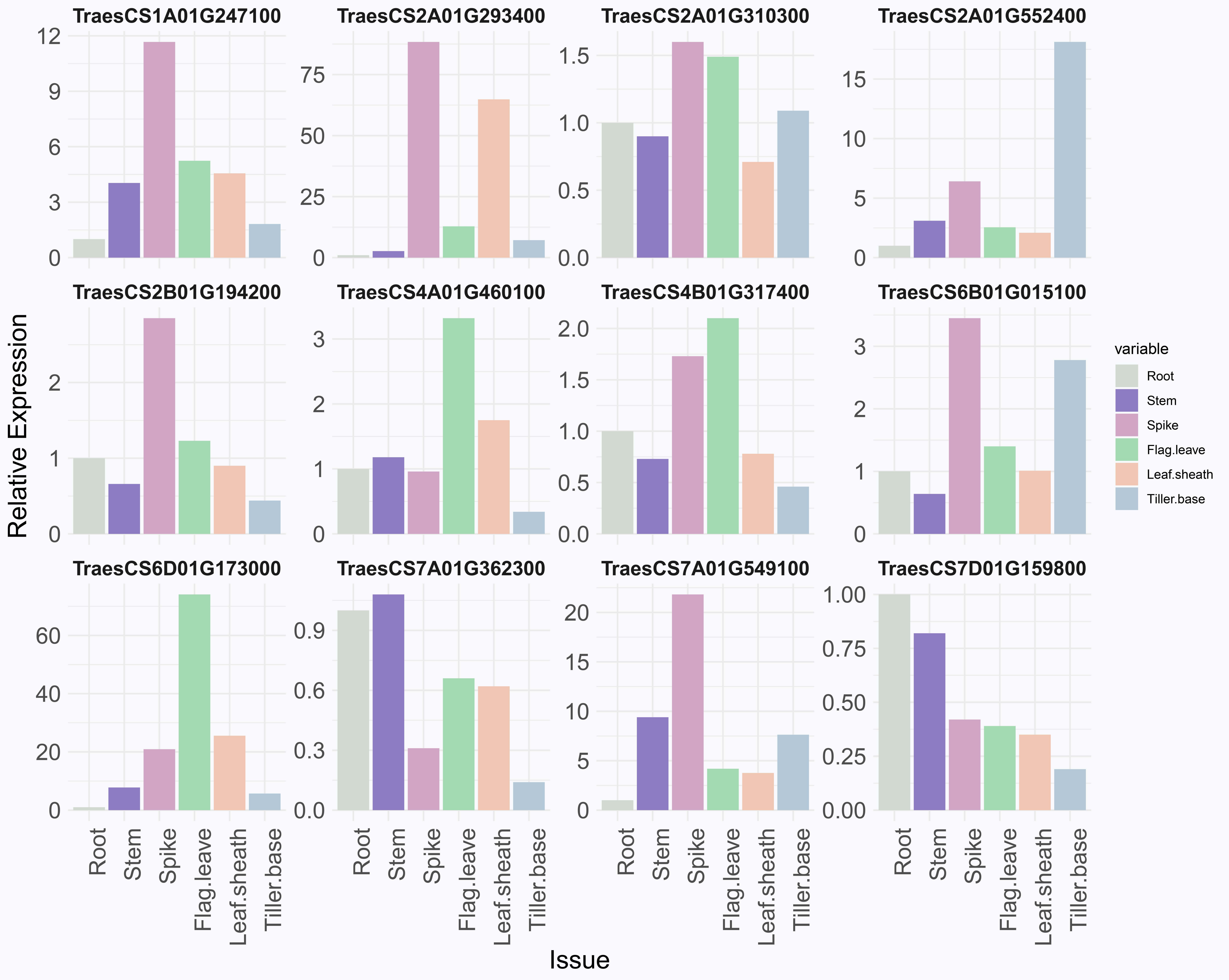

TraesCS1A01G247100
TraesCS2A01G293400
TraesCS2A01G310300
TraesCS2A01G552400
TraesCS2B01G194200
TraesCS4A01G460100
TraesCS4B01G317400
TraesCS6B01G015100
TraesCS6D01G173000
TraesCS7A01G362300
TraesCS7A01G549100
TraesCS7D01G159800
Relative Expression
Issue
Root
Stem
Spike
Flag.leave
Leaf.sheath
Tiller.base
variable

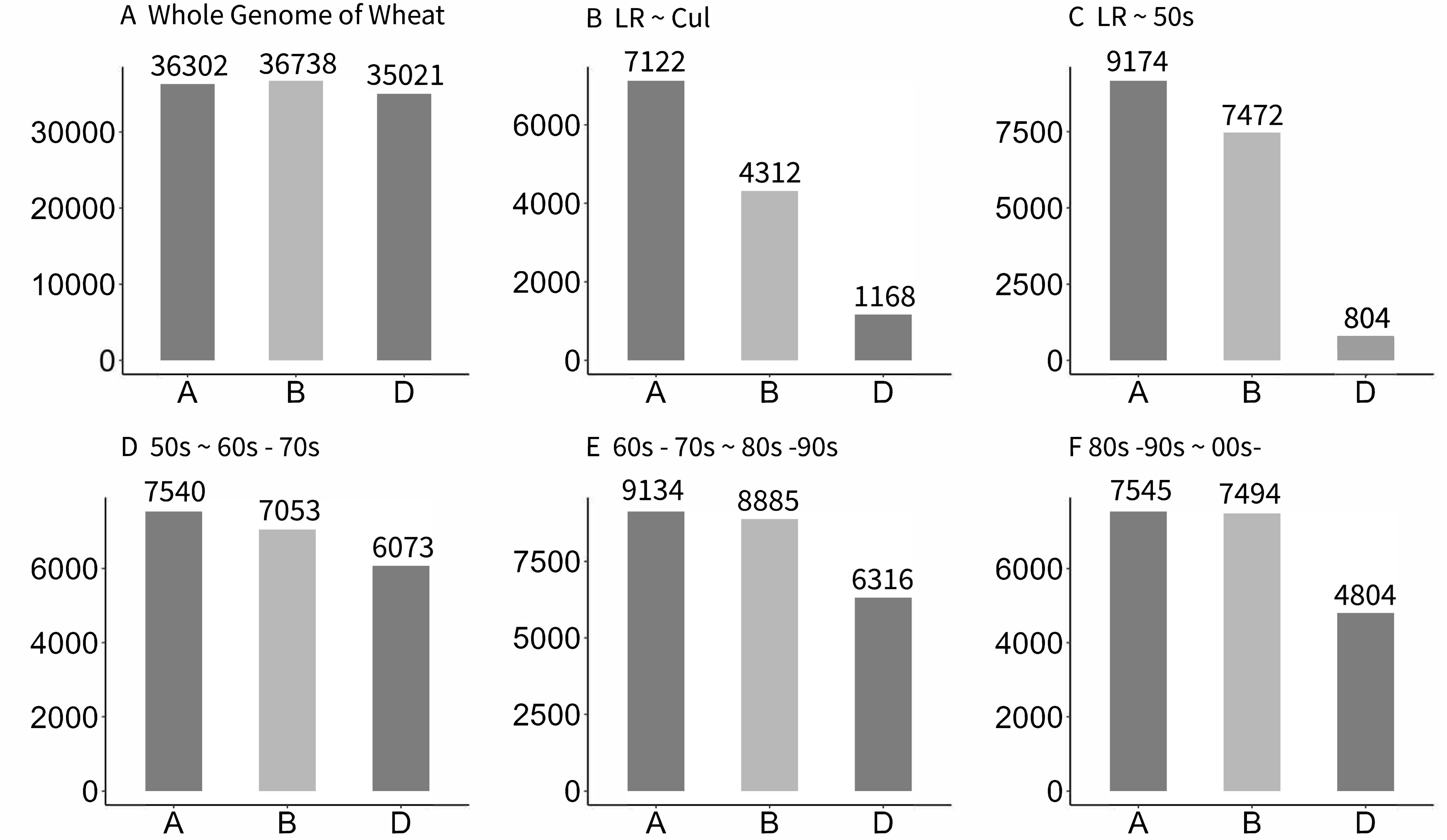
A Whole Genome of Wheat
36302
36738
35021
30000
20000
10000
0
A
B
D
B LR ~ Cul
7122
4312
1168
6000
4000
2000
0
A
B
D
C LR ~ 50s
9174
7472
804
7500
5000
2500
0
A
B
D
D 50s ~ 60s - 70s
7540
7053
6073
6000
4000
2000
0
A
B
D
E 60s - 70s ~ 80s -90s
9134
8885
6316
7500
5000
2500
0
A
B
D
F 80s -90s ~ 00s-
7545
7494
4804
6000
4000
2000
0
A
B
D
