## Supplementary material for "Genome-wide analysis reveals the genetic basis of key agronomic traits and modern wheat breeding in Henan Province": Table S

**Supplementary Table S1**. List of known genes in selection sweeps between landraces and cultivars with different periods

| Gene | Symbol | Chr | Group | Trait | Reference |
|---|---|---|---|---|---|
| TraesCS1A01G007934 | *Glu-A3* | 1A | LR~50s | low molecular weight glutenin subunit | Zhao et al., 2023 |
| TraesCS1A01G008000 | *Glu-A3* | 1A | LR~50s | low molecular weight glutenin subunit | Elhadi et al., 2021 |
| TraesCS1A01G008100 | *Pm3* | 1A | LR~50s | Powdery mildew resistance | Koller et al., 2018 |
| TraesCS1A01G010500 | *Csl-1A* | 1A | LR~Cul; LR~50s | tiller number and stem cell wall composition | Hyles et al., 2017 |
| TraesCS1A01G010900 | *Glu-A3* | 1A | LR~Cul; LR~50s | low molecular weight glutenin subunit | Zhao et al., 2023 |
| TraesCS1A01G010905 | *Glu-A3* | 1A | LR~Cul; LR~50s | low molecular weight glutenin subunit | Zhao et al., 2023 |
| TraesCS1A01G018600 | *Lr10* | 1A | LR~Cul; LR~50s; 60s-70s~80s-90s | Leaf rust resistance | Feuille et al., 1997 |
| TraesCS1A01G031600 | *Rf1* | 1A | LR~Cul; LR~50s | male sterility and fertility | Melonek et al., 2021 |
| TraesCS1A01G058400 | *TaTOE-A1* | 1A | LR~50s | flowering time | Zikhali et al., 2017 |
| TraesCS1A01G061500 | *Yr15-1A* | 1A | LR~Cul; LR~50s | Stripe rust | Klymiuk et al., 2018 |
| TraesCS1A01G220300 | *WCO-1* | 1A | LR~Cul | Flowering time | Shaw et al., 2020 |
| TraesCS1A01G255100 | *TaGID1-1A* | 1A | 60s-70s~80s-90s; 80s-90s~00s- | plant height and coleoptile growth | Li., et al 2013 |
| TraesCS1A01G264300 | *WPI-1-1A* | 1A | 60s-70s~80s-90s; 80s-90s~00s- | wheat spike and spikelet/florets development | Hama et al., 2004 |
| TraesCS1A01G314200 | *TaMFS1* | 1A | LR~50s; 60s-70s~80s-90s | spike inflorescence architecture; mild supernumerary spikelets; grain number per spike | Liu et al., 2021 |
| TraesCS1A01G317311 | *Glu-1Ax* | 1A | LR~50s; 60s-70s~80s-90s | high molecular weight glutenin subunit 1Ax (Glu-1Ax) | Shewry et al., 2002 |
| TraesCS1A01G338600 | *TaFT3-A1* | 1A | LR~50s | flowering time | Zikhali et al., 2017 |
| TraesCS1A01G353300 | *TaSS4-1A* | 1A | LR~50s; 50s~60s-70s | starch | Hawkins et al., 2021 |
| TraesCS1A01G382800 | *TaASN2-1A* | 1A | LR~Cul; LR~50s | free asparagine | Raffan et al., 2021 |
| TraesCS1A01G420000 | *TaRec8-A1* | 1A | 50s~60s-70s; 80s-90s~00s- | Chromosome cohesion and kinetochore orientation in meiosis | Ma et al., 2018 |
| TraesCS1A01G466500 | *Glu-1Ay* | 1A | LR~50s; 60s-70s~80s-90s; 80s-90s~00s- | high molecular weight glutenin subunit 1Ay (Glu-1Ay) | Shewry et al., 2002 |
| TraesCS1B01G038500 | *Rf3* | 1B | LR~Cul; LR~50s | male sterility and fertility | Melonek et al., 2021 |
| TraesCS1B01G042700 | *Glu-B3* | 1B | LR~50s; 50s~60s-70s | low molecular weight glutenin subunit | Zhao et al., 2023 |
| TraesCS1B01G144800 | *AG1-1B* | 1B | 60s-70s~80s-90s | wheat spike and spikelet/florets development | Meguro et al., 2023 |
| TraesCS1B01G326500 | *DUO-B1* | 1B | 50s~60s-70s | spike inflorescence architecture; mild supernumerary spikelets; grain number per spike | Wang et al., 2022 |
| TraesCS1B01G329711 | *Glu-1Bx* | 1B | 50s~60s-70s | high molecular weight glutenin subunit 1Bx (Glu-1Bx) | Shewry et al., 2002 |
| TraesCS1B01G329992 | *Glu-1By* | 1B | 50s~60s-70s | high molecular weight glutenin subunit 1By (Glu-1By) | Shewry et al., 2002 |

| | | | | | |
|---|---|---|---|---|---|
| TraesCS1B01G351100 | *TaFT3-B1* | 1B | 50s~60s-70s; 80s-90s~00s- | flowering time | Zikhali et al., 2017 |
| TraesCS1B01G368500 | *TaSS4-1B* | 1B | LR~50s | starch | Hawkins et al., 2021 |
| TraesCS1B01G408200 | *TaASN2-1B* | 1B | 80s-90s~00s- | free asparagine | Raffan et al., 2021 |
| TraesCS1B01G449700 | *TaAGPL1* | 1B | LR~Cul | Starch synthesis | Gu et al., 2021 |
| TraesCS1B01G450400 | *TaRec8-B1* | 1B | 50s~60s-70s; 60s-70s~80s-90s | Chromosome cohesion and kinetochore orientation in meiosis | Ma et al., 2018 |
| TraesCS1D01G000300 | *Glu-D3* | 1D | 60s-70s~80s-90s | low molecular weight glutenin subunit | Zhao et al., 2023 |
| TraesCS1D01G001200 | *LGP1* | 1D | 60s-70s~80s-90s; 80s-90s~00s- | gluten accumulation | Chen et al., 2021 |
| TraesCS1D01G005400 | *Lr21* | 1D | 60s-70s~80s-90s | Leaf rust resistance | Huang et al., 2003 |
| TraesCS1D01G007400 | *Glu-D3* | 1D | 50s~60s-70s | low molecular weight glutenin subunit | Zhao et al., 2023 |
| TraesCS1D01G008600 | *Glu-D3* | 1D | 50s~60s-70s; 60s-70s~80s-90s | low molecular weight glutenin subunit | Zhao et al., 2023 |
| TraesCS1D01G009400 | *Glu-D3* | 1D | 50s~60s-70s; 60s-70s~80s-90s | low molecular weight glutenin subunit | Zhao et al., 2023 |
| TraesCS1D01G009900 | *Glu-D3* | 1D | 50s~60s-70s | low molecular weight glutenin subunit | Zhao et al., 2023 |
| TraesCS1D01G015100 | *Glu-D3* | 1D | 60s-70s~80s-90s | low molecular weight glutenin subunit | Zhao et al., 2023 |
| TraesCS1D01G029200 | *SrTA1662* | 1D | 60s-70s~80s-90s | stem rust | Arora et al., 2019 |
| TraesCS1D01G094000 | *TaPGS1* | 1D | 60s-70s~80s-90s | Grain size; Grain weight; Carbohydrate and total protein; Grain yield | Guo et al., 2022 |
| TraesCS1D01G155200 | *TaHB-D2* | 1D | 80s-90s~00s- | inflorescence branching; grain protein content | Dixon et al., 2022 |
| TraesCS1D01G195300 | *TaPYL1-1D* | 1D | 50s~60s-70s | drought tolerance and water-use efficiency | Mao et al., 2022 |
| TraesCS1D01G254500 | *TaGID1-1D* | 1D | 60s-70s~80s-90s | plant height and coleoptile growth | Li., et al 2013 |
| TraesCS1D01G264500 | *PI1-1D* | 1D | 80s-90s~00s- | wheat spike and spikelet/florets development | Raza et al., 2022 |
| TraesCS1D01G314700 | *DUO-D1* | 1D | 50s~60s-70s; 60s-70s~80s-90s | spike inflorescence architecture; mild supernumerary spikelets; grain number per spike | Wang et al., 2022 |
| TraesCS1D01G356900 | *TaSS4-1D* | 1D | 50s~60s-70s | starch | Hawkins et al., 2021 |
| TraesCS1D01G390500 | *TaASN2-1D* | 1D | LR~Cul; LR~50s | free asparagine | Raffan et al., 2021 |
| TraesCS2A01G059900 | *Rht8* | 2A | 80s-90s~00s- | plant height | Chai et al., 2022 |
| TraesCS2A01G095700 | *TmNAS3-2A* | 2A | LR~Cul | Fe content of wheat grain | Wang et al., 2022 |
| TraesCS2A01G121000 | *TaNHX2-2A* | 2A | 60s-70s~80s-90s | salt | Singh et al., 2021 |
| TraesCS2A01G168200 | *TaSuSy2* | 2A | LR~50s | starch | Mukherjee et al., 2015 |
| TraesCS2A01G261200 | *FUL2-2A* | 2A | 60s-70s~80s-90s | wheat spike and spikelet development/flowering/plant height | Li et al., 2019 |
| TraesCS2A01G464900 | *Sr21* | 2A | LR~50s | stem rust | Chen et al., 2018 |
| TraesCS2A01G468200 | *PPO-A1* | 2A | LR~Cul; LR~50s; 60s-70s~80s-90s; 80s-90s~00s- | PPO activity/end-use quality/wheat products colour | Brian and Skinner 2011 |

| | | | | | |
|---|---|---|---|---|---|
| TraesCS2A01G468500 | *PPO-A2* | 2A | LR~Cul; LR~50s; 60s-70s~80s-90s; 80s-90s~00s- | PPO activity/end-use quality/wheat products colour | Brian and Skinner 2011 |
| TraesCS2A01G502300 | *TaSPL-A* | 2A | LR~Cul | Flag leaf angle | Niu et al., 2023 |
| TraesCS2A01G514200 | *TaAP2* | 2A | LR~Cul | Flowering time | Ning et al., 2003 |
| TraesCS2A01G555600 | *TaSTT3b-2A* | 2A | LR~Cul | sharp eyespot and increases grain weight | Zhu et al., 2022 |
| TraesCS2B01G488000 | *YrSP* | 2B | 60s-70s~80s-90s | Stripe rust | Marchal et al., 2018 |
| TraesCS2B01G034300 | *Sm1* | 2B | LR~50s | orange wheat blossom midge (OWBM) | Walkowiak et al., 2020 |
| TraesCS2B01G111100 | *TmNAS3-2B* | 2B | 60s-70s~80s-90s | Fe content of wheat grain | Wang et al., 2022 |
| TraesCS2B01G141900 | *TaNHX2-2B* | 2B | LR~Cul; LR~50s; 50s~60s-70s; 60s-70s~80s-90s | salt | Xu et al., 2013 |
| TraesCS2B01G182800 | *Lr13* | 2B | LR~50s; 50s~60s-70s | leaf rust; early leaf senescence;necrosis; stripe rust | Hewitt et al., 2021 |
| TraesCS2B01G194200 | *TaSuSy2* | 2B | LR~50s | starch | Mukherjee et al., 2015 |
| TraesCS2B01G200800 | *FUL3-2B* | 2B | LR~Cul; LR~50s | wheat spike and spikelet development/flowering/plant height | Li et al., 2019 |
| TraesCS2B01G201900 | *TaCYP78A5-2B* | 2B | 60s-70s~80s-90s | grain weight; grain yield per plant | Guo et al., 2022 |
| TraesCS2B01G202300 | *TaGW7* | 2B | 60s-70s~80s-90s | grain shape and grain weight | Wang et al., 2019 |
| TraesCS2B01G202700 | *TaAFP-B* | 2B | 50s~60s-70s; 60s-70s~80s-90s | Seed Dormancy | Feng et al., 2022 |
| TraesCS2B01G281000 | *FUL2-2B* | 2B | LR~50s | wheat spike and spikelet development/flowering/plant height | Li et al., 2019 |
| TraesCS2B01G309500 | *TaSBEIIa* | 2B | LR~50s; 50s~60s-70s; 60s-70s~80s-90s | starch | Pei et al., 2023 |
| TraesCS2B01G327300 | *TaSBEIIb* | 2B | LR~Cul; LR~50s; 50s~60s-70s; 80s-90s~00s- | starch | Baysal et al., 2020 |
| TraesCS2B01G343600 | *TaNAC100-2B* | 2B | LR~50s; 50s~60s-70s | total seed proteins; plant height; heading date; seed size; thousand kernel weight | Li et al., 2021 |
| TraesCS2B01G405700 | *GNI-B1* | 2B | 50s~60s-70s; 60s-70s~80s-90s | grain number; grain weight | Sakuma et al., 2019 |
| TraesCS2B01G578500 | *TaARF12-2B* | 2B | LR~50s | plant height; spike length; seeds per spike; grain weight | Kong et al., 2023 |
| TraesCS2B01G587900 | *TaSTT3b-2B* | 2B | 60s-70s~80s-90s | sharp eyespot and increases grain weight | Zhu et al., 2022 |
| TraesCSU01G196100 | *PPD-B1* | 2B | 60s-70s~80s-90s | photoperiod | Bealeset al., 2007 |
| TraesCS2D01G006600 | *TaHXK3-2D* | 2D | 60s-70s~80s-90s | stomatal index | Li et al., 2022 |
| TraesCS2D01G016400 | *Sr46* | 2D | 50s~60s-70s; 80s-90s~00s- | stem rust | Arora et al., 2019 |
| TraesCS2D01G016900 | *TaDA1* | 2D | 50s~60s-70s; 80s-90s~00s- | grain size | Liu et al., 2020 |
| TraesCS2D01G118200 | *trs2* | 2D | 80s-90s~00s- | spikelet formation and awn elongation | Du et al., 2021 |
| TraesCS2D01G123000 | *TaNHX2-2D* | 2D | 80s-90s~00s- | salt | Xu et al., 2013 |
| TraesCS2D01G181400 | *FUL3-2D* | 2D | 60s-70s~80s-90s | wheat spike and spikelet development/flowering/plant height | Li et al., 2019 |
| TraesCS2D01G324700 | *TaNAC100-2D* | 2D | 80s-90s~00s- | total seed proteins; plant height; heading date; seed size; thousand kernel weight | Li et al., 2021 |

| | | | | | |
|---|---|---|---|---|---|
| TraesCS2D01G385500 | *GNI-D1* | 2D | 50s~60s-70s; 60s-70s~80s-90s | grain number; grain weight | Sakuma et al., 2019 |
| TraesCS2D01G548900 | *TaARF12-2D* | 2D | 50s~60s-70s | plant height; spike length; seeds per spike; grain weight | Kong et al., 2023 |
| TraesCS2D01G558800 | *TaSTT3b-2D* | 2D | LR~50s | sharp eyespot and increases grain weight | Zhu et al., 2022 |
| TraesCS2D01G567000 | *TaSPR-D* | 2D | 50s~60s-70s | seed storage protein synthesis | Shen et al., 2021 |
| TraesCSU01G024900 | *Rht8* | 2D | 60s-70s~80s-90s | plant height | Chai et al., 2022 |
| TraesCS3A01G006600 | *TaPHS1* | 3A | LR~Cul; LR~50s; 50s~60s-70s | Pre-harvest sprouting | Liu et al., 2013 |
| TraesCS3A01G049500 | *Stb6* | 3A | LR~Cul; LR~50s | Septoria tritici blotch disease | Saintenac et al., 2018 |
| TraesCS3A01G077900 | *TaNAC019-3A* | 3A | 60s-70s~80s-90s; 80s-90s~00s- | glutenin and starch accumulation | Gao et al., 2021 |
| TraesCS3A01G116300 | *TaGI-3A* | 3A | 80s-90s~00s- | flowering; photoperiods | Zhao et al., 2005 |
| TraesCS3A01G143100 | *FT-A2* | 3A | 60s-70s~80s-90s; 80s-90s~00s- | grain number per spike | Glenn et al., 2022 |
| TraesCS3A01G155200 | *TaIAA17-3A* | 3A | 80s-90s~00s- | tiller angle | Zhao et al., 2023 |
| TraesCS3A01G249100 | *TaFUSCA3-3A* | 3A | LR~50s; 60s-70s~80s-90s | end-use quality/seed protein | Sun et al., 2017 |
| TraesCS3A01G311100 | *CKX2.2.1_3A* | 3A | LR~Cul | Grain Size; Filling Rate and weight | Jain et al., 2022 |
| TraesCS3A01G314300 | *AG2-3A* | 3A | LR~50s; 60s-70s~80s-90s | wheat spike and spikelet/florets development | Meguro et al., 2023 |
| TraesCS3A01G328000 | *TaERF3-3A* | 3A | 80s-90s~00s- | grain size, grain weight | Jia et all., 2021 |
| TraesCS3A01G336500 | *TaGAMYB3* | 3A | LR~Cul; 50s~60s-70s; 60s-70s~80s-90s; 80s-90s~00s-; | plant architecture | Jian et al., 2022 |
| TraesCS3A01G401700 | *TaZIP4-A1* | 3A | 50s~60s-70s; 60s-70s~80s-90s | synapsis and crossover of homologous chromosomes | Rey et al., 2017 |
| TraesCS3A01G519000 | *TaDWF4-3A* | 3A | 50s~60s-70s; 80s-90s~00s- | BR synthesis pathway; carbon assimilation; wheat productivity; N use efficiency | Milner et al., 2022 |
| TraesCS3B01G007400 | *TaMFT_3B2* | 3B | LR~Cul | Flowering time | Vetch et al., 2022 |
| TraesCS3B01G010100 | *TaMFT_3B1* | 3B | LR~Cul | Seed dormancy | Vetch et al., 2022 |
| TraesCS3B01G019900 | *Fhb1* | 3B | LR~50s; 50s~60s-70s; 60s-70s~80s-90s | Fusarium head blight | Li et al., 2019 |
| TraesCS3B01G092800 | *TaNAC019-3B* | 3B | 80s-90s~00s- | glutenin and starch accumulation | Gao et al., 2021 |
| TraesCS3B01G135400 | *TaGI-3B* | 3B | 50s~60s-70s | flowering; photoperiods | Zhao et al., 2005 |
| TraesCS3B01G157500 | *AG2-3B* | 3B | LR~50s; 50s~60s-70s | wheat spike and spikelet/florets development | Meguro et al., 2023 |
| TraesCS3B01G162000 | *FT-B2* | 3B | 50s~60s-70s | grain number per spike | Glenn et al., 2022 |
| TraesCS3B01G181500 | *TaIAA17-3B* | 3B | 50s~60s-70s | tiller angle | Zhao et al., 2023 |
| TraesCS3B01G277100 | *TaGS5-3B* | 3B | LR~50s | grain weight | Ma et al., 2016 |
| TraesCS3B01G392800 | *TaMs5-B* | 3B | 50s~60s-70s | male sterility | Pallotta et al., 2019 |
| TraesCS3B01G399800 | *TaWOX5-3B* | 3B | LR~Cul; 50s~60s-70s; 60s-70s~80s-90s; 80s-90s~00s- | genetic transformation; transformation efficiency; flag leaf width | Wang et al., 2022 |

| | | | | | |
|---|---|---|---|---|---|
| TraesCS3B01G515900 | *Tamyb10-B1* | 3B | 50s~60s-70s; 60s-70s~80s-90s; 80s-90s~00s- | Red grain colour | Himi et al., 2005 |
| TraesCS3B01G572900 | *TaSINA* | 3B | 60s-70s~80s-90s; 80s-90s~00s- | increased yield in hot and dry climates | Thomelin et al., 2021 |
| TraesCS3B01G586500 | *TaDWF4-3B* | 3B | 50s~60s-70s; 60s-70s~80s-90s | BR synthesis pathway; carbon assimilation; wheat productivity; N use efficiency | Milner et al., 2022 |
| TraesCS3B01G594300 | *WPCL1-3B* | 3B | 50s~60s-70s; 60s-70s~80s-90s | Flowering time; heading time | Mizuno et al., 2016 |
| TraesCS3B01G608800 | *TdDof* | 3B | LR~Cul; LR~50s; 50s~60s-70s | solid stem | Nilsen et al., 2020 |
| TraesCS3D01G078500 | *TaNAC019-3D* | 3D | LR~Cul; 80s-90s~00s- | glutenin and starch accumulation | Gao et al., 2021 |
| TraesCS3D01G118200 | *TaGI-3D* | 3D | 50s~60s-70s | flowering; photoperiods | Zhao et al., 2005 |
| TraesCS3D01G144500 | *FT-D2* | 3D | 50s~60s-70s | grain number per spike | Glenn et al., 2022 |
| TraesCS3D01G162700 | *TaIAA17-3D* | 3D | 80s-90s~00s- | tiller angle | Zhao et al., 2023 |
| TraesCS3D01G396500 | *TaZIP4-D1* | 3D | 50s~60s-70s | synapsis and crossover of homologous chromosomes | Rey et al., 2017 |
| TraesCS3D01G500800 | *Stb16q* | 3D | 60s-70s~80s-90s | Septoria tritici blotch | Saintenac et al., 2021 |
| TraesCS3D01G526300 | *TaDWF4-3D1* | 3D | LR~Cul; 50s~60s-70s; | BR synthesis pathway; carbon assimilation; wheat productivity; N use efficiency | Milner et al., 2022 |
| TraesCS3D01G526400 | *TaDWF4-3D2* | 3D | LR~Cul; 50s~60s-70s; | BR synthesis pathway;carbon assimilation;wheat productivity;N use efficiency | Milner et al., 2022 |
| TraesCS5B01G255100 | *TaZIP4-B2* | 3D | 50s~60s-70s | synapsis and crossover of homologous chromosomes | Rey et al., 2017 |
| TraesCS4A01G002600 | *SVP3-4A* | 4A | 50s~60s-70s | wheat spike and spikelet development/flowering/plant height | Li et al., 2021 |
| TraesCS4A01G005600 | *TaSRL1-4A* | 4A | LR~Cul; LR~50s; 50s~60s-70s | root length; plant height | Zhuang et al., 2021 |
| TraesCS4A01G066200 | *Lr67* | 4A | 80s-90s~00s- | leaf rust; stripe rust; powdery mildew; stem rust | Moore et al., 2015 |
| TraesCS4A01G078000 | *TaDWF4-4A* | 4A | 80s-90s~00s- | BR synthesis pathway; carbon assimilation; wheat productivity; N use efficiency | Milner et al., 2022 |
| TraesCS4A01G105300 | *TaPRR73-4A* | 4A | 60s-70s~80s-90s | heading date and plant height | Zhang et al., 2016 |
| TraesCS4A01G106400 | *TaMPK3-4A* | 4A | 60s-70s~80s-90s | ABA; Drought tolerance | Liu et al., 2022 |
| TraesCS4A01G145300 | *TaNHX1-4A* | 4A | LR~50s | salt | Singh et al., 2021 |
| TraesCS4A01G201900 | *TaFROG* | 4A | LR~Cul | Fusarium head blight resistance | Perochon et al., 2015 |
| TraesCS4A01G271300 | *TaTB1-4A* | 4A | 50s~60s-70s; 60s-70s~80s-90s; 80s-90s~00s- | inflorescence architecture | Dixon et al., 2018 |
| TraesCS4A01G275900 | *Lpx-3* | 4A | 60s-70s~80s-90s | yellow color of pasta;yellow pigment content | Carrera et al., 2007 |
| TraesCS4A01G284000 | *BGC1* | 4A | 80s-90s~00s- | starch synthesis | Chia et al., 2020 |
| TraesCS4A01G347000 | *TaIRI9* | 4A | LR~Cul; LR~50s; 50s~60s-70s | vernalization | Cao et al ., 2021 |
| TraesCS4A01G418200 | *Wx* | 4A | 50s~60s-70s | amylose synthesis | Yamamori et al., 1994 |
| TraesCS4A01G446700 | *TaSuSy4* | 4A | 50s~60s-70s; 60s-70s~80s-90s | starch | Han et al., 2023 |

| | | | | | |
|---|---|---|---|---|---|
| TraesCS4B01G029700 | *BGC1* | 4B | LR~50s; 60s-70s~80s-90s; 80s-90s~00s- | starch synthesis | Chia et al., 2020 |
| TraesCS4B01G037700 | *Lpx-B1.2* | 4B | 80s-90s~00s- | yellow color of pasta; yellow pigment content | Carrera et al., 2007 |
| TraesCS4B01G037800 | *Lpx-3* | 4B | 80s-90s~00s- | yellow color of pasta; yellow pigment content | Carrera et al., 2007 |
| TraesCS4B01G042700 | *TaTB1-4B* | 4B | 80s-90s~00s- | inflorescence architecture | Dixon et al., 2018 |
| TraesCS4B01G043100 | *Rht-B1* | 4B | 60s-70s~80s-90s; 80s-90s~00s- | plant height | Peng et al., 1999 |
| TraesCS4B01G098200 | *TaNAC071-B* | 4B | 80s-90s~00s- | drought tolerance | Lv et al., 2021 |
| TraesCS4B01G125700 | *TaNHX1-4B* | 4B | 80s-90s~00s- | salt | Singh et al., 2021 |
| TraesCS4B01G198700 | *TaPRR73-4B* | 4B | 60s-70s~80s-90s | heading date and plant height | Zhang et al., 2016 |
| TraesCS4B01G234100 | *TaDWF4-4B* | 4B | LR~50s | BR synthesis pathway; carbon assimilation; wheat productivity; N use efficiency | Milner et al., 2022 |
| TraesCS4B01G243500 | *Lr67* | 4B | 60s-70s~80s-90s | leaf rust; stripe rust; powdery mildew; stem rust | Moore et al., 2015 |
| TraesCS4B01G299400 | *TaSRL1-4B* | 4B | 80s-90s~00s- | root length; plant height | Zhuang et al., 2021 |
| TraesCS4B01G317000 | *TaPHT1-9-4B* | 4B | 50s~60s-70s; 60s-70s~80s-90s | Pi uptake and translocation | Wang et al., 2021 |
| TraesCS4B01G322000 | *TaTMT3B* | 4B | 50s~60s-70s | alleviates growth and yield penalties associated with MLO disruption | Li et al., 2022 |
| TraesCS4B01G330000 | *TaBot1-4B* | 4B | 50s~60s-70s; 60s-70s~80s-90s; 80s-90s~00s- | boron toxicity tolerance | Sutton et al., 2007 |
| TraesCS4B01G346700 | *WSOC1-4B* | 4B | 80s-90s~00s- | flowering | Shitsukawa et al., 2007 |
| TraesCS4B01G372700 | *VRN-B2* | 4B | LR~Cul; LR~50s; 50s~60s-70s; | vernalization | Kippes et al., 2016 |
| TraesCS4B01G385500 | *TaDIR-B1* | 4B | 60s-70s~80s-90s | Fusarium crown rot | Yang et al., 2021 |
| TraesCS4D01G040100 | *TaTB1-4D* | 4D | 80s-90s~00s- | inflorescence architecture | Dixon et al., 2018 |
| TraesCS4D01G040400 | *Rht-D1* | 4D | LR~Cul; 80s-90s~00s- | plant height | Peng et al., 1999 |
| TraesCS4D01G147600 | *TaNHX1-4D* | 4D | 80s-90s~00s- | salt | Singh et al., 2021 |
| TraesCS4D01G243100 | *Lr67* | 4D | 80s-90s~00s- | leaf rust; stripe rust; powdery mildew; stem rust | Moore et al., 2015 |
| TraesCS4D01G319100 | *TaMlo-D1* | 4D | LR~Cul; 80s-90s~00s- | Powdery mildew resistance | Acevedo-Garcia et al., 2017 |
| TraesCS4D01G341700 | *WSOC1-4D* | 4D | 80s-90s~00s- | flowering | Shitsukawa et al., 2007 |
| TraesCS5A01G005400 | *Sr60* | 5A | LR~Cul; LR~50s | stem rust | Chen et al., 2020 |
| TraesCS5A01G038300 | *TaARF25-5A* | 5A | LR~50s; 50s~60s-70s; 60s-70s~80s-90s | grain size, grain weight | Jia et all., 2021 |
| TraesCS5A01G215100 | *TaDEP1-5A* | 5A | LR~50s | plant height | Saintenac et al., 2021 |
| TraesCS5A01G216200 | *TaQsd1-5A* | 5A | 50s~60s-70s; 80s-90s~00s- | Pre-harvest sprouting | Onishi et al., 2017 |
| TraesCS5A01G311600 | *CBF-A12* | 5A | LR~50s | Winter hardiness; cold | Vágújfalvi et al., 2005 |
| TraesCS5A01G344100 | *YrU1* | 5A | LR~Cul; 60s-70s~80s-90s | stripe rust | Wang et al., 2020 |

| | | | | | |
|---|---|---|---|---|---|
| TraesCS5A01G376500 | *TaPIL1* | 5A | LR~50s | tiller number | Zhang et al., 2022 |
| TraesCS5A01G378700 | *Lpx-A2* | 5A | LR~Cul; LR~50s; 50s~60s-70s; | yellow color of pasta; yellow pigment content | Meng et al., 2020 |
| TraesCS5A01G383400 | *TaHMA3-5A* | 5A | LR~50s; 80s-90s~00s- | cadmium | Zhang et al., 2020 |
| TraesCS5A01G460000 | *TaIPT8-5A* | 5A | 60s-70s~80s-90s; 80s-90s~00s- | drought tolerance | Wang et al., 2023 |
| TraesCS5A01G472000 | *TaAGPS2* | 5A | LR~Cul | Starch synthesis | Mukherjee et al., 2022 |
| TraesCS5A01G473800 | *Q* | 5A | 60s-70s~80s-90s | confers the free-threshing character and influences many other agronomically important traits | Faris et al., 2003 |
| TraesCS5A01G494800 | *TaMlo-A1* | 5A | 50s~60s-70s | Powdery mildew resistance | Acevedo-Garcia et al., 2017 |
| TraesCS5A01G501500 | *TaBot1* | 5A | 60s-70s~80s-90s; 80s-90s~00s- | boron toxicity tolerance | Sutton et al., 2007 |
| TraesCS5A01G515500 | *WSOC1-5A* | 5A | 50s~60s-70s; 60s-70s~80s-90s; 80s-90s~00s- | flowering | Shitsukawa et al., 2007 |
| TraesCS5A01G541300 | *VRN2-5A* | 5A | LR~Cul; LR~50s | vernalization | Yan et al., 2004 |
| TraesCS5A01G542800 | *ALI-1* | 5A | LR~50s; 80s-90s~00s- | awn | Wang et al., 2020 |
| TraesCS5A01G543100 | *Rht12* | 5A | LR~50s; 80s-90s~00s- | plant height | Sun et al., 2019 |
| TraesCS5A01G549300 | *TaDIR-A1* | 5A | LR~Cul; 60s-70s~80s-90s; 80s-90s~00s- | Fusarium crown rot | Yang et al., 2021 |
| TraesCS5B01G039800 | *TaARF25-5B* | 5B | 60s-70s~80s-90s | grain size, grain weight | Jia et all., 2021 |
| TraesCS5B01G154100 | *LYS3* | 5B | 50s~60s-70s | embryo growth | Orman-Ligeza et sl., 2020 |
| TraesCS5B01G202100 | *TaVIT2-5B* | 5B | 50s~60s-70s | Iron transporter | Connorton et al., 2017 |
| TraesCS5B01G208700 | *TaDEP1-5B* | 5B | LR~Cul;50s~60s-70s; 60s-70s~80s-90s | plant height | Zhang et al., 2016 |
| TraesCS5B01G214700 | *TaQsd1-5B* | 5B | LR~Cul | Pre-harvest sprouting | Abe et al., 2019 |
| TraesCS5B01G312200 | *CBF-B12* | 5B | LR~Cul; LR~50s; 60s-70s~80s-90s; 80s-90s~00s- | Winter hardiness; cold | Pearce et al., 2013 |
| TraesCS5B01G316900 | *TaHST1L-5B* | 5B | 50s~60s-70s; 60s-70s~80s-90s | tiller angle | Zhao et al., 2023 |
| TraesCS5B01G344200 | *YrU1-5B* | 5B | 60s-70s~80s-90s | stripe rust | Wang et al., 2020 |
| TraesCS5B01G382300 | *Lpx-A2* | 5B | LR~Cul; LR~50s; 50s~60s-70s | yellow color of pasta; yellow pigment content | Carrera et al., 2007 |
| TraesCS5B01G388000 | *TaHMA3-5B* | 5B | 50s~60s-70s; 60s-70s~80s-90s; 80s-90s~00s- | cadmium | Maccaferri et al., 2019 |
| TraesCS5B01G396200 | *PhyC* | 5B | LR~Cul; LR~50s | heading date; flowering time | Mizuno et al., 2012 |
| TraesCS5B01G486900 | *Q* | 5B | LR~50s; 80s-90s~00s- | confers the free-threshing character and influences many other agronomically important traits | Faris et al., 2003 |
| TraesCS5B01G565100 | *TaMKK3-B* | 5B | LR~Cul; LR~50s; 50s~60s-70s | pre-harvest sprouting; seed dormancy | Torada et al., 2016 |
| TraesCS5D01G322600 | *TaHST1L-5D* | 5D | 60s-70s~80s-90s | tiller angle | Zhao et al., 2023 |
| TraesCS5D01G349800 | *YrU1-5D* | 5D | 60s-70s~80s-90s | stripe rust | Wang et al., 2020 |
| TraesCS5D01G549600 | *TaMKK3-D* | 5D | 50s~60s-70s | pre-harvest sprouting; seed dormancy | Torada et al., 2016 |

| | | | | | |
|---|---|---|---|---|---|
| TraesCS6A01G091300 | *TaOGT-6A* | 6A | LR~Cul | Flowering time; Vernalization | Fan et al., 2021 |
| TraesCS6A01G259000 | *TaAGL6-A* | 6A | 60s-70s~80s-90s | floral organ; spikelet meristem; spikelet number per spike; grain number per spike | Kong et al., 2022 |
| TraesCS6A01G392400 | *KAT-2A* | 6A | 60s-70s~80s-90s | Grain weight | Chen et al., 2020 |
| TraesCS6B01G029600 | *Yr15-6B* | 6B | LR~Cul; LR~50s | Stripe rust | Klymiuk et al., 2018 |
| TraesCS6B01G119500 | *TaOGT-6B* | 6B | LR~Cul | Flowering time; Vernalization | Fan et al., 2021 |
| TraesCS6B01G167900 | *TaPsIPK1-6B* | 6B | 50s~60s-70s; 60s-70s~80s-90s | stripe rust | Wang et al., 2022 |
| TraesCS6B01G185700 | *TaCBF1b* | 6B | 60s-70s~80s-90s | stripe rust | Wang et al., 2022 |
| TraesCS6B01G207500 | *Gpc-B1* | 6B | 60s-70s~80s-90s | grain protein, zinc, and iron content | Uauy et al., 2006 |
| TraesCS6B01G286400 | *TaAGL6-B* | 6B | LR~Cul | floral organ; spikelet meristem; spikelet number per spike; grain number per spike | Kong et al., 2022 |
| TraesCS6B01G319500 | *TaHd1-6B* | 6B | 80s-90s~00s- | flowering | Nemoto et al., 2003 |
| TraesCS6B01G337300 | *TaPIN1-6B1* | 6B | LR~Cul; LR~50s; 60s-70s~80s-90s; 80s-90s~00s- | tiller number; spikelet number; grain number per panicle; 1000-grain weight | Yao et al., 2021 |
| TraesCS6D01G129900 | *TaPsIPK1-6D* | 6D | 50s~60s-70s | stripe rust | Wang et al., 2022 |
| TraesCS6D01G168200 | *TaBT1-6D* | 6D | 50s~60s-70s | starch synthesis and thousand kernel weight | Wang et al., 2019 |
| TraesCS6D01G176900 | *TaGW2-D1* | 6D | 80s-90s~00s- | grain weight; grain size; grain width | Zhang et al., 2018 |
| TraesCS6D01G207100 | *TaPRR1-D1* | 6D | 60s-70s~80s-90s | heading date, plant height and thousand grain weight | Sun et al., 2020 |
| TraesCS6D01G240200 | *TaAGL6-D* | 6D | 60s-70s~80s-90s | floral organ; spikelet meristem; spikelet number per spike; grain number per spike | Kong et al., 2022 |
| TraesCS6D01G287800 | *TaPIN1-6D* | 6D | 50s~60s-70s; 60s-70s~80s-90s; 80s-90s~00s- | tiller number; spikelet number; grain number per panicle;1000-grain weight | Yao et al., 2021 |
| TraesCS6D01G493900 | *KAT-2D* | 6D | 60s-70s~80s-90s | Grain weight | Chen et al., 2022 |
| TraesCS7A01G070100 | *Wx* | 7A | LR~Cul; LR~50s | amylose synthesis | Yamamori et al., 1994 |
| TraesCS7A01G115400 | *VRN-A3* | 7A | LR~50s; 50s~60s-70s; 60s-70s~80s-90s; 80s-90s~00s- | Vernalization; flowering | Yan et al., 2006 |
| TraesCS7A01G140500 | *TaSRG1A* | 7A | LR~50s; 50s~60s-70s; 60s-70s~80s-90s | cadmium accumulation | Wei et al., 2021 |
| TraesCS7A01G141300 | *TaAVP1-7A* | 7A | LR~50s; 50s~60s-70s; 60s-70s~80s-90s | salt | Menadue et al., 2021 |
| TraesCS7A01G151900 | *TaGW2L-7A* | 7A | 50s~60s-70s; 60s-70s~80s-90s; 80s-90s~00s- | heading date | Zhang et al., 2022 |
| TraesCS7A01G152500 | *TaNAC77-7A* | 7A | 60s-70s~80s-90s | seed storage protein | Luo et al., 2021 |
| TraesCS7A01G263900 | *TaOTUB1* | 7A | LR~50s | tiller number | Jiang et al., 2021 |
| TraesCS7A01G369500 | *TaMBF1c* | 7A | 60s-70s~80s-90s | heat | Tian et al., 2022 |
| TraesCS7A01G418900 | *TaD27-A* | 7A | 60s-70s~80s-90s; 80s-90s~00s- | tiller number | Zhao et al., 2020 |
| TraesCS7A01G423100 | *TaZIM-A1* | 7A | 50s~60s-70s | flowering time; thousand-kernel weight | Liu et al., 2019 |

| | | | | | |
|---|---|---|---|---|---|
| TraesCS7A01G497200 | *TaCol-B5-7A* | 7A | LR~50s | spike architecture; grain yield | Zhang et al., 2022 |
| TraesCS7A01G499600 | *Sr22* | 7A | LR~50s | stem rust | Karelov et al., 2022 |
| TraesCS7A01G514400 | *Pinb2-1* | 7A | LR~50s | grain hardness; kernel texture | Li et al., 2020 |
| TraesCS7A01G514500 | *Pinb2-2* | 7A | LR~50s | grain hardness; kernel texture | Li et al., 2020 |
| TraesCS7A01G514505 | *Pinb2-3* | 7A | LR~50s | grain hardness; kernel texture | Li et al., 2020 |
| TraesCS7A01G540000 | *SuSr-D1-7A* | 7A | LR~Cul; LR~50s | suppression of stem rust resistance | Hiebert et al., 2020 |
| TraesCS7A01G549100 | *TaSBEIa* | 7A | LR~Cul; LR~50s; 50s~60s-70s; 60s-70s~80s-90s; 80s-90s~00s- | starch | Liu et al., 2022 |
| TraesCS7A01G549300 | *TaSBEIb* | 7A | LR~Cul; LR~50s; 50s~60s-70s; 60s-70s~80s-90s; 80s-90s~00s- | starch | Pei et al., 2023 |
| TraesCS7A01G557300 | *Psy1* | 7A | LR~50s | Phytoene synthase; yellow pigment content | He et al., 2008 |
| TraesCS7B01G010000 | *TaGW2L-7B* | 7B | LR~50s | heading date | Zhang et al., 2022 |
| TraesCS7B01G013100 | *TaVRN3-7B* | 7B | LR~Cul; LR~50s | Vernalization | Yan et al., 2006 |
| TraesCS7B01G020800 | *SEP1-4-7B* | 7B | LR~50s | flower development and morphogenesis | Paolacci et al., 2007 |
| TraesCS7B01G118300 | *TaHd1-7B* | 7B | 50s~60s-70s; 80s-90s~00s- | flowering | Shimada et al., 2009 |
| TraesCS7B01G196800 | *TaARE1-7B* | 7B | 60s-70s~80s-90s | nitrogen utilization | Zhang et al., 2021 |
| TraesCS7B01G259000 | *TaMBF1c* | 7B | 80s-90s~00s- | heat | Tian et al., 2022 |
| TraesCS7B01G286600 | *AP3-1-7B* | 7B | 80s-90s~00s- | wheat spike and spikelet/florets development | Hama et al., 2004 |
| TraesCS7B01G319100 | *TaD27-B* | 7B | 80s-90s~00s- | tiller number | Zhao et al., 2020 |
| TraesCS7B01G323600 | *TaZIM-B1* | 7B | 80s-90s~00s- | flowering time; thousand-kernel weight | Liu et al., 2019 |
| TraesCS7B01G400600 | *TaCol-B5* | 7B | 60s-70s~80s-90s; 80s-90s~00s- | spike architecture; grain yield | Zhang et al., 2022 |
| TraesCS7B01G441700 | *Pm5e* | 7B | LR~50s | Powdery mildew resistance | Xie et al., 2020 |
| TraesCS7B01G460900 | *SuSr-D1-7B-1* | 7B | LR~50s | suppression of stem rust resistance | Hiebert et al., 2020 |
| TraesCS7B01G472300 | *TaSBEIb* | 7B | LR~50s | starch | Pei et al., 2023 |
| TraesCS7B01G475400 | *Bo1* | 7B | LR~50s; 50s~60s-70s | boron tolerance | Pallotta et al., 2014 |
| TraesCS7B01G482000 | *Psy1* | 7B | LR~50s | Phytoene synthase; yellow pigment content | He et al., 2008 |
| TraesCS7D01G036600 | *TaSuSy4* | 7D | 60s-70s~80s-90s | starch | Han et al., 2023 |
| TraesCS7D01G080300 | *Lr34* | 7D | 80s-90s~00s- | leaf rust; stripe rust; powdery mildew; leaf tip necrosis | Krattinger et al., 2009 |
| TraesCS7D01G107500 | *TaGW2L-7D* | 7D | 50s~60s-70s | heading date | Zhang et al., 2022 |
| TraesCS7D01G111600 | *VRN-D3* | 7D | 50s~60s-70s | Vernalization; flowering | Yan et al., 2006 |
| TraesCS7D01G120500 | *SEP1-4-7D* | 7D | 50s~60s-70s; 80s-90s~00s- | flower development and morphogenesis | Schilling et al., 2020 |
| TraesCS7D01G154200 | *TaNAC77-7D* | 7D | 60s-70s~80s-90s | seed storage protein | Luo et al., 2021 |

| | | | | | |
|---|---|---|---|---|---|
| TraesCS7D01G176700 | *TaVRT-2* | 7D | 80s-90s~00s- | flowering | Kane et al., 2005 |
| TraesCS7D01G283700 | *TaARE1-7D* | 7D | 50s~60s-70s | nitrogen utilization | Zhang et al., 2021 |
| TraesCS7D01G353800 | *TaMBF1c* | 7D | 80s-90s~00s- | heat | Tian et al., 2022 |
| TraesCS7D01G416000 | *TaZIM-D1* | 7D | 60s-70s~80s-90s | flowering time; thousand-kernel weight | Liu et al., 2019 |
| TraesCS7D01G526100 | *SuSr-D1* | 7D | 60s-70s~80s-90s | suppression of stem rust resistance | Hiebert et al., 2020 |
| TraesCS7D01G553300 | *Psy1* | 7D | 60s-70s~80s-90s | Phytoene synthase; yellow pigment content | He et al., 2008 |

**Supplementary Table S2**. Genes in the selection intervals associated with wheat trait changes

| Trait type | Gene | Symbol | Chr | Group | Trait | XP-CLR | SweeD | $F_{st}$ | Tajima's D | $\pi_i/\pi_j$ |
|---|---|---|---|---|---|---|---|---|---|---|
| Morphology | TraesCS7D01G120500 | *SEP1-4-7D* | 7D | 50s ~ 60s-70s | Flower Development And Morphogenesis | | 75.12 | | 4.55 | |
| | TraesCS2B01G405700 | *GNI-B1* | 2B | | Grain Number | | 22.96 | | | |
| | TraesCS4A01G271300 | *TaTB1-4A* | 4A | | Inflorescence Architecture | 5.75 | 142.10 | | | |
| | TraesCS3A01G336500 | *TaGAMYB3* | 3A | | Plant Architecture | 4.04 | 213.78 | | | |
| | TraesCS2B01G343600 | *TaNAC100-2B* | 2B | | Plant Height | | 213.78 | 0.25 | | |
| | TraesCS4A01G002600 | *SVP3-4A* | 4A | | Plant Height | | 9.96 | | | |
| | TraesCS5B01G208700 | *TaDEP1-5B* | 5B | | Plant Height | 8.88 | 28.19511 | 0.269148 | | |
| | TraesCS2D01G548900 | *TaARF12-2D* | 2D | | Plant Height; Spike Length | | | 0.69 | 6.15 | |
| | TraesCS4A01G005600 | *TaSRL1-4A* | 4A | | Root Length; Plant Height | 6.85 | 49.97 | | | |
| | TraesCS3B01G608800 | *TdDof* | 3B | | Solid Stem | 45.55 | 8.22 | | | |
| | TraesCS3B01G181500 | *TaIAA17-3B* | 3B | | Tiller Angle | | 61.11 | | 4.35 | |
| | TraesCS5B01G316900 | *TaHST1L-5B* | 5B | | Tiller Angle | 9.14 | 213.14 | | | |
| Yield | TraesCS2B01G343600 | *TaNAC100-2B* | 2B | 50s ~ 60s-70s | Grain Weight | | 213.78 | 0.25 | | |
| | TraesCS2B01G405700 | *GNI-B1* | 2B | | Grain Weight | | 22.96 | | | |
| | TraesCS2D01G385500 | *GNI-D1* | 2D | | Grain Weight | | 58.13 | | | 3.27 |
| | TraesCS2D01G548900 | *TaARF12-2D* | 2D | | Grain Weight | | | 0.69 | 6.15 | |
| | TraesCS5A01G038300 | *TaARF25-5A* | 5A | | Grain Weight | 4.17 | 64.91 | | | |
| | TraesCS6D01G168200 | *TaBT1-6D* | 6D | | Grain Weight | 4.02 | | 0.11 | | 2.27 |
| | TraesCS6D01G287800 | *TaPIN1-6D* | 6D | | Grain Weight | | | | | |
| | TraesCS7A01G423100 | *TaZIM-A1* | 7A | | Grain Weight | 3.06 | 200.86 | | 6.95 | |
| | TraesCS3A01G143100 | *FT-A2* | 3A | | Grain Number Per Spike | | 17.65 | | | 6.73 |

| | | | | | | | | | | |
|---|---|---|---|---|---|---|---|---|---|---|
| | TraesCS1A01G264300 | *WPI-1-1A* | 1A | 80s-90s ~ 00s- | Spike And Spikelet/Florets Development | 15.52 | 54.79 | | 4.53 | |
| | TraesCS1D01G264500 | *PI1-1D* | 1D | | Spike And Spikelet/Florets Development | 39.00 | 21.29 | | | |
| | TraesCS7B01G286600 | *AP3-1-7B* | 7B | | Spike And Spikelet/Florets Development | | 22.92 | | 4.74 | |
| | TraesCS7B01G400600 | *TaCol-B5* | 7B | | Spike Architecture | 10.55 | 14.38 | | | |
| | TraesCS2D01G118200 | *trs2* | 2D | | Spikelet Formation | | 15.15 | | 5.03 | |
| | TraesCS6B01G337300 | *TaPIN1-6B1* | 6B | | Spikelet Number | | 17.91 | | | |
| | TraesCS6D01G287800 | *TaPIN1-6D* | 6D | | Spikelet Number | 7.97 | 48.64 | | | |
| | TraesCS5A01G038300 | *TaARF25-5A* | 5A | LR~50s | Grain Size | 1579.64 | 43.03 | | 4.59 | |
| Grain-filling | TraesCS1A01G353300 | *TaSS4-1A* | 1A | 50s ~ 60s-70s | Starch | | | 0.48 | | |
| | TraesCS1D01G356900 | *TaSS4-1D* | 1D | | Starch | 5.50 | 29.78 | | | |
| | TraesCS2B01G309500 | *TaSBEIIa* | 2B | | Starch | 10.65 | 16.05 | | | |
| | TraesCS2B01G327300 | *TaSBEIIb* | 2B | | Starch | | 29.99 | | 6.07 | |
| | TraesCS4A01G446700 | *TaSuSy4* | 4A | | Starch | | 54.88 | | | |
| | TraesCS6D01G168200 | *TaBT1-6D* | 6D | | Starch | 4.018615 | | 0.111305 | | 2.27 |
| | TraesCS7A01G549100 | *TaSBEIa* | 7A | | Starch | | 54.88 | | | 2.17 |
| | TraesCS7A01G549300 | *TaSBEIb* | 7A | | Starch | | 65.21 | | 5.36 | 10.72 |
| | TraesCS2B01G309500 | *TaSBEIIa* | 2B | 60s-70s ~ 80s-90s | Starch | 3.93 | 128.60 | | 4.38 | |
| | TraesCS3A01G077900 | *TaNAC019-3A* | 3A | | Starch | | | | 4.79 | |
| | TraesCS4A01G446700 | *TaSuSy4* | 4A | | Starch | 3.57 | 173.10 | | | |
| | TraesCS4B01G029700 | *BGC1* | 4B | | Starch | 4.25 | 15.53 | | | |
| | TraesCS7A01G549100 | *TaSBEIa* | 7A | | Starch | | 15.54 | | 4.69 | 2.17 |
| | TraesCS7A01G549300 | *TaSBEIb* | 7A | | Starch | | 23.47 | | 5.95 | 10.72 |
| | TraesCS7D01G036600 | *TaSuSy4* | 7D | | Starch | 8.51 | 57.90 | 0.23 | | |
| Resistance | TraesCS4B01G243500 | *Lr67* | 4B | 60s-70s ~ | Powdery Mildew | | 19.20868 | | 6.12263 | |
| | TraesCS2B01G488000 | *YrSP* | 2B | | Stripe Rust | | 57.90 | | | |

| | | | | | | | | | | |
|---|---|---|---|---|---|---|---|---|---|---|
| | TraesCS4B01G243500 | *Lr67* | 4B | 80s-90s | Stripe Rust | | 19.21 | | 6.12 | |
| | TraesCS5A01G344100 | *YrU1* | 5A | | Stripe Rust | 4.77 | 16.73 | | | 2.26 |
| | TraesCS5B01G344200 | *YrU1-5B* | 5B | | Stripe Rust | 31.72 | 36.02 | | 4.49 | |
| | TraesCS5D01G349800 | *YrU1-5D* | 5D | | Stripe Rust | 22.78 | 139.38 | | 5.27 | |
| | TraesCS6B01G167900 | *TaPsIPK1-6B* | 6B | | Stripe Rust | 13.85 | 21.22 | | | |
| | TraesCS6B01G185700 | *TaCBF1b* | 6B | | Stripe Rust | 11.17 | 240.77 | | 4.99 | |
| | TraesCS4A01G066200 | *Lr67* | 4A | 80s-90s ~ 00s- | Powdery Mildew | 15.817214 | 11.98242 | | | |
| | TraesCS4D01G243100 | *Lr67* | 4D | | Powdery Mildew | 8.30076 | 20.813 | | | |
| | TraesCS4D01G319100 | *TaMlo-D1* | 4D | | Powdery Mildew | 8.28 | 33.35 | | | |
| | TraesCS7D01G080300 | *Lr34* | 7D | | Powdery Mildew | 10.177659 | 23.51569 | 0.108456 | | |
| | TraesCS4A01G066200 | *Lr67* | 4A | | Stripe Rust | 15.82 | 11.98 | | | |
| | TraesCS4D01G243100 | *Lr67* | 4D | | Stripe Rust | 8.30 | 20.81 | | | |
| | TraesCS7D01G080300 | *Lr34* | 7D | | Stripe Rust | 10.18 | 23.52 | 0.11 | | |

**Supplementary Table S3**. Five selection signal genes out of 28 important genes in Zhengmai 7698

| **Gene/subunit** | **Group** | **Superior traits** | **Reference** |
|---|---|---|---|
| TraesCS1A01G317311 | 60s-70s~80s-90s; 80s-90s~00s- | High molecular weight glutenin subunits | Liu et al., 2008 |
| TraesCS1B01G329711 | 50s~60s-70s | | Schwarz et al., 2004 |
| TraesCS1B01G329992 | 50s~60s-70s | | Lei et al., 2006 |
| TraesCS2A01G468200 | 60s-70s~80s-90s; 80s-90s~00s- | Polyphenol oxidase activity | He et al., 2007 |
| TraesCS3A01G006600 | 50s~60-70s | Pre-harvest sprouting resistant | Liu et al., 2013 |

**Supplementary Table S4**. All QTNs identified to be associated with wheat plant height.

| Trait | Marker | Chr | Position | Add.effect | Dom.effect | P-value | Method | Sig. |
|---|---|---|---|---|---|---|---|---|
| PH_17AY | AX-110548369 | 1A | 13882546 | 1.5788 | | 2.53E-05 | BLUPmrMLM | SUG |
| PH_17AY | AX-109577721 | 1A | 13916589 | -1.7082 | 0.6206 | 6.90E-11 | IIIVmrMLM | SIG |
| PH_17AY | AX-109832236 | 1A | 15375351 | 5.3209 | | 6.12E-07 | FarmCPU | SUG |
| PH_17AY | AX-109827473 | 1A | 412791189 | 1.6840 | | 1.36E-04 | BLUPmrMLM | SUG |
| PH_17AY | AX-109064427 | 1B | 133161502 | 3.6933 | 0.1971 | 6.44E-35 | IIIVmrMLM | SIG |
| PH_17AY | AX-110568993 | 1B | 483581912 | -1.9629 | | 4.76E-06 | BLUPmrMLM | SUG |
| PH_17AY | AX-110012360 | 1B | 551118742 | -5.1866 | | 1.27E-06 | FarmCPU | SUG |
| PH_17AY | AX-108934179 | 1D | 20310288 | -1.3996 | 2.9418 | 2.39E-08 | IIIVmrMLM | SIG |
| PH_17AY | AX-111544586 | 2A | 76511277 | 6.0479 | | 1.44E-07 | FarmCPU | SUG |
| PH_17AY | AX-109512157 | 2A | 661485453 | -2.0098 | | 5.37E-07 | BLUPmrMLM | SUG |
| PH_17AY | AX-94429380 | 2A | 748542204 | -1.8028 | | 2.76E-13 | IIIVmrMLM | SIG |
| PH_17AY | AX-109940975 | 2B | 29113742 | -1.8482 | -3.5915 | 1.02E-22 | IIIVmrMLM | SIG |
| PH_17AY | AX-110931530 | 2B | 30133814 | -0.9296 | 7.4012 | 1.94E-53 | IIIVmrMLM | SIG |
| PH_17AY | AX-108831673 | 2B | 116849598 | 2.1884 | | 1.64E-18 | IIIVmrMLM | SIG |
| PH_17AY | AX-110990968 | 2B | 394112535 | 0.6179 | 16.6534 | 2.10E-10 | IIIVmrMLM | SIG |
| PH_17AY | AX-109406538 | 2D | 357488969 | 2.9135 | | 3.18E-30 | IIIVmrMLM | SIG |
| PH_17AY | AX-110914132 | 2D | 601962924 | -0.3340 | 8.4732 | 8.42E-07 | IIIVmrMLM | SUG |
| PH_17AY | AX-110486574 | 2D | 634143501 | 2.0480 | 4.7095 | 4.25E-16 | IIIVmrMLM | SIG |
| PH_17AY | AX-108734284 | 3A | 727001039 | -2.4286 | | 1.59E-07 | BLUPmrMLM | SUG |
| PH_17AY | AX-109935848 | 3B | 41828409 | 0.0839 | 5.2403 | 1.43E-11 | IIIVmrMLM | SIG |
| PH_17AY | AX-110369750 | 3B | 327881587 | 2.1860 | -2.2481 | 1.40E-17 | IIIVmrMLM | SIG |
| PH_17AY | AX-111030059 | 3B | 513548749 | -1.9743 | | 2.25E-05 | BLUPmrMLM | SUG |
| PH_17AY | AX-94503581 | 3B | 680243929 | -2.8711 | | 2.23E-06 | BLUPmrMLM | SUG |
| PH_17AY | AX-109292211 | 3B | 780616084 | -4.2657 | | 8.91E-07 | FarmCPU | SUG |
| PH_17AY | AX-110712777 | 3B | 821704357 | 0.3350 | 3.0649 | 1.93E-07 | IIIVmrMLM | SUG |
| PH_17AY | AX-94958584 | 3D | 21005788 | -2.2438 | | 1.02E-07 | BLUPmrMLM | SIG |
| PH_17AY | AX-111697670 | 3D | 611314553 | -0.3468 | -6.8453 | 1.43E-06 | IIIVmrMLM | SUG |
| PH_17AY | AX-108965701 | 4A | 147629644 | 2.1501 | | 5.59E-05 | BLUPmrMLM | SUG |
| PH_17AY | AX-94995565 | 4A | 636678133 | 6.0580 | 2.451 | 3.81E-51 | IIIVmrMLM | SIG |
| PH_17AY | AX-94746133 | 4A | 727055286 | 3.4269 | | 2.78E-09 | BLUPmrMLM | SIG |
| PH_17AY | AX-109945280 | 4A | 742841666 | -2.0998 | | 1.44E-05 | BLUPmrMLM | SUG |
| PH_17AY | AX-109434530 | 4B | 93539612 | 4.0813 | | 8.48E-53 | IIIVmrMLM | SIG |
| PH_17AY | AX-94477205 | 4B | 385424629 | 2.3149 | | 2.18E-20 | IIIVmrMLM | SIG |
| PH_17AY | AX-94464927 | 4B | 667680347 | -7.5358 | | 1.21E-10 | BLUPmrMLM | SIG |
| PH_17AY | AX-108804341 | 4D | 4137420 | 2.9258 | | 5.28E-06 | BLUPmrMLM | SUG |
| PH_17AY | AX-89593703 | 4D | 15958106 | 5.8181 | | 1.71E-06 | EMMAX | SUG |
| PH_17AY | AX-89593703 | 4D | 15958106 | -6.0658 | | 2.10E-07 | GEMMA | SUG |
| PH_17AY | AX-111743329 | 4D | 16360654 | 0.3557 | 8.6341 | 4.58E-07 | IIIVmrMLM | SUG |
| PH_17AY | AX-89703298 | 4D | 16926596 | 6.7316 | | 4.84E-07 | EMMAX | SUG |
| PH_17AY | AX-89703298 | 4D | 16926596 | -6.9433 | | 6.73E-08 | GEMMA | SIG |
| PH_17AY | AX-109281717 | 4D | 17613651 | 6.9439 | | 2.78E-07 | EMMAX | SUG |

| | | | | | | | | |
|---|---|---|---|---|---|---|---|---|
| PH_17AY | AX-109281717 | 4D | 17613651 | -7.1788 | | 3.16E-08 | GEMMA | SIG |
| PH_17AY | AX-109665328 | 4D | 18539670 | 6.4829 | | 1.16E-06 | EMMAX | SUG |
| PH_17AY | AX-109665328 | 4D | 18539670 | -6.7682 | | 1.28E-07 | GEMMA | SUG |
| PH_17AY | AX-108916749 | 4D | 19090268 | 6.3767 | | 4.93E-08 | EMMAX | SIG |
| PH_17AY | AX-108916749 | 4D | 19090268 | -6.5413 | | 6.94E-09 | GEMMA | SIG |
| PH_17AY | AX-108916749 | 4D | 19090268 | 4.3155 | 2.2266 | 3.67E-56 | IIIVmrMLM | SIG |
| PH_17AY | AX-89421921 | 4D | 19288898 | 6.8314 | | 3.66E-07 | EMMAX | SUG |
| PH_17AY | AX-89421921 | 4D | 19288898 | -7.0641 | | 4.35E-08 | GEMMA | SIG |
| PH_17AY | AX-95153659 | 4D | 301516416 | 1.8336 | | 1.83E-04 | BLUPmrMLM | SUG |
| PH_17AY | AX-110547306 | 5A | 20461065 | -0.5959 | -3.9244 | 2.22E-08 | IIIVmrMLM | SIG |
| PH_17AY | AX-109075553 | 5A | 50292393 | 1.9860 | | 4.85E-07 | BLUPmrMLM | SUG |
| PH_17AY | AX-111650496 | 5A | 197374930 | 2.2234 | -3.8468 | 2.03E-24 | IIIVmrMLM | SIG |
| PH_17AY | AX-109476851 | 5A | 568227919 | 4.3071 | | 1.37E-07 | BLUPmrMLM | SUG |
| PH_17AY | AX-108803976 | 5A | 578063291 | 1.1323 | 6.6775 | 9.24E-19 | IIIVmrMLM | SIG |
| PH_17AY | AX-108886743 | 5A | 641422431 | 0.3946 | -3.3056 | 2.90E-11 | IIIVmrMLM | SIG |
| PH_17AY | AX-110642982 | 5A | 703199986 | -8.2759 | | 1.54E-06 | GEMMA | SUG |
| PH_17AY | AX-110577776 | 5B | 66027364 | 3.2104 | | 4.04E-06 | BLUPmrMLM | SUG |
| PH_17AY | AX-108822938 | 5B | 66339657 | 8.3288 | | 3.05E-07 | FarmCPU | SUG |
| PH_17AY | AX-110016236 | 5B | 298198172 | -0.1918 | -8.4443 | 1.69E-11 | IIIVmrMLM | SIG |
| PH_17AY | AX-108747423 | 5B | 406217264 | -1.7377 | 2.3726 | 2.74E-12 | IIIVmrMLM | SIG |
| PH_17AY | AX-110371594 | 5B | 416378795 | -0.2564 | 4.3458 | 5.00E-07 | IIIVmrMLM | SUG |
| PH_17AY | AX-109349107 | 5B | 487601762 | 0.3226 | -4.1527 | 4.00E-06 | IIIVmrMLM | SUG |
| PH_17AY | AX-110018488 | 5B | 528535571 | 7.4778 | | 9.89E-07 | EMMAX | SUG |
| PH_17AY | AX-110018488 | 5B | 528535571 | -7.6263 | | 2.08E-07 | GEMMA | SUG |
| PH_17AY | AX-110018488 | 5B | 528535571 | 3.2459 | | 2.89E-36 | IIIVmrMLM | SIG |
| PH_17AY | AX-109934606 | 5B | 536575934 | 8.7689 | | 1.24E-06 | GEMMA | SUG |
| PH_17AY | AX-110443142 | 5B | 536591031 | -8.9947 | | 1.99E-06 | EMMAX | SUG |
| PH_17AY | AX-110443142 | 5B | 536591031 | 8.9406 | | 9.98E-07 | GEMMA | SUG |
| PH_17AY | AX-110921767 | 5B | 536795156 | 4.1873 | | 1.03E-06 | GEMMA | SUG |
| PH_17AY | AX-110385481 | 5B | 551505030 | 0.3569 | 4.7702 | 8.73E-15 | IIIVmrMLM | SIG |
| PH_17AY | AX-110991199 | 5B | 611650575 | 0.0206 | 5.5507 | 2.83E-07 | IIIVmrMLM | SUG |
| PH_17AY | AX-94503623 | 5B | 619135083 | -0.0682 | 3.7636 | 1.69E-13 | IIIVmrMLM | SIG |
| PH_17AY | AX-111582260 | 5B | 686426668 | -1.8931 | | 9.10E-05 | BLUPmrMLM | SUG |
| PH_17AY | AX-109567209 | 5D | 106520306 | -1.7201 | | 2.86E-12 | IIIVmrMLM | SIG |
| PH_17AY | AX-111529279 | 6A | 16437705 | 2.0214 | | 1.19E-05 | BLUPmrMLM | SUG |
| PH_17AY | AX-109019711 | 6A | 26975126 | 2.4186 | -1.1545 | 8.54E-12 | IIIVmrMLM | SIG |
| PH_17AY | AX-109947841 | 6B | 93435503 | -0.8577 | 7.2284 | 9.11E-08 | IIIVmrMLM | SIG |
| PH_17AY | AX-111195087 | 6B | 226358232 | -2.0613 | | 5.05E-06 | BLUPmrMLM | SUG |
| PH_17AY | AX-111006686 | 6B | 686803165 | -0.0531 | -17.543 | 9.38E-12 | IIIVmrMLM | SIG |
| PH_17AY | AX-94809200 | 6D | 15807973 | 2.6963 | -5.0792 | 4.27E-28 | IIIVmrMLM | SIG |
| PH_17AY | AX-95137140 | 7A | 15086668 | 1.8145 | | 1.97E-13 | IIIVmrMLM | SIG |
| PH_17AY | AX-108916605 | 7A | 125708264 | 3.4758 | | 3.30E-08 | BLUPmrMLM | SIG |
| PH_17AY | AX-111502341 | 7A | 198199217 | 6.1412 | | 3.21E-07 | FarmCPU | SUG |
| PH_17AY | AX-110381728 | 7A | 535901170 | -4.1764 | -1.0154 | 1.08E-48 | IIIVmrMLM | SIG |

| | | | | | | | | |
|---|---|---|---|---|---|---|---|---|
| PH_17AY | AX-94381170 | 7A | 636901870 | -1.0442 | 4.8151 | 3.09E-11 | IIIVmrMLM | SIG |
| PH_17AY | AX-111460480 | 7A | 688619689 | -3.7193 | | 9.92E-11 | BLUPmrMLM | SIG |
| PH_17AY | AX-111280522 | 7B | 130477199 | 0.2159 | -17.9745 | 1.42E-10 | IIIVmrMLM | SIG |
| PH_17AY | AX-111038845 | 7B | 328525711 | -2.6497 | -1.4832 | 2.42E-15 | IIIVmrMLM | SIG |
| PH_17AY | AX-111480183 | 7B | 328703745 | -2.3212 | | 3.31E-07 | BLUPmrMLM | SUG |
| PH_17AY | AX-110995160 | 7B | 554341864 | -3.3007 | | 3.28E-10 | BLUPmrMLM | SIG |
| PH_17AY | AX-110574804 | 7B | 605657821 | 0.7443 | -3.2614 | 9.31E-21 | IIIVmrMLM | SIG |
| PH_17AY | AX-94999858 | 7B | 706813719 | -1.8149 | | 1.03E-05 | BLUPmrMLM | SUG |
| PH_17AY | AX-110698849 | 7D | 351016239 | 3.1005 | | 1.31E-07 | BLUPmrMLM | SUG |
| PH_17AY | AX-110636845 | 7D | 617078025 | 0.7733 | -7.1209 | 3.66E-07 | IIIVmrMLM | SUG |
| PH_17JY | AX-94572346 | 1A | 14255910 | 0.3190 | -22.8765 | 1.09E-08 | IIIVmrMLM | SIG |
| PH_17JY | AX-110429109 | 1A | 189643659 | -0.7768 | 2.3994 | 1.30E-05 | IIIVmrMLM | SUG |
| PH_17JY | AX-110474000 | 1A | 504509704 | 4.0303 | | 9.75E-12 | BLUPmrMLM | SIG |
| PH_17JY | AX-109313554 | 1A | 508130904 | -2.1725 | | 1.08E-20 | IIIVmrMLM | SIG |
| PH_17JY | AX-110958868 | 1A | 569134121 | 2.6913 | | 8.19E-09 | BLUPmrMLM | SIG |
| PH_17JY | AX-111700376 | 1B | 45986664 | -0.3886 | -18.0595 | 1.14E-10 | IIIVmrMLM | SIG |
| PH_17JY | AX-94621343 | 1B | 158545260 | 1.9009 | | 5.39E-05 | BLUPmrMLM | SUG |
| PH_17JY | AX-109886737 | 1B | 390282044 | 1.7937 | | 5.35E-05 | BLUPmrMLM | SUG |
| PH_17JY | AX-109878755 | 1B | 395529482 | -0.9593 | -1.393 | 6.88E-05 | IIIVmrMLM | SUG |
| PH_17JY | AX-109878526 | 1B | 415386611 | 0.2712 | 4.5497 | 3.53E-08 | IIIVmrMLM | SIG |
| PH_17JY | AX-94656925 | 1D | 2485672 | 7.9474 | | 1.20E-10 | FarmCPU | SIG |
| PH_17JY | AX-110559092 | 1D | 387638588 | -2.5166 | | 7.50E-08 | BLUPmrMLM | SIG |
| PH_17JY | AX-94751461 | 1D | 419473390 | -1.3006 | | 1.07E-08 | IIIVmrMLM | SIG |
| PH_17JY | AX-110557158 | 2A | 189920245 | -3.2123 | | 1.36E-08 | BLUPmrMLM | SIG |
| PH_17JY | AX-110734128 | 2A | 307330154 | 0.0011 | -6.9042 | 5.90E-09 | IIIVmrMLM | SIG |
| PH_17JY | AX-110198456 | 2A | 317805831 | 3.0074 | | 4.41E-06 | BLUPmrMLM | SUG |
| PH_17JY | AX-109828660 | 2A | 413180560 | -2.0256 | | 1.90E-06 | BLUPmrMLM | SUG |
| PH_17JY | AX-109273607 | 2A | 483674366 | 3.2603 | | 1.42E-07 | BLUPmrMLM | SUG |
| PH_17JY | AX-109476100 | 2A | 699589854 | -2.5566 | 0.6234 | 1.40E-16 | IIIVmrMLM | SIG |
| PH_17JY | AX-111536980 | 2A | 735296764 | 0.4102 | -7.9308 | 3.34E-04 | IIIVmrMLM | SUG |
| PH_17JY | AX-111729566 | 2A | 753018070 | -0.1529 | 9.7562 | 1.05E-07 | IIIVmrMLM | SIG |
| PH_17JY | AX-110440215 | 2A | 753567811 | 2.3281 | | 5.06E-05 | BLUPmrMLM | SUG |
| PH_17JY | AX-111216759 | 2A | 758935907 | 1.5274 | -2.9681 | 2.14E-11 | IIIVmrMLM | SIG |
| PH_17JY | AX-109563795 | 2A | 768528507 | 0.0797 | 8.8402 | 1.24E-08 | IIIVmrMLM | SIG |
| PH_17JY | AX-108748416 | 2B | 115909069 | 9.9232 | | 2.53E-09 | FarmCPU | SIG |
| PH_17JY | AX-109351901 | 2B | 568443398 | 2.5608 | | 8.00E-07 | BLUPmrMLM | SUG |
| PH_17JY | AX-94622096 | 2B | 596927760 | 1.9361 | -0.9292 | 3.56E-15 | IIIVmrMLM | SIG |
| PH_17JY | AX-111450272 | 2B | 663839215 | 3.4961 | | 2.34E-15 | BLUPmrMLM | SIG |
| PH_17JY | AX-111450272 | 2B | 663839215 | -1.8029 | 0.9241 | 2.01E-12 | IIIVmrMLM | SIG |
| PH_17JY | AX-94494401 | 2B | 695375149 | -1.7105 | | 9.57E-14 | IIIVmrMLM | SIG |
| PH_17JY | AX-109000516 | 2B | 727685553 | 6.7133 | | 4.14E-08 | FarmCPU | SIG |
| PH_17JY | AX-94749804 | 2B | 762518824 | -0.0183 | 4.0247 | 9.25E-17 | IIIVmrMLM | SIG |
| PH_17JY | AX-109422526 | 2D | 35022365 | 0.9567 | 12.3067 | 5.02E-15 | IIIVmrMLM | SIG |
| PH_17JY | AX-110235512 | 2D | 129981951 | 2.4048 | 3.9443 | 4.68E-24 | IIIVmrMLM | SIG |

| | | | | | | | | |
|---|---|---|---|---|---|---|---|---|
| PH_17JY | AX-108868100 | 3A | 24773177 | 2.6984 | | 1.64E-07 | BLUPmrMLM | SUG |
| PH_17JY | AX-109487960 | 3A | 51379419 | 1.7397 | | 2.64E-05 | BLUPmrMLM | SUG |
| PH_17JY | AX-108941426 | 3A | 404993065 | 16.3875 | | 8.03E-07 | FarmCPU | SUG |
| PH_17JY | AX-108972490 | 3A | 512601505 | 1.8688 | -0.4921 | 1.53E-14 | IIIVmrMLM | SIG |
| PH_17JY | AX-110003269 | 3A | 660423901 | -0.7496 | 3.2301 | 1.90E-09 | IIIVmrMLM | SIG |
| PH_17JY | AX-111064156 | 3A | 730876525 | -0.3742 | 2.6281 | 8.36E-06 | IIIVmrMLM | SUG |
| PH_17JY | AX-95191276 | 3A | 735891482 | 0.9160 | 3.8979 | 2.89E-14 | IIIVmrMLM | SIG |
| PH_17JY | AX-111130626 | 3B | 352937637 | 1.1216 | -5.1635 | 1.94E-09 | IIIVmrMLM | SIG |
| PH_17JY | AX-94936967 | 3B | 660703071 | 3.5476 | | 1.26E-07 | BLUPmrMLM | SUG |
| PH_17JY | AX-111528181 | 3B | 684040548 | -0.1861 | 33.4855 | 6.16E-09 | IIIVmrMLM | SIG |
| PH_17JY | AX-111563398 | 3B | 740713005 | 0.9580 | -3.3137 | 1.26E-08 | IIIVmrMLM | SIG |
| PH_17JY | AX-95109003 | 3D | 571493682 | 3.1578 | | 1.93E-07 | BLUPmrMLM | SUG |
| PH_17JY | AX-89706785 | 4A | 7412218 | 4.1932 | | 2.50E-11 | BLUPmrMLM | SIG |
| PH_17JY | AX-110394147 | 4A | 17797531 | 3.0697 | | 1.36E-10 | BLUPmrMLM | SIG |
| PH_17JY | AX-108999122 | 4A | 68459986 | -0.2680 | -7.6751 | 4.31E-13 | IIIVmrMLM | SIG |
| PH_17JY | AX-94698407 | 4A | 201759277 | 0.3770 | 2.5877 | 7.01E-21 | IIIVmrMLM | SIG |
| PH_17JY | AX-108772267 | 4A | 626170283 | 3.1049 | | 3.88E-08 | BLUPmrMLM | SIG |
| PH_17JY | AX-110489987 | 4A | 684386085 | -1.8374 | | 1.39E-05 | BLUPmrMLM | SUG |
| PH_17JY | AX-110402155 | 4A | 739152247 | 4.7331 | | 1.04E-06 | FarmCPU | SUG |
| PH_17JY | AX-89728715 | 4B | 12893649 | 0.0311 | 13.7029 | 7.74E-09 | IIIVmrMLM | SIG |
| PH_17JY | AX-111829807 | 4B | 603046843 | -0.2126 | 9.1245 | 1.42E-11 | IIIVmrMLM | SIG |
| PH_17JY | AX-111544338 | 4B | 641174612 | 1.7487 | | 5.85E-05 | BLUPmrMLM | SUG |
| PH_17JY | AX-111069203 | 4B | 643918661 | 1.0073 | 0.8546 | 5.75E-05 | IIIVmrMLM | SUG |
| PH_17JY | AX-89703298 | 4D | 16926596 | 7.7969 | | 1.14E-07 | EMMAX | SIG |
| PH_17JY | AX-89703298 | 4D | 16926596 | -7.9474 | | 2.06E-08 | GEMMA | SIG |
| PH_17JY | AX-109281717 | 4D | 17613651 | 8.0243 | | 5.52E-08 | EMMAX | SIG |
| PH_17JY | AX-109281717 | 4D | 17613651 | -8.1714 | | 9.37E-09 | GEMMA | SIG |
| PH_17JY | AX-109281717 | 4D | 17613651 | 2.1481 | -6.9929 | 1.22E-20 | IIIVmrMLM | SIG |
| PH_17JY | AX-109665328 | 4D | 18539670 | 7.2417 | | 6.38E-07 | EMMAX | SUG |
| PH_17JY | AX-109665328 | 4D | 18539670 | -7.4937 | | 8.75E-08 | GEMMA | SIG |
| PH_17JY | AX-108916749 | 4D | 19090268 | 6.4171 | | 4.64E-07 | EMMAX | SUG |
| PH_17JY | AX-108916749 | 4D | 19090268 | -6.5120 | | 1.23E-07 | GEMMA | SUG |
| PH_17JY | AX-89421921 | 4D | 19288898 | 7.9995 | | 4.98E-08 | EMMAX | SIG |
| PH_17JY | AX-89421921 | 4D | 19288898 | -8.1337 | | 9.01E-09 | GEMMA | SIG |
| PH_17JY | AX-89445201 | 4D | 19459579 | -6.2214 | | 1.20E-06 | GEMMA | SUG |
| PH_17JY | AX-109756732 | 4D | 253134678 | -0.3774 | 2.1611 | 7.55E-09 | IIIVmrMLM | SIG |
| PH_17JY | AX-95186747 | 4D | 446428878 | -51.8260 | | 6.53E-08 | EMMAX | SIG |
| PH_17JY | AX-109903760 | 5A | 30056842 | -8.5767 | | 1.94E-06 | GEMMA | SUG |
| PH_17JY | AX-111694389 | 5A | 37966512 | -0.1415 | 2.3628 | 1.23E-12 | IIIVmrMLM | SIG |
| PH_17JY | AX-110570771 | 5A | 83292772 | 2.1364 | 0.9413 | 1.94E-12 | IIIVmrMLM | SIG |
| PH_17JY | AX-110380328 | 5A | 272133446 | 4.4545 | 0.2218 | 7.70E-62 | IIIVmrMLM | SIG |
| PH_17JY | AX-109454024 | 5A | 488638025 | 6.2799 | | 2.69E-10 | FarmCPU | SIG |
| PH_17JY | AX-108870201 | 5A | 567895675 | -0.6724 | 15.9642 | 6.69E-10 | IIIVmrMLM | SIG |
| PH_17JY | AX-110360409 | 5A | 578215897 | 1.1413 | -5.1563 | 2.13E-18 | IIIVmrMLM | SIG |

| | | | | | | | | |
|---|---|---|---|---|---|---|---|---|
| PH_17JY | AX-108896801 | 5A | 704839430 | -4.9046 | | 4.24E-08 | BLUPmrMLM | SIG |
| PH_17JY | AX-110999817 | 5A | 708416688 | -0.1888 | -6.0677 | 3.78E-48 | IIIVmrMLM | SIG |
| PH_17JY | AX-111719711 | 5B | 34227773 | -1.7863 | | 1.70E-05 | BLUPmrMLM | SUG |
| PH_17JY | AX-111013696 | 5B | 410725022 | 2.9161 | | 4.13E-06 | BLUPmrMLM | SUG |
| PH_17JY | AX-109387489 | 5B | 411743978 | -6.1674 | | 7.90E-07 | EMMAX | SUG |
| PH_17JY | AX-109387489 | 5B | 411743978 | 6.0716 | | 4.17E-07 | GEMMA | SUG |
| PH_17JY | AX-111552930 | 5B | 517219107 | 1.9509 | -2.286 | 7.74E-17 | IIIVmrMLM | SIG |
| PH_17JY | AX-110018488 | 5B | 528535571 | -7.6018 | | 2.20E-06 | GEMMA | SUG |
| PH_17JY | AX-109463034 | 5B | 544994414 | -21.1160 | | 7.92E-07 | EMMAX | SUG |
| PH_17JY | AX-109463034 | 5B | 544994414 | 21.1531 | | 4.44E-07 | GEMMA | SUG |
| PH_17JY | AX-109975908 | 5B | 582902069 | -4.8505 | | 9.78E-07 | FarmCPU | SUG |
| PH_17JY | AX-110442272 | 5B | 605610876 | -3.5314 | -2.849 | 7.18E-45 | IIIVmrMLM | SIG |
| PH_17JY | AX-94573733 | 5B | 687394963 | -0.2573 | -16.7769 | 1.61E-18 | IIIVmrMLM | SIG |
| PH_17JY | AX-111712700 | 5D | 541942893 | -0.2249 | -7.1541 | 1.50E-21 | IIIVmrMLM | SIG |
| PH_17JY | AX-110431858 | 6A | 26973059 | 3.6207 | | 3.84E-14 | BLUPmrMLM | SIG |
| PH_17JY | AX-109422357 | 6A | 341727858 | -1.5617 | 2.6605 | 1.91E-11 | IIIVmrMLM | SIG |
| PH_17JY | AX-108853441 | 6B | 197242252 | 2.9131 | | 7.58E-07 | BLUPmrMLM | SUG |
| PH_17JY | AX-108972994 | 6B | 292819957 | -0.4069 | -6.8168 | 8.14E-05 | IIIVmrMLM | SUG |
| PH_17JY | AX-109932835 | 6B | 492900231 | -0.2091 | -3.6905 | 3.85E-10 | IIIVmrMLM | SIG |
| PH_17JY | AX-110129520 | 6B | 529056319 | 1.7667 | | 1.71E-04 | BLUPmrMLM | SUG |
| PH_17JY | AX-112288906 | 6B | 688607609 | 1.8780 | | 1.71E-06 | BLUPmrMLM | SUG |
| PH_17JY | AX-109364183 | 6B | 694461959 | -10.6108 | | 2.21E-08 | FarmCPU | SIG |
| PH_17JY | AX-111776691 | 7A | 47499427 | 2.1032 | | 2.11E-07 | BLUPmrMLM | SUG |
| PH_17JY | AX-110475510 | 7A | 204182749 | -9.8235 | | 1.93E-06 | FarmCPU | SUG |
| PH_17JY | AX-111043307 | 7A | 234554138 | -1.5333 | -9.0476 | 8.58E-15 | IIIVmrMLM | SIG |
| PH_17JY | AX-110499112 | 7A | 448484396 | 0.6176 | 3.5786 | 1.88E-09 | IIIVmrMLM | SIG |
| PH_17JY | AX-109980760 | 7A | 456667106 | -7.8881 | | 3.89E-07 | FarmCPU | SUG |
| PH_17JY | AX-110991488 | 7A | 734924694 | -0.1432 | 14.1948 | 3.38E-20 | IIIVmrMLM | SIG |
| PH_17JY | AX-109841665 | 7B | 135522860 | -1.0588 | -6.6817 | 4.58E-10 | IIIVmrMLM | SIG |
| PH_17JY | AX-109347055 | 7B | 160613546 | -0.3808 | -5.4301 | 9.37E-13 | IIIVmrMLM | SIG |
| PH_17JY | AX-109993291 | 7B | 335788474 | 1.9420 | 2.5006 | 1.07E-16 | IIIVmrMLM | SIG |
| PH_17JY | AX-108726644 | 7B | 607426744 | 0.2389 | 6.1508 | 4.12E-09 | IIIVmrMLM | SIG |
| PH_17JY | AX-109167420 | 7D | 72299265 | -3.6940 | | 1.88E-08 | BLUPmrMLM | SIG |
| PH_17JY | AX-94510310 | 7D | 504831110 | -20.9804 | | 7.25E-07 | FarmCPU | SUG |
| PH_17ZMD | AX-109538413 | 1A | 53023601 | 2.4207 | | 1.07E-04 | BLUPmrMLM | SUG |
| PH_17ZMD | AX-111635819 | 1A | 351890717 | -11.8075 | | 4.72E-07 | GEMMA | SUG |
| PH_17ZMD | AX-110594323 | 1A | 351890921 | -11.8075 | | 4.72E-07 | GEMMA | SUG |
| PH_17ZMD | AX-109839041 | 1A | 351966228 | -11.8075 | | 4.72E-07 | GEMMA | SUG |
| PH_17ZMD | AX-111215880 | 1A | 351966755 | -11.8075 | | 4.72E-07 | GEMMA | SUG |
| PH_17ZMD | AX-109061216 | 1A | 351967379 | -11.8075 | | 4.72E-07 | GEMMA | SUG |
| PH_17ZMD | AX-109845602 | 1A | 352133191 | -11.2588 | | 1.08E-06 | GEMMA | SUG |
| PH_17ZMD | AX-108746718 | 1A | 352141947 | -11.1620 | | 1.92E-06 | GEMMA | SUG |
| PH_17ZMD | AX-109939645 | 1A | 352142757 | -11.8075 | | 4.72E-07 | GEMMA | SUG |
| PH_17ZMD | AX-110034268 | 1A | 352143071 | -11.1620 | | 1.92E-06 | GEMMA | SUG |

| | | | | | | | | |
|---|---|---|---|---|---|---|---|---|
| PH_17ZMD | AX-108885180 | 1A | 352171771 | -11.8075 | | 4.72E-07 | GEMMA | SUG |
| PH_17ZMD | AX-111509016 | 1A | 352852530 | -11.8075 | | 4.72E-07 | GEMMA | SUG |
| PH_17ZMD | AX-110983499 | 1A | 352858301 | -11.3221 | | 8.34E-07 | GEMMA | SUG |
| PH_17ZMD | AX-111469642 | 1A | 352862364 | -11.8075 | | 4.72E-07 | GEMMA | SUG |
| PH_17ZMD | AX-109360835 | 1A | 370002467 | 2.1025 | 19.5923 | 1.29E-28 | IIIVmrMLM | SIG |
| PH_17ZMD | AX-109379218 | 1A | 388947077 | -11.4677 | | 2.08E-06 | GEMMA | SUG |
| PH_17ZMD | AX-111819445 | 1A | 388947113 | -11.4677 | | 2.08E-06 | GEMMA | SUG |
| PH_17ZMD | AX-108742536 | 1A | 389029091 | -11.4677 | | 2.08E-06 | GEMMA | SUG |
| PH_17ZMD | AX-110403537 | 1A | 389061020 | -11.4677 | | 2.08E-06 | GEMMA | SUG |
| PH_17ZMD | AX-110448433 | 1A | 389061946 | -22.9213 | | 2.11E-06 | GEMMA | SUG |
| PH_17ZMD | AX-111106333 | 1A | 389200944 | 11.5420 | | 2.30E-06 | EMMAX | SUG |
| PH_17ZMD | AX-111106333 | 1A | 389200944 | -11.2822 | | 1.19E-06 | GEMMA | SUG |
| PH_17ZMD | AX-108850470 | 1A | 389228613 | -11.4607 | | 2.11E-06 | GEMMA | SUG |
| PH_17ZMD | AX-110492346 | 1A | 389252927 | -11.4607 | | 2.11E-06 | GEMMA | SUG |
| PH_17ZMD | AX-109509868 | 1A | 389363391 | -11.4607 | | 2.11E-06 | GEMMA | SUG |
| PH_17ZMD | AX-110628967 | 1A | 389803509 | -11.4607 | | 2.11E-06 | GEMMA | SUG |
| PH_17ZMD | AX-110629172 | 1A | 403654793 | 2.0289 | 4.3487 | 1.69E-14 | IIIVmrMLM | SIG |
| PH_17ZMD | AX-110530998 | 1A | 508146494 | -4.3144 | -6.3343 | 7.97E-54 | IIIVmrMLM | SIG |
| PH_17ZMD | AX-94934987 | 1A | 534043666 | -54.0600 | | 5.52E-08 | EMMAX | SIG |
| PH_17ZMD | AX-108966467 | 1B | 35973726 | 3.0244 | -3.4736 | 2.32E-30 | IIIVmrMLM | SIG |
| PH_17ZMD | AX-109372658 | 1B | 41084324 | 0.2544 | 6.2286 | 1.24E-12 | IIIVmrMLM | SIG |
| PH_17ZMD | AX-94819044 | 1B | 209521689 | -4.8343 | | 1.58E-10 | BLUPmrMLM | SIG |
| PH_17ZMD | AX-108775679 | 1B | 377830921 | 2.7338 | | 3.85E-07 | BLUPmrMLM | SUG |
| PH_17ZMD | AX-95160816 | 1D | 9765802 | -2.4312 | | 5.20E-06 | BLUPmrMLM | SUG |
| PH_17ZMD | AX-94420233 | 1D | 155927416 | -54.0600 | | 5.52E-08 | EMMAX | SIG |
| PH_17ZMD | AX-111345949 | 1D | 253194257 | -1.4893 | 3.2652 | 3.06E-09 | IIIVmrMLM | SIG |
| PH_17ZMD | AX-110526102 | 2A | 186397106 | -7.6260 | | 1.98E-06 | FarmCPU | SUG |
| PH_17ZMD | AX-110456050 | 2A | 751211338 | -3.4449 | | 1.79E-08 | BLUPmrMLM | SIG |
| PH_17ZMD | AX-110910161 | 2B | 116213727 | -7.8729 | | 1.63E-07 | FarmCPU | SUG |
| PH_17ZMD | AX-110556938 | 2B | 599853018 | 0.1078 | -3.5156 | 1.09E-04 | IIIVmrMLM | SUG |
| PH_17ZMD | AX-110393538 | 2B | 647706241 | -0.0155 | 12.9355 | 2.04E-09 | IIIVmrMLM | SIG |
| PH_17ZMD | AX-110035685 | 2B | 736246538 | -0.1677 | -9.5075 | 1.17E-12 | IIIVmrMLM | SIG |
| PH_17ZMD | AX-109420136 | 2B | 776411192 | -2.4574 | | 1.84E-07 | BLUPmrMLM | SUG |
| PH_17ZMD | AX-109899210 | 2B | 785883839 | 0.2490 | -8.9412 | 3.44E-13 | IIIVmrMLM | SIG |
| PH_17ZMD | AX-111218213 | 2D | 20190462 | -0.1581 | 4.5123 | 9.05E-32 | IIIVmrMLM | SIG |
| PH_17ZMD | AX-95209398 | 2D | 405408295 | -45.5220 | | 9.93E-08 | EMMAX | SIG |
| PH_17ZMD | AX-94834922 | 2D | 419030489 | -0.9739 | 8.0615 | 2.75E-07 | IIIVmrMLM | SUG |
| PH_17ZMD | AX-94427799 | 2D | 523357562 | -9.8209 | | 7.29E-07 | EMMAX | SUG |
| PH_17ZMD | AX-94427799 | 2D | 523357562 | 9.7684 | | 5.69E-07 | GEMMA | SUG |
| PH_17ZMD | AX-111336862 | 2D | 629367397 | -4.0927 | | 2.53E-10 | BLUPmrMLM | SIG |
| PH_17ZMD | AX-111336862 | 2D | 629367397 | -3.9473 | -1.2657 | 2.53E-44 | IIIVmrMLM | SIG |
| PH_17ZMD | AX-110388901 | 3A | 502915293 | 0.2894 | 21.3163 | 6.87E-09 | IIIVmrMLM | SIG |
| PH_17ZMD | AX-89475213 | 3A | 511995321 | 2.2867 | | 1.78E-06 | BLUPmrMLM | SUG |
| PH_17ZMD | AX-109361997 | 3A | 655338013 | -2.6509 | | 6.56E-07 | BLUPmrMLM | SUG |

| | | | | | | | | |
|---|---|---|---|---|---|---|---|---|
| PH_17ZMD | AX-111483830 | 3A | 718947465 | -0.0588 | 9.2731 | 9.34E-07 | IIIVmrMLM | SUG |
| PH_17ZMD | AX-111737513 | 3B | 126920865 | 0.7003 | 6.2381 | 2.37E-17 | IIIVmrMLM | SIG |
| PH_17ZMD | AX-108941515 | 3B | 469534228 | 3.8616 | | 3.45E-12 | BLUPmrMLM | SIG |
| PH_17ZMD | AX-108941515 | 3B | 469534228 | -1.9467 | -2.6461 | 1.55E-14 | IIIVmrMLM | SIG |
| PH_17ZMD | AX-108839193 | 3B | 562784942 | -1.8696 | | 2.55E-05 | BLUPmrMLM | SUG |
| PH_17ZMD | AX-110149046 | 3B | 643275270 | -3.4452 | | 2.52E-07 | BLUPmrMLM | SUG |
| PH_17ZMD | AX-111673406 | 3B | 760046216 | 1.2223 | 4.6376 | 1.98E-17 | IIIVmrMLM | SIG |
| PH_17ZMD | AX-109354595 | 3D | 459785395 | -1.9000 | | 8.97E-06 | BLUPmrMLM | SUG |
| PH_17ZMD | AX-95005610 | 3D | 546331575 | -37.5130 | | 2.13E-06 | EMMAX | SUG |
| PH_17ZMD | AX-94653465 | 3D | 555796772 | 2.9655 | | 3.25E-29 | IIIVmrMLM | SIG |
| PH_17ZMD | AX-111204246 | 3D | 571500279 | -2.3865 | -1.5579 | 7.48E-18 | IIIVmrMLM | SIG |
| PH_17ZMD | AX-111585637 | 4A | 2960053 | 2.0786 | | 8.86E-05 | BLUPmrMLM | SUG |
| PH_17ZMD | AX-94495955 | 4A | 37033089 | -18.2574 | | 2.18E-06 | FarmCPU | SUG |
| PH_17ZMD | AX-109500876 | 4A | 109106855 | 2.4467 | | 4.89E-05 | BLUPmrMLM | SUG |
| PH_17ZMD | AX-109380181 | 4B | 25421015 | 2.3258 | 11.353 | 3.84E-20 | IIIVmrMLM | SIG |
| PH_17ZMD | AX-108831701 | 4B | 191790892 | -2.0876 | | 3.55E-06 | BLUPmrMLM | SUG |
| PH_17ZMD | AX-108831701 | 4B | 191790892 | 0.2392 | -2.8483 | 1.17E-08 | IIIVmrMLM | SIG |
| PH_17ZMD | AX-108853535 | 4B | 411919572 | -1.5627 | | 1.31E-04 | BLUPmrMLM | SUG |
| PH_17ZMD | AX-109371081 | 4B | 527611179 | 0.5840 | -4.716 | 1.15E-10 | IIIVmrMLM | SIG |
| PH_17ZMD | AX-89593703 | 4D | 15958106 | -6.4138 | | 2.22E-06 | GEMMA | SUG |
| PH_17ZMD | AX-89703298 | 4D | 16926596 | 7.8128 | | 6.42E-07 | EMMAX | SUG |
| PH_17ZMD | AX-89703298 | 4D | 16926596 | -7.9817 | | 1.20E-07 | GEMMA | SUG |
| PH_17ZMD | AX-109281717 | 4D | 17613651 | 7.9760 | | 5.14E-07 | EMMAX | SUG |
| PH_17ZMD | AX-109281717 | 4D | 17613651 | -8.1702 | | 8.28E-08 | GEMMA | SIG |
| PH_17ZMD | AX-109665328 | 4D | 18539670 | 7.7125 | | 8.10E-07 | EMMAX | SUG |
| PH_17ZMD | AX-109665328 | 4D | 18539670 | -7.9690 | | 1.12E-07 | GEMMA | SIG |
| PH_17ZMD | AX-108916749 | 4D | 19090268 | 7.0900 | | 1.74E-07 | EMMAX | SUG |
| PH_17ZMD | AX-108916749 | 4D | 19090268 | -7.1749 | | 4.50E-08 | GEMMA | SIG |
| PH_17ZMD | AX-89421921 | 4D | 19288898 | 8.1120 | | 2.99E-07 | EMMAX | SUG |
| PH_17ZMD | AX-89421921 | 4D | 19288898 | -8.2917 | | 4.76E-08 | GEMMA | SIG |
| PH_17ZMD | AX-89421921 | 4D | 19288898 | 2.9544 | -7.4757 | 2.48E-28 | IIIVmrMLM | SIG |
| PH_17ZMD | AX-89445201 | 4D | 19459579 | 6.9371 | | 7.12E-07 | EMMAX | SUG |
| PH_17ZMD | AX-89445201 | 4D | 19459579 | -7.1510 | | 1.12E-07 | GEMMA | SIG |
| PH_17ZMD | AX-94484740 | 4D | 57072762 | -16.1997 | | 1.45E-09 | BLUPmrMLM | SIG |
| PH_17ZMD | AX-111416995 | 4D | 334598743 | -64.1980 | | 3.65E-07 | EMMAX | SUG |
| PH_17ZMD | AX-95186747 | 4D | 446428878 | -50.3190 | | 7.70E-07 | EMMAX | SUG |
| PH_17ZMD | AX-110622186 | 4D | 509433509 | -4.7374 | | 7.22E-09 | BLUPmrMLM | SIG |
| PH_17ZMD | AX-111122310 | 5A | 511619430 | -1.1370 | -0.3799 | 8.24E-05 | IIIVmrMLM | SUG |
| PH_17ZMD | AX-111487734 | 5A | 521724613 | 2.5295 | | 3.59E-07 | BLUPmrMLM | SUG |
| PH_17ZMD | AX-111036679 | 5A | 549208939 | 1.7758 | | 1.60E-04 | BLUPmrMLM | SUG |
| PH_17ZMD | AX-111077995 | 5A | 583590027 | -1.5203 | | 6.65E-05 | BLUPmrMLM | SUG |
| PH_17ZMD | AX-110124007 | 5A | 659400260 | -15.2077 | | 1.13E-06 | GEMMA | SUG |
| PH_17ZMD | AX-94752094 | 5A | 673539685 | 3.2709 | | 1.53E-06 | BLUPmrMLM | SUG |
| PH_17ZMD | AX-109271018 | 5A | 706243655 | 2.8750 | 0.4306 | 1.53E-22 | IIIVmrMLM | SIG |

| | | | | | | | | |
|---|---|---|---|---|---|---|---|---|
| PH_17ZMD | AX-95151481 | 5A | 706427281 | 4.7758 | | 3.14E-06 | BLUPmrMLM | SUG |
| PH_17ZMD | AX-94928671 | 5B | 383510019 | -2.2984 | | 1.22E-05 | BLUPmrMLM | SUG |
| PH_17ZMD | AX-108899168 | 5B | 410604227 | 0.2893 | 19.6436 | 4.14E-14 | IIIVmrMLM | SIG |
| PH_17ZMD | AX-109394866 | 5B | 513849166 | 1.7593 | -0.6954 | 1.39E-08 | IIIVmrMLM | SIG |
| PH_17ZMD | AX-111036555 | 5B | 513978487 | -7.8028 | | 1.33E-06 | EMMAX | SUG |
| PH_17ZMD | AX-111036555 | 5B | 513978487 | 7.8392 | | 6.80E-07 | GEMMA | SUG |
| PH_17ZMD | AX-109325020 | 5B | 517196591 | 5.4699 | 4.4312 | 7.49E-76 | IIIVmrMLM | SIG |
| PH_17ZMD | AX-111512697 | 5B | 527774551 | 8.4743 | | 2.00E-06 | EMMAX | SUG |
| PH_17ZMD | AX-111512697 | 5B | 527774551 | -8.6328 | | 4.96E-07 | GEMMA | SUG |
| PH_17ZMD | AX-110018488 | 5B | 528535571 | 9.2838 | | 8.19E-08 | EMMAX | SIG |
| PH_17ZMD | AX-110018488 | 5B | 528535571 | -9.6231 | | 6.98E-09 | GEMMA | SIG |
| PH_17ZMD | AX-111147486 | 5B | 528700469 | -7.7223 | | 7.54E-07 | GEMMA | SUG |
| PH_17ZMD | AX-108833640 | 5B | 528701508 | -7.3210 | | 1.40E-06 | GEMMA | SUG |
| PH_17ZMD | AX-110969237 | 5B | 529044398 | -7.7184 | | 7.51E-07 | GEMMA | SUG |
| PH_17ZMD | AX-111157103 | 5B | 583289315 | 2.6773 | | 5.37E-08 | BLUPmrMLM | SIG |
| PH_17ZMD | AX-111571815 | 5B | 595471198 | -1.9604 | | 1.83E-04 | BLUPmrMLM | SUG |
| PH_17ZMD | AX-109911942 | 5B | 692397094 | 0.0673 | -3.3773 | 1.10E-05 | IIIVmrMLM | SUG |
| PH_17ZMD | AX-94728034 | 5D | 434481168 | -73.2097 | -76.2953 | 0.00E+00 | IIIVmrMLM | SIG |
| PH_17ZMD | AX-110724909 | 6A | 59732085 | -3.0913 | | 2.43E-31 | IIIVmrMLM | SIG |
| PH_17ZMD | AX-112285858 | 6A | 84968797 | 2.0105 | | 1.61E-05 | BLUPmrMLM | SUG |
| PH_17ZMD | AX-95162142 | 6B | 199761147 | -26.8100 | | 2.35E-07 | EMMAX | SUG |
| PH_17ZMD | AX-94746925 | 6B | 577363493 | -64.1980 | | 3.65E-07 | EMMAX | SUG |
| PH_17ZMD | AX-111529494 | 6B | 711629196 | 0.5164 | -3.6329 | 3.77E-08 | IIIVmrMLM | SIG |
| PH_17ZMD | AX-111211808 | 6D | 5913263 | -0.7228 | -5.3241 | 3.32E-17 | IIIVmrMLM | SIG |
| PH_17ZMD | AX-94809200 | 6D | 15807973 | 2.6263 | -4.1496 | 4.33E-24 | IIIVmrMLM | SIG |
| PH_17ZMD | AX-95007223 | 6D | 36801236 | 4.5084 | | 1.14E-07 | BLUPmrMLM | SIG |
| PH_17ZMD | AX-89661736 | 6D | 86551098 | 1.1375 | 1.6274 | 3.41E-05 | IIIVmrMLM | SUG |
| PH_17ZMD | AX-94460648 | 6D | 462355369 | 1.6358 | | 8.53E-06 | BLUPmrMLM | SUG |
| PH_17ZMD | AX-94907690 | 6D | 471004496 | 2.9106 | | 2.57E-05 | BLUPmrMLM | SUG |
| PH_17ZMD | AX-109099689 | 7A | 167956145 | 0.0488 | -8.649 | 1.09E-07 | IIIVmrMLM | SIG |
| PH_17ZMD | AX-111759717 | 7A | 198671663 | 7.4762 | | 1.83E-06 | FarmCPU | SUG |
| PH_17ZMD | AX-111050220 | 7A | 256816439 | 2.8929 | | 1.53E-07 | BLUPmrMLM | SUG |
| PH_17ZMD | AX-95124326 | 7A | 505017486 | -64.1980 | | 3.65E-07 | EMMAX | SUG |
| PH_17ZMD | AX-111537467 | 7A | 581846079 | 1.5258 | | 1.90E-09 | IIIVmrMLM | SIG |
| PH_17ZMD | AX-110531923 | 7A | 620187306 | 13.8910 | | 1.86E-06 | GEMMA | SUG |
| PH_17ZMD | AX-109967948 | 7A | 636201281 | 0.1253 | 10.3178 | 3.51E-15 | IIIVmrMLM | SIG |
| PH_17ZMD | AX-110443657 | 7A | 666987276 | 0.1990 | -3.0882 | 1.47E-08 | IIIVmrMLM | SIG |
| PH_17ZMD | AX-108735789 | 7A | 714598577 | -0.0155 | 35.4418 | 1.18E-27 | IIIVmrMLM | SIG |
| PH_17ZMD | AX-110395012 | 7B | 66499791 | -0.8112 | -5.0774 | 1.52E-19 | IIIVmrMLM | SIG |
| PH_17ZMD | AX-95088118 | 7B | 94994860 | 3.0106 | | 1.75E-06 | BLUPmrMLM | SUG |
| PH_17ZMD | AX-94509715 | 7B | 445668577 | -64.1980 | | 3.65E-07 | EMMAX | SUG |
| PH_17ZMD | AX-94699640 | 7B | 516802754 | 3.0189 | | 4.70E-08 | BLUPmrMLM | SIG |
| PH_17ZMD | AX-111473859 | 7B | 609406824 | 2.0292 | | 1.16E-04 | BLUPmrMLM | SUG |

| | | | | | | | | |
|---|---|---|---|---|---|---|---|---|
| PH_17ZMD | AX-95226642 | 7B | 701188341 | -8.6151 | -19.4835 | 6.16E-147 | IIIVmrMLM | SIG |
| PH_17ZMD | AX-111091808 | 7D | 309028081 | 1.5024 | -12.8318 | 2.72E-16 | IIIVmrMLM | SIG |
| PH_17ZMD | AX-110440288 | 7D | 623527823 | -1.8966 | | 6.23E-05 | BLUPmrMLM | SUG |
| PH_18AY | AX-109832236 | 1A | 15375351 | 5.0512 | | 1.55E-07 | FarmCPU | SUG |
| PH_18AY | AX-109337531 | 1A | 51225450 | -0.3744 | -4.9756 | 4.53E-09 | IIIVmrMLM | SIG |
| PH_18AY | AX-108901390 | 1A | 546225444 | 7.1657 | | 1.20E-07 | FarmCPU | SUG |
| PH_18AY | AX-111655094 | 1A | 553066811 | 1.8962 | | 1.39E-07 | BLUPmrMLM | SUG |
| PH_18AY | AX-109406736 | 1A | 569381386 | -1.2964 | | 6.07E-10 | IIIVmrMLM | SIG |
| PH_18AY | AX-95095232 | 1B | 2056936 | 7.4013 | | 7.39E-10 | FarmCPU | SIG |
| PH_18AY | AX-108864425 | 1B | 14450964 | 0.3656 | -5.4567 | 3.60E-20 | IIIVmrMLM | SIG |
| PH_18AY | AX-110920573 | 1B | 26284397 | -1.8833 | | 3.66E-10 | BLUPmrMLM | SIG |
| PH_18AY | AX-109522858 | 1B | 65774858 | 1.5864 | | 5.67E-14 | IIIVmrMLM | SIG |
| PH_18AY | AX-111828510 | 1B | 548979604 | -4.5573 | | 1.64E-06 | FarmCPU | SUG |
| PH_18AY | AX-111761207 | 1B | 586110156 | 2.0449 | | 1.75E-04 | BLUPmrMLM | SUG |
| PH_18AY | AX-109006966 | 1B | 588148040 | -1.7621 | | 2.34E-07 | BLUPmrMLM | SUG |
| PH_18AY | AX-109821280 | 1B | 593812877 | -0.2013 | -13.6663 | 2.18E-14 | IIIVmrMLM | SIG |
| PH_18AY | AX-110369260 | 1B | 636032921 | -1.4577 | -3.0291 | 8.18E-12 | IIIVmrMLM | SIG |
| PH_18AY | AX-111522318 | 1D | 46395622 | 1.8265 | | 3.80E-07 | BLUPmrMLM | SUG |
| PH_18AY | AX-111482260 | 1D | 243707050 | 0.1983 | 9.1501 | 1.88E-06 | IIIVmrMLM | SUG |
| PH_18AY | AX-94665179 | 1D | 470896292 | 18.2413 | | 6.01E-07 | FarmCPU | SUG |
| PH_18AY | AX-111048393 | 2A | 75655049 | -1.8308 | | 7.95E-05 | BLUPmrMLM | SUG |
| PH_18AY | AX-110123869 | 2A | 284265347 | 0.4277 | 9.9049 | 1.98E-13 | IIIVmrMLM | SIG |
| PH_18AY | AX-110611858 | 2A | 568994034 | 1.5051 | | 2.07E-05 | BLUPmrMLM | SUG |
| PH_18AY | AX-111177381 | 2A | 717170801 | 1.0994 | | 4.28E-05 | BLUPmrMLM | SUG |
| PH_18AY | AX-108928895 | 2A | 728029569 | 0.1236 | 2.7514 | 1.46E-05 | IIIVmrMLM | SUG |
| PH_18AY | AX-109072035 | 2A | 728356544 | -1.2088 | | 7.37E-09 | IIIVmrMLM | SIG |
| PH_18AY | AX-109367336 | 2B | 29253872 | -1.5135 | 2.5177 | 1.20E-17 | IIIVmrMLM | SIG |
| PH_18AY | AX-110925634 | 2B | 30496700 | -4.4371 | | 1.29E-06 | GEMMA | SUG |
| PH_18AY | AX-109281639 | 2B | 30549242 | 4.5883 | | 1.79E-06 | EMMAX | SUG |
| PH_18AY | AX-109281639 | 2B | 30549242 | -4.6341 | | 9.19E-07 | GEMMA | SUG |
| PH_18AY | AX-94625058 | 2B | 742524167 | -2.3979 | | 1.08E-09 | BLUPmrMLM | SIG |
| PH_18AY | AX-94463704 | 2D | 16909807 | -9.2054 | | 1.67E-07 | EMMAX | SUG |
| PH_18AY | AX-94463704 | 2D | 16909807 | 9.3500 | | 5.30E-08 | GEMMA | SIG |
| PH_18AY | AX-95235724 | 2D | 16909856 | 8.1581 | | 1.06E-06 | GEMMA | SUG |
| PH_18AY | AX-94620138 | 2D | 16909870 | -9.3153 | | 2.00E-07 | EMMAX | SUG |
| PH_18AY | AX-94620138 | 2D | 16909870 | 9.4786 | | 6.70E-08 | GEMMA | SIG |
| PH_18AY | AX-94672317 | 2D | 16909999 | -8.3067 | | 6.76E-07 | EMMAX | SUG |
| PH_18AY | AX-94672317 | 2D | 16909999 | 8.3762 | | 2.98E-07 | GEMMA | SUG |
| PH_18AY | AX-95221455 | 2D | 16910027 | -8.2452 | | 6.05E-07 | EMMAX | SUG |
| PH_18AY | AX-95221455 | 2D | 16910027 | 8.4109 | | 2.18E-07 | GEMMA | SUG |
| PH_18AY | AX-109009107 | 2D | 16910108 | -8.2364 | | 1.82E-06 | EMMAX | SUG |
| PH_18AY | AX-109009107 | 2D | 16910108 | 8.3776 | | 7.14E-07 | GEMMA | SUG |
| PH_18AY | AX-110627083 | 2D | 17452055 | -0.9878 | 3.2915 | 3.14E-23 | IIIVmrMLM | SIG |

| | | | | | | | | |
|---|---|---|---|---|---|---|---|---|
| PH_18AY | AX-110467557 | 3A | 507462215 | -2.5268 | | 1.63E-04 | BLUPmrMLM | SUG |
| PH_18AY | AX-109835025 | 3A | 624764357 | 1.8757 | -0.6391 | 2.41E-14 | IIIVmrMLM | SIG |
| PH_18AY | AX-108871922 | 3A | 718861816 | -0.4978 | 3.4893 | 7.04E-12 | IIIVmrMLM | SIG |
| PH_18AY | AX-110431349 | 3A | 726696769 | -1.2281 | | 5.13E-05 | BLUPmrMLM | SUG |
| PH_18AY | AX-108778800 | 3A | 727635905 | 2.1520 | | 5.21E-08 | BLUPmrMLM | SIG |
| PH_18AY | AX-109967016 | 3A | 734470629 | -0.0908 | -3.4024 | 2.09E-06 | IIIVmrMLM | SUG |
| PH_18AY | AX-109387101 | 3B | 17916598 | 1.6165 | | 3.47E-06 | BLUPmrMLM | SUG |
| PH_18AY | AX-89699865 | 3B | 29727439 | -0.3242 | -3.9434 | 5.30E-14 | IIIVmrMLM | SIG |
| PH_18AY | AX-111023983 | 3B | 224470110 | -2.1512 | -0.5162 | 6.43E-15 | IIIVmrMLM | SIG |
| PH_18AY | AX-109383200 | 3B | 327649349 | -0.2648 | -2.4917 | 3.33E-06 | IIIVmrMLM | SUG |
| PH_18AY | AX-109372906 | 3B | 425534653 | -1.6791 | 7.0698 | 1.39E-17 | IIIVmrMLM | SIG |
| PH_18AY | AX-110984156 | 3B | 520333477 | -1.5430 | 1.3822 | 2.50E-12 | IIIVmrMLM | SIG |
| PH_18AY | AX-108749526 | 3B | 545045263 | 1.9971 | | 1.63E-04 | BLUPmrMLM | SUG |
| PH_18AY | AX-110560967 | 3B | 598860826 | -0.9116 | | 1.36E-04 | BLUPmrMLM | SUG |
| PH_18AY | AX-95089162 | 3B | 811446460 | -2.0126 | | 2.33E-07 | BLUPmrMLM | SUG |
| PH_18AY | AX-108883163 | 3B | 819506306 | -2.7832 | | 9.83E-12 | BLUPmrMLM | SIG |
| PH_18AY | AX-110523727 | 3D | 260181109 | 52.7260 | | 2.13E-06 | EMMAX | SUG |
| PH_18AY | AX-94907389 | 3D | 389191643 | -6.1541 | | 3.84E-17 | BLUPmrMLM | SIG |
| PH_18AY | AX-94907389 | 3D | 389191643 | 6.0227 | | 8.98E-07 | EMMAX | SUG |
| PH_18AY | AX-94907389 | 3D | 389191643 | -6.1699 | | 3.90E-07 | GEMMA | SUG |
| PH_18AY | AX-111551242 | 3D | 603273440 | 1.1853 | -0.7154 | 3.85E-07 | IIIVmrMLM | SUG |
| PH_18AY | AX-110562653 | 4A | 613407426 | -1.8863 | | 1.34E-04 | BLUPmrMLM | SUG |
| PH_18AY | AX-108772042 | 4A | 660646409 | -1.8276 | | 8.41E-07 | BLUPmrMLM | SUG |
| PH_18AY | AX-111041462 | 4A | 667316125 | 0.0687 | -2.0489 | 1.52E-07 | IIIVmrMLM | SUG |
| PH_18AY | AX-110429305 | 4A | 718875514 | 1.8719 | -0.8655 | 1.21E-13 | IIIVmrMLM | SIG |
| PH_18AY | AX-109923509 | 4A | 742832076 | -0.1807 | -4.0531 | 8.88E-16 | IIIVmrMLM | SIG |
| PH_18AY | AX-111523628 | 4B | 265309614 | 0.2899 | -7.263 | 2.20E-11 | IIIVmrMLM | SIG |
| PH_18AY | AX-111107034 | 4B | 392372140 | 4.1665 | | 5.22E-07 | EMMAX | SUG |
| PH_18AY | AX-111107034 | 4B | 392372140 | -4.1929 | | 2.88E-07 | GEMMA | SUG |
| PH_18AY | AX-110667361 | 4B | 618526394 | 0.1566 | 2.9803 | 6.53E-07 | IIIVmrMLM | SUG |
| PH_18AY | AX-89593703 | 4D | 15958106 | -4.9056 | | 1.76E-06 | GEMMA | SUG |
| PH_18AY | AX-89703298 | 4D | 16926596 | 6.3018 | | 9.17E-08 | EMMAX | SIG |
| PH_18AY | AX-89703298 | 4D | 16926596 | -6.5290 | | 9.02E-09 | GEMMA | SIG |
| PH_18AY | AX-109281717 | 4D | 17613651 | 6.5464 | | 4.23E-08 | EMMAX | SIG |
| PH_18AY | AX-109281717 | 4D | 17613651 | -6.7672 | | 3.81E-09 | GEMMA | SIG |
| PH_18AY | AX-109665328 | 4D | 18539670 | 6.0663 | | 2.28E-07 | EMMAX | SUG |
| PH_18AY | AX-109665328 | 4D | 18539670 | -6.3483 | | 2.00E-08 | GEMMA | SIG |
| PH_18AY | AX-108916749 | 4D | 19090268 | 5.1936 | | 3.59E-07 | EMMAX | SUG |
| PH_18AY | AX-108916749 | 4D | 19090268 | -5.3327 | | 6.47E-08 | GEMMA | SIG |
| PH_18AY | AX-108916749 | 4D | 19090268 | 4.1613 | 1.3154 | 4.82E-68 | IIIVmrMLM | SIG |
| PH_18AY | AX-89421921 | 4D | 19288898 | 6.5285 | | 3.76E-08 | EMMAX | SIG |
| PH_18AY | AX-89421921 | 4D | 19288898 | -6.7520 | | 3.30E-09 | GEMMA | SIG |
| PH_18AY | AX-89445201 | 4D | 19459579 | 5.0132 | | 1.99E-06 | EMMAX | SUG |
| PH_18AY | AX-89445201 | 4D | 19459579 | -5.2186 | | 2.96E-07 | GEMMA | SUG |

| | | | | | | | | |
|---|---|---|---|---|---|---|---|---|
| PH_18AY | AX-111970247 | 4D | 32225033 | 3.6486 | | 1.01E-06 | EMMAX | SUG |
| PH_18AY | AX-111970247 | 4D | 32225033 | -3.7095 | | 3.51E-07 | GEMMA | SUG |
| PH_18AY | AX-108860446 | 5A | 31100599 | -1.7060 | | 3.04E-05 | BLUPmrMLM | SUG |
| PH_18AY | AX-109454024 | 5A | 488638025 | 4.3624 | | 2.75E-08 | FarmCPU | SIG |
| PH_18AY | AX-109347539 | 5A | 538001845 | 0.0814 | -7.5008 | 1.14E-17 | IIIVmrMLM | SIG |
| PH_18AY | AX-110999817 | 5A | 708416688 | -0.0208 | -6.6745 | 5.58E-64 | IIIVmrMLM | SIG |
| PH_18AY | AX-110482793 | 5B | 506782550 | -4.5219 | | 3.18E-10 | BLUPmrMLM | SIG |
| PH_18AY | AX-110369376 | 5B | 507953336 | 3.8763 | 5.9225 | 2.19E-61 | IIIVmrMLM | SIG |
| PH_18AY | AX-109324011 | 5B | 536847515 | -0.3456 | 4.5555 | 4.20E-07 | IIIVmrMLM | SUG |
| PH_18AY | AX-110956454 | 5B | 612832259 | -1.2027 | | 1.52E-04 | BLUPmrMLM | SUG |
| PH_18AY | AX-110428229 | 5B | 662031427 | 1.9615 | | 7.80E-06 | BLUPmrMLM | SUG |
| PH_18AY | AX-94860490 | 5D | 555928841 | 4.7267 | 15.9719 | 7.10E-96 | IIIVmrMLM | SIG |
| PH_18AY | AX-111084482 | 6A | 2949306 | -2.1120 | | 1.38E-06 | BLUPmrMLM | SUG |
| PH_18AY | AX-110360902 | 6A | 587378637 | 0.9084 | | 1.83E-04 | BLUPmrMLM | SUG |
| PH_18AY | AX-110360902 | 6A | 587378637 | -1.4944 | 0.5016 | 8.13E-11 | IIIVmrMLM | SIG |
| PH_18AY | AX-110007903 | 6B | 22553595 | -1.6062 | | 1.48E-08 | BLUPmrMLM | SIG |
| PH_18AY | AX-108829072 | 6B | 44285007 | -1.2764 | | 7.73E-05 | BLUPmrMLM | SUG |
| PH_18AY | AX-109439156 | 6B | 488537393 | -1.7003 | | 5.66E-06 | BLUPmrMLM | SUG |
| PH_18AY | AX-110948737 | 6B | 582403139 | 1.4820 | | 1.12E-06 | BLUPmrMLM | SUG |
| PH_18AY | AX-110360201 | 6B | 712454322 | -0.6656 | -4.6102 | 1.82E-12 | IIIVmrMLM | SIG |
| PH_18AY | AX-109295742 | 6D | 419017728 | 0.3156 | -12.6597 | 1.36E-11 | IIIVmrMLM | SIG |
| PH_18AY | AX-109913379 | 6D | 469341593 | 2.8978 | | 5.54E-08 | BLUPmrMLM | SIG |
| PH_18AY | AX-109913379 | 6D | 469341593 | -1.3995 | 0.1373 | 7.47E-09 | IIIVmrMLM | SIG |
| PH_18AY | AX-110078977 | 7A | 48093982 | -2.3790 | | 3.30E-11 | BLUPmrMLM | SIG |
| PH_18AY | AX-110927892 | 7A | 65332331 | 0.0856 | 8.2279 | 2.34E-06 | IIIVmrMLM | SUG |
| PH_18AY | AX-109273777 | 7A | 201227626 | -4.4534 | | 7.07E-07 | FarmCPU | SUG |
| PH_18AY | AX-109412145 | 7A | 533552194 | 2.2839 | | 3.52E-07 | BLUPmrMLM | SUG |
| PH_18AY | AX-109880303 | 7A | 569120372 | -0.0594 | 4.7586 | 2.59E-13 | IIIVmrMLM | SIG |
| PH_18AY | AX-109620440 | 7A | 619621671 | 1.3878 | | 1.36E-04 | BLUPmrMLM | SUG |
| PH_18AY | AX-111524448 | 7A | 713176845 | -0.1915 | 4.6854 | 1.47E-04 | IIIVmrMLM | SUG |
| PH_18AY | AX-108883403 | 7B | 45629721 | 0.1365 | -3.1914 | 7.12E-04 | IIIVmrMLM | SUG |
| PH_18AY | AX-111152023 | 7B | 334196810 | -1.4744 | | 9.76E-07 | BLUPmrMLM | SUG |
| PH_18AY | AX-111575791 | 7B | 361027957 | -1.6772 | | 2.29E-15 | IIIVmrMLM | SIG |
| PH_18AY | AX-109938058 | 7B | 642748561 | 1.4007 | | 1.37E-05 | BLUPmrMLM | SUG |
| PH_18AY | AX-111635537 | 7B | 714298036 | -1.4953 | | 1.42E-06 | BLUPmrMLM | SUG |
| PH_18AY | AX-108886058 | 7D | 70057841 | 2.5709 | | 2.12E-06 | BLUPmrMLM | SUG |
| PH_18AY | AX-110522071 | 7D | 518743241 | 1.0185 | 3.8197 | 6.59E-07 | IIIVmrMLM | SUG |
| PH_18AY | AX-110357839 | 7D | 574869865 | 52.7260 | | 2.13E-06 | EMMAX | SUG |
| PH_18AY | AX-110908361 | 7D | 574876786 | 52.7260 | | 2.13E-06 | EMMAX | SUG |
| PH_18JY | AX-110073653 | 1A | 15344362 | -0.1775 | 19.1615 | 2.03E-18 | IIIVmrMLM | SIG |
| PH_18JY | AX-111634381 | 1A | 30131954 | -13.7740 | | 1.56E-06 | EMMAX | SUG |
| PH_18JY | AX-111634381 | 1A | 30131954 | 13.7852 | | 1.11E-06 | GEMMA | SUG |
| PH_18JY | AX-111039702 | 1A | 35545627 | -1.6959 | | 4.12E-05 | BLUPmrMLM | SUG |
| PH_18JY | AX-109032430 | 1A | 336134629 | -1.0230 | -5.8864 | 1.05E-13 | IIIVmrMLM | SIG |

| | | | | | | | | |
|---|---|---|---|---|---|---|---|---|
| PH_18JY | AX-109355416 | 1A | 494727689 | 5.5914 | | 1.98E-120 | IIIVmrMLM | SIG |
| PH_18JY | AX-109817102 | 1A | 495013803 | 8.9946 | | 2.08E-06 | EMMAX | SUG |
| PH_18JY | AX-109817102 | 1A | 495013803 | -9.0440 | | 8.26E-07 | GEMMA | SUG |
| PH_18JY | AX-111609168 | 1A | 495014994 | -8.6842 | | 1.38E-06 | GEMMA | SUG |
| PH_18JY | AX-110444939 | 1A | 495134348 | 7.8762 | | 2.18E-06 | EMMAX | SUG |
| PH_18JY | AX-110444939 | 1A | 495134348 | -7.9321 | | 1.09E-06 | GEMMA | SUG |
| PH_18JY | AX-109423357 | 1A | 516313166 | -2.6076 | | 1.66E-06 | BLUPmrMLM | SUG |
| PH_18JY | AX-94525724 | 1A | 564749741 | -29.9029 | | 7.11E-07 | FarmCPU | SUG |
| PH_18JY | AX-109985162 | 1B | 9177234 | -2.7079 | | 2.91E-06 | BLUPmrMLM | SUG |
| PH_18JY | AX-111217304 | 1B | 26260503 | -2.0967 | | 1.16E-09 | BLUPmrMLM | SIG |
| PH_18JY | AX-110612345 | 1B | 26631475 | 1.7176 | -0.5035 | 5.96E-15 | IIIVmrMLM | SIG |
| PH_18JY | AX-111026252 | 1B | 83660266 | 1.7374 | | 1.97E-04 | BLUPmrMLM | SUG |
| PH_18JY | AX-109299876 | 1B | 456999202 | 1.3985 | | 4.09E-05 | BLUPmrMLM | SUG |
| PH_18JY | AX-94795975 | 1B | 658482894 | 0.3019 | 38.4625 | 1.58E-15 | IIIVmrMLM | SIG |
| PH_18JY | AX-108873354 | 1B | 658536771 | 1.6644 | | 3.95E-07 | BLUPmrMLM | SUG |
| PH_18JY | AX-111157582 | 1B | 672880906 | 1.8579 | | 1.81E-04 | BLUPmrMLM | SUG |
| PH_18JY | AX-109079032 | 1D | 31059478 | -1.9412 | | 1.82E-06 | BLUPmrMLM | SUG |
| PH_18JY | AX-109372602 | 1D | 347285960 | 36.2500 | | 7.80E-08 | EMMAX | SIG |
| PH_18JY | AX-110733343 | 1D | 400880030 | 59.2950 | | 6.16E-12 | EMMAX | SIG |
| PH_18JY | AX-111030152 | 2A | 31820968 | -7.1254 | | 1.61E-06 | FarmCPU | SUG |
| PH_18JY | AX-111633619 | 2A | 496352754 | 11.4239 | | 2.28E-06 | GEMMA | SUG |
| PH_18JY | AX-111519971 | 2A | 692354010 | 1.2164 | | 1.02E-06 | BLUPmrMLM | SUG |
| PH_18JY | AX-111163674 | 2A | 703863086 | -1.6668 | | 2.30E-06 | BLUPmrMLM | SUG |
| PH_18JY | AX-111188192 | 2A | 734154299 | -15.0070 | | 1.70E-06 | EMMAX | SUG |
| PH_18JY | AX-111188192 | 2A | 734154299 | 14.9813 | | 2.05E-06 | GEMMA | SUG |
| PH_18JY | AX-94409599 | 2A | 734256911 | -15.0070 | | 1.70E-06 | EMMAX | SUG |
| PH_18JY | AX-94409599 | 2A | 734256911 | 14.9813 | | 2.05E-06 | GEMMA | SUG |
| PH_18JY | AX-110084699 | 2A | 734596092 | -15.4100 | | 1.39E-06 | EMMAX | SUG |
| PH_18JY | AX-110084699 | 2A | 734596092 | 15.3312 | | 1.72E-06 | GEMMA | SUG |
| PH_18JY | AX-109448328 | 2A | 768536052 | 0.5922 | 10.493 | 1.97E-12 | IIIVmrMLM | SIG |
| PH_18JY | AX-111111902 | 2B | 28415726 | 0.4231 | 3.9796 | 1.45E-10 | IIIVmrMLM | SIG |
| PH_18JY | AX-108748416 | 2B | 115909069 | 11.3292 | | 1.69E-08 | FarmCPU | SIG |
| PH_18JY | AX-110093244 | 2B | 150937460 | 0.1971 | 3.8026 | 1.48E-25 | IIIVmrMLM | SIG |
| PH_18JY | AX-94750463 | 2B | 524178912 | -2.9519 | | 6.87E-47 | IIIVmrMLM | SIG |
| PH_18JY | AX-111640532 | 2B | 682801798 | -0.6143 | 2.2886 | 1.28E-15 | IIIVmrMLM | SIG |
| PH_18JY | AX-110612651 | 2B | 703717563 | 3.5803 | | 1.40E-08 | BLUPmrMLM | SIG |
| PH_18JY | AX-110103130 | 2B | 767028694 | -1.4056 | | 9.47E-06 | BLUPmrMLM | SUG |
| PH_18JY | AX-94478750 | 2D | 136167994 | 10.9199 | | 2.12E-06 | GEMMA | SUG |
| PH_18JY | AX-94427799 | 2D | 523357562 | 3.3201 | | 1.47E-04 | BLUPmrMLM | SUG |
| PH_18JY | AX-109271432 | 3A | 20737483 | -1.1479 | | 1.36E-05 | BLUPmrMLM | SUG |
| PH_18JY | AX-94766803 | 3A | 26040630 | -0.2565 | -6.1811 | 1.86E-06 | IIIVmrMLM | SUG |
| PH_18JY | AX-110037431 | 3A | 32230885 | 1.7058 | | 5.26E-06 | BLUPmrMLM | SUG |
| PH_18JY | AX-94598560 | 3A | 90857824 | -12.9374 | | 1.59E-06 | FarmCPU | SUG |

| | | | | | | | | |
|---|---|---|---|---|---|---|---|---|
| PH_18JY | AX-109887132 | 3A | 128162161 | 0.0310 | 9.7225 | 4.77E-10 | IIIVmrMLM | SIG |
| PH_18JY | AX-109477152 | 3A | 433767107 | 3.6744 | 1.8457 | 3.48E-60 | IIIVmrMLM | SIG |
| PH_18JY | AX-110966197 | 3A | 486746660 | 1.4357 | | 1.83E-04 | BLUPmrMLM | SUG |
| PH_18JY | AX-110467557 | 3A | 507462215 | 5.9334 | | 3.99E-07 | EMMAX | SUG |
| PH_18JY | AX-110467557 | 3A | 507462215 | -5.9655 | | 2.20E-07 | GEMMA | SUG |
| PH_18JY | AX-111517150 | 3A | 537747224 | -1.0610 | -2.1492 | 6.87E-10 | IIIVmrMLM | SIG |
| PH_18JY | AX-95258822 | 3A | 586407724 | 59.2950 | | 6.16E-12 | EMMAX | SIG |
| PH_18JY | AX-95094987 | 3A | 625393271 | 8.2683 | | 7.54E-07 | EMMAX | SUG |
| PH_18JY | AX-95094987 | 3A | 625393271 | -8.4304 | | 5.27E-07 | GEMMA | SUG |
| PH_18JY | AX-110711989 | 3B | 269949176 | 10.3490 | | 4.70E-09 | EMMAX | SIG |
| PH_18JY | AX-110711989 | 3B | 269949176 | -10.3896 | | 5.63E-09 | GEMMA | SIG |
| PH_18JY | AX-109389434 | 3B | 303101605 | -11.0080 | | 5.88E-08 | EMMAX | SIG |
| PH_18JY | AX-109389434 | 3B | 303101605 | 11.1851 | | 3.97E-08 | GEMMA | SIG |
| PH_18JY | AX-111798171 | 3B | 499749960 | 0.6773 | -10.117 | 1.72E-09 | IIIVmrMLM | SIG |
| PH_18JY | AX-110107828 | 3B | 542511339 | -49.2490 | | 2.02E-06 | EMMAX | SUG |
| PH_18JY | AX-94486578 | 3B | 581636747 | -49.2490 | | 2.02E-06 | EMMAX | SUG |
| PH_18JY | AX-110494045 | 3B | 747550627 | -3.1149 | | 3.00E-13 | BLUPmrMLM | SIG |
| PH_18JY | AX-94777755 | 3D | 7972995 | -3.3200 | | 2.13E-05 | BLUPmrMLM | SUG |
| PH_18JY | AX-95109848 | 3D | 25124764 | 38.4610 | | 2.41E-08 | EMMAX | SIG |
| PH_18JY | AX-94549097 | 3D | 70712813 | 1.4888 | | 2.09E-05 | BLUPmrMLM | SUG |
| PH_18JY | AX-112289757 | 3D | 410520869 | -3.2533 | | 2.42E-05 | BLUPmrMLM | SUG |
| PH_18JY | AX-94832342 | 3D | 447990281 | -26.1290 | | 3.89E-07 | EMMAX | SUG |
| PH_18JY | AX-109932274 | 4A | 60712364 | -2.2319 | | 4.62E-06 | BLUPmrMLM | SUG |
| PH_18JY | AX-109392302 | 4A | 249562048 | 0.6880 | 4.2878 | 5.81E-07 | IIIVmrMLM | SUG |
| PH_18JY | AX-111717201 | 4A | 562402398 | -2.2448 | | 5.29E-06 | BLUPmrMLM | SUG |
| PH_18JY | AX-111717201 | 4A | 562402398 | 1.3735 | 0.3535 | 1.52E-10 | IIIVmrMLM | SIG |
| PH_18JY | AX-110562653 | 4A | 613407426 | -2.2063 | | 8.43E-05 | BLUPmrMLM | SUG |
| PH_18JY | AX-109380181 | 4B | 25421015 | 1.9824 | -2.4575 | 5.91E-23 | IIIVmrMLM | SIG |
| PH_18JY | AX-108944764 | 4D | 15911901 | 6.0780 | | 5.50E-07 | EMMAX | SUG |
| PH_18JY | AX-108944764 | 4D | 15911901 | -6.1198 | | 2.45E-07 | GEMMA | SUG |
| PH_18JY | AX-89593703 | 4D | 15958106 | 6.7754 | | 1.31E-08 | EMMAX | SIG |
| PH_18JY | AX-89593703 | 4D | 15958106 | -7.0571 | | 1.28E-09 | GEMMA | SIG |
| PH_18JY | AX-89703298 | 4D | 16926596 | -6.0009 | | 1.66E-15 | BLUPmrMLM | SIG |
| PH_18JY | AX-89703298 | 4D | 16926596 | 8.0983 | | 1.12E-09 | EMMAX | SIG |
| PH_18JY | AX-89703298 | 4D | 16926596 | -8.3538 | | 1.12E-10 | GEMMA | SIG |
| PH_18JY | AX-109281717 | 4D | 17613651 | 8.5397 | | 2.93E-10 | EMMAX | SIG |
| PH_18JY | AX-109281717 | 4D | 17613651 | -8.7447 | | 3.08E-11 | GEMMA | SIG |
| PH_18JY | AX-109665328 | 4D | 18539670 | 8.0123 | | 7.77E-10 | EMMAX | SIG |
| PH_18JY | AX-109665328 | 4D | 18539670 | -8.3576 | | 6.02E-11 | GEMMA | SIG |
| PH_18JY | AX-108916749 | 4D | 19090268 | 5.7130 | | 4.76E-07 | EMMAX | SUG |
| PH_18JY | AX-108916749 | 4D | 19090268 | -5.8221 | | 1.50E-07 | GEMMA | SUG |
| PH_18JY | AX-89421921 | 4D | 19288898 | 8.4953 | | 2.79E-10 | EMMAX | SIG |
| PH_18JY | AX-89421921 | 4D | 19288898 | -8.7254 | | 2.57E-11 | GEMMA | SIG |
| PH_18JY | AX-89421921 | 4D | 19288898 | 4.2858 | -4.9486 | 3.11E-82 | IIIVmrMLM | SIG |

| | | | | | | | | |
|---|---|---|---|---|---|---|---|---|
| PH_18JY | AX-89445201 | 4D | 19459579 | 5.7656 | | 7.55E-07 | EMMAX | SUG |
| PH_18JY | AX-89445201 | 4D | 19459579 | -5.9779 | | 1.43E-07 | GEMMA | SUG |
| PH_18JY | AX-94719027 | 4D | 37514818 | 36.3980 | | 3.18E-08 | EMMAX | SIG |
| PH_18JY | AX-109761244 | 4D | 132218554 | -36.7150 | | 4.00E-07 | EMMAX | SUG |
| PH_18JY | AX-108958305 | 4D | 198613145 | -49.2490 | | 2.02E-06 | EMMAX | SUG |
| PH_18JY | AX-110047101 | 4D | 224594707 | -61.9320 | | 2.43E-09 | EMMAX | SIG |
| PH_18JY | AX-108841393 | 4D | 229817746 | 31.3080 | | 8.37E-07 | EMMAX | SUG |
| PH_18JY | AX-95186747 | 4D | 446428878 | -47.6390 | | 9.56E-09 | EMMAX | SIG |
| PH_18JY | AX-110387369 | 4D | 467431300 | -41.5660 | | 7.68E-07 | EMMAX | SUG |
| PH_18JY | AX-109385490 | 5A | 26753680 | 2.3412 | -3.1779 | 2.38E-31 | IIIVmrMLM | SIG |
| PH_18JY | AX-109903760 | 5A | 30056842 | 7.8350 | | 9.27E-07 | EMMAX | SUG |
| PH_18JY | AX-109903760 | 5A | 30056842 | -7.8585 | | 5.20E-07 | GEMMA | SUG |
| PH_18JY | AX-108754051 | 5A | 30504616 | -3.7783 | | 1.40E-10 | BLUPmrMLM | SIG |
| PH_18JY | AX-94694183 | 5A | 109304236 | -10.9940 | | 1.15E-07 | FarmCPU | SIG |
| PH_18JY | AX-109880983 | 5A | 298010264 | -2.5173 | | 3.13E-05 | BLUPmrMLM | SUG |
| PH_18JY | AX-111687675 | 5A | 395679502 | -1.0641 | -3.0981 | 8.66E-10 | IIIVmrMLM | SIG |
| PH_18JY | AX-95122272 | 5A | 398278397 | 42.1250 | | 1.50E-08 | EMMAX | SIG |
| PH_18JY | AX-108943812 | 5A | 503632674 | -2.1701 | | 1.01E-08 | BLUPmrMLM | SIG |
| PH_18JY | AX-111531124 | 5A | 508333653 | -0.5824 | -5.3453 | 8.28E-08 | IIIVmrMLM | SIG |
| PH_18JY | AX-110963728 | 5A | 693675632 | -3.1003 | | 5.05E-07 | BLUPmrMLM | SUG |
| PH_18JY | AX-110014427 | 5A | 697488076 | 0.7840 | -7.7191 | 9.76E-17 | IIIVmrMLM | SIG |
| PH_18JY | AX-110417678 | 5A | 701292911 | 12.7550 | | 2.73E-08 | EMMAX | SIG |
| PH_18JY | AX-110417678 | 5A | 701292911 | -12.9075 | | 2.18E-08 | GEMMA | SIG |
| PH_18JY | AX-108954010 | 5A | 702001853 | -0.3819 | 6.2187 | 1.79E-08 | IIIVmrMLM | SIG |
| PH_18JY | AX-94509069 | 5B | 211148106 | -2.5334 | 11.5594 | 4.10E-40 | IIIVmrMLM | SIG |
| PH_18JY | AX-94430767 | 5B | 243433888 | 2.6572 | 0.3989 | 4.34E-33 | IIIVmrMLM | SIG |
| PH_18JY | AX-109394686 | 5B | 406071467 | -3.0025 | 2.7383 | 7.13E-47 | IIIVmrMLM | SIG |
| PH_18JY | AX-110018488 | 5B | 528535571 | 4.8894 | | 1.16E-100 | IIIVmrMLM | SIG |
| PH_18JY | AX-108932221 | 5B | 536054310 | -1.6462 | | 2.94E-07 | BLUPmrMLM | SUG |
| PH_18JY | AX-110538089 | 5B | 539858157 | -1.6473 | | 7.41E-06 | BLUPmrMLM | SUG |
| PH_18JY | AX-110064454 | 5D | 2254066 | 32.8340 | | 1.20E-06 | EMMAX | SUG |
| PH_18JY | AX-110365570 | 5D | 285354818 | 24.1850 | | 1.50E-07 | EMMAX | SUG |
| PH_18JY | AX-110930042 | 5D | 362887848 | -44.2630 | | 5.24E-09 | EMMAX | SIG |
| PH_18JY | AX-94668949 | 5D | 454183213 | 35.0650 | | 1.44E-07 | EMMAX | SUG |
| PH_18JY | AX-94963508 | 5D | 484426813 | -61.9320 | | 2.43E-09 | EMMAX | SIG |
| PH_18JY | AX-109081370 | 5D | 493585976 | 42.1250 | | 1.50E-08 | EMMAX | SIG |
| PH_18JY | AX-108830877 | 5D | 541798506 | -0.8638 | -4.7068 | 1.74E-18 | IIIVmrMLM | SIG |
| PH_18JY | AX-94860490 | 5D | 555928841 | 2.4002 | 10.8216 | 3.23E-43 | IIIVmrMLM | SIG |
| PH_18JY | AX-109970551 | 6A | 330915971 | -1.6533 | | 1.08E-04 | BLUPmrMLM | SUG |
| PH_18JY | AX-108991161 | 6A | 610198092 | 2.2489 | | 8.80E-30 | IIIVmrMLM | SIG |
| PH_18JY | AX-110063373 | 6A | 617402073 | -1.3932 | | 8.85E-05 | BLUPmrMLM | SUG |
| PH_18JY | AX-109932518 | 6B | 26518151 | -1.5429 | | 9.68E-08 | BLUPmrMLM | SIG |
| PH_18JY | AX-110198219 | 6B | 49829896 | -0.1065 | 7.5664 | 4.52E-07 | IIIVmrMLM | SUG |

| | | | | | | | | |
|---|---|---|---|---|---|---|---|---|
| PH_18JY | AX-109508116 | 6B | 147675562 | 1.5993 | | 4.09E-05 | BLUPmrMLM | SUG |
| PH_18JY | AX-109508116 | 6B | 147675562 | -1.3367 | 1.5708 | 1.47E-11 | IIIVmrMLM | SIG |
| PH_18JY | AX-109364183 | 6B | 694461959 | -12.7395 | | 2.26E-08 | FarmCPU | SIG |
| PH_18JY | AX-110403555 | 6B | 714780774 | 1.9755 | | 6.22E-05 | BLUPmrMLM | SUG |
| PH_18JY | AX-108963827 | 6D | 63463118 | -2.1131 | | 1.84E-04 | BLUPmrMLM | SUG |
| PH_18JY | AX-108963827 | 6D | 63463118 | 1.6555 | 1.5292 | 1.49E-16 | IIIVmrMLM | SIG |
| PH_18JY | AX-110184372 | 6D | 299221902 | 30.2360 | | 1.99E-06 | FarmCPU | SUG |
| PH_18JY | AX-111157636 | 6D | 370683123 | 4.2327 | | 9.45E-07 | EMMAX | SUG |
| PH_18JY | AX-111157636 | 6D | 370683123 | -4.2767 | | 5.04E-07 | GEMMA | SUG |
| PH_18JY | AX-110575536 | 6D | 380176310 | 5.0114 | | 1.29E-07 | EMMAX | SUG |
| PH_18JY | AX-110575536 | 6D | 380176310 | -5.0480 | | 6.86E-08 | GEMMA | SIG |
| PH_18JY | AX-110575536 | 6D | 380176310 | 2.2064 | 6.0093 | 5.36E-30 | IIIVmrMLM | SIG |
| PH_18JY | AX-95141417 | 6D | 402119936 | 27.9490 | | 1.71E-06 | EMMAX | SUG |
| PH_18JY | AX-109744641 | 6D | 464830893 | -0.8360 | | 1.02E-05 | IIIVmrMLM | SUG |
| PH_18JY | AX-110692160 | 7A | 34748485 | 0.7850 | -3.3819 | 3.06E-23 | IIIVmrMLM | SIG |
| PH_18JY | AX-109332374 | 7A | 166274186 | 2.0445 | 3.266 | 3.01E-24 | IIIVmrMLM | SIG |
| PH_18JY | AX-108759658 | 7A | 339515663 | -30.9530 | | 3.83E-09 | EMMAX | SIG |
| PH_18JY | AX-94470116 | 7A | 341434034 | -14.1300 | | 2.02E-06 | EMMAX | SUG |
| PH_18JY | AX-94470116 | 7A | 341434034 | 13.8745 | | 2.08E-06 | GEMMA | SUG |
| PH_18JY | AX-110649535 | 7A | 362743928 | 0.1954 | 10.6171 | 1.11E-07 | IIIVmrMLM | SIG |
| PH_18JY | AX-109505823 | 7A | 622135874 | 0.5426 | 9.8521 | 2.19E-22 | IIIVmrMLM | SIG |
| PH_18JY | AX-110362833 | 7A | 697765578 | -0.0022 | -2.4524 | 3.86E-07 | IIIVmrMLM | SUG |
| PH_18JY | AX-110545817 | 7A | 711657696 | 1.8748 | | 8.17E-22 | IIIVmrMLM | SIG |
| PH_18JY | AX-111524448 | 7A | 713176845 | -0.2327 | 6.0613 | 2.50E-08 | IIIVmrMLM | SIG |
| PH_18JY | AX-111037532 | 7B | 130502745 | -1.9015 | | 1.24E-07 | BLUPmrMLM | SUG |
| PH_18JY | AX-111037532 | 7B | 130502745 | 1.8696 | 0.9677 | 1.60E-18 | IIIVmrMLM | SIG |
| PH_18JY | AX-109362596 | 7B | 142929759 | -2.6057 | | 5.85E-08 | BLUPmrMLM | SIG |
| PH_18JY | AX-111575791 | 7B | 361027957 | -1.3801 | | 6.62E-13 | IIIVmrMLM | SIG |
| PH_18JY | AX-109923902 | 7B | 629937286 | 1.2555 | 4.2182 | 1.86E-11 | IIIVmrMLM | SIG |
| PH_18JY | AX-109458469 | 7B | 715365394 | 9.6766 | | 2.04E-06 | GEMMA | SUG |
| PH_18JY | AX-112286117 | 7B | 716041172 | -12.6070 | | 5.47E-07 | EMMAX | SUG |
| PH_18JY | AX-112286117 | 7B | 716041172 | 12.5231 | | 4.20E-07 | GEMMA | SUG |
| PH_18JY | AX-94654805 | 7D | 27097795 | -58.4010 | | 1.03E-14 | EMMAX | SIG |
| PH_18JY | AX-95099853 | 7D | 65036126 | -2.0529 | | 1.62E-05 | BLUPmrMLM | SUG |
| PH_18JY | AX-95099853 | 7D | 65036126 | 0.8502 | | 7.23E-06 | IIIVmrMLM | SUG |
| PH_18JY | AX-111713225 | 7D | 71863395 | -3.2143 | | 5.70E-08 | BLUPmrMLM | SIG |
| PH_18JY | AX-95156310 | 7D | 129781287 | -35.1550 | | 7.82E-09 | EMMAX | SIG |
| PH_18JY | AX-110925192 | 7D | 197297065 | -46.4200 | | 3.56E-12 | EMMAX | SIG |
| PH_18JY | AX-94915838 | 7D | 508614230 | -24.0450 | | 1.71E-06 | EMMAX | SUG |
| PH_18JY | AX-109858608 | 7D | 557455518 | -44.2920 | | 3.42E-10 | EMMAX | SIG |
| PH_18XX | AX-109832236 | 1A | 15375351 | 5.8997 | | 4.34E-08 | FarmCPU | SIG |
| PH_18XX | AX-111015018 | 1A | 385277535 | -1.0130 | | 6.19E-07 | BLUPmrMLM | SUG |
| PH_18XX | AX-109902759 | 1A | 388463098 | 1.8797 | -1.6669 | 3.11E-35 | IIIVmrMLM | SIG |
| PH_18XX | AX-109328604 | 1A | 552160033 | -0.9893 | | 3.57E-06 | BLUPmrMLM | SUG |

| | | | | | | | | |
|---|---|---|---|---|---|---|---|---|
| PH_18XX | AX-109322584 | 1A | 569146363 | -1.1765 | 1.134 | 2.19E-15 | IIIVmrMLM | SIG |
| PH_18XX | AX-111216240 | 1A | 569424493 | 1.2626 | | 8.18E-06 | BLUPmrMLM | SUG |
| PH_18XX | AX-111154144 | 1A | 574832295 | -3.2503 | | 3.90E-08 | BLUPmrMLM | SIG |
| PH_18XX | AX-108738714 | 1B | 90991683 | 2.8347 | -0.5541 | 4.68E-65 | IIIVmrMLM | SIG |
| PH_18XX | AX-109431794 | 1B | 356886594 | -1.9061 | | 4.73E-08 | BLUPmrMLM | SIG |
| PH_18XX | AX-109446199 | 1B | 579938012 | 6.0399 | | 1.28E-08 | FarmCPU | SIG |
| PH_18XX | AX-108837702 | 1B | 590066339 | -1.4101 | | 1.88E-06 | BLUPmrMLM | SUG |
| PH_18XX | AX-94949295 | 1B | 645708529 | -0.5963 | 9.7221 | 7.62E-16 | IIIVmrMLM | SIG |
| PH_18XX | AX-111030152 | 2A | 31820968 | -5.6027 | | 2.95E-09 | FarmCPU | SIG |
| PH_18XX | AX-89621258 | 2A | 49851341 | 1.7664 | -1.5747 | 2.96E-31 | IIIVmrMLM | SIG |
| PH_18XX | AX-110123869 | 2A | 284265347 | -1.2962 | 7.422 | 1.41E-30 | IIIVmrMLM | SIG |
| PH_18XX | AX-111129529 | 2A | 375188307 | 1.3167 | 2.3711 | 3.13E-20 | IIIVmrMLM | SIG |
| PH_18XX | AX-111633619 | 2A | 496352754 | -9.7271 | | 2.55E-07 | EMMAX | SUG |
| PH_18XX | AX-111633619 | 2A | 496352754 | 9.5947 | | 2.86E-07 | GEMMA | SUG |
| PH_18XX | AX-108934390 | 2A | 563945960 | 1.5992 | | 7.56E-28 | IIIVmrMLM | SIG |
| PH_18XX | AX-94779131 | 2A | 607830166 | -13.0920 | | 1.05E-06 | EMMAX | SUG |
| PH_18XX | AX-94779131 | 2A | 607830166 | 13.2298 | | 6.49E-07 | GEMMA | SUG |
| PH_18XX | AX-111068539 | 2A | 753153585 | 0.7284 | | 4.11E-05 | BLUPmrMLM | SUG |
| PH_18XX | AX-94488783 | 2A | 755788522 | -2.3172 | | 5.36E-07 | BLUPmrMLM | SUG |
| PH_18XX | AX-94537654 | 2B | 24853000 | 0.3883 | -2.4608 | 2.49E-41 | IIIVmrMLM | SIG |
| PH_18XX | AX-109405720 | 2B | 25980102 | -1.5231 | -0.084 | 7.15E-19 | IIIVmrMLM | SIG |
| PH_18XX | AX-111562074 | 2B | 74685875 | 1.5975 | | 2.36E-09 | BLUPmrMLM | SIG |
| PH_18XX | AX-109322878 | 2B | 89423615 | 1.1291 | | 8.35E-07 | BLUPmrMLM | SUG |
| PH_18XX | AX-94749804 | 2B | 762518824 | 2.2327 | | 6.68E-07 | BLUPmrMLM | SUG |
| PH_18XX | AX-94804912 | 2B | 784551326 | 12.5247 | | 3.05E-08 | FarmCPU | SIG |
| PH_18XX | AX-94934049 | 2B | 788244037 | 1.2463 | | 8.65E-06 | BLUPmrMLM | SUG |
| PH_18XX | AX-109946720 | 2D | 110666440 | -1.6610 | | 6.78E-05 | BLUPmrMLM | SUG |
| PH_18XX | AX-109627287 | 2D | 116801772 | -6.8665 | | 5.70E-07 | FarmCPU | SUG |
| PH_18XX | AX-109844299 | 2D | 491947833 | -5.4101 | | 1.25E-06 | EMMAX | SUG |
| PH_18XX | AX-109844299 | 2D | 491947833 | 5.4513 | | 6.77E-07 | GEMMA | SUG |
| PH_18XX | AX-110552619 | 2D | 509540852 | 1.2703 | | 6.33E-07 | BLUPmrMLM | SUG |
| PH_18XX | AX-94939797 | 2D | 635950351 | 1.9041 | | 5.77E-08 | BLUPmrMLM | SIG |
| PH_18XX | AX-109840736 | 3A | 50643363 | -0.9986 | -1.4683 | 1.60E-11 | IIIVmrMLM | SIG |
| PH_18XX | AX-110467557 | 3A | 507462215 | 4.9562 | | 4.49E-08 | EMMAX | SIG |
| PH_18XX | AX-110467557 | 3A | 507462215 | -4.9415 | | 3.15E-08 | GEMMA | SIG |
| PH_18XX | AX-111512557 | 3A | 512859054 | 1.3439 | 0.9488 | 2.18E-18 | IIIVmrMLM | SIG |
| PH_18XX | AX-111579126 | 3A | 587965191 | 1.4452 | | 5.23E-07 | BLUPmrMLM | SUG |
| PH_18XX | AX-109887519 | 3A | 654584870 | 0.8656 | | 9.14E-05 | BLUPmrMLM | SUG |
| PH_18XX | AX-111152038 | 3A | 655581373 | -1.0539 | 5.6498 | 1.29E-20 | IIIVmrMLM | SIG |
| PH_18XX | AX-110573094 | 3A | 703209853 | -0.8845 | 1.9848 | 3.42E-10 | IIIVmrMLM | SIG |
| PH_18XX | AX-110491100 | 3A | 705251779 | -7.8344 | | 1.03E-07 | EMMAX | SIG |
| PH_18XX | AX-110491100 | 3A | 705251779 | 7.7575 | | 7.29E-08 | GEMMA | SIG |
| PH_18XX | AX-108941177 | 3A | 705259848 | -7.1423 | | 3.06E-07 | EMMAX | SUG |
| PH_18XX | AX-108941177 | 3A | 705259848 | 7.1035 | | 1.84E-07 | GEMMA | SUG |

| | | | | | | | | |
|---|---|---|---|---|---|---|---|---|
| PH_18XX | AX-111695263 | 3A | 705274739 | -7.3084 | | 3.21E-07 | EMMAX | SUG |
| PH_18XX | AX-111695263 | 3A | 705274739 | 7.3026 | | 1.77E-07 | GEMMA | SUG |
| PH_18XX | AX-110976174 | 3A | 705274782 | -14.0450 | | 4.31E-07 | EMMAX | SUG |
| PH_18XX | AX-110976174 | 3A | 705274782 | 13.8782 | | 3.29E-07 | GEMMA | SUG |
| PH_18XX | AX-110484538 | 3A | 705296840 | -17.2220 | | 3.91E-08 | EMMAX | SIG |
| PH_18XX | AX-110484538 | 3A | 705296840 | 16.9472 | | 2.92E-08 | GEMMA | SIG |
| PH_18XX | AX-111121746 | 3A | 720214589 | -1.7206 | | 1.18E-06 | BLUPmrMLM | SUG |
| PH_18XX | AX-110461664 | 3A | 735270884 | 1.4793 | | 4.36E-07 | BLUPmrMLM | SUG |
| PH_18XX | AX-110070120 | 3B | 2885668 | -0.4711 | 3.3073 | 3.16E-06 | IIIVmrMLM | SUG |
| PH_18XX | AX-94624961 | 3B | 7605311 | 1.4644 | 1.979 | 3.60E-36 | IIIVmrMLM | SIG |
| PH_18XX | AX-110960188 | 3B | 20818271 | -1.2294 | -1.1233 | 1.19E-16 | IIIVmrMLM | SIG |
| PH_18XX | AX-111700618 | 3B | 345531099 | 0.1138 | -4.3876 | 2.77E-12 | IIIVmrMLM | SIG |
| PH_18XX | AX-111756000 | 3B | 470096153 | -0.1001 | -5.1347 | 2.62E-08 | IIIVmrMLM | SIG |
| PH_18XX | AX-111654766 | 3B | 738656110 | 0.9248 | | 1.31E-04 | BLUPmrMLM | SUG |
| PH_18XX | AX-109310521 | 3B | 741653603 | -3.2877 | | 2.46E-09 | BLUPmrMLM | SIG |
| PH_18XX | AX-108923152 | 3B | 773728815 | -4.4105 | | 1.95E-06 | FarmCPU | SUG |
| PH_18XX | AX-94827431 | 3D | 7972918 | -2.7191 | | 2.04E-05 | BLUPmrMLM | SUG |
| PH_18XX | AX-94907389 | 3D | 389191643 | -2.3949 | | 1.28E-06 | BLUPmrMLM | SUG |
| PH_18XX | AX-94518159 | 3D | 561100857 | 1.5042 | | 4.50E-25 | IIIVmrMLM | SIG |
| PH_18XX | AX-94845094 | 3D | 600423999 | -12.7057 | | 1.60E-07 | FarmCPU | SUG |
| PH_18XX | AX-108751023 | 4A | 60921930 | 1.4074 | 0.8051 | 3.28E-21 | IIIVmrMLM | SIG |
| PH_18XX | AX-110124454 | 4A | 87357972 | -0.3380 | 1.4591 | 2.83E-04 | IIIVmrMLM | SUG |
| PH_18XX | AX-109572358 | 4A | 547520547 | 0.6715 | 2.3303 | 2.35E-21 | IIIVmrMLM | SIG |
| PH_18XX | AX-109970485 | 4A | 627949032 | -0.2498 | -2.8885 | 2.41E-13 | IIIVmrMLM | SIG |
| PH_18XX | AX-109299526 | 4A | 725843038 | -2.4133 | | 3.33E-12 | BLUPmrMLM | SIG |
| PH_18XX | AX-111473821 | 4B | 643918109 | 0.1081 | -2.9799 | 1.05E-06 | IIIVmrMLM | SUG |
| PH_18XX | AX-108944764 | 4D | 15911901 | 4.4739 | | 1.52E-06 | EMMAX | SUG |
| PH_18XX | AX-108944764 | 4D | 15911901 | -4.4987 | | 7.60E-07 | GEMMA | SUG |
| PH_18XX | AX-89593703 | 4D | 15958106 | 4.9691 | | 5.35E-08 | EMMAX | SIG |
| PH_18XX | AX-89593703 | 4D | 15958106 | -5.1439 | | 7.96E-09 | GEMMA | SIG |
| PH_18XX | AX-89703298 | 4D | 16926596 | 7.3864 | | 2.85E-13 | EMMAX | SIG |
| PH_18XX | AX-89703298 | 4D | 16926596 | -7.5398 | | 1.79E-14 | GEMMA | SIG |
| PH_18XX | AX-109281717 | 4D | 17613651 | 7.7063 | | 7.02E-14 | EMMAX | SIG |
| PH_18XX | AX-109281717 | 4D | 17613651 | -7.8457 | | 3.86E-15 | GEMMA | SIG |
| PH_18XX | AX-109665328 | 4D | 18539670 | 7.7486 | | 5.24E-15 | EMMAX | SIG |
| PH_18XX | AX-109665328 | 4D | 18539670 | -7.8892 | | 3.52E-16 | GEMMA | SIG |
| PH_18XX | AX-109665328 | 4D | 18539670 | 3.9784 | 0.1471 | 9.11E-113 | IIIVmrMLM | SIG |
| PH_18XX | AX-108916749 | 4D | 19090268 | 5.7652 | | 2.83E-11 | EMMAX | SIG |
| PH_18XX | AX-108916749 | 4D | 19090268 | -5.8332 | | 5.30E-12 | GEMMA | SIG |
| PH_18XX | AX-89421921 | 4D | 19288898 | 7.7135 | | 4.47E-14 | EMMAX | SIG |
| PH_18XX | AX-89421921 | 4D | 19288898 | -7.8527 | | 2.39E-15 | GEMMA | SIG |
| PH_18XX | AX-89445201 | 4D | 19459579 | 4.8527 | | 2.83E-08 | EMMAX | SIG |
| PH_18XX | AX-89445201 | 4D | 19459579 | -5.0957 | | 2.72E-09 | GEMMA | SIG |

| | | | | | | | | |
|---|---|---|---|---|---|---|---|---|
| PH_18XX | AX-111970247 | 4D | 32225033 | -2.1082 | | 1.29E-08 | BLUPmrMLM | SIG |
| PH_18XX | AX-110184307 | 4D | 220588272 | -1.0347 | | 3.43E-13 | IIIVmrMLM | SIG |
| PH_18XX | AX-110047101 | 4D | 224594707 | -46.4660 | | 9.36E-09 | EMMAX | SIG |
| PH_18XX | AX-95186747 | 4D | 446428878 | -39.9120 | | 4.79E-10 | EMMAX | SIG |
| PH_18XX | AX-110387369 | 4D | 467431300 | -33.5420 | | 2.55E-07 | EMMAX | SUG |
| PH_18XX | AX-110485186 | 5A | 8120257 | -0.6608 | 1.0492 | 4.05E-06 | IIIVmrMLM | SUG |
| PH_18XX | AX-110440664 | 5A | 18893163 | -1.3632 | 2.0002 | 2.70E-20 | IIIVmrMLM | SIG |
| PH_18XX | AX-109946449 | 5A | 18905296 | 1.3285 | | 6.98E-06 | BLUPmrMLM | SUG |
| PH_18XX | AX-111726524 | 5A | 29628692 | -2.1519 | | 1.59E-08 | BLUPmrMLM | SIG |
| PH_18XX | AX-111726524 | 5A | 29628692 | 2.1209 | 1.0134 | 2.22E-43 | IIIVmrMLM | SIG |
| PH_18XX | AX-109903760 | 5A | 30056842 | 6.6181 | | 5.99E-08 | EMMAX | SIG |
| PH_18XX | AX-109903760 | 5A | 30056842 | -6.6054 | | 3.15E-08 | GEMMA | SIG |
| PH_18XX | AX-109820064 | 5A | 703212889 | 6.6800 | | 3.97E-07 | EMMAX | SUG |
| PH_18XX | AX-109820064 | 5A | 703212889 | -6.6590 | | 2.05E-07 | GEMMA | SUG |
| PH_18XX | AX-110999817 | 5A | 708416688 | 0.8526 | -6.379 | 9.81E-109 | IIIVmrMLM | SIG |
| PH_18XX | AX-110694654 | 5A | 708860993 | -0.0503 | -1.4768 | 7.52E-10 | IIIVmrMLM | SIG |
| PH_18XX | AX-110477837 | 5B | 64025780 | 6.7371 | | 3.54E-07 | FarmCPU | SUG |
| PH_18XX | AX-94509069 | 5B | 211148106 | -0.4759 | 10.6988 | 4.32E-13 | IIIVmrMLM | SIG |
| PH_18XX | AX-111523050 | 5B | 337060642 | 0.0966 | -6.6269 | 2.57E-06 | IIIVmrMLM | SUG |
| PH_18XX | AX-110018488 | 5B | 528535571 | -3.3583 | | 1.20E-06 | BLUPmrMLM | SUG |
| PH_18XX | AX-110018488 | 5B | 528535571 | -5.3899 | | 1.72E-06 | GEMMA | SUG |
| PH_18XX | AX-110018488 | 5B | 528535571 | 2.1898 | | 3.57E-47 | IIIVmrMLM | SIG |
| PH_18XX | AX-110706119 | 5B | 686315496 | -1.2521 | | 2.63E-06 | BLUPmrMLM | SUG |
| PH_18XX | AX-110930042 | 5D | 362887848 | -36.2240 | | 7.60E-10 | EMMAX | SIG |
| PH_18XX | AX-109666105 | 5D | 473534891 | 1.3898 | 1.3183 | 9.30E-21 | IIIVmrMLM | SIG |
| PH_18XX | AX-94963508 | 5D | 484426813 | -46.4660 | | 9.36E-09 | EMMAX | SIG |
| PH_18XX | AX-111456331 | 6A | 17395657 | 1.5604 | | 3.92E-05 | BLUPmrMLM | SUG |
| PH_18XX | AX-109553965 | 6A | 26932963 | -0.8088 | | 1.08E-04 | BLUPmrMLM | SUG |
| PH_18XX | AX-108803390 | 6A | 559008014 | 1.1212 | | 3.00E-06 | BLUPmrMLM | SUG |
| PH_18XX | AX-111450720 | 6B | 25972891 | -1.7181 | | 4.38E-08 | BLUPmrMLM | SIG |
| PH_18XX | AX-111471655 | 6B | 582865287 | 1.4122 | | 2.35E-06 | BLUPmrMLM | SUG |
| PH_18XX | AX-95631878 | 6B | 645353503 | 0.5207 | 5.0015 | 1.21E-07 | IIIVmrMLM | SUG |
| PH_18XX | AX-109364183 | 6B | 694461959 | -9.1944 | | 1.84E-07 | FarmCPU | SUG |
| PH_18XX | AX-95006509 | 6B | 708020637 | 0.0703 | -1.4394 | 3.65E-10 | IIIVmrMLM | SIG |
| PH_18XX | AX-89463101 | 6D | 17946690 | 0.7338 | -1.7176 | 5.73E-07 | IIIVmrMLM | SUG |
| PH_18XX | AX-109492844 | 6D | 89236767 | -2.2214 | -3.8417 | 1.46E-65 | IIIVmrMLM | SIG |
| PH_18XX | AX-109447460 | 6D | 89262091 | 1.5011 | | 1.02E-06 | BLUPmrMLM | SUG |
| PH_18XX | AX-94623747 | 6D | 466737822 | -0.4512 | -1.8383 | 2.84E-09 | IIIVmrMLM | SIG |
| PH_18XX | AX-109409627 | 6D | 468841296 | 1.2611 | | 4.90E-06 | BLUPmrMLM | SUG |
| PH_18XX | AX-108825706 | 6D | 469812057 | -1.2911 | -0.4575 | 1.80E-10 | IIIVmrMLM | SIG |
| PH_18XX | AX-111489417 | 7A | 514670037 | -0.3149 | 1.816 | 5.44E-06 | IIIVmrMLM | SUG |
| PH_18XX | AX-111685547 | 7A | 545890031 | 1.0738 | | 9.56E-05 | BLUPmrMLM | SUG |
| PH_18XX | AX-108799475 | 7A | 581690737 | -1.2935 | | 1.22E-04 | BLUPmrMLM | SUG |

| | | | | | | | | |
|---|---|---|---|---|---|---|---|---|
| PH_18XX | AX-109505823 | 7A | 622135874 | 3.9100 | 9.9597 | 1.35E-125 | IIIVmrMLM | SIG |
| PH_18XX | AX-109841908 | 7A | 680112565 | 0.9880 | | 5.46E-06 | BLUPmrMLM | SUG |
| PH_18XX | AX-108962229 | 7A | 708912808 | 2.6028 | | 1.30E-09 | BLUPmrMLM | SIG |
| PH_18XX | AX-110054532 | 7A | 713759224 | -0.0582 | -3.6833 | 9.15E-11 | IIIVmrMLM | SIG |
| PH_18XX | AX-109437769 | 7A | 733984502 | -0.1071 | -4.124 | 6.06E-28 | IIIVmrMLM | SIG |
| PH_18XX | AX-110488268 | 7B | 42147774 | -3.1646 | -1.5357 | 1.28E-75 | IIIVmrMLM | SIG |
| PH_18XX | AX-111220072 | 7B | 63936379 | 0.8717 | | 1.45E-04 | BLUPmrMLM | SUG |
| PH_18XX | AX-111141626 | 7B | 244631972 | -0.1647 | 2.9271 | 1.50E-07 | IIIVmrMLM | SUG |
| PH_18XX | AX-95112092 | 7B | 581893312 | -0.7626 | | 8.83E-05 | BLUPmrMLM | SUG |
| PH_18XX | AX-112286117 | 7B | 716041172 | -9.9213 | | 5.62E-08 | EMMAX | SIG |
| PH_18XX | AX-112286117 | 7B | 716041172 | 9.9293 | | 3.20E-08 | GEMMA | SIG |
| PH_18XX | AX-110493018 | 7B | 717630916 | 0.1879 | 13.7176 | 8.58E-08 | IIIVmrMLM | SIG |
| PH_18XX | AX-110275961 | 7D | 23192143 | 2.3910 | 2.3351 | 5.28E-53 | IIIVmrMLM | SIG |
| PH_18XX | AX-94654805 | 7D | 27097795 | -35.1070 | | 8.13E-09 | EMMAX | SIG |
| PH_18XX | AX-111061288 | 7D | 66537922 | 1.1118 | | 8.23E-06 | BLUPmrMLM | SUG |
| PH_18XX | AX-111061288 | 7D | 66537922 | -0.9046 | -0.5075 | 5.12E-09 | IIIVmrMLM | SIG |
| PH_18XX | AX-94997278 | 7D | 74467098 | 3.3219 | | 1.61E-08 | BLUPmrMLM | SIG |
| PH_18XX | AX-94997278 | 7D | 74467098 | -0.9335 | 0.3369 | 6.56E-04 | IIIVmrMLM | SUG |
| PH_18XX | AX-110925192 | 7D | 197297065 | -28.7030 | | 6.99E-08 | EMMAX | SIG |
| PH_18ZMD | AX-109358987 | 1A | 8289762 | -11.3929 | | 1.41E-06 | FarmCPU | SUG |
| PH_18ZMD | AX-111663403 | 1A | 11413174 | 0.0109 | -4.6233 | 4.01E-09 | IIIVmrMLM | SIG |
| PH_18ZMD | AX-109832236 | 1A | 15375351 | 6.3924 | | 5.56E-07 | FarmCPU | SUG |
| PH_18ZMD | AX-110917340 | 1A | 239888558 | 2.6824 | | 1.93E-06 | BLUPmrMLM | SUG |
| PH_18ZMD | AX-110983499 | 1A | 352858301 | -9.7572 | | 2.07E-06 | GEMMA | SUG |
| PH_18ZMD | AX-111029436 | 1A | 372764935 | -1.6999 | | 1.13E-04 | BLUPmrMLM | SUG |
| PH_18ZMD | AX-108840039 | 1A | 387936105 | 10.0520 | | 1.27E-06 | EMMAX | SUG |
| PH_18ZMD | AX-108840039 | 1A | 387936105 | -10.3402 | | 2.89E-07 | GEMMA | SUG |
| PH_18ZMD | AX-111668333 | 1A | 388937021 | -10.4222 | | 2.20E-06 | GEMMA | SUG |
| PH_18ZMD | AX-109379218 | 1A | 388947077 | 11.2430 | | 1.30E-06 | EMMAX | SUG |
| PH_18ZMD | AX-109379218 | 1A | 388947077 | -10.8742 | | 8.70E-07 | GEMMA | SUG |
| PH_18ZMD | AX-111819445 | 1A | 388947113 | 11.2430 | | 1.30E-06 | EMMAX | SUG |
| PH_18ZMD | AX-111819445 | 1A | 388947113 | -10.8742 | | 8.70E-07 | GEMMA | SUG |
| PH_18ZMD | AX-108742536 | 1A | 389029091 | 11.2430 | | 1.30E-06 | EMMAX | SUG |
| PH_18ZMD | AX-108742536 | 1A | 389029091 | -10.8742 | | 8.70E-07 | GEMMA | SUG |
| PH_18ZMD | AX-110403537 | 1A | 389061020 | 11.2430 | | 1.30E-06 | EMMAX | SUG |
| PH_18ZMD | AX-110403537 | 1A | 389061020 | -10.8742 | | 8.70E-07 | GEMMA | SUG |
| PH_18ZMD | AX-110448433 | 1A | 389061946 | 21.9560 | | 2.26E-06 | EMMAX | SUG |
| PH_18ZMD | AX-110448433 | 1A | 389061946 | -21.2204 | | 1.61E-06 | GEMMA | SUG |
| PH_18ZMD | AX-111106333 | 1A | 389200944 | -10.0587 | | 2.05E-06 | GEMMA | SUG |
| PH_18ZMD | AX-108850470 | 1A | 389228613 | 10.9780 | | 2.26E-06 | EMMAX | SUG |
| PH_18ZMD | AX-108850470 | 1A | 389228613 | -10.6102 | | 1.61E-06 | GEMMA | SUG |
| PH_18ZMD | AX-110492346 | 1A | 389252927 | 10.9780 | | 2.26E-06 | EMMAX | SUG |
| PH_18ZMD | AX-110492346 | 1A | 389252927 | -10.6102 | | 1.61E-06 | GEMMA | SUG |

| | | | | | | | | |
|---|---|---|---|---|---|---|---|---|
| PH_18ZMD | AX-109509868 | 1A | 389363391 | 10.9780 | | 2.26E-06 | EMMAX | SUG |
| PH_18ZMD | AX-109509868 | 1A | 389363391 | -10.6102 | | 1.61E-06 | GEMMA | SUG |
| PH_18ZMD | AX-110628967 | 1A | 389803509 | 10.9780 | | 2.26E-06 | EMMAX | SUG |
| PH_18ZMD | AX-110628967 | 1A | 389803509 | -10.6102 | | 1.61E-06 | GEMMA | SUG |
| PH_18ZMD | AX-110152611 | 1A | 390698547 | -2.0619 | -4.108 | 6.43E-16 | IIIVmrMLM | SIG |
| PH_18ZMD | AX-110554073 | 1A | 460637199 | -11.1462 | | 3.49E-07 | FarmCPU | SUG |
| PH_18ZMD | AX-109887708 | 1A | 494715990 | 10.7130 | | 2.17E-06 | EMMAX | SUG |
| PH_18ZMD | AX-109887708 | 1A | 494715990 | -10.7104 | | 8.84E-07 | GEMMA | SUG |
| PH_18ZMD | AX-111505010 | 1A | 494742630 | -10.4675 | | 1.02E-06 | GEMMA | SUG |
| PH_18ZMD | AX-110501782 | 1A | 494752723 | -10.4675 | | 1.02E-06 | GEMMA | SUG |
| PH_18ZMD | AX-109817102 | 1A | 495013803 | 9.8957 | | 1.36E-06 | EMMAX | SUG |
| PH_18ZMD | AX-109817102 | 1A | 495013803 | -9.9019 | | 5.15E-07 | GEMMA | SUG |
| PH_18ZMD | AX-111107506 | 1A | 515578994 | 1.7103 | | 6.75E-05 | BLUPmrMLM | SUG |
| PH_18ZMD | AX-94934987 | 1A | 534043666 | -49.3710 | | 3.73E-08 | EMMAX | SIG |
| PH_18ZMD | AX-108901390 | 1A | 546225444 | 8.6653 | | 1.40E-06 | FarmCPU | SUG |
| PH_18ZMD | AX-95149778 | 1B | 15745786 | 0.0385 | 6.3103 | 2.95E-06 | IIIVmrMLM | SUG |
| PH_18ZMD | AX-109372658 | 1B | 41084324 | 0.1735 | 4.1005 | 1.92E-06 | IIIVmrMLM | SUG |
| PH_18ZMD | AX-110935835 | 1B | 620555612 | 1.4531 | 32.4082 | 1.26E-14 | IIIVmrMLM | SIG |
| PH_18ZMD | AX-94656925 | 1D | 2485672 | 7.6661 | | 1.33E-07 | FarmCPU | SUG |
| PH_18ZMD | AX-94420233 | 1D | 155927416 | -49.3710 | | 3.73E-08 | EMMAX | SIG |
| PH_18ZMD | AX-111090826 | 1D | 316571808 | 5.5047 | | 7.58E-10 | BLUPmrMLM | SIG |
| PH_18ZMD | AX-111554249 | 1D | 409853326 | 0.9895 | -4.7105 | 2.48E-11 | IIIVmrMLM | SIG |
| PH_18ZMD | AX-111030152 | 2A | 31820968 | -8.6255 | | 4.89E-09 | FarmCPU | SIG |
| PH_18ZMD | AX-110513895 | 2A | 61480507 | -1.6520 | | 2.85E-06 | BLUPmrMLM | SUG |
| PH_18ZMD | AX-109403848 | 2A | 444792470 | -2.4859 | | 9.11E-05 | BLUPmrMLM | SUG |
| PH_18ZMD | AX-108816023 | 2A | 555397721 | 1.7547 | 7.4969 | 2.00E-12 | IIIVmrMLM | SIG |
| PH_18ZMD | AX-110456050 | 2A | 751211338 | -2.0897 | | 1.02E-04 | BLUPmrMLM | SUG |
| PH_18ZMD | AX-109518247 | 2B | 48289705 | 2.0178 | 0.1775 | 5.56E-14 | IIIVmrMLM | SIG |
| PH_18ZMD | AX-109459500 | 2B | 117106337 | -2.7458 | | 3.98E-07 | BLUPmrMLM | SUG |
| PH_18ZMD | AX-109459500 | 2B | 117106337 | 2.0562 | 2.0497 | 1.10E-15 | IIIVmrMLM | SIG |
| PH_18ZMD | AX-111523828 | 2B | 522723421 | 2.4516 | | 6.91E-06 | BLUPmrMLM | SUG |
| PH_18ZMD | AX-109408715 | 2B | 532494104 | 11.9325 | | 2.05E-06 | GEMMA | SUG |
| PH_18ZMD | AX-109294263 | 2B | 619754236 | 2.7018 | | 3.61E-05 | BLUPmrMLM | SUG |
| PH_18ZMD | AX-111483077 | 2B | 796331961 | 3.1369 | 0.4881 | 1.99E-29 | IIIVmrMLM | SIG |
| PH_18ZMD | AX-95209398 | 2D | 405408295 | -37.4380 | | 8.15E-07 | EMMAX | SUG |
| PH_18ZMD | AX-109844299 | 2D | 491947833 | 5.3005 | | 7.73E-10 | BLUPmrMLM | SIG |
| PH_18ZMD | AX-109844299 | 2D | 491947833 | 7.4531 | | 2.01E-06 | GEMMA | SUG |
| PH_18ZMD | AX-94427799 | 2D | 523357562 | -9.4526 | | 2.21E-07 | EMMAX | SUG |
| PH_18ZMD | AX-94427799 | 2D | 523357562 | 9.4064 | | 1.50E-07 | GEMMA | SUG |
| PH_18ZMD | AX-109473421 | 2D | 542135244 | 2.0425 | -7.4305 | 4.09E-18 | IIIVmrMLM | SIG |
| PH_18ZMD | AX-111336862 | 2D | 629367397 | -2.5644 | -0.6411 | 1.55E-22 | IIIVmrMLM | SIG |
| PH_18ZMD | AX-109911657 | 2D | 640389871 | 2.8937 | | 1.03E-07 | BLUPmrMLM | SIG |
| PH_18ZMD | AX-95240191 | 3A | 7920892 | -7.8594 | -3.8997 | 5.52E-97 | IIIVmrMLM | SIG |
| PH_18ZMD | AX-110436755 | 3A | 19504804 | 2.5902 | | 1.52E-07 | BLUPmrMLM | SUG |

| | | | | | | | | |
|---|---|---|---|---|---|---|---|---|
| PH_18ZMD | AX-89453831 | 3A | 724699913 | 2.5029 | -8.5642 | 1.47E-25 | IIIVmrMLM | SIG |
| PH_18ZMD | AX-110914455 | 3B | 20381356 | -1.6903 | -0.1708 | 8.16E-11 | IIIVmrMLM | SIG |
| PH_18ZMD | AX-110039198 | 3B | 69719049 | 0.0123 | 7.7644 | 5.37E-14 | IIIVmrMLM | SIG |
| PH_18ZMD | AX-111526364 | 3B | 117577172 | -1.9707 | | 6.44E-06 | BLUPmrMLM | SUG |
| PH_18ZMD | AX-108941515 | 3B | 469534228 | -2.1269 | -2.2793 | 5.98E-17 | IIIVmrMLM | SIG |
| PH_18ZMD | AX-110149046 | 3B | 643275270 | -2.3244 | | 1.58E-06 | BLUPmrMLM | SUG |
| PH_18ZMD | AX-111673406 | 3B | 760046216 | 0.0155 | 4.0915 | 5.65E-12 | IIIVmrMLM | SIG |
| PH_18ZMD | AX-109292211 | 3B | 780616084 | -6.1646 | | 1.83E-08 | FarmCPU | SIG |
| PH_18ZMD | AX-94607567 | 3D | 285781654 | -45.7220 | | 2.23E-06 | EMMAX | SUG |
| PH_18ZMD | AX-110437801 | 4A | 11172466 | -0.1294 | 12.6277 | 8.29E-07 | IIIVmrMLM | SUG |
| PH_18ZMD | AX-108831701 | 4B | 191790892 | 0.3358 | -1.7421 | 3.73E-04 | IIIVmrMLM | SUG |
| PH_18ZMD | AX-94521391 | 4B | 650730152 | -1.1248 | | 2.72E-05 | BLUPmrMLM | SUG |
| PH_18ZMD | AX-108798487 | 4B | 664199032 | -5.5008 | | 5.38E-09 | BLUPmrMLM | SIG |
| PH_18ZMD | AX-89593703 | 4D | 15958106 | -5.9341 | | 1.49E-06 | GEMMA | SUG |
| PH_18ZMD | AX-89703298 | 4D | 16926596 | 7.5937 | | 7.89E-08 | EMMAX | SIG |
| PH_18ZMD | AX-89703298 | 4D | 16926596 | -7.7586 | | 1.41E-08 | GEMMA | SIG |
| PH_18ZMD | AX-109281717 | 4D | 17613651 | 7.9126 | | 3.23E-08 | EMMAX | SIG |
| PH_18ZMD | AX-109281717 | 4D | 17613651 | -8.0855 | | 5.01E-09 | GEMMA | SIG |
| PH_18ZMD | AX-109665328 | 4D | 18539670 | 7.7112 | | 3.93E-08 | EMMAX | SIG |
| PH_18ZMD | AX-109665328 | 4D | 18539670 | -7.9117 | | 5.84E-09 | GEMMA | SIG |
| PH_18ZMD | AX-108916749 | 4D | 19090268 | 6.6867 | | 8.01E-08 | EMMAX | SIG |
| PH_18ZMD | AX-108916749 | 4D | 19090268 | -6.7925 | | 1.94E-08 | GEMMA | SIG |
| PH_18ZMD | AX-89421921 | 4D | 19288898 | 7.7584 | | 4.92E-08 | EMMAX | SIG |
| PH_18ZMD | AX-89421921 | 4D | 19288898 | -7.9264 | | 8.12E-09 | GEMMA | SIG |
| PH_18ZMD | AX-89445201 | 4D | 19459579 | 6.6704 | | 1.39E-07 | EMMAX | SUG |
| PH_18ZMD | AX-89445201 | 4D | 19459579 | -6.8539 | | 2.30E-08 | GEMMA | SIG |
| PH_18ZMD | AX-89445201 | 4D | 19459579 | 2.4954 | -2.2945 | 3.65E-22 | IIIVmrMLM | SIG |
| PH_18ZMD | AX-111416995 | 4D | 334598743 | -57.3670 | | 3.68E-07 | EMMAX | SUG |
| PH_18ZMD | AX-111562024 | 5A | 9388502 | 2.3864 | | 2.74E-05 | BLUPmrMLM | SUG |
| PH_18ZMD | AX-110951363 | 5A | 377327509 | -0.4214 | -3.5906 | 8.51E-12 | IIIVmrMLM | SIG |
| PH_18ZMD | AX-111032889 | 5A | 519890740 | -0.3106 | 7.0432 | 6.28E-13 | IIIVmrMLM | SIG |
| PH_18ZMD | AX-111031992 | 5A | 540369331 | 2.3827 | | 7.61E-06 | BLUPmrMLM | SUG |
| PH_18ZMD | AX-111077995 | 5A | 583590027 | -2.8184 | | 2.39E-07 | BLUPmrMLM | SUG |
| PH_18ZMD | AX-108892661 | 5A | 687687823 | -2.4281 | | 6.27E-05 | BLUPmrMLM | SUG |
| PH_18ZMD | AX-108728422 | 5A | 706669730 | -2.5151 | 1.4425 | 3.24E-21 | IIIVmrMLM | SIG |
| PH_18ZMD | AX-110999817 | 5A | 708416688 | 2.7098 | -7.6419 | 6.00E-73 | IIIVmrMLM | SIG |
| PH_18ZMD | AX-111755953 | 5B | 26010172 | -1.7370 | | 5.10E-05 | BLUPmrMLM | SUG |
| PH_18ZMD | AX-111036555 | 5B | 513978487 | -7.3097 | | 8.02E-07 | EMMAX | SUG |
| PH_18ZMD | AX-111036555 | 5B | 513978487 | 7.3787 | | 3.61E-07 | GEMMA | SUG |
| PH_18ZMD | AX-109325020 | 5B | 517196591 | -7.1424 | | 1.04E-06 | GEMMA | SUG |
| PH_18ZMD | AX-111492401 | 5B | 522009317 | -2.5393 | | 2.66E-05 | BLUPmrMLM | SUG |
| PH_18ZMD | AX-111492401 | 5B | 522009317 | 1.6203 | -2.2305 | 3.10E-11 | IIIVmrMLM | SIG |
| PH_18ZMD | AX-111512697 | 5B | 527774551 | -7.6946 | | 9.83E-07 | GEMMA | SUG |
| PH_18ZMD | AX-110018488 | 5B | 528535571 | 8.3215 | | 1.71E-07 | EMMAX | SUG |

| | | | | | | | | |
|---|---|---|---|---|---|---|---|---|
| PH_18ZMD | AX-110018488 | 5B | 528535571 | -8.6456 | | 1.90E-08 | GEMMA | SIG |
| PH_18ZMD | AX-110985763 | 5B | 529624248 | -8.5951 | | 1.50E-06 | GEMMA | SUG |
| PH_18ZMD | AX-110933239 | 5B | 529693265 | 2.4894 | 4.1387 | 4.11E-24 | IIIVmrMLM | SIG |
| PH_18ZMD | AX-109276514 | 5B | 568384010 | 0.0142 | -6.0726 | 1.86E-14 | IIIVmrMLM | SIG |
| PH_18ZMD | AX-94728034 | 5D | 434481168 | -3.6723 | | 7.30E-05 | BLUPmrMLM | SUG |
| PH_18ZMD | AX-109453419 | 5D | 451166746 | -0.4523 | 12.1942 | 3.66E-06 | IIIVmrMLM | SUG |
| PH_18ZMD | AX-108951505 | 5D | 486241658 | 0.7138 | 6.2308 | 4.96E-09 | IIIVmrMLM | SIG |
| PH_18ZMD | AX-94860490 | 5D | 555928841 | 3.4703 | 14.8035 | 1.05E-50 | IIIVmrMLM | SIG |
| PH_18ZMD | AX-111185152 | 6A | 2400640 | 0.3975 | 8.9857 | 2.24E-07 | IIIVmrMLM | SUG |
| PH_18ZMD | AX-110425274 | 6A | 26963931 | -0.0014 | 3.1434 | 4.04E-04 | IIIVmrMLM | SUG |
| PH_18ZMD | AX-94453644 | 6A | 55013752 | -1.7417 | | 1.34E-12 | IIIVmrMLM | SIG |
| PH_18ZMD | AX-112285858 | 6A | 84968797 | 1.6730 | | 9.51E-06 | BLUPmrMLM | SUG |
| PH_18ZMD | AX-108758970 | 6B | 47137302 | 0.1354 | -2.9184 | 2.13E-17 | IIIVmrMLM | SIG |
| PH_18ZMD | AX-95162142 | 6B | 199761147 | -26.7160 | | 3.42E-07 | EMMAX | SUG |
| PH_18ZMD | AX-109979570 | 6B | 334222226 | 0.8580 | 3.8328 | 1.89E-07 | IIIVmrMLM | SUG |
| PH_18ZMD | AX-94746925 | 6B | 577363493 | -57.3670 | | 3.68E-07 | EMMAX | SUG |
| PH_18ZMD | AX-94809200 | 6D | 15807973 | -0.1250 | -10.3117 | 3.08E-19 | IIIVmrMLM | SIG |
| PH_18ZMD | AX-89661736 | 6D | 86551098 | -2.4467 | | 3.11E-06 | BLUPmrMLM | SUG |
| PH_18ZMD | AX-109748962 | 6D | 161324640 | -1.8048 | | 2.20E-13 | IIIVmrMLM | SIG |
| PH_18ZMD | AX-108760312 | 7A | 68024239 | -4.1013 | | 1.57E-09 | BLUPmrMLM | SIG |
| PH_18ZMD | AX-111759717 | 7A | 198671663 | 8.2841 | | 1.43E-08 | FarmCPU | SIG |
| PH_18ZMD | AX-95633120 | 7A | 237517975 | 1.8035 | | 2.29E-13 | IIIVmrMLM | SIG |
| PH_18ZMD | AX-111050220 | 7A | 256816439 | 2.2485 | | 1.42E-04 | BLUPmrMLM | SUG |
| PH_18ZMD | AX-95124326 | 7A | 505017486 | -57.3670 | | 3.68E-07 | EMMAX | SUG |
| PH_18ZMD | AX-111688785 | 7A | 710995296 | -2.4318 | 2.3629 | 3.46E-21 | IIIVmrMLM | SIG |
| PH_18ZMD | AX-94509715 | 7B | 445668577 | -57.3670 | | 3.68E-07 | EMMAX | SUG |
| PH_18ZMD | AX-94502864 | 7B | 591399395 | -3.1225 | 1.0405 | 8.74E-12 | IIIVmrMLM | SIG |
| PH_18ZMD | AX-111473859 | 7B | 609406824 | 2.5420 | | 9.61E-07 | BLUPmrMLM | SUG |
| PH_18ZMD | AX-94740041 | 7B | 670253799 | -0.2391 | 1.7075 | 4.39E-04 | IIIVmrMLM | SUG |
| PH_18ZMD | AX-108855434 | 7B | 704825109 | 1.1315 | | 7.59E-05 | BLUPmrMLM | SUG |
| PH_18ZMD | AX-110468801 | 7D | 32160814 | -26.4720 | | 2.10E-06 | EMMAX | SUG |
| PH_18ZMD | AX-110814077 | 7D | 255221053 | 6.3899 | 2.4253 | 1.02E-80 | IIIVmrMLM | SIG |
| PH_BLUP | AX-111583730 | 1A | 14072756 | -0.8378 | -0.4436 | 2.17E-09 | IIIVmrMLM | SIG |
| PH_BLUP | AX-110985598 | 1A | 494728164 | 2.0377 | -0.3267 | 7.52E-44 | IIIVmrMLM | SIG |
| PH_BLUP | AX-109472932 | 1A | 569447719 | 1.1734 | | 5.05E-07 | BLUPmrMLM | SUG |
| PH_BLUP | AX-109472932 | 1A | 569447719 | -1.4378 | 0.2413 | 1.27E-25 | IIIVmrMLM | SIG |
| PH_BLUP | AX-109323248 | 1A | 574935647 | 1.1394 | 1.2539 | 3.17E-17 | IIIVmrMLM | SIG |
| PH_BLUP | AX-94861274 | 1B | 10069740 | -1.6806 | | 6.06E-06 | BLUPmrMLM | SUG |
| PH_BLUP | AX-95250058 | 1B | 16672239 | -1.1434 | | 1.55E-04 | BLUPmrMLM | SUG |
| PH_BLUP | AX-110057531 | 1B | 17068225 | 2.2059 | 0.1624 | 1.37E-49 | IIIVmrMLM | SIG |
| PH_BLUP | AX-111217304 | 1B | 26260503 | -1.1922 | | 4.31E-06 | BLUPmrMLM | SUG |
| PH_BLUP | AX-108738714 | 1B | 90991683 | -2.3536 | | 6.81E-07 | BLUPmrMLM | SUG |
| PH_BLUP | AX-108775679 | 1B | 377830921 | 1.1979 | | 2.57E-05 | BLUPmrMLM | SUG |
| PH_BLUP | AX-111695254 | 1B | 562468363 | 0.7342 | | 1.50E-04 | BLUPmrMLM | SUG |

| | | | | | | | | |
|---|---|---|---|---|---|---|---|---|
| PH_BLUP | AX-89322086 | 1B | 660531516 | 0.0115 | 12.1477 | 6.37E-04 | IIIVmrMLM | SUG |
| PH_BLUP | AX-111038951 | 1B | 660565770 | -0.0067 | 12.1477 | 6.39E-04 | IIIVmrMLM | SUG |
| PH_BLUP | AX-111030152 | 2A | 31820968 | -5.5468 | | 9.34E-09 | FarmCPU | SIG |
| PH_BLUP | AX-110914499 | 2A | 485779404 | -9.2586 | | 6.09E-07 | EMMAX | SUG |
| PH_BLUP | AX-110914499 | 2A | 485779404 | 8.9732 | | 1.29E-06 | GEMMA | SUG |
| PH_BLUP | AX-111633619 | 2A | 496352754 | -9.3354 | | 7.76E-07 | EMMAX | SUG |
| PH_BLUP | AX-111633619 | 2A | 496352754 | 9.1992 | | 8.75E-07 | GEMMA | SUG |
| PH_BLUP | AX-109536075 | 2A | 676246515 | -0.7759 | | 1.16E-04 | BLUPmrMLM | SUG |
| PH_BLUP | AX-111794227 | 2A | 693038669 | -1.5550 | | 4.37E-31 | IIIVmrMLM | SIG |
| PH_BLUP | AX-108730116 | 2A | 721573234 | -0.2802 | 8.1535 | 3.59E-10 | IIIVmrMLM | SIG |
| PH_BLUP | AX-110666955 | 2B | 29001163 | -2.8064 | -4.5992 | 9.68E-109 | IIIVmrMLM | SIG |
| PH_BLUP | AX-109459500 | 2B | 117106337 | -1.0910 | | 2.16E-06 | BLUPmrMLM | SUG |
| PH_BLUP | AX-110093244 | 2B | 150937460 | -0.6882 | 1.7939 | 1.89E-17 | IIIVmrMLM | SIG |
| PH_BLUP | AX-111218213 | 2D | 20190462 | 0.2091 | 2.3391 | 1.48E-33 | IIIVmrMLM | SIG |
| PH_BLUP | AX-109844299 | 2D | 491947833 | -5.6475 | | 4.75E-07 | EMMAX | SUG |
| PH_BLUP | AX-109844299 | 2D | 491947833 | 5.6878 | | 2.54E-07 | GEMMA | SUG |
| PH_BLUP | AX-110552619 | 2D | 509540852 | 1.1615 | | 7.63E-06 | BLUPmrMLM | SUG |
| PH_BLUP | AX-94427799 | 2D | 523357562 | -7.1534 | | 6.49E-09 | EMMAX | SIG |
| PH_BLUP | AX-94427799 | 2D | 523357562 | 7.1177 | | 5.18E-09 | GEMMA | SIG |
| PH_BLUP | AX-94427799 | 2D | 523357562 | -1.8578 | 3.4603 | 7.97E-41 | IIIVmrMLM | SIG |
| PH_BLUP | AX-94452673 | 3A | 7928453 | -2.9988 | -0.483 | 1.77E-42 | IIIVmrMLM | SIG |
| PH_BLUP | AX-94543536 | 3A | 10217238 | -2.5218 | | 9.82E-08 | BLUPmrMLM | SIG |
| PH_BLUP | AX-111493296 | 3A | 19503535 | -0.7075 | | 1.39E-04 | BLUPmrMLM | SUG |
| PH_BLUP | AX-109313160 | 3A | 23369035 | -0.3014 | 2.3955 | 1.54E-28 | IIIVmrMLM | SIG |
| PH_BLUP | AX-110519127 | 3A | 58837750 | 0.1619 | 15.8356 | 1.31E-11 | IIIVmrMLM | SIG |
| PH_BLUP | AX-110467557 | 3A | 507462215 | -3.3405 | | 2.11E-13 | BLUPmrMLM | SIG |
| PH_BLUP | AX-110467557 | 3A | 507462215 | 4.4036 | | 7.78E-07 | EMMAX | SUG |
| PH_BLUP | AX-110467557 | 3A | 507462215 | -4.3965 | | 5.12E-07 | GEMMA | SUG |
| PH_BLUP | AX-111495554 | 3A | 510693632 | -0.2416 | -2.5975 | 1.28E-13 | IIIVmrMLM | SIG |
| PH_BLUP | AX-111512557 | 3A | 512859054 | 1.6552 | 0.2023 | 1.83E-29 | IIIVmrMLM | SIG |
| PH_BLUP | AX-108761792 | 3A | 549266002 | 2.2012 | -0.1038 | 5.09E-52 | IIIVmrMLM | SIG |
| PH_BLUP | AX-110516751 | 3A | 587558820 | 1.1721 | | 8.25E-06 | BLUPmrMLM | SUG |
| PH_BLUP | AX-94714911 | 3A | 682976438 | -0.7974 | -2.587 | 4.91E-10 | IIIVmrMLM | SIG |
| PH_BLUP | AX-89364000 | 3A | 688683634 | -1.8500 | | 1.51E-06 | BLUPmrMLM | SUG |
| PH_BLUP | AX-110491100 | 3A | 705251779 | -7.5099 | | 4.25E-07 | EMMAX | SUG |
| PH_BLUP | AX-110491100 | 3A | 705251779 | 7.4814 | | 2.83E-07 | GEMMA | SUG |
| PH_BLUP | AX-108941177 | 3A | 705259848 | -7.8014 | | 2.72E-08 | EMMAX | SIG |
| PH_BLUP | AX-108941177 | 3A | 705259848 | 7.7708 | | 1.63E-08 | GEMMA | SIG |
| PH_BLUP | AX-111695263 | 3A | 705274739 | -7.5469 | | 1.38E-07 | EMMAX | SUG |
| PH_BLUP | AX-111695263 | 3A | 705274739 | 7.5382 | | 8.23E-08 | GEMMA | SIG |
| PH_BLUP | AX-110976174 | 3A | 705274782 | -14.3190 | | 3.25E-07 | EMMAX | SUG |
| PH_BLUP | AX-110976174 | 3A | 705274782 | 14.1824 | | 2.56E-07 | GEMMA | SUG |
| PH_BLUP | AX-110484538 | 3A | 705296840 | -18.2620 | | 7.58E-09 | EMMAX | SIG |

| | | | | | | | | |
|---|---|---|---|---|---|---|---|---|
| PH_BLUP | AX-110484538 | 3A | 705296840 | 18.1072 | | 4.72E-09 | GEMMA | SIG |
| PH_BLUP | AX-111483830 | 3A | 718947465 | 0.7741 | 4.6144 | 1.31E-13 | IIIVmrMLM | SIG |
| PH_BLUP | AX-94631122 | 3A | 727224468 | -0.6832 | -3.794 | 2.93E-42 | IIIVmrMLM | SIG |
| PH_BLUP | AX-111043811 | 3B | 41983207 | -0.9566 | | 5.23E-05 | BLUPmrMLM | SUG |
| PH_BLUP | AX-108778384 | 3B | 63399491 | -0.5325 | 3.9742 | 5.61E-15 | IIIVmrMLM | SIG |
| PH_BLUP | AX-109431668 | 3B | 120569866 | -0.9309 | 0.4869 | 6.64E-11 | IIIVmrMLM | SIG |
| PH_BLUP | AX-111725339 | 3B | 203343072 | -1.3988 | -3.551 | 4.68E-52 | IIIVmrMLM | SIG |
| PH_BLUP | AX-110711989 | 3B | 269949176 | 6.5940 | | 1.21E-06 | EMMAX | SUG |
| PH_BLUP | AX-110711989 | 3B | 269949176 | -6.5459 | | 1.38E-06 | GEMMA | SUG |
| PH_BLUP | AX-108941515 | 3B | 469534228 | 1.4185 | | 2.41E-08 | BLUPmrMLM | SIG |
| PH_BLUP | AX-111150856 | 3B | 542680107 | -1.3955 | | 1.83E-08 | BLUPmrMLM | SIG |
| PH_BLUP | AX-110375698 | 3B | 546063769 | 1.6352 | | 1.66E-09 | BLUPmrMLM | SIG |
| PH_BLUP | AX-110420253 | 3B | 713545679 | -0.5541 | -7.1002 | 8.84E-22 | IIIVmrMLM | SIG |
| PH_BLUP | AX-111760526 | 3B | 761709231 | -0.5544 | | 9.97E-05 | BLUPmrMLM | SUG |
| PH_BLUP | AX-109292211 | 3B | 780616084 | -4.3993 | | 2.03E-06 | FarmCPU | SUG |
| PH_BLUP | AX-109270557 | 3D | 571821753 | 1.0271 | | 1.89E-06 | BLUPmrMLM | SUG |
| PH_BLUP | AX-109913089 | 4A | 24900535 | 0.2617 | -2.281 | 2.37E-09 | IIIVmrMLM | SIG |
| PH_BLUP | AX-109380181 | 4B | 25421015 | -2.8653 | | 1.53E-11 | BLUPmrMLM | SIG |
| PH_BLUP | AX-109434530 | 4B | 93539612 | 3.9923 | | 2.03E-130 | IIIVmrMLM | SIG |
| PH_BLUP | AX-111107034 | 4B | 392372140 | -0.9970 | | 1.09E-04 | BLUPmrMLM | SUG |
| PH_BLUP | AX-108944764 | 4D | 15911901 | 4.7103 | | 5.47E-07 | EMMAX | SUG |
| PH_BLUP | AX-108944764 | 4D | 15911901 | -4.7516 | | 2.65E-07 | GEMMA | SUG |
| PH_BLUP | AX-89593703 | 4D | 15958106 | 5.2885 | | 9.86E-09 | EMMAX | SIG |
| PH_BLUP | AX-89593703 | 4D | 15958106 | -5.4976 | | 1.16E-09 | GEMMA | SIG |
| PH_BLUP | AX-89703298 | 4D | 16926596 | 7.0787 | | 5.71E-12 | EMMAX | SIG |
| PH_BLUP | AX-89703298 | 4D | 16926596 | -7.2711 | | 4.67E-13 | GEMMA | SIG |
| PH_BLUP | AX-109281717 | 4D | 17613651 | 7.4239 | | 1.49E-12 | EMMAX | SIG |
| PH_BLUP | AX-109281717 | 4D | 17613651 | -7.6204 | | 9.42E-14 | GEMMA | SIG |
| PH_BLUP | AX-109665328 | 4D | 18539670 | 6.5216 | | 8.29E-11 | EMMAX | SIG |
| PH_BLUP | AX-109665328 | 4D | 18539670 | -6.9153 | | 3.28E-12 | GEMMA | SIG |
| PH_BLUP | AX-108916749 | 4D | 19090268 | 6.0069 | | 5.72E-12 | EMMAX | SIG |
| PH_BLUP | AX-108916749 | 4D | 19090268 | -6.1046 | | 8.72E-13 | GEMMA | SIG |
| PH_BLUP | AX-89421921 | 4D | 19288898 | 7.3929 | | 1.31E-12 | EMMAX | SIG |
| PH_BLUP | AX-89421921 | 4D | 19288898 | -7.5848 | | 8.66E-14 | GEMMA | SIG |
| PH_BLUP | AX-89421921 | 4D | 19288898 | 5.4539 | 1.1601 | 1.05E-185 | IIIVmrMLM | SIG |
| PH_BLUP | AX-89445201 | 4D | 19459579 | -3.4208 | | 4.12E-15 | BLUPmrMLM | SIG |
| PH_BLUP | AX-89445201 | 4D | 19459579 | 4.6051 | | 1.41E-07 | EMMAX | SUG |
| PH_BLUP | AX-89445201 | 4D | 19459579 | -4.8787 | | 1.65E-08 | GEMMA | SIG |
| PH_BLUP | AX-110047101 | 4D | 224594707 | -54.6480 | | 7.57E-12 | EMMAX | SIG |
| PH_BLUP | AX-95186747 | 4D | 446428878 | -44.6130 | | 3.31E-12 | EMMAX | SIG |
| PH_BLUP | AX-110387369 | 4D | 467431300 | -37.9600 | | 5.44E-09 | EMMAX | SIG |
| PH_BLUP | AX-95684258 | 5A | 2122207 | 2.6091 | | 4.91E-08 | BLUPmrMLM | SIG |

| | | | | | | | | |
|---|---|---|---|---|---|---|---|---|
| PH_BLUP | AX-110485186 | 5A | 8120257 | -0.3384 | 3.0286 | 9.98E-12 | IIIVmrMLM | SIG |
| PH_BLUP | AX-108884355 | 5A | 9656059 | -2.6574 | 4.9242 | 7.48E-75 | IIIVmrMLM | SIG |
| PH_BLUP | AX-109324359 | 5A | 518296146 | -1.1270 | | 2.35E-06 | BLUPmrMLM | SUG |
| PH_BLUP | AX-109447549 | 5A | 537015637 | 1.3493 | 5.0642 | 1.02E-23 | IIIVmrMLM | SIG |
| PH_BLUP | AX-110477265 | 5A | 579166659 | 1.3818 | -0.6767 | 3.43E-22 | IIIVmrMLM | SIG |
| PH_BLUP | AX-110042101 | 5A | 584413423 | -0.0176 | -1.8137 | 1.89E-12 | IIIVmrMLM | SIG |
| PH_BLUP | AX-109842338 | 5A | 641048380 | 1.8523 | -1.4607 | 2.25E-40 | IIIVmrMLM | SIG |
| PH_BLUP | AX-110714077 | 5A | 697053429 | -2.8950 | | 2.62E-12 | BLUPmrMLM | SIG |
| PH_BLUP | AX-108952168 | 5A | 698619167 | -0.2997 | -3.9246 | 1.28E-10 | IIIVmrMLM | SIG |
| PH_BLUP | AX-110417678 | 5A | 701292911 | 8.8215 | | 6.97E-07 | EMMAX | SUG |
| PH_BLUP | AX-110417678 | 5A | 701292911 | -8.8549 | | 6.17E-07 | GEMMA | SUG |
| PH_BLUP | AX-109967112 | 5B | 404684284 | -1.9956 | -0.6109 | 6.22E-43 | IIIVmrMLM | SIG |
| PH_BLUP | AX-111156055 | 5B | 416156777 | -2.2347 | | 1.67E-13 | BLUPmrMLM | SIG |
| PH_BLUP | AX-108755343 | 5B | 459492140 | 1.7066 | 2.4177 | 1.26E-35 | IIIVmrMLM | SIG |
| PH_BLUP | AX-109975159 | 5B | 527774613 | -3.4404 | | 5.22E-10 | BLUPmrMLM | SIG |
| PH_BLUP | AX-110018488 | 5B | 528535571 | 6.0067 | | 2.45E-07 | EMMAX | SUG |
| PH_BLUP | AX-110018488 | 5B | 528535571 | -6.0829 | | 1.03E-07 | GEMMA | SIG |
| PH_BLUP | AX-110018488 | 5B | 528535571 | 4.5609 | | 1.64E-153 | IIIVmrMLM | SIG |
| PH_BLUP | AX-94782021 | 5B | 531583413 | 1.6968 | | 1.03E-05 | BLUPmrMLM | SUG |
| PH_BLUP | AX-109452394 | 5B | 536518634 | 1.3611 | | 5.13E-06 | BLUPmrMLM | SUG |
| PH_BLUP | AX-108944059 | 5B | 536843401 | 0.7847 | 2.6639 | 1.42E-16 | IIIVmrMLM | SIG |
| PH_BLUP | AX-108879525 | 5B | 566972413 | -0.1895 | -3.4384 | 1.21E-10 | IIIVmrMLM | SIG |
| PH_BLUP | AX-86170796 | 5D | 3609894 | -1.2274 | | 5.24E-07 | BLUPmrMLM | SUG |
| PH_BLUP | AX-110930042 | 5D | 362887848 | -39.0800 | | 2.35E-11 | EMMAX | SIG |
| PH_BLUP | AX-94963508 | 5D | 484426813 | -54.6480 | | 7.57E-12 | EMMAX | SIG |
| PH_BLUP | AX-109471937 | 6A | 23602043 | -1.6466 | | 3.52E-06 | BLUPmrMLM | SUG |
| PH_BLUP | AX-95165798 | 6A | 46975506 | -1.3035 | | 8.94E-08 | BLUPmrMLM | SIG |
| PH_BLUP | AX-109970551 | 6A | 330915971 | 1.3326 | -0.0455 | 1.80E-22 | IIIVmrMLM | SIG |
| PH_BLUP | AX-108758970 | 6B | 47137302 | -0.7864 | | 8.23E-05 | BLUPmrMLM | SUG |
| PH_BLUP | AX-109847757 | 6B | 67109005 | 1.0830 | | 1.05E-16 | IIIVmrMLM | SIG |
| PH_BLUP | AX-86173709 | 6B | 136114962 | 1.3256 | | 5.34E-08 | BLUPmrMLM | SIG |
| PH_BLUP | AX-110432637 | 6B | 136498311 | -1.1680 | 0.6384 | 1.22E-17 | IIIVmrMLM | SIG |
| PH_BLUP | AX-110570556 | 6B | 521446569 | 0.2383 | 1.2637 | 1.10E-09 | IIIVmrMLM | SIG |
| PH_BLUP | AX-108796810 | 6B | 525541129 | -1.3721 | | 5.92E-08 | BLUPmrMLM | SIG |
| PH_BLUP | AX-110925969 | 6B | 655780737 | -0.3652 | -3.1419 | 1.29E-13 | IIIVmrMLM | SIG |
| PH_BLUP | AX-109364183 | 6B | 694461959 | -9.8047 | | 1.70E-07 | FarmCPU | SUG |
| PH_BLUP | AX-94509919 | 6B | 707382514 | 0.6545 | | 1.29E-05 | BLUPmrMLM | SUG |
| PH_BLUP | AX-94803991 | 6D | 2246424 | -0.8982 | | 3.85E-12 | IIIVmrMLM | SIG |
| PH_BLUP | AX-111560737 | 6D | 2716398 | -0.4921 | | 7.01E-05 | BLUPmrMLM | SUG |
| PH_BLUP | AX-110898999 | 6D | 9020295 | 0.3651 | 7.3098 | 8.49E-05 | IIIVmrMLM | SUG |
| PH_BLUP | AX-94745565 | 6D | 25760471 | 2.7202 | | 1.12E-76 | IIIVmrMLM | SIG |
| PH_BLUP | AX-95007223 | 6D | 36801236 | -0.6502 | | 3.94E-07 | IIIVmrMLM | SUG |
| PH_BLUP | AX-109897288 | 6D | 194038026 | 1.6010 | | 2.83E-06 | BLUPmrMLM | SUG |

| | | | | | | | | |
|---|---|---|---|---|---|---|---|---|
| PH_BLUP | AX-109435951 | 6D | 469993277 | 0.6327 | | 2.41E-05 | BLUPmrMLM | SUG |
| PH_BLUP | AX-94642959 | 7A | 14345684 | 0.5895 | | 4.09E-06 | IIIVmrMLM | SUG |
| PH_BLUP | AX-109484742 | 7A | 17455949 | 1.3181 | | 1.64E-10 | BLUPmrMLM | SIG |
| PH_BLUP | AX-111692733 | 7A | 54820735 | 0.0711 | 4.8112 | 9.09E-06 | IIIVmrMLM | SUG |
| PH_BLUP | AX-108914852 | 7A | 237935325 | -2.0467 | | 1.58E-04 | BLUPmrMLM | SUG |
| PH_BLUP | AX-111050220 | 7A | 256816439 | 1.7516 | | 5.89E-09 | BLUPmrMLM | SIG |
| PH_BLUP | AX-111605089 | 7A | 538250230 | -0.0847 | 8.5768 | 4.11E-07 | IIIVmrMLM | SUG |
| PH_BLUP | AX-109583446 | 7A | 652602859 | -0.8507 | 0.5257 | 2.24E-08 | IIIVmrMLM | SIG |
| PH_BLUP | AX-111460480 | 7A | 688619689 | -1.3062 | | 7.60E-06 | BLUPmrMLM | SUG |
| PH_BLUP | AX-108874534 | 7A | 703703465 | -0.1763 | -3.2769 | 4.62E-05 | IIIVmrMLM | SUG |
| PH_BLUP | AX-108962229 | 7A | 708912808 | -1.4512 | -0.3677 | 2.20E-25 | IIIVmrMLM | SIG |
| PH_BLUP | AX-94794309 | 7B | 3795409 | 5.4943 | | 3.84E-09 | BLUPmrMLM | SIG |
| PH_BLUP | AX-111079048 | 7B | 50297875 | 1.2110 | | 6.43E-05 | BLUPmrMLM | SUG |
| PH_BLUP | AX-108784045 | 7B | 160448849 | -1.6831 | | 1.07E-12 | BLUPmrMLM | SIG |
| PH_BLUP | AX-111505923 | 7B | 715541984 | -1.0202 | -0.2618 | 9.53E-12 | IIIVmrMLM | SIG |
| PH_BLUP | AX-112286117 | 7B | 716041172 | -9.0002 | | 1.04E-06 | EMMAX | SUG |
| PH_BLUP | AX-112286117 | 7B | 716041172 | 8.9711 | | 7.62E-07 | GEMMA | SUG |
| PH_BLUP | AX-94654805 | 7D | 27097795 | -29.6980 | | 2.55E-07 | EMMAX | SUG |
| PH_BLUP | AX-111713225 | 7D | 71863395 | -2.4508 | | 3.73E-10 | BLUPmrMLM | SIG |
| PH_BLUP | AX-94997278 | 7D | 74467098 | 3.3496 | | 3.31E-11 | BLUPmrMLM | SIG |
| PH_BLUP | AX-94997278 | 7D | 74467098 | -1.5079 | 0.2814 | 2.13E-08 | IIIVmrMLM | SIG |
| PH_BLUP | AX-111013473 | 7D | 104628775 | -1.2530 | | 1.33E-07 | BLUPmrMLM | SUG |
| PH_BLUP | AX-110925192 | 7D | 197297065 | -25.1930 | | 8.78E-07 | EMMAX | SUG |

PH: plant height

**Supplementary Table S5**. All QTNs identified as associated with spike length in wheat.

| Trait | Marker | Chr | Position | Add.effect | Dom.effect | P-value | Method | Sig. |
|---|---|---|---|---|---|---|---|---|
| SL_17AY | AX-111035740 | 1A | 22345029 | 0.0134 | -0.4522 | 1.42E-05 | IIIVmrMLM | SUG |
| SL_17AY | AX-111726814 | 1A | 504281689 | 0.0132 | 1.1537 | 1.62E-07 | IIIVmrMLM | SUG |
| SL_17AY | AX-109891745 | 1A | 543950795 | -0.1264 | -0.4293 | 1.82E-11 | IIIVmrMLM | SIG |
| SL_17AY | AX-110482470 | 1B | 27282733 | -0.3197 | | 3.12E-08 | GEMMA | SIG |
| SL_17AY | AX-94585083 | 1B | 626085377 | -0.6256 | | 1.91E-04 | BLUPmrMLM | SUG |
| SL_17AY | AX-109393726 | 1B | 668430767 | -0.6958 | | 4.29E-07 | GEMMA | SUG |
| SL_17AY | AX-109926505 | 1B | 668655094 | -0.3429 | | 3.45E-07 | GEMMA | SUG |
| SL_17AY | AX-108877140 | 1B | 680813935 | 0.1809 | -0.0917 | 1.82E-10 | IIIVmrMLM | SIG |
| SL_17AY | AX-94792955 | 1D | 9873097 | -0.0362 | 0.41 | 1.97E-11 | IIIVmrMLM | SIG |
| SL_17AY | AX-111192792 | 1D | 273815679 | 0.3244 | | 3.98E-07 | GEMMA | SUG |
| SL_17AY | AX-95257101 | 1D | 273906197 | 0.5892 | | 2.10E-06 | GEMMA | SUG |
| SL_17AY | AX-109969365 | 1D | 273911025 | 0.3037 | | 1.39E-06 | GEMMA | SUG |
| SL_17AY | AX-109311974 | 2A | 3110094 | 0.3854 | | 1.52E-06 | GEMMA | SUG |
| SL_17AY | AX-108848462 | 2A | 4110206 | -0.0203 | -0.5199 | 1.52E-07 | IIIVmrMLM | SUG |
| SL_17AY | AX-109364347 | 2A | 7301119 | 0.1752 | | 1.61E-04 | BLUPmrMLM | SUG |
| SL_17AY | AX-108820240 | 2A | 36379654 | -0.0067 | -1.3857 | 2.00E-09 | IIIVmrMLM | SIG |
| SL_17AY | AX-109856737 | 2A | 90504929 | 0.0574 | -0.3856 | 5.68E-08 | IIIVmrMLM | SIG |
| SL_17AY | AX-110547358 | 2A | 183025587 | -0.0388 | -0.6239 | 1.45E-04 | IIIVmrMLM | SUG |
| SL_17AY | AX-109427494 | 2A | 196289225 | 0.0828 | -0.5593 | 1.65E-08 | IIIVmrMLM | SIG |
| SL_17AY | AX-109044308 | 2A | 683709401 | -0.3660 | | 1.23E-05 | BLUPmrMLM | SUG |
| SL_17AY | AX-109044308 | 2A | 683709401 | 0.2025 | | 6.59E-14 | IIIVmrMLM | SIG |
| SL_17AY | AX-111230613 | 2B | 50740746 | 0.0151 | 0.3878 | 1.43E-11 | IIIVmrMLM | SIG |
| SL_17AY | AX-109022733 | 2B | 109932031 | -0.1271 | | 1.84E-04 | BLUPmrMLM | SUG |
| SL_17AY | AX-109356749 | 2B | 116762983 | 0.0982 | 0.7524 | 3.11E-09 | IIIVmrMLM | SIG |
| SL_17AY | AX-110460935 | 2B | 767897927 | 0.0028 | 0.8118 | 9.10E-06 | IIIVmrMLM | SUG |
| SL_17AY | AX-110010922 | 2B | 779367412 | 0.1278 | 0.3329 | 4.25E-06 | IIIVmrMLM | SUG |
| SL_17AY | AX-94398733 | 2D | 523625859 | 0.3152 | | 1.11E-29 | IIIVmrMLM | SIG |
| SL_17AY | AX-109318386 | 3A | 682359154 | 0.3355 | | 4.95E-07 | GEMMA | SUG |
| SL_17AY | AX-110081775 | 3A | 682360038 | 0.3368 | | 5.15E-07 | GEMMA | SUG |
| SL_17AY | AX-109835734 | 3A | 682360670 | 0.3368 | | 5.15E-07 | GEMMA | SUG |
| SL_17AY | AX-111528868 | 3A | 682361343 | 0.3244 | | 1.28E-06 | GEMMA | SUG |
| SL_17AY | AX-111534220 | 3A | 682361650 | 0.3244 | | 1.28E-06 | GEMMA | SUG |
| SL_17AY | AX-109543005 | 3A | 682363552 | 0.3244 | | 1.28E-06 | GEMMA | SUG |
| SL_17AY | AX-110467047 | 3A | 682366220 | 0.3244 | | 1.28E-06 | GEMMA | SUG |
| SL_17AY | AX-111092799 | 3A | 682367130 | 0.3302 | | 8.80E-07 | GEMMA | SUG |
| SL_17AY | AX-110477879 | 3A | 682426565 | 0.3409 | | 4.20E-07 | GEMMA | SUG |
| SL_17AY | AX-110032485 | 3A | 682482137 | 0.3244 | | 1.28E-06 | GEMMA | SUG |
| SL_17AY | AX-111649590 | 3A | 682577354 | 0.3243 | | 1.25E-06 | GEMMA | SUG |
| SL_17AY | AX-108930803 | 3A | 682600010 | 0.3241 | | 9.12E-07 | GEMMA | SUG |
| SL_17AY | AX-110042634 | 3A | 682600120 | 0.3243 | | 1.25E-06 | GEMMA | SUG |
| SL_17AY | AX-111076184 | 3A | 682649491 | 0.3176 | | 2.07E-06 | GEMMA | SUG |

| | | | | | | | |
|---|---|---|---|---|---|---|---|
| SL_17AY | AX-110410110 | 3A | 682659316 | 0.3243 | 1.25E-06 | GEMMA | SUG |
| SL_17AY | AX-110488147 | 3A | 682659638 | 0.3440 | 3.74E-07 | GEMMA | SUG |
| SL_17AY | AX-111459742 | 3A | 682661654 | 0.3185 | 2.05E-06 | GEMMA | SUG |
| SL_17AY | AX-109639643 | 3A | 682711980 | 0.3243 | 1.25E-06 | GEMMA | SUG |
| SL_17AY | AX-109595870 | 3A | 682712836 | 0.3230 | 1.40E-06 | GEMMA | SUG |
| SL_17AY | AX-111657715 | 3A | 682733325 | 0.3243 | 1.25E-06 | GEMMA | SUG |
| SL_17AY | AX-111583020 | 3A | 682733436 | 0.3243 | 1.25E-06 | GEMMA | SUG |
| SL_17AY | AX-108796495 | 3A | 682733623 | 0.3243 | 1.25E-06 | GEMMA | SUG |
| SL_17AY | AX-110413799 | 3A | 682733669 | 0.3243 | 1.25E-06 | GEMMA | SUG |
| SL_17AY | AX-111652000 | 3A | 682745804 | 0.3322 | 5.54E-07 | GEMMA | SUG |
| SL_17AY | AX-111476952 | 3A | 682746359 | 0.3243 | 1.25E-06 | GEMMA | SUG |
| SL_17AY | AX-109946485 | 3A | 682746465 | 0.3243 | 1.25E-06 | GEMMA | SUG |
| SL_17AY | AX-110978118 | 3A | 682746510 | 0.3339 | 7.47E-07 | GEMMA | SUG |
| SL_17AY | AX-109828410 | 3A | 682746575 | 0.3275 | 1.20E-06 | GEMMA | SUG |
| SL_17AY | AX-110428166 | 3A | 682748332 | 0.3301 | 8.59E-07 | GEMMA | SUG |
| SL_17AY | AX-111058941 | 3A | 682748426 | 0.3243 | 1.25E-06 | GEMMA | SUG |
| SL_17AY | AX-111087111 | 3A | 682782228 | 0.3243 | 1.25E-06 | GEMMA | SUG |
| SL_17AY | AX-111450715 | 3A | 682939904 | 0.3301 | 8.59E-07 | GEMMA | SUG |
| SL_17AY | AX-110496222 | 3A | 682940185 | 0.3245 | 1.21E-06 | GEMMA | SUG |
| SL_17AY | AX-111548921 | 3A | 682940818 | 0.3214 | 1.52E-06 | GEMMA | SUG |
| SL_17AY | AX-110972183 | 3A | 682971074 | 0.3243 | 1.25E-06 | GEMMA | SUG |
| SL_17AY | AX-108927537 | 3A | 682974852 | 0.3274 | 9.87E-07 | GEMMA | SUG |
| SL_17AY | AX-110945844 | 3A | 682978148 | 0.3392 | 6.35E-07 | GEMMA | SUG |
| SL_17AY | AX-111457327 | 3A | 682987484 | 0.3104 | 1.20E-06 | GEMMA | SUG |
| SL_17AY | AX-109441500 | 3A | 682992689 | -0.2616 | 1.32E-06 | GEMMA | SUG |
| SL_17AY | AX-109919209 | 3A | 682995323 | -0.2619 | 1.53E-07 | GEMMA | SUG |
| SL_17AY | AX-109889130 | 3A | 683037703 | -0.2695 | 1.38E-07 | GEMMA | SUG |
| SL_17AY | AX-109392160 | 3A | 683062079 | -0.2612 | 1.61E-07 | GEMMA | SUG |
| SL_17AY | AX-109316757 | 3A | 683066117 | -0.2636 | 1.79E-07 | GEMMA | SUG |
| SL_17AY | AX-95232814 | 3A | 683071252 | -0.5138 | 1.88E-07 | GEMMA | SUG |
| SL_17AY | AX-111478900 | 3A | 683181728 | -0.2642 | 1.07E-07 | GEMMA | SIG |
| SL_17AY | AX-109823297 | 3A | 683186492 | -0.2414 | 9.40E-07 | GEMMA | SUG |
| SL_17AY | AX-95132491 | 3A | 683187888 | -0.2714 | 6.93E-08 | GEMMA | SIG |
| SL_17AY | AX-89533807 | 3A | 683192534 | -0.2648 | 1.55E-07 | GEMMA | SUG |
| SL_17AY | AX-94455123 | 3A | 683193285 | -0.2701 | 7.86E-08 | GEMMA | SIG |
| SL_17AY | AX-109311882 | 3A | 683194731 | -0.2928 | 1.96E-08 | GEMMA | SIG |
| SL_17AY | AX-95658166 | 3A | 683195193 | -0.2701 | 7.86E-08 | GEMMA | SIG |
| SL_17AY | AX-86184063 | 3A | 683249793 | -0.2830 | 2.18E-08 | GEMMA | SIG |
| SL_17AY | AX-110458513 | 3A | 683259201 | -0.2831 | 1.98E-08 | GEMMA | SIG |
| SL_17AY | AX-110188010 | 3A | 683261490 | -0.2880 | 1.34E-08 | GEMMA | SIG |
| SL_17AY | AX-109989456 | 3A | 683266127 | -0.2868 | 1.76E-08 | GEMMA | SIG |
| SL_17AY | AX-109647843 | 3A | 683297803 | -0.3692 | 2.07E-06 | GEMMA | SUG |
| SL_17AY | AX-110076300 | 3A | 683354584 | -0.2855 | 1.68E-08 | GEMMA | SIG |
| SL_17AY | AX-108981334 | 3A | 683354722 | -0.2790 | 3.32E-08 | GEMMA | SIG |

| | | | | | | | | |
|---|---|---|---|---|---|---|---|---|
| SL_17AY | AX-111008285 | 3A | 683359493 | -0.2623 | | 7.21E-07 | GEMMA | SUG |
| SL_17AY | AX-95219519 | 3A | 683364619 | -0.6044 | | 3.15E-08 | GEMMA | SIG |
| SL_17AY | AX-108999044 | 3A | 683364762 | -0.3027 | | 3.63E-08 | GEMMA | SIG |
| SL_17AY | AX-86177295 | 3A | 683515653 | -0.3026 | | 8.15E-08 | GEMMA | SIG |
| SL_17AY | AX-109276728 | 3A | 683516864 | -0.3001 | | 4.59E-08 | GEMMA | SIG |
| SL_17AY | AX-110655327 | 3A | 683518749 | -0.5888 | | 6.17E-08 | GEMMA | SIG |
| SL_17AY | AX-110495706 | 3A | 683519275 | -0.2969 | | 5.93E-08 | GEMMA | SIG |
| SL_17AY | AX-109827120 | 3A | 683524484 | -0.2998 | | 4.35E-08 | GEMMA | SIG |
| SL_17AY | AX-110922577 | 3A | 683573600 | 0.2914 | | 1.41E-06 | EMMAX | SUG |
| SL_17AY | AX-110922577 | 3A | 683573600 | -0.3065 | | 4.88E-09 | GEMMA | SIG |
| SL_17AY | AX-111561745 | 3A | 684045758 | 0.6385 | | 1.25E-07 | EMMAX | SUG |
| SL_17AY | AX-111561745 | 3A | 684045758 | -0.6637 | | 2.35E-10 | GEMMA | SIG |
| SL_17AY | AX-111099482 | 3A | 684291195 | 0.2926 | | 1.37E-06 | EMMAX | SUG |
| SL_17AY | AX-111099482 | 3A | 684291195 | -0.3066 | | 4.00E-09 | GEMMA | SIG |
| SL_17AY | AX-110904850 | 3A | 684294910 | -0.3084 | | 9.04E-09 | GEMMA | SIG |
| SL_17AY | AX-109295307 | 3A | 684296831 | 0.3085 | | 4.04E-07 | EMMAX | SUG |
| SL_17AY | AX-109295307 | 3A | 684296831 | -0.3224 | | 8.04E-10 | GEMMA | SIG |
| SL_17AY | AX-112288904 | 3A | 684297263 | 0.3610 | | 9.89E-08 | EMMAX | SIG |
| SL_17AY | AX-112288904 | 3A | 684297263 | -0.3815 | | 1.48E-10 | GEMMA | SIG |
| SL_17AY | AX-95628824 | 3A | 684297438 | 0.3052 | | 5.45E-07 | EMMAX | SUG |
| SL_17AY | AX-95628824 | 3A | 684297438 | -0.3177 | | 1.34E-09 | GEMMA | SIG |
| SL_17AY | AX-109849557 | 3A | 684297680 | 0.3084 | | 4.35E-07 | EMMAX | SUG |
| SL_17AY | AX-109849557 | 3A | 684297680 | -0.3204 | | 1.10E-09 | GEMMA | SIG |
| SL_17AY | AX-109463080 | 3A | 684298993 | 0.3052 | | 5.45E-07 | EMMAX | SUG |
| SL_17AY | AX-109463080 | 3A | 684298993 | -0.3177 | | 1.34E-09 | GEMMA | SIG |
| SL_17AY | AX-108936105 | 3A | 684299388 | 0.6034 | | 6.76E-07 | EMMAX | SUG |
| SL_17AY | AX-108936105 | 3A | 684299388 | -0.6316 | | 1.44E-09 | GEMMA | SIG |
| SL_17AY | AX-110080445 | 3A | 684301379 | 0.3012 | | 7.57E-07 | EMMAX | SUG |
| SL_17AY | AX-110080445 | 3A | 684301379 | -0.3133 | | 2.10E-09 | GEMMA | SIG |
| SL_17AY | AX-110488808 | 3A | 684301883 | -0.3507 | | 1.40E-06 | GEMMA | SUG |
| SL_17AY | AX-109462935 | 3A | 684303011 | 0.4312 | | 7.39E-08 | EMMAX | SIG |
| SL_17AY | AX-109462935 | 3A | 684303011 | -0.4490 | | 1.45E-10 | GEMMA | SIG |
| SL_17AY | AX-108911182 | 3A | 684303851 | -0.2983 | | 3.72E-10 | BLUPmrMLM | SIG |
| SL_17AY | AX-108911182 | 3A | 684303851 | 0.3348 | | 2.46E-08 | EMMAX | SIG |
| SL_17AY | AX-108911182 | 3A | 684303851 | -0.3411 | | 1.27E-10 | GEMMA | SIG |
| SL_17AY | AX-108911182 | 3A | 684303851 | 0.1131 | -0.1991 | 2.50E-05 | IIIVmrMLM | SUG |
| SL_17AY | AX-108780807 | 3A | 684322462 | 0.3077 | | 6.01E-07 | EMMAX | SUG |
| SL_17AY | AX-108780807 | 3A | 684322462 | -0.3207 | | 1.42E-09 | GEMMA | SIG |
| SL_17AY | AX-109493018 | 3A | 684393993 | 0.2999 | | 1.03E-06 | EMMAX | SUG |
| SL_17AY | AX-109493018 | 3A | 684393993 | -0.3106 | | 3.86E-09 | GEMMA | SIG |
| SL_17AY | AX-109884438 | 3A | 684394403 | 0.3052 | | 5.45E-07 | EMMAX | SUG |
| SL_17AY | AX-109884438 | 3A | 684394403 | -0.3177 | | 1.34E-09 | GEMMA | SIG |
| SL_17AY | AX-110722061 | 3A | 684395471 | 0.3052 | | 5.45E-07 | EMMAX | SUG |
| SL_17AY | AX-110722061 | 3A | 684395471 | -0.3177 | | 1.34E-09 | GEMMA | SIG |

| | | | | | | | |
|---|---|---|---|---|---|---|---|
| SL_17AY | AX-110123926 | 3A | 684397823 | 0.6195 | 2.96E-07 | EMMAX | SUG |
| SL_17AY | AX-110123926 | 3A | 684397823 | -0.6455 | 6.19E-10 | GEMMA | SIG |
| SL_17AY | AX-109329838 | 3A | 684578788 | 0.3052 | 5.45E-07 | EMMAX | SUG |
| SL_17AY | AX-109329838 | 3A | 684578788 | -0.3177 | 1.34E-09 | GEMMA | SIG |
| SL_17AY | AX-109991839 | 3A | 684578837 | 0.3052 | 5.45E-07 | EMMAX | SUG |
| SL_17AY | AX-109991839 | 3A | 684578837 | -0.3177 | 1.34E-09 | GEMMA | SIG |
| SL_17AY | AX-110535607 | 3A | 684579307 | 0.3018 | 9.39E-07 | EMMAX | SUG |
| SL_17AY | AX-110535607 | 3A | 684579307 | -0.3147 | 2.51E-09 | GEMMA | SIG |
| SL_17AY | AX-110656725 | 3A | 684579543 | 0.3052 | 5.45E-07 | EMMAX | SUG |
| SL_17AY | AX-110656725 | 3A | 684579543 | -0.3177 | 1.34E-09 | GEMMA | SIG |
| SL_17AY | AX-110637660 | 3A | 684579820 | 0.3037 | 6.73E-07 | EMMAX | SUG |
| SL_17AY | AX-110637660 | 3A | 684579820 | -0.3167 | 1.60E-09 | GEMMA | SIG |
| SL_17AY | AX-111665968 | 3A | 684581252 | 0.3052 | 5.45E-07 | EMMAX | SUG |
| SL_17AY | AX-111665968 | 3A | 684581252 | -0.3177 | 1.34E-09 | GEMMA | SIG |
| SL_17AY | AX-108841461 | 3A | 684584230 | 0.3055 | 5.70E-07 | EMMAX | SUG |
| SL_17AY | AX-108841461 | 3A | 684584230 | -0.3188 | 1.31E-09 | GEMMA | SIG |
| SL_17AY | AX-109324697 | 3A | 684586755 | 0.3052 | 5.45E-07 | EMMAX | SUG |
| SL_17AY | AX-109324697 | 3A | 684586755 | -0.3177 | 1.34E-09 | GEMMA | SIG |
| SL_17AY | AX-108789451 | 3A | 684588483 | 0.3013 | 8.14E-07 | EMMAX | SUG |
| SL_17AY | AX-108789451 | 3A | 684588483 | -0.3141 | 2.09E-09 | GEMMA | SIG |
| SL_17AY | AX-110466677 | 3A | 684592398 | 0.3052 | 5.45E-07 | EMMAX | SUG |
| SL_17AY | AX-110466677 | 3A | 684592398 | -0.3177 | 1.34E-09 | GEMMA | SIG |
| SL_17AY | AX-111170176 | 3A | 684594036 | -0.3287 | 7.86E-09 | GEMMA | SIG |
| SL_17AY | AX-110435652 | 3A | 684594079 | 0.3052 | 5.45E-07 | EMMAX | SUG |
| SL_17AY | AX-110435652 | 3A | 684594079 | -0.3177 | 1.34E-09 | GEMMA | SIG |
| SL_17AY | AX-108747994 | 3A | 684751521 | 0.3118 | 2.69E-07 | EMMAX | SUG |
| SL_17AY | AX-108747994 | 3A | 684751521 | -0.3228 | 7.18E-10 | GEMMA | SIG |
| SL_17AY | AX-110516971 | 3A | 684911677 | 0.3104 | 2.02E-06 | GEMMA | SUG |
| SL_17AY | AX-109831155 | 3A | 685014559 | -0.2277 | 1.31E-06 | GEMMA | SUG |
| SL_17AY | AX-111176368 | 3A | 685124835 | -0.2176 | 1.66E-06 | GEMMA | SUG |
| SL_17AY | AX-86176907 | 3A | 685357896 | -0.2176 | 1.66E-06 | GEMMA | SUG |
| SL_17AY | AX-94396499 | 3A | 685357955 | -0.2176 | 1.66E-06 | GEMMA | SUG |
| SL_17AY | AX-86174490 | 3A | 685360270 | -0.2755 | 2.24E-08 | GEMMA | SIG |
| SL_17AY | AX-109440476 | 3A | 685436902 | -0.2301 | 5.69E-07 | GEMMA | SUG |
| SL_17AY | AX-110545140 | 3A | 685436965 | -0.2262 | 7.55E-07 | GEMMA | SUG |
| SL_17AY | AX-111624763 | 3A | 685437330 | -0.2262 | 7.55E-07 | GEMMA | SUG |
| SL_17AY | AX-109819999 | 3A | 685438888 | -0.2262 | 7.55E-07 | GEMMA | SUG |
| SL_17AY | AX-109313738 | 3A | 685439507 | -0.2262 | 7.55E-07 | GEMMA | SUG |
| SL_17AY | AX-110496125 | 3A | 685442935 | -0.2213 | 1.18E-06 | GEMMA | SUG |
| SL_17AY | AX-111575018 | 3A | 685582352 | 0.3211 | 1.51E-06 | GEMMA | SUG |
| SL_17AY | AX-108753431 | 3A | 685702163 | 0.3348 | 9.18E-08 | GEMMA | SIG |
| SL_17AY | AX-111182359 | 3A | 685780310 | -0.2267 | 7.22E-07 | GEMMA | SUG |
| SL_17AY | AX-109840140 | 3A | 685781388 | -0.2281 | 6.40E-07 | GEMMA | SUG |
| SL_17AY | AX-111802405 | 3A | 686075960 | -0.4179 | 1.96E-06 | GEMMA | SUG |

| | | | | | | | | |
|---|---|---|---|---|---|---|---|---|
| SL_17AY | AX-95003297 | 3A | 686121498 | 0.2724 | | 2.13E-06 | EMMAX | SUG |
| SL_17AY | AX-95003297 | 3A | 686121498 | -0.2844 | | 9.29E-09 | GEMMA | SIG |
| SL_17AY | AX-109065107 | 3A | 686125876 | -0.2194 | | 1.31E-06 | GEMMA | SUG |
| SL_17AY | AX-94432010 | 3B | 349947257 | 1.9502 | | 4.93E-07 | GEMMA | SUG |
| SL_17AY | AX-111035357 | 3B | 557266528 | 0.0457 | -0.3259 | 1.27E-10 | IIIVmrMLM | SIG |
| SL_17AY | AX-94949839 | 3B | 723276180 | -0.5035 | | 2.95E-07 | GEMMA | SUG |
| SL_17AY | AX-110275510 | 3D | 474995279 | -0.2385 | -0.0759 | 2.91E-17 | IIIVmrMLM | SIG |
| SL_17AY | AX-95139890 | 3D | 546339375 | -0.5311 | | 1.33E-07 | GEMMA | SUG |
| SL_17AY | AX-95083270 | 3D | 547627077 | -0.6101 | | 4.61E-07 | EMMAX | SUG |
| SL_17AY | AX-95083270 | 3D | 547627077 | 0.6340 | | 1.21E-09 | GEMMA | SIG |
| SL_17AY | AX-109344376 | 4A | 707314889 | 0.0851 | -0.6374 | 3.49E-06 | IIIVmrMLM | SUG |
| SL_17AY | AX-110068755 | 4B | 1254779 | 0.0384 | 1.3118 | 4.04E-08 | IIIVmrMLM | SIG |
| SL_17AY | AX-109050465 | 4B | 257506502 | -1.7362 | | 6.22E-07 | GEMMA | SUG |
| SL_17AY | AX-110020229 | 4B | 537093052 | 0.0266 | 0.2722 | 9.74E-14 | IIIVmrMLM | SIG |
| SL_17AY | AX-111094296 | 5A | 374629792 | -0.0289 | -0.9953 | 5.94E-09 | IIIVmrMLM | SIG |
| SL_17AY | AX-111603118 | 5B | 66112219 | 0.0849 | -0.3847 | 5.63E-11 | IIIVmrMLM | SIG |
| SL_17AY | AX-110590093 | 5B | 421095571 | -0.0450 | 0.4949 | 3.48E-17 | IIIVmrMLM | SIG |
| SL_17AY | AX-108862689 | 5B | 565771746 | -0.0396 | -0.2728 | 1.43E-06 | IIIVmrMLM | SUG |
| SL_17AY | AX-109111875 | 5D | 318805394 | -0.3555 | | 2.21E-06 | GEMMA | SUG |
| SL_17AY | AX-109218363 | 5D | 344397400 | -0.1860 | | 4.95E-12 | IIIVmrMLM | SIG |
| SL_17AY | AX-110051508 | 5D | 553481210 | 0.1504 | -2.0268 | 5.57E-21 | IIIVmrMLM | SIG |
| SL_17AY | AX-111539735 | 6A | 105129957 | -0.1407 | | 1.37E-07 | IIIVmrMLM | SUG |
| SL_17AY | AX-94764073 | 6B | 23658714 | 0.1701 | | 2.31E-10 | IIIVmrMLM | SIG |
| SL_17AY | AX-109337169 | 6B | 635020099 | 0.1824 | | 1.24E-04 | BLUPmrMLM | SUG |
| SL_17AY | AX-108870504 | 7A | 73732383 | 0.0963 | -0.3254 | 4.09E-09 | IIIVmrMLM | SIG |
| SL_17AY | AX-110717549 | 7A | 549334917 | 0.0258 | 1.489 | 8.01E-10 | IIIVmrMLM | SIG |
| SL_17AY | AX-109904065 | 7A | 645274854 | -0.1128 | 0.6106 | 1.15E-10 | IIIVmrMLM | SIG |
| SL_17AY | AX-110190562 | 7B | 136321516 | -0.0834 | -0.7124 | 1.54E-11 | IIIVmrMLM | SIG |
| SL_17AY | AX-111052202 | 7B | 592194365 | -0.1486 | 0.0633 | 2.83E-07 | IIIVmrMLM | SUG |
| SL_17AY | AX-110515695 | 7D | 37973588 | -0.2082 | | 5.30E-05 | BLUPmrMLM | SUG |
| SL_17AY | AX-110515695 | 7D | 37973588 | -0.3033 | | 1.71E-06 | GEMMA | SUG |
| SL_17AY | AX-110515695 | 7D | 37973588 | 0.1959 | 0.3223 | 1.29E-12 | IIIVmrMLM | SIG |
| SL_17AY | AX-110059680 | 7D | 191107706 | -0.3521 | | 1.70E-06 | GEMMA | SUG |
| SL_17AY | AX-109276948 | 7D | 220973440 | -0.3830 | | 1.07E-07 | GEMMA | SIG |
| SL_17AY | AX-111891203 | 7D | 221698858 | -0.3596 | | 4.63E-07 | GEMMA | SUG |
| SL_17AY | AX-110818469 | 7D | 221993803 | -0.3726 | | 2.06E-07 | GEMMA | SUG |
| SL_17AY | AX-111003388 | 7D | 223844284 | -0.3489 | | 1.00E-06 | GEMMA | SUG |
| SL_17AY | AX-109342179 | 7D | 224450406 | -0.3412 | | 1.71E-06 | GEMMA | SUG |
| SL_17AY | AX-110564468 | 7D | 227401276 | -0.3537 | | 6.74E-07 | GEMMA | SUG |
| SL_17AY | AX-109041335 | 7D | 230591444 | -0.3571 | | 5.56E-07 | GEMMA | SUG |
| SL_17AY | AX-109099091 | 7D | 266688807 | -0.7365 | | 4.48E-07 | GEMMA | SUG |
| SL_17AY | AX-110278422 | 7D | 344668648 | -0.3515 | | 2.23E-06 | GEMMA | SUG |
| SL_17AY | AX-110986792 | 7D | 357040286 | -0.3636 | | 1.05E-06 | GEMMA | SUG |
| SL_17AY | AX-109032020 | 7D | 359245146 | -0.3499 | | 2.17E-06 | GEMMA | SUG |

| | | | | | | | | |
|---|---|---|---|---|---|---|---|---|
| SL_17AY | AX-111261154 | 7D | 363818801 | -0.3544 | | 1.90E-06 | GEMMA | SUG |
| SL_17AY | AX-109583235 | 7D | 370800158 | -0.3517 | | 2.24E-06 | GEMMA | SUG |
| SL_17AY | AX-111911849 | 7D | 384087880 | -0.3571 | | 2.00E-06 | GEMMA | SUG |
| SL_17AY | AX-89385155 | 7D | 390903608 | -0.2871 | | 1.96E-06 | GEMMA | SUG |
| SL_17AY | AX-109318566 | 7D | 394236920 | -0.2645 | | 2.87E-07 | BLUPmrMLM | SUG |
| SL_17AY | AX-109318566 | 7D | 394236920 | 0.3402 | | 5.36E-07 | EMMAX | SUG |
| SL_17AY | AX-109318566 | 7D | 394236920 | -0.3401 | | 5.45E-08 | GEMMA | SIG |
| SL_17AY | AX-110946735 | 7D | 395846311 | -0.3785 | | 4.12E-07 | GEMMA | SUG |
| SL_17AY | AX-108867989 | 7D | 617295694 | 0.0670 | -0.6388 | 2.14E-08 | IIIVmrMLM | SIG |
| SL_17JY | AX-89478051 | 1A | 11566853 | -0.2988 | | 1.58E-04 | BLUPmrMLM | SUG |
| SL_17JY | AX-111503088 | 1A | 333509365 | -0.3521 | | 6.93E-07 | BLUPmrMLM | SUG |
| SL_17JY | AX-108950691 | 1A | 394123467 | -0.0289 | 1.2744 | 4.27E-05 | IIIVmrMLM | SUG |
| SL_17JY | AX-108728065 | 1A | 497085247 | -0.1401 | -0.7984 | 5.59E-12 | IIIVmrMLM | SIG |
| SL_17JY | AX-94413745 | 1A | 512666565 | -0.3672 | | 6.39E-16 | IIIVmrMLM | SIG |
| SL_17JY | AX-108945848 | 1B | 166056418 | 2.1388 | | 1.84E-06 | FarmCPU | SUG |
| SL_17JY | AX-109009115 | 1B | 167107973 | 2.1388 | | 1.84E-06 | FarmCPU | SUG |
| SL_17JY | AX-110435866 | 1B | 668724693 | -0.0929 | 0.8923 | 1.13E-08 | IIIVmrMLM | SIG |
| SL_17JY | AX-109367768 | 2A | 179304500 | -0.0568 | -0.8701 | 2.10E-07 | IIIVmrMLM | SUG |
| SL_17JY | AX-111004089 | 2A | 550689364 | -0.1393 | -1.0739 | 4.27E-08 | IIIVmrMLM | SIG |
| SL_17JY | AX-110200150 | 2A | 654174838 | 0.3469 | | 1.11E-07 | BLUPmrMLM | SIG |
| SL_17JY | AX-110200150 | 2A | 654174838 | -0.3126 | -0.1399 | 1.23E-10 | IIIVmrMLM | SIG |
| SL_17JY | AX-108817857 | 2B | 102105764 | -0.1822 | | 1.08E-04 | BLUPmrMLM | SUG |
| SL_17JY | AX-108834775 | 2B | 450549625 | -0.1281 | -1.9145 | 1.31E-06 | IIIVmrMLM | SUG |
| SL_17JY | AX-94902423 | 2B | 793582292 | -0.3427 | | 1.07E-07 | BLUPmrMLM | SIG |
| SL_17JY | AX-94902423 | 2B | 793582292 | 0.3034 | 0.0879 | 4.31E-09 | IIIVmrMLM | SIG |
| SL_17JY | AX-111793952 | 2D | 421261826 | 0.2841 | | 1.00E-04 | BLUPmrMLM | SUG |
| SL_17JY | AX-111211642 | 3A | 67907762 | 0.1640 | | 1.90E-04 | BLUPmrMLM | SUG |
| SL_17JY | AX-86171064 | 3A | 647478803 | 0.2791 | -0.3227 | 3.76E-09 | IIIVmrMLM | SIG |
| SL_17JY | AX-109401618 | 3A | 730336396 | -0.2738 | 2.2916 | 1.80E-12 | IIIVmrMLM | SIG |
| SL_17JY | AX-111121866 | 3B | 758420925 | 0.2614 | | 1.21E-05 | BLUPmrMLM | SUG |
| SL_17JY | AX-110025946 | 4A | 47151264 | -0.4120 | | 6.13E-07 | BLUPmrMLM | SUG |
| SL_17JY | AX-110640156 | 4A | 621478918 | -0.2251 | | 2.12E-05 | BLUPmrMLM | SUG |
| SL_17JY | AX-109433861 | 4B | 656032772 | 0.1888 | | 1.51E-05 | BLUPmrMLM | SUG |
| SL_17JY | AX-111065075 | 5A | 537589010 | 0.1999 | -3.1648 | 1.63E-10 | IIIVmrMLM | SIG |
| SL_17JY | AX-108870666 | 5B | 34771161 | 0.2099 | | 1.07E-04 | BLUPmrMLM | SUG |
| SL_17JY | AX-109975446 | 5B | 82233019 | -0.2605 | | 2.84E-05 | BLUPmrMLM | SUG |
| SL_17JY | AX-95125792 | 6A | 13072033 | 0.0332 | 0.5425 | 7.02E-05 | IIIVmrMLM | SUG |
| SL_17JY | AX-110726653 | 6A | 549032014 | 0.0490 | 0.6741 | 5.45E-05 | IIIVmrMLM | SUG |
| SL_17JY | AX-94799263 | 6A | 596905361 | -0.4385 | -0.0195 | 1.94E-11 | IIIVmrMLM | SIG |
| SL_17JY | AX-108780338 | 6B | 28000315 | 0.2747 | | 6.08E-06 | BLUPmrMLM | SUG |
| SL_17JY | AX-108728767 | 6B | 54662178 | 0.2824 | 0.1403 | 3.49E-09 | IIIVmrMLM | SIG |
| SL_17JY | AX-109067887 | 6B | 481677515 | -0.0951 | 0.8643 | 2.52E-05 | IIIVmrMLM | SUG |
| SL_17JY | AX-109438457 | 6B | 708222748 | -0.1547 | | 6.88E-05 | BLUPmrMLM | SUG |
| SL_17JY | AX-110986707 | 6D | 463797427 | 0.0130 | -0.8446 | 5.79E-06 | IIIVmrMLM | SUG |

| | | | | | | | | |
|---|---|---|---|---|---|---|---|---|
| SL_17JY | AX-94527341 | 6D | 468836635 | 0.5389 | | 2.88E-06 | BLUPmrMLM | SUG |
| SL_17JY | AX-111604502 | 7A | 19191724 | 0.2129 | | 1.86E-04 | BLUPmrMLM | SUG |
| SL_17JY | AX-109992872 | 7A | 110070206 | -0.0519 | 1.7415 | 2.45E-08 | IIIVmrMLM | SIG |
| SL_17JY | AX-110067376 | 7A | 138692082 | 0.4629 | | 1.40E-06 | GEMMA | SUG |
| SL_17JY | AX-111100853 | 7A | 138957200 | -0.5105 | | 6.61E-07 | GEMMA | SUG |
| SL_17JY | AX-109009201 | 7A | 145462820 | 0.5861 | | 1.54E-05 | BLUPmrMLM | SUG |
| SL_17JY | AX-109009201 | 7A | 145462820 | 0.8031 | | 1.59E-06 | GEMMA | SUG |
| SL_17JY | AX-108786152 | 7A | 606324724 | -0.2768 | | 7.15E-06 | BLUPmrMLM | SUG |
| SL_17JY | AX-110914020 | 7A | 701406187 | 0.3030 | | 2.61E-05 | BLUPmrMLM | SUG |
| SL_17JY | AX-110914020 | 7A | 701406187 | -0.2987 | 0.0518 | 6.30E-08 | IIIVmrMLM | SIG |
| SL_17JY | AX-109380821 | 7B | 705322984 | 0.0702 | -0.7388 | 2.01E-17 | IIIVmrMLM | SIG |
| SL_18AY | AX-108791871 | 1A | 53673511 | 0.0762 | -0.2862 | 3.75E-10 | IIIVmrMLM | SIG |
| SL_18AY | AX-110095467 | 1A | 498137970 | -0.1407 | | 1.84E-04 | BLUPmrMLM | SUG |
| SL_18AY | AX-109329888 | 1A | 572210434 | -0.0847 | -0.7127 | 8.18E-11 | IIIVmrMLM | SIG |
| SL_18AY | AX-109078095 | 1A | 578202540 | -0.5964 | | 3.78E-10 | BLUPmrMLM | SIG |
| SL_18AY | AX-108829737 | 1A | 589628965 | -0.1809 | | 2.84E-05 | BLUPmrMLM | SUG |
| SL_18AY | AX-111668935 | 1B | 4531482 | 0.1247 | -0.0237 | 8.35E-08 | IIIVmrMLM | SIG |
| SL_18AY | AX-109941451 | 1B | 302503642 | -0.1070 | -0.2221 | 2.02E-14 | IIIVmrMLM | SIG |
| SL_18AY | AX-110484579 | 1B | 409636535 | 0.1171 | 0.2182 | 1.47E-12 | IIIVmrMLM | SIG |
| SL_18AY | AX-110938847 | 2A | 1497437 | 0.0286 | 0.4562 | 4.25E-08 | IIIVmrMLM | SIG |
| SL_18AY | AX-110482921 | 2A | 7689866 | 0.1184 | | 3.22E-05 | BLUPmrMLM | SUG |
| SL_18AY | AX-110482921 | 2A | 7689866 | -0.1212 | -0.9038 | 5.75E-13 | IIIVmrMLM | SIG |
| SL_18AY | AX-111156761 | 2A | 76894114 | -0.1082 | -0.3373 | 2.58E-13 | IIIVmrMLM | SIG |
| SL_18AY | AX-108758737 | 2A | 192062894 | -0.0374 | -1.1719 | 1.18E-09 | IIIVmrMLM | SIG |
| SL_18AY | AX-111613861 | 2A | 605000807 | 0.2796 | | 1.94E-09 | BLUPmrMLM | SIG |
| SL_18AY | AX-109283226 | 2A | 611880470 | -0.2369 | 0.0047 | 1.77E-27 | IIIVmrMLM | SIG |
| SL_18AY | AX-110579581 | 2A | 703875578 | -0.0018 | 1.5698 | 3.47E-13 | IIIVmrMLM | SIG |
| SL_18AY | AX-94622565 | 2A | 779673707 | 0.1391 | 0.8713 | 1.87E-15 | IIIVmrMLM | SIG |
| SL_18AY | AX-108860387 | 2B | 595707451 | 0.1993 | | 4.93E-06 | BLUPmrMLM | SUG |
| SL_18AY | AX-110911725 | 2B | 755341817 | -0.2951 | | 1.99E-06 | GEMMA | SUG |
| SL_18AY | AX-109860979 | 2B | 791411098 | -0.0210 | -0.5671 | 3.96E-08 | IIIVmrMLM | SIG |
| SL_18AY | AX-94406090 | 2D | 9393902 | 0.3262 | | 8.98E-06 | BLUPmrMLM | SUG |
| SL_18AY | AX-108991361 | 2D | 23304090 | -0.2285 | 0.0807 | 1.74E-25 | IIIVmrMLM | SIG |
| SL_18AY | AX-94507834 | 2D | 43369642 | -0.7587 | 0.2428 | 5.33E-158 | IIIVmrMLM | SIG |
| SL_18AY | AX-111929126 | 2D | 597686819 | -0.1086 | -0.6264 | 1.31E-22 | IIIVmrMLM | SIG |
| SL_18AY | AX-94883443 | 2D | 634296171 | -0.0801 | | 7.72E-05 | IIIVmrMLM | SUG |
| SL_18AY | AX-110984001 | 3A | 1520159 | -0.0604 | 0.7859 | 2.17E-07 | IIIVmrMLM | SUG |
| SL_18AY | AX-109579575 | 3A | 196180365 | 0.1184 | 0.5182 | 2.04E-12 | IIIVmrMLM | SIG |
| SL_18AY | AX-111495554 | 3A | 510693632 | -0.2148 | | 5.56E-06 | BLUPmrMLM | SUG |
| SL_18AY | AX-109477623 | 3A | 624851337 | 0.3032 | 0.0018 | 1.43E-35 | IIIVmrMLM | SIG |
| SL_18AY | AX-112288904 | 3A | 684297263 | 0.2082 | -0.0999 | 1.70E-21 | IIIVmrMLM | SIG |
| SL_18AY | AX-110390328 | 3B | 2882721 | 0.2384 | | 1.17E-07 | BLUPmrMLM | SUG |
| SL_18AY | AX-110587489 | 3B | 14253549 | 0.1186 | | 7.82E-05 | BLUPmrMLM | SUG |

| | | | | | | | | |
|---|---|---|---|---|---|---|---|---|
| SL_18AY | AX-94615079 | 3B | 232567634 | -0.1863 | | 1.33E-04 | BLUPmrMLM | SUG |
| SL_18AY | AX-108941515 | 3B | 469534228 | 0.1822 | | 1.52E-06 | BLUPmrMLM | SUG |
| SL_18AY | AX-110515200 | 3B | 511050987 | 0.0559 | 0.2905 | 6.99E-17 | IIIVmrMLM | SIG |
| SL_18AY | AX-111040970 | 3B | 562918451 | 0.2102 | | 1.39E-23 | IIIVmrMLM | SIG |
| SL_18AY | AX-111758074 | 3B | 768451386 | 0.1308 | | 1.64E-04 | BLUPmrMLM | SUG |
| SL_18AY | AX-109497989 | 3B | 779139167 | 1.3165 | | 8.14E-07 | GEMMA | SUG |
| SL_18AY | AX-110996288 | 3D | 294047172 | -0.1670 | | 6.12E-07 | BLUPmrMLM | SUG |
| SL_18AY | AX-110996288 | 3D | 294047172 | 0.1501 | 0.2257 | 4.57E-13 | IIIVmrMLM | SIG |
| SL_18AY | AX-110105874 | 4A | 19655948 | -0.3985 | | 9.31E-07 | EMMAX | SUG |
| SL_18AY | AX-110105874 | 4A | 19655948 | 0.3986 | | 6.41E-07 | GEMMA | SUG |
| SL_18AY | AX-109329316 | 4A | 21222835 | 0.2308 | | 1.93E-07 | BLUPmrMLM | SUG |
| SL_18AY | AX-111583398 | 4A | 109217253 | 0.2027 | | 9.77E-06 | BLUPmrMLM | SUG |
| SL_18AY | AX-108795868 | 4A | 673679914 | 0.1922 | -0.0424 | 1.74E-15 | IIIVmrMLM | SIG |
| SL_18AY | AX-108877201 | 4B | 655489692 | 0.0004 | -0.4838 | 8.96E-08 | IIIVmrMLM | SIG |
| SL_18AY | AX-110660970 | 4D | 179851231 | -0.2125 | | 5.00E-24 | IIIVmrMLM | SIG |
| SL_18AY | AX-94520301 | 5A | 445264623 | -0.1013 | | 1.08E-04 | BLUPmrMLM | SUG |
| SL_18AY | AX-109277958 | 5A | 566989983 | -0.1464 | -0.0314 | 2.33E-10 | IIIVmrMLM | SIG |
| SL_18AY | AX-109375326 | 5A | 609951432 | 0.2884 | | 1.07E-06 | FarmCPU | SUG |
| SL_18AY | AX-109818044 | 5A | 609951936 | 0.2850 | | 1.45E-06 | FarmCPU | SUG |
| SL_18AY | AX-108939870 | 5A | 610082004 | 0.2813 | | 2.02E-06 | FarmCPU | SUG |
| SL_18AY | AX-110594079 | 5A | 610180225 | 0.2813 | | 2.02E-06 | FarmCPU | SUG |
| SL_18AY | AX-111254684 | 5A | 610211291 | 0.2942 | | 1.16E-06 | FarmCPU | SUG |
| SL_18AY | AX-111465776 | 5A | 610219559 | 0.2842 | | 1.77E-06 | FarmCPU | SUG |
| SL_18AY | AX-111240636 | 5A | 653884368 | 0.2371 | | 5.62E-29 | IIIVmrMLM | SIG |
| SL_18AY | AX-94780072 | 5B | 13428632 | 0.1960 | | 6.35E-21 | IIIVmrMLM | SIG |
| SL_18AY | AX-110441595 | 5B | 417880578 | -0.1774 | | 1.27E-04 | BLUPmrMLM | SUG |
| SL_18AY | AX-109597529 | 5B | 531039580 | -0.0111 | 1.5368 | 7.90E-11 | IIIVmrMLM | SIG |
| SL_18AY | AX-109912680 | 5B | 534756220 | 0.1146 | | 4.09E-05 | BLUPmrMLM | SUG |
| SL_18AY | AX-110373709 | 5B | 603897101 | 0.1585 | | 1.33E-06 | BLUPmrMLM | SUG |
| SL_18AY | AX-95217894 | 5B | 631888512 | -0.2055 | | 1.10E-22 | IIIVmrMLM | SIG |
| SL_18AY | AX-110072244 | 5D | 57230257 | -0.0724 | 1.3002 | 3.27E-09 | IIIVmrMLM | SIG |
| SL_18AY | AX-110225350 | 5D | 459062642 | 0.1538 | -0.1471 | 8.20E-13 | IIIVmrMLM | SIG |
| SL_18AY | AX-95188660 | 5D | 528764624 | -0.4240 | | 8.79E-06 | BLUPmrMLM | SUG |
| SL_18AY | AX-110919411 | 6A | 1690595 | -0.0027 | -0.4229 | 9.37E-07 | IIIVmrMLM | SUG |
| SL_18AY | AX-111513752 | 6A | 445007549 | -0.1435 | 0.3815 | 8.37E-20 | IIIVmrMLM | SIG |
| SL_18AY | AX-95000292 | 6A | 617658645 | -0.2310 | | 3.24E-05 | BLUPmrMLM | SUG |
| SL_18AY | AX-94677670 | 6B | 22819515 | 0.0073 | 0.5468 | 3.37E-07 | IIIVmrMLM | SUG |
| SL_18AY | AX-109936451 | 6B | 143200777 | -0.0079 | -0.3253 | 4.36E-11 | IIIVmrMLM | SIG |
| SL_18AY | AX-94771536 | 6B | 143661387 | -0.4095 | | 7.79E-17 | BLUPmrMLM | SIG |
| SL_18AY | AX-109998444 | 6B | 303181854 | 0.0051 | 0.4089 | 2.23E-11 | IIIVmrMLM | SIG |
| SL_18AY | AX-108725833 | 6D | 5134596 | 0.1469 | | 4.16E-06 | BLUPmrMLM | SUG |
| SL_18AY | AX-111037646 | 6D | 17127837 | 0.2782 | 0.6081 | 3.00E-55 | IIIVmrMLM | SIG |
| SL_18AY | AX-95230036 | 6D | 17299082 | -0.1054 | | 5.26E-05 | BLUPmrMLM | SUG |
| SL_18AY | AX-111463721 | 6D | 433401548 | -0.1456 | -0.1287 | 1.42E-11 | IIIVmrMLM | SIG |

| | | | | | | | | |
|---|---|---|---|---|---|---|---|---|
| SL_18AY | AX-109879989 | 7A | 13838274 | -0.1306 | | 6.41E-06 | BLUPmrMLM | SUG |
| SL_18AY | AX-109879989 | 7A | 13838274 | 0.1279 | 0.2844 | 3.61E-10 | IIIVmrMLM | SIG |
| SL_18AY | AX-110677194 | 7A | 68327395 | -0.2670 | 0.0444 | 4.01E-31 | IIIVmrMLM | SIG |
| SL_18AY | AX-109334699 | 7A | 69362215 | 0.2473 | 0.6052 | 4.56E-51 | IIIVmrMLM | SIG |
| SL_18AY | AX-109413541 | 7A | 134416041 | -1.6863 | | 1.32E-06 | EMMAX | SUG |
| SL_18AY | AX-111645267 | 7A | 196669501 | -0.1568 | | 2.74E-05 | BLUPmrMLM | SUG |
| SL_18AY | AX-110561326 | 7A | 513412334 | 0.0190 | 0.7075 | 7.83E-05 | IIIVmrMLM | SUG |
| SL_18AY | AX-109324892 | 7A | 625193469 | -0.1896 | -0.032 | 4.78E-17 | IIIVmrMLM | SIG |
| SL_18AY | AX-110511070 | 7A | 639546012 | 0.1723 | 0.1775 | 8.33E-16 | IIIVmrMLM | SIG |
| SL_18AY | AX-109279314 | 7B | 62784350 | 0.0471 | -0.7135 | 2.84E-06 | IIIVmrMLM | SUG |
| SL_18AY | AX-110140515 | 7B | 337533448 | -0.2381 | | 1.47E-07 | BLUPmrMLM | SUG |
| SL_18AY | AX-110140515 | 7B | 337533448 | 0.0678 | -0.1719 | 1.67E-04 | IIIVmrMLM | SUG |
| SL_18AY | AX-111044293 | 7B | 714766596 | 0.1511 | -0.281 | 1.41E-17 | IIIVmrMLM | SIG |
| SL_18AY | AX-111685391 | 7B | 725264838 | -0.0749 | 1.1757 | 2.28E-08 | IIIVmrMLM | SIG |
| SL_18AY | AX-111713035 | 7B | 743147463 | -0.3019 | -0.113 | 6.99E-38 | IIIVmrMLM | SIG |
| SL_18AY | AX-94614496 | 7D | 262286238 | -0.1195 | | 4.86E-09 | IIIVmrMLM | SIG |
| SL_18AY | AX-110946735 | 7D | 395846311 | -0.2788 | | 3.62E-07 | BLUPmrMLM | SUG |
| SL_18AY | AX-111160590 | 7D | 611751703 | -0.0424 | -0.4082 | 5.35E-09 | IIIVmrMLM | SIG |
| SL_18JY | AX-94530450 | 1A | 1174820 | 0.2200 | | 1.28E-05 | BLUPmrMLM | SUG |
| SL_18JY | AX-111081504 | 1A | 27327553 | 0.0078 | 0.4198 | 2.72E-07 | IIIVmrMLM | SUG |
| SL_18JY | AX-111015470 | 1A | 29787383 | -0.2245 | -0.018 | 1.56E-13 | IIIVmrMLM | SIG |
| SL_18JY | AX-108978402 | 1A | 485014584 | 0.0425 | 1.1261 | 1.16E-13 | IIIVmrMLM | SIG |
| SL_18JY | AX-110023784 | 1A | 520606961 | 0.0730 | -0.2172 | 4.64E-06 | IIIVmrMLM | SUG |
| SL_18JY | AX-110394578 | 1A | 522239093 | -0.1464 | | 7.60E-05 | BLUPmrMLM | SUG |
| SL_18JY | AX-109390078 | 1A | 543177661 | -0.0010 | 0.3466 | 1.12E-11 | IIIVmrMLM | SIG |
| SL_18JY | AX-109078095 | 1A | 578202540 | -0.5449 | | 1.15E-06 | GEMMA | SUG |
| SL_18JY | AX-108954721 | 1A | 587368067 | -0.1427 | | 1.20E-05 | BLUPmrMLM | SUG |
| SL_18JY | AX-111477855 | 1B | 348480772 | 0.2482 | -0.8981 | 1.09E-31 | IIIVmrMLM | SIG |
| SL_18JY | AX-110933759 | 1B | 377675920 | -0.0402 | -0.2158 | 1.52E-05 | IIIVmrMLM | SUG |
| SL_18JY | AX-109870951 | 1B | 577655889 | -0.2739 | | 2.23E-07 | BLUPmrMLM | SUG |
| SL_18JY | AX-111096015 | 1B | 613727594 | -0.1982 | | 3.15E-06 | BLUPmrMLM | SUG |
| SL_18JY | AX-109372293 | 1B | 635920800 | 0.1340 | | 1.73E-05 | BLUPmrMLM | SUG |
| SL_18JY | AX-108750918 | 1D | 4596218 | 0.0849 | -0.7355 | 3.33E-09 | IIIVmrMLM | SIG |
| SL_18JY | AX-94893335 | 1D | 58757024 | 0.2419 | | 2.08E-05 | BLUPmrMLM | SUG |
| SL_18JY | AX-109746270 | 1D | 433528632 | -0.1013 | | 4.90E-05 | BLUPmrMLM | SUG |
| SL_18JY | AX-95255103 | 2A | 67521674 | 0.0341 | 0.8322 | 4.89E-08 | IIIVmrMLM | SIG |
| SL_18JY | AX-108834206 | 2A | 502270483 | 0.0028 | -0.712 | 2.62E-08 | IIIVmrMLM | SIG |
| SL_18JY | AX-110497091 | 2A | 753704714 | 0.1207 | | 2.09E-05 | BLUPmrMLM | SUG |
| SL_18JY | AX-111252001 | 2A | 778870108 | -0.1069 | | 8.89E-06 | BLUPmrMLM | SUG |
| SL_18JY | AX-94601893 | 2B | 34084963 | -0.0335 | 0.2533 | 3.37E-05 | IIIVmrMLM | SUG |
| SL_18JY | AX-110017170 | 2B | 464442409 | -0.0158 | 0.2474 | 2.34E-15 | IIIVmrMLM | SIG |
| SL_18JY | AX-111055170 | 2B | 753805731 | -0.1443 | | 4.45E-07 | BLUPmrMLM | SUG |
| SL_18JY | AX-110617369 | 2B | 773092166 | 0.1251 | | 6.36E-05 | BLUPmrMLM | SUG |
| SL_18JY | AX-110617369 | 2B | 773092166 | -0.2137 | 0.0523 | 4.53E-21 | IIIVmrMLM | SIG |

| | | | | | | | | |
|---|---|---|---|---|---|---|---|---|
| SL_18JY | AX-110038796 | 2B | 799533170 | 0.0245 | -0.7223 | 2.62E-04 | IIIVmrMLM | SUG |
| SL_18JY | AX-109741923 | 2D | 17947900 | 0.1536 | -0.189 | 4.56E-13 | IIIVmrMLM | SIG |
| SL_18JY | AX-111002945 | 2D | 19505527 | -0.2424 | | 1.14E-07 | BLUPmrMLM | SIG |
| SL_18JY | AX-111002945 | 2D | 19505527 | 0.1812 | 0.1289 | 7.36E-17 | IIIVmrMLM | SIG |
| SL_18JY | AX-110487796 | 2D | 21809735 | -0.2627 | | 1.21E-06 | EMMAX | SUG |
| SL_18JY | AX-110487796 | 2D | 21809735 | 0.2640 | | 4.36E-07 | GEMMA | SUG |
| SL_18JY | AX-110780489 | 2D | 21810300 | 0.2582 | | 9.08E-07 | GEMMA | SUG |
| SL_18JY | AX-111585353 | 2D | 21909785 | -0.2927 | | 6.00E-07 | EMMAX | SUG |
| SL_18JY | AX-111585353 | 2D | 21909785 | 0.2941 | | 2.04E-07 | GEMMA | SUG |
| SL_18JY | AX-109727939 | 2D | 21911024 | -0.2694 | | 9.33E-07 | EMMAX | SUG |
| SL_18JY | AX-109727939 | 2D | 21911024 | 0.2700 | | 3.50E-07 | GEMMA | SUG |
| SL_18JY | AX-111491285 | 2D | 21911067 | 0.2572 | | 9.99E-07 | GEMMA | SUG |
| SL_18JY | AX-111220034 | 2D | 21911258 | -0.2856 | | 7.42E-07 | EMMAX | SUG |
| SL_18JY | AX-111220034 | 2D | 21911258 | 0.2868 | | 2.91E-07 | GEMMA | SUG |
| SL_18JY | AX-111574942 | 2D | 21911320 | -0.2585 | | 1.96E-06 | EMMAX | SUG |
| SL_18JY | AX-111574942 | 2D | 21911320 | 0.2596 | | 7.52E-07 | GEMMA | SUG |
| SL_18JY | AX-109844617 | 2D | 21911530 | 0.1890 | | 6.94E-07 | BLUPmrMLM | SUG |
| SL_18JY | AX-109844617 | 2D | 21911530 | -0.2692 | | 3.40E-07 | EMMAX | SUG |
| SL_18JY | AX-109844617 | 2D | 21911530 | 0.2694 | | 1.38E-07 | GEMMA | SUG |
| SL_18JY | AX-109785758 | 2D | 21915115 | -0.2656 | | 8.73E-07 | EMMAX | SUG |
| SL_18JY | AX-109785758 | 2D | 21915115 | 0.2664 | | 3.22E-07 | GEMMA | SUG |
| SL_18JY | AX-110516159 | 2D | 21915211 | -0.2637 | | 6.68E-07 | EMMAX | SUG |
| SL_18JY | AX-110516159 | 2D | 21915211 | 0.2647 | | 2.45E-07 | GEMMA | SUG |
| SL_18JY | AX-110499484 | 2D | 21915260 | -0.2666 | | 6.34E-07 | EMMAX | SUG |
| SL_18JY | AX-110499484 | 2D | 21915260 | 0.2674 | | 2.31E-07 | GEMMA | SUG |
| SL_18JY | AX-110370402 | 2D | 21915541 | -0.2611 | | 1.34E-06 | EMMAX | SUG |
| SL_18JY | AX-110370402 | 2D | 21915541 | 0.2623 | | 4.85E-07 | GEMMA | SUG |
| SL_18JY | AX-109286507 | 2D | 21915778 | -0.2713 | | 4.05E-07 | EMMAX | SUG |
| SL_18JY | AX-109286507 | 2D | 21915778 | 0.2719 | | 1.50E-07 | GEMMA | SUG |
| SL_18JY | AX-110059604 | 2D | 21919402 | -0.2604 | | 1.44E-06 | EMMAX | SUG |
| SL_18JY | AX-110059604 | 2D | 21919402 | 0.2613 | | 5.51E-07 | GEMMA | SUG |
| SL_18JY | AX-111264534 | 2D | 21924723 | -0.2694 | | 8.38E-07 | EMMAX | SUG |
| SL_18JY | AX-111264534 | 2D | 21924723 | 0.2707 | | 2.94E-07 | GEMMA | SUG |
| SL_18JY | AX-110921284 | 2D | 21925408 | -0.2619 | | 1.72E-06 | EMMAX | SUG |
| SL_18JY | AX-110921284 | 2D | 21925408 | 0.2632 | | 6.17E-07 | GEMMA | SUG |
| SL_18JY | AX-110413771 | 2D | 21927487 | 0.2499 | | 1.56E-06 | GEMMA | SUG |
| SL_18JY | AX-110891163 | 2D | 21934578 | 0.2570 | | 1.18E-06 | GEMMA | SUG |
| SL_18JY | AX-109153441 | 2D | 22099623 | 0.2536 | | 1.50E-06 | GEMMA | SUG |
| SL_18JY | AX-86180719 | 2D | 22110951 | -0.2598 | | 1.85E-06 | EMMAX | SUG |
| SL_18JY | AX-86180719 | 2D | 22110951 | 0.2610 | | 6.76E-07 | GEMMA | SUG |
| SL_18JY | AX-110494885 | 2D | 22123193 | -0.2581 | | 2.14E-06 | EMMAX | SUG |
| SL_18JY | AX-110494885 | 2D | 22123193 | 0.2593 | | 7.82E-07 | GEMMA | SUG |
| SL_18JY | AX-109548559 | 2D | 22153165 | 0.2512 | | 1.66E-06 | GEMMA | SUG |
| SL_18JY | AX-111565649 | 2D | 22154507 | 0.2554 | | 1.32E-06 | GEMMA | SUG |

| | | | | | | | | |
|---|---|---|---|---|---|---|---|---|
| SL_18JY | AX-108727836 | 2D | 22163334 | 0.2480 | | 1.77E-06 | GEMMA | SUG |
| SL_18JY | AX-89576820 | 2D | 22176723 | 0.2497 | | 1.72E-06 | GEMMA | SUG |
| SL_18JY | AX-109842248 | 2D | 23102530 | -0.2843 | | 2.53E-08 | EMMAX | SIG |
| SL_18JY | AX-109842248 | 2D | 23102530 | 0.2891 | | 4.53E-09 | GEMMA | SIG |
| SL_18JY | AX-108991361 | 2D | 23304090 | -0.2894 | | 1.54E-08 | EMMAX | SIG |
| SL_18JY | AX-108991361 | 2D | 23304090 | 0.2938 | | 2.78E-09 | GEMMA | SIG |
| SL_18JY | AX-108991361 | 2D | 23304090 | -0.1156 | -0.1479 | 5.96E-08 | IIIVmrMLM | SIG |
| SL_18JY | AX-111561744 | 2D | 23416219 | -0.2721 | | 9.38E-08 | EMMAX | SIG |
| SL_18JY | AX-111561744 | 2D | 23416219 | 0.2771 | | 1.81E-08 | GEMMA | SIG |
| SL_18JY | AX-111038058 | 2D | 23417635 | -0.2712 | | 1.04E-07 | EMMAX | SIG |
| SL_18JY | AX-111038058 | 2D | 23417635 | 0.2763 | | 2.00E-08 | GEMMA | SIG |
| SL_18JY | AX-111439345 | 2D | 458519561 | 0.1991 | | 7.45E-06 | BLUPmrMLM | SUG |
| SL_18JY | AX-110399950 | 3A | 682940543 | 0.1824 | | 1.76E-06 | BLUPmrMLM | SUG |
| SL_18JY | AX-108840020 | 3B | 12405885 | -0.0127 | 0.3738 | 6.46E-05 | IIIVmrMLM | SUG |
| SL_18JY | AX-110496417 | 3B | 20840296 | -0.1502 | | 1.93E-06 | BLUPmrMLM | SUG |
| SL_18JY | AX-110602961 | 3B | 204614320 | 0.1515 | | 4.75E-06 | BLUPmrMLM | SUG |
| SL_18JY | AX-94633748 | 3B | 628722797 | 0.1950 | | 7.78E-06 | BLUPmrMLM | SUG |
| SL_18JY | AX-94633748 | 3B | 628722797 | -0.1075 | | 1.01E-07 | IIIVmrMLM | SIG |
| SL_18JY | AX-110386923 | 3B | 771552082 | -0.3749 | | 2.42E-62 | IIIVmrMLM | SIG |
| SL_18JY | AX-94541915 | 3D | 496785393 | -1.2410 | | 1.07E-06 | EMMAX | SUG |
| SL_18JY | AX-94541915 | 3D | 496785393 | 1.2405 | | 5.79E-07 | GEMMA | SUG |
| SL_18JY | AX-111146251 | 3D | 588999575 | -0.0326 | -0.6524 | 1.48E-06 | IIIVmrMLM | SUG |
| SL_18JY | AX-109407502 | 4A | 69508934 | -0.1851 | | 6.76E-07 | BLUPmrMLM | SUG |
| SL_18JY | AX-111077888 | 4A | 99022748 | -0.0185 | -0.471 | 2.84E-21 | IIIVmrMLM | SIG |
| SL_18JY | AX-94536350 | 4A | 676421370 | -0.0037 | -0.6103 | 2.96E-19 | IIIVmrMLM | SIG |
| SL_18JY | AX-94454177 | 4B | 636916350 | 0.1526 | 0.3826 | 1.88E-14 | IIIVmrMLM | SIG |
| SL_18JY | AX-89739690 | 4D | 32216233 | 0.1842 | -0.0109 | 1.75E-17 | IIIVmrMLM | SIG |
| SL_18JY | AX-110040600 | 5A | 115742178 | 0.1367 | | 4.36E-06 | BLUPmrMLM | SUG |
| SL_18JY | AX-110422571 | 5A | 465690125 | -0.1180 | -0.0463 | 8.41E-08 | IIIVmrMLM | SIG |
| SL_18JY | AX-109517804 | 5A | 502827669 | -0.0317 | 1.7676 | 8.69E-07 | IIIVmrMLM | SUG |
| SL_18JY | AX-108873512 | 5B | 68025825 | -0.1440 | | 2.67E-05 | BLUPmrMLM | SUG |
| SL_18JY | AX-86175133 | 5B | 437802180 | -0.1054 | | 1.24E-05 | BLUPmrMLM | SUG |
| SL_18JY | AX-109899485 | 5B | 465052250 | -0.2334 | | 2.27E-09 | BLUPmrMLM | SIG |
| SL_18JY | AX-109463404 | 5B | 557902007 | -0.0789 | -0.3829 | 1.56E-09 | IIIVmrMLM | SIG |
| SL_18JY | AX-111056297 | 5B | 591667741 | -0.0473 | 0.575 | 1.73E-10 | IIIVmrMLM | SIG |
| SL_18JY | AX-109424073 | 5B | 700224630 | -0.0364 | -0.259 | 6.46E-08 | IIIVmrMLM | SIG |
| SL_18JY | AX-109074279 | 5D | 425625579 | 0.1521 | | 9.78E-14 | IIIVmrMLM | SIG |
| SL_18JY | AX-110745765 | 5D | 560189984 | 0.2008 | | 2.06E-06 | BLUPmrMLM | SUG |
| SL_18JY | AX-111085933 | 6A | 16226560 | 0.0272 | -0.7875 | 2.65E-10 | IIIVmrMLM | SIG |
| SL_18JY | AX-110733835 | 6A | 360493631 | 0.6274 | | 7.47E-08 | FarmCPU | SIG |
| SL_18JY | AX-109831410 | 6B | 646615984 | -0.1422 | 0.0777 | 1.84E-10 | IIIVmrMLM | SIG |
| SL_18JY | AX-110634426 | 6B | 694400585 | 0.2694 | 0.2868 | 1.88E-35 | IIIVmrMLM | SIG |
| SL_18JY | AX-111629501 | 6B | 702825965 | 0.6603 | | 2.28E-06 | EMMAX | SUG |
| SL_18JY | AX-111629501 | 6B | 702825965 | -0.6684 | | 8.68E-07 | GEMMA | SUG |

| | | | | | | | | |
|---|---|---|---|---|---|---|---|---|
| SL_18JY | AX-108880725 | 6B | 704735685 | 0.1738 | | 5.42E-06 | BLUPmrMLM | SUG |
| SL_18JY | AX-111568358 | 6B | 712253937 | -0.0258 | -0.4485 | 2.82E-07 | IIIVmrMLM | SUG |
| SL_18JY | AX-111151831 | 6D | 463747526 | 0.0478 | -0.2223 | 1.10E-14 | IIIVmrMLM | SIG |
| SL_18JY | AX-110643450 | 7A | 558399409 | 0.0207 | 0.9229 | 1.05E-07 | IIIVmrMLM | SIG |
| SL_18JY | AX-110919181 | 7B | 41287511 | -0.2161 | | 3.37E-06 | BLUPmrMLM | SUG |
| SL_18JY | AX-109960483 | 7B | 328921297 | -0.1163 | | 2.67E-06 | BLUPmrMLM | SUG |
| SL_18JY | AX-95133915 | 7B | 706815412 | 0.0919 | | 5.42E-05 | BLUPmrMLM | SUG |
| SL_18JY | AX-110986792 | 7D | 357040286 | 0.2160 | -0.3424 | 3.94E-24 | IIIVmrMLM | SIG |
| SL_18JY | AX-109190790 | 7D | 509835162 | -0.1360 | | 1.59E-05 | BLUPmrMLM | SUG |
| SL_18XX | AX-111535253 | 1A | 15525424 | 0.0182 | -0.5475 | 4.83E-09 | IIIVmrMLM | SIG |
| SL_18XX | AX-111006831 | 1A | 26971944 | -0.1014 | | 3.93E-07 | BLUPmrMLM | SUG |
| SL_18XX | AX-111006831 | 1A | 26971944 | 0.1203 | 0.0445 | 1.77E-11 | IIIVmrMLM | SIG |
| SL_18XX | AX-111118356 | 1A | 53040561 | -0.0590 | -0.8332 | 5.13E-13 | IIIVmrMLM | SIG |
| SL_18XX | AX-108790034 | 1A | 270864158 | -0.1083 | | 1.40E-05 | BLUPmrMLM | SUG |
| SL_18XX | AX-108790034 | 1A | 270864158 | 0.1490 | -0.0866 | 1.29E-17 | IIIVmrMLM | SIG |
| SL_18XX | AX-108943520 | 1A | 375447935 | -0.1384 | 0.1843 | 1.71E-16 | IIIVmrMLM | SIG |
| SL_18XX | AX-95150513 | 1A | 500827314 | -0.1857 | | 7.36E-05 | BLUPmrMLM | SUG |
| SL_18XX | AX-110487442 | 1A | 555719834 | 0.1063 | -0.0242 | 3.01E-09 | IIIVmrMLM | SIG |
| SL_18XX | AX-108844267 | 1A | 568785749 | 0.1172 | | 8.72E-05 | BLUPmrMLM | SUG |
| SL_18XX | AX-110606195 | 1A | 574477470 | -0.1061 | -0.4787 | 1.25E-15 | IIIVmrMLM | SIG |
| SL_18XX | AX-109078095 | 1A | 578202540 | 4.6900 | 4.3788 | 0.00E+00 | IIIVmrMLM | SIG |
| SL_18XX | AX-111064668 | 1A | 585753393 | 0.1325 | | 8.04E-06 | BLUPmrMLM | SUG |
| SL_18XX | AX-111064668 | 1A | 585753393 | -0.1417 | -0.0095 | 2.82E-11 | IIIVmrMLM | SIG |
| SL_18XX | AX-111612963 | 1B | 333599799 | -0.0943 | | 1.48E-04 | BLUPmrMLM | SUG |
| SL_18XX | AX-111083420 | 1B | 418826302 | -0.0710 | -0.6382 | 1.32E-26 | IIIVmrMLM | SIG |
| SL_18XX | AX-108903748 | 1B | 445743398 | 0.1345 | | 6.70E-05 | BLUPmrMLM | SUG |
| SL_18XX | AX-108903748 | 1B | 445743398 | -0.2189 | -0.0724 | 5.44E-23 | IIIVmrMLM | SIG |
| SL_18XX | AX-109030186 | 1B | 660454234 | -0.2020 | | 1.57E-31 | IIIVmrMLM | SIG |
| SL_18XX | AX-108881574 | 1D | 9857772 | 0.1557 | | 3.64E-20 | IIIVmrMLM | SIG |
| SL_18XX | AX-94859184 | 1D | 427478466 | -0.0234 | 0.2679 | 7.97E-18 | IIIVmrMLM | SIG |
| SL_18XX | AX-111754415 | 2A | 74143363 | 0.4457 | | 3.33E-07 | FarmCPU | SUG |
| SL_18XX | AX-109858598 | 2A | 198849085 | -0.1245 | 0.7092 | 3.02E-16 | IIIVmrMLM | SIG |
| SL_18XX | AX-110079477 | 2A | 432588841 | -0.1210 | | 1.30E-05 | BLUPmrMLM | SUG |
| SL_18XX | AX-109462547 | 2A | 573337130 | 0.1063 | | 8.67E-06 | BLUPmrMLM | SUG |
| SL_18XX | AX-110697462 | 2A | 581455118 | -0.0018 | 0.3367 | 3.81E-22 | IIIVmrMLM | SIG |
| SL_18XX | AX-108975725 | 2A | 710376132 | -0.1253 | | 5.75E-05 | BLUPmrMLM | SUG |
| SL_18XX | AX-94687724 | 2A | 715336986 | 0.3250 | | 5.05E-07 | BLUPmrMLM | SUG |
| SL_18XX | AX-109985301 | 2A | 750615055 | 0.1222 | -0.0164 | 2.33E-11 | IIIVmrMLM | SIG |
| SL_18XX | AX-109914564 | 2A | 750822455 | 0.1258 | | 9.46E-06 | BLUPmrMLM | SUG |
| SL_18XX | AX-94793700 | 2B | 88840036 | 0.1177 | | 8.31E-07 | BLUPmrMLM | SUG |
| SL_18XX | AX-94793700 | 2B | 88840036 | -0.1234 | -0.0257 | 2.35E-12 | IIIVmrMLM | SIG |
| SL_18XX | AX-109321174 | 2B | 170418359 | 0.1791 | | 5.81E-10 | BLUPmrMLM | SIG |
| SL_18XX | AX-111017539 | 2B | 171424483 | 0.1387 | 0.1831 | 2.82E-16 | IIIVmrMLM | SIG |
| SL_18XX | AX-110423885 | 2B | 766172990 | -0.1072 | | 1.53E-04 | BLUPmrMLM | SUG |

| | | | | | | | | |
|---|---|---|---|---|---|---|---|---|
| SL_18XX | AX-110055167 | 2B | 779224260 | 0.0029 | 0.511 | 1.16E-11 | IIIVmrMLM | SIG |
| SL_18XX | AX-110315690 | 2D | 21532303 | 0.2579 | | 1.13E-12 | BLUPmrMLM | SIG |
| SL_18XX | AX-109836946 | 2D | 32795318 | -0.1563 | 0.2634 | 3.58E-20 | IIIVmrMLM | SIG |
| SL_18XX | AX-110901179 | 2D | 649655239 | -0.0712 | | 1.07E-04 | BLUPmrMLM | SUG |
| SL_18XX | AX-109291377 | 3A | 231687378 | 0.2133 | 0.1049 | 3.27E-31 | IIIVmrMLM | SIG |
| SL_18XX | AX-110474993 | 3A | 317874072 | 0.0479 | 0.4559 | 1.42E-07 | IIIVmrMLM | SUG |
| SL_18XX | AX-108736778 | 3A | 510851811 | -0.0263 | 0.2465 | 3.42E-09 | IIIVmrMLM | SIG |
| SL_18XX | AX-109844590 | 3A | 655992019 | -0.1719 | | 4.83E-10 | BLUPmrMLM | SIG |
| SL_18XX | AX-109844590 | 3A | 655992019 | 0.1231 | -0.0353 | 1.09E-11 | IIIVmrMLM | SIG |
| SL_18XX | AX-109856164 | 3A | 691218520 | -0.1169 | 0.2087 | 7.54E-13 | IIIVmrMLM | SIG |
| SL_18XX | AX-110947728 | 3B | 18502138 | -0.0475 | 0.8174 | 5.82E-11 | IIIVmrMLM | SIG |
| SL_18XX | AX-109315378 | 3B | 396713084 | -0.1646 | -0.0208 | 1.10E-16 | IIIVmrMLM | SIG |
| SL_18XX | AX-109474531 | 3B | 440856985 | 0.1631 | | 5.00E-07 | BLUPmrMLM | SUG |
| SL_18XX | AX-111567612 | 3B | 494527545 | -0.1161 | | 6.75E-05 | BLUPmrMLM | SUG |
| SL_18XX | AX-111567612 | 3B | 494527545 | 0.1349 | 0.0466 | 2.28E-11 | IIIVmrMLM | SIG |
| SL_18XX | AX-111135161 | 3B | 581490510 | -0.0998 | | 4.21E-06 | BLUPmrMLM | SUG |
| SL_18XX | AX-109617898 | 3B | 711619155 | 0.0294 | 0.429 | 2.58E-07 | IIIVmrMLM | SUG |
| SL_18XX | AX-89532021 | 3B | 828920628 | -0.1166 | | 1.10E-04 | BLUPmrMLM | SUG |
| SL_18XX | AX-110275510 | 3D | 474995279 | -0.1490 | -0.789 | 1.99E-20 | IIIVmrMLM | SIG |
| SL_18XX | AX-110623976 | 4A | 60929707 | 0.1889 | | 1.11E-06 | BLUPmrMLM | SUG |
| SL_18XX | AX-110122516 | 4A | 103329003 | -0.0777 | -0.7878 | 3.48E-25 | IIIVmrMLM | SIG |
| SL_18XX | AX-108965701 | 4A | 147629644 | 0.0575 | 0.2622 | 1.91E-14 | IIIVmrMLM | SIG |
| SL_18XX | AX-109383852 | 4A | 198122747 | 0.3099 | | 4.26E-06 | BLUPmrMLM | SUG |
| SL_18XX | AX-110019748 | 4A | 698898096 | 0.0446 | 0.3061 | 7.64E-17 | IIIVmrMLM | SIG |
| SL_18XX | AX-111048537 | 4A | 726969325 | 0.1136 | | 2.10E-05 | BLUPmrMLM | SUG |
| SL_18XX | AX-111701101 | 4B | 519351016 | -0.1887 | | 2.10E-07 | BLUPmrMLM | SUG |
| SL_18XX | AX-111073447 | 4B | 636965370 | 0.0671 | | 4.57E-05 | IIIVmrMLM | SUG |
| SL_18XX | AX-95653095 | 5A | 2646301 | -0.0735 | | 1.89E-04 | BLUPmrMLM | SUG |
| SL_18XX | AX-94471636 | 5A | 33012106 | 0.1813 | | 3.74E-07 | BLUPmrMLM | SUG |
| SL_18XX | AX-111013276 | 5A | 506159283 | -0.1579 | -0.3499 | 3.88E-22 | IIIVmrMLM | SIG |
| SL_18XX | AX-111622382 | 5A | 508210451 | 0.1387 | | 1.94E-06 | BLUPmrMLM | SUG |
| SL_18XX | AX-109069602 | 5A | 567211658 | -0.1026 | | 6.31E-10 | IIIVmrMLM | SIG |
| SL_18XX | AX-108838394 | 5B | 61801648 | 0.0829 | -0.4678 | 2.73E-15 | IIIVmrMLM | SIG |
| SL_18XX | AX-110442887 | 5B | 652372001 | -0.2629 | | 1.28E-05 | BLUPmrMLM | SUG |
| SL_18XX | AX-111639475 | 5D | 429713292 | -0.3818 | -0.7452 | 2.05E-99 | IIIVmrMLM | SIG |
| SL_18XX | AX-109053938 | 5D | 560217732 | 0.0990 | | 1.34E-04 | BLUPmrMLM | SUG |
| SL_18XX | AX-109873685 | 6A | 907392 | -0.1185 | | 7.73E-05 | BLUPmrMLM | SUG |
| SL_18XX | AX-110389554 | 6A | 16703640 | 0.1150 | | 1.70E-05 | BLUPmrMLM | SUG |
| SL_18XX | AX-94505018 | 6A | 612327597 | 0.1175 | 0.5094 | 2.32E-17 | IIIVmrMLM | SIG |
| SL_18XX | AX-110617639 | 6B | 9058948 | 0.1276 | | 3.12E-05 | BLUPmrMLM | SUG |
| SL_18XX | AX-111523255 | 6B | 28003889 | 0.0140 | 0.3563 | 8.20E-06 | IIIVmrMLM | SUG |
| SL_18XX | AX-109482802 | 6B | 93842716 | 0.0090 | -0.376 | 8.68E-16 | IIIVmrMLM | SIG |
| SL_18XX | AX-94609323 | 6B | 218656255 | 0.1092 | | 3.20E-05 | BLUPmrMLM | SUG |
| SL_18XX | AX-89421346 | 6B | 515058787 | 0.1287 | | 5.10E-06 | BLUPmrMLM | SUG |

| | | | | | | | | |
|---|---|---|---|---|---|---|---|---|
| SL_18XX | AX-109435951 | 6D | 469993277 | 0.0227 | -0.879 | 4.10E-07 | IIIVmrMLM | SUG |
| SL_18XX | AX-110169597 | 7A | 133025679 | 0.0581 | -1.3987 | 2.08E-08 | IIIVmrMLM | SIG |
| SL_18XX | AX-110431062 | 7A | 438698521 | -0.1103 | 0.2616 | 4.04E-12 | IIIVmrMLM | SIG |
| SL_18XX | AX-109488223 | 7A | 442279147 | 0.3076 | | 6.42E-14 | BLUPmrMLM | SIG |
| SL_18XX | AX-111594910 | 7B | 19687108 | -0.1468 | | 2.81E-07 | BLUPmrMLM | SUG |
| SL_18XX | AX-109134548 | 7D | 33182408 | 0.1274 | | 1.74E-07 | BLUPmrMLM | SUG |
| SL_18XX | AX-110297437 | 7D | 185890693 | 0.6170 | | 3.16E-07 | EMMAX | SUG |
| SL_18XX | AX-110297437 | 7D | 185890693 | -0.6192 | | 1.18E-07 | GEMMA | SUG |
| SL_18XX | AX-110297437 | 7D | 185890693 | 0.1390 | | 1.37E-16 | IIIVmrMLM | SIG |
| SL_18XX | AX-94764986 | 7D | 186792308 | -0.6067 | | 1.20E-06 | GEMMA | SUG |
| SL_18XX | AX-109332089 | 7D | 187434448 | 0.3115 | | 7.89E-07 | EMMAX | SUG |
| SL_18XX | AX-109332089 | 7D | 187434448 | -0.3114 | | 2.23E-07 | GEMMA | SUG |
| SL_18XX | AX-109035303 | 7D | 188187300 | 0.3140 | | 8.37E-07 | EMMAX | SUG |
| SL_18XX | AX-109035303 | 7D | 188187300 | -0.3128 | | 2.60E-07 | GEMMA | SUG |
| SL_18XX | AX-110015666 | 7D | 188399769 | -0.3026 | | 1.06E-06 | GEMMA | SUG |
| SL_18XX | AX-110750491 | 7D | 189001615 | -0.2885 | | 1.29E-06 | GEMMA | SUG |
| SL_18XX | AX-110010796 | 7D | 189090567 | 0.2898 | | 2.24E-06 | EMMAX | SUG |
| SL_18XX | AX-110010796 | 7D | 189090567 | -0.2879 | | 8.48E-07 | GEMMA | SUG |
| SL_18XX | AX-111044062 | 7D | 189128925 | -0.2899 | | 1.23E-06 | GEMMA | SUG |
| SL_18XX | AX-110057871 | 7D | 190184754 | 0.3047 | | 1.02E-06 | EMMAX | SUG |
| SL_18XX | AX-110057871 | 7D | 190184754 | -0.3034 | | 3.23E-07 | GEMMA | SUG |
| SL_18XX | AX-111108835 | 7D | 191401968 | 0.3053 | | 1.13E-06 | EMMAX | SUG |
| SL_18XX | AX-111108835 | 7D | 191401968 | -0.3062 | | 2.77E-07 | GEMMA | SUG |
| SL_18XX | AX-111076518 | 7D | 191629005 | 0.3063 | | 1.97E-06 | EMMAX | SUG |
| SL_18XX | AX-111076518 | 7D | 191629005 | -0.3075 | | 4.75E-07 | GEMMA | SUG |
| SL_18XX | AX-109915283 | 7D | 191631703 | -0.2799 | | 1.70E-06 | GEMMA | SUG |
| SL_18XX | AX-109732612 | 7D | 191812509 | -0.2799 | | 1.70E-06 | GEMMA | SUG |
| SL_18XX | AX-111374877 | 7D | 191815180 | -0.2778 | | 1.86E-06 | GEMMA | SUG |
| SL_18XX | AX-110239974 | 7D | 191824680 | -0.2780 | | 1.91E-06 | GEMMA | SUG |
| SL_18XX | AX-111057482 | 7D | 191825383 | -0.2763 | | 2.08E-06 | GEMMA | SUG |
| SL_18XX | AX-110053968 | 7D | 191831926 | -0.2775 | | 1.86E-06 | GEMMA | SUG |
| SL_18XX | AX-111542234 | 7D | 191869153 | -0.2803 | | 1.65E-06 | GEMMA | SUG |
| SL_18XX | AX-110417203 | 7D | 191872348 | -0.2798 | | 1.65E-06 | GEMMA | SUG |
| SL_18XX | AX-109877293 | 7D | 192018380 | -0.2781 | | 1.78E-06 | GEMMA | SUG |
| SL_18XX | AX-111673313 | 7D | 192020178 | -0.2781 | | 1.78E-06 | GEMMA | SUG |
| SL_18XX | AX-109428601 | 7D | 192022692 | -0.2832 | | 1.12E-06 | GEMMA | SUG |
| SL_18XX | AX-108904641 | 7D | 192027937 | -0.2832 | | 1.12E-06 | GEMMA | SUG |
| SL_18XX | AX-111684035 | 7D | 192028016 | -0.2807 | | 1.25E-06 | GEMMA | SUG |
| SL_18XX | AX-109917900 | 7D | 192054373 | -0.2800 | | 1.19E-06 | GEMMA | SUG |
| SL_18XX | AX-111607467 | 7D | 222521294 | 0.2826 | | 1.51E-06 | EMMAX | SUG |
| SL_18XX | AX-111607467 | 7D | 222521294 | -0.2844 | | 4.23E-07 | GEMMA | SUG |
| SL_18XX | AX-110254132 | 7D | 227074119 | 0.2801 | | 1.62E-06 | EMMAX | SUG |
| SL_18XX | AX-110254132 | 7D | 227074119 | -0.2807 | | 4.91E-07 | GEMMA | SUG |
| SL_18XX | AX-111038335 | 7D | 227614380 | 0.2803 | | 1.67E-06 | EMMAX | SUG |

| | | | | | | | | |
|---|---|---|---|---|---|---|---|---|
| SL_18XX | AX-111038335 | 7D | 227614380 | -0.2814 | | 4.89E-07 | GEMMA | SUG |
| SL_18XX | AX-94734044 | 7D | 535728143 | 0.0687 | -0.1665 | 7.47E-15 | IIIVmrMLM | SIG |
| SL_18XX | AX-94567102 | 7D | 575864262 | -0.4890 | | 9.58E-07 | BLUPmrMLM | SUG |
| SL_18ZMD | AX-111160407 | 1A | 15430755 | -0.2284 | | 7.20E-05 | BLUPmrMLM | SUG |
| SL_18ZMD | AX-111652275 | 1A | 551979324 | 0.3259 | -0.1157 | 5.68E-14 | IIIVmrMLM | SIG |
| SL_18ZMD | AX-110403844 | 1A | 566562487 | -0.4381 | -0.081 | 1.22E-23 | IIIVmrMLM | SIG |
| SL_18ZMD | AX-109886705 | 1B | 24633219 | -0.0268 | -0.6052 | 2.95E-12 | IIIVmrMLM | SIG |
| SL_18ZMD | AX-108879680 | 1B | 328646759 | 0.4291 | -0.0345 | 4.69E-14 | IIIVmrMLM | SIG |
| SL_18ZMD | AX-109986674 | 1B | 566716390 | 0.0700 | -0.8067 | 5.74E-06 | IIIVmrMLM | SUG |
| SL_18ZMD | AX-111505779 | 1B | 689380879 | 0.2441 | | 3.99E-05 | BLUPmrMLM | SUG |
| SL_18ZMD | AX-109412379 | 1D | 46215959 | 0.0681 | 2.7969 | 3.69E-13 | IIIVmrMLM | SIG |
| SL_18ZMD | AX-110943965 | 2A | 83202479 | -0.0145 | 0.8011 | 4.84E-06 | IIIVmrMLM | SUG |
| SL_18ZMD | AX-109816405 | 2B | 41149142 | 0.6083 | | 2.86E-06 | BLUPmrMLM | SUG |
| SL_18ZMD | AX-109962437 | 2B | 671254874 | -0.2194 | | 1.40E-05 | BLUPmrMLM | SUG |
| SL_18ZMD | AX-109583977 | 2D | 18703610 | 0.0289 | 1.192 | 5.93E-12 | IIIVmrMLM | SIG |
| SL_18ZMD | AX-109896082 | 3A | 10221647 | 0.3507 | -0.142 | 1.65E-15 | IIIVmrMLM | SIG |
| SL_18ZMD | AX-109848595 | 3A | 24792711 | 0.2745 | | 1.31E-04 | BLUPmrMLM | SUG |
| SL_18ZMD | AX-94434803 | 3A | 57568570 | 0.1972 | | 2.15E-05 | BLUPmrMLM | SUG |
| SL_18ZMD | AX-109424866 | 3A | 547474078 | 0.1069 | 0.8525 | 1.42E-09 | IIIVmrMLM | SIG |
| SL_18ZMD | AX-109401618 | 3A | 730336396 | 0.3194 | 2.7977 | 8.87E-20 | IIIVmrMLM | SIG |
| SL_18ZMD | AX-111152411 | 3A | 733185446 | -0.1051 | -0.4957 | 1.84E-08 | IIIVmrMLM | SIG |
| SL_18ZMD | AX-108761982 | 3B | 18823307 | -0.1290 | -0.8459 | 2.87E-08 | IIIVmrMLM | SIG |
| SL_18ZMD | AX-109033369 | 3B | 62744016 | -0.1246 | -1.8155 | 5.67E-12 | IIIVmrMLM | SIG |
| SL_18ZMD | AX-89582767 | 4A | 584936217 | 0.3285 | | 2.73E-08 | BLUPmrMLM | SIG |
| SL_18ZMD | AX-89582767 | 4A | 584936217 | -0.2982 | -0.0262 | 1.47E-10 | IIIVmrMLM | SIG |
| SL_18ZMD | AX-111596465 | 4A | 627052040 | 0.0633 | 0.5286 | 3.27E-06 | IIIVmrMLM | SUG |
| SL_18ZMD | AX-111104520 | 4B | 612372577 | 0.0219 | 0.5715 | 2.23E-05 | IIIVmrMLM | SUG |
| SL_18ZMD | AX-109998028 | 4B | 642461414 | -0.0821 | 2.2218 | 6.48E-05 | IIIVmrMLM | SUG |
| SL_18ZMD | AX-108968377 | 4D | 506823199 | 0.2916 | | 2.55E-07 | BLUPmrMLM | SUG |
| SL_18ZMD | AX-111560076 | 5A | 384951371 | -0.0713 | -0.5039 | 1.61E-10 | IIIVmrMLM | SIG |
| SL_18ZMD | AX-109430223 | 5A | 439829477 | -0.0936 | -1.3062 | 1.70E-07 | IIIVmrMLM | SUG |
| SL_18ZMD | AX-109391838 | 5A | 472252124 | 0.0045 | 0.73 | 2.94E-08 | IIIVmrMLM | SIG |
| SL_18ZMD | AX-111622382 | 5A | 508210451 | -0.2661 | 0.0441 | 6.60E-08 | IIIVmrMLM | SIG |
| SL_18ZMD | AX-110948070 | 5B | 382735889 | -0.0272 | -0.7971 | 3.21E-05 | IIIVmrMLM | SUG |
| SL_18ZMD | AX-111643097 | 5B | 637390014 | 0.3888 | | 7.43E-07 | GEMMA | SUG |
| SL_18ZMD | AX-111103826 | 5B | 638671923 | 0.6591 | | 2.22E-06 | GEMMA | SUG |
| SL_18ZMD | AX-111632368 | 5B | 640396356 | -0.3722 | | 2.35E-07 | GEMMA | SUG |
| SL_18ZMD | AX-111632368 | 5B | 640396356 | 0.3478 | -0.2656 | 1.35E-15 | IIIVmrMLM | SIG |
| SL_18ZMD | AX-110448811 | 5B | 640411790 | 0.5130 | | 8.59E-07 | GEMMA | SUG |
| SL_18ZMD | AX-111584392 | 5B | 640412308 | 0.7934 | | 1.95E-06 | GEMMA | SUG |
| SL_18ZMD | AX-109375179 | 5B | 640425026 | 0.4381 | | 4.48E-07 | GEMMA | SUG |
| SL_18ZMD | AX-108806708 | 5B | 640427610 | 0.4381 | | 4.48E-07 | GEMMA | SUG |
| SL_18ZMD | AX-110023063 | 5B | 640427816 | 0.4381 | | 4.48E-07 | GEMMA | SUG |
| SL_18ZMD | AX-109358615 | 5B | 640432324 | 0.4454 | | 3.44E-07 | GEMMA | SUG |

| | | | | | | | | |
|---|---|---|---|---|---|---|---|---|
| SL_18ZMD | AX-109941289 | 5B | 640443228 | 0.4271 | | 8.81E-07 | GEMMA | SUG |
| SL_18ZMD | AX-111509106 | 5B | 640462164 | 0.4381 | | 4.48E-07 | GEMMA | SUG |
| SL_18ZMD | AX-110552173 | 5B | 640473538 | 0.4381 | | 4.48E-07 | GEMMA | SUG |
| SL_18ZMD | AX-110650282 | 5B | 640548389 | 0.4178 | | 1.38E-06 | GEMMA | SUG |
| SL_18ZMD | AX-110672325 | 5B | 640549863 | 0.5073 | | 9.12E-07 | GEMMA | SUG |
| SL_18ZMD | AX-111083236 | 5B | 640557054 | 0.4353 | | 4.43E-07 | GEMMA | SUG |
| SL_18ZMD | AX-111684814 | 5B | 641024280 | 0.4013 | | 2.29E-06 | GEMMA | SUG |
| SL_18ZMD | AX-110386773 | 5B | 641060386 | 0.4110 | | 1.55E-06 | GEMMA | SUG |
| SL_18ZMD | AX-109442841 | 5B | 641102195 | 0.4414 | | 4.05E-07 | GEMMA | SUG |
| SL_18ZMD | AX-110046475 | 5B | 641107486 | 0.4090 | | 1.67E-06 | GEMMA | SUG |
| SL_18ZMD | AX-110668137 | 5B | 641107622 | 0.4152 | | 1.29E-06 | GEMMA | SUG |
| SL_18ZMD | AX-108768159 | 5B | 641109188 | 0.4223 | | 7.75E-07 | GEMMA | SUG |
| SL_18ZMD | AX-108740514 | 5B | 641134634 | 0.4060 | | 1.57E-06 | GEMMA | SUG |
| SL_18ZMD | AX-109987277 | 5B | 641135358 | 0.4028 | | 2.10E-06 | GEMMA | SUG |
| SL_18ZMD | AX-109447725 | 5B | 641553808 | 0.4132 | | 1.35E-06 | GEMMA | SUG |
| SL_18ZMD | AX-110399021 | 5B | 644850463 | 0.4828 | | 4.09E-08 | GEMMA | SIG |
| SL_18ZMD | AX-109475699 | 5B | 644853684 | 0.4502 | | 1.85E-07 | GEMMA | SUG |
| SL_18ZMD | AX-108993341 | 5B | 644854785 | 0.9083 | | 1.27E-07 | GEMMA | SUG |
| SL_18ZMD | AX-109317067 | 5B | 644856940 | 0.4214 | | 1.99E-06 | GEMMA | SUG |
| SL_18ZMD | AX-111508432 | 5B | 644857108 | 0.5289 | | 9.82E-07 | GEMMA | SUG |
| SL_18ZMD | AX-111457323 | 5D | 477875557 | -0.2472 | | 1.52E-06 | BLUPmrMLM | SUG |
| SL_18ZMD | AX-108980620 | 6A | 1807418 | -0.2788 | 3.026 | 3.06E-16 | IIIVmrMLM | SIG |
| SL_18ZMD | AX-111101623 | 6A | 61891179 | -0.2049 | -0.0996 | 3.36E-05 | IIIVmrMLM | SUG |
| SL_18ZMD | AX-94876976 | 6A | 614669489 | 0.3294 | | 5.26E-05 | BLUPmrMLM | SUG |
| SL_18ZMD | AX-110374034 | 6B | 20077005 | -0.0269 | 1.4555 | 1.29E-04 | IIIVmrMLM | SUG |
| SL_18ZMD | AX-110012043 | 6D | 137016269 | -0.2976 | | 6.36E-13 | IIIVmrMLM | SIG |
| SL_18ZMD | AX-94640722 | 7A | 5051737 | 0.0515 | -0.6134 | 1.14E-04 | IIIVmrMLM | SUG |
| SL_18ZMD | AX-89639361 | 7A | 129867656 | -0.2711 | | 2.84E-06 | BLUPmrMLM | SUG |
| SL_18ZMD | AX-111760021 | 7A | 695638939 | 0.2322 | | 9.36E-05 | BLUPmrMLM | SUG |
| SL_18ZMD | AX-110500747 | 7B | 683398840 | 0.4480 | | 4.44E-08 | BLUPmrMLM | SIG |
| SL_18ZMD | AX-110467729 | 7D | 43386933 | 0.0031 | -0.5854 | 1.07E-05 | IIIVmrMLM | SUG |
| SL_18ZMD | AX-111266686 | 7D | 638157164 | -6.1509 | | 6.34E-08 | EMMAX | SIG |
| SL_BLUP | AX-111535253 | 1A | 15525424 | 0.0297 | -0.3182 | 2.16E-05 | IIIVmrMLM | SUG |
| SL_BLUP | AX-110362532 | 1A | 552126201 | -0.0439 | -0.2895 | 2.24E-13 | IIIVmrMLM | SIG |
| SL_BLUP | AX-111064668 | 1A | 585753393 | 0.1179 | | 3.56E-06 | BLUPmrMLM | SUG |
| SL_BLUP | AX-111586591 | 1B | 94979342 | -0.0725 | | 2.06E-05 | BLUPmrMLM | SUG |
| SL_BLUP | AX-109330456 | 1B | 449932934 | -0.0851 | | 1.04E-04 | BLUPmrMLM | SUG |
| SL_BLUP | AX-110975095 | 1B | 623679022 | -0.0103 | 0.4294 | 5.14E-06 | IIIVmrMLM | SUG |
| SL_BLUP | AX-110435866 | 1B | 668724693 | -0.0350 | 0.2109 | 7.47E-06 | IIIVmrMLM | SUG |
| SL_BLUP | AX-110373686 | 1D | 8234578 | -0.0990 | | 3.62E-05 | BLUPmrMLM | SUG |
| SL_BLUP | AX-109600192 | 1D | 283189988 | -0.1189 | | 7.41E-06 | BLUPmrMLM | SUG |
| SL_BLUP | AX-108873616 | 2A | 193732765 | -0.1129 | -0.0917 | 6.96E-14 | IIIVmrMLM | SIG |
| SL_BLUP | AX-110399256 | 2A | 542661733 | 0.0084 | -0.3437 | 5.37E-07 | IIIVmrMLM | SUG |
| SL_BLUP | AX-108837102 | 2A | 611821955 | -0.0851 | 0.4496 | 3.81E-09 | IIIVmrMLM | SIG |

| | | | | | | | | |
|---|---|---|---|---|---|---|---|---|
| SL_BLUP | AX-110530703 | 2A | 617725615 | -0.0956 | | 7.98E-05 | BLUPmrMLM | SUG |
| SL_BLUP | AX-110200150 | 2A | 654174838 | 0.1028 | | 3.61E-05 | BLUPmrMLM | SUG |
| SL_BLUP | AX-111193990 | 2B | 9724504 | -0.0692 | | 1.05E-06 | IIIVmrMLM | SUG |
| SL_BLUP | AX-108985575 | 2B | 703665690 | 0.0638 | -0.084 | 3.29E-05 | IIIVmrMLM | SUG |
| SL_BLUP | AX-109844617 | 2D | 21911530 | 0.1959 | | 1.58E-15 | BLUPmrMLM | SIG |
| SL_BLUP | AX-109842248 | 2D | 23102530 | 0.1619 | | 1.38E-06 | GEMMA | SUG |
| SL_BLUP | AX-108991361 | 2D | 23304090 | 0.1654 | | 8.90E-07 | GEMMA | SUG |
| SL_BLUP | AX-111038058 | 2D | 23417635 | -0.1253 | 0.0373 | 1.07E-16 | IIIVmrMLM | SIG |
| SL_BLUP | AX-109627287 | 2D | 116801772 | -0.0071 | -0.5872 | 2.10E-07 | IIIVmrMLM | SUG |
| SL_BLUP | AX-108882839 | 3A | 12323991 | 0.1319 | | 5.99E-08 | BLUPmrMLM | SIG |
| SL_BLUP | AX-111531488 | 3A | 22533581 | -0.0134 | 0.5879 | 1.40E-04 | IIIVmrMLM | SUG |
| SL_BLUP | AX-110679559 | 3A | 485912014 | 0.0639 | -0.3457 | 5.05E-11 | IIIVmrMLM | SIG |
| SL_BLUP | AX-110561877 | 3A | 726127025 | 0.1658 | | 3.41E-09 | BLUPmrMLM | SIG |
| SL_BLUP | AX-109033369 | 3B | 62744016 | -0.0415 | -0.4763 | 1.93E-08 | IIIVmrMLM | SIG |
| SL_BLUP | AX-94615079 | 3B | 232567634 | -0.0893 | | 1.95E-04 | BLUPmrMLM | SUG |
| SL_BLUP | AX-109404313 | 3B | 468041202 | 0.1644 | -0.0529 | 3.65E-25 | IIIVmrMLM | SIG |
| SL_BLUP | AX-110934736 | 3B | 470495728 | 0.1245 | | 1.57E-07 | BLUPmrMLM | SUG |
| SL_BLUP | AX-109447402 | 3B | 538215295 | 0.0474 | -0.1663 | 4.72E-06 | IIIVmrMLM | SUG |
| SL_BLUP | AX-108781578 | 3B | 662523075 | 0.0009 | -0.2603 | 6.59E-05 | IIIVmrMLM | SUG |
| SL_BLUP | AX-110275510 | 3D | 474995279 | -0.0840 | -0.0435 | 2.94E-08 | IIIVmrMLM | SIG |
| SL_BLUP | AX-94541915 | 3D | 496785393 | -0.2668 | | 8.22E-64 | IIIVmrMLM | SIG |
| SL_BLUP | AX-95083270 | 3D | 547627077 | 0.3052 | | 1.31E-10 | BLUPmrMLM | SIG |
| SL_BLUP | AX-111488831 | 3D | 584977626 | 0.6817 | 1.2113 | 8.76E-230 | IIIVmrMLM | SIG |
| SL_BLUP | AX-108968290 | 4A | 11295242 | -0.0092 | 0.7913 | 3.87E-04 | IIIVmrMLM | SUG |
| SL_BLUP | AX-108921906 | 4A | 29020483 | 0.1089 | | 3.51E-14 | IIIVmrMLM | SIG |
| SL_BLUP | AX-111676988 | 4A | 40194335 | 0.0346 | 0.2831 | 7.68E-11 | IIIVmrMLM | SIG |
| SL_BLUP | AX-111800028 | 4A | 97542828 | 0.0386 | -0.3781 | 3.47E-09 | IIIVmrMLM | SIG |
| SL_BLUP | AX-108781086 | 4A | 106364121 | -0.1222 | | 3.51E-05 | BLUPmrMLM | SUG |
| SL_BLUP | AX-108781086 | 4A | 106364121 | 0.1550 | 0.075 | 5.01E-16 | IIIVmrMLM | SIG |
| SL_BLUP | AX-95023120 | 4A | 443441677 | 0.0841 | | 3.35E-05 | BLUPmrMLM | SUG |
| SL_BLUP | AX-110551959 | 4B | 601196352 | -0.0853 | | 5.36E-05 | BLUPmrMLM | SUG |
| SL_BLUP | AX-110516351 | 4B | 637364174 | -0.0814 | | 9.75E-06 | BLUPmrMLM | SUG |
| SL_BLUP | AX-110368142 | 5A | 444523191 | 0.1260 | 0.1406 | 3.03E-17 | IIIVmrMLM | SIG |
| SL_BLUP | AX-110954024 | 5A | 475270864 | -0.0723 | | 1.72E-04 | BLUPmrMLM | SUG |
| SL_BLUP | AX-111622382 | 5A | 508210451 | 0.0962 | | 1.42E-05 | BLUPmrMLM | SUG |
| SL_BLUP | AX-95096584 | 5A | 564533573 | -0.0841 | | 3.51E-09 | IIIVmrMLM | SIG |
| SL_BLUP | AX-111540500 | 5A | 565491466 | 0.0796 | | 7.83E-05 | BLUPmrMLM | SUG |
| SL_BLUP | AX-110425511 | 5B | 418009115 | -0.0221 | 0.3152 | 2.11E-04 | IIIVmrMLM | SUG |
| SL_BLUP | AX-109929907 | 5B | 522109479 | 0.0837 | | 1.64E-04 | BLUPmrMLM | SUG |
| SL_BLUP | AX-110363386 | 5B | 612680877 | 0.1151 | | 8.03E-06 | BLUPmrMLM | SUG |
| SL_BLUP | AX-111676716 | 5B | 669607184 | -0.0690 | | 1.53E-04 | BLUPmrMLM | SUG |
| SL_BLUP | AX-110625313 | 5D | 478883726 | -0.1241 | | 1.90E-09 | BLUPmrMLM | SIG |
| SL_BLUP | AX-108801122 | 6A | 328393744 | 0.1512 | | 2.13E-08 | BLUPmrMLM | SIG |

| | | | | | | | | |
|---|---|---|---|---|---|---|---|---|
| SL_BLUP | AX-110733835 | 6A | 360493631 | 0.4044 | | 3.65E-07 | FarmCPU | SUG |
| SL_BLUP | AX-108753591 | 6A | 612660211 | 0.0015 | 0.3633 | 1.23E-14 | IIIVmrMLM | SIG |
| SL_BLUP | AX-108884781 | 6B | 30299119 | 0.0632 | | 1.47E-04 | BLUPmrMLM | SUG |
| SL_BLUP | AX-109646483 | 6B | 703265547 | -0.0311 | 0.1817 | 1.94E-05 | IIIVmrMLM | SUG |
| SL_BLUP | AX-109397671 | 6B | 718924441 | 0.1811 | 0.029 | 4.45E-16 | IIIVmrMLM | SIG |
| SL_BLUP | AX-94745553 | 6D | 6276611 | -0.0088 | 0.1109 | 2.70E-06 | IIIVmrMLM | SUG |
| SL_BLUP | AX-95257962 | 6D | 28409241 | -0.0730 | | 6.84E-05 | BLUPmrMLM | SUG |
| SL_BLUP | AX-111644101 | 6D | 380442210 | -0.0755 | 0.2514 | 9.53E-08 | IIIVmrMLM | SIG |
| SL_BLUP | AX-108844053 | 7A | 85338508 | 0.0767 | | 8.80E-07 | BLUPmrMLM | SUG |
| SL_BLUP | AX-111631763 | 7A | 159325506 | -0.0566 | -0.2311 | 4.87E-10 | IIIVmrMLM | SIG |
| SL_BLUP | AX-111590324 | 7B | 150886421 | 0.0028 | -0.3949 | 6.62E-10 | IIIVmrMLM | SIG |
| SL_BLUP | AX-94448771 | 7B | 684397290 | 0.1082 | | 4.60E-06 | BLUPmrMLM | SUG |
| SL_BLUP | AX-108843451 | 7B | 730422639 | -0.1044 | -0.2869 | 2.62E-44 | IIIVmrMLM | SIG |
| SL_BLUP | AX-111598705 | 7B | 733609157 | -0.0850 | | 2.67E-05 | BLUPmrMLM | SUG |
| SL_BLUP | AX-111061288 | 7D | 66537922 | -0.0726 | -0.0362 | 5.27E-06 | IIIVmrMLM | SUG |
| SL_BLUP | AX-110898194 | 7D | 168777588 | 0.1217 | | 6.04E-05 | BLUPmrMLM | SUG |
| SL_BLUP | AX-110946735 | 7D | 395846311 | 0.0883 | 0.0216 | 6.10E-09 | IIIVmrMLM | SIG |
| SL_BLUP | AX-109962102 | 7D | 605157582 | -0.1172 | -0.446 | 3.79E-17 | IIIVmrMLM | SIG |

SL: spike length

**Supplementary Table S6**. All QTNs identified as associated with ratio of peduncle length to plant height in wheat.

| Trait | Marker | Chr | Position | Add.effect | Dom.effect | P-value | Method | Sig. |
|---|---|---|---|---|---|---|---|---|
| PPR_18AY | AX-109479907 | 1D | 46323974 | -0.0012 | -0.0459 | 1.51E-10 | IIIVmrMLM | SIG |
| PPR_18AY | AX-110113706 | 2A | 8758726 | -0.0070 | 0.0015 | 1.54E-07 | IIIVmrMLM | SUG |
| PPR_18AY | AX-111741582 | 2A | 414988352 | 0.0010 | 0.0245 | 1.21E-06 | IIIVmrMLM | SUG |
| PPR_18AY | AX-109576912 | 2B | 131302364 | 0.0011 | 0.0341 | 4.57E-07 | IIIVmrMLM | SUG |
| PPR_18AY | AX-95215702 | 2B | 260648430 | -0.0099 | 0.0057 | 1.09E-18 | IIIVmrMLM | SIG |
| PPR_18AY | AX-111577035 | 2B | 415307095 | -0.0026 | 0.0368 | 3.95E-09 | IIIVmrMLM | SIG |
| PPR_18AY | AX-110528509 | 2B | 540438405 | -0.0039 | -0.0179 | 1.97E-10 | IIIVmrMLM | SIG |
| PPR_18AY | AX-111476923 | 2B | 743728971 | -0.0137 | | 1.03E-06 | GEMMA | SUG |
| PPR_18AY | AX-111476923 | 2B | 743728971 | 0.0106 | 0.0020 | 4.35E-11 | IIIVmrMLM | SIG |
| PPR_18AY | AX-110924593 | 2B | 743803249 | 0.0091 | | 1.49E-06 | GEMMA | SUG |
| PPR_18AY | AX-108826933 | 2D | 506202563 | -0.0074 | -0.0191 | 7.93E-12 | IIIVmrMLM | SIG |
| PPR_18AY | AX-111211824 | 2D | 647455478 | 0.0052 | | 6.49E-07 | IIIVmrMLM | SUG |
| PPR_18AY | AX-94727962 | 3A | 461393820 | 0.0070 | | 7.62E-06 | BLUPmrMLM | SUG |
| PPR_18AY | AX-111491995 | 3A | 715280633 | 0.0055 | | 9.15E-05 | BLUPmrMLM | SUG |
| PPR_18AY | AX-110923489 | 3B | 15009564 | -0.0063 | | 1.13E-05 | BLUPmrMLM | SUG |
| PPR_18AY | AX-110923489 | 3B | 15009564 | 0.0068 | 0.0050 | 9.01E-10 | IIIVmrMLM | SIG |
| PPR_18AY | AX-110368038 | 3B | 216895237 | -0.0021 | -0.0246 | 2.22E-04 | IIIVmrMLM | SUG |
| PPR_18AY | AX-111780324 | 4A | 13285565 | -0.0045 | -0.0160 | 1.23E-09 | IIIVmrMLM | SIG |
| PPR_18AY | AX-111486923 | 4B | 407072254 | -0.0001 | 0.0362 | 1.22E-04 | IIIVmrMLM | SUG |
| PPR_18AY | AX-111484642 | 5A | 6482214 | 0.0090 | | 8.10E-06 | BLUPmrMLM | SUG |
| PPR_18AY | AX-110433504 | 5A | 588845013 | 0.0073 | | 1.86E-05 | BLUPmrMLM | SUG |
| PPR_18AY | AX-110510319 | 5A | 660186076 | 0.0070 | | 3.32E-11 | IIIVmrMLM | SIG |
| PPR_18AY | AX-111611290 | 5B | 13187554 | 0.0015 | -0.0322 | 2.97E-06 | IIIVmrMLM | SUG |
| PPR_18AY | AX-111524067 | 5B | 34291504 | 0.0084 | | 6.52E-07 | BLUPmrMLM | SUG |
| PPR_18AY | AX-109909648 | 5B | 34782236 | -0.0066 | 0.0012 | 2.10E-07 | IIIVmrMLM | SUG |
| PPR_18AY | AX-109907795 | 5B | 314351620 | 0.0004 | 0.0100 | 4.17E-05 | IIIVmrMLM | SUG |
| PPR_18AY | AX-111685579 | 5B | 594930571 | 0.0040 | 0.0198 | 9.58E-14 | IIIVmrMLM | SIG |
| PPR_18AY | AX-111677549 | 5D | 549578793 | 0.0086 | | 4.23E-07 | BLUPmrMLM | SUG |
| PPR_18AY | AX-110286706 | 5D | 550421747 | -0.0028 | -0.0449 | 9.34E-09 | IIIVmrMLM | SIG |
| PPR_18AY | AX-109418119 | 5D | 559683036 | 0.0015 | 0.0264 | 1.92E-06 | IIIVmrMLM | SUG |
| PPR_18AY | AX-108746385 | 6B | 30171150 | -0.0004 | -0.0132 | 8.83E-04 | IIIVmrMLM | SUG |
| PPR_18AY | AX-110994392 | 6B | 260295797 | 0.0188 | | 1.24E-07 | FarmCPU | SUG |
| PPR_18AY | AX-109422752 | 7A | 20604036 | -0.0046 | 0.0009 | 9.07E-05 | IIIVmrMLM | SUG |
| PPR_18AY | AX-109383924 | 7A | 70501352 | -0.0161 | | 9.63E-07 | FarmCPU | SUG |
| PPR_18AY | AX-110167659 | 7A | 448713163 | 0.0022 | 0.0125 | 6.08E-05 | IIIVmrMLM | SUG |
| PPR_18AY | AX-109275127 | 7A | 646589336 | 0.0012 | 0.0257 | 1.41E-05 | IIIVmrMLM | SUG |
| PPR_18AY | AX-108875352 | 7A | 670975129 | -0.0019 | 0.0223 | 8.06E-09 | IIIVmrMLM | SIG |
| PPR_18AY | AX-110959031 | 7B | 57845846 | 0.0005 | 0.0168 | 2.04E-05 | IIIVmrMLM | SUG |
| PPR_18AY | AX-110589973 | 7B | 594394840 | 0.0092 | | 4.59E-05 | BLUPmrMLM | SUG |
| PPR_18AY | AX-110589973 | 7B | 594394840 | -0.0110 | 0.0042 | 1.28E-20 | IIIVmrMLM | SIG |

| | | | | | | | | |
|---|---|---|---|---|---|---|---|---|
| PPR_18JY | AX-111119411 | 1A | 544922567 | -0.0073 | | 7.16E-07 | FarmCPU | SUG |
| PPR_18JY | AX-110986258 | 1B | 65908256 | -0.0029 | | 1.25E-04 | BLUPmrMLM | SUG |
| PPR_18JY | AX-111495399 | 1B | 299717566 | -0.0011 | -0.0078 | 4.91E-08 | IIIVmrMLM | SIG |
| PPR_18JY | AX-94688951 | 1B | 317013369 | -0.0054 | | 7.74E-05 | BLUPmrMLM | SUG |
| PPR_18JY | AX-109858910 | 1B | 483878226 | -0.0038 | | 1.80E-06 | BLUPmrMLM | SUG |
| PPR_18JY | AX-111022013 | 1B | 549361711 | 0.0089 | | 1.88E-06 | FarmCPU | SUG |
| PPR_18JY | AX-110741578 | 1B | 655092881 | -0.0032 | 0.0008 | 8.06E-07 | IIIVmrMLM | SUG |
| PPR_18JY | AX-110077846 | 2A | 88983293 | 0.0021 | 0.0140 | 4.06E-10 | IIIVmrMLM | SIG |
| PPR_18JY | AX-111601071 | 2A | 382184186 | 0.0012 | 0.0115 | 3.84E-13 | IIIVmrMLM | SIG |
| PPR_18JY | AX-111559860 | 2A | 407372832 | 0.0073 | | 3.38E-07 | BLUPmrMLM | SUG |
| PPR_18JY | AX-111514059 | 2A | 500708451 | 0.0019 | -0.0182 | 2.10E-15 | IIIVmrMLM | SIG |
| PPR_18JY | AX-110374260 | 2A | 515451216 | 0.0066 | 0.0028 | 3.32E-26 | IIIVmrMLM | SIG |
| PPR_18JY | AX-110593881 | 2A | 641798939 | 0.0004 | -0.0195 | 4.39E-06 | IIIVmrMLM | SUG |
| PPR_18JY | AX-109477982 | 2A | 755120274 | 0.0042 | 0.0011 | 4.44E-10 | IIIVmrMLM | SIG |
| PPR_18JY | AX-94385196 | 2A | 775619943 | 0.0506 | | 4.77E-07 | FarmCPU | SUG |
| PPR_18JY | AX-111289464 | 2B | 157856040 | -0.0040 | -0.0004 | 2.78E-09 | IIIVmrMLM | SIG |
| PPR_18JY | AX-109311497 | 2B | 158776798 | -0.0033 | | 1.54E-05 | BLUPmrMLM | SUG |
| PPR_18JY | AX-109815095 | 2D | 628941279 | -0.0011 | 0.0097 | 4.08E-08 | IIIVmrMLM | SIG |
| PPR_18JY | AX-111147717 | 3A | 560292782 | 0.0038 | | 7.34E-05 | BLUPmrMLM | SUG |
| PPR_18JY | AX-109917524 | 3A | 657302203 | 0.0045 | | 4.01E-06 | BLUPmrMLM | SUG |
| PPR_18JY | AX-110675156 | 3B | 14327321 | -0.0077 | 0.0004 | 1.30E-24 | IIIVmrMLM | SIG |
| PPR_18JY | AX-94687599 | 3B | 64699148 | 0.0039 | -0.0019 | 1.96E-08 | IIIVmrMLM | SIG |
| PPR_18JY | AX-108782041 | 3B | 103032791 | 0.0008 | 0.0197 | 4.18E-15 | IIIVmrMLM | SIG |
| PPR_18JY | AX-86166069 | 3B | 699673316 | -0.0010 | 0.0124 | 3.87E-04 | IIIVmrMLM | SUG |
| PPR_18JY | AX-110461838 | 3B | 760437784 | 0.0121 | | 1.65E-07 | FarmCPU | SUG |
| PPR_18JY | AX-109306055 | 4A | 106583729 | -0.0043 | | 1.44E-04 | BLUPmrMLM | SUG |
| PPR_18JY | AX-110033540 | 4A | 467299944 | 0.0064 | 0.0026 | 1.58E-24 | IIIVmrMLM | SIG |
| PPR_18JY | AX-110370878 | 4A | 509595685 | -0.0023 | | 6.60E-05 | IIIVmrMLM | SUG |
| PPR_18JY | AX-108747090 | 4A | 708706473 | 0.0020 | 0.0345 | 1.69E-16 | IIIVmrMLM | SIG |
| PPR_18JY | AX-109593911 | 4D | 156408099 | -0.0041 | | 4.15E-12 | IIIVmrMLM | SIG |
| PPR_18JY | AX-109934789 | 5A | 7715630 | 0.0041 | | 8.38E-05 | BLUPmrMLM | SUG |
| PPR_18JY | AX-111043720 | 5A | 438638250 | 0.0020 | 0.0230 | 2.53E-08 | IIIVmrMLM | SIG |
| PPR_18JY | AX-109476993 | 5A | 592681648 | 0.0054 | | 1.15E-08 | BLUPmrMLM | SIG |
| PPR_18JY | AX-110415945 | 5A | 593164247 | 0.0065 | | 2.02E-06 | GEMMA | SUG |
| PPR_18JY | AX-94535815 | 5A | 593172184 | 0.0066 | | 1.83E-06 | GEMMA | SUG |
| PPR_18JY | AX-110966473 | 5A | 595376010 | -0.0044 | -0.0041 | 9.26E-13 | IIIVmrMLM | SIG |
| PPR_18JY | AX-110422393 | 5B | 242188636 | -0.0003 | -0.0074 | 4.42E-07 | IIIVmrMLM | SUG |
| PPR_18JY | AX-109381150 | 5B | 685072957 | -0.0007 | -0.0136 | 4.26E-06 | IIIVmrMLM | SUG |
| PPR_18JY | AX-111200760 | 5B | 707134710 | 0.0034 | | 7.75E-05 | BLUPmrMLM | SUG |
| PPR_18JY | AX-111200760 | 5B | 707134710 | -0.0039 | 0.0019 | 3.07E-10 | IIIVmrMLM | SIG |
| PPR_18JY | AX-94736165 | 5D | 6355274 | -0.0188 | | 1.25E-04 | BLUPmrMLM | SUG |
| PPR_18JY | AX-110549552 | 5D | 26827571 | 0.0031 | | 1.18E-06 | BLUPmrMLM | SUG |
| PPR_18JY | AX-111030610 | 5D | 106818950 | -0.0043 | | 4.89E-05 | BLUPmrMLM | SUG |
| PPR_18JY | AX-94880993 | 5D | 362885728 | 0.0071 | | 5.45E-05 | BLUPmrMLM | SUG |

| | | | | | | | | |
|---|---|---|---|---|---|---|---|---|
| PPR_18JY | AX-89417887 | 5D | 480184657 | -0.0041 | | 5.95E-06 | BLUPmrMLM | SUG |
| PPR_18JY | AX-110020513 | 6A | 2268652 | -0.0052 | | 3.77E-05 | BLUPmrMLM | SUG |
| PPR_18JY | AX-109548493 | 6A | 2954769 | 0.0014 | 0.0093 | 2.88E-07 | IIIVmrMLM | SUG |
| PPR_18JY | AX-110636510 | 6A | 55010969 | 0.0000 | 0.0212 | 6.22E-07 | IIIVmrMLM | SUG |
| PPR_18JY | AX-109529400 | 6A | 422940016 | -0.0067 | | 2.89E-11 | BLUPmrMLM | SIG |
| PPR_18JY | AX-109529400 | 6A | 422940016 | -0.0077 | | 2.08E-06 | GEMMA | SUG |
| PPR_18JY | AX-110462127 | 6A | 608335749 | -0.0039 | -0.0006 | 2.34E-08 | IIIVmrMLM | SIG |
| PPR_18JY | AX-94932242 | 6B | 4877531 | 0.0059 | -0.0020 | 3.20E-17 | IIIVmrMLM | SIG |
| PPR_18JY | AX-108762584 | 6B | 43470393 | 0.0038 | | 3.53E-05 | BLUPmrMLM | SUG |
| PPR_18JY | AX-110433547 | 6B | 81027892 | -0.0007 | -0.0085 | 8.85E-10 | IIIVmrMLM | SIG |
| PPR_18JY | AX-111475876 | 6B | 296408471 | 0.0054 | | 5.06E-08 | BLUPmrMLM | SIG |
| PPR_18JY | AX-94909830 | 6B | 470809494 | 0.0040 | | 1.53E-11 | IIIVmrMLM | SIG |
| PPR_18JY | AX-110966467 | 6B | 480376599 | -0.0022 | 0.0088 | 6.94E-15 | IIIVmrMLM | SIG |
| PPR_18JY | AX-109503672 | 6B | 527841682 | 0.0038 | | 1.13E-04 | BLUPmrMLM | SUG |
| PPR_18JY | AX-109859491 | 6B | 679615860 | -0.0009 | -0.0190 | 6.15E-07 | IIIVmrMLM | SUG |
| PPR_18JY | AX-111665888 | 6B | 696557776 | -0.0032 | | 1.48E-05 | BLUPmrMLM | SUG |
| PPR_18JY | AX-109468042 | 6D | 429133286 | 0.0239 | | 1.51E-06 | FarmCPU | SUG |
| PPR_18JY | AX-111978566 | 6D | 463878938 | 0.0041 | | 9.46E-06 | BLUPmrMLM | SUG |
| PPR_18JY | AX-95201556 | 7A | 2717009 | 0.0033 | | 1.78E-05 | BLUPmrMLM | SUG |
| PPR_18JY | AX-111514591 | 7A | 27904256 | -0.0047 | 0.0033 | 2.85E-14 | IIIVmrMLM | SIG |
| PPR_18JY | AX-109403952 | 7A | 65686854 | -0.0011 | -0.0241 | 4.53E-10 | IIIVmrMLM | SIG |
| PPR_18JY | AX-111226547 | 7A | 677580690 | 0.0048 | | 2.94E-06 | BLUPmrMLM | SUG |
| PPR_18JY | AX-110430158 | 7B | 10965869 | -0.0391 | | 1.73E-06 | FarmCPU | SUG |
| PPR_18JY | AX-111042153 | 7B | 41287475 | 0.0048 | 0.0021 | 3.97E-13 | IIIVmrMLM | SIG |
| PPR_18JY | AX-110953520 | 7B | 51348206 | 0.0043 | 0.0038 | 1.61E-11 | IIIVmrMLM | SIG |
| PPR_18JY | AX-109908748 | 7B | 53114653 | 0.0059 | | 2.04E-07 | BLUPmrMLM | SUG |
| PPR_18JY | AX-109908748 | 7B | 53114653 | -0.0069 | 0.0010 | 3.80E-24 | IIIVmrMLM | SIG |
| PPR_18XX | AX-94810188 | 1A | 319320199 | 0.0027 | | 1.90E-06 | IIIVmrMLM | SUG |
| PPR_18XX | AX-110361855 | 1A | 486822582 | 0.0053 | | 4.09E-06 | BLUPmrMLM | SUG |
| PPR_18XX | AX-109823728 | 1B | 536591695 | -0.0047 | | 5.30E-06 | BLUPmrMLM | SUG |
| PPR_18XX | AX-109788356 | 1D | 147718657 | -0.0083 | 0.0026 | 4.21E-40 | IIIVmrMLM | SIG |
| PPR_18XX | AX-111064741 | 2A | 43050297 | -0.0021 | -0.0153 | 2.00E-12 | IIIVmrMLM | SIG |
| PPR_18XX | AX-109466795 | 2A | 182685104 | 0.0029 | -0.0075 | 2.44E-09 | IIIVmrMLM | SIG |
| PPR_18XX | AX-109537275 | 2A | 333741957 | 0.0011 | -0.0137 | 9.42E-12 | IIIVmrMLM | SIG |
| PPR_18XX | AX-111097244 | 2A | 475814810 | -0.0211 | | 5.98E-07 | EMMAX | SUG |
| PPR_18XX | AX-111097244 | 2A | 475814810 | 0.0202 | | 1.17E-06 | GEMMA | SUG |
| PPR_18XX | AX-109511786 | 2A | 641604707 | -0.0031 | -0.0122 | 5.39E-14 | IIIVmrMLM | SIG |
| PPR_18XX | AX-109312185 | 2A | 758923227 | 0.0003 | 0.0078 | 3.64E-04 | IIIVmrMLM | SUG |
| PPR_18XX | AX-108989039 | 2B | 26098046 | 0.0061 | | 8.38E-05 | BLUPmrMLM | SUG |
| PPR_18XX | AX-111597275 | 2B | 41414402 | 0.0012 | 0.0432 | 1.65E-11 | IIIVmrMLM | SIG |
| PPR_18XX | AX-111182005 | 2B | 291614871 | -0.0001 | 0.0258 | 2.85E-15 | IIIVmrMLM | SIG |
| PPR_18XX | AX-111680356 | 2B | 385175775 | -0.0065 | | 3.83E-06 | BLUPmrMLM | SUG |
| PPR_18XX | AX-89399113 | 2B | 546587929 | 0.0007 | 0.0347 | 5.45E-10 | IIIVmrMLM | SIG |
| PPR_18XX | AX-108877101 | 2D | 350616746 | 0.0079 | | 2.01E-04 | BLUPmrMLM | SUG |

| | | | | | | | | |
|---|---|---|---|---|---|---|---|---|
| PPR_18XX | AX-109927541 | 2D | 600440215 | -0.0030 | | 1.04E-04 | BLUPmrMLM | SUG |
| PPR_18XX | AX-110553011 | 3A | 226802 | 0.0040 | | 1.58E-04 | BLUPmrMLM | SUG |
| PPR_18XX | AX-110444413 | 3A | 57012474 | 0.0008 | -0.0427 | 1.37E-12 | IIIVmrMLM | SIG |
| PPR_18XX | AX-109983001 | 3A | 691350457 | 0.0023 | 0.0138 | 1.98E-08 | IIIVmrMLM | SIG |
| PPR_18XX | AX-109966075 | 3A | 726521491 | 0.0044 | | 1.34E-04 | BLUPmrMLM | SUG |
| PPR_18XX | AX-111541115 | 3B | 283126328 | -0.0020 | 0.0318 | 6.22E-15 | IIIVmrMLM | SIG |
| PPR_18XX | AX-109924687 | 3B | 285094647 | 0.0256 | | 9.72E-07 | EMMAX | SUG |
| PPR_18XX | AX-109924687 | 3B | 285094647 | -0.0251 | | 2.08E-06 | GEMMA | SUG |
| PPR_18XX | AX-108967270 | 3B | 581128508 | -0.0114 | | 1.64E-69 | IIIVmrMLM | SIG |
| PPR_18XX | AX-108918805 | 3B | 655638189 | -0.0470 | | 3.54E-07 | EMMAX | SUG |
| PPR_18XX | AX-108918805 | 3B | 655638189 | 0.0468 | | 2.62E-07 | GEMMA | SUG |
| PPR_18XX | AX-110053308 | 3B | 761333202 | 0.0104 | | 1.02E-06 | EMMAX | SUG |
| PPR_18XX | AX-110072018 | 3D | 438947232 | -0.0035 | | 1.12E-04 | BLUPmrMLM | SUG |
| PPR_18XX | AX-94541915 | 3D | 496785393 | -0.0446 | | 1.11E-06 | EMMAX | SUG |
| PPR_18XX | AX-94541915 | 3D | 496785393 | 0.0444 | | 7.54E-07 | GEMMA | SUG |
| PPR_18XX | AX-110160363 | 3D | 516427576 | 0.0053 | -0.0124 | 3.41E-19 | IIIVmrMLM | SIG |
| PPR_18XX | AX-111397740 | 3D | 568832931 | 0.0196 | | 9.15E-08 | FarmCPU | SIG |
| PPR_18XX | AX-111167627 | 4A | 82255635 | 0.0002 | -0.0087 | 1.69E-08 | IIIVmrMLM | SIG |
| PPR_18XX | AX-109293726 | 4A | 707514936 | -0.0014 | -0.0484 | 3.36E-14 | IIIVmrMLM | SIG |
| PPR_18XX | AX-86178637 | 4A | 739525562 | -0.0032 | -0.0118 | 1.66E-17 | IIIVmrMLM | SIG |
| PPR_18XX | AX-108812567 | 4B | 3886625 | 0.0062 | | 1.03E-25 | IIIVmrMLM | SIG |
| PPR_18XX | AX-110669437 | 4B | 194001678 | 0.0135 | | 1.72E-07 | BLUPmrMLM | SUG |
| PPR_18XX | AX-110363537 | 4B | 533984924 | -0.0174 | | 2.24E-06 | EMMAX | SUG |
| PPR_18XX | AX-110363537 | 4B | 533984924 | 0.0174 | | 1.92E-06 | GEMMA | SUG |
| PPR_18XX | AX-110030611 | 4B | 534066903 | -0.0219 | | 7.19E-07 | EMMAX | SUG |
| PPR_18XX | AX-110030611 | 4B | 534066903 | 0.0224 | | 4.60E-07 | GEMMA | SUG |
| PPR_18XX | AX-109874653 | 4B | 534407516 | 0.0044 | | 1.12E-04 | BLUPmrMLM | SUG |
| PPR_18XX | AX-109606142 | 4D | 428962895 | -0.0041 | | 2.31E-12 | IIIVmrMLM | SIG |
| PPR_18XX | AX-111579762 | 4D | 459272818 | 0.0046 | -0.0019 | 3.34E-11 | IIIVmrMLM | SIG |
| PPR_18XX | AX-111480164 | 4D | 503600992 | -0.0038 | 0.0432 | 3.58E-12 | IIIVmrMLM | SIG |
| PPR_18XX | AX-110135066 | 5A | 559841777 | 0.0264 | | 1.67E-06 | GEMMA | SUG |
| PPR_18XX | AX-110362233 | 5B | 239401583 | 0.0031 | -0.0129 | 1.13E-27 | IIIVmrMLM | SIG |
| PPR_18XX | AX-109866169 | 5B | 487730650 | 0.0012 | -0.0168 | 1.14E-11 | IIIVmrMLM | SIG |
| PPR_18XX | AX-109009157 | 5B | 505564435 | 0.0182 | | 4.20E-08 | EMMAX | SIG |
| PPR_18XX | AX-109009157 | 5B | 505564435 | -0.0177 | | 5.35E-08 | GEMMA | SIG |
| PPR_18XX | AX-94764537 | 5B | 527058542 | -0.0007 | -0.0080 | 7.62E-11 | IIIVmrMLM | SIG |
| PPR_18XX | AX-109819320 | 5B | 556962611 | 0.0101 | | 1.16E-12 | BLUPmrMLM | SIG |
| PPR_18XX | AX-110577576 | 5B | 557582172 | -0.0054 | 0.0015 | 1.03E-17 | IIIVmrMLM | SIG |
| PPR_18XX | AX-108925664 | 5B | 558746971 | -0.0066 | -0.0123 | 1.07E-27 | IIIVmrMLM | SIG |
| PPR_18XX | AX-110593193 | 5B | 645519567 | -0.0049 | | 3.85E-17 | IIIVmrMLM | SIG |
| PPR_18XX | AX-108849241 | 5B | 672119507 | 0.0038 | -0.0004 | 5.41E-09 | IIIVmrMLM | SIG |
| PPR_18XX | AX-111832199 | 5D | 414530479 | -0.0043 | 0.0027 | 2.58E-12 | IIIVmrMLM | SIG |
| PPR_18XX | AX-110044398 | 5D | 556556558 | 0.0101 | | 1.05E-08 | BLUPmrMLM | SIG |
| PPR_18XX | AX-94666339 | 6A | 616913 | 0.0072 | | 5.58E-06 | BLUPmrMLM | SUG |

| | | | | | | | | |
|---|---|---|---|---|---|---|---|---|
| PPR_18XX | AX-109314545 | 6A | 183377839 | 0.0092 | | 6.79E-06 | BLUPmrMLM | SUG |
| PPR_18XX | AX-94800757 | 6A | 588604246 | -0.0007 | -0.0137 | 5.99E-06 | IIIVmrMLM | SUG |
| PPR_18XX | AX-94389933 | 6B | 3889628 | 0.0010 | -0.0203 | 1.19E-12 | IIIVmrMLM | SIG |
| PPR_18XX | AX-111697596 | 6B | 25936344 | 0.0051 | | 2.36E-06 | BLUPmrMLM | SUG |
| PPR_18XX | AX-110594254 | 6B | 69961231 | -0.0075 | | 1.73E-35 | IIIVmrMLM | SIG |
| PPR_18XX | AX-110496614 | 6B | 113765457 | -0.0009 | -0.0062 | 5.97E-10 | IIIVmrMLM | SIG |
| PPR_18XX | AX-110935567 | 6B | 141098799 | -0.0001 | 0.0086 | 3.73E-07 | IIIVmrMLM | SUG |
| PPR_18XX | AX-110520005 | 6B | 141099926 | -0.0067 | | 3.56E-06 | BLUPmrMLM | SUG |
| PPR_18XX | AX-108850595 | 6B | 572294975 | -0.0005 | -0.0190 | 2.60E-09 | IIIVmrMLM | SIG |
| PPR_18XX | AX-109983953 | 6B | 620793910 | -0.0011 | 0.0193 | 1.71E-07 | IIIVmrMLM | SUG |
| PPR_18XX | AX-109339400 | 6B | 630778734 | -0.0181 | | 1.52E-06 | EMMAX | SUG |
| PPR_18XX | AX-109339400 | 6B | 630778734 | 0.0173 | | 2.31E-06 | GEMMA | SUG |
| PPR_18XX | AX-108827730 | 6B | 700530538 | 0.0008 | 0.0123 | 9.71E-06 | IIIVmrMLM | SUG |
| PPR_18XX | AX-109053164 | 6B | 718364001 | -0.0006 | 0.0195 | 4.53E-07 | IIIVmrMLM | SUG |
| PPR_18XX | AX-108793603 | 6D | 54031416 | 0.0051 | | 1.29E-06 | BLUPmrMLM | SUG |
| PPR_18XX | AX-108793603 | 6D | 54031416 | -0.0039 | -0.0006 | 1.78E-10 | IIIVmrMLM | SIG |
| PPR_18XX | AX-110315493 | 6D | 179068242 | -0.0056 | | 2.55E-21 | IIIVmrMLM | SIG |
| PPR_18XX | AX-110314464 | 6D | 442937103 | -0.0040 | | 2.90E-12 | IIIVmrMLM | SIG |
| PPR_18XX | AX-111484978 | 7A | 532712286 | 0.0011 | -0.0331 | 4.03E-05 | IIIVmrMLM | SUG |
| PPR_18XX | AX-110434343 | 7A | 699798065 | -0.0044 | | 2.70E-05 | BLUPmrMLM | SUG |
| PPR_18XX | AX-109485882 | 7A | 719568648 | 0.0000 | 0.0111 | 3.21E-04 | IIIVmrMLM | SUG |
| PPR_18XX | AX-110626661 | 7B | 154812167 | 0.0066 | | 7.85E-06 | BLUPmrMLM | SUG |
| PPR_18XX | AX-109053711 | 7B | 546825731 | -0.0043 | | 1.13E-04 | BLUPmrMLM | SUG |
| PPR_18XX | AX-109949985 | 7B | 593399989 | -0.0049 | -0.0007 | 7.88E-13 | IIIVmrMLM | SIG |
| PPR_18XX | AX-108744845 | 7B | 643629221 | 0.0014 | 0.0131 | 9.58E-12 | IIIVmrMLM | SIG |
| PPR_18XX | AX-110939648 | 7B | 689857436 | -0.0043 | | 1.78E-05 | BLUPmrMLM | SUG |
| PPR_18XX | AX-110939648 | 7B | 689857436 | 0.0044 | -0.0034 | 1.87E-12 | IIIVmrMLM | SIG |
| PPR_18XX | AX-110673140 | 7D | 56212382 | 0.0021 | -0.0489 | 1.30E-20 | IIIVmrMLM | SIG |
| PPR_18XX | AX-110283607 | 7D | 546441919 | -0.0132 | -0.0049 | 2.38E-71 | IIIVmrMLM | SIG |
| PPR_18XX | AX-89389306 | 7D | 571274697 | -0.0057 | | 8.25E-08 | BLUPmrMLM | SIG |
| PPR_18ZMD | AX-109973169 | 1A | 522960363 | -0.0021 | 0.0785 | 1.60E-05 | IIIVmrMLM | SUG |
| PPR_18ZMD | AX-109960413 | 1A | 552166396 | 0.0389 | | 9.47E-07 | GEMMA | SUG |
| PPR_18ZMD | AX-110095531 | 1A | 552210497 | -0.0758 | | 2.32E-06 | EMMAX | SUG |
| PPR_18ZMD | AX-110095531 | 1A | 552210497 | 0.0760 | | 6.58E-07 | GEMMA | SUG |
| PPR_18ZMD | AX-111139210 | 1A | 567233776 | 0.0031 | 0.0168 | 3.31E-08 | IIIVmrMLM | SIG |
| PPR_18ZMD | AX-109750301 | 1D | 52710153 | 0.0040 | 0.1158 | 7.58E-09 | IIIVmrMLM | SIG |
| PPR_18ZMD | AX-108797247 | 2A | 475809387 | -0.0044 | -0.0170 | 4.56E-16 | IIIVmrMLM | SIG |
| PPR_18ZMD | AX-110006113 | 2A | 548325116 | -0.0095 | | 1.72E-05 | BLUPmrMLM | SUG |
| PPR_18ZMD | AX-110951550 | 2A | 578455046 | 0.0123 | 0.0088 | 9.17E-18 | IIIVmrMLM | SIG |
| PPR_18ZMD | AX-94545636 | 2A | 745408732 | 0.0114 | | 1.64E-05 | BLUPmrMLM | SUG |
| PPR_18ZMD | AX-109842599 | 2B | 4838565 | 0.0113 | | 1.46E-05 | BLUPmrMLM | SUG |
| PPR_18ZMD | AX-111459232 | 2B | 26020709 | 0.0022 | -0.0196 | 1.29E-08 | IIIVmrMLM | SIG |
| PPR_18ZMD | AX-111167392 | 2B | 236168397 | 0.0012 | -0.0510 | 3.08E-06 | IIIVmrMLM | SUG |
| PPR_18ZMD | AX-108988828 | 2D | 258796332 | 0.0262 | 0.0060 | 6.67E-22 | IIIVmrMLM | SIG |

| | | | | | | | | |
|---|---|---|---|---|---|---|---|---|
| PPR_18ZMD | AX-109506378 | 3A | 15784862 | -0.0026 | -0.0351 | 2.03E-11 | IIIVmrMLM | SIG |
| PPR_18ZMD | AX-108752799 | 3A | 36495647 | -0.0011 | -0.0328 | 4.95E-06 | IIIVmrMLM | SUG |
| PPR_18ZMD | AX-109940555 | 3A | 540216255 | 0.0074 | | 5.53E-05 | BLUPmrMLM | SUG |
| PPR_18ZMD | AX-110939635 | 3A | 646557815 | 0.0223 | 0.0091 | 1.71E-50 | IIIVmrMLM | SIG |
| PPR_18ZMD | AX-111148800 | 3A | 682574896 | -0.0141 | | 3.03E-06 | BLUPmrMLM | SUG |
| PPR_18ZMD | AX-108844766 | 3A | 722344584 | 0.0107 | 0.0053 | 2.71E-15 | IIIVmrMLM | SIG |
| PPR_18ZMD | AX-111154447 | 3A | 737599077 | -0.0084 | | 5.76E-05 | BLUPmrMLM | SUG |
| PPR_18ZMD | AX-111154447 | 3A | 737599077 | 0.0155 | 0.0044 | 1.21E-29 | IIIVmrMLM | SIG |
| PPR_18ZMD | AX-109378500 | 3A | 746319006 | -0.0070 | 0.0054 | 1.60E-06 | IIIVmrMLM | SUG |
| PPR_18ZMD | AX-111623897 | 3B | 60397394 | 0.0017 | -0.0289 | 5.57E-09 | IIIVmrMLM | SIG |
| PPR_18ZMD | AX-111487717 | 3B | 327848987 | 0.0062 | 0.0191 | 1.12E-10 | IIIVmrMLM | SIG |
| PPR_18ZMD | AX-110461838 | 3B | 760437784 | 0.0163 | | 9.83E-07 | FarmCPU | SUG |
| PPR_18ZMD | AX-95068735 | 3D | 570885824 | -0.0091 | | 1.31E-06 | BLUPmrMLM | SUG |
| PPR_18ZMD | AX-108827331 | 4A | 90506902 | -0.0052 | 0.0073 | 3.27E-05 | IIIVmrMLM | SUG |
| PPR_18ZMD | AX-109039692 | 4A | 544237195 | -0.0113 | | 2.86E-09 | BLUPmrMLM | SIG |
| PPR_18ZMD | AX-109466920 | 4B | 6142772 | 0.0156 | | 2.08E-06 | FarmCPU | SUG |
| PPR_18ZMD | AX-110099973 | 4B | 60423687 | 0.0016 | 0.0275 | 3.59E-09 | IIIVmrMLM | SIG |
| PPR_18ZMD | AX-109452428 | 4B | 101021017 | -0.0122 | | 5.42E-07 | BLUPmrMLM | SUG |
| PPR_18ZMD | AX-110385313 | 5A | 554190156 | -0.0066 | | 2.63E-07 | IIIVmrMLM | SUG |
| PPR_18ZMD | AX-108801363 | 5A | 555065862 | 0.0135 | | 1.87E-06 | FarmCPU | SUG |
| PPR_18ZMD | AX-109467309 | 5A | 562109645 | 0.0025 | 0.0188 | 1.84E-07 | IIIVmrMLM | SUG |
| PPR_18ZMD | AX-109313824 | 5B | 23684766 | 0.0111 | | 8.48E-05 | BLUPmrMLM | SUG |
| PPR_18ZMD | AX-89671930 | 5B | 45384652 | -0.0067 | | 7.50E-05 | BLUPmrMLM | SUG |
| PPR_18ZMD | AX-110049944 | 5B | 422126350 | -0.0003 | -0.0357 | 4.61E-15 | IIIVmrMLM | SIG |
| PPR_18ZMD | AX-111505028 | 5B | 609600665 | -0.0054 | -0.0239 | 4.82E-08 | IIIVmrMLM | SIG |
| PPR_18ZMD | AX-111464841 | 5B | 642325284 | 0.0060 | -0.0132 | 2.79E-06 | IIIVmrMLM | SUG |
| PPR_18ZMD | AX-109531765 | 5D | 32153643 | 0.0046 | 0.0333 | 2.27E-13 | IIIVmrMLM | SIG |
| PPR_18ZMD | AX-111582202 | 5D | 265753708 | 0.0736 | | 1.89E-06 | FarmCPU | SUG |
| PPR_18ZMD | AX-111907758 | 5D | 279839132 | -0.0213 | | 6.61E-08 | BLUPmrMLM | SIG |
| PPR_18ZMD | AX-95149766 | 5D | 302350886 | 0.0070 | | 4.95E-08 | IIIVmrMLM | SIG |
| PPR_18ZMD | AX-94388289 | 5D | 319222357 | -0.0114 | | 1.80E-05 | BLUPmrMLM | SUG |
| PPR_18ZMD | AX-95175066 | 5D | 494473072 | -0.0086 | | 5.49E-06 | BLUPmrMLM | SUG |
| PPR_18ZMD | AX-94702220 | 5D | 534865577 | -0.0182 | | 3.67E-40 | IIIVmrMLM | SIG |
| PPR_18ZMD | AX-94960903 | 6B | 8417892 | 0.0101 | | 3.97E-08 | BLUPmrMLM | SIG |
| PPR_18ZMD | AX-111588006 | 6B | 691636764 | 0.0124 | | 4.54E-08 | BLUPmrMLM | SIG |
| PPR_18ZMD | AX-110364023 | 6D | 2753518 | -0.0009 | -0.0124 | 4.85E-08 | IIIVmrMLM | SIG |
| PPR_18ZMD | AX-111027087 | 7A | 440359149 | 0.0034 | 0.0168 | 1.92E-05 | IIIVmrMLM | SUG |
| PPR_18ZMD | AX-109412056 | 7A | 676176608 | 0.0113 | | 7.96E-07 | BLUPmrMLM | SUG |
| PPR_18ZMD | AX-111014499 | 7A | 686241718 | -0.0068 | | 1.27E-04 | BLUPmrMLM | SUG |
| PPR_18ZMD | AX-109312726 | 7B | 589706480 | -0.0108 | | 1.92E-06 | BLUPmrMLM | SUG |
| PPR_18ZMD | AX-108899549 | 7B | 726880710 | 0.0088 | | 3.49E-06 | BLUPmrMLM | SUG |
| PPR_18ZMD | AX-95103080 | 7D | 208402 | -0.0055 | | 9.48E-05 | BLUPmrMLM | SUG |
| PPR_18ZMD | AX-95103080 | 7D | 208402 | 0.0068 | -0.0019 | 9.83E-06 | IIIVmrMLM | SUG |
| PPR_BLUP | AX-110493113 | 1A | 483034220 | 0.0005 | -0.0267 | 4.89E-17 | IIIVmrMLM | SIG |

| | | | | | | | | |
|---|---|---|---|---|---|---|---|---|
| PPR_BLUP | AX-108905435 | 1B | 57013700 | 0.0063 | -0.0005 | 9.43E-42 | IIIVmrMLM | SIG |
| PPR_BLUP | AX-109984183 | 1B | 304176354 | 0.0042 | | 9.92E-06 | BLUPmrMLM | SUG |
| PPR_BLUP | AX-109553065 | 1B | 530782054 | 0.0002 | -0.0069 | 9.25E-05 | IIIVmrMLM | SUG |
| PPR_BLUP | AX-94700454 | 1D | 7948235 | 0.0003 | 0.0045 | 4.17E-11 | IIIVmrMLM | SIG |
| PPR_BLUP | AX-111531363 | 1D | 24584257 | -0.0028 | -0.0068 | 2.61E-12 | IIIVmrMLM | SIG |
| PPR_BLUP | AX-109750301 | 1D | 52710153 | -0.0095 | 0.0299 | 5.95E-95 | IIIVmrMLM | SIG |
| PPR_BLUP | AX-109824680 | 2A | 43817622 | -0.0002 | -0.0074 | 9.91E-08 | IIIVmrMLM | SIG |
| PPR_BLUP | AX-111559860 | 2A | 407372832 | 0.0061 | | 1.41E-08 | BLUPmrMLM | SIG |
| PPR_BLUP | AX-111741582 | 2A | 414988352 | 0.0017 | 0.0117 | 1.10E-13 | IIIVmrMLM | SIG |
| PPR_BLUP | AX-94640748 | 2A | 475876556 | -0.0602 | | 1.72E-06 | EMMAX | SUG |
| PPR_BLUP | AX-110951550 | 2A | 578455046 | -0.0039 | | 2.49E-08 | BLUPmrMLM | SIG |
| PPR_BLUP | AX-110951550 | 2A | 578455046 | 0.0034 | 0.0010 | 2.20E-13 | IIIVmrMLM | SIG |
| PPR_BLUP | AX-108870652 | 2A | 617718562 | 0.0007 | 0.0109 | 4.48E-10 | IIIVmrMLM | SIG |
| PPR_BLUP | AX-111572089 | 2A | 751192255 | 0.0105 | | 1.39E-06 | FarmCPU | SUG |
| PPR_BLUP | AX-109474840 | 2A | 775756324 | 0.0092 | | 1.82E-07 | FarmCPU | SUG |
| PPR_BLUP | AX-110543864 | 2B | 4780929 | -0.0024 | 0.0026 | 8.43E-09 | IIIVmrMLM | SIG |
| PPR_BLUP | AX-111651927 | 2B | 4836307 | -0.0024 | 0.0026 | 8.43E-09 | IIIVmrMLM | SIG |
| PPR_BLUP | AX-110700714 | 2B | 10064484 | 0.0001 | 0.0116 | 1.73E-05 | IIIVmrMLM | SUG |
| PPR_BLUP | AX-109835973 | 2B | 11139880 | 0.0092 | | 1.90E-06 | GEMMA | SUG |
| PPR_BLUP | AX-110595438 | 2B | 546420600 | -0.0007 | 0.0271 | 1.67E-11 | IIIVmrMLM | SIG |
| PPR_BLUP | AX-110533196 | 2B | 595092535 | 0.0047 | | 3.34E-11 | BLUPmrMLM | SIG |
| PPR_BLUP | AX-109880475 | 2B | 746717847 | 0.0007 | -0.0104 | 1.86E-08 | IIIVmrMLM | SIG |
| PPR_BLUP | AX-111922990 | 2D | 633692091 | -0.0015 | -0.0153 | 4.56E-08 | IIIVmrMLM | SIG |
| PPR_BLUP | AX-94745115 | 2D | 646032417 | 0.0070 | 0.0164 | 2.46E-65 | IIIVmrMLM | SIG |
| PPR_BLUP | AX-111211824 | 2D | 647455478 | 0.0025 | | 1.55E-10 | IIIVmrMLM | SIG |
| PPR_BLUP | AX-110922437 | 3A | 771574 | 0.0028 | | 4.75E-06 | BLUPmrMLM | SUG |
| PPR_BLUP | AX-108752799 | 3A | 36495647 | -0.0005 | -0.0166 | 4.54E-14 | IIIVmrMLM | SIG |
| PPR_BLUP | AX-110459044 | 3A | 540045215 | 0.0044 | | 1.87E-05 | BLUPmrMLM | SUG |
| PPR_BLUP | AX-110461119 | 3A | 726457659 | -0.0043 | -0.0063 | 1.92E-26 | IIIVmrMLM | SIG |
| PPR_BLUP | AX-109351407 | 3A | 733963954 | 0.0030 | -0.0044 | 1.97E-13 | IIIVmrMLM | SIG |
| PPR_BLUP | AX-108802615 | 3B | 78575200 | -0.0057 | | 3.97E-08 | BLUPmrMLM | SIG |
| PPR_BLUP | AX-109417535 | 3B | 386234241 | -0.0150 | | 2.77E-10 | FarmCPU | SIG |
| PPR_BLUP | AX-111143552 | 3B | 588877197 | 0.0006 | 0.0163 | 9.79E-10 | IIIVmrMLM | SIG |
| PPR_BLUP | AX-110461838 | 3B | 760437784 | 0.0081 | | 2.59E-07 | FarmCPU | SUG |
| PPR_BLUP | AX-94438223 | 3D | 127335244 | 0.0024 | | 1.51E-09 | IIIVmrMLM | SIG |
| PPR_BLUP | AX-94926805 | 4A | 81817244 | -0.0015 | -0.0066 | 1.86E-12 | IIIVmrMLM | SIG |
| PPR_BLUP | AX-110033540 | 4A | 467299944 | 0.0040 | 0.0025 | 9.52E-22 | IIIVmrMLM | SIG |
| PPR_BLUP | AX-108825392 | 4A | 601380776 | 0.0017 | -0.0005 | 9.94E-05 | IIIVmrMLM | SUG |
| PPR_BLUP | AX-94577688 | 4B | 91957485 | 0.0028 | | 2.72E-07 | BLUPmrMLM | SUG |
| PPR_BLUP | AX-109395015 | 4B | 650737952 | 0.0052 | | 1.68E-08 | BLUPmrMLM | SIG |
| PPR_BLUP | AX-109838169 | 5A | 7803356 | 0.0047 | 0.0105 | 4.03E-40 | IIIVmrMLM | SIG |
| PPR_BLUP | AX-110594343 | 5A | 119571142 | 0.0002 | 0.0087 | 1.20E-10 | IIIVmrMLM | SIG |
| PPR_BLUP | AX-111002112 | 5A | 559783966 | -0.0003 | -0.0086 | 7.03E-13 | IIIVmrMLM | SIG |
| PPR_BLUP | AX-86170514 | 5A | 617236443 | -0.0036 | | 1.38E-08 | BLUPmrMLM | SIG |

| | | | | | | | | |
|---|---|---|---|---|---|---|---|---|
| PPR_BLUP | AX-111696799 | 5B | 32669779 | -0.0030 | | 1.79E-04 | BLUPmrMLM | SUG |
| PPR_BLUP | AX-109909648 | 5B | 34782236 | -0.0042 | -0.0001 | 1.14E-19 | IIIVmrMLM | SIG |
| PPR_BLUP | AX-109918369 | 5B | 220814297 | 0.0043 | 0.0142 | 8.37E-35 | IIIVmrMLM | SIG |
| PPR_BLUP | AX-111250458 | 5B | 225590461 | -0.0034 | -0.0012 | 6.53E-15 | IIIVmrMLM | SIG |
| PPR_BLUP | AX-110374631 | 5B | 400681209 | 0.0003 | -0.0190 | 2.47E-12 | IIIVmrMLM | SIG |
| PPR_BLUP | AX-111127545 | 5B | 565317118 | 0.0006 | 0.0086 | 1.15E-10 | IIIVmrMLM | SIG |
| PPR_BLUP | AX-108849241 | 5B | 672119507 | -0.0036 | | 5.30E-08 | BLUPmrMLM | SIG |
| PPR_BLUP | AX-109988793 | 5D | 556856383 | 0.0053 | | 4.62E-09 | BLUPmrMLM | SIG |
| PPR_BLUP | AX-110092474 | 6A | 611312714 | 0.0001 | -0.0079 | 2.72E-06 | IIIVmrMLM | SUG |
| PPR_BLUP | AX-109456453 | 6A | 612123939 | -0.0041 | | 9.14E-25 | IIIVmrMLM | SIG |
| PPR_BLUP | AX-109928626 | 6B | 667313 | 0.0058 | | 4.00E-11 | BLUPmrMLM | SIG |
| PPR_BLUP | AX-111144797 | 6B | 3623048 | 0.0008 | 0.0164 | 3.13E-07 | IIIVmrMLM | SUG |
| PPR_BLUP | AX-94389933 | 6B | 3889628 | 0.0019 | -0.0113 | 2.84E-13 | IIIVmrMLM | SIG |
| PPR_BLUP | AX-94960903 | 6B | 8417892 | -0.0022 | -0.0044 | 4.51E-08 | IIIVmrMLM | SIG |
| PPR_BLUP | AX-110599738 | 6B | 117802572 | 0.0002 | -0.0061 | 1.10E-11 | IIIVmrMLM | SIG |
| PPR_BLUP | AX-94602984 | 6B | 462170032 | 0.0070 | | 1.47E-07 | BLUPmrMLM | SUG |
| PPR_BLUP | AX-108894246 | 6B | 522002006 | -0.0052 | -0.0003 | 2.41E-32 | IIIVmrMLM | SIG |
| PPR_BLUP | AX-109604473 | 6B | 633119899 | 0.0029 | | 9.67E-06 | BLUPmrMLM | SUG |
| PPR_BLUP | AX-95115165 | 6D | 11059665 | 0.0084 | 0.0049 | 4.02E-49 | IIIVmrMLM | SIG |
| PPR_BLUP | AX-94420296 | 6D | 382589362 | -0.0045 | | 1.50E-05 | BLUPmrMLM | SUG |
| PPR_BLUP | AX-111056118 | 7A | 206690963 | 0.0059 | | 1.40E-08 | FarmCPU | SIG |
| PPR_BLUP | AX-111647572 | 7A | 598219776 | 0.0009 | 0.0082 | 4.00E-09 | IIIVmrMLM | SIG |
| PPR_BLUP | AX-109431727 | 7A | 679749940 | 0.0033 | | 2.01E-05 | BLUPmrMLM | SUG |
| PPR_BLUP | AX-109955950 | 7B | 50312986 | -0.0032 | | 2.93E-05 | BLUPmrMLM | SUG |
| PPR_BLUP | AX-111529876 | 7B | 594367483 | 0.0067 | | 1.03E-10 | BLUPmrMLM | SIG |
| PPR_BLUP | AX-111529876 | 7B | 594367483 | -0.0047 | 0.0005 | 8.98E-21 | IIIVmrMLM | SIG |
| PPR_BLUP | AX-95103080 | 7D | 208402 | -0.0033 | | 9.46E-08 | BLUPmrMLM | SIG |

PPR: ratio of peduncle length to plant height

**Supplementary Table S7.** All the known genes for three morphological traits in genome-wide association studies and their pieces of evidence.

| Trait | Genome-wide association study | | | | | | | Gene | Symbol | Reference |
|---|---|---|---|---|---|---|---|---|---|---|
| | Environment | Marker | Chr | Position | Add/Dom | P-value | Method | | | |
| PH | PH_18JY | AX-109372602 | 1D | 347285960 | 36.25/-- | 7.80E-08 | EMMAX | TraesCS1D01G254500 | *TaGID1-1D* | Li., et al 2013 |
| PH | PH_17JY | AX-109828660 | 2A | 413180560 | -2.0256/-- | 1.90E-06 | BLUPmrMLM | TraesCS2A01G261200 | *FUL2-2A* | Li et al., 2019 |
| PH | PH_18XX | AX-94488783 | 2A | 755788522 | -2.3172/-- | 5.36E-07 | BLUPmrMLM | TraesCS2A01G547800 | *TaARF12-2A* | Kong et al., 2023 |
| PH | PH_18JY | AX-110103130 | 2B | 767028694 | -1.4056/-- | 9.47E-06 | BLUPmrMLM | TraesCS2B01G578500 | *TaARF12-2B* | Kong et al., 2023 |
| PH | PH_17ZMD | AX-94834922 | 2D | 419030489 | -0.120808782 | 2.75E-07 | IIIVmrMLM | TraesCS2D01G324700 | *TaNAC100-2D* | Li et al., 2021 |
| PH | PH_17ZMD | AX-111585637 | 4A | 2960053 | 2.0786/-- | 8.86E-05 | BLUPmrMLM | TraesCS4A01G002600 | *SVP3-4A* | Li et al., 2021 |
| PH | PH_17ZMD | AX-111585637 | 4A | 2960053 | 2.0786/-- | 8.86E-05 | BLUPmrMLM | TraesCS4A01G005600 | *TaSRL1-4A* | Zhuang et al., 2021 |
| PH | PH_17AY | AX-108916749 | 4D | 19090268 | 6.3767/-- | 4.93E-08 | EMMAX | TraesCS4D01G040400 | *Rht-D1* | Peng et al., 1999 |
| PH | PH_17AY | AX-108916749 | 4D | 19090268 | -6.541293/-- | 6.94E-09 | GEMMA | TraesCS4D01G040400 | *Rht-D1* | Peng et al., 1999 |
| PH | PH_17AY | AX-108916749 | 4D | 19090268 | 4.3155/2.2266 | 3.67E-56 | IIIVmrMLM | TraesCS4D01G040400 | *Rht-D1* | Peng et al., 1999 |
| PH | PH_17JY | AX-108916749 | 4D | 19090268 | 6.4171/-- | 4.64E-07 | EMMAX | TraesCS4D01G040400 | *Rht-D1* | Peng et al., 1999 |
| PH | PH_17JY | AX-108916749 | 4D | 19090268 | -6.512045/-- | 1.23E-07 | GEMMA | TraesCS4D01G040400 | *Rht-D1* | Peng et al., 1999 |
| PH | PH_17ZMD | AX-108916749 | 4D | 19090268 | 7.09/-- | 1.74E-07 | EMMAX | TraesCS4D01G040400 | *Rht-D1* | Peng et al., 1999 |
| PH | PH_17ZMD | AX-108916749 | 4D | 19090268 | -7.174857/-- | 4.50E-08 | GEMMA | TraesCS4D01G040400 | *Rht-D1* | Peng et al., 1999 |
| PH | PH_18AY | AX-108916749 | 4D | 19090268 | 5.1936/-- | 3.59E-07 | EMMAX | TraesCS4D01G040400 | *Rht-D1* | Peng et al., 1999 |
| PH | PH_18AY | AX-108916749 | 4D | 19090268 | -5.332692/-- | 6.47E-08 | GEMMA | TraesCS4D01G040400 | *Rht-D1* | Peng et al., 1999 |
| PH | PH_18AY | AX-108916749 | 4D | 19090268 | 4.1613/1.3154 | 4.82E-68 | IIIVmrMLM | TraesCS4D01G040400 | *Rht-D1* | Peng et al., 1999 |
| PH | PH_18JY | AX-108916749 | 4D | 19090268 | 5.713/-- | 4.76E-07 | EMMAX | TraesCS4D01G040400 | *Rht-D1* | Peng et al., 1999 |
| PH | PH_18JY | AX-108916749 | 4D | 19090268 | -5.822133/-- | 1.50E-07 | GEMMA | TraesCS4D01G040400 | *Rht-D1* | Peng et al., 1999 |
| PH | PH_18XX | AX-108916749 | 4D | 19090268 | 5.7652/-- | 2.83E-11 | EMMAX | TraesCS4D01G040400 | *Rht-D1* | Peng et al., 1999 |
| PH | PH_18XX | AX-108916749 | 4D | 19090268 | -5.833156/-- | 5.30E-12 | GEMMA | TraesCS4D01G040400 | *Rht-D1* | Peng et al., 1999 |
| PH | PH_18ZMD | AX-108916749 | 4D | 19090268 | 6.6867/-- | 8.01E-08 | EMMAX | TraesCS4D01G040400 | *Rht-D1* | Peng et al., 1999 |

| | | | | | | | | | | |
|---|---|---|---|---|---|---|---|---|---|---|
| PH | PH_18ZMD | AX-108916749 | 4D | 19090268 | -6.792505/-- | 1.94E-08 | GEMMA | TraesCS4D01G040400 | *Rht-D1* | Peng et al., 1999 |
| PH | PH_BLUP | AX-108916749 | 4D | 19090268 | 6.0069/-- | 5.72E-12 | EMMAX | TraesCS4D01G040400 | *Rht-D1* | Peng et al., 1999 |
| PH | PH_BLUP | AX-108916749 | 4D | 19090268 | -6.104553/-- | 8.72E-13 | GEMMA | TraesCS4D01G040400 | *Rht-D1* | Peng et al., 1999 |
| PH | PH_17AY | AX-109281717 | 4D | 17613651 | 6.9439/-- | 2.78E-07 | EMMAX | TraesCS4D01G040400 | *Rht-D1* | Peng et al., 1999 |
| PH | PH_17AY | AX-109281717 | 4D | 17613651 | -7.178824/-- | 3.16E-08 | GEMMA | TraesCS4D01G040400 | *Rht-D1* | Peng et al., 1999 |
| PH | PH_17JY | AX-109281717 | 4D | 17613651 | 8.0243/-- | 5.52E-08 | EMMAX | TraesCS4D01G040400 | *Rht-D1* | Peng et al., 1999 |
| PH | PH_17JY | AX-109281717 | 4D | 17613651 | -8.171364/-- | 9.37E-09 | GEMMA | TraesCS4D01G040400 | *Rht-D1* | Peng et al., 1999 |
| PH | PH_17JY | AX-109281717 | 4D | 17613651 | 2.1481/-6.9929 | 1.22E-20 | IIIVmrMLM | TraesCS4D01G040400 | *Rht-D1* | Peng et al., 1999 |
| PH | PH_17ZMD | AX-109281717 | 4D | 17613651 | 7.976/-- | 5.14E-07 | EMMAX | TraesCS4D01G040400 | *Rht-D1* | Peng et al., 1999 |
| PH | PH_17ZMD | AX-109281717 | 4D | 17613651 | -8.170171/-- | 8.28E-08 | GEMMA | TraesCS4D01G040400 | *Rht-D1* | Peng et al., 1999 |
| PH | PH_18AY | AX-109281717 | 4D | 17613651 | 6.5464/-- | 4.23E-08 | EMMAX | TraesCS4D01G040400 | *Rht-D1* | Peng et al., 1999 |
| PH | PH_18AY | AX-109281717 | 4D | 17613651 | -6.767167/-- | 3.81E-09 | GEMMA | TraesCS4D01G040400 | *Rht-D1* | Peng et al., 1999 |
| PH | PH_18JY | AX-109281717 | 4D | 17613651 | 8.5397/-- | 2.93E-10 | EMMAX | TraesCS4D01G040400 | *Rht-D1* | Peng et al., 1999 |
| PH | PH_18JY | AX-109281717 | 4D | 17613651 | -8.744742/-- | 3.08E-11 | GEMMA | TraesCS4D01G040400 | *Rht-D1* | Peng et al., 1999 |
| PH | PH_18XX | AX-109281717 | 4D | 17613651 | 7.7063/-- | 7.02E-14 | EMMAX | TraesCS4D01G040400 | *Rht-D1* | Peng et al., 1999 |
| PH | PH_18XX | AX-109281717 | 4D | 17613651 | -7.845674/-- | 3.86E-15 | GEMMA | TraesCS4D01G040400 | *Rht-D1* | Peng et al., 1999 |
| PH | PH_18ZMD | AX-109281717 | 4D | 17613651 | 7.9126/-- | 3.23E-08 | EMMAX | TraesCS4D01G040400 | *Rht-D1* | Peng et al., 1999 |
| PH | PH_18ZMD | AX-109281717 | 4D | 17613651 | -8.085452/-- | 5.01E-09 | GEMMA | TraesCS4D01G040400 | *Rht-D1* | Peng et al., 1999 |
| PH | PH_BLUP | AX-109281717 | 4D | 17613651 | 7.4239/-- | 1.49E-12 | EMMAX | TraesCS4D01G040400 | *Rht-D1* | Peng et al., 1999 |
| PH | PH_BLUP | AX-109281717 | 4D | 17613651 | -7.620443/-- | 9.42E-14 | GEMMA | TraesCS4D01G040400 | *Rht-D1* | Peng et al., 1999 |
| PH | PH_17AY | AX-109665328 | 4D | 18539670 | 6.4829/-- | 1.16E-06 | EMMAX | TraesCS4D01G040400 | *Rht-D1* | Peng et al., 1999 |
| PH | PH_17AY | AX-109665328 | 4D | 18539670 | -6.768205/-- | 1.28E-07 | GEMMA | TraesCS4D01G040400 | *Rht-D1* | Peng et al., 1999 |
| PH | PH_17JY | AX-109665328 | 4D | 18539670 | 7.2417/-- | 6.38E-07 | EMMAX | TraesCS4D01G040400 | *Rht-D1* | Peng et al., 1999 |
| PH | PH_17JY | AX-109665328 | 4D | 18539670 | -7.493711/-- | 8.75E-08 | GEMMA | TraesCS4D01G040400 | *Rht-D1* | Peng et al., 1999 |
| PH | PH_17ZMD | AX-109665328 | 4D | 18539670 | 7.7125/-- | 8.10E-07 | EMMAX | TraesCS4D01G040400 | *Rht-D1* | Peng et al., 1999 |
| PH | PH_17ZMD | AX-109665328 | 4D | 18539670 | -7.968959/-- | 1.12E-07 | GEMMA | TraesCS4D01G040400 | *Rht-D1* | Peng et al., 1999 |

| | | | | | | | | | | |
|---|---|---|---|---|---|---|---|---|---|---|
| PH | PH_18AY | AX-109665328 | 4D | 18539670 | 6.0663/-- | 2.28E-07 | EMMAX | TraesCS4D01G040400 | *Rht-D1* | Peng et al., 1999 |
| PH | PH_18AY | AX-109665328 | 4D | 18539670 | -6.348341/-- | 2.00E-08 | GEMMA | TraesCS4D01G040400 | *Rht-D1* | Peng et al., 1999 |
| PH | PH_18JY | AX-109665328 | 4D | 18539670 | 8.0123/-- | 7.77E-10 | EMMAX | TraesCS4D01G040400 | *Rht-D1* | Peng et al., 1999 |
| PH | PH_18JY | AX-109665328 | 4D | 18539670 | -8.357627/-- | 6.02E-11 | GEMMA | TraesCS4D01G040400 | *Rht-D1* | Peng et al., 1999 |
| PH | PH_18XX | AX-109665328 | 4D | 18539670 | 7.7486/-- | 5.24E-15 | EMMAX | TraesCS4D01G040400 | *Rht-D1* | Peng et al., 1999 |
| PH | PH_18XX | AX-109665328 | 4D | 18539670 | -7.889161/-- | 3.52E-16 | GEMMA | TraesCS4D01G040400 | *Rht-D1* | Peng et al., 1999 |
| PH | PH_18XX | AX-109665328 | 4D | 18539670 | 3.9784/0.1471 | 9.11E-113 | IIIVmrMLM | TraesCS4D01G040400 | *Rht-D1* | Peng et al., 1999 |
| PH | PH_18ZMD | AX-109665328 | 4D | 18539670 | 7.7112/-- | 3.93E-08 | EMMAX | TraesCS4D01G040400 | *Rht-D1* | Peng et al., 1999 |
| PH | PH_18ZMD | AX-109665328 | 4D | 18539670 | -7.911671/-- | 5.84E-09 | GEMMA | TraesCS4D01G040400 | *Rht-D1* | Peng et al., 1999 |
| PH | PH_BLUP | AX-109665328 | 4D | 18539670 | 6.5216/-- | 8.29E-11 | EMMAX | TraesCS4D01G040400 | *Rht-D1* | Peng et al., 1999 |
| PH | PH_BLUP | AX-109665328 | 4D | 18539670 | -6.915289/-- | 3.28E-12 | GEMMA | TraesCS4D01G040400 | *Rht-D1* | Peng et al., 1999 |
| PH | PH_18JY | AX-110387369 | 4D | 467431300 | -41.566/-- | 7.68E-07 | EMMAX | TraesCS4D01G298400 | *TaSRL1-4D* | Zhuang et al., 2021 |
| PH | PH_18JY | AX-110387369 | 4D | 467431300 | -41.566/-- | 7.68E-07 | EMMAX | TraesCS4D01G301100 | *SVP3-4D* | Li et al., 2021 |
| PH | PH_18XX | AX-110387369 | 4D | 467431300 | -33.542/-- | 2.55E-07 | EMMAX | TraesCS4D01G298400 | *TaSRL1-4D* | Zhuang et al., 2021 |
| PH | PH_18XX | AX-110387369 | 4D | 467431300 | -33.542/-- | 2.55E-07 | EMMAX | TraesCS4D01G301100 | *SVP3-4D* | Li et al., 2021 |
| PH | PH_BLUP | AX-110387369 | 4D | 467431300 | -37.96/-- | 5.44E-09 | EMMAX | TraesCS4D01G298400 | *TaSRL1-4D* | Zhuang et al., 2021 |
| PH | PH_BLUP | AX-110387369 | 4D | 467431300 | -37.96/-- | 5.44E-09 | EMMAX | TraesCS4D01G301100 | *SVP3-4D* | Li et al., 2021 |
| PH | PH_17AY | AX-89421921 | 4D | 19288898 | 6.8314/-- | 3.66E-07 | EMMAX | TraesCS4D01G040400 | *Rht-D1* | Peng et al., 1999 |
| PH | PH_17AY | AX-89421921 | 4D | 19288898 | -7.064118/-- | 4.35E-08 | GEMMA | TraesCS4D01G040400 | *Rht-D1* | Peng et al., 1999 |
| PH | PH_17JY | AX-89421921 | 4D | 19288898 | 7.9995/-- | 4.98E-08 | EMMAX | TraesCS4D01G040400 | *Rht-D1* | Peng et al., 1999 |
| PH | PH_17JY | AX-89421921 | 4D | 19288898 | -8.133686/-- | 9.01E-09 | GEMMA | TraesCS4D01G040400 | *Rht-D1* | Peng et al., 1999 |
| PH | PH_17ZMD | AX-89421921 | 4D | 19288898 | 8.112/-- | 2.99E-07 | EMMAX | TraesCS4D01G040400 | *Rht-D1* | Peng et al., 1999 |
| PH | PH_17ZMD | AX-89421921 | 4D | 19288898 | -8.291709/-- | 4.76E-08 | GEMMA | TraesCS4D01G040400 | *Rht-D1* | Peng et al., 1999 |
| PH | PH_17ZMD | AX-89421921 | 4D | 19288898 | 2.9544/-7.4757 | 2.48E-28 | IIIVmrMLM | TraesCS4D01G040400 | *Rht-D1* | Peng et al., 1999 |
| PH | PH_18AY | AX-89421921 | 4D | 19288898 | 6.5285/-- | 3.76E-08 | EMMAX | TraesCS4D01G040400 | *Rht-D1* | Peng et al., 1999 |
| PH | PH_18AY | AX-89421921 | 4D | 19288898 | -6.751967/-- | 3.30E-09 | GEMMA | TraesCS4D01G040400 | *Rht-D1* | Peng et al., 1999 |

| | | | | | | | | | | |
|---|---|---|---|---|---|---|---|---|---|---|
| PH | PH_18JY | AX-89421921 | 4D | 19288898 | 8.4953/-- | 2.79E-10 | EMMAX | TraesCS4D01G040400 | *Rht-D1* | Peng et al., 1999 |
| PH | PH_18JY | AX-89421921 | 4D | 19288898 | -8.725351/-- | 2.57E-11 | GEMMA | TraesCS4D01G040400 | *Rht-D1* | Peng et al., 1999 |
| PH | PH_18JY | AX-89421921 | 4D | 19288898 | 4.2858/-4.9486 | 3.11E-82 | IIIVmrMLM | TraesCS4D01G040400 | *Rht-D1* | Peng et al., 1999 |
| PH | PH_18XX | AX-89421921 | 4D | 19288898 | 7.7135/-- | 4.47E-14 | EMMAX | TraesCS4D01G040400 | *Rht-D1* | Peng et al., 1999 |
| PH | PH_18XX | AX-89421921 | 4D | 19288898 | -7.852742/-- | 2.39E-15 | GEMMA | TraesCS4D01G040400 | *Rht-D1* | Peng et al., 1999 |
| PH | PH_18ZMD | AX-89421921 | 4D | 19288898 | 7.7584/-- | 4.92E-08 | EMMAX | TraesCS4D01G040400 | *Rht-D1* | Peng et al., 1999 |
| PH | PH_18ZMD | AX-89421921 | 4D | 19288898 | -7.926403/-- | 8.12E-09 | GEMMA | TraesCS4D01G040400 | *Rht-D1* | Peng et al., 1999 |
| PH | PH_BLUP | AX-89421921 | 4D | 19288898 | 7.3929/-- | 1.31E-12 | EMMAX | TraesCS4D01G040400 | *Rht-D1* | Peng et al., 1999 |
| PH | PH_BLUP | AX-89421921 | 4D | 19288898 | -7.584804/-- | 8.66E-14 | GEMMA | TraesCS4D01G040400 | *Rht-D1* | Peng et al., 1999 |
| PH | PH_BLUP | AX-89421921 | 4D | 19288898 | 5.4539/1.1601 | 1.05E-185 | IIIVmrMLM | TraesCS4D01G040400 | *Rht-D1* | Peng et al., 1999 |
| PH | PH_17JY | AX-89445201 | 4D | 19459579 | -6.221432/-- | 1.20E-06 | GEMMA | TraesCS4D01G040400 | *Rht-D1* | Peng et al., 1999 |
| PH | PH_17ZMD | AX-89445201 | 4D | 19459579 | 6.9371/-- | 7.12E-07 | EMMAX | TraesCS4D01G040400 | *Rht-D1* | Peng et al., 1999 |
| PH | PH_17ZMD | AX-89445201 | 4D | 19459579 | -7.150951/-- | 1.12E-07 | GEMMA | TraesCS4D01G040400 | *Rht-D1* | Peng et al., 1999 |
| PH | PH_18AY | AX-89445201 | 4D | 19459579 | 5.0132/-- | 1.99E-06 | EMMAX | TraesCS4D01G040400 | *Rht-D1* | Peng et al., 1999 |
| PH | PH_18AY | AX-89445201 | 4D | 19459579 | -5.218602/-- | 2.96E-07 | GEMMA | TraesCS4D01G040400 | *Rht-D1* | Peng et al., 1999 |
| PH | PH_18JY | AX-89445201 | 4D | 19459579 | 5.7656/-- | 7.55E-07 | EMMAX | TraesCS4D01G040400 | *Rht-D1* | Peng et al., 1999 |
| PH | PH_18JY | AX-89445201 | 4D | 19459579 | -5.97794/-- | 1.43E-07 | GEMMA | TraesCS4D01G040400 | *Rht-D1* | Peng et al., 1999 |
| PH | PH_18XX | AX-89445201 | 4D | 19459579 | 4.8527/-- | 2.83E-08 | EMMAX | TraesCS4D01G040400 | *Rht-D1* | Peng et al., 1999 |
| PH | PH_18XX | AX-89445201 | 4D | 19459579 | -5.095674/-- | 2.72E-09 | GEMMA | TraesCS4D01G040400 | *Rht-D1* | Peng et al., 1999 |
| PH | PH_18ZMD | AX-89445201 | 4D | 19459579 | 6.6704/-- | 1.39E-07 | EMMAX | TraesCS4D01G040400 | *Rht-D1* | Peng et al., 1999 |
| PH | PH_18ZMD | AX-89445201 | 4D | 19459579 | -6.853899/-- | 2.30E-08 | GEMMA | TraesCS4D01G040400 | *Rht-D1* | Peng et al., 1999 |
| PH | PH_18ZMD | AX-89445201 | 4D | 19459579 | 2.4954/-2.2945 | 3.65E-22 | IIIVmrMLM | TraesCS4D01G040400 | *Rht-D1* | Peng et al., 1999 |
| PH | PH_BLUP | AX-89445201 | 4D | 19459579 | -3.4208/-- | 4.12E-15 | BLUPmrMLM | TraesCS4D01G040400 | *Rht-D1* | Peng et al., 1999 |
| PH | PH_BLUP | AX-89445201 | 4D | 19459579 | 4.6051/-- | 1.41E-07 | EMMAX | TraesCS4D01G040400 | *Rht-D1* | Peng et al., 1999 |
| PH | PH_BLUP | AX-89445201 | 4D | 19459579 | -4.878713/-- | 1.65E-08 | GEMMA | TraesCS4D01G040400 | *Rht-D1* | Peng et al., 1999 |
| PH | PH_17AY | AX-89703298 | 4D | 16926596 | 6.7316/-- | 4.84E-07 | EMMAX | TraesCS4D01G040400 | *Rht-D1* | Peng et al., 1999 |

| | | | | | | | | | | |
|---|---|---|---|---|---|---|---|---|---|---|
| PH | PH_17AY | AX-89703298 | 4D | 16926596 | -6.943309/-- | 6.73E-08 | GEMMA | TraesCS4D01G040400 | *Rht-D1* | Peng et al., 1999 |
| PH | PH_17JY | AX-89703298 | 4D | 16926596 | 7.7969/-- | 1.14E-07 | EMMAX | TraesCS4D01G040400 | *Rht-D1* | Peng et al., 1999 |
| PH | PH_17JY | AX-89703298 | 4D | 16926596 | -7.947411/-- | 2.06E-08 | GEMMA | TraesCS4D01G040400 | *Rht-D1* | Peng et al., 1999 |
| PH | PH_17ZMD | AX-89703298 | 4D | 16926596 | 7.8128/-- | 6.42E-07 | EMMAX | TraesCS4D01G040400 | *Rht-D1* | Peng et al., 1999 |
| PH | PH_17ZMD | AX-89703298 | 4D | 16926596 | -7.981668/-- | 1.20E-07 | GEMMA | TraesCS4D01G040400 | *Rht-D1* | Peng et al., 1999 |
| PH | PH_18AY | AX-89703298 | 4D | 16926596 | 6.3018/-- | 9.17E-08 | EMMAX | TraesCS4D01G040400 | *Rht-D1* | Peng et al., 1999 |
| PH | PH_18AY | AX-89703298 | 4D | 16926596 | -6.529017/-- | 9.02E-09 | GEMMA | TraesCS4D01G040400 | *Rht-D1* | Peng et al., 1999 |
| PH | PH_18JY | AX-89703298 | 4D | 16926596 | -6.0009/-- | 1.66E-15 | BLUPmrMLM | TraesCS4D01G040400 | *Rht-D1* | Peng et al., 1999 |
| PH | PH_18JY | AX-89703298 | 4D | 16926596 | 8.0983/-- | 1.12E-09 | EMMAX | TraesCS4D01G040400 | *Rht-D1* | Peng et al., 1999 |
| PH | PH_18JY | AX-89703298 | 4D | 16926596 | -8.353808/-- | 1.12E-10 | GEMMA | TraesCS4D01G040400 | *Rht-D1* | Peng et al., 1999 |
| PH | PH_18XX | AX-89703298 | 4D | 16926596 | 7.3864/-- | 2.85E-13 | EMMAX | TraesCS4D01G040400 | *Rht-D1* | Peng et al., 1999 |
| PH | PH_18XX | AX-89703298 | 4D | 16926596 | -7.53979/-- | 1.79E-14 | GEMMA | TraesCS4D01G040400 | *Rht-D1* | Peng et al., 1999 |
| PH | PH_18ZMD | AX-89703298 | 4D | 16926596 | 7.5937/-- | 7.89E-08 | EMMAX | TraesCS4D01G040400 | *Rht-D1* | Peng et al., 1999 |
| PH | PH_18ZMD | AX-89703298 | 4D | 16926596 | -7.758605/-- | 1.41E-08 | GEMMA | TraesCS4D01G040400 | *Rht-D1* | Peng et al., 1999 |
| PH | PH_BLUP | AX-89703298 | 4D | 16926596 | 7.0787/-- | 5.71E-12 | EMMAX | TraesCS4D01G040400 | *Rht-D1* | Peng et al., 1999 |
| PH | PH_BLUP | AX-89703298 | 4D | 16926596 | -7.271137/-- | 4.67E-13 | GEMMA | TraesCS4D01G040400 | *Rht-D1* | Peng et al., 1999 |
| PH | PH_BLUP | AX-108952168 | 5A | 698619167 | 0.07636447 | 1.28E-10 | IIIVmrMLM | TraesCS5A01G543100 | *Rht12* | Sun et al., 2019 |
| PH | PH_18JY | AX-110014427 | 5A | 697488076 | 0.784/-7.7191 | 9.76E-17 | IIIVmrMLM | TraesCS5A01G543100 | *Rht12* | Sun et al., 2019 |
| PH | PH_BLUP | AX-110714077 | 5A | 697053429 | -2.895/-- | 2.62E-12 | BLUPmrMLM | TraesCS5A01G543100 | *Rht12* | Sun et al., 2019 |
| PH | PH_17AY | AX-109567209 | 5D | 106520306 | -1.7201/-- | 2.86E-12 | IIIVmrMLM | TraesCS5D01G097400 | *TaWOX2-5D* | Si et al., 2021 |
| SL | SL_18XX | AX-110423885 | 2B | 766172990 | -0.1072/-- | 1.53E-04 | BLUPmrMLM | TraesCS2B01G578500 | *TaARF12-2B* | Kong et al., 2023 |
| SL | SL_17AY | AX-110460935 | 2B | 767897927 | 0.0028/0.8118 | 9.10E-06 | IIIVmrMLM | TraesCS2B01G578500 | *TaARF12-2B* | Kong et al., 2023 |
| SL | SL_17JY | AX-110726653 | 6A | 549032014 | 0.049/0.6741 | 5.45E-05 | IIIVmrMLM | TraesCS6A01G313800 | *SVP1-6A* | Li et al., 2021 |
| SL | SL_18JY | AX-110643450 | 7A | 558399409 | 0.0207/0.9229 | 1.05E-07 | IIIVmrMLM | TraesCS7A01G383800 | *AP3-1-7A* | Hama et al., 2004 |
| PPR | PPR_18JY | AX-109477982 | 2A | 755120274 | 0.0042/0.0011 | 4.44E-10 | IIIVmrMLM | TraesCS2A01G547800 | *TaARF12-2A* | Kong et al., 2023 |
| PPR | PPR_18XX | AX-111597275 | 2B | 41414402 | 0.0012/0.0432 | 1.65E-11 | IIIVmrMLM | TraesCS2B01G073600 | *Rht8* | Chai et al., 2022 |

| | | | | | | | | | | |
|---|---|---|---|---|---|---|---|---|---|---|
| PPR | PPR_18JY | AX-111030610 | 5D | 106818950 | -0.0043/-- | 4.89E-05 | BLUPmrMLM | TraesCS5D01G097400 | *TaWOX2-5D* | Si et al., 2021 |

PH: plant height, SL: spike length, PPR: ratio of peduncle length to plant height.

**Supplementary Table S8.** Candidate genes for three morphological traits in genome-wide association studies and their pieces of evidence

| Trait | Genome-wide association studies | | | | | | | Differential expression analysis | | | Homologous genes/Genes reported in Rice/wheat | GO annotation analysis | | P-value in haplotype analysis | Reference |
|---|---|---|---|---|---|---|---|---|---|---|---|---|---|---|---|
| | **Environment** | **Marker** | **Chr** | **Position** | **Add/Dom** | **P-value** | **Method** | **Gene** | **Log$_2$\|FC\|** | **P. adj** | | **ID** | **Annotation** | | |
| PH | PH_18ZMD | AX-108742536 | 1A | 389029091 | 11.243/-- | 1.30E-06 | EMMAX | TraesCS1A01G218800 | -1.54$^{PH3}$; -1.97$^{PH2}$ | 2.69E-08$^{PH3}$; 1.61E-04$^{PH2}$ | | GO:0009739 | response to gibberellin | 8.37E-05 | |
| PH | PH_17ZMD | AX-108742536 | 1A | 389029091 | -11.46766/-- | 2.08E-06 | GEMMA | TraesCS1A01G218800 | -1.54$^{PH3}$; -1.97$^{PH2}$ | 2.69E-08$^{PH3}$; 1.61E-04$^{PH2}$ | | GO:0009739 | response to gibberellin | 7.06E-04 | |
| PH | PH_18ZMD | AX-108742536 | 1A | 389029091 | -10.87423/-- | 8.70E-07 | GEMMA | TraesCS1A01G218800 | -1.54$^{PH3}$; -1.97$^{PH2}$ | 2.69E-08$^{PH3}$; 1.61E-04$^{PH2}$ | | GO:0009739 | response to gibberellin | 8.37E-05 | |
| PH | PH_18ZMD | AX-108840039 | 1A | 387936105 | 10.052/-- | 1.27E-06 | EMMAX | TraesCS1A01G218800 | -1.54$^{PH3}$; -1.97$^{PH2}$ | 2.69E-08$^{PH3}$; 1.61E-04$^{PH2}$ | | GO:0009739 | response to gibberellin | 8.37E-05 | |
| PH | PH_18ZMD | AX-108840039 | 1A | 387936105 | -10.34022/-- | 2.89E-07 | GEMMA | TraesCS1A01G218800 | -1.54$^{PH3}$; -1.97$^{PH2}$ | 2.69E-08$^{PH3}$; 1.61E-04$^{PH2}$ | | GO:0009739 | response to gibberellin | 8.37E-05 | |
| PH | PH_18ZMD | AX-108850470 | 1A | 389228613 | 10.978/-- | 2.26E-06 | EMMAX | TraesCS1A01G218800 | -1.54$^{PH3}$; -1.97$^{PH2}$ | 2.69E-08$^{PH3}$; 1.61E-04$^{PH2}$ | | GO:0009739 | response to gibberellin | 8.37E-05 | |
| PH | PH_18ZMD | AX-108850470 | 1A | 389228613 | -10.61021/-- | 1.61E-06 | GEMMA | TraesCS1A01G218800 | -1.54$^{PH3}$; -1.97$^{PH2}$ | 2.69E-08$^{PH3}$; 1.61E-04$^{PH2}$ | | GO:0009739 | response to gibberellin | 8.37E-05 | |
| PH | PH_17ZMD | AX-108850470 | 1A | 389228613 | -11.46067/-- | 2.11E-06 | GEMMA | TraesCS1A01G218800 | -1.54$^{PH3}$; -1.97$^{PH2}$ | 2.69E-08$^{PH3}$; 1.61E-04$^{PH2}$ | | GO:0009739 | response to gibberellin | 7.06E-04 | |
| PH | PH_18XX | AX-109902759 | 1A | 388463098 | 1.8797/-1.6669 | 3.11E-35 | IIIVmrMLM | TraesCS1A01G218800 | -1.54$^{PH3}$; -1.97$^{PH2}$ | 2.69E-08$^{PH3}$; 1.61E-04$^{PH2}$ | | GO:0009739 | response to gibberellin | 3.75E-05 | |
| PH | PH_18ZMD | AX-109379218 | 1A | 388947077 | -10.87423/-- | 8.70E-07 | GEMMA | TraesCS1A01G218800 | -1.54$^{PH3}$; -1.97$^{PH2}$ | 2.69E-08$^{PH3}$; 1.61E-04$^{PH2}$ | | GO:0009739 | response to gibberellin | 8.37E-05 | |
| PH | PH_17ZMD | AX-109379218 | 1A | 388947077 | -11.46766/-- | 2.08E-06 | GEMMA | TraesCS1A01G218800 | -1.54$^{PH3}$; -1.97$^{PH2}$ | 2.69E-08$^{PH3}$; 1.61E-04$^{PH2}$ | | GO:0009739 | response to gibberellin | 7.06E-04 | |
| PH | PH_18ZMD | AX-109379218 | 1A | 388947077 | 11.243/-- | 1.30E-06 | EMMAX | TraesCS1A01G218800 | -1.54$^{PH3}$; -1.97$^{PH2}$ | 2.69E-08$^{PH3}$; 1.61E-04$^{PH2}$ | | GO:0009739 | response to gibberellin | 8.37E-05 | |
| PH | PH_18ZMD | AX-111819445 | 1A | 388947113 | -10.87423/-- | 8.70E-07 | GEMMA | TraesCS1A01G218800 | -1.54$^{PH3}$; -1.97$^{PH2}$ | 2.69E-08$^{PH3}$; 1.61E-04$^{PH2}$ | | GO:0009739 | response to gibberellin | 8.37E-05 | |
| PH | PH_18ZMD | AX-110492346 | 1A | 389252927 | -10.61021/-- | 1.61E-06 | GEMMA | TraesCS1A01G218800 | -1.54$^{PH3}$; -1.97$^{PH2}$ | 2.69E-08$^{PH3}$; 1.61E-04$^{PH2}$ | | GO:0009739 | response to gibberellin | 8.37E-05 | |
| PH | PH_17ZMD | AX-110492346 | 1A | 389252927 | -11.46067/-- | 2.11E-06 | GEMMA | TraesCS1A01G218800 | -1.54$^{PH3}$; -1.97$^{PH2}$ | 2.69E-08$^{PH3}$; 1.61E-04$^{PH2}$ | | GO:0009739 | response to gibberellin | 7.06E-04 | |
| PH | PH_18ZMD | AX-110492346 | 1A | 389252927 | 10.978/-- | 2.26E-06 | EMMAX | TraesCS1A01G218800 | -1.54$^{PH3}$; -1.97$^{PH2}$ | 2.69E-08$^{PH3}$; 1.61E-04$^{PH2}$ | | GO:0009739 | response to gibberellin | 8.37E-05 | |
| PH | PH_18ZMD | AX-110403537 | 1A | 389061020 | 11.243/-- | 1.30E-06 | EMMAX | TraesCS1A01G218800 | -1.54$^{PH3}$; -1.97$^{PH2}$ | 2.69E-08$^{PH3}$; 1.61E-04$^{PH2}$ | | GO:0009739 | response to gibberellin | 8.37E-05 | |
| PH | PH_18ZMD | AX-110403537 | 1A | 389061020 | -10.87423/-- | 8.70E-07 | GEMMA | TraesCS1A01G218800 | -1.54$^{PH3}$; -1.97$^{PH2}$ | 2.69E-08$^{PH3}$; 1.61E-04$^{PH2}$ | | GO:0009739 | response to gibberellin | 8.37E-05 | |
| PH | PH_17ZMD | AX-110403537 | 1A | 389061020 | -11.46766/-- | 2.08E-06 | GEMMA | TraesCS1A01G218800 | -1.54$^{PH3}$; -1.97$^{PH2}$ | 2.69E-08$^{PH3}$; 1.61E-04$^{PH2}$ | | GO:0009739 | response to gibberellin | 7.06E-04 | |
| PH | PH_17ZMD | AX-109509868 | 1A | 389363391 | -11.46067/-- | 2.11E-06 | GEMMA | TraesCS1A01G218800 | -1.54$^{PH3}$; -1.97$^{PH2}$ | 2.69E-08$^{PH3}$; 1.61E-04$^{PH2}$ | | GO:0009739 | response to gibberellin | 7.06E-04 | |
| PH | PH_18ZMD | AX-109509868 | 1A | 389363391 | 10.978/-- | 2.26E-06 | EMMAX | TraesCS1A01G218800 | -1.54$^{PH3}$; -1.97$^{PH2}$ | 2.69E-08$^{PH3}$; 1.61E-04$^{PH2}$ | | GO:0009739 | response to gibberellin | 8.37E-05 | |

| | | | | | | | | | | | | | |
|---|---|---|---|---|---|---|---|---|---|---|---|---|---|
| PH | PH_18ZMD | AX-109509868 | 1A | 389363391 | -10.61021/-- | 1.61E-06 | GEMMA | TraesCS1A01G218800 | -1.54 [PH3]; -1.97 [PH2] | 2.69E-08 [PH3]; 1.61E-04 [PH2] | GO:0009739 | response to gibberellin | 8.37E-05 |
| PH | PH_17ZMD | AX-110448433 | 1A | 389061946 | -22.92134/-- | 2.11E-06 | GEMMA | TraesCS1A01G218800 | -1.54 [PH3]; -1.97 [PH2] | 2.69E-08 [PH3]; 1.61E-04 [PH2] | GO:0009739 | response to gibberellin | 7.06E-04 |
| PH | PH_18ZMD | AX-110448433 | 1A | 389061946 | -21.22042/-- | 1.61E-06 | GEMMA | TraesCS1A01G218800 | -1.54 [PH3]; -1.97 [PH2] | 2.69E-08 [PH3]; 1.61E-04 [PH2] | GO:0009739 | response to gibberellin | 8.37E-05 |
| PH | PH_17ZMD | AX-111106333 | 1A | 389200944 | -11.28217/-- | 1.19E-06 | GEMMA | TraesCS1A01G218800 | -1.54 [PH3]; -1.97 [PH2] | 2.69E-08 [PH3]; 1.61E-04 [PH2] | GO:0009739 | response to gibberellin | 7.06E-04 |
| PH | PH_17ZMD | AX-111819445 | 1A | 388947113 | -11.46766/-- | 2.08E-06 | GEMMA | TraesCS1A01G218800 | -1.54 [PH3]; -1.97 [PH2] | 2.69E-08 [PH3]; 1.61E-04 [PH2] | GO:0009739 | response to gibberellin | 7.06E-04 |
| PH | PH_18ZMD | AX-111819445 | 1A | 388947113 | 11.243/-- | 1.30E-06 | EMMAX | TraesCS1A01G218800 | -1.54 [PH3]; -1.97 [PH2] | 2.69E-08 [PH3]; 1.61E-04 [PH2] | GO:0009739 | response to gibberellin | 8.37E-05 |
| PH | PH_18ZMD | AX-111668333 | 1A | 388937021 | -10.42218/-- | 2.20E-06 | GEMMA | TraesCS1A01G218800 | -1.54 [PH3]; -1.97 [PH2] | 2.69E-08 [PH3]; 1.61E-04 [PH2] | GO:0009739 | response to gibberellin | 8.37E-05 |
| PH | PH_18ZMD | AX-110448433 | 1A | 389061946 | 21.956/-- | 2.26E-06 | EMMAX | TraesCS1A01G218800 | -1.54 [PH3]; -1.97 [PH2] | 2.69E-08 [PH3]; 1.61E-04 [PH2] | GO:0009739 | response to gibberellin | 8.37E-05 |
| PH | PH_17ZMD | AX-111106333 | 1A | 389200944 | 11.542/-- | 2.30E-06 | EMMAX | TraesCS1A01G218800 | -1.54 [PH3]; -1.97 [PH2] | 2.69E-08 [PH3]; 1.61E-04 [PH2] | GO:0009739 | response to gibberellin | 7.06E-04 |
| PH | PH_18ZMD | AX-111106333 | 1A | 389200944 | -10.05872/-- | 2.05E-06 | GEMMA | TraesCS1A01G218800 | -1.54 [PH3]; -1.97 [PH2] | 2.69E-08 [PH3]; 1.61E-04 [PH2] | GO:0009739 | response to gibberellin | 8.37E-05 |
| PH | PH_18ZMD | AX-110554073 | 1A | 460637199 | -11.14624815/-- | 3.49E-07 | FarmCPU | TraesCS1A01G263600 | -1.92 [PH3] | 4.32E-03 [PH3] | GO:0010476 | gibberellin mediated signaling pathway | 3.19E-138 |
| PH | PH_18ZMD | AX-110554073 | 1A | 460637199 | -11.14624815/-- | 3.49E-07 | FarmCPU | TraesCS1A01G263600 | -1.92 [PH3] | 4.32E-03 [PH3] | GO:0009685 | gibberellin metabolic process | 3.19E-138 |
| PH | PH_18ZMD | AX-110554073 | 1A | 460637199 | -11.14624815/-- | 3.49E-07 | FarmCPU | TraesCS1A01G263600 | -1.92 [PH3] | 4.32E-03 [PH3] | GO:0009739 | response to gibberellin | 3.19E-138 |
| PH | PH_18ZMD | AX-110554073 | 1A | 460637199 | -11.14624815/-- | 3.49E-07 | FarmCPU | TraesCS1A01G263600 | -1.92 [PH3] | 4.32E-03 [PH3] | GO:0009686 | gibberellin biosynthetic process | 3.19E-138 |
| PH | PH_18ZMD | AX-110554073 | 1A | 460637199 | -11.14624815/-- | 3.49E-07 | FarmCPU | TraesCS1A01G263600 | -1.92 [PH3] | 4.32E-03 [PH3] | GO:0071370 | cellular response to gibberellin stimulus | 3.19E-138 |
| PH | PH_18JY | AX-111157582 | 1B | 672880906 | 1.8579/-- | 1.81E-04 | BLUPmrMLM | TraesCS1B01G462300 | 4.21 [PH2] | 4.33E-06 [PH2] | GO:0071367 | cellular response to brassinosteroid stimulus | 8.73E-15 |
| PH | PH_18JY | AX-111157582 | 1B | 672880906 | 1.8579/-- | 1.81E-04 | BLUPmrMLM | TraesCS1B01G462300 | 4.21 [PH2] | 4.33E-06 [PH2] | GO:0009733 | response to auxin | 8.73E-15 |
| PH | PH_18JY | AX-111157582 | 1B | 672880906 | 1.8579/-- | 1.81E-04 | BLUPmrMLM | TraesCS1B01G462300 | 4.21 [PH2] | 4.33E-06 [PH2] | GO:0009734 | auxin-activated signaling pathway | 8.73E-15 |
| PH | PH_18JY | AX-111157582 | 1B | 672880906 | 1.8579/-- | 1.81E-04 | BLUPmrMLM | TraesCS1B01G462300 | 4.21 [PH2] | 4.33E-06 [PH2] | GO:0009742 | brassinosteroid mediated signaling pathway | 8.73E-15 |
| PH | PH_18JY | AX-111157582 | 1B | 672880906 | 1.8579/-- | 1.81E-04 | BLUPmrMLM | TraesCS1B01G462300 | 4.21 [PH2] | 4.33E-06 [PH2] | GO:0009741 | response to brassinosteroid | 8.73E-15 |
| PH | PH_18JY | AX-111157582 | 1B | 672880906 | 1.8579/-- | 1.81E-04 | BLUPmrMLM | TraesCS1B01G462300 | 4.21 [PH2] | 4.33E-06 [PH2] | GO:0071365 | cellular response to auxin stimulus | 8.73E-15 |
| PH | PH_18JY | AX-111163674 | 2A | 703863086 | -1.6668/-- | 2.30E-06 | BLUPmrMLM | TraesCS2A01G453300 | -2.16 [PH1] | 1.18E-03 [PH1] | GO:0009741 | response to brassinosteroid | 5.54E-04 |
| PH | PH_18XX | AX-111068539 | 2A | 753153585 | 0.7284/-- | 4.11E-05 | BLUPmrMLM | TraesCS2A01G545400 | 1.16 [PH2] | 1.76E-03 [PH2] | GO:0009741 | response to brassinosteroid | 3.53E-06 |
| PH | PH_18XX | AX-94488783 | 2A | 755788522 | -2.3172/-- | 5.36E-07 | BLUPmrMLM | TraesCS2A01G545400 | 1.16 [PH2] | 1.76E-03 [PH2] | GO:0009741 | response to brassinosteroid | 3.53E-06 |

| | | | | | | | | | | | | | | | |
|---|---|---|---|---|---|---|---|---|---|---|---|---|---|---|---|
| PH | PH_17JY | AX-110440215 | 2A | 753567811 | 2.3281/-- | 5.06E-05 | BLUPmrMLM | TraesCS2A01G545400 | 1.16 $^{PH2}$ | 1.76E-03 $^{PH2}$ | | GO:0009741 | response to brassinosteroid | 5.68E-08 | |
| PH | PH_17JY | AX-111729566 | 2A | 753018070 | -0.015672085 | 1.05E-07 | IIIVmrMLM | TraesCS2A01G545400 | 1.16 $^{PH2}$ | 1.76E-03 $^{PH2}$ | | GO:0009741 | response to brassinosteroid | 5.68E-08 | |
| PH | PH_17JY | AX-111216759 | 2A | 758935907 | 1.5274/-2.9681 | 2.14E-11 | IIIVmrMLM | TraesCS2A01G552400 | 1.39 $^{PH2}$; 1.44 $^{PH3}$ | 1.32E-02 $^{PH2}$; 1.38E-03 $^{PH3}$ | *OsMYB106* | GO:0009739 | response to gibberellin | 3.12E-21 | Wang et al., 2020 |
| PH | PH_17JY | AX-111216759 | 2A | 758935907 | 1.5274/-2.9681 | 2.14E-11 | IIIVmrMLM | TraesCS2A01G552400 | 1.39 $^{PH2}$; 1.44 $^{PH3}$ | 1.32E-02 $^{PH2}$; 1.38E-03 $^{PH3}$ | *OsMYB106* | GO:0009733 | response to auxin | 3.12E-21 | Wang et al., 2020 |
| PH | PH_18JY | AX-110103130 | 2B | 767028694 | -1.4056/-- | 9.47E-06 | BLUPmrMLM | TraesCS2B01G575900 | 1.32 $^{PH2}$ | 1.41E-02 $^{PH2}$ | | GO:0009741 | response to brassinosteroid | 9.98E-08 | |
| PH | PH_18ZMD | AX-110914455 | 3B | 20381356 | 9.896370023 | 8.16E-11 | IIIVmrMLM | TraesCS3B01G039100 | -1.58 $^{PH2}$ | 3.34E-02 $^{PH2}$ | | GO:0009739 | response to gibberellin | 7.18E-26 | |
| PH | PH_18AY | AX-109387101 | 3B | 17916598 | 1.6165/-- | 3.47E-06 | BLUPmrMLM | TraesCS3B01G039100 | -1.58 $^{PH2}$ | 3.34E-02 $^{PH2}$ | | GO:0071370 | cellular response to gibberellin stimulus | 9.43E-25 | |
| PH | PH_18AY | AX-109387101 | 3B | 17916598 | 1.6165/-- | 3.47E-06 | BLUPmrMLM | TraesCS3B01G039100 | -1.58 $^{PH2}$ | 3.34E-02 $^{PH2}$ | | GO:0009739 | response to gibberellin | 9.43E-25 | |
| PH | PH_18XX | AX-110960188 | 3B | 20818271 | 1.094453841 | 1.19E-16 | IIIVmrMLM | TraesCS3B01G039100 | -1.58 $^{PH2}$ | 3.34E-02 $^{PH2}$ | | GO:0010371 | regulation of gibberellin biosynthetic process | 1.26E-26 | |
| PH | PH_18ZMD | AX-110914455 | 3B | 20381356 | 9.896370023 | 8.16E-11 | IIIVmrMLM | TraesCS3B01G039100 | -1.58 $^{PH2}$ | 3.34E-02 $^{PH2}$ | | GO:0010371 | regulation of gibberellin biosynthetic process | 7.18E-26 | |
| PH | PH_18AY | AX-109387101 | 3B | 17916598 | 1.6165/-- | 3.47E-06 | BLUPmrMLM | TraesCS3B01G039100 | -1.58 $^{PH2}$ | 3.34E-02 $^{PH2}$ | | GO:0010371 | regulation of gibberellin biosynthetic process | 9.43E-25 | |
| PH | PH_18ZMD | AX-110914455 | 3B | 20381356 | 9.896370023 | 8.16E-11 | IIIVmrMLM | TraesCS3B01G039100 | -1.58 $^{PH2}$ | 3.34E-02 $^{PH2}$ | | GO:0071370 | cellular response to gibberellin stimulus | 7.18E-26 | |
| PH | PH_18AY | AX-109387101 | 3B | 17916598 | 1.6165/-- | 3.47E-06 | BLUPmrMLM | TraesCS3B01G039100 | -1.58 $^{PH2}$ | 3.34E-02 $^{PH2}$ | | GO:0010476 | gibberellin mediated signaling pathway | 9.43E-25 | |
| PH | PH_18AY | AX-109387101 | 3B | 17916598 | 1.6165/-- | 3.47E-06 | BLUPmrMLM | TraesCS3B01G039100 | -1.58 $^{PH2}$ | 3.34E-02 $^{PH2}$ | | GO:0009686 | gibberellin biosynthetic process | 9.43E-25 | |
| PH | PH_18ZMD | AX-110914455 | 3B | 20381356 | 9.896370023 | 8.16E-11 | IIIVmrMLM | TraesCS3B01G039100 | -1.58 $^{PH2}$ | 3.34E-02 $^{PH2}$ | | GO:0009685 | gibberellin metabolic process | 7.18E-26 | |
| PH | PH_18XX | AX-110960188 | 3B | 20818271 | 1.094453841 | 1.19E-16 | IIIVmrMLM | TraesCS3B01G039100 | -1.58 $^{PH2}$ | 3.34E-02 $^{PH2}$ | | GO:0010476 | gibberellin mediated signaling pathway | 1.26E-26 | |
| PH | PH_18XX | AX-110960188 | 3B | 20818271 | 1.094453841 | 1.19E-16 | IIIVmrMLM | TraesCS3B01G039100 | -1.58 $^{PH2}$ | 3.34E-02 $^{PH2}$ | | GO:0009685 | gibberellin metabolic process | 1.26E-26 | |
| PH | PH_18ZMD | AX-110914455 | 3B | 20381356 | 9.896370023 | 8.16E-11 | IIIVmrMLM | TraesCS3B01G039100 | -1.58 $^{PH2}$ | 3.34E-02 $^{PH2}$ | | GO:0009686 | gibberellin biosynthetic process | 7.18E-26 | |
| PH | PH_18AY | AX-109387101 | 3B | 17916598 | 1.6165/-- | 3.47E-06 | BLUPmrMLM | TraesCS3B01G039100 | -1.58 $^{PH2}$ | 3.34E-02 $^{PH2}$ | | GO:0010373 | negative regulation of gibberellin biosynthetic process | 9.43E-25 | |
| PH | PH_18AY | AX-109387101 | 3B | 17916598 | 1.6165/-- | 3.47E-06 | BLUPmrMLM | TraesCS3B01G039100 | -1.58 $^{PH2}$ | 3.34E-02 $^{PH2}$ | | GO:0009685 | gibberellin metabolic process | 9.43E-25 | |
| PH | PH_18ZMD | AX-110914455 | 3B | 20381356 | 9.896370023 | 8.16E-11 | IIIVmrMLM | TraesCS3B01G039100 | -1.58 $^{PH2}$ | 3.34E-02 $^{PH2}$ | | GO:0010373 | negative regulation of gibberellin biosynthetic process | 7.18E-26 | |
| PH | PH_18XX | AX-110960188 | 3B | 20818271 | 1.094453841 | 1.19E-16 | IIIVmrMLM | TraesCS3B01G039100 | -1.58 $^{PH2}$ | 3.34E-02 $^{PH2}$ | | GO:0009739 | response to gibberellin | 1.26E-26 | |
| PH | PH_18XX | AX-110960188 | 3B | 20818271 | 1.094453841 | 1.19E-16 | IIIVmrMLM | TraesCS3B01G039100 | -1.58 $^{PH2}$ | 3.34E-02 $^{PH2}$ | | GO:0009686 | gibberellin biosynthetic process | 1.26E-26 | |
| PH | PH_18XX | AX-110960188 | 3B | 20818271 | 1.094453841 | 1.19E-16 | IIIVmrMLM | TraesCS3B01G039100 | -1.58 $^{PH2}$ | 3.34E-02 $^{PH2}$ | | GO:0071370 | cellular response to gibberellin stimulus | 1.26E-26 | |

| | | | | | | | | | | | | | | | |
|---|---|---|---|---|---|---|---|---|---|---|---|---|---|---|---|
| PH | PH_18XX | AX-110960188 | 3B | 20818271 | 1.094453841 | 1.19E-16 | IIIVmrMLM | TraesCS3B01G039100 | -1.58 $^{PH2}$ | 3.34E-02 $^{PH2}$ | | GO:0010373 | negative regulation of gibberellin biosynthetic process | 1.26E-26 | |
| PH | PH_18ZMD | AX-110914455 | 3B | 20381356 | 9.896370023 | 8.16E-11 | IIIVmrMLM | TraesCS3B01G039100 | -1.58 $^{PH2}$ | 3.34E-02 $^{PH2}$ | | GO:0010476 | gibberellin mediated signaling pathway | 7.18E-26 | |
| PH | PH_17AY | AX-111697670 | 3D | 611314553 | 0.050662498 | 1.43E-06 | IIIVmrMLM | TraesCS3D01G540600 | -2.32 $^{PH2}$ | 7.26E-04 $^{PH2}$ | | GO:0009739 | response to gibberellin | 4.03E-16 | |
| PH | PH_18ZMD | AX-110437801 | 4A | 11172466 | -0.010247313 | 8.29E-07 | IIIVmrMLM | TraesCS4A01G016600 | 1.23 $^{PH3}$ | 4.10E-06 $^{PH3}$ | | GO:0009739 | response to gibberellin | 5.84E-74 | |
| PH | PH_18AY | AX-110429305 | 4A | 718875514 | 1.8719/-0.8655 | 1.21E-13 | IIIVmrMLM | TraesCS4A01G455200 | 3.8 $^{PH3}$ | 2.09E-03 $^{PH3}$ | | GO:0071365 | cellular response to auxin stimulus | 1.48E-21 | |
| PH | PH_18AY | AX-110429305 | 4A | 718875514 | 1.8719/-0.8655 | 1.21E-13 | IIIVmrMLM | TraesCS4A01G455200 | 3.8 $^{PH3}$ | 2.09E-03 $^{PH3}$ | | GO:0009734 | auxin-activated signaling pathway | 1.48E-21 | |
| PH | PH_18AY | AX-110429305 | 4A | 718875514 | 1.8719/-0.8655 | 1.21E-13 | IIIVmrMLM | TraesCS4A01G455200 | 3.8 $^{PH3}$ | 2.09E-03 $^{PH3}$ | | GO:0009742 | brassinosteroid mediated signaling pathway | 1.48E-21 | |
| PH | PH_18AY | AX-110429305 | 4A | 718875514 | 1.8719/-0.8655 | 1.21E-13 | IIIVmrMLM | TraesCS4A01G455200 | 3.8 $^{PH3}$ | 2.09E-03 $^{PH3}$ | | GO:0071367 | cellular response to brassinosteroid stimulus | 1.48E-21 | |
| PH | PH_18AY | AX-110429305 | 4A | 718875514 | 1.8719/-0.8655 | 1.21E-13 | IIIVmrMLM | TraesCS4A01G455200 | 3.8 $^{PH3}$ | 2.09E-03 $^{PH3}$ | | GO:0009733 | response to auxin | 1.48E-21 | |
| PH | PH_18AY | AX-110429305 | 4A | 718875514 | 1.8719/-0.8655 | 1.21E-13 | IIIVmrMLM | TraesCS4A01G455200 | 3.8 $^{PH3}$ | 2.09E-03 $^{PH3}$ | | GO:0009741 | response to brassinosteroid | 1.48E-21 | |
| PH | PH_18XX | AX-109299526 | 4A | 725843038 | -2.4133/-- | 3.33E-12 | BLUPmrMLM | TraesCS4A01G460100 | -1.22 $^{PH3}$; 2.96 $^{PH2}$ | 2.54E-02 $^{PH3}$; 2.63E-03 $^{PH2}$ | *KAO* | GO:0016131 | brassinosteroid metabolic process | 5.33E-24 | Miao et al., 2019 |
| PH | PH_17AY | AX-94746133 | 4A | 727055286 | 3.4269/-- | 2.78E-09 | BLUPmrMLM | TraesCS4A01G460100 | -1.22 $^{PH3}$; 2.96 $^{PH2}$ | 2.54E-02 $^{PH3}$; 2.63E-03 $^{PH2}$ | *KAO* | GO:0016131 | brassinosteroid metabolic process | 2.29E-17 | Miao et al., 2019 |
| PH | PH_18XX | AX-109299526 | 4A | 725843038 | -2.4133/-- | 3.33E-12 | BLUPmrMLM | TraesCS4A01G460100 | -1.22 $^{PH3}$; 2.96 $^{PH2}$ | 2.54E-02 $^{PH3}$; 2.63E-03 $^{PH2}$ | *KAO* | GO:0009686 | gibberellin biosynthetic process | 5.33E-24 | Miao et al., 2019 |
| PH | PH_18XX | AX-109299526 | 4A | 725843038 | -2.4133/-- | 3.33E-12 | BLUPmrMLM | TraesCS4A01G460100 | -1.22 $^{PH3}$; 2.96 $^{PH2}$ | 2.54E-02 $^{PH3}$; 2.63E-03 $^{PH2}$ | *KAO* | GO:0016132 | brassinosteroid biosynthetic process | 5.33E-24 | Miao et al., 2019 |
| PH | PH_17AY | AX-94746133 | 4A | 727055286 | 3.4269/-- | 2.78E-09 | BLUPmrMLM | TraesCS4A01G460100 | -1.22 $^{PH3}$; 2.96 $^{PH2}$ | 2.54E-02 $^{PH3}$; 2.63E-03 $^{PH2}$ | *KAO* | GO:0010268 | brassinosteroid homeostasis | 2.29E-17 | Miao et al., 2019 |
| PH | PH_17AY | AX-94746133 | 4A | 727055286 | 3.4269/-- | 2.78E-09 | BLUPmrMLM | TraesCS4A01G460100 | -1.22 $^{PH3}$; 2.96 $^{PH2}$ | 2.54E-02 $^{PH3}$; 2.63E-03 $^{PH2}$ | *KAO* | GO:0009686 | gibberellin biosynthetic process | 2.29E-17 | Miao et al., 2019 |
| PH | PH_17AY | AX-94746133 | 4A | 727055286 | 3.4269/-- | 2.78E-09 | BLUPmrMLM | TraesCS4A01G460100 | -1.22 $^{PH3}$; 2.96 $^{PH2}$ | 2.54E-02 $^{PH3}$; 2.63E-03 $^{PH2}$ | *KAO* | GO:0016132 | brassinosteroid biosynthetic process | 2.29E-17 | Miao et al., 2019 |
| PH | PH_17AY | AX-94746133 | 4A | 727055286 | 3.4269/-- | 2.78E-09 | BLUPmrMLM | TraesCS4A01G460100 | -1.22 $^{PH3}$; 2.96 $^{PH2}$ | 2.54E-02 $^{PH3}$; 2.63E-03 $^{PH2}$ | *KAO* | GO:0009685 | gibberellin metabolic process | 2.29E-17 | Miao et al., 2019 |
| PH | PH_18XX | AX-109299526 | 4A | 725843038 | -2.4133/-- | 3.33E-12 | BLUPmrMLM | TraesCS4A01G460100 | -1.22 $^{PH3}$; 2.96 $^{PH2}$ | 2.54E-02 $^{PH3}$; 2.63E-03 $^{PH2}$ | *KAO* | GO:0010268 | brassinosteroid homeostasis | 5.33E-24 | Miao et al., 2019 |
| PH | PH_18XX | AX-109299526 | 4A | 725843038 | -2.4133/-- | 3.33E-12 | BLUPmrMLM | TraesCS4A01G460100 | -1.22 $^{PH3}$; 2.96 $^{PH2}$ | 2.54E-02 $^{PH3}$; 2.63E-03 $^{PH2}$ | *KAO* | GO:0009685 | gibberellin metabolic process | 5.33E-24 | Miao et al., 2019 |
| PH | PH_18ZMD | AX-111031992 | 5A | 540369331 | 2.3827/-- | 7.61E-06 | BLUPmrMLM | TraesCS5A01G330500 | -1.69 $^{PH1}$ | 4.37E-03 $^{PH1}$ | | GO:0009741 | response to brassinosteroid | 1.44E-15 | |
| PH | PH_18XX | AX-110694654 | 5A | 708860993 | 0.03406013 | 7.52E-10 | IIIVmrMLM | TraesCS5A01G558600 | 1.72 $^{PH3}$ | 2.62E-03 $^{PH3}$ | | GO:0009739 | response to gibberellin | 7.02E-233 | |
| PH | PH_18ZMD | AX-110999817 | 5A | 708416688 | 2.7098/-7.6419 | 6.00E-73 | IIIVmrMLM | TraesCS5A01G558600 | 1.72 $^{PH3}$ | 2.62E-03 $^{PH3}$ | | GO:0009739 | response to gibberellin | 1.34E-192 | |
| PH | PH_17JY | AX-110999817 | 5A | 708416688 | 0.031115579 | 3.78E-48 | IIIVmrMLM | TraesCS5A01G558600 | 1.72 $^{PH3}$ | 2.62E-03 $^{PH3}$ | | GO:0009739 | response to gibberellin | 7.61E-159 | |

| | | | | | | | | | | | | | | | |
|---|---|---|---|---|---|---|---|---|---|---|---|---|---|---|---|
| PH | PH_18XX | AX-110999817 | 5A | 708416688 | 0.8526/-6.379 | 9.81E-109 | IIIVmrMLM | TraesCS5A01G558600 | 1.72 [PH3] | 2.62E-03 [PH3] | | GO:0009739 | response to gibberellin | 7.02E-233 | |
| PH | PH_18AY | AX-110999817 | 5A | 708416688 | 0.003116338 | 5.58E-64 | IIIVmrMLM | TraesCS5A01G558600 | 1.72 [PH3] | 2.62E-03 [PH3] | | GO:0009739 | response to gibberellin | 4.07E-177 | |
| PH | PH_17JY | AX-111013696 | 5B | 410725022 | 2.9161/-- | 4.13E-06 | BLUPmrMLM | TraesCS5B01G232100 | 5.38 [PH3] | 2.89E-03 [PH3] | | GO:0009741 | response to brassinosteroid | 1.41E-83 | |
| PH | PH_17JY | AX-111013696 | 5B | 410725022 | 2.9161/-- | 4.13E-06 | BLUPmrMLM | TraesCS5B01G232100 | 5.38 [PH3] | 2.89E-03 [PH3] | | GO:0071367 | cellular response to brassinosteroid stimulus | 1.41E-83 | |
| PH | PH_17ZMD | AX-108899168 | 5B | 410604227 | 0.2893/19.6436 | 4.14E-14 | IIIVmrMLM | TraesCS5B01G232100 | 5.38 [PH3] | 2.89E-03 [PH3] | | GO:0071367 | cellular response to brassinosteroid stimulus | 2.99E-100 | |
| PH | PH_17ZMD | AX-108899168 | 5B | 410604227 | 0.2893/19.6436 | 4.14E-14 | IIIVmrMLM | TraesCS5B01G232100 | 5.38 [PH3] | 2.89E-03 [PH3] | | GO:0009741 | response to brassinosteroid | 2.99E-100 | |
| PH | PH_17AY | AX-110371594 | 5B | 416378795 | -0.058999494 | 5.00E-07 | IIIVmrMLM | TraesCS5B01G236500 | -1.6 [PH3] | 1.92E-02 [PH3] | | GO:0016131 | brassinosteroid metabolic process | 9.77E-76 | |
| PH | PH_17AY | AX-110371594 | 5B | 416378795 | -0.058999494 | 5.00E-07 | IIIVmrMLM | TraesCS5B01G236500 | -1.6 [PH3] | 1.92E-02 [PH3] | | GO:0010268 | brassinosteroid homeostasis | 9.77E-76 | |
| PH | PH_BLUP | AX-111156055 | 5B | 416156777 | -2.2347/-- | 1.67E-13 | BLUPmrMLM | TraesCS5B01G236500 | -1.6 [PH3] | 1.92E-02 [PH3] | | GO:0016132 | brassinosteroid biosynthetic process | 1.24E-102 | |
| PH | PH_BLUP | AX-111156055 | 5B | 416156777 | -2.2347/-- | 1.67E-13 | BLUPmrMLM | TraesCS5B01G236500 | -1.6 [PH3] | 1.92E-02 [PH3] | | GO:0016131 | brassinosteroid metabolic process | 1.24E-102 | |
| PH | PH_17AY | AX-110371594 | 5B | 416378795 | -0.058999494 | 5.00E-07 | IIIVmrMLM | TraesCS5B01G236500 | -1.6 [PH3] | 1.92E-02 [PH3] | | GO:0016132 | brassinosteroid biosynthetic process | 9.77E-76 | |
| PH | PH_BLUP | AX-111156055 | 5B | 416156777 | -2.2347/-- | 1.67E-13 | BLUPmrMLM | TraesCS5B01G236500 | -1.6 [PH3] | 1.92E-02 [PH3] | | GO:0010268 | brassinosteroid homeostasis | 1.24E-102 | |
| PH | PH_18XX | AX-110706119 | 5B | 686315496 | -1.2521/-- | 2.63E-06 | BLUPmrMLM | TraesCS5B01G523100 | -2.66 [PH2] | 4.44E-02 [PH2] | | GO:0009739 | response to gibberellin | 1.50E-123 | |
| PH | PH_17ZMD | AX-95162142 | 6B | 199761147 | -26.81/-- | 2.35E-07 | EMMAX | TraesCS6B01G178300 | 1.67 [PH3] | 6.97E-03 [PH3] | | GO:0071367 | cellular response to brassinosteroid stimulus | 2.22E-25 | |
| PH | PH_17JY | AX-108853441 | 6B | 197242252 | 2.9131/-- | 7.58E-07 | BLUPmrMLM | TraesCS6B01G178300 | 1.67 [PH3] | 6.97E-03 [PH3] | | GO:0009742 | brassinosteroid mediated signaling pathway | 3.00E-26 | |
| PH | PH_17ZMD | AX-95162142 | 6B | 199761147 | -26.81/-- | 2.35E-07 | EMMAX | TraesCS6B01G178300 | 1.67 [PH3] | 6.97E-03 [PH3] | | GO:0009741 | response to brassinosteroid | 2.22E-25 | |
| PH | PH_17JY | AX-108853441 | 6B | 197242252 | 2.9131/-- | 7.58E-07 | BLUPmrMLM | TraesCS6B01G178300 | 1.67 [PH3] | 6.97E-03 [PH3] | | GO:0071367 | cellular response to brassinosteroid stimulus | 3.00E-26 | |
| PH | PH_18ZMD | AX-95162142 | 6B | 199761147 | -26.716/-- | 3.42E-07 | EMMAX | TraesCS6B01G178300 | 1.67 [PH3] | 6.97E-03 [PH3] | | GO:0009742 | brassinosteroid mediated signaling pathway | 2.59E-27 | |
| PH | PH_18ZMD | AX-95162142 | 6B | 199761147 | -26.716/-- | 3.42E-07 | EMMAX | TraesCS6B01G178300 | 1.67 [PH3] | 6.97E-03 [PH3] | | GO:0071367 | cellular response to brassinosteroid stimulus | 2.59E-27 | |
| PH | PH_17ZMD | AX-95162142 | 6B | 199761147 | -26.81/-- | 2.35E-07 | EMMAX | TraesCS6B01G178300 | 1.67 [PH3] | 6.97E-03 [PH3] | | GO:0009742 | brassinosteroid mediated signaling pathway | 2.22E-25 | |
| PH | PH_18ZMD | AX-95162142 | 6B | 199761147 | -26.716/-- | 3.42E-07 | EMMAX | TraesCS6B01G178300 | 1.67 [PH3] | 6.97E-03 [PH3] | | GO:0009741 | response to brassinosteroid | 2.59E-27 | |
| PH | PH_17JY | AX-108853441 | 6B | 197242252 | 2.9131/-- | 7.58E-07 | BLUPmrMLM | TraesCS6B01G178300 | 1.67 [PH3] | 6.97E-03 [PH3] | | GO:0009741 | response to brassinosteroid | 3.00E-26 | |
| PH | PH_17JY | AX-110129520 | 6B | 529056319 | 1.7667/-- | 1.71E-04 | BLUPmrMLM | TraesCS6B01G293600 | 3.07 [PH2] | 2.44E-04 [PH2] | | GO:0009741 | response to brassinosteroid | 1.01E-61 | |
| PH | PH_18ZMD | AX-109748962 | 6D | 161324640 | -1.8048/-- | 2.20E-13 | IIIVmrMLM | TraesCS6D01G173000 | 1.86 [PH3] | 3.41E-06 [PH3] | *OsCGA1* | GO:0010476 | gibberellin mediated signaling pathway | 5.66E-104 | Hudson et al. 2013 |

| | | | | | | | | | | | | | | | |
|---|---|---|---|---|---|---|---|---|---|---|---|---|---|---|---|
| PH | PH_18ZMD | AX-109748962 | 6D | 161324640 | -1.8048/-- | 2.20E-13 | IIIVmrMLM | TraesCS6D01G173000 | $1.86^{PH3}$ | $3.41E-06^{PH3}$ | *OsCGA1* | GO:0071370 | cellular response to gibberellin stimulus | 5.66E-104 | Hudson et al. 2013 |
| PH | PH_18ZMD | AX-109748962 | 6D | 161324640 | -1.8048/-- | 2.20E-13 | IIIVmrMLM | TraesCS6D01G173000 | $1.86^{PH3}$ | $3.41E-06^{PH3}$ | *OsCGA1* | GO:0009739 | response to gibberellin | 5.66E-104 | Hudson et al. 2013 |
| PH | PH_17AY | AX-110381728 | 7A | 535901170 | 4.113058893 | 1.08E-48 | IIIVmrMLM | TraesCS7A01G362300 | $-2.59^{PH2}$ | $2.90E-15^{PH2}$ | *d35*; *OsKOS4* | GO:0009739 | response to gibberellin | 6.26E-83 | Itoh et al. 2004; Zhu et al., 2005 |
| PH | PH_17AY | AX-110381728 | 7A | 535901170 | 4.113058893 | 1.08E-48 | IIIVmrMLM | TraesCS7A01G362300 | $-2.59^{PH2}$ | $2.90E-15^{PH2}$ | *d35*; *OsKOS4* | GO:0071370 | cellular response to gibberellin stimulus | 6.26E-83 | Itoh et al. 2004; Zhu et al., 2005 |
| PH | PH_BLUP | AX-111605089 | 7A | 538250230 | -0.009875478 | 4.11E-07 | IIIVmrMLM | TraesCS7A01G362300 | $-2.59^{PH2}$ | $2.90E-15^{PH2}$ | *d35*; *OsKOS4* | GO:0009686 | gibberellin biosynthetic process | 1.31E-110 | Itoh et al. 2004; Zhu et al., 2005 |
| PH | PH_17AY | AX-110381728 | 7A | 535901170 | 4.113058893 | 1.08E-48 | IIIVmrMLM | TraesCS7A01G362300 | $-2.59^{PH2}$ | $2.90E-15^{PH2}$ | *d35*; *OsKOS4* | GO:0009686 | gibberellin biosynthetic process | 6.26E-83 | Itoh et al. 2004; Zhu et al., 2005 |
| PH | PH_BLUP | AX-111605089 | 7A | 538250230 | -0.009875478 | 4.11E-07 | IIIVmrMLM | TraesCS7A01G362300 | $-2.59^{PH2}$ | $2.90E-15^{PH2}$ | *d35*; *OsKOS4* | GO:0009685 | gibberellin metabolic process | 1.31E-110 | Itoh et al. 2004; Zhu et al., 2005 |
| PH | PH_17AY | AX-110381728 | 7A | 535901170 | 4.113058893 | 1.08E-48 | IIIVmrMLM | TraesCS7A01G362300 | $-2.59^{PH2}$ | $2.90E-15^{PH2}$ | *d35*; *OsKOS4* | GO:0010476 | gibberellin mediated signaling pathway | 6.26E-83 | Itoh et al. 2004; Zhu et al., 2005 |
| PH | PH_BLUP | AX-111605089 | 7A | 538250230 | -0.009875478 | 4.11E-07 | IIIVmrMLM | TraesCS7A01G362300 | $-2.59^{PH2}$ | $2.90E-15^{PH2}$ | *d35*; *OsKOS4* | GO:0009739 | response to gibberellin | 1.31E-110 | Itoh et al. 2004; Zhu et al., 2005 |
| PH | PH_17AY | AX-110381728 | 7A | 535901170 | 4.113058893 | 1.08E-48 | IIIVmrMLM | TraesCS7A01G362300 | $-2.59^{PH2}$ | $2.90E-15^{PH2}$ | *d35*; *OsKOS4* | GO:0009685 | gibberellin metabolic process | 6.26E-83 | Itoh et al. 2004; Zhu et al., 2005 |
| PH | PH_BLUP | AX-111605089 | 7A | 538250230 | -0.009875478 | 4.11E-07 | IIIVmrMLM | TraesCS7A01G362300 | $-2.59^{PH2}$ | $2.90E-15^{PH2}$ | *d35*; *OsKOS4* | GO:0010476 | gibberellin mediated signaling pathway | 1.31E-110 | Itoh et al. 2004; Zhu et al., 2005 |
| PH | PH_BLUP | AX-111605089 | 7A | 538250230 | -0.009875478 | 4.11E-07 | IIIVmrMLM | TraesCS7A01G362300 | $-2.59^{PH2}$ | $2.90E-15^{PH2}$ | *d35*; *OsKOS4* | GO:0071370 | cellular response to gibberellin stimulus | 1.31E-110 | Itoh et al. 2004; Zhu et al., 2005 |
| PH | PH_18AY | AX-109620440 | 7A | 619621671 | 1.3878/-- | 1.36E-04 | BLUPmrMLM | TraesCS7A01G424700 | $-1.32^{PH1}$ | $6.31E-03^{PH1}$ | | GO:0009742 | brassinosteroid mediated signaling pathway | 9.63E-27 | |
| PH | PH_18AY | AX-109620440 | 7A | 619621671 | 1.3878/-- | 1.36E-04 | BLUPmrMLM | TraesCS7A01G424700 | $-1.32^{PH1}$ | $6.31E-03^{PH1}$ | | GO:0071367 | cellular response to brassinosteroid stimulus | 9.63E-27 | |
| PH | PH_18AY | AX-109620440 | 7A | 619621671 | 1.3878/-- | 1.36E-04 | BLUPmrMLM | TraesCS7A01G424700 | $-1.32^{PH1}$ | $6.31E-03^{PH1}$ | | GO:0009741 | response to brassinosteroid | 9.63E-27 | |
| SL | SL_17JY | AX-94413745 | 1A | 512666565 | -0.3672/-- | 6.39E-16 | IIIVmrMLM | TraesCS1A01G320200 | $-1.3^{SL}$ | $1.32E-03^{SL}$ | | GO:0009733 | response to auxin | 2.51E-02 | |
| SL | SL_17JY | AX-94413745 | 1A | 512666565 | -0.3672/-- | 6.39E-16 | IIIVmrMLM | TraesCS1A01G320200 | $-1.3^{SL}$ | $1.32E-03^{SL}$ | | GO:0010252 | auxin homeostasis | 2.51E-02 | |
| SL | SL_17JY | AX-94413745 | 1A | 512666565 | -0.3672/-- | 6.39E-16 | IIIVmrMLM | TraesCS1A01G320200 | $-1.3^{SL}$ | $1.32E-03^{SL}$ | | GO:0071365 | cellular response to auxin stimulus | 2.51E-02 | |
| SL | SL_17JY | AX-94413745 | 1A | 512666565 | -0.3672/-- | 6.39E-16 | IIIVmrMLM | TraesCS1A01G320200 | $-1.3^{SL}$ | $1.32E-03^{SL}$ | | GO:0009734 | auxin-activated signaling pathway | 2.51E-02 | |
| SL | SL_18JY | AX-110023784 | 1A | 520606961 | 0.073/-0.2172 | 4.64E-06 | IIIVmrMLM | TraesCS1A01G334200 | $-2.06^{SL}$ | $2.21E-03^{SL}$ | | GO:2000012 | regulation of auxin polar transport | 5.83E-05 | |
| SL | SL_18JY | AX-110394578 | 1A | 522239093 | -0.1464/-- | 7.60E-05 | BLUPmrMLM | TraesCS1A01G334200 | $-2.06^{SL}$ | $2.21E-03^{SL}$ | | GO:2000012 | regulation of auxin polar transport | 5.83E-05 | |
| SL | SL_18JY | AX-110023784 | 1A | 520606961 | 0.073/-0.2172 | 4.64E-06 | IIIVmrMLM | TraesCS1A01G334900 | $1.35^{SL}$ | $3.45E-07^{SL}$ | | GO:0009733 | response to auxin | 4.07E-03 | |
| SL | SL_18JY | AX-110023784 | 1A | 520606961 | 0.073/-0.2172 | 4.64E-06 | IIIVmrMLM | TraesCS1A01G334900 | $1.35^{SL}$ | $3.45E-07^{SL}$ | | GO:0009850 | auxin metabolic process | 4.07E-03 | |
| SL | SL_18JY | AX-110394578 | 1A | 522239093 | -0.1464/-- | 7.60E-05 | BLUPmrMLM | TraesCS1A01G334900 | $1.35^{SL}$ | $3.45E-07^{SL}$ | | GO:0009733 | response to auxin | 4.07E-03 | |

| | | | | | | | | | | | | | |
|---|---|---|---|---|---|---|---|---|---|---|---|---|---|
| SL | SL_18JY | AX-110394578 | 1A | 522239093 | -0.1464/-- | 7.60E-05 | BLUPmrMLM | TraesCS1A01G334900 | 1.35 $^{SL}$ | 3.45E-07 $^{SL}$ | GO:0009850 | auxin metabolic process | 4.07E-03 |
| SL | SL_18AY | AX-111668935 | 1B | 4531482 | 0.1247/-0.0237 | 8.35E-08 | IIIVmrMLM | TraesCS1B01G005300 | -4.98 $^{SL}$ | 2.38E-06 $^{SL}$ | GO:0010252 | auxin homeostasis | 1.54E-02 |
| SL | SL_17AY | AX-108848462 | 2A | 4110206 | 0.03904597 | 1.52E-07 | IIIVmrMLM | TraesCS2A01G009800 | -6.14 $^{SL}$ | 2.61E-03 $^{SL}$ | GO:0009851 | auxin biosynthetic process | 1.17E-08 |
| SL | SL_17AY | AX-108848462 | 2A | 4110206 | 0.03904597 | 1.52E-07 | IIIVmrMLM | TraesCS2A01G009800 | -6.14 $^{SL}$ | 2.61E-03 $^{SL}$ | GO:0009850 | auxin metabolic process | 1.17E-08 |
| SL | SL_17AY | AX-109311974 | 2A | 3110094 | 0.3853832/-- | 1.52E-06 | GEMMA | TraesCS2A01G009800 | -6.14 $^{SL}$ | 2.61E-03 $^{SL}$ | GO:0009850 | auxin metabolic process | 1.17E-08 |
| SL | SL_17AY | AX-109311974 | 2A | 3110094 | 0.3853832/-- | 1.52E-06 | GEMMA | TraesCS2A01G009800 | -6.14 $^{SL}$ | 2.61E-03 $^{SL}$ | GO:0009851 | auxin biosynthetic process | 1.17E-08 |
| SL | SL_17AY | AX-108848462 | 2A | 4110206 | 0.03904597 | 1.52E-07 | IIIVmrMLM | TraesCS2A01G013000 | 1.36 $^{SL}$ | 1.53E-03 $^{SL}$ | GO:0016132 | brassinosteroid biosynthetic process | 1.18E-04 |
| SL | SL_17AY | AX-108848462 | 2A | 4110206 | 0.03904597 | 1.52E-07 | IIIVmrMLM | TraesCS2A01G013000 | 1.36 $^{SL}$ | 1.53E-03 $^{SL}$ | GO:0016131 | brassinosteroid metabolic process | 1.18E-04 |
| SL | SL_17AY | AX-108848462 | 2A | 4110206 | 0.03904597 | 1.52E-07 | IIIVmrMLM | TraesCS2A01G013000 | 1.36 $^{SL}$ | 1.53E-03 $^{SL}$ | GO:0010268 | brassinosteroid homeostasis | 1.18E-04 |
| SL | SL_BLUP | AX-108882839 | 3A | 12323991 | 0.1319/-- | 5.99E-08 | BLUPmrMLM | TraesCS3A01G019600 | 2.72 $^{SL}$ | 2.77E-12 $^{SL}$ | GO:0071365 | cellular response to auxin stimulus | 7.57E-04 |
| SL | SL_18ZMD | AX-109896082 | 3A | 10221647 | 0.3507/-0.142 | 1.65E-15 | IIIVmrMLM | TraesCS3A01G019600 | 2.72 $^{SL}$ | 2.77E-12 $^{SL}$ | GO:0071365 | cellular response to auxin stimulus | 9.09E-28 |
| SL | SL_BLUP | AX-108882839 | 3A | 12323991 | 0.1319/-- | 5.99E-08 | BLUPmrMLM | TraesCS3A01G019600 | 2.72 $^{SL}$ | 2.77E-12 $^{SL}$ | GO:0009734 | auxin-activated signaling pathway | 7.57E-04 |
| SL | SL_BLUP | AX-108882839 | 3A | 12323991 | 0.1319/-- | 5.99E-08 | BLUPmrMLM | TraesCS3A01G019600 | 2.72 $^{SL}$ | 2.77E-12 $^{SL}$ | GO:0009733 | response to auxin | 7.57E-04 |
| SL | SL_18ZMD | AX-109896082 | 3A | 10221647 | 0.3507/-0.142 | 1.65E-15 | IIIVmrMLM | TraesCS3A01G019600 | 2.72 $^{SL}$ | 2.77E-12 $^{SL}$ | GO:0009733 | response to auxin | 9.09E-28 |
| SL | SL_18ZMD | AX-109896082 | 3A | 10221647 | 0.3507/-0.142 | 1.65E-15 | IIIVmrMLM | TraesCS3A01G019600 | 2.72 $^{SL}$ | 2.77E-12 $^{SL}$ | GO:0009734 | auxin-activated signaling pathway | 9.09E-28 |
| SL | SL_17JY | AX-111211642 | 3A | 67907762 | 0.164/-- | 1.90E-04 | BLUPmrMLM | TraesCS3A01G103800 | -1.58 $^{SL}$ | 3.23E-06 $^{SL}$ | GO:0016131 | brassinosteroid metabolic process | 1.10E-02 |
| SL | SL_17JY | AX-111211642 | 3A | 67907762 | 0.164/-- | 1.90E-04 | BLUPmrMLM | TraesCS3A01G103800 | -1.58 $^{SL}$ | 3.23E-06 $^{SL}$ | GO:0016132 | brassinosteroid biosynthetic process | 1.10E-02 |
| SL | SL_17JY | AX-111211642 | 3A | 67907762 | 0.164/-- | 1.90E-04 | BLUPmrMLM | TraesCS3A01G103800 | -1.58 $^{SL}$ | 3.23E-06 $^{SL}$ | GO:0010268 | brassinosteroid homeostasis | 1.10E-02 |
| SL | SL_BLUP | AX-108968290 | 4A | 11295242 | -0.011626438 | 3.87E-04 | IIIVmrMLM | TraesCS4A01G016600 | 1.23 $^{SL}$ | 4.10E-06 $^{SL}$ | GO:0009739 | response to gibberellin | 1.23E-03 |
| SL | SL_18JY | AX-109517804 | 5A | 502827669 | -0.017933922 | 8.69E-07 | IIIVmrMLM | TraesCS5A01G295800 | -4.63 $^{SL}$ | 3.52E-03 $^{SL}$ | GO:0009733 | response to auxin | 3.42E-02 |
| SL | SL_18JY | AX-109517804 | 5A | 502827669 | -0.017933922 | 8.69E-07 | IIIVmrMLM | TraesCS5A01G295800 | -4.63 $^{SL}$ | 3.52E-03 $^{SL}$ | GO:0009734 | auxin-activated signaling pathway | 3.42E-02 |
| SL | SL_18JY | AX-109517804 | 5A | 502827669 | -0.017933922 | 8.69E-07 | IIIVmrMLM | TraesCS5A01G295800 | -4.63 $^{SL}$ | 3.52E-03 $^{SL}$ | GO:0071365 | cellular response to auxin stimulus | 3.42E-02 |
| SL | SL_18AY | AX-94780072 | 5B | 13428632 | 0.196/-- | 6.35E-21 | IIIVmrMLM | TraesCS5B01G014100 | -2.38 $^{SL}$ | 4.69E-02 $^{SL}$ | GO:0009733 | response to auxin | 4.04E-03 |
| SL | SL_18AY | AX-94780072 | 5B | 13428632 | 0.196/-- | 6.35E-21 | IIIVmrMLM | TraesCS5B01G014100 | -2.38 $^{SL}$ | 4.69E-02 $^{SL}$ | GO:0009734 | auxin-activated signaling pathway | 4.04E-03 |

| | | | | | | | | | | | | | | | |
|---|---|---|---|---|---|---|---|---|---|---|---|---|---|---|---|
| SL | SL_18AY | AX-94780072 | 5B | 13428632 | 0.196/-- | 6.35E-21 | IIIVmrMLM | TraesCS5B01G014100 | -2.38$^{SL}$ | 4.69E-02$^{SL}$ | | GO:0071365 | cellular response to auxin stimulus | 4.04E-03 | |
| SL | SL_18AY | AX-110441595 | 5B | 417880578 | -0.1774/-- | 1.27E-04 | BLUPmrMLM | TraesCS5B01G236500 | -1.6$^{SL}$ | 1.92E-02$^{SL}$ | | GO:0016132 | brassinosteroid biosynthetic process | 1.43E-03 | |
| SL | SL_18AY | AX-110441595 | 5B | 417880578 | -0.1774/-- | 1.27E-04 | BLUPmrMLM | TraesCS5B01G236500 | -1.6$^{SL}$ | 1.92E-02$^{SL}$ | | GO:0016131 | brassinosteroid metabolic process | 1.43E-03 | |
| SL | SL_18AY | AX-110441595 | 5B | 417880578 | -0.1774/-- | 1.27E-04 | BLUPmrMLM | TraesCS5B01G236500 | -1.6$^{SL}$ | 1.92E-02$^{SL}$ | | GO:0010268 | brassinosteroid homeostasis | 1.43E-03 | |
| SL | SL_18JY | AX-109899485 | 5B | 465052250 | -0.2334/-- | 2.27E-09 | BLUPmrMLM | TraesCS5B01G277500 | -1.78$^{SL}$ | 1.75E-03$^{SL}$ | | GO:0010252 | auxin homeostasis | 9.78E-03 | |
| SL | SL_BLUP | AX-111598705 | 7B | 733609157 | -0.085/-- | 2.67E-05 | BLUPmrMLM | TraesCS7B01G478200 | 1.79$^{SL}$ | 8.52E-03$^{SL}$ | | GO:0009733 | response to auxin | 5.02E-03 | |
| SL | SL_BLUP | AX-111598705 | 7B | 733609157 | -0.085/-- | 2.67E-05 | BLUPmrMLM | TraesCS7B01G478200 | 1.79$^{SL}$ | 8.52E-03$^{SL}$ | | GO:0009734 | auxin-activated signaling pathway | 5.02E-03 | |
| SL | SL_BLUP | AX-111598705 | 7B | 733609157 | -0.085/-- | 2.67E-05 | BLUPmrMLM | TraesCS7B01G478200 | 1.79$^{SL}$ | 8.52E-03$^{SL}$ | | GO:0010600 | regulation of auxin biosynthetic process | 5.02E-03 | |
| SL | SL_BLUP | AX-111598705 | 7B | 733609157 | -0.085/-- | 2.67E-05 | BLUPmrMLM | TraesCS7B01G478200 | 1.79$^{SL}$ | 8.52E-03$^{SL}$ | | GO:0009739 | response to gibberellin | 5.02E-03 | |
| SL | SL_BLUP | AX-111598705 | 7B | 733609157 | -0.085/-- | 2.67E-05 | BLUPmrMLM | TraesCS7B01G478200 | 1.79$^{SL}$ | 8.52E-03$^{SL}$ | | GO:0090354 | regulation of auxin metabolic process | 5.02E-03 | |
| SL | SL_BLUP | AX-111598705 | 7B | 733609157 | -0.085/-- | 2.67E-05 | BLUPmrMLM | TraesCS7B01G478200 | 1.79$^{SL}$ | 8.52E-03$^{SL}$ | | GO:0071365 | cellular response to auxin stimulus | 5.02E-03 | |
| SL | SL_18AY | AX-111713035 | 7B | 743147463 | 2.671681416 | 6.99E-38 | IIIVmrMLM | TraesCS7B01G484200 | 1.69$^{SL}$ | 2.09E-10$^{SL}$ | | GO:0016131 | brassinosteroid metabolic process | 2.78E-04 | |
| SL | SL_18AY | AX-111713035 | 7B | 743147463 | 2.671681416 | 6.99E-38 | IIIVmrMLM | TraesCS7B01G484200 | 1.69$^{SL}$ | 2.09E-10$^{SL}$ | | GO:0016132 | brassinosteroid biosynthetic process | 2.78E-04 | |
| PPR | PPR_18ZMD | AX-94545636 | 2A | 745408732 | 0.0114/-- | 1.64E-05 | BLUPmrMLM | TraesCS2A01G521500 | 4.3$^{PPR1}$ | 4.72E-03$^{PPR1}$ | | GO:0009741 | response to brassinosteroid | 4.75E-05 | |
| PPR | PPR_18XX | AX-109312185 | 2A | 758923227 | 0.0003/0.0078 | 3.64E-04 | IIIVmrMLM | TraesCS2A01G552400 | 1.39$^{PPR1}$; 1.44$^{PPR2}$ | 1.32E-02$^{PPR1}$; 1.38E-03$^{PPR2}$ | *OsMYB106* | GO:0009739 | response to gibberellin | 8.44E-11 | Wang et al., 2020 |
| PPR | PPR_18XX | AX-109312185 | 2A | 758923227 | 0.0003/0.0078 | 3.64E-04 | IIIVmrMLM | TraesCS2A01G552400 | 1.39$^{PPR1}$; 1.44$^{PPR2}$ | 1.32E-02$^{PPR1}$; 1.38E-03$^{PPR2}$ | *OsMYB106* | GO:0009733 | response to auxin | 8.44E-11 | Wang et al., 2020 |
| PPR | PPR_18ZMD | AX-109842599 | 2B | 4838565 | 0.0113/-- | 1.46E-05 | BLUPmrMLM | TraesCS2B01G010100 | 2.83$^{PPR1}$ | 2.41E-02$^{PPR1}$ | | GO:0009851 | auxin biosynthetic process | 8.39E-04 | |
| PPR | PPR_18ZMD | AX-109842599 | 2B | 4838565 | 0.0113/-- | 1.46E-05 | BLUPmrMLM | TraesCS2B01G010100 | 2.83$^{PPR1}$ | 2.41E-02$^{PPR1}$ | | GO:0009742 | brassinosteroid mediated signaling pathway | 8.39E-04 | |
| PPR | PPR_18ZMD | AX-109842599 | 2B | 4838565 | 0.0113/-- | 1.46E-05 | BLUPmrMLM | TraesCS2B01G010100 | 2.83$^{PPR1}$ | 2.41E-02$^{PPR1}$ | | GO:0009741 | response to brassinosteroid | 8.39E-04 | |
| PPR | PPR_18ZMD | AX-109842599 | 2B | 4838565 | 0.0113/-- | 1.46E-05 | BLUPmrMLM | TraesCS2B01G010100 | 2.83$^{PPR1}$ | 2.41E-02$^{PPR1}$ | | GO:0071367 | cellular response to brassinosteroid stimulus | 8.39E-04 | |
| PPR | PPR_18ZMD | AX-109842599 | 2B | 4838565 | 0.0113/-- | 1.46E-05 | BLUPmrMLM | TraesCS2B01G010100 | 2.83$^{PPR1}$ | 2.41E-02$^{PPR1}$ | | GO:0090354 | regulation of auxin metabolic process | 8.39E-04 | |
| PPR | PPR_18ZMD | AX-109842599 | 2B | 4838565 | 0.0113/-- | 1.46E-05 | BLUPmrMLM | TraesCS2B01G010100 | 2.83$^{PPR1}$ | 2.41E-02$^{PPR1}$ | | GO:0009850 | auxin metabolic process | 8.39E-04 | |
| PPR | PPR_18ZMD | AX-109842599 | 2B | 4838565 | 0.0113/-- | 1.46E-05 | BLUPmrMLM | TraesCS2B01G010100 | 2.83$^{PPR1}$ | 2.41E-02$^{PPR1}$ | | GO:0010600 | regulation of auxin biosynthetic process | 8.39E-04 | |
| PPR | PPR_BLUP | AX-110543864 | 2B | 4780929 | -0.923076923 | 8.43E-09 | IIIVmrMLM | TraesCS2B01G010100 | 2.83$^{PPR1}$ | 2.41E-02$^{PPR1}$ | | GO:0009850 | auxin metabolic process | 1.43E-15 | |

| | | | | | | | | | | | | | |
|---|---|---|---|---|---|---|---|---|---|---|---|---|---|
| PPR | PPR_BLUP | AX-110543864 | 2B | 4780929 | -0.923076923 | 8.43E-09 | IIIVmrMLM | TraesCS2B01G010100 | 2.83 [PPR1] | 2.41E-02 [PPR1] | GO:0009742 | brassinosteroid mediated signaling pathway | 1.43E-15 |
| PPR | PPR_BLUP | AX-110543864 | 2B | 4780929 | -0.923076923 | 8.43E-09 | IIIVmrMLM | TraesCS2B01G010100 | 2.83 [PPR1] | 2.41E-02 [PPR1] | GO:0090354 | regulation of auxin metabolic process | 1.43E-15 |
| PPR | PPR_BLUP | AX-110543864 | 2B | 4780929 | -0.923076923 | 8.43E-09 | IIIVmrMLM | TraesCS2B01G010100 | 2.83 [PPR1] | 2.41E-02 [PPR1] | GO:0071367 | cellular response to brassinosteroid stimulus | 1.43E-15 |
| PPR | PPR_BLUP | AX-110543864 | 2B | 4780929 | -0.923076923 | 8.43E-09 | IIIVmrMLM | TraesCS2B01G010100 | 2.83 [PPR1] | 2.41E-02 [PPR1] | GO:0010600 | regulation of auxin biosynthetic process | 1.43E-15 |
| PPR | PPR_BLUP | AX-111651927 | 2B | 4836307 | -0.923076923 | 8.43E-09 | IIIVmrMLM | TraesCS2B01G010100 | 2.83 [PPR1] | 2.41E-02 [PPR1] | GO:0009741 | response to brassinosteroid | 1.43E-15 |
| PPR | PPR_BLUP | AX-110543864 | 2B | 4780929 | -0.923076923 | 8.43E-09 | IIIVmrMLM | TraesCS2B01G010100 | 2.83 [PPR1] | 2.41E-02 [PPR1] | GO:0009741 | response to brassinosteroid | 1.43E-15 |
| PPR | PPR_BLUP | AX-110543864 | 2B | 4780929 | -0.923076923 | 8.43E-09 | IIIVmrMLM | TraesCS2B01G010100 | 2.83 [PPR1] | 2.41E-02 [PPR1] | GO:0009851 | auxin biosynthetic process | 1.43E-15 |
| PPR | PPR_BLUP | AX-111651927 | 2B | 4836307 | -0.923076923 | 8.43E-09 | IIIVmrMLM | TraesCS2B01G010100 | 2.83 [PPR1] | 2.41E-02 [PPR1] | GO:0071367 | cellular response to brassinosteroid stimulus | 1.43E-15 |
| PPR | PPR_BLUP | AX-111651927 | 2B | 4836307 | -0.923076923 | 8.43E-09 | IIIVmrMLM | TraesCS2B01G010100 | 2.83 [PPR1] | 2.41E-02 [PPR1] | GO:0010600 | regulation of auxin biosynthetic process | 1.43E-15 |
| PPR | PPR_BLUP | AX-111651927 | 2B | 4836307 | -0.923076923 | 8.43E-09 | IIIVmrMLM | TraesCS2B01G010100 | 2.83 [PPR1] | 2.41E-02 [PPR1] | GO:0009851 | auxin biosynthetic process | 1.43E-15 |
| PPR | PPR_BLUP | AX-111651927 | 2B | 4836307 | -0.923076923 | 8.43E-09 | IIIVmrMLM | TraesCS2B01G010100 | 2.83 [PPR1] | 2.41E-02 [PPR1] | GO:0090354 | regulation of auxin metabolic process | 1.43E-15 |
| PPR | PPR_BLUP | AX-111651927 | 2B | 4836307 | -0.923076923 | 8.43E-09 | IIIVmrMLM | TraesCS2B01G010100 | 2.83 [PPR1] | 2.41E-02 [PPR1] | GO:0009850 | auxin metabolic process | 1.43E-15 |
| PPR | PPR_BLUP | AX-111651927 | 2B | 4836307 | -0.923076923 | 8.43E-09 | IIIVmrMLM | TraesCS2B01G010100 | 2.83 [PPR1] | 2.41E-02 [PPR1] | GO:0009742 | brassinosteroid mediated signaling pathway | 1.43E-15 |
| PPR | PPR_18ZMD | AX-109378500 | 3A | 746319006 | -1.296296296 | 1.60E-06 | IIIVmrMLM | TraesCS3A01G535100 | -2.37 [PPR1] | 3.91E-04 [PPR1] | GO:0009739 | response to gibberellin | 1.99E-12 |
| PPR | PPR_18AY | AX-111780324 | 4A | 13285565 | 0.28125 | 1.23E-09 | IIIVmrMLM | TraesCS4A01G016600 | 1.23 [PPR2] | 4.10E-06 [PPR2] | GO:0009739 | response to gibberellin | 8.35E-13 |
| PPR | PPR_18XX | AX-111167627 | 4A | 82255635 | 0.0002/-0.0087 | 1.69E-08 | IIIVmrMLM | TraesCS4A01G079300 | 4.3 [PPR2]; 4.9 [PPR1] | 1.02E-02 [PPR2]; 4.04E-02 [PPR1] | GO:0009739 | response to gibberellin | 2.64E-02 |
| PPR | PPR_18XX | AX-111167627 | 4A | 82255635 | 0.0002/-0.0087 | 1.69E-08 | IIIVmrMLM | TraesCS4A01G079300 | 4.3 [PPR2]; 4.9 [PPR1] | 1.02E-02 [PPR2]; 4.04E-02 [PPR1] | GO:0071370 | cellular response to gibberellin stimulus | 2.64E-02 |
| PPR | PPR_BLUP | AX-94926805 | 4A | 81817244 | 0.227272727 | 1.86E-12 | IIIVmrMLM | TraesCS4A01G079300 | 4.3 [PPR2]; 4.9 [PPR1] | 1.02E-02 [PPR2]; 4.04E-02 [PPR1] | GO:0009739 | response to gibberellin | 1.33E-07 |
| PPR | PPR_BLUP | AX-94926805 | 4A | 81817244 | 0.227272727 | 1.86E-12 | IIIVmrMLM | TraesCS4A01G079300 | 4.3 [PPR2]; 4.9 [PPR1] | 1.02E-02 [PPR2]; 4.04E-02 [PPR1] | GO:0071370 | cellular response to gibberellin stimulus | 1.33E-07 |
| PPR | PPR_BLUP | AX-94926805 | 4A | 81817244 | 0.227272727 | 1.86E-12 | IIIVmrMLM | TraesCS4A01G079300 | 4.3 [PPR2]; 4.9 [PPR1] | 1.02E-02 [PPR2]; 4.04E-02 [PPR1] | GO:0010476 | gibberellin mediated signaling pathway | 1.33E-07 |
| PPR | PPR_18XX | AX-111167627 | 4A | 82255635 | 0.0002/-0.0087 | 1.69E-08 | IIIVmrMLM | TraesCS4A01G079300 | 4.3 [PPR2]; 4.9 [PPR1] | 1.02E-02 [PPR2]; 4.04E-02 [PPR1] | GO:0010476 | gibberellin mediated signaling pathway | 2.64E-02 |
| PPR | PPR_18XX | AX-109874653 | 4B | 534407516 | 0.0044/-- | 1.12E-04 | BLUPmrMLM | TraesCS4B01G263400 | 3.87 [PPR1] | 1.12E-02 [PPR1] | GO:0009739 | response to gibberellin | 5.29E-27 |
| PPR | PPR_18XX | AX-110363537 | 4B | 533984924 | 0.01740832/-- | 1.92E-06 | GEMMA | TraesCS4B01G263400 | 3.87 [PPR1] | 1.12E-02 [PPR1] | GO:0009739 | response to gibberellin | 5.29E-27 |
| PPR | PPR_18XX | AX-109874653 | 4B | 534407516 | 0.0044/-- | 1.12E-04 | BLUPmrMLM | TraesCS4B01G263400 | 3.87 [PPR1] | 1.12E-02 [PPR1] | GO:0071370 | cellular response to gibberellin stimulus | 5.29E-27 |

| | | | | | | | | | | | | | | | |
|---|---|---|---|---|---|---|---|---|---|---|---|---|---|---|---|
| PPR | PPR_18XX | AX-110363537 | 4B | 533984924 | 0.01740832/-- | 1.92E-06 | GEMMA | TraesCS4B01G263400 | 3.87 $^{PPR1}$ | 1.12E-02 $^{PPR1}$ | | GO:0071370 | cellular response to gibberellin stimulus | 5.29E-27 | |
| PPR | PPR_18XX | AX-110030611 | 4B | 534066903 | -0.021945/-- | 7.19E-07 | EMMAX | TraesCS4B01G263400 | 3.87 $^{PPR1}$ | 1.12E-02 $^{PPR1}$ | | GO:0010476 | gibberellin mediated signaling pathway | 5.29E-27 | |
| PPR | PPR_18XX | AX-109874653 | 4B | 534407516 | 0.0044/-- | 1.12E-04 | BLUPmrMLM | TraesCS4B01G263400 | 3.87 $^{PPR1}$ | 1.12E-02 $^{PPR1}$ | | GO:0010476 | gibberellin mediated signaling pathway | 5.29E-27 | |
| PPR | PPR_18XX | AX-110030611 | 4B | 534066903 | -0.021945/-- | 7.19E-07 | EMMAX | TraesCS4B01G263400 | 3.87 $^{PPR1}$ | 1.12E-02 $^{PPR1}$ | | GO:0009739 | response to gibberellin | 5.29E-27 | |
| PPR | PPR_18XX | AX-110030611 | 4B | 534066903 | -0.021945/-- | 7.19E-07 | EMMAX | TraesCS4B01G263400 | 3.87 $^{PPR1}$ | 1.12E-02 $^{PPR1}$ | | GO:0071370 | cellular response to gibberellin stimulus | 5.29E-27 | |
| PPR | PPR_18XX | AX-110363537 | 4B | 533984924 | -0.017432/-- | 2.24E-06 | EMMAX | TraesCS4B01G263400 | 3.87 $^{PPR1}$ | 1.12E-02 $^{PPR1}$ | | GO:0010476 | gibberellin mediated signaling pathway | 5.29E-27 | |
| PPR | PPR_18XX | AX-110363537 | 4B | 533984924 | -0.017432/-- | 2.24E-06 | EMMAX | TraesCS4B01G263400 | 3.87 $^{PPR1}$ | 1.12E-02 $^{PPR1}$ | | GO:0071370 | cellular response to gibberellin stimulus | 5.29E-27 | |
| PPR | PPR_18XX | AX-110363537 | 4B | 533984924 | -0.017432/-- | 2.24E-06 | EMMAX | TraesCS4B01G263400 | 3.87 $^{PPR1}$ | 1.12E-02 $^{PPR1}$ | | GO:0009739 | response to gibberellin | 5.29E-27 | |
| PPR | PPR_18XX | AX-110363537 | 4B | 533984924 | 0.01740832/-- | 1.92E-06 | GEMMA | TraesCS4B01G263400 | 3.87 $^{PPR1}$ | 1.12E-02 $^{PPR1}$ | | GO:0010476 | gibberellin mediated signaling pathway | 5.29E-27 | |
| PPR | PPR_18XX | AX-110030611 | 4B | 534066903 | 0.02243506/-- | 4.60E-07 | GEMMA | TraesCS4B01G263400 | 3.87 $^{PPR1}$ | 1.12E-02 $^{PPR1}$ | | GO:0010476 | gibberellin mediated signaling pathway | 5.29E-27 | |
| PPR | PPR_18XX | AX-110030611 | 4B | 534066903 | 0.02243506/-- | 4.60E-07 | GEMMA | TraesCS4B01G263400 | 3.87 $^{PPR1}$ | 1.12E-02 $^{PPR1}$ | | GO:0009739 | response to gibberellin | 5.29E-27 | |
| PPR | PPR_18XX | AX-110030611 | 4B | 534066903 | 0.02243506/-- | 4.60E-07 | GEMMA | TraesCS4B01G263400 | 3.87 $^{PPR1}$ | 1.12E-02 $^{PPR1}$ | | GO:0071370 | cellular response to gibberellin stimulus | 5.29E-27 | |
| PPR | PPR_18XX | AX-109819320 | 5B | 556962611 | 0.0101/-- | 1.16E-12 | BLUPmrMLM | TraesCS5B01G377500 | -2.25 $^{PPR1}$ | 1.76E-03 $^{PPR1}$ | | GO:0009733 | response to auxin | 3.32E-24 | |
| PPR | PPR_18XX | AX-109819320 | 5B | 556962611 | 0.0101/-- | 1.16E-12 | BLUPmrMLM | TraesCS5B01G377500 | -2.25 $^{PPR1}$ | 1.76E-03 $^{PPR1}$ | | GO:0009739 | response to gibberellin | 3.32E-24 | |
| PPR | PPR_18JY | AX-109381150 | 5B | 685072957 | 0.051470588 | 4.26E-06 | IIIVmrMLM | TraesCS5B01G523100 | -2.66 $^{PPR1}$ | 4.44E-02 $^{PPR1}$ | | GO:0009739 | response to gibberellin | 3.46E-56 | |
| PPR | PPR_18JY | AX-109503672 | 6B | 527841682 | 0.0038/-- | 1.13E-04 | BLUPmrMLM | TraesCS6B01G293600 | 3.07 $^{PPR1}$ | 2.44E-04 $^{PPR1}$ | | GO:0009741 | response to brassinosteroid | 7.24E-65 | |
| PPR | PPR_18JY | AX-111514591 | 7A | 27904256 | -1.424242424 | 2.85E-14 | IIIVmrMLM | TraesCS7A01G059000 | 1.54 $^{PPR2}$; 4.84 $^{PPR1}$ | 5.72E-03 $^{PPR2}$; 8.79E-10 $^{PPR1}$ | *DLT* | GO:1900459 | positive regulation of brassinosteroid mediated signaling pathway | 1.14E-03 | Niu et al. 2022 |
| PPR | PPR_18JY | AX-111514591 | 7A | 27904256 | -1.424242424 | 2.85E-14 | IIIVmrMLM | TraesCS7A01G059000 | 1.54 $^{PPR2}$; 4.84 $^{PPR1}$ | 5.72E-03 $^{PPR2}$; 8.79E-10 $^{PPR1}$ | *DLT* | GO:1900457 | regulation of brassinosteroid mediated signaling pathway | 1.14E-03 | Niu et al. 2022 |

**Supplementary Table S9**. Table S9 All QTNs identified as associated with grain length in wheat

| Trait | Marker | Chr | Position | Add.effect | Dom.effect | P-value | Method | Sig. |
|---|---|---|---|---|---|---|---|---|
| GL_17AY | AX-86170993 | 1A | 301130580 | 0.0548 | | 2.00E-04 | BLUPmrMLM | SUG |
| GL_17AY | AX-94723352 | 1A | 571580530 | -0.0873 | | 6.50E-18 | IIIVmrMLM | SIG |
| GL_17AY | AX-109064815 | 1B | 69180085 | -0.0304 | -0.6068 | 2.93E-13 | IIIVmrMLM | SIG |
| GL_17AY | AX-111600456 | 1D | 330935948 | 0.0910 | -0.0175 | 5.43E-18 | IIIVmrMLM | SIG |
| GL_17AY | AX-110598923 | 2A | 187199075 | 0.0976 | | 1.53E-06 | BLUPmrMLM | SUG |
| GL_17AY | AX-110672921 | 2A | 488861670 | 0.0708 | | 1.56E-12 | IIIVmrMLM | SIG |
| GL_17AY | AX-109621306 | 2A | 554044925 | -0.0322 | 0.1082 | 1.05E-12 | IIIVmrMLM | SIG |
| GL_17AY | AX-109458770 | 2A | 716126215 | 0.1573 | | 2.07E-06 | GEMMA | SUG |
| GL_17AY | AX-109994714 | 2A | 716177205 | 0.1275 | | 6.35E-06 | BLUPmrMLM | SUG |
| GL_17AY | AX-108921258 | 2A | 716236583 | 0.1907 | | 2.63E-07 | GEMMA | SUG |
| GL_17AY | AX-110936584 | 2A | 716491516 | 0.1811 | | 1.67E-06 | GEMMA | SUG |
| GL_17AY | AX-109949360 | 2A | 716919824 | 0.2035 | | 5.50E-08 | GEMMA | SIG |
| GL_17AY | AX-111097278 | 2A | 717021012 | 0.1438 | | 1.81E-06 | GEMMA | SUG |
| GL_17AY | AX-110606231 | 2A | 717141146 | 0.2817 | | 1.79E-06 | GEMMA | SUG |
| GL_17AY | AX-109507407 | 2A | 717389779 | 0.1464 | | 1.82E-06 | GEMMA | SUG |
| GL_17AY | AX-110575805 | 2A | 717423753 | 0.1430 | | 1.10E-06 | GEMMA | SUG |
| GL_17AY | AX-111719891 | 2A | 717564817 | 0.1418 | | 1.80E-06 | GEMMA | SUG |
| GL_17AY | AX-110601263 | 2A | 717571266 | 0.1517 | | 1.53E-06 | GEMMA | SUG |
| GL_17AY | AX-110427653 | 2A | 717743589 | 0.1432 | | 1.67E-06 | GEMMA | SUG |
| GL_17AY | AX-108963673 | 2A | 717754883 | 0.1432 | | 1.67E-06 | GEMMA | SUG |
| GL_17AY | AX-109344129 | 2A | 717803976 | 0.1432 | | 1.67E-06 | GEMMA | SUG |
| GL_17AY | AX-111611081 | 2A | 717816882 | 0.1432 | | 1.67E-06 | GEMMA | SUG |
| GL_17AY | AX-111021808 | 2A | 718158173 | 0.1475 | | 1.82E-06 | GEMMA | SUG |
| GL_17AY | AX-109462354 | 2A | 718166340 | 0.1473 | | 2.00E-06 | GEMMA | SUG |
| GL_17AY | AX-109295270 | 2A | 718227374 | -0.0689 | 0.0723 | 4.39E-11 | IIIVmrMLM | SIG |
| GL_17AY | AX-111720998 | 2A | 719404597 | 0.1983 | | 1.33E-06 | GEMMA | SUG |
| GL_17AY | AX-109311877 | 2B | 23309728 | 0.0874 | 0.1466 | 3.10E-17 | IIIVmrMLM | SIG |
| GL_17AY | AX-111477705 | 2B | 303301761 | -0.0612 | | 8.43E-10 | IIIVmrMLM | SIG |
| GL_17AY | AX-110479087 | 2B | 754670959 | 0.0376 | 0.152 | 3.40E-04 | IIIVmrMLM | SUG |
| GL_17AY | AX-109813791 | 2D | 533205451 | 0.4475 | | 1.32E-06 | GEMMA | SUG |
| GL_17AY | AX-109875391 | 3A | 12330274 | 0.0520 | -0.2146 | 5.34E-07 | IIIVmrMLM | SUG |
| GL_17AY | AX-109947185 | 3A | 210249582 | -0.0081 | 0.1558 | 3.16E-08 | IIIVmrMLM | SIG |
| GL_17AY | AX-111079116 | 3A | 665749032 | -0.0170 | 0.3416 | 5.27E-09 | IIIVmrMLM | SIG |
| GL_17AY | AX-111587576 | 3A | 718337057 | 0.0226 | -0.7616 | 3.30E-08 | IIIVmrMLM | SIG |
| GL_17AY | AX-94498390 | 3A | 737969643 | 0.0065 | 1.3835 | 7.80E-08 | IIIVmrMLM | SIG |
| GL_17AY | AX-108913515 | 3B | 262878478 | 0.0328 | -0.1858 | 3.34E-12 | IIIVmrMLM | SIG |
| GL_17AY | AX-111701658 | 3B | 294269604 | -0.3664 | | 3.63E-07 | FarmCPU | SUG |
| GL_17AY | AX-109907632 | 3B | 297212638 | -0.4005 | | 3.25E-08 | FarmCPU | SIG |
| GL_17AY | AX-110017186 | 3B | 298569111 | -0.3135 | | 5.10E-07 | FarmCPU | SUG |
| GL_17AY | AX-111697025 | 3B | 780885069 | -0.0790 | | 4.23E-15 | IIIVmrMLM | SIG |
| GL_17AY | AX-109315197 | 4A | 625104917 | 0.0182 | 0.7363 | 1.80E-05 | IIIVmrMLM | SUG |

| | | | | | | | | |
|---|---|---|---|---|---|---|---|---|
| GL_17AY | AX-109633846 | 4A | 707250680 | 0.0353 | 0.5806 | 4.67E-13 | IIIVmrMLM | SIG |
| GL_17AY | AX-111558259 | 4B | 602408602 | -0.0187 | 1.0564 | 1.77E-13 | IIIVmrMLM | SIG |
| GL_17AY | AX-110461516 | 5B | 13367717 | -0.1111 | -0.1023 | 6.06E-26 | IIIVmrMLM | SIG |
| GL_17AY | AX-110624525 | 5B | 115411485 | -0.0017 | 0.3667 | 1.05E-05 | IIIVmrMLM | SUG |
| GL_17AY | AX-110400528 | 5B | 383029696 | -0.0305 | 0.5187 | 1.36E-09 | IIIVmrMLM | SIG |
| GL_17AY | AX-95172045 | 5B | 406063074 | -0.0546 | -0.4537 | 1.18E-13 | IIIVmrMLM | SIG |
| GL_17AY | AX-109319325 | 5B | 429791616 | 0.0608 | 0.045 | 2.32E-08 | IIIVmrMLM | SIG |
| GL_17AY | AX-110519538 | 5B | 523182125 | 0.0151 | -0.1207 | 2.00E-05 | IIIVmrMLM | SUG |
| GL_17AY | AX-108896792 | 5B | 692310285 | 0.0012 | -0.2739 | 1.12E-08 | IIIVmrMLM | SIG |
| GL_17AY | AX-110076701 | 5D | 133535094 | -0.0292 | 0.2136 | 1.36E-05 | IIIVmrMLM | SUG |
| GL_17AY | AX-109213583 | 5D | 288600581 | -1.5616 | | 1.20E-06 | EMMAX | SUG |
| GL_17AY | AX-110413677 | 6A | 2058555 | 0.0008 | 0.1784 | 2.53E-05 | IIIVmrMLM | SUG |
| GL_17AY | AX-110511492 | 6B | 437452265 | 0.0159 | 0.2684 | 2.86E-05 | IIIVmrMLM | SUG |
| GL_17AY | AX-110997312 | 6B | 524524383 | -0.0563 | | 1.98E-04 | BLUPmrMLM | SUG |
| GL_17AY | AX-110987911 | 6B | 685197457 | -0.0603 | 0.0689 | 2.71E-09 | IIIVmrMLM | SIG |
| GL_17AY | AX-111370974 | 6D | 137893383 | -0.1098 | | 4.25E-06 | BLUPmrMLM | SUG |
| GL_17AY | AX-95107462 | 6D | 318348669 | 0.0800 | | 1.62E-04 | BLUPmrMLM | SUG |
| GL_17AY | AX-111550880 | 6D | 432873020 | -0.0938 | -0.0001 | 2.50E-13 | IIIVmrMLM | SIG |
| GL_17AY | AX-110074625 | 6D | 462630227 | -0.0356 | 0.1295 | 3.65E-08 | IIIVmrMLM | SIG |
| GL_17AY | AX-109869515 | 6D | 469164504 | 0.0263 | -0.2432 | 6.32E-12 | IIIVmrMLM | SIG |
| GL_17AY | AX-109829501 | 7A | 31195495 | -0.0130 | -0.6111 | 1.34E-06 | IIIVmrMLM | SUG |
| GL_17AY | AX-109855240 | 7A | 675486517 | -0.0067 | -0.1757 | 4.63E-06 | IIIVmrMLM | SUG |
| GL_17AY | AX-109279123 | 7B | 42931162 | -0.0055 | 0.1908 | 6.73E-06 | IIIVmrMLM | SUG |
| GL_17AY | AX-111573433 | 7D | 621486358 | -1.5616 | | 1.20E-06 | EMMAX | SUG |
| GL_17SQ | AX-111144479 | 1A | 9639133 | -0.1179 | -0.2548 | 3.96E-43 | IIIVmrMLM | SIG |
| GL_17SQ | AX-111770952 | 1A | 462335079 | 0.0467 | | 5.90E-06 | BLUPmrMLM | SUG |
| GL_17SQ | AX-110054683 | 1B | 42171178 | -0.1653 | -0.2607 | 1.46E-78 | IIIVmrMLM | SIG |
| GL_17SQ | AX-108858909 | 1B | 648990743 | -0.0517 | | 4.20E-05 | BLUPmrMLM | SUG |
| GL_17SQ | AX-95681612 | 1D | 48401161 | 0.3367 | | 1.38E-06 | FarmCPU | SUG |
| GL_17SQ | AX-94780647 | 1D | 86925751 | -0.1785 | | 6.63E-10 | BLUPmrMLM | SIG |
| GL_17SQ | AX-95180170 | 1D | 216400005 | -0.0592 | | 1.44E-04 | BLUPmrMLM | SUG |
| GL_17SQ | AX-94516823 | 2A | 191878516 | -0.0610 | | 1.82E-05 | BLUPmrMLM | SUG |
| GL_17SQ | AX-110914958 | 2A | 681477062 | 0.0858 | | 2.41E-05 | BLUPmrMLM | SUG |
| GL_17SQ | AX-109471825 | 2A | 693262436 | 0.1024 | 0.7267 | 4.24E-31 | IIIVmrMLM | SIG |
| GL_17SQ | AX-109978759 | 2A | 712315567 | -0.0441 | | 7.56E-06 | BLUPmrMLM | SUG |
| GL_17SQ | AX-109361285 | 2B | 764735126 | -0.0568 | | 2.48E-07 | BLUPmrMLM | SUG |
| GL_17SQ | AX-109951325 | 2B | 776816206 | -0.5438 | | 2.10E-07 | EMMAX | SUG |
| GL_17SQ | AX-95070110 | 2D | 122703116 | -1.4881 | | 1.97E-07 | EMMAX | SUG |
| GL_17SQ | AX-111570261 | 3B | 5497775 | -0.1178 | 0.109 | 1.54E-32 | IIIVmrMLM | SIG |
| GL_17SQ | AX-109274638 | 3B | 237325305 | -0.2409 | | 1.08E-06 | FarmCPU | SUG |
| GL_17SQ | AX-110913674 | 3B | 423621581 | -0.0613 | | 1.84E-05 | BLUPmrMLM | SUG |
| GL_17SQ | AX-111479930 | 3B | 490588211 | 0.4817 | | 9.32E-07 | GEMMA | SUG |
| GL_17SQ | AX-95023938 | 3B | 542735535 | -0.4077 | | 2.02E-08 | EMMAX | SIG |
| GL_17SQ | AX-95023938 | 3B | 542735535 | 0.4113 | | 3.33E-09 | GEMMA | SIG |

| | | | | | | | | |
|---|---|---|---|---|---|---|---|---|
| GL_17SQ | AX-108824541 | 3B | 673850284 | -1.4881 | | 1.97E-07 | EMMAX | SUG |
| GL_17SQ | AX-108794942 | 3B | 794344602 | 0.0047 | 0.4259 | 1.13E-04 | IIIVmrMLM | SUG |
| GL_17SQ | AX-111278919 | 3D | 104593669 | -1.4881 | | 1.97E-07 | EMMAX | SUG |
| GL_17SQ | AX-95234844 | 3D | 185792116 | -1.5424 | | 8.78E-11 | EMMAX | SIG |
| GL_17SQ | AX-109736551 | 3D | 371754400 | 0.0583 | | 3.76E-06 | BLUPmrMLM | SUG |
| GL_17SQ | AX-108956954 | 4A | 579840479 | 0.3675 | | 1.99E-06 | GEMMA | SUG |
| GL_17SQ | AX-94728117 | 4B | 232103485 | -1.5424 | | 8.78E-11 | EMMAX | SIG |
| GL_17SQ | AX-94547062 | 4B | 612846887 | -0.2796 | -0.1464 | 1.90E-98 | IIIVmrMLM | SIG |
| GL_17SQ | AX-108823697 | 4B | 661144811 | -0.0630 | | 7.61E-07 | BLUPmrMLM | SUG |
| GL_17SQ | AX-109042400 | 4D | 127693435 | -1.4881 | | 1.97E-07 | EMMAX | SUG |
| GL_17SQ | AX-109318201 | 4D | 197019433 | 0.2892 | | 6.77E-07 | FarmCPU | SUG |
| GL_17SQ | AX-110970840 | 4D | 197285998 | -1.4881 | | 1.97E-07 | EMMAX | SUG |
| GL_17SQ | AX-110357870 | 4D | 220975170 | -1.4881 | | 1.97E-07 | EMMAX | SUG |
| GL_17SQ | AX-110737051 | 4D | 294359371 | -1.4881 | | 1.97E-07 | EMMAX | SUG |
| GL_17SQ | AX-111489178 | 4D | 425247926 | -1.4881 | | 1.97E-07 | EMMAX | SUG |
| GL_17SQ | AX-109061821 | 5A | 502822955 | -0.0209 | -0.1549 | 2.07E-08 | IIIVmrMLM | SIG |
| GL_17SQ | AX-95006111 | 5A | 521934541 | -0.0452 | | 6.23E-05 | BLUPmrMLM | SUG |
| GL_17SQ | AX-111531103 | 5B | 466935159 | 0.0759 | -0.0782 | 5.36E-15 | IIIVmrMLM | SIG |
| GL_17SQ | AX-109839495 | 5B | 466991414 | -0.2611 | | 1.56E-06 | GEMMA | SUG |
| GL_17SQ | AX-94875579 | 5D | 50921989 | -1.4881 | | 1.97E-07 | EMMAX | SUG |
| GL_17SQ | AX-111059304 | 5D | 207969939 | -1.4881 | | 1.97E-07 | EMMAX | SUG |
| GL_17SQ | AX-110447123 | 6A | 16703177 | -0.0353 | | 1.54E-04 | BLUPmrMLM | SUG |
| GL_17SQ | AX-111766874 | 6A | 445114359 | -0.0576 | -0.0059 | 8.21E-09 | IIIVmrMLM | SIG |
| GL_17SQ | AX-111051197 | 6A | 445357012 | 0.2202 | | 8.51E-07 | GEMMA | SUG |
| GL_17SQ | AX-108831348 | 6A | 446302585 | -0.0812 | | 2.66E-09 | BLUPmrMLM | SIG |
| GL_17SQ | AX-108831348 | 6A | 446302585 | -0.1003 | | 2.24E-06 | GEMMA | SUG |
| GL_17SQ | AX-109421708 | 6B | 467871355 | 0.0306 | | 9.46E-05 | BLUPmrMLM | SUG |
| GL_17SQ | AX-110460521 | 6B | 468028930 | 0.0611 | 0.0797 | 3.24E-10 | IIIVmrMLM | SIG |
| GL_17SQ | AX-112289648 | 6B | 470830331 | -0.1247 | -0.4335 | 5.86E-45 | IIIVmrMLM | SIG |
| GL_17SQ | AX-95013272 | 6B | 477225270 | 0.2025 | | 7.74E-07 | EMMAX | SUG |
| GL_17SQ | AX-95013272 | 6B | 477225270 | -0.2003 | | 5.58E-07 | GEMMA | SUG |
| GL_17SQ | AX-94588720 | 6B | 477790406 | 0.2137 | | 5.95E-07 | EMMAX | SUG |
| GL_17SQ | AX-94588720 | 6B | 477790406 | -0.2118 | | 3.38E-07 | GEMMA | SUG |
| GL_17SQ | AX-111550880 | 6D | 432873020 | 0.0477 | | 7.78E-05 | BLUPmrMLM | SUG |
| GL_17SQ | AX-108725320 | 7A | 669775668 | -0.1025 | 0.061 | 1.13E-24 | IIIVmrMLM | SIG |
| GL_17SQ | AX-110033032 | 7A | 716262864 | -0.0116 | -1.2501 | 5.35E-14 | IIIVmrMLM | SIG |
| GL_17SQ | AX-109460479 | 7B | 628754127 | 0.0261 | -1.2217 | 9.23E-09 | IIIVmrMLM | SIG |
| GL_17SQ | AX-110028290 | 7D | 4630546 | 0.1483 | 0.0587 | 6.58E-48 | IIIVmrMLM | SIG |
| GL_17SQ | AX-111985769 | 7D | 270599780 | -1.4881 | | 1.97E-07 | EMMAX | SUG |
| GL_17SQ | AX-109057282 | 7D | 546415051 | -0.0915 | | 4.93E-22 | IIIVmrMLM | SIG |
| GL_17SQ | AX-108877934 | 7D | 615887708 | 0.0634 | | 4.37E-05 | BLUPmrMLM | SUG |
| GL_17XX | AX-109836265 | 1A | 577989829 | 0.1197 | | 8.80E-09 | BLUPmrMLM | SIG |
| GL_17XX | AX-111008033 | 1B | 15169796 | 0.1963 | | 1.50E-06 | GEMMA | SUG |
| GL_17XX | AX-109420526 | 1B | 29217823 | -0.2361 | | 2.57E-07 | EMMAX | SUG |

| | | | | | | | |
|---|---|---|---|---|---|---|---|
| GL_17XX | AX-109420526 | 1B | 29217823 | 0.2341 | | 5.56E-07 GEMMA | SUG |
| GL_17XX | AX-110591874 | 1B | 29408765 | -0.1931 | | 2.05E-06 EMMAX | SUG |
| GL_17XX | AX-109885673 | 1B | 31048864 | -0.2176 | | 4.37E-07 EMMAX | SUG |
| GL_17XX | AX-109885673 | 1B | 31048864 | 0.2143 | | 2.21E-07 GEMMA | SUG |
| GL_17XX | AX-94615994 | 1B | 33979041 | 0.4303 | | 9.32E-07 GEMMA | SUG |
| GL_17XX | AX-111069652 | 1B | 36086820 | 0.2271 | | 4.76E-07 GEMMA | SUG |
| GL_17XX | AX-94698017 | 1B | 86708972 | -0.4744 | | 2.80E-08 EMMAX | SIG |
| GL_17XX | AX-94698017 | 1B | 86708972 | 0.4732 | | 2.05E-08 GEMMA | SIG |
| GL_17XX | AX-95242102 | 1B | 90699644 | 0.2103 | | 6.71E-07 GEMMA | SUG |
| GL_17XX | AX-95171244 | 1B | 90992756 | 0.1763 | | 1.03E-06 GEMMA | SUG |
| GL_17XX | AX-95010697 | 1B | 90992761 | 0.1768 | | 1.37E-06 GEMMA | SUG |
| GL_17XX | AX-95173174 | 1B | 90993055 | -0.1902 | | 1.08E-06 EMMAX | SUG |
| GL_17XX | AX-95173174 | 1B | 90993055 | 0.1895 | | 1.09E-07 GEMMA | SIG |
| GL_17XX | AX-111174986 | 1B | 119881507 | 0.4406 | | 3.59E-07 GEMMA | SUG |
| GL_17XX | AX-111541173 | 1B | 136627719 | 0.2286 | | 1.14E-06 GEMMA | SUG |
| GL_17XX | AX-111116469 | 1B | 143057215 | 0.3842 | | 4.28E-07 GEMMA | SUG |
| GL_17XX | AX-94390293 | 1B | 176996634 | 0.4249 | | 1.48E-06 GEMMA | SUG |
| GL_17XX | AX-108931371 | 1B | 185429686 | -0.3123 | | 1.04E-08 EMMAX | SIG |
| GL_17XX | AX-108931371 | 1B | 185429686 | 0.3099 | | 2.53E-09 GEMMA | SIG |
| GL_17XX | AX-108931371 | 1B | 185429686 | -0.0437 | 0.2824 | 1.93E-22 IIIVmrMLM | SIG |
| GL_17XX | AX-110481673 | 1B | 640606889 | -0.0424 | | 2.24E-05 BLUPmrMLM | SUG |
| GL_17XX | AX-95129810 | 1D | 513067 | -0.0793 | | 6.31E-05 BLUPmrMLM | SUG |
| GL_17XX | AX-109860229 | 1D | 2156335 | -0.0616 | | 1.59E-05 BLUPmrMLM | SUG |
| GL_17XX | AX-94786159 | 1D | 79116398 | -0.4331 | | 1.17E-08 EMMAX | SIG |
| GL_17XX | AX-94786159 | 1D | 79116398 | 0.4274 | | 1.06E-08 GEMMA | SIG |
| GL_17XX | AX-110917931 | 1D | 175622526 | 2.4149 | | 1.87E-12 EMMAX | SIG |
| GL_17XX | AX-111089849 | 2A | 748701272 | 0.0272 | | 1.41E-04 BLUPmrMLM | SUG |
| GL_17XX | AX-108781105 | 2A | 752793414 | -0.0580 | | 7.76E-06 BLUPmrMLM | SUG |
| GL_17XX | AX-110975896 | 2B | 293220306 | 0.7658 | | 4.94E-07 EMMAX | SUG |
| GL_17XX | AX-109435051 | 2B | 315096464 | -0.9845 | | 1.45E-10 EMMAX | SIG |
| GL_17XX | AX-108735017 | 2D | 5761352 | 1.2917 | | 1.17E-07 EMMAX | SUG |
| GL_17XX | AX-94804217 | 2D | 8478708 | -0.0594 | | 5.11E-05 BLUPmrMLM | SUG |
| GL_17XX | AX-110999397 | 2D | 75749348 | -0.0392 | | 3.73E-05 BLUPmrMLM | SUG |
| GL_17XX | AX-95118616 | 2D | 544781641 | 0.1408 | | 5.87E-05 BLUPmrMLM | SUG |
| GL_17XX | AX-111176610 | 3B | 440679552 | -0.8529 | | 4.34E-08 EMMAX | SIG |
| GL_17XX | AX-111150243 | 3B | 788378866 | -0.0808 | | 1.40E-06 BLUPmrMLM | SUG |
| GL_17XX | AX-109112674 | 3D | 90767918 | 2.4149 | | 1.87E-12 EMMAX | SIG |
| GL_17XX | AX-95234844 | 3D | 185792116 | -1.4219 | | 4.68E-07 EMMAX | SUG |
| GL_17XX | AX-108933702 | 3D | 445322655 | -0.4610 | | 5.13E-09 EMMAX | SIG |
| GL_17XX | AX-108933702 | 3D | 445322655 | 0.5138 | | 6.89E-09 GEMMA | SIG |
| GL_17XX | AX-110419491 | 4A | 28563674 | -0.0447 | | 1.13E-04 BLUPmrMLM | SUG |
| GL_17XX | AX-94758025 | 4A | 38840188 | 2.4149 | | 1.87E-12 EMMAX | SIG |
| GL_17XX | AX-109509823 | 4A | 54984149 | -0.3416 | | 1.39E-06 EMMAX | SUG |
| GL_17XX | AX-94543301 | 4A | 539756222 | -0.4502 | | 6.57E-07 EMMAX | SUG |

| | | | | | | | | |
|---|---|---|---|---|---|---|---|---|
| GL_17XX | AX-94543301 | 4A | 539756222 | 0.4428 | | 1.05E-06 | GEMMA | SUG |
| GL_17XX | AX-109305727 | 4A | 616674326 | -0.0918 | | 3.56E-08 | BLUPmrMLM | SIG |
| GL_17XX | AX-108791521 | 4B | 10003294 | -0.1265 | | 6.47E-07 | FarmCPU | SUG |
| GL_17XX | AX-109541464 | 4B | 15750376 | 0.0606 | | 3.64E-05 | BLUPmrMLM | SUG |
| GL_17XX | AX-94728117 | 4B | 232103485 | -1.4219 | | 4.68E-07 | EMMAX | SUG |
| GL_17XX | AX-94581617 | 4B | 560014564 | 2.4149 | | 1.87E-12 | EMMAX | SIG |
| GL_17XX | AX-111483036 | 4B | 581503751 | 0.0913 | | 2.05E-07 | BLUPmrMLM | SUG |
| GL_17XX | AX-111879954 | 4D | 150774424 | 1.0931 | | 3.07E-07 | EMMAX | SUG |
| GL_17XX | AX-111903053 | 4D | 484511212 | -0.0438 | | 8.24E-05 | BLUPmrMLM | SUG |
| GL_17XX | AX-109382475 | 5A | 595390927 | 0.0381 | | 2.96E-05 | BLUPmrMLM | SUG |
| GL_17XX | AX-109473519 | 5A | 622528405 | -0.2011 | | 7.01E-07 | EMMAX | SUG |
| GL_17XX | AX-109473519 | 5A | 622528405 | 0.2024 | | 5.44E-07 | GEMMA | SUG |
| GL_17XX | AX-110997479 | 5A | 622669723 | 0.1668 | | 2.02E-06 | GEMMA | SUG |
| GL_17XX | AX-111575882 | 5A | 622679543 | 0.1575 | | 2.04E-06 | GEMMA | SUG |
| GL_17XX | AX-111470748 | 5B | 555215171 | -0.1349 | | 9.17E-12 | BLUPmrMLM | SIG |
| GL_17XX | AX-111019304 | 5B | 708222456 | -0.0620 | | 6.78E-06 | BLUPmrMLM | SUG |
| GL_17XX | AX-109482637 | 5D | 396416138 | 2.4149 | | 1.87E-12 | EMMAX | SIG |
| GL_17XX | AX-94497850 | 5D | 498186974 | -0.1610 | | 6.14E-06 | BLUPmrMLM | SUG |
| GL_17XX | AX-109851112 | 6A | 21639408 | 0.0608 | | 5.16E-05 | BLUPmrMLM | SUG |
| GL_17XX | AX-111706207 | 6A | 446904387 | 0.1310 | | 1.98E-06 | GEMMA | SUG |
| GL_17XX | AX-110934121 | 6A | 446904603 | 0.1310 | | 1.98E-06 | GEMMA | SUG |
| GL_17XX | AX-109920660 | 6A | 446933412 | 0.1305 | | 1.99E-06 | GEMMA | SUG |
| GL_17XX | AX-109901887 | 6A | 447115011 | -0.1346 | | 1.85E-06 | EMMAX | SUG |
| GL_17XX | AX-109901887 | 6A | 447115011 | 0.1340 | | 1.11E-06 | GEMMA | SUG |
| GL_17XX | AX-108878021 | 6A | 447118643 | 0.1310 | | 1.98E-06 | GEMMA | SUG |
| GL_17XX | AX-111564812 | 6A | 447122365 | -0.1334 | | 1.89E-06 | EMMAX | SUG |
| GL_17XX | AX-111564812 | 6A | 447122365 | 0.1327 | | 1.19E-06 | GEMMA | SUG |
| GL_17XX | AX-109876479 | 6A | 447161894 | 0.1310 | | 1.98E-06 | GEMMA | SUG |
| GL_17XX | AX-110375269 | 6A | 447188041 | 0.1347 | | 1.80E-06 | GEMMA | SUG |
| GL_17XX | AX-111025071 | 6A | 448053662 | -0.1385 | | 1.53E-06 | EMMAX | SUG |
| GL_17XX | AX-111025071 | 6A | 448053662 | 0.1371 | | 1.20E-06 | GEMMA | SUG |
| GL_17XX | AX-109513951 | 6B | 21804452 | 0.0523 | | 3.29E-06 | BLUPmrMLM | SUG |
| GL_17XX | AX-109343240 | 6B | 37046972 | 0.0590 | | 5.28E-05 | BLUPmrMLM | SUG |
| GL_17XX | AX-111633191 | 6B | 95490391 | -1.2906 | | 1.07E-10 | EMMAX | SIG |
| GL_17XX | AX-111472510 | 6B | 343787184 | 0.1935 | | 6.23E-07 | FarmCPU | SUG |
| GL_17XX | AX-110507415 | 6B | 352596924 | 0.1927 | | 7.92E-07 | FarmCPU | SUG |
| GL_17XX | AX-95197065 | 6B | 558925064 | 0.0657 | | 1.83E-06 | BLUPmrMLM | SUG |
| GL_17XX | AX-110627311 | 6B | 565704627 | 0.0624 | | 8.71E-05 | BLUPmrMLM | SUG |
| GL_17XX | AX-108739611 | 6B | 633895937 | -0.1977 | | 1.18E-07 | EMMAX | SUG |
| GL_17XX | AX-108739611 | 6B | 633895937 | 0.1944 | | 6.29E-08 | GEMMA | SIG |
| GL_17XX | AX-110509650 | 6B | 642922912 | -0.0764 | 1.9764 | 9.76E-19 | IIIVmrMLM | SIG |
| GL_17XX | AX-94965223 | 6D | 97015100 | 1.1690 | | 1.85E-06 | EMMAX | SUG |
| GL_17XX | AX-110454170 | 7A | 8375277 | 0.0068 | 1.9764 | 1.48E-10 | IIIVmrMLM | SIG |
| GL_17XX | AX-111823943 | 7A | 664700587 | 0.0482 | | 2.25E-05 | BLUPmrMLM | SUG |

| | | | | | | | | |
|---|---|---|---|---|---|---|---|---|
| GL_17XX | AX-108745065 | 7B | 84619559 | -0.0893 | -1.1087 | 2.30E-18 | IIIVmrMLM | SIG |
| GL_17XX | AX-94563183 | 7B | 354697105 | -0.3732 | | 1.13E-07 | EMMAX | SIG |
| GL_17XX | AX-94563183 | 7B | 354697105 | 0.3828 | | 1.26E-07 | GEMMA | SUG |
| GL_17XX | AX-94618304 | 7B | 470523833 | -0.6386 | | 2.04E-11 | EMMAX | SIG |
| GL_17XX | AX-94618304 | 7B | 470523833 | 0.6755 | | 2.03E-11 | GEMMA | SIG |
| GL_17XX | AX-111221981 | 7D | 2315652 | -0.8873 | | 1.44E-07 | EMMAX | SUG |
| GL_17XX | AX-110062884 | 7D | 20623856 | -0.5175 | | 1.78E-06 | EMMAX | SUG |
| GL_17XX | AX-94850331 | 7D | 31884381 | 0.4330 | | 5.97E-07 | GEMMA | SUG |
| GL_17XX | AX-109459815 | 7D | 392632007 | -1.0382 | | 3.21E-09 | EMMAX | SIG |
| GL_17XX | AX-110960135 | 7D | 486590501 | -2.8315 | | 1.58E-27 | EMMAX | SIG |
| GL_17XX | AX-111656854 | 7D | 597477552 | 1.2870 | | 3.35E-07 | EMMAX | SUG |
| GL_17ZMD | AX-110959647 | 1B | 593251091 | -0.1045 | 0.6908 | 4.74E-12 | IIIVmrMLM | SIG |
| GL_17ZMD | AX-108743703 | 2A | 299305162 | -0.0291 | 0.9846 | 3.13E-05 | IIIVmrMLM | SUG |
| GL_17ZMD | AX-110452207 | 2A | 539399868 | -0.1863 | | 1.61E-05 | BLUPmrMLM | sig |
| GL_17ZMD | AX-110465290 | 2B | 425955203 | 0.0189 | -0.5952 | 1.13E-05 | IIIVmrMLM | SUG |
| GL_17ZMD | AX-94869937 | 3D | 45803410 | -0.1756 | | 3.20E-16 | IIIVmrMLM | SIG |
| GL_17ZMD | AX-111516401 | 4A | 5831749 | 0.0833 | | 1.49E-04 | BLUPmrMLM | sig |
| GL_17ZMD | AX-110146053 | 4B | 421230568 | -0.0558 | 0.7462 | 5.64E-06 | IIIVmrMLM | SUG |
| GL_17ZMD | AX-108727418 | 5B | 405018149 | 0.2789 | | 9.83E-07 | FarmCPU | SUG |
| GL_17ZMD | AX-110567206 | 5B | 692632242 | 0.0795 | -0.3042 | 2.07E-06 | IIIVmrMLM | SUG |
| GL_17ZMD | AX-110373355 | 6A | 65145862 | 0.0086 | 0.5907 | 6.83E-05 | IIIVmrMLM | SUG |
| GL_17ZMD | AX-111062605 | 6A | 592289164 | -0.1409 | | 8.54E-06 | BLUPmrMLM | sig |
| GL_17ZMD | AX-108748193 | 7A | 703912060 | 0.0377 | -0.7392 | 1.19E-04 | IIIVmrMLM | SUG |
| GL_18AY | AX-94828051 | 1A | 94103812 | 0.1245 | | 1.86E-06 | BLUPmrMLM | SUG |
| GL_18AY | AX-94850477 | 1A | 108692813 | 0.1750 | | 4.13E-07 | GEMMA | SUG |
| GL_18AY | AX-110622929 | 1A | 540414611 | 0.0630 | | 9.73E-05 | BLUPmrMLM | SUG |
| GL_18AY | AX-111193032 | 1B | 4340360 | -0.1015 | | 2.40E-06 | BLUPmrMLM | SUG |
| GL_18AY | AX-109420526 | 1B | 29217823 | 0.1149 | | 8.64E-05 | BLUPmrMLM | SUG |
| GL_18AY | AX-94689485 | 1B | 92871645 | 0.1523 | | 1.80E-06 | GEMMA | SUG |
| GL_18AY | AX-111075801 | 1B | 95328846 | -0.1885 | | 1.61E-07 | FarmCPU | SUG |
| GL_18AY | AX-109309965 | 1B | 451543923 | -0.1352 | | 1.19E-06 | FarmCPU | SUG |
| GL_18AY | AX-94392216 | 1D | 11498625 | 0.2648 | | 8.44E-09 | FarmCPU | SIG |
| GL_18AY | AX-111800704 | 1D | 441090985 | 0.4120 | | 4.29E-09 | FarmCPU | SIG |
| GL_18AY | AX-109987438 | 2A | 4295985 | 0.0681 | | 2.49E-07 | BLUPmrMLM | SUG |
| GL_18AY | AX-109960770 | 2A | 142320921 | -0.0222 | 0.1737 | 3.12E-05 | IIIVmrMLM | SUG |
| GL_18AY | AX-111492478 | 2A | 715970302 | 0.0848 | | 5.01E-06 | BLUPmrMLM | SUG |
| GL_18AY | AX-109507603 | 2B | 99798122 | 0.0056 | -0.1286 | 1.71E-04 | IIIVmrMLM | SUG |
| GL_18AY | AX-111105137 | 2B | 117689246 | 0.0935 | | 1.90E-05 | BLUPmrMLM | SUG |
| GL_18AY | AX-111451431 | 2B | 205643210 | 0.0314 | 0.4916 | 5.16E-08 | IIIVmrMLM | SIG |
| GL_18AY | AX-94799956 | 2B | 532081152 | 0.0170 | 0.1444 | 1.89E-06 | IIIVmrMLM | SUG |
| GL_18AY | AX-94452919 | 2D | 77381522 | 0.1170 | 0.3025 | 2.02E-43 | IIIVmrMLM | SIG |
| GL_18AY | AX-109844715 | 2D | 151436768 | -0.0656 | | 6.85E-05 | BLUPmrMLM | SUG |
| GL_18AY | AX-110133132 | 2D | 604508871 | 0.0704 | | 1.14E-04 | BLUPmrMLM | SUG |
| GL_18AY | AX-110940849 | 3A | 532641574 | 0.0789 | | 4.40E-05 | BLUPmrMLM | SUG |

| | | | | | | | |
|---|---|---|---|---|---|---|---|
| GL_18AY | AX-109306485 | 3A | 716981865 | -0.0230 | -0.442 | 9.05E-07 IIIVmrMLM | SUG |
| GL_18AY | AX-111756784 | 3A | 719647215 | -0.0678 | -0.0319 | 4.95E-10 IIIVmrMLM | SIG |
| GL_18AY | AX-108969038 | 3B | 200466510 | -0.2051 | | 1.51E-06 FarmCPU | SUG |
| GL_18AY | AX-110038137 | 3B | 390111472 | 0.0346 | -0.1292 | 3.41E-09 IIIVmrMLM | SIG |
| GL_18AY | AX-111478013 | 3B | 466621685 | 0.0191 | -0.1578 | 1.37E-08 IIIVmrMLM | SIG |
| GL_18AY | AX-109479189 | 3B | 752269146 | -0.0802 | | 1.46E-06 BLUPmrMLM | SUG |
| GL_18AY | AX-109737295 | 3D | 185495307 | -0.3075 | | 2.05E-08 FarmCPU | SIG |
| GL_18AY | AX-109445819 | 4A | 123928604 | 0.0440 | -0.1738 | 1.21E-07 IIIVmrMLM | SUG |
| GL_18AY | AX-95202867 | 4A | 277889699 | -0.0923 | | 6.43E-07 BLUPmrMLM | SUG |
| GL_18AY | AX-94520475 | 4A | 597715406 | 0.1336 | | 2.76E-07 BLUPmrMLM | SUG |
| GL_18AY | AX-110477036 | 4A | 711398837 | 0.0019 | 0.3082 | 3.46E-05 IIIVmrMLM | SUG |
| GL_18AY | AX-110951049 | 4A | 725460139 | -0.1146 | -0.0055 | 1.10E-25 IIIVmrMLM | SIG |
| GL_18AY | AX-111135595 | 4B | 35175628 | 0.1359 | | 2.15E-11 FarmCPU | SIG |
| GL_18AY | AX-110452997 | 4B | 594261603 | -0.0400 | -0.3668 | 1.92E-08 IIIVmrMLM | SIG |
| GL_18AY | AX-94410603 | 5A | 5719660 | -0.1112 | -0.0324 | 7.67E-14 IIIVmrMLM | SIG |
| GL_18AY | AX-94987862 | 5A | 596369178 | 0.5085 | | 7.62E-07 GEMMA | SUG |
| GL_18AY | AX-109473519 | 5A | 622528405 | 0.1071 | | 2.23E-05 BLUPmrMLM | SUG |
| GL_18AY | AX-111250136 | 5B | 63739515 | 0.0837 | -0.0164 | 5.09E-12 IIIVmrMLM | SIG |
| GL_18AY | AX-110529773 | 5B | 86361024 | 0.0233 | -0.0886 | 1.99E-04 IIIVmrMLM | SUG |
| GL_18AY | AX-109327599 | 5B | 101151654 | -0.0121 | -0.154 | 8.58E-06 IIIVmrMLM | SUG |
| GL_18AY | AX-108931712 | 5B | 583475063 | 0.4691 | | 2.29E-06 GEMMA | SUG |
| GL_18AY | AX-111196036 | 5B | 584142374 | 0.4691 | | 2.29E-06 GEMMA | SUG |
| GL_18AY | AX-108833623 | 5B | 584796716 | 0.4691 | | 2.29E-06 GEMMA | SUG |
| GL_18AY | AX-110497421 | 5B | 665964553 | -0.0138 | -0.6614 | 2.15E-07 IIIVmrMLM | SUG |
| GL_18AY | AX-110555872 | 5B | 678708904 | 0.0448 | 0.1839 | 1.42E-16 IIIVmrMLM | SIG |
| GL_18AY | AX-110644337 | 5D | 284714723 | -0.0682 | -0.241 | 1.54E-16 IIIVmrMLM | SIG |
| GL_18AY | AX-110550202 | 6A | 153671397 | -0.0005 | 0.144 | 6.97E-04 IIIVmrMLM | SUG |
| GL_18AY | AX-86179015 | 6A | 437702676 | -0.0966 | -0.0534 | 2.31E-12 IIIVmrMLM | SIG |
| GL_18AY | AX-110703929 | 6A | 485164913 | -0.0128 | 0.5259 | 6.79E-10 IIIVmrMLM | SIG |
| GL_18AY | AX-109856815 | 6A | 564900753 | 0.0203 | -0.3211 | 1.33E-04 IIIVmrMLM | SUG |
| GL_18AY | AX-95239046 | 6A | 565467912 | 0.0205 | 0.1559 | 4.84E-15 IIIVmrMLM | SIG |
| GL_18AY | AX-108732926 | 6A | 600953819 | 0.0468 | | 7.84E-05 BLUPmrMLM | SUG |
| GL_18AY | AX-110147058 | 6B | 181990507 | -0.0797 | | 3.93E-15 IIIVmrMLM | SIG |
| GL_18AY | AX-109960324 | 6B | 609851458 | -0.0107 | -0.1864 | 4.46E-06 IIIVmrMLM | SUG |
| GL_18AY | AX-110509650 | 6B | 642922912 | -0.0166 | 1.5581 | 6.51E-10 IIIVmrMLM | SIG |
| GL_18AY | AX-110454170 | 7A | 8375277 | 0.0112 | 1.5581 | 1.32E-09 IIIVmrMLM | SIG |
| GL_18AY | AX-110957429 | 7A | 62754691 | -0.0831 | -0.2695 | 1.33E-27 IIIVmrMLM | SIG |
| GL_18AY | AX-110475510 | 7A | 204182749 | 0.1540 | | 4.87E-07 FarmCPU | SUG |
| GL_18AY | AX-111462234 | 7A | 700068738 | -0.0098 | 0.5413 | 2.94E-10 IIIVmrMLM | SIG |
| GL_18AY | AX-108963772 | 7B | 1022770 | -0.1132 | | 1.73E-06 FarmCPU | SUG |
| GL_18JY | AX-95227604 | 1A | 22288849 | 0.3064 | | 1.44E-06 GEMMA | SUG |
| GL_18JY | AX-110504088 | 1A | 468686364 | -0.0883 | 0.0221 | 1.64E-27 IIIVmrMLM | SIG |
| GL_18JY | AX-94710984 | 1B | 1421110 | -0.0251 | 0.1605 | 3.69E-14 IIIVmrMLM | SIG |
| GL_18JY | AX-111008033 | 1B | 15169796 | 0.1691 | | 4.04E-08 GEMMA | SIG |

| | | | | | | | | |
|---|---|---|---|---|---|---|---|---|
| GL_18JY | AX-110075386 | 1B | 34252695 | 0.1489 | | 2.20E-06 | GEMMA | SUG |
| GL_18JY | AX-111069652 | 1B | 36086820 | 0.1615 | | 8.28E-07 | GEMMA | SUG |
| GL_18JY | AX-94698017 | 1B | 86708972 | -0.3859 | | 9.81E-07 | EMMAX | SUG |
| GL_18JY | AX-94698017 | 1B | 86708972 | 0.3833 | | 2.95E-08 | GEMMA | SIG |
| GL_18JY | AX-111174986 | 1B | 119881507 | 0.3486 | | 1.66E-07 | GEMMA | SUG |
| GL_18JY | AX-111116469 | 1B | 143057215 | 0.3265 | | 2.34E-08 | GEMMA | SIG |
| GL_18JY | AX-94418929 | 1B | 157280398 | 0.2651 | | 2.30E-06 | GEMMA | SUG |
| GL_18JY | AX-108726211 | 1B | 171639599 | 0.0226 | 0.1705 | 2.64E-08 | IIIVmrMLM | SIG |
| GL_18JY | AX-94390293 | 1B | 176996634 | 0.3428 | | 2.31E-07 | GEMMA | SUG |
| GL_18JY | AX-108931371 | 1B | 185429686 | -0.2550 | | 9.91E-07 | EMMAX | SUG |
| GL_18JY | AX-108931371 | 1B | 185429686 | 0.2530 | | 8.41E-08 | GEMMA | SIG |
| GL_18JY | AX-111668617 | 1B | 191316160 | 0.3168 | | 1.34E-07 | GEMMA | SUG |
| GL_18JY | AX-95013113 | 1B | 212427766 | 0.2999 | | 3.78E-07 | GEMMA | SUG |
| GL_18JY | AX-111579663 | 1B | 426078236 | 0.0213 | 0.8825 | 2.05E-16 | IIIVmrMLM | SIG |
| GL_18JY | AX-94955220 | 1D | 42136124 | -0.0684 | | 1.44E-04 | BLUPmrMLM | SUG |
| GL_18JY | AX-94786159 | 1D | 79116398 | 0.3231 | | 3.14E-07 | GEMMA | SUG |
| GL_18JY | AX-111103449 | 1D | 133946396 | 0.1977 | | 1.29E-06 | GEMMA | SUG |
| GL_18JY | AX-109911526 | 2A | 721520986 | -0.0675 | | 3.80E-06 | BLUPmrMLM | SUG |
| GL_18JY | AX-109056359 | 2A | 750776356 | 0.0114 | -0.089 | 1.31E-07 | IIIVmrMLM | SUG |
| GL_18JY | AX-110978901 | 2A | 768728670 | 0.3811 | | 8.77E-07 | GEMMA | SUG |
| GL_18JY | AX-110987396 | 2A | 769221290 | 0.3749 | | 7.48E-07 | GEMMA | SUG |
| GL_18JY | AX-109435051 | 2B | 315096464 | -0.9853 | | 2.08E-10 | EMMAX | SIG |
| GL_18JY | AX-109284635 | 2B | 453269142 | -0.2710 | | 1.09E-07 | EMMAX | SIG |
| GL_18JY | AX-109284635 | 2B | 453269142 | 0.2741 | | 7.11E-08 | GEMMA | SIG |
| GL_18JY | AX-109392513 | 2B | 789601180 | -0.0649 | | 1.30E-05 | BLUPmrMLM | SUG |
| GL_18JY | AX-109854558 | 2B | 789873244 | 0.3429 | | 1.89E-06 | GEMMA | SUG |
| GL_18JY | AX-94985548 | 2B | 790278482 | 0.3429 | | 1.89E-06 | GEMMA | SUG |
| GL_18JY | AX-108821475 | 2B | 792989744 | 0.3429 | | 1.89E-06 | GEMMA | SUG |
| GL_18JY | AX-111587937 | 2B | 793047984 | 0.3429 | | 1.89E-06 | GEMMA | SUG |
| GL_18JY | AX-94516132 | 2B | 793168560 | 0.3429 | | 1.89E-06 | GEMMA | SUG |
| GL_18JY | AX-94697254 | 2D | 33710135 | 0.2393 | | 2.00E-06 | FarmCPU | SUG |
| GL_18JY | AX-109886994 | 2D | 548974432 | 0.0945 | 0.5325 | 2.78E-35 | IIIVmrMLM | SIG |
| GL_18JY | AX-94968179 | 2D | 574110628 | -0.0504 | | 4.49E-11 | IIIVmrMLM | SIG |
| GL_18JY | AX-95072346 | 2D | 586363215 | -0.1915 | | 2.08E-06 | GEMMA | SUG |
| GL_18JY | AX-109185488 | 2D | 642969692 | 0.0994 | 0.2588 | 2.94E-48 | IIIVmrMLM | SIG |
| GL_18JY | AX-94405846 | 2D | 644094043 | 0.3429 | | 1.89E-06 | GEMMA | SUG |
| GL_18JY | AX-111058596 | 2D | 644095171 | 0.3749 | | 7.48E-07 | GEMMA | SUG |
| GL_18JY | AX-94597784 | 3A | 13354919 | 0.1079 | | 2.40E-06 | BLUPmrMLM | SUG |
| GL_18JY | AX-111620802 | 3A | 15430673 | 0.0609 | | 3.14E-06 | BLUPmrMLM | SUG |
| GL_18JY | AX-109489505 | 3A | 38238882 | 0.6324 | | 1.38E-06 | FarmCPU | SUG |
| GL_18JY | AX-95151919 | 3A | 689498082 | 0.0497 | | 3.22E-05 | BLUPmrMLM | SUG |
| GL_18JY | AX-109274638 | 3B | 237325305 | -0.2404 | | 1.14E-06 | FarmCPU | SUG |
| GL_18JY | AX-111176610 | 3B | 440679552 | -0.9847 | | 1.45E-10 | EMMAX | SIG |
| GL_18JY | AX-95234844 | 3D | 185792116 | -1.4559 | | 1.05E-07 | EMMAX | SIG |

| | | | | | | | | |
|---|---|---|---|---|---|---|---|---|
| GL_18JY | AX-108933702 | 3D | 445322655 | -0.4660 | | 4.48E-10 | EMMAX | SIG |
| GL_18JY | AX-108933702 | 3D | 445322655 | 0.4931 | | 7.22E-10 | GEMMA | SIG |
| GL_18JY | AX-110634570 | 4A | 17784797 | -0.0428 | 0.0094 | 4.04E-06 | IIIVmrMLM | SUG |
| GL_18JY | AX-94677750 | 4A | 78285942 | 0.3427 | | 2.20E-06 | GEMMA | SUG |
| GL_18JY | AX-109305727 | 4A | 616674326 | 0.0508 | -0.0401 | 5.30E-09 | IIIVmrMLM | SIG |
| GL_18JY | AX-110933833 | 4A | 640836816 | 0.0520 | 0.4122 | 2.89E-20 | IIIVmrMLM | SIG |
| GL_18JY | AX-94728117 | 4B | 232103485 | -1.4559 | | 1.05E-07 | EMMAX | SIG |
| GL_18JY | AX-111590062 | 4B | 643333990 | -0.0097 | 0.7491 | 1.29E-04 | IIIVmrMLM | SUG |
| GL_18JY | AX-109414446 | 4B | 644796811 | -0.0163 | 1.9929 | 8.79E-25 | IIIVmrMLM | SIG |
| GL_18JY | AX-94558702 | 4B | 649603400 | -2.0437 | | 3.13E-10 | EMMAX | SIG |
| GL_18JY | AX-110622250 | 4D | 416766247 | -0.0384 | 0.094 | 6.85E-07 | IIIVmrMLM | SUG |
| GL_18JY | AX-110669567 | 5A | 595089097 | 0.0642 | | 1.48E-04 | BLUPmrMLM | SUG |
| GL_18JY | AX-94430561 | 5A | 596013453 | 0.4481 | | 1.79E-06 | GEMMA | SUG |
| GL_18JY | AX-111101649 | 5B | 185470339 | -0.0295 | 0.2731 | 1.20E-07 | IIIVmrMLM | SUG |
| GL_18JY | AX-109303205 | 5B | 223992784 | 0.0168 | 0.277 | 2.13E-13 | IIIVmrMLM | SIG |
| GL_18JY | AX-94826983 | 5B | 247255788 | -0.0633 | | 2.79E-16 | IIIVmrMLM | SIG |
| GL_18JY | AX-109038640 | 5B | 491066755 | -0.0304 | -0.1113 | 1.20E-15 | IIIVmrMLM | SIG |
| GL_18JY | AX-108931712 | 5B | 583475063 | 0.4692 | | 8.72E-08 | GEMMA | SIG |
| GL_18JY | AX-108946482 | 5B | 583616090 | 0.4481 | | 1.79E-06 | GEMMA | SUG |
| GL_18JY | AX-110627615 | 5B | 583714194 | 0.4481 | | 1.79E-06 | GEMMA | SUG |
| GL_18JY | AX-111742138 | 5B | 583911580 | 0.4481 | | 1.79E-06 | GEMMA | SUG |
| GL_18JY | AX-94973664 | 5B | 583913905 | 0.4481 | | 1.79E-06 | GEMMA | SUG |
| GL_18JY | AX-110447642 | 5B | 584068823 | 0.2240 | | 1.79E-06 | GEMMA | SUG |
| GL_18JY | AX-110402121 | 5B | 584068898 | 0.2246 | | 1.27E-06 | GEMMA | SUG |
| GL_18JY | AX-86180204 | 5B | 584069243 | 0.4335 | | 8.76E-07 | GEMMA | SUG |
| GL_18JY | AX-94457982 | 5B | 584070960 | 0.4481 | | 1.79E-06 | GEMMA | SUG |
| GL_18JY | AX-110740302 | 5B | 584108556 | 0.4481 | | 1.79E-06 | GEMMA | SUG |
| GL_18JY | AX-111675975 | 5B | 584109393 | 0.2240 | | 1.79E-06 | GEMMA | SUG |
| GL_18JY | AX-111019260 | 5B | 584110237 | 0.4481 | | 1.79E-06 | GEMMA | SUG |
| GL_18JY | AX-109879550 | 5B | 584110814 | 0.4481 | | 1.79E-06 | GEMMA | SUG |
| GL_18JY | AX-109529914 | 5B | 584111028 | 0.4481 | | 1.79E-06 | GEMMA | SUG |
| GL_18JY | AX-111196036 | 5B | 584142374 | 0.4692 | | 8.72E-08 | GEMMA | SIG |
| GL_18JY | AX-111542649 | 5B | 584582150 | 0.4481 | | 1.79E-06 | GEMMA | SUG |
| GL_18JY | AX-108945007 | 5B | 584616343 | 0.4481 | | 1.79E-06 | GEMMA | SUG |
| GL_18JY | AX-109907290 | 5B | 584639041 | 0.4481 | | 1.79E-06 | GEMMA | SUG |
| GL_18JY | AX-108916036 | 5B | 584644380 | 0.3174 | | 1.04E-07 | GEMMA | SIG |
| GL_18JY | AX-109373963 | 5B | 584796270 | 0.4481 | | 1.79E-06 | GEMMA | SUG |
| GL_18JY | AX-109443147 | 5B | 584796550 | 0.4481 | | 1.79E-06 | GEMMA | SUG |
| GL_18JY | AX-108833623 | 5B | 584796716 | 0.4692 | | 8.72E-08 | GEMMA | SIG |
| GL_18JY | AX-110904949 | 5B | 584825940 | 0.4421 | | 2.35E-07 | GEMMA | SUG |
| GL_18JY | AX-110269040 | 5D | 428345291 | 0.0685 | | 4.41E-06 | BLUPmrMLM | SUG |
| GL_18JY | AX-111177872 | 5D | 476671897 | 0.3955 | | 8.81E-07 | GEMMA | SUG |
| GL_18JY | AX-94497850 | 5D | 498186974 | -0.1277 | | 7.84E-05 | BLUPmrMLM | SUG |
| GL_18JY | AX-95188660 | 5D | 528764624 | 0.1811 | -0.0174 | 1.67E-13 | IIIVmrMLM | SIG |

| | | | | | | | | |
|---|---|---|---|---|---|---|---|---|
| GL_18JY | AX-110431858 | 6A | 26973059 | -0.0569 | | 4.03E-05 | BLUPmrMLM | SUG |
| GL_18JY | AX-108817049 | 6A | 416728745 | -0.0568 | -0.1042 | 3.55E-16 | IIIVmrMLM | SIG |
| GL_18JY | AX-110075161 | 6A | 452958151 | -0.0357 | -0.0047 | 3.42E-05 | IIIVmrMLM | SUG |
| GL_18JY | AX-94978834 | 6A | 594531897 | 0.0569 | | 1.28E-04 | BLUPmrMLM | SUG |
| GL_18JY | AX-94978834 | 6A | 594531897 | -0.0443 | 0.0259 | 2.03E-05 | IIIVmrMLM | SUG |
| GL_18JY | AX-111633191 | 6B | 95490391 | -1.5007 | | 1.77E-13 | EMMAX | SIG |
| GL_18JY | AX-111472510 | 6B | 343787184 | 0.1892 | | 2.24E-06 | FarmCPU | SUG |
| GL_18JY | AX-110166683 | 6B | 632989068 | 0.0640 | 0.0097 | 1.20E-10 | IIIVmrMLM | SIG |
| GL_18JY | AX-110409139 | 6D | 421889524 | 0.0497 | | 1.69E-04 | BLUPmrMLM | SUG |
| GL_18JY | AX-110454170 | 7A | 8375277 | 0.0107 | 4.1823 | 9.54E-82 | IIIVmrMLM | SIG |
| GL_18JY | AX-94498129 | 7A | 628867462 | -0.1192 | | 1.62E-04 | BLUPmrMLM | SUG |
| GL_18JY | AX-94842225 | 7A | 641317155 | 0.2642 | | 7.26E-07 | FarmCPU | SUG |
| GL_18JY | AX-109883719 | 7A | 690581796 | 0.0760 | | 2.24E-05 | BLUPmrMLM | SUG |
| GL_18JY | AX-111490522 | 7A | 707103478 | 0.3193 | | 1.30E-07 | FarmCPU | SUG |
| GL_18JY | AX-111635682 | 7B | 14763864 | -0.0450 | | 3.62E-09 | IIIVmrMLM | SIG |
| GL_18JY | AX-94563183 | 7B | 354697105 | -0.3265 | | 1.16E-06 | EMMAX | SUG |
| GL_18JY | AX-94563183 | 7B | 354697105 | 0.3337 | | 1.77E-06 | GEMMA | SUG |
| GL_18JY | AX-94618304 | 7B | 470523833 | -0.6035 | | 2.05E-11 | EMMAX | SIG |
| GL_18JY | AX-94618304 | 7B | 470523833 | 0.6319 | | 2.73E-11 | GEMMA | SIG |
| GL_18JY | AX-111221981 | 7D | 2315652 | -1.2578 | | 5.62E-13 | EMMAX | SIG |
| GL_18JY | AX-94850331 | 7D | 31884381 | 0.3069 | | 1.16E-06 | GEMMA | SUG |
| GL_18JY | AX-111124984 | 7D | 115543219 | 0.0525 | | 6.61E-07 | BLUPmrMLM | SUG |
| GL_18JY | AX-111100483 | 7D | 255225881 | 0.0355 | -0.2093 | 3.92E-10 | IIIVmrMLM | SIG |
| GL_18JY | AX-109459815 | 7D | 392632007 | -0.8175 | | 1.76E-06 | EMMAX | SUG |
| GL_18JY | AX-110960135 | 7D | 486590501 | -2.9674 | | 1.17E-31 | EMMAX | SIG |
| GL_18XX | AX-111519905 | 1A | 9974565 | -0.0478 | | 2.55E-06 | BLUPmrMLM | SUG |
| GL_18XX | AX-95173174 | 1B | 90993055 | 0.1609 | | 5.77E-07 | GEMMA | SUG |
| GL_18XX | AX-111116469 | 1B | 143057215 | -0.3759 | | 5.95E-07 | EMMAX | SUG |
| GL_18XX | AX-111116469 | 1B | 143057215 | 0.3892 | | 1.47E-08 | GEMMA | SIG |
| GL_18XX | AX-94390293 | 1B | 176996634 | 0.3915 | | 1.10E-06 | GEMMA | SUG |
| GL_18XX | AX-94488179 | 1B | 474450661 | -0.4384 | | 5.62E-10 | FarmCPU | SIG |
| GL_18XX | AX-94786159 | 1D | 79116398 | -0.3294 | | 7.23E-07 | EMMAX | SUG |
| GL_18XX | AX-94786159 | 1D | 79116398 | 0.3289 | | 9.00E-07 | GEMMA | SUG |
| GL_18XX | AX-110619791 | 2A | 68887889 | -0.0641 | | 4.42E-05 | BLUPmrMLM | SUG |
| GL_18XX | AX-111739920 | 2B | 159096863 | -0.0398 | | 1.92E-05 | BLUPmrMLM | SUG |
| GL_18XX | AX-109435051 | 2B | 315096464 | -0.7817 | | 4.30E-09 | EMMAX | SIG |
| GL_18XX | AX-111439519 | 3D | 8530236 | 0.1410 | | 3.78E-07 | FarmCPU | SUG |
| GL_18XX | AX-95234844 | 3D | 185792116 | -1.4994 | | 8.58E-10 | EMMAX | SIG |
| GL_18XX | AX-94709485 | 3D | 361376741 | 0.1189 | | 1.89E-06 | BLUPmrMLM | SUG |
| GL_18XX | AX-108933702 | 3D | 445322655 | -0.3516 | | 3.06E-07 | EMMAX | SUG |
| GL_18XX | AX-108933702 | 3D | 445322655 | 0.3624 | | 2.77E-07 | GEMMA | SUG |
| GL_18XX | AX-94429052 | 4A | 2794203 | -0.4517 | | 4.23E-06 | BLUPmrMLM | SUG |
| GL_18XX | AX-109509823 | 4A | 54984149 | -0.2989 | | 1.25E-06 | EMMAX | SUG |
| GL_18XX | AX-109509823 | 4A | 54984149 | 0.2976 | | 1.34E-06 | GEMMA | SUG |

| | | | | | | | | |
|---|---|---|---|---|---|---|---|---|
| GL_18XX | AX-108791521 | 4B | 10003294 | -0.1184 | | 4.96E-10 | FarmCPU | SIG |
| GL_18XX | AX-94728117 | 4B | 232103485 | -1.4994 | | 8.58E-10 | EMMAX | SIG |
| GL_18XX | AX-111483036 | 4B | 581503751 | 0.0595 | | 2.50E-05 | BLUPmrMLM | SUG |
| GL_18XX | AX-109013070 | 4B | 667887477 | -0.0471 | | 3.46E-06 | BLUPmrMLM | SUG |
| GL_18XX | AX-108957073 | 4D | 418943220 | 0.2512 | | 1.84E-07 | FarmCPU | SUG |
| GL_18XX | AX-94653997 | 4D | 485522383 | -0.1511 | | 6.51E-07 | FarmCPU | SUG |
| GL_18XX | AX-110050904 | 5A | 34327685 | 0.0408 | | 1.28E-04 | BLUPmrMLM | SUG |
| GL_18XX | AX-111592289 | 5A | 532449987 | 0.0900 | | 1.08E-09 | BLUPmrMLM | SIG |
| GL_18XX | AX-111482807 | 5A | 576373436 | 0.0638 | | 1.21E-07 | BLUPmrMLM | SUG |
| GL_18XX | AX-110669567 | 5A | 595089097 | 0.0533 | | 3.10E-05 | BLUPmrMLM | SUG |
| GL_18XX | AX-108869061 | 5A | 631632727 | 0.0843 | | 6.91E-07 | BLUPmrMLM | SUG |
| GL_18XX | AX-111592014 | 5B | 50502310 | 0.0829 | | 7.30E-09 | BLUPmrMLM | SIG |
| GL_18XX | AX-94497850 | 5D | 498186974 | -0.0927 | | 9.34E-05 | BLUPmrMLM | SUG |
| GL_18XX | AX-95079914 | 5D | 551826420 | -0.0764 | | 9.85E-06 | BLUPmrMLM | SUG |
| GL_18XX | AX-111572602 | 6A | 2293409 | 0.0418 | | 8.47E-05 | BLUPmrMLM | SUG |
| GL_18XX | AX-111633191 | 6B | 95490391 | -0.9727 | | 1.81E-08 | EMMAX | SIG |
| GL_18XX | AX-109856732 | 6B | 476031162 | -0.0843 | | 1.47E-10 | BLUPmrMLM | SIG |
| GL_18XX | AX-110933813 | 6B | 486229851 | 0.0939 | | 6.29E-11 | BLUPmrMLM | SIG |
| GL_18XX | AX-110166683 | 6B | 632989068 | -0.0999 | | 7.42E-08 | BLUPmrMLM | SIG |
| GL_18XX | AX-110509650 | 6B | 642922912 | -0.0719 | 1.5822 | 2.53E-17 | IIIVmrMLM | SIG |
| GL_18XX | AX-108948736 | 6D | 85202080 | 0.2185 | | 6.09E-08 | FarmCPU | SIG |
| GL_18XX | AX-110454170 | 7A | 8375277 | 0.0253 | 1.5822 | 1.91E-09 | IIIVmrMLM | SIG |
| GL_18XX | AX-111566293 | 7A | 10802325 | 0.0883 | | 3.11E-07 | BLUPmrMLM | SUG |
| GL_18XX | AX-110540637 | 7A | 670832511 | 0.0310 | | 7.79E-05 | BLUPmrMLM | SUG |
| GL_18XX | AX-94618304 | 7B | 470523833 | -0.4628 | | 3.00E-08 | EMMAX | SIG |
| GL_18XX | AX-94618304 | 7B | 470523833 | 0.4752 | | 2.80E-08 | GEMMA | SIG |
| GL_18XX | AX-109391485 | 7B | 693307347 | -0.0679 | | 2.14E-10 | BLUPmrMLM | SIG |
| GL_18XX | AX-111221981 | 7D | 2315652 | -0.8465 | | 7.09E-09 | EMMAX | SIG |
| GL_18XX | AX-94850331 | 7D | 31884381 | -0.9906 | -0.6272 | 0.00E+00 | IIIVmrMLM | SIG |
| GL_18XX | AX-110898194 | 7D | 168777588 | 0.0491 | | 1.22E-04 | BLUPmrMLM | SUG |
| GL_18XX | AX-94526971 | 7D | 387335431 | 0.7259 | | 2.35E-09 | FarmCPU | SIG |
| GL_18XX | AX-110960135 | 7D | 486590501 | -2.3032 | | 4.53E-24 | EMMAX | SIG |
| GL_18ZMD | AX-108895113 | 1B | 40660487 | 0.1769 | | 1.23E-06 | EMMAX | SUG |
| GL_18ZMD | AX-108895113 | 1B | 40660487 | -0.1802 | | 2.60E-07 | GEMMA | SUG |
| GL_18ZMD | AX-109868465 | 1B | 40724908 | 0.1734 | | 1.99E-06 | EMMAX | SUG |
| GL_18ZMD | AX-109868465 | 1B | 40724908 | -0.1769 | | 4.30E-07 | GEMMA | SUG |
| GL_18ZMD | AX-108947433 | 1B | 40807024 | 0.3601 | | 6.96E-07 | EMMAX | SUG |
| GL_18ZMD | AX-108947433 | 1B | 40807024 | -0.3662 | | 1.43E-07 | GEMMA | SUG |
| GL_18ZMD | AX-109343600 | 1B | 40809362 | 0.1792 | | 8.68E-07 | EMMAX | SUG |
| GL_18ZMD | AX-109343600 | 1B | 40809362 | -0.1824 | | 1.81E-07 | GEMMA | SUG |
| GL_18ZMD | AX-94965878 | 1B | 40878719 | 0.1726 | | 2.10E-06 | EMMAX | SUG |
| GL_18ZMD | AX-94965878 | 1B | 40878719 | -0.1761 | | 4.55E-07 | GEMMA | SUG |
| GL_18ZMD | AX-111648440 | 1B | 41085538 | -0.1621 | | 2.22E-06 | GEMMA | SUG |
| GL_18ZMD | AX-110373430 | 1B | 41095679 | 0.3591 | | 3.26E-07 | EMMAX | SUG |

| | | | | | | | | |
|---|---|---|---|---|---|---|---|---|
| GL_18ZMD | AX-110373430 | 1B | 41095679 | -0.3649 | | 6.45E-08 | GEMMA | SIG |
| GL_18ZMD | AX-109283974 | 3A | 648222853 | -0.0958 | | 1.64E-04 | BLUPmrMLM | SUG |
| GL_18ZMD | AX-109839043 | 3B | 226484519 | -0.2850 | | 8.25E-07 | FarmCPU | SUG |
| GL_18ZMD | AX-109274631 | 3B | 556050943 | 0.0863 | | 6.31E-05 | BLUPmrMLM | SUG |
| GL_18ZMD | AX-111501984 | 5B | 614983305 | 0.3023 | | 3.27E-07 | FarmCPU | SUG |
| GL_18ZMD | AX-109516013 | 5D | 209826171 | -0.1399 | | 4.23E-05 | BLUPmrMLM | SUG |
| GL_18ZMD | AX-109792794 | 5D | 398425491 | -0.3811 | | 6.34E-07 | EMMAX | SUG |
| GL_18ZMD | AX-109792794 | 5D | 398425491 | 0.3748 | | 2.55E-07 | GEMMA | SUG |
| GL_18ZMD | AX-94755484 | 6B | 41920027 | -0.2984 | | 1.01E-04 | BLUPmrMLM | SUG |
| GL_18ZMD | AX-111153914 | 7A | 507877239 | 0.3003 | | 1.71E-06 | FarmCPU | SUG |
| GL_18ZMD | AX-95017844 | 7B | 568488282 | -0.0706 | | 1.47E-04 | BLUPmrMLM | SUG |
| GL_19XX | AX-108968207 | 1A | 532777479 | -0.2147 | -0.4671 | 3.25E-95 | IIIVmrMLM | SIG |
| GL_19XX | AX-94817273 | 1A | 586512733 | 0.0594 | | 4.98E-06 | BLUPmrMLM | SUG |
| GL_19XX | AX-111019319 | 1B | 37797055 | -0.0789 | 1.1411 | 1.35E-18 | IIIVmrMLM | SIG |
| GL_19XX | AX-109329289 | 1B | 90544558 | 0.1728 | | 5.75E-07 | GEMMA | SUG |
| GL_19XX | AX-95173174 | 1B | 90993055 | 0.1460 | | 1.12E-06 | GEMMA | SUG |
| GL_19XX | AX-111381972 | 1D | 235027073 | 0.2923 | 0.582 | 2.55E-129 | IIIVmrMLM | SIG |
| GL_19XX | AX-110987396 | 2A | 769221290 | 0.3664 | | 1.92E-06 | GEMMA | SUG |
| GL_19XX | AX-108858584 | 2B | 750007634 | 0.1487 | | 5.24E-07 | GEMMA | SUG |
| GL_19XX | AX-95150022 | 2B | 761960730 | -0.0564 | | 8.32E-06 | BLUPmrMLM | SUG |
| GL_19XX | AX-94804217 | 2D | 8478708 | -0.0447 | | 9.73E-05 | BLUPmrMLM | SUG |
| GL_19XX | AX-109590718 | 2D | 81836279 | 0.0140 | -0.7777 | 7.09E-10 | IIIVmrMLM | SIG |
| GL_19XX | AX-111058596 | 2D | 644095171 | 0.3664 | | 1.92E-06 | GEMMA | SUG |
| GL_19XX | AX-108871202 | 3A | 683730766 | -0.0344 | | 1.93E-04 | BLUPmrMLM | SUG |
| GL_19XX | AX-109444679 | 3B | 140491453 | 0.0327 | | 5.73E-05 | BLUPmrMLM | SUG |
| GL_19XX | AX-109561781 | 4A | 100558685 | 0.0576 | | 2.03E-06 | BLUPmrMLM | SUG |
| GL_19XX | AX-112287420 | 4B | 623022977 | -0.0469 | | 9.38E-07 | BLUPmrMLM | SUG |
| GL_19XX | AX-108894317 | 4B | 664057292 | -0.0640 | | 1.25E-08 | BLUPmrMLM | SIG |
| GL_19XX | AX-111523027 | 5A | 631665321 | -0.0514 | | 4.05E-06 | BLUPmrMLM | SUG |
| GL_19XX | AX-95149626 | 5B | 63362235 | 0.5364 | | 1.29E-08 | BLUPmrMLM | SIG |
| GL_19XX | AX-110004271 | 5D | 13987424 | 0.3436 | | 2.00E-06 | GEMMA | SUG |
| GL_19XX | AX-95188660 | 5D | 528764624 | -0.1663 | | 1.43E-07 | BLUPmrMLM | SUG |
| GL_19XX | AX-110428682 | 6A | 13674435 | -0.0401 | | 3.88E-05 | BLUPmrMLM | SUG |
| GL_19XX | AX-109868965 | 6A | 17661065 | 0.0430 | | 2.86E-05 | BLUPmrMLM | SUG |
| GL_19XX | AX-111472510 | 6B | 343787184 | 0.1656 | | 1.59E-06 | FarmCPU | SUG |
| GL_19XX | AX-110509650 | 6B | 642922912 | -0.0386 | 0.814 | 1.73E-06 | IIIVmrMLM | SUG |
| GL_19XX | AX-109432737 | 6B | 683151377 | -0.0378 | | 5.28E-06 | BLUPmrMLM | SUG |
| GL_19XX | AX-111188882 | 6B | 696132549 | 0.0450 | | 1.08E-05 | BLUPmrMLM | SUG |
| GL_19XX | AX-110570445 | 7A | 568913192 | -0.0814 | | 3.52E-09 | BLUPmrMLM | SIG |
| GL_19XX | AX-89376898 | 7B | 9801996 | -0.0450 | | 1.97E-05 | BLUPmrMLM | SUG |
| GL_19XX | AX-110122829 | 7B | 128824116 | -0.0962 | | 1.61E-06 | BLUPmrMLM | SUG |
| GL_19XX | AX-94526971 | 7D | 387335431 | 0.7954 | | 2.55E-07 | FarmCPU | SUG |
| GL_19XX | AX-111835948 | 7D | 400486341 | -0.0603 | | 1.10E-05 | BLUPmrMLM | SUG |
| GL_19XX | AX-110670911 | 7D | 608863909 | -0.1374 | -0.2462 | 8.58E-52 | IIIVmrMLM | SIG |

| | | | | | | | | |
|---|---|---|---|---|---|---|---|---|
| GL_BLUP | AX-109948031 | 1A | 53187171 | -0.0886 | | 1.51E-05 | BLUPmrMLM | SUG |
| GL_BLUP | AX-109640257 | 1A | 564781279 | 0.0676 | | 2.11E-06 | FarmCPU | SUG |
| GL_BLUP | AX-111116469 | 1B | 143057215 | 0.2562 | | 1.41E-07 | GEMMA | SUG |
| GL_BLUP | AX-108931371 | 1B | 185429686 | 0.1595 | | 9.49E-07 | GEMMA | SUG |
| GL_BLUP | AX-95681612 | 1D | 48401161 | 0.1570 | | 1.64E-06 | FarmCPU | SUG |
| GL_BLUP | AX-94786159 | 1D | 79116398 | -0.2525 | | 8.91E-08 | EMMAX | SIG |
| GL_BLUP | AX-94786159 | 1D | 79116398 | 0.2512 | | 1.40E-07 | GEMMA | SUG |
| GL_BLUP | AX-109849099 | 2A | 703661456 | 0.0475 | | 2.17E-07 | BLUPmrMLM | SUG |
| GL_BLUP | AX-109435051 | 2B | 315096464 | -0.5742 | | 1.18E-09 | EMMAX | SIG |
| GL_BLUP | AX-109016211 | 2D | 164285537 | -0.7851 | | 1.20E-06 | EMMAX | SUG |
| GL_BLUP | AX-109400512 | 2D | 529605010 | 0.0611 | | 3.47E-07 | BLUPmrMLM | SUG |
| GL_BLUP | AX-95118616 | 2D | 544781641 | 0.1089 | | 9.61E-07 | BLUPmrMLM | SUG |
| GL_BLUP | AX-111929073 | 2D | 571885425 | 0.1320 | | 6.91E-08 | FarmCPU | SIG |
| GL_BLUP | AX-89565232 | 3A | 556354785 | -0.0450 | | 6.23E-07 | BLUPmrMLM | SUG |
| GL_BLUP | AX-110448131 | 3B | 225261015 | -0.1524 | | 5.72E-10 | FarmCPU | SIG |
| GL_BLUP | AX-111176610 | 3B | 440679552 | -0.5434 | | 1.37E-08 | EMMAX | SIG |
| GL_BLUP | AX-111567612 | 3B | 494527545 | -0.0405 | | 7.38E-06 | BLUPmrMLM | SUG |
| GL_BLUP | AX-95234844 | 3D | 185792116 | -1.0778 | | 5.55E-10 | EMMAX | SIG |
| GL_BLUP | AX-111193837 | 3D | 238620709 | -0.1842 | | 2.62E-08 | FarmCPU | SIG |
| GL_BLUP | AX-108933702 | 3D | 445322655 | -0.2948 | | 1.36E-09 | EMMAX | SIG |
| GL_BLUP | AX-108933702 | 3D | 445322655 | 0.3080 | | 1.23E-09 | GEMMA | SIG |
| GL_BLUP | AX-111439128 | 3D | 572240905 | -0.0343 | | 1.96E-07 | BLUPmrMLM | SUG |
| GL_BLUP | AX-109509823 | 4A | 54984149 | -0.2230 | | 3.58E-07 | EMMAX | SUG |
| GL_BLUP | AX-109509823 | 4A | 54984149 | 0.2219 | | 4.84E-07 | GEMMA | SUG |
| GL_BLUP | AX-109561781 | 4A | 100558685 | 0.0339 | | 1.15E-04 | BLUPmrMLM | SUG |
| GL_BLUP | AX-111519175 | 4A | 590942121 | 0.0256 | | 2.75E-05 | BLUPmrMLM | SUG |
| GL_BLUP | AX-95134639 | 4A | 726299982 | 0.0441 | | 2.68E-08 | BLUPmrMLM | SIG |
| GL_BLUP | AX-111467178 | 4B | 13947273 | -0.0266 | | 5.01E-05 | BLUPmrMLM | SUG |
| GL_BLUP | AX-94728117 | 4B | 232103485 | -1.0778 | | 5.55E-10 | EMMAX | SIG |
| GL_BLUP | AX-111483036 | 4B | 581503751 | 0.0382 | | 2.65E-05 | BLUPmrMLM | SUG |
| GL_BLUP | AX-86179393 | 5A | 610217061 | -0.0358 | | 1.23E-06 | BLUPmrMLM | SUG |
| GL_BLUP | AX-94837642 | 5B | 566500338 | 0.0200 | | 1.16E-04 | BLUPmrMLM | SUG |
| GL_BLUP | AX-109039546 | 5B | 694314697 | 0.0301 | | 8.16E-06 | BLUPmrMLM | SUG |
| GL_BLUP | AX-95023275 | 6A | 417443293 | 0.9566 | | 1.33E-12 | FarmCPU | SIG |
| GL_BLUP | AX-111633191 | 6B | 95490391 | -0.7381 | | 1.60E-09 | EMMAX | SIG |
| GL_BLUP | AX-110166683 | 6B | 632989068 | -0.0555 | | 8.74E-06 | BLUPmrMLM | SUG |
| GL_BLUP | AX-108739611 | 6B | 633895937 | -0.1233 | | 8.05E-08 | EMMAX | SIG |
| GL_BLUP | AX-108739611 | 6B | 633895937 | 0.1219 | | 4.65E-08 | GEMMA | SIG |
| GL_BLUP | AX-110509650 | 6B | 642922912 | -0.0638 | 1.1439 | 2.83E-20 | IIIVmrMLM | SIG |
| GL_BLUP | AX-111370974 | 6D | 137893383 | -0.0667 | | 9.63E-09 | BLUPmrMLM | SIG |
| GL_BLUP | AX-108995772 | 6D | 142860586 | -0.0648 | | 1.39E-06 | FarmCPU | SUG |
| GL_BLUP | AX-109065407 | 6D | 256578442 | -0.1491 | | 1.62E-06 | FarmCPU | SUG |
| GL_BLUP | AX-110454170 | 7A | 8375277 | 0.0224 | 1.1439 | 2.28E-09 | IIIVmrMLM | SIG |
| GL_BLUP | AX-109402351 | 7A | 203527622 | -0.0320 | | 7.68E-06 | BLUPmrMLM | SUG |

| | | | | | | | |
|---|---|---|---|---|---|---|---|
| GL_BLUP | AX-109838679 | 7A | 327307051 | -0.0288 | 1.01E-04 | BLUPmrMLM | SUG |
| GL_BLUP | AX-94563183 | 7B | 354697105 | -0.2272 | 9.10E-08 | EMMAX | SIG |
| GL_BLUP | AX-94563183 | 7B | 354697105 | 0.2312 | 1.11E-07 | GEMMA | SIG |
| GL_BLUP | AX-109335081 | 7B | 397233152 | -0.1774 | 2.22E-06 | EMMAX | SUG |
| GL_BLUP | AX-109335081 | 7B | 397233152 | 0.1849 | 2.20E-06 | GEMMA | SUG |
| GL_BLUP | AX-94618304 | 7B | 470523833 | -0.3612 | 1.08E-09 | EMMAX | SIG |
| GL_BLUP | AX-94618304 | 7B | 470523833 | 0.3668 | 7.41E-10 | GEMMA | SIG |
| GL_BLUP | AX-95017844 | 7B | 568488282 | -0.0362 | 1.20E-05 | BLUPmrMLM | SUG |
| GL_BLUP | AX-111221981 | 7D | 2315652 | -0.5861 | 1.77E-08 | EMMAX | SIG |
| GL_BLUP | AX-110062884 | 7D | 20623856 | -0.3188 | 1.84E-06 | EMMAX | SUG |
| GL_BLUP | AX-110888456 | 7D | 41407631 | 0.0228 | 9.67E-05 | BLUPmrMLM | SUG |
| GL_BLUP | AX-109303704 | 7D | 52947227 | -0.0858 | 4.24E-07 | FarmCPU | SUG |
| GL_BLUP | AX-110960135 | 7D | 486590501 | -1.6838 | 1.67E-25 | EMMAX | SIG |

GL: grain length.

**Supplementary Table S10**. All QTNs identified as associated with grain width in wheat

| Trait | Marker | Chr | Position | Add.effect | Dom.effect | P-value | Method | Sig. |
|---|---|---|---|---|---|---|---|---|
| GW_17AY | AX-108780402 | 1A | 564570992 | 0.1377 | | 6.33E-08 | FarmCPU | SIG |
| GW_17AY | AX-109427328 | 1A | 564586780 | 0.1829 | | 1.21E-06 | FarmCPU | SUG |
| GW_17AY | AX-111610087 | 1A | 564865778 | -0.1071 | | 1.07E-06 | FarmCPU | SUG |
| GW_17AY | AX-94966528 | 1D | 392147568 | -0.0771 | | 1.60E-04 | BLUPmrMLM | SIG |
| GW_17AY | AX-108764267 | 2A | 754034071 | 0.0518 | | 4.19E-05 | BLUPmrMLM | SIG |
| GW_17AY | AX-110600488 | 4A | 352261301 | -0.0495 | | 1.07E-04 | BLUPmrMLM | SIG |
| GW_17AY | AX-111662946 | 6B | 714127224 | 0.0661 | | 1.13E-06 | BLUPmrMLM | SIG |
| GW_17AY | AX-109539083 | 7A | 675502040 | -0.0360 | | 8.93E-05 | BLUPmrMLM | SIG |
| GW_17AY | AX-110194437 | 7B | 742435904 | -0.0328 | | 9.89E-05 | BLUPmrMLM | SIG |
| GW_17SQ | AX-111612789 | 1A | 11891744 | -0.0246 | | 2.47E-05 | BLUPmrMLM | SUG |
| GW_17SQ | AX-111578899 | 1A | 104739832 | -0.0089 | 0.0487 | 4.05E-06 | IIIVmrMLM | SUG |
| GW_17SQ | AX-110420370 | 1A | 489413809 | 0.0651 | | 1.64E-06 | GEMMA | SUG |
| GW_17SQ | AX-111021731 | 1A | 492713762 | 0.0656 | | 1.49E-06 | GEMMA | SUG |
| GW_17SQ | AX-109447934 | 1A | 492766573 | 0.1285 | | 1.77E-06 | GEMMA | SUG |
| GW_17SQ | AX-110990331 | 1A | 492780215 | 0.1285 | | 1.77E-06 | GEMMA | SUG |
| GW_17SQ | AX-109423019 | 1A | 493204191 | 0.0657 | | 1.88E-06 | GEMMA | SUG |
| GW_17SQ | AX-111521837 | 1A | 493218248 | -0.0693 | | 1.94E-06 | EMMAX | SUG |
| GW_17SQ | AX-111521837 | 1A | 493218248 | 0.0690 | | 6.37E-07 | GEMMA | SUG |
| GW_17SQ | AX-108725777 | 1A | 493219294 | 0.0706 | | 4.13E-07 | GEMMA | SUG |
| GW_17SQ | AX-109396023 | 1A | 493220926 | -0.0652 | | 1.35E-06 | EMMAX | SUG |
| GW_17SQ | AX-109396023 | 1A | 493220926 | 0.0658 | | 2.93E-07 | GEMMA | SUG |
| GW_17SQ | AX-110635453 | 1A | 493513970 | -0.0826 | | 4.80E-07 | EMMAX | SUG |
| GW_17SQ | AX-110635453 | 1A | 493513970 | 0.0819 | | 9.44E-08 | GEMMA | SIG |
| GW_17SQ | AX-109422899 | 1A | 493527588 | -0.0811 | | 8.37E-07 | EMMAX | SUG |
| GW_17SQ | AX-109422899 | 1A | 493527588 | 0.0805 | | 1.55E-07 | GEMMA | SUG |
| GW_17SQ | AX-111009245 | 1A | 493546679 | -0.1760 | | 5.95E-08 | EMMAX | SIG |
| GW_17SQ | AX-111009245 | 1A | 493546679 | 0.1744 | | 1.09E-08 | GEMMA | SIG |
| GW_17SQ | AX-110134130 | 1A | 493579083 | -0.0843 | | 3.29E-07 | EMMAX | SUG |
| GW_17SQ | AX-110134130 | 1A | 493579083 | 0.0834 | | 6.22E-08 | GEMMA | SIG |
| GW_17SQ | AX-108789421 | 1A | 493581377 | -0.0818 | | 8.08E-07 | EMMAX | SUG |
| GW_17SQ | AX-108789421 | 1A | 493581377 | 0.0813 | | 1.45E-07 | GEMMA | SUG |
| GW_17SQ | AX-110409474 | 1A | 493581540 | -0.0843 | | 3.29E-07 | EMMAX | SUG |
| GW_17SQ | AX-110409474 | 1A | 493581540 | 0.0834 | | 6.22E-08 | GEMMA | SIG |
| GW_17SQ | AX-111061283 | 1A | 493582041 | -0.0843 | | 3.29E-07 | EMMAX | SUG |
| GW_17SQ | AX-111061283 | 1A | 493582041 | 0.0834 | | 6.22E-08 | GEMMA | SIG |
| GW_17SQ | AX-110513757 | 1A | 493582085 | -0.0806 | | 9.49E-07 | EMMAX | SUG |
| GW_17SQ | AX-110513757 | 1A | 493582085 | 0.0803 | | 1.47E-07 | GEMMA | SUG |
| GW_17SQ | AX-109404268 | 1A | 493582117 | -0.0826 | | 6.92E-07 | EMMAX | SUG |
| GW_17SQ | AX-109404268 | 1A | 493582117 | 0.0819 | | 1.28E-07 | GEMMA | SUG |
| GW_17SQ | AX-89438125 | 1A | 493585276 | -0.0843 | | 3.29E-07 | EMMAX | SUG |
| GW_17SQ | AX-89438125 | 1A | 493585276 | 0.0834 | | 6.22E-08 | GEMMA | SIG |

| | | | | | | | |
|---|---|---|---|---|---|---|---|
| GW_17SQ | AX-109318573 | 1A | 493586035 | -0.0838 | 2.62E-07 | EMMAX | SUG |
| GW_17SQ | AX-109318573 | 1A | 493586035 | 0.0835 | 3.40E-08 | GEMMA | SIG |
| GW_17SQ | AX-109448378 | 1A | 493586243 | -0.0831 | 4.39E-07 | EMMAX | SUG |
| GW_17SQ | AX-109448378 | 1A | 493586243 | 0.0823 | 8.29E-08 | GEMMA | SIG |
| GW_17SQ | AX-110414710 | 1A | 493588655 | 0.0742 | 1.08E-06 | GEMMA | SUG |
| GW_17SQ | AX-111113867 | 1A | 493629274 | -0.0861 | 1.66E-07 | EMMAX | SUG |
| GW_17SQ | AX-111113867 | 1A | 493629274 | 0.0852 | 3.27E-08 | GEMMA | SIG |
| GW_17SQ | AX-111013613 | 1A | 493629438 | -0.0869 | 1.02E-07 | EMMAX | SIG |
| GW_17SQ | AX-111013613 | 1A | 493629438 | 0.0860 | 1.86E-08 | GEMMA | SIG |
| GW_17SQ | AX-110436018 | 1A | 493630508 | -0.0836 | 4.11E-07 | EMMAX | SUG |
| GW_17SQ | AX-110436018 | 1A | 493630508 | 0.0830 | 7.29E-08 | GEMMA | SIG |
| GW_17SQ | AX-110406533 | 1A | 493663494 | -0.0837 | 3.44E-07 | EMMAX | SUG |
| GW_17SQ | AX-110406533 | 1A | 493663494 | 0.0829 | 6.91E-08 | GEMMA | SIG |
| GW_17SQ | AX-109937131 | 1A | 493664011 | 0.0712 | 1.33E-06 | GEMMA | SUG |
| GW_17SQ | AX-110433017 | 1A | 493664714 | -0.0777 | 9.47E-08 | EMMAX | SIG |
| GW_17SQ | AX-110433017 | 1A | 493664714 | 0.0773 | 2.52E-08 | GEMMA | SIG |
| GW_17SQ | AX-108762436 | 1A | 493665599 | -0.0867 | 1.58E-07 | EMMAX | SUG |
| GW_17SQ | AX-108762436 | 1A | 493665599 | 0.0857 | 3.14E-08 | GEMMA | SIG |
| GW_17SQ | AX-109818522 | 1A | 493666056 | -0.0870 | 1.50E-07 | EMMAX | SUG |
| GW_17SQ | AX-109818522 | 1A | 493666056 | 0.0862 | 3.18E-08 | GEMMA | SIG |
| GW_17SQ | AX-108956416 | 1A | 493666566 | -0.0791 | 1.60E-06 | EMMAX | SUG |
| GW_17SQ | AX-108956416 | 1A | 493666566 | 0.0787 | 3.07E-07 | GEMMA | SUG |
| GW_17SQ | AX-110960642 | 1A | 493734728 | -0.0867 | 1.58E-07 | EMMAX | SUG |
| GW_17SQ | AX-110960642 | 1A | 493734728 | 0.0857 | 3.14E-08 | GEMMA | SIG |
| GW_17SQ | AX-111637826 | 1A | 493734853 | -0.0812 | 8.58E-07 | EMMAX | SUG |
| GW_17SQ | AX-111637826 | 1A | 493734853 | 0.0805 | 1.67E-07 | GEMMA | SUG |
| GW_17SQ | AX-111085923 | 1A | 493735347 | -0.0842 | 3.97E-07 | EMMAX | SUG |
| GW_17SQ | AX-111085923 | 1A | 493735347 | 0.0836 | 7.46E-08 | GEMMA | SIG |
| GW_17SQ | AX-108875380 | 1A | 493735495 | -0.0862 | 1.66E-07 | EMMAX | SUG |
| GW_17SQ | AX-108875380 | 1A | 493735495 | 0.0852 | 3.51E-08 | GEMMA | SIG |
| GW_17SQ | AX-110636811 | 1A | 493735845 | -0.0842 | 3.97E-07 | EMMAX | SUG |
| GW_17SQ | AX-110636811 | 1A | 493735845 | 0.0836 | 7.46E-08 | GEMMA | SIG |
| GW_17SQ | AX-111557604 | 1A | 493738685 | -0.0862 | 1.66E-07 | EMMAX | SUG |
| GW_17SQ | AX-111557604 | 1A | 493738685 | 0.0852 | 3.51E-08 | GEMMA | SIG |
| GW_17SQ | AX-110088318 | 1A | 493738732 | -0.0849 | 2.74E-07 | EMMAX | SUG |
| GW_17SQ | AX-110088318 | 1A | 493738732 | 0.0840 | 5.58E-08 | GEMMA | SIG |
| GW_17SQ | AX-111154927 | 1A | 493813029 | -0.0843 | 3.29E-07 | EMMAX | SUG |
| GW_17SQ | AX-111154927 | 1A | 493813029 | 0.0834 | 6.22E-08 | GEMMA | SIG |
| GW_17SQ | AX-109018278 | 1A | 493813818 | -0.0817 | 7.12E-07 | EMMAX | SUG |
| GW_17SQ | AX-109018278 | 1A | 493813818 | 0.0812 | 1.27E-07 | GEMMA | SUG |
| GW_17SQ | AX-109940181 | 1A | 493933165 | -0.0818 | 8.08E-07 | EMMAX | SUG |
| GW_17SQ | AX-109940181 | 1A | 493933165 | 0.0813 | 1.45E-07 | GEMMA | SUG |
| GW_17SQ | AX-108745005 | 1A | 493936294 | -0.0802 | 7.93E-07 | EMMAX | SUG |
| GW_17SQ | AX-108745005 | 1A | 493936294 | 0.0797 | 1.56E-07 | GEMMA | SUG |

| | | | | | | | |
|---|---|---|---|---|---|---|---|
| GW_17SQ | AX-110603959 | 1A | 493937326 | -0.0797 | 1.31E-06 | EMMAX | SUG |
| GW_17SQ | AX-110603959 | 1A | 493937326 | 0.0794 | 2.26E-07 | GEMMA | SUG |
| GW_17SQ | AX-110967232 | 1A | 493937846 | -0.0788 | 1.76E-06 | EMMAX | SUG |
| GW_17SQ | AX-110967232 | 1A | 493937846 | 0.0777 | 4.67E-07 | GEMMA | SUG |
| GW_17SQ | AX-110666814 | 1A | 493938742 | -0.0818 | 8.08E-07 | EMMAX | SUG |
| GW_17SQ | AX-110666814 | 1A | 493938742 | 0.0813 | 1.45E-07 | GEMMA | SUG |
| GW_17SQ | AX-109999875 | 1A | 493940225 | -0.0866 | 1.86E-07 | EMMAX | SUG |
| GW_17SQ | AX-109999875 | 1A | 493940225 | 0.0859 | 3.77E-08 | GEMMA | SIG |
| GW_17SQ | AX-110449932 | 1A | 493970291 | -0.0818 | 8.08E-07 | EMMAX | SUG |
| GW_17SQ | AX-110449932 | 1A | 493970291 | 0.0813 | 1.45E-07 | GEMMA | SUG |
| GW_17SQ | AX-109540497 | 1A | 494021785 | -0.0818 | 8.08E-07 | EMMAX | SUG |
| GW_17SQ | AX-109540497 | 1A | 494021785 | 0.0813 | 1.45E-07 | GEMMA | SUG |
| GW_17SQ | AX-109010375 | 1A | 494023468 | -0.0807 | 8.39E-07 | EMMAX | SUG |
| GW_17SQ | AX-109010375 | 1A | 494023468 | 0.0802 | 1.58E-07 | GEMMA | SUG |
| GW_17SQ | AX-108789443 | 1A | 494029776 | 0.0761 | 7.92E-07 | GEMMA | SUG |
| GW_17SQ | AX-110391607 | 1A | 494029827 | -0.0802 | 9.23E-07 | EMMAX | SUG |
| GW_17SQ | AX-110391607 | 1A | 494029827 | 0.0802 | 1.41E-07 | GEMMA | SUG |
| GW_17SQ | AX-109942628 | 1A | 494031359 | 0.0745 | 8.54E-07 | GEMMA | SUG |
| GW_17SQ | AX-110453457 | 1A | 494031771 | -0.0805 | 6.98E-07 | EMMAX | SUG |
| GW_17SQ | AX-110453457 | 1A | 494031771 | 0.0805 | 1.13E-07 | GEMMA | SIG |
| GW_17SQ | AX-111543279 | 1A | 494034314 | 0.0742 | 1.16E-06 | GEMMA | SUG |
| GW_17SQ | AX-109965289 | 1A | 494064737 | 0.0755 | 8.66E-07 | GEMMA | SUG |
| GW_17SQ | AX-110452798 | 1A | 494066706 | -0.1658 | 2.43E-07 | EMMAX | SUG |
| GW_17SQ | AX-110452798 | 1A | 494066706 | 0.1646 | 4.53E-08 | GEMMA | SIG |
| GW_17SQ | AX-111007830 | 1A | 494067998 | 0.0755 | 8.66E-07 | GEMMA | SUG |
| GW_17SQ | AX-110121603 | 1A | 494069829 | -0.1597 | 8.05E-07 | EMMAX | SUG |
| GW_17SQ | AX-110121603 | 1A | 494069829 | 0.1590 | 1.41E-07 | GEMMA | SUG |
| GW_17SQ | AX-111596405 | 1A | 494071526 | -0.1028 | 4.37E-08 | EMMAX | SIG |
| GW_17SQ | AX-111596405 | 1A | 494071526 | 0.1037 | 5.00E-09 | GEMMA | SIG |
| GW_17SQ | AX-110375342 | 1A | 494145687 | -0.0788 | 1.76E-06 | EMMAX | SUG |
| GW_17SQ | AX-110375342 | 1A | 494145687 | 0.0777 | 4.67E-07 | GEMMA | SUG |
| GW_17SQ | AX-108735378 | 1A | 494145936 | -0.0814 | 9.05E-07 | EMMAX | SUG |
| GW_17SQ | AX-108735378 | 1A | 494145936 | 0.0810 | 1.49E-07 | GEMMA | SUG |
| GW_17SQ | AX-111047488 | 1A | 494146274 | -0.0818 | 8.08E-07 | EMMAX | SUG |
| GW_17SQ | AX-111047488 | 1A | 494146274 | 0.0813 | 1.45E-07 | GEMMA | SUG |
| GW_17SQ | AX-110608035 | 1A | 494186447 | -0.0842 | 3.97E-07 | EMMAX | SUG |
| GW_17SQ | AX-110608035 | 1A | 494186447 | 0.0836 | 7.46E-08 | GEMMA | SIG |
| GW_17SQ | AX-110497883 | 1A | 494256666 | -0.0842 | 3.97E-07 | EMMAX | SUG |
| GW_17SQ | AX-110497883 | 1A | 494256666 | 0.0836 | 7.46E-08 | GEMMA | SIG |
| GW_17SQ | AX-111213027 | 1A | 494392823 | -0.0818 | 8.08E-07 | EMMAX | SUG |
| GW_17SQ | AX-111213027 | 1A | 494392823 | 0.0813 | 1.45E-07 | GEMMA | SUG |
| GW_17SQ | AX-108972380 | 1A | 494413908 | -0.0818 | 8.08E-07 | EMMAX | SUG |
| GW_17SQ | AX-108972380 | 1A | 494413908 | 0.0813 | 1.45E-07 | GEMMA | SUG |
| GW_17SQ | AX-110521184 | 1A | 494429823 | -0.0796 | 1.18E-06 | EMMAX | SUG |

| | | | | | | | |
|---|---|---|---|---|---|---|---|
| GW_17SQ | AX-110521184 | 1A | 494429823 | 0.0787 | 2.91E-07 | GEMMA | SUG |
| GW_17SQ | AX-109862881 | 1A | 494430364 | 0.0701 | 2.29E-06 | GEMMA | SUG |
| GW_17SQ | AX-109944702 | 1A | 494431149 | -0.0842 | 3.97E-07 | EMMAX | SUG |
| GW_17SQ | AX-109944702 | 1A | 494431149 | 0.0836 | 7.46E-08 | GEMMA | SIG |
| GW_17SQ | AX-108980074 | 1A | 494439319 | 0.1418 | 1.80E-06 | GEMMA | SUG |
| GW_17SQ | AX-111048985 | 1A | 494448229 | -0.1658 | 2.43E-07 | EMMAX | SUG |
| GW_17SQ | AX-111048985 | 1A | 494448229 | 0.1646 | 4.53E-08 | GEMMA | SIG |
| GW_17SQ | AX-108772354 | 1A | 494448372 | -0.1373 | 1.24E-06 | EMMAX | SUG |
| GW_17SQ | AX-108772354 | 1A | 494448372 | 0.1367 | 3.39E-07 | GEMMA | SUG |
| GW_17SQ | AX-108980349 | 1A | 494448409 | 0.0750 | 7.33E-07 | GEMMA | SUG |
| GW_17SQ | AX-110619847 | 1A | 494450065 | -0.0818 | 8.08E-07 | EMMAX | SUG |
| GW_17SQ | AX-110619847 | 1A | 494450065 | 0.0813 | 1.45E-07 | GEMMA | SUG |
| GW_17SQ | AX-95125485 | 1A | 494450453 | -0.1773 | 3.64E-08 | EMMAX | SIG |
| GW_17SQ | AX-95125485 | 1A | 494450453 | 0.1755 | 6.42E-09 | GEMMA | SIG |
| GW_17SQ | AX-108993531 | 1A | 494450592 | -0.1539 | 1.13E-06 | EMMAX | SUG |
| GW_17SQ | AX-108993531 | 1A | 494450592 | 0.1549 | 1.32E-07 | GEMMA | SUG |
| GW_17SQ | AX-109276242 | 1A | 494453464 | -0.0800 | 1.65E-06 | EMMAX | SUG |
| GW_17SQ | AX-109276242 | 1A | 494453464 | 0.0797 | 2.81E-07 | GEMMA | SUG |
| GW_17SQ | AX-89349301 | 1A | 494715565 | -0.0791 | 2.12E-06 | EMMAX | SUG |
| GW_17SQ | AX-89349301 | 1A | 494715565 | 0.0779 | 5.57E-07 | GEMMA | SUG |
| GW_17SQ | AX-110463408 | 1A | 494722975 | -0.0834 | 6.26E-07 | EMMAX | SUG |
| GW_17SQ | AX-110463408 | 1A | 494722975 | 0.0829 | 1.04E-07 | GEMMA | SIG |
| GW_17SQ | AX-111144937 | 1A | 494726460 | 0.0769 | 6.52E-07 | GEMMA | SUG |
| GW_17SQ | AX-111144497 | 1A | 494726576 | 0.0769 | 6.52E-07 | GEMMA | SUG |
| GW_17SQ | AX-111602720 | 1A | 494726699 | 0.0769 | 6.52E-07 | GEMMA | SUG |
| GW_17SQ | AX-110670628 | 1A | 494739964 | -0.0804 | 1.39E-06 | EMMAX | SUG |
| GW_17SQ | AX-110670628 | 1A | 494739964 | 0.0792 | 3.47E-07 | GEMMA | SUG |
| GW_17SQ | AX-110675153 | 1A | 494741947 | 0.0769 | 6.52E-07 | GEMMA | SUG |
| GW_17SQ | AX-109932787 | 1A | 494803158 | -0.0860 | 3.02E-07 | EMMAX | SUG |
| GW_17SQ | AX-109932787 | 1A | 494803158 | 0.0852 | 5.28E-08 | GEMMA | SIG |
| GW_17SQ | AX-110525994 | 1A | 494810533 | -0.0812 | 1.19E-06 | EMMAX | SUG |
| GW_17SQ | AX-110525994 | 1A | 494810533 | 0.0809 | 1.94E-07 | GEMMA | SUG |
| GW_17SQ | AX-110940414 | 1A | 494816188 | -0.0848 | 3.21E-07 | EMMAX | SUG |
| GW_17SQ | AX-110940414 | 1A | 494816188 | 0.0842 | 5.28E-08 | GEMMA | SIG |
| GW_17SQ | AX-108740070 | 1A | 494852292 | -0.0817 | 7.51E-07 | EMMAX | SUG |
| GW_17SQ | AX-108740070 | 1A | 494852292 | 0.0812 | 1.31E-07 | GEMMA | SUG |
| GW_17SQ | AX-108881137 | 1A | 494942998 | -0.0860 | 3.02E-07 | EMMAX | SUG |
| GW_17SQ | AX-108881137 | 1A | 494942998 | 0.0852 | 5.28E-08 | GEMMA | SIG |
| GW_17SQ | AX-110602614 | 1A | 495037397 | -0.0885 | 8.26E-08 | EMMAX | SIG |
| GW_17SQ | AX-110602614 | 1A | 495037397 | 0.0874 | 1.70E-08 | GEMMA | SIG |
| GW_17SQ | AX-108787343 | 1A | 495134000 | -0.0808 | 1.26E-06 | EMMAX | SUG |
| GW_17SQ | AX-108787343 | 1A | 495134000 | 0.0803 | 2.39E-07 | GEMMA | SUG |
| GW_17SQ | AX-108758902 | 1A | 495135932 | -0.0834 | 6.26E-07 | EMMAX | SUG |
| GW_17SQ | AX-108758902 | 1A | 495135932 | 0.0829 | 1.04E-07 | GEMMA | SIG |

| | | | | | | | | |
|---|---|---|---|---|---|---|---|---|
| GW_17SQ | AX-108738443 | 1A | 495195036 | 0.0777 | | 5.24E-07 | GEMMA | SUG |
| GW_17SQ | AX-110594847 | 1A | 495275003 | -0.1628 | | 6.28E-07 | EMMAX | SUG |
| GW_17SQ | AX-110594847 | 1A | 495275003 | 0.1620 | | 1.02E-07 | GEMMA | SIG |
| GW_17SQ | AX-110919077 | 1A | 495283358 | -0.0777 | | 2.29E-06 | EMMAX | SUG |
| GW_17SQ | AX-110919077 | 1A | 495283358 | 0.0777 | | 3.40E-07 | GEMMA | SUG |
| GW_17SQ | AX-109410486 | 1A | 495283987 | -0.0856 | | 2.62E-07 | EMMAX | SUG |
| GW_17SQ | AX-109410486 | 1A | 495283987 | 0.0849 | | 4.24E-08 | GEMMA | SIG |
| GW_17SQ | AX-108968174 | 1A | 495285947 | -0.0834 | | 6.26E-07 | EMMAX | SUG |
| GW_17SQ | AX-108968174 | 1A | 495285947 | 0.0829 | | 1.04E-07 | GEMMA | SIG |
| GW_17SQ | AX-89348509 | 1A | 495287232 | -0.0804 | | 9.60E-07 | EMMAX | SUG |
| GW_17SQ | AX-89348509 | 1A | 495287232 | 0.0799 | | 1.70E-07 | GEMMA | SUG |
| GW_17SQ | AX-108927914 | 1A | 495287952 | -0.0834 | | 6.26E-07 | EMMAX | SUG |
| GW_17SQ | AX-108927914 | 1A | 495287952 | 0.0829 | | 1.04E-07 | GEMMA | SIG |
| GW_17SQ | AX-111697444 | 1A | 495290142 | -0.0842 | | 4.25E-07 | EMMAX | SUG |
| GW_17SQ | AX-111697444 | 1A | 495290142 | 0.0836 | | 6.81E-08 | GEMMA | SIG |
| GW_17SQ | AX-108783272 | 1A | 495302918 | -0.0834 | | 6.26E-07 | EMMAX | SUG |
| GW_17SQ | AX-108783272 | 1A | 495302918 | 0.0829 | | 1.04E-07 | GEMMA | SIG |
| GW_17SQ | AX-109520189 | 1A | 495303344 | -0.0794 | | 9.94E-07 | EMMAX | SUG |
| GW_17SQ | AX-109520189 | 1A | 495303344 | 0.0792 | | 1.52E-07 | GEMMA | SUG |
| GW_17SQ | AX-89481437 | 1A | 496088699 | -0.0808 | | 3.52E-07 | EMMAX | SUG |
| GW_17SQ | AX-89481437 | 1A | 496088699 | 0.0817 | | 3.25E-08 | GEMMA | SIG |
| GW_17SQ | AX-109375456 | 1A | 496145066 | -0.0899 | | 1.70E-08 | EMMAX | SIG |
| GW_17SQ | AX-109375456 | 1A | 496145066 | 0.0892 | | 2.36E-09 | GEMMA | SIG |
| GW_17SQ | AX-109924172 | 1A | 577887450 | -0.0118 | 0.0694 | 7.70E-13 | IIIVmrMLM | SIG |
| GW_17SQ | AX-95150104 | 1A | 593297263 | 0.0148 | -0.0496 | 7.56E-06 | IIIVmrMLM | SUG |
| GW_17SQ | AX-111203541 | 1B | 74248438 | -0.0003 | 0.1354 | 2.27E-06 | IIIVmrMLM | SUG |
| GW_17SQ | AX-109156332 | 1D | 469527442 | -0.0214 | | 3.08E-09 | IIIVmrMLM | SIG |
| GW_17SQ | AX-111093750 | 2A | 8729197 | 0.0187 | -0.0145 | 4.48E-06 | IIIVmrMLM | SUG |
| GW_17SQ | AX-111078345 | 2A | 15482218 | 0.0380 | | 1.24E-06 | BLUPmrMLM | SUG |
| GW_17SQ | AX-110416871 | 2A | 709414300 | -0.0041 | -0.1107 | 8.39E-09 | IIIVmrMLM | SIG |
| GW_17SQ | AX-111005120 | 2A | 745221303 | -0.0312 | -0.014 | 6.56E-14 | IIIVmrMLM | SIG |
| GW_17SQ | AX-108830144 | 2B | 23271184 | -0.0024 | -0.2859 | 5.18E-12 | IIIVmrMLM | SIG |
| GW_17SQ | AX-111024765 | 2B | 680074379 | -0.0162 | | 1.11E-04 | BLUPmrMLM | SUG |
| GW_17SQ | AX-110598837 | 2B | 746717981 | -0.0096 | 0.6724 | 1.75E-14 | IIIVmrMLM | SIG |
| GW_17SQ | AX-94868242 | 2B | 761989101 | -0.0019 | 0.1032 | 2.65E-11 | IIIVmrMLM | SIG |
| GW_17SQ | AX-110094837 | 2B | 793533510 | 0.0046 | -0.0521 | 5.02E-07 | IIIVmrMLM | SUG |
| GW_17SQ | AX-110035891 | 2D | 62290445 | -0.0048 | 0.1345 | 1.10E-04 | IIIVmrMLM | SUG |
| GW_17SQ | AX-110896230 | 2D | 73209127 | -0.0318 | -0.014 | 1.34E-13 | IIIVmrMLM | SIG |
| GW_17SQ | AX-110554761 | 2D | 600065511 | -0.0169 | | 1.34E-04 | BLUPmrMLM | SUG |
| GW_17SQ | AX-111786186 | 3A | 22840693 | -0.0233 | | 1.57E-04 | BLUPmrMLM | SUG |
| GW_17SQ | AX-109032308 | 3A | 532615629 | 0.0798 | | 1.73E-06 | GEMMA | SUG |
| GW_17SQ | AX-111731681 | 3A | 567562297 | -0.0050 | -0.0632 | 1.95E-07 | IIIVmrMLM | SUG |
| GW_17SQ | AX-109519759 | 3A | 661498544 | 0.0037 | 0.0809 | 1.20E-04 | IIIVmrMLM | SUG |
| GW_17SQ | AX-89451205 | 3A | 709192992 | -0.0321 | | 8.62E-07 | BLUPmrMLM | SUG |

| | | | | | | | | |
|---|---|---|---|---|---|---|---|---|
| GW_17SQ | AX-89451205 | 3A | 709192992 | 0.0231 | 0.0554 | 5.17E-10 | IIIVmrMLM | SIG |
| GW_17SQ | AX-109375863 | 3A | 731084608 | -0.0241 | 0.0718 | 5.68E-11 | IIIVmrMLM | SIG |
| GW_17SQ | AX-94957235 | 3A | 746051958 | -0.0285 | | 5.75E-05 | BLUPmrMLM | SUG |
| GW_17SQ | AX-109345791 | 3B | 432839506 | -0.0162 | -0.0613 | 6.75E-17 | IIIVmrMLM | SIG |
| GW_17SQ | AX-110362815 | 3B | 475656251 | -0.0274 | | 1.13E-05 | BLUPmrMLM | SUG |
| GW_17SQ | AX-110362815 | 3B | 475656251 | 0.0336 | 0.0124 | 2.74E-17 | IIIVmrMLM | SIG |
| GW_17SQ | AX-111115844 | 4A | 728852358 | -0.0202 | | 3.66E-05 | BLUPmrMLM | SUG |
| GW_17SQ | AX-109465587 | 4A | 732734279 | -0.0351 | 0.0012 | 3.39E-15 | IIIVmrMLM | SIG |
| GW_17SQ | AX-111135528 | 4B | 670443225 | -0.0296 | | 5.71E-16 | IIIVmrMLM | SIG |
| GW_17SQ | AX-110946030 | 4D | 208560799 | 0.0559 | | 1.19E-04 | BLUPmrMLM | SUG |
| GW_17SQ | AX-111216033 | 5A | 11934234 | 0.0286 | -0.0327 | 3.01E-14 | IIIVmrMLM | SIG |
| GW_17SQ | AX-110984007 | 5A | 479048638 | 0.0058 | 0.0727 | 1.43E-06 | IIIVmrMLM | SUG |
| GW_17SQ | AX-110125815 | 5A | 526247469 | 0.0226 | | 1.44E-04 | BLUPmrMLM | SUG |
| GW_17SQ | AX-111058773 | 5A | 667028379 | -0.0386 | | 4.58E-07 | BLUPmrMLM | SUG |
| GW_17SQ | AX-109500949 | 5B | 30994623 | -0.0041 | -0.0733 | 1.01E-08 | IIIVmrMLM | SIG |
| GW_17SQ | AX-109401420 | 5B | 309413507 | -0.0074 | -0.0908 | 1.48E-09 | IIIVmrMLM | SIG |
| GW_17SQ | AX-111041800 | 5B | 458144181 | -0.0411 | | 1.68E-07 | BLUPmrMLM | SUG |
| GW_17SQ | AX-109292201 | 5B | 685087539 | -0.0265 | | 1.35E-04 | BLUPmrMLM | SUG |
| GW_17SQ | AX-111544197 | 5B | 694006788 | 0.0181 | | 1.39E-04 | BLUPmrMLM | SUG |
| GW_17SQ | AX-110808055 | 5D | 162559013 | 0.0080 | -0.079 | 1.82E-07 | IIIVmrMLM | SUG |
| GW_17SQ | AX-108821206 | 5D | 247780961 | -0.1092 | | 2.43E-07 | EMMAX | SUG |
| GW_17SQ | AX-108821206 | 5D | 247780961 | 0.1086 | | 1.18E-07 | GEMMA | SUG |
| GW_17SQ | AX-111815966 | 5D | 300882214 | -0.2017 | | 2.30E-06 | EMMAX | SUG |
| GW_17SQ | AX-111815966 | 5D | 300882214 | 0.1956 | | 1.37E-06 | GEMMA | SUG |
| GW_17SQ | AX-94894465 | 5D | 393907372 | -0.0399 | | 9.75E-27 | IIIVmrMLM | SIG |
| GW_17SQ | AX-110696799 | 5D | 439085044 | 0.0252 | | 3.93E-12 | IIIVmrMLM | SIG |
| GW_17SQ | AX-110974080 | 5D | 539222152 | 0.0186 | 0.0078 | 1.80E-06 | IIIVmrMLM | SUG |
| GW_17SQ | AX-108938900 | 6A | 73821085 | 0.0171 | 0.0204 | 8.77E-06 | IIIVmrMLM | SUG |
| GW_17SQ | AX-111797071 | 6A | 262800331 | 0.0282 | -0.0055 | 5.09E-13 | IIIVmrMLM | SIG |
| GW_17SQ | AX-109095739 | 6A | 289117086 | -0.0773 | | 1.62E-13 | BLUPmrMLM | SIG |
| GW_17SQ | AX-109095739 | 6A | 289117086 | 0.0816 | | 5.99E-07 | EMMAX | SUG |
| GW_17SQ | AX-109095739 | 6A | 289117086 | -0.0806 | | 2.50E-07 | GEMMA | SUG |
| GW_17SQ | AX-110933316 | 6A | 599177104 | 0.0366 | | 5.47E-08 | BLUPmrMLM | SIG |
| GW_17SQ | AX-111130984 | 6A | 599676451 | -0.0289 | -0.0534 | 5.52E-15 | IIIVmrMLM | SIG |
| GW_17SQ | AX-89672871 | 6B | 9098279 | 0.0247 | 0.0252 | 7.61E-11 | IIIVmrMLM | SIG |
| GW_17SQ | AX-109585557 | 6B | 153407756 | 0.0104 | 0.0677 | 9.94E-09 | IIIVmrMLM | SIG |
| GW_17SQ | AX-110482223 | 6B | 223657898 | -0.0793 | 0.0877 | 1.46E-79 | IIIVmrMLM | SIG |
| GW_17SQ | AX-111566512 | 6B | 490901658 | 0.0006 | -0.0653 | 4.82E-12 | IIIVmrMLM | SIG |
| GW_17SQ | AX-95097406 | 6B | 559356967 | 0.0170 | | 2.27E-06 | IIIVmrMLM | SUG |
| GW_17SQ | AX-111481557 | 6D | 86651424 | 0.0334 | | 2.10E-07 | BLUPmrMLM | SUG |
| GW_17SQ | AX-109956383 | 7A | 20494842 | 0.0309 | | 1.22E-06 | BLUPmrMLM | SUG |
| GW_17SQ | AX-95631593 | 7A | 140994052 | -0.0134 | -0.1678 | 4.33E-12 | IIIVmrMLM | SIG |
| GW_17SQ | AX-94754948 | 7A | 730431498 | -0.1306 | | 4.57E-08 | BLUPmrMLM | SIG |
| GW_17SQ | AX-109453615 | 7B | 40356595 | -0.0580 | -0.0182 | 1.10E-34 | IIIVmrMLM | SIG |

| | | | | | | | | |
|---|---|---|---|---|---|---|---|---|
| GW_17SQ | AX-111568366 | 7B | 73137862 | 0.0685 | | 1.09E-06 | FarmCPU | SUG |
| GW_17SQ | AX-109860705 | 7B | 124006510 | 0.0140 | -0.1006 | 2.38E-12 | IIIVmrMLM | SIG |
| GW_17SQ | AX-111578778 | 7D | 632838751 | 0.0077 | 0.0892 | 1.45E-09 | IIIVmrMLM | SIG |
| GW_17XX | AX-111055136 | 1A | 353250144 | -0.0045 | 1.0091 | 1.52E-32 | IIIVmrMLM | SIG |
| GW_17XX | AX-108755149 | 1A | 366981972 | -0.0083 | 0.5447 | 4.74E-12 | IIIVmrMLM | SIG |
| GW_17XX | AX-111013589 | 1A | 428673616 | -0.7285 | | 8.65E-15 | EMMAX | SIG |
| GW_17XX | AX-109427328 | 1A | 564586780 | 0.1030 | | 2.11E-06 | FarmCPU | SUG |
| GW_17XX | AX-94432470 | 1B | 297113879 | -0.3146 | | 5.52E-07 | EMMAX | SUG |
| GW_17XX | AX-110514172 | 1B | 559333678 | -0.7696 | | 8.85E-10 | EMMAX | SIG |
| GW_17XX | AX-111796916 | 1B | 605987118 | 0.0339 | | 5.92E-07 | BLUPmrMLM | SUG |
| GW_17XX | AX-95083211 | 1B | 672928015 | -0.0324 | | 5.41E-06 | BLUPmrMLM | SUG |
| GW_17XX | AX-95083211 | 1B | 672928015 | 0.0220 | -0.0035 | 3.79E-04 | IIIVmrMLM | SUG |
| GW_17XX | AX-94896359 | 1D | 7210992 | -0.0383 | | 3.21E-05 | BLUPmrMLM | SUG |
| GW_17XX | AX-94896359 | 1D | 7210992 | 0.0734 | 0.0289 | 9.27E-63 | IIIVmrMLM | SIG |
| GW_17XX | AX-109249234 | 1D | 122142604 | 0.0163 | | 4.88E-07 | IIIVmrMLM | SUG |
| GW_17XX | AX-94564632 | 1D | 124344059 | -0.3146 | | 5.52E-07 | EMMAX | SUG |
| GW_17XX | AX-111150614 | 1D | 124345536 | -0.3146 | | 5.52E-07 | EMMAX | SUG |
| GW_17XX | AX-111801848 | 1D | 200463811 | -0.6941 | | 1.33E-14 | EMMAX | SIG |
| GW_17XX | AX-95188544 | 2A | 59554321 | -0.2072 | | 1.71E-06 | EMMAX | SUG |
| GW_17XX | AX-95188544 | 2A | 59554321 | 0.2047 | | 9.59E-07 | GEMMA | SUG |
| GW_17XX | AX-109446972 | 2A | 187199191 | -0.0232 | | 2.99E-05 | BLUPmrMLM | SUG |
| GW_17XX | AX-94764362 | 2A | 692531292 | -0.0946 | | 2.06E-06 | EMMAX | SUG |
| GW_17XX | AX-94452059 | 2A | 692531374 | -0.0994 | | 8.54E-07 | EMMAX | SUG |
| GW_17XX | AX-94452059 | 2A | 692531374 | 0.1001 | | 1.76E-06 | GEMMA | SUG |
| GW_17XX | AX-108957347 | 2A | 692937617 | -0.1127 | | 1.95E-06 | EMMAX | SUG |
| GW_17XX | AX-110396904 | 2A | 758270594 | -0.1776 | | 2.32E-07 | EMMAX | SUG |
| GW_17XX | AX-110396904 | 2A | 758270594 | 0.1821 | | 4.32E-07 | GEMMA | SUG |
| GW_17XX | AX-110575220 | 2B | 88839532 | -0.4430 | | 1.08E-10 | EMMAX | SIG |
| GW_17XX | AX-110407522 | 2B | 88843147 | -0.3442 | | 5.53E-09 | EMMAX | SIG |
| GW_17XX | AX-109918309 | 2B | 88846689 | -0.2972 | | 3.17E-07 | EMMAX | SUG |
| GW_17XX | AX-109842008 | 2B | 88848753 | -0.4430 | | 1.08E-10 | EMMAX | SIG |
| GW_17XX | AX-111504331 | 2B | 88927001 | -0.1987 | | 1.74E-09 | EMMAX | SIG |
| GW_17XX | AX-111504331 | 2B | 88927001 | 0.2023 | | 2.53E-09 | GEMMA | SIG |
| GW_17XX | AX-111553102 | 2B | 88932703 | -0.4430 | | 1.08E-10 | EMMAX | SIG |
| GW_17XX | AX-110491472 | 2B | 88972848 | -0.1987 | | 1.74E-09 | EMMAX | SIG |
| GW_17XX | AX-110491472 | 2B | 88972848 | 0.2023 | | 2.53E-09 | GEMMA | SIG |
| GW_17XX | AX-108802135 | 2B | 88974337 | -0.1995 | | 1.42E-09 | EMMAX | SIG |
| GW_17XX | AX-108802135 | 2B | 88974337 | 0.2003 | | 1.63E-09 | GEMMA | SIG |
| GW_17XX | AX-111729936 | 2B | 88975938 | -0.5519 | | 2.72E-13 | EMMAX | SIG |
| GW_17XX | AX-111039155 | 2B | 89435305 | -0.3104 | | 1.11E-07 | EMMAX | SIG |
| GW_17XX | AX-95220376 | 2B | 91547976 | -0.2164 | | 1.26E-06 | EMMAX | SUG |
| GW_17XX | AX-111026696 | 2B | 91612859 | -0.2164 | | 1.26E-06 | EMMAX | SUG |
| GW_17XX | AX-110995859 | 2B | 94926376 | -0.1185 | | 5.88E-07 | EMMAX | SUG |
| GW_17XX | AX-110995859 | 2B | 94926376 | 0.1188 | | 3.34E-07 | GEMMA | SUG |

| | | | | | | | | |
|---|---|---|---|---|---|---|---|---|
| GW_17XX | AX-94585598 | 2B | 105375977 | 0.1379 | | 1.89E-06 | GEMMA | SUG |
| GW_17XX | AX-109941896 | 2B | 118320500 | -0.1818 | | 2.67E-07 | EMMAX | SUG |
| GW_17XX | AX-109941896 | 2B | 118320500 | 0.1831 | | 7.98E-08 | GEMMA | SIG |
| GW_17XX | AX-110623031 | 2B | 118603313 | -0.3634 | | 8.86E-08 | EMMAX | SIG |
| GW_17XX | AX-111208632 | 2B | 245006974 | 0.1147 | | 1.90E-06 | GEMMA | SUG |
| GW_17XX | AX-112288222 | 2B | 426478450 | -0.0019 | 0.0633 | 9.52E-08 | IIIVmrMLM | SIG |
| GW_17XX | AX-94554516 | 2D | 2363017 | -0.8849 | | 2.82E-22 | EMMAX | SIG |
| GW_17XX | AX-95097309 | 2D | 56202857 | -0.4430 | | 1.08E-10 | EMMAX | SIG |
| GW_17XX | AX-110563788 | 2D | 73019968 | 0.0006 | -0.0586 | 2.75E-04 | IIIVmrMLM | SUG |
| GW_17XX | AX-94581002 | 2D | 201943345 | -0.5099 | | 1.10E-06 | EMMAX | SUG |
| GW_17XX | AX-111084803 | 2D | 328355468 | -0.4060 | | 5.01E-09 | EMMAX | SIG |
| GW_17XX | AX-94591324 | 2D | 473276253 | -0.4406 | | 2.38E-09 | EMMAX | SIG |
| GW_17XX | AX-111068326 | 2D | 601273284 | 0.0035 | -0.0805 | 1.04E-06 | IIIVmrMLM | SUG |
| GW_17XX | AX-111562358 | 3A | 147799164 | -0.2119 | | 8.22E-07 | EMMAX | SUG |
| GW_17XX | AX-111562358 | 3A | 147799164 | 0.2126 | | 2.61E-07 | GEMMA | SUG |
| GW_17XX | AX-111575018 | 3A | 685582352 | 0.0249 | 0.0009 | 5.73E-12 | IIIVmrMLM | SIG |
| GW_17XX | AX-108815613 | 3A | 695753359 | 0.0199 | | 1.02E-09 | IIIVmrMLM | SIG |
| GW_17XX | AX-111013308 | 3B | 497425509 | 0.0001 | 0.1098 | 1.84E-10 | IIIVmrMLM | SIG |
| GW_17XX | AX-109932984 | 3B | 713542828 | 0.0053 | -0.0374 | 7.56E-18 | IIIVmrMLM | SIG |
| GW_17XX | AX-95099439 | 3B | 768746333 | -0.7061 | | 1.98E-14 | EMMAX | SIG |
| GW_17XX | AX-95083173 | 3D | 126945778 | -0.7696 | | 8.85E-10 | EMMAX | SIG |
| GW_17XX | AX-111046642 | 3D | 143003646 | -1.4370 | | 1.95E-30 | EMMAX | SIG |
| GW_17XX | AX-109644087 | 4A | 168870504 | 0.0303 | | 7.05E-20 | IIIVmrMLM | SIG |
| GW_17XX | AX-95015737 | 4A | 346060786 | -0.1849 | | 4.16E-08 | EMMAX | SIG |
| GW_17XX | AX-95015737 | 4A | 346060786 | 0.1945 | | 9.08E-08 | GEMMA | SIG |
| GW_17XX | AX-94563592 | 4A | 356553186 | -0.1764 | | 1.33E-07 | EMMAX | SUG |
| GW_17XX | AX-94563592 | 4A | 356553186 | 0.1855 | | 3.04E-07 | GEMMA | SUG |
| GW_17XX | AX-109023322 | 4A | 687312211 | -0.3152 | | 5.02E-07 | EMMAX | SUG |
| GW_17XX | AX-111079457 | 4A | 722423137 | -0.0059 | -0.0298 | 2.07E-04 | IIIVmrMLM | SUG |
| GW_17XX | AX-109295508 | 4A | 739598992 | -0.0045 | -0.0975 | 5.64E-07 | IIIVmrMLM | SUG |
| GW_17XX | AX-94438527 | 4B | 26491532 | -0.0663 | | 3.87E-07 | GEMMA | SUG |
| GW_17XX | AX-94508980 | 4B | 27522453 | -0.1150 | | 2.12E-06 | GEMMA | SUG |
| GW_17XX | AX-108946619 | 4B | 263254047 | -0.0242 | | 1.49E-13 | IIIVmrMLM | SIG |
| GW_17XX | AX-109032341 | 4B | 295963534 | -0.1849 | | 4.16E-08 | EMMAX | SIG |
| GW_17XX | AX-109032341 | 4B | 295963534 | 0.1945 | | 9.08E-08 | GEMMA | SIG |
| GW_17XX | AX-111157728 | 4B | 296009502 | -0.1384 | | 1.87E-06 | EMMAX | SUG |
| GW_17XX | AX-109383374 | 4B | 296406286 | -0.1849 | | 4.16E-08 | EMMAX | SIG |
| GW_17XX | AX-109383374 | 4B | 296406286 | 0.1945 | | 9.08E-08 | GEMMA | SIG |
| GW_17XX | AX-111533236 | 4B | 299257109 | -0.1633 | | 3.16E-07 | EMMAX | SUG |
| GW_17XX | AX-111533236 | 4B | 299257109 | 0.1702 | | 6.68E-07 | GEMMA | SUG |
| GW_17XX | AX-94477702 | 4B | 300055236 | -0.1720 | | 1.82E-07 | EMMAX | SUG |
| GW_17XX | AX-94477702 | 4B | 300055236 | 0.1814 | | 4.08E-07 | GEMMA | SUG |
| GW_17XX | AX-95159519 | 4B | 300055714 | -0.2552 | | 1.24E-10 | EMMAX | SIG |
| GW_17XX | AX-95159519 | 4B | 300055714 | 0.2590 | | 2.20E-10 | GEMMA | SIG |

| | | | | | | | | |
|---|---|---|---|---|---|---|---|---|
| GW_17XX | AX-94722710 | 4B | 300810457 | -0.1909 | | 2.10E-08 | EMMAX | SIG |
| GW_17XX | AX-94722710 | 4B | 300810457 | 0.2062 | | 4.75E-08 | GEMMA | SIG |
| GW_17XX | AX-109579982 | 4B | 534331888 | 0.0230 | -0.021 | 2.32E-11 | IIIVmrMLM | SIG |
| GW_17XX | AX-95259438 | 4B | 579406257 | -0.1760 | | 1.41E-06 | EMMAX | SUG |
| GW_17XX | AX-95259438 | 4B | 579406257 | 0.1767 | | 1.94E-06 | GEMMA | SUG |
| GW_17XX | AX-109560277 | 4D | 113169995 | -1.4370 | | 1.95E-30 | EMMAX | SIG |
| GW_17XX | AX-109098724 | 4D | 158293010 | -0.4163 | | 1.77E-12 | EMMAX | SIG |
| GW_17XX | AX-111259868 | 4D | 165577168 | -1.4370 | | 1.95E-30 | EMMAX | SIG |
| GW_17XX | AX-109862804 | 4D | 202968848 | -0.3279 | | 3.26E-07 | EMMAX | SUG |
| GW_17XX | AX-112287379 | 4D | 241796879 | -0.1720 | | 1.82E-07 | EMMAX | SUG |
| GW_17XX | AX-112287379 | 4D | 241796879 | 0.1814 | | 4.08E-07 | GEMMA | SUG |
| GW_17XX | AX-109501527 | 4D | 246375877 | -0.7696 | | 8.85E-10 | EMMAX | SIG |
| GW_17XX | AX-109011177 | 4D | 346462443 | -0.2247 | | 1.69E-06 | EMMAX | SUG |
| GW_17XX | AX-110415726 | 4D | 502594399 | -0.6227 | | 2.50E-13 | EMMAX | SIG |
| GW_17XX | AX-94546265 | 5A | 46232449 | 0.0260 | | 5.54E-06 | BLUPmrMLM | SUG |
| GW_17XX | AX-110529122 | 5A | 48746269 | -0.0343 | -0.008 | 6.89E-23 | IIIVmrMLM | SIG |
| GW_17XX | AX-111498225 | 5A | 692364590 | -0.0101 | -0.0401 | 1.70E-07 | IIIVmrMLM | SUG |
| GW_17XX | AX-110397398 | 5B | 463130232 | -0.0259 | | 5.10E-05 | BLUPmrMLM | SUG |
| GW_17XX | AX-109468686 | 5B | 584894906 | -0.0118 | -0.3137 | 6.12E-07 | IIIVmrMLM | SUG |
| GW_17XX | AX-109390035 | 5B | 679136987 | -0.0218 | -0.0006 | 2.81E-07 | IIIVmrMLM | SUG |
| GW_17XX | AX-111668610 | 5D | 61072160 | -0.4745 | | 2.70E-12 | EMMAX | SIG |
| GW_17XX | AX-110051439 | 5D | 72467149 | -0.3355 | | 3.30E-07 | EMMAX | SUG |
| GW_17XX | AX-111939538 | 5D | 160286069 | -0.6230 | | 3.72E-14 | EMMAX | SIG |
| GW_17XX | AX-109332091 | 5D | 385608552 | -0.0254 | -0.0924 | 1.96E-14 | IIIVmrMLM | SIG |
| GW_17XX | AX-95199226 | 6A | 125497524 | -1.4370 | | 1.95E-30 | EMMAX | SIG |
| GW_17XX | AX-94492233 | 6A | 544478755 | -0.3952 | | 1.74E-09 | EMMAX | SIG |
| GW_17XX | AX-111262274 | 6A | 579131834 | -0.0167 | 0.025 | 1.17E-06 | IIIVmrMLM | SUG |
| GW_17XX | AX-111520069 | 6B | 175719106 | 0.0070 | 0.0604 | 5.12E-11 | IIIVmrMLM | SIG |
| GW_17XX | AX-108842364 | 6B | 451402100 | 0.0003 | 0.0618 | 7.47E-07 | IIIVmrMLM | SUG |
| GW_17XX | AX-109960216 | 6B | 461846325 | 0.0222 | | 1.93E-05 | BLUPmrMLM | SUG |
| GW_17XX | AX-111627601 | 6B | 473259388 | -0.0199 | | 1.99E-04 | BLUPmrMLM | SUG |
| GW_17XX | AX-108925647 | 6B | 645815399 | -0.1911 | | 3.51E-09 | EMMAX | SIG |
| GW_17XX | AX-108925647 | 6B | 645815399 | 0.1904 | | 1.43E-09 | GEMMA | SIG |
| GW_17XX | AX-111540249 | 6B | 646132357 | -0.1462 | | 1.10E-06 | EMMAX | SUG |
| GW_17XX | AX-111540249 | 6B | 646132357 | 0.1458 | | 3.74E-07 | GEMMA | SUG |
| GW_17XX | AX-110515672 | 6B | 646133256 | -0.1705 | | 8.23E-08 | EMMAX | SIG |
| GW_17XX | AX-110515672 | 6B | 646133256 | 0.1697 | | 3.63E-08 | GEMMA | SIG |
| GW_17XX | AX-110931236 | 6B | 646134232 | -0.1705 | | 8.23E-08 | EMMAX | SIG |
| GW_17XX | AX-110931236 | 6B | 646134232 | 0.1697 | | 3.63E-08 | GEMMA | SIG |
| GW_17XX | AX-110573643 | 6B | 646579323 | -0.1770 | | 2.99E-08 | EMMAX | SIG |
| GW_17XX | AX-110573643 | 6B | 646579323 | 0.1760 | | 1.01E-08 | GEMMA | SIG |
| GW_17XX | AX-109580754 | 6B | 647150561 | -0.0229 | 0.3497 | 3.23E-15 | IIIVmrMLM | SIG |
| GW_17XX | AX-108971391 | 6B | 652064807 | -0.1948 | | 1.07E-06 | EMMAX | SUG |
| GW_17XX | AX-108971391 | 6B | 652064807 | 0.1929 | | 2.50E-07 | GEMMA | SUG |

| | | | | | | | | |
|---|---|---|---|---|---|---|---|---|
| GW_17XX | AX-94862958 | 6D | 2103224 | 0.0047 | -0.0366 | 6.33E-18 | IIIVmrMLM | SIG |
| GW_17XX | AX-109091335 | 6D | 109485655 | -1.4370 | | 1.95E-30 | EMMAX | SIG |
| GW_17XX | AX-108815731 | 6D | 127137109 | -1.4370 | | 1.95E-30 | EMMAX | SIG |
| GW_17XX | AX-110952407 | 6D | 428638114 | -1.4370 | | 1.95E-30 | EMMAX | SIG |
| GW_17XX | AX-109907254 | 7A | 127844349 | 0.1225 | | 1.08E-06 | GEMMA | SUG |
| GW_17XX | AX-110379194 | 7A | 720621953 | -0.1086 | | 2.33E-06 | EMMAX | SUG |
| GW_17XX | AX-110379194 | 7A | 720621953 | 0.1084 | | 1.37E-06 | GEMMA | SUG |
| GW_17XX | AX-108757205 | 7B | 85311709 | -0.0146 | 0.0969 | 3.80E-09 | IIIVmrMLM | SIG |
| GW_17XX | AX-108756319 | 7B | 679786935 | -1.4370 | | 1.95E-30 | EMMAX | SIG |
| GW_17XX | AX-95239089 | 7B | 699375329 | 0.0044 | 0.1008 | 1.17E-06 | IIIVmrMLM | SUG |
| GW_17XX | AX-108964563 | 7B | 712076548 | -0.0266 | | 9.19E-09 | BLUPmrMLM | SIG |
| GW_17XX | AX-94626341 | 7D | 110286295 | -0.2314 | | 2.66E-07 | EMMAX | SUG |
| GW_17XX | AX-94626341 | 7D | 110286295 | 0.2319 | | 1.78E-07 | GEMMA | SUG |
| GW_17XX | AX-111259919 | 7D | 359814722 | -0.5611 | | 7.80E-12 | EMMAX | SIG |
| GW_17XX | AX-95135172 | 7D | 363044822 | -0.2866 | | 1.37E-06 | EMMAX | SUG |
| GW_17XX | AX-109632206 | 7D | 372244929 | -1.4370 | | 1.95E-30 | EMMAX | SIG |
| GW_17XX | AX-95257251 | 7D | 512281202 | -0.7696 | | 8.85E-10 | EMMAX | SIG |
| GW_17XX | AX-111783605 | 7D | 528053219 | -0.3355 | | 1.65E-08 | EMMAX | SIG |
| GW_17XX | AX-109517671 | 7D | 565202537 | 0.0245 | -0.0422 | 4.91E-13 | IIIVmrMLM | SIG |
| GW_17XX | AX-111066405 | 7D | 592553571 | -0.6505 | | 1.95E-12 | EMMAX | SIG |
| GW_17XX | AX-109281960 | 7D | 592616061 | -0.4938 | | 6.21E-10 | EMMAX | SIG |
| GW_17XX | AX-109578957 | 7D | 592618737 | -0.6505 | | 1.95E-12 | EMMAX | SIG |
| GW_17XX | AX-110642814 | 7D | 592675625 | -0.6505 | | 1.95E-12 | EMMAX | SIG |
| GW_17XX | AX-110074623 | 7D | 592675666 | -0.6505 | | 1.95E-12 | EMMAX | SIG |
| GW_17XX | AX-111041751 | 7D | 592763081 | -0.6505 | | 1.95E-12 | EMMAX | SIG |
| GW_17XX | AX-110385111 | 7D | 592856686 | -0.6505 | | 1.95E-12 | EMMAX | SIG |
| GW_17XX | AX-94545540 | 7D | 600453744 | -0.6969 | | 1.80E-14 | EMMAX | SIG |
| GW_17XX | AX-95118138 | 7D | 600455779 | -0.4873 | | 4.74E-10 | EMMAX | SIG |
| GW_17XX | AX-110018185 | 7D | 611351361 | -0.0098 | 0.0983 | 8.66E-10 | IIIVmrMLM | SIG |
| GW_17ZMD | AX-109351496 | 1B | 623830408 | -0.0078 | -0.7085 | 3.89E-05 | IIIVmrMLM | SUG |
| GW_17ZMD | AX-109872000 | 2A | 380566504 | 0.0154 | 0.4167 | 5.76E-07 | IIIVmrMLM | SUG |
| GW_17ZMD | AX-109313463 | 2A | 678038940 | 0.0736 | -0.0503 | 1.22E-05 | IIIVmrMLM | SUG |
| GW_17ZMD | AX-94873196 | 2A | 744081290 | 0.0567 | | 1.65E-05 | IIIVmrMLM | SUG |
| GW_17ZMD | AX-110967584 | 2B | 765577401 | -0.0270 | -0.5308 | 2.26E-09 | IIIVmrMLM | SIG |
| GW_17ZMD | AX-111651751 | 3A | 46832855 | -0.0759 | | 4.88E-05 | BLUPmrMLM | SIG |
| GW_17ZMD | AX-111097121 | 3B | 246500476 | 0.0179 | -0.2082 | 1.40E-05 | IIIVmrMLM | SUG |
| GW_17ZMD | AX-111477466 | 3B | 341919432 | 0.0783 | -0.3217 | 3.73E-15 | IIIVmrMLM | SIG |
| GW_17ZMD | AX-108813535 | 3B | 678549369 | -0.1172 | | 1.69E-05 | BLUPmrMLM | SIG |
| GW_17ZMD | AX-109835036 | 3B | 817321911 | 0.0732 | | 2.13E-05 | BLUPmrMLM | SIG |
| GW_17ZMD | AX-111010012 | 4A | 673355278 | -0.0177 | 0.1783 | 4.00E-05 | IIIVmrMLM | SUG |
| GW_17ZMD | AX-111015646 | 4A | 742968720 | -0.0015 | 0.1922 | 1.13E-06 | IIIVmrMLM | SUG |
| GW_17ZMD | AX-94430295 | 4D | 7758059 | -0.0889 | -0.2554 | 1.90E-21 | IIIVmrMLM | SIG |
| GW_17ZMD | AX-111035309 | 5B | 589890159 | -0.0699 | -0.1344 | 4.32E-07 | IIIVmrMLM | SUG |
| GW_17ZMD | AX-95092536 | 5D | 437345420 | 0.1429 | | 1.18E-04 | BLUPmrMLM | SIG |

| | | | | | | | | |
|---|---|---|---|---|---|---|---|---|
| GW_17ZMD | AX-111101623 | 6A | 61891179 | -0.0682 | -0.0357 | 1.65E-05 | IIIVmrMLM | SUG |
| GW_17ZMD | AX-110989117 | 6A | 615770287 | 0.0248 | 0.4625 | 2.65E-08 | IIIVmrMLM | SIG |
| GW_17ZMD | AX-109916648 | 6B | 364121279 | -0.0123 | 0.2337 | 4.95E-05 | IIIVmrMLM | SUG |
| GW_17ZMD | AX-95166654 | 6B | 467460492 | 0.2588 | | 1.09E-06 | FarmCPU | SUG |
| GW_17ZMD | AX-109513780 | 6B | 719761000 | 0.0338 | -0.1287 | 9.13E-08 | IIIVmrMLM | SIG |
| GW_17ZMD | AX-111237845 | 6D | 3843656 | -0.0118 | 0.1241 | 1.23E-05 | IIIVmrMLM | SUG |
| GW_17ZMD | AX-95102532 | 7A | 6760426 | -0.1251 | | 4.28E-05 | BLUPmrMLM | SIG |
| GW_17ZMD | AX-110457352 | 7A | 13741290 | -0.0319 | -0.2307 | 8.78E-11 | IIIVmrMLM | SIG |
| GW_17ZMD | AX-110135909 | 7D | 408267277 | -0.0746 | | 1.80E-08 | IIIVmrMLM | SIG |
| GW_18AY | AX-109370500 | 1B | 304379381 | 0.0599 | | 1.21E-05 | BLUPmrMLM | SUG |
| GW_18AY | AX-111495168 | 3A | 507115740 | 0.0748 | | 4.84E-06 | BLUPmrMLM | SUG |
| GW_18AY | AX-110099879 | 3A | 742440500 | 0.0413 | | 1.02E-04 | BLUPmrMLM | SUG |
| GW_18AY | AX-94818869 | 3B | 354429149 | 0.0849 | | 2.46E-05 | BLUPmrMLM | SUG |
| GW_18AY | AX-110364457 | 5A | 547357382 | -0.0529 | | 3.81E-06 | BLUPmrMLM | SUG |
| GW_18AY | AX-110391003 | 5B | 580519842 | -0.0674 | | 9.45E-07 | BLUPmrMLM | SUG |
| GW_18AY | AX-108848017 | 6D | 15419960 | -0.0756 | | 3.80E-07 | BLUPmrMLM | SUG |
| GW_18AY | AX-110411257 | 7A | 660443419 | 0.0459 | | 3.24E-05 | BLUPmrMLM | SUG |
| GW_18AY | AX-109205685 | 7D | 31414542 | -0.0364 | | 6.04E-05 | BLUPmrMLM | SUG |
| GW_18JY | AX-94603312 | 1A | 497227861 | 0.1895 | | 1.81E-07 | EMMAX | SUG |
| GW_18JY | AX-94603312 | 1A | 497227861 | -0.1919 | | 6.13E-08 | GEMMA | SIG |
| GW_18JY | AX-110964252 | 1A | 497280829 | 0.1908 | | 7.51E-07 | EMMAX | SUG |
| GW_18JY | AX-110964252 | 1A | 497280829 | -0.1931 | | 2.62E-07 | GEMMA | SUG |
| GW_18JY | AX-111289455 | 1A | 497316631 | 0.1908 | | 7.51E-07 | EMMAX | SUG |
| GW_18JY | AX-111289455 | 1A | 497316631 | -0.1931 | | 2.62E-07 | GEMMA | SUG |
| GW_18JY | AX-111607699 | 1A | 497907702 | -0.0472 | | 1.66E-08 | BLUPmrMLM | SIG |
| GW_18JY | AX-94722837 | 1A | 549840341 | -0.0289 | | 4.03E-15 | IIIVmrMLM | SIG |
| GW_18JY | AX-109288858 | 1A | 567973333 | 0.0079 | -0.0921 | 8.82E-16 | IIIVmrMLM | SIG |
| GW_18JY | AX-111596938 | 1B | 505054918 | -0.0176 | | 8.92E-05 | BLUPmrMLM | SUG |
| GW_18JY | AX-109353433 | 2A | 44839327 | 0.0036 | 0.0795 | 8.30E-06 | IIIVmrMLM | SUG |
| GW_18JY | AX-111786868 | 2A | 64358348 | 0.0337 | | 1.11E-19 | IIIVmrMLM | SIG |
| GW_18JY | AX-110429503 | 2A | 70944071 | -0.0059 | 0.0504 | 1.19E-06 | IIIVmrMLM | SUG |
| GW_18JY | AX-110385298 | 2A | 88293768 | -0.0035 | 0.0955 | 5.78E-05 | IIIVmrMLM | SUG |
| GW_18JY | AX-110198456 | 2A | 317805831 | -0.1079 | | 2.09E-07 | FarmCPU | SUG |
| GW_18JY | AX-108901846 | 2A | 517756458 | 0.0292 | -0.0182 | 7.79E-14 | IIIVmrMLM | SIG |
| GW_18JY | AX-109339780 | 2A | 608599107 | 0.0223 | | 5.89E-05 | BLUPmrMLM | SUG |
| GW_18JY | AX-110965993 | 2A | 639930670 | -0.0071 | 0.0736 | 4.52E-12 | IIIVmrMLM | SIG |
| GW_18JY | AX-108860920 | 2B | 8771575 | 0.0101 | 0.0496 | 1.02E-17 | IIIVmrMLM | SIG |
| GW_18JY | AX-94975480 | 2B | 95770643 | 0.0269 | | 2.81E-05 | BLUPmrMLM | SUG |
| GW_18JY | AX-109952335 | 2B | 141563005 | 0.0208 | | 8.83E-05 | BLUPmrMLM | SUG |
| GW_18JY | AX-111763398 | 2B | 682929794 | 0.0406 | 0.0161 | 1.96E-20 | IIIVmrMLM | SIG |
| GW_18JY | AX-110406718 | 2B | 717792580 | 0.0065 | -0.1975 | 7.70E-09 | IIIVmrMLM | SIG |
| GW_18JY | AX-109901197 | 2D | 14886925 | 0.0216 | -0.0292 | 1.80E-08 | IIIVmrMLM | SIG |
| GW_18JY | AX-111880735 | 2D | 17306612 | 0.0007 | -0.0856 | 1.00E-08 | IIIVmrMLM | SIG |
| GW_18JY | AX-108883295 | 2D | 27264461 | -0.0178 | -0.1186 | 4.42E-11 | IIIVmrMLM | SIG |

| | | | | | | | | |
|---|---|---|---|---|---|---|---|---|
| GW_18JY | AX-94697254 | 2D | 33710135 | 0.1178 | | 1.34E-07 | FarmCPU | SUG |
| GW_18JY | AX-108970944 | 2D | 341580209 | 0.0200 | | 1.09E-05 | BLUPmrMLM | SUG |
| GW_18JY | AX-109973125 | 3A | 478680500 | -0.0378 | | 7.99E-07 | BLUPmrMLM | SUG |
| GW_18JY | AX-111052121 | 3A | 651324077 | 0.0218 | | 1.73E-05 | BLUPmrMLM | SUG |
| GW_18JY | AX-108987955 | 3A | 667353880 | 0.0534 | | 3.71E-09 | BLUPmrMLM | SIG |
| GW_18JY | AX-108919606 | 3A | 717309925 | -0.0118 | -0.3109 | 2.71E-08 | IIIVmrMLM | SIG |
| GW_18JY | AX-111118989 | 3B | 15131859 | -0.0098 | -0.0416 | 1.31E-11 | IIIVmrMLM | SIG |
| GW_18JY | AX-110481377 | 3B | 309384755 | 0.0075 | -0.055 | 8.89E-14 | IIIVmrMLM | SIG |
| GW_18JY | AX-110379827 | 3B | 333704358 | 0.0123 | 0.0695 | 4.51E-15 | IIIVmrMLM | SIG |
| GW_18JY | AX-111274915 | 3B | 394791258 | -0.0032 | 0.0693 | 7.84E-21 | IIIVmrMLM | SIG |
| GW_18JY | AX-111010161 | 3B | 414752378 | -0.0127 | 0.0654 | 7.53E-16 | IIIVmrMLM | SIG |
| GW_18JY | AX-111048076 | 3B | 570208555 | 0.0073 | -0.0644 | 1.18E-06 | IIIVmrMLM | SUG |
| GW_18JY | AX-94603300 | 3D | 546581282 | 0.0243 | | 3.04E-11 | IIIVmrMLM | SIG |
| GW_18JY | AX-109285866 | 4A | 613569939 | 0.0376 | -0.007 | 4.91E-14 | IIIVmrMLM | SIG |
| GW_18JY | AX-109486950 | 4A | 636056314 | 0.0215 | | 4.02E-05 | BLUPmrMLM | SUG |
| GW_18JY | AX-109373245 | 4A | 681706805 | -0.0451 | | 1.68E-05 | BLUPmrMLM | SUG |
| GW_18JY | AX-109835724 | 4B | 3904654 | -0.0005 | -0.1869 | 1.72E-06 | IIIVmrMLM | SUG |
| GW_18JY | AX-111052220 | 4B | 99051321 | -0.0241 | | 3.96E-06 | BLUPmrMLM | SUG |
| GW_18JY | AX-110485186 | 5A | 8120257 | -0.0192 | | 9.16E-06 | BLUPmrMLM | SUG |
| GW_18JY | AX-110485186 | 5A | 8120257 | 0.0280 | -0.0101 | 1.61E-12 | IIIVmrMLM | SIG |
| GW_18JY | AX-109903760 | 5A | 30056842 | -0.1149 | | 9.95E-08 | EMMAX | SIG |
| GW_18JY | AX-109903760 | 5A | 30056842 | 0.1157 | | 4.91E-08 | GEMMA | SIG |
| GW_18JY | AX-108840641 | 5A | 586673355 | 0.0219 | -0.2011 | 9.24E-14 | IIIVmrMLM | SIG |
| GW_18JY | AX-110979406 | 5A | 638901408 | 0.0211 | | 3.73E-05 | BLUPmrMLM | SUG |
| GW_18JY | AX-110613342 | 5B | 82631608 | -0.0269 | | 1.71E-06 | BLUPmrMLM | SUG |
| GW_18JY | AX-108934677 | 5B | 520119491 | 0.1357 | | 1.33E-06 | FarmCPU | SUG |
| GW_18JY | AX-111684648 | 5B | 520178595 | 0.1262 | | 6.90E-07 | FarmCPU | SUG |
| GW_18JY | AX-111047480 | 5B | 569827971 | 0.0238 | 0.0061 | 3.76E-08 | IIIVmrMLM | SIG |
| GW_18JY | AX-109354765 | 5B | 678620791 | -0.0308 | | 6.82E-17 | IIIVmrMLM | SIG |
| GW_18JY | AX-109407131 | 5D | 512550680 | -0.0206 | | 1.53E-05 | BLUPmrMLM | SUG |
| GW_18JY | AX-111701819 | 6A | 103294811 | -0.0216 | | 1.78E-04 | BLUPmrMLM | SUG |
| GW_18JY | AX-89563509 | 6A | 107211012 | 0.0395 | 0.0187 | 1.75E-24 | IIIVmrMLM | SIG |
| GW_18JY | AX-110066270 | 6B | 289301998 | 0.0082 | -0.0964 | 8.02E-10 | IIIVmrMLM | SIG |
| GW_18JY | AX-109391962 | 6B | 339268873 | 0.1295 | | 2.31E-06 | FarmCPU | SUG |
| GW_18JY | AX-108928674 | 6D | 8248078 | -0.0010 | -0.154 | 1.32E-06 | IIIVmrMLM | SUG |
| GW_18JY | AX-109551952 | 6D | 19141283 | 0.0078 | -0.0985 | 2.88E-07 | IIIVmrMLM | SUG |
| GW_18JY | AX-110955782 | 6D | 457299721 | 0.0362 | | 3.20E-07 | BLUPmrMLM | SUG |
| GW_18JY | AX-109298719 | 7A | 78843415 | 0.0020 | -0.0808 | 2.59E-09 | IIIVmrMLM | SIG |
| GW_18JY | AX-110686637 | 7A | 130100315 | -0.0082 | 0.0658 | 5.00E-09 | IIIVmrMLM | SIG |
| GW_18JY | AX-109392740 | 7A | 136163106 | -0.0223 | | 1.12E-05 | BLUPmrMLM | SUG |
| GW_18JY | AX-109982504 | 7A | 523194047 | 0.0475 | | 7.10E-07 | BLUPmrMLM | SUG |
| GW_18JY | AX-111078799 | 7A | 679865448 | -0.0065 | -0.1559 | 1.75E-12 | IIIVmrMLM | SIG |
| GW_18JY | AX-109815745 | 7B | 252686467 | -0.0227 | | 4.69E-10 | IIIVmrMLM | SIG |
| GW_18JY | AX-109995248 | 7B | 329513938 | 0.0038 | 0.0674 | 3.18E-09 | IIIVmrMLM | SIG |

| | | | | | | | | |
|---|---|---|---|---|---|---|---|---|
| GW_18JY | AX-86166823 | 7B | 657933319 | 0.0268 | | 7.25E-07 | BLUPmrMLM | SUG |
| GW_18JY | AX-111007605 | 7D | 65009144 | 0.0879 | | 6.04E-18 | BLUPmrMLM | SIG |
| GW_18JY | AX-111007605 | 7D | 65009144 | -0.0793 | | 8.43E-08 | EMMAX | SIG |
| GW_18JY | AX-111007605 | 7D | 65009144 | 0.0796 | | 2.61E-08 | GEMMA | SIG |
| GW_18JY | AX-109666241 | 7D | 66441478 | 0.0671 | | 1.15E-06 | GEMMA | SUG |
| GW_18JY | AX-111184541 | 7D | 66833728 | -0.0756 | | 1.08E-06 | EMMAX | SUG |
| GW_18JY | AX-111184541 | 7D | 66833728 | 0.0763 | | 2.76E-07 | GEMMA | SUG |
| GW_18JY | AX-111843581 | 7D | 67448053 | -0.0812 | | 2.27E-07 | EMMAX | SUG |
| GW_18JY | AX-111843581 | 7D | 67448053 | 0.0814 | | 7.20E-08 | GEMMA | SIG |
| GW_18JY | AX-110387533 | 7D | 131093753 | -0.0361 | | 1.95E-07 | BLUPmrMLM | SUG |
| GW_18JY | AX-111656957 | 7D | 498264066 | -0.0210 | | 3.89E-05 | BLUPmrMLM | SUG |
| GW_18XX | AX-109834552 | 1A | 10425012 | 0.0077 | -0.0754 | 3.36E-08 | IIIVmrMLM | SIG |
| GW_18XX | AX-111186786 | 1A | 518515638 | 0.0984 | | 1.50E-07 | FarmCPU | SUG |
| GW_18XX | AX-110436591 | 1A | 531641304 | -0.0453 | | 2.21E-06 | GEMMA | SUG |
| GW_18XX | AX-111086613 | 1A | 567994548 | 0.0089 | -0.0421 | 4.75E-10 | IIIVmrMLM | SIG |
| GW_18XX | AX-109403122 | 1B | 4443782 | 0.0182 | 0.0096 | 1.42E-11 | IIIVmrMLM | SIG |
| GW_18XX | AX-95242102 | 1B | 90699644 | -0.0401 | 0.0415 | 3.26E-47 | IIIVmrMLM | SIG |
| GW_18XX | AX-94924134 | 1B | 104770438 | 0.0418 | 0.0404 | 2.48E-50 | IIIVmrMLM | SIG |
| GW_18XX | AX-95234705 | 1B | 122968516 | -0.0130 | | 2.86E-07 | IIIVmrMLM | SUG |
| GW_18XX | AX-109843642 | 1B | 394675284 | -0.0252 | | 1.00E-07 | BLUPmrMLM | SIG |
| GW_18XX | AX-110423274 | 1B | 652482645 | 0.0078 | -0.1268 | 3.65E-12 | IIIVmrMLM | SIG |
| GW_18XX | AX-111114795 | 1D | 45065040 | 0.0235 | 0.0177 | 1.30E-17 | IIIVmrMLM | SIG |
| GW_18XX | AX-111194284 | 1D | 67002589 | 0.0141 | | 1.75E-04 | BLUPmrMLM | SUG |
| GW_18XX | AX-109632950 | 1D | 95587796 | -0.0370 | | 2.98E-07 | BLUPmrMLM | SUG |
| GW_18XX | AX-109632950 | 1D | 95587796 | 0.0156 | | 8.81E-10 | IIIVmrMLM | SIG |
| GW_18XX | AX-95152087 | 1D | 459028000 | 0.0257 | | 7.70E-23 | IIIVmrMLM | SIG |
| GW_18XX | AX-111557864 | 2A | 7720330 | -0.0018 | 0.0686 | 2.06E-12 | IIIVmrMLM | SIG |
| GW_18XX | AX-110376679 | 2A | 150121738 | 0.0003 | 0.0626 | 1.15E-13 | IIIVmrMLM | SIG |
| GW_18XX | AX-110936759 | 2A | 767331683 | -0.0153 | | 2.01E-04 | BLUPmrMLM | SUG |
| GW_18XX | AX-108920640 | 2B | 65107033 | -0.0173 | 0.0044 | 1.93E-08 | IIIVmrMLM | SIG |
| GW_18XX | AX-111037038 | 2B | 572269390 | -0.0343 | | 2.06E-11 | BLUPmrMLM | SIG |
| GW_18XX | AX-110629565 | 2B | 609958684 | -0.0205 | | 4.32E-06 | BLUPmrMLM | SUG |
| GW_18XX | AX-111684525 | 2B | 786122899 | 0.0212 | | 5.40E-06 | BLUPmrMLM | SUG |
| GW_18XX | AX-89728114 | 2D | 26344368 | -0.0203 | | 1.13E-06 | BLUPmrMLM | SUG |
| GW_18XX | AX-110400558 | 2D | 118532047 | -0.0347 | | 4.67E-38 | IIIVmrMLM | SIG |
| GW_18XX | AX-110533343 | 2D | 529855718 | -0.0210 | -0.0705 | 5.49E-16 | IIIVmrMLM | SIG |
| GW_18XX | AX-94563003 | 2D | 603139541 | 0.0032 | -0.0245 | 6.18E-05 | IIIVmrMLM | SUG |
| GW_18XX | AX-109505314 | 3A | 11553836 | -0.0079 | 0.0268 | 1.06E-06 | IIIVmrMLM | SUG |
| GW_18XX | AX-109417605 | 3A | 22426877 | -0.0011 | 0.0739 | 5.73E-24 | IIIVmrMLM | SIG |
| GW_18XX | AX-108778023 | 3A | 683114405 | -0.0286 | | 1.03E-08 | BLUPmrMLM | SIG |
| GW_18XX | AX-108778023 | 3A | 683114405 | 0.0076 | -0.0362 | 2.25E-08 | IIIVmrMLM | SIG |
| GW_18XX | AX-110922076 | 3B | 475650825 | 0.0336 | 0.0778 | 2.43E-49 | IIIVmrMLM | SIG |
| GW_18XX | AX-110603940 | 3B | 749990662 | -0.0165 | | 7.61E-05 | BLUPmrMLM | SUG |
| GW_18XX | AX-111756515 | 4A | 13752499 | 0.0091 | 0.0411 | 4.03E-13 | IIIVmrMLM | SIG |

| | | | | | | | | |
|---|---|---|---|---|---|---|---|---|
| GW_18XX | AX-109358749 | 4A | 41030315 | 0.0093 | 0.0358 | 5.44E-07 | IIIVmrMLM | SUG |
| GW_18XX | AX-109272207 | 4A | 133050401 | 0.0090 | -0.0531 | 1.60E-09 | IIIVmrMLM | SIG |
| GW_18XX | AX-109286260 | 4A | 503210659 | -0.0690 | | 2.35E-08 | FarmCPU | SIG |
| GW_18XX | AX-94701059 | 4A | 515660406 | 0.0542 | 0.022 | 8.77E-19 | IIIVmrMLM | SIG |
| GW_18XX | AX-109386882 | 4A | 663440698 | -0.0016 | -0.1018 | 2.13E-11 | IIIVmrMLM | SIG |
| GW_18XX | AX-110364037 | 4B | 37921389 | -0.0148 | | 7.70E-05 | BLUPmrMLM | SUG |
| GW_18XX | AX-94823601 | 4D | 77026743 | 0.1983 | | 3.13E-10 | FarmCPU | SIG |
| GW_18XX | AX-109515348 | 4D | 502034827 | -0.0106 | 0.122 | 9.73E-08 | IIIVmrMLM | SIG |
| GW_18XX | AX-109276523 | 5A | 36328550 | -0.0137 | -0.0383 | 7.93E-20 | IIIVmrMLM | SIG |
| GW_18XX | AX-94443922 | 5A | 578061400 | 0.0294 | | 1.29E-06 | BLUPmrMLM | SUG |
| GW_18XX | AX-109974112 | 5B | 316380076 | 0.0047 | 0.053 | 6.27E-12 | IIIVmrMLM | SIG |
| GW_18XX | AX-109018584 | 5B | 461544397 | -0.0193 | -0.0351 | 2.08E-13 | IIIVmrMLM | SIG |
| GW_18XX | AX-110398849 | 5B | 477507219 | -0.0301 | | 1.90E-04 | BLUPmrMLM | SUG |
| GW_18XX | AX-111449127 | 5B | 477759059 | 0.0275 | | 1.82E-05 | BLUPmrMLM | SUG |
| GW_18XX | AX-111684719 | 5B | 477953826 | -0.0045 | -0.0409 | 3.06E-05 | IIIVmrMLM | SUG |
| GW_18XX | AX-110477945 | 5B | 516214162 | -0.0816 | | 1.83E-07 | FarmCPU | SUG |
| GW_18XX | AX-109325336 | 5B | 586924065 | 0.0033 | 0.0366 | 1.45E-11 | IIIVmrMLM | SIG |
| GW_18XX | AX-108767013 | 5B | 671471152 | 0.0038 | -0.0319 | 5.37E-06 | IIIVmrMLM | SUG |
| GW_18XX | AX-109935774 | 5B | 694009767 | -0.0152 | -0.0413 | 4.24E-09 | IIIVmrMLM | SIG |
| GW_18XX | AX-111165832 | 5D | 367422540 | 0.0358 | 0.0388 | 7.84E-39 | IIIVmrMLM | SIG |
| GW_18XX | AX-111663394 | 6A | 76089539 | 0.0327 | | 7.98E-11 | BLUPmrMLM | SIG |
| GW_18XX | AX-111493035 | 6A | 293145390 | -0.0285 | -0.0038 | 1.53E-24 | IIIVmrMLM | SIG |
| GW_18XX | AX-111171333 | 6A | 545837569 | 0.0060 | -0.0396 | 1.48E-06 | IIIVmrMLM | SUG |
| GW_18XX | AX-110048670 | 6B | 345210653 | -0.0171 | | 1.41E-05 | BLUPmrMLM | SUG |
| GW_18XX | AX-94881276 | 6B | 492492883 | 0.0221 | -0.0714 | 1.35E-18 | IIIVmrMLM | SIG |
| GW_18XX | AX-111496655 | 6B | 591241100 | 0.0243 | | 4.33E-06 | BLUPmrMLM | SUG |
| GW_18XX | AX-111496655 | 6B | 591241100 | -0.0173 | 0.2145 | 5.32E-13 | IIIVmrMLM | SIG |
| GW_18XX | AX-111647379 | 6B | 624798547 | 0.0028 | 0.0634 | 2.12E-08 | IIIVmrMLM | SIG |
| GW_18XX | AX-94788204 | 6D | 2891109 | -0.0289 | | 4.72E-05 | BLUPmrMLM | SUG |
| GW_18XX | AX-94612915 | 6D | 464704396 | 0.0201 | | 5.48E-05 | BLUPmrMLM | SUG |
| GW_18XX | AX-94612915 | 6D | 464704396 | -0.0234 | -0.0042 | 8.81E-09 | IIIVmrMLM | SIG |
| GW_18XX | AX-111164358 | 7A | 89973428 | 0.0191 | | 4.43E-06 | BLUPmrMLM | SUG |
| GW_18XX | AX-111759717 | 7A | 198671663 | -0.0531 | | 1.56E-06 | FarmCPU | SUG |
| GW_18XX | AX-94965000 | 7B | 559146431 | 0.0255 | | 1.66E-05 | BLUPmrMLM | SUG |
| GW_18XX | AX-108732300 | 7B | 605649033 | -0.0007 | 0.0573 | 3.56E-16 | IIIVmrMLM | SIG |
| GW_18XX | AX-108876626 | 7B | 739105825 | 0.0174 | | 1.67E-05 | BLUPmrMLM | SUG |
| GW_18ZMD | AX-94599240 | 1A | 57397380 | -0.0981 | | 1.55E-04 | BLUPmrMLM | SUG |
| GW_18ZMD | AX-94595297 | 3B | 696778434 | 0.0656 | | 1.33E-04 | BLUPmrMLM | SUG |
| GW_18ZMD | AX-109436801 | 5A | 579319031 | -0.0178 | 1.0433 | 6.82E-08 | IIIVmrMLM | SIG |
| GW_19XX | AX-110418620 | 1A | 11553367 | -0.0161 | | 1.61E-04 | BLUPmrMLM | SUG |
| GW_19XX | AX-111737467 | 1A | 49095127 | 0.0043 | 0.1661 | 1.07E-05 | IIIVmrMLM | SUG |
| GW_19XX | AX-111551807 | 1B | 17885289 | 0.0156 | | 7.45E-05 | BLUPmrMLM | SUG |
| GW_19XX | AX-109882166 | 1B | 22123302 | -0.0224 | 0.0649 | 4.52E-14 | IIIVmrMLM | SIG |
| GW_19XX | AX-110953597 | 1B | 303664637 | -0.0429 | | 3.88E-08 | BLUPmrMLM | SIG |

| | | | | | | | | |
|---|---|---|---|---|---|---|---|---|
| GW_19XX | AX-110933719 | 1B | 643323713 | -0.0015 | 0.0432 | 7.80E-09 | IIIVmrMLM | SIG |
| GW_19XX | AX-110580330 | 1B | 647287399 | -0.0258 | -0.0407 | 2.66E-17 | IIIVmrMLM | SIG |
| GW_19XX | AX-110581400 | 1D | 479582629 | -0.0275 | | 1.34E-04 | BLUPmrMLM | SUG |
| GW_19XX | AX-109873296 | 2A | 68474281 | -0.0521 | | 2.09E-06 | GEMMA | SUG |
| GW_19XX | AX-108766887 | 2A | 68811399 | -0.0529 | | 1.62E-06 | GEMMA | SUG |
| GW_19XX | AX-108885595 | 2A | 69310224 | -0.0535 | | 2.13E-06 | GEMMA | SUG |
| GW_19XX | AX-111053248 | 2A | 71359445 | 0.0523 | | 7.26E-07 | EMMAX | SUG |
| GW_19XX | AX-111053248 | 2A | 71359445 | -0.0523 | | 4.95E-07 | GEMMA | SUG |
| GW_19XX | AX-109487855 | 2A | 178428180 | 0.0213 | 0.0263 | 5.87E-12 | IIIVmrMLM | SIG |
| GW_19XX | AX-94588403 | 2A | 476226618 | 0.4424 | | 1.01E-06 | EMMAX | SUG |
| GW_19XX | AX-94578630 | 2A | 615742043 | 0.4424 | | 1.01E-06 | EMMAX | SUG |
| GW_19XX | AX-108766462 | 2A | 646850152 | -0.0964 | | 5.93E-07 | EMMAX | SUG |
| GW_19XX | AX-108766462 | 2A | 646850152 | 0.0957 | | 2.80E-07 | GEMMA | SUG |
| GW_19XX | AX-109413019 | 2A | 765543864 | -0.0419 | | 1.59E-06 | FarmCPU | SUG |
| GW_19XX | AX-109997877 | 2A | 772968383 | -0.0006 | -0.1219 | 6.11E-08 | IIIVmrMLM | SIG |
| GW_19XX | AX-94478206 | 2B | 19415103 | -0.0394 | | 9.07E-08 | BLUPmrMLM | SIG |
| GW_19XX | AX-111652786 | 2B | 44902446 | -0.0062 | -0.0764 | 4.97E-10 | IIIVmrMLM | SIG |
| GW_19XX | AX-95137042 | 2B | 631801052 | 0.1147 | | 1.36E-07 | EMMAX | SUG |
| GW_19XX | AX-95137042 | 2B | 631801052 | -0.1142 | | 8.57E-08 | GEMMA | SIG |
| GW_19XX | AX-110948588 | 2B | 786699693 | 0.0341 | | 4.92E-07 | BLUPmrMLM | SUG |
| GW_19XX | AX-94918279 | 2D | 17605421 | 0.0301 | -0.0025 | 5.44E-10 | IIIVmrMLM | SIG |
| GW_19XX | AX-94501483 | 2D | 102543433 | 0.4866 | | 7.29E-07 | EMMAX | SUG |
| GW_19XX | AX-109115125 | 2D | 524347420 | -0.0546 | | 8.11E-07 | EMMAX | SUG |
| GW_19XX | AX-109115125 | 2D | 524347420 | 0.0548 | | 4.20E-07 | GEMMA | SUG |
| GW_19XX | AX-109864938 | 2D | 524465433 | 0.0520 | | 2.11E-06 | GEMMA | SUG |
| GW_19XX | AX-111494833 | 2D | 526125751 | -0.0548 | | 8.88E-07 | EMMAX | SUG |
| GW_19XX | AX-111494833 | 2D | 526125751 | 0.0549 | | 5.01E-07 | GEMMA | SUG |
| GW_19XX | AX-111455970 | 2D | 527684581 | -0.0569 | | 3.89E-07 | EMMAX | SUG |
| GW_19XX | AX-111455970 | 2D | 527684581 | 0.0571 | | 1.95E-07 | GEMMA | SUG |
| GW_19XX | AX-109905424 | 2D | 527924859 | -0.0554 | | 7.57E-07 | EMMAX | SUG |
| GW_19XX | AX-109905424 | 2D | 527924859 | 0.0555 | | 3.85E-07 | GEMMA | SUG |
| GW_19XX | AX-109783146 | 2D | 528197326 | -0.0563 | | 5.49E-07 | EMMAX | SUG |
| GW_19XX | AX-109783146 | 2D | 528197326 | 0.0564 | | 2.70E-07 | GEMMA | SUG |
| GW_19XX | AX-111838051 | 2D | 528599809 | -0.0565 | | 4.28E-07 | EMMAX | SUG |
| GW_19XX | AX-111838051 | 2D | 528599809 | 0.0565 | | 2.38E-07 | GEMMA | SUG |
| GW_19XX | AX-111491026 | 2D | 529540670 | -0.0556 | | 1.41E-06 | EMMAX | SUG |
| GW_19XX | AX-111491026 | 2D | 529540670 | 0.0556 | | 9.35E-07 | GEMMA | SUG |
| GW_19XX | AX-110002863 | 2D | 530092677 | 0.0548 | | 1.71E-06 | GEMMA | SUG |
| GW_19XX | AX-110007515 | 2D | 532168715 | -0.0558 | | 1.99E-06 | EMMAX | SUG |
| GW_19XX | AX-110007515 | 2D | 532168715 | 0.0559 | | 1.02E-06 | GEMMA | SUG |
| GW_19XX | AX-109645731 | 2D | 533202879 | -0.0549 | | 1.92E-06 | EMMAX | SUG |
| GW_19XX | AX-109645731 | 2D | 533202879 | 0.0550 | | 9.35E-07 | GEMMA | SUG |
| GW_19XX | AX-94593438 | 2D | 651610906 | -0.0742 | | 1.78E-06 | FarmCPU | SUG |
| GW_19XX | AX-108745355 | 3A | 550292294 | -0.0295 | -0.0707 | 1.38E-23 | IIIVmrMLM | SIG |

| | | | | | | | | |
|---|---|---|---|---|---|---|---|---|
| GW_19XX | AX-109000915 | 3B | 344983164 | -0.0108 | -0.0859 | 1.05E-11 | IIIVmrMLM | SIG |
| GW_19XX | AX-109359048 | 3B | 595556323 | -0.0223 | | 7.65E-05 | BLUPmrMLM | SUG |
| GW_19XX | AX-110123177 | 3B | 758202413 | -0.0489 | | 9.22E-07 | FarmCPU | SUG |
| GW_19XX | AX-111506118 | 3B | 776357065 | -0.0010 | 0.0985 | 6.09E-04 | IIIVmrMLM | SUG |
| GW_19XX | AX-110015231 | 3D | 23555106 | -0.6186 | | 1.58E-06 | EMMAX | SUG |
| GW_19XX | AX-95130892 | 3D | 143957102 | -0.0259 | | 6.71E-18 | IIIVmrMLM | SIG |
| GW_19XX | AX-110983530 | 4A | 24901573 | 0.0820 | | 2.93E-09 | FarmCPU | SIG |
| GW_19XX | AX-109920329 | 4B | 139621756 | -0.1582 | | 2.08E-06 | EMMAX | SUG |
| GW_19XX | AX-109920329 | 4B | 139621756 | 0.1546 | | 1.88E-06 | GEMMA | SUG |
| GW_19XX | AX-109920329 | 4B | 139621756 | 0.0151 | 0.1439 | 1.26E-19 | IIIVmrMLM | SIG |
| GW_19XX | AX-109961307 | 4B | 247563885 | -0.0628 | -0.0377 | 9.64E-66 | IIIVmrMLM | SIG |
| GW_19XX | AX-108745161 | 4B | 643490681 | 0.0185 | | 1.22E-04 | BLUPmrMLM | SUG |
| GW_19XX | AX-110487810 | 4B | 659055657 | 0.0067 | 0.2017 | 5.66E-10 | IIIVmrMLM | SIG |
| GW_19XX | AX-111022427 | 4D | 137024457 | -0.6710 | | 3.24E-07 | EMMAX | SUG |
| GW_19XX | AX-110601834 | 4D | 308655751 | -0.4949 | | 5.34E-08 | EMMAX | SIG |
| GW_19XX | AX-94696492 | 4D | 466131867 | 0.3218 | | 5.42E-07 | FarmCPU | SUG |
| GW_19XX | AX-94404950 | 4D | 497036741 | -0.0006 | 0.241 | 4.74E-10 | IIIVmrMLM | SIG |
| GW_19XX | AX-110560415 | 5A | 152687523 | -0.0299 | -0.1974 | 1.07E-29 | IIIVmrMLM | SIG |
| GW_19XX | AX-108848326 | 5A | 395682258 | 0.0067 | 0.1358 | 1.56E-15 | IIIVmrMLM | SIG |
| GW_19XX | AX-110125815 | 5A | 526247469 | 0.0213 | | 2.51E-05 | BLUPmrMLM | SUG |
| GW_19XX | AX-111764163 | 5A | 584395322 | -0.0015 | -0.1332 | 1.51E-06 | IIIVmrMLM | SUG |
| GW_19XX | AX-109470637 | 5B | 362847828 | 0.0496 | | 1.56E-06 | FarmCPU | SUG |
| GW_19XX | AX-109305733 | 5B | 435668404 | 0.0586 | | 1.45E-06 | FarmCPU | SUG |
| GW_19XX | AX-111684719 | 5B | 477953826 | -0.0247 | -0.0735 | 8.54E-23 | IIIVmrMLM | SIG |
| GW_19XX | AX-111684648 | 5B | 520178595 | 0.1022 | | 6.52E-07 | FarmCPU | SUG |
| GW_19XX | AX-111171810 | 5B | 589375693 | 0.0338 | | 1.01E-09 | BLUPmrMLM | SIG |
| GW_19XX | AX-109331761 | 5B | 592858122 | -0.0261 | | 6.56E-06 | BLUPmrMLM | SUG |
| GW_19XX | AX-110989430 | 5B | 646240562 | -0.0038 | -0.0956 | 3.48E-05 | IIIVmrMLM | SUG |
| GW_19XX | AX-108976584 | 5B | 680811883 | 0.0022 | -0.2028 | 7.05E-06 | IIIVmrMLM | SUG |
| GW_19XX | AX-94474207 | 5D | 488428762 | -0.0175 | | 3.10E-09 | IIIVmrMLM | SIG |
| GW_19XX | AX-111595127 | 6A | 291076212 | 0.1726 | | 1.07E-09 | FarmCPU | SIG |
| GW_19XX | AX-108817811 | 6B | 55343145 | -0.0391 | -0.0119 | 6.13E-31 | IIIVmrMLM | SIG |
| GW_19XX | AX-110467577 | 6B | 127324922 | -0.0032 | 0.0839 | 5.76E-07 | IIIVmrMLM | SUG |
| GW_19XX | AX-111555407 | 6B | 631848292 | 0.0055 | -0.0798 | 4.11E-10 | IIIVmrMLM | SIG |
| GW_19XX | AX-108912664 | 6B | 711707364 | -0.0050 | 0.0675 | 3.81E-06 | IIIVmrMLM | SUG |
| GW_19XX | AX-110558093 | 6D | 315986033 | 0.0130 | | 1.62E-04 | BLUPmrMLM | SUG |
| GW_19XX | AX-109305993 | 6D | 470387179 | 0.0162 | -0.1401 | 3.79E-16 | IIIVmrMLM | SIG |
| GW_19XX | AX-109338248 | 7A | 708435360 | 0.0255 | | 2.30E-05 | BLUPmrMLM | SUG |
| GW_19XX | AX-108725863 | 7B | 135773615 | 0.0200 | | 1.63E-04 | BLUPmrMLM | SUG |
| GW_19XX | AX-110413774 | 7B | 217481381 | -0.0043 | 0.0356 | 1.15E-07 | IIIVmrMLM | SIG |
| GW_19XX | AX-109193759 | 7D | 81228758 | 0.0292 | | 5.81E-22 | IIIVmrMLM | SIG |
| GW_19XX | AX-110110668 | 7D | 119568417 | -0.6710 | | 3.24E-07 | EMMAX | SUG |
| GW_19XX | AX-94669864 | 7D | 140320279 | -0.6710 | | 3.24E-07 | EMMAX | SUG |
| GW_19XX | AX-94687235 | 7D | 140320388 | -0.6710 | | 3.24E-07 | EMMAX | SUG |

| | | | | | | | | |
|---|---|---|---|---|---|---|---|---|
| GW_19XX | AX-94950388 | 7D | 140320392 | -0.4871 | | 7.04E-07 | EMMAX | SUG |
| GW_19XX | AX-109365787 | 7D | 140321161 | -0.6710 | | 3.24E-07 | EMMAX | SUG |
| GW_19XX | AX-111536967 | 7D | 140328640 | -0.6710 | | 3.24E-07 | EMMAX | SUG |
| GW_19XX | AX-111053722 | 7D | 140329511 | -0.6710 | | 3.24E-07 | EMMAX | SUG |
| GW_19XX | AX-94737486 | 7D | 550214563 | 0.0839 | | 6.89E-07 | FarmCPU | SUG |
| GW_BLUP | AX-110710487 | 1A | 291982509 | 0.0435 | | 8.39E-07 | FarmCPU | SUG |
| GW_BLUP | AX-111614749 | 1A | 439669876 | -0.0150 | -0.0004 | 1.69E-12 | IIIVmrMLM | SIG |
| GW_BLUP | AX-111171313 | 1A | 464909104 | 0.0005 | -0.0285 | 5.17E-05 | IIIVmrMLM | SUG |
| GW_BLUP | AX-111183991 | 1A | 482970383 | -0.0041 | -0.0268 | 1.02E-07 | IIIVmrMLM | SIG |
| GW_BLUP | AX-94691261 | 1A | 505524324 | -0.0087 | | 1.58E-06 | IIIVmrMLM | SUG |
| GW_BLUP | AX-111460358 | 1A | 531593814 | 0.0364 | | 1.68E-70 | IIIVmrMLM | SIG |
| GW_BLUP | AX-108780402 | 1A | 564570992 | 0.0562 | | 6.82E-07 | FarmCPU | SUG |
| GW_BLUP | AX-109427328 | 1A | 564586780 | 0.0825 | | 7.52E-07 | FarmCPU | SUG |
| GW_BLUP | AX-109508933 | 1B | 6937851 | -0.0136 | | 2.96E-06 | BLUPmrMLM | SUG |
| GW_BLUP | AX-94381509 | 1B | 263312785 | -0.0112 | | 7.87E-10 | IIIVmrMLM | SIG |
| GW_BLUP | AX-111451216 | 1B | 637880634 | -0.0109 | | 1.83E-05 | BLUPmrMLM | SUG |
| GW_BLUP | AX-110991342 | 1B | 686855959 | -0.0176 | | 6.47E-06 | BLUPmrMLM | SUG |
| GW_BLUP | AX-111170771 | 2A | 765013989 | 0.0042 | 0.0535 | 2.31E-07 | IIIVmrMLM | SUG |
| GW_BLUP | AX-108773328 | 2A | 766996924 | 0.0031 | -0.0338 | 4.91E-17 | IIIVmrMLM | SIG |
| GW_BLUP | AX-109877161 | 2A | 770510238 | 0.0006 | -0.0735 | 1.31E-12 | IIIVmrMLM | SIG |
| GW_BLUP | AX-108790634 | 2B | 158907312 | -0.0116 | | 2.08E-05 | BLUPmrMLM | SUG |
| GW_BLUP | AX-111208632 | 2B | 245006974 | -0.0150 | | 3.69E-16 | IIIVmrMLM | SIG |
| GW_BLUP | AX-111548943 | 2B | 611309341 | 0.0014 | -0.0286 | 2.85E-06 | IIIVmrMLM | SUG |
| GW_BLUP | AX-94697254 | 2D | 33710135 | 0.0681 | | 1.18E-06 | FarmCPU | SUG |
| GW_BLUP | AX-108993647 | 2D | 157136755 | 0.0153 | | 8.11E-06 | BLUPmrMLM | SUG |
| GW_BLUP | AX-94442651 | 2D | 507050616 | 0.0860 | | 2.08E-07 | FarmCPU | SUG |
| GW_BLUP | AX-111661053 | 2D | 543644929 | -0.0118 | | 1.03E-10 | IIIVmrMLM | SIG |
| GW_BLUP | AX-110511281 | 2D | 599866981 | 0.0096 | -0.0148 | 7.87E-07 | IIIVmrMLM | SUG |
| GW_BLUP | AX-110498729 | 3A | 68418974 | -0.0115 | 0.0035 | 1.29E-08 | IIIVmrMLM | SIG |
| GW_BLUP | AX-110180986 | 3A | 253461561 | -0.0191 | | 5.34E-05 | BLUPmrMLM | SUG |
| GW_BLUP | AX-111532665 | 3B | 235900924 | 0.0009 | 0.0372 | 3.19E-12 | IIIVmrMLM | SIG |
| GW_BLUP | AX-110367865 | 3B | 349977878 | 0.0117 | -0.024 | 1.06E-11 | IIIVmrMLM | SIG |
| GW_BLUP | AX-110442134 | 3B | 400299836 | -0.0059 | 0.2216 | 3.49E-13 | IIIVmrMLM | SIG |
| GW_BLUP | AX-111618044 | 3B | 750117968 | -0.0142 | | 1.55E-05 | BLUPmrMLM | SUG |
| GW_BLUP | AX-94566074 | 3D | 94239601 | 0.0016 | 0.0549 | 8.90E-05 | IIIVmrMLM | SUG |
| GW_BLUP | AX-109438396 | 4A | 91313202 | -0.0079 | 0.0587 | 6.39E-15 | IIIVmrMLM | SIG |
| GW_BLUP | AX-111501520 | 4A | 140362105 | 0.0021 | -0.0304 | 2.42E-06 | IIIVmrMLM | SUG |
| GW_BLUP | AX-109062834 | 4A | 171201070 | 0.0124 | -0.0085 | 1.87E-08 | IIIVmrMLM | SIG |
| GW_BLUP | AX-111010012 | 4A | 673355278 | -0.0001 | 0.0476 | 2.70E-15 | IIIVmrMLM | SIG |
| GW_BLUP | AX-109320186 | 4A | 725494675 | -0.0083 | -0.035 | 3.43E-14 | IIIVmrMLM | SIG |
| GW_BLUP | AX-111582892 | 4B | 298005511 | -0.0073 | 0.0487 | 4.86E-10 | IIIVmrMLM | SIG |
| GW_BLUP | AX-110380040 | 4B | 495141579 | 0.0060 | -0.0473 | 6.14E-09 | IIIVmrMLM | SIG |
| GW_BLUP | AX-110565007 | 4B | 635267501 | -0.0142 | 0.0185 | 8.64E-14 | IIIVmrMLM | SIG |
| GW_BLUP | AX-110604883 | 4D | 177489950 | 0.0826 | | 4.14E-07 | FarmCPU | SUG |

| | | | | | | | | |
|---|---|---|---|---|---|---|---|---|
| GW_BLUP | AX-109944695 | 5A | 11671147 | -0.0099 | -0.0185 | 2.14E-07 | IIIVmrMLM | SUG |
| GW_BLUP | AX-109877098 | 5A | 526589781 | -0.0029 | 0.0271 | 6.30E-10 | IIIVmrMLM | SIG |
| GW_BLUP | AX-89598451 | 5A | 580797853 | 0.0219 | | 1.26E-07 | BLUPmrMLM | SUG |
| GW_BLUP | AX-89598451 | 5A | 580797853 | -0.0273 | | 1.72E-06 | EMMAX | SUG |
| GW_BLUP | AX-89598451 | 5A | 580797853 | 0.0272 | | 7.79E-07 | GEMMA | SUG |
| GW_BLUP | AX-111004666 | 5B | 418015438 | 0.0024 | -0.0534 | 2.82E-06 | IIIVmrMLM | SUG |
| GW_BLUP | AX-109301788 | 5B | 457457841 | -0.0074 | -0.0296 | 1.77E-10 | IIIVmrMLM | SIG |
| GW_BLUP | AX-111213950 | 5B | 502014259 | -0.0036 | -0.0582 | 6.09E-06 | IIIVmrMLM | SUG |
| GW_BLUP | AX-110369150 | 5B | 563135936 | 0.0194 | | 5.33E-07 | BLUPmrMLM | SUG |
| GW_BLUP | AX-109917396 | 5B | 589356944 | -0.0131 | 0.0178 | 5.22E-13 | IIIVmrMLM | SIG |
| GW_BLUP | AX-94704170 | 5D | 550191254 | -0.0037 | 0.0134 | 3.39E-05 | IIIVmrMLM | SUG |
| GW_BLUP | AX-110483958 | 6A | 17937038 | 0.0144 | 0.0115 | 2.57E-13 | IIIVmrMLM | SIG |
| GW_BLUP | AX-109938404 | 6A | 78474027 | 0.0092 | -0.016 | 1.68E-06 | IIIVmrMLM | SUG |
| GW_BLUP | AX-110596509 | 6A | 100303261 | 0.0064 | 0.097 | 2.04E-07 | IIIVmrMLM | SUG |
| GW_BLUP | AX-94614015 | 6A | 259207357 | -0.0060 | -0.0294 | 1.21E-26 | IIIVmrMLM | SIG |
| GW_BLUP | AX-94541154 | 6B | 64183305 | -0.0238 | | 1.03E-05 | BLUPmrMLM | SUG |
| GW_BLUP | AX-111170401 | 6B | 212566503 | -0.0046 | 0.1042 | 2.62E-12 | IIIVmrMLM | SIG |
| GW_BLUP | AX-108887099 | 6B | 276337569 | 0.0002 | 0.0829 | 6.81E-06 | IIIVmrMLM | SUG |
| GW_BLUP | AX-109391962 | 6B | 339268873 | 0.0906 | | 1.27E-07 | FarmCPU | SUG |
| GW_BLUP | AX-111123882 | 6D | 272759669 | 0.0799 | | 1.13E-06 | FarmCPU | SUG |
| GW_BLUP | AX-108905623 | 7A | 22938419 | 0.0118 | | 8.30E-06 | BLUPmrMLM | SUG |
| GW_BLUP | AX-110453704 | 7A | 265761400 | -0.0001 | -0.0464 | 1.06E-06 | IIIVmrMLM | SUG |
| GW_BLUP | AX-111510467 | 7B | 65367921 | -0.0038 | 0.0379 | 1.62E-09 | IIIVmrMLM | SIG |
| GW_BLUP | AX-110737362 | 7D | 101030120 | -0.0049 | -0.0872 | 1.64E-09 | IIIVmrMLM | SIG |
| GW_BLUP | AX-94751715 | 7D | 634695054 | 0.0005 | 0.041 | 5.60E-06 | IIIVmrMLM | SUG |

GW: grain width.

**Supplementary Table S11**. All QTNs identified as associated with ratio of grain width to length in wheat.

| Trait | Marker | Chr | Position | Add.effect | Dom.effect | P-value | Method | Sig. |
|---|---|---|---|---|---|---|---|---|
| GWLR_17AY | AX-109085065 | 1A | 267791375 | 0.0075 | | 6.50E-05 | BLUPmrMLM | SUG |
| GWLR_17AY | AX-111099425 | 1A | 373320690 | 0.0104 | | 2.08E-07 | BLUPmrMLM | SUG |
| GWLR_17AY | AX-109448581 | 1A | 434112696 | -0.0067 | 0.001 | 1.92E-07 | IIIVmrMLM | SUG |
| GWLR_17AY | AX-109401574 | 1B | 675797677 | -0.0069 | | 2.38E-05 | BLUPmrMLM | SUG |
| GWLR_17AY | AX-111045818 | 2A | 70975917 | 0.0004 | -0.0323 | 4.48E-06 | IIIVmrMLM | SUG |
| GWLR_17AY | AX-108810502 | 2A | 520320857 | 0.0027 | -0.0361 | 4.62E-08 | IIIVmrMLM | SIG |
| GWLR_17AY | AX-94668849 | 2B | 701095953 | 0.0086 | | 8.20E-13 | IIIVmrMLM | SIG |
| GWLR_17AY | AX-110735756 | 2B | 754107847 | -0.0031 | -0.0327 | 3.18E-06 | IIIVmrMLM | SUG |
| GWLR_17AY | AX-109343971 | 2B | 754661631 | 0.0093 | -0.0027 | 1.68E-13 | IIIVmrMLM | SIG |
| GWLR_17AY | AX-94804217 | 2D | 8478708 | -0.0006 | 0.01 | 2.98E-08 | IIIVmrMLM | SIG |
| GWLR_17AY | AX-109259942 | 2D | 649743378 | -0.0054 | | 2.53E-05 | BLUPmrMLM | SUG |
| GWLR_17AY | AX-110676576 | 3A | 559000020 | 0.0006 | -0.0186 | 3.77E-06 | IIIVmrMLM | SUG |
| GWLR_17AY | AX-109278562 | 3B | 71131760 | -0.0021 | -0.0198 | 2.60E-13 | IIIVmrMLM | SIG |
| GWLR_17AY | AX-109363484 | 3B | 221341812 | -0.0126 | | 7.46E-06 | BLUPmrMLM | SUG |
| GWLR_17AY | AX-111118034 | 3B | 341060375 | 0.0288 | | 1.41E-06 | GEMMA | SUG |
| GWLR_17AY | AX-111118034 | 3B | 341060375 | -0.0075 | -0.0285 | 1.64E-16 | IIIVmrMLM | SIG |
| GWLR_17AY | AX-110920233 | 3B | 741368334 | -0.0019 | -0.0213 | 2.89E-09 | IIIVmrMLM | SIG |
| GWLR_17AY | AX-108738871 | 4A | 595394490 | 0.0065 | | 4.58E-05 | BLUPmrMLM | SUG |
| GWLR_17AY | AX-111734312 | 4A | 657195042 | -0.0024 | 0.0478 | 3.82E-05 | IIIVmrMLM | SUG |
| GWLR_17AY | AX-110942925 | 4A | 661733742 | 0.0024 | 0.0392 | 1.10E-06 | IIIVmrMLM | SUG |
| GWLR_17AY | AX-109473770 | 4B | 492526207 | 0.0003 | -0.0425 | 7.99E-06 | IIIVmrMLM | SUG |
| GWLR_17AY | AX-109846838 | 4B | 657290599 | 0.0119 | | 1.49E-09 | BLUPmrMLM | SIG |
| GWLR_17AY | AX-109846838 | 4B | 657290599 | 0.0121 | | 8.00E-07 | GEMMA | SUG |
| GWLR_17AY | AX-109311348 | 5A | 11560749 | 0.0001 | 0.0206 | 5.24E-04 | IIIVmrMLM | SUG |
| GWLR_17AY | AX-111659501 | 5A | 529807948 | -0.0183 | -0.0053 | 1.31E-42 | IIIVmrMLM | SIG |
| GWLR_17AY | AX-110512488 | 5A | 531401988 | 0.0190 | 0.0031 | 7.65E-43 | IIIVmrMLM | SIG |
| GWLR_17AY | AX-108948625 | 5A | 617346045 | 0.0118 | | 1.27E-06 | BLUPmrMLM | SUG |
| GWLR_17AY | AX-109018768 | 5B | 616657148 | 0.0001 | 0.043 | 3.27E-08 | IIIVmrMLM | SIG |
| GWLR_17AY | AX-108937910 | 5B | 634934797 | 0.0074 | 0.0081 | 5.84E-09 | IIIVmrMLM | SIG |
| GWLR_17AY | AX-111761006 | 5B | 635620467 | -0.0110 | | 1.03E-06 | GEMMA | SUG |
| GWLR_17AY | AX-89360769 | 5B | 635768143 | -0.0110 | | 1.03E-06 | GEMMA | SUG |
| GWLR_17AY | AX-109846975 | 5B | 696718392 | -0.0013 | 0.0273 | 1.39E-07 | IIIVmrMLM | SUG |
| GWLR_17AY | AX-108751726 | 5D | 507914070 | -0.0080 | | 2.73E-05 | BLUPmrMLM | SUG |
| GWLR_17AY | AX-108959026 | 6A | 97592599 | 0.0075 | | 8.30E-06 | BLUPmrMLM | SUG |
| GWLR_17AY | AX-109515669 | 6A | 523554249 | -0.0013 | -0.0433 | 8.88E-06 | IIIVmrMLM | SUG |
| GWLR_17AY | AX-108787520 | 6B | 125488307 | 0.0082 | | 1.88E-05 | BLUPmrMLM | SUG |
| GWLR_17AY | AX-95117119 | 6B | 236576788 | -0.0061 | | 2.96E-07 | IIIVmrMLM | SUG |
| GWLR_17AY | AX-108894746 | 6B | 424426865 | -0.0020 | 0.0617 | 3.66E-07 | IIIVmrMLM | SUG |
| GWLR_17AY | AX-110448763 | 6B | 450228007 | -0.0007 | 0.1884 | 9.85E-10 | IIIVmrMLM | SIG |
| GWLR_17AY | AX-94975297 | 6B | 659640022 | -0.0084 | | 3.76E-12 | IIIVmrMLM | SIG |
| GWLR_17AY | AX-111516302 | 6B | 660644327 | -0.0017 | 0.0193 | 1.76E-04 | IIIVmrMLM | SUG |

| | | | | | | | | |
|---|---|---|---|---|---|---|---|---|
| GWLR_17AY | AX-110837283 | 6D | 425297368 | 0.0002 | 0.0205 | 1.76E-05 | IIIVmrMLM | SUG |
| GWLR_17AY | AX-110935561 | 7A | 437402981 | 0.0052 | 0.0377 | 2.09E-09 | IIIVmrMLM | SIG |
| GWLR_17AY | AX-95087469 | 7B | 146817012 | 0.0076 | 0.0541 | 1.70E-15 | IIIVmrMLM | SIG |
| GWLR_17AY | AX-111207452 | 7B | 179918520 | -0.0010 | -0.0241 | 2.45E-05 | IIIVmrMLM | SUG |
| GWLR_17AY | AX-109941342 | 7B | 382672646 | 0.0251 | | 8.26E-07 | BLUPmrMLM | SUG |
| GWLR_17AY | AX-111229720 | 7D | 489320042 | 0.0187 | | 1.86E-09 | BLUPmrMLM | SIG |
| GWLR_17SQ | AX-109815199 | 1A | 32553176 | 0.0072 | -0.0117 | 1.03E-20 | IIIVmrMLM | SIG |
| GWLR_17SQ | AX-109918247 | 1A | 505922717 | -0.0001 | 0.0327 | 4.51E-05 | IIIVmrMLM | SUG |
| GWLR_17SQ | AX-111523370 | 1A | 558978393 | -0.0116 | | 1.23E-05 | BLUPmrMLM | SUG |
| GWLR_17SQ | AX-111530819 | 1A | 559030252 | -0.0028 | 0.0143 | 2.93E-13 | IIIVmrMLM | SIG |
| GWLR_17SQ | AX-111801586 | 1A | 578331842 | 0.0067 | | 6.16E-05 | BLUPmrMLM | SUG |
| GWLR_17SQ | AX-109335072 | 1B | 636895338 | -0.0042 | | 1.77E-05 | BLUPmrMLM | SUG |
| GWLR_17SQ | AX-111621453 | 1B | 668652787 | 0.0043 | | 6.58E-05 | BLUPmrMLM | SUG |
| GWLR_17SQ | AX-86165507 | 1D | 616307 | -0.0018 | -0.0077 | 1.32E-11 | IIIVmrMLM | SIG |
| GWLR_17SQ | AX-108844277 | 1D | 13353971 | -0.0077 | | 2.89E-05 | BLUPmrMLM | SUG |
| GWLR_17SQ | AX-111187068 | 1D | 18998107 | 0.0030 | | 5.98E-05 | BLUPmrMLM | SUG |
| GWLR_17SQ | AX-111334634 | 1D | 223370329 | -0.0004 | 0.0314 | 4.13E-06 | IIIVmrMLM | SUG |
| GWLR_17SQ | AX-95123147 | 1D | 465078260 | 0.0348 | 0.0506 | 3.78E-222 | IIIVmrMLM | SIG |
| GWLR_17SQ | AX-94464913 | 2A | 754361180 | -0.0407 | | 2.99E-08 | GEMMA | SIG |
| GWLR_17SQ | AX-109306017 | 2B | 18950926 | -0.0007 | -0.0134 | 4.98E-07 | IIIVmrMLM | SUG |
| GWLR_17SQ | AX-109959348 | 2B | 57146811 | -0.0052 | -0.0638 | 6.69E-19 | IIIVmrMLM | SIG |
| GWLR_17SQ | AX-111113361 | 2B | 155573935 | 0.0036 | 0.0115 | 2.08E-06 | IIIVmrMLM | SUG |
| GWLR_17SQ | AX-109868941 | 2B | 682929028 | 0.0076 | -0.0082 | 6.51E-23 | IIIVmrMLM | SIG |
| GWLR_17SQ | AX-109852161 | 2B | 719169266 | -0.0059 | | 2.82E-05 | BLUPmrMLM | SUG |
| GWLR_17SQ | AX-109852161 | 2B | 719169266 | 0.0078 | -0.0017 | 3.97E-13 | IIIVmrMLM | SIG |
| GWLR_17SQ | AX-110598837 | 2B | 746717981 | 0.0007 | 0.1793 | 2.46E-21 | IIIVmrMLM | SIG |
| GWLR_17SQ | AX-111882497 | 2D | 193161430 | 0.0003 | -0.2885 | 2.78E-48 | IIIVmrMLM | SIG |
| GWLR_17SQ | AX-111762258 | 2D | 328693208 | 0.0042 | | 3.05E-06 | BLUPmrMLM | SUG |
| GWLR_17SQ | AX-110003269 | 3A | 660423901 | 0.0050 | | 3.03E-06 | BLUPmrMLM | SUG |
| GWLR_17SQ | AX-110074147 | 3A | 660853725 | -0.0040 | 0.0038 | 3.53E-07 | IIIVmrMLM | SUG |
| GWLR_17SQ | AX-108917972 | 3A | 686761161 | 0.0007 | -0.0339 | 2.52E-06 | IIIVmrMLM | SUG |
| GWLR_17SQ | AX-108759526 | 3B | 17633131 | 0.0048 | -0.002 | 2.00E-08 | IIIVmrMLM | SIG |
| GWLR_17SQ | AX-111490934 | 3B | 49286684 | -0.0271 | | 7.62E-07 | GEMMA | SUG |
| GWLR_17SQ | AX-110532482 | 3B | 330597973 | -0.0027 | 0.0194 | 6.23E-14 | IIIVmrMLM | SIG |
| GWLR_17SQ | AX-111517177 | 3B | 672777988 | 0.0001 | -0.0217 | 4.71E-07 | IIIVmrMLM | SUG |
| GWLR_17SQ | AX-111002730 | 3B | 739123259 | -0.0105 | | 1.01E-05 | BLUPmrMLM | SUG |
| GWLR_17SQ | AX-108940066 | 3B | 795180455 | -0.0403 | | 1.67E-06 | GEMMA | SUG |
| GWLR_17SQ | AX-111123890 | 3B | 804181056 | -0.0121 | | 6.76E-08 | BLUPmrMLM | SIG |
| GWLR_17SQ | AX-111078608 | 3D | 550456660 | -0.0353 | | 1.40E-07 | GEMMA | SUG |
| GWLR_17SQ | AX-95632946 | 3D | 551998548 | -0.0322 | | 1.21E-06 | GEMMA | SUG |
| GWLR_17SQ | AX-95074536 | 4A | 175855598 | 0.0071 | | 3.63E-05 | BLUPmrMLM | SUG |
| GWLR_17SQ | AX-109970783 | 4A | 631234330 | -0.0047 | 0.0075 | 2.23E-11 | IIIVmrMLM | SIG |
| GWLR_17SQ | AX-108730857 | 4A | 695773000 | -0.0391 | | 5.51E-07 | FarmCPU | SUG |
| GWLR_17SQ | AX-110999700 | 4A | 696005293 | -0.0364 | | 5.93E-07 | FarmCPU | SUG |

| | | | | | | | | |
|---|---|---|---|---|---|---|---|---|
| GWLR_17SQ | AX-94957899 | 4B | 808894 | -0.0081 | | 1.31E-07 | BLUPmrMLM | SUG |
| GWLR_17SQ | AX-109382842 | 4B | 536427404 | -0.0021 | 0.0216 | 4.19E-08 | IIIVmrMLM | SIG |
| GWLR_17SQ | AX-111974781 | 4D | 457509876 | -0.0009 | 0.0181 | 2.51E-13 | IIIVmrMLM | SIG |
| GWLR_17SQ | AX-110409742 | 4D | 501122912 | 0.0062 | | 2.18E-05 | BLUPmrMLM | SUG |
| GWLR_17SQ | AX-110445929 | 5A | 676377 | 0.0169 | | 5.34E-08 | GEMMA | SIG |
| GWLR_17SQ | AX-108793692 | 5A | 689654 | 0.0147 | | 1.96E-06 | GEMMA | SUG |
| GWLR_17SQ | AX-110633655 | 5A | 690369 | 0.0277 | | 2.28E-06 | GEMMA | SUG |
| GWLR_17SQ | AX-109554869 | 5A | 692719 | 0.0113 | | 3.83E-08 | BLUPmrMLM | SIG |
| GWLR_17SQ | AX-109554869 | 5A | 692719 | 0.0166 | | 6.48E-08 | GEMMA | SIG |
| GWLR_17SQ | AX-89542609 | 5A | 695030 | 0.0151 | | 9.35E-07 | GEMMA | SUG |
| GWLR_17SQ | AX-110169646 | 5A | 10556613 | -0.0359 | | 1.26E-07 | GEMMA | SUG |
| GWLR_17SQ | AX-111613760 | 5A | 480784411 | 0.0057 | | 3.52E-05 | BLUPmrMLM | SUG |
| GWLR_17SQ | AX-109977398 | 5B | 495019337 | -0.0065 | | 8.15E-05 | BLUPmrMLM | SUG |
| GWLR_17SQ | AX-111612039 | 5B | 576576939 | -0.0055 | | 3.35E-06 | BLUPmrMLM | SUG |
| GWLR_17SQ | AX-111168362 | 5B | 707174563 | -0.0467 | | 1.71E-06 | GEMMA | SUG |
| GWLR_17SQ | AX-110696546 | 5D | 131254330 | 0.0059 | | 2.77E-15 | IIIVmrMLM | SIG |
| GWLR_17SQ | AX-111540558 | 6A | 85825381 | -0.0073 | -0.0022 | 4.22E-20 | IIIVmrMLM | SIG |
| GWLR_17SQ | AX-109480438 | 6A | 483592148 | -0.0487 | | 4.07E-08 | GEMMA | SIG |
| GWLR_17SQ | AX-108889767 | 6A | 516399422 | 0.0052 | | 5.53E-05 | BLUPmrMLM | SUG |
| GWLR_17SQ | AX-94865371 | 6B | 35823256 | 0.0062 | | 3.98E-06 | BLUPmrMLM | SUG |
| GWLR_17SQ | AX-108751258 | 6B | 133363863 | 0.0071 | | 1.65E-06 | BLUPmrMLM | SUG |
| GWLR_17SQ | AX-110439928 | 6B | 485720321 | 0.0009 | -0.0189 | 8.10E-06 | IIIVmrMLM | SUG |
| GWLR_17SQ | AX-94689724 | 6B | 575638302 | -0.0044 | | 9.67E-05 | BLUPmrMLM | SUG |
| GWLR_17SQ | AX-108787334 | 6B | 697514868 | 0.0073 | | 1.81E-04 | BLUPmrMLM | SUG |
| GWLR_17SQ | AX-110022559 | 6D | 65764837 | -0.0081 | | 2.24E-26 | IIIVmrMLM | SIG |
| GWLR_17SQ | AX-111465035 | 7A | 11266938 | -0.0095 | | 9.07E-11 | BLUPmrMLM | SIG |
| GWLR_17SQ | AX-110534960 | 7A | 708224422 | -0.0061 | | 7.80E-06 | BLUPmrMLM | SUG |
| GWLR_17SQ | AX-110033032 | 7A | 716262864 | 0.0012 | 0.0654 | 4.68E-07 | IIIVmrMLM | SUG |
| GWLR_17SQ | AX-110714393 | 7B | 591673028 | 0.0054 | | 1.11E-05 | BLUPmrMLM | SUG |
| GWLR_17SQ | AX-109599178 | 7B | 629034998 | 0.0023 | 0.0171 | 1.93E-22 | IIIVmrMLM | SIG |
| GWLR_17SQ | AX-108813158 | 7B | 688919772 | -0.0040 | 0.0076 | 5.25E-08 | IIIVmrMLM | SIG |
| GWLR_17SQ | AX-110671782 | 7B | 741939156 | -0.0129 | 0.0008 | 1.29E-37 | IIIVmrMLM | SIG |
| GWLR_17SQ | AX-111613093 | 7D | 19390238 | -0.0098 | | 2.61E-08 | BLUPmrMLM | SIG |
| GWLR_17SQ | AX-109363059 | 7D | 202229258 | 0.0061 | | 6.89E-05 | BLUPmrMLM | SUG |
| GWLR_17SQ | AX-109844669 | 7D | 538895752 | -0.0358 | | 1.78E-08 | GEMMA | SIG |
| GWLR_17XX | AX-94610374 | 1A | 9642956 | -0.0063 | | 3.62E-07 | BLUPmrMLM | SUG |
| GWLR_17XX | AX-94610374 | 1A | 9642956 | 0.0062 | 0.0033 | 1.17E-15 | IIIVmrMLM | SIG |
| GWLR_17XX | AX-111055136 | 1A | 353250144 | -0.0004 | 0.1588 | 7.77E-22 | IIIVmrMLM | SIG |
| GWLR_17XX | AX-108755149 | 1A | 366981972 | 0.0010 | 0.2198 | 7.69E-39 | IIIVmrMLM | SIG |
| GWLR_17XX | AX-108825668 | 1A | 424236416 | 0.0067 | 0.0048 | 7.13E-23 | IIIVmrMLM | SIG |
| GWLR_17XX | AX-111013589 | 1A | 428673616 | -0.1508 | | 3.76E-10 | EMMAX | SIG |
| GWLR_17XX | AX-109289583 | 1A | 557848861 | -0.0044 | 0.0012 | 6.70E-11 | IIIVmrMLM | SIG |
| GWLR_17XX | AX-108758503 | 1B | 87284436 | 0.0040 | -0.0063 | 4.35E-12 | IIIVmrMLM | SIG |
| GWLR_17XX | AX-94879861 | 1B | 188461387 | 0.0306 | | 4.12E-10 | BLUPmrMLM | SIG |

| | | | | | | | | |
|---|---|---|---|---|---|---|---|---|
| GWLR_17XX | AX-94879861 | 1B | 188461387 | 0.0411 | | 7.27E-07 | GEMMA | SUG |
| GWLR_17XX | AX-94700372 | 1B | 188461659 | 0.0411 | | 7.27E-07 | GEMMA | SUG |
| GWLR_17XX | AX-94463026 | 1B | 193287431 | 0.0000 | -0.0188 | 1.91E-05 | IIIVmrMLM | SUG |
| GWLR_17XX | AX-110564293 | 1B | 553167208 | 0.0059 | | 6.66E-09 | BLUPmrMLM | SIG |
| GWLR_17XX | AX-110514172 | 1B | 559333678 | -0.2865 | | 3.12E-21 | EMMAX | SIG |
| GWLR_17XX | AX-111796916 | 1B | 605987118 | 0.0075 | | 1.50E-06 | BLUPmrMLM | SUG |
| GWLR_17XX | AX-108757556 | 1B | 631966587 | 0.0015 | -0.0281 | 4.17E-08 | IIIVmrMLM | SIG |
| GWLR_17XX | AX-110917931 | 1D | 175622526 | -0.2947 | | 2.47E-19 | EMMAX | SIG |
| GWLR_17XX | AX-111801848 | 1D | 200463811 | -0.1484 | | 5.52E-11 | EMMAX | SIG |
| GWLR_17XX | AX-95217624 | 1D | 466493835 | 0.0095 | | 7.06E-44 | IIIVmrMLM | SIG |
| GWLR_17XX | AX-94804524 | 1D | 476569531 | -0.0439 | | 2.20E-06 | EMMAX | SUG |
| GWLR_17XX | AX-94804524 | 1D | 476569531 | 0.0441 | | 1.82E-06 | GEMMA | SUG |
| GWLR_17XX | AX-94804524 | 1D | 476569531 | -0.0088 | | 1.03E-38 | IIIVmrMLM | SIG |
| GWLR_17XX | AX-111609009 | 2A | 190579415 | 0.0000 | 0.0096 | 4.50E-19 | IIIVmrMLM | SIG |
| GWLR_17XX | AX-111019160 | 2A | 594434944 | -0.0060 | | 9.70E-05 | BLUPmrMLM | SUG |
| GWLR_17XX | AX-94452059 | 2A | 692531374 | -0.0250 | | 1.36E-06 | EMMAX | SUG |
| GWLR_17XX | AX-94452059 | 2A | 692531374 | 0.0251 | | 1.29E-06 | GEMMA | SUG |
| GWLR_17XX | AX-108957347 | 2A | 692937617 | -0.0301 | | 5.78E-07 | EMMAX | SUG |
| GWLR_17XX | AX-108957347 | 2A | 692937617 | 0.0303 | | 4.73E-07 | GEMMA | SUG |
| GWLR_17XX | AX-110374373 | 2A | 715380531 | -0.0008 | 0.0167 | 1.01E-06 | IIIVmrMLM | SUG |
| GWLR_17XX | AX-111729936 | 2B | 88975938 | -0.1063 | | 1.12E-07 | EMMAX | SIG |
| GWLR_17XX | AX-110975896 | 2B | 293220306 | -0.0909 | | 4.98E-10 | EMMAX | SIG |
| GWLR_17XX | AX-111467522 | 2B | 546767454 | -0.0054 | 0.0145 | 9.47E-17 | IIIVmrMLM | SIG |
| GWLR_17XX | AX-109392460 | 2B | 682175602 | 0.0019 | 0.0987 | 5.10E-11 | IIIVmrMLM | SIG |
| GWLR_17XX | AX-94808054 | 2B | 686834920 | -0.0063 | | 8.25E-22 | IIIVmrMLM | SIG |
| GWLR_17XX | AX-95116633 | 2B | 744691043 | 0.0044 | | 5.87E-12 | IIIVmrMLM | SIG |
| GWLR_17XX | AX-111165576 | 2B | 751257300 | 0.0172 | | 9.97E-07 | GEMMA | SUG |
| GWLR_17XX | AX-108981879 | 2B | 751297287 | 0.0082 | | 4.63E-07 | BLUPmrMLM | SUG |
| GWLR_17XX | AX-94554516 | 2D | 2363017 | -0.1732 | | 7.84E-14 | EMMAX | SIG |
| GWLR_17XX | AX-108735017 | 2D | 5761352 | -0.1767 | | 3.45E-14 | EMMAX | SIG |
| GWLR_17XX | AX-94581002 | 2D | 201943345 | -0.1811 | | 4.98E-12 | EMMAX | SIG |
| GWLR_17XX | AX-109726263 | 2D | 259479144 | -0.0175 | | 2.02E-06 | EMMAX | SUG |
| GWLR_17XX | AX-109726263 | 2D | 259479144 | 0.0003 | 0.0284 | 3.13E-09 | IIIVmrMLM | SIG |
| GWLR_17XX | AX-110466106 | 2D | 427632009 | -0.1252 | | 6.94E-10 | EMMAX | SIG |
| GWLR_17XX | AX-94591324 | 2D | 473276253 | -0.0934 | | 2.69E-07 | EMMAX | SUG |
| GWLR_17XX | AX-111064992 | 2D | 606650039 | -0.0003 | -0.0318 | 3.73E-12 | IIIVmrMLM | SIG |
| GWLR_17XX | AX-111694324 | 3A | 9546253 | 0.0076 | | 1.19E-07 | BLUPmrMLM | SUG |
| GWLR_17XX | AX-94837598 | 3A | 491184465 | -0.0239 | | 8.64E-07 | BLUPmrMLM | SUG |
| GWLR_17XX | AX-108835108 | 3A | 547369516 | -0.0047 | 0.0155 | 9.22E-17 | IIIVmrMLM | SIG |
| GWLR_17XX | AX-111512877 | 3A | 716564054 | 0.0077 | | 7.02E-06 | BLUPmrMLM | SUG |
| GWLR_17XX | AX-111512877 | 3A | 716564054 | -0.0029 | | 3.65E-06 | IIIVmrMLM | SUG |
| GWLR_17XX | AX-111106184 | 3B | 248317860 | 0.0287 | | 1.43E-06 | FarmCPU | SUG |
| GWLR_17XX | AX-110556790 | 3B | 617698541 | -0.0045 | | 1.57E-05 | BLUPmrMLM | SUG |
| GWLR_17XX | AX-109532821 | 3B | 720236933 | -0.0003 | 0.0069 | 4.11E-05 | IIIVmrMLM | SUG |

| | | | | | | | | |
|---|---|---|---|---|---|---|---|---|
| GWLR_17XX | AX-95099439 | 3B | 768746333 | -0.1423 | | 8.73E-10 | EMMAX | SIG |
| GWLR_17XX | AX-94567805 | 3B | 779542533 | 0.0015 | -0.0456 | 3.03E-09 | IIIVmrMLM | SIG |
| GWLR_17XX | AX-109112674 | 3D | 90767918 | -0.2947 | | 2.47E-19 | EMMAX | SIG |
| GWLR_17XX | AX-95083173 | 3D | 126945778 | -0.2865 | | 3.12E-21 | EMMAX | SIG |
| GWLR_17XX | AX-111046642 | 3D | 143003646 | -0.2809 | | 2.71E-18 | EMMAX | SIG |
| GWLR_17XX | AX-94837065 | 3D | 389076127 | -0.0802 | | 7.60E-07 | EMMAX | SUG |
| GWLR_17XX | AX-94945211 | 4A | 21210752 | 0.0892 | | 1.78E-07 | FarmCPU | SUG |
| GWLR_17XX | AX-94758025 | 4A | 38840188 | -0.2947 | | 2.47E-19 | EMMAX | SIG |
| GWLR_17XX | AX-109820200 | 4A | 73548087 | -0.0051 | 0.0038 | 3.16E-13 | IIIVmrMLM | SIG |
| GWLR_17XX | AX-95015737 | 4A | 346060786 | -0.0446 | | 3.40E-07 | EMMAX | SUG |
| GWLR_17XX | AX-95015737 | 4A | 346060786 | 0.0477 | | 4.15E-07 | GEMMA | SUG |
| GWLR_17XX | AX-109305727 | 4A | 616674326 | 0.0067 | | 3.32E-05 | BLUPmrMLM | SUG |
| GWLR_17XX | AX-110364963 | 4A | 633372083 | 0.0185 | 0.0106 | ######## | IIIVmrMLM | SIG |
| GWLR_17XX | AX-110633996 | 4A | 636102418 | 0.0039 | | 5.68E-05 | BLUPmrMLM | SUG |
| GWLR_17XX | AX-111523628 | 4B | 265309614 | -0.0186 | | 7.79E-07 | EMMAX | SUG |
| GWLR_17XX | AX-111523628 | 4B | 265309614 | 0.0193 | | 1.20E-06 | GEMMA | SUG |
| GWLR_17XX | AX-109032341 | 4B | 295963534 | -0.0446 | | 3.40E-07 | EMMAX | SUG |
| GWLR_17XX | AX-109032341 | 4B | 295963534 | 0.0477 | | 4.15E-07 | GEMMA | SUG |
| GWLR_17XX | AX-109383374 | 4B | 296406286 | -0.0446 | | 3.40E-07 | EMMAX | SUG |
| GWLR_17XX | AX-109383374 | 4B | 296406286 | 0.0477 | | 4.15E-07 | GEMMA | SUG |
| GWLR_17XX | AX-109622186 | 4B | 297349753 | 0.0098 | | 5.06E-06 | BLUPmrMLM | SUG |
| GWLR_17XX | AX-109622186 | 4B | 297349753 | -0.0206 | | 1.15E-07 | EMMAX | SIG |
| GWLR_17XX | AX-109622186 | 4B | 297349753 | 0.0218 | | 1.79E-07 | GEMMA | SUG |
| GWLR_17XX | AX-108968956 | 4B | 298179735 | -0.0152 | | 1.59E-06 | EMMAX | SUG |
| GWLR_17XX | AX-108968956 | 4B | 298179735 | 0.0156 | | 2.19E-06 | GEMMA | SUG |
| GWLR_17XX | AX-111533236 | 4B | 299257109 | -0.0394 | | 1.93E-06 | EMMAX | SUG |
| GWLR_17XX | AX-94477702 | 4B | 300055236 | -0.0411 | | 1.47E-06 | EMMAX | SUG |
| GWLR_17XX | AX-94477702 | 4B | 300055236 | 0.0441 | | 1.86E-06 | GEMMA | SUG |
| GWLR_17XX | AX-95159519 | 4B | 300055714 | -0.0540 | | 1.17E-07 | EMMAX | SUG |
| GWLR_17XX | AX-95159519 | 4B | 300055714 | 0.0562 | | 1.79E-07 | GEMMA | SUG |
| GWLR_17XX | AX-94722710 | 4B | 300810457 | -0.0453 | | 3.03E-07 | EMMAX | SUG |
| GWLR_17XX | AX-94722710 | 4B | 300810457 | 0.0495 | | 3.72E-07 | GEMMA | SUG |
| GWLR_17XX | AX-110534731 | 4B | 302558741 | -0.0180 | | 1.05E-06 | EMMAX | SUG |
| GWLR_17XX | AX-110534731 | 4B | 302558741 | 0.0188 | | 1.53E-06 | GEMMA | SUG |
| GWLR_17XX | AX-108888815 | 4B | 346030545 | -0.0545 | | 2.45E-07 | EMMAX | SUG |
| GWLR_17XX | AX-95180258 | 4B | 475168218 | -0.0011 | 0.0177 | 1.04E-04 | IIIVmrMLM | SUG |
| GWLR_17XX | AX-111467588 | 4B | 534727035 | 0.0045 | 0.0146 | 1.18E-11 | IIIVmrMLM | SIG |
| GWLR_17XX | AX-94581617 | 4B | 560014564 | -0.2947 | | 2.47E-19 | EMMAX | SIG |
| GWLR_17XX | AX-109560277 | 4D | 113169995 | -0.2809 | | 2.71E-18 | EMMAX | SIG |
| GWLR_17XX | AX-94572577 | 4D | 139210415 | 0.0044 | | 6.81E-12 | IIIVmrMLM | SIG |
| GWLR_17XX | AX-111879954 | 4D | 150774424 | -0.1174 | | 1.21E-08 | EMMAX | SIG |
| GWLR_17XX | AX-109098724 | 4D | 158293010 | -0.0727 | | 8.08E-07 | EMMAX | SUG |
| GWLR_17XX | AX-111259868 | 4D | 165577168 | -0.2809 | | 2.71E-18 | EMMAX | SIG |
| GWLR_17XX | AX-112287379 | 4D | 241796879 | -0.0411 | | 1.47E-06 | EMMAX | SUG |

| | | | | | | | | |
|---|---|---|---|---|---|---|---|---|
| GWLR_17XX | AX-112287379 | 4D | 241796879 | 0.0441 | | 1.86E-06 | GEMMA | SUG |
| GWLR_17XX | AX-109501527 | 4D | 246375877 | -0.2865 | | 3.12E-21 | EMMAX | SIG |
| GWLR_17XX | AX-95114857 | 4D | 305984617 | 0.0050 | | 2.69E-05 | BLUPmrMLM | SUG |
| GWLR_17XX | AX-110334400 | 4D | 465319486 | -0.0077 | | 1.85E-05 | BLUPmrMLM | SUG |
| GWLR_17XX | AX-109801669 | 4D | 498976584 | -0.0036 | 0.01 | 1.12E-07 | IIIVmrMLM | SIG |
| GWLR_17XX | AX-110409742 | 4D | 501122912 | -0.0045 | -0.0032 | 2.98E-11 | IIIVmrMLM | SIG |
| GWLR_17XX | AX-110415726 | 4D | 502594399 | -0.1320 | | 1.52E-09 | EMMAX | SIG |
| GWLR_17XX | AX-109472629 | 5A | 44991597 | 0.0047 | | 8.36E-05 | BLUPmrMLM | SUG |
| GWLR_17XX | AX-109315829 | 5A | 596386006 | 0.0004 | 0.0435 | 1.89E-07 | IIIVmrMLM | SUG |
| GWLR_17XX | AX-110925012 | 5A | 657111831 | -0.0047 | | 1.99E-06 | BLUPmrMLM | SUG |
| GWLR_17XX | AX-94798669 | 5B | 454895066 | -0.1305 | | 1.34E-08 | EMMAX | SIG |
| GWLR_17XX | AX-111470748 | 5B | 555215171 | 0.0085 | | 1.41E-06 | BLUPmrMLM | SUG |
| GWLR_17XX | AX-111819528 | 5B | 594496980 | 0.0052 | | 6.99E-06 | BLUPmrMLM | SUG |
| GWLR_17XX | AX-108789375 | 5B | 605797966 | -0.0044 | | 6.25E-05 | BLUPmrMLM | SUG |
| GWLR_17XX | AX-94615915 | 5B | 658917932 | -0.0673 | | 1.10E-06 | EMMAX | SUG |
| GWLR_17XX | AX-94615915 | 5B | 658917932 | 0.0714 | | 5.43E-07 | GEMMA | SUG |
| GWLR_17XX | AX-111668610 | 5D | 61072160 | -0.0876 | | 2.36E-07 | EMMAX | SUG |
| GWLR_17XX | AX-110051439 | 5D | 72467149 | -0.0873 | | 1.17E-07 | EMMAX | SUG |
| GWLR_17XX | AX-109805442 | 5D | 76042021 | -0.1053 | | 3.47E-08 | EMMAX | SIG |
| GWLR_17XX | AX-111939538 | 5D | 160286069 | -0.1039 | | 1.27E-06 | EMMAX | SUG |
| GWLR_17XX | AX-109214449 | 5D | 291032010 | -0.1156 | | 1.05E-09 | EMMAX | SIG |
| GWLR_17XX | AX-109482637 | 5D | 396416138 | -0.2947 | | 2.47E-19 | EMMAX | SIG |
| GWLR_17XX | AX-110976243 | 5D | 443882218 | -0.0725 | -0.0897 | 0.00E+00 | IIIVmrMLM | SIG |
| GWLR_17XX | AX-109308225 | 5D | 456705502 | -0.0491 | | 2.28E-08 | EMMAX | SIG |
| GWLR_17XX | AX-109308225 | 5D | 456705502 | 0.0493 | | 2.33E-08 | GEMMA | SIG |
| GWLR_17XX | AX-111734641 | 5D | 562100952 | 0.0062 | | 7.32E-06 | BLUPmrMLM | SUG |
| GWLR_17XX | AX-95199226 | 6A | 125497524 | -0.2809 | | 2.71E-18 | EMMAX | SIG |
| GWLR_17XX | AX-110592354 | 6A | 351999178 | -0.0055 | | 3.27E-05 | BLUPmrMLM | SUG |
| GWLR_17XX | AX-109444511 | 6B | 77775216 | -0.0047 | -0.0007 | 4.96E-11 | IIIVmrMLM | SIG |
| GWLR_17XX | AX-110573643 | 6B | 646579323 | -0.0407 | | 8.07E-07 | EMMAX | SUG |
| GWLR_17XX | AX-110573643 | 6B | 646579323 | 0.0407 | | 4.71E-07 | GEMMA | SUG |
| GWLR_17XX | AX-108928674 | 6D | 8248078 | -0.0036 | | 1.54E-04 | BLUPmrMLM | SUG |
| GWLR_17XX | AX-94996931 | 6D | 16295781 | 0.0077 | | 1.57E-05 | BLUPmrMLM | SUG |
| GWLR_17XX | AX-94965223 | 6D | 97015100 | -0.1527 | | 8.74E-11 | EMMAX | SIG |
| GWLR_17XX | AX-109091335 | 6D | 109485655 | -0.2809 | | 2.71E-18 | EMMAX | SIG |
| GWLR_17XX | AX-108815731 | 6D | 127137109 | -0.2809 | | 2.71E-18 | EMMAX | SIG |
| GWLR_17XX | AX-110567255 | 6D | 328832605 | 0.0948 | | 8.14E-07 | FarmCPU | SUG |
| GWLR_17XX | AX-110952407 | 6D | 428638114 | -0.2809 | | 2.71E-18 | EMMAX | SIG |
| GWLR_17XX | AX-110454170 | 7A | 8375277 | -0.0013 | -0.157 | 3.88E-22 | IIIVmrMLM | SIG |
| GWLR_17XX | AX-111631588 | 7A | 41798967 | 0.0042 | | 1.51E-04 | BLUPmrMLM | SUG |
| GWLR_17XX | AX-110957702 | 7A | 81138369 | 0.0082 | | 8.84E-05 | BLUPmrMLM | SUG |
| GWLR_17XX | AX-111594470 | 7A | 81231649 | -0.0043 | 0.0027 | 1.28E-09 | IIIVmrMLM | SIG |
| GWLR_17XX | AX-111630457 | 7A | 128827864 | 0.0007 | 0.0136 | 9.34E-09 | IIIVmrMLM | SIG |
| GWLR_17XX | AX-111504797 | 7A | 683987459 | 0.0031 | -0.0178 | 5.61E-11 | IIIVmrMLM | SIG |

| | | | | | | | | |
|---|---|---|---|---|---|---|---|---|
| GWLR_17XX | AX-111119424 | 7A | 708549994 | 0.0413 | | 1.10E-06 | FarmCPU | SUG |
| GWLR_17XX | AX-110462755 | 7A | 714044893 | -0.0011 | 0.0258 | 4.16E-06 | IIIVmrMLM | SUG |
| GWLR_17XX | AX-94778006 | 7B | 660247200 | 0.0086 | | 5.68E-37 | IIIVmrMLM | SIG |
| GWLR_17XX | AX-108756319 | 7B | 679786935 | -0.2809 | | 2.71E-18 | EMMAX | SIG |
| GWLR_17XX | AX-110619032 | 7B | 706862136 | 0.0035 | | 1.25E-04 | BLUPmrMLM | SUG |
| GWLR_17XX | AX-108971956 | 7D | 69282824 | -0.0121 | | 6.84E-65 | IIIVmrMLM | SIG |
| GWLR_17XX | AX-110577815 | 7D | 140321360 | -0.1532 | | 4.17E-10 | EMMAX | SIG |
| GWLR_17XX | AX-111259919 | 7D | 359814722 | -0.1293 | | 8.37E-10 | EMMAX | SIG |
| GWLR_17XX | AX-109632206 | 7D | 372244929 | -0.2809 | | 2.71E-18 | EMMAX | SIG |
| GWLR_17XX | AX-110501789 | 7D | 437915263 | -0.0019 | 0.1589 | 2.02E-23 | IIIVmrMLM | SIG |
| GWLR_17XX | AX-110404977 | 7D | 442309684 | -0.1181 | | 9.07E-10 | EMMAX | SIG |
| GWLR_17XX | AX-95257251 | 7D | 512281202 | -0.2865 | | 3.12E-21 | EMMAX | SIG |
| GWLR_17XX | AX-111066405 | 7D | 592553571 | -0.1230 | | 1.46E-07 | EMMAX | SUG |
| GWLR_17XX | AX-109578957 | 7D | 592618737 | -0.1230 | | 1.46E-07 | EMMAX | SUG |
| GWLR_17XX | AX-110642814 | 7D | 592675625 | -0.1230 | | 1.46E-07 | EMMAX | SUG |
| GWLR_17XX | AX-110074623 | 7D | 592675666 | -0.1230 | | 1.46E-07 | EMMAX | SUG |
| GWLR_17XX | AX-111041751 | 7D | 592763081 | -0.1230 | | 1.46E-07 | EMMAX | SUG |
| GWLR_17XX | AX-110385111 | 7D | 592856686 | -0.1230 | | 1.46E-07 | EMMAX | SUG |
| GWLR_17XX | AX-111656854 | 7D | 597477552 | -0.1634 | | 1.61E-11 | EMMAX | SIG |
| GWLR_17XX | AX-94545540 | 7D | 600453744 | -0.1377 | | 2.03E-09 | EMMAX | SIG |
| GWLR_17XX | AX-95118138 | 7D | 600455779 | -0.1034 | | 1.37E-07 | EMMAX | SUG |
| GWLR_17ZMD | AX-110587466 | 1A | 586771318 | 0.0061 | 0.1084 | 1.65E-08 | IIIVmrMLM | SIG |
| GWLR_17ZMD | AX-94450612 | 1B | 193626164 | -0.0014 | -0.0241 | 4.81E-05 | IIIVmrMLM | SUG |
| GWLR_17ZMD | AX-111106287 | 1D | 20332338 | -0.0045 | 0.0419 | 1.67E-06 | IIIVmrMLM | SUG |
| GWLR_17ZMD | AX-111736577 | 2A | 515324896 | -0.0010 | 0.0622 | 1.73E-05 | IIIVmrMLM | SUG |
| GWLR_17ZMD | AX-111215241 | 2A | 734098919 | 0.0006 | 0.036 | 3.92E-10 | IIIVmrMLM | SIG |
| GWLR_17ZMD | AX-111451722 | 2A | 759647902 | 0.0047 | 0.0742 | 4.34E-07 | IIIVmrMLM | SUG |
| GWLR_17ZMD | AX-108759338 | 2B | 686454964 | -0.0162 | -0.0351 | 1.96E-13 | IIIVmrMLM | SIG |
| GWLR_17ZMD | AX-95244014 | 2B | 737048967 | 0.0115 | | 6.52E-05 | BLUPmrMLM | SUG |
| GWLR_17ZMD | AX-109206489 | 2D | 228209817 | -0.0404 | | 5.91E-07 | BLUPmrMLM | SUG |
| GWLR_17ZMD | AX-94782143 | 2D | 551926536 | 0.0010 | -0.0281 | 7.92E-06 | IIIVmrMLM | SUG |
| GWLR_17ZMD | AX-95188817 | 2D | 638367648 | 0.0005 | -0.0245 | 1.46E-14 | IIIVmrMLM | SIG |
| GWLR_17ZMD | AX-110180986 | 3A | 253461561 | -0.0261 | -0.0525 | 4.03E-46 | IIIVmrMLM | SIG |
| GWLR_17ZMD | AX-94439666 | 3A | 651498746 | -0.0180 | | 3.74E-05 | BLUPmrMLM | SUG |
| GWLR_17ZMD | AX-111124996 | 3B | 11109220 | 0.0731 | | 1.82E-06 | GEMMA | SUG |
| GWLR_17ZMD | AX-109032617 | 3B | 11166445 | 0.0368 | | 2.13E-06 | GEMMA | SUG |
| GWLR_17ZMD | AX-110499341 | 3B | 11305936 | 0.0368 | | 2.13E-06 | GEMMA | SUG |
| GWLR_17ZMD | AX-110571710 | 3B | 13969303 | 0.0370 | | 2.05E-06 | GEMMA | SUG |
| GWLR_17ZMD | AX-109987987 | 3B | 33339246 | -0.0068 | -0.0292 | 1.55E-15 | IIIVmrMLM | SIG |
| GWLR_17ZMD | AX-111583818 | 3B | 564298699 | 0.0020 | -0.0211 | 2.34E-04 | IIIVmrMLM | SUG |
| GWLR_17ZMD | AX-111032304 | 4A | 594658813 | 0.0000 | -0.051 | 1.33E-07 | IIIVmrMLM | SUG |
| GWLR_17ZMD | AX-111479683 | 4A | 668647454 | 0.0166 | | 4.24E-05 | BLUPmrMLM | SUG |
| GWLR_17ZMD | AX-110994389 | 4A | 716791718 | -0.0194 | | 3.45E-06 | BLUPmrMLM | SUG |
| GWLR_17ZMD | AX-110994389 | 4A | 716791718 | 0.0448 | 0.0293 | 1.80E-57 | IIIVmrMLM | SIG |

| | | | | | | | | |
|---|---|---|---|---|---|---|---|---|
| GWLR_17ZMD | AX-94807935 | 4A | 733912425 | -0.0076 | | 6.42E-05 | BLUPmrMLM | SUG |
| GWLR_17ZMD | AX-110942419 | 4B | 535338782 | -0.0034 | 0.0261 | 7.58E-06 | IIIVmrMLM | SUG |
| GWLR_17ZMD | AX-111254684 | 5A | 610211291 | 0.0128 | | 3.11E-07 | BLUPmrMLM | SUG |
| GWLR_17ZMD | AX-110468168 | 5B | 533494422 | 0.0161 | | 7.28E-06 | BLUPmrMLM | SUG |
| GWLR_17ZMD | AX-110426463 | 5B | 684610407 | -0.0163 | | 4.65E-06 | BLUPmrMLM | SUG |
| GWLR_17ZMD | AX-111486410 | 6A | 3008615 | 0.0023 | 0.0245 | 6.15E-05 | IIIVmrMLM | SUG |
| GWLR_17ZMD | AX-89705852 | 6A | 3288377 | -0.0117 | | 1.05E-04 | BLUPmrMLM | SUG |
| GWLR_17ZMD | AX-94522668 | 6A | 614491460 | -0.0001 | 0.0386 | 1.72E-05 | IIIVmrMLM | SUG |
| GWLR_17ZMD | AX-109299489 | 6B | 518027353 | -0.0144 | -0.0459 | 3.46E-15 | IIIVmrMLM | SIG |
| GWLR_17ZMD | AX-111692750 | 6B | 667963982 | -0.0051 | -0.0295 | 2.38E-05 | IIIVmrMLM | SUG |
| GWLR_17ZMD | AX-111213544 | 7A | 13909034 | -0.0031 | -0.0346 | 2.65E-10 | IIIVmrMLM | SIG |
| GWLR_17ZMD | AX-110425902 | 7A | 14290080 | 0.0127 | | 6.84E-06 | BLUPmrMLM | SUG |
| GWLR_17ZMD | AX-110367912 | 7A | 33058374 | 0.0106 | | 1.50E-04 | BLUPmrMLM | SUG |
| GWLR_17ZMD | AX-111290109 | 7A | 646677262 | -0.0144 | | 5.97E-05 | BLUPmrMLM | SUG |
| GWLR_17ZMD | AX-109860542 | 7B | 3865196 | -0.0034 | -0.0625 | 5.51E-08 | IIIVmrMLM | SIG |
| GWLR_17ZMD | AX-110910992 | 7B | 686142950 | -0.0192 | -0.0045 | 1.74E-12 | IIIVmrMLM | SIG |
| GWLR_17ZMD | AX-109894909 | 7B | 686660826 | -0.0120 | | 4.26E-05 | BLUPmrMLM | SUG |
| GWLR_17ZMD | AX-94876950 | 7B | 708953325 | 0.0074 | 0.0676 | 5.18E-19 | IIIVmrMLM | SIG |
| GWLR_17ZMD | AX-109319113 | 7B | 709199006 | 0.0089 | | 7.25E-05 | BLUPmrMLM | SUG |
| GWLR_17ZMD | AX-111007948 | 7B | 712896019 | 0.0109 | | 4.87E-06 | BLUPmrMLM | SUG |
| GWLR_17ZMD | AX-94712028 | 7B | 733596226 | 0.0008 | 0.0838 | 1.11E-08 | IIIVmrMLM | SIG |
| GWLR_17ZMD | AX-110426870 | 7B | 734135345 | -0.0080 | 0.0998 | 2.75E-07 | IIIVmrMLM | SUG |
| GWLR_17ZMD | AX-110079643 | 7D | 99958426 | -0.0191 | -0.1104 | 1.48E-22 | IIIVmrMLM | SIG |
| GWLR_18AY | AX-109477406 | 1A | 121874362 | 0.0135 | | 2.30E-05 | BLUPmrMLM | SUG |
| GWLR_18AY | AX-111821585 | 1D | 468321305 | 0.0106 | | 7.30E-05 | BLUPmrMLM | SUG |
| GWLR_18AY | AX-94463012 | 2A | 238530713 | -0.0086 | | 5.03E-05 | BLUPmrMLM | SUG |
| GWLR_18AY | AX-94415912 | 2A | 768546472 | -0.0177 | | 1.81E-07 | BLUPmrMLM | SUG |
| GWLR_18AY | AX-111029439 | 2D | 632289579 | -0.0077 | | 1.30E-04 | BLUPmrMLM | SUG |
| GWLR_18AY | AX-111584482 | 3A | 613176720 | -0.0102 | | 2.40E-05 | BLUPmrMLM | SUG |
| GWLR_18AY | AX-108785400 | 3B | 42157700 | 0.0088 | | 1.71E-06 | BLUPmrMLM | SUG |
| GWLR_18AY | AX-109373029 | 5A | 11149914 | -0.0107 | | 1.39E-06 | BLUPmrMLM | SUG |
| GWLR_18AY | AX-108925480 | 6B | 668323785 | -0.0097 | | 1.14E-04 | BLUPmrMLM | SUG |
| GWLR_18AY | AX-109322729 | 7D | 25416415 | -0.0116 | 0.1502 | 4.20E-19 | IIIVmrMLM | SIG |
| GWLR_18JY | AX-108939528 | 1A | 435452647 | 0.0056 | | 1.40E-05 | BLUPmrMLM | SUG |
| GWLR_18JY | AX-108877137 | 1A | 438306058 | -0.0049 | 0.0034 | 1.61E-11 | IIIVmrMLM | SIG |
| GWLR_18JY | AX-94603312 | 1A | 497227861 | -0.0324 | | 2.14E-06 | GEMMA | SUG |
| GWLR_18JY | AX-94603312 | 1A | 497227861 | 0.0140 | | 1.20E-73 | IIIVmrMLM | SIG |
| GWLR_18JY | AX-109272292 | 1A | 534945325 | 0.0061 | | 6.12E-05 | BLUPmrMLM | SUG |
| GWLR_18JY | AX-109314743 | 1B | 4223111 | 0.0050 | | 2.07E-06 | BLUPmrMLM | SUG |
| GWLR_18JY | AX-109839498 | 1B | 198552272 | 0.0098 | | 2.23E-06 | FarmCPU | SUG |
| GWLR_18JY | AX-94618671 | 1D | 9762532 | 0.0061 | | 1.41E-18 | IIIVmrMLM | SIG |
| GWLR_18JY | AX-94811311 | 1D | 16380996 | -0.0255 | | 2.07E-10 | FarmCPU | SIG |
| GWLR_18JY | AX-111606028 | 1D | 413326536 | 0.0029 | -0.0293 | 7.52E-10 | IIIVmrMLM | SIG |
| GWLR_18JY | AX-111133786 | 2A | 443765558 | -0.0055 | | 7.73E-16 | IIIVmrMLM | SIG |

| | | | | | | | | |
|---|---|---|---|---|---|---|---|---|
| GWLR_18JY | AX-111190966 | 2A | 543567658 | -0.0025 | -0.0102 | 5.53E-18 | IIIVmrMLM | SIG |
| GWLR_18JY | AX-110375794 | 2A | 750754084 | 0.0039 | | 6.89E-05 | BLUPmrMLM | SUG |
| GWLR_18JY | AX-111518998 | 2B | 18950849 | 0.0073 | | 5.98E-07 | BLUPmrMLM | SUG |
| GWLR_18JY | AX-111515897 | 2B | 182019414 | -0.0053 | -0.0064 | 4.31E-14 | IIIVmrMLM | SIG |
| GWLR_18JY | AX-95184473 | 2D | 618149805 | 0.0056 | | 2.20E-16 | IIIVmrMLM | SIG |
| GWLR_18JY | AX-95168570 | 2D | 635667657 | -0.0102 | | 4.25E-07 | FarmCPU | SUG |
| GWLR_18JY | AX-108945354 | 3A | 574283260 | 0.0059 | -0.0016 | 2.29E-15 | IIIVmrMLM | SIG |
| GWLR_18JY | AX-94779233 | 3A | 682977788 | -0.0043 | | 2.09E-10 | IIIVmrMLM | SIG |
| GWLR_18JY | AX-110532482 | 3B | 330597973 | -0.0033 | 0.0127 | 8.95E-11 | IIIVmrMLM | SIG |
| GWLR_18JY | AX-110379827 | 3B | 333704358 | 0.0128 | | 7.07E-07 | BLUPmrMLM | SUG |
| GWLR_18JY | AX-109582763 | 3B | 469652148 | 0.0082 | | 2.06E-08 | BLUPmrMLM | SIG |
| GWLR_18JY | AX-109582763 | 3B | 469652148 | -0.0044 | 0.0031 | 1.07E-08 | IIIVmrMLM | SIG |
| GWLR_18JY | AX-111092550 | 4A | 590620068 | -0.0323 | | 1.48E-08 | FarmCPU | SIG |
| GWLR_18JY | AX-94790047 | 4A | 595847438 | 0.0037 | | 4.26E-08 | IIIVmrMLM | SIG |
| GWLR_18JY | AX-111257442 | 4A | 695385272 | -0.0041 | | 9.46E-05 | BLUPmrMLM | SUG |
| GWLR_18JY | AX-111257442 | 4A | 695385272 | 0.0046 | -0.0024 | 6.85E-10 | IIIVmrMLM | SIG |
| GWLR_18JY | AX-111527822 | 4A | 707831647 | 0.0067 | -0.0053 | 4.10E-19 | IIIVmrMLM | SIG |
| GWLR_18JY | AX-109295896 | 4A | 726439863 | 0.0041 | 0.013 | 6.59E-10 | IIIVmrMLM | SIG |
| GWLR_18JY | AX-108906159 | 4B | 13947313 | 0.0087 | | 4.71E-07 | BLUPmrMLM | SUG |
| GWLR_18JY | AX-110070095 | 4B | 32251278 | -0.0058 | | 2.85E-05 | BLUPmrMLM | SUG |
| GWLR_18JY | AX-110070095 | 4B | 32251278 | 0.0051 | -0.0013 | 4.22E-08 | IIIVmrMLM | SIG |
| GWLR_18JY | AX-108802832 | 4B | 658950619 | 0.0265 | | 2.69E-09 | FarmCPU | SIG |
| GWLR_18JY | AX-108906113 | 4B | 665289833 | -0.0071 | | 6.85E-07 | BLUPmrMLM | SUG |
| GWLR_18JY | AX-95248024 | 4D | 67325911 | -0.0063 | | 4.10E-05 | BLUPmrMLM | SUG |
| GWLR_18JY | AX-95248024 | 4D | 67325911 | 0.0100 | 0.0025 | 4.38E-25 | IIIVmrMLM | SIG |
| GWLR_18JY | AX-110587222 | 5A | 331392 | -0.0065 | -0.0125 | 5.35E-20 | IIIVmrMLM | SIG |
| GWLR_18JY | AX-109874921 | 5A | 392831638 | -0.0012 | -0.0127 | 1.41E-05 | IIIVmrMLM | SUG |
| GWLR_18JY | AX-108840641 | 5A | 586673355 | 0.0183 | -0.019 | ######## | IIIVmrMLM | SIG |
| GWLR_18JY | AX-110669567 | 5A | 595089097 | -0.0064 | | 7.38E-06 | BLUPmrMLM | SUG |
| GWLR_18JY | AX-110386045 | 5A | 614997519 | -0.0008 | -0.0464 | 8.82E-08 | IIIVmrMLM | SIG |
| GWLR_18JY | AX-110413599 | 5B | 8090266 | 0.0046 | | 1.28E-11 | IIIVmrMLM | SIG |
| GWLR_18JY | AX-94847013 | 5B | 531612029 | 0.0070 | | 1.36E-07 | BLUPmrMLM | SUG |
| GWLR_18JY | AX-109881727 | 5B | 552787014 | -0.0098 | | 1.97E-07 | BLUPmrMLM | SUG |
| GWLR_18JY | AX-111014460 | 5B | 556173478 | 0.0070 | -0.0053 | 2.95E-22 | IIIVmrMLM | SIG |
| GWLR_18JY | AX-111601445 | 5B | 637836174 | 0.0009 | 0.0196 | 7.85E-07 | IIIVmrMLM | SUG |
| GWLR_18JY | AX-95200157 | 5D | 100126521 | -0.0089 | | 2.35E-05 | BLUPmrMLM | SUG |
| GWLR_18JY | AX-95200157 | 5D | 100126521 | -0.0113 | -0.0246 | 5.33E-78 | IIIVmrMLM | SIG |
| GWLR_18JY | AX-109809706 | 5D | 476819446 | 0.0032 | | 1.03E-05 | BLUPmrMLM | SUG |
| GWLR_18JY | AX-111415450 | 5D | 536756895 | -0.0057 | 0.0011 | 2.41E-15 | IIIVmrMLM | SIG |
| GWLR_18JY | AX-110916616 | 6A | 16415635 | 0.0001 | 0.0255 | 2.24E-08 | IIIVmrMLM | SIG |
| GWLR_18JY | AX-109471106 | 6A | 19041391 | -0.0038 | | 7.87E-06 | BLUPmrMLM | SUG |
| GWLR_18JY | AX-111072344 | 6A | 479725444 | 0.0011 | 0.0112 | 8.46E-07 | IIIVmrMLM | SUG |
| GWLR_18JY | AX-108848119 | 6B | 199078108 | -0.0157 | | 4.47E-07 | FarmCPU | SUG |
| GWLR_18JY | AX-109340938 | 6B | 234420881 | -0.0003 | 0.0127 | 2.26E-06 | IIIVmrMLM | SUG |

| | | | | | | | | |
|---|---|---|---|---|---|---|---|---|
| GWLR_18JY | AX-108793108 | 6B | 643788449 | -0.0042 | | 9.42E-05 | BLUPmrMLM | SUG |
| GWLR_18JY | AX-108766710 | 6B | 706897139 | -0.0045 | | 4.29E-05 | BLUPmrMLM | SUG |
| GWLR_18JY | AX-108766710 | 6B | 706897139 | 0.0067 | -0.0226 | 1.49E-21 | IIIVmrMLM | SIG |
| GWLR_18JY | AX-108928674 | 6D | 8248078 | -0.0001 | -0.035 | 2.46E-09 | IIIVmrMLM | SIG |
| GWLR_18JY | AX-94562028 | 6D | 468842206 | 0.0001 | -0.0401 | 6.37E-12 | IIIVmrMLM | SIG |
| GWLR_18JY | AX-110454170 | 7A | 8375277 | -0.0003 | -0.1436 | 1.98E-16 | IIIVmrMLM | SIG |
| GWLR_18JY | AX-108738966 | 7A | 8898126 | -0.0007 | 0.005 | 1.37E-04 | IIIVmrMLM | SUG |
| GWLR_18JY | AX-110489580 | 7A | 522365900 | -0.0103 | | 1.10E-06 | FarmCPU | SUG |
| GWLR_18JY | AX-109430738 | 7A | 696675658 | -0.0192 | | 2.27E-06 | FarmCPU | SUG |
| GWLR_18JY | AX-94799978 | 7B | 179255425 | -0.0367 | | 1.63E-06 | GEMMA | SUG |
| GWLR_18JY | AX-111636661 | 7B | 209694308 | -0.0023 | 0.0127 | 4.43E-12 | IIIVmrMLM | SIG |
| GWLR_18JY | AX-94563183 | 7B | 354697105 | 0.0289 | | 1.07E-06 | EMMAX | SUG |
| GWLR_18JY | AX-94563183 | 7B | 354697105 | -0.0290 | | 5.49E-07 | GEMMA | SUG |
| GWLR_18JY | AX-94618304 | 7B | 470523833 | -0.0380 | | 1.33E-06 | GEMMA | SUG |
| GWLR_18JY | AX-109913972 | 7B | 482025378 | 0.0403 | | 7.71E-07 | GEMMA | SUG |
| GWLR_18JY | AX-109648945 | 7B | 512475806 | 0.0047 | -0.0309 | 3.05E-15 | IIIVmrMLM | SIG |
| GWLR_18JY | AX-111079170 | 7B | 706604828 | 0.0018 | -0.0125 | 1.26E-10 | IIIVmrMLM | SIG |
| GWLR_18JY | AX-111007605 | 7D | 65009144 | -0.0136 | | 1.81E-06 | EMMAX | SUG |
| GWLR_18JY | AX-111007605 | 7D | 65009144 | 0.0138 | | 5.94E-07 | GEMMA | SUG |
| GWLR_18JY | AX-111843581 | 7D | 67448053 | 0.0140 | | 1.37E-06 | GEMMA | SUG |
| GWLR_18JY | AX-110960135 | 7D | 486590501 | 0.1252 | | 3.53E-08 | EMMAX | SIG |
| GWLR_18XX | AX-110580754 | 1A | 30138311 | -0.0162 | -0.0242 | 1.34E-84 | IIIVmrMLM | SIG |
| GWLR_18XX | AX-109531732 | 1A | 32376278 | -0.0044 | | 1.11E-04 | BLUPmrMLM | SUG |
| GWLR_18XX | AX-110596531 | 1D | 98228 | 0.0056 | | 1.34E-07 | BLUPmrMLM | SUG |
| GWLR_18XX | AX-111195521 | 1D | 10669555 | -0.0051 | | 1.06E-05 | BLUPmrMLM | SUG |
| GWLR_18XX | AX-109947512 | 2A | 23960807 | 0.0036 | -0.1151 | 6.03E-12 | IIIVmrMLM | SIG |
| GWLR_18XX | AX-109816238 | 2A | 485257001 | -0.0069 | | 1.69E-05 | BLUPmrMLM | SUG |
| GWLR_18XX | AX-109899833 | 2A | 509101562 | 0.0077 | 0.02 | 6.90E-24 | IIIVmrMLM | SIG |
| GWLR_18XX | AX-94901744 | 2B | 746037076 | 0.0065 | | 1.95E-15 | IIIVmrMLM | SIG |
| GWLR_18XX | AX-110632670 | 2D | 592784219 | -0.0089 | | 2.43E-06 | BLUPmrMLM | SUG |
| GWLR_18XX | AX-109489505 | 3A | 38238882 | -0.0369 | | 8.21E-08 | FarmCPU | SIG |
| GWLR_18XX | AX-110702581 | 3A | 73564564 | -0.0237 | | 1.35E-06 | FarmCPU | SUG |
| GWLR_18XX | AX-86175243 | 3A | 569427117 | -0.0080 | | 1.83E-07 | BLUPmrMLM | SUG |
| GWLR_18XX | AX-110114596 | 3A | 570798486 | -0.0194 | -0.0305 | 8.84E-98 | IIIVmrMLM | SIG |
| GWLR_18XX | AX-109956562 | 3A | 694685883 | 0.0044 | | 3.39E-06 | BLUPmrMLM | SUG |
| GWLR_18XX | AX-110022644 | 3B | 367481142 | -0.0245 | | 1.64E-06 | GEMMA | SUG |
| GWLR_18XX | AX-110022644 | 3B | 367481142 | 0.0218 | -0.0119 | 5.40E-71 | IIIVmrMLM | SIG |
| GWLR_18XX | AX-110563139 | 3D | 169041250 | 0.0123 | | 5.84E-05 | BLUPmrMLM | SUG |
| GWLR_18XX | AX-94407690 | 4A | 583953494 | 0.0530 | | 8.92E-07 | EMMAX | SUG |
| GWLR_18XX | AX-109303311 | 4D | 502834145 | -0.0040 | | 4.21E-05 | BLUPmrMLM | SUG |
| GWLR_18XX | AX-110669567 | 5A | 595089097 | -0.0054 | | 1.17E-05 | BLUPmrMLM | SUG |
| GWLR_18XX | AX-110616089 | 5B | 458387843 | -0.0042 | | 2.36E-05 | BLUPmrMLM | SUG |
| GWLR_18XX | AX-111729579 | 6A | 2398289 | -0.0073 | | 6.33E-12 | BLUPmrMLM | SIG |
| GWLR_18XX | AX-109856732 | 6B | 476031162 | 0.0045 | | 2.95E-06 | BLUPmrMLM | SUG |

| | | | | | | | | |
|---|---|---|---|---|---|---|---|---|
| GWLR_18XX | AX-110509650 | 6B | 642922912 | 0.0028 | -0.0859 | 3.02E-07 | IIIVmrMLM | SUG |
| GWLR_18XX | AX-94822594 | 6D | 422606296 | -0.0215 | -0.0283 | ######## | IIIVmrMLM | SIG |
| GWLR_18XX | AX-94463922 | 6D | 473075864 | -0.0075 | | 4.24E-08 | BLUPmrMLM | SIG |
| GWLR_18XX | AX-110454170 | 7A | 8375277 | -0.0008 | -0.0859 | 5.35E-05 | IIIVmrMLM | SUG |
| GWLR_18XX | AX-109413541 | 7A | 134416041 | 0.0370 | | 5.87E-07 | EMMAX | SUG |
| GWLR_18XX | AX-109913972 | 7B | 482025378 | 0.0232 | | 1.45E-07 | BLUPmrMLM | SUG |
| GWLR_18XX | AX-110960135 | 7D | 486590501 | 0.1229 | | 9.89E-12 | EMMAX | SIG |
| GWLR_18XX | AX-110504990 | 7D | 637972551 | 0.0342 | -0.0745 | ######## | IIIVmrMLM | SIG |
| GWLR_18ZMD | AX-110431731 | 1A | 9586281 | 0.0052 | -0.0687 | 2.34E-06 | IIIVmrMLM | SUG |
| GWLR_18ZMD | AX-110070596 | 1B | 123962879 | 0.0027 | 0.106 | 1.05E-07 | IIIVmrMLM | SIG |
| GWLR_18ZMD | AX-89596928 | 1D | 412030250 | 0.0071 | | 8.18E-05 | BLUPmrMLM | SUG |
| GWLR_18ZMD | AX-111232822 | 2A | 605245121 | -0.0079 | | 4.26E-05 | BLUPmrMLM | SUG |
| GWLR_18ZMD | AX-109051532 | 2B | 35627353 | -0.0029 | -0.0906 | 4.60E-06 | IIIVmrMLM | SUG |
| GWLR_18ZMD | AX-109894174 | 2D | 369685465 | -0.0085 | 0.0457 | 2.99E-09 | IIIVmrMLM | SIG |
| GWLR_18ZMD | AX-110547501 | 3B | 7147674 | -0.0083 | -0.0967 | 1.28E-09 | IIIVmrMLM | SIG |
| GWLR_18ZMD | AX-111657883 | 3B | 628802454 | 0.0097 | | 2.24E-05 | BLUPmrMLM | SUG |
| GWLR_18ZMD | AX-110035340 | 3B | 704328530 | 0.0064 | -0.0584 | 8.93E-11 | IIIVmrMLM | SIG |
| GWLR_18ZMD | AX-110481013 | 4A | 73180517 | -0.0067 | 0.0686 | 1.24E-09 | IIIVmrMLM | SIG |
| GWLR_18ZMD | AX-111137061 | 4A | 537963260 | 0.0136 | | 6.80E-08 | BLUPmrMLM | SIG |
| GWLR_18ZMD | AX-108908546 | 4B | 601770481 | -0.0086 | | 2.15E-05 | BLUPmrMLM | SUG |
| GWLR_18ZMD | AX-111575370 | 6B | 34934461 | -0.0002 | -0.0621 | 6.80E-04 | IIIVmrMLM | SUG |
| GWLR_18ZMD | AX-110294892 | 6D | 275464980 | 0.0175 | | 2.62E-06 | BLUPmrMLM | SUG |
| GWLR_18ZMD | AX-109281742 | 7A | 723960193 | -0.0062 | 0.1002 | 7.94E-07 | IIIVmrMLM | SUG |
| GWLR_18ZMD | AX-95017844 | 7B | 568488282 | 0.0081 | | 9.32E-05 | BLUPmrMLM | SUG |
| GWLR_18ZMD | AX-108953139 | 7D | 513815828 | -0.0016 | -0.0904 | 1.35E-06 | IIIVmrMLM | SUG |
| GWLR_19XX | AX-109414019 | 1A | 354941498 | -0.0047 | -0.0119 | 4.29E-24 | IIIVmrMLM | SIG |
| GWLR_19XX | AX-111676427 | 1A | 558171385 | -0.0013 | 0.0094 | 1.01E-11 | IIIVmrMLM | SIG |
| GWLR_19XX | AX-94425596 | 1B | 6350605 | 0.0081 | | 9.89E-06 | BLUPmrMLM | SUG |
| GWLR_19XX | AX-110923727 | 1B | 641865930 | -0.0038 | | 1.45E-06 | BLUPmrMLM | SUG |
| GWLR_19XX | AX-110923727 | 1B | 641865930 | 0.0049 | -0.0018 | 2.50E-15 | IIIVmrMLM | SIG |
| GWLR_19XX | AX-111787325 | 2A | 192587807 | -0.0021 | 0.0176 | 5.86E-10 | IIIVmrMLM | SIG |
| GWLR_19XX | AX-108952549 | 2A | 193545114 | 0.0036 | 0.0113 | 3.70E-17 | IIIVmrMLM | SIG |
| GWLR_19XX | AX-109919012 | 2A | 292816265 | -0.0019 | 0.0052 | 2.97E-06 | IIIVmrMLM | SUG |
| GWLR_19XX | AX-94640748 | 2A | 475876556 | 0.0730 | | 9.17E-07 | EMMAX | SUG |
| GWLR_19XX | AX-95629338 | 2A | 735564561 | -0.0012 | -0.009 | 1.33E-18 | IIIVmrMLM | SIG |
| GWLR_19XX | AX-111020262 | 2A | 761486483 | -0.0030 | -0.0181 | 3.42E-19 | IIIVmrMLM | SIG |
| GWLR_19XX | AX-111464039 | 2B | 1509007 | -0.0001 | 0.006 | 2.73E-07 | IIIVmrMLM | SUG |
| GWLR_19XX | AX-94681430 | 2B | 18941839 | 0.0063 | | 3.11E-09 | BLUPmrMLM | SIG |
| GWLR_19XX | AX-110432736 | 2B | 422506230 | -0.0007 | -0.0229 | 2.30E-07 | IIIVmrMLM | SUG |
| GWLR_19XX | AX-108999424 | 2B | 546587812 | 0.0016 | 0.0222 | 8.29E-12 | IIIVmrMLM | SIG |
| GWLR_19XX | AX-111470278 | 2B | 584897538 | -0.0044 | 0.0044 | 7.40E-15 | IIIVmrMLM | SIG |
| GWLR_19XX | AX-109281528 | 2B | 729896769 | 0.0030 | 0.0018 | 3.36E-07 | IIIVmrMLM | SUG |
| GWLR_19XX | AX-108914802 | 2B | 753952960 | -0.0029 | 0.0113 | 6.07E-11 | IIIVmrMLM | SIG |
| GWLR_19XX | AX-111009498 | 2D | 67178390 | 0.0017 | 0.0178 | 2.65E-10 | IIIVmrMLM | SIG |

| | | | | | | | | |
|---|---|---|---|---|---|---|---|---|
| GWLR_19XX | AX-95129757 | 2D | 82227388 | 0.0006 | 0.0079 | 1.93E-10 | IIIVmrMLM | SIG |
| GWLR_19XX | AX-95133267 | 2D | 300437030 | 0.0039 | | 3.22E-05 | BLUPmrMLM | SUG |
| GWLR_19XX | AX-109886994 | 2D | 548974432 | -0.0124 | -0.0641 | 2.10E-92 | IIIVmrMLM | SIG |
| GWLR_19XX | AX-110633040 | 3A | 8327407 | -0.0056 | | 9.89E-06 | BLUPmrMLM | SUG |
| GWLR_19XX | AX-110702581 | 3A | 73564564 | -0.0240 | | 8.21E-07 | FarmCPU | SUG |
| GWLR_19XX | AX-94789190 | 3B | 148999445 | 0.0024 | -0.0041 | 4.15E-08 | IIIVmrMLM | SIG |
| GWLR_19XX | AX-109310034 | 3B | 313175284 | 0.0118 | | 1.07E-07 | BLUPmrMLM | SIG |
| GWLR_19XX | AX-109319585 | 3B | 409819680 | -0.0020 | -0.0127 | 1.05E-07 | IIIVmrMLM | SIG |
| GWLR_19XX | AX-110939022 | 3B | 567246558 | 0.0006 | -0.0253 | 1.17E-08 | IIIVmrMLM | SIG |
| GWLR_19XX | AX-111626425 | 3B | 616168048 | 0.0026 | -0.0067 | 5.11E-06 | IIIVmrMLM | SUG |
| GWLR_19XX | AX-111124226 | 3B | 644325254 | 0.0038 | | 7.91E-05 | BLUPmrMLM | SUG |
| GWLR_19XX | AX-111475434 | 3B | 725566727 | 0.0041 | | 6.16E-06 | BLUPmrMLM | SUG |
| GWLR_19XX | AX-110445450 | 4A | 687159341 | -0.0073 | | 2.67E-06 | BLUPmrMLM | SUG |
| GWLR_19XX | AX-109024221 | 4A | 699901753 | 0.0060 | | 2.15E-05 | BLUPmrMLM | SUG |
| GWLR_19XX | AX-108957818 | 4B | 12146320 | -0.0025 | | 1.30E-04 | BLUPmrMLM | SUG |
| GWLR_19XX | AX-95169614 | 5A | 288906811 | 0.0043 | | 5.17E-15 | IIIVmrMLM | SIG |
| GWLR_19XX | AX-94465397 | 5B | 217913372 | -0.0124 | | 1.42E-11 | BLUPmrMLM | SIG |
| GWLR_19XX | AX-94465397 | 5B | 217913372 | 0.0042 | | 3.26E-14 | IIIVmrMLM | SIG |
| GWLR_19XX | AX-109853735 | 5B | 478924301 | -0.0032 | -0.0062 | 3.78E-09 | IIIVmrMLM | SIG |
| GWLR_19XX | AX-109963680 | 5B | 576924815 | -0.0055 | | 3.27E-06 | BLUPmrMLM | SUG |
| GWLR_19XX | AX-94818739 | 5D | 7412672 | -0.0104 | | 1.36E-05 | BLUPmrMLM | SUG |
| GWLR_19XX | AX-110598930 | 5D | 443881916 | 0.0044 | | 8.66E-16 | IIIVmrMLM | SIG |
| GWLR_19XX | AX-111109562 | 6B | 22244496 | -0.0040 | | 5.71E-07 | BLUPmrMLM | SUG |
| GWLR_19XX | AX-109852941 | 6B | 28981065 | 0.0007 | 0.0228 | 2.80E-10 | IIIVmrMLM | SIG |
| GWLR_19XX | AX-110076928 | 6B | 30064465 | 0.0049 | -0.0007 | 3.60E-15 | IIIVmrMLM | SIG |
| GWLR_19XX | AX-109349909 | 6B | 573302920 | -0.0026 | | 5.87E-05 | BLUPmrMLM | SUG |
| GWLR_19XX | AX-111568425 | 6B | 614953169 | 0.0031 | | 8.16E-05 | BLUPmrMLM | SUG |
| GWLR_19XX | AX-94820365 | 6B | 657090667 | 0.0141 | -0.0263 | 2.59E-110 | IIIVmrMLM | SIG |
| GWLR_19XX | AX-110711998 | 6B | 657203665 | -0.0150 | -0.012 | 2.30E-110 | IIIVmrMLM | SIG |
| GWLR_19XX | AX-111683171 | 6D | 78546393 | -0.0054 | | 1.43E-22 | IIIVmrMLM | SIG |
| GWLR_19XX | AX-111678494 | 6D | 138913532 | -0.1352 | -0.1159 | 0.00E+00 | IIIVmrMLM | SIG |
| GWLR_19XX | AX-109489004 | 6D | 467009560 | -0.0055 | | 1.42E-05 | BLUPmrMLM | SUG |
| GWLR_19XX | AX-109489004 | 6D | 467009560 | 0.0050 | 0.0012 | 6.59E-08 | IIIVmrMLM | SIG |
| GWLR_19XX | AX-111759717 | 7A | 198671663 | -0.0082 | | 3.73E-14 | BLUPmrMLM | SIG |
| GWLR_19XX | AX-108927508 | 7A | 573438061 | -0.0011 | -0.014 | 2.31E-06 | IIIVmrMLM | SUG |
| GWLR_19XX | AX-111002093 | 7B | 135496351 | 0.0064 | 0.0031 | 5.14E-27 | IIIVmrMLM | SIG |
| GWLR_19XX | AX-108725863 | 7B | 135773615 | 0.0083 | | 1.14E-06 | GEMMA | SUG |
| GWLR_19XX | AX-108899392 | 7B | 392035430 | -0.0047 | | 2.90E-06 | BLUPmrMLM | SUG |
| GWLR_19XX | AX-109865611 | 7B | 453339045 | 0.0017 | -0.0115 | 2.30E-06 | IIIVmrMLM | SUG |
| GWLR_19XX | AX-110565424 | 7D | 170281523 | 0.0086 | | 1.35E-06 | BLUPmrMLM | SUG |
| GWLR_19XX | AX-94739446 | 7D | 446989673 | 0.0004 | -0.0644 | 1.08E-10 | IIIVmrMLM | SIG |
| GWLR_BLUP | AX-109531732 | 1A | 32376278 | 0.0040 | -0.0014 | 6.91E-26 | IIIVmrMLM | SIG |
| GWLR_BLUP | AX-108749396 | 1A | 474229744 | 0.0001 | -0.0078 | 5.05E-09 | IIIVmrMLM | SIG |
| GWLR_BLUP | AX-111014568 | 1A | 535855735 | 0.0016 | -0.0105 | 2.43E-08 | IIIVmrMLM | SIG |

| | | | | | | | | |
|---|---|---|---|---|---|---|---|---|
| GWLR_BLUP | AX-109422886 | 1A | 544147942 | -0.0003 | -0.0107 | 3.45E-08 | IIIVmrMLM | SIG |
| GWLR_BLUP | AX-94478603 | 1B | 583686187 | -0.0002 | 0.0104 | 1.20E-05 | IIIVmrMLM | SUG |
| GWLR_BLUP | AX-110923727 | 1B | 641865930 | -0.0019 | | 1.01E-04 | BLUPmrMLM | SUG |
| GWLR_BLUP | AX-110923727 | 1B | 641865930 | 0.0024 | -0.0021 | 4.02E-11 | IIIVmrMLM | SIG |
| GWLR_BLUP | AX-109946817 | 1B | 670803285 | 0.0012 | 0.0123 | 1.40E-09 | IIIVmrMLM | SIG |
| GWLR_BLUP | AX-109860229 | 1D | 2156335 | 0.0026 | | 1.32E-04 | BLUPmrMLM | SUG |
| GWLR_BLUP | AX-109926928 | 1D | 13890082 | -0.0022 | | 1.82E-05 | BLUPmrMLM | SUG |
| GWLR_BLUP | AX-109224093 | 1D | 21878667 | 0.0021 | | 5.20E-10 | IIIVmrMLM | SIG |
| GWLR_BLUP | AX-110352936 | 1D | 427154358 | 0.0001 | -0.0503 | 3.46E-30 | IIIVmrMLM | SIG |
| GWLR_BLUP | AX-111212906 | 2A | 716044360 | 0.0003 | 0.0067 | 6.95E-07 | IIIVmrMLM | SUG |
| GWLR_BLUP | AX-111560255 | 2D | 9080514 | -0.0030 | 0.0225 | 1.52E-23 | IIIVmrMLM | SIG |
| GWLR_BLUP | AX-109286507 | 2D | 21915778 | 0.0000 | 0.0129 | 1.45E-06 | IIIVmrMLM | SUG |
| GWLR_BLUP | AX-108941505 | 2D | 328928390 | -0.0018 | -0.0083 | 9.11E-10 | IIIVmrMLM | SIG |
| GWLR_BLUP | AX-94987429 | 2D | 587166298 | 0.0005 | -0.0176 | 9.55E-13 | IIIVmrMLM | SIG |
| GWLR_BLUP | AX-109489505 | 3A | 38238882 | -0.0203 | | 5.35E-07 | FarmCPU | SUG |
| GWLR_BLUP | AX-111526867 | 3A | 550655603 | 0.0044 | | 1.01E-04 | BLUPmrMLM | SUG |
| GWLR_BLUP | AX-109919341 | 3A | 574323687 | -0.0038 | | 4.44E-07 | BLUPmrMLM | SUG |
| GWLR_BLUP | AX-109371068 | 3A | 697622547 | -0.0017 | | 4.13E-05 | BLUPmrMLM | SUG |
| GWLR_BLUP | AX-110379061 | 3A | 741166296 | 0.0016 | | 1.35E-04 | BLUPmrMLM | SUG |
| GWLR_BLUP | AX-111516534 | 3B | 13251156 | -0.0002 | 0.0057 | 2.45E-06 | IIIVmrMLM | SUG |
| GWLR_BLUP | AX-110610514 | 3B | 107772536 | 0.0018 | | 1.52E-04 | BLUPmrMLM | SUG |
| GWLR_BLUP | AX-111087934 | 3B | 390236503 | -0.0031 | -0.0109 | 2.99E-32 | IIIVmrMLM | SIG |
| GWLR_BLUP | AX-111688132 | 3B | 416086191 | 0.0004 | -0.0059 | 8.66E-16 | IIIVmrMLM | SIG |
| GWLR_BLUP | AX-109341354 | 3B | 465961303 | -0.0032 | | 5.48E-07 | BLUPmrMLM | SUG |
| GWLR_BLUP | AX-110563790 | 3B | 618003208 | 0.0024 | | 6.70E-07 | BLUPmrMLM | SUG |
| GWLR_BLUP | AX-110563790 | 3B | 618003208 | -0.0018 | -0.0007 | 1.51E-06 | IIIVmrMLM | SUG |
| GWLR_BLUP | AX-94709485 | 3D | 361376741 | 0.0106 | 0.0035 | 2.66E-67 | IIIVmrMLM | SIG |
| GWLR_BLUP | AX-110481013 | 4A | 73180517 | -0.0015 | 0.0177 | 9.32E-16 | IIIVmrMLM | SIG |
| GWLR_BLUP | AX-109629235 | 4A | 130216350 | 0.0002 | -0.0051 | 3.08E-07 | IIIVmrMLM | SUG |
| GWLR_BLUP | AX-111519175 | 4A | 590942121 | -0.0028 | | 1.38E-08 | BLUPmrMLM | SIG |
| GWLR_BLUP | AX-111467178 | 4B | 13947273 | 0.0015 | | 6.95E-05 | BLUPmrMLM | SUG |
| GWLR_BLUP | AX-108946619 | 4B | 263254047 | -0.0026 | | 5.35E-15 | IIIVmrMLM | SIG |
| GWLR_BLUP | AX-109843364 | 4B | 414634903 | -0.0004 | 0.0093 | 9.50E-07 | IIIVmrMLM | SUG |
| GWLR_BLUP | AX-109551817 | 4B | 416643468 | -0.0029 | | 3.30E-05 | BLUPmrMLM | SUG |
| GWLR_BLUP | AX-109310564 | 4B | 435640906 | -0.0010 | -0.0277 | 5.07E-19 | IIIVmrMLM | SIG |
| GWLR_BLUP | AX-110036417 | 5A | 35022242 | -0.0023 | | 1.70E-04 | BLUPmrMLM | SUG |
| GWLR_BLUP | AX-95653009 | 5A | 549188587 | 0.0034 | | 7.92E-06 | BLUPmrMLM | SUG |
| GWLR_BLUP | AX-110009183 | 5A | 560339571 | 0.0019 | -0.0071 | 4.35E-12 | IIIVmrMLM | SIG |
| GWLR_BLUP | AX-110465539 | 5B | 49713646 | -0.0030 | | 2.49E-05 | BLUPmrMLM | SUG |
| GWLR_BLUP | AX-110491310 | 5B | 323857400 | -0.0044 | 0.0006 | 1.58E-20 | IIIVmrMLM | SIG |
| GWLR_BLUP | AX-110557214 | 5B | 530738787 | -0.0015 | -0.0125 | 2.09E-09 | IIIVmrMLM | SIG |
| GWLR_BLUP | AX-111612039 | 5B | 576576939 | 0.0031 | 0.0018 | 4.61E-17 | IIIVmrMLM | SIG |
| GWLR_BLUP | AX-109048495 | 5D | 21731782 | -0.0053 | -0.0011 | 2.21E-07 | IIIVmrMLM | SUG |
| GWLR_BLUP | AX-108821106 | 5D | 107147713 | -0.0129 | | 1.37E-06 | FarmCPU | SUG |

| | | | | | | | | |
|---|---|---|---|---|---|---|---|---|
| GWLR_BLUP | AX-110585829 | 5D | 537362100 | 0.0025 | | 2.92E-05 | BLUPmrMLM | SUG |
| GWLR_BLUP | AX-94446985 | 6B | 7137766 | -0.0031 | | 5.61E-06 | BLUPmrMLM | SUG |
| GWLR_BLUP | AX-111633191 | 6B | 95490391 | 0.0388 | | 1.38E-06 | EMMAX | SUG |
| GWLR_BLUP | AX-110945404 | 6B | 625067880 | -0.0042 | | 3.60E-05 | BLUPmrMLM | SUG |
| GWLR_BLUP | AX-110945404 | 6B | 625067880 | 0.0038 | 0.0056 | 2.16E-30 | IIIVmrMLM | SIG |
| GWLR_BLUP | AX-109338425 | 6D | 463760824 | -0.0016 | | 1.65E-04 | BLUPmrMLM | SUG |
| GWLR_BLUP | AX-89511804 | 7A | 79637810 | 0.0022 | | 1.19E-04 | BLUPmrMLM | SUG |
| GWLR_BLUP | AX-108910815 | 7A | 156689806 | 0.0021 | 0.0203 | 9.77E-16 | IIIVmrMLM | SIG |
| GWLR_BLUP | AX-94563183 | 7B | 354697105 | 0.0132 | | 1.49E-06 | EMMAX | SUG |
| GWLR_BLUP | AX-94563183 | 7B | 354697105 | -0.0132 | | 7.16E-07 | GEMMA | SUG |
| GWLR_BLUP | AX-94618304 | 7B | 470523833 | 0.0182 | | 2.07E-06 | EMMAX | SUG |
| GWLR_BLUP | AX-94618304 | 7B | 470523833 | -0.0183 | | 1.10E-06 | GEMMA | SUG |
| GWLR_BLUP | AX-109913972 | 7B | 482025378 | 0.0136 | | 3.29E-11 | BLUPmrMLM | SIG |
| GWLR_BLUP | AX-109913972 | 7B | 482025378 | 0.0184 | | 2.19E-06 | GEMMA | SUG |
| GWLR_BLUP | AX-111987065 | 7D | 267005167 | -0.0006 | 0.0097 | 8.63E-05 | IIIVmrMLM | SUG |
| GWLR_BLUP | AX-110960135 | 7D | 486590501 | 0.0548 | | 3.76E-07 | EMMAX | SUG |

GWLR: ratio of grain width to length

**Supplementary Table S12**. All QTNs identified as associated with thousand grain weight in wheat.

| Trait | Marker | Chr | Position | Add.effect | Dom.effect | P-value | Method | Sig. |
|---|---|---|---|---|---|---|---|---|
| TGW_17AY | AX-110994042 | 1A | 31549776 | 0.1419 | 2.0594 | 9.34E-08 | IIIVmrMLM | SIG |
| TGW_17AY | AX-111590048 | 1A | 38727469 | 0.6075 | 0.0335 | 8.05E-05 | IIIVmrMLM | SUG |
| TGW_17AY | AX-110604530 | 1A | 475742613 | -0.0477 | -1.3367 | 4.74E-06 | IIIVmrMLM | SUG |
| TGW_17AY | AX-109472062 | 1A | 579859461 | -0.8122 | | 7.85E-06 | BLUPmrMLM | SUG |
| TGW_17AY | AX-111515958 | 2A | 70985299 | -1.6436 | | 1.59E-06 | FarmCPU | SUG |
| TGW_17AY | AX-110449519 | 2A | 71051123 | -3.2571 | | 1.68E-06 | FarmCPU | SUG |
| TGW_17AY | AX-109501313 | 2A | 71106087 | -3.2571 | | 1.68E-06 | FarmCPU | SUG |
| TGW_17AY | AX-110988056 | 2A | 71107811 | -1.6436 | | 1.59E-06 | FarmCPU | SUG |
| TGW_17AY | AX-110127392 | 2A | 71119182 | -1.6436 | | 1.59E-06 | FarmCPU | SUG |
| TGW_17AY | AX-110979159 | 2A | 71139554 | -1.6282 | | 1.92E-06 | FarmCPU | SUG |
| TGW_17AY | AX-111115411 | 2A | 71140350 | -1.6436 | | 1.59E-06 | FarmCPU | SUG |
| TGW_17AY | AX-110917681 | 2A | 71145388 | -1.6436 | | 1.59E-06 | FarmCPU | SUG |
| TGW_17AY | AX-111673388 | 2A | 71331270 | -1.6436 | | 1.59E-06 | FarmCPU | SUG |
| TGW_17AY | AX-108944857 | 2A | 71331390 | -1.6880 | | 1.23E-06 | FarmCPU | SUG |
| TGW_17AY | AX-111741806 | 2A | 71331980 | -1.6436 | | 1.59E-06 | FarmCPU | SUG |
| TGW_17AY | AX-110563221 | 2A | 71346290 | -1.7385 | | 3.17E-07 | FarmCPU | SUG |
| TGW_17AY | AX-110626211 | 2A | 71346379 | -1.6352 | | 1.99E-06 | FarmCPU | SUG |
| TGW_17AY | AX-109423823 | 2A | 71351492 | -3.2571 | | 1.68E-06 | FarmCPU | SUG |
| TGW_17AY | AX-111646775 | 2A | 71353445 | -3.2551 | | 1.90E-06 | FarmCPU | SUG |
| TGW_17AY | AX-109872156 | 2A | 71354930 | -1.6436 | | 1.59E-06 | FarmCPU | SUG |
| TGW_17AY | AX-111601526 | 2A | 71364224 | -1.6436 | | 1.59E-06 | FarmCPU | SUG |
| TGW_17AY | AX-111100439 | 2A | 71366142 | -1.7929 | | 5.19E-07 | FarmCPU | SUG |
| TGW_17AY | AX-109867877 | 2A | 71382389 | -1.6436 | | 1.59E-06 | FarmCPU | SUG |
| TGW_17AY | AX-111491711 | 2A | 71387666 | -1.7263 | | 6.19E-07 | FarmCPU | SUG |
| TGW_17AY | AX-111037204 | 2A | 71392536 | -1.7619 | | 6.52E-07 | FarmCPU | SUG |
| TGW_17AY | AX-111060532 | 2A | 71434236 | -3.3057 | | 1.35E-06 | FarmCPU | SUG |
| TGW_17AY | AX-111513449 | 2A | 71486263 | -3.2027 | | 2.11E-06 | FarmCPU | SUG |
| TGW_17AY | AX-109965157 | 2A | 71612965 | -1.6044 | | 2.24E-06 | FarmCPU | SUG |
| TGW_17AY | AX-110078849 | 2A | 71616345 | -1.6427 | | 1.26E-06 | FarmCPU | SUG |
| TGW_17AY | AX-110064196 | 2A | 71627782 | -3.3025 | | 6.99E-07 | FarmCPU | SUG |
| TGW_17AY | AX-108804889 | 2A | 72607507 | -3.3945 | | 7.51E-07 | FarmCPU | SUG |
| TGW_17AY | AX-94590897 | 2A | 72742731 | -3.3945 | | 7.51E-07 | FarmCPU | SUG |
| TGW_17AY | AX-94400797 | 2A | 72931314 | -3.3416 | | 1.03E-06 | FarmCPU | SUG |
| TGW_17AY | AX-94430919 | 2A | 723098907 | -0.5110 | 1.2338 | 1.99E-11 | IIIVmrMLM | SIG |
| TGW_17AY | AX-94570561 | 2B | 31622205 | 3.4770 | | 2.94E-07 | EMMAX | SUG |
| TGW_17AY | AX-94570561 | 2B | 31622205 | -3.4714 | | 9.68E-08 | GEMMA | SIG |
| TGW_17AY | AX-109520325 | 2B | 32159981 | 2.7933 | | 2.40E-08 | EMMAX | SIG |
| TGW_17AY | AX-109520325 | 2B | 32159981 | -2.7846 | | 1.01E-08 | GEMMA | SIG |
| TGW_17AY | AX-94449117 | 2B | 32884866 | -3.1702 | | 1.10E-06 | GEMMA | SUG |
| TGW_17AY | AX-94769911 | 2B | 32912527 | -3.1572 | | 1.45E-06 | GEMMA | SUG |
| TGW_17AY | AX-109905516 | 2B | 33382382 | 3.5089 | | 3.46E-07 | EMMAX | SUG |

| | | | | | | | | |
|---|---|---|---|---|---|---|---|---|
| TGW_17AY | AX-109905516 | 2B | 33382382 | -3.4981 | | 1.37E-07 | GEMMA | SUG |
| TGW_17AY | AX-109654211 | 2B | 33460387 | 1.8270 | | 1.64E-06 | EMMAX | SUG |
| TGW_17AY | AX-109654211 | 2B | 33460387 | -1.8223 | | 8.49E-07 | GEMMA | SUG |
| TGW_17AY | AX-110059594 | 2B | 33492749 | 1.8270 | | 1.64E-06 | EMMAX | SUG |
| TGW_17AY | AX-110059594 | 2B | 33492749 | -1.8223 | | 8.49E-07 | GEMMA | SUG |
| TGW_17AY | AX-110483781 | 2B | 33513487 | 2.6182 | | 4.42E-08 | EMMAX | SIG |
| TGW_17AY | AX-110483781 | 2B | 33513487 | -2.6103 | | 1.73E-08 | GEMMA | SIG |
| TGW_17AY | AX-111069224 | 2B | 33517434 | -1.7900 | | 1.65E-06 | GEMMA | SUG |
| TGW_17AY | AX-109478745 | 2B | 141328725 | 1.0559 | | 2.65E-05 | BLUPmrMLM | SUG |
| TGW_17AY | AX-109478745 | 2B | 141328725 | -0.6338 | | 2.07E-08 | IIIVmrMLM | SIG |
| TGW_17AY | AX-111067313 | 3A | 31949850 | 0.7311 | -2.4976 | 2.86E-11 | IIIVmrMLM | SIG |
| TGW_17AY | AX-110055912 | 3A | 599867639 | 0.2029 | -1.0781 | 2.29E-07 | IIIVmrMLM | SUG |
| TGW_17AY | AX-109435843 | 3A | 645097084 | -0.8651 | | 2.30E-06 | GEMMA | SUG |
| TGW_17AY | AX-89450957 | 3A | 645097367 | -0.8851 | | 1.39E-06 | GEMMA | SUG |
| TGW_17AY | AX-89450957 | 3A | 645097367 | 0.4415 | 2.1231 | 1.53E-05 | IIIVmrMLM | SUG |
| TGW_17AY | AX-111556318 | 3A | 732639440 | -0.6660 | | 1.98E-04 | BLUPmrMLM | SUG |
| TGW_17AY | AX-109582977 | 3B | 7110270 | -0.6816 | -10.9838 | 5.25E-15 | IIIVmrMLM | SIG |
| TGW_17AY | AX-111832753 | 3B | 13807795 | 0.7430 | -0.3397 | 1.40E-05 | IIIVmrMLM | SUG |
| TGW_17AY | AX-109860599 | 3B | 17668406 | 0.0432 | -2.1223 | 3.03E-04 | IIIVmrMLM | SUG |
| TGW_17AY | AX-108796899 | 3B | 383826470 | -0.9607 | -3.0168 | 6.57E-23 | IIIVmrMLM | SIG |
| TGW_17AY | AX-109855276 | 3B | 406663792 | -0.0630 | 3.1684 | 1.30E-07 | IIIVmrMLM | SUG |
| TGW_17AY | AX-110563032 | 3B | 800177834 | 0.3510 | -4.1556 | 1.30E-08 | IIIVmrMLM | SIG |
| TGW_17AY | AX-108893223 | 3D | 5407251 | 0.4370 | 4.0834 | 1.66E-09 | IIIVmrMLM | SIG |
| TGW_17AY | AX-109532774 | 3D | 123688688 | -1.2045 | | 6.45E-05 | BLUPmrMLM | SUG |
| TGW_17AY | AX-94964849 | 3D | 282938013 | 2.6234 | | 5.19E-07 | BLUPmrMLM | SUG |
| TGW_17AY | AX-94964849 | 3D | 282938013 | -3.3017 | | 1.26E-06 | EMMAX | SUG |
| TGW_17AY | AX-94964849 | 3D | 282938013 | 3.2913 | | 5.36E-07 | GEMMA | SUG |
| TGW_17AY | AX-108961317 | 4A | 9905667 | -1.0611 | | 5.01E-05 | BLUPmrMLM | SUG |
| TGW_17AY | AX-108961317 | 4A | 9905667 | 0.6995 | | 6.84E-10 | IIIVmrMLM | SIG |
| TGW_17AY | AX-110613866 | 4A | 11678769 | -0.8442 | | 3.18E-05 | BLUPmrMLM | SUG |
| TGW_17AY | AX-110918202 | 4A | 83319693 | -0.5326 | -2.9283 | 6.30E-12 | IIIVmrMLM | SIG |
| TGW_17AY | AX-111241403 | 4B | 420802503 | 0.3260 | 4.1517 | 8.45E-09 | IIIVmrMLM | SIG |
| TGW_17AY | AX-109412822 | 4B | 637316848 | -0.0731 | -5.5835 | 3.84E-06 | IIIVmrMLM | SUG |
| TGW_17AY | AX-109269882 | 4D | 483601767 | 0.8674 | | 3.25E-14 | IIIVmrMLM | SIG |
| TGW_17AY | AX-111524849 | 5A | 10211267 | -0.3250 | 3.0044 | 7.73E-06 | IIIVmrMLM | SUG |
| TGW_17AY | AX-110566136 | 5A | 583533588 | -0.8800 | | 1.15E-05 | BLUPmrMLM | SUG |
| TGW_17AY | AX-109904805 | 5A | 665887714 | -0.8167 | | 2.81E-06 | BLUPmrMLM | SUG |
| TGW_17AY | AX-111574749 | 5B | 415835414 | -0.1642 | -6.4824 | 8.54E-05 | IIIVmrMLM | SUG |
| TGW_17AY | AX-108773010 | 5B | 557955798 | 0.1508 | -4.8349 | 6.72E-11 | IIIVmrMLM | SIG |
| TGW_17AY | AX-110565030 | 5B | 592803107 | -0.0913 | 1.5371 | 5.99E-07 | IIIVmrMLM | SUG |
| TGW_17AY | AX-110563852 | 5B | 696716309 | 0.1718 | -1.5905 | 2.15E-10 | IIIVmrMLM | SIG |
| TGW_17AY | AX-94914486 | 5D | 188336419 | -0.9763 | | 1.43E-05 | BLUPmrMLM | SUG |
| TGW_17AY | AX-110372528 | 6A | 597948910 | -0.4594 | -0.2007 | 3.06E-04 | IIIVmrMLM | SUG |
| TGW_17AY | AX-109412729 | 6B | 18146046 | 0.2495 | 2.3253 | 7.87E-08 | IIIVmrMLM | SIG |

| | | | | | | | | |
|---|---|---|---|---|---|---|---|---|
| TGW_17AY | AX-111091344 | 6B | 27999926 | 1.8699 | | 8.26E-07 | FarmCPU | SUG |
| TGW_17AY | AX-94956128 | 6B | 560801582 | 0.5913 | -0.2242 | 6.93E-06 | IIIVmrMLM | SUG |
| TGW_17AY | AX-110416142 | 6B | 706649015 | 0.6856 | | 4.54E-05 | BLUPmrMLM | SUG |
| TGW_17AY | AX-111645830 | 7A | 264758772 | 0.6490 | | 5.33E-05 | BLUPmrMLM | SUG |
| TGW_17AY | AX-110724172 | 7D | 438298662 | 0.6870 | | 1.34E-09 | IIIVmrMLM | SIG |
| TGW_17AY | AX-111460719 | 7D | 524913472 | 0.7228 | 7.4313 | 2.87E-22 | IIIVmrMLM | SIG |
| TGW_17JY | AX-110994541 | 1A | 358655877 | 0.0677 | 3.09 | 1.51E-05 | IIIVmrMLM | SUG |
| TGW_17JY | AX-109347609 | 1A | 462803258 | 0.5925 | 4.4768 | 3.48E-12 | IIIVmrMLM | SIG |
| TGW_17JY | AX-110369023 | 1A | 516623756 | 0.4284 | -2.7549 | 1.09E-16 | IIIVmrMLM | SIG |
| TGW_17JY | AX-94425619 | 1A | 545531217 | -30.2810 | | 1.20E-07 | EMMAX | SUG |
| TGW_17JY | AX-109427328 | 1A | 564586780 | 5.3258 | | 7.90E-07 | FarmCPU | SUG |
| TGW_17JY | AX-94525724 | 1A | 564749741 | 10.7443 | | 1.23E-06 | FarmCPU | SUG |
| TGW_17JY | AX-111631366 | 1A | 573715763 | 1.8201 | | 4.39E-08 | BLUPmrMLM | SIG |
| TGW_17JY | AX-111631366 | 1A | 573715763 | -0.9025 | 0.1773 | 2.46E-09 | IIIVmrMLM | SIG |
| TGW_17JY | AX-109965064 | 1A | 580958888 | -0.0803 | -5.948 | 9.94E-09 | IIIVmrMLM | SIG |
| TGW_17JY | AX-95174049 | 1A | 582842690 | -2.4124 | | 1.85E-06 | GEMMA | SUG |
| TGW_17JY | AX-110588448 | 1A | 582858807 | -2.3306 | | 2.09E-06 | GEMMA | SUG |
| TGW_17JY | AX-110563876 | 1A | 582868976 | 5.1923 | | 4.58E-07 | EMMAX | SUG |
| TGW_17JY | AX-110563876 | 1A | 582868976 | -5.1514 | | 3.66E-07 | GEMMA | SUG |
| TGW_17JY | AX-108921894 | 1A | 583128744 | 2.4589 | | 2.22E-06 | EMMAX | SUG |
| TGW_17JY | AX-108921894 | 1A | 583128744 | -2.4422 | | 1.69E-06 | GEMMA | SUG |
| TGW_17JY | AX-108835837 | 1A | 583172068 | -2.4124 | | 1.85E-06 | GEMMA | SUG |
| TGW_17JY | AX-108826270 | 1A | 583246065 | -2.4124 | | 1.85E-06 | GEMMA | SUG |
| TGW_17JY | AX-111627822 | 1A | 583256261 | 2.4407 | | 2.17E-06 | EMMAX | SUG |
| TGW_17JY | AX-111627822 | 1A | 583256261 | -2.4214 | | 1.53E-06 | GEMMA | SUG |
| TGW_17JY | AX-111173346 | 1A | 583287789 | -2.4124 | | 1.85E-06 | GEMMA | SUG |
| TGW_17JY | AX-110463611 | 1A | 583289369 | -2.3861 | | 2.05E-06 | GEMMA | SUG |
| TGW_17JY | AX-109418659 | 1A | 583292411 | 4.8331 | | 1.45E-06 | EMMAX | SUG |
| TGW_17JY | AX-109418659 | 1A | 583292411 | -4.7841 | | 1.19E-06 | GEMMA | SUG |
| TGW_17JY | AX-108887078 | 1A | 584583811 | 0.1187 | -1.8489 | 2.25E-06 | IIIVmrMLM | SUG |
| TGW_17JY | AX-95220493 | 1B | 16850430 | -4.1523 | | 2.28E-06 | EMMAX | SUG |
| TGW_17JY | AX-109402278 | 1B | 118518098 | -0.5807 | -3.3746 | 8.06E-09 | IIIVmrMLM | SIG |
| TGW_17JY | AX-94771017 | 1B | 166078289 | 3.3110 | | 2.28E-06 | GEMMA | SUG |
| TGW_17JY | AX-109442564 | 1B | 374863817 | 0.7851 | 4.2264 | 9.17E-18 | IIIVmrMLM | SIG |
| TGW_17JY | AX-111705530 | 1B | 465975682 | 0.7789 | -0.03 | 3.87E-07 | IIIVmrMLM | SUG |
| TGW_17JY | AX-110149284 | 1B | 472446338 | -1.1995 | | 4.26E-07 | BLUPmrMLM | SUG |
| TGW_17JY | AX-94781792 | 2A | 13849176 | 2.4408 | | 2.92E-05 | BLUPmrMLM | SUG |
| TGW_17JY | AX-110572850 | 2A | 288889596 | 0.2304 | -3.1688 | 3.17E-09 | IIIVmrMLM | SIG |
| TGW_17JY | AX-109893551 | 2A | 748199036 | -0.1210 | -2.6826 | 7.17E-04 | IIIVmrMLM | SUG |
| TGW_17JY | AX-109022397 | 2A | 758561818 | -0.0387 | 2.6469 | 7.49E-11 | IIIVmrMLM | SIG |
| TGW_17JY | AX-109890405 | 2B | 25579188 | -0.8669 | | 7.04E-05 | BLUPmrMLM | SUG |
| TGW_17JY | AX-89454891 | 2B | 106829802 | 1.6335 | | 7.17E-07 | BLUPmrMLM | SUG |
| TGW_17JY | AX-110611059 | 2B | 132410331 | -0.0492 | -5.5678 | 5.72E-15 | IIIVmrMLM | SIG |
| TGW_17JY | AX-111066087 | 2B | 453828149 | 1.0357 | | 6.23E-06 | BLUPmrMLM | SUG |

| | | | | | | | | |
|---|---|---|---|---|---|---|---|---|
| TGW_17JY | AX-111066087 | 2B | 453828149 | -0.9733 | -0.8296 | 9.49E-13 | IIIVmrMLM | SIG |
| TGW_17JY | AX-108862764 | 2B | 606618022 | 0.2593 | -6.1819 | 1.93E-08 | IIIVmrMLM | SIG |
| TGW_17JY | AX-111543331 | 2B | 770436325 | -1.1169 | -0.0272 | 8.98E-09 | IIIVmrMLM | SIG |
| TGW_17JY | AX-109400512 | 2D | 529605010 | 2.3911 | | 1.47E-06 | GEMMA | SUG |
| TGW_17JY | AX-109945455 | 2D | 546174846 | -1.0162 | 2.6377 | 3.97E-14 | IIIVmrMLM | SIG |
| TGW_17JY | AX-94802493 | 2D | 627166647 | -3.6289 | | 2.21E-05 | BLUPmrMLM | SUG |
| TGW_17JY | AX-110363555 | 3A | 17026703 | 0.2959 | 4.1904 | 4.96E-10 | IIIVmrMLM | SIG |
| TGW_17JY | AX-95184677 | 3A | 510007314 | -1.8758 | | 4.24E-07 | BLUPmrMLM | SUG |
| TGW_17JY | AX-95184677 | 3A | 510007314 | 1.3767 | 0.6075 | 1.66E-16 | IIIVmrMLM | SIG |
| TGW_17JY | AX-109821800 | 3B | 264012735 | -0.9274 | 3.8417 | 7.36E-21 | IIIVmrMLM | SIG |
| TGW_17JY | AX-110466741 | 3B | 545010427 | -0.5678 | 4.0836 | 3.66E-32 | IIIVmrMLM | SIG |
| TGW_17JY | AX-109294992 | 3B | 705194695 | -1.4145 | 0.5924 | 4.85E-24 | IIIVmrMLM | SIG |
| TGW_17JY | AX-108801106 | 3B | 734160439 | 0.7455 | -9.1967 | 4.18E-15 | IIIVmrMLM | SIG |
| TGW_17JY | AX-110127873 | 3B | 734485466 | 0.7348 | | 3.68E-05 | BLUPmrMLM | SUG |
| TGW_17JY | AX-110127873 | 3B | 734485466 | -1.1356 | -1.7513 | 1.85E-17 | IIIVmrMLM | SIG |
| TGW_17JY | AX-94888678 | 3D | 3298556 | 2.3010 | 0.5516 | 1.17E-21 | IIIVmrMLM | SIG |
| TGW_17JY | AX-108812655 | 3D | 557465592 | 0.5269 | 10.7941 | 3.70E-19 | IIIVmrMLM | SIG |
| TGW_17JY | AX-111031267 | 4A | 398033542 | -0.7871 | 6.4668 | 3.02E-18 | IIIVmrMLM | SIG |
| TGW_17JY | AX-109270508 | 4B | 643325849 | 0.7733 | | 1.13E-04 | BLUPmrMLM | SUG |
| TGW_17JY | AX-109817614 | 4B | 659590826 | -0.4851 | 1.0066 | 9.59E-06 | IIIVmrMLM | SUG |
| TGW_17JY | AX-109431851 | 4D | 107277810 | -30.2810 | | 1.20E-07 | EMMAX | SUG |
| TGW_17JY | AX-110457081 | 4D | 503958928 | 5.2225 | | 8.86E-07 | GEMMA | SUG |
| TGW_17JY | AX-109541070 | 5A | 549207473 | 0.3113 | 4.0105 | 2.16E-07 | IIIVmrMLM | SUG |
| TGW_17JY | AX-109892293 | 5A | 563481040 | 0.9805 | | 3.43E-05 | BLUPmrMLM | SUG |
| TGW_17JY | AX-109449120 | 5B | 78719818 | 0.4779 | 4.1633 | 5.25E-19 | IIIVmrMLM | SIG |
| TGW_17JY | AX-109872845 | 5B | 486483108 | -0.9773 | | 1.47E-05 | BLUPmrMLM | SUG |
| TGW_17JY | AX-110669023 | 5B | 691863507 | 0.9074 | -0.6545 | 1.12E-10 | IIIVmrMLM | SIG |
| TGW_17JY | AX-110524707 | 5B | 701764232 | -1.0022 | | 1.57E-14 | IIIVmrMLM | SIG |
| TGW_17JY | AX-108986508 | 6A | 595180495 | -1.4829 | 4.8635 | 1.85E-28 | IIIVmrMLM | SIG |
| TGW_17JY | AX-111695521 | 6B | 595295134 | -30.2810 | | 1.20E-07 | EMMAX | SUG |
| TGW_17JY | AX-110962724 | 6B | 669496280 | -0.8861 | 2.4316 | 2.65E-14 | IIIVmrMLM | SIG |
| TGW_17JY | AX-110871088 | 6D | 375214100 | -0.7646 | 1.7733 | 4.60E-09 | IIIVmrMLM | SIG |
| TGW_17JY | AX-109895604 | 7B | 24050619 | 1.0918 | | 3.33E-06 | BLUPmrMLM | SUG |
| TGW_17JY | AX-110978879 | 7B | 233380660 | -1.3308 | -0.9504 | 8.74E-18 | IIIVmrMLM | SIG |
| TGW_17JY | AX-109936890 | 7B | 594951127 | 0.0518 | -1.6139 | 4.46E-05 | IIIVmrMLM | SUG |
| TGW_17JY | AX-110937795 | 7B | 639236098 | 0.8827 | 1.5048 | 2.76E-11 | IIIVmrMLM | SIG |
| TGW_17JY | AX-108762309 | 7B | 654326461 | 0.1612 | -7.0752 | 4.14E-12 | IIIVmrMLM | SIG |
| TGW_17JY | AX-110676928 | 7D | 619079252 | -30.2810 | | 1.20E-07 | EMMAX | SUG |
| TGW_17SQ | AX-109427328 | 1A | 564586780 | 3.5450 | | 1.41E-06 | FarmCPU | SUG |
| TGW_17SQ | AX-110987235 | 1A | 584755997 | -0.8679 | | 1.62E-06 | BLUPmrMLM | SUG |
| TGW_17SQ | AX-109903479 | 1B | 394073619 | 3.7275 | | 4.13E-07 | FarmCPU | SUG |
| TGW_17SQ | AX-94684170 | 1B | 597537011 | 3.5115 | | 7.57E-07 | FarmCPU | SUG |
| TGW_17SQ | AX-111585185 | 1B | 670919594 | -1.3865 | | 6.64E-07 | EMMAX | SUG |
| TGW_17SQ | AX-111585185 | 1B | 670919594 | 1.3861 | | 3.10E-07 | GEMMA | SUG |

| | | | | | | | | |
|---|---|---|---|---|---|---|---|---|
| TGW_17SQ | AX-94790358 | 1D | 11409748 | -0.4372 | | 5.00E-05 | BLUPmrMLM | SUG |
| TGW_17SQ | AX-111441361 | 1D | 391278422 | 1.3596 | | 1.52E-05 | BLUPmrMLM | SUG |
| TGW_17SQ | AX-108748677 | 2A | 1444321 | -0.4974 | | 3.79E-05 | BLUPmrMLM | SUG |
| TGW_17SQ | AX-94707348 | 2A | 535713056 | 1.1370 | | 1.65E-06 | BLUPmrMLM | SUG |
| TGW_17SQ | AX-111566543 | 2A | 602781339 | 0.7584 | | 5.38E-08 | BLUPmrMLM | SIG |
| TGW_17SQ | AX-109536319 | 2A | 608601284 | 0.3849 | | 9.39E-05 | BLUPmrMLM | SUG |
| TGW_17SQ | AX-111513541 | 2A | 753092366 | 0.4206 | | 1.60E-04 | BLUPmrMLM | SUG |
| TGW_17SQ | AX-111021032 | 2B | 780817099 | -0.4663 | | 3.92E-08 | BLUPmrMLM | SIG |
| TGW_17SQ | AX-89333763 | 2B | 788028356 | -0.4131 | | 2.45E-06 | BLUPmrMLM | SUG |
| TGW_17SQ | AX-94697254 | 2D | 33710135 | 3.0090 | | 1.20E-06 | FarmCPU | SUG |
| TGW_17SQ | AX-94442651 | 2D | 507050616 | 3.9716 | | 5.63E-08 | FarmCPU | SIG |
| TGW_17SQ | AX-110447745 | 3A | 647695584 | -0.3466 | | 1.97E-04 | BLUPmrMLM | SUG |
| TGW_17SQ | AX-108811358 | 3A | 732400432 | -0.3310 | | 3.24E-06 | BLUPmrMLM | SUG |
| TGW_17SQ | AX-110524117 | 3B | 41037082 | 0.2882 | | 6.41E-05 | BLUPmrMLM | SUG |
| TGW_17SQ | AX-109274638 | 3B | 237325305 | -3.1101 | | 3.04E-07 | FarmCPU | SUG |
| TGW_17SQ | AX-111503859 | 3B | 749670621 | -0.8667 | | 2.78E-06 | BLUPmrMLM | SUG |
| TGW_17SQ | AX-109553432 | 3B | 756108575 | 0.6490 | | 1.05E-04 | BLUPmrMLM | SUG |
| TGW_17SQ | AX-109420906 | 4A | 115634351 | 3.5752 | | 5.18E-07 | FarmCPU | SUG |
| TGW_17SQ | AX-94942161 | 4A | 136928119 | 0.7445 | | 1.40E-05 | BLUPmrMLM | SUG |
| TGW_17SQ | AX-95145873 | 4D | 138432501 | -0.3934 | | 1.89E-05 | BLUPmrMLM | SUG |
| TGW_17SQ | AX-110604883 | 4D | 177489950 | 3.7204 | | 2.36E-07 | FarmCPU | SUG |
| TGW_17SQ | AX-109318201 | 4D | 197019433 | 3.5389 | | 7.75E-07 | FarmCPU | SUG |
| TGW_17SQ | AX-110946030 | 4D | 208560799 | 1.1217 | | 1.25E-04 | BLUPmrMLM | SUG |
| TGW_17SQ | AX-110528135 | 4D | 445004978 | 3.4412 | | 8.76E-07 | FarmCPU | SUG |
| TGW_17SQ | AX-111112946 | 5A | 424008295 | 0.9389 | | 6.79E-10 | BLUPmrMLM | SIG |
| TGW_17SQ | AX-109471979 | 5A | 528825589 | -0.9854 | | 8.99E-10 | BLUPmrMLM | SIG |
| TGW_17SQ | AX-109460863 | 5A | 594957849 | -0.4519 | | 2.52E-05 | BLUPmrMLM | SUG |
| TGW_17SQ | AX-111587325 | 5B | 463222171 | 1.7403 | | 6.56E-07 | EMMAX | SUG |
| TGW_17SQ | AX-111587325 | 5B | 463222171 | -1.7377 | | 2.37E-07 | GEMMA | SUG |
| TGW_17SQ | AX-111027966 | 5B | 463222844 | -1.6078 | | 8.61E-07 | GEMMA | SUG |
| TGW_17SQ | AX-109052486 | 5B | 463223004 | 1.3763 | | 1.87E-06 | EMMAX | SUG |
| TGW_17SQ | AX-109052486 | 5B | 463223004 | -1.3835 | | 6.42E-07 | GEMMA | SUG |
| TGW_17SQ | AX-110917592 | 5B | 463244055 | -1.6751 | | 2.27E-06 | GEMMA | SUG |
| TGW_17SQ | AX-109814822 | 5B | 463328296 | -1.3796 | | 2.14E-06 | GEMMA | SUG |
| TGW_17SQ | AX-108838136 | 5B | 463343339 | -1.3796 | | 2.14E-06 | GEMMA | SUG |
| TGW_17SQ | AX-109900805 | 5B | 463526656 | -1.3796 | | 2.14E-06 | GEMMA | SUG |
| TGW_17SQ | AX-110512002 | 5B | 463528661 | -1.3796 | | 2.14E-06 | GEMMA | SUG |
| TGW_17SQ | AX-111038776 | 5B | 463777757 | -3.3607 | | 1.96E-06 | GEMMA | SUG |
| TGW_17SQ | AX-108957255 | 5B | 464384033 | 1.3792 | | 2.23E-06 | GEMMA | SUG |
| TGW_17SQ | AX-108957255 | 5B | 464384033 | -1.5833 | -4.3308 | 7.87E-31 | IIIVmrMLM | SIG |
| TGW_17SQ | AX-110568495 | 5B | 485994328 | -0.5405 | | 2.67E-05 | BLUPmrMLM | SUG |
| TGW_17SQ | AX-111684648 | 5B | 520178595 | 3.4156 | | 1.17E-06 | FarmCPU | SUG |
| TGW_17SQ | AX-111518462 | 5B | 678319620 | 1.1159 | | 6.30E-08 | BLUPmrMLM | SIG |
| TGW_17SQ | AX-109416491 | 5B | 690126821 | 1.6019 | | 1.49E-07 | BLUPmrMLM | SUG |

| | | | | | | | | |
|---|---|---|---|---|---|---|---|---|
| TGW_17SQ | AX-111464653 | 5D | 486138536 | 3.2595 | | 2.07E-06 | FarmCPU | SUG |
| TGW_17SQ | AX-109407131 | 5D | 512550680 | -0.7007 | | 8.97E-09 | BLUPmrMLM | SIG |
| TGW_17SQ | AX-110486124 | 6A | 317789634 | -0.4098 | | 1.33E-04 | BLUPmrMLM | SUG |
| TGW_17SQ | AX-111544007 | 6B | 204513322 | -0.4767 | | 2.00E-05 | BLUPmrMLM | SUG |
| TGW_17SQ | AX-109391962 | 6B | 339268873 | 4.0797 | | 7.04E-08 | FarmCPU | SIG |
| TGW_17SQ | AX-110371715 | 6B | 641868401 | -0.7142 | | 4.76E-05 | BLUPmrMLM | SUG |
| TGW_17SQ | AX-111123882 | 6D | 272759669 | 3.5972 | | 8.02E-07 | FarmCPU | SUG |
| TGW_17SQ | AX-111153914 | 7A | 507877239 | 3.8796 | | 3.95E-07 | FarmCPU | SUG |
| TGW_17SQ | AX-94842225 | 7A | 641317155 | 3.1655 | | 1.45E-06 | FarmCPU | SUG |
| TGW_17SQ | AX-110835487 | 7D | 426153254 | 3.8960 | | 8.30E-07 | FarmCPU | SUG |
| TGW_17SQ | AX-111065461 | 7D | 590865283 | 4.3969 | | 1.32E-06 | FarmCPU | SUG |
| TGW_17SQ | AX-94965167 | 7D | 616505700 | 1.1532 | | 6.50E-06 | BLUPmrMLM | SUG |
| TGW_17XX | AX-109493578 | 1A | 23000066 | 1.0342 | | 1.45E-07 | BLUPmrMLM | SIG |
| TGW_17XX | AX-109420526 | 1B | 29217823 | 1.5501 | | 2.20E-06 | BLUPmrMLM | SIG |
| TGW_17XX | AX-86183921 | 1B | 673745594 | -3.2014 | | 2.25E-06 | FarmCPU | SUG |
| TGW_17XX | AX-94983738 | 1D | 5357782 | -1.2086 | | 5.76E-06 | BLUPmrMLM | SIG |
| TGW_17XX | AX-108949727 | 2A | 727614699 | -0.4730 | | 8.76E-05 | BLUPmrMLM | SIG |
| TGW_17XX | AX-110595017 | 2A | 744599180 | -0.4781 | | 1.07E-04 | BLUPmrMLM | SIG |
| TGW_17XX | AX-110524795 | 2B | 25077954 | -0.4882 | | 1.80E-04 | BLUPmrMLM | SIG |
| TGW_17XX | AX-111575313 | 2B | 244273019 | 0.4929 | | 1.06E-04 | BLUPmrMLM | SIG |
| TGW_17XX | AX-110958786 | 2B | 762448602 | 0.4690 | | 1.82E-04 | BLUPmrMLM | SIG |
| TGW_17XX | AX-94442651 | 2D | 507050616 | 3.7430 | | 1.44E-06 | FarmCPU | SUG |
| TGW_17XX | AX-110448131 | 3B | 225261015 | -3.0497 | | 4.15E-07 | FarmCPU | SUG |
| TGW_17XX | AX-109479590 | 3B | 237320573 | -3.0690 | | 1.94E-06 | FarmCPU | SUG |
| TGW_17XX | AX-109274638 | 3B | 237325305 | -3.2071 | | 5.36E-07 | FarmCPU | SUG |
| TGW_17XX | AX-110604883 | 4D | 177489950 | 3.8077 | | 6.00E-07 | FarmCPU | SUG |
| TGW_17XX | AX-109318201 | 4D | 197019433 | 3.9225 | | 1.88E-07 | FarmCPU | SUG |
| TGW_17XX | AX-110502948 | 5A | 28495425 | 1.4787 | | 8.87E-08 | BLUPmrMLM | SIG |
| TGW_17XX | AX-110530649 | 5B | 69640482 | -1.0293 | | 3.41E-06 | BLUPmrMLM | SIG |
| TGW_17XX | AX-110012552 | 5B | 680681780 | -0.4186 | | 8.61E-05 | BLUPmrMLM | SIG |
| TGW_17XX | AX-94511633 | 5B | 681168512 | 3.5862 | | 6.75E-07 | FarmCPU | SUG |
| TGW_17XX | AX-109391962 | 6B | 339268873 | 3.9079 | | 1.16E-06 | FarmCPU | SUG |
| TGW_17XX | AX-110509650 | 6B | 642922912 | 0.0425 | 15.4894 | 5.19E-04 | IIIVmrMLM | SUG |
| TGW_17XX | AX-111123882 | 6D | 272759669 | 3.7718 | | 8.74E-07 | FarmCPU | SUG |
| TGW_17XX | AX-110833215 | 6D | 457281781 | 0.7185 | | 1.35E-05 | BLUPmrMLM | SIG |
| TGW_17XX | AX-110474471 | 6D | 470685649 | -0.6619 | | 7.73E-06 | BLUPmrMLM | SIG |
| TGW_17XX | AX-110454170 | 7A | 8375277 | -0.1564 | 15.4894 | 3.40E-04 | IIIVmrMLM | SUG |
| TGW_17XX | AX-110387533 | 7D | 131093753 | -0.9368 | | 3.39E-06 | BLUPmrMLM | SIG |
| TGW_17XX | AX-109459815 | 7D | 392632007 | 2.0165 | | 1.46E-06 | EMMAX | SUG |
| TGW_17XX | AX-110835487 | 7D | 426153254 | 4.4302 | | 1.17E-07 | FarmCPU | SUG |
| TGW_17XX | AX-110960135 | 7D | 486590501 | 3.0819 | | 2.65E-11 | EMMAX | SIG |
| TGW_17ZMD | AX-111553431 | 1A | 14517999 | 0.9638 | | 8.07E-05 | BLUPmrMLM | SUG |
| TGW_17ZMD | AX-109927876 | 1A | 529259749 | 0.5540 | 3.8219 | 3.48E-09 | IIIVmrMLM | SIG |
| TGW_17ZMD | AX-110685978 | 1B | 330793428 | 0.9110 | | 3.60E-14 | IIIVmrMLM | SIG |

| | | | | | | | | |
|---|---|---|---|---|---|---|---|---|
| TGW_17ZMD | AX-108752498 | 1B | 644404296 | 0.3277 | 2.2324 | 4.90E-06 | IIIVmrMLM | SUG |
| TGW_17ZMD | AX-109421874 | 1B | 653506543 | -0.5652 | 2.1382 | 6.44E-11 | IIIVmrMLM | SIG |
| TGW_17ZMD | AX-109260317 | 1D | 38861232 | -0.3799 | -5.8593 | 5.62E-11 | IIIVmrMLM | SIG |
| TGW_17ZMD | AX-94678100 | 2A | 65183989 | -1.2770 | | 1.36E-05 | BLUPmrMLM | SUG |
| TGW_17ZMD | AX-110377545 | 2A | 464955280 | 0.5783 | 3.2096 | 1.20E-23 | IIIVmrMLM | SIG |
| TGW_17ZMD | AX-109491430 | 2A | 472802867 | -0.2211 | -5.4827 | 2.73E-06 | IIIVmrMLM | SUG |
| TGW_17ZMD | AX-94641362 | 2A | 723824451 | 0.7331 | | 1.25E-05 | BLUPmrMLM | SUG |
| TGW_17ZMD | AX-111122462 | 2B | 4610921 | 0.2710 | -2.26 | 4.52E-11 | IIIVmrMLM | SIG |
| TGW_17ZMD | AX-110477125 | 2B | 90188763 | -0.9149 | | 2.89E-06 | BLUPmrMLM | SUG |
| TGW_17ZMD | AX-111695208 | 2B | 103922204 | 0.7861 | -3.4976 | 1.97E-16 | IIIVmrMLM | SIG |
| TGW_17ZMD | AX-110381269 | 2B | 713549487 | -1.6438 | | 4.05E-07 | EMMAX | SUG |
| TGW_17ZMD | AX-110381269 | 2B | 713549487 | 1.6341 | | 3.78E-07 | GEMMA | SUG |
| TGW_17ZMD | AX-110381269 | 2B | 713549487 | -1.1831 | -0.2045 | 1.09E-17 | IIIVmrMLM | SIG |
| TGW_17ZMD | AX-94819224 | 2B | 754546200 | 1.0090 | | 7.14E-17 | IIIVmrMLM | SIG |
| TGW_17ZMD | AX-109320687 | 3A | 142902377 | -0.6803 | 2.051 | 1.08E-10 | IIIVmrMLM | SIG |
| TGW_17ZMD | AX-109063550 | 3A | 244300002 | -0.4621 | | 9.10E-05 | IIIVmrMLM | SUG |
| TGW_17ZMD | AX-109287306 | 3A | 403110157 | -0.3287 | 3.2732 | 4.42E-06 | IIIVmrMLM | SUG |
| TGW_17ZMD | AX-94556128 | 3A | 523550891 | 1.0616 | | 6.51E-05 | BLUPmrMLM | SUG |
| TGW_17ZMD | AX-109607606 | 3A | 551506355 | 0.5207 | -5.0637 | 4.27E-12 | IIIVmrMLM | SIG |
| TGW_17ZMD | AX-109330600 | 3A | 700247598 | 1.0428 | 0.7443 | 1.50E-14 | IIIVmrMLM | SIG |
| TGW_17ZMD | AX-111597853 | 3B | 145905834 | 0.2515 | 2.4207 | 1.20E-06 | IIIVmrMLM | SUG |
| TGW_17ZMD | AX-111584538 | 3B | 315182563 | 0.6106 | 2.5244 | 1.24E-10 | IIIVmrMLM | SIG |
| TGW_17ZMD | AX-111091215 | 3B | 687860739 | 0.1358 | -3.1593 | 6.60E-09 | IIIVmrMLM | SIG |
| TGW_17ZMD | AX-94692827 | 3B | 751058717 | 0.6584 | | 3.03E-08 | IIIVmrMLM | SIG |
| TGW_17ZMD | AX-109388600 | 3B | 763693546 | 0.6829 | 0.8775 | 2.43E-08 | IIIVmrMLM | SIG |
| TGW_17ZMD | AX-109842520 | 4A | 531309587 | 0.6627 | 2.2856 | 5.48E-14 | IIIVmrMLM | SIG |
| TGW_17ZMD | AX-109960485 | 4A | 639621154 | -0.1781 | 1.2292 | 2.21E-05 | IIIVmrMLM | SUG |
| TGW_17ZMD | AX-110174336 | 5A | 18922911 | 1.5063 | | 1.89E-06 | GEMMA | SUG |
| TGW_17ZMD | AX-94816504 | 5A | 18922927 | 1.4700 | | 1.67E-08 | BLUPmrMLM | SIG |
| TGW_17ZMD | AX-94816504 | 5A | 18922927 | -1.7054 | | 1.63E-06 | EMMAX | SUG |
| TGW_17ZMD | AX-94816504 | 5A | 18922927 | 1.7041 | | 1.14E-07 | GEMMA | SIG |
| TGW_17ZMD | AX-109389585 | 5A | 18948563 | 3.1404 | | 4.09E-07 | GEMMA | SUG |
| TGW_17ZMD | AX-111012018 | 5A | 19066884 | 1.4861 | | 2.23E-06 | GEMMA | SUG |
| TGW_17ZMD | AX-108739759 | 5A | 19069586 | 3.0927 | | 7.22E-07 | GEMMA | SUG |
| TGW_17ZMD | AX-109365612 | 5A | 19107969 | 1.6086 | | 4.36E-07 | GEMMA | SUG |
| TGW_17ZMD | AX-109454736 | 5A | 19108636 | 1.5896 | | 3.38E-07 | GEMMA | SUG |
| TGW_17ZMD | AX-111575289 | 5A | 19222566 | 3.0776 | | 1.05E-06 | GEMMA | SUG |
| TGW_17ZMD | AX-111171059 | 5A | 19223635 | 3.2805 | | 2.30E-07 | GEMMA | SUG |
| TGW_17ZMD | AX-111054067 | 5A | 19230597 | 1.6347 | | 2.48E-07 | GEMMA | SUG |
| TGW_17ZMD | AX-109419088 | 5A | 19231120 | 2.5162 | | 2.56E-07 | GEMMA | SUG |
| TGW_17ZMD | AX-110458949 | 5A | 19233051 | 3.1463 | | 2.18E-07 | GEMMA | SUG |
| TGW_17ZMD | AX-94915147 | 5A | 450463750 | -1.6969 | | 2.68E-06 | BLUPmrMLM | SUG |
| TGW_17ZMD | AX-94913721 | 5A | 490318770 | 0.3693 | 2.6512 | 6.14E-08 | IIIVmrMLM | SIG |
| TGW_17ZMD | AX-110029006 | 5A | 511915557 | 0.0510 | -5.961 | 2.22E-07 | IIIVmrMLM | SUG |

| | | | | | | | | |
|---|---|---|---|---|---|---|---|---|
| TGW_17ZMD | AX-109512815 | 5A | 537373893 | -0.3661 | 3.4066 | 2.68E-10 | IIIVmrMLM | SIG |
| TGW_17ZMD | AX-111801605 | 5A | 552900757 | -1.2635 | | 1.88E-05 | BLUPmrMLM | SUG |
| TGW_17ZMD | AX-108805993 | 5B | 55370824 | 1.2640 | -0.3764 | 7.09E-23 | IIIVmrMLM | SIG |
| TGW_17ZMD | AX-111663927 | 5B | 299242437 | -0.3912 | -2.8885 | 1.26E-12 | IIIVmrMLM | SIG |
| TGW_17ZMD | AX-94853278 | 5B | 536516419 | -0.7317 | | 1.47E-04 | BLUPmrMLM | SUG |
| TGW_17ZMD | AX-94663230 | 5D | 28447906 | -3.3325 | | 1.41E-06 | EMMAX | SUG |
| TGW_17ZMD | AX-94663230 | 5D | 28447906 | 3.3402 | | 6.97E-08 | GEMMA | SIG |
| TGW_17ZMD | AX-111122961 | 6A | 145150018 | 0.7874 | | 1.17E-04 | BLUPmrMLM | SUG |
| TGW_17ZMD | AX-110032892 | 6B | 652320197 | 2.0033 | | 3.01E-07 | GEMMA | SUG |
| TGW_17ZMD | AX-110585464 | 7A | 701050963 | -0.4698 | 4.8337 | 3.36E-07 | IIIVmrMLM | SUG |
| TGW_18JY | AX-110958079 | 1A | 45542200 | 0.3399 | 1.3931 | 9.98E-13 | IIIVmrMLM | SIG |
| TGW_18JY | AX-109969772 | 1A | 442817137 | -0.8307 | | 2.36E-23 | IIIVmrMLM | SIG |
| TGW_18JY | AX-110108217 | 1A | 463154024 | 1.0189 | | 4.46E-33 | IIIVmrMLM | SIG |
| TGW_18JY | AX-111186786 | 1A | 518515638 | 2.8875 | | 4.68E-07 | FarmCPU | SUG |
| TGW_18JY | AX-94636510 | 1A | 522190455 | -0.6742 | | 5.63E-07 | BLUPmrMLM | SUG |
| TGW_18JY | AX-109414513 | 1A | 540629580 | -0.7876 | 0.2874 | 2.60E-19 | IIIVmrMLM | SIG |
| TGW_18JY | AX-94525724 | 1A | 564749741 | 6.3472 | | 2.06E-06 | FarmCPU | SUG |
| TGW_18JY | AX-111541662 | 1B | 1429808 | 0.1930 | 7.1935 | 1.22E-07 | IIIVmrMLM | SUG |
| TGW_18JY | AX-109882166 | 1B | 22123302 | 0.5637 | | 3.20E-05 | BLUPmrMLM | SUG |
| TGW_18JY | AX-111597039 | 1B | 332383749 | -0.1277 | -1.8777 | 1.51E-05 | IIIVmrMLM | SUG |
| TGW_18JY | AX-95093502 | 1B | 554586510 | -0.6634 | 0.6852 | 4.82E-15 | IIIVmrMLM | SIG |
| TGW_18JY | AX-110427177 | 1B | 582927346 | -1.1996 | | 2.63E-07 | BLUPmrMLM | SUG |
| TGW_18JY | AX-108884734 | 1D | 46457630 | 0.3101 | -1.8691 | 8.12E-08 | IIIVmrMLM | SIG |
| TGW_18JY | AX-111074100 | 1D | 68446394 | -0.8634 | | 4.98E-07 | BLUPmrMLM | SUG |
| TGW_18JY | AX-108798166 | 1D | 468399780 | 6.0856 | | 3.81E-08 | EMMAX | SIG |
| TGW_18JY | AX-108798166 | 1D | 468399780 | -6.0304 | | 3.68E-08 | GEMMA | SIG |
| TGW_18JY | AX-110533557 | 2A | 663947 | -0.5489 | | 1.04E-04 | BLUPmrMLM | SUG |
| TGW_18JY | AX-109980275 | 2A | 8160814 | 0.5233 | | 5.22E-05 | BLUPmrMLM | SUG |
| TGW_18JY | AX-111786868 | 2A | 64358348 | 0.8537 | | 1.80E-24 | IIIVmrMLM | SIG |
| TGW_18JY | AX-108731586 | 2A | 86676532 | -0.0428 | 2.7449 | 1.46E-12 | IIIVmrMLM | SIG |
| TGW_18JY | AX-111051158 | 2A | 158803109 | 0.0847 | -1.4937 | 4.86E-08 | IIIVmrMLM | SIG |
| TGW_18JY | AX-108886473 | 2A | 475771811 | -0.1847 | 1.2924 | 1.13E-12 | IIIVmrMLM | SIG |
| TGW_18JY | AX-108790363 | 2A | 708186516 | -0.4746 | -0.099 | 6.05E-08 | IIIVmrMLM | SIG |
| TGW_18JY | AX-108949727 | 2A | 727614699 | -0.8410 | | 3.76E-07 | BLUPmrMLM | SUG |
| TGW_18JY | AX-109330937 | 2A | 755378747 | -0.1825 | -1.3875 | 5.98E-08 | IIIVmrMLM | SIG |
| TGW_18JY | AX-111489027 | 2A | 758274417 | -0.6165 | | 3.33E-06 | BLUPmrMLM | SUG |
| TGW_18JY | AX-108922838 | 2B | 3329464 | 0.1219 | 2.8862 | 3.61E-07 | IIIVmrMLM | SUG |
| TGW_18JY | AX-110531005 | 2B | 146640562 | -0.6350 | | 1.21E-05 | BLUPmrMLM | SUG |
| TGW_18JY | AX-108825946 | 2B | 716995051 | -0.4389 | | 4.91E-05 | BLUPmrMLM | SUG |
| TGW_18JY | AX-95235288 | 2B | 722346909 | -0.4229 | | 1.23E-04 | BLUPmrMLM | SUG |
| TGW_18JY | AX-111078139 | 2B | 725985214 | -0.6119 | -2.6224 | 2.28E-19 | IIIVmrMLM | SIG |
| TGW_18JY | AX-109427839 | 2B | 728763995 | 0.5343 | 1.8962 | 1.12E-14 | IIIVmrMLM | SIG |
| TGW_18JY | AX-111767820 | 2D | 17303983 | 0.0754 | -1.4351 | 6.81E-04 | IIIVmrMLM | SUG |
| TGW_18JY | AX-110531728 | 2D | 618249171 | -1.0458 | | 3.40E-06 | BLUPmrMLM | SUG |

| | | | | | | | | |
|---|---|---|---|---|---|---|---|---|
| TGW_18JY | AX-111640432 | 3A | 728419576 | 0.6661 | | 4.49E-07 | BLUPmrMLM | SUG |
| TGW_18JY | AX-109019801 | 3A | 747613640 | 0.0186 | 1.7461 | 1.60E-21 | IIIVmrMLM | SIG |
| TGW_18JY | AX-111212607 | 3B | 13651258 | -0.3383 | -2.4885 | 8.50E-15 | IIIVmrMLM | SIG |
| TGW_18JY | AX-109435664 | 3B | 40536093 | 0.3327 | -1.14 | 6.68E-08 | IIIVmrMLM | SIG |
| TGW_18JY | AX-94462585 | 3B | 141816928 | 0.5411 | -0.2955 | 8.44E-09 | IIIVmrMLM | SIG |
| TGW_18JY | AX-109274638 | 3B | 237325305 | -4.5824 | | 5.41E-15 | FarmCPU | SIG |
| TGW_18JY | AX-110999727 | 3B | 719933688 | 0.1287 | 1.5624 | 4.23E-06 | IIIVmrMLM | SUG |
| TGW_18JY | AX-94840959 | 3D | 611918568 | -0.1987 | 2.9179 | 1.60E-08 | IIIVmrMLM | SIG |
| TGW_18JY | AX-89323121 | 4A | 734256538 | 0.8519 | | 5.20E-05 | BLUPmrMLM | SUG |
| TGW_18JY | AX-110909865 | 4B | 16389880 | -0.5041 | | 1.08E-04 | BLUPmrMLM | SUG |
| TGW_18JY | AX-109090170 | 4B | 87023106 | 0.1854 | 5.4789 | 8.01E-07 | IIIVmrMLM | SUG |
| TGW_18JY | AX-111554742 | 4B | 393344292 | 1.7611 | | 1.40E-07 | BLUPmrMLM | SUG |
| TGW_18JY | AX-111554742 | 4B | 393344292 | -0.5613 | | 6.13E-12 | IIIVmrMLM | SIG |
| TGW_18JY | AX-111611789 | 5A | 29963648 | -3.0140 | | 2.95E-07 | EMMAX | SUG |
| TGW_18JY | AX-111611789 | 5A | 29963648 | 3.0174 | | 2.41E-07 | GEMMA | SUG |
| TGW_18JY | AX-109903760 | 5A | 30056842 | 2.8903 | | 2.55E-12 | BLUPmrMLM | SIG |
| TGW_18JY | AX-109903760 | 5A | 30056842 | -3.8767 | | 3.30E-10 | EMMAX | SIG |
| TGW_18JY | AX-109903760 | 5A | 30056842 | 3.8963 | | 2.11E-10 | GEMMA | SIG |
| TGW_18JY | AX-94543078 | 5A | 99595622 | -1.8612 | | 1.01E-07 | FarmCPU | SIG |
| TGW_18JY | AX-111231015 | 5A | 448044954 | -0.8326 | | 1.05E-06 | BLUPmrMLM | SUG |
| TGW_18JY | AX-110675322 | 5A | 480425843 | 1.7894 | 0.6181 | 1.24E-38 | IIIVmrMLM | SIG |
| TGW_18JY | AX-111239641 | 5A | 531656356 | 0.0340 | 2.1503 | 1.31E-08 | IIIVmrMLM | SIG |
| TGW_18JY | AX-110530894 | 5A | 584590793 | 3.8739 | | 1.00E-06 | EMMAX | SUG |
| TGW_18JY | AX-110530894 | 5A | 584590793 | -3.8286 | | 5.86E-07 | GEMMA | SUG |
| TGW_18JY | AX-112288451 | 5A | 584614282 | 7.5079 | | 1.94E-06 | EMMAX | SUG |
| TGW_18JY | AX-112288451 | 5A | 584614282 | -7.3826 | | 1.32E-06 | GEMMA | SUG |
| TGW_18JY | AX-109528616 | 5A | 585095529 | 7.5079 | | 1.94E-06 | EMMAX | SUG |
| TGW_18JY | AX-109528616 | 5A | 585095529 | -7.3826 | | 1.32E-06 | GEMMA | SUG |
| TGW_18JY | AX-110404585 | 5A | 585212941 | 7.5650 | | 1.65E-06 | EMMAX | SUG |
| TGW_18JY | AX-110404585 | 5A | 585212941 | -7.4572 | | 1.08E-06 | GEMMA | SUG |
| TGW_18JY | AX-108779357 | 5B | 30653646 | -0.4149 | -5.2983 | 3.52E-15 | IIIVmrMLM | SIG |
| TGW_18JY | AX-110368784 | 5B | 405768546 | 1.2509 | | 2.73E-08 | BLUPmrMLM | SIG |
| TGW_18JY | AX-110946166 | 5B | 455354873 | 2.2609 | | 6.02E-07 | FarmCPU | SUG |
| TGW_18JY | AX-109838322 | 5B | 556973087 | 3.2785 | | 2.07E-06 | GEMMA | SUG |
| TGW_18JY | AX-109373858 | 5B | 665939342 | 0.1133 | 3.0659 | 9.24E-25 | IIIVmrMLM | SIG |
| TGW_18JY | AX-109816855 | 5B | 684384173 | -1.2710 | -0.0763 | 4.59E-34 | IIIVmrMLM | SIG |
| TGW_18JY | AX-110534182 | 5B | 699694182 | -0.5079 | 0.1312 | 8.50E-09 | IIIVmrMLM | SIG |
| TGW_18JY | AX-108759986 | 5B | 700008579 | 0.4403 | | 1.55E-04 | BLUPmrMLM | SUG |
| TGW_18JY | AX-110616258 | 5D | 541612891 | -1.8261 | | 1.18E-06 | EMMAX | SUG |
| TGW_18JY | AX-110616258 | 5D | 541612891 | 1.8194 | | 4.33E-07 | GEMMA | SUG |
| TGW_18JY | AX-110616258 | 5D | 541612891 | -0.9535 | -0.4527 | 9.22E-28 | IIIVmrMLM | SIG |
| TGW_18JY | AX-109864870 | 6A | 32792795 | -1.2581 | | 2.31E-06 | BLUPmrMLM | SUG |
| TGW_18JY | AX-111533185 | 6A | 104527839 | 0.6788 | | 3.03E-05 | BLUPmrMLM | SUG |
| TGW_18JY | AX-108817049 | 6A | 416728745 | -1.0902 | 0.1151 | 1.25E-31 | IIIVmrMLM | SIG |

| | | | | | | | | |
|---|---|---|---|---|---|---|---|---|
| TGW_18JY | AX-109373499 | 6A | 574243605 | 0.1051 | 1.4254 | 1.09E-11 | IIIVmrMLM | SIG |
| TGW_18JY | AX-111473756 | 6B | 7304167 | 0.1835 | -4.4872 | 2.29E-07 | IIIVmrMLM | SUG |
| TGW_18JY | AX-109375407 | 6B | 18281605 | -0.6602 | | 9.28E-06 | BLUPmrMLM | SUG |
| TGW_18JY | AX-108841575 | 6B | 86645945 | -0.5077 | -1.06 | 3.42E-10 | IIIVmrMLM | SIG |
| TGW_18JY | AX-109370945 | 6B | 456774277 | 1.0622 | | 1.64E-07 | BLUPmrMLM | SUG |
| TGW_18JY | AX-108733460 | 6B | 488557165 | 1.6359 | | 1.06E-06 | FarmCPU | SUG |
| TGW_18JY | AX-108835866 | 6B | 573596942 | 0.2442 | 1.5778 | 2.32E-13 | IIIVmrMLM | SIG |
| TGW_18JY | AX-110489846 | 6B | 678162403 | -0.0194 | -1.918 | 2.00E-06 | IIIVmrMLM | SUG |
| TGW_18JY | AX-110992019 | 6B | 706700338 | -0.8887 | | 2.87E-08 | BLUPmrMLM | SIG |
| TGW_18JY | AX-108932028 | 6D | 3317339 | 0.2677 | -1.0407 | 6.07E-07 | IIIVmrMLM | SUG |
| TGW_18JY | AX-109967269 | 6D | 22183383 | 0.6695 | | 2.38E-06 | BLUPmrMLM | SUG |
| TGW_18JY | AX-110425881 | 6D | 26850629 | 0.0320 | -2.6957 | 3.41E-05 | IIIVmrMLM | SUG |
| TGW_18JY | AX-109724395 | 6D | 470727348 | 0.7855 | 0.5315 | 8.02E-20 | IIIVmrMLM | SIG |
| TGW_18JY | AX-111038802 | 6D | 470752471 | -1.6977 | | 2.90E-05 | BLUPmrMLM | SUG |
| TGW_18JY | AX-110454170 | 7A | 8375277 | -0.1334 | 15.6934 | 1.78E-14 | IIIVmrMLM | SIG |
| TGW_18JY | AX-95075921 | 7A | 19007631 | -0.8837 | | 4.10E-06 | BLUPmrMLM | SUG |
| TGW_18JY | AX-109356150 | 7A | 89971814 | -1.2365 | | 3.52E-09 | BLUPmrMLM | SIG |
| TGW_18JY | AX-110990714 | 7A | 113244887 | -1.4217 | 0.2807 | 4.49E-36 | IIIVmrMLM | SIG |
| TGW_18JY | AX-108965103 | 7A | 529983946 | -0.2479 | 2.5897 | 1.26E-09 | IIIVmrMLM | SIG |
| TGW_18JY | AX-109990643 | 7A | 534004060 | -0.0155 | -2.1088 | 1.56E-18 | IIIVmrMLM | SIG |
| TGW_18JY | AX-108839894 | 7A | 678153714 | 1.3089 | | 2.11E-07 | BLUPmrMLM | SUG |
| TGW_18JY | AX-110906405 | 7B | 709469414 | -0.7333 | -0.8974 | 6.27E-18 | IIIVmrMLM | SIG |
| TGW_18JY | AX-108917551 | 7B | 717344698 | -0.5852 | | 1.62E-07 | BLUPmrMLM | SUG |
| TGW_18JY | AX-110960135 | 7D | 486590501 | -16.9920 | | 3.26E-07 | EMMAX | SUG |
| TGW_18XX | AX-95198554 | 1A | 12304373 | -0.1576 | 2.1404 | 1.01E-10 | IIIVmrMLM | SIG |
| TGW_18XX | AX-110514823 | 1A | 43330926 | 0.0443 | 1.8406 | 1.24E-11 | IIIVmrMLM | SIG |
| TGW_18XX | AX-111511896 | 1A | 488155945 | -0.0226 | 2.9531 | 3.71E-14 | IIIVmrMLM | SIG |
| TGW_18XX | AX-109427893 | 1A | 540502254 | -0.3835 | 0.2161 | 1.45E-08 | IIIVmrMLM | SIG |
| TGW_18XX | AX-111017649 | 1A | 546488635 | 0.2053 | 2.7685 | 1.33E-09 | IIIVmrMLM | SIG |
| TGW_18XX | AX-109427328 | 1A | 564586780 | 3.7972 | | 2.04E-06 | FarmCPU | SUG |
| TGW_18XX | AX-109832076 | 1A | 583460518 | -0.5023 | | 1.52E-06 | BLUPmrMLM | SUG |
| TGW_18XX | AX-109981359 | 1A | 584807135 | 0.6502 | | 4.36E-08 | BLUPmrMLM | SIG |
| TGW_18XX | AX-109944767 | 1B | 332565438 | 0.4287 | | 1.88E-05 | BLUPmrMLM | SUG |
| TGW_18XX | AX-111077704 | 1B | 340424510 | 0.8218 | 0.1766 | 1.53E-32 | IIIVmrMLM | SIG |
| TGW_18XX | AX-109903479 | 1B | 394073619 | 3.7811 | | 1.89E-06 | FarmCPU | SUG |
| TGW_18XX | AX-108980623 | 1B | 409273408 | 0.5965 | | 3.45E-08 | BLUPmrMLM | SIG |
| TGW_18XX | AX-94684170 | 1B | 597537011 | 3.7911 | | 8.46E-07 | FarmCPU | SUG |
| TGW_18XX | AX-111157053 | 1B | 643365932 | -0.4468 | 0.5538 | 4.26E-12 | IIIVmrMLM | SIG |
| TGW_18XX | AX-110043099 | 1D | 258660 | -0.2550 | 4.3892 | 2.04E-12 | IIIVmrMLM | SIG |
| TGW_18XX | AX-110294806 | 1D | 42827184 | -0.5981 | | 1.38E-08 | BLUPmrMLM | SIG |
| TGW_18XX | AX-111194284 | 1D | 67002589 | 0.4247 | | 9.09E-06 | BLUPmrMLM | SUG |
| TGW_18XX | AX-110592232 | 1D | 486365358 | 0.0792 | 1.7759 | 2.09E-07 | IIIVmrMLM | SUG |
| TGW_18XX | AX-108739692 | 2A | 60280251 | -1.7719 | | 2.13E-06 | EMMAX | SUG |
| TGW_18XX | AX-108739692 | 2A | 60280251 | 1.7679 | | 1.16E-06 | GEMMA | SUG |

| | | | | | | | | |
|---|---|---|---|---|---|---|---|---|
| TGW_18XX | AX-108739692 | 2A | 60280251 | -1.0946 | -0.2518 | 2.76E-54 | IIIVmrMLM | SIG |
| TGW_18XX | AX-110926144 | 2A | 614413443 | -0.1090 | 1.5323 | 1.20E-27 | IIIVmrMLM | SIG |
| TGW_18XX | AX-108940264 | 2A | 677175739 | -0.5045 | | 6.07E-06 | BLUPmrMLM | SUG |
| TGW_18XX | AX-110640235 | 2A | 703124456 | -0.0216 | -0.893 | 7.68E-05 | IIIVmrMLM | SUG |
| TGW_18XX | AX-94496472 | 2A | 729199454 | 0.7068 | | 1.78E-06 | BLUPmrMLM | SUG |
| TGW_18XX | AX-109947604 | 2A | 752854585 | -0.4031 | | 7.04E-05 | BLUPmrMLM | SUG |
| TGW_18XX | AX-109344204 | 2A | 759309971 | 0.0573 | 1.1996 | 1.35E-10 | IIIVmrMLM | SIG |
| TGW_18XX | AX-94548253 | 2B | 16272965 | -0.3766 | | 3.19E-09 | IIIVmrMLM | SIG |
| TGW_18XX | AX-108824752 | 2B | 66805981 | -0.6521 | 0.7661 | 1.83E-22 | IIIVmrMLM | SIG |
| TGW_18XX | AX-109286668 | 2B | 181773892 | 0.3641 | -0.5819 | 6.34E-09 | IIIVmrMLM | SIG |
| TGW_18XX | AX-108862346 | 2B | 327656136 | 3.4870 | | 2.06E-06 | FarmCPU | SUG |
| TGW_18XX | AX-111022990 | 2B | 596836513 | 0.1885 | -1.6442 | 1.02E-15 | IIIVmrMLM | SIG |
| TGW_18XX | AX-95167880 | 2B | 673708364 | -0.4865 | 0.0886 | 5.60E-13 | IIIVmrMLM | SIG |
| TGW_18XX | AX-110675881 | 2B | 732203884 | 5.2854 | | 8.19E-07 | GEMMA | SUG |
| TGW_18XX | AX-110555481 | 2B | 786772802 | 0.5447 | | 8.49E-07 | BLUPmrMLM | SUG |
| TGW_18XX | AX-95205943 | 2D | 12552398 | -0.8324 | | 8.78E-06 | BLUPmrMLM | SUG |
| TGW_18XX | AX-94697254 | 2D | 33710135 | 3.3826 | | 4.60E-07 | FarmCPU | SUG |
| TGW_18XX | AX-110412786 | 2D | 38767905 | 0.7123 | | 9.74E-07 | BLUPmrMLM | SUG |
| TGW_18XX | AX-110412786 | 2D | 38767905 | -0.7060 | -0.4129 | 6.96E-25 | IIIVmrMLM | SIG |
| TGW_18XX | AX-94628670 | 2D | 123215140 | 0.9462 | | 1.97E-04 | BLUPmrMLM | SUG |
| TGW_18XX | AX-94442651 | 2D | 507050616 | 4.3766 | | 3.34E-08 | FarmCPU | SIG |
| TGW_18XX | AX-109286245 | 3A | 421119830 | 3.0837 | | 1.33E-06 | FarmCPU | SUG |
| TGW_18XX | AX-110534996 | 3A | 427186899 | 1.1252 | | 1.61E-08 | BLUPmrMLM | SIG |
| TGW_18XX | AX-110378966 | 3B | 10368223 | -0.6903 | | 2.91E-06 | BLUPmrMLM | SUG |
| TGW_18XX | AX-109565183 | 3B | 14368020 | -0.5764 | | 3.61E-06 | BLUPmrMLM | SUG |
| TGW_18XX | AX-109849144 | 3B | 39542024 | -0.6264 | | 4.10E-07 | BLUPmrMLM | SUG |
| TGW_18XX | AX-110448131 | 3B | 225261015 | -2.9651 | | 1.68E-06 | FarmCPU | SUG |
| TGW_18XX | AX-109274638 | 3B | 237325305 | -3.1284 | | 1.98E-06 | FarmCPU | SUG |
| TGW_18XX | AX-110686903 | 3B | 341059440 | -0.1045 | -2.7959 | 3.92E-08 | IIIVmrMLM | SIG |
| TGW_18XX | AX-111282456 | 3B | 367786072 | 0.8018 | | 2.34E-07 | BLUPmrMLM | SUG |
| TGW_18XX | AX-109839667 | 3B | 469646958 | 0.0285 | -1.9679 | 8.14E-11 | IIIVmrMLM | SIG |
| TGW_18XX | AX-110366839 | 3B | 744363110 | 0.6813 | -0.0946 | 2.35E-15 | IIIVmrMLM | SIG |
| TGW_18XX | AX-110603940 | 3B | 749990662 | -0.5648 | | 4.01E-07 | BLUPmrMLM | SUG |
| TGW_18XX | AX-110433065 | 3D | 573431301 | 0.4860 | 0.1265 | 1.03E-07 | IIIVmrMLM | SIG |
| TGW_18XX | AX-109420906 | 4A | 115634351 | 3.7605 | | 1.15E-06 | FarmCPU | SUG |
| TGW_18XX | AX-111024713 | 4A | 594665324 | -0.6178 | | 1.25E-10 | BLUPmrMLM | SIG |
| TGW_18XX | AX-108833696 | 4A | 712602406 | -0.4616 | | 2.55E-05 | BLUPmrMLM | SUG |
| TGW_18XX | AX-109501835 | 4B | 20590398 | 0.8724 | 0.2371 | 1.64E-34 | IIIVmrMLM | SIG |
| TGW_18XX | AX-109920329 | 4B | 139621756 | -1.0115 | 0.7188 | 2.56E-47 | IIIVmrMLM | SIG |
| TGW_18XX | AX-110945021 | 4B | 658945428 | -0.8526 | | 8.57E-07 | BLUPmrMLM | SUG |
| TGW_18XX | AX-111714870 | 4D | 68054117 | 3.9830 | | 1.74E-06 | FarmCPU | SUG |
| TGW_18XX | AX-110604883 | 4D | 177489950 | 4.2522 | | 4.94E-08 | FarmCPU | SIG |
| TGW_18XX | AX-109318201 | 4D | 197019433 | 3.9986 | | 1.94E-07 | FarmCPU | SUG |
| TGW_18XX | AX-108890143 | 5A | 31511922 | -0.5456 | 0.1842 | 2.89E-09 | IIIVmrMLM | SIG |

| | | | | | | | | |
|---|---|---|---|---|---|---|---|---|
| TGW_18XX | AX-109376414 | 5A | 32972598 | -0.3468 | | 1.59E-05 | BLUPmrMLM | SUG |
| TGW_18XX | AX-94762802 | 5A | 37620417 | 0.7500 | | 8.39E-05 | BLUPmrMLM | SUG |
| TGW_18XX | AX-110532263 | 5A | 122023525 | 0.1550 | -2.0658 | 1.44E-12 | IIIVmrMLM | SIG |
| TGW_18XX | AX-109512930 | 5A | 447158258 | 0.2293 | -10.7178 | 2.47E-13 | IIIVmrMLM | SIG |
| TGW_18XX | AX-111578813 | 5A | 705565384 | -0.1974 | 2.5121 | 5.70E-12 | IIIVmrMLM | SIG |
| TGW_18XX | AX-110933310 | 5B | 31438637 | -0.3245 | -0.7825 | 3.46E-12 | IIIVmrMLM | SIG |
| TGW_18XX | AX-109373007 | 5B | 397780675 | -2.5850 | | 1.46E-06 | FarmCPU | SUG |
| TGW_18XX | AX-111684719 | 5B | 477953826 | -0.0325 | -1.2522 | 1.44E-06 | IIIVmrMLM | SUG |
| TGW_18XX | AX-111684648 | 5B | 520178595 | 3.8718 | | 3.54E-07 | FarmCPU | SUG |
| TGW_18XX | AX-110471122 | 5B | 532602914 | 0.2980 | 4.4138 | 2.35E-18 | IIIVmrMLM | SIG |
| TGW_18XX | AX-94511633 | 5B | 681168512 | 3.9000 | | 1.27E-07 | FarmCPU | SUG |
| TGW_18XX | AX-111458073 | 5B | 694033860 | 0.7863 | | 2.79E-12 | BLUPmrMLM | SIG |
| TGW_18XX | AX-111139651 | 5B | 699616555 | -0.0985 | -0.8252 | 6.10E-21 | IIIVmrMLM | SIG |
| TGW_18XX | AX-109465280 | 5D | 207252679 | -1.0299 | | 2.39E-08 | BLUPmrMLM | SIG |
| TGW_18XX | AX-95082229 | 5D | 416207438 | -1.0510 | | 2.85E-52 | IIIVmrMLM | SIG |
| TGW_18XX | AX-111464653 | 5D | 486138536 | 3.5865 | | 1.52E-06 | FarmCPU | SUG |
| TGW_18XX | AX-111133055 | 5D | 490598269 | -0.6505 | | 2.00E-05 | BLUPmrMLM | SUG |
| TGW_18XX | AX-94754448 | 6A | 31355561 | -0.7269 | | 2.85E-28 | IIIVmrMLM | SIG |
| TGW_18XX | AX-110088164 | 6A | 119789966 | -0.5944 | | 2.44E-05 | BLUPmrMLM | SUG |
| TGW_18XX | AX-108739437 | 6A | 127098136 | 0.8833 | 0.238 | 4.72E-35 | IIIVmrMLM | SIG |
| TGW_18XX | AX-109341101 | 6A | 271258432 | -0.7361 | | 7.68E-07 | BLUPmrMLM | SUG |
| TGW_18XX | AX-109299648 | 6A | 595030062 | 0.5622 | | 2.00E-04 | BLUPmrMLM | SUG |
| TGW_18XX | AX-109037034 | 6A | 603809881 | -0.5066 | | 4.60E-05 | BLUPmrMLM | SUG |
| TGW_18XX | AX-109861336 | 6A | 610284657 | -0.3899 | | 1.16E-05 | BLUPmrMLM | SUG |
| TGW_18XX | AX-111000497 | 6A | 613355611 | 0.9581 | | 2.23E-14 | BLUPmrMLM | SIG |
| TGW_18XX | AX-110063373 | 6A | 617402073 | 0.1456 | 1.3183 | 3.87E-07 | IIIVmrMLM | SUG |
| TGW_18XX | AX-94931823 | 6B | 201205658 | 0.1051 | 0.803 | 4.77E-07 | IIIVmrMLM | SUG |
| TGW_18XX | AX-109391962 | 6B | 339268873 | 4.5750 | | 2.42E-08 | FarmCPU | SIG |
| TGW_18XX | AX-110479567 | 6B | 574416964 | -0.1600 | 3.8765 | 2.30E-07 | IIIVmrMLM | SUG |
| TGW_18XX | AX-111123882 | 6D | 272759669 | 4.2621 | | 5.51E-08 | FarmCPU | SIG |
| TGW_18XX | AX-110046771 | 6D | 465447468 | -0.2807 | | 1.54E-04 | BLUPmrMLM | SUG |
| TGW_18XX | AX-86176335 | 6D | 473418687 | 0.9695 | | 4.84E-10 | BLUPmrMLM | SIG |
| TGW_18XX | AX-110454170 | 7A | 8375277 | 0.2523 | 9.5068 | 7.04E-12 | IIIVmrMLM | SIG |
| TGW_18XX | AX-111153914 | 7A | 507877239 | 4.1038 | | 6.81E-07 | FarmCPU | SUG |
| TGW_18XX | AX-109081729 | 7A | 648700205 | -0.3032 | 2.1221 | 2.55E-11 | IIIVmrMLM | SIG |
| TGW_18XX | AX-111770216 | 7A | 717443302 | -0.2423 | -9.5088 | 1.25E-11 | IIIVmrMLM | SIG |
| TGW_18XX | AX-109290188 | 7A | 736522870 | -0.5086 | | 1.83E-05 | BLUPmrMLM | SUG |
| TGW_18XX | AX-111112635 | 7B | 19891825 | 0.0022 | 0.8772 | 7.93E-27 | IIIVmrMLM | SIG |
| TGW_18XX | AX-94821564 | 7B | 41524564 | -0.1942 | 1.0106 | 2.58E-06 | IIIVmrMLM | SUG |
| TGW_18XX | AX-109885210 | 7B | 632324556 | 0.4223 | -0.2055 | 8.41E-09 | IIIVmrMLM | SIG |
| TGW_18XX | AX-111235052 | 7B | 742261618 | -0.0803 | -3.3055 | 4.02E-14 | IIIVmrMLM | SIG |
| TGW_18XX | AX-111918328 | 7D | 93673586 | 1.2682 | | 2.79E-08 | BLUPmrMLM | SIG |
| TGW_18XX | AX-110835487 | 7D | 426153254 | 4.5303 | | 1.33E-07 | FarmCPU | SUG |
| TGW_18XX | AX-111065461 | 7D | 590865283 | 4.6825 | | 2.19E-06 | FarmCPU | SUG |

| | | | | | | | | |
|---|---|---|---|---|---|---|---|---|
| TGW_18XX | AX-110614870 | 7D | 608124620 | 0.4073 | | 3.00E-05 | BLUPmrMLM | SUG |
| TGW_18XX | AX-109938735 | 7D | 614187479 | -0.2216 | 0.9089 | 1.40E-08 | IIIVmrMLM | SIG |
| TGW_18ZMD | AX-108780402 | 1A | 564570992 | 2.5622 | | 5.49E-07 | FarmCPU | SUG |
| TGW_18ZMD | AX-109005639 | 2B | 788519649 | 1.6381 | | 8.00E-08 | BLUPmrMLM | SIG |
| TGW_18ZMD | AX-94579852 | 2D | 189601211 | -29.1930 | | 6.78E-09 | EMMAX | SIG |
| TGW_18ZMD | AX-94442651 | 2D | 507050616 | 3.6252 | | 2.18E-06 | FarmCPU | SUG |
| TGW_18ZMD | AX-110942302 | 3B | 380728885 | 3.8861 | | 1.78E-06 | FarmCPU | SUG |
| TGW_18ZMD | AX-111216811 | 3B | 792106339 | -0.6573 | | 5.31E-05 | BLUPmrMLM | SUG |
| TGW_18ZMD | AX-94968108 | 3D | 596890133 | -13.8910 | | 2.25E-07 | EMMAX | SUG |
| TGW_18ZMD | AX-110021031 | 4D | 155135769 | -29.1930 | | 6.78E-09 | EMMAX | SIG |
| TGW_18ZMD | AX-109329176 | 6B | 4885850 | -0.8543 | | 6.92E-05 | BLUPmrMLM | SUG |
| TGW_18ZMD | AX-109391962 | 6B | 339268873 | 3.9365 | | 6.94E-07 | FarmCPU | SUG |
| TGW_18ZMD | AX-109319706 | 6B | 704705182 | -0.8149 | | 2.96E-05 | BLUPmrMLM | SUG |
| TGW_18ZMD | AX-109372256 | 7A | 437403649 | 2.9219 | | 7.46E-07 | FarmCPU | SUG |
| TGW_18ZMD | AX-108773677 | 7A | 444503341 | 2.8756 | | 1.20E-06 | FarmCPU | SUG |
| TGW_18ZMD | AX-110185538 | 7A | 444964265 | 2.7441 | | 1.67E-06 | FarmCPU | SUG |
| TGW_18ZMD | AX-109531112 | 7A | 446361440 | 2.5667 | | 8.69E-07 | FarmCPU | SUG |
| TGW_18ZMD | AX-95100648 | 7A | 572323579 | -19.4880 | | 9.56E-07 | EMMAX | SUG |
| TGW_18ZMD | AX-110386995 | 7A | 641398757 | -1.3360 | | 3.77E-08 | BLUPmrMLM | SIG |
| TGW_18ZMD | AX-108873400 | 7A | 644877210 | 0.9091 | | 6.91E-05 | BLUPmrMLM | SUG |
| TGW_18ZMD | AX-111543985 | 7A | 709627123 | -0.9624 | | 1.13E-04 | BLUPmrMLM | SUG |
| TGW_18ZMD | AX-109314090 | 7B | 653889269 | 2.1295 | | 1.91E-06 | FarmCPU | SUG |
| TGW_18ZMD | AX-95632322 | 7B | 653889814 | 3.8524 | | 1.85E-06 | FarmCPU | SUG |
| TGW_18ZMD | AX-108895074 | 7B | 653891315 | 2.0654 | | 1.51E-06 | FarmCPU | SUG |
| TGW_18ZMD | AX-109433389 | 7D | 504750431 | -2.6083 | 25.8491 | 7.81E-50 | IIIVmrMLM | SIG |
| TGW_19XX | AX-109427328 | 1A | 564586780 | 4.4114 | | 9.60E-07 | FarmCPU | SUG |
| TGW_19XX | AX-94525724 | 1A | 564749741 | 8.8080 | | 1.93E-06 | FarmCPU | SUG |
| TGW_19XX | AX-110953597 | 1B | 303664637 | 1.9361 | | 1.85E-06 | EMMAX | SUG |
| TGW_19XX | AX-110953597 | 1B | 303664637 | -1.9202 | | 2.14E-06 | GEMMA | SUG |
| TGW_19XX | AX-94684170 | 1B | 597537011 | 4.1841 | | 1.44E-06 | FarmCPU | SUG |
| TGW_19XX | AX-108934860 | 2A | 70146016 | -1.4680 | | 1.34E-06 | GEMMA | SUG |
| TGW_19XX | AX-111053248 | 2A | 71359445 | -1.1029 | | 2.45E-08 | BLUPmrMLM | SIG |
| TGW_19XX | AX-111053248 | 2A | 71359445 | -1.4883 | | 2.08E-06 | GEMMA | SUG |
| TGW_19XX | AX-109580490 | 2A | 477281002 | -2.3124 | | 1.71E-06 | GEMMA | SUG |
| TGW_19XX | AX-110386092 | 2A | 733923313 | -1.1957 | | 1.85E-05 | BLUPmrMLM | SUG |
| TGW_19XX | AX-94488783 | 2A | 755788522 | 0.8834 | | 6.64E-05 | BLUPmrMLM | SUG |
| TGW_19XX | AX-111021677 | 2B | 5767670 | 0.7824 | | 1.40E-06 | BLUPmrMLM | SUG |
| TGW_19XX | AX-110524795 | 2B | 25077954 | -0.4496 | | 2.65E-05 | BLUPmrMLM | SUG |
| TGW_19XX | AX-94442651 | 2D | 507050616 | 4.6699 | | 1.79E-07 | FarmCPU | SUG |
| TGW_19XX | AX-109947185 | 3A | 210249582 | 0.7408 | | 1.45E-04 | BLUPmrMLM | SUG |
| TGW_19XX | AX-109286245 | 3A | 421119830 | 3.4524 | | 1.58E-06 | FarmCPU | SUG |
| TGW_19XX | AX-109992721 | 3A | 685701474 | -0.8790 | | 4.90E-06 | BLUPmrMLM | SUG |
| TGW_19XX | AX-110641031 | 3B | 7601649 | 0.4127 | | 1.03E-04 | BLUPmrMLM | SUG |
| TGW_19XX | AX-109821909 | 4A | 686376485 | -0.7722 | | 2.38E-06 | BLUPmrMLM | SUG |

| | | | | | | | |
|---|---|---|---|---|---|---|---|
| TGW_19XX | AX-108913575 | 4A | 695293726 | -1.1635 | 4.32E-10 | BLUPmrMLM | SIG |
| TGW_19XX | AX-95148161 | 4A | 725839035 | -1.1484 | 1.10E-04 | BLUPmrMLM | SUG |
| TGW_19XX | AX-94508980 | 4B | 27522453 | -4.2743 | 1.65E-06 | GEMMA | SUG |
| TGW_19XX | AX-110980542 | 4B | 588429891 | -1.0355 | 3.58E-09 | BLUPmrMLM | SIG |
| TGW_19XX | AX-110604883 | 4D | 177489950 | 4.5062 | 3.07E-07 | FarmCPU | SUG |
| TGW_19XX | AX-109318201 | 4D | 197019433 | 4.3765 | 4.40E-07 | FarmCPU | SUG |
| TGW_19XX | AX-110853864 | 4D | 371786062 | -10.1380 | 1.33E-06 | EMMAX | SUG |
| TGW_19XX | AX-109275686 | 5A | 527997344 | 0.6736 | 3.53E-05 | BLUPmrMLM | SUG |
| TGW_19XX | AX-94420859 | 5B | 326086222 | -1.2102 | 1.63E-06 | BLUPmrMLM | SUG |
| TGW_19XX | AX-109880599 | 5B | 397831475 | -0.6107 | 3.55E-05 | BLUPmrMLM | SUG |
| TGW_19XX | AX-111494028 | 5B | 411541368 | 1.2850 | 1.69E-08 | BLUPmrMLM | SIG |
| TGW_19XX | AX-109402674 | 5B | 601331641 | -0.5495 | 2.76E-05 | BLUPmrMLM | SUG |
| TGW_19XX | AX-94511633 | 5B | 681168512 | 4.0385 | 1.28E-06 | FarmCPU | SUG |
| TGW_19XX | AX-94728376 | 5B | 703096796 | 1.3629 | 1.38E-08 | BLUPmrMLM | SIG |
| TGW_19XX | AX-109323599 | 5D | 541953656 | 0.9452 | 1.84E-06 | BLUPmrMLM | SUG |
| TGW_19XX | AX-111071251 | 6B | 11318645 | -4.3808 | 1.37E-06 | FarmCPU | SUG |
| TGW_19XX | AX-110429558 | 6B | 11319311 | -8.7531 | 1.70E-06 | FarmCPU | SUG |
| TGW_19XX | AX-109391962 | 6B | 339268873 | 5.1121 | 3.23E-08 | FarmCPU | SIG |
| TGW_19XX | AX-109364183 | 6B | 694461959 | 2.9106 | 1.91E-06 | FarmCPU | SUG |
| TGW_19XX | AX-111123882 | 6D | 272759669 | 4.5541 | 2.68E-07 | FarmCPU | SUG |
| TGW_19XX | AX-94562739 | 7A | 31876082 | -1.5931 | 2.55E-06 | BLUPmrMLM | SUG |
| TGW_19XX | AX-111153914 | 7A | 507877239 | 4.6572 | 5.71E-07 | FarmCPU | SUG |
| TGW_19XX | AX-109504215 | 7A | 638487210 | 1.5805 | 7.11E-07 | EMMAX | SUG |
| TGW_19XX | AX-109504215 | 7A | 638487210 | -1.5755 | 5.69E-07 | GEMMA | SUG |
| TGW_19XX | AX-111675730 | 7A | 638488309 | 1.5138 | 1.64E-06 | EMMAX | SUG |
| TGW_19XX | AX-111675730 | 7A | 638488309 | -1.4968 | 1.72E-06 | GEMMA | SUG |
| TGW_19XX | AX-108729470 | 7A | 638869864 | -0.8930 | 7.01E-06 | BLUPmrMLM | SUG |
| TGW_19XX | AX-108729470 | 7A | 638869864 | 1.6769 | 3.73E-07 | EMMAX | SUG |
| TGW_19XX | AX-108729470 | 7A | 638869864 | -1.6586 | 4.07E-07 | GEMMA | SUG |
| TGW_19XX | AX-109904527 | 7A | 640557150 | -1.5136 | 2.03E-06 | GEMMA | SUG |
| TGW_19XX | AX-110045543 | 7A | 640612936 | 1.5322 | 2.30E-06 | EMMAX | SUG |
| TGW_19XX | AX-110045543 | 7A | 640612936 | -1.5148 | 1.89E-06 | GEMMA | SUG |
| TGW_19XX | AX-108752431 | 7A | 641013507 | 1.4800 | 1.29E-06 | EMMAX | SUG |
| TGW_19XX | AX-108752431 | 7A | 641013507 | -1.4687 | 1.13E-06 | GEMMA | SUG |
| TGW_19XX | AX-109861376 | 7A | 699150090 | 0.6255 | 1.20E-04 | BLUPmrMLM | SUG |
| TGW_19XX | AX-108885233 | 7B | 84634645 | 0.7126 | 3.15E-06 | BLUPmrMLM | SUG |
| TGW_19XX | AX-94880394 | 7B | 253644552 | -5.3031 | 7.13E-07 | EMMAX | SUG |
| TGW_19XX | AX-94880394 | 7B | 253644552 | 5.2857 | 5.56E-07 | GEMMA | SUG |
| TGW_19XX | AX-95007233 | 7B | 253644571 | -5.3031 | 7.13E-07 | EMMAX | SUG |
| TGW_19XX | AX-95007233 | 7B | 253644571 | 5.2857 | 5.56E-07 | GEMMA | SUG |
| TGW_19XX | AX-94547633 | 7B | 253644650 | -5.3031 | 7.13E-07 | EMMAX | SUG |
| TGW_19XX | AX-94547633 | 7B | 253644650 | 5.2857 | 5.56E-07 | GEMMA | SUG |
| TGW_19XX | AX-111659217 | 7B | 254093708 | -5.4287 | 2.18E-07 | EMMAX | SUG |
| TGW_19XX | AX-111659217 | 7B | 254093708 | 5.4324 | 1.26E-07 | GEMMA | SUG |

| | | | | | | | | |
|---|---|---|---|---|---|---|---|---|
| TGW_19XX | AX-111659217 | 7B | 254093708 | -2.2554 | | 1.69E-43 | IIIVmrMLM | SIG |
| TGW_19XX | AX-94393280 | 7B | 254461692 | -5.3031 | | 7.13E-07 | EMMAX | SUG |
| TGW_19XX | AX-94393280 | 7B | 254461692 | 5.2857 | | 5.56E-07 | GEMMA | SUG |
| TGW_19XX | AX-110628362 | 7B | 334457457 | -5.8524 | | 1.96E-07 | EMMAX | SUG |
| TGW_19XX | AX-110628362 | 7B | 334457457 | 5.8167 | | 1.38E-07 | GEMMA | SUG |
| TGW_19XX | AX-95201674 | 7D | 277723188 | -5.3031 | | 7.13E-07 | EMMAX | SUG |
| TGW_19XX | AX-95201674 | 7D | 277723188 | 5.2857 | | 5.56E-07 | GEMMA | SUG |
| TGW_19XX | AX-110835487 | 7D | 426153254 | 4.9170 | | 3.89E-07 | FarmCPU | SUG |
| TGW_BLUP | AX-109005483 | 1A | 324487743 | -0.1604 | -1.2983 | 6.84E-07 | IIIVmrMLM | SUG |
| TGW_BLUP | AX-108797269 | 1A | 504613819 | -0.2016 | 0.9497 | 4.50E-16 | IIIVmrMLM | SIG |
| TGW_BLUP | AX-109427328 | 1A | 564586780 | 2.8285 | | 2.90E-09 | FarmCPU | SIG |
| TGW_BLUP | AX-111163116 | 1B | 297729910 | -0.3492 | | 7.60E-05 | BLUPmrMLM | SUG |
| TGW_BLUP | AX-108810059 | 1B | 300651321 | 0.4422 | 1.0295 | 3.73E-28 | IIIVmrMLM | SIG |
| TGW_BLUP | AX-109423631 | 1B | 334475559 | 0.3642 | -0.9551 | 9.28E-34 | IIIVmrMLM | SIG |
| TGW_BLUP | AX-110427177 | 1B | 582927346 | -0.3161 | | 1.83E-04 | BLUPmrMLM | SUG |
| TGW_BLUP | AX-108752498 | 1B | 644404296 | -0.2910 | | 5.87E-05 | BLUPmrMLM | SUG |
| TGW_BLUP | AX-95122382 | 1D | 6366932 | 0.2591 | | 4.83E-09 | IIIVmrMLM | SIG |
| TGW_BLUP | AX-110083733 | 1D | 41593584 | -0.4750 | | 7.43E-07 | BLUPmrMLM | SUG |
| TGW_BLUP | AX-109593928 | 1D | 68310644 | 0.4319 | -0.3765 | 1.97E-20 | IIIVmrMLM | SIG |
| TGW_BLUP | AX-95220243 | 1D | 95794331 | -1.1910 | | 1.02E-07 | BLUPmrMLM | SIG |
| TGW_BLUP | AX-108934860 | 2A | 70146016 | 0.4536 | 0.2605 | 3.05E-21 | IIIVmrMLM | SIG |
| TGW_BLUP | AX-111053248 | 2A | 71359445 | -0.7488 | | 1.86E-09 | BLUPmrMLM | SIG |
| TGW_BLUP | AX-109496621 | 2A | 148591856 | -0.1815 | 2.503 | 3.38E-15 | IIIVmrMLM | SIG |
| TGW_BLUP | AX-110175271 | 2A | 538007453 | -0.0769 | 1.9448 | 7.59E-07 | IIIVmrMLM | SUG |
| TGW_BLUP | AX-108859953 | 2A | 638302213 | 0.3979 | | 9.71E-06 | BLUPmrMLM | SUG |
| TGW_BLUP | AX-110600844 | 2A | 699606252 | -0.4421 | | 9.99E-07 | BLUPmrMLM | SUG |
| TGW_BLUP | AX-109386507 | 2A | 708050255 | 0.3801 | | 1.82E-06 | BLUPmrMLM | SUG |
| TGW_BLUP | AX-110989770 | 2A | 713246413 | -0.2573 | -2.1669 | 3.29E-18 | IIIVmrMLM | SIG |
| TGW_BLUP | AX-109951988 | 2A | 750561236 | -0.3515 | | 4.67E-07 | BLUPmrMLM | SUG |
| TGW_BLUP | AX-111559233 | 2A | 771062769 | -0.2642 | | 1.63E-04 | BLUPmrMLM | SUG |
| TGW_BLUP | AX-94794635 | 2B | 8592734 | 0.3603 | | 9.06E-16 | IIIVmrMLM | SIG |
| TGW_BLUP | AX-110524795 | 2B | 25077954 | -0.3794 | | 1.19E-05 | BLUPmrMLM | SUG |
| TGW_BLUP | AX-108824752 | 2B | 66805981 | 0.7532 | | 2.09E-09 | BLUPmrMLM | SIG |
| TGW_BLUP | AX-111066087 | 2B | 453828149 | -0.2964 | -0.2061 | 3.05E-10 | IIIVmrMLM | SIG |
| TGW_BLUP | AX-109304083 | 2B | 753972322 | -0.2789 | -1.1856 | 6.03E-15 | IIIVmrMLM | SIG |
| TGW_BLUP | AX-94750470 | 2D | 16344570 | 0.1990 | -0.6414 | 6.53E-12 | IIIVmrMLM | SIG |
| TGW_BLUP | AX-94628670 | 2D | 123215140 | 1.2790 | | 2.77E-07 | BLUPmrMLM | SUG |
| TGW_BLUP | AX-94628670 | 2D | 123215140 | -0.8686 | -0.1335 | 6.07E-11 | IIIVmrMLM | SIG |
| TGW_BLUP | AX-94577499 | 2D | 571882959 | -0.1460 | 0.344 | 3.48E-08 | IIIVmrMLM | SIG |
| TGW_BLUP | AX-109744601 | 2D | 614844617 | 0.3638 | 1.3098 | 1.43E-27 | IIIVmrMLM | SIG |
| TGW_BLUP | AX-109185488 | 2D | 642969692 | -2.0411 | | 1.98E-06 | EMMAX | SUG |
| TGW_BLUP | AX-109185488 | 2D | 642969692 | 2.0393 | | 1.31E-06 | GEMMA | SUG |
| TGW_BLUP | AX-109185488 | 2D | 642969692 | -0.2944 | 1.2256 | 1.36E-22 | IIIVmrMLM | SIG |
| TGW_BLUP | AX-111050852 | 3A | 208354 | 0.0115 | 3.22 | 3.35E-08 | IIIVmrMLM | SIG |

| | | | | | | | | |
|---|---|---|---|---|---|---|---|---|
| TGW_BLUP | AX-108997118 | 3A | 7260087 | -0.4041 | 0.313 | 1.04E-17 | IIIVmrMLM | SIG |
| TGW_BLUP | AX-95228570 | 3A | 586408529 | -0.4602 | | 5.78E-05 | BLUPmrMLM | SUG |
| TGW_BLUP | AX-110975740 | 3A | 656099548 | 0.2763 | | 2.09E-05 | BLUPmrMLM | SUG |
| TGW_BLUP | AX-110581785 | 3A | 695679602 | 1.0181 | 0.2827 | 1.42E-81 | IIIVmrMLM | SIG |
| TGW_BLUP | AX-109019801 | 3A | 747613640 | 0.1486 | 0.8118 | 8.80E-18 | IIIVmrMLM | SIG |
| TGW_BLUP | AX-95247325 | 3B | 66056980 | -0.4515 | 0.0451 | 1.18E-09 | IIIVmrMLM | SIG |
| TGW_BLUP | AX-109274638 | 3B | 237325305 | -2.3543 | | 2.05E-10 | FarmCPU | SIG |
| TGW_BLUP | AX-109095830 | 3B | 410811276 | -0.2666 | -1.2582 | 9.23E-14 | IIIVmrMLM | SIG |
| TGW_BLUP | AX-108878936 | 3B | 558567987 | -0.4295 | | 4.33E-05 | BLUPmrMLM | SUG |
| TGW_BLUP | AX-108878936 | 3B | 558567987 | 0.5982 | 0.0318 | 8.21E-25 | IIIVmrMLM | SIG |
| TGW_BLUP | AX-109923823 | 3B | 749771886 | -0.3630 | | 1.68E-06 | BLUPmrMLM | SUG |
| TGW_BLUP | AX-111618044 | 3B | 750117968 | 0.3334 | -1.0778 | 6.90E-14 | IIIVmrMLM | SIG |
| TGW_BLUP | AX-110426026 | 3B | 762431803 | 0.1831 | -1.5172 | 2.06E-12 | IIIVmrMLM | SIG |
| TGW_BLUP | AX-110502956 | 4A | 152193158 | -0.5070 | 0.1811 | 3.43E-25 | IIIVmrMLM | SIG |
| TGW_BLUP | AX-110058508 | 4A | 728090480 | 0.3087 | | 1.92E-05 | BLUPmrMLM | SUG |
| TGW_BLUP | AX-95203615 | 4B | 561516400 | -0.4019 | -0.1215 | 1.54E-10 | IIIVmrMLM | SIG |
| TGW_BLUP | AX-110169646 | 5A | 10556613 | 2.3792 | | 2.04E-06 | GEMMA | SUG |
| TGW_BLUP | AX-111112946 | 5A | 424008295 | -0.3386 | -0.2363 | 5.61E-13 | IIIVmrMLM | SIG |
| TGW_BLUP | AX-110201131 | 5A | 448597261 | 0.5970 | | 2.61E-12 | BLUPmrMLM | SIG |
| TGW_BLUP | AX-109287745 | 5A | 526617830 | 0.7950 | | 2.61E-10 | BLUPmrMLM | SIG |
| TGW_BLUP | AX-94686717 | 5A | 533072113 | -0.5970 | -0.4396 | 9.16E-34 | IIIVmrMLM | SIG |
| TGW_BLUP | AX-110400975 | 5A | 567921322 | 0.0945 | -2.8197 | 1.41E-11 | IIIVmrMLM | SIG |
| TGW_BLUP | AX-89598451 | 5A | 580797853 | 0.4150 | | 1.69E-06 | BLUPmrMLM | SUG |
| TGW_BLUP | AX-109496894 | 5B | 7736591 | 0.7800 | | 6.89E-58 | IIIVmrMLM | SIG |
| TGW_BLUP | AX-111533119 | 5B | 27876582 | -0.2096 | 2.1213 | 7.47E-12 | IIIVmrMLM | SIG |
| TGW_BLUP | AX-109018708 | 5B | 405722389 | -0.5390 | 0.4524 | 4.68E-30 | IIIVmrMLM | SIG |
| TGW_BLUP | AX-110125323 | 5B | 409803018 | 0.4547 | | 1.86E-05 | BLUPmrMLM | SUG |
| TGW_BLUP | AX-109052486 | 5B | 463223004 | -0.3504 | | 3.55E-05 | BLUPmrMLM | SUG |
| TGW_BLUP | AX-110385695 | 5B | 690089042 | 0.6060 | | 8.91E-08 | BLUPmrMLM | SIG |
| TGW_BLUP | AX-109935774 | 5B | 694009767 | -0.3097 | -1.1486 | 6.81E-13 | IIIVmrMLM | SIG |
| TGW_BLUP | AX-94504308 | 5D | 74467144 | 0.0511 | 2.2529 | 3.39E-09 | IIIVmrMLM | SIG |
| TGW_BLUP | AX-109407131 | 5D | 512550680 | 0.3920 | 0.8834 | 1.64E-18 | IIIVmrMLM | SIG |
| TGW_BLUP | AX-109823970 | 6A | 1040181 | -0.1325 | 1.0171 | 1.20E-05 | IIIVmrMLM | SUG |
| TGW_BLUP | AX-95247642 | 6A | 21442944 | -0.1487 | 1.0291 | 7.98E-22 | IIIVmrMLM | SIG |
| TGW_BLUP | AX-110511426 | 6A | 73837265 | -0.1330 | -1.1062 | 6.69E-08 | IIIVmrMLM | SIG |
| TGW_BLUP | AX-110991104 | 6A | 79266467 | 0.3558 | 0.1436 | 6.26E-14 | IIIVmrMLM | SIG |
| TGW_BLUP | AX-110366132 | 6A | 617809451 | 0.5808 | | 9.64E-08 | BLUPmrMLM | SIG |
| TGW_BLUP | AX-109375407 | 6B | 18281605 | -0.4434 | | 2.95E-07 | BLUPmrMLM | SUG |
| TGW_BLUP | AX-110493821 | 6B | 29671148 | 0.3646 | | 3.88E-07 | BLUPmrMLM | SUG |
| TGW_BLUP | AX-109391962 | 6B | 339268873 | 3.5652 | | 1.84E-07 | FarmCPU | SUG |
| TGW_BLUP | AX-108762235 | 6B | 679245634 | 0.0519 | -0.8328 | 6.47E-06 | IIIVmrMLM | SUG |
| TGW_BLUP | AX-109847836 | 6D | 2372237 | -0.0401 | -2.265 | 2.78E-08 | IIIVmrMLM | SIG |
| TGW_BLUP | AX-108897553 | 6D | 470390265 | 0.0170 | -2.103 | 6.73E-22 | IIIVmrMLM | SIG |
| TGW_BLUP | AX-109081729 | 7A | 648700205 | 0.5432 | 2.0223 | 7.01E-39 | IIIVmrMLM | SIG |

| | | | | | | | | |
|---|---|---|---|---|---|---|---|---|
| TGW_BLUP | AX-110984004 | 7A | 673020654 | -0.4250 | | 2.83E-05 | BLUPmrMLM | SUG |
| TGW_BLUP | AX-108839894 | 7A | 678153714 | -0.0130 | -1.0361 | 2.21E-06 | IIIVmrMLM | SUG |
| TGW_BLUP | AX-108777232 | 7B | 67698816 | -0.0316 | 0.7649 | 3.22E-12 | IIIVmrMLM | SIG |
| TGW_BLUP | AX-109524696 | 7B | 734407845 | -0.9608 | | 6.20E-11 | BLUPmrMLM | SIG |
| TGW_BLUP | AX-89403396 | 7D | 124291890 | -0.2266 | | 2.81E-07 | IIIVmrMLM | SUG |
| TGW_BLUP | AX-110730551 | 7D | 344688778 | -0.0016 | 1.776 | 7.06E-06 | IIIVmrMLM | SUG |
| TGW_BLUP | AX-111353936 | 7D | 566214823 | 0.5012 | | 2.40E-06 | BLUPmrMLM | SUG |

TGW: thousand grain weight.

**Supplementary Table S13**. All QTNs identified to be associated with spikelet number per spike in wheat.

| Trait | Marker | Chr | Position | Add.effect | Dom.effect | P-value | Method | Sig. |
|---|---|---|---|---|---|---|---|---|
| SNS_17AY | AX-110006279 | 1A | 323184555 | -0.5346 | -0.3613 | 5.37E-23 | IIIVmrMLM | SIG |
| SNS_17AY | AX-110587308 | 1A | 587844131 | 0.9633 | -0.0111 | 2.7E-64 | IIIVmrMLM | SIG |
| SNS_17AY | AX-108854586 | 1B | 335880361 | -0.1512 | -0.568 | 5.75E-06 | IIIVmrMLM | SUG |
| SNS_17AY | AX-110663509 | 1D | 22380094 | 0.4445 | | 4.57E-05 | BLUPmrMLM | SUG |
| SNS_17AY | AX-110663509 | 1D | 22380094 | -0.5456 | -0.1039 | 2.91E-18 | IIIVmrMLM | SIG |
| SNS_17AY | AX-109314532 | 1D | 427488585 | 0.3603 | | 0.000162 | BLUPmrMLM | SUG |
| SNS_17AY | AX-94401351 | 2A | 24029377 | -0.9089 | | 7.21E-07 | GEMMA | SUG |
| SNS_17AY | AX-95126447 | 2A | 24060452 | 0.9806 | | 8.98E-07 | EMMAX | SUG |
| SNS_17AY | AX-95126447 | 2A | 24060452 | -0.9818 | | 9.46E-08 | GEMMA | SIG |
| SNS_17AY | AX-109084387 | 2A | 312488660 | -0.0134 | 1.1442 | 0.000128 | IIIVmrMLM | SUG |
| SNS_17AY | AX-94758817 | 2A | 740873756 | 0.4976 | | 0.000103 | BLUPmrMLM | SUG |
| SNS_17AY | AX-110990619 | 2B | 34283542 | -0.5565 | | 1.06E-06 | GEMMA | SUG |
| SNS_17AY | AX-89763994 | 2B | 34284408 | -0.5598 | | 6.73E-07 | GEMMA | SUG |
| SNS_17AY | AX-95087661 | 2B | 284934379 | -0.2735 | | 0.000124 | BLUPmrMLM | SUG |
| SNS_17AY | AX-111486438 | 2D | 20866150 | -0.5200 | | 1.96E-06 | GEMMA | SUG |
| SNS_17AY | AX-110377320 | 2D | 20940622 | -0.4789 | | 2.06E-06 | GEMMA | SUG |
| SNS_17AY | AX-111772631 | 2D | 20982871 | -0.4971 | | 1.02E-06 | GEMMA | SUG |
| SNS_17AY | AX-109817136 | 2D | 20983979 | -0.5100 | | 1.32E-06 | GEMMA | SUG |
| SNS_17AY | AX-110919716 | 2D | 21002736 | -0.4838 | | 1.51E-06 | GEMMA | SUG |
| SNS_17AY | AX-110411772 | 2D | 21003083 | -0.4900 | | 2.12E-06 | GEMMA | SUG |
| SNS_17AY | AX-110373790 | 2D | 21004025 | -0.4890 | | 1.25E-06 | GEMMA | SUG |
| SNS_17AY | AX-110926124 | 2D | 21004115 | -0.5691 | | 7.59E-07 | GEMMA | SUG |
| SNS_17AY | AX-109917920 | 2D | 21004362 | -0.4736 | | 2.31E-06 | GEMMA | SUG |
| SNS_17AY | AX-110967278 | 2D | 21004963 | -0.5593 | | 1.56E-06 | GEMMA | SUG |
| SNS_17AY | AX-111589614 | 2D | 21006215 | -0.5623 | | 7.34E-07 | GEMMA | SUG |
| SNS_17AY | AX-108922534 | 2D | 21007167 | -0.4777 | | 2.02E-06 | GEMMA | SUG |
| SNS_17AY | AX-110920552 | 2D | 21009425 | -0.4901 | | 1.37E-06 | GEMMA | SUG |
| SNS_17AY | AX-109347692 | 2D | 21009516 | -0.5311 | | 7.6E-07 | GEMMA | SUG |
| SNS_17AY | AX-110286662 | 2D | 21009875 | -0.5640 | | 1.64E-06 | GEMMA | SUG |
| SNS_17AY | AX-109679370 | 2D | 21012260 | -0.5825 | | 1.65E-06 | GEMMA | SUG |
| SNS_17AY | AX-109847823 | 2D | 21012368 | -0.9697 | | 1.81E-06 | GEMMA | SUG |
| SNS_17AY | AX-109404810 | 2D | 21014695 | 0.5507 | | 2.16E-06 | EMMAX | SUG |
| SNS_17AY | AX-109404810 | 2D | 21014695 | -0.5517 | | 2.73E-07 | GEMMA | SUG |
| SNS_17AY | AX-109313347 | 2D | 21014765 | -0.5445 | | 5.38E-07 | GEMMA | SUG |
| SNS_17AY | AX-111476707 | 2D | 21026487 | 0.5889 | | 2.09E-06 | EMMAX | SUG |
| SNS_17AY | AX-111476707 | 2D | 21026487 | -0.5890 | | 3.59E-07 | GEMMA | SUG |
| SNS_17AY | AX-108769343 | 2D | 21027592 | -1.1493 | | 1.65E-06 | GEMMA | SUG |
| SNS_17AY | AX-111117886 | 2D | 21028457 | -0.5065 | | 4.34E-07 | GEMMA | SUG |
| SNS_17AY | AX-110459643 | 2D | 21051652 | -0.5278 | | 5.77E-07 | GEMMA | SUG |
| SNS_17AY | AX-94782932 | 2D | 21061255 | -0.9985 | | 9.57E-07 | GEMMA | SUG |
| SNS_17AY | AX-111124839 | 2D | 21087438 | -1.0018 | | 8E-07 | GEMMA | SUG |

| | | | | | | | | |
|---|---|---|---|---|---|---|---|---|
| SNS_17AY | AX-111814682 | 2D | 21398067 | -0.5155 | | 1.1E-06 | GEMMA | SUG |
| SNS_17AY | AX-110018799 | 2D | 21534949 | -0.4931 | | 1.29E-06 | GEMMA | SUG |
| SNS_17AY | AX-109825834 | 2D | 21554281 | -0.5795 | | 5.23E-07 | GEMMA | SUG |
| SNS_17AY | AX-111574942 | 2D | 21911320 | -0.4524 | | 1.3E-06 | GEMMA | SUG |
| SNS_17AY | AX-110516159 | 2D | 21915211 | -0.4173 | | 3.06E-10 | BLUPmrMLM | SIG |
| SNS_17AY | AX-110516159 | 2D | 21915211 | -0.4482 | | 1.17E-06 | GEMMA | SUG |
| SNS_17AY | AX-110266075 | 2D | 22176657 | -0.4416 | | 1.92E-06 | GEMMA | SUG |
| SNS_17AY | AX-89576820 | 2D | 22176723 | -0.4386 | | 2.14E-06 | GEMMA | SUG |
| SNS_17AY | AX-108740376 | 2D | 22203367 | -0.4578 | | 7.72E-07 | GEMMA | SUG |
| SNS_17AY | AX-110553852 | 2D | 22203646 | -0.4429 | | 1.81E-06 | GEMMA | SUG |
| SNS_17AY | AX-89702253 | 2D | 22203843 | -0.4578 | | 7.72E-07 | GEMMA | SUG |
| SNS_17AY | AX-110058858 | 2D | 22237096 | -0.4575 | | 7.89E-07 | GEMMA | SUG |
| SNS_17AY | AX-111444111 | 2D | 22255143 | -0.4651 | | 5.77E-07 | GEMMA | SUG |
| SNS_17AY | AX-109880678 | 2D | 22281780 | -0.4604 | | 6.69E-07 | GEMMA | SUG |
| SNS_17AY | AX-111634202 | 2D | 22288039 | -0.4526 | | 1.05E-06 | GEMMA | SUG |
| SNS_17AY | AX-110712373 | 2D | 22288246 | -0.4575 | | 7.89E-07 | GEMMA | SUG |
| SNS_17AY | AX-109363180 | 2D | 22349863 | -0.4733 | | 3.32E-07 | GEMMA | SUG |
| SNS_17AY | AX-110329747 | 2D | 22350020 | -0.4613 | | 6.58E-07 | GEMMA | SUG |
| SNS_17AY | AX-109972094 | 2D | 22350525 | -0.4664 | | 4.99E-07 | GEMMA | SUG |
| SNS_17AY | AX-110348073 | 2D | 22350570 | -0.4635 | | 5.57E-07 | GEMMA | SUG |
| SNS_17AY | AX-110873504 | 2D | 22351495 | -0.4712 | | 3.92E-07 | GEMMA | SUG |
| SNS_17AY | AX-109839882 | 2D | 22357144 | -0.4659 | | 5.48E-07 | GEMMA | SUG |
| SNS_17AY | AX-111467599 | 2D | 22392429 | -0.4653 | | 5.54E-07 | GEMMA | SUG |
| SNS_17AY | AX-110966612 | 2D | 22419146 | -0.4573 | | 8.15E-07 | GEMMA | SUG |
| SNS_17AY | AX-111990365 | 2D | 22419647 | -0.4642 | | 5.62E-07 | GEMMA | SUG |
| SNS_17AY | AX-94745278 | 2D | 22451354 | -0.4772 | | 3.9E-07 | GEMMA | SUG |
| SNS_17AY | AX-109410970 | 2D | 22460303 | -0.4679 | | 4.31E-07 | GEMMA | SUG |
| SNS_17AY | AX-111227264 | 2D | 22460692 | -0.4773 | | 2.82E-07 | GEMMA | SUG |
| SNS_17AY | AX-111105994 | 2D | 22461473 | -0.4715 | | 3.85E-07 | GEMMA | SUG |
| SNS_17AY | AX-95075528 | 2D | 468097701 | -1.9206 | | 7.09E-05 | BLUPmrMLM | SUG |
| SNS_17AY | AX-109394789 | 3A | 695517857 | 0.1115 | -0.5738 | 2.4E-06 | IIIVmrMLM | SUG |
| SNS_17AY | AX-110974574 | 3B | 223514983 | -0.2706 | | 0.000109 | BLUPmrMLM | SUG |
| SNS_17AY | AX-108909296 | 3B | 285415864 | 0.4308 | | 3E-05 | BLUPmrMLM | SUG |
| SNS_17AY | AX-94570847 | 3B | 419308930 | 0.2708 | | 2.28E-08 | IIIVmrMLM | SIG |
| SNS_17AY | AX-108892190 | 3B | 507303122 | 0.0171 | -0.9355 | 5.08E-08 | IIIVmrMLM | SIG |
| SNS_17AY | AX-111800988 | 3B | 586755775 | -0.0046 | -1.0242 | 4.38E-09 | IIIVmrMLM | SIG |
| SNS_17AY | AX-110561020 | 3B | 712554456 | 0.3563 | | 1.15E-05 | BLUPmrMLM | SUG |
| SNS_17AY | AX-109653319 | 3B | 717368674 | 0.2147 | 1.2342 | 5.72E-22 | IIIVmrMLM | SIG |
| SNS_17AY | AX-109473335 | 3B | 729668082 | 0.3668 | -0.9727 | 1.17E-14 | IIIVmrMLM | SIG |
| SNS_17AY | AX-94557205 | 3B | 816264746 | 0.5235 | | 4.93E-05 | BLUPmrMLM | SUG |
| SNS_17AY | AX-110931712 | 3D | 609949401 | -0.0839 | 0.8944 | 0.000101 | IIIVmrMLM | SUG |
| SNS_17AY | AX-111746554 | 3D | 613384351 | 0.4062 | | 1.54E-16 | IIIVmrMLM | SIG |
| SNS_17AY | AX-89363204 | 4A | 4827482 | -0.1196 | -1.1636 | 5.03E-08 | IIIVmrMLM | SIG |
| SNS_17AY | AX-109460280 | 4A | 51131830 | 0.0810 | -2.0882 | 4.98E-08 | IIIVmrMLM | SIG |

| | | | | | | | | |
|---|---|---|---|---|---|---|---|---|
| SNS_17AY | AX-110964870 | 4A | 78729700 | -0.1782 | -0.7463 | 1.75E-13 | IIIVmrMLM | SIG |
| SNS_17AY | AX-110008956 | 4B | 2148230 | -0.4396 | | 7.7E-10 | BLUPmrMLM | SIG |
| SNS_17AY | AX-110112546 | 4B | 627321890 | 0.3947 | 0.2596 | 8.18E-14 | IIIVmrMLM | SIG |
| SNS_17AY | AX-109295342 | 4B | 659152345 | -0.3250 | | 9.56E-05 | BLUPmrMLM | SUG |
| SNS_17AY | AX-108942840 | 5B | 38032372 | 0.1055 | 0.6942 | 1.03E-14 | IIIVmrMLM | SIG |
| SNS_17AY | AX-110477130 | 5B | 420367588 | 0.3890 | | 1.07E-06 | BLUPmrMLM | SUG |
| SNS_17AY | AX-109004813 | 5B | 462729389 | -0.1202 | 1.1709 | 2.69E-07 | IIIVmrMLM | SUG |
| SNS_17AY | AX-111469888 | 5B | 500533382 | -0.2964 | 0.304 | 6.94E-09 | IIIVmrMLM | SIG |
| SNS_17AY | AX-109890969 | 5B | 532821470 | 0.1060 | 1.1574 | 2.67E-07 | IIIVmrMLM | SUG |
| SNS_17AY | AX-111806645 | 5D | 323622 | 0.0130 | -1.493 | 6.23E-10 | IIIVmrMLM | SIG |
| SNS_17AY | AX-94966145 | 6A | 5221726 | -0.1381 | 2.3518 | 4.79E-10 | IIIVmrMLM | SIG |
| SNS_17AY | AX-111739834 | 6A | 595061673 | -0.3124 | | 0.000129 | BLUPmrMLM | SUG |
| SNS_17AY | AX-111564326 | 6B | 461588956 | -0.0673 | -1.167 | 3.21E-07 | IIIVmrMLM | SUG |
| SNS_17AY | AX-111529384 | 6B | 643439687 | -0.1381 | 0.4665 | 1.26E-12 | IIIVmrMLM | SIG |
| SNS_17AY | AX-111564275 | 6D | 109894053 | -0.5004 | | 8.37E-07 | GEMMA | SUG |
| SNS_17AY | AX-110436028 | 6D | 109896038 | -0.4809 | | 2.1E-06 | GEMMA | SUG |
| SNS_17AY | AX-109395167 | 6D | 465577457 | -0.2358 | | 9.1E-05 | BLUPmrMLM | SUG |
| SNS_17AY | AX-110567419 | 7A | 556169911 | 0.5006 | | 2.31E-06 | EMMAX | SUG |
| SNS_17AY | AX-110567419 | 7A | 556169911 | -0.5001 | | 2.13E-06 | GEMMA | SUG |
| SNS_17AY | AX-110019761 | 7A | 556177761 | 0.4910 | | 2.11E-06 | EMMAX | SUG |
| SNS_17AY | AX-109964606 | 7A | 556178563 | 0.4982 | | 1.43E-06 | EMMAX | SUG |
| SNS_17AY | AX-109964606 | 7A | 556178563 | -0.4999 | | 2.2E-06 | GEMMA | SUG |
| SNS_17AY | AX-95120206 | 7A | 556504067 | 0.5086 | | 1.95E-06 | EMMAX | SUG |
| SNS_17AY | AX-95120206 | 7A | 556504067 | -0.5091 | | 2.25E-06 | GEMMA | SUG |
| SNS_17AY | AX-108737878 | 7A | 556504864 | 0.5093 | | 1.87E-06 | EMMAX | SUG |
| SNS_17AY | AX-108737878 | 7A | 556504864 | -0.5099 | | 2.21E-06 | GEMMA | SUG |
| SNS_17AY | AX-110480283 | 7A | 556504926 | 0.5093 | | 1.87E-06 | EMMAX | SUG |
| SNS_17AY | AX-110480283 | 7A | 556504926 | -0.5099 | | 2.21E-06 | GEMMA | SUG |
| SNS_17AY | AX-108775166 | 7A | 556505602 | 0.5299 | | 4.02E-07 | EMMAX | SUG |
| SNS_17AY | AX-108775166 | 7A | 556505602 | -0.5299 | | 4.23E-07 | GEMMA | SUG |
| SNS_17AY | AX-108923531 | 7A | 556506636 | 0.5093 | | 1.87E-06 | EMMAX | SUG |
| SNS_17AY | AX-108923531 | 7A | 556506636 | -0.5099 | | 2.21E-06 | GEMMA | SUG |
| SNS_17AY | AX-109432838 | 7A | 556506760 | 0.5093 | | 1.87E-06 | EMMAX | SUG |
| SNS_17AY | AX-109432838 | 7A | 556506760 | -0.5099 | | 2.21E-06 | GEMMA | SUG |
| SNS_17AY | AX-111477381 | 7A | 556569046 | 0.5305 | | 1.05E-06 | EMMAX | SUG |
| SNS_17AY | AX-111477381 | 7A | 556569046 | -0.5326 | | 1.45E-06 | GEMMA | SUG |
| SNS_17AY | AX-94680052 | 7A | 675594466 | -0.2823 | | 5.87E-09 | IIIVmrMLM | SIG |
| SNS_17AY | AX-111198368 | 7A | 706250295 | -0.4229 | 0.3597 | 1.21E-16 | IIIVmrMLM | SIG |
| SNS_17AY | AX-109961681 | 7B | 328341775 | 0.0436 | -1.4696 | 1.41E-06 | IIIVmrMLM | SUG |
| SNS_17AY | AX-111289984 | 7B | 595860938 | 0.0677 | 1.5019 | 0.000256 | IIIVmrMLM | SUG |
| SNS_17AY | AX-109580774 | 7B | 713458863 | -0.0872 | -2.1363 | 2.27E-06 | IIIVmrMLM | SUG |
| SNS_17AY | AX-108928296 | 7D | 448853259 | -0.3383 | | 1.94E-05 | BLUPmrMLM | SUG |
| SNS_17JY | AX-109278778 | 1A | 43954590 | 1.0115 | 6.3757 | 1.52E-48 | IIIVmrMLM | SIG |
| SNS_17JY | AX-108903063 | 1A | 45630334 | -0.8194 | 1.6305 | 1.2E-31 | IIIVmrMLM | SIG |

| | | | | | | | | |
|---|---|---|---|---|---|---|---|---|
| SNS_17JY | AX-109879655 | 1A | 58842504 | 1.0511 | | 1.16E-06 | GEMMA | SUG |
| SNS_17JY | AX-109946801 | 1A | 474316304 | -0.5613 | | 4.34E-06 | BLUPmrMLM | SUG |
| SNS_17JY | AX-110960919 | 1A | 564128400 | -0.0070 | -2.3183 | 3.16E-06 | IIIVmrMLM | SUG |
| SNS_17JY | AX-111229470 | 1B | 521058454 | -0.0313 | -1.0539 | 2.89E-16 | IIIVmrMLM | SIG |
| SNS_17JY | AX-109464119 | 1B | 627661839 | -0.0801 | 0.928 | 0.00021 | IIIVmrMLM | SUG |
| SNS_17JY | AX-111175329 | 2A | 538368699 | -0.0384 | 1.3097 | 3.35E-06 | IIIVmrMLM | SUG |
| SNS_17JY | AX-111255386 | 2A | 648353981 | -0.1078 | 4.1922 | 4.3E-10 | IIIVmrMLM | SIG |
| SNS_17JY | AX-111075043 | 2B | 68176680 | -0.3914 | | 0.000118 | BLUPmrMLM | SUG |
| SNS_17JY | AX-112285965 | 2B | 214593242 | 0.0769 | -0.8322 | 3.52E-07 | IIIVmrMLM | SUG |
| SNS_17JY | AX-111187069 | 2B | 425436545 | -0.5294 | 2.6211 | 6.77E-17 | IIIVmrMLM | SIG |
| SNS_17JY | AX-110390570 | 2B | 539699807 | 0.0142 | -1.4483 | 4.66E-10 | IIIVmrMLM | SIG |
| SNS_17JY | AX-111439177 | 2D | 601511174 | -0.6996 | | 2.9E-06 | BLUPmrMLM | SUG |
| SNS_17JY | AX-110679559 | 3A | 485912014 | -0.7113 | -0.0633 | 1.57E-21 | IIIVmrMLM | SIG |
| SNS_17JY | AX-94451753 | 3B | 3271249 | -0.4220 | 0.5419 | 5.06E-09 | IIIVmrMLM | SIG |
| SNS_17JY | AX-109960381 | 3B | 481926049 | -0.1419 | 1.4095 | 4.22E-09 | IIIVmrMLM | SIG |
| SNS_17JY | AX-109939994 | 3B | 499530064 | 0.7395 | | 3.11E-07 | BLUPmrMLM | SUG |
| SNS_17JY | AX-109939994 | 3B | 499530064 | -0.6183 | 0.2282 | 5.23E-11 | IIIVmrMLM | SIG |
| SNS_17JY | AX-109379765 | 3B | 767539773 | 0.5926 | | 2.99E-06 | BLUPmrMLM | SUG |
| SNS_17JY | AX-111705710 | 3D | 6411526 | -0.1154 | -3.2748 | 6.45E-06 | IIIVmrMLM | SUG |
| SNS_17JY | AX-94465953 | 3D | 591878287 | 1.2617 | 0.7714 | 8.26E-56 | IIIVmrMLM | SIG |
| SNS_17JY | AX-109080534 | 4A | 261106855 | -0.1208 | 7.7139 | 1.81E-06 | IIIVmrMLM | SUG |
| SNS_17JY | AX-110433115 | 4A | 743849873 | -0.2318 | -0.7718 | 2.65E-10 | IIIVmrMLM | SIG |
| SNS_17JY | AX-111173338 | 4B | 498406298 | 0.1259 | -1.8741 | 1.23E-11 | IIIVmrMLM | SIG |
| SNS_17JY | AX-109622624 | 4D | 392756716 | 0.6391 | | 0.000157 | BLUPmrMLM | SUG |
| SNS_17JY | AX-109297427 | 5A | 480786023 | -0.6084 | 0.0252 | 4.45E-17 | IIIVmrMLM | SIG |
| SNS_17JY | AX-108757990 | 5A | 545877401 | 0.9272 | | 6.26E-05 | BLUPmrMLM | SUG |
| SNS_17JY | AX-111064823 | 5A | 562204317 | 0.2227 | -2.8955 | 6.03E-08 | IIIVmrMLM | SIG |
| SNS_17JY | AX-110979025 | 5A | 587979513 | 0.5543 | | 1.08E-06 | BLUPmrMLM | SUG |
| SNS_17JY | AX-111131744 | 5B | 69667894 | 0.1783 | -1.611 | 6.38E-10 | IIIVmrMLM | SIG |
| SNS_17JY | AX-108863198 | 5B | 84714081 | -0.2543 | 0.7478 | 3.38E-07 | IIIVmrMLM | SUG |
| SNS_17JY | AX-109292922 | 5B | 297269030 | -0.1458 | 2.1886 | 1.27E-11 | IIIVmrMLM | SIG |
| SNS_17JY | AX-110065016 | 5B | 393575896 | 0.0716 | 2.0369 | 0.000357 | IIIVmrMLM | SUG |
| SNS_17JY | AX-111051520 | 5B | 562278179 | -0.5875 | | 5.18E-05 | BLUPmrMLM | SUG |
| SNS_17JY | AX-109016185 | 6A | 556094742 | -0.3734 | | 5.1E-05 | BLUPmrMLM | SUG |
| SNS_17JY | AX-110396365 | 6A | 594523770 | 0.4603 | | 7.49E-05 | BLUPmrMLM | SUG |
| SNS_17JY | AX-109922096 | 6A | 594739977 | -0.9650 | 0.6368 | 2.42E-34 | IIIVmrMLM | SIG |
| SNS_17JY | AX-108848956 | 6B | 301568609 | 0.0707 | 5.5584 | 3.17E-06 | IIIVmrMLM | SUG |
| SNS_17JY | AX-109986863 | 6B | 655886208 | 0.8274 | | 2.21E-06 | BLUPmrMLM | SUG |
| SNS_17JY | AX-94502175 | 6B | 660380975 | 0.3002 | | 1.08E-05 | IIIVmrMLM | SUG |
| SNS_17JY | AX-110349778 | 6D | 117374296 | -0.4216 | -1.2943 | 3.58E-11 | IIIVmrMLM | SIG |
| SNS_17JY | AX-111055096 | 7A | 128117104 | -0.0596 | 0.9228 | 3.6E-06 | IIIVmrMLM | SUG |
| SNS_17JY | AX-94670021 | 7D | 4005207 | 0.4860 | -1.0214 | 1.14E-13 | IIIVmrMLM | SIG |
| SNS_17JY | AX-111131217 | 7D | 8179821 | 0.0203 | 4.762 | 3.02E-10 | IIIVmrMLM | SIG |
| SNS_17JY | AX-94565322 | 7D | 475727021 | 0.0075 | -2.5192 | 2.84E-09 | IIIVmrMLM | SIG |

| | | | | | | | | |
|---|---|---|---|---|---|---|---|---|
| SNS_17JY | AX-95189491 | 7D | 495142627 | -0.9640 | | 3.55E-07 | BLUPmrMLM | SUG |
| SNS_17JY | AX-95189491 | 7D | 495142627 | 0.3408 | | 6.32E-07 | IIIVmrMLM | SUG |
| SNS_17JY | AX-94893648 | 7D | 616256903 | 0.7379 | | 8.26E-05 | BLUPmrMLM | SUG |
| SNS_17JY | AX-94893648 | 7D | 616256903 | -0.5200 | 0.2706 | 5.26E-10 | IIIVmrMLM | SIG |
| SNS_18AY | AX-109281308 | 1A | 39306547 | -0.3319 | | 3.61E-05 | BLUPmrMLM | SUG |
| SNS_18AY | AX-110983558 | 1A | 536975551 | 0.2126 | | 2.19E-05 | BLUPmrMLM | SUG |
| SNS_18AY | AX-111175338 | 1A | 581439146 | 0.4073 | | 3.98E-07 | BLUPmrMLM | SUG |
| SNS_18AY | AX-111022083 | 1B | 564718030 | 0.0890 | -2.6878 | 4.97E-09 | IIIVmrMLM | SIG |
| SNS_18AY | AX-108953140 | 2A | 115123884 | 0.2794 | | 2.53E-05 | BLUPmrMLM | SUG |
| SNS_18AY | AX-109480830 | 2A | 189167477 | -0.4513 | | 4.58E-30 | IIIVmrMLM | SIG |
| SNS_18AY | AX-110966443 | 2A | 719720650 | -0.3180 | 0.0137 | 1.14E-11 | IIIVmrMLM | SIG |
| SNS_18AY | AX-111561722 | 2A | 721022021 | 0.2049 | | 6.88E-05 | BLUPmrMLM | SUG |
| SNS_18AY | AX-94852581 | 2A | 745670462 | 0.0334 | 1.4887 | 1.05E-11 | IIIVmrMLM | SIG |
| SNS_18AY | AX-110089929 | 2B | 361128622 | -0.0972 | -0.7523 | 7.4E-07 | IIIVmrMLM | SUG |
| SNS_18AY | AX-111063611 | 2D | 34279198 | 0.1440 | 1.0408 | 8.22E-09 | IIIVmrMLM | SIG |
| SNS_18AY | AX-109755758 | 2D | 156358859 | -0.2869 | | 8.09E-06 | BLUPmrMLM | SUG |
| SNS_18AY | AX-109289848 | 3A | 268219162 | 0.2662 | 0.3272 | 2.64E-11 | IIIVmrMLM | SIG |
| SNS_18AY | AX-94522205 | 3A | 638010015 | -0.6313 | | 7.21E-07 | FarmCPU | SUG |
| SNS_18AY | AX-110484538 | 3A | 705296840 | 2.5931 | | 7.71E-07 | GEMMA | SUG |
| SNS_18AY | AX-109305421 | 3B | 757013 | -0.0629 | 1.0165 | 5.98E-06 | IIIVmrMLM | SUG |
| SNS_18AY | AX-110408539 | 3B | 242704361 | 0.3728 | | 2.08E-06 | BLUPmrMLM | SUG |
| SNS_18AY | AX-110408539 | 3B | 242704361 | -0.4607 | 0.0136 | 9.04E-25 | IIIVmrMLM | SIG |
| SNS_18AY | AX-110976026 | 3B | 721555659 | -0.1686 | 2.2658 | 9.25E-09 | IIIVmrMLM | SIG |
| SNS_18AY | AX-94834894 | 3D | 563860653 | -0.2787 | | 3.65E-13 | IIIVmrMLM | SIG |
| SNS_18AY | AX-110518599 | 4A | 4838274 | 0.2889 | 0.5291 | 4.22E-13 | IIIVmrMLM | SIG |
| SNS_18AY | AX-94564593 | 4A | 45109525 | 0.0579 | -0.3098 | 4.61E-08 | IIIVmrMLM | SIG |
| SNS_18AY | AX-109416710 | 4A | 579541337 | -0.3855 | 2.096 | 2.21E-26 | IIIVmrMLM | SIG |
| SNS_18AY | AX-108947790 | 4A | 624658137 | -0.2817 | | 3.38E-05 | BLUPmrMLM | SUG |
| SNS_18AY | AX-110047769 | 4A | 628336734 | 0.2863 | | 8.53E-05 | BLUPmrMLM | SUG |
| SNS_18AY | AX-108762439 | 4A | 731411254 | -0.3714 | | 2.91E-07 | BLUPmrMLM | SUG |
| SNS_18AY | AX-110392557 | 4B | 29257749 | 0.0601 | 1.3084 | 4.01E-05 | IIIVmrMLM | SUG |
| SNS_18AY | AX-110582885 | 5A | 13345117 | -0.1532 | | 8.51E-05 | BLUPmrMLM | SUG |
| SNS_18AY | AX-108936505 | 5A | 30025037 | 0.3366 | | 1.17E-07 | BLUPmrMLM | SUG |
| SNS_18AY | AX-109903760 | 5A | 30056842 | 1.1663 | | 7.04E-08 | EMMAX | SIG |
| SNS_18AY | AX-109903760 | 5A | 30056842 | -1.1562 | | 5.15E-08 | GEMMA | SIG |
| SNS_18AY | AX-111623606 | 5A | 70576582 | 0.0106 | -0.4758 | 3.44E-08 | IIIVmrMLM | SIG |
| SNS_18AY | AX-110441490 | 5A | 476834942 | 0.1910 | 0.8397 | 1.3E-12 | IIIVmrMLM | SIG |
| SNS_18AY | AX-109408359 | 5A | 512124032 | 0.2447 | 0.1153 | 2.59E-09 | IIIVmrMLM | SIG |
| SNS_18AY | AX-110929330 | 5A | 532560676 | -0.3147 | | 1.54E-06 | BLUPmrMLM | SUG |
| SNS_18AY | AX-110929330 | 5A | 532560676 | 0.2783 | -0.1494 | 1.17E-09 | IIIVmrMLM | SIG |
| SNS_18AY | AX-108989100 | 5A | 559820786 | -1.0647 | | 1.99E-07 | FarmCPU | SUG |
| SNS_18AY | AX-111620792 | 5A | 699482546 | -0.0375 | 0.697 | 1.01E-06 | IIIVmrMLM | SUG |
| SNS_18AY | AX-108744609 | 5B | 63269770 | -0.1080 | -1.155 | 2.1E-07 | IIIVmrMLM | SUG |
| SNS_18AY | AX-108937022 | 5B | 81214131 | 0.1786 | 0.5491 | 2.87E-13 | IIIVmrMLM | SIG |

| | | | | | | | | |
|---|---|---|---|---|---|---|---|---|
| SNS_18AY | AX-109018708 | 5B | 405722389 | 0.0705 | 0.9125 | 1.29E-05 | IIIVmrMLM | SUG |
| SNS_18AY | AX-108857615 | 5B | 430360253 | 0.1239 | -0.7204 | 2.64E-10 | IIIVmrMLM | SIG |
| SNS_18AY | AX-94443687 | 5B | 431798839 | -0.3668 | | 4.43E-21 | IIIVmrMLM | SIG |
| SNS_18AY | AX-111087937 | 5B | 709939439 | 0.5319 | | 3.46E-06 | BLUPmrMLM | SUG |
| SNS_18AY | AX-94951915 | 6A | 2178020 | 0.2772 | | 1.47E-06 | BLUPmrMLM | SUG |
| SNS_18AY | AX-109873889 | 6A | 580215729 | -0.2249 | | 1.29E-05 | BLUPmrMLM | SUG |
| SNS_18AY | AX-111613086 | 6A | 607966360 | -0.3413 | | 2.42E-06 | BLUPmrMLM | SUG |
| SNS_18AY | AX-109398347 | 6A | 608976305 | 0.3046 | 0.2773 | 3.72E-14 | IIIVmrMLM | SIG |
| SNS_18AY | AX-111533554 | 6B | 125404774 | -0.2292 | | 1.78E-09 | IIIVmrMLM | SIG |
| SNS_18AY | AX-94815662 | 6B | 127320485 | -0.4258 | | 4.22E-08 | BLUPmrMLM | SIG |
| SNS_18AY | AX-110582998 | 6B | 256656242 | -0.1487 | -2.6287 | 5.7E-14 | IIIVmrMLM | SIG |
| SNS_18AY | AX-110018821 | 6B | 516085553 | -0.3347 | | 5.36E-07 | BLUPmrMLM | SUG |
| SNS_18AY | AX-108848769 | 6D | 16655862 | -4.5744 | -4.0861 | 0 | IIIVmrMLM | SIG |
| SNS_18AY | AX-111123882 | 6D | 272759669 | -1.2300 | | 1.55E-06 | FarmCPU | SUG |
| SNS_18AY | AX-110923614 | 7A | 28352822 | -0.1378 | -3.8503 | 2.92E-07 | IIIVmrMLM | SUG |
| SNS_18AY | AX-111566717 | 7A | 52244453 | -0.2439 | | 1.64E-10 | IIIVmrMLM | SIG |
| SNS_18AY | AX-111590334 | 7A | 512089130 | 0.0263 | 2.7191 | 2.22E-13 | IIIVmrMLM | SIG |
| SNS_18AY | AX-108762572 | 7A | 627149804 | -0.2646 | | 2.15E-05 | BLUPmrMLM | SUG |
| SNS_18AY | AX-110460526 | 7A | 647929754 | -0.3212 | | 6.58E-06 | BLUPmrMLM | SUG |
| SNS_18AY | AX-110976213 | 7A | 675483958 | 0.8458 | | 1.74E-06 | FarmCPU | SUG |
| SNS_18AY | AX-111577052 | 7A | 695896051 | 0.0395 | 0.7647 | 9.06E-06 | IIIVmrMLM | SUG |
| SNS_18AY | AX-109430147 | 7B | 220721692 | 0.0390 | -1.2448 | 1.34E-05 | IIIVmrMLM | SUG |
| SNS_18AY | AX-111174514 | 7B | 506023046 | -0.0126 | -0.4838 | 1.33E-05 | IIIVmrMLM | SUG |
| SNS_18AY | AX-110457590 | 7B | 613193601 | 0.4315 | -0.0151 | 2.75E-26 | IIIVmrMLM | SIG |
| SNS_18AY | AX-111521286 | 7B | 701923851 | -0.0276 | -1.4898 | 4.76E-09 | IIIVmrMLM | SIG |
| SNS_18AY | AX-110373318 | 7B | 702247074 | 0.3545 | | 4.03E-08 | BLUPmrMLM | SIG |
| SNS_18AY | AX-111832758 | 7D | 66483796 | -0.6927 | | 1.52E-06 | GEMMA | SUG |
| SNS_18JY | AX-94596123 | 1B | 548970174 | 0.2264 | | 2.8E-05 | BLUPmrMLM | SUG |
| SNS_18JY | AX-111182155 | 1D | 282279896 | 0.4214 | | 2.21E-06 | BLUPmrMLM | SUG |
| SNS_18JY | AX-111026713 | 2A | 715823420 | 0.2356 | | 5.12E-07 | BLUPmrMLM | SUG |
| SNS_18JY | AX-110692318 | 2A | 744754960 | -0.2841 | | 3.1E-06 | BLUPmrMLM | SUG |
| SNS_18JY | AX-110499313 | 2B | 67685795 | -0.1524 | | 6.89E-06 | BLUPmrMLM | SUG |
| SNS_18JY | AX-109891725 | 2D | 616994681 | -0.2113 | | 8.7E-06 | BLUPmrMLM | SUG |
| SNS_18JY | AX-94389846 | 3A | 700422181 | -0.1207 | | 0.000103 | BLUPmrMLM | SUG |
| SNS_18JY | AX-109496510 | 3A | 723348441 | -0.2529 | | 1.39E-06 | BLUPmrMLM | SUG |
| SNS_18JY | AX-111066590 | 3A | 741983597 | -0.2035 | | 0.000168 | BLUPmrMLM | SUG |
| SNS_18JY | AX-109896354 | 3B | 168122143 | -1.0475 | | 1.07E-06 | FarmCPU | SUG |
| SNS_18JY | AX-110366998 | 3B | 364345388 | 0.9185 | | 6.15E-07 | FarmCPU | SUG |
| SNS_18JY | AX-110463606 | 3B | 656916252 | 0.2337 | | 7.49E-06 | BLUPmrMLM | SUG |
| SNS_18JY | AX-109468100 | 3B | 744516874 | 0.1792 | | 4.57E-05 | BLUPmrMLM | SUG |
| SNS_18JY | AX-111548257 | 3D | 12733555 | 0.2116 | | 2.46E-05 | BLUPmrMLM | SUG |
| SNS_18JY | AX-111101026 | 4A | 78942874 | -0.4027 | | 1.55E-06 | BLUPmrMLM | SUG |
| SNS_18JY | AX-110455433 | 4B | 660337364 | 0.4625 | | 9.08E-07 | FarmCPU | SUG |
| SNS_18JY | AX-108968377 | 4D | 506823199 | -0.1740 | | 9.34E-06 | BLUPmrMLM | SUG |

| | | | | | | | | |
|---|---|---|---|---|---|---|---|---|
| SNS_18JY | AX-109934166 | 5B | 60876473 | -0.1599 | | 0.000201 | BLUPmrMLM | SUG |
| SNS_18JY | AX-111590173 | 5B | 421191385 | 0.3861 | | 2.22E-06 | GEMMA | SUG |
| SNS_18JY | AX-109582173 | 5B | 421192328 | 0.3948 | | 1.41E-06 | GEMMA | SUG |
| SNS_18JY | AX-111055741 | 5B | 421274258 | 0.3948 | | 1.41E-06 | GEMMA | SUG |
| SNS_18JY | AX-95194941 | 5B | 421275897 | -0.7591 | | 2.06E-06 | GEMMA | SUG |
| SNS_18JY | AX-109036790 | 5B | 421276751 | 0.3866 | | 1.71E-06 | GEMMA | SUG |
| SNS_18JY | AX-110020458 | 5B | 592862695 | 0.1967 | | 3.1E-05 | BLUPmrMLM | SUG |
| SNS_18JY | AX-111489741 | 5B | 642047731 | 0.1643 | | 6.12E-05 | BLUPmrMLM | SUG |
| SNS_18JY | AX-108840042 | 5D | 42961528 | -0.1476 | | 2.22E-05 | BLUPmrMLM | SUG |
| SNS_18JY | AX-109376104 | 6B | 696027028 | -0.1283 | | 8.88E-05 | BLUPmrMLM | SUG |
| SNS_18JY | AX-109281272 | 7A | 6508743 | 0.2339 | | 6.85E-05 | BLUPmrMLM | SUG |
| SNS_18JY | AX-111652976 | 7B | 329514338 | -0.1996 | | 1.05E-06 | BLUPmrMLM | SUG |
| SNS_18JY | AX-111475158 | 7B | 680235293 | 0.1441 | | 0.000124 | BLUPmrMLM | SUG |
| SNS_18XX | AX-110944959 | 1A | 339895079 | 0.0323 | 0.5749 | 2.08E-07 | IIIVmrMLM | SUG |
| SNS_18XX | AX-111662360 | 1A | 489402037 | 0.7887 | | 2.25E-06 | GEMMA | SUG |
| SNS_18XX | AX-110648456 | 1B | 389226819 | 0.1903 | 1.3804 | 2.39E-24 | IIIVmrMLM | SIG |
| SNS_18XX | AX-110707054 | 1B | 673916692 | 0.2148 | | 4.48E-05 | BLUPmrMLM | SUG |
| SNS_18XX | AX-94652478 | 1D | 3958384 | 0.3354 | | 1.74E-06 | BLUPmrMLM | SUG |
| SNS_18XX | AX-109349477 | 1D | 15363542 | 0.1920 | | 1.43E-06 | BLUPmrMLM | SUG |
| SNS_18XX | AX-94571943 | 1D | 492987095 | -0.0434 | 0.7737 | 5.82E-08 | IIIVmrMLM | SIG |
| SNS_18XX | AX-94918359 | 2A | 66809150 | -0.2257 | 0.0352 | 1.44E-17 | IIIVmrMLM | SIG |
| SNS_18XX | AX-109479111 | 2A | 476792209 | 0.1277 | 0.3729 | 9.21E-10 | IIIVmrMLM | SIG |
| SNS_18XX | AX-111073917 | 2A | 715899397 | 0.1511 | | 2.97E-05 | BLUPmrMLM | SUG |
| SNS_18XX | AX-109451281 | 2B | 104594417 | -0.3278 | -0.1184 | 1.34E-36 | IIIVmrMLM | SIG |
| SNS_18XX | AX-110634537 | 2B | 122385222 | 0.4462 | 0.8851 | 1.31E-82 | IIIVmrMLM | SIG |
| SNS_18XX | AX-110083263 | 2B | 529098177 | -0.3769 | -0.0849 | 5.11E-47 | IIIVmrMLM | SIG |
| SNS_18XX | AX-111589395 | 2B | 742819261 | -0.0165 | 0.9102 | 1.45E-07 | IIIVmrMLM | SUG |
| SNS_18XX | AX-110970217 | 2D | 470906707 | -0.3375 | -1.5073 | 1.13E-49 | IIIVmrMLM | SIG |
| SNS_18XX | AX-111480094 | 3A | 485684195 | 0.3064 | | 5.52E-37 | IIIVmrMLM | SIG |
| SNS_18XX | AX-111673190 | 3A | 746146319 | 0.1990 | 0.0119 | 5.7E-10 | IIIVmrMLM | SIG |
| SNS_18XX | AX-111612104 | 3B | 116188801 | -0.1839 | | 4.03E-06 | BLUPmrMLM | SUG |
| SNS_18XX | AX-110068969 | 3B | 658061367 | -0.0279 | -0.37 | 1.79E-13 | IIIVmrMLM | SIG |
| SNS_18XX | AX-110073738 | 3B | 668888194 | -0.0041 | 0.6573 | 1.35E-08 | IIIVmrMLM | SIG |
| SNS_18XX | AX-109978455 | 3B | 749637598 | 0.2491 | 0.0402 | 2.66E-14 | IIIVmrMLM | SIG |
| SNS_18XX | AX-110471093 | 3B | 782032804 | -0.0106 | -0.7836 | 5.73E-05 | IIIVmrMLM | SUG |
| SNS_18XX | AX-110405928 | 4A | 117331175 | 0.0017 | -0.7111 | 6.64E-08 | IIIVmrMLM | SIG |
| SNS_18XX | AX-95202867 | 4A | 277889699 | -0.1891 | | 5.07E-06 | BLUPmrMLM | SUG |
| SNS_18XX | AX-111717386 | 4A | 606517595 | -0.0638 | -0.3081 | 4.19E-07 | IIIVmrMLM | SUG |
| SNS_18XX | AX-109893421 | 4A | 688839204 | 0.0498 | 0.5493 | 7.74E-07 | IIIVmrMLM | SUG |
| SNS_18XX | AX-111510412 | 4B | 158546987 | -0.1436 | -0.449 | 1.62E-12 | IIIVmrMLM | SIG |
| SNS_18XX | AX-94802502 | 4B | 371977654 | 0.4884 | 0.0762 | 6.86E-26 | IIIVmrMLM | SIG |
| SNS_18XX | AX-94590603 | 5A | 384635052 | 0.1418 | | 2.75E-05 | BLUPmrMLM | SUG |
| SNS_18XX | AX-111594426 | 5A | 512054146 | -0.1816 | | 3.24E-06 | BLUPmrMLM | SUG |
| SNS_18XX | AX-94414339 | 5A | 556006342 | -0.2840 | -0.1955 | 1.47E-30 | IIIVmrMLM | SIG |

| | | | | | | | | |
|---|---|---|---|---|---|---|---|---|
| SNS_18XX | AX-111578637 | 5A | 563767692 | -0.0113 | 0.3082 | 8.06E-06 | IIIVmrMLM | SUG |
| SNS_18XX | AX-110606073 | 5A | 701962724 | -0.1564 | | 2.92E-05 | BLUPmrMLM | SUG |
| SNS_18XX | AX-111656147 | 5B | 84476546 | 0.0199 | -1.172 | 9.81E-13 | IIIVmrMLM | SIG |
| SNS_18XX | AX-110610000 | 5B | 711310110 | -0.1860 | | 8.38E-05 | BLUPmrMLM | SUG |
| SNS_18XX | AX-110584740 | 5B | 712411959 | 0.2678 | 0.0525 | 3.86E-24 | IIIVmrMLM | SIG |
| SNS_18XX | AX-111556923 | 6A | 9326702 | -0.0173 | -0.3824 | 0.000237 | IIIVmrMLM | SUG |
| SNS_18XX | AX-111632133 | 6A | 33556088 | 0.1766 | | 5.99E-05 | BLUPmrMLM | SUG |
| SNS_18XX | AX-109952070 | 6A | 593423250 | -0.2332 | | 6.44E-08 | BLUPmrMLM | SIG |
| SNS_18XX | AX-109352913 | 6B | 202133169 | 0.0080 | 0.4482 | 9.57E-13 | IIIVmrMLM | SIG |
| SNS_18XX | AX-109270492 | 6B | 679282033 | -0.1116 | | 0.000115 | BLUPmrMLM | SUG |
| SNS_18XX | AX-94426347 | 6B | 707110794 | -0.0908 | | 0.000132 | BLUPmrMLM | SUG |
| SNS_18XX | AX-110633938 | 6D | 470602132 | -0.0127 | -0.8029 | 1.3E-08 | IIIVmrMLM | SIG |
| SNS_18XX | AX-108885503 | 7A | 98026967 | -0.2528 | | 4.67E-07 | BLUPmrMLM | SUG |
| SNS_18XX | AX-108885503 | 7A | 98026967 | 0.3509 | | 9.37E-07 | EMMAX | SUG |
| SNS_18XX | AX-108885503 | 7A | 98026967 | -0.3535 | | 4.39E-07 | GEMMA | SUG |
| SNS_18XX | AX-108885503 | 7A | 98026967 | 0.1690 | -0.6025 | 8.07E-13 | IIIVmrMLM | SIG |
| SNS_18XX | AX-111808701 | 7A | 116003403 | -0.1442 | | 3.8E-05 | BLUPmrMLM | SUG |
| SNS_18XX | AX-111808701 | 7A | 116003403 | 0.1843 | -0.0409 | 1.01E-09 | IIIVmrMLM | SIG |
| SNS_18XX | AX-110077338 | 7A | 254388927 | -0.0567 | -0.4526 | 6.91E-14 | IIIVmrMLM | SIG |
| SNS_18XX | AX-109491276 | 7A | 706280025 | -0.0738 | -0.4135 | 6.5E-08 | IIIVmrMLM | SIG |
| SNS_18XX | AX-108897063 | 7A | 718203771 | 0.0197 | -2.0815 | 2.13E-12 | IIIVmrMLM | SIG |
| SNS_18XX | AX-111471715 | 7B | 129525956 | -0.1324 | | 6.32E-05 | BLUPmrMLM | SUG |
| SNS_18XX | AX-111450131 | 7B | 134017227 | -0.1449 | 0.3641 | 5.93E-17 | IIIVmrMLM | SIG |
| SNS_18XX | AX-94770214 | 7B | 498523685 | -0.1655 | | 6.72E-13 | IIIVmrMLM | SIG |
| SNS_18XX | AX-111155391 | 7B | 687856955 | 0.0189 | 0.9965 | 2.82E-06 | IIIVmrMLM | SUG |
| SNS_18XX | AX-111551257 | 7B | 732422423 | -0.2587 | | 9.26E-06 | BLUPmrMLM | SUG |
| SNS_18XX | AX-108883474 | 7D | 615934069 | -0.1614 | | 3.86E-06 | BLUPmrMLM | SUG |
| SNS_BLUP | AX-111524523 | 1A | 22938066 | -0.0516 | | 1.57E-05 | BLUPmrMLM | SUG |
| SNS_BLUP | AX-111607717 | 1A | 36844582 | 0.0439 | 0.0023 | 2.47E-23 | IIIVmrMLM | SIG |
| SNS_BLUP | AX-110982739 | 1A | 38723482 | -0.0774 | | 5.02E-06 | BLUPmrMLM | SUG |
| SNS_BLUP | AX-111650973 | 1A | 512491608 | -0.0398 | | 3.42E-23 | IIIVmrMLM | SIG |
| SNS_BLUP | AX-110960919 | 1A | 564128400 | -0.0116 | -0.168 | 2.3E-10 | IIIVmrMLM | SIG |
| SNS_BLUP | AX-110541070 | 1B | 640921845 | -0.0011 | -0.0343 | 7.52E-07 | IIIVmrMLM | SUG |
| SNS_BLUP | AX-110557135 | 2A | 31942667 | -0.0066 | -0.0797 | 2.4E-07 | IIIVmrMLM | SUG |
| SNS_BLUP | AX-108916615 | 2A | 35954359 | 0.0350 | -0.0034 | 1.7E-14 | IIIVmrMLM | SIG |
| SNS_BLUP | AX-109469379 | 2A | 539402856 | 0.0777 | | 2.52E-05 | BLUPmrMLM | SUG |
| SNS_BLUP | AX-109971726 | 2B | 439530 | -0.0583 | | 6.51E-07 | BLUPmrMLM | SUG |
| SNS_BLUP | AX-111149302 | 2B | 26148281 | -0.0123 | 0.1369 | 3.56E-16 | IIIVmrMLM | SIG |
| SNS_BLUP | AX-110945395 | 2B | 68582225 | 0.0710 | | 1.62E-07 | BLUPmrMLM | SUG |
| SNS_BLUP | AX-111218699 | 2B | 410921125 | 0.0193 | -0.0818 | 8.34E-13 | IIIVmrMLM | SIG |
| SNS_BLUP | AX-109518784 | 2B | 659086334 | 0.0089 | 0.1089 | 1.17E-08 | IIIVmrMLM | SIG |
| SNS_BLUP | AX-108774705 | 2B | 667247976 | -0.0520 | | 1.33E-05 | BLUPmrMLM | SUG |
| SNS_BLUP | AX-111513301 | 2D | 15756024 | -0.0008 | -0.3539 | 1.46E-06 | IIIVmrMLM | SUG |
| SNS_BLUP | AX-109217069 | 2D | 170808466 | 0.0297 | 0.3433 | 5.31E-18 | IIIVmrMLM | SIG |

| | | | | | | | | |
|---|---|---|---|---|---|---|---|---|
| SNS_BLUP | AX-111149345 | 3A | 12157923 | -0.0040 | 0.1289 | 0.000183 | IIIVmrMLM | SUG |
| SNS_BLUP | AX-109394789 | 3A | 695517857 | 0.0077 | -0.0707 | 3.92E-12 | IIIVmrMLM | SIG |
| SNS_BLUP | AX-111490915 | 3A | 720420925 | 0.0012 | -0.1138 | 3.58E-06 | IIIVmrMLM | SUG |
| SNS_BLUP | AX-108973844 | 3A | 725194673 | 0.0488 | | 5.69E-06 | BLUPmrMLM | SUG |
| SNS_BLUP | AX-110103877 | 3B | 70427006 | 0.0053 | 0.0992 | 3.19E-11 | IIIVmrMLM | SIG |
| SNS_BLUP | AX-108918111 | 3B | 265713103 | 0.0306 | -0.0799 | 5.28E-37 | IIIVmrMLM | SIG |
| SNS_BLUP | AX-110930630 | 3B | 411339587 | 0.0038 | 0.1942 | 1.36E-09 | IIIVmrMLM | SIG |
| SNS_BLUP | AX-111602251 | 3B | 565277238 | 0.0143 | 0.0732 | 3.69E-12 | IIIVmrMLM | SIG |
| SNS_BLUP | AX-109036640 | 3B | 749671030 | 0.0330 | 0.0087 | 7.59E-13 | IIIVmrMLM | SIG |
| SNS_BLUP | AX-108903841 | 3B | 819941849 | 0.0422 | | 0.000139 | BLUPmrMLM | SUG |
| SNS_BLUP | AX-109638284 | 3D | 220068635 | 0.0328 | | 1.18E-16 | IIIVmrMLM | SIG |
| SNS_BLUP | AX-111589696 | 4A | 684908443 | 0.0570 | | 1.45E-05 | BLUPmrMLM | SUG |
| SNS_BLUP | AX-110973979 | 4A | 729900597 | -0.0528 | | 8.89E-07 | BLUPmrMLM | SUG |
| SNS_BLUP | AX-110372696 | 4B | 660656121 | 0.0204 | -0.0372 | 6.68E-07 | IIIVmrMLM | SUG |
| SNS_BLUP | AX-108855539 | 5A | 529846821 | -0.0354 | -0.0191 | 1.45E-11 | IIIVmrMLM | SIG |
| SNS_BLUP | AX-109436301 | 5B | 65965632 | -0.0403 | | 0.000185 | BLUPmrMLM | SUG |
| SNS_BLUP | AX-110168965 | 5B | 434057190 | 0.0391 | -0.0274 | 2.52E-21 | IIIVmrMLM | SIG |
| SNS_BLUP | AX-89569962 | 5B | 572955946 | -0.0137 | -0.0634 | 5.55E-07 | IIIVmrMLM | SUG |
| SNS_BLUP | AX-110424395 | 5B | 646278664 | -0.0033 | -0.2096 | 1.89E-18 | IIIVmrMLM | SIG |
| SNS_BLUP | AX-95227398 | 5D | 398782794 | 0.0593 | | 1.16E-06 | BLUPmrMLM | SUG |
| SNS_BLUP | AX-111758704 | 5D | 541366083 | 0.0563 | | 2.36E-06 | BLUPmrMLM | SUG |
| SNS_BLUP | AX-111758704 | 5D | 541366083 | -0.0197 | -0.016 | 3.17E-06 | IIIVmrMLM | SUG |
| SNS_BLUP | AX-94966145 | 6A | 5221726 | -0.0057 | 0.2047 | 7.76E-10 | IIIVmrMLM | SIG |
| SNS_BLUP | AX-110593028 | 6A | 564974058 | 0.0375 | | 8.52E-05 | BLUPmrMLM | SUG |
| SNS_BLUP | AX-89322586 | 6B | 42680851 | -0.0044 | 0.0367 | 1.19E-06 | IIIVmrMLM | SUG |
| SNS_BLUP | AX-111500287 | 6B | 136503264 | 0.0463 | | 0.000085 | BLUPmrMLM | SUG |
| SNS_BLUP | AX-94502175 | 6B | 660380975 | -0.0689 | | 0.000172 | BLUPmrMLM | SUG |
| SNS_BLUP | AX-94502175 | 6B | 660380975 | 0.0191 | | 9.36E-07 | IIIVmrMLM | SUG |
| SNS_BLUP | AX-109483048 | 7A | 13780580 | -0.0219 | -0.179 | 1.57E-14 | IIIVmrMLM | SIG |
| SNS_BLUP | AX-111055096 | 7A | 128117104 | -0.0011 | 0.0465 | 0.000039 | IIIVmrMLM | SUG |
| SNS_BLUP | AX-111174514 | 7B | 506023046 | 0.0003 | -0.0683 | 1.01E-09 | IIIVmrMLM | SIG |
| SNS_BLUP | AX-110373318 | 7B | 702247074 | 0.0583 | | 2.54E-07 | BLUPmrMLM | SUG |
| SNS_BLUP | AX-110373318 | 7B | 702247074 | -0.0222 | 0.0061 | 3.87E-06 | IIIVmrMLM | SUG |
| SNS_BLUP | AX-95215050 | 7B | 706857327 | -0.0038 | -0.0429 | 0.000105 | IIIVmrMLM | SUG |
| SNS_BLUP | AX-111061288 | 7D | 66537922 | -0.0320 | 0.0161 | 5.89E-14 | IIIVmrMLM | SIG |
| SNS_BLUP | AX-108928265 | 7D | 118292316 | -0.0633 | | 9.38E-08 | BLUPmrMLM | SIG |
| SNS_BLUP | AX-108787886 | 7D | 552272241 | 0.0035 | 0.1836 | 1.98E-09 | IIIVmrMLM | SIG |

SNS: spikelet number per spike.

**Supplementary Table S14**. All the known genes for five yield-related traits in genome-wide association studies and their pieces of evidence

| Trait | Genome-wide association studies | | | | | | | Gene | Symbol | Reference |
|---|---|---|---|---|---|---|---|---|---|---|
| | Environment | Marker | Chr | Position | Add/Dom | P-value | Method | | | |
| GL | GL_17XX | AX-94786159 | chr1D | 79116398 | 0.4274/-- | 1.06E-08 | GEMMA | TraesCS1D01G094000 | *TaPGS1* | Guo et al., 2022 |
| GL | GL_18JY | AX-94786159 | chr1D | 79116398 | 0.3231/-- | 3.14E-07 | GEMMA | TraesCS1D01G094000 | *TaPGS1* | Guo et al., 2022 |
| GL | GL_BLUP | AX-94786159 | chr1D | 79116398 | 0.2512/-- | 1.40E-07 | GEMMA | TraesCS1D01G094000 | *TaPGS1* | Guo et al., 2022 |
| GL | GL_18XX | AX-94786159 | chr1D | 79116398 | -0.3294/-- | 7.23E-07 | EMMAX | TraesCS1D01G094000 | *TaPGS1* | Guo et al., 2022 |
| GL | GL_17XX | AX-94786159 | chr1D | 79116398 | -0.4331/-- | 1.17E-08 | EMMAX | TraesCS1D01G094000 | *TaPGS1* | Guo et al., 2022 |
| GL | GL_BLUP | AX-94786159 | chr1D | 79116398 | -0.2525/-- | 8.91E-08 | EMMAX | TraesCS1D01G094000 | *TaPGS1* | Guo et al., 2022 |
| GL | GL_18XX | AX-94786159 | chr1D | 79116398 | 0.3289/-- | 9.00E-07 | GEMMA | TraesCS1D01G094000 | *TaPGS1* | Guo et al., 2022 |
| GL | GL_19XX | AX-94804217 | chr2D | 8478708 | -0.0447/-- | 9.73E-05 | BLUPmrMLM | TraesCS2D01G016900 | *TaDA1* | Liu et al., 2020 |
| GL | GL_17XX | AX-94804217 | chr2D | 8478708 | -0.0594/-- | 5.11E-05 | BLUPmrMLM | TraesCS2D01G016900 | *TaDA1* | Liu et al., 2020 |
| GL | GL_18XX | AX-110050904 | chr5A | 34327685 | 0.0408/-- | 0.000128 | BLUPmrMLM | TraesCS5A01G038300 | *TaARF25-5A* | Jia et al., 2021 |
| GL | GL_17XX | AX-111470748 | chr5B | 555215171 | -0.1349/-- | 9.17E-12 | BLUPmrMLM | TraesCS5B01G375800 | *TaGL3-5B* | Yang et al.,2019 |
| GWLR | GWLR_17AY | AX-94804217 | chr2D | 8478708 | -0.06 | 2.98E-08 | IIIVmrMLM | TraesCS2D01G016900 | *TaDA1* | Liu et al., 2020 |
| GWLR | GWLR_BLUP | AX-111560255 | chr2D | 9080514 | -0.133333333 | 1.52E-23 | IIIVmrMLM | TraesCS2D01G016900 | *TaDA1* | Liu et al., 2020 |
| GWLR | GWLR_BLUP | AX-111526867 | chr3A | 550655603 | 0.0044/-- | 0.000101 | BLUPmrMLM | TraesCS3A01G311100 | *TaCKX6a02-3A* | Lu et al.,2015 |
| GWLR | GWLR_18JY | AX-108945354 | chr3A | 574283260 | 0.0059/-0.0016 | 2.29E-15 | IIIVmrMLM | TraesCS3A01G328000 | *TaERF3-3A* | Jia et al., 2021 |
| GWLR | GWLR_BLUP | AX-109919341 | chr3A | 574323687 | -0.0038/-- | 4.44E-07 | BLUPmrMLM | TraesCS3A01G328000 | *TaERF3-3A* | Jia et al., 2021 |
| GWLR | GWLR_19XX | AX-110939022 | chr3B | 567246558 | 0.0006/-0.0253 | 1.17E-08 | IIIVmrMLM | TraesCS3B01G357500 | *TaERF3-3B* | Jia et al., 2021 |
| GWLR | GWLR_BLUP | AX-110036417 | chr5A | 35022242 | -0.0023/-- | 0.00017 | BLUPmrMLM | TraesCS5A01G038300 | *TaARF25-5A* | Jia et al., 2021 |
| GWLR | GWLR_18JY | AX-109881727 | chr5B | 552787014 | -0.0098/-- | 1.97E-07 | BLUPmrMLM | TraesCS5B01G375800 | *TaGL3-5B* | Yang et al., 2019 |
| GWLR | GWLR_17XX | AX-111470748 | chr5B | 555215171 | 0.0085/-- | 1.41E-06 | BLUPmrMLM | TraesCS5B01G375800 | *TaGL3-5B* | Yang et al., 2019 |
| GWLR | GWLR_17SQ | AX-94689724 | chr6B | 575638302 | -0.0044/-- | 9.67E-05 | BLUPmrMLM | TraesCS6B01G327500 | *TaGS1a-6B* | Guo et al., 2013 |

| | | | | | | | | | | |
|---|---|---|---|---|---|---|---|---|---|---|
| GW | GW_19XX | AX-108745355 | chr3A | 550292294 | 0.417256011 | 1.38E-23 | IIIVmrMLM | TraesCS3A01G311100 | *TaCKX6a02-3A* | Lu et al., 2015 |
| GW | GW_BLUP | AX-94566074 | chr3D | 94239601 | 0.0016/0.0549 | 8.90E-05 | IIIVmrMLM | TraesCS3D01G137200 | *Tasg-D1/S1* | Gupta et al., 2020 |
| GW | GW_18XX | AX-109276523 | chr5A | 36328550 | 0.35770235 | 7.93E-20 | IIIVmrMLM | TraesCS5A01G038300 | *TaARF25-5A* | Jia et al., 2021 |
| TGW | TGW_18JY | AX-109330937 | chr2A | 755378747 | 0.131531532 | 5.98E-08 | IIIVmrMLM | TraesCS2A01G547800 | *TaARF12-2A* | Li et al., 2022 |
| TGW | TGW_19XX | AX-94488783 | chr2A | 755788522 | 0.8834/-- | 6.64E-05 | BLUPmrMLM | TraesCS2A01G547800 | *TaARF12-2A* | Li et al., 2022 |
| TGW | TGW_18JY | AX-111489027 | chr2A | 758274417 | -0.6165/-- | 3.33E-06 | BLUPmrMLM | TraesCS2A01G555600 | *TaSTT3b-2A* | Zhu et al., 2022 |
| TGW | TGW_17JY | AX-109022397 | chr2A | 758561818 | -0.014620877 | 7.49E-11 | IIIVmrMLM | TraesCS2A01G555600 | *TaSTT3b-2A* | Zhu et al., 2022 |
| TGW | TGW_18XX | AX-109344204 | chr2A | 759309971 | 0.0573/1.1996 | 1.35E-10 | IIIVmrMLM | TraesCS2A01G555600 | *TaSTT3b-2A* | Zhu et al., 2022 |
| TGW | TGW_18XX | AX-109286668 | chr2B | 181773892 | 0.3641/-0.5819 | 6.34E-09 | IIIVmrMLM | TraesCS2B01G201900 | *TaCYP78A5-2B* | Guo et al., 2021 |
| TGW | TGW_18XX | AX-109286668 | chr2B | 181773892 | 0.3641/-0.5819 | 6.34E-09 | IIIVmrMLM | TraesCS2B01G202300 | *TaGW7* | Wang et al., 2019 |
| TGW | TGW_17ZMD | AX-109607606 | chr3A | 551506355 | 0.5207/-5.0637 | 4.27E-12 | IIIVmrMLM | TraesCS3A01G311100 | *TaCKX6a02-3A* | Lu et al., 2015 |
| TGW | TGW_18XX | AX-94762802 | chr5A | 37620417 | 0.75/-- | 8.39E-05 | BLUPmrMLM | TraesCS5A01G038300 | *TaARF25-5A* | Jia et al., 2021 |
| TGW | TGW_18JY | AX-109370945 | chr6B | 456774277 | 1.0622/-- | 1.64E-07 | BLUPmrMLM | TraesCS6B01G253900 | *TaPRR1-B1* | Sun et al., 2020 |
| TGW | TGW_17JY | AX-111695521 | chr6B | 595295134 | -30.281/-- | 1.20E-07 | EMMAX | TraesCS6B01G337300 | *TaPIN1-6B1* | Yao et al., 2021 |
| SNS | SNS_18AY | AX-110018821 | chr6B | 516085553 | -0.33 /-- | 5.36E-07 | BLUPmrMLM | TraesCS6B01G286400 | *TaAGL6-B* | Kong et al.,2022 |
| SNS | SNS_17AY | AX-94680052 | chr7A | 675594466 | -0.28 /-- | 5.87E-09 | IIIVmrMLM | TraesCS7A01G481600 | *WAPO1* | Kuzay et al.,2019 |
| SNS | SNS_18AY | AX-110976213 | chr7A | 675483958 | 0.85 /-- | 1.74E-06 | FarmCPU | TraesCS7A01G481600 | *WAPO1* | Kuzay et al.,2019 |

SNS: spikelet number per spike, TGW: thousand grain weight, GL: grain length, GW: grain width, GWLR: ratio of grain width to length.

**Supplementary Table S15**. Candidate genes for five yield-related traits in genome-wide association studies and their pieces of evidence

| Trait | Genome-wide association studies | | | | | | | Differential expression analysis | | | Homologous genes/Genes reported in Rice | GO annotation analysis | | P-value in haplotype analysis | Reference |
|---|---|---|---|---|---|---|---|---|---|---|---|---|---|---|---|
| | Environment | Marker | Chr | Position | Add/Dom | P-value | Method | Gene | $Log_2$\|FC\| | P. adj | | ID | Annotation | | |
| GL | GL_18AY | AX-108732926 | 6A | 600953819 | 0.0468/-- | 7.84E-05 | BLUPmrMLM | TraesCS6A01G382400 | -13.21 [G] | 8.87e-19 [G] | | GO:0015770 | sucrose transport | 1.80E-30 | |
| GL | GL_18AY | AX-108732926 | 6A | 600953819 | 0.0468/-- | 7.84E-05 | BLUPmrMLM | TraesCS6A01G382500 | -4.01 [G] | 2.94e-12 [G] | | GO:0015770 | sucrose transport | 9.84E-31 | |
| GL | GL_18AY | AX-108732926 | 6A | 600953819 | 0.0468/-- | 7.84E-05 | BLUPmrMLM | TraesCS6A01G382600 | -10.09 [G] | 1.35e-87 [G] | | GO:0015770 | sucrose transport | 3.84E-32 | |
| GL | GL_18XX | AX-111572602 | 6A | 2293409 | 0.0418/-- | 8.47E-05 | BLUPmrMLM | TraesCS6A01G009100 | 8.26 [G] | 3.69e-07 [G] | | GO:0015770 | sucrose transport | 1.70E-08 | |
| GL | GL_17AY | AX-110413677 | 6A | 2058555 | 0.0008/0.1784 | 2.53E-05 | IIIVmrMLM | TraesCS6A01G009100 | 8.26 [G] | 3.69e-07 [G] | | GO:0015770 | sucrose transport | 4.21E-07 | |
| GL | GL_17XX | AX-110062884 | 7D | 20623856 | -0.5175/-- | 1.78E-06 | EMMAX | TraesCS7D01G039500 | -1.41 [G] | 5.79e-06 [G] | | GO:0005982 | starch metabolic process | 1.40E-09 | |
| GL | GL_BLUP | AX-110062884 | 7D | 20623856 | -0.3188/-- | 1.84E-06 | EMMAX | TraesCS7D01G039500 | -1.41 [G] | 5.79e-06 [G] | | GO:0005982 | starch metabolic process | 1.03E-13 | |
| GL | GL_18JY | AX-94558702 | 4B | 649603400 | -2.0437/-- | 3.13E-10 | EMMAX | TraesCS4B01G360800 | -2.45 [G] | 1.39e-43 [G] | | GO:0005982 | starch metabolic process | 1.14E-21 | |
| GL | GL_18AY | AX-110477036 | 4A | 711398837 | 0.0019/0.3082 | 3.46E-05 | IIIVmrMLM | TraesCS4A01G444300 | -1.32 [G] | 7.51e-07 [G] | | GO:0005982 | starch metabolic process | 1.56E-10 | |
| GL | GL_18JY | AX-110504088 | 1A | 468686364 | -0.0883/0.0221 | 1.64E-27 | IIIVmrMLM | TraesCS1A01G273900 | -2.6 [G] | 9.91e-06 [G] | | GO:0005982 | starch metabolic process | 1.50E-12 | |
| GL | GL_18JY | AX-111635682 | 7B | 14763864 | -0.045/-- | 3.62E-09 | IIIVmrMLM | TraesCS7B01G018600 | 1.6 [G] | 1.21e-07 [G] | | GO:0005982 | starch metabolic process | 1.50E-12 | |
| GL | GL_17XX | AX-111903053 | 4D | 484511212 | -0.0438/-- | 8.24E-05 | BLUPmrMLM | TraesCS4D01G318400 | 7.37 [G] | 4.88e-106 [G] | *OsCDPK1* | | | 1.84E-11 | Ho et al., 2013 |
| GL | GL_17XX | AX-111903053 | 4D | 484511212 | -0.0438/-- | 8.24E-05 | BLUPmrMLM | TraesCS4D01G325200 | -1.59 [G] | 6.12e-05 [G] | *OsBSK1-1* | | | 6.38E-10 | Tian et al., 2022 |
| GL | GL_18XX | AX-94653997 | 4D | 485522383 | -0.1511/-- | 6.51E-07 | FarmCPU | TraesCS4D01G325200 | -1.59 [G] | 6.12e-05 [G] | *OsBSK1-1* | | | 6.06E-14 | Tian et al., 2022 |
| GL | GL_18AY | AX-109987438 | 2A | 4295985 | 0.0681/-- | 2.49E-07 | BLUPmrMLM | TraesCS2A01G010600 | 9.23 [G] | 3.45e-12 [G] | *OsMED15a* | | | 1.47E-04 | Dwivedi et al., 2019 |
| GL | GL_18XX | AX-111482807 | 5A | 576373436 | 0.0638/-- | 1.21E-07 | BLUPmrMLM | TraesCS5A01G377700 | 6.46 [G] | 1.52e-04 [G] | *RGG1* | | | 2.62E-26 | Tao et al., 2020 |
| GL | GL_17XX | AX-108735017 | 2D | 5761352 | 1.2917/-- | 1.17E-07 | EMMAX | TraesCS2D01G011200 | 12.06 [G] | 5.52e-21 [G] | *OsMED15a* | | | 5.70E-08 | Dwivedi et al., 2019 |
| GL | GL_18AY | AX-111135595 | 4B | 35175628 | 0.1359/-- | 2.15E-11 | FarmCPU | TraesCS4B01G047400 | 10.0 [G] | 1.17e-14 [G] | *OsGS1* | | | 2.69E-05 | Liu et al., 2022 |
| GL | GL_18JY | AX-94697254 | 2D | 33710135 | 0.2393/-- | 2.00E-06 | FarmCPU | TraesCS2D01G082000 | -2.45 [G] | 3.38e-23 [G] | *OsPLC1* | | | 7.87E-03 | Wu et al., 2022 |
| GL | GL_17AY | AX-111097278 | 2A | 717021012 | 0.1438/-- | 1.81E-06 | GEMMA | TraesCS2A01G482400 | 3.91 [G] | 1.26e-07 [G] | *OsMED15a* | | | 3.35E-24 | Dwivedi et al., 2019 |

| | | | | | | | | | | | | | | | |
|---|---|---|---|---|---|---|---|---|---|---|---|---|---|---|---|
| GL | GL_17AY | AX-110606231 | 2A | 717141146 | 0.2817/-- | 1.79E-06 | GEMMA | TraesCS2A01G482400 | 3.91[G] | 1.26e-07[G] | *OsMED15a* | | | 3.35E-24 | Dwivedi et al., 2019 |
| GL | GL_17AY | AX-109507407 | 2A | 717389779 | 0.1464/-- | 1.82E-06 | GEMMA | TraesCS2A01G482400 | 3.91[G] | 1.26e-07[G] | *OsMED15a* | | | 3.35E-24 | Dwivedi et al., 2019 |
| GL | GL_17AY | AX-110575805 | 2A | 717423753 | 0.143/-- | 1.10E-06 | GEMMA | TraesCS2A01G482400 | 3.91[G] | 1.26e-07[G] | *OsMED15a* | | | 3.35E-24 | Dwivedi et al., 2019 |
| GL | GL_17AY | AX-111719891 | 2A | 717564817 | 0.1418/-- | 1.80E-06 | GEMMA | TraesCS2A01G482400 | 3.91[G] | 1.26e-07[G] | *OsMED15a* | | | 3.35E-24 | Dwivedi et al., 2019 |
| GL | GL_17AY | AX-110601263 | 2A | 717571266 | 0.1517/-- | 1.53E-06 | GEMMA | TraesCS2A01G482400 | 3.91[G] | 1.26e-07[G] | *OsMED15a* | | | 3.35E-24 | Dwivedi et al., 2019 |
| GL | GL_17AY | AX-110427653 | 2A | 717743589 | 0.1432/-- | 1.67E-06 | GEMMA | TraesCS2A01G482400 | 3.91[G] | 1.26e-07[G] | *OsMED15a* | | | 3.35E-24 | Dwivedi et al., 2019 |
| GL | GL_17AY | AX-108963673 | 2A | 717754883 | 0.1432/-- | 1.67E-06 | GEMMA | TraesCS2A01G482400 | 3.91[G] | 1.26e-07[G] | *OsMED15a* | | | 3.35E-24 | Dwivedi et al., 2019 |
| GL | GL_17AY | AX-109344129 | 2A | 717803976 | 0.1432/-- | 1.67E-06 | GEMMA | TraesCS2A01G482400 | 3.91[G] | 1.26e-07[G] | *OsMED15a* | | | 3.35E-24 | Dwivedi et al., 2019 |
| GL | GL_17AY | AX-111611081 | 2A | 717816882 | 0.1432/-- | 1.67E-06 | GEMMA | TraesCS2A01G482400 | 3.91[G] | 1.26e-07[G] | *OsMED15a* | | | 3.35E-24 | Dwivedi et al., 2019 |
| GL | GL_17AY | AX-111021808 | 2A | 718158173 | 0.1475/-- | 1.82E-06 | GEMMA | TraesCS2A01G482400 | 3.91[G] | 1.26e-07[G] | *OsMED15a* | | | 3.35E-24 | Dwivedi et al., 2019 |
| GL | GL_17AY | AX-109462354 | 2A | 718166340 | 0.1473/-- | 2.00E-06 | GEMMA | TraesCS2A01G482400 | 3.91[G] | 1.26e-07[G] | *OsMED15a* | | | 3.35E-24 | Dwivedi et al., 2019 |
| GL | GL_17AY | AX-111720998 | 2A | 719404597 | 0.1983/-- | 1.33E-06 | GEMMA | TraesCS2A01G482400 | 3.91[G] | 1.26e-07[G] | *OsMED15a* | | | 3.35E-24 | Dwivedi et al., 2019 |
| GL | GL_17AY | AX-109295270 | 2A | 718227374 | -0.0689/0.0723 | 4.39E-11 | IIIVmrMLM | TraesCS2A01G482400 | 3.91[G] | 1.26e-07[G] | *OsMED15a* | | | 3.35E-24 | Dwivedi et al., 2019 |
| GL | GL_18JY | AX-95072346 | 2D | 586363215 | -0.1915/-- | 2.08E-06 | GEMMA | TraesCS2D01G482500 | 3.2[G] | 2.38e-30[G] | *OsMED15a* | | | 1.19E-57 | Dwivedi et al., 2019 |
| GL | GL_17SQ | AX-108956954 | 4A | 579840479 | 0.3675/-- | 1.99E-06 | GEMMA | TraesCS4A01G266900 | 5.87[G] | 4.21e-23[G] | *OsGS1* | | | 1.05E-05 | Liu et al., 2022 |
| GL | GL_17AY | AX-110074625 | 6D | 462630227 | -0.0356/0.1295 | 3.65E-08 | IIIVmrMLM | TraesCS6D01G378300 | -1.1[G] | 6.14e-05[G] | *OsIAA10* | | | 2.29E-12 | Ma et al., 2023 |
| GL | GL_18JY | AX-109038640 | 5B | 491066755 | -0.0304/-0.1113 | 1.20E-15 | IIIVmrMLM | TraesCS5B01G307600 | 3.29[G] | 7.01e-05[G] | *OsSCP46* | | | 8.02E-05 | Li et al., 2016 |
| GL | GL_17SQ | AX-108794942 | 3B | 794344602 | 0.0047/0.4259 | 1.13E-04 | IIIVmrMLM | TraesCS3B01G560400 | 5.73[G] | 2.39e-02[G] | *OsFl3* | | | 6.00E-14 | Guo et al., 2022 |
| GL | GL_17SQ | AX-108794942 | 3B | 794344602 | 0.0047/0.4259 | 1.13E-04 | IIIVmrMLM | TraesCS3B01G561000 | 13.87[G] | 8.05e-28[G] | *OsFl3* | | | 3.83E-14 | Guo et al., 2022 |
| GL | GL_17SQ | AX-94547062 | 4B | 612846887 | -0.2796/-0.1464 | 1.90E-98 | IIIVmrMLM | TraesCS4B01G321800 | 9.65[G] | 4.00e-165[G] | *OsCDPK1* | | | 1.39E-41 | Ho et al., 2013 |
| GWLR | GWLR_17SQ | AX-110409742 | 4D | 501122912 | 0.0062/-- | 2.18E-05 | BLUPmrMLM | TraesCS4D01G350500 | -9.66[G] | 1.48e-33[G] | | GO:0005982 | starch metabolic process | 2.02E-08 | |
| GWLR | GWLR_18XX | AX-109303311 | 4D | 502834145 | -0.004/-- | 4.21E-05 | BLUPmrMLM | TraesCS4D01G350500 | -9.66[G] | 1.48e-33[G] | | GO:0005982 | starch metabolic process | 1.24E-17 | |
| GWLR | GWLR_17XX | AX-110415726 | 4D | 502594399 | -0.132/-- | 1.52E-09 | EMMAX | TraesCS4D01G350500 | -9.66[G] | 1.48e-33[G] | | GO:0005982 | starch metabolic process | 4.25E-08 | |
| GWLR | GWLR_17XX | AX-110409742 | 4D | 501122912 | -0.0045/-0.0032 | 2.98E-11 | IIIVmrMLM | TraesCS4D01G350500 | -9.66[G] | 1.48e-33[G] | | GO:0005982 | starch metabolic process | 4.25E-08 | |

| | | | | | | | | | | | | | | | |
|---|---|---|---|---|---|---|---|---|---|---|---|---|---|---|---|
| GWLR | GWLR_17SQ | AX-111613093 | 7D | 19390238 | -0.0098/-- | 2.61E-08 | BLUPmrMLM | TraesCS7D01G039500 | -1.41 [G] | 5.79e-06 [G] | | GO:0005982 | starch metabolic process | 1.35E-04 | |
| GWLR | GWLR_18XX | AX-109303311 | 4D | 502834145 | -0.004/-- | 4.21E-05 | BLUPmrMLM | TraesCS4D01G353800 | -2.84 [G] | 3.00e-32 [G] | | GO:0005982 | starch metabolic process | 1.28E-14 | |
| GWLR | GWLR_17XX | AX-110415726 | 4D | 502594399 | -0.132/-- | 1.52E-09 | EMMAX | TraesCS4D01G353800 | -2.84 [G] | 3.00e-32 [G] | | GO:0005982 | starch metabolic process | 1.16E-07 | |
| GWLR | GWLR_18XX | AX-111729579 | 6A | 2398289 | -0.0073/-- | 6.33E-12 | BLUPmrMLM | TraesCS6A01G009100 | 8.26 [G] | 3.69e-07 [G] | | GO:0015770 | sucrose transport | 2.90E-03 | |
| GWLR | GWLR_19XX | AX-110445450 | 4A | 687159341 | -0.0073/-- | 2.67E-06 | BLUPmrMLM | TraesCS4A01G418200 | 16.28 [G] | 2.95e-39 [G] | | GO:0005982 | starch metabolic process | 1.46E-02 | |
| GWLR | GWLR_BLUP | AX-94446985 | 6B | 7137766 | -0.0031/-- | 5.61E-06 | BLUPmrMLM | TraesCS6B01G015100 | 6.15 [G] | 6.14e-04 [G] | | GO:0015770 | sucrose transport | 4.82E-14 | |
| GWLR | GWLR_18XX | AX-109899833 | 2A | 509101562 | 0.0077/0.02 | 6.90E-24 | IIIVmrMLM | TraesCS2A01G295400 | 3.54 [G] | 1.84e-10 [G] | | GO:0005982 | starch metabolic process | 1.23E-02 | |
| GWLR | GWLR_18ZMD | AX-109281742 | 7A | 723960193 | -0.0062/0.1002 | 7.94E-07 | IIIVmrMLM | TraesCS7A01G549100 | 7.04 [G] | 6.55e-43 [G] | | GO:0005982 | starch metabolic process | 7.65E-09 | |
| GWLR | GWLR_18ZMD | AX-109281742 | 7A | 723960193 | -0.0062/0.1002 | 7.94E-07 | IIIVmrMLM | TraesCS7A01G549200 | 13.66 [G] | 4.55e-28 [G] | | GO:0005982 | starch metabolic process | 1.70E-09 | |
| GWLR | GWLR_18ZMD | AX-109281742 | 7A | 723960193 | -0.0062/0.1002 | 7.94E-07 | IIIVmrMLM | TraesCS7A01G549300 | 12.05 [G] | 6.10e-22 [G] | | GO:0005982 | starch metabolic process | 3.52E-05 | |
| GWLR | GWLR_BLUP | AX-111014568 | 1A | 535855735 | 0.0016/-0.0105 | 2.43E-08 | IIIVmrMLM | TraesCS1A01G353300 | -1.3 [G] | 9.91e-03 [G] | | GO:0005982 | starch metabolic process | 1.27E-13 | |
| GWLR | GWLR_17SQ | AX-94689724 | 6B | 575638302 | -0.0044/-- | 9.67E-05 | BLUPmrMLM | TraesCS6B01G327500 | -1.89 [G] | 1.40e-13 [G] | *OsGS1* | | | 8.10E-04 | Liu et al., 2022 |
| GWLR | GWLR_18ZMD | AX-111137061 | 4A | 537963260 | 0.0136/-- | 6.80E-08 | BLUPmrMLM | TraesCS4A01G227900 | -1.12 [G] | 7.22e-08 [G] | *OsAL11* | | | 2.74E-02 | Yang et al., 2022 |
| GWLR | GWLR_BLUP | AX-110610514 | 3B | 107772536 | 0.0018/-- | 1.52E-04 | BLUPmrMLM | TraesCS3B01G128700 | 8.23 [G] | 1.06e-09 [G] | *OsCKX1* | | | 4.74E-03 | Rong et al., 2022 |
| GWLR | GWLR_17XX | AX-108735017 | 2D | 5761352 | -0.1767/-- | 3.45E-14 | EMMAX | TraesCS2D01G011200 | 12.06 [G] | 5.52e-21 [G] | *OsMED15a* | | | 1.17E-03 | Dwivedi et al., 2019 |
| GWLR | GWLR_17XX | AX-109308225 | 5D | 456705502 | -0.0491/-- | 2.28E-08 | EMMAX | TraesCS5D01G387700 | 5.01 [G] | 3.22e-17 [G] | *RGG1* | | | 7.91E-09 | Tao et al., 2020 |
| GWLR | GWLR_17XX | AX-109308225 | 5D | 456705502 | 0.0493/-- | 2.33E-08 | GEMMA | TraesCS5D01G387700 | 5.01 [G] | 3.22e-17 [G] | *RGG1* | | | 7.91E-09 | Tao et al., 2020 |
| GWLR | GWLR_17SQ | AX-94464913 | 2A | 754361180 | -0.0407/-- | 2.99E-08 | GEMMA | TraesCS2A01G548100 | 4.71 [G] | 6.71e-18 [G] | *OsPAO5* | | | 1.11E-17 | Lv et al., 2021 |
| GWLR | GWLR_17SQ | AX-109868941 | 2B | 682929028 | 0.0076/-0.0082 | 6.51E-23 | IIIVmrMLM | TraesCS2B01G485700 | 8.12 [G] | 2.06e-08 [G] | *GSD1* | | | 8.35E-05 | Gui et al., 2014 |
| GWLR | GWLR_17XX | AX-109392460 | 2B | 682175602 | 0.0019/0.0987 | 5.10E-11 | IIIVmrMLM | TraesCS2B01G485700 | 8.12 [G] | 2.06e-08 [G] | *GSD1* | | | 5.84E-04 | Gui et al., 2014 |
| GWLR | GWLR_18ZMD | AX-110481013 | 4A | 73180517 | -0.0067/0.0686 | 1.24E-09 | IIIVmrMLM | TraesCS4A01G074300 | 1.71 [G] | 1.62e-07 [G] | *OsGBP3* | | | 1.41E-03 | Gong et al., 2018 |
| GWLR | GWLR_17XX | AX-109820200 | 4A | 73548087 | -0.0051/0.0038 | 3.16E-13 | IIIVmrMLM | TraesCS4A01G074300 | 1.71 [G] | 1.62e-07 [G] | *OsGBP3* | | | 3.74E-03 | Gong et al., 2018 |
| GWLR | GWLR_BLUP | AX-110481013 | 4A | 73180517 | -0.0015/0.0177 | 9.32E-16 | IIIVmrMLM | TraesCS4A01G074300 | 1.71 [G] | 1.62e-07 [G] | *OsGBP3* | | | 3.65E-03 | Gong et al., 2018 |
| GW | GW_17SQ | AX-110554761 | 2D | 600065511 | -0.0169/-- | 1.34E-04 | BLUPmrMLM | TraesCS2D01G505700 | 9.36 [G] | 4.36e-12 [G] | | GO:0015770 | sucrose transport | 9.65E-09 | |

| | | | | | | | | | | | | | | | |
|---|---|---|---|---|---|---|---|---|---|---|---|---|---|---|---|
| GW | GW_17XX | AX-111068326 | 2D | 601273284 | 0.0035/-0.0805 | 1.04E-06 | IIIVmrMLM | TraesCS2D01G505700 | 9.36 $^{G}$ | 4.36e-12 $^{G}$ | | GO:0015770 | sucrose transport | 2.12E-09 | |
| GW | GW_BLUP | AX-110511281 | 2D | 599866981 | 0.0096/-0.0148 | 7.87E-07 | IIIVmrMLM | TraesCS2D01G505700 | 9.36 $^{G}$ | 4.36e-12 $^{G}$ | | GO:0015770 | sucrose transport | 1.43E-16 | |
| GW | GW_17SQ | AX-109956383 | 7A | 20494842 | 0.0309/-- | 1.22E-06 | BLUPmrMLM | TraesCS7A01G043700 | -1.21 $^{G}$ | 3.78e-08 $^{G}$ | | GO:0005982 | starch metabolic process | 4.02E-17 | |
| GW | GW_18XX | AX-94788204 | 6D | 2891109 | -0.0289/-- | 4.72E-05 | BLUPmrMLM | TraesCS6D01G009700 | 5.88 $^{G}$ | 1.87e-02 $^{G}$ | | GO:0015770 | sucrose transport | 1.02E-02 | |
| GW | GW_17XX | AX-94862958 | 6D | 2103224 | 0.0047/-0.0366 | 6.33E-18 | IIIVmrMLM | TraesCS6D01G009700 | 5.88 $^{G}$ | 1.87e-02 $^{G}$ | | GO:0015770 | sucrose transport | 4.20E-02 | |
| GW | GW_18XX | AX-94788204 | 6D | 2891109 | -0.0289/-- | 4.72E-05 | BLUPmrMLM | TraesCS6D01G012100 | 10.27 $^{G}$ | 1.28e-12 $^{G}$ | | GO:0015770 | sucrose transport | 3.89E-09 | |
| GW | GW_18XX | AX-111186786 | 1A | 518515638 | 0.0984/-- | 1.50E-07 | FarmCPU | TraesCS1A01G329900 | 10.27 $^{G}$ | 8.24e-16 $^{G}$ | | GO:0010581 | starch biosynthetic process | 5.76E-107 | |
| GW | GW_17SQ | AX-111130984 | 6A | 599676451 | -0.0289/-0.0534 | 5.52E-15 | IIIVmrMLM | TraesCS6A01G382400 | -13.21 $^{G}$ | 8.87e-19 $^{G}$ | | GO:0015770 | sucrose transport | 4.19E-30 | |
| GW | GW_17SQ | AX-89672871 | 6B | 9098279 | 0.0247/0.0252 | 7.61E-11 | IIIVmrMLM | TraesCS6B01G015100 | 6.15 $^{G}$ | 6.14e-04 $^{G}$ | | GO:0015770 | sucrose transport | 5.38E-28 | |
| GW | GW_18JY | AX-109298719 | 7A | 78843415 | 0.002/-0.0808 | 2.59E-09 | IIIVmrMLM | TraesCS7A01G120300 | 1.37 $^{G}$ | 6.97e-07 $^{G}$ | | GO:0005982 | starch metabolic process | 1.92E-45 | |
| GW | GW_19XX | AX-110467577 | 6B | 127324922 | -0.0032/0.0839 | 5.76E-07 | IIIVmrMLM | TraesCS6B01G130700 | 1.41 $^{G}$ | 1.99e-04 $^{G}$ | | GO:0005982 | starch metabolic process | 9.86E-13 | |
| GW | GW_19XX | AX-110413774 | 7B | 217481381 | -0.0043/0.0356 | 1.15E-07 | IIIVmrMLM | TraesCS7B01G160000 | 12.52 $^{G}$ | 1.03e-21 $^{G}$ | | GO:0015770 | sucrose transport | 5.03E-37 | |
| GW | GW_BLUP | AX-111614749 | 1A | 439669876 | -0.015/-0.0004 | 1.69E-12 | IIIVmrMLM | TraesCS1A01G247100 | 3.44 $^{G}$ | 2.77e-21 $^{G}$ | | GO:0005982 | starch metabolic process | 3.21E-45 | |
| GW | GW_BLUP | AX-94381509 | 1B | 263312785 | -0.0112/nan | 7.87E-10 | IIIVmrMLM | TraesCS1B01G155700 | -1.73 $^{G}$ | 1.14e-13 $^{G}$ | | GO:0005982 | starch metabolic process | 3.29E-03 | |
| GW | GW_BLUP | AX-111170401 | 6B | 212566503 | -0.0046/0.1042 | 2.62E-12 | IIIVmrMLM | TraesCS6B01G186700 | 5.54 $^{G}$ | 4.13e-07 $^{G}$ | | GO:0005982 | starch metabolic process | 3.53E-51 | |
| GW | GW_17XX | AX-95083211 | 1B | 672928015 | -0.0324/-- | 5.41E-06 | BLUPmrMLM | TraesCS1B01G462000 | 1.7 $^{G}$ | 5.26e-10 $^{G}$ | *OsCTPS1* | | | 6.32E-10 | Yoon et al., 2021 |
| GW | GW_17XX | AX-95083211 | 1B | 672928015 | 0.022/-0.0035 | 3.79E-04 | IIIVmrMLM | TraesCS1B01G462000 | 1.7 $^{G}$ | 5.26e-10 $^{G}$ | *OsCTPS1* | | | 6.32E-10 | Yoon et al., 2021 |
| GW | GW_19XX | AX-108725863 | 7B | 135773615 | 0.02/-- | 1.63E-04 | BLUPmrMLM | TraesCS7B01G115300 | 9.73 $^{G}$ | 9.44e-11 $^{G}$ | *OsGSR1* | | | 1.95E-54 | Li et al., 2021 |
| GW | GW_17SQ | AX-110416871 | 2A | 709414300 | -0.0041/-0.1107 | 8.39E-09 | IIIVmrMLM | TraesCS2A01G464000 | 5.27 $^{G}$ | 4.05e-06 $^{G}$ | *GSD1* | | | 4.99E-15 | Gui et al., 2014 |
| GW | GW_18JY | AX-111763398 | 2B | 682929794 | 0.0406/0.0161 | 1.96E-20 | IIIVmrMLM | TraesCS2B01G485700 | 8.12 $^{G}$ | 2.06e-08 $^{G}$ | *GSD1* | | | 4.70E-12 | Gui et al., 2014 |
| TGW | TGW_17SQ | AX-94707348 | 2A | 535713056 | 1.137/-- | 1.65E-06 | BLUPmrMLM | TraesCS2A01G310300 | 13.53 $^{G}$ | 1.03e-43 $^{G}$ | *OsBEIIb* | GO:0005982 | starch metabolic process | 1.26E-14 | Chen et al., 2022 |
| TGW | TGW_18JY | AX-95075921 | 7A | 19007631 | -0.8837/-- | 4.10E-06 | BLUPmrMLM | TraesCS7A01G043700 | -1.21 $^{G}$ | 3.78e-08 $^{G}$ | | GO:0005982 | starch metabolic process | 5.77E-23 | |
| TGW | TGW_18XX | AX-108833696 | 4A | 712602406 | -0.4616/-- | 2.55E-05 | BLUPmrMLM | TraesCS4A01G444300 | -1.32 $^{G}$ | 7.51e-07 $^{G}$ | | GO:0005982 | starch metabolic process | 7.96E-23 | |
| TGW | TGW_18XX | AX-109037034 | 6A | 603809881 | -0.5066/-- | 4.60E-05 | BLUPmrMLM | TraesCS6A01G382500 | -4.01 $^{G}$ | 2.94e-12 $^{G}$ | | GO:0015770 | sucrose transport | 1.21E-45 | |

| | | | | | | | | | | | | | | | |
|---|---|---|---|---|---|---|---|---|---|---|---|---|---|---|---|
| TGW | TGW_18XX | AX-109037034 | 6A | 603809881 | -0.5066/-- | 4.60E-05 | BLUPmrMLM | TraesCS6A01G382600 | -10.09$^{G}$ | 1.35e-87$^{G}$ | | GO:0015770 | sucrose transport | 3.36E-47 | |
| TGW | TGW_17JY | AX-110676928 | 7D | 619079252 | -30.281/-- | 1.20E-07 | EMMAX | TraesCS7D01G525100 | 5.23$^{G}$ | 5.36e-03$^{G}$ | | GO:0005985 | sucrose metabolic process | 3.50E-02 | |
| TGW | TGW_17SQ | AX-94684170 | 1B | 597537011 | 3.5115/-- | 7.57E-07 | FarmCPU | TraesCS1B01G368500 | -2.32$^{G}$ | 4.05e-16$^{G}$ | | GO:0005982 | starch metabolic process | 6.98E-05 | |
| TGW | TGW_18XX | AX-94684170 | 1B | 597537011 | 3.7911/-- | 8.46E-07 | FarmCPU | TraesCS1B01G368500 | -2.32$^{G}$ | 4.05e-16$^{G}$ | | GO:0005982 | starch metabolic process | 1.14E-04 | |
| TGW | TGW_19XX | AX-94684170 | 1B | 597537011 | 4.1841/-- | 1.44E-06 | FarmCPU | TraesCS1B01G368500 | -2.32$^{G}$ | 4.05e-16$^{G}$ | | GO:0005982 | starch metabolic process | 3.91E-04 | |
| TGW | TGW_18JY | AX-111186786 | 1A | 518515638 | 2.8875/-- | 4.68E-07 | FarmCPU | TraesCS1A01G329900 | 10.27$^{G}$ | 8.24e-16$^{G}$ | | GO:0010581 | starch biosynthetic process | 1.15E-89 | |
| TGW | TGW_17JY | AX-110457081 | 4D | 503958928 | 5.2225/-- | 8.86E-07 | GEMMA | TraesCS4D01G350500 | -9.66$^{G}$ | 1.48e-33$^{G}$ | | GO:0005982 | starch metabolic process | 3.76E-02 | |
| TGW | TGW_17JY | AX-110457081 | 4D | 503958928 | 5.2225/-- | 8.86E-07 | GEMMA | TraesCS4D01G353800 | -2.84$^{G}$ | 3.00e-32$^{G}$ | | GO:0005982 | starch metabolic process | 1.03E-02 | |
| TGW | TGW_17AY | AX-110994042 | 1A | 31549776 | 0.1419/2.0594 | 9.34E-08 | IIIVmrMLM | TraesCS1A01G050400 | 10.73$^{G}$ | 1.43e-17$^{G}$ | | GO:0015770 | sucrose transport | 4.43E-14 | |
| TGW | TGW_18JY | AX-109427839 | 2B | 728763995 | 0.5343/1.8962 | 1.12E-14 | IIIVmrMLM | TraesCS2B01G533300 | 5.28$^{G}$ | 2.94e-05$^{G}$ | | GO:0015770 | sucrose transport | 1.97E-23 | |
| TGW | TGW_18JY | AX-111473756 | 6B | 7304167 | 0.1835/-4.4872 | 2.29E-07 | IIIVmrMLM | TraesCS6B01G015100 | 6.15$^{G}$ | 6.14e-04$^{G}$ | *Os-11N3* | GO:0015770 | sucrose transport | 7.54E-36 | Luo et al., 2021 |
| TGW | TGW_18JY | AX-108932028 | 6D | 3317339 | 0.2677/-1.0407 | 6.07E-07 | IIIVmrMLM | TraesCS6D01G012100 | 10.27$^{G}$ | 1.28e-12$^{G}$ | *Os-11N3* | GO:0015770 | sucrose transport | 1.22E-10 | Luo et al., 2021 |
| TGW | TGW_18XX | AX-109501835 | 4B | 20590398 | 0.8724/0.2371 | 1.64E-34 | IIIVmrMLM | TraesCS4B01G029700 | 1.25$^{G}$ | 1.23e-05$^{G}$ | | GO:0005982 | starch metabolic process | 2.72E-31 | |
| TGW | TGW_BLUP | AX-109847836 | 6D | 2372237 | -0.0401/-2.265 | 2.78E-08 | IIIVmrMLM | TraesCS6D01G009700 | 5.88$^{G}$ | 1.87e-02$^{G}$ | | GO:0015770 | sucrose transport | 6.18E-03 | |
| TGW | TGW_17SQ | AX-110987235 | 1A | 584755997 | -0.8679/-- | 1.62E-06 | BLUPmrMLM | TraesCS1A01G435000 | -2.31$^{G}$ | 3.61e-03$^{G}$ | *STH1* | | | 2.24E-21 | Xiang et al., 2022 |
| TGW | TGW_18XX | AX-109981359 | 1A | 584807135 | 0.6502/-- | 4.36E-08 | BLUPmrMLM | TraesCS1A01G435000 | -2.31$^{G}$ | 3.61e-03$^{G}$ | *STH1* | | | 2.15E-24 | Xiang et al., 2022 |
| TGW | TGW_17JY | AX-108887078 | 1A | 584583811 | 0.1187/-1.8489 | 2.25E-06 | IIIVmrMLM | TraesCS1A01G435000 | -2.31$^{G}$ | 3.61e-03$^{G}$ | *STH1* | | | 4.65E-16 | Xiang et al., 2022 |
| TGW | TGW_18JY | AX-108917551 | 7B | 717344698 | -0.5852/-- | 1.62E-07 | BLUPmrMLM | TraesCS7B01G455000 | -3.24$^{G}$ | 1.74e-07$^{G}$ | *OsCKX11* | | | 2.23E-12 | Rong et al., 2022 |
| TGW | TGW_18XX | AX-110294806 | 1D | 42827184 | -0.5981/-- | 1.38E-08 | BLUPmrMLM | TraesCS1D01G063500 | 9.94$^{G}$ | 3.29e-06$^{G}$ | *OsLTPL36* | | | 1.70E-03 | Wang et al., 2015 |
| TGW | TGW_BLUP | AX-110083733 | 1D | 41593584 | -0.475/-- | 7.43E-07 | BLUPmrMLM | TraesCS1D01G063500 | 9.94$^{G}$ | 3.29e-06$^{G}$ | *OsLTPL36* | | | 1.75E-03 | Wang et al., 2015 |
| TGW | TGW_18XX | AX-110294806 | 1D | 42827184 | -0.5981/-- | 1.38E-08 | BLUPmrMLM | TraesCS1D01G063900 | 12.49$^{G}$ | 3.00e-20$^{G}$ | *OsLTPL36* | | | 1.88E-04 | Wang et al., 2015 |
| TGW | TGW_18JY | AX-108884734 | 1D | 46457630 | 0.3101/-1.8691 | 8.12E-08 | IIIVmrMLM | TraesCS1D01G063900 | 12.49$^{G}$ | 3.00e-20$^{G}$ | *OsLTPL36* | | | 7.26E-04 | Wang et al., 2015 |
| TGW | TGW_18XX | AX-111024713 | 4A | 594665324 | -0.6178/-- | 1.25E-10 | BLUPmrMLM | TraesCS4A01G289300 | 1.54$^{G}$ | 4.48e-11$^{G}$ | *OsSCL30* | | | 2.65E-05 | Zhang et al., 2022 |
| TGW | TGW_18XX | AX-109341101 | 6A | 271258432 | -0.7361/-- | 7.68E-07 | BLUPmrMLM | TraesCS6A01G194600 | 4.63$^{G}$ | 4.03e-16$^{G}$ | *FUWA* | | | 3.37E-02 | Chen et al., 2015 |

| | | | | | | | | | | | | | | | |
|---|---|---|---|---|---|---|---|---|---|---|---|---|---|---|---|
| TGW | TGW_19XX | AX-94488783 | 2A | 755788522 | 0.8834/-- | 6.64E-05 | BLUPmrMLM | TraesCS2A01G548100 | 4.71$^{G}$ | 6.71e-18$^{G}$ | *OsPAO5* | | | 2.18E-49 | Lv et al., 2021 |
| TGW | TGW_18JY | AX-109330937 | 2A | 755378747 | -0.1825/-1.3875 | 5.98E-08 | IIIVmrMLM | TraesCS2A01G548100 | 4.71$^{G}$ | 6.71e-18$^{G}$ | *OsPAO5* | | | 8.98E-34 | Lv et al., 2021 |
| TGW | TGW_19XX | AX-111021677 | 2B | 5767670 | 0.7824/-- | 1.40E-06 | BLUPmrMLM | TraesCS2B01G010800 | 7.55$^{G}$ | 9.29e-08$^{G}$ | *OsMED15a* | | | 3.30E-44 | Dwivedi et al., 2019 |
| TGW | TGW_17ZMD | AX-111122462 | 2B | 4610921 | 0.271/-2.26 | 4.52E-11 | IIIVmrMLM | TraesCS2B01G010800 | 7.55$^{G}$ | 9.29e-08$^{G}$ | *OsMED15a* | | | 2.27E-14 | Dwivedi et al., 2019 |
| TGW | TGW_19XX | AX-111021677 | 2B | 5767670 | 0.7824/-- | 1.40E-06 | BLUPmrMLM | TraesCS2B01G012500 | -2.22$^{G}$ | 1.48e-18$^{G}$ | *FLO19* | | | 1.86E-49 | Lei et al., 2021 |
| TGW | TGW_17ZMD | AX-111122462 | 2B | 4610921 | 0.271/-2.26 | 4.52E-11 | IIIVmrMLM | TraesCS2B01G012500 | -2.22$^{G}$ | 1.48e-18$^{G}$ | *FLO19* | | | 4.22E-15 | Lei et al., 2021 |
| TGW | TGW_17XX | AX-109459815 | 7D | 392632007 | 2.0165/-- | 1.46E-06 | EMMAX | TraesCS7D01G309400 | -7.1$^{G}$ | 1.03e-61$^{G}$ | *OsHT* | | | 4.58E-10 | Guo et al., 2023 |
| TGW | TGW_17SQ | AX-110528135 | 4D | 445004978 | 3.4412/-- | 8.76E-07 | FarmCPU | TraesCS4D01G274200 | -2.38$^{G}$ | 2.60e-03$^{G}$ | *OsNBL3* | | | 7.06E-05 | Qiu et al., 2021 |
| TGW | TGW_17XX | AX-86183921 | 1B | 673745594 | -3.2014/-- | 2.25E-06 | FarmCPU | TraesCS1B01G462000 | 1.7$^{G}$ | 5.26e-10$^{G}$ | *OsCTPS1* | | | 7.21E-15 | Yoon et al., 2021 |
| TGW | TGW_17AY | AX-111590048 | 1A | 38727469 | 0.6075/0.0335 | 8.05E-05 | IIIVmrMLM | TraesCS1A01G058400 | -2.65$^{G}$ | 2.99e-11$^{G}$ | *RSR1* | | | 1.62E-17 | Fu et al., 2010 |
| TGW | TGW_17AY | AX-109855276 | 3B | 406663792 | -0.063/3.1684 | 1.30E-07 | IIIVmrMLM | TraesCS3B01G252000 | -2.41$^{G}$ | 5.84e-08$^{G}$ | *OsCTPS1* | | | 8.67E-28 | Yoon et al., 2021 |
| TGW | TGW_18XX | AX-110514823 | 1A | 43330926 | 0.0443/1.8406 | 1.24E-11 | IIIVmrMLM | TraesCS1A01G063200 | 5.21$^{G}$ | 1.02e-02$^{G}$ | *OsLTPL36* | | | 1.88E-08 | Wang et al., 2015 |
| TGW | TGW_18XX | AX-111684719 | 5B | 477953826 | -0.0325/-1.2522 | 1.44E-06 | IIIVmrMLM | TraesCS5B01G292100 | 5.46$^{G}$ | 3.67e-03$^{G}$ | *OsBC1* | | | 4.81E-67 | Ruan et al., 2018 |
| TGW | TGW_17ZMD | AX-109260317 | 1D | 38861232 | -0.3799/-5.8593 | 5.62E-11 | IIIVmrMLM | TraesCS1D01G059200 | -3.09$^{G}$ | 1.86e-14$^{G}$ | *RSR1* | | | 5.03E-11 | Fu et al., 2010 |
| TGW | TGW_17ZMD | AX-109607606 | 3A | 551506355 | 0.5207/-5.0637 | 4.27E-12 | IIIVmrMLM | TraesCS3A01G311000 | 10.12$^{G}$ | 2.19e-12$^{G}$ | *Gn1a* | | | 1.21E-29 | Rong et al., 2022 |
| SNS | SNS_BLUP | AX-109518784 | 2B | 659086334 | 0.0089/0.1089 | 1.17E-08 | IIIVmrMLM | TraesCS2B01G464900 | 4.91$^{SNS1}$; -8.84$^{SNS2}$ | 3.57E-27$^{SNS1}$; 1.33E-30$^{SNS2}$ | *OsNPF7.6* | | | 1.95E-05 | Zhang et al.,2022 |
| SNS | SNS_BLUP | AX-111149345 | 3A | 12157923 | -0.03103 | 0.000183 | IIIVmrMLM | TraesCS3A01G022700 | 1.54$^{SNS2}$; -1.73$^{SNS1}$ | 1.03E-05$^{SNS2}$; 1.62E-04$^{SNS1}$ | | GO:2000028 | regulation of photoperiodism, flowering | 1.35E-05 | |
| SNS | SNS_BLUP | AX-111149345 | 3A | 12157923 | -0.03103 | 0.000183 | IIIVmrMLM | TraesCS3A01G026900 | -2.26$^{SNS2}$ | 5.84E-12$^{SNS2}$ | | GO:0048573 | photoperiodism, flowering | 3.82E-02 | |
| SNS | SNS_BLUP | AX-111149345 | 3A | 12157923 | -0.03103 | 0.000183 | IIIVmrMLM | TraesCS3A01G026900 | -2.26$^{SNS2}$ | 5.84E-12$^{SNS2}$ | | GO:0009648 | photoperiodism | 3.82E-02 | |
| SNS | SNS_BLUP | AX-111149345 | 3A | 12157923 | -0.03103 | 0.000183 | IIIVmrMLM | TraesCS3A01G026900 | -2.26$^{SNS2}$ | 5.84E-12$^{SNS2}$ | | GO:0048574 | long-day photoperiodism, flowering | 3.82E-02 | |
| SNS | SNS_BLUP | AX-111149345 | 3A | 12157923 | -0.03103 | 0.000183 | IIIVmrMLM | TraesCS3A01G026900 | -2.26$^{SNS2}$ | 5.84E-12$^{SNS2}$ | | GO:0048571 | long-day photoperiodism | 3.82E-02 | |
| SNS | SNS_18AY | AX-94522205 | 3A | 638010015 | -0.631292298783947/-- | 7.21E-07 | FarmCPU | TraesCS3A01G391600 | -1.31$^{SNS1}$; 1.06$^{SNS2}$ | 4.00E-08$^{SNS1}$; 1.13E-02$^{SNS2}$ | | GO:0009735 | response to cytokinin | 1.44E-02 | |
| SNS | SNS_18AY | AX-94522205 | 3A | 638010015 | -0.631292298783947/-- | 7.21E-07 | FarmCPU | TraesCS3A01G391600 | -1.31$^{SNS1}$; 1.06$^{SNS2}$ | 4.00E-08$^{SNS1}$; 1.13E-02$^{SNS2}$ | | GO:0071368 | cellular response to cytokinin stimulus | 1.44E-02 | |
| SNS | SNS_18AY | AX-94522205 | 3A | 638010015 | -0.631292298783947/-- | 7.21E-07 | FarmCPU | TraesCS3A01G391600 | -1.31$^{SNS1}$; 1.06$^{SNS2}$ | 4.00E-08$^{SNS1}$; 1.13E-02$^{SNS2}$ | | GO:0009736 | cytokinin-activated signaling pathway | 1.44E-02 | |

| | | | | | | | | | | | | | | | |
|---|---|---|---|---|---|---|---|---|---|---|---|---|---|---|---|
| SNS | SNS_BLUP | AX-108973844 | 3A | 725194673 | 0.0488/-- | 5.69E-06 | BLUPmrMLM | TraesCS3A01G502900 | 6.15 $^{SNS1}$; -5.79 $^{SNS2}$ | 1.50E-04 $^{SNS1}$; 4.99E-04 $^{SNS2}$ | | GO:0009735 | response to cytokinin | 8.88E-04 | |
| SNS | SNS_18XX | AX-110073738 | 3B | 668888194 | -0.00624 | 1.35E-08 | IIIVmrMLM | TraesCS3B01G428100 | 4.62 $^{SNS2}$; -6.47 $^{SNS1}$ | 2.79E-08 $^{SNS2}$; 3.44E-12 $^{SNS1}$ | | GO:2000028 | regulation of photoperiodism, flowering | 1.21E-02 | |
| SNS | SNS_17JY | AX-109379765 | 3B | 767539773 | 0.5926/-- | 2.99E-06 | BLUPmrMLM | TraesCS3B01G525300 | 1.79 $^{SNS1}$ | 3.27E-03 $^{SNS1}$ | | GO:0009690 | cytokinin metabolic process | 4.75E-07 | |
| SNS | SNS_17JY | AX-109379765 | 3B | 767539773 | 0.5926/-- | 2.99E-06 | BLUPmrMLM | TraesCS3B01G525300 | 1.79 $^{SNS1}$ | 3.27E-03 $^{SNS1}$ | | GO:0009823 | cytokinin catabolic process | 4.75E-07 | |
| SNS | SNS_18JY | AX-111101026 | 4A | 78942874 | -0.4027/-- | 1.55E-06 | BLUPmrMLM | TraesCS4A01G079300 | 5.62 $^{SNS2}$ | 6.46E-04 $^{SNS2}$ | | GO:0009736 | cytokinin-activated signaling pathway | 6.95E-04 | |
| SNS | SNS_18JY | AX-111101026 | 4A | 78942874 | -0.4027/-- | 1.55E-06 | BLUPmrMLM | TraesCS4A01G079300 | 5.62 $^{SNS2}$ | 6.46E-04 $^{SNS2}$ | | GO:0071368 | cellular response to cytokinin stimulus | 6.95E-04 | |
| SNS | SNS_18JY | AX-111101026 | 4A | 78942874 | -0.4027/-- | 1.55E-06 | BLUPmrMLM | TraesCS4A01G079300 | 5.62 $^{SNS2}$ | 6.46E-04 $^{SNS2}$ | | GO:0010476 | gibberellin mediated signaling pathway | 6.95E-04 | |
| SNS | SNS_18JY | AX-111101026 | 4A | 78942874 | -0.4027/-- | 1.55E-06 | BLUPmrMLM | TraesCS4A01G079300 | 5.62 $^{SNS2}$ | 6.46E-04 $^{SNS2}$ | | GO:0009735 | response to cytokinin | 6.95E-04 | |
| SNS | SNS_18JY | AX-111101026 | 4A | 78942874 | -0.4027/-- | 1.55E-06 | BLUPmrMLM | TraesCS4A01G079300 | 5.62 $^{SNS2}$ | 6.46E-04 $^{SNS2}$ | | GO:0071370 | cellular response to gibberellin stimulus | 6.95E-04 | |
| SNS | SNS_18JY | AX-111101026 | 4A | 78942874 | -0.4027/-- | 1.55E-06 | BLUPmrMLM | TraesCS4A01G079300 | 5.62 $^{SNS2}$ | 6.46E-04 $^{SNS2}$ | | GO:0009739 | response to gibberellin | 6.95E-04 | |
| SNS | SNS_18AY | AX-109416710 | 4A | 579541337 | -0.18392 | 2.21E-26 | IIIVmrMLM | TraesCS4A01G266900 | -2.64 $^{SNS1}$ | 1.07E-11 $^{SNS1}$ | *OsGS1* | | | 8.24E-03 | Ohashi et al.,2018 |
| SNS | SNS_17JY | AX-109297427 | 5A | 480786023 | -24.1429 | 4.45E-17 | IIIVmrMLM | TraesCS5A01G272400 | 2.72 $^{SNS2}$; -3.57 $^{SNS1}$ | 1.01E-24 $^{SNS2}$; 1.37E-64 $^{SNS1}$ | | GO:0009735 | response to cytokinin | 3.79E-07 | |
| SNS | SNS_18JY | AX-111489741 | 5B | 642047731 | 0.1643/-- | 6.12E-05 | BLUPmrMLM | TraesCS5B01G469600 | 5.60 $^{SNS1}$ | 2.86E-03 $^{SNS1}$ | | GO:0009691 | cytokinin biosynthetic process | 4.64E-09 | |
| SNS | SNS_18JY | AX-111489741 | 5B | 642047731 | 0.1643/-- | 6.12E-05 | BLUPmrMLM | TraesCS5B01G469600 | 5.60 $^{SNS1}$ | 2.86E-03 $^{SNS1}$ | | GO:0009690 | cytokinin metabolic process | 4.64E-09 | |
| SNS | SNS_18AY | AX-108762572 | 7A | 627149804 | -0.2646/-- | 2.15E-05 | BLUPmrMLM | TraesCS7A01G433000 | -5.53 $^{SNS1}$; 7.77 $^{SNS2}$ | 2.19E-14 $^{SNS1}$; 1.64E-09 $^{SNS2}$ | | GO:0048587 | regulation of short-day photoperiodism, flowering | 9.44E-31 | |
| SNS | SNS_18AY | AX-108762572 | 7A | 627149804 | -0.2646/-- | 2.15E-05 | BLUPmrMLM | TraesCS7A01G433000 | -5.53 $^{SNS1}$; 7.77 $^{SNS2}$ | 2.19E-14 $^{SNS1}$; 1.64E-09 $^{SNS2}$ | | GO:2000028 | regulation of photoperiodism, flowering | 9.44E-31 | |
| SNS | SNS_18XX | AX-94770214 | 7B | 498523685 | -0.1655/-- | 6.72E-13 | IIIVmrMLM | TraesCS7B01G271200 | -3.05 $^{SNS2}$ | 4.45E-15 $^{SNS2}$ | *OsAAP3* | | | 4.67E-03 | Lu et al.,2018 |

**Supplementary Table S16**. All QTNs identified as associated with active filling period in wheat.

| Trait | Marker | Chr | Position | Add.effect | Dom.effect | P-value | Method | Sig. |
|---|---|---|---|---|---|---|---|---|
| T_17AY | AX-95177876 | 2A | 88009461 | -25.407 | | 2.34E-07 | EMMAX | SUG |
| T_17AY | AX-110049347 | 3B | 493392497 | -1.027 | | 1.71E-05 | BLUPmrMLM | SUG |
| T_17AY | AX-110460659 | 3B | 739908219 | 1.473 | | 2.74E-05 | BLUPmrMLM | SUG |
| T_17AY | AX-111464003 | 4B | 399825077 | -1.141 | | 1.27E-05 | BLUPmrMLM | SUG |
| T_17AY | AX-109515751 | 4B | 665916709 | 1.341 | | 1.56E-05 | BLUPmrMLM | SUG |
| T_17AY | AX-109515751 | 4B | 665916709 | -1.919 | | 1.75E-06 | EMMAX | SUG |
| T_17AY | AX-109515751 | 4B | 665916709 | 1.919 | | 1.09E-06 | GEMMA | SUG |
| T_17AY | AX-109866446 | 5B | 27479946 | -1.029 | | 2.86E-05 | BLUPmrMLM | SUG |
| T_17AY | AX-111135879 | 5B | 466629644 | -1.813 | | 4.29E-06 | BLUPmrMLM | SUG |
| T_17AY | AX-110112057 | 5B | 572145204 | -2.618 | | 2.28E-06 | FarmCPU | SUG |
| T_17AY | AX-111718398 | 5B | 572394959 | -2.647 | | 2.32E-06 | FarmCPU | SUG |
| T_17AY | AX-94876976 | 6A | 614669489 | -1.360 | | 1.26E-04 | BLUPmrMLM | SUG |
| T_17AY | AX-108805321 | 7D | 591007232 | -28.777 | -0.9412 | 0.00E+00 | IIIVmrMLM | SIG |
| T_17JY | AX-111258241 | 1A | 98490306 | -28.498 | | 2.26E-06 | EMMAX | SUG |
| T_17JY | AX-109913297 | 1A | 302636102 | -28.498 | | 2.26E-06 | EMMAX | SUG |
| T_17JY | AX-94642388 | 1A | 308787331 | -28.498 | | 2.26E-06 | EMMAX | SUG |
| T_17JY | AX-109494630 | 1A | 319319948 | -28.498 | | 2.26E-06 | EMMAX | SUG |
| T_17JY | AX-109622845 | 1A | 319441335 | -28.498 | | 2.26E-06 | EMMAX | SUG |
| T_17JY | AX-109018008 | 1A | 319888131 | -28.498 | | 2.26E-06 | EMMAX | SUG |
| T_17JY | AX-110510731 | 1A | 320892031 | -28.498 | | 2.26E-06 | EMMAX | SUG |
| T_17JY | AX-94764575 | 1A | 321113972 | -28.498 | | 2.26E-06 | EMMAX | SUG |
| T_17JY | AX-108932890 | 1A | 321408443 | -28.498 | | 2.26E-06 | EMMAX | SUG |
| T_17JY | AX-111536488 | 1A | 326320422 | 0.843 | -2.8797 | 1.92E-15 | IIIVmrMLM | SIG |
| T_17JY | AX-108845125 | 1A | 327542942 | -28.498 | | 2.26E-06 | EMMAX | SUG |
| T_17JY | AX-108999147 | 1A | 333921129 | -28.498 | | 2.26E-06 | EMMAX | SUG |
| T_17JY | AX-111468889 | 1A | 334200480 | -28.498 | | 2.26E-06 | EMMAX | SUG |
| T_17JY | AX-95235095 | 1A | 337010898 | -28.498 | | 2.26E-06 | EMMAX | SUG |
| T_17JY | AX-109559815 | 1A | 337151148 | -28.498 | | 2.26E-06 | EMMAX | SUG |
| T_17JY | AX-94652579 | 1A | 341868519 | -28.498 | | 2.26E-06 | EMMAX | SUG |
| T_17JY | AX-94903766 | 1A | 341871247 | -28.498 | | 2.26E-06 | EMMAX | SUG |
| T_17JY | AX-109333935 | 1A | 348684553 | 0.389 | -5.3229 | 1.94E-06 | IIIVmrMLM | SUG |
| T_17JY | AX-109346027 | 1A | 570424040 | 0.694 | -11.6977 | 4.23E-14 | IIIVmrMLM | SIG |
| T_17JY | AX-109528352 | 1A | 574336949 | -1.633 | | 1.44E-22 | IIIVmrMLM | SIG |
| T_17JY | AX-111133395 | 1A | 585352170 | 0.872 | | 1.26E-04 | BLUPmrMLM | SUG |
| T_17JY | AX-110512163 | 1B | 227720400 | 3.111 | | 2.14E-06 | GEMMA | SUG |
| T_17JY | AX-95213636 | 1B | 329945077 | -28.498 | | 2.26E-06 | EMMAX | SUG |
| T_17JY | AX-95081345 | 1B | 330507911 | -28.498 | | 2.26E-06 | EMMAX | SUG |
| T_17JY | AX-110909537 | 1B | 332138774 | -28.498 | | 2.26E-06 | EMMAX | SUG |
| T_17JY | AX-108794824 | 1B | 338007453 | -28.498 | | 2.26E-06 | EMMAX | SUG |
| T_17JY | AX-109827182 | 1B | 344373169 | -28.498 | | 2.26E-06 | EMMAX | SUG |
| T_17JY | AX-110601945 | 1B | 348474818 | -28.498 | | 2.26E-06 | EMMAX | SUG |

| | | | | | | | | |
|---|---|---|---|---|---|---|---|---|
| T_17JY | AX-110412833 | 1B | 350630782 | -28.498 | | 2.26E-06 | EMMAX | SUG |
| T_17JY | AX-109001855 | 1B | 357425696 | -28.498 | | 2.26E-06 | EMMAX | SUG |
| T_17JY | AX-94982009 | 1B | 357642256 | -28.498 | | 2.26E-06 | EMMAX | SUG |
| T_17JY | AX-94836816 | 1B | 361015117 | -28.498 | | 2.26E-06 | EMMAX | SUG |
| T_17JY | AX-94435895 | 1B | 364419847 | -28.498 | | 2.26E-06 | EMMAX | SUG |
| T_17JY | AX-95094416 | 1B | 372219987 | -28.498 | | 2.26E-06 | EMMAX | SUG |
| T_17JY | AX-109328971 | 1B | 374345603 | -28.498 | | 2.26E-06 | EMMAX | SUG |
| T_17JY | AX-94847341 | 1B | 613001140 | -3.541 | | 1.61E-06 | FarmCPU | SUG |
| T_17JY | AX-108736850 | 1D | 235572490 | -28.498 | | 2.26E-06 | EMMAX | SUG |
| T_17JY | AX-111098190 | 1D | 235672276 | -28.498 | | 2.26E-06 | EMMAX | SUG |
| T_17JY | AX-108924022 | 1D | 235677547 | -28.498 | | 2.26E-06 | EMMAX | SUG |
| T_17JY | AX-111465797 | 1D | 235677641 | -28.498 | | 2.26E-06 | EMMAX | SUG |
| T_17JY | AX-108846020 | 1D | 235980289 | -28.498 | | 2.26E-06 | EMMAX | SUG |
| T_17JY | AX-108951507 | 1D | 236176294 | -28.498 | | 2.26E-06 | EMMAX | SUG |
| T_17JY | AX-109345900 | 1D | 236312913 | -28.498 | | 2.26E-06 | EMMAX | SUG |
| T_17JY | AX-95245271 | 1D | 236438901 | -28.498 | | 2.26E-06 | EMMAX | SUG |
| T_17JY | AX-109409323 | 1D | 236440506 | -28.498 | | 2.26E-06 | EMMAX | SUG |
| T_17JY | AX-109583947 | 1D | 236446487 | -28.498 | | 2.26E-06 | EMMAX | SUG |
| T_17JY | AX-109866596 | 1D | 236676919 | -28.498 | | 2.26E-06 | EMMAX | SUG |
| T_17JY | AX-109868389 | 1D | 236808415 | -28.498 | | 2.26E-06 | EMMAX | SUG |
| T_17JY | AX-109318471 | 1D | 236857328 | -28.498 | | 2.26E-06 | EMMAX | SUG |
| T_17JY | AX-111541002 | 1D | 237147144 | -28.498 | | 2.26E-06 | EMMAX | SUG |
| T_17JY | AX-110433409 | 1D | 237322173 | -28.498 | | 2.26E-06 | EMMAX | SUG |
| T_17JY | AX-110413020 | 1D | 237322245 | -28.498 | | 2.26E-06 | EMMAX | SUG |
| T_17JY | AX-109910884 | 1D | 237322283 | -28.498 | | 2.26E-06 | EMMAX | SUG |
| T_17JY | AX-111199976 | 1D | 237322427 | -28.498 | | 2.26E-06 | EMMAX | SUG |
| T_17JY | AX-95184815 | 1D | 237325061 | -28.498 | | 2.26E-06 | EMMAX | SUG |
| T_17JY | AX-110973913 | 1D | 237363130 | -28.498 | | 2.26E-06 | EMMAX | SUG |
| T_17JY | AX-109828320 | 1D | 237363887 | -28.498 | | 2.26E-06 | EMMAX | SUG |
| T_17JY | AX-111392715 | 1D | 237749068 | -28.498 | | 2.26E-06 | EMMAX | SUG |
| T_17JY | AX-109364218 | 1D | 237804432 | -28.498 | | 2.26E-06 | EMMAX | SUG |
| T_17JY | AX-109998839 | 1D | 237918034 | -28.498 | | 2.26E-06 | EMMAX | SUG |
| T_17JY | AX-94948053 | 1D | 238069148 | -28.498 | | 2.26E-06 | EMMAX | SUG |
| T_17JY | AX-108784410 | 1D | 238072079 | -28.498 | | 2.26E-06 | EMMAX | SUG |
| T_17JY | AX-94949091 | 1D | 238074596 | -28.498 | | 2.26E-06 | EMMAX | SUG |
| T_17JY | AX-94725024 | 1D | 238075659 | -28.498 | | 2.26E-06 | EMMAX | SUG |
| T_17JY | AX-109939375 | 1D | 238323928 | -28.498 | | 2.26E-06 | EMMAX | SUG |
| T_17JY | AX-110418021 | 1D | 239151464 | -28.498 | | 2.26E-06 | EMMAX | SUG |
| T_17JY | AX-111594811 | 1D | 239196897 | -28.498 | | 2.26E-06 | EMMAX | SUG |
| T_17JY | AX-109625802 | 1D | 239839864 | -28.498 | | 2.26E-06 | EMMAX | SUG |
| T_17JY | AX-111454504 | 1D | 240127132 | -28.498 | | 2.26E-06 | EMMAX | SUG |
| T_17JY | AX-109676476 | 1D | 240306208 | -28.498 | | 2.26E-06 | EMMAX | SUG |
| T_17JY | AX-108736538 | 1D | 240597042 | -28.498 | | 2.26E-06 | EMMAX | SUG |
| T_17JY | AX-111010735 | 1D | 240597387 | -28.498 | | 2.26E-06 | EMMAX | SUG |

| | | | | | | | |
|---|---|---|---|---|---|---|---|
| T_17JY | AX-110108957 | 1D | 240989008 | -28.498 | 2.26E-06 | EMMAX | SUG |
| T_17JY | AX-110519414 | 1D | 241479210 | -28.498 | 2.26E-06 | EMMAX | SUG |
| T_17JY | AX-109882700 | 1D | 241878143 | -28.498 | 2.26E-06 | EMMAX | SUG |
| T_17JY | AX-110155982 | 1D | 241919780 | -28.498 | 2.26E-06 | EMMAX | SUG |
| T_17JY | AX-109212477 | 1D | 242026791 | -28.498 | 2.26E-06 | EMMAX | SUG |
| T_17JY | AX-109365163 | 1D | 242846619 | -28.498 | 2.26E-06 | EMMAX | SUG |
| T_17JY | AX-109641665 | 1D | 243808816 | -28.498 | 2.26E-06 | EMMAX | SUG |
| T_17JY | AX-111769865 | 1D | 244882828 | -28.498 | 2.26E-06 | EMMAX | SUG |
| T_17JY | AX-111495445 | 1D | 245106447 | -28.498 | 2.26E-06 | EMMAX | SUG |
| T_17JY | AX-110969582 | 1D | 246067633 | -28.498 | 2.26E-06 | EMMAX | SUG |
| T_17JY | AX-111545673 | 1D | 246391570 | -28.498 | 2.26E-06 | EMMAX | SUG |
| T_17JY | AX-111120255 | 1D | 246651022 | -28.498 | 2.26E-06 | EMMAX | SUG |
| T_17JY | AX-110815749 | 1D | 246679975 | -28.498 | 2.26E-06 | EMMAX | SUG |
| T_17JY | AX-109410891 | 1D | 246944230 | -28.498 | 2.26E-06 | EMMAX | SUG |
| T_17JY | AX-108783978 | 1D | 246944357 | -28.498 | 2.26E-06 | EMMAX | SUG |
| T_17JY | AX-110451197 | 1D | 247243061 | -28.498 | 2.26E-06 | EMMAX | SUG |
| T_17JY | AX-94388421 | 1D | 247302322 | -28.498 | 2.26E-06 | EMMAX | SUG |
| T_17JY | AX-110363370 | 1D | 247520027 | -28.498 | 2.26E-06 | EMMAX | SUG |
| T_17JY | AX-110957015 | 1D | 247521899 | -28.498 | 2.26E-06 | EMMAX | SUG |
| T_17JY | AX-111467050 | 1D | 247841054 | -28.498 | 2.26E-06 | EMMAX | SUG |
| T_17JY | AX-109260881 | 1D | 248971368 | -28.498 | 2.26E-06 | EMMAX | SUG |
| T_17JY | AX-109192445 | 1D | 249033515 | -28.498 | 2.26E-06 | EMMAX | SUG |
| T_17JY | AX-109646563 | 1D | 249033553 | -28.498 | 2.26E-06 | EMMAX | SUG |
| T_17JY | AX-111029003 | 1D | 249033760 | -28.498 | 2.26E-06 | EMMAX | SUG |
| T_17JY | AX-94756225 | 1D | 249377789 | -28.498 | 2.26E-06 | EMMAX | SUG |
| T_17JY | AX-109942431 | 1D | 249559851 | -28.498 | 2.26E-06 | EMMAX | SUG |
| T_17JY | AX-110126093 | 1D | 249569138 | -28.498 | 2.26E-06 | EMMAX | SUG |
| T_17JY | AX-108915716 | 1D | 249754077 | -28.498 | 2.26E-06 | EMMAX | SUG |
| T_17JY | AX-111804188 | 1D | 249940526 | -28.498 | 2.26E-06 | EMMAX | SUG |
| T_17JY | AX-110674821 | 1D | 249940962 | -14.522 | 1.69E-06 | EMMAX | SUG |
| T_17JY | AX-111799824 | 1D | 250077275 | -28.498 | 2.26E-06 | EMMAX | SUG |
| T_17JY | AX-111167326 | 1D | 250139641 | -28.498 | 2.26E-06 | EMMAX | SUG |
| T_17JY | AX-94958554 | 1D | 250139862 | -28.498 | 2.26E-06 | EMMAX | SUG |
| T_17JY | AX-94490718 | 1D | 250139967 | -28.498 | 2.26E-06 | EMMAX | SUG |
| T_17JY | AX-95162815 | 1D | 250140008 | -28.498 | 2.26E-06 | EMMAX | SUG |
| T_17JY | AX-95219882 | 1D | 250140395 | -28.498 | 2.26E-06 | EMMAX | SUG |
| T_17JY | AX-110089462 | 1D | 250140685 | -28.498 | 2.26E-06 | EMMAX | SUG |
| T_17JY | AX-108852205 | 1D | 250143864 | -28.498 | 2.26E-06 | EMMAX | SUG |
| T_17JY | AX-109623944 | 1D | 250213215 | -28.498 | 2.26E-06 | EMMAX | SUG |
| T_17JY | AX-111529542 | 1D | 250213701 | -28.498 | 2.26E-06 | EMMAX | SUG |
| T_17JY | AX-109374947 | 1D | 250213880 | -28.498 | 2.26E-06 | EMMAX | SUG |
| T_17JY | AX-94381613 | 1D | 250215461 | -28.498 | 2.26E-06 | EMMAX | SUG |
| T_17JY | AX-109166702 | 1D | 250324378 | -28.498 | 2.26E-06 | EMMAX | SUG |
| T_17JY | AX-111548765 | 1D | 250417047 | -28.498 | 2.26E-06 | EMMAX | SUG |

| | | | | | | | |
|---|---|---|---|---|---|---|---|
| T_17JY | AX-111615193 | 1D | 250427323 | -28.498 | 2.26E-06 | EMMAX | SUG |
| T_17JY | AX-111556355 | 1D | 250427768 | -28.498 | 2.26E-06 | EMMAX | SUG |
| T_17JY | AX-108823685 | 1D | 250744706 | -28.498 | 2.26E-06 | EMMAX | SUG |
| T_17JY | AX-109171296 | 1D | 251299487 | -28.498 | 2.26E-06 | EMMAX | SUG |
| T_17JY | AX-110041225 | 1D | 251307401 | -28.498 | 2.26E-06 | EMMAX | SUG |
| T_17JY | AX-110350667 | 1D | 251509994 | -28.498 | 2.26E-06 | EMMAX | SUG |
| T_17JY | AX-111488847 | 1D | 251711625 | -28.498 | 2.26E-06 | EMMAX | SUG |
| T_17JY | AX-111616512 | 1D | 252529720 | -28.498 | 2.26E-06 | EMMAX | SUG |
| T_17JY | AX-111947944 | 1D | 252565359 | -28.498 | 2.26E-06 | EMMAX | SUG |
| T_17JY | AX-109900503 | 1D | 252618669 | -28.498 | 2.26E-06 | EMMAX | SUG |
| T_17JY | AX-110070403 | 1D | 253199987 | -28.498 | 2.26E-06 | EMMAX | SUG |
| T_17JY | AX-94725591 | 1D | 253207333 | -28.498 | 2.26E-06 | EMMAX | SUG |
| T_17JY | AX-111259231 | 1D | 253518297 | -28.498 | 2.26E-06 | EMMAX | SUG |
| T_17JY | AX-94454946 | 1D | 254610655 | -28.498 | 2.26E-06 | EMMAX | SUG |
| T_17JY | AX-95116750 | 1D | 254610848 | -28.498 | 2.26E-06 | EMMAX | SUG |
| T_17JY | AX-110476430 | 1D | 254615121 | -28.498 | 2.26E-06 | EMMAX | SUG |
| T_17JY | AX-111456432 | 1D | 254790091 | -28.498 | 2.26E-06 | EMMAX | SUG |
| T_17JY | AX-110153362 | 1D | 255282797 | -28.498 | 2.26E-06 | EMMAX | SUG |
| T_17JY | AX-109518046 | 1D | 255426656 | -28.498 | 2.26E-06 | EMMAX | SUG |
| T_17JY | AX-95204038 | 1D | 255491391 | -28.498 | 2.26E-06 | EMMAX | SUG |
| T_17JY | AX-94525776 | 1D | 255493522 | -28.498 | 2.26E-06 | EMMAX | SUG |
| T_17JY | AX-109453816 | 1D | 255832696 | -28.498 | 2.26E-06 | EMMAX | SUG |
| T_17JY | AX-94458590 | 1D | 255834514 | -28.498 | 2.26E-06 | EMMAX | SUG |
| T_17JY | AX-109975146 | 1D | 256280210 | -28.498 | 2.26E-06 | EMMAX | SUG |
| T_17JY | AX-110559185 | 1D | 257017309 | -28.498 | 2.26E-06 | EMMAX | SUG |
| T_17JY | AX-109523705 | 1D | 257351979 | -28.498 | 2.26E-06 | EMMAX | SUG |
| T_17JY | AX-108962788 | 1D | 257353834 | -28.498 | 2.26E-06 | EMMAX | SUG |
| T_17JY | AX-111184272 | 1D | 257366039 | -28.498 | 2.26E-06 | EMMAX | SUG |
| T_17JY | AX-111087251 | 1D | 260322158 | -28.498 | 2.26E-06 | EMMAX | SUG |
| T_17JY | AX-109471332 | 1D | 260404019 | -28.498 | 2.26E-06 | EMMAX | SUG |
| T_17JY | AX-109556460 | 1D | 261225150 | -21.674 | 3.97E-07 | EMMAX | SUG |
| T_17JY | AX-108958481 | 1D | 261880606 | -28.498 | 2.26E-06 | EMMAX | SUG |
| T_17JY | AX-111019612 | 1D | 261894948 | -28.498 | 2.26E-06 | EMMAX | SUG |
| T_17JY | AX-108809138 | 1D | 262033961 | -28.498 | 2.26E-06 | EMMAX | SUG |
| T_17JY | AX-110056705 | 1D | 262034125 | -28.498 | 2.26E-06 | EMMAX | SUG |
| T_17JY | AX-109414051 | 1D | 262037491 | -28.498 | 2.26E-06 | EMMAX | SUG |
| T_17JY | AX-111179253 | 1D | 262249791 | -28.498 | 2.26E-06 | EMMAX | SUG |
| T_17JY | AX-109355923 | 1D | 262471995 | -28.498 | 2.26E-06 | EMMAX | SUG |
| T_17JY | AX-111065611 | 1D | 262522529 | -28.498 | 2.26E-06 | EMMAX | SUG |
| T_17JY | AX-109419591 | 1D | 262568229 | -28.498 | 2.26E-06 | EMMAX | SUG |
| T_17JY | AX-94560123 | 1D | 262706031 | -28.498 | 2.26E-06 | EMMAX | SUG |
| T_17JY | AX-108770408 | 1D | 262709255 | -28.498 | 2.26E-06 | EMMAX | SUG |
| T_17JY | AX-111746820 | 1D | 263073152 | -28.498 | 2.26E-06 | EMMAX | SUG |
| T_17JY | AX-111491952 | 1D | 263168997 | -28.498 | 2.26E-06 | EMMAX | SUG |

| | | | | | | | |
|---|---|---|---|---|---|---|---|
| T_17JY | AX-109095866 | 1D | 263466276 | -28.498 | 2.26E-06 | EMMAX | SUG |
| T_17JY | AX-111646030 | 1D | 263487130 | -28.498 | 2.26E-06 | EMMAX | SUG |
| T_17JY | AX-111664263 | 1D | 263646754 | -28.498 | 2.26E-06 | EMMAX | SUG |
| T_17JY | AX-110897576 | 1D | 263754340 | -28.498 | 2.26E-06 | EMMAX | SUG |
| T_17JY | AX-94438221 | 1D | 263876916 | -28.498 | 2.26E-06 | EMMAX | SUG |
| T_17JY | AX-94541147 | 1D | 263878254 | -28.498 | 2.26E-06 | EMMAX | SUG |
| T_17JY | AX-94709761 | 1D | 263879114 | -28.498 | 2.26E-06 | EMMAX | SUG |
| T_17JY | AX-109506760 | 1D | 263880499 | -28.498 | 2.26E-06 | EMMAX | SUG |
| T_17JY | AX-109592790 | 1D | 264435808 | -28.498 | 2.26E-06 | EMMAX | SUG |
| T_17JY | AX-111628442 | 1D | 264812105 | -28.498 | 2.26E-06 | EMMAX | SUG |
| T_17JY | AX-111581734 | 1D | 265293969 | -28.498 | 2.26E-06 | EMMAX | SUG |
| T_17JY | AX-109396383 | 1D | 265309952 | -28.498 | 2.26E-06 | EMMAX | SUG |
| T_17JY | AX-111514578 | 1D | 266304169 | -28.498 | 2.26E-06 | EMMAX | SUG |
| T_17JY | AX-110532422 | 1D | 266812138 | -28.498 | 2.26E-06 | EMMAX | SUG |
| T_17JY | AX-110926459 | 1D | 267339889 | -28.498 | 2.26E-06 | EMMAX | SUG |
| T_17JY | AX-109953993 | 1D | 267591659 | -28.498 | 2.26E-06 | EMMAX | SUG |
| T_17JY | AX-110419604 | 1D | 267730660 | -28.498 | 2.26E-06 | EMMAX | SUG |
| T_17JY | AX-108981058 | 1D | 267984226 | -28.498 | 2.26E-06 | EMMAX | SUG |
| T_17JY | AX-109901764 | 1D | 268011111 | -28.498 | 2.26E-06 | EMMAX | SUG |
| T_17JY | AX-109986249 | 1D | 268536229 | -28.498 | 2.26E-06 | EMMAX | SUG |
| T_17JY | AX-108870847 | 1D | 268928937 | -28.498 | 2.26E-06 | EMMAX | SUG |
| T_17JY | AX-108809828 | 1D | 268949498 | -28.498 | 2.26E-06 | EMMAX | SUG |
| T_17JY | AX-111138052 | 1D | 268985436 | -28.498 | 2.26E-06 | EMMAX | SUG |
| T_17JY | AX-110510356 | 1D | 269523446 | -28.498 | 2.26E-06 | EMMAX | SUG |
| T_17JY | AX-108747138 | 1D | 269549740 | -28.498 | 2.26E-06 | EMMAX | SUG |
| T_17JY | AX-110854128 | 1D | 269739412 | -28.498 | 2.26E-06 | EMMAX | SUG |
| T_17JY | AX-109935464 | 1D | 270017696 | -28.498 | 2.26E-06 | EMMAX | SUG |
| T_17JY | AX-95255262 | 1D | 270018569 | -28.498 | 2.26E-06 | EMMAX | SUG |
| T_17JY | AX-95200634 | 1D | 270021061 | -28.498 | 2.26E-06 | EMMAX | SUG |
| T_17JY | AX-111991086 | 1D | 270031122 | -28.498 | 2.26E-06 | EMMAX | SUG |
| T_17JY | AX-110549521 | 1D | 270055949 | -28.498 | 2.26E-06 | EMMAX | SUG |
| T_17JY | AX-111439704 | 1D | 270056764 | -28.498 | 2.26E-06 | EMMAX | SUG |
| T_17JY | AX-94749696 | 1D | 270225970 | -28.498 | 2.26E-06 | EMMAX | SUG |
| T_17JY | AX-95117626 | 1D | 270225999 | -28.498 | 2.26E-06 | EMMAX | SUG |
| T_17JY | AX-108923886 | 1D | 270253723 | -28.498 | 2.26E-06 | EMMAX | SUG |
| T_17JY | AX-109532125 | 1D | 270290910 | -28.498 | 2.26E-06 | EMMAX | SUG |
| T_17JY | AX-111661240 | 1D | 270738550 | -28.498 | 2.26E-06 | EMMAX | SUG |
| T_17JY | AX-111392373 | 1D | 271138174 | -28.498 | 2.26E-06 | EMMAX | SUG |
| T_17JY | AX-111555592 | 1D | 271688618 | -19.273 | 3.52E-08 | EMMAX | SIG |
| T_17JY | AX-109484115 | 1D | 271795205 | -28.498 | 2.26E-06 | EMMAX | SUG |
| T_17JY | AX-109806921 | 1D | 271839490 | -28.498 | 2.26E-06 | EMMAX | SUG |
| T_17JY | AX-109268376 | 1D | 271842148 | -28.498 | 2.26E-06 | EMMAX | SUG |
| T_17JY | AX-110895847 | 1D | 271844257 | -28.498 | 2.26E-06 | EMMAX | SUG |
| T_17JY | AX-111992616 | 1D | 272187008 | -28.498 | 2.26E-06 | EMMAX | SUG |

| | | | | | | | | |
|---|---|---|---|---|---|---|---|---|
| T_17JY | AX-95248648 | 1D | 272191130 | -28.498 | | 2.26E-06 | EMMAX | SUG |
| T_17JY | AX-110426070 | 1D | 274211484 | -28.498 | | 2.26E-06 | EMMAX | SUG |
| T_17JY | AX-109760822 | 1D | 274250650 | -28.498 | | 2.26E-06 | EMMAX | SUG |
| T_17JY | AX-111784781 | 1D | 274261764 | -28.498 | | 2.26E-06 | EMMAX | SUG |
| T_17JY | AX-109934942 | 1D | 274262027 | -28.498 | | 2.26E-06 | EMMAX | SUG |
| T_17JY | AX-111037808 | 1D | 274412002 | -28.498 | | 2.26E-06 | EMMAX | SUG |
| T_17JY | AX-109925995 | 1D | 274412281 | -28.498 | | 2.26E-06 | EMMAX | SUG |
| T_17JY | AX-109743825 | 1D | 274468154 | -28.498 | | 2.26E-06 | EMMAX | SUG |
| T_17JY | AX-110426230 | 1D | 274469480 | -28.498 | | 2.26E-06 | EMMAX | SUG |
| T_17JY | AX-111914971 | 1D | 274509037 | -28.498 | | 2.26E-06 | EMMAX | SUG |
| T_17JY | AX-111976657 | 1D | 274558716 | -28.498 | | 2.26E-06 | EMMAX | SUG |
| T_17JY | AX-111561415 | 1D | 274936304 | -28.498 | | 2.26E-06 | EMMAX | SUG |
| T_17JY | AX-111148304 | 1D | 274936340 | -28.498 | | 2.26E-06 | EMMAX | SUG |
| T_17JY | AX-110514071 | 2A | 481809576 | -1.418 | -12.3263 | 5.00E-26 | IIIVmrMLM | SIG |
| T_17JY | AX-109483644 | 2A | 618833837 | -1.487 | | 1.70E-08 | BLUPmrMLM | SIG |
| T_17JY | AX-110171846 | 2A | 635099979 | -3.218 | | 2.96E-07 | EMMAX | SUG |
| T_17JY | AX-110171846 | 2A | 635099979 | 3.227 | | 1.83E-07 | GEMMA | SUG |
| T_17JY | AX-109358833 | 2A | 745677157 | 0.172 | 22.9382 | 1.50E-15 | IIIVmrMLM | SIG |
| T_17JY | AX-109518247 | 2B | 48289705 | 1.356 | | 1.00E-05 | BLUPmrMLM | SUG |
| T_17JY | AX-110941362 | 2B | 69006681 | -0.059 | 6.4087 | 4.42E-06 | IIIVmrMLM | SUG |
| T_17JY | AX-111463303 | 2B | 169865292 | -1.042 | | 1.83E-05 | BLUPmrMLM | SUG |
| T_17JY | AX-111463303 | 2B | 169865292 | 1.513 | | 6.71E-07 | EMMAX | SUG |
| T_17JY | AX-111463303 | 2B | 169865292 | -1.512 | | 4.24E-07 | GEMMA | SUG |
| T_17JY | AX-94815181 | 2B | 324657994 | -1.335 | | 5.59E-16 | IIIVmrMLM | SIG |
| T_17JY | AX-95133585 | 2B | 792768564 | 1.307 | | 6.55E-05 | BLUPmrMLM | SUG |
| T_17JY | AX-111152847 | 2D | 166001639 | 0.671 | 14.4234 | 1.72E-18 | IIIVmrMLM | SIG |
| T_17JY | AX-108747515 | 2D | 286019462 | -3.966 | 22.7638 | 7.11E-95 | IIIVmrMLM | SIG |
| T_17JY | AX-110580566 | 3A | 672183135 | 1.159 | | 4.07E-05 | BLUPmrMLM | SUG |
| T_17JY | AX-111000991 | 3A | 686745027 | 1.534 | | 2.94E-06 | BLUPmrMLM | SUG |
| T_17JY | AX-109312388 | 3A | 690787355 | 0.362 | -1.9821 | 7.28E-04 | IIIVmrMLM | SUG |
| T_17JY | AX-94505389 | 3B | 273249792 | -28.498 | | 2.26E-06 | EMMAX | SUG |
| T_17JY | AX-109971913 | 3B | 390677445 | 0.816 | 6.8098 | 1.61E-12 | IIIVmrMLM | SIG |
| T_17JY | AX-109386317 | 3B | 494167022 | 0.378 | 4.2882 | 1.24E-06 | IIIVmrMLM | SUG |
| T_17JY | AX-95197028 | 3B | 581645670 | -0.790 | | 2.37E-05 | BLUPmrMLM | SUG |
| T_17JY | AX-94459925 | 3D | 220863718 | 1.345 | | 3.37E-05 | BLUPmrMLM | SUG |
| T_17JY | AX-108879375 | 4A | 722842236 | -1.647 | | 1.87E-07 | BLUPmrMLM | SUG |
| T_17JY | AX-109601703 | 4B | 2234842 | -0.837 | 0.8755 | 1.60E-06 | IIIVmrMLM | SUG |
| T_17JY | AX-109280048 | 4B | 10834564 | 0.063 | 17.9114 | 7.30E-10 | IIIVmrMLM | SIG |
| T_17JY | AX-95150109 | 4B | 25482066 | -1.548 | | 1.49E-04 | BLUPmrMLM | SUG |
| T_17JY | AX-110005661 | 4B | 543576848 | -0.466 | 5.9588 | 1.45E-07 | IIIVmrMLM | SUG |
| T_17JY | AX-110377469 | 5A | 650641925 | -35.051 | | 9.31E-09 | EMMAX | SIG |
| T_17JY | AX-111668132 | 5A | 655047712 | -28.498 | | 2.26E-06 | EMMAX | SUG |
| T_17JY | AX-110464156 | 5A | 708525444 | -0.531 | 8.2001 | 6.73E-07 | IIIVmrMLM | SUG |
| T_17JY | AX-111499177 | 5D | 526567959 | -0.152 | 9.4598 | 8.33E-05 | IIIVmrMLM | SUG |

| | | | | | | | | |
|---|---|---|---|---|---|---|---|---|
| T_17JY | AX-95133516 | 6A | 564717997 | 0.493 | 3.9852 | 2.20E-09 | IIIVmrMLM | SIG |
| T_17JY | AX-111536252 | 6B | 55201213 | -0.512 | 10.1943 | 2.42E-07 | IIIVmrMLM | SUG |
| T_17JY | AX-110435804 | 6B | 506842685 | 0.008 | 14.7249 | 5.93E-10 | IIIVmrMLM | SIG |
| T_17JY | AX-110931881 | 6B | 712348786 | 0.123 | 19.0711 | 4.32E-06 | IIIVmrMLM | SUG |
| T_17JY | AX-108893208 | 6D | 3920459 | 0.666 | | 1.43E-04 | BLUPmrMLM | SUG |
| T_17JY | AX-111694520 | 6D | 224751142 | -0.016 | -5.0833 | 1.78E-04 | IIIVmrMLM | SUG |
| T_17JY | AX-94497426 | 6D | 468912303 | 1.079 | | 8.74E-07 | BLUPmrMLM | SUG |
| T_17JY | AX-111537729 | 7A | 139672190 | -0.769 | -4.6453 | 1.02E-11 | IIIVmrMLM | SIG |
| T_17JY | AX-111108137 | 7B | 138472730 | 0.152 | 12.8193 | 1.16E-05 | IIIVmrMLM | SUG |
| T_17JY | AX-111550314 | 7B | 612289854 | 0.391 | 3.9423 | 6.34E-08 | IIIVmrMLM | SIG |
| T_17JY | AX-109941199 | 7D | 108407039 | -28.498 | | 2.26E-06 | EMMAX | SUG |
| T_17SQ | AX-110699024 | 1B | 93853770 | 2.541 | | 1.38E-04 | BLUPmrMLM | SUG |
| T_17SQ | AX-110603097 | 7A | 236312639 | -1.337 | | 7.46E-05 | BLUPmrMLM | SUG |
| T_18AY | AX-109938873 | 1A | 15323295 | 1.354 | | 2.72E-06 | BLUPmrMLM | SUG |
| T_18AY | AX-111506896 | 1A | 151767824 | 1.550 | | 8.71E-06 | BLUPmrMLM | SUG |
| T_18AY | AX-94788541 | 1B | 3272100 | 1.087 | | 1.03E-05 | BLUPmrMLM | SUG |
| T_18AY | AX-111089269 | 1B | 653185396 | 1.601 | | 3.49E-05 | BLUPmrMLM | SUG |
| T_18AY | AX-94668915 | 2D | 317864658 | 0.949 | | 6.40E-05 | BLUPmrMLM | SUG |
| T_18AY | AX-108922737 | 2D | 362777975 | 17.119 | | 4.45E-07 | FarmCPU | SUG |
| T_18AY | AX-110811553 | 2D | 490799621 | 15.922 | | 2.20E-07 | FarmCPU | SUG |
| T_18AY | AX-110421140 | 2D | 536173633 | 17.119 | | 4.45E-07 | FarmCPU | SUG |
| T_18AY | AX-110429039 | 2D | 574389639 | 3.028 | | 7.32E-07 | GEMMA | SUG |
| T_18AY | AX-109276344 | 2D | 574403656 | 2.999 | | 9.44E-07 | GEMMA | SUG |
| T_18AY | AX-111238436 | 3B | 463843723 | -4.389 | | 9.84E-07 | GEMMA | SUG |
| T_18AY | AX-108813535 | 3B | 678549369 | 2.666 | | 3.26E-07 | BLUPmrMLM | SUG |
| T_18AY | AX-109761402 | 3D | 22306875 | 9.839 | | 4.46E-07 | FarmCPU | SUG |
| T_18AY | AX-111728339 | 3D | 22328926 | 9.112 | | 1.23E-06 | FarmCPU | SUG |
| T_18AY | AX-110728609 | 3D | 22936338 | 14.097 | | 1.66E-06 | FarmCPU | SUG |
| T_18AY | AX-94858769 | 3D | 548554219 | 0.919 | | 2.98E-05 | BLUPmrMLM | SUG |
| T_18AY | AX-109225795 | 3D | 569469438 | 17.794 | | 1.13E-06 | FarmCPU | SUG |
| T_18AY | AX-109088681 | 4D | 16362217 | 17.794 | | 1.13E-06 | FarmCPU | SUG |
| T_18AY | AX-110916364 | 4D | 28242114 | 17.794 | | 1.13E-06 | FarmCPU | SUG |
| T_18AY | AX-110967663 | 4D | 65221764 | 17.794 | | 1.13E-06 | FarmCPU | SUG |
| T_18AY | AX-110974080 | 5D | 539222152 | -0.923 | | 2.56E-05 | BLUPmrMLM | SUG |
| T_18AY | AX-94547086 | 6A | 1879928 | 1.505 | | 3.37E-05 | BLUPmrMLM | SUG |
| T_18AY | AX-109334445 | 6B | 101661935 | 2.085 | | 9.28E-06 | BLUPmrMLM | SUG |
| T_18AY | AX-109334445 | 6B | 101661935 | 2.748 | | 3.72E-07 | GEMMA | SUG |
| T_18AY | AX-111835071 | 6B | 163755834 | 1.407 | | 2.00E-05 | BLUPmrMLM | SUG |
| T_18AY | AX-94383040 | 6D | 441619718 | 13.117 | | 2.37E-08 | FarmCPU | SIG |
| T_18AY | AX-108997750 | 6D | 462701843 | -1.334 | | 3.24E-05 | BLUPmrMLM | SUG |
| T_18AY | AX-110920055 | 7A | 13182341 | 1.246 | | 1.11E-04 | BLUPmrMLM | SUG |
| T_18AY | AX-111505923 | 7B | 715541984 | -1.303 | | 2.50E-05 | BLUPmrMLM | SUG |
| T_18AY | AX-109426738 | 7D | 493660501 | 10.731 | | 1.63E-07 | FarmCPU | SUG |
| T_18JY | AX-111506559 | 1B | 1201184 | 0.011 | 4.6614 | 2.00E-04 | IIIVmrMLM | SUG |

| | | | | | | | | |
|---|---|---|---|---|---|---|---|---|
| T_18JY | AX-94395854 | 1B | 465099326 | 15.806 | | 8.76E-08 | FarmCPU | SIG |
| T_18JY | AX-110971985 | 1B | 490045031 | -1.175 | | 1.22E-06 | FarmCPU | SUG |
| T_18JY | AX-109593928 | 1D | 68310644 | -0.298 | 24.2991 | 1.93E-07 | IIIVmrMLM | SUG |
| T_18JY | AX-109626297 | 2A | 104924459 | 0.831 | | 1.53E-04 | BLUPmrMLM | SUG |
| T_18JY | AX-111643003 | 2A | 581168171 | 1.035 | | 1.90E-05 | BLUPmrMLM | SUG |
| T_18JY | AX-94706344 | 2A | 760582990 | 3.092 | | 9.08E-09 | FarmCPU | SIG |
| T_18JY | AX-109941581 | 2B | 117419951 | 0.251 | 8.5756 | 5.40E-05 | IIIVmrMLM | SUG |
| T_18JY | AX-94412862 | 2B | 206111249 | -1.977 | | 4.07E-23 | IIIVmrMLM | SIG |
| T_18JY | AX-110977304 | 2B | 687335451 | -0.926 | -7.4698 | 7.75E-12 | IIIVmrMLM | SIG |
| T_18JY | AX-109755758 | 2D | 156358859 | -1.661 | 0.3088 | 5.74E-16 | IIIVmrMLM | SIG |
| T_18JY | AX-110811553 | 2D | 490799621 | -19.689 | | 3.22E-07 | EMMAX | SUG |
| T_18JY | AX-109883386 | 3A | 21784828 | -0.139 | 5.6031 | 7.77E-05 | IIIVmrMLM | SUG |
| T_18JY | AX-109034917 | 3A | 457482289 | 3.251 | | 1.02E-06 | FarmCPU | SUG |
| T_18JY | AX-94929472 | 3D | 254365479 | 1.593 | | 4.80E-05 | BLUPmrMLM | SUG |
| T_18JY | AX-108896957 | 3D | 612011590 | -0.410 | 21.1429 | 2.05E-06 | IIIVmrMLM | SUG |
| T_18JY | AX-109499259 | 4A | 689435245 | 1.886 | | 1.13E-08 | BLUPmrMLM | SIG |
| T_18JY | AX-108772059 | 4B | 1087568 | -0.978 | | 8.09E-05 | BLUPmrMLM | SUG |
| T_18JY | AX-86177725 | 4B | 519257712 | 0.086 | 13.9185 | 7.41E-09 | IIIVmrMLM | SIG |
| T_18JY | AX-94912623 | 4D | 4353773 | -15.601 | | 1.40E-06 | EMMAX | SUG |
| T_18JY | AX-94880334 | 4D | 15557562 | -16.795 | | 5.41E-12 | FarmCPU | SIG |
| T_18JY | AX-110033925 | 5B | 513812397 | -2.378 | | 4.41E-08 | FarmCPU | SIG |
| T_18JY | AX-95141629 | 5D | 348262228 | -23.877 | | 1.44E-07 | EMMAX | SUG |
| T_18JY | AX-110528706 | 5D | 392073733 | 2.897 | | 1.72E-07 | FarmCPU | SUG |
| T_18JY | AX-94769320 | 5D | 546689801 | 0.693 | 12.2815 | 3.20E-08 | IIIVmrMLM | SIG |
| T_18JY | AX-110469613 | 6A | 594626432 | 1.077 | | 2.86E-05 | BLUPmrMLM | SUG |
| T_18JY | AX-109287410 | 6B | 145342595 | -0.128 | 11.8237 | 2.89E-05 | IIIVmrMLM | SUG |
| T_18JY | AX-111017528 | 6B | 474771011 | 0.605 | -3.6764 | 3.87E-07 | IIIVmrMLM | SUG |
| T_18JY | AX-110955032 | 6B | 686736979 | 0.065 | 4.1635 | 1.03E-08 | IIIVmrMLM | SIG |
| T_18JY | AX-110931881 | 6B | 712348786 | -0.236 | 34.6968 | 2.27E-13 | IIIVmrMLM | SIG |
| T_18JY | AX-110898890 | 6D | 22156695 | 1.380 | | 6.82E-05 | BLUPmrMLM | SUG |
| T_18JY | AX-89690864 | 7A | 8254170 | 0.479 | 17.314 | 2.55E-08 | IIIVmrMLM | SIG |
| T_18JY | AX-110024144 | 7A | 34962296 | -0.089 | 12.3079 | 5.32E-04 | IIIVmrMLM | SUG |
| T_18JY | AX-110906041 | 7A | 161218255 | 1.660 | | 9.94E-08 | FarmCPU | SIG |
| T_18JY | AX-108874812 | 7B | 732599380 | -0.342 | 5.8059 | 1.03E-06 | IIIVmrMLM | SUG |
| T_18JY | AX-109339626 | 7D | 383088654 | 3.003 | 22.5266 | 2.32E-52 | IIIVmrMLM | SIG |
| T_18JY | AX-110042864 | 7D | 519149589 | -2.481 | | 3.97E-08 | BLUPmrMLM | SIG |
| T_18JY | AX-110671304 | 7D | 532629865 | -0.273 | 13.9783 | 2.70E-09 | IIIVmrMLM | SIG |
| T_19XX | AX-86177248 | 1A | 298117391 | 1.764 | | 1.37E-06 | GEMMA | SUG |
| T_19XX | AX-111692565 | 1A | 322091173 | -2.964 | | 1.79E-06 | GEMMA | SUG |
| T_19XX | AX-109942506 | 1A | 565247584 | 0.354 | 2.7053 | 1.10E-12 | IIIVmrMLM | SIG |
| T_19XX | AX-95174049 | 1A | 582842690 | -2.075 | | 1.81E-06 | GEMMA | SUG |
| T_19XX | AX-111638016 | 1A | 582856718 | -2.100 | | 1.52E-06 | GEMMA | SUG |
| T_19XX | AX-110563876 | 1A | 582868976 | -4.096 | | 2.14E-06 | GEMMA | SUG |
| T_19XX | AX-110687192 | 1A | 582872235 | -2.103 | | 1.49E-06 | GEMMA | SUG |

| | | | | | | | | |
|---|---|---|---|---|---|---|---|---|
| T_19XX | AX-110481982 | 1A | 583126295 | -1.875 | | 1.30E-06 | GEMMA | SUG |
| T_19XX | AX-108835837 | 1A | 583172068 | -2.075 | | 1.81E-06 | GEMMA | SUG |
| T_19XX | AX-108826270 | 1A | 583246065 | -2.075 | | 1.81E-06 | GEMMA | SUG |
| T_19XX | AX-111627822 | 1A | 583256261 | -2.091 | | 1.49E-06 | GEMMA | SUG |
| T_19XX | AX-111173346 | 1A | 583287789 | -2.075 | | 1.81E-06 | GEMMA | SUG |
| T_19XX | AX-110463611 | 1A | 583289369 | -2.097 | | 1.28E-06 | GEMMA | SUG |
| T_19XX | AX-109418659 | 1A | 583292411 | -4.091 | | 1.47E-06 | GEMMA | SUG |
| T_19XX | AX-109489638 | 1A | 583294542 | -2.029 | | 1.36E-06 | GEMMA | SUG |
| T_19XX | AX-110414136 | 1A | 583325952 | -2.147 | | 8.08E-07 | GEMMA | SUG |
| T_19XX | AX-109910467 | 1A | 583363808 | -2.159 | | 4.56E-07 | GEMMA | SUG |
| T_19XX | AX-110363533 | 1A | 583453964 | -3.840 | | 9.36E-07 | GEMMA | SUG |
| T_19XX | AX-109576446 | 1A | 583458061 | -1.881 | | 1.52E-06 | GEMMA | SUG |
| T_19XX | AX-109968922 | 1A | 583470268 | -2.062 | | 3.79E-07 | GEMMA | SUG |
| T_19XX | AX-109317151 | 1A | 584070547 | -2.106 | | 1.13E-06 | GEMMA | SUG |
| T_19XX | AX-108790938 | 1A | 592315185 | 0.083 | -3.2914 | 5.98E-04 | IIIVmrMLM | SUG |
| T_19XX | AX-110521285 | 1B | 483878109 | -0.324 | 1.9182 | 3.18E-08 | IIIVmrMLM | SIG |
| T_19XX | AX-94568274 | 1B | 633120777 | -1.572 | | 5.31E-05 | BLUPmrMLM | SUG |
| T_19XX | AX-111132504 | 1D | 271284376 | -4.127 | | 2.17E-06 | GEMMA | SUG |
| T_19XX | AX-109893346 | 1D | 474064100 | 1.397 | 0.351 | 1.43E-17 | IIIVmrMLM | SIG |
| T_19XX | AX-109527900 | 2A | 83989988 | -0.766 | | 1.37E-04 | BLUPmrMLM | SUG |
| T_19XX | AX-111031818 | 2A | 136838918 | 0.919 | 0.1737 | 3.29E-07 | IIIVmrMLM | SUG |
| T_19XX | AX-109496621 | 2A | 148591856 | -0.179 | 6.0987 | 1.02E-06 | IIIVmrMLM | SUG |
| T_19XX | AX-109832463 | 2A | 358906450 | -0.078 | -2.596 | 1.01E-07 | IIIVmrMLM | SIG |
| T_19XX | AX-109398144 | 2A | 448832136 | 0.572 | 8.5 | 2.61E-12 | IIIVmrMLM | SIG |
| T_19XX | AX-109353419 | 2A | 618980931 | -0.347 | -5.7794 | 1.89E-13 | IIIVmrMLM | SIG |
| T_19XX | AX-108956123 | 2A | 691378369 | 0.337 | 7.2581 | 3.31E-07 | IIIVmrMLM | SUG |
| T_19XX | AX-111600569 | 2A | 735693118 | 0.397 | -2.3429 | 2.46E-17 | IIIVmrMLM | SIG |
| T_19XX | AX-110500540 | 2B | 137979605 | 1.626 | | 6.09E-06 | BLUPmrMLM | SUG |
| T_19XX | AX-109656267 | 2B | 422878071 | -1.548 | | 8.98E-05 | BLUPmrMLM | SUG |
| T_19XX | AX-95234063 | 2B | 449841621 | 0.481 | 2.3561 | 4.72E-06 | IIIVmrMLM | SUG |
| T_19XX | AX-108937517 | 2B | 708865726 | 0.009 | 3.2959 | 5.45E-05 | IIIVmrMLM | SUG |
| T_19XX | AX-94520912 | 2D | 218378125 | 2.643 | | 1.61E-06 | FarmCPU | SUG |
| T_19XX | AX-110965386 | 2D | 609954770 | 0.157 | 4.3142 | 7.10E-08 | IIIVmrMLM | SIG |
| T_19XX | AX-110731790 | 3A | 215110746 | 5.383 | | 1.57E-09 | GEMMA | SIG |
| T_19XX | AX-109418776 | 3A | 514522678 | -1.050 | | 6.76E-06 | BLUPmrMLM | SUG |
| T_19XX | AX-94488815 | 3A | 701140704 | 2.817 | | 7.82E-07 | GEMMA | SUG |
| T_19XX | AX-94795166 | 3B | 78598490 | -3.443 | | 9.09E-10 | GEMMA | SIG |
| T_19XX | AX-110471408 | 3B | 124812293 | -0.781 | -5.9372 | 2.10E-11 | IIIVmrMLM | SIG |
| T_19XX | AX-109279712 | 3B | 153953795 | 0.343 | 3.8203 | 7.40E-06 | IIIVmrMLM | SUG |
| T_19XX | AX-109883053 | 3B | 651275582 | -0.060 | -2.3604 | 7.53E-05 | IIIVmrMLM | SUG |
| T_19XX | AX-108917304 | 3B | 754446743 | 4.492 | | 2.31E-07 | GEMMA | SUG |
| T_19XX | AX-108742030 | 3B | 757468725 | 0.888 | | 1.95E-04 | BLUPmrMLM | SUG |
| T_19XX | AX-111594823 | 3D | 463540916 | -1.800 | | 1.32E-06 | BLUPmrMLM | SUG |
| T_19XX | AX-94578015 | 4A | 17979191 | -1.748 | | 3.34E-30 | IIIVmrMLM | SIG |

| | | | | | | | | |
|---|---|---|---|---|---|---|---|---|
| T_19XX | AX-111045478 | 4A | 28544433 | 0.031 | -4.6419 | 7.26E-06 | IIIVmrMLM | SUG |
| T_19XX | AX-94910710 | 4A | 601366543 | 5.048 | | 1.84E-06 | FarmCPU | SUG |
| T_19XX | AX-94635019 | 4A | 601366905 | 5.048 | | 1.84E-06 | FarmCPU | SUG |
| T_19XX | AX-108956803 | 4A | 662961097 | 0.223 | 3.4751 | 2.50E-05 | IIIVmrMLM | SUG |
| T_19XX | AX-110940812 | 4B | 3853762 | 2.533 | | 1.99E-06 | FarmCPU | SUG |
| T_19XX | AX-108812567 | 4B | 3886625 | 4.630 | | 5.84E-07 | FarmCPU | SUG |
| T_19XX | AX-86178836 | 4B | 6084593 | 2.573 | | 1.66E-06 | FarmCPU | SUG |
| T_19XX | AX-94724471 | 4B | 6084778 | 2.044 | | 4.11E-07 | FarmCPU | SUG |
| T_19XX | AX-95630739 | 4B | 6149863 | 5.048 | | 1.84E-06 | FarmCPU | SUG |
| T_19XX | AX-109145302 | 4D | 17208503 | 4.062 | | 3.12E-07 | FarmCPU | SUG |
| T_19XX | AX-108901371 | 4D | 401635796 | 7.008 | | 1.23E-06 | GEMMA | SUG |
| T_19XX | AX-109920270 | 5A | 17377939 | -0.003 | 1.7562 | 2.68E-05 | IIIVmrMLM | SUG |
| T_19XX | AX-94714888 | 5A | 37619362 | -1.338 | | 4.28E-19 | IIIVmrMLM | SIG |
| T_19XX | AX-108739557 | 5B | 214182236 | -2.477 | | 1.84E-06 | GEMMA | SUG |
| T_19XX | AX-110116446 | 5B | 239437975 | -2.650 | | 4.34E-07 | GEMMA | SUG |
| T_19XX | AX-110616657 | 5B | 239438625 | -2.560 | | 1.40E-06 | GEMMA | SUG |
| T_19XX | AX-111187755 | 5B | 239438747 | -2.690 | | 4.55E-07 | GEMMA | SUG |
| T_19XX | AX-111069506 | 5B | 243768703 | -2.644 | | 4.74E-07 | GEMMA | SUG |
| T_19XX | AX-110115020 | 5B | 304656472 | -2.433 | | 2.14E-06 | GEMMA | SUG |
| T_19XX | AX-110085797 | 5B | 498876732 | -2.479 | | 7.99E-07 | GEMMA | SUG |
| T_19XX | AX-111453863 | 5B | 499383561 | -2.385 | | 1.62E-06 | GEMMA | SUG |
| T_19XX | AX-109559492 | 5B | 501960566 | -2.712 | | 9.04E-08 | GEMMA | SIG |
| T_19XX | AX-110052990 | 5B | 501960790 | -2.605 | | 2.32E-07 | GEMMA | SUG |
| T_19XX | AX-94429757 | 5B | 502377400 | -4.497 | | 3.30E-07 | GEMMA | SUG |
| T_19XX | AX-94456306 | 5B | 503093380 | -2.639 | | 8.68E-08 | GEMMA | SIG |
| T_19XX | AX-110083362 | 5B | 504775261 | -2.304 | | 2.14E-06 | GEMMA | SUG |
| T_19XX | AX-94737861 | 5B | 505806528 | -2.458 | | 7.30E-07 | GEMMA | SUG |
| T_19XX | AX-111036857 | 5B | 505808757 | -2.429 | | 1.61E-06 | GEMMA | SUG |
| T_19XX | AX-109909239 | 5B | 512987171 | 0.234 | 2.7081 | 5.53E-06 | IIIVmrMLM | SUG |
| T_19XX | AX-111023077 | 5B | 514047363 | -3.041 | | 3.98E-08 | GEMMA | SIG |
| T_19XX | AX-111469168 | 5B | 517202211 | -2.513 | | 1.39E-06 | GEMMA | SUG |
| T_19XX | AX-108868151 | 5B | 517585608 | -3.031 | | 2.96E-08 | GEMMA | SIG |
| T_19XX | AX-109406233 | 5B | 521141085 | -2.990 | | 2.12E-08 | GEMMA | SIG |
| T_19XX | AX-94612811 | 5B | 522082107 | 1.925 | | 5.86E-05 | BLUPmrMLM | SUG |
| T_19XX | AX-111723333 | 5B | 681780729 | 0.312 | 3.5502 | 3.59E-06 | IIIVmrMLM | SUG |
| T_19XX | AX-110912272 | 5B | 707131823 | -0.052 | 9.8023 | 1.17E-05 | IIIVmrMLM | SUG |
| T_19XX | AX-111950468 | 5D | 23053957 | 8.486 | 10.5348 | ######## | IIIVmrMLM | SIG |
| T_19XX | AX-110454851 | 5D | 414749247 | -3.013 | | 7.49E-09 | GEMMA | SIG |
| T_19XX | AX-111161727 | 5D | 423840514 | -4.451 | | 2.16E-06 | GEMMA | SUG |
| T_19XX | AX-109871224 | 6A | 18732298 | 1.434 | | 1.33E-07 | BLUPmrMLM | SUG |
| T_19XX | AX-110660489 | 6A | 63589264 | -0.175 | -2.3917 | 1.10E-05 | IIIVmrMLM | SUG |
| T_19XX | AX-110966462 | 6A | 595608240 | -0.113 | -2.4482 | 3.46E-09 | IIIVmrMLM | SIG |
| T_19XX | AX-110449653 | 6B | 363315600 | -0.200 | -4.3414 | 2.40E-12 | IIIVmrMLM | SIG |
| T_19XX | AX-95094378 | 6B | 679242840 | -4.570 | | 4.55E-07 | GEMMA | SUG |

| | | | | | | | | |
|---|---|---|---|---|---|---|---|---|
| T_19XX | AX-111072520 | 7A | 40434342 | 6.464 | | 1.88E-06 | GEMMA | SUG |
| T_19XX | AX-111123810 | 7A | 533571893 | -3.144 | | 5.27E-07 | GEMMA | SUG |
| T_19XX | AX-110451142 | 7A | 634931164 | -0.219 | 11.5541 | 3.92E-10 | IIIVmrMLM | SIG |
| T_19XX | AX-110976332 | 7A | 636894164 | -0.124 | 3.7675 | 1.80E-05 | IIIVmrMLM | SUG |
| T_19XX | AX-109357238 | 7A | 693956603 | 0.837 | | 2.50E-05 | BLUPmrMLM | SUG |
| T_19XX | AX-111701625 | 7B | 155790987 | 0.955 | | 3.74E-05 | BLUPmrMLM | SUG |
| T_19XX | AX-111546920 | 7B | 627954523 | -0.093 | 3.8432 | 2.51E-06 | IIIVmrMLM | SUG |
| T_19XX | AX-111675649 | 7B | 628783891 | 0.014 | 10.2614 | 4.28E-06 | IIIVmrMLM | SUG |
| T_19XX | AX-110987749 | 7B | 641149328 | 0.180 | 15.6786 | 2.68E-09 | IIIVmrMLM | SIG |
| T_BLUP | AX-94621343 | 1B | 158545260 | -0.033 | | 2.16E-06 | BLUPmrMLM | SUG |
| T_BLUP | AX-108871949 | 2A | 75143860 | 0.007 | -0.2098 | 8.60E-12 | IIIVmrMLM | SIG |
| T_BLUP | AX-109923782 | 2A | 82180795 | 0.005 | -0.073 | 7.03E-06 | IIIVmrMLM | SUG |
| T_BLUP | AX-111125305 | 2A | 675093713 | -0.029 | -0.0068 | 7.32E-12 | IIIVmrMLM | SIG |
| T_BLUP | AX-94924770 | 2A | 735564525 | -0.013 | 0.0999 | 9.26E-13 | IIIVmrMLM | SIG |
| T_BLUP | AX-111758213 | 2B | 245146722 | -0.024 | | 1.49E-05 | BLUPmrMLM | SUG |
| T_BLUP | AX-109841923 | 2B | 573752048 | -0.002 | -0.0723 | 2.07E-08 | IIIVmrMLM | SIG |
| T_BLUP | AX-109824281 | 2B | 634649436 | 0.010 | 0.0923 | 4.39E-07 | IIIVmrMLM | SUG |
| T_BLUP | AX-110952281 | 2B | 765280430 | 0.004 | 0.3783 | 4.73E-11 | IIIVmrMLM | SIG |
| T_BLUP | AX-110856438 | 2D | 16730289 | 0.007 | 0.2099 | 7.19E-09 | IIIVmrMLM | SIG |
| T_BLUP | AX-110358286 | 2D | 450730539 | 0.040 | | 1.96E-05 | BLUPmrMLM | SUG |
| T_BLUP | AX-109463920 | 2D | 608771854 | 0.006 | 0.0733 | 1.07E-06 | IIIVmrMLM | SUG |
| T_BLUP | AX-108921723 | 2D | 639633193 | 0.008 | -0.0703 | 4.32E-10 | IIIVmrMLM | SIG |
| T_BLUP | AX-110942379 | 3A | 12419154 | 0.034 | | 6.36E-18 | IIIVmrMLM | SIG |
| T_BLUP | AX-111833519 | 3B | 239031838 | -0.016 | 0.0457 | 5.76E-07 | IIIVmrMLM | SUG |
| T_BLUP | AX-110362195 | 3B | 823527068 | -0.013 | 0.0961 | 3.84E-09 | IIIVmrMLM | SIG |
| T_BLUP | AX-110375029 | 4A | 59681391 | 0.024 | 0.0892 | 2.07E-10 | IIIVmrMLM | SIG |
| T_BLUP | AX-109356483 | 4A | 127432909 | 0.009 | -0.0943 | 5.67E-09 | IIIVmrMLM | SIG |
| T_BLUP | AX-111017431 | 4A | 597898318 | -0.025 | 0.0059 | 1.05E-07 | IIIVmrMLM | SIG |
| T_BLUP | AX-94923774 | 4A | 733460243 | -0.023 | | 1.01E-04 | BLUPmrMLM | SUG |
| T_BLUP | AX-94923774 | 4A | 733460243 | 0.017 | 0.0085 | 4.19E-04 | IIIVmrMLM | SUG |
| T_BLUP | AX-111022167 | 4B | 605201873 | -0.004 | 0.0428 | 5.31E-06 | IIIVmrMLM | SUG |
| T_BLUP | AX-111941553 | 4D | 221830526 | 0.030 | | 1.81E-14 | IIIVmrMLM | SIG |
| T_BLUP | AX-95075181 | 5A | 14266468 | -0.025 | | 1.41E-04 | BLUPmrMLM | SUG |
| T_BLUP | AX-94818122 | 5A | 122997638 | -0.023 | | 4.79E-09 | IIIVmrMLM | SIG |
| T_BLUP | AX-109840368 | 5B | 699599828 | 0.008 | -0.0764 | 1.46E-06 | IIIVmrMLM | SUG |
| T_BLUP | AX-111497901 | 6A | 36988422 | -0.010 | 0.1036 | 6.57E-05 | IIIVmrMLM | SUG |
| T_BLUP | AX-108799643 | 6B | 369957174 | 0.006 | -0.1364 | 7.67E-05 | IIIVmrMLM | SUG |
| T_BLUP | AX-110926064 | 6B | 491951337 | -0.008 | -0.0796 | 9.33E-08 | IIIVmrMLM | SIG |
| T_BLUP | AX-112290431 | 6B | 703287322 | -0.002 | -0.2406 | 4.75E-09 | IIIVmrMLM | SIG |
| T_BLUP | AX-111761852 | 6D | 124124843 | -0.030 | 0.0215 | 2.32E-13 | IIIVmrMLM | SIG |
| T_BLUP | AX-94641271 | 6D | 333804220 | 0.032 | | 3.70E-05 | BLUPmrMLM | SUG |
| T_BLUP | AX-109339986 | 6D | 461334374 | 0.053 | | 6.35E-06 | BLUPmrMLM | SUG |
| T_BLUP | AX-110098562 | 7A | 617694655 | -0.023 | | 1.43E-04 | BLUPmrMLM | SUG |
| T_BLUP | AX-111613533 | 7A | 697401406 | -0.019 | | 2.29E-05 | BLUPmrMLM | SUG |

| | | | | | | | | |
|---|---|---|---|---|---|---|---|---|
| T_BLUP | AX-109270483 | 7B | 681578739 | 0.003 | 0.2086 | 5.00E-05 | IIIVmrMLM | SUG |
| T_BLUP | AX-111455256 | 7B | 740013976 | 0.022 | | 9.44E-05 | BLUPmrMLM | SUG |
| T_BLUP | AX-109873875 | 7D | 115493089 | -0.002 | 0.0983 | 1.86E-05 | IIIVmrMLM | SUG |
| T_BLUP | AX-109933656 | 7D | 612929882 | -0.011 | -0.0355 | 2.97E-07 | IIIVmrMLM | SUG |

T: active filling period.

**Supplementary Table S17.** All QTNs identified as associated with average grain filling rate in wheat.

| Trait | Marker | Chr | Position | Add.effect | Dom.effect | P-value | Method | Sig. |
|---|---|---|---|---|---|---|---|---|
| V_17AY | AX-111031083 | 1B | 50023282 | 0.0055 | -0.3237 | 4.08E-07 | IIIVmrMLM | SUG |
| V_17AY | AX-109519577 | 1B | 98910393 | 0.0742 | | 4.79E-07 | BLUPmrMLM | SUG |
| V_17AY | AX-109109867 | 1B | 110950982 | -0.0135 | -0.4117 | 4.08E-08 | IIIVmrMLM | SIG |
| V_17AY | AX-111540876 | 1B | 543473980 | 0.0161 | 0.1345 | 3.31E-13 | IIIVmrMLM | SIG |
| V_17AY | AX-109433340 | 1B | 623738154 | 0.0140 | 0.1485 | 2.09E-05 | IIIVmrMLM | SUG |
| V_17AY | AX-108788117 | 1B | 654531498 | -0.0945 | | 7.82E-08 | BLUPmrMLM | SIG |
| V_17AY | AX-94966722 | 1B | 670557461 | -0.1077 | | 4.11E-05 | BLUPmrMLM | SUG |
| V_17AY | AX-111056687 | 1B | 673412687 | 0.0539 | | 3.99E-05 | BLUPmrMLM | SUG |
| V_17AY | AX-110047281 | 1D | 595098 | -2.0441 | | 1.16E-08 | EMMAX | SIG |
| V_17AY | AX-110392337 | 1D | 73679336 | -0.4989 | | 2.06E-09 | EMMAX | SIG |
| V_17AY | AX-109495041 | 2A | 144658876 | -0.0196 | -0.1825 | 8.36E-08 | IIIVmrMLM | SIG |
| V_17AY | AX-109644557 | 2A | 602699209 | -0.0355 | -0.1802 | 3.70E-11 | IIIVmrMLM | SIG |
| V_17AY | AX-109273856 | 2A | 771343034 | -0.0502 | | 1.02E-05 | BLUPmrMLM | SUG |
| V_17AY | AX-94971846 | 2B | 30499258 | 0.0234 | 0.1441 | 1.62E-18 | IIIVmrMLM | SIG |
| V_17AY | AX-111559203 | 2B | 154588905 | 0.0844 | -0.3446 | 1.40E-26 | IIIVmrMLM | SIG |
| V_17AY | AX-109974976 | 2B | 421251405 | 0.0633 | | 3.42E-05 | BLUPmrMLM | SUG |
| V_17AY | AX-109499517 | 2B | 427082691 | -0.0231 | 0.493 | 4.74E-09 | IIIVmrMLM | SIG |
| V_17AY | AX-109426196 | 2B | 537338926 | -0.0206 | -0.1867 | 1.07E-15 | IIIVmrMLM | SIG |
| V_17AY | AX-94618830 | 2B | 546979309 | -2.0441 | | 1.16E-08 | EMMAX | SIG |
| V_17AY | AX-109919029 | 2B | 549204515 | 0.3378 | | 3.89E-07 | GEMMA | SUG |
| V_17AY | AX-109412939 | 2B | 550847190 | -0.3983 | | 5.09E-07 | EMMAX | SUG |
| V_17AY | AX-109412939 | 2B | 550847190 | 0.4038 | | 1.35E-08 | GEMMA | SIG |
| V_17AY | AX-111058323 | 2B | 551398006 | 0.2713 | | 2.16E-06 | GEMMA | SUG |
| V_17AY | AX-111714730 | 2B | 551445878 | -0.4004 | | 2.16E-07 | EMMAX | SUG |
| V_17AY | AX-111714730 | 2B | 551445878 | 0.4046 | | 6.54E-09 | GEMMA | SIG |
| V_17AY | AX-111065061 | 2B | 743546442 | 0.0344 | -0.0133 | 8.25E-05 | IIIVmrMLM | SUG |
| V_17AY | AX-110098913 | 2D | 317585511 | -2.0441 | | 1.16E-08 | EMMAX | SIG |
| V_17AY | AX-108787002 | 2D | 326202355 | -2.0441 | | 1.16E-08 | EMMAX | SIG |
| V_17AY | AX-109926918 | 2D | 494312828 | 0.0543 | | 1.71E-04 | BLUPmrMLM | SUG |
| V_17AY | AX-109328166 | 2D | 508469235 | -2.0441 | | 1.16E-08 | EMMAX | SIG |
| V_17AY | AX-108759657 | 2D | 618658081 | -0.0685 | | 5.33E-06 | BLUPmrMLM | SUG |
| V_17AY | AX-111255774 | 3A | 1025920 | 0.0011 | 0.1423 | 3.23E-05 | IIIVmrMLM | SUG |
| V_17AY | AX-111628091 | 3A | 12785355 | 0.1782 | | 2.00E-07 | GEMMA | SUG |
| V_17AY | AX-111265000 | 3A | 12928360 | -0.0068 | -0.1802 | 5.43E-08 | IIIVmrMLM | SIG |
| V_17AY | AX-108845007 | 3A | 12997270 | -0.1837 | | 1.87E-06 | EMMAX | SUG |
| V_17AY | AX-108845007 | 3A | 12997270 | 0.1865 | | 3.01E-08 | GEMMA | SIG |
| V_17AY | AX-95143496 | 3A | 12997719 | 0.1834 | | 4.01E-08 | GEMMA | SIG |
| V_17AY | AX-110913445 | 3A | 12998870 | -0.1837 | | 1.87E-06 | EMMAX | SUG |
| V_17AY | AX-110913445 | 3A | 12998870 | 0.1865 | | 3.01E-08 | GEMMA | SIG |
| V_17AY | AX-89450584 | 3A | 12999035 | -0.1837 | | 1.87E-06 | EMMAX | SUG |
| V_17AY | AX-89450584 | 3A | 12999035 | 0.1865 | | 3.01E-08 | GEMMA | SIG |

| | | | | | | | | |
|---|---|---|---|---|---|---|---|---|
| V_17AY | AX-109441093 | 3A | 13001330 | 0.1646 | | 2.38E-07 | GEMMA | SUG |
| V_17AY | AX-111582837 | 3A | 13009071 | 0.1472 | | 1.39E-06 | GEMMA | SUG |
| V_17AY | AX-111460884 | 3A | 13025627 | 0.1754 | | 5.59E-07 | GEMMA | SUG |
| V_17AY | AX-110955079 | 3A | 13025941 | 0.1754 | | 5.59E-07 | GEMMA | SUG |
| V_17AY | AX-86184828 | 3A | 13038241 | -0.1860 | | 1.63E-06 | EMMAX | SUG |
| V_17AY | AX-86184828 | 3A | 13038241 | 0.1874 | | 3.75E-08 | GEMMA | SIG |
| V_17AY | AX-86166665 | 3A | 13038544 | 0.1798 | | 1.53E-07 | GEMMA | SUG |
| V_17AY | AX-86184778 | 3A | 13038599 | 0.1856 | | 7.86E-08 | GEMMA | SIG |
| V_17AY | AX-111589685 | 3A | 13042584 | 0.1904 | | 4.60E-08 | GEMMA | SIG |
| V_17AY | AX-108775363 | 3A | 13044766 | 0.1812 | | 1.01E-07 | GEMMA | SIG |
| V_17AY | AX-111065185 | 3A | 13078767 | 0.1675 | | 2.05E-07 | GEMMA | SUG |
| V_17AY | AX-110909396 | 3A | 13081297 | 0.3740 | | 7.66E-08 | GEMMA | SIG |
| V_17AY | AX-109346552 | 3A | 13190883 | -0.1877 | | 8.86E-07 | EMMAX | SUG |
| V_17AY | AX-109346552 | 3A | 13190883 | 0.1913 | | 1.01E-08 | GEMMA | SIG |
| V_17AY | AX-109878083 | 3A | 13269613 | 0.1210 | | 1.70E-06 | GEMMA | SUG |
| V_17AY | AX-109992464 | 3A | 13281571 | 0.1198 | | 8.84E-07 | GEMMA | SUG |
| V_17AY | AX-110417013 | 3A | 13317686 | 0.1157 | | 2.16E-06 | GEMMA | SUG |
| V_17AY | AX-109992759 | 3A | 13336538 | 0.1184 | | 2.17E-06 | GEMMA | SUG |
| V_17AY | AX-108729182 | 3A | 13345844 | 0.1198 | | 1.44E-06 | GEMMA | SUG |
| V_17AY | AX-110936651 | 3A | 13351271 | 0.1234 | | 9.32E-07 | GEMMA | SUG |
| V_17AY | AX-109890717 | 3A | 13353457 | 0.1197 | | 1.52E-06 | GEMMA | SUG |
| V_17AY | AX-109322880 | 3A | 13358458 | 0.1219 | | 1.18E-06 | GEMMA | SUG |
| V_17AY | AX-110516835 | 3A | 13359361 | 0.1171 | | 2.31E-06 | GEMMA | SUG |
| V_17AY | AX-111217068 | 3A | 13365393 | 0.2534 | | 5.35E-07 | GEMMA | SUG |
| V_17AY | AX-111450027 | 3A | 13367521 | 0.1157 | | 1.23E-06 | GEMMA | SUG |
| V_17AY | AX-111499928 | 3A | 13394372 | 0.1412 | | 5.95E-08 | GEMMA | SIG |
| V_17AY | AX-108733909 | 3A | 13394434 | 0.1143 | | 1.09E-06 | GEMMA | SUG |
| V_17AY | AX-109465246 | 3A | 13399366 | 0.1373 | | 1.17E-07 | GEMMA | SUG |
| V_17AY | AX-108951347 | 3A | 13399645 | 0.1412 | | 5.95E-08 | GEMMA | SIG |
| V_17AY | AX-95023272 | 3A | 13405077 | 0.0039 | 0.1901 | 2.49E-06 | IIIVmrMLM | SUG |
| V_17AY | AX-95098281 | 3A | 13405093 | 0.2736 | | 1.49E-07 | GEMMA | SUG |
| V_17AY | AX-110649779 | 3A | 13495516 | 0.2639 | | 1.14E-07 | GEMMA | SIG |
| V_17AY | AX-94523509 | 3A | 13496628 | 0.1381 | | 1.29E-07 | GEMMA | SUG |
| V_17AY | AX-111483558 | 3A | 13501374 | 0.2649 | | 2.33E-07 | GEMMA | SUG |
| V_17AY | AX-111730267 | 3A | 13503787 | 0.1381 | | 1.29E-07 | GEMMA | SUG |
| V_17AY | AX-89339551 | 3A | 13504175 | 0.1381 | | 1.29E-07 | GEMMA | SUG |
| V_17AY | AX-110535074 | 3A | 13511474 | 0.2773 | | 1.86E-07 | GEMMA | SUG |
| V_17AY | AX-110584359 | 3A | 13516636 | 0.2885 | | 3.27E-08 | GEMMA | SIG |
| V_17AY | AX-111062701 | 3A | 13517880 | 0.2485 | | 4.49E-07 | GEMMA | SUG |
| V_17AY | AX-110963955 | 3A | 13557708 | 0.2518 | | 9.34E-07 | GEMMA | SUG |
| V_17AY | AX-109977764 | 3A | 13559291 | 0.2489 | | 1.13E-06 | GEMMA | SUG |
| V_17AY | AX-94583236 | 3A | 13739466 | 0.1827 | | 3.77E-08 | GEMMA | SIG |
| V_17AY | AX-109362869 | 3A | 13739946 | -0.3697 | | 2.18E-06 | EMMAX | SUG |
| V_17AY | AX-109362869 | 3A | 13739946 | 0.3813 | | 2.05E-08 | GEMMA | SIG |

| | | | | | | | | |
|---|---|---|---|---|---|---|---|---|
| V_17AY | AX-109975558 | 3A | 13788160 | 0.2434 | | 3.56E-08 | GEMMA | SIG |
| V_17AY | AX-109285709 | 3A | 13901742 | 0.4005 | | 1.21E-06 | GEMMA | SUG |
| V_17AY | AX-111490647 | 3A | 13901791 | 0.4131 | | 2.82E-07 | GEMMA | SUG |
| V_17AY | AX-109839370 | 3A | 13971877 | 0.2074 | | 6.90E-07 | GEMMA | SUG |
| V_17AY | AX-109281788 | 3A | 14012613 | 0.1968 | | 1.31E-06 | GEMMA | SUG |
| V_17AY | AX-110415319 | 3A | 14013078 | 0.2074 | | 6.90E-07 | GEMMA | SUG |
| V_17AY | AX-108725274 | 3A | 14014210 | 0.2181 | | 2.38E-07 | GEMMA | SUG |
| V_17AY | AX-111714168 | 3A | 14042236 | 0.1968 | | 1.31E-06 | GEMMA | SUG |
| V_17AY | AX-111502587 | 3A | 14042372 | 0.1968 | | 1.31E-06 | GEMMA | SUG |
| V_17AY | AX-110470380 | 3A | 14042450 | 0.1968 | | 1.31E-06 | GEMMA | SUG |
| V_17AY | AX-111543947 | 3A | 114792980 | -0.1101 | | 6.33E-05 | BLUPmrMLM | SUG |
| V_17AY | AX-111543947 | 3A | 114792980 | 0.0397 | | 2.60E-07 | IIIVmrMLM | SUG |
| V_17AY | AX-95023926 | 3A | 653243442 | 0.0745 | | 6.57E-21 | IIIVmrMLM | SIG |
| V_17AY | AX-110391933 | 3A | 738905274 | -0.0108 | 0.3913 | 2.94E-07 | IIIVmrMLM | SUG |
| V_17AY | AX-95110648 | 3A | 739559346 | -0.9080 | | 9.38E-07 | EMMAX | SUG |
| V_17AY | AX-108904779 | 3B | 398377287 | 0.0403 | -0.0706 | 3.74E-08 | IIIVmrMLM | SIG |
| V_17AY | AX-111236342 | 3B | 439703056 | -0.0848 | -0.0048 | 2.76E-20 | IIIVmrMLM | SIG |
| V_17AY | AX-110049347 | 3B | 493392497 | 0.0572 | | 1.65E-06 | BLUPmrMLM | SUG |
| V_17AY | AX-109481030 | 3B | 499814950 | -0.0601 | 0.6421 | 2.11E-18 | IIIVmrMLM | SIG |
| V_17AY | AX-109427069 | 3B | 584577513 | -0.0005 | 0.1771 | 6.09E-05 | IIIVmrMLM | SUG |
| V_17AY | AX-108873971 | 3B | 798550115 | -2.0441 | | 1.16E-08 | EMMAX | SIG |
| V_17AY | AX-94662210 | 3B | 812572565 | -2.0441 | | 1.16E-08 | EMMAX | SIG |
| V_17AY | AX-94487727 | 3B | 817009854 | -1.0478 | | 6.37E-09 | EMMAX | SIG |
| V_17AY | AX-95254443 | 3D | 418396767 | -2.0441 | | 1.16E-08 | EMMAX | SIG |
| V_17AY | AX-109450588 | 3D | 445486559 | -2.0441 | | 1.16E-08 | EMMAX | SIG |
| V_17AY | AX-110392272 | 3D | 594847340 | -1.2776 | | 9.49E-07 | EMMAX | SUG |
| V_17AY | AX-95232627 | 3D | 594891831 | -1.2776 | | 9.49E-07 | EMMAX | SUG |
| V_17AY | AX-94871493 | 3D | 594891928 | -1.2776 | | 9.49E-07 | EMMAX | SUG |
| V_17AY | AX-95238470 | 3D | 594891932 | -1.2776 | | 9.49E-07 | EMMAX | SUG |
| V_17AY | AX-94928843 | 3D | 595066148 | -1.2776 | | 9.49E-07 | EMMAX | SUG |
| V_17AY | AX-94856639 | 3D | 595066164 | -1.2776 | | 9.49E-07 | EMMAX | SUG |
| V_17AY | AX-109834003 | 3D | 595115084 | -1.2776 | | 9.49E-07 | EMMAX | SUG |
| V_17AY | AX-111697259 | 3D | 595187135 | -1.2776 | | 9.49E-07 | EMMAX | SUG |
| V_17AY | AX-94603168 | 3D | 595271058 | -0.5868 | | 1.99E-06 | EMMAX | SUG |
| V_17AY | AX-94924379 | 3D | 595271201 | -1.2776 | | 9.49E-07 | EMMAX | SUG |
| V_17AY | AX-95180413 | 3D | 595271202 | -1.2776 | | 9.49E-07 | EMMAX | SUG |
| V_17AY | AX-109008383 | 3D | 595273845 | -1.2776 | | 9.49E-07 | EMMAX | SUG |
| V_17AY | AX-109394070 | 3D | 595279092 | -0.5868 | | 1.99E-06 | EMMAX | SUG |
| V_17AY | AX-94874313 | 3D | 598011380 | -2.0441 | | 1.16E-08 | EMMAX | SIG |
| V_17AY | AX-94569728 | 4B | 518641201 | -2.0441 | | 1.16E-08 | EMMAX | SIG |
| V_17AY | AX-108790830 | 4D | 6672205 | -2.0441 | | 1.16E-08 | EMMAX | SIG |
| V_17AY | AX-109498125 | 4D | 168900845 | -2.0441 | | 1.16E-08 | EMMAX | SIG |
| V_17AY | AX-108756835 | 4D | 323706230 | -2.0441 | | 1.16E-08 | EMMAX | SIG |
| V_17AY | AX-108994164 | 4D | 327427960 | -2.0441 | | 1.16E-08 | EMMAX | SIG |

| | | | | | | | | |
|---|---|---|---|---|---|---|---|---|
| V_17AY | AX-111745748 | 4D | 340296689 | -2.0441 | | 1.16E-08 | EMMAX | SIG |
| V_17AY | AX-109534277 | 4D | 378260308 | -2.0441 | | 1.16E-08 | EMMAX | SIG |
| V_17AY | AX-94967011 | 5A | 70498269 | -0.0526 | | 1.37E-11 | IIIVmrMLM | SIG |
| V_17AY | AX-110167972 | 5A | 396754324 | -2.0441 | | 1.16E-08 | EMMAX | SIG |
| V_17AY | AX-110710951 | 5A | 472547282 | -0.5112 | | 3.26E-08 | EMMAX | SIG |
| V_17AY | AX-110962408 | 5A | 476135701 | -0.0324 | -0.2332 | 4.28E-13 | IIIVmrMLM | SIG |
| V_17AY | AX-94808117 | 5A | 476439578 | 0.1031 | | 6.00E-06 | BLUPmrMLM | SUG |
| V_17AY | AX-110541590 | 5A | 548579019 | -0.0323 | 0.691 | 1.14E-13 | IIIVmrMLM | SIG |
| V_17AY | AX-109857340 | 5A | 548862216 | 0.0081 | 0.4313 | 4.96E-13 | IIIVmrMLM | SIG |
| V_17AY | AX-109962402 | 5A | 666725153 | 0.0662 | | 4.35E-17 | IIIVmrMLM | SIG |
| V_17AY | AX-110389287 | 5B | 391903643 | 0.0045 | 0.1245 | 1.63E-11 | IIIVmrMLM | SIG |
| V_17AY | AX-111067739 | 5B | 476797802 | -0.0451 | | 5.81E-09 | IIIVmrMLM | SIG |
| V_17AY | AX-111151328 | 5B | 612941748 | -0.0502 | 1.1226 | 1.73E-17 | IIIVmrMLM | SIG |
| V_17AY | AX-111073414 | 5B | 679746034 | -0.0279 | -0.1669 | 1.82E-15 | IIIVmrMLM | SIG |
| V_17AY | AX-111699741 | 5B | 681078007 | 0.0339 | 0.5963 | 8.98E-12 | IIIVmrMLM | SIG |
| V_17AY | AX-95658487 | 5B | 682699266 | 0.0830 | | 3.62E-08 | BLUPmrMLM | SIG |
| V_17AY | AX-111577029 | 5B | 692522362 | 0.1444 | | 2.88E-06 | BLUPmrMLM | SUG |
| V_17AY | AX-109312560 | 5D | 21550414 | 0.0171 | 0.5366 | 3.52E-11 | IIIVmrMLM | SIG |
| V_17AY | AX-110313779 | 5D | 286455125 | -1.2888 | | 5.87E-07 | EMMAX | SUG |
| V_17AY | AX-111216290 | 5D | 421594485 | 0.4059 | | 1.42E-07 | GEMMA | SUG |
| V_17AY | AX-110616307 | 5D | 421619502 | 0.2062 | | 8.27E-08 | GEMMA | SIG |
| V_17AY | AX-109509060 | 6A | 320926179 | 0.0085 | 0.2071 | 4.39E-08 | IIIVmrMLM | SIG |
| V_17AY | AX-110412537 | 6B | 127612354 | 0.1339 | | 4.46E-06 | BLUPmrMLM | SUG |
| V_17AY | AX-94470972 | 6B | 226395363 | 0.7587 | 0.9158 | 0.00E+00 | IIIVmrMLM | SIG |
| V_17AY | AX-109945036 | 6B | 446297923 | 0.0124 | 0.1663 | 6.82E-06 | IIIVmrMLM | SUG |
| V_17AY | AX-109575981 | 6D | 331255908 | 0.0496 | | 1.68E-10 | IIIVmrMLM | SIG |
| V_17AY | AX-110981313 | 6D | 423790976 | -2.0441 | | 1.16E-08 | EMMAX | SIG |
| V_17AY | AX-94827671 | 7A | 11087367 | -0.0563 | | 3.21E-05 | BLUPmrMLM | SUG |
| V_17AY | AX-110371827 | 7A | 200060765 | 0.0505 | | 4.53E-05 | BLUPmrMLM | SUG |
| V_17AY | AX-108799491 | 7A | 267434161 | 0.0114 | 0.2817 | 5.33E-07 | IIIVmrMLM | SUG |
| V_17AY | AX-109949014 | 7A | 511922399 | 0.0093 | 0.1621 | 5.47E-08 | IIIVmrMLM | SIG |
| V_17AY | AX-112288950 | 7A | 538693397 | -0.1122 | 0.6222 | 5.94E-44 | IIIVmrMLM | SIG |
| V_17AY | AX-109924132 | 7A | 594163470 | -0.0077 | 0.0749 | 4.46E-08 | IIIVmrMLM | SIG |
| V_17AY | AX-111602188 | 7B | 63937125 | -2.0441 | | 1.16E-08 | EMMAX | SIG |
| V_17AY | AX-94549402 | 7B | 641806996 | 0.0369 | | 1.68E-06 | IIIVmrMLM | SUG |
| V_17AY | AX-109357973 | 7B | 642568775 | 0.0457 | | 2.80E-05 | BLUPmrMLM | SUG |
| V_17AY | AX-108769514 | 7B | 643700521 | 0.0451 | | 1.91E-04 | BLUPmrMLM | SUG |
| V_17AY | AX-111094331 | 7B | 723958130 | 0.3538 | | 1.86E-07 | GEMMA | SUG |
| V_17AY | AX-95684144 | 7B | 725385135 | -0.9321 | | 3.27E-08 | EMMAX | SIG |
| V_17AY | AX-94468441 | 7B | 725965878 | 0.1334 | | 2.69E-07 | GEMMA | SUG |
| V_17AY | AX-110445652 | 7B | 726646189 | -0.9321 | | 3.27E-08 | EMMAX | SIG |
| V_17AY | AX-95219160 | 7B | 727529846 | -0.9321 | | 3.27E-08 | EMMAX | SIG |
| V_17AY | AX-108728735 | 7B | 727652888 | -0.6023 | | 3.55E-07 | EMMAX | SUG |
| V_17AY | AX-108730446 | 7B | 729251407 | -0.9321 | | 3.27E-08 | EMMAX | SIG |

| | | | | | | | | |
|---|---|---|---|---|---|---|---|---|
| V_17AY | AX-111080832 | 7D | 83765607 | -2.0441 | | 1.16E-08 | EMMAX | SIG |
| V_17AY | AX-108933675 | 7D | 309758149 | 0.3669 | | 1.65E-06 | GEMMA | SUG |
| V_17AY | AX-110138076 | 7D | 390663125 | -2.0441 | | 1.16E-08 | EMMAX | SIG |
| V_17AY | AX-111022532 | 7D | 577019498 | 0.0509 | -0.0327 | 6.20E-10 | IIIVmrMLM | SIG |
| V_17AY | AX-94892803 | 7D | 625911375 | 0.0119 | 0.1496 | 9.69E-08 | IIIVmrMLM | SIG |
| V_17JY | AX-111093289 | 1A | 4207407 | -0.0229 | -0.1348 | 1.23E-11 | IIIVmrMLM | SIG |
| V_17JY | AX-109515249 | 1A | 320389780 | -0.0059 | 0.2659 | 1.77E-05 | IIIVmrMLM | SUG |
| V_17JY | AX-109823646 | 1A | 507751349 | -0.0173 | 0.1734 | 4.19E-05 | IIIVmrMLM | SUG |
| V_17JY | AX-109965064 | 1A | 580958888 | 0.0314 | -0.3836 | 5.68E-08 | IIIVmrMLM | SIG |
| V_17JY | AX-110932359 | 1A | 584539181 | 0.0552 | | 2.43E-05 | BLUPmrMLM | SUG |
| V_17JY | AX-110020113 | 1A | 591298823 | -0.0590 | 0.0742 | 1.76E-08 | IIIVmrMLM | SIG |
| V_17JY | AX-108811255 | 1B | 215088284 | -0.0041 | -0.1936 | 2.76E-05 | IIIVmrMLM | SUG |
| V_17JY | AX-111467003 | 1B | 642958515 | -0.0143 | 0.3287 | 8.63E-06 | IIIVmrMLM | SUG |
| V_17JY | AX-108802980 | 1B | 654101240 | -0.0349 | -0.4739 | 1.46E-08 | IIIVmrMLM | SIG |
| V_17JY | AX-110393744 | 1B | 662836423 | -0.0099 | -0.2161 | 3.06E-07 | IIIVmrMLM | SUG |
| V_17JY | AX-94526625 | 1D | 262341601 | -0.0842 | | 5.88E-17 | IIIVmrMLM | SIG |
| V_17JY | AX-111101313 | 1D | 329717031 | -0.1364 | -0.191 | 2.80E-37 | IIIVmrMLM | SIG |
| V_17JY | AX-110385168 | 2A | 440082976 | 0.0947 | | 7.99E-06 | BLUPmrMLM | SUG |
| V_17JY | AX-110921739 | 2A | 753113832 | -0.0461 | 0.2077 | 8.57E-06 | IIIVmrMLM | SUG |
| V_17JY | AX-89625389 | 3A | 12974622 | -0.1325 | | 1.27E-36 | IIIVmrMLM | SIG |
| V_17JY | AX-94653799 | 3A | 556359643 | 0.0559 | | 3.21E-06 | BLUPmrMLM | SUG |
| V_17JY | AX-109449758 | 3B | 717049513 | 0.0129 | -0.2682 | 4.13E-06 | IIIVmrMLM | SUG |
| V_17JY | AX-111196555 | 3B | 718286807 | -0.0506 | | 5.37E-05 | BLUPmrMLM | SUG |
| V_17JY | AX-94569370 | 3D | 4394608 | 0.1297 | | 7.71E-07 | BLUPmrMLM | SUG |
| V_17JY | AX-94569370 | 3D | 4394608 | -0.1035 | 0.0128 | 3.47E-20 | IIIVmrMLM | SIG |
| V_17JY | AX-110023899 | 3D | 556830798 | -0.0148 | 0.7929 | 4.69E-13 | IIIVmrMLM | SIG |
| V_17JY | AX-109584387 | 4B | 540408156 | 0.0261 | 0.8507 | 4.58E-05 | IIIVmrMLM | SUG |
| V_17JY | AX-109310695 | 5A | 566636735 | -0.1215 | | 1.13E-31 | IIIVmrMLM | SIG |
| V_17JY | AX-111117839 | 5B | 56823336 | 0.0030 | 0.3095 | 3.03E-10 | IIIVmrMLM | SIG |
| V_17JY | AX-110092729 | 5B | 85918342 | -0.0093 | -0.1738 | 1.99E-07 | IIIVmrMLM | SUG |
| V_17JY | AX-110961703 | 5B | 572668935 | 0.0877 | -0.2787 | 2.28E-17 | IIIVmrMLM | SIG |
| V_17JY | AX-111691372 | 5B | 641800860 | 0.0548 | -0.0941 | 1.30E-07 | IIIVmrMLM | SUG |
| V_17JY | AX-111476842 | 5B | 641801084 | -0.0505 | | 3.74E-05 | BLUPmrMLM | SUG |
| V_17JY | AX-110992345 | 6A | 20872523 | -0.0527 | | 1.45E-05 | BLUPmrMLM | SUG |
| V_17JY | AX-110992345 | 6A | 20872523 | 0.0398 | -0.0565 | 2.00E-04 | IIIVmrMLM | SUG |
| V_17JY | AX-111545041 | 6B | 31732845 | 0.0621 | | 6.90E-05 | BLUPmrMLM | SUG |
| V_17JY | AX-109979570 | 6B | 334222226 | 0.0159 | -0.1353 | 9.60E-05 | IIIVmrMLM | SUG |
| V_17JY | AX-110599656 | 6B | 520286398 | 0.0272 | -0.311 | 6.50E-08 | IIIVmrMLM | SIG |
| V_17JY | AX-108893208 | 6D | 3920459 | -0.0661 | | 2.25E-06 | BLUPmrMLM | SUG |
| V_17JY | AX-111988548 | 6D | 126021896 | 0.2901 | | 1.94E-06 | FarmCPU | SUG |
| V_17JY | AX-111734855 | 6D | 140217420 | 0.2781 | | 1.55E-06 | FarmCPU | SUG |
| V_17JY | AX-110650710 | 6D | 148794642 | 0.0817 | | 4.37E-16 | IIIVmrMLM | SIG |
| V_17JY | AX-110530035 | 7A | 528933513 | -0.0625 | | 1.89E-04 | BLUPmrMLM | SUG |
| V_17JY | AX-110575127 | 7A | 713173974 | 0.0028 | 0.6707 | 5.55E-06 | IIIVmrMLM | SUG |

| | | | | | | | | |
|---|---|---|---|---|---|---|---|---|
| V_17JY | AX-89526681 | 7B | 34668546 | -0.0597 | | 8.36E-05 | BLUPmrMLM | SUG |
| V_17JY | AX-94965000 | 7B | 559146431 | -0.0097 | 0.1355 | 3.08E-08 | IIIVmrMLM | SIG |
| V_17JY | AX-111987065 | 7D | 267005167 | 0.0509 | 0.5416 | 5.11E-16 | IIIVmrMLM | SIG |
| V_17JY | AX-95107918 | 7D | 576285018 | 0.1302 | | 1.25E-07 | BLUPmrMLM | SUG |
| V_17JY | AX-110765221 | 7D | 616377712 | -0.0517 | | 1.64E-07 | IIIVmrMLM | SUG |
| V_17JY | AX-110395618 | 7D | 619989528 | 0.0107 | 0.143 | 2.11E-04 | IIIVmrMLM | SUG |
| V_17JY | AX-110395618 | 7D | 619989528 | 0.2772 | | 1.88E-06 | GEMMA | SUG |
| V_17SQ | AX-109100815 | 1A | 435081406 | 0.0040 | 0.1571 | 2.53E-18 | IIIVmrMLM | SIG |
| V_17SQ | AX-108765513 | 2A | 98899087 | -0.0039 | -0.7322 | 8.04E-06 | IIIVmrMLM | SUG |
| V_17SQ | AX-109523266 | 2A | 740857412 | -0.0674 | | 1.23E-04 | BLUPmrMLM | SUG |
| V_17SQ | AX-109533291 | 2A | 765374350 | -0.0465 | 0.04 | 2.36E-11 | IIIVmrMLM | SIG |
| V_17SQ | AX-109902042 | 2B | 721280529 | 0.0186 | 0.1452 | 9.67E-10 | IIIVmrMLM | SIG |
| V_17SQ | AX-94442651 | 2D | 507050616 | 0.1873 | | 4.56E-07 | FarmCPU | SUG |
| V_17SQ | AX-111478343 | 3A | 224260678 | -0.0095 | 0.1263 | 5.94E-09 | IIIVmrMLM | SIG |
| V_17SQ | AX-108893324 | 3A | 485014357 | 0.0372 | | 6.26E-05 | BLUPmrMLM | SUG |
| V_17SQ | AX-108893324 | 3A | 485014357 | -0.0341 | 0.0127 | 3.48E-05 | IIIVmrMLM | SUG |
| V_17SQ | AX-110570407 | 3B | 410892577 | 0.0202 | 0.1214 | 4.61E-06 | IIIVmrMLM | SUG |
| V_17SQ | AX-111501616 | 3D | 599430827 | 0.0082 | 0.4288 | 3.75E-11 | IIIVmrMLM | SIG |
| V_17SQ | AX-94402318 | 3D | 609626811 | 0.0587 | -0.0851 | 7.49E-19 | IIIVmrMLM | SIG |
| V_17SQ | AX-109420906 | 4A | 115634351 | 0.1784 | | 8.52E-07 | FarmCPU | SUG |
| V_17SQ | AX-109092307 | 4A | 485173712 | -0.0389 | | 2.23E-09 | IIIVmrMLM | SIG |
| V_17SQ | AX-110409613 | 4B | 536088638 | 0.0238 | 0.2889 | 9.78E-10 | IIIVmrMLM | SIG |
| V_17SQ | AX-110449465 | 4B | 579884727 | -0.1779 | | 1.74E-06 | GEMMA | SUG |
| V_17SQ | AX-110996750 | 4D | 43397801 | -0.5774 | -0.4834 | 0.00E+00 | IIIVmrMLM | SIG |
| V_17SQ | AX-110604883 | 4D | 177489950 | 0.1760 | | 1.61E-06 | FarmCPU | SUG |
| V_17SQ | AX-110528135 | 4D | 445004978 | 0.1720 | | 1.03E-06 | FarmCPU | SUG |
| V_17SQ | AX-110686922 | 5A | 3402300 | -0.1540 | | 9.63E-07 | FarmCPU | SUG |
| V_17SQ | AX-109845940 | 5A | 414824229 | -0.0075 | -0.1027 | 9.69E-08 | IIIVmrMLM | SIG |
| V_17SQ | AX-108917596 | 5A | 518602100 | 0.2441 | | 1.52E-06 | GEMMA | SUG |
| V_17SQ | AX-109652474 | 5A | 519654975 | 0.2388 | | 1.76E-06 | GEMMA | SUG |
| V_17SQ | AX-111156514 | 5A | 519713020 | 0.2388 | | 1.76E-06 | GEMMA | SUG |
| V_17SQ | AX-111462197 | 5A | 519932603 | 0.2491 | | 1.59E-06 | GEMMA | SUG |
| V_17SQ | AX-111740483 | 5A | 520093132 | 0.2388 | | 1.76E-06 | GEMMA | SUG |
| V_17SQ | AX-110468146 | 5A | 520095740 | 0.2388 | | 1.76E-06 | GEMMA | SUG |
| V_17SQ | AX-108734074 | 5A | 520120297 | 0.2388 | | 1.76E-06 | GEMMA | SUG |
| V_17SQ | AX-109967095 | 5A | 520377086 | 0.2388 | | 1.76E-06 | GEMMA | SUG |
| V_17SQ | AX-109376123 | 5A | 520686648 | 0.2388 | | 1.76E-06 | GEMMA | SUG |
| V_17SQ | AX-109620761 | 5B | 84374458 | 0.0250 | 0.112 | 4.46E-13 | IIIVmrMLM | SIG |
| V_17SQ | AX-111676830 | 5B | 302107387 | -0.0359 | -0.0455 | 1.41E-08 | IIIVmrMLM | SIG |
| V_17SQ | AX-109867574 | 5B | 458117972 | 0.0069 | -0.2854 | 2.22E-11 | IIIVmrMLM | SIG |
| V_17SQ | AX-94700083 | 5B | 597813028 | 0.0100 | 0.0836 | 6.96E-08 | IIIVmrMLM | SIG |
| V_17SQ | AX-109537450 | 5B | 692629908 | -0.0115 | 0.226 | 5.03E-08 | IIIVmrMLM | SIG |
| V_17SQ | AX-111486228 | 5B | 693705288 | 0.0469 | | 1.32E-05 | BLUPmrMLM | SUG |
| V_17SQ | AX-110910508 | 5B | 703099269 | -0.0054 | -0.1284 | 1.18E-05 | IIIVmrMLM | SUG |

| | | | | | | | | |
|---|---|---|---|---|---|---|---|---|
| V_17SQ | AX-111186232 | 5B | 711269599 | -0.0222 | 1.1795 | 1.15E-15 | IIIVmrMLM | SIG |
| V_17SQ | AX-110460040 | 5D | 247618289 | -0.0730 | | 4.71E-06 | BLUPmrMLM | SUG |
| V_17SQ | AX-109512365 | 6B | 529713648 | 0.0124 | 0.1341 | 2.12E-09 | IIIVmrMLM | SIG |
| V_17SQ | AX-109315599 | 6B | 615533663 | 0.0550 | | 4.49E-05 | BLUPmrMLM | SUG |
| V_17SQ | AX-110211576 | 6D | 438817120 | -0.0501 | | 6.64E-05 | BLUPmrMLM | SUG |
| V_17SQ | AX-110954658 | 7A | 206721763 | 0.0032 | 0.4454 | 6.75E-10 | IIIVmrMLM | SIG |
| V_17SQ | AX-110624620 | 7A | 247065085 | 0.0051 | -0.1002 | 1.18E-07 | IIIVmrMLM | SUG |
| V_17SQ | AX-108908590 | 7A | 721572672 | 0.0162 | -0.368 | 6.43E-12 | IIIVmrMLM | SIG |
| V_17SQ | AX-109117982 | 7D | 41453283 | 0.0621 | | 1.82E-04 | BLUPmrMLM | SUG |
| V_17SQ | AX-109117982 | 7D | 41453283 | -0.0457 | | 2.64E-12 | IIIVmrMLM | SIG |
| V_17SQ | AX-110736759 | 7D | 173900640 | -0.0469 | -0.0315 | 1.78E-11 | IIIVmrMLM | SIG |
| V_18AY | AX-111706902 | 1A | 534940298 | -0.0279 | | 1.06E-04 | BLUPmrMLM | SUG |
| V_18AY | AX-110977273 | 1B | 329377173 | -0.0222 | 0.0734 | 2.04E-05 | IIIVmrMLM | SUG |
| V_18AY | AX-108837702 | 1B | 590066339 | 0.0429 | | 4.13E-05 | BLUPmrMLM | SUG |
| V_18AY | AX-109822794 | 2A | 74561389 | 0.0009 | -0.0842 | 1.59E-07 | IIIVmrMLM | SUG |
| V_18AY | AX-110123869 | 2A | 284265347 | -0.0197 | -0.2346 | 1.15E-10 | IIIVmrMLM | SIG |
| V_18AY | AX-111161668 | 2A | 699577780 | -0.0007 | -0.1398 | 1.62E-04 | IIIVmrMLM | SUG |
| V_18AY | AX-111634969 | 2B | 18263358 | -0.0379 | 0.0145 | 5.03E-07 | IIIVmrMLM | SUG |
| V_18AY | AX-111134442 | 2B | 183950769 | -0.0163 | -0.1055 | 7.35E-10 | IIIVmrMLM | SIG |
| V_18AY | AX-110409334 | 2B | 655971308 | -0.0141 | 0.631 | 1.03E-05 | IIIVmrMLM | SUG |
| V_18AY | AX-86179787 | 2D | 12786080 | -0.0300 | 0.137 | 1.39E-08 | IIIVmrMLM | SIG |
| V_18AY | AX-109831876 | 3A | 587903751 | -0.0394 | | 1.34E-04 | BLUPmrMLM | SUG |
| V_18AY | AX-108750895 | 3B | 60378096 | -0.0053 | -0.1141 | 2.03E-04 | IIIVmrMLM | SUG |
| V_18AY | AX-109854926 | 3B | 216379514 | -0.0040 | 0.208 | 4.80E-07 | IIIVmrMLM | SUG |
| V_18AY | AX-108912639 | 3B | 385142555 | -0.0022 | 0.2387 | 5.11E-12 | IIIVmrMLM | SIG |
| V_18AY | AX-110101772 | 3D | 566486214 | 0.0032 | 0.2353 | 4.07E-06 | IIIVmrMLM | SUG |
| V_18AY | AX-109950845 | 4A | 165384884 | 0.1114 | | 1.57E-06 | GEMMA | SUG |
| V_18AY | AX-109406924 | 4A | 222426885 | 0.1283 | | 2.08E-06 | GEMMA | SUG |
| V_18AY | AX-108849069 | 4A | 235835183 | 0.1114 | | 1.57E-06 | GEMMA | SUG |
| V_18AY | AX-111148636 | 4A | 241488851 | 0.1723 | | 2.07E-07 | GEMMA | SUG |
| V_18AY | AX-109416191 | 4A | 257588388 | 0.1292 | | 7.44E-07 | GEMMA | SUG |
| V_18AY | AX-110958757 | 4A | 325474739 | 0.1429 | | 6.59E-07 | GEMMA | SUG |
| V_18AY | AX-108878378 | 4A | 365471237 | 0.2207 | | 5.28E-07 | GEMMA | SUG |
| V_18AY | AX-108843439 | 4A | 367914721 | 0.1093 | | 2.24E-06 | GEMMA | SUG |
| V_18AY | AX-109970978 | 4A | 404973142 | 0.1875 | | 5.94E-07 | GEMMA | SUG |
| V_18AY | AX-109861259 | 4A | 408289061 | 0.1167 | | 1.39E-06 | GEMMA | SUG |
| V_18AY | AX-110597403 | 4A | 410678628 | 0.1268 | | 1.86E-06 | GEMMA | SUG |
| V_18AY | AX-111637659 | 4A | 414720487 | 0.1219 | | 1.01E-06 | GEMMA | SUG |
| V_18AY | AX-110132259 | 4A | 415855084 | 0.1230 | | 9.31E-07 | GEMMA | SUG |
| V_18AY | AX-110126645 | 4A | 427783849 | 0.1093 | | 2.24E-06 | GEMMA | SUG |
| V_18AY | AX-110471402 | 4A | 437431457 | 0.1082 | | 2.33E-06 | GEMMA | SUG |
| V_18AY | AX-109058806 | 4A | 441126306 | 0.1620 | | 1.68E-06 | GEMMA | SUG |
| V_18AY | AX-111080240 | 4A | 449889292 | -0.0110 | -0.1069 | 1.77E-09 | IIIVmrMLM | SIG |
| V_18AY | AX-110541157 | 4A | 626510311 | -0.0169 | 0.4361 | 1.48E-09 | IIIVmrMLM | SIG |

| | | | | | | | | |
|---|---|---|---|---|---|---|---|---|
| V_18AY | AX-89362528 | 4B | 637390229 | -0.0471 | -0.0546 | 3.76E-13 | IIIVmrMLM | SIG |
| V_18AY | AX-108976919 | 4B | 672563232 | -0.0322 | | 5.24E-06 | BLUPmrMLM | SUG |
| V_18AY | AX-108976919 | 4B | 672563232 | 0.0366 | -0.1692 | 2.35E-10 | IIIVmrMLM | SIG |
| V_18AY | AX-111176492 | 5A | 364981589 | 0.0430 | | 1.04E-04 | BLUPmrMLM | SUG |
| V_18AY | AX-111480358 | 5A | 521939210 | -0.0138 | -0.1064 | 2.10E-08 | IIIVmrMLM | SIG |
| V_18AY | AX-110044503 | 5A | 528288061 | -0.0142 | -0.2791 | 1.08E-10 | IIIVmrMLM | SIG |
| V_18AY | AX-110941281 | 5A | 588022791 | 0.0337 | 0.0107 | 8.03E-07 | IIIVmrMLM | SUG |
| V_18AY | AX-110123916 | 5A | 588401903 | 0.0100 | 0.1597 | 1.12E-06 | IIIVmrMLM | SUG |
| V_18AY | AX-111465232 | 5A | 698495514 | -0.0130 | 0.0858 | 2.99E-13 | IIIVmrMLM | SIG |
| V_18AY | AX-111557902 | 5B | 257910180 | 0.0715 | | 2.20E-05 | BLUPmrMLM | SUG |
| V_18AY | AX-111557902 | 5B | 257910180 | -0.0400 | | 1.09E-10 | IIIVmrMLM | SIG |
| V_18AY | AX-109542966 | 5B | 470978512 | -0.0994 | | 8.83E-09 | BLUPmrMLM | SIG |
| V_18AY | AX-108840042 | 5D | 42961528 | 0.0356 | 0.045 | 5.25E-08 | IIIVmrMLM | SIG |
| V_18AY | AX-94460640 | 5D | 220098422 | 0.0341 | | 3.30E-08 | IIIVmrMLM | SIG |
| V_18AY | AX-110471646 | 6A | 5327988 | 0.0465 | | 7.57E-05 | BLUPmrMLM | SUG |
| V_18AY | AX-111548703 | 6A | 13901452 | -0.0048 | 0.1769 | 2.82E-06 | IIIVmrMLM | SUG |
| V_18AY | AX-111696088 | 6B | 191610061 | -0.0397 | | 1.48E-10 | IIIVmrMLM | SIG |
| V_18AY | AX-110414931 | 7A | 663225859 | 0.0000 | 0.5159 | 7.37E-06 | IIIVmrMLM | SUG |
| V_18AY | AX-111583409 | 7A | 696648523 | -0.0907 | -0.1372 | 4.44E-42 | IIIVmrMLM | SIG |
| V_18AY | AX-111728563 | 7A | 703564535 | 0.0062 | -0.4351 | 2.07E-06 | IIIVmrMLM | SUG |
| V_18AY | AX-110451297 | 7B | 70528757 | 0.0413 | 0.0827 | 1.21E-10 | IIIVmrMLM | SIG |
| V_18AY | AX-109045123 | 7B | 134224402 | 0.0315 | | 3.21E-07 | IIIVmrMLM | SUG |
| V_18AY | AX-94403275 | 7B | 718148246 | -0.0177 | -0.1828 | 5.88E-07 | IIIVmrMLM | SUG |
| V_18JY | AX-110982739 | 1A | 38723482 | -0.0488 | 0.0555 | 4.05E-11 | IIIVmrMLM | SIG |
| V_18JY | AX-111140620 | 1B | 488654477 | -0.0132 | -0.226 | 2.57E-05 | IIIVmrMLM | SUG |
| V_18JY | AX-108735443 | 1B | 614592227 | 0.0054 | 0.1108 | 9.53E-06 | IIIVmrMLM | SUG |
| V_18JY | AX-111511089 | 1B | 651889889 | -0.0181 | 0.1072 | 2.07E-10 | IIIVmrMLM | SIG |
| V_18JY | AX-110656149 | 1D | 28582081 | 0.0421 | | 2.63E-09 | IIIVmrMLM | SIG |
| V_18JY | AX-94767334 | 1D | 30107708 | 0.0350 | | 6.54E-07 | IIIVmrMLM | SUG |
| V_18JY | AX-110479317 | 2A | 461400524 | -0.0610 | | 5.14E-05 | BLUPmrMLM | SUG |
| V_18JY | AX-108726222 | 2B | 665195 | -0.0355 | -0.0111 | 5.90E-06 | IIIVmrMLM | SUG |
| V_18JY | AX-110021725 | 2D | 433016946 | -0.0422 | -0.0021 | 2.48E-08 | IIIVmrMLM | SIG |
| V_18JY | AX-108795296 | 3A | 15284502 | 0.0081 | 0.1743 | 1.51E-06 | IIIVmrMLM | SUG |
| V_18JY | AX-109376551 | 3A | 702010254 | -0.0370 | -0.0337 | 1.21E-06 | IIIVmrMLM | SUG |
| V_18JY | AX-110472635 | 3A | 709437566 | 0.0081 | -0.1049 | 4.40E-09 | IIIVmrMLM | SIG |
| V_18JY | AX-108959271 | 3A | 738751751 | -0.0617 | | 4.84E-06 | BLUPmrMLM | SUG |
| V_18JY | AX-94437164 | 3B | 112281346 | -0.0468 | | 4.41E-11 | IIIVmrMLM | SIG |
| V_18JY | AX-110384434 | 3B | 224311404 | 0.0443 | -0.0153 | 1.38E-06 | IIIVmrMLM | SUG |
| V_18JY | AX-109274638 | 3B | 237325305 | -0.1785 | | 3.41E-07 | FarmCPU | SUG |
| V_18JY | AX-111154757 | 3B | 362049509 | -0.0143 | -0.1272 | 2.95E-17 | IIIVmrMLM | SIG |
| V_18JY | AX-111571461 | 3B | 398344047 | -0.0214 | 0.2497 | 7.89E-08 | IIIVmrMLM | SIG |
| V_18JY | AX-94482301 | 3B | 550646472 | -0.0637 | -0.0507 | 2.43E-17 | IIIVmrMLM | SIG |
| V_18JY | AX-110374458 | 3B | 762803100 | 0.0668 | | 1.32E-06 | BLUPmrMLM | SUG |
| V_18JY | AX-111598404 | 3B | 828424807 | 0.0058 | -0.1381 | 5.11E-07 | IIIVmrMLM | SUG |

| | | | | | | | | |
|---|---|---|---|---|---|---|---|---|
| V_18JY | AX-110613795 | 3D | 6016156 | -0.0008 | -0.2272 | 1.02E-06 | IIIVmrMLM | SUG |
| V_18JY | AX-94434096 | 4A | 11843465 | 0.0581 | | 5.39E-06 | BLUPmrMLM | SUG |
| V_18JY | AX-111010398 | 4A | 566568943 | 0.0277 | 0.1074 | 2.31E-12 | IIIVmrMLM | SIG |
| V_18JY | AX-111101603 | 4B | 426825943 | -0.0491 | | 1.48E-04 | BLUPmrMLM | SUG |
| V_18JY | AX-109876373 | 4B | 545026906 | 0.0431 | | 7.81E-05 | BLUPmrMLM | SUG |
| V_18JY | AX-111475478 | 4D | 15241363 | -0.0346 | | 1.64E-04 | BLUPmrMLM | SUG |
| V_18JY | AX-109861283 | 5A | 18754105 | -0.0018 | 0.2766 | 4.49E-07 | IIIVmrMLM | SUG |
| V_18JY | AX-110185399 | 5A | 84307738 | -0.1151 | | 8.56E-07 | FarmCPU | SUG |
| V_18JY | AX-99612213 | 5A | 472549846 | -0.0335 | 0.0485 | 1.22E-06 | IIIVmrMLM | SUG |
| V_18JY | AX-110929258 | 5B | 635500 | -0.0319 | | 5.51E-05 | BLUPmrMLM | SUG |
| V_18JY | AX-109366230 | 5B | 11763565 | 0.0038 | 0.103 | 1.65E-06 | IIIVmrMLM | SUG |
| V_18JY | AX-110942490 | 5B | 303439742 | -0.0468 | | 1.61E-05 | BLUPmrMLM | SUG |
| V_18JY | AX-110942490 | 5B | 303439742 | 0.0357 | -0.0521 | 1.17E-06 | IIIVmrMLM | SUG |
| V_18JY | AX-111611710 | 5B | 456592153 | -0.0143 | -0.1286 | 3.62E-05 | IIIVmrMLM | SUG |
| V_18JY | AX-109308779 | 5B | 686549526 | -0.0734 | | 1.09E-06 | GEMMA | SUG |
| V_18JY | AX-110000074 | 5D | 510903531 | -0.0106 | 0.5645 | 5.54E-14 | IIIVmrMLM | SIG |
| V_18JY | AX-109988806 | 6A | 244806022 | -0.0099 | 0.0977 | 2.11E-07 | IIIVmrMLM | SUG |
| V_18JY | AX-110518201 | 6B | 479077712 | 0.0549 | | 3.59E-06 | BLUPmrMLM | SUG |
| V_18JY | AX-110536078 | 6B | 482068535 | 0.0104 | 0.2308 | 5.61E-16 | IIIVmrMLM | SIG |
| V_18JY | AX-95113401 | 6B | 716437491 | -0.0524 | | 1.31E-05 | BLUPmrMLM | SUG |
| V_18JY | AX-95113401 | 6B | 716437491 | 0.0401 | -0.0097 | 1.93E-04 | IIIVmrMLM | SUG |
| V_18JY | AX-108744137 | 6D | 86602960 | -0.0359 | | 8.65E-05 | BLUPmrMLM | SUG |
| V_18JY | AX-109577332 | 7A | 12712818 | 0.0014 | 0.0789 | 2.98E-05 | IIIVmrMLM | SUG |
| V_18JY | AX-109335759 | 7A | 19419576 | 0.0367 | | 1.31E-04 | BLUPmrMLM | SUG |
| V_18JY | AX-110373987 | 7A | 149497638 | 0.0371 | -0.2975 | 3.14E-14 | IIIVmrMLM | SIG |
| V_18JY | AX-95239643 | 7A | 253801294 | 0.0575 | | 1.18E-04 | BLUPmrMLM | SUG |
| V_18JY | AX-110670722 | 7B | 687559113 | -0.0416 | | 5.87E-05 | BLUPmrMLM | SUG |
| V_18JY | AX-108876019 | 7B | 705812472 | 0.0357 | -0.0052 | 3.49E-05 | IIIVmrMLM | SUG |
| V_18JY | AX-109660239 | 7D | 529065509 | -0.2006 | | 8.72E-08 | FarmCPU | SIG |
| V_18JY | AX-110043120 | 7D | 529718874 | 0.1695 | | 1.37E-06 | FarmCPU | SUG |
| V_19XX | AX-94885115 | 1A | 574935641 | 0.0140 | -0.107 | 2.81E-05 | IIIVmrMLM | SUG |
| V_19XX | AX-111076068 | 1B | 641626988 | -0.0025 | -0.1663 | 1.79E-05 | IIIVmrMLM | SUG |
| V_19XX | AX-110951240 | 1B | 667471175 | 0.0754 | -0.0109 | 3.86E-10 | IIIVmrMLM | SIG |
| V_19XX | AX-108977627 | 1D | 18434579 | -0.0349 | | 1.42E-04 | BLUPmrMLM | SUG |
| V_19XX | AX-108903031 | 2A | 246750499 | -0.0152 | -0.1208 | 6.09E-05 | IIIVmrMLM | SUG |
| V_19XX | AX-111756851 | 2A | 365311747 | 0.0627 | | 7.00E-06 | BLUPmrMLM | SUG |
| V_19XX | AX-109329398 | 2B | 30467393 | 0.0056 | 0.2115 | 9.70E-07 | IIIVmrMLM | SUG |
| V_19XX | AX-111616011 | 2D | 431005994 | -0.0977 | | 3.67E-05 | BLUPmrMLM | SUG |
| V_19XX | AX-111157535 | 3A | 246411867 | -0.0012 | 0.1372 | 5.80E-08 | IIIVmrMLM | SIG |
| V_19XX | AX-111100825 | 3A | 513431817 | -0.0324 | -0.1905 | 2.31E-10 | IIIVmrMLM | SIG |
| V_19XX | AX-111668345 | 3A | 547375427 | 0.1713 | | 1.93E-06 | GEMMA | SUG |
| V_19XX | AX-108837444 | 3A | 701883881 | -0.0236 | 0.0969 | 8.16E-05 | IIIVmrMLM | SUG |
| V_19XX | AX-111613821 | 3A | 743233091 | -0.0411 | | 1.24E-05 | BLUPmrMLM | SUG |
| V_19XX | AX-111075723 | 3B | 189491216 | -0.0658 | | 1.53E-04 | BLUPmrMLM | SUG |

| | | | | | | | | |
|---|---|---|---|---|---|---|---|---|
| V_19XX | AX-109627460 | 3B | 361746252 | 0.1960 | | 2.17E-06 | GEMMA | SUG |
| V_19XX | AX-108730133 | 3B | 796472256 | 0.0486 | | 5.41E-06 | BLUPmrMLM | SUG |
| V_19XX | AX-111474562 | 3D | 561907763 | 0.0562 | -0.2606 | 6.31E-12 | IIIVmrMLM | SIG |
| V_19XX | AX-109519754 | 4A | 530911336 | -0.0287 | -0.1206 | 4.81E-07 | IIIVmrMLM | SUG |
| V_19XX | AX-108989194 | 4A | 539749095 | 0.0192 | -0.2076 | 2.16E-06 | IIIVmrMLM | SUG |
| V_19XX | AX-108730857 | 4A | 695773000 | 0.0743 | -0.6091 | 1.70E-17 | IIIVmrMLM | SIG |
| V_19XX | AX-109375803 | 4B | 19760967 | 0.0830 | | 3.07E-05 | BLUPmrMLM | SUG |
| V_19XX | AX-109375803 | 4B | 19760967 | -0.0770 | 0.0684 | 1.99E-13 | IIIVmrMLM | SIG |
| V_19XX | AX-110119356 | 4D | 206916951 | 0.0945 | | 6.01E-05 | BLUPmrMLM | SUG |
| V_19XX | AX-110119356 | 4D | 206916951 | -0.0516 | | 8.19E-08 | IIIVmrMLM | SIG |
| V_19XX | AX-111615781 | 5A | 622545850 | -0.0133 | 0.4754 | 1.76E-09 | IIIVmrMLM | SIG |
| V_19XX | AX-109408919 | 5A | 631660443 | 0.0106 | 0.2925 | 1.27E-05 | IIIVmrMLM | SUG |
| V_19XX | AX-111092536 | 5A | 668350592 | -0.0330 | 0.1662 | 1.45E-09 | IIIVmrMLM | SIG |
| V_19XX | AX-108834834 | 5A | 672345719 | 0.0662 | | 1.63E-05 | BLUPmrMLM | SUG |
| V_19XX | AX-95119812 | 5B | 679135614 | -0.0004 | -0.134 | 1.91E-05 | IIIVmrMLM | SUG |
| V_19XX | AX-94649090 | 5B | 695660124 | 0.0246 | -0.1013 | 7.87E-07 | IIIVmrMLM | SUG |
| V_19XX | AX-111755433 | 5D | 117243708 | 0.0512 | | 2.00E-04 | BLUPmrMLM | SUG |
| V_19XX | AX-110463548 | 6A | 2788772 | -0.0802 | | 1.04E-05 | BLUPmrMLM | SUG |
| V_19XX | AX-110550202 | 6A | 153671397 | -0.0585 | | 3.58E-05 | BLUPmrMLM | SUG |
| V_19XX | AX-95139273 | 6B | 30011557 | 0.0158 | -0.1545 | 4.94E-06 | IIIVmrMLM | SUG |
| V_19XX | AX-109964404 | 7A | 50020938 | 0.0070 | 0.1711 | 5.37E-05 | IIIVmrMLM | SUG |
| V_19XX | AX-108764784 | 7A | 682777496 | -0.0313 | -0.1027 | 1.06E-13 | IIIVmrMLM | SIG |
| V_19XX | AX-110014352 | 7B | 99800781 | 0.0497 | | 6.10E-06 | BLUPmrMLM | SUG |
| V_19XX | AX-110906456 | 7B | 478466747 | 0.0551 | | 1.17E-04 | BLUPmrMLM | SUG |
| V_19XX | AX-95160924 | 7B | 601117142 | -0.0772 | | 8.53E-05 | BLUPmrMLM | SUG |
| V_19XX | AX-108993899 | 7B | 702236106 | 0.0479 | | 6.06E-07 | IIIVmrMLM | SUG |
| V_BLUP | AX-108787384 | 1A | 376508048 | -0.0125 | -0.0405 | 5.58E-14 | IIIVmrMLM | SIG |
| V_BLUP | AX-110485334 | 1A | 410678115 | -0.0206 | | 3.71E-05 | BLUPmrMLM | SUG |
| V_BLUP | AX-111470849 | 1A | 473906588 | 0.0034 | -0.0812 | 8.62E-11 | IIIVmrMLM | SIG |
| V_BLUP | AX-110037375 | 1B | 386964234 | 0.0038 | -0.0886 | 2.44E-11 | IIIVmrMLM | SIG |
| V_BLUP | AX-110394593 | 1B | 463530278 | 0.0159 | | 3.32E-05 | BLUPmrMLM | SUG |
| V_BLUP | AX-110394593 | 1B | 463530278 | -0.0024 | 0.0364 | 1.91E-06 | IIIVmrMLM | SUG |
| V_BLUP | AX-111511089 | 1B | 651889889 | -0.0011 | 0.0395 | 4.94E-08 | IIIVmrMLM | SIG |
| V_BLUP | AX-111038629 | 2A | 70646959 | 0.0127 | | 6.72E-05 | BLUPmrMLM | SUG |
| V_BLUP | AX-109383749 | 2A | 483937237 | 0.0061 | -0.0664 | 3.90E-07 | IIIVmrMLM | SUG |
| V_BLUP | AX-109973922 | 2A | 504467482 | -0.0064 | -0.0986 | 1.43E-06 | IIIVmrMLM | SUG |
| V_BLUP | AX-111071776 | 2B | 18193542 | 0.0164 | | 8.33E-05 | BLUPmrMLM | SUG |
| V_BLUP | AX-109356374 | 2B | 654528306 | 0.0082 | 0.2322 | 1.24E-09 | IIIVmrMLM | SIG |
| V_BLUP | AX-94442651 | 2D | 507050616 | 0.1018 | | 1.62E-07 | FarmCPU | SUG |
| V_BLUP | AX-110000073 | 2D | 606688607 | -0.0036 | 0.0548 | 2.80E-04 | IIIVmrMLM | SUG |
| V_BLUP | AX-110922479 | 3A | 8795309 | 0.0022 | 0.0862 | 1.85E-06 | IIIVmrMLM | SUG |
| V_BLUP | AX-111582837 | 3A | 13009071 | 0.0445 | | 9.34E-07 | GEMMA | SUG |
| V_BLUP | AX-109286245 | 3A | 421119830 | 0.0767 | | 9.05E-07 | FarmCPU | SUG |
| V_BLUP | AX-95184677 | 3A | 510007314 | -0.0385 | | 2.29E-06 | GEMMA | SUG |

| | | | | | | | | |
|---|---|---|---|---|---|---|---|---|
| V_BLUP | AX-109274638 | 3B | 237325305 | -0.0849 | | 1.26E-07 | FarmCPU | SUG |
| V_BLUP | AX-111600364 | 3B | 795031754 | -0.0178 | -0.0182 | 2.76E-09 | IIIVmrMLM | SIG |
| V_BLUP | AX-109288929 | 3D | 374023593 | 0.0301 | | 1.49E-05 | BLUPmrMLM | SUG |
| V_BLUP | AX-109420906 | 4A | 115634351 | 0.0942 | | 6.43E-07 | FarmCPU | SUG |
| V_BLUP | AX-94942161 | 4A | 136928119 | 0.0357 | | 5.12E-07 | BLUPmrMLM | SUG |
| V_BLUP | AX-94942161 | 4A | 136928119 | -0.0185 | | 8.25E-11 | IIIVmrMLM | SIG |
| V_BLUP | AX-111031267 | 4A | 398033542 | -0.0054 | 0.1018 | 1.50E-06 | IIIVmrMLM | SUG |
| V_BLUP | AX-111065828 | 4A | 634112640 | 0.0025 | 0.0667 | 5.12E-10 | IIIVmrMLM | SIG |
| V_BLUP | AX-110449465 | 4B | 579884727 | -0.0793 | | 5.49E-07 | GEMMA | SUG |
| V_BLUP | AX-110604883 | 4D | 177489950 | 0.0935 | | 1.02E-06 | FarmCPU | SUG |
| V_BLUP | AX-108820823 | 5A | 473425215 | -0.0217 | | 7.33E-06 | BLUPmrMLM | SUG |
| V_BLUP | AX-109039273 | 5B | 14371594 | -0.0027 | -0.0354 | 3.92E-07 | IIIVmrMLM | SUG |
| V_BLUP | AX-110413869 | 5B | 462559785 | 0.0013 | 0.0698 | 5.70E-06 | IIIVmrMLM | SUG |
| V_BLUP | AX-108957573 | 5B | 476812795 | 0.0245 | 0.0113 | 8.06E-14 | IIIVmrMLM | SIG |
| V_BLUP | AX-111591776 | 5B | 476928425 | -0.0022 | -0.1447 | 1.99E-06 | IIIVmrMLM | SUG |
| V_BLUP | AX-110398849 | 5B | 477507219 | -0.0390 | | 2.06E-06 | GEMMA | SUG |
| V_BLUP | AX-111581527 | 5B | 485410065 | 0.0010 | -0.0736 | 4.21E-05 | IIIVmrMLM | SUG |
| V_BLUP | AX-109824744 | 5B | 545398047 | -0.0337 | | 1.78E-07 | BLUPmrMLM | SUG |
| V_BLUP | AX-111528735 | 5B | 574063657 | 0.0918 | | 1.84E-06 | FarmCPU | SUG |
| V_BLUP | AX-111691372 | 5B | 641800860 | 0.0182 | -0.0512 | 1.17E-10 | IIIVmrMLM | SIG |
| V_BLUP | AX-111031169 | 5B | 669905984 | 0.0070 | 0.2116 | 7.18E-10 | IIIVmrMLM | SIG |
| V_BLUP | AX-95658487 | 5B | 682699266 | -0.0140 | -0.1133 | 9.55E-08 | IIIVmrMLM | SIG |
| V_BLUP | AX-109437383 | 5B | 690078365 | 0.0335 | | 7.32E-07 | GEMMA | SUG |
| V_BLUP | AX-109026256 | 5B | 690078401 | 0.0325 | | 1.57E-06 | GEMMA | SUG |
| V_BLUP | AX-109271003 | 5B | 690078593 | 0.0328 | | 1.29E-06 | GEMMA | SUG |
| V_BLUP | AX-109298088 | 5B | 690078627 | 0.0324 | | 1.99E-06 | GEMMA | SUG |
| V_BLUP | AX-94464717 | 5B | 690471344 | 0.0695 | | 1.70E-06 | GEMMA | SUG |
| V_BLUP | AX-110598090 | 5B | 690699628 | 0.0331 | | 1.74E-06 | GEMMA | SUG |
| V_BLUP | AX-111583446 | 5B | 691954571 | 0.0258 | | 6.49E-08 | BLUPmrMLM | SIG |
| V_BLUP | AX-111583446 | 5B | 691954571 | 0.0304 | | 4.80E-07 | GEMMA | SUG |
| V_BLUP | AX-108840042 | 5D | 42961528 | -0.0160 | | 3.98E-05 | BLUPmrMLM | SUG |
| V_BLUP | AX-108725335 | 6A | 14101606 | -0.0018 | 0.078 | 2.38E-04 | IIIVmrMLM | SUG |
| V_BLUP | AX-111453496 | 6A | 30431324 | -0.0001 | 0.0368 | 1.11E-06 | IIIVmrMLM | SUG |
| V_BLUP | AX-111620734 | 6B | 4125682 | -0.0138 | 0.0263 | 4.23E-06 | IIIVmrMLM | SUG |
| V_BLUP | AX-110430140 | 6B | 4548176 | 0.0175 | | 3.02E-05 | BLUPmrMLM | SUG |
| V_BLUP | AX-111726688 | 6B | 11300863 | -0.1522 | | 3.24E-07 | FarmCPU | SUG |
| V_BLUP | AX-111071251 | 6B | 11318645 | -0.0963 | | 9.77E-07 | FarmCPU | SUG |
| V_BLUP | AX-110429558 | 6B | 11319311 | -0.1940 | | 9.96E-07 | FarmCPU | SUG |
| V_BLUP | AX-111569081 | 6B | 11587124 | -0.0703 | | 2.28E-06 | FarmCPU | SUG |
| V_BLUP | AX-109391962 | 6B | 339268873 | 0.1033 | | 2.85E-07 | FarmCPU | SUG |
| V_BLUP | AX-110527512 | 6B | 447180159 | -0.0080 | -0.0616 | 6.45E-07 | IIIVmrMLM | SUG |
| V_BLUP | AX-111571572 | 6B | 679301596 | 0.0216 | -0.0053 | 4.81E-06 | IIIVmrMLM | SUG |
| V_BLUP | AX-94420296 | 6D | 382589362 | 0.0158 | | 2.09E-08 | IIIVmrMLM | SIG |
| V_BLUP | AX-111759310 | 6D | 410108635 | -0.0678 | | 1.29E-06 | FarmCPU | SUG |

| | | | | | | | | |
|---|---|---|---|---|---|---|---|---|
| V_BLUP | AX-109305993 | 6D | 470387179 | -0.0666 | | 3.58E-07 | GEMMA | SUG |
| V_BLUP | AX-110526933 | 6D | 470387275 | -0.0544 | | 9.55E-07 | GEMMA | SUG |
| V_BLUP | AX-109299321 | 6D | 470387591 | -0.0666 | | 3.58E-07 | GEMMA | SUG |
| V_BLUP | AX-111038802 | 6D | 470752471 | -0.0522 | | 1.61E-06 | GEMMA | SUG |
| V_BLUP | AX-110857795 | 6D | 470847028 | -0.0553 | | 7.34E-07 | GEMMA | SUG |
| V_BLUP | AX-111010395 | 6D | 470864026 | -0.0510 | | 2.25E-06 | GEMMA | SUG |
| V_BLUP | AX-109304810 | 7A | 211734058 | -0.0061 | -0.063 | 6.07E-06 | IIIVmrMLM | SUG |
| V_BLUP | AX-111809963 | 7A | 227086699 | 0.0053 | 0.0523 | 4.73E-08 | IIIVmrMLM | SIG |
| V_BLUP | AX-94889159 | 7B | 144733129 | 0.0175 | | 7.54E-10 | IIIVmrMLM | SIG |

V: average grain filling rate.

**Supplementary Table S18.** All QTNs identified as associated with maximum filling rate in wheat.

| Trait | Marker | Chr | Position | Add.effect | Dom.effect | P-value | Method | Sig. |
|---|---|---|---|---|---|---|---|---|
| Vmax_17AY | AX-111031083 | 1B | 50023282 | -0.0012 | -0.5881 | 2.23E-10 | IIIVmrMLM | SIG |
| Vmax_17AY | AX-109519577 | 1B | 98910393 | 0.1218 | | 3.36E-08 | BLUPmrMLM | SIG |
| Vmax_17AY | AX-109109867 | 1B | 110950982 | -0.0153 | -0.6471 | 5.88E-09 | IIIVmrMLM | SIG |
| Vmax_17AY | AX-111540876 | 1B | 543473980 | 0.0309 | 0.1946 | 1.32E-13 | IIIVmrMLM | SIG |
| Vmax_17AY | AX-108788117 | 1B | 654531498 | -0.1262 | | 4.74E-07 | BLUPmrMLM | SUG |
| Vmax_17AY | AX-111056687 | 1B | 673412687 | 0.0727 | | 3.29E-05 | BLUPmrMLM | SUG |
| Vmax_17AY | AX-110047281 | 1D | 595098 | -3.0604 | | 1.24E-08 | EMMAX | SIG |
| Vmax_17AY | AX-110392337 | 1D | 73679336 | -0.7459 | | 2.35E-09 | EMMAX | SIG |
| Vmax_17AY | AX-109495041 | 2A | 144658876 | -0.0458 | -0.3052 | 1.45E-11 | IIIVmrMLM | SIG |
| Vmax_17AY | AX-109644557 | 2A | 602699209 | -0.0462 | -0.2872 | 1.63E-11 | IIIVmrMLM | SIG |
| Vmax_17AY | AX-109273856 | 2A | 771343034 | -0.0710 | | 2.14E-05 | BLUPmrMLM | SUG |
| Vmax_17AY | AX-94971846 | 2B | 30499258 | 0.0249 | 0.2254 | 3.19E-20 | IIIVmrMLM | SIG |
| Vmax_17AY | AX-111559203 | 2B | 154588905 | 0.0928 | -0.3814 | 3.42E-16 | IIIVmrMLM | SIG |
| Vmax_17AY | AX-109974976 | 2B | 421251405 | 0.1116 | | 2.72E-06 | BLUPmrMLM | SUG |
| Vmax_17AY | AX-109499517 | 2B | 427082691 | -0.0438 | 0.5677 | 5.15E-08 | IIIVmrMLM | SIG |
| Vmax_17AY | AX-109426196 | 2B | 537338926 | -0.0369 | -0.2645 | 1.18E-15 | IIIVmrMLM | SIG |
| Vmax_17AY | AX-94618830 | 2B | 546979309 | -3.0604 | | 1.24E-08 | EMMAX | SIG |
| Vmax_17AY | AX-109919029 | 2B | 549204515 | 0.5045 | | 4.37E-07 | GEMMA | SUG |
| Vmax_17AY | AX-109412939 | 2B | 550847190 | -0.5951 | | 5.70E-07 | EMMAX | SUG |
| Vmax_17AY | AX-109412939 | 2B | 550847190 | 0.6032 | | 1.54E-08 | GEMMA | SIG |
| Vmax_17AY | AX-111058323 | 2B | 551398006 | 0.4058 | | 2.31E-06 | GEMMA | SUG |
| Vmax_17AY | AX-111714730 | 2B | 551445878 | -0.5986 | | 2.40E-07 | EMMAX | SUG |
| Vmax_17AY | AX-111714730 | 2B | 551445878 | 0.6048 | | 7.38E-09 | GEMMA | SIG |
| Vmax_17AY | AX-108891116 | 2B | 743332515 | -0.0857 | 0.2103 | 1.30E-16 | IIIVmrMLM | SIG |
| Vmax_17AY | AX-111065061 | 2B | 743546442 | 0.1267 | 0.01 | 1.56E-24 | IIIVmrMLM | SIG |
| Vmax_17AY | AX-110098913 | 2D | 317585511 | -3.0604 | | 1.24E-08 | EMMAX | SIG |
| Vmax_17AY | AX-108787002 | 2D | 326202355 | -3.0604 | | 1.24E-08 | EMMAX | SIG |
| Vmax_17AY | AX-109926918 | 2D | 494312828 | 0.0981 | | 1.04E-05 | BLUPmrMLM | SUG |
| Vmax_17AY | AX-109328166 | 2D | 508469235 | -3.0604 | | 1.24E-08 | EMMAX | SIG |
| Vmax_17AY | AX-111628091 | 3A | 12785355 | 0.2662 | | 2.23E-07 | GEMMA | SUG |
| Vmax_17AY | AX-111265000 | 3A | 12928360 | -0.0189 | -0.2581 | 3.53E-08 | IIIVmrMLM | SIG |
| Vmax_17AY | AX-108845007 | 3A | 12997270 | -0.2745 | | 2.08E-06 | EMMAX | SUG |
| Vmax_17AY | AX-108845007 | 3A | 12997270 | 0.2787 | | 3.38E-08 | GEMMA | SIG |
| Vmax_17AY | AX-95143496 | 3A | 12997719 | 0.2739 | | 4.55E-08 | GEMMA | SIG |
| Vmax_17AY | AX-110913445 | 3A | 12998870 | -0.2745 | | 2.08E-06 | EMMAX | SUG |
| Vmax_17AY | AX-110913445 | 3A | 12998870 | 0.2787 | | 3.38E-08 | GEMMA | SIG |
| Vmax_17AY | AX-89450584 | 3A | 12999035 | -0.2745 | | 2.08E-06 | EMMAX | SUG |
| Vmax_17AY | AX-89450584 | 3A | 12999035 | 0.2787 | | 3.38E-08 | GEMMA | SIG |
| Vmax_17AY | AX-109441093 | 3A | 13001330 | 0.2458 | | 2.66E-07 | GEMMA | SUG |
| Vmax_17AY | AX-111582837 | 3A | 13009071 | 0.2195 | | 1.57E-06 | GEMMA | SUG |
| Vmax_17AY | AX-111460884 | 3A | 13025627 | 0.2618 | | 6.28E-07 | GEMMA | SUG |

| | | | | | | | |
|---|---|---|---|---|---|---|---|
| Vmax_17AY | AX-110955079 | 3A | 13025941 | 0.2618 | 6.28E-07 | GEMMA | SUG |
| Vmax_17AY | AX-86184828 | 3A | 13038241 | -0.2780 | 1.80E-06 | EMMAX | SUG |
| Vmax_17AY | AX-86184828 | 3A | 13038241 | 0.2801 | 4.17E-08 | GEMMA | SIG |
| Vmax_17AY | AX-86166665 | 3A | 13038544 | 0.2686 | 1.69E-07 | GEMMA | SUG |
| Vmax_17AY | AX-86184778 | 3A | 13038599 | 0.2772 | 8.79E-08 | GEMMA | SIG |
| Vmax_17AY | AX-111589685 | 3A | 13042584 | 0.2844 | 5.14E-08 | GEMMA | SIG |
| Vmax_17AY | AX-108775363 | 3A | 13044766 | 0.2707 | 1.13E-07 | GEMMA | SIG |
| Vmax_17AY | AX-111065185 | 3A | 13078767 | 0.2503 | 2.25E-07 | GEMMA | SUG |
| Vmax_17AY | AX-110909396 | 3A | 13081297 | 0.5589 | 8.50E-08 | GEMMA | SIG |
| Vmax_17AY | AX-109346552 | 3A | 13190883 | -0.2804 | 9.94E-07 | EMMAX | SUG |
| Vmax_17AY | AX-109346552 | 3A | 13190883 | 0.2858 | 1.15E-08 | GEMMA | SIG |
| Vmax_17AY | AX-109878083 | 3A | 13269613 | 0.1805 | 1.89E-06 | GEMMA | SUG |
| Vmax_17AY | AX-109992464 | 3A | 13281571 | 0.1789 | 9.85E-07 | GEMMA | SUG |
| Vmax_17AY | AX-108729182 | 3A | 13345844 | 0.1787 | 1.63E-06 | GEMMA | SUG |
| Vmax_17AY | AX-110936651 | 3A | 13351271 | 0.1841 | 1.06E-06 | GEMMA | SUG |
| Vmax_17AY | AX-109890717 | 3A | 13353457 | 0.1785 | 1.73E-06 | GEMMA | SUG |
| Vmax_17AY | AX-109322880 | 3A | 13358458 | 0.1818 | 1.34E-06 | GEMMA | SUG |
| Vmax_17AY | AX-111217068 | 3A | 13365393 | 0.3785 | 5.86E-07 | GEMMA | SUG |
| Vmax_17AY | AX-111450027 | 3A | 13367521 | 0.1731 | 1.30E-06 | GEMMA | SUG |
| Vmax_17AY | AX-111499928 | 3A | 13394372 | 0.2109 | 6.68E-08 | GEMMA | SIG |
| Vmax_17AY | AX-108733909 | 3A | 13394434 | 0.1710 | 1.17E-06 | GEMMA | SUG |
| Vmax_17AY | AX-109465246 | 3A | 13399366 | 0.2050 | 1.31E-07 | GEMMA | SUG |
| Vmax_17AY | AX-108951347 | 3A | 13399645 | 0.2109 | 6.68E-08 | GEMMA | SIG |
| Vmax_17AY | AX-95098281 | 3A | 13405093 | 0.4086 | 1.66E-07 | GEMMA | SUG |
| Vmax_17AY | AX-110649779 | 3A | 13495516 | 0.3942 | 1.27E-07 | GEMMA | SUG |
| Vmax_17AY | AX-94523509 | 3A | 13496628 | 0.2062 | 1.44E-07 | GEMMA | SUG |
| Vmax_17AY | AX-111483558 | 3A | 13501374 | 0.3955 | 2.60E-07 | GEMMA | SUG |
| Vmax_17AY | AX-111730267 | 3A | 13503787 | 0.2062 | 1.44E-07 | GEMMA | SUG |
| Vmax_17AY | AX-89339551 | 3A | 13504175 | 0.2062 | 1.44E-07 | GEMMA | SUG |
| Vmax_17AY | AX-110535074 | 3A | 13511474 | 0.4140 | 2.08E-07 | GEMMA | SUG |
| Vmax_17AY | AX-110584359 | 3A | 13516636 | 0.4309 | 3.68E-08 | GEMMA | SIG |
| Vmax_17AY | AX-111062701 | 3A | 13517880 | 0.3711 | 4.95E-07 | GEMMA | SUG |
| Vmax_17AY | AX-110963955 | 3A | 13557708 | 0.3760 | 1.03E-06 | GEMMA | SUG |
| Vmax_17AY | AX-109977764 | 3A | 13559291 | 0.3714 | 1.25E-06 | GEMMA | SUG |
| Vmax_17AY | AX-94583236 | 3A | 13739466 | 0.2729 | 4.25E-08 | GEMMA | SIG |
| Vmax_17AY | AX-109362869 | 3A | 13739946 | 0.5695 | 2.32E-08 | GEMMA | SIG |
| Vmax_17AY | AX-109975558 | 3A | 13788160 | 0.3636 | 4.01E-08 | GEMMA | SIG |
| Vmax_17AY | AX-109285709 | 3A | 13901742 | 0.5966 | 1.41E-06 | GEMMA | SUG |
| Vmax_17AY | AX-111490647 | 3A | 13901791 | 0.6151 | 3.38E-07 | GEMMA | SUG |
| Vmax_17AY | AX-109839370 | 3A | 13971877 | 0.3090 | 8.12E-07 | GEMMA | SUG |
| Vmax_17AY | AX-109281788 | 3A | 14012613 | 0.2931 | 1.53E-06 | GEMMA | SUG |
| Vmax_17AY | AX-110415319 | 3A | 14013078 | 0.3090 | 8.12E-07 | GEMMA | SUG |
| Vmax_17AY | AX-108725274 | 3A | 14014210 | 0.3249 | 2.85E-07 | GEMMA | SUG |
| Vmax_17AY | AX-111714168 | 3A | 14042236 | 0.2931 | 1.53E-06 | GEMMA | SUG |

| | | | | | | | | |
|---|---|---|---|---|---|---|---|---|
| Vmax_17AY | AX-111502587 | 3A | 14042372 | 0.2931 | | 1.53E-06 | GEMMA | SUG |
| Vmax_17AY | AX-110470380 | 3A | 14042450 | 0.2931 | | 1.53E-06 | GEMMA | SUG |
| Vmax_17AY | AX-111543947 | 3A | 114792980 | -0.1603 | | 1.18E-04 | BLUPmrMLM | SUG |
| Vmax_17AY | AX-111543947 | 3A | 114792980 | 0.0669 | | 3.05E-09 | IIIVmrMLM | SIG |
| Vmax_17AY | AX-110130505 | 3A | 645611934 | 0.0165 | -0.4095 | 1.61E-09 | IIIVmrMLM | SIG |
| Vmax_17AY | AX-95023926 | 3A | 653243442 | 0.1291 | | 3.85E-28 | IIIVmrMLM | SIG |
| Vmax_17AY | AX-109458644 | 3A | 709377347 | -0.0127 | 0.4645 | 8.96E-06 | IIIVmrMLM | SUG |
| Vmax_17AY | AX-110391933 | 3A | 738905274 | -0.0123 | 0.6175 | 3.83E-08 | IIIVmrMLM | SIG |
| Vmax_17AY | AX-95110648 | 3A | 739559346 | -1.3597 | | 9.84E-07 | EMMAX | SUG |
| Vmax_17AY | AX-108904779 | 3B | 398377287 | 0.0595 | -0.1343 | 6.89E-10 | IIIVmrMLM | SIG |
| Vmax_17AY | AX-111236342 | 3B | 439703056 | -0.1017 | 0.0205 | 1.81E-14 | IIIVmrMLM | SIG |
| Vmax_17AY | AX-109481030 | 3B | 499814950 | -0.0981 | 0.8454 | 1.63E-20 | IIIVmrMLM | SIG |
| Vmax_17AY | AX-109427069 | 3B | 584577513 | -0.0091 | 0.3186 | 3.06E-07 | IIIVmrMLM | SUG |
| Vmax_17AY | AX-108873971 | 3B | 798550115 | -3.0604 | | 1.24E-08 | EMMAX | SIG |
| Vmax_17AY | AX-94662210 | 3B | 812572565 | -3.0604 | | 1.24E-08 | EMMAX | SIG |
| Vmax_17AY | AX-94487727 | 3B | 817009854 | -1.5688 | | 6.87E-09 | EMMAX | SIG |
| Vmax_17AY | AX-95254443 | 3D | 418396767 | -3.0604 | | 1.24E-08 | EMMAX | SIG |
| Vmax_17AY | AX-109450588 | 3D | 445486559 | -3.0604 | | 1.24E-08 | EMMAX | SIG |
| Vmax_17AY | AX-110392272 | 3D | 594847340 | -1.9135 | | 9.94E-07 | EMMAX | SUG |
| Vmax_17AY | AX-95232627 | 3D | 594891831 | -1.9135 | | 9.94E-07 | EMMAX | SUG |
| Vmax_17AY | AX-94871493 | 3D | 594891928 | -1.9135 | | 9.94E-07 | EMMAX | SUG |
| Vmax_17AY | AX-95238470 | 3D | 594891932 | -1.9135 | | 9.94E-07 | EMMAX | SUG |
| Vmax_17AY | AX-94928843 | 3D | 595066148 | -1.9135 | | 9.94E-07 | EMMAX | SUG |
| Vmax_17AY | AX-94856639 | 3D | 595066164 | -1.9135 | | 9.94E-07 | EMMAX | SUG |
| Vmax_17AY | AX-109834003 | 3D | 595115084 | -1.9135 | | 9.94E-07 | EMMAX | SUG |
| Vmax_17AY | AX-111697259 | 3D | 595187135 | -1.9135 | | 9.94E-07 | EMMAX | SUG |
| Vmax_17AY | AX-94603168 | 3D | 595271058 | -0.8788 | | 2.08E-06 | EMMAX | SUG |
| Vmax_17AY | AX-94924379 | 3D | 595271201 | -1.9135 | | 9.94E-07 | EMMAX | SUG |
| Vmax_17AY | AX-95180413 | 3D | 595271202 | -1.9135 | | 9.94E-07 | EMMAX | SUG |
| Vmax_17AY | AX-109008383 | 3D | 595273845 | -1.9135 | | 9.94E-07 | EMMAX | SUG |
| Vmax_17AY | AX-109394070 | 3D | 595279092 | -0.8788 | | 2.08E-06 | EMMAX | SUG |
| Vmax_17AY | AX-94874313 | 3D | 598011380 | -3.0604 | | 1.24E-08 | EMMAX | SIG |
| Vmax_17AY | AX-94959684 | 3D | 614366361 | 0.1900 | | 1.63E-05 | BLUPmrMLM | SUG |
| Vmax_17AY | AX-94569728 | 4B | 518641201 | -3.0604 | | 1.24E-08 | EMMAX | SIG |
| Vmax_17AY | AX-108999430 | 4B | 667683709 | -0.0783 | | 6.08E-05 | BLUPmrMLM | SUG |
| Vmax_17AY | AX-108790830 | 4D | 6672205 | -3.0604 | | 1.24E-08 | EMMAX | SIG |
| Vmax_17AY | AX-109498125 | 4D | 168900845 | -3.0604 | | 1.24E-08 | EMMAX | SIG |
| Vmax_17AY | AX-108756835 | 4D | 323706230 | -3.0604 | | 1.24E-08 | EMMAX | SIG |
| Vmax_17AY | AX-108994164 | 4D | 327427960 | -3.0604 | | 1.24E-08 | EMMAX | SIG |
| Vmax_17AY | AX-111745748 | 4D | 340296689 | -3.0604 | | 1.24E-08 | EMMAX | SIG |
| Vmax_17AY | AX-109534277 | 4D | 378260308 | -3.0604 | | 1.24E-08 | EMMAX | SIG |
| Vmax_17AY | AX-94967011 | 5A | 70498269 | -0.1075 | | 1.37E-20 | IIIVmrMLM | SIG |
| Vmax_17AY | AX-110167972 | 5A | 396754324 | -3.0604 | | 1.24E-08 | EMMAX | SIG |
| Vmax_17AY | AX-109318142 | 5A | 448043894 | -0.0677 | | 1.78E-04 | BLUPmrMLM | SUG |

| | | | | | | | | |
|---|---|---|---|---|---|---|---|---|
| Vmax_17AY | AX-110710951 | 5A | 472547282 | -0.7650 | | 3.54E-08 | EMMAX | SIG |
| Vmax_17AY | AX-110962408 | 5A | 476135701 | -0.0196 | -0.3233 | 1.77E-09 | IIIVmrMLM | SIG |
| Vmax_17AY | AX-94808117 | 5A | 476439578 | 0.2177 | | 3.12E-07 | BLUPmrMLM | SUG |
| Vmax_17AY | AX-110541590 | 5A | 548579019 | -0.0654 | 0.9416 | 8.09E-16 | IIIVmrMLM | SIG |
| Vmax_17AY | AX-109857340 | 5A | 548862216 | 0.0111 | 0.6037 | 3.83E-12 | IIIVmrMLM | SIG |
| Vmax_17AY | AX-110389287 | 5B | 391903643 | 0.0231 | 0.225 | 1.44E-17 | IIIVmrMLM | SIG |
| Vmax_17AY | AX-111151328 | 5B | 612941748 | -0.0234 | 1.7126 | 9.93E-11 | IIIVmrMLM | SIG |
| Vmax_17AY | AX-111073414 | 5B | 679746034 | -0.0461 | -0.2504 | 7.99E-17 | IIIVmrMLM | SIG |
| Vmax_17AY | AX-111699741 | 5B | 681078007 | 0.0448 | 0.9154 | 8.00E-12 | IIIVmrMLM | SIG |
| Vmax_17AY | AX-95658487 | 5B | 682699266 | 0.0988 | | 4.19E-06 | BLUPmrMLM | SUG |
| Vmax_17AY | AX-111577029 | 5B | 692522362 | 0.2479 | | 7.19E-08 | BLUPmrMLM | SIG |
| Vmax_17AY | AX-109312560 | 5D | 21550414 | 0.0310 | 0.7395 | 8.58E-11 | IIIVmrMLM | SIG |
| Vmax_17AY | AX-110313779 | 5D | 286455125 | -1.9300 | | 6.18E-07 | EMMAX | SUG |
| Vmax_17AY | AX-111216290 | 5D | 421594485 | 0.6074 | | 1.51E-07 | GEMMA | SUG |
| Vmax_17AY | AX-110616307 | 5D | 421619502 | 0.3086 | | 8.84E-08 | GEMMA | SIG |
| Vmax_17AY | AX-109509060 | 6A | 320926179 | 0.0034 | 0.353 | 2.12E-10 | IIIVmrMLM | SIG |
| Vmax_17AY | AX-108968801 | 6A | 599304441 | 0.0009 | 0.4101 | 5.38E-05 | IIIVmrMLM | SUG |
| Vmax_17AY | AX-94470972 | 6B | 226395363 | 0.9356 | 1.1779 | 0.00E+00 | IIIVmrMLM | SIG |
| Vmax_17AY | AX-109575981 | 6D | 331255908 | 0.0984 | | 1.07E-17 | IIIVmrMLM | SIG |
| Vmax_17AY | AX-110981313 | 6D | 423790976 | -3.0604 | | 1.24E-08 | EMMAX | SIG |
| Vmax_17AY | AX-108919084 | 6D | 470477263 | 0.0035 | -0.4612 | 2.01E-07 | IIIVmrMLM | SUG |
| Vmax_17AY | AX-94827671 | 7A | 11087367 | -0.0976 | | 2.71E-06 | BLUPmrMLM | SUG |
| Vmax_17AY | AX-108799491 | 7A | 267434161 | 0.0261 | 0.3759 | 9.16E-07 | IIIVmrMLM | SUG |
| Vmax_17AY | AX-109949014 | 7A | 511922399 | -0.0275 | 0.3537 | 5.86E-17 | IIIVmrMLM | SIG |
| Vmax_17AY | AX-112288950 | 7A | 538693397 | -0.2470 | 0.6369 | 1.98E-78 | IIIVmrMLM | SIG |
| Vmax_17AY | AX-109924132 | 7A | 594163470 | 0.0133 | 0.1119 | 1.63E-08 | IIIVmrMLM | SIG |
| Vmax_17AY | AX-94751384 | 7B | 35080930 | 0.2991 | | 6.28E-11 | BLUPmrMLM | SIG |
| Vmax_17AY | AX-111602188 | 7B | 63937125 | -3.0604 | | 1.24E-08 | EMMAX | SIG |
| Vmax_17AY | AX-94549402 | 7B | 641806996 | 0.0481 | | 1.71E-05 | IIIVmrMLM | SUG |
| Vmax_17AY | AX-108879449 | 7B | 643356884 | -0.0747 | | 6.11E-05 | BLUPmrMLM | SUG |
| Vmax_17AY | AX-111094331 | 7B | 723958130 | 0.5301 | | 1.91E-07 | GEMMA | SUG |
| Vmax_17AY | AX-95684144 | 7B | 725385135 | -1.3958 | | 3.47E-08 | EMMAX | SIG |
| Vmax_17AY | AX-94468441 | 7B | 725965878 | 0.2006 | | 2.49E-07 | GEMMA | SUG |
| Vmax_17AY | AX-110445652 | 7B | 726646189 | -1.3958 | | 3.47E-08 | EMMAX | SIG |
| Vmax_17AY | AX-95219160 | 7B | 727529846 | -1.3958 | | 3.47E-08 | EMMAX | SIG |
| Vmax_17AY | AX-108728735 | 7B | 727652888 | -0.9028 | | 3.65E-07 | EMMAX | SUG |
| Vmax_17AY | AX-108730446 | 7B | 729251407 | -1.3958 | | 3.47E-08 | EMMAX | SIG |
| Vmax_17AY | AX-109894291 | 7B | 734140942 | -0.0468 | 0.2895 | 1.54E-08 | IIIVmrMLM | SIG |
| Vmax_17AY | AX-111080832 | 7D | 83765607 | -3.0604 | | 1.24E-08 | EMMAX | SIG |
| Vmax_17AY | AX-108933675 | 7D | 309758149 | 0.5495 | | 1.71E-06 | GEMMA | SUG |
| Vmax_17AY | AX-110138076 | 7D | 390663125 | -3.0604 | | 1.24E-08 | EMMAX | SIG |
| Vmax_17AY | AX-111022532 | 7D | 577019498 | 0.0759 | -0.0575 | 2.16E-10 | IIIVmrMLM | SIG |
| Vmax_17AY | AX-94892803 | 7D | 625911375 | -0.0087 | 0.2689 | 1.07E-10 | IIIVmrMLM | SIG |
| Vmax_17JY | AX-94420005 | 1A | 3846233 | 0.1197 | | 7.60E-05 | BLUPmrMLM | SUG |

| | | | | | | | | |
|---|---|---|---|---|---|---|---|---|
| Vmax_17JY | AX-111090830 | 1A | 386777934 | -0.0266 | -0.4377 | 2.68E-06 | IIIVmrMLM | SUG |
| Vmax_17JY | AX-109965064 | 1A | 580958888 | 0.0366 | -0.5804 | 3.62E-07 | IIIVmrMLM | SUG |
| Vmax_17JY | AX-110932359 | 1A | 584539181 | 0.0778 | | 1.18E-04 | BLUPmrMLM | SUG |
| Vmax_17JY | AX-110020113 | 1A | 591298823 | -0.1256 | -0.0113 | 2.00E-15 | IIIVmrMLM | SIG |
| Vmax_17JY | AX-108811255 | 1B | 215088284 | 0.0382 | -0.2733 | 4.92E-06 | IIIVmrMLM | SUG |
| Vmax_17JY | AX-108786849 | 1B | 507809130 | 0.0646 | -0.4088 | 4.21E-08 | IIIVmrMLM | SIG |
| Vmax_17JY | AX-111467003 | 1B | 642958515 | -0.0241 | 0.5885 | 9.75E-08 | IIIVmrMLM | SIG |
| Vmax_17JY | AX-108802980 | 1B | 654101240 | -0.0217 | -0.9249 | 8.34E-10 | IIIVmrMLM | SIG |
| Vmax_17JY | AX-110393744 | 1B | 662836423 | -0.0247 | -0.3474 | 2.01E-08 | IIIVmrMLM | SIG |
| Vmax_17JY | AX-110399529 | 1B | 665798565 | -0.1464 | | 2.98E-07 | BLUPmrMLM | SUG |
| Vmax_17JY | AX-94526625 | 1D | 262341601 | -0.1455 | | 2.11E-21 | IIIVmrMLM | SIG |
| Vmax_17JY | AX-111101313 | 1D | 329717031 | -0.1839 | -0.2441 | 9.45E-31 | IIIVmrMLM | SIG |
| Vmax_17JY | AX-111059046 | 2A | 215822064 | 0.1816 | 0.549 | 1.09E-40 | IIIVmrMLM | SIG |
| Vmax_17JY | AX-110385168 | 2A | 440082976 | 0.1622 | | 1.20E-06 | BLUPmrMLM | SUG |
| Vmax_17JY | AX-111568790 | 2B | 49491732 | -0.0862 | | 1.08E-04 | BLUPmrMLM | SUG |
| Vmax_17JY | AX-109326360 | 2B | 450428262 | -0.0192 | -0.436 | 6.79E-07 | IIIVmrMLM | SUG |
| Vmax_17JY | AX-111010133 | 2D | 303387364 | -0.0444 | 0.5025 | 9.71E-08 | IIIVmrMLM | SIG |
| Vmax_17JY | AX-89625389 | 3A | 12974622 | -0.1382 | | 1.32E-19 | IIIVmrMLM | SIG |
| Vmax_17JY | AX-108844494 | 3A | 716095009 | 0.0222 | -0.3251 | 1.21E-04 | IIIVmrMLM | SUG |
| Vmax_17JY | AX-109449758 | 3B | 717049513 | 0.0429 | -0.3677 | 1.30E-06 | IIIVmrMLM | SUG |
| Vmax_17JY | AX-94569370 | 3D | 4394608 | 0.1938 | | 9.75E-07 | BLUPmrMLM | SUG |
| Vmax_17JY | AX-94569370 | 3D | 4394608 | -0.1737 | -0.0118 | 2.22E-24 | IIIVmrMLM | SIG |
| Vmax_17JY | AX-95148741 | 3D | 389631381 | -0.1371 | | 1.40E-04 | BLUPmrMLM | SUG |
| Vmax_17JY | AX-110023899 | 3D | 556830798 | -0.0350 | 1.041 | 5.81E-11 | IIIVmrMLM | SIG |
| Vmax_17JY | AX-109310695 | 5A | 566636735 | -0.4850 | | 1.18E-134 | IIIVmrMLM | SIG |
| Vmax_17JY | AX-111117839 | 5B | 56823336 | 0.0033 | 0.413 | 3.09E-08 | IIIVmrMLM | SIG |
| Vmax_17JY | AX-110961703 | 5B | 572668935 | 0.1629 | -0.1553 | 7.59E-25 | IIIVmrMLM | SIG |
| Vmax_17JY | AX-111476842 | 5B | 641801084 | -0.0796 | | 9.33E-06 | BLUPmrMLM | SUG |
| Vmax_17JY | AX-110992345 | 6A | 20872523 | -0.0665 | | 1.46E-04 | BLUPmrMLM | SUG |
| Vmax_17JY | AX-110196566 | 6A | 454111039 | 0.1361 | | 1.03E-07 | BLUPmrMLM | SIG |
| Vmax_17JY | AX-108891961 | 6A | 524764415 | -0.0166 | 0.4495 | 1.83E-04 | IIIVmrMLM | SUG |
| Vmax_17JY | AX-110566974 | 6A | 595069271 | 0.3211 | -0.0154 | 7.91E-72 | IIIVmrMLM | SIG |
| Vmax_17JY | AX-109979570 | 6B | 334222226 | 0.0136 | -0.1835 | 9.74E-04 | IIIVmrMLM | SUG |
| Vmax_17JY | AX-110599656 | 6B | 520286398 | 0.0438 | -0.3975 | 1.37E-06 | IIIVmrMLM | SUG |
| Vmax_17JY | AX-108893208 | 6D | 3920459 | -0.0730 | | 8.24E-05 | BLUPmrMLM | SUG |
| Vmax_17JY | AX-111988548 | 6D | 126021896 | 0.4352 | | 1.92E-06 | FarmCPU | SUG |
| Vmax_17JY | AX-111734855 | 6D | 140217420 | 0.4172 | | 1.53E-06 | FarmCPU | SUG |
| Vmax_17JY | AX-110650710 | 6D | 148794642 | 0.1116 | | 1.35E-13 | IIIVmrMLM | SIG |
| Vmax_17JY | AX-110575127 | 7A | 713173974 | 0.0282 | 1.0938 | 1.39E-07 | IIIVmrMLM | SUG |
| Vmax_17JY | AX-94965000 | 7B | 559146431 | -0.0256 | 0.148 | 3.72E-05 | IIIVmrMLM | SUG |
| Vmax_17JY | AX-86165716 | 7D | 146626182 | -0.4672 | -0.6318 | 3.11E-135 | IIIVmrMLM | SIG |
| Vmax_17JY | AX-111987065 | 7D | 267005167 | 0.0087 | 0.7219 | 1.02E-08 | IIIVmrMLM | SIG |
| Vmax_17JY | AX-95107918 | 7D | 576285018 | 0.1229 | | 1.27E-04 | BLUPmrMLM | SUG |
| Vmax_17JY | AX-110765221 | 7D | 616377712 | -0.0637 | | 1.78E-05 | IIIVmrMLM | SUG |

| | | | | | | | | |
|---|---|---|---|---|---|---|---|---|
| Vmax_17JY | AX-110395618 | 7D | 619989528 | 0.1023 | 0.333 | 1.48E-17 | IIIVmrMLM | SIG |
| Vmax_17JY | AX-110395618 | 7D | 619989528 | 0.4162 | | 1.86E-06 | GEMMA | SUG |
| Vmax_17SQ | AX-109100815 | 1A | 435081406 | 0.0224 | 0.2139 | 3.78E-17 | IIIVmrMLM | SIG |
| Vmax_17SQ | AX-111079784 | 1A | 446143086 | -0.0412 | | 1.15E-05 | IIIVmrMLM | SUG |
| Vmax_17SQ | AX-108765513 | 2A | 98899087 | -0.0114 | -0.935 | 6.33E-05 | IIIVmrMLM | SUG |
| Vmax_17SQ | AX-109533291 | 2A | 765374350 | -0.0677 | 0.0348 | 8.68E-11 | IIIVmrMLM | SIG |
| Vmax_17SQ | AX-109902042 | 2B | 721280529 | 0.0256 | 0.2133 | 1.02E-09 | IIIVmrMLM | SIG |
| Vmax_17SQ | AX-95128214 | 2D | 424000265 | 0.1093 | | 1.43E-04 | BLUPmrMLM | SUG |
| Vmax_17SQ | AX-94442651 | 2D | 507050616 | 0.2812 | | 4.35E-07 | FarmCPU | SUG |
| Vmax_17SQ | AX-111478343 | 3A | 224260678 | -0.0052 | 0.1746 | 7.32E-08 | IIIVmrMLM | SIG |
| Vmax_17SQ | AX-108893324 | 3A | 485014357 | -0.0609 | 0.0044 | 5.09E-07 | IIIVmrMLM | SUG |
| Vmax_17SQ | AX-110570407 | 3B | 410892577 | 0.0355 | 0.2078 | 3.41E-08 | IIIVmrMLM | SIG |
| Vmax_17SQ | AX-111501616 | 3D | 599430827 | 0.0315 | 0.5975 | 2.79E-12 | IIIVmrMLM | SIG |
| Vmax_17SQ | AX-94402318 | 3D | 609626811 | 0.0997 | -0.1061 | 1.43E-23 | IIIVmrMLM | SIG |
| Vmax_17SQ | AX-109420906 | 4A | 115634351 | 0.2678 | | 8.25E-07 | FarmCPU | SUG |
| Vmax_17SQ | AX-109092307 | 4A | 485173712 | -0.0583 | | 7.11E-10 | IIIVmrMLM | SIG |
| Vmax_17SQ | AX-111052325 | 4B | 282114533 | 0.1026 | | 2.19E-06 | BLUPmrMLM | SUG |
| Vmax_17SQ | AX-110409613 | 4B | 536088638 | 0.0303 | 0.4801 | 7.11E-11 | IIIVmrMLM | SIG |
| Vmax_17SQ | AX-110449465 | 4B | 579884727 | -0.2661 | | 1.82E-06 | GEMMA | SUG |
| Vmax_17SQ | AX-110604883 | 4D | 177489950 | 0.2641 | | 1.58E-06 | FarmCPU | SUG |
| Vmax_17SQ | AX-110528135 | 4D | 445004978 | 0.2574 | | 1.07E-06 | FarmCPU | SUG |
| Vmax_17SQ | AX-110686922 | 5A | 3402300 | -0.2315 | | 8.98E-07 | FarmCPU | SUG |
| Vmax_17SQ | AX-109845940 | 5A | 414824229 | -0.0140 | -0.1418 | 3.07E-07 | IIIVmrMLM | SUG |
| Vmax_17SQ | AX-108917596 | 5A | 518602100 | 0.3678 | | 1.34E-06 | GEMMA | SUG |
| Vmax_17SQ | AX-109652474 | 5A | 519654975 | 0.3599 | | 1.53E-06 | GEMMA | SUG |
| Vmax_17SQ | AX-111156514 | 5A | 519713020 | 0.3599 | | 1.53E-06 | GEMMA | SUG |
| Vmax_17SQ | AX-111462197 | 5A | 519932603 | 0.3758 | | 1.36E-06 | GEMMA | SUG |
| Vmax_17SQ | AX-111740483 | 5A | 520093132 | 0.3599 | | 1.53E-06 | GEMMA | SUG |
| Vmax_17SQ | AX-110468146 | 5A | 520095740 | 0.3599 | | 1.53E-06 | GEMMA | SUG |
| Vmax_17SQ | AX-108734074 | 5A | 520120297 | 0.3599 | | 1.53E-06 | GEMMA | SUG |
| Vmax_17SQ | AX-109967095 | 5A | 520377086 | 0.3599 | | 1.53E-06 | GEMMA | SUG |
| Vmax_17SQ | AX-109376123 | 5A | 520686648 | 0.3599 | | 1.53E-06 | GEMMA | SUG |
| Vmax_17SQ | AX-109620761 | 5B | 84374458 | 0.0278 | 0.1353 | 4.22E-09 | IIIVmrMLM | SIG |
| Vmax_17SQ | AX-111676830 | 5B | 302107387 | 0.0884 | | 2.39E-05 | BLUPmrMLM | SUG |
| Vmax_17SQ | AX-111676830 | 5B | 302107387 | -0.0596 | -0.0553 | 5.81E-09 | IIIVmrMLM | SIG |
| Vmax_17SQ | AX-109867574 | 5B | 458117972 | 0.0044 | -0.4462 | 9.83E-13 | IIIVmrMLM | SIG |
| Vmax_17SQ | AX-94700083 | 5B | 597813028 | 0.0287 | 0.1494 | 2.65E-12 | IIIVmrMLM | SIG |
| Vmax_17SQ | AX-109537450 | 5B | 692629908 | -0.0128 | 0.2921 | 2.22E-06 | IIIVmrMLM | SUG |
| Vmax_17SQ | AX-110910508 | 5B | 703099269 | -0.0155 | -0.1872 | 4.47E-06 | IIIVmrMLM | SUG |
| Vmax_17SQ | AX-111186232 | 5B | 711269599 | -0.0624 | 1.8212 | 3.27E-23 | IIIVmrMLM | SIG |
| Vmax_17SQ | AX-109831979 | 6B | 199930566 | -0.0599 | | 2.43E-05 | BLUPmrMLM | SUG |
| Vmax_17SQ | AX-109512365 | 6B | 529713648 | 0.0266 | 0.1867 | 1.44E-09 | IIIVmrMLM | SIG |
| Vmax_17SQ | AX-109315599 | 6B | 615533663 | 0.0821 | | 4.39E-05 | BLUPmrMLM | SUG |
| Vmax_17SQ | AX-109315599 | 6B | 615533663 | -0.0680 | -0.0105 | 4.87E-10 | IIIVmrMLM | SIG |

| | | | | | | | | |
|---|---|---|---|---|---|---|---|---|
| Vmax_17SQ | AX-89498965 | 6D | 471008558 | -0.0077 | 0.1633 | 4.53E-04 | IIIVmrMLM | SUG |
| Vmax_17SQ | AX-110954658 | 7A | 206721763 | -0.0080 | 0.6005 | 9.53E-09 | IIIVmrMLM | SIG |
| Vmax_17SQ | AX-110624620 | 7A | 247065085 | 0.0088 | -0.1847 | 1.28E-11 | IIIVmrMLM | SIG |
| Vmax_17SQ | AX-108908590 | 7A | 721572672 | 0.0245 | -0.4956 | 1.18E-10 | IIIVmrMLM | SIG |
| Vmax_17SQ | AX-109117982 | 7D | 41453283 | 0.1057 | | 3.57E-05 | BLUPmrMLM | SUG |
| Vmax_17SQ | AX-109117982 | 7D | 41453283 | -0.0688 | | 5.25E-13 | IIIVmrMLM | SIG |
| Vmax_17SQ | AX-110736759 | 7D | 173900640 | -0.0734 | -0.0648 | 2.42E-13 | IIIVmrMLM | SIG |
| Vmax_18AY | AX-94486559 | 1A | 15437125 | 0.0255 | -0.1081 | 1.85E-09 | IIIVmrMLM | SIG |
| Vmax_18AY | AX-109462275 | 1A | 15485453 | -0.0575 | | 4.89E-05 | BLUPmrMLM | SUG |
| Vmax_18AY | AX-111538498 | 1A | 460936979 | -0.0882 | | 8.10E-05 | BLUPmrMLM | SUG |
| Vmax_18AY | AX-111706902 | 1A | 534940298 | -0.0449 | | 5.92E-05 | BLUPmrMLM | SUG |
| Vmax_18AY | AX-95095232 | 1B | 2056936 | -0.0735 | | 2.90E-06 | BLUPmrMLM | SUG |
| Vmax_18AY | AX-111109428 | 1B | 332803747 | 0.1023 | | 2.00E-09 | BLUPmrMLM | SIG |
| Vmax_18AY | AX-89729848 | 2A | 36878667 | 0.0542 | | 4.86E-05 | BLUPmrMLM | SUG |
| Vmax_18AY | AX-110123869 | 2A | 284265347 | 0.0019 | -0.3042 | 2.44E-06 | IIIVmrMLM | SUG |
| Vmax_18AY | AX-109907808 | 2A | 752609186 | 0.0620 | | 1.35E-04 | BLUPmrMLM | SUG |
| Vmax_18AY | AX-111134442 | 2B | 183950769 | -0.0095 | -0.1608 | 1.14E-08 | IIIVmrMLM | SIG |
| Vmax_18AY | AX-109481993 | 2B | 388000896 | -0.0567 | | 2.06E-05 | BLUPmrMLM | SUG |
| Vmax_18AY | AX-94898754 | 2B | 461522925 | 0.0931 | | 1.02E-21 | IIIVmrMLM | SIG |
| Vmax_18AY | AX-109854926 | 3B | 216379514 | -0.0137 | 0.422 | 1.01E-11 | IIIVmrMLM | SIG |
| Vmax_18AY | AX-108912639 | 3B | 385142555 | -0.0106 | 0.2854 | 7.47E-08 | IIIVmrMLM | SIG |
| Vmax_18AY | AX-110098997 | 4A | 89943894 | 0.1733 | | 2.04E-06 | GEMMA | SUG |
| Vmax_18AY | AX-109627752 | 4A | 118747670 | 0.3213 | | 1.82E-06 | GEMMA | SUG |
| Vmax_18AY | AX-109348640 | 4A | 121207271 | 0.1605 | | 2.12E-06 | GEMMA | SUG |
| Vmax_18AY | AX-111140963 | 4A | 152807896 | 0.3297 | | 1.35E-06 | GEMMA | SUG |
| Vmax_18AY | AX-110132766 | 4A | 154908204 | 0.1644 | | 1.60E-06 | GEMMA | SUG |
| Vmax_18AY | AX-109950845 | 4A | 165384884 | 0.1723 | | 7.24E-07 | GEMMA | SUG |
| Vmax_18AY | AX-109862441 | 4A | 165707854 | 0.3297 | | 1.35E-06 | GEMMA | SUG |
| Vmax_18AY | AX-108917086 | 4A | 165753947 | 0.1644 | | 1.60E-06 | GEMMA | SUG |
| Vmax_18AY | AX-110556624 | 4A | 165832051 | 0.3054 | | 2.29E-06 | GEMMA | SUG |
| Vmax_18AY | AX-110524678 | 4A | 168795479 | 0.1660 | | 1.91E-06 | GEMMA | SUG |
| Vmax_18AY | AX-108804939 | 4A | 170832027 | 0.2160 | | 1.73E-06 | GEMMA | SUG |
| Vmax_18AY | AX-109934614 | 4A | 173925034 | 0.1644 | | 1.60E-06 | GEMMA | SUG |
| Vmax_18AY | AX-109576373 | 4A | 178621641 | 0.1644 | | 1.60E-06 | GEMMA | SUG |
| Vmax_18AY | AX-109850133 | 4A | 179500490 | 0.1670 | | 1.17E-06 | GEMMA | SUG |
| Vmax_18AY | AX-111137232 | 4A | 182924399 | 0.1644 | | 1.60E-06 | GEMMA | SUG |
| Vmax_18AY | AX-109903953 | 4A | 183698008 | 0.1644 | | 1.60E-06 | GEMMA | SUG |
| Vmax_18AY | AX-108814975 | 4A | 191126654 | 0.3297 | | 1.35E-06 | GEMMA | SUG |
| Vmax_18AY | AX-108764932 | 4A | 199388135 | 0.1644 | | 1.60E-06 | GEMMA | SUG |
| Vmax_18AY | AX-109881634 | 4A | 200166777 | 0.1644 | | 1.60E-06 | GEMMA | SUG |
| Vmax_18AY | AX-111048599 | 4A | 202117228 | 0.1644 | | 1.60E-06 | GEMMA | SUG |
| Vmax_18AY | AX-109417746 | 4A | 202608040 | 0.1644 | | 1.60E-06 | GEMMA | SUG |
| Vmax_18AY | AX-111496705 | 4A | 202952284 | 0.1644 | | 1.60E-06 | GEMMA | SUG |
| Vmax_18AY | AX-108756180 | 4A | 205235766 | 0.1644 | | 1.60E-06 | GEMMA | SUG |

| | | | | | | | |
|---|---|---|---|---|---|---|---|
| Vmax_18AY | AX-109048683 | 4A | 205377946 | 0.1644 | 1.60E-06 | GEMMA | SUG |
| Vmax_18AY | AX-109037943 | 4A | 210391364 | 0.1644 | 1.60E-06 | GEMMA | SUG |
| Vmax_18AY | AX-111597218 | 4A | 216177716 | 0.1644 | 1.60E-06 | GEMMA | SUG |
| Vmax_18AY | AX-109832235 | 4A | 216752232 | 0.1644 | 1.60E-06 | GEMMA | SUG |
| Vmax_18AY | AX-111485889 | 4A | 216760504 | 0.1644 | 1.60E-06 | GEMMA | SUG |
| Vmax_18AY | AX-108737265 | 4A | 219810302 | 0.3297 | 1.35E-06 | GEMMA | SUG |
| Vmax_18AY | AX-109436260 | 4A | 220767423 | 0.3297 | 1.35E-06 | GEMMA | SUG |
| Vmax_18AY | AX-109406924 | 4A | 222426885 | 0.1979 | 9.99E-07 | GEMMA | SUG |
| Vmax_18AY | AX-110947583 | 4A | 224071762 | 0.1670 | 1.17E-06 | GEMMA | SUG |
| Vmax_18AY | AX-108827766 | 4A | 226165765 | 0.1644 | 1.60E-06 | GEMMA | SUG |
| Vmax_18AY | AX-111003891 | 4A | 231847821 | 0.1644 | 1.60E-06 | GEMMA | SUG |
| Vmax_18AY | AX-110027861 | 4A | 234264820 | 0.3297 | 1.35E-06 | GEMMA | SUG |
| Vmax_18AY | AX-108849069 | 4A | 235835183 | 0.1723 | 7.24E-07 | GEMMA | SUG |
| Vmax_18AY | AX-109348374 | 4A | 240962502 | 0.1644 | 1.60E-06 | GEMMA | SUG |
| Vmax_18AY | AX-110994487 | 4A | 241399485 | 0.3297 | 1.35E-06 | GEMMA | SUG |
| Vmax_18AY | AX-111148636 | 4A | 241488851 | 0.2649 | 9.83E-08 | GEMMA | SIG |
| Vmax_18AY | AX-111080469 | 4A | 242675554 | 0.2107 | 1.70E-06 | GEMMA | SUG |
| Vmax_18AY | AX-109989814 | 4A | 247563880 | 0.1644 | 1.60E-06 | GEMMA | SUG |
| Vmax_18AY | AX-111234288 | 4A | 247742583 | 0.1644 | 1.60E-06 | GEMMA | SUG |
| Vmax_18AY | AX-111069459 | 4A | 248354619 | 0.1644 | 1.60E-06 | GEMMA | SUG |
| Vmax_18AY | AX-111786156 | 4A | 249516948 | 0.1670 | 1.17E-06 | GEMMA | SUG |
| Vmax_18AY | AX-110621563 | 4A | 250433654 | 0.1644 | 1.60E-06 | GEMMA | SUG |
| Vmax_18AY | AX-111254728 | 4A | 253472715 | 0.2943 | 2.31E-06 | GEMMA | SUG |
| Vmax_18AY | AX-111451653 | 4A | 253968218 | 0.1644 | 1.60E-06 | GEMMA | SUG |
| Vmax_18AY | AX-108848781 | 4A | 254047465 | 0.1644 | 1.60E-06 | GEMMA | SUG |
| Vmax_18AY | AX-110717688 | 4A | 254262580 | 0.1685 | 1.37E-06 | GEMMA | SUG |
| Vmax_18AY | AX-110504125 | 4A | 255733621 | 0.1651 | 1.64E-06 | GEMMA | SUG |
| Vmax_18AY | AX-109656061 | 4A | 256010896 | 0.1616 | 1.86E-06 | GEMMA | SUG |
| Vmax_18AY | AX-109416191 | 4A | 257588388 | 0.1990 | 3.52E-07 | GEMMA | SUG |
| Vmax_18AY | AX-110499119 | 4A | 257904346 | 0.1644 | 1.60E-06 | GEMMA | SUG |
| Vmax_18AY | AX-111618334 | 4A | 258155728 | 0.3297 | 1.35E-06 | GEMMA | SUG |
| Vmax_18AY | AX-109584923 | 4A | 260956090 | 0.1644 | 1.60E-06 | GEMMA | SUG |
| Vmax_18AY | AX-110507379 | 4A | 261266052 | 0.1644 | 1.60E-06 | GEMMA | SUG |
| Vmax_18AY | AX-111513759 | 4A | 261374169 | 0.3297 | 1.35E-06 | GEMMA | SUG |
| Vmax_18AY | AX-110058021 | 4A | 298848071 | 0.3297 | 1.35E-06 | GEMMA | SUG |
| Vmax_18AY | AX-110479627 | 4A | 299565039 | 0.1644 | 1.60E-06 | GEMMA | SUG |
| Vmax_18AY | AX-110533996 | 4A | 300798690 | 0.1687 | 1.81E-06 | GEMMA | SUG |
| Vmax_18AY | AX-111031413 | 4A | 300922187 | 0.1644 | 1.60E-06 | GEMMA | SUG |
| Vmax_18AY | AX-111777520 | 4A | 301102577 | 0.3297 | 1.35E-06 | GEMMA | SUG |
| Vmax_18AY | AX-110473064 | 4A | 311014961 | 0.1670 | 1.17E-06 | GEMMA | SUG |
| Vmax_18AY | AX-109290754 | 4A | 312684859 | 0.1670 | 1.17E-06 | GEMMA | SUG |
| Vmax_18AY | AX-111001770 | 4A | 322563896 | 0.3297 | 1.35E-06 | GEMMA | SUG |
| Vmax_18AY | AX-110958757 | 4A | 325474739 | 0.2199 | 3.22E-07 | GEMMA | SUG |
| Vmax_18AY | AX-111693754 | 4A | 327393524 | 0.1644 | 1.60E-06 | GEMMA | SUG |

| | | | | | | | |
|---|---|---|---|---|---|---|---|
| Vmax_18AY | AX-108846717 | 4A | 347384663 | 0.1644 | 1.60E-06 | GEMMA | SUG |
| Vmax_18AY | AX-111508837 | 4A | 347669842 | 0.1644 | 1.60E-06 | GEMMA | SUG |
| Vmax_18AY | AX-108814165 | 4A | 349145038 | 0.1644 | 1.60E-06 | GEMMA | SUG |
| Vmax_18AY | AX-110490021 | 4A | 351585543 | 0.1644 | 1.60E-06 | GEMMA | SUG |
| Vmax_18AY | AX-111610688 | 4A | 351793750 | 0.1644 | 1.60E-06 | GEMMA | SUG |
| Vmax_18AY | AX-108784614 | 4A | 352465279 | 0.1644 | 1.60E-06 | GEMMA | SUG |
| Vmax_18AY | AX-111002327 | 4A | 354312000 | 0.1663 | 1.88E-06 | GEMMA | SUG |
| Vmax_18AY | AX-109966106 | 4A | 354398208 | 0.1644 | 1.60E-06 | GEMMA | SUG |
| Vmax_18AY | AX-111113593 | 4A | 354639045 | 0.1644 | 1.60E-06 | GEMMA | SUG |
| Vmax_18AY | AX-111476244 | 4A | 355218984 | 0.1644 | 1.60E-06 | GEMMA | SUG |
| Vmax_18AY | AX-109834401 | 4A | 355542458 | 0.1644 | 1.60E-06 | GEMMA | SUG |
| Vmax_18AY | AX-108974178 | 4A | 361328820 | 0.1644 | 1.60E-06 | GEMMA | SUG |
| Vmax_18AY | AX-109841914 | 4A | 364921317 | 0.1644 | 1.60E-06 | GEMMA | SUG |
| Vmax_18AY | AX-108878378 | 4A | 365471237 | 0.3397 | 2.52E-07 | GEMMA | SUG |
| Vmax_18AY | AX-108843439 | 4A | 367914721 | 0.1691 | 1.05E-06 | GEMMA | SUG |
| Vmax_18AY | AX-110996352 | 4A | 372078526 | 0.3297 | 1.35E-06 | GEMMA | SUG |
| Vmax_18AY | AX-109301315 | 4A | 377616381 | 0.1776 | 1.23E-06 | GEMMA | SUG |
| Vmax_18AY | AX-109315481 | 4A | 378362609 | 0.1644 | 1.60E-06 | GEMMA | SUG |
| Vmax_18AY | AX-109417345 | 4A | 382707757 | 0.1652 | 1.32E-06 | GEMMA | SUG |
| Vmax_18AY | AX-108917729 | 4A | 383303509 | 0.3297 | 1.35E-06 | GEMMA | SUG |
| Vmax_18AY | AX-111546349 | 4A | 391298614 | 0.3297 | 1.35E-06 | GEMMA | SUG |
| Vmax_18AY | AX-110477191 | 4A | 391553751 | 0.1644 | 1.60E-06 | GEMMA | SUG |
| Vmax_18AY | AX-109396498 | 4A | 392090783 | 0.1644 | 1.60E-06 | GEMMA | SUG |
| Vmax_18AY | AX-111684041 | 4A | 398662457 | 0.1644 | 1.60E-06 | GEMMA | SUG |
| Vmax_18AY | AX-108821857 | 4A | 398891912 | 0.1644 | 1.60E-06 | GEMMA | SUG |
| Vmax_18AY | AX-109436757 | 4A | 401917799 | 0.1644 | 1.60E-06 | GEMMA | SUG |
| Vmax_18AY | AX-109032606 | 4A | 402103534 | 0.2883 | 1.19E-06 | GEMMA | SUG |
| Vmax_18AY | AX-109970978 | 4A | 404973142 | 0.2891 | 2.76E-07 | GEMMA | SUG |
| Vmax_18AY | AX-109861259 | 4A | 408289061 | 0.1807 | 6.22E-07 | GEMMA | SUG |
| Vmax_18AY | AX-110912869 | 4A | 410455724 | 0.1664 | 1.53E-06 | GEMMA | SUG |
| Vmax_18AY | AX-109417161 | 4A | 410673530 | 0.1644 | 1.60E-06 | GEMMA | SUG |
| Vmax_18AY | AX-110597403 | 4A | 410678628 | 0.1947 | 9.89E-07 | GEMMA | SUG |
| Vmax_18AY | AX-111505744 | 4A | 411525949 | 0.1644 | 1.60E-06 | GEMMA | SUG |
| Vmax_18AY | AX-111637659 | 4A | 414720487 | 0.1879 | 4.96E-07 | GEMMA | SUG |
| Vmax_18AY | AX-110376664 | 4A | 415037069 | 0.1644 | 1.60E-06 | GEMMA | SUG |
| Vmax_18AY | AX-109852990 | 4A | 415180213 | 0.3291 | 1.06E-06 | GEMMA | SUG |
| Vmax_18AY | AX-110132259 | 4A | 415855084 | 0.1896 | 4.48E-07 | GEMMA | SUG |
| Vmax_18AY | AX-110465853 | 4A | 419441081 | 0.3297 | 1.35E-06 | GEMMA | SUG |
| Vmax_18AY | AX-110716657 | 4A | 422612361 | 0.3297 | 1.35E-06 | GEMMA | SUG |
| Vmax_18AY | AX-111222147 | 4A | 427268358 | 0.1644 | 1.60E-06 | GEMMA | SUG |
| Vmax_18AY | AX-110126645 | 4A | 427783849 | 0.1691 | 1.05E-06 | GEMMA | SUG |
| Vmax_18AY | AX-111112454 | 4A | 430045992 | 0.1692 | 2.18E-06 | GEMMA | SUG |
| Vmax_18AY | AX-111555450 | 4A | 434439205 | 0.3297 | 1.35E-06 | GEMMA | SUG |
| Vmax_18AY | AX-110471402 | 4A | 437431457 | 0.1673 | 1.12E-06 | GEMMA | SUG |

| | | | | | | | | |
|---|---|---|---|---|---|---|---|---|
| Vmax_18AY | AX-111651709 | 4A | 438249689 | 0.1644 | | 1.60E-06 | GEMMA | SUG |
| Vmax_18AY | AX-111558966 | 4A | 438622700 | 0.1644 | | 1.60E-06 | GEMMA | SUG |
| Vmax_18AY | AX-108809010 | 4A | 438993497 | 0.1644 | | 1.60E-06 | GEMMA | SUG |
| Vmax_18AY | AX-111461142 | 4A | 439495774 | 0.3297 | | 1.35E-06 | GEMMA | SUG |
| Vmax_18AY | AX-111666849 | 4A | 440161449 | 0.1644 | | 1.60E-06 | GEMMA | SUG |
| Vmax_18AY | AX-108798226 | 4A | 440920222 | 0.1644 | | 1.60E-06 | GEMMA | SUG |
| Vmax_18AY | AX-109058806 | 4A | 441126306 | 0.2500 | | 8.06E-07 | GEMMA | SUG |
| Vmax_18AY | AX-109901098 | 4A | 448108988 | 0.1644 | | 1.60E-06 | GEMMA | SUG |
| Vmax_18AY | AX-110522872 | 4A | 448656377 | 0.1639 | | 1.50E-06 | GEMMA | SUG |
| Vmax_18AY | AX-110541157 | 4A | 626510311 | -0.0132 | 0.634 | 1.33E-07 | IIIVmrMLM | SUG |
| Vmax_18AY | AX-108943948 | 4B | 431262827 | 0.0266 | 0.2608 | 1.02E-05 | IIIVmrMLM | SUG |
| Vmax_18AY | AX-89362528 | 4B | 637390229 | 0.0641 | | 1.29E-05 | BLUPmrMLM | SUG |
| Vmax_18AY | AX-89362528 | 4B | 637390229 | -0.0726 | -0.1231 | 1.98E-13 | IIIVmrMLM | SIG |
| Vmax_18AY | AX-109302905 | 4B | 669662753 | -0.0793 | 0.0348 | 4.92E-11 | IIIVmrMLM | SIG |
| Vmax_18AY | AX-108976919 | 4B | 672563232 | 0.0802 | -0.1589 | 2.57E-16 | IIIVmrMLM | SIG |
| Vmax_18AY | AX-111176492 | 5A | 364981589 | 0.0685 | | 2.64E-05 | BLUPmrMLM | SUG |
| Vmax_18AY | AX-111480358 | 5A | 521939210 | -0.0133 | -0.198 | 9.34E-11 | IIIVmrMLM | SIG |
| Vmax_18AY | AX-110044503 | 5A | 528288061 | -0.0211 | -0.4485 | 2.51E-11 | IIIVmrMLM | SIG |
| Vmax_18AY | AX-110941281 | 5A | 588022791 | 0.0557 | -0.0113 | 1.07E-07 | IIIVmrMLM | SIG |
| Vmax_18AY | AX-110123916 | 5A | 588401903 | 0.0197 | 0.2041 | 2.37E-05 | IIIVmrMLM | SUG |
| Vmax_18AY | AX-111465232 | 5A | 698495514 | -0.0053 | 0.11 | 4.29E-09 | IIIVmrMLM | SIG |
| Vmax_18AY | AX-109542966 | 5B | 470978512 | -0.1045 | | 6.28E-06 | BLUPmrMLM | SUG |
| Vmax_18AY | AX-108866638 | 5B | 514929856 | 0.0577 | | 1.31E-09 | IIIVmrMLM | SIG |
| Vmax_18AY | AX-108840042 | 5D | 42961528 | 0.0535 | 0.188 | 8.70E-09 | IIIVmrMLM | SIG |
| Vmax_18AY | AX-94460640 | 5D | 220098422 | 0.0558 | | 4.08E-09 | IIIVmrMLM | SIG |
| Vmax_18AY | AX-111548703 | 6A | 13901452 | 0.0085 | 0.2941 | 3.66E-07 | IIIVmrMLM | SUG |
| Vmax_18AY | AX-109846550 | 6A | 79410206 | -0.0630 | | 6.25E-05 | BLUPmrMLM | SUG |
| Vmax_18AY | AX-109872058 | 6A | 97892055 | -0.0171 | 0.1584 | 9.33E-06 | IIIVmrMLM | SUG |
| Vmax_18AY | AX-109840004 | 6B | 290011424 | 0.0130 | 0.5274 | 1.22E-05 | IIIVmrMLM | SUG |
| Vmax_18AY | AX-109528715 | 6B | 630548604 | -0.0306 | 0.1685 | 1.27E-07 | IIIVmrMLM | SUG |
| Vmax_18AY | AX-108758805 | 6B | 643139508 | -0.0044 | 0.4929 | 1.09E-05 | IIIVmrMLM | SUG |
| Vmax_18AY | AX-109403214 | 7A | 103149209 | 0.0033 | 0.1677 | 9.41E-08 | IIIVmrMLM | SIG |
| Vmax_18AY | AX-111031010 | 7A | 604702756 | -0.0614 | -0.1903 | 8.63E-19 | IIIVmrMLM | SIG |
| Vmax_18AY | AX-110414931 | 7A | 663225859 | -0.0032 | 0.8457 | 1.60E-06 | IIIVmrMLM | SUG |
| Vmax_18AY | AX-109045123 | 7B | 134224402 | 0.0749 | | 5.90E-15 | IIIVmrMLM | SIG |
| Vmax_18AY | AX-94403275 | 7B | 718148246 | 0.0150 | -0.2398 | 9.19E-05 | IIIVmrMLM | SUG |
| Vmax_18JY | AX-111733802 | 1A | 353914935 | 0.0027 | 0.405 | 3.22E-10 | IIIVmrMLM | SIG |
| Vmax_18JY | AX-110991546 | 1A | 549300554 | -0.0148 | 0.1891 | 1.08E-07 | IIIVmrMLM | SIG |
| Vmax_18JY | AX-110397108 | 1B | 661046954 | 0.0296 | -0.1181 | 7.46E-10 | IIIVmrMLM | SIG |
| Vmax_18JY | AX-109299102 | 1D | 25233854 | -1.7052 | | 7.71E-08 | EMMAX | SIG |
| Vmax_18JY | AX-110656149 | 1D | 28582081 | 0.0868 | | 7.02E-15 | IIIVmrMLM | SIG |
| Vmax_18JY | AX-109960546 | 2A | 716906727 | 0.0416 | -0.1786 | 1.87E-07 | IIIVmrMLM | SUG |
| Vmax_18JY | AX-109844474 | 2B | 29469163 | 0.0647 | -0.0677 | 2.18E-08 | IIIVmrMLM | SIG |
| Vmax_18JY | AX-109926170 | 2B | 131130216 | -0.0189 | 0.1876 | 2.94E-06 | IIIVmrMLM | SUG |

| | | | | | | | | |
|---|---|---|---|---|---|---|---|---|
| Vmax_18JY | AX-110530420 | 2B | 193629245 | 0.0172 | 0.3752 | 2.12E-09 | IIIVmrMLM | SIG |
| Vmax_18JY | AX-110571449 | 2B | 705724336 | 0.0720 | 0.3926 | 1.59E-17 | IIIVmrMLM | SIG |
| Vmax_18JY | AX-108804627 | 2B | 749138296 | 0.0353 | -0.2008 | 6.97E-09 | IIIVmrMLM | SIG |
| Vmax_18JY | AX-94761755 | 2D | 76896040 | -2.3690 | | 1.15E-07 | EMMAX | SIG |
| Vmax_18JY | AX-110572284 | 2D | 119668606 | -2.3690 | | 1.15E-07 | EMMAX | SIG |
| Vmax_18JY | AX-89685490 | 2D | 152627670 | 0.0771 | -0.0936 | 2.57E-11 | IIIVmrMLM | SIG |
| Vmax_18JY | AX-108855254 | 2D | 402851647 | 0.0964 | | 8.84E-18 | IIIVmrMLM | SIG |
| Vmax_18JY | AX-108974236 | 2D | 433222791 | 0.0823 | | 2.31E-05 | BLUPmrMLM | SUG |
| Vmax_18JY | AX-110721742 | 3A | 101275805 | 0.1328 | | 6.08E-05 | BLUPmrMLM | SUG |
| Vmax_18JY | AX-110472635 | 3A | 709437566 | 0.0396 | -0.1197 | 1.94E-07 | IIIVmrMLM | SUG |
| Vmax_18JY | AX-110668812 | 3A | 712465609 | 0.2854 | | 1.29E-06 | FarmCPU | SUG |
| Vmax_18JY | AX-110992633 | 3A | 738753561 | 0.0651 | 0.1506 | 2.29E-08 | IIIVmrMLM | SIG |
| Vmax_18JY | AX-94437164 | 3B | 112281346 | -0.0979 | | 2.91E-18 | IIIVmrMLM | SIG |
| Vmax_18JY | AX-111154757 | 3B | 362049509 | -0.0303 | -0.1713 | 1.15E-13 | IIIVmrMLM | SIG |
| Vmax_18JY | AX-110438296 | 3B | 386149088 | -0.0054 | -0.2846 | 6.94E-09 | IIIVmrMLM | SIG |
| Vmax_18JY | AX-94734909 | 3B | 417714428 | -2.3690 | | 1.15E-07 | EMMAX | SIG |
| Vmax_18JY | AX-111711959 | 3B | 547054567 | -0.1020 | -0.0015 | 2.72E-17 | IIIVmrMLM | SIG |
| Vmax_18JY | AX-111745200 | 3B | 595268176 | -2.3690 | | 1.15E-07 | EMMAX | SIG |
| Vmax_18JY | AX-109889124 | 3B | 809339558 | -2.3690 | | 1.15E-07 | EMMAX | SIG |
| Vmax_18JY | AX-111598404 | 3B | 828424807 | -0.0188 | -0.1683 | 6.01E-05 | IIIVmrMLM | SUG |
| Vmax_18JY | AX-94688609 | 3D | 322439775 | -2.3690 | | 1.15E-07 | EMMAX | SIG |
| Vmax_18JY | AX-94902788 | 3D | 445021233 | -2.3690 | | 1.15E-07 | EMMAX | SIG |
| Vmax_18JY | AX-110960487 | 3D | 450883789 | -1.7943 | | 1.08E-08 | EMMAX | SIG |
| Vmax_18JY | AX-108775096 | 3D | 601186984 | 0.0061 | -0.5734 | 3.31E-07 | IIIVmrMLM | SUG |
| Vmax_18JY | AX-94419071 | 3D | 603536051 | -2.3690 | | 1.15E-07 | EMMAX | SIG |
| Vmax_18JY | AX-109283680 | 4A | 92725776 | -0.0019 | 0.6553 | 2.43E-06 | IIIVmrMLM | SUG |
| Vmax_18JY | AX-111572808 | 4A | 404228421 | -0.0084 | -0.5918 | 2.59E-04 | IIIVmrMLM | SUG |
| Vmax_18JY | AX-111010398 | 4A | 566568943 | 0.0125 | 0.2386 | 4.73E-20 | IIIVmrMLM | SIG |
| Vmax_18JY | AX-109440771 | 4D | 31447593 | -2.3690 | | 1.15E-07 | EMMAX | SIG |
| Vmax_18JY | AX-111053158 | 4D | 176701905 | -2.3690 | | 1.15E-07 | EMMAX | SIG |
| Vmax_18JY | AX-111488058 | 4D | 326770951 | -2.3690 | | 1.15E-07 | EMMAX | SIG |
| Vmax_18JY | AX-110967587 | 4D | 403181432 | -2.3690 | | 1.15E-07 | EMMAX | SIG |
| Vmax_18JY | AX-111639412 | 5A | 679775437 | 0.0127 | -0.0924 | 5.32E-04 | IIIVmrMLM | SUG |
| Vmax_18JY | AX-108882297 | 5B | 66663744 | 0.0353 | 0.252 | 1.28E-06 | IIIVmrMLM | SUG |
| Vmax_18JY | AX-110060332 | 5B | 70091302 | 0.0400 | 0.2546 | 1.28E-11 | IIIVmrMLM | SIG |
| Vmax_18JY | AX-110046875 | 5B | 531406678 | -0.0216 | 0.1625 | 1.79E-06 | IIIVmrMLM | SUG |
| Vmax_18JY | AX-111764169 | 5B | 669925043 | 0.0127 | 0.6705 | 2.75E-08 | IIIVmrMLM | SIG |
| Vmax_18JY | AX-94568716 | 5D | 278636565 | -2.3690 | | 1.15E-07 | EMMAX | SIG |
| Vmax_18JY | AX-111547894 | 6A | 16378761 | 0.0090 | 0.3394 | 2.52E-08 | IIIVmrMLM | SIG |
| Vmax_18JY | AX-111728091 | 6A | 599718262 | 0.2131 | | 7.44E-07 | FarmCPU | SUG |
| Vmax_18JY | AX-110536078 | 6B | 482068535 | -0.0134 | 0.3609 | 1.01E-15 | IIIVmrMLM | SIG |
| Vmax_18JY | AX-94707298 | 6B | 662560780 | -2.3690 | | 1.15E-07 | EMMAX | SIG |
| Vmax_18JY | AX-95113401 | 6B | 716437491 | -0.0962 | | 9.52E-06 | BLUPmrMLM | SUG |
| Vmax_18JY | AX-95113401 | 6B | 716437491 | 0.0920 | 0.038 | 1.93E-09 | IIIVmrMLM | SIG |

| | | | | | | | | |
|---|---|---|---|---|---|---|---|---|
| Vmax_18JY | AX-110065115 | 6D | 85790623 | 0.0121 | 0.8254 | 1.28E-07 | IIIVmrMLM | SUG |
| Vmax_18JY | AX-94502471 | 6D | 453347870 | -2.3690 | | 1.15E-07 | EMMAX | SIG |
| Vmax_18JY | AX-110376969 | 6D | 454221557 | -2.3690 | | 1.15E-07 | EMMAX | SIG |
| Vmax_18JY | AX-111235557 | 7A | 451904323 | 0.0284 | -0.3899 | 5.38E-09 | IIIVmrMLM | SIG |
| Vmax_18JY | AX-94743071 | 7A | 541439144 | -2.3690 | | 1.15E-07 | EMMAX | SIG |
| Vmax_18JY | AX-108765333 | 7B | 4908706 | -0.0745 | | 7.39E-05 | BLUPmrMLM | SUG |
| Vmax_18JY | AX-109478037 | 7B | 159009520 | 0.0247 | 0.2282 | 1.42E-16 | IIIVmrMLM | SIG |
| Vmax_18JY | AX-110380171 | 7B | 239172527 | 0.0360 | 1.0635 | 9.64E-07 | IIIVmrMLM | SUG |
| Vmax_18JY | AX-109868959 | 7B | 704252664 | 0.0585 | 0.0299 | 2.35E-06 | IIIVmrMLM | SUG |
| Vmax_18JY | AX-109297242 | 7D | 27656271 | -2.3690 | | 1.15E-07 | EMMAX | SIG |
| Vmax_18JY | AX-110949339 | 7D | 119758051 | -0.0090 | 0.706 | 2.38E-07 | IIIVmrMLM | SUG |
| Vmax_18JY | AX-111011064 | 7D | 388130551 | -2.3690 | | 1.15E-07 | EMMAX | SIG |
| Vmax_18JY | AX-109660239 | 7D | 529065509 | -0.2720 | | 1.12E-08 | FarmCPU | SIG |
| Vmax_18JY | AX-94503003 | 7D | 602183791 | -2.3690 | | 1.15E-07 | EMMAX | SIG |
| Vmax_19XX | AX-109281152 | 1A | 9546451 | 0.0616 | -0.0789 | 1.39E-05 | IIIVmrMLM | SUG |
| Vmax_19XX | AX-109831162 | 1A | 101789877 | -0.0942 | -0.1953 | 1.90E-10 | IIIVmrMLM | SIG |
| Vmax_19XX | AX-109989800 | 1A | 327541838 | -0.0181 | 0.6446 | 4.60E-07 | IIIVmrMLM | SUG |
| Vmax_19XX | AX-111091775 | 1A | 436447126 | 0.0215 | 0.2335 | 4.57E-07 | IIIVmrMLM | SUG |
| Vmax_19XX | AX-111800966 | 1A | 491390772 | 0.1556 | | 6.66E-06 | BLUPmrMLM | SUG |
| Vmax_19XX | AX-108887078 | 1A | 584583811 | -0.0240 | -0.2252 | 2.19E-07 | IIIVmrMLM | SUG |
| Vmax_19XX | AX-95659604 | 1B | 645572033 | 0.0267 | 0.7139 | 4.23E-06 | IIIVmrMLM | SUG |
| Vmax_19XX | AX-111756851 | 2A | 365311747 | 0.0961 | | 1.73E-05 | BLUPmrMLM | SUG |
| Vmax_19XX | AX-110959473 | 2B | 29559393 | 0.0987 | | 9.34E-06 | BLUPmrMLM | SUG |
| Vmax_19XX | AX-110959473 | 2B | 29559393 | -0.0706 | 0.0794 | 3.03E-06 | IIIVmrMLM | SUG |
| Vmax_19XX | AX-111002945 | 2D | 19505527 | -0.0926 | 0.0743 | 1.33E-09 | IIIVmrMLM | SIG |
| Vmax_19XX | AX-111540433 | 3A | 506400798 | -0.0128 | 0.2794 | 9.56E-04 | IIIVmrMLM | SUG |
| Vmax_19XX | AX-109288496 | 3A | 515727225 | -0.0115 | -0.3534 | 1.68E-06 | IIIVmrMLM | SUG |
| Vmax_19XX | AX-109902599 | 3A | 584738239 | 0.1435 | 0.4412 | 1.85E-22 | IIIVmrMLM | SIG |
| Vmax_19XX | AX-111561859 | 3A | 712203223 | -0.0269 | -0.2494 | 1.40E-07 | IIIVmrMLM | SUG |
| Vmax_19XX | AX-94795166 | 3B | 78598490 | 0.2460 | | 3.87E-07 | GEMMA | SUG |
| Vmax_19XX | AX-111142307 | 3B | 289732545 | -0.0247 | 0.1767 | 3.22E-05 | IIIVmrMLM | SUG |
| Vmax_19XX | AX-110632987 | 4A | 13251112 | 0.0074 | -0.2032 | 1.37E-06 | IIIVmrMLM | SUG |
| Vmax_19XX | AX-110538379 | 4A | 540539813 | -0.0209 | -0.377 | 1.50E-04 | IIIVmrMLM | SUG |
| Vmax_19XX | AX-111021866 | 4A | 615159971 | 0.1250 | 0.0835 | 2.99E-16 | IIIVmrMLM | SIG |
| Vmax_19XX | AX-94657107 | 4B | 78485318 | -0.1212 | | 1.83E-04 | BLUPmrMLM | SUG |
| Vmax_19XX | AX-89696150 | 5A | 6725669 | 0.0911 | | 4.47E-05 | BLUPmrMLM | SUG |
| Vmax_19XX | AX-95092715 | 5A | 99416023 | 0.1349 | | 1.97E-20 | IIIVmrMLM | SIG |
| Vmax_19XX | AX-95134882 | 5A | 309683378 | -0.0898 | | 3.08E-10 | IIIVmrMLM | SIG |
| Vmax_19XX | AX-111526971 | 5A | 476652631 | -0.0957 | | 2.26E-05 | BLUPmrMLM | SUG |
| Vmax_19XX | AX-108954804 | 5A | 587653258 | -0.0093 | -0.1579 | 1.46E-07 | IIIVmrMLM | SUG |
| Vmax_19XX | AX-111615781 | 5A | 622545850 | -0.0086 | 0.592 | 1.69E-06 | IIIVmrMLM | SUG |
| Vmax_19XX | AX-94657293 | 5B | 45392491 | 0.0346 | 0.2801 | 3.06E-08 | IIIVmrMLM | SIG |
| Vmax_19XX | AX-110537485 | 5B | 517694271 | -0.0669 | 0.401 | 2.18E-13 | IIIVmrMLM | SIG |
| Vmax_19XX | AX-111764169 | 5B | 669925043 | -0.0413 | 0.469 | 5.87E-05 | IIIVmrMLM | SUG |

| | | | | | | | | |
|---|---|---|---|---|---|---|---|---|
| Vmax_19XX | AX-110616489 | 5B | 689901902 | 0.0829 | | 2.86E-05 | BLUPmrMLM | SUG |
| Vmax_19XX | AX-110923367 | 5B | 691503490 | 0.0218 | 0.2748 | 5.74E-08 | IIIVmrMLM | SIG |
| Vmax_19XX | AX-111620734 | 6B | 4125682 | -0.0154 | 0.5942 | 4.36E-06 | IIIVmrMLM | SUG |
| Vmax_19XX | AX-111523255 | 6B | 28003889 | 0.0291 | 0.3154 | 5.71E-07 | IIIVmrMLM | SUG |
| Vmax_19XX | AX-110095302 | 6D | 464345566 | 0.0022 | 0.2031 | 2.14E-11 | IIIVmrMLM | SIG |
| Vmax_19XX | AX-109876286 | 7A | 20672125 | -0.0902 | | 3.29E-05 | BLUPmrMLM | SUG |
| Vmax_19XX | AX-109964404 | 7A | 50020938 | 0.0606 | 0.2565 | 4.41E-09 | IIIVmrMLM | SIG |
| Vmax_19XX | AX-111631026 | 7A | 561690250 | 0.0011 | -0.1644 | 4.82E-05 | IIIVmrMLM | SUG |
| Vmax_19XX | AX-111657234 | 7A | 642884672 | 0.0737 | | 1.35E-04 | BLUPmrMLM | SUG |
| Vmax_19XX | AX-108880866 | 7B | 51627614 | -0.0873 | -0.1157 | 3.17E-09 | IIIVmrMLM | SIG |
| Vmax_19XX | AX-108814125 | 7B | 156735386 | 0.0224 | 1.0866 | 1.26E-07 | IIIVmrMLM | SUG |
| Vmax_19XX | AX-108906914 | 7B | 604035802 | 0.0262 | 0.413 | 2.25E-06 | IIIVmrMLM | SUG |
| Vmax_BLUP | AX-110020173 | 1A | 493588916 | -0.0124 | 0.0842 | 2.86E-07 | IIIVmrMLM | SUG |
| Vmax_BLUP | AX-110932359 | 1A | 584539181 | 0.0325 | | 7.04E-09 | BLUPmrMLM | SIG |
| Vmax_BLUP | AX-94684170 | 1B | 597537011 | 0.1489 | | 1.10E-06 | FarmCPU | SUG |
| Vmax_BLUP | AX-109915945 | 1B | 666642701 | -0.0237 | | 4.04E-05 | BLUPmrMLM | SUG |
| Vmax_BLUP | AX-109915945 | 1B | 666642701 | 0.0352 | -0.0113 | 4.72E-13 | IIIVmrMLM | SIG |
| Vmax_BLUP | AX-110991342 | 1B | 686855959 | 0.0339 | 0.0122 | 6.15E-07 | IIIVmrMLM | SUG |
| Vmax_BLUP | AX-110965430 | 1D | 88379613 | -0.0097 | 0.2557 | 3.88E-07 | IIIVmrMLM | SUG |
| Vmax_BLUP | AX-94490686 | 1D | 493705285 | 0.0127 | 0.086 | 1.01E-06 | IIIVmrMLM | SUG |
| Vmax_BLUP | AX-112287088 | 2A | 14940390 | -0.0001 | -0.051 | 3.13E-04 | IIIVmrMLM | SUG |
| Vmax_BLUP | AX-109973922 | 2A | 504467482 | 0.0006 | -0.2058 | 1.86E-08 | IIIVmrMLM | SIG |
| Vmax_BLUP | AX-111071776 | 2B | 18193542 | 0.0240 | | 1.48E-04 | BLUPmrMLM | SUG |
| Vmax_BLUP | AX-108862346 | 2B | 327656136 | 0.1379 | | 2.26E-06 | FarmCPU | SUG |
| Vmax_BLUP | AX-110009814 | 2B | 433967938 | 0.0060 | 0.0421 | 1.02E-04 | IIIVmrMLM | SUG |
| Vmax_BLUP | AX-110370957 | 2B | 603911965 | 0.0285 | | 4.69E-05 | BLUPmrMLM | SUG |
| Vmax_BLUP | AX-109036011 | 2D | 289381694 | 0.1481 | | 1.33E-06 | FarmCPU | SUG |
| Vmax_BLUP | AX-94442651 | 2D | 507050616 | 0.1738 | | 3.27E-08 | FarmCPU | SIG |
| Vmax_BLUP | AX-110546630 | 3A | 7581946 | -0.0067 | 0.0773 | 1.54E-06 | IIIVmrMLM | SUG |
| Vmax_BLUP | AX-95215706 | 3A | 311986520 | 0.1484 | | 1.84E-06 | FarmCPU | SUG |
| Vmax_BLUP | AX-109286245 | 3A | 421119830 | 0.1257 | | 6.75E-07 | FarmCPU | SUG |
| Vmax_BLUP | AX-95184677 | 3A | 510007314 | -0.0639 | | 2.13E-06 | GEMMA | SUG |
| Vmax_BLUP | AX-109274638 | 3B | 237325305 | -0.1318 | | 4.24E-07 | FarmCPU | SUG |
| Vmax_BLUP | AX-109880624 | 3B | 370136318 | 0.1668 | | 1.31E-06 | FarmCPU | SUG |
| Vmax_BLUP | AX-109338715 | 3B | 555074471 | 0.0461 | | 1.77E-22 | IIIVmrMLM | SIG |
| Vmax_BLUP | AX-110960487 | 3D | 450883789 | -0.5704 | | 2.06E-06 | EMMAX | SUG |
| Vmax_BLUP | AX-110491129 | 3D | 541615575 | -0.0328 | | 9.56E-05 | BLUPmrMLM | SUG |
| Vmax_BLUP | AX-109541921 | 4A | 100102114 | -0.0317 | | 6.27E-06 | BLUPmrMLM | SUG |
| Vmax_BLUP | AX-109420906 | 4A | 115634351 | 0.1601 | | 1.73E-07 | FarmCPU | SUG |
| Vmax_BLUP | AX-94942161 | 4A | 136928119 | 0.0533 | | 3.95E-06 | BLUPmrMLM | SUG |
| Vmax_BLUP | AX-94942161 | 4A | 136928119 | -0.0280 | | 1.30E-09 | IIIVmrMLM | SIG |
| Vmax_BLUP | AX-111065828 | 4A | 634112640 | -0.0071 | 0.1168 | 1.06E-11 | IIIVmrMLM | SIG |
| Vmax_BLUP | AX-111657963 | 4A | 704758691 | 0.0004 | -0.115 | 2.57E-10 | IIIVmrMLM | SIG |
| Vmax_BLUP | AX-110962740 | 4B | 326500336 | -0.0054 | -0.1364 | 8.99E-05 | IIIVmrMLM | SUG |

| | | | | | | | | |
|---|---|---|---|---|---|---|---|---|
| Vmax_BLUP | AX-109012419 | 4B | 526966444 | 0.0055 | -0.0787 | 2.74E-06 | IIIVmrMLM | SUG |
| Vmax_BLUP | AX-111714870 | 4D | 68054117 | 0.1645 | | 6.46E-07 | FarmCPU | SUG |
| Vmax_BLUP | AX-110604883 | 4D | 177489950 | 0.1629 | | 1.43E-07 | FarmCPU | SUG |
| Vmax_BLUP | AX-109318201 | 4D | 197019433 | 0.1533 | | 4.96E-07 | FarmCPU | SUG |
| Vmax_BLUP | AX-109448373 | 5A | 7850856 | 0.0121 | -0.0491 | 2.28E-14 | IIIVmrMLM | SIG |
| Vmax_BLUP | AX-108785128 | 5A | 69797566 | -0.0036 | 0.0978 | 2.69E-06 | IIIVmrMLM | SUG |
| Vmax_BLUP | AX-112288451 | 5A | 584614282 | -0.2132 | | 2.32E-06 | GEMMA | SUG |
| Vmax_BLUP | AX-109528616 | 5A | 585095529 | -0.2132 | | 2.32E-06 | GEMMA | SUG |
| Vmax_BLUP | AX-110416216 | 5A | 596525894 | -0.0247 | -0.0265 | 5.36E-07 | IIIVmrMLM | SUG |
| Vmax_BLUP | AX-108882297 | 5B | 66663744 | 0.0117 | 0.1448 | 1.14E-08 | IIIVmrMLM | SIG |
| Vmax_BLUP | AX-111534633 | 5B | 70735587 | 0.0199 | -0.0584 | 3.39E-08 | IIIVmrMLM | SIG |
| Vmax_BLUP | AX-110167816 | 5B | 75310707 | -0.0200 | -0.0913 | 4.13E-08 | IIIVmrMLM | SIG |
| Vmax_BLUP | AX-111139645 | 5B | 414729885 | -0.0309 | | 5.78E-06 | BLUPmrMLM | SUG |
| Vmax_BLUP | AX-108957573 | 5B | 476812795 | 0.0434 | 0.0028 | 3.42E-15 | IIIVmrMLM | SIG |
| Vmax_BLUP | AX-108994649 | 5B | 576225314 | 0.0384 | | 4.84E-05 | BLUPmrMLM | SUG |
| Vmax_BLUP | AX-109468686 | 5B | 584894906 | -0.0033 | 0.6014 | 7.09E-07 | IIIVmrMLM | SUG |
| Vmax_BLUP | AX-109509597 | 5B | 608678516 | 0.0222 | 0.3617 | 5.26E-10 | IIIVmrMLM | SIG |
| Vmax_BLUP | AX-110391970 | 5B | 669923358 | 0.0098 | 0.3695 | 2.84E-06 | IIIVmrMLM | SUG |
| Vmax_BLUP | AX-111674614 | 5B | 681034572 | -0.0391 | | 9.90E-05 | BLUPmrMLM | SUG |
| Vmax_BLUP | AX-94511633 | 5B | 681168512 | 0.1466 | | 5.80E-07 | FarmCPU | SUG |
| Vmax_BLUP | AX-109507390 | 5B | 684226311 | 0.0540 | | 1.96E-06 | GEMMA | SUG |
| Vmax_BLUP | AX-109437383 | 5B | 690078365 | -0.0575 | | 1.12E-06 | EMMAX | SUG |
| Vmax_BLUP | AX-109437383 | 5B | 690078365 | 0.0574 | | 1.24E-07 | GEMMA | SUG |
| Vmax_BLUP | AX-109026256 | 5B | 690078401 | -0.0571 | | 1.08E-06 | EMMAX | SUG |
| Vmax_BLUP | AX-109026256 | 5B | 690078401 | 0.0570 | | 1.24E-07 | GEMMA | SUG |
| Vmax_BLUP | AX-109271003 | 5B | 690078593 | -0.0576 | | 8.27E-07 | EMMAX | SUG |
| Vmax_BLUP | AX-109271003 | 5B | 690078593 | 0.0575 | | 1.02E-07 | GEMMA | SIG |
| Vmax_BLUP | AX-109298088 | 5B | 690078627 | -0.0572 | | 1.16E-06 | EMMAX | SUG |
| Vmax_BLUP | AX-109298088 | 5B | 690078627 | 0.0572 | | 1.40E-07 | GEMMA | SUG |
| Vmax_BLUP | AX-111642516 | 5B | 690087901 | 0.0573 | | 2.75E-07 | GEMMA | SUG |
| Vmax_BLUP | AX-110656582 | 5B | 690088227 | 0.0913 | | 9.96E-07 | GEMMA | SUG |
| Vmax_BLUP | AX-110977457 | 5B | 690088574 | 0.0451 | | 1.52E-06 | GEMMA | SUG |
| Vmax_BLUP | AX-110385695 | 5B | 690089042 | 0.0566 | | 3.13E-07 | GEMMA | SUG |
| Vmax_BLUP | AX-110491508 | 5B | 690089701 | 0.0541 | | 9.95E-07 | GEMMA | SUG |
| Vmax_BLUP | AX-95137418 | 5B | 690093273 | 0.1028 | | 2.19E-06 | GEMMA | SUG |
| Vmax_BLUP | AX-111527603 | 5B | 690109922 | 0.0532 | | 9.25E-07 | GEMMA | SUG |
| Vmax_BLUP | AX-109416491 | 5B | 690126821 | 0.0975 | | 1.18E-06 | GEMMA | SUG |
| Vmax_BLUP | AX-109827487 | 5B | 690176896 | 0.0527 | | 2.12E-06 | GEMMA | SUG |
| Vmax_BLUP | AX-109496546 | 5B | 690178038 | 0.0530 | | 1.92E-06 | GEMMA | SUG |
| Vmax_BLUP | AX-108814536 | 5B | 690219918 | 0.0540 | | 1.35E-06 | GEMMA | SUG |
| Vmax_BLUP | AX-108814586 | 5B | 690237485 | 0.0583 | | 7.42E-07 | GEMMA | SUG |
| Vmax_BLUP | AX-110458804 | 5B | 690240811 | 0.0594 | | 2.88E-07 | GEMMA | SUG |
| Vmax_BLUP | AX-108727124 | 5B | 690259043 | 0.1178 | | 4.35E-07 | GEMMA | SUG |
| Vmax_BLUP | AX-111487380 | 5B | 690272780 | 0.1164 | | 3.56E-07 | GEMMA | SUG |

| | | | | | | | | |
|---|---|---|---|---|---|---|---|---|
| Vmax_BLUP | AX-110396012 | 5B | 690318937 | -0.0606 | | 1.78E-06 | EMMAX | SUG |
| Vmax_BLUP | AX-110396012 | 5B | 690318937 | 0.0606 | | 1.75E-07 | GEMMA | SUG |
| Vmax_BLUP | AX-110675241 | 5B | 690319061 | 0.1164 | | 3.56E-07 | GEMMA | SUG |
| Vmax_BLUP | AX-94579483 | 5B | 690331051 | -0.1195 | | 1.40E-06 | EMMAX | SUG |
| Vmax_BLUP | AX-94579483 | 5B | 690331051 | 0.1191 | | 1.62E-07 | GEMMA | SUG |
| Vmax_BLUP | AX-95226271 | 5B | 690331087 | -0.1195 | | 1.40E-06 | EMMAX | SUG |
| Vmax_BLUP | AX-95226271 | 5B | 690331087 | 0.1191 | | 1.62E-07 | GEMMA | SUG |
| Vmax_BLUP | AX-111035704 | 5B | 690361150 | -0.0598 | | 2.14E-06 | EMMAX | SUG |
| Vmax_BLUP | AX-111035704 | 5B | 690361150 | 0.0597 | | 2.28E-07 | GEMMA | SUG |
| Vmax_BLUP | AX-111003577 | 5B | 690362025 | -0.0598 | | 2.14E-06 | EMMAX | SUG |
| Vmax_BLUP | AX-111003577 | 5B | 690362025 | 0.0597 | | 2.28E-07 | GEMMA | SUG |
| Vmax_BLUP | AX-94464717 | 5B | 690471344 | -0.1236 | | 1.17E-06 | EMMAX | SUG |
| Vmax_BLUP | AX-94464717 | 5B | 690471344 | 0.1235 | | 1.10E-07 | GEMMA | SIG |
| Vmax_BLUP | AX-111107686 | 5B | 690660732 | 0.0588 | | 2.56E-07 | GEMMA | SUG |
| Vmax_BLUP | AX-110945406 | 5B | 690676794 | 0.0588 | | 2.56E-07 | GEMMA | SUG |
| Vmax_BLUP | AX-109954516 | 5B | 690696584 | -0.0595 | | 2.22E-06 | EMMAX | SUG |
| Vmax_BLUP | AX-109954516 | 5B | 690696584 | 0.0595 | | 2.42E-07 | GEMMA | SUG |
| Vmax_BLUP | AX-110598090 | 5B | 690699628 | 0.0564 | | 4.23E-07 | GEMMA | SUG |
| Vmax_BLUP | AX-111006648 | 5B | 690700621 | 0.0583 | | 6.61E-07 | GEMMA | SUG |
| Vmax_BLUP | AX-94734778 | 5B | 691135060 | -0.1495 | | 1.34E-06 | EMMAX | SUG |
| Vmax_BLUP | AX-94734778 | 5B | 691135060 | 0.1494 | | 1.59E-07 | GEMMA | SUG |
| Vmax_BLUP | AX-94531988 | 5B | 691152152 | 0.1407 | | 1.83E-06 | GEMMA | SUG |
| Vmax_BLUP | AX-111583446 | 5B | 691954571 | 0.0468 | | 1.47E-06 | GEMMA | SUG |
| Vmax_BLUP | AX-109689624 | 5D | 17780809 | -0.0301 | | 4.41E-05 | BLUPmrMLM | SUG |
| Vmax_BLUP | AX-108840042 | 5D | 42961528 | -0.0239 | | 5.20E-05 | BLUPmrMLM | SUG |
| Vmax_BLUP | AX-111464653 | 5D | 486138536 | 0.1515 | | 2.94E-07 | FarmCPU | SUG |
| Vmax_BLUP | AX-110643388 | 5D | 547051110 | 0.1073 | | 1.45E-06 | GEMMA | SUG |
| Vmax_BLUP | AX-109846583 | 6A | 3773384 | 0.0068 | -0.1291 | 2.72E-07 | IIIVmrMLM | SUG |
| Vmax_BLUP | AX-111256802 | 6A | 260660724 | 0.1270 | | 1.44E-06 | FarmCPU | SUG |
| Vmax_BLUP | AX-111726688 | 6B | 11300863 | -0.2543 | | 1.38E-07 | FarmCPU | SUG |
| Vmax_BLUP | AX-111071251 | 6B | 11318645 | -0.1560 | | 9.92E-07 | FarmCPU | SUG |
| Vmax_BLUP | AX-110429558 | 6B | 11319311 | -0.3142 | | 1.02E-06 | FarmCPU | SUG |
| Vmax_BLUP | AX-111076083 | 6B | 250340828 | 0.0031 | -0.0841 | 2.05E-07 | IIIVmrMLM | SUG |
| Vmax_BLUP | AX-109391962 | 6B | 339268873 | 0.1797 | | 3.39E-08 | FarmCPU | SIG |
| Vmax_BLUP | AX-110675643 | 6B | 568273150 | -0.0072 | 0.037 | 1.95E-05 | IIIVmrMLM | SUG |
| Vmax_BLUP | AX-111759310 | 6D | 410108635 | -0.1103 | | 1.18E-06 | FarmCPU | SUG |
| Vmax_BLUP | AX-109305993 | 6D | 470387179 | -0.1040 | | 1.64E-06 | GEMMA | SUG |
| Vmax_BLUP | AX-109299321 | 6D | 470387591 | -0.1040 | | 1.64E-06 | GEMMA | SUG |
| Vmax_BLUP | AX-109304810 | 7A | 211734058 | 0.0079 | -0.0982 | 3.06E-05 | IIIVmrMLM | SUG |
| Vmax_BLUP | AX-111153914 | 7A | 507877239 | 0.1577 | | 1.52E-06 | FarmCPU | SUG |
| Vmax_BLUP | AX-94842225 | 7A | 641317155 | 0.1430 | | 4.21E-07 | FarmCPU | SUG |
| Vmax_BLUP | AX-111254487 | 7A | 643075152 | 0.0201 | | 1.60E-04 | BLUPmrMLM | SUG |
| Vmax_BLUP | AX-109822861 | 7B | 44878319 | -0.0270 | | 8.55E-06 | BLUPmrMLM | SUG |
| Vmax_BLUP | AX-108906914 | 7B | 604035802 | 0.0113 | 0.1217 | 3.20E-06 | IIIVmrMLM | SUG |

| | | | | | | | | |
|---|---|---|---|---|---|---|---|---|
| Vmax_BLUP | AX-111923285 | 7D | 178141446 | 0.0094 | 0.0724 | 4.84E-06 | IIIVmrMLM | SUG |

Vmax: maximum filling rate.

**Supplementary Table S19**. Candidate genes for three grain filling-related traits in genome-wide association studies and their pieces of evidence

| Trait | Genome-wide association studies | | | | | | | Differential expression analysis | | | Homologous genes/Genes reported in Rice/wheat | GO annotation analysis | | P-value in haplotype analysis | Reference |
|---|---|---|---|---|---|---|---|---|---|---|---|---|---|---|---|
| | **Environment** | **Marker** | **Chr** | **Position** | **Add/Dom** | **P-value** | **Method** | **Gene** | **$Log_2\lvert FC\rvert$** | **P. adj** | | **ID** | **Annotation** | | |
| T | T_18AY | AX-109938873 | 1A | 15323295 | 1.3544/-- | 2.72E-06 | BLUPmrMLM | TraesCS1A02G033500 | -2.42[V2] | 6.68E-15[V2] | | GO:0009744 | response to sucrose | 2.77E-08 | |
| T | T_18AY | AX-109938873 | 1A | 15323295 | 1.3544/-- | 2.72E-06 | BLUPmrMLM | TraesCS1A02G033600 | 4.08[V1]; -4.89[V2] | 9.78E-221[V1]; 2.59E-49[V2] | | GO:0009744 | response to sucrose | 1.14E-04 | |
| T | T_19XX | AX-111031818 | 2A | 136838918 | 0.9185/0.1737 | 3.29E-07 | IIIVmrMLM | TraesCS2A02G176000 | -3.53[V2] | 2.30E-2[V2] | *TaGW7* | | | 1.80E-15 | Wang et al., 2019 |
| T | T_17JY | AX-111463303 | 2B | 169865292 | -1.042/-- | 1.83E-05 | BLUPmrMLM | TraesCS2B02G194200 | 3.03[V1]; -7.73[V2] | 4.06E-230[V1]; 3.5E-136[V2] | *TaSuSy2* | GO:0016157 | sucrose synthase activity | 5.73E-03 | Mukherjee et al., 2015 |
| T | T_17JY | AX-111463303 | 2B | 169865292 | 1.5132/-- | 6.71E-07 | EMMAX | TraesCS2B02G194200 | 3.03[V1]; -7.73[V2] | 4.06E-230[V1]; 3.5E-136[V2] | *TaSuSy2* | GO:0016157 | sucrose synthase activity | 5.73E-03 | Mukherjee et al., 2015 |
| T | T_17JY | AX-111463303 | 2B | 169865292 | -1.51223/-- | 4.24E-07 | GEMMA | TraesCS2B02G194200 | 3.03[V1]; -7.73[V2] | 4.06E-230[V1]; 3.5E-136[V2] | *TaSuSy2* | GO:0016157 | sucrose synthase activity | 5.73E-03 | Mukherjee et al., 2015 |
| T | T_BLUP | AX-110952281 | 2B | 765280430 | 0.0036/0.3783 | 4.73E-11 | IIIVmrMLM | TraesCS2B01G578500 | -1.42[V1]; -1.42[V2] | 5.86E-06[V1]; 5.84E-07[V2] | *TaARF12-2B* | | | 3.79E-05 | Kong et al., 2023 |
| T | T_BLUP | AX-111017431 | 4A | 597898318 | -0.0247/0.0059 | 1.05E-07 | IIIVmrMLM | TraesCS4A01G302000 | 1.63[V2] | 3.14E-28[V2] | | GO:0009744 | response to sucrose | 5.20E-03 | |
| T | T_17AY | AX-109515751 | 4B | 665916709 | 1.3412/-- | 1.56E-05 | BLUPmrMLM | TraesCS4B01G390500 | -1.07[V1]; -3.22[V2] | 1.81E-3[V1]; 4.6E-20[V2] | | GO:0009744 | response to sucrose | 2.48E-03 | |
| T | T_17AY | AX-109515751 | 4B | 665916709 | -1.9191/-- | 1.75E-06 | EMMAX | TraesCS4B01G390500 | -1.07[V1]; -3.22[V2] | 0.00181[V1]; 4.6E-20[V2] | | GO:0009744 | response to sucrose | 2.48E-03 | |
| T | T_17AY | AX-109515751 | 4B | 665916709 | 1.918766/-- | 1.09E-06 | GEMMA | TraesCS4B01G390500 | -1.07[V1]; -3.22[V2] | 1.81E-3[V1]; 4.6E-20[V2] | | GO:0009744 | response to sucrose | 2.48E-03 | |
| T | T_17JY | AX-109941199 | 7D | 108407039 | -28.498/-- | 2.26E-06 | EMMAX | TraesCS7D01G159800 | 1.22[V1]; -2.46[V2] | 4.87E-47[V1]; 1.72E-174[V2] | *TaSuSy1* | GO:0016157 | sucrose synthase activity | 1.20E-02 | Maraña et al., 1990 |
| T | T_BLUP | AX-109933656 | 7D | 612929882 | -0.0113/-0.0355 | 2.97E-07 | IIIVmrMLM | TraesCS7D01G512100 | 3.49[V2] | 1.27E-48[V2] | | GO:0009744 | response to sucrose | 2.36E-07 | |
| V | V_BLUP | AX-108787384 | 1A | 376508048 | -0.0125/-0.0405 | 5.58E-14 | IIIVmrMLM | TraesCS1A01G214700 | -7.51[V2] | 1.41E-05[V2] | | GO:0005982 | starch metabolic process | 3.53E-42 | |
| V | V_17AY | AX-111031083 | 1B | 50023282 | 0.0055/-0.3237 | 4.08E-07 | IIIVmrMLM | TraesCS1B01G066900 | 11.03[V1]; 5.64[V2] | 1.69E-12[V1]; 7.43E-4[V2] | | GO:0015770 | sucrose transport | 4.80E-05 | |
| V | V_BLUP | AX-109973922 | 2A | 504467482 | -0.0064/-0.0986 | 1.43E-06 | IIIVmrMLM | TraesCS2A01G293400 | -2.36[V2] | 4.86E-128[V2] | *TaSBEIIa* | GO:0005982 | starch metabolic process | 2.55E-61 | Kumar et al., 2022 |
| V | V_18AY | AX-111134442 | 2B | 183950769 | -0.0163/-0.1055 | 7.35E-10 | IIIVmrMLM | TraesCS2B01G202300 | -1.91[V2] | 3.33E-2[V2] | *TaGW7* | | | 4.77E-14 | Wang et al., 2019 |
| V | V_BLUP | AX-110922479 | 3A | 8795309 | 0.0022/0.0862 | 1.85E-06 | IIIVmrMLM | TraesCS3A01G015500 | 0.81[V1]; 1.19[V2] | 5.96E-14[V1]; 5.3E-31[V2] | | GO:0046524 | sucrose-phosphate synthase activity | 4.44E-65 | |
| V | V_17JY | AX-109449758 | 3B | 717049513 | 0.0129/-0.2682 | 4.13E-06 | IIIVmrMLM | TraesCS3B01G469000 | -1.95[V1]; 1.7[V2] | 3.86E-18[V1]; 3.56E-26[V2] | | GO:0009744 | response to sucrose | 4.45E-05 | |
| V | V_19XX | AX-108730133 | 3B | 796472256 | 0.0486/-- | 5.41E-06 | BLUPmrMLM | TraesCS3B01G560400 | 3.85[V1]; -6.41[V2] | 1.74E-20[V1]; 5.69E-05[V2] | *OsIF3* | | | 4.40E-05 | Guo et al., 2022 |

| | | | | | | | | | | | | | | | |
|---|---|---|---|---|---|---|---|---|---|---|---|---|---|---|---|
| V | V_BLUP | AX-111600364 | 3B | 795031754 | -0.0178/-0.0182 | 2.76E-09 | IIIVmrMLM | TraesCS3B01G560400 | 3.85 $^{V1}$; -6.41 $^{V2}$ | 1.74E-20 $^{V1}$; 5.69E-05 $^{V2}$ | *OsIF3* | | | 5.52E-16 | Guo et al., 2022 |
| V | V_19XX | AX-108730133 | 3B | 796472256 | 0.0486/-- | 5.41E-06 | BLUPmrMLM | TraesCS3B01G561000 | 3.04 $^{V1}$; -10.54 $^{V2}$ | 4.9E-48 $^{V1}$; 2.96E-12 $^{V2}$ | *OsIF3* | | | 4.48E-05 | Guo et al., 2022 |
| V | V_BLUP | AX-111600364 | 3B | 795031754 | -0.0178/-0.0182 | 2.76E-09 | IIIVmrMLM | TraesCS3B01G561000 | 3.04 $^{V1}$; -10.54 $^{V2}$ | 4.9E-48 $^{V1}$; 2.96E-12 $^{V2}$ | *OsIF3* | | | 4.30E-16 | Guo et al., 2022 |
| V | V_18JY | AX-94434096 | 4A | 11843465 | 0.0581/-- | 5.39E-06 | BLUPmrMLM | TraesCS4A01G016600 | -5.08 $^{V1}$; -7.98 $^{V2}$ | 6.8E-28 $^{V1}$; 4.53E-24 $^{V2}$ | | GO:0009744 | response to sucrose | 2.43E-31 | |
| V | V_18JY | AX-110929258 | 5B | 635500 | -0.0319/-- | 5.51E-05 | BLUPmrMLM | TraesCS5B01G000100 | -0.65 $^{V1}$; 1.76 $^{V2}$ | 1.22E-05 $^{V1}$; 8.92E-47 $^{V2}$ | | GO:0015770 | sucrose transport | 6.49E-14 | |
| V | V_BLUP | AX-110430140 | 6B | 4548176 | 0.0175/-- | 3.02E-05 | BLUPmrMLM | TraesCS6B01G009600 | -2.87 $^{V2}$ | 1.19E-20 $^{V2}$ | | GO:0009744 | response to sucrose | 5.84E-24 | |
| V | V_BLUP | AX-111620734 | 6B | 4125682 | -0.0138/0.0263 | 4.23E-06 | IIIVmrMLM | TraesCS6B01G009600 | -2.87 $^{V2}$ | 1.19E-20 $^{V2}$ | | GO:0009744 | response to sucrose | 5.84E-24 | |
| V | V_BLUP | AX-109305993 | 6D | 470387179 | -0.06663336/-- | 3.58E-07 | GEMMA | TraesCS6D01G403800 | 0.94 $^{V1}$; -1.01 $^{V2}$ | 4.86E-2 $^{V1}$; 3.32E-2 $^{V2}$ | | GO:0016157 | sucrose synthase activity | 1.27E-20 | |
| V | V_BLUP | AX-110526933 | 6D | 470387275 | -0.05438403/-- | 9.55E-07 | GEMMA | TraesCS6D01G403800 | 0.94 $^{V1}$; -1.01 $^{V2}$ | 4.86E-2 $^{V1}$; 3.32E-2 $^{V2}$ | | GO:0016157 | sucrose synthase activity | 1.27E-20 | |
| V | V_BLUP | AX-109299321 | 6D | 470387591 | -0.06663336/-- | 3.58E-07 | GEMMA | TraesCS6D01G403800 | 0.94 $^{V1}$; -1.01 $^{V2}$ | 4.86E-2 $^{V1}$; 3.32E-2 $^{V2}$ | | GO:0016157 | sucrose synthase activity | 1.27E-20 | |
| V | V_BLUP | AX-111038802 | 6D | 470752471 | -0.05224823/-- | 1.61E-06 | GEMMA | TraesCS6D01G403800 | 0.94 $^{V1}$; -1.01 $^{V2}$ | 4.86E-2 $^{V1}$; 3.32E-2 $^{V2}$ | | GO:0016157 | sucrose synthase activity | 1.27E-20 | |
| V | V_BLUP | AX-110857795 | 6D | 470847028 | -0.05526586/-- | 7.34E-07 | GEMMA | TraesCS6D01G403800 | 0.94 $^{V1}$; -1.01 $^{V2}$ | 4.86E-2 $^{V1}$; 3.32E-2 $^{V2}$ | | GO:0016157 | sucrose synthase activity | 1.27E-20 | |
| V | V_BLUP | AX-111010395 | 6D | 470864026 | -0.0510118/-- | 2.25E-06 | GEMMA | TraesCS6D01G403800 | 0.94 $^{V1}$; -1.01 $^{V2}$ | 4.86E-2 $^{V1}$; 3.32E-2 $^{V2}$ | | GO:0016157 | sucrose synthase activity | 1.27E-20 | |
| V | V_18JY | AX-109335759 | 7A | 19419576 | 0.0367/-- | 0.000131 | BLUPmrMLM | TraesCS7A01G040900 | 0.71 $^{V1}$; -4.99 $^{V2}$ | 3.6E-05 $^{V1}$; 8.9E-88 $^{V2}$ | | GO:0016157 | sucrose synthase activity | 5.62E-10 | |
| V | V_18AY | AX-111728563 | 7A | 703564535 | 0.0062/-0.4351 | 2.07E-06 | IIIVmrMLM | TraesCS7A01G518200 | 1.07 $^{V2}$ | 1.49E-14 $^{V2}$ | | GO:0046524 | sucrose-phosphate synthase activity | 1.30E-13 | |
| V | V_18AY | AX-111728563 | 7A | 703564535 | 0.0062/-0.4351 | 2.07E-06 | IIIVmrMLM | TraesCS7A01G518300 | -5.07 $^{V2}$ | 1.23E-45 $^{V2}$ | | GO:0009744 | response to sucrose | 1.17E-33 | |
| V | V_17SQ | AX-108908590 | 7A | 721572672 | 0.0162/-0.368 | 6.43E-12 | IIIVmrMLM | TraesCS7A01G549100 | 6.30 $^{V1}$; -3.94 $^{V2}$ | 0.00 $^{V1}$; 5.02E-55 $^{V2}$ | *TaSBEIa* | GO:0005982 | starch metabolic process | 1.74E-13 | Liu et al., 2022 |
| V | V_17SQ | AX-108908590 | 7A | 721572672 | 0.0162/-0.368 | 6.43E-12 | IIIVmrMLM | TraesCS7A01G549200 | 8.53 $^{V1}$; -3.69 $^{V2}$ | 6.78E-52 $^{V1}$; 3.40E-2 $^{V2}$ | | GO:0005982 | starch metabolic process | 1.16E-14 | |
| V | V_17SQ | AX-108908590 | 7A | 721572672 | 0.0162/-0.368 | 6.43E-12 | IIIVmrMLM | TraesCS7A01G549300 | 3.52 $^{V1}$; -7.53 $^{V2}$ | 1.28E-30 $^{V1}$; 1.2E-06 $^{V2}$ | *TaSBEIb* | GO:0005982 | starch metabolic process | 1.10E-13 | Pei et al., 2023 |
| V | V_17JY | AX-89526681 | 7B | 34668546 | -0.0597/-- | 8.36E-05 | BLUPmrMLM | TraesCS7B01G034600 | 2.70 $^{V1}$; -3.48 $^{V2}$ | 4.82E-47 $^{V1}$; 1.88E-31 $^{V2}$ | | GO:0019252 | starch biosynthetic process | 1.09E-13 | |
| V | V_18AY | AX-109045123 | 7B | 134224402 | 0.0315/-- | 3.21E-07 | IIIVmrMLM | TraesCS7B01G115300 | -5.40 $^{V1}$; -9.00 $^{V2}$ | 2.92E-27 $^{V1}$; 1.3E-16 $^{V2}$ | *TaGASR7-B1* | | | 4.01E-14 | Zhang et al., 2016 |
| V | V_18JY | AX-108876019 | 7B | 705812472 | 0.0357/-0.0052 | 3.49E-05 | IIIVmrMLM | TraesCS7B01G439900 | 2.32 $^{V2}$ | 7.09E-17 $^{V2}$ | | GO:0009744 | response to sucrose | 1.01E-15 | |
| V | V_17AY | AX-95219160 | 7B | 727529846 | -0.93205/-- | 3.27E-08 | EMMAX | TraesCS7B01G472500 | 4.71 $^{V1}$; -2.47 $^{V2}$ | 0.00 $^{V1}$; 2.58E-81 $^{V2}$ | | GO:0005982 | starch metabolic process | 8.32E-12 | |
| V | V_17AY | AX-108728735 | 7B | 727652888 | -0.6023/-- | 3.55E-07 | EMMAX | TraesCS7B01G472500 | 4.71 $^{V1}$; -2.47 $^{V2}$ | 0.00 $^{V1}$; 2.58E-81 $^{V2}$ | | GO:0005982 | starch metabolic process | 8.32E-12 | |

| | | | | | | | | | | | | | | | |
|---|---|---|---|---|---|---|---|---|---|---|---|---|---|---|---|
| V | V_17AY | AX-108730446 | 7B | 729251407 | -0.93205/-- | 3.27E-08 | EMMAX | TraesCS7B01G472500 | 4.71 $^{V1}$; -2.47 $^{V2}$ | 0.00 $^{V1}$; 2.58E-81 $^{V2}$ | | GO:0005982 | starch metabolic process | 8.32E-12 | |
| V | V_17SQ | AX-110736759 | 7D | 173900640 | -0.0469/-0.0315 | 1.78E-11 | IIIVmrMLM | TraesCS7D01G215600 | -8.63 $^{V2}$ | 6.78E-08 $^{V2}$ | | GO:0009744 | response to sucrose | 2.28E-17 | |
| Vmax | Vmax_18AY | AX-109462275 | 1A | 15485453 | -0.0575/-- | 4.89E-05 | BLUPmrMLM | TraesCS1A01G033500 | -2.42 $^{V2}$ | 6.68E-15 $^{V2}$ | | GO:0009744 | response to sucrose | 3.38E-21 | |
| Vmax | Vmax_18AY | AX-94486559 | 1A | 15437125 | 0.0255/-0.1081 | 1.85E-09 | IIIVmrMLM | TraesCS1A01G033500 | -2.42 $^{V2}$ | 6.68E-15 $^{V2}$ | | GO:0009744 | response to sucrose | 3.38E-21 | |
| Vmax | Vmax_18AY | AX-109462275 | 1A | 15485453 | -0.0575/-- | 4.89E-05 | BLUPmrMLM | TraesCS1A01G033600 | 4.08 $^{V1}$; -4.89 $^{V2}$ | 9.78E-221 $^{V1}$;2.59E-49 $^{V2}$ | | GO:0009744 | response to sucrose | 4.11E-10 | |
| Vmax | Vmax_18AY | AX-94486559 | 1A | 15437125 | 0.0255/-0.1081 | 1.85E-09 | IIIVmrMLM | TraesCS1A01G033600 | 4.08 $^{V1}$; -4.89 $^{V2}$ | 9.78E-221 $^{V1}$; 2.59E-49 $^{V2}$ | | GO:0009744 | response to sucrose | 4.11E-10 | |
| Vmax | Vmax_17JY | AX-111090830 | 1A | 386777934 | -0.0266/-0.4377 | 2.68E-06 | IIIVmrMLM | TraesCS1A01G219400 | 0.61 $^{V1}$; 1.85 $^{V2}$ | 3.77E-4 $^{V1}$; 1.06E-36 $^{V2}$ | | GO:0009744 | response to sucrose | 1.03E-39 | |
| Vmax | Vmax_19XX | AX-111091775 | 1A | 436447126 | 0.0215/0.2335 | 4.57E-07 | IIIVmrMLM | TraesCS1A01G247100 | 0.72 $^{V1}$; -2.19 $^{V2}$ | 8.03E-05 $^{V1}$; 2.95E-30 $^{V2}$ | *TaISA2* | GO:0005982 | starch metabolic process | 9.40E-11 | Pei et al., 2023 |
| Vmax | Vmax_17AY | AX-111031083 | 1B | 50023282 | -0.0012/-0.5881 | 2.23E-10 | IIIVmrMLM | TraesCS1B01G066900 | 11.03 $^{V1}$; 5.64 $^{V2}$ | 1.69E-12 $^{V1}$; 7.43E-4 $^{V2}$ | | GO:0015770 | sucrose transport | 4.74E-05 | |
| Vmax | Vmax_19XX | AX-95659604 | 1B | 645572033 | 0.0267/0.7139 | 4.23E-06 | IIIVmrMLM | TraesCS1B01G422700 | -10.01 $^{V1}$; -10.91 $^{V2}$ | 1.29E-10 $^{V1}$; 9.58E-13 $^{V2}$ | | GO:0005982 | starch metabolic process | 3.66E-15 | |
| Vmax | Vmax_BLUP | AX-109973922 | 2A | 504467482 | 6e-04/-0.2058 | 1.86E-08 | IIIVmrMLM | TraesCS2A01G293400 | -2.36 $^{V2}$ | 4.86E-128 $^{V2}$ | *TaSBEIIa* | GO:0005982 | starch metabolic process | 3.53E-62 | Kumar et al., 2022 |
| Vmax | Vmax_18AY | AX-111134442 | 2B | 183950769 | -0.0095/-0.1608 | 1.14E-08 | IIIVmrMLM | TraesCS2B01G202300 | -1.91 $^{V2}$ | 3.33E-2 $^{V2}$ | *TaGW7* | | | 1.07E-13 | Wang et al., 2019 |
| Vmax | Vmax_17JY | AX-109449758 | 3B | 717049513 | 0.0429/-0.3677 | 1.3E-06 | IIIVmrMLM | TraesCS3B01G469000 | -1.95 $^{V1}$; 1.70 $^{V2}$ | 3.86E-18 $^{V1}$; 3.56E-26 $^{V2}$ | | GO:0009744 | response to sucrose | 4.58E-05 | |
| Vmax | Vmax_19XX | AX-110632987 | 4A | 13251112 | 0.0074/-0.2032 | 1.37E-06 | IIIVmrMLM | TraesCS4A01G016600 | -5.08 $^{V1}$; -7.98 $^{V2}$ | 6.8E-28 $^{V1}$; 4.53E-24 $^{V2}$ | | GO:0009744 | response to sucrose | 9.52E-15 | |
| Vmax | Vmax_BLUP | AX-109541921 | 4A | 100102114 | -0.0317/-- | 6.27E-06 | BLUPmrMLM | TraesCS4A01G093100 | -2.35 $^{V1}$; -11.58 $^{V2}$ | 1.74E-17 $^{V1}$; 1.73E-14 $^{V2}$ | | GO:0005986 | sucrose biosynthetic process | 2.78E-03 | |
| Vmax | Vmax_17AY | AX-108999430 | 4B | 667683709 | -0.0783/-- | 6.08E-05 | BLUPmrMLM | TraesCS4B01G390500 | -1.07 $^{V1}$; -3.22 $^{V2}$ | 1.81E-3 $^{V1}$; 4.6E-20 $^{V2}$ | | GO:0009744 | response to sucrose | 1.37E-09 | |
| Vmax | Vmax_19XX | AX-94657293 | 5B | 45392491 | 0.0346/0.2801 | 3.06E-08 | IIIVmrMLM | TraesCS5B01G039800 | -1.70 $^{V1}$; -1.64 $^{V2}$ | 3.34E-19 $^{V1}$; 5.18E-20 $^{V2}$ | *TaARF25-5B* | | | 1.79E-05 | Jia et al., 2021 |
| Vmax | Vmax_BLUP | AX-108994649 | 5B | 576225314 | 0.0384/-- | 4.84E-05 | BLUPmrMLM | TraesCS5B01G400000 | 1.54 $^{V1}$; -4.94 $^{V2}$ | 7.33E-30 $^{V1}$; 1.06E-157 $^{V2}$ | *TaPHOL* | | | 4.93E-68 | Miao et al., 2022 |
| Vmax | Vmax_19XX | AX-111620734 | 6B | 4125682 | -0.0154/0.5942 | 4.36E-06 | IIIVmrMLM | TraesCS6B01G009600 | -2.87 $^{V2}$ | 1.19E-20 $^{V2}$ | | GO:0009744 | response to sucrose | 2.97E-10 | |
| Vmax | Vmax_18AY | AX-108758805 | 6B | 643139508 | -0.0044/0.4929 | 1.09E-05 | IIIVmrMLM | TraesCS6B01G369700 | 0.57 $^{V1}$; 1.72 $^{V2}$ | 3.44E-2 $^{V1}$; 1.21E-14 $^{V2}$ | | GO:0009744 | response to sucrose | 4.78E-17 | |
| Vmax | Vmax_17AY | AX-108919084 | 6D | 470477263 | 0.0035/-0.4612 | 2.01E-07 | IIIVmrMLM | TraesCS6D01G403800 | 0.94 $^{V1}$; -1.01 $^{V2}$ | 4.86E-2 $^{V1}$; 3.32E-2 $^{V2}$ | | GO:0016157 | sucrose synthase activity | 2.57E-03 | |
| Vmax | Vmax_17SQ | AX-89498965 | 6D | 471008558 | -0.0077/0.1633 | 0.000453 | IIIVmrMLM | TraesCS6D01G403800 | 0.94 $^{V1}$; -1.01 $^{V2}$ | 0.0486 $^{V1}$; 3.32E-2 $^{V2}$ | | GO:0016157 | sucrose synthase activity | 9.07E-09 | |
| Vmax | Vmax_BLUP | AX-109305993 | 6D | 470387179 | -0.1040199/-- | 1.64E-06 | GEMMA | TraesCS6D01G403800 | 0.94 $^{V1}$; -1.01 $^{V2}$ | 4.86E-2 $^{V1}$; 3.32E-2 $^{V2}$ | | GO:0016157 | sucrose synthase activity | 6.47E-21 | |
| Vmax | Vmax_BLUP | AX-109299321 | 6D | 470387591 | -0.1040199/-- | 1.64E-06 | GEMMA | TraesCS6D01G403800 | 0.94 $^{V1}$; -1.01 $^{V2}$ | 4.86E-2 $^{V1}$; 3.32E-2 $^{V2}$ | | GO:0016157 | sucrose synthase activity | 6.47E-21 | |

| | | | | | | | | | | | | | | | |
|---|---|---|---|---|---|---|---|---|---|---|---|---|---|---|---|
| Vmax | Vmax_17SQ | AX-108908590 | 7A | 721572672 | 0.0245/-0.4956 | 1.18E-10 | IIIVmrMLM | TraesCS7A01G549100 | 6.30 $^{V1}$; -3.94 $^{V2}$ | 0.00 $^{V1}$; 5.02E-55 $^{V2}$ | *TaSBEIa* | GO:0005982 | starch metabolic process | 1.56E-13 | Liu et al., 2022 |
| Vmax | Vmax_17SQ | AX-108908590 | 7A | 721572672 | 0.0245/-0.4956 | 1.18E-10 | IIIVmrMLM | TraesCS7A01G549200 | 8.53 $^{V1}$; -3.69 $^{V2}$ | 6.78E-52 $^{V1}$; 3.40E-2 $^{V2}$ | | GO:0005982 | starch metabolic process | 1.00E-14 | |
| Vmax | Vmax_17SQ | AX-108908590 | 7A | 721572672 | 0.0245/-0.4956 | 1.18E-10 | IIIVmrMLM | TraesCS7A01G549300 | 3.52 $^{V1}$; -7.53 $^{V2}$ | 1.28E-30 $^{V1}$; 1.2E-06 $^{V2}$ | *TaSBEIb* | GO:0005982 | starch metabolic process | 1.04E-13 | Pei et al., 2023 |
| Vmax | Vmax_17AY | AX-94751384 | 7B | 35080930 | 0.2991/-- | 6.28E-11 | BLUPmrMLM | TraesCS7B01G034600 | 2.70 $^{V1}$; -3.48 $^{V2}$ | 4.82E-47 $^{V1}$; 1.88E-31 $^{V2}$ | | GO:0019252 | starch biosynthetic process | 1.46E-02 | |
| Vmax | Vmax_18AY | AX-109045123 | 7B | 134224402 | 0.0749/-- | 5.9E-15 | IIIVmrMLM | TraesCS7B01G115300 | -5.40 $^{V1}$; -9.00 $^{V2}$ | 2.92E-27 $^{V1}$; 1.3E-16 $^{V2}$ | *TaGASR7-B1* | | | 2.62E-14 | Zhang et al., 2016 |
| Vmax | Vmax_18JY | AX-109868959 | 7B | 704252664 | 0.0585/0.0299 | 2.35E-06 | IIIVmrMLM | TraesCS7B01G439900 | 2.32 $^{V2}$ | 7.09E-17 $^{V2}$ | | GO:0009744 | response to sucrose | 1.41E-17 | |
| Vmax | Vmax_17AY | AX-95219160 | 7B | 727529846 | -1.3958/-- | 3.47E-08 | EMMAX | TraesCS7B01G472500 | 4.71 $^{V1}$; -2.47 $^{V2}$ | 0.00 $^{V1}$; 2.58E-81 $^{V2}$ | | GO:0005982 | starch metabolic process | 9.14E-12 | |
| Vmax | Vmax_17AY | AX-108728735 | 7B | 727652888 | -0.9028/-- | 3.65E-07 | EMMAX | TraesCS7B01G472500 | 4.71 $^{V1}$; -2.47 $^{V2}$ | 0.00 $^{V1}$; 2.58E-81 $^{V2}$ | | GO:0005982 | starch metabolic process | 9.14E-12 | |
| Vmax | Vmax_17AY | AX-108730446 | 7B | 729251407 | -1.3958/-- | 3.47E-08 | EMMAX | TraesCS7B01G472500 | 4.71 $^{V1}$; -2.47 $^{V2}$ | 0.00 $^{V1}$; 2.58E-81 $^{V2}$ | | GO:0005982 | starch metabolic process | 9.14E-12 | |
| Vmax | Vmax_17SQ | AX-110736759 | 7D | 173900640 | -0.0734/-0.0648 | 2.42E-13 | IIIVmrMLM | TraesCS7D01G215600 | -8.63 $^{V2}$ | 6.78E-08 $^{V2}$ | | GO:0009744 | response to sucrose | 2.21E-17 | |

**Supplementary Table S20**. All QTNs identified as associated with flowering date in wheat.

| Trait | Marker | Chr | Position | Add.effect | Dom.effect | P-value | Method | Sig. |
|---|---|---|---|---|---|---|---|---|
| FD_17AY | AX-94849470 | 1A | 551308340 | -0.3474 | | 2.96E-05 | BLUPmrMLM | SUG |
| FD_17AY | AX-111700863 | 1A | 557968013 | -0.3949 | | 9.17E-07 | BLUPmrMLM | SUG |
| FD_17AY | AX-109283335 | 1B | 323660235 | 0.3072 | | 1.89E-04 | BLUPmrMLM | SUG |
| FD_17AY | AX-111503667 | 1B | 485802556 | -0.2517 | | 1.34E-04 | BLUPmrMLM | SUG |
| FD_17AY | AX-95074606 | 1B | 661502620 | -0.8006 | | 2.21E-06 | BLUPmrMLM | SUG |
| FD_17AY | AX-95074606 | 1B | 661502620 | -0.1337 | -0.7241 | 1.62E-09 | IIIVmrMLM | SIG |
| FD_17AY | AX-108750878 | 1D | 14147616 | 0.0144 | 0.7996 | 5.90E-08 | IIIVmrMLM | SIG |
| FD_17AY | AX-108898996 | 1D | 454766541 | -0.0156 | 1.9019 | 1.02E-11 | IIIVmrMLM | SIG |
| FD_17AY | AX-94793594 | 1D | 462488082 | -0.4980 | 0.2247 | 6.88E-16 | IIIVmrMLM | SIG |
| FD_17AY | AX-110089297 | 1D | 462996873 | -0.5275 | | 5.95E-07 | BLUPmrMLM | SUG |
| FD_17AY | AX-110731402 | 2A | 308034792 | -0.3762 | -1.269 | 4.10E-35 | IIIVmrMLM | SIG |
| FD_17AY | AX-108912390 | 2A | 428552045 | 0.0135 | 0.9391 | 1.43E-11 | IIIVmrMLM | SIG |
| FD_17AY | AX-109518634 | 2A | 632498489 | 0.1632 | 0.678 | 1.36E-20 | IIIVmrMLM | SIG |
| FD_17AY | AX-108864407 | 2B | 242535487 | -0.0192 | 0.7973 | 2.22E-06 | IIIVmrMLM | SUG |
| FD_17AY | AX-110993119 | 2B | 290868022 | 0.0337 | 0.7204 | 6.47E-09 | IIIVmrMLM | SIG |
| FD_17AY | AX-111537086 | 2B | 601183765 | 0.3683 | | 2.05E-13 | IIIVmrMLM | SIG |
| FD_17AY | AX-111468726 | 2B | 657791701 | -0.4506 | | 1.09E-06 | BLUPmrMLM | SUG |
| FD_17AY | AX-111585974 | 2B | 765900766 | -0.3254 | 0.2469 | 7.13E-08 | IIIVmrMLM | SIG |
| FD_17AY | AX-111151139 | 2D | 411476541 | -0.4630 | | 2.08E-07 | BLUPmrMLM | SUG |
| FD_17AY | AX-110579075 | 3A | 686193501 | -0.3003 | | 8.23E-05 | BLUPmrMLM | SUG |
| FD_17AY | AX-110491100 | 3A | 705251779 | 2.0536 | | 8.62E-09 | GEMMA | SIG |
| FD_17AY | AX-108941177 | 3A | 705259848 | 1.9732 | | 3.97E-09 | GEMMA | SIG |
| FD_17AY | AX-111695263 | 3A | 705274739 | 1.9966 | | 1.02E-08 | GEMMA | SIG |
| FD_17AY | AX-110976174 | 3A | 705274782 | 3.8381 | | 7.62E-09 | GEMMA | SIG |
| FD_17AY | AX-110484538 | 3A | 705296840 | 4.2541 | | 1.57E-08 | GEMMA | SIG |
| FD_17AY | AX-110390871 | 3B | 7252439 | 0.5327 | | 4.63E-08 | BLUPmrMLM | SIG |
| FD_17AY | AX-110390871 | 3B | 7252439 | 0.7323 | | 7.35E-07 | GEMMA | SUG |
| FD_17AY | AX-111081536 | 3B | 8806517 | -0.2955 | 0.3106 | 1.63E-08 | IIIVmrMLM | SIG |
| FD_17AY | AX-111135864 | 3B | 50193392 | -0.3820 | | 2.41E-05 | BLUPmrMLM | SUG |
| FD_17AY | AX-110989677 | 3B | 61104501 | -0.1080 | -0.7183 | 2.18E-08 | IIIVmrMLM | SIG |
| FD_17AY | AX-111073617 | 3B | 262742817 | 0.1959 | 0.937 | 9.06E-14 | IIIVmrMLM | SIG |
| FD_17AY | AX-109967710 | 3B | 469873446 | 0.4955 | | 1.96E-07 | BLUPmrMLM | SUG |
| FD_17AY | AX-109967710 | 3B | 469873446 | -0.4690 | 0.006 | 3.08E-17 | IIIVmrMLM | SIG |
| FD_17AY | AX-108910474 | 3B | 685981381 | 0.3894 | | 6.40E-07 | BLUPmrMLM | SUG |
| FD_17AY | AX-95112826 | 3B | 696673313 | 0.3673 | | 2.36E-13 | IIIVmrMLM | SIG |
| FD_17AY | AX-111576602 | 3B | 727153104 | 0.1950 | 2.012 | 1.93E-10 | IIIVmrMLM | SIG |
| FD_17AY | AX-111543008 | 3B | 727794913 | -0.4001 | | 5.14E-05 | BLUPmrMLM | SUG |
| FD_17AY | AX-110397603 | 3B | 779581070 | 0.4806 | | 8.32E-11 | BLUPmrMLM | SIG |
| FD_17AY | AX-110918080 | 3B | 781851318 | -0.0342 | -0.7337 | 2.02E-04 | IIIVmrMLM | SUG |
| FD_17AY | AX-108951591 | 3D | 420455318 | -0.2956 | | 4.88E-05 | BLUPmrMLM | SUG |
| FD_17AY | AX-110701651 | 4A | 230241219 | -0.0212 | -2.9396 | 1.29E-04 | IIIVmrMLM | SUG |

| | | | | | | | | |
|---|---|---|---|---|---|---|---|---|
| FD_17AY | AX-111190872 | 4A | 540416431 | -0.3537 | | 1.59E-12 | IIIVmrMLM | SIG |
| FD_17AY | AX-108881020 | 4B | 8412450 | 0.4267 | | 6.74E-07 | BLUPmrMLM | SUG |
| FD_17AY | AX-108838330 | 4B | 465981268 | -0.0188 | -0.8297 | 2.44E-06 | IIIVmrMLM | SUG |
| FD_17AY | AX-109934789 | 5A | 7715630 | -0.0322 | -1.0078 | 3.00E-09 | IIIVmrMLM | SIG |
| FD_17AY | AX-111689418 | 5A | 540026591 | -0.0717 | 2.4267 | 5.25E-13 | IIIVmrMLM | SIG |
| FD_17AY | AX-108814780 | 5A | 654921809 | 0.2338 | -0.449 | 6.44E-06 | IIIVmrMLM | SUG |
| FD_17AY | AX-111682490 | 5B | 59932775 | 1.1447 | | 1.42E-06 | FarmCPU | SUG |
| FD_17AY | AX-108822938 | 5B | 66339657 | 1.3141 | | 1.79E-06 | FarmCPU | SUG |
| FD_17AY | AX-109480591 | 5B | 460248328 | 0.0707 | -1.1744 | 2.68E-11 | IIIVmrMLM | SIG |
| FD_17AY | AX-111161208 | 5B | 529989075 | 0.1302 | 4.7512 | 1.45E-05 | IIIVmrMLM | SUG |
| FD_17AY | AX-95215490 | 5B | 649490548 | 0.3649 | | 3.31E-13 | IIIVmrMLM | SIG |
| FD_17AY | AX-94582317 | 5D | 29585874 | -0.0155 | 1.2795 | 2.08E-08 | IIIVmrMLM | SIG |
| FD_17AY | AX-109369920 | 5D | 249650807 | -0.5896 | | 4.49E-05 | BLUPmrMLM | SUG |
| FD_17AY | AX-109859450 | 5D | 529225487 | 0.1058 | 1.0044 | 9.09E-19 | IIIVmrMLM | SIG |
| FD_17AY | AX-110167616 | 5D | 532965771 | 0.4146 | | 6.33E-07 | BLUPmrMLM | SUG |
| FD_17AY | AX-111486410 | 6A | 3008615 | -0.0020 | -0.9088 | 8.80E-11 | IIIVmrMLM | SIG |
| FD_17AY | AX-109840305 | 6A | 106196611 | 0.3498 | 0.0481 | 7.53E-10 | IIIVmrMLM | SIG |
| FD_17AY | AX-95153605 | 6B | 31516855 | -0.2823 | | 1.32E-08 | IIIVmrMLM | SIG |
| FD_17AY | AX-109840463 | 6B | 127611268 | 0.3212 | | 3.36E-05 | BLUPmrMLM | SUG |
| FD_17AY | AX-110360699 | 6B | 201254117 | -0.7439 | | 9.84E-12 | BLUPmrMLM | SIG |
| FD_17AY | AX-110360699 | 6B | 201254117 | 0.5481 | -0.2655 | 1.95E-21 | IIIVmrMLM | SIG |
| FD_17AY | AX-108779531 | 6B | 686383283 | -0.4234 | | 6.46E-05 | BLUPmrMLM | SUG |
| FD_17AY | AX-108901786 | 6B | 712475275 | -0.0352 | -1.1221 | 4.01E-05 | IIIVmrMLM | SUG |
| FD_17AY | AX-89424484 | 6D | 96823283 | -0.2366 | | 1.43E-04 | BLUPmrMLM | SUG |
| FD_17AY | AX-109833655 | 7A | 10110972 | 0.5119 | | 8.69E-07 | BLUPmrMLM | SUG |
| FD_17AY | AX-110977044 | 7A | 12725608 | -0.2435 | -0.3252 | 5.24E-06 | IIIVmrMLM | SUG |
| FD_17AY | AX-109068845 | 7A | 117272978 | 0.1902 | 2.7686 | 1.90E-09 | IIIVmrMLM | SIG |
| FD_17AY | AX-108940753 | 7A | 541337103 | 0.0180 | 0.5679 | 2.22E-05 | IIIVmrMLM | SUG |
| FD_17AY | AX-111051951 | 7A | 642035607 | 0.3154 | | 4.38E-06 | BLUPmrMLM | SUG |
| FD_17AY | AX-109520645 | 7A | 652603840 | -0.3362 | -0.0269 | 5.18E-10 | IIIVmrMLM | SIG |
| FD_17AY | AX-110576947 | 7D | 65436205 | -0.6965 | | 1.78E-08 | BLUPmrMLM | SIG |
| FD_17AY | AX-111184541 | 7D | 66833728 | -1.0440 | | 1.37E-06 | GEMMA | SUG |
| FD_17AY | AX-111843581 | 7D | 67448053 | -1.0513 | | 1.41E-06 | GEMMA | SUG |
| FD_17AY | AX-110875275 | 7D | 257526220 | -0.4879 | 0.4462 | 1.35E-20 | IIIVmrMLM | SIG |
| FD_17AY | AX-109824355 | 7D | 529964800 | -0.1363 | -0.4427 | 1.33E-05 | IIIVmrMLM | SUG |
| FD_17ZMD | AX-94704697 | 1A | 4812682 | -0.4235 | | 2.23E-16 | IIIVmrMLM | SIG |
| FD_17ZMD | AX-111125508 | 1A | 8634079 | -0.7019 | -0.2025 | 2.00E-37 | IIIVmrMLM | SIG |
| FD_17ZMD | AX-94411371 | 1A | 13670297 | 0.1583 | -0.8712 | 1.77E-07 | IIIVmrMLM | SUG |
| FD_17ZMD | AX-110505731 | 1A | 324499550 | -0.9207 | | 1.42E-06 | BLUPmrMLM | SUG |
| FD_17ZMD | AX-109374474 | 1A | 404541202 | 0.8572 | | 2.69E-12 | BLUPmrMLM | SIG |
| FD_17ZMD | AX-109993427 | 1B | 386626322 | 0.2048 | 0.904 | 3.06E-13 | IIIVmrMLM | SIG |
| FD_17ZMD | AX-110503854 | 1B | 398790050 | -0.0434 | 0.8973 | 7.03E-12 | IIIVmrMLM | SIG |
| FD_17ZMD | AX-94496522 | 1B | 622886614 | 2.6785 | 1.1603 | 4.37E-91 | IIIVmrMLM | SIG |
| FD_17ZMD | AX-108732493 | 1B | 640851255 | 0.3475 | | 7.38E-05 | BLUPmrMLM | SUG |

| | | | | | | | | |
|---|---|---|---|---|---|---|---|---|
| FD_17ZMD | AX-110967256 | 1B | 677617120 | -0.0776 | -0.6484 | 2.18E-09 | IIIVmrMLM | SIG |
| FD_17ZMD | AX-108952278 | 1B | 687536032 | -0.3850 | | 5.91E-06 | BLUPmrMLM | SUG |
| FD_17ZMD | AX-109366846 | 1D | 35753416 | 0.4922 | | 2.32E-07 | BLUPmrMLM | SUG |
| FD_17ZMD | AX-109142090 | 1D | 476910568 | -0.7971 | | 6.72E-10 | BLUPmrMLM | SIG |
| FD_17ZMD | AX-109142090 | 1D | 476910568 | 0.5650 | 0.979 | 4.21E-26 | IIIVmrMLM | SIG |
| FD_17ZMD | AX-110374023 | 2A | 71438950 | 0.4007 | | 2.37E-05 | BLUPmrMLM | SUG |
| FD_17ZMD | AX-110922941 | 2A | 73551783 | -0.0507 | -1.9451 | 2.50E-07 | IIIVmrMLM | SUG |
| FD_17ZMD | AX-111674236 | 2A | 466161184 | 0.1232 | 1.1758 | 6.59E-11 | IIIVmrMLM | SIG |
| FD_17ZMD | AX-109924835 | 2A | 535801035 | -0.0517 | -0.8635 | 9.22E-15 | IIIVmrMLM | SIG |
| FD_17ZMD | AX-109537967 | 2A | 619207292 | -0.4942 | | 1.29E-06 | BLUPmrMLM | SUG |
| FD_17ZMD | AX-110468896 | 2A | 753567489 | -0.0259 | -3.0333 | 1.99E-13 | IIIVmrMLM | SIG |
| FD_17ZMD | AX-110014288 | 2B | 164175967 | 0.1324 | -0.7253 | 1.78E-11 | IIIVmrMLM | SIG |
| FD_17ZMD | AX-111775165 | 2B | 285238717 | 0.0884 | -3.5868 | 1.38E-08 | IIIVmrMLM | SIG |
| FD_17ZMD | AX-111213446 | 2B | 490313471 | -0.1699 | 1.6147 | 1.46E-08 | IIIVmrMLM | SIG |
| FD_17ZMD | AX-108763630 | 2B | 573724064 | 0.0334 | -0.8457 | 2.00E-08 | IIIVmrMLM | SIG |
| FD_17ZMD | AX-109323364 | 2B | 742472864 | 0.6254 | | 2.43E-06 | BLUPmrMLM | SUG |
| FD_17ZMD | AX-110476283 | 2B | 758901285 | 0.7181 | | 1.01E-07 | BLUPmrMLM | SIG |
| FD_17ZMD | AX-108904634 | 2B | 796344276 | -0.1194 | -0.7976 | 2.06E-06 | IIIVmrMLM | SUG |
| FD_17ZMD | AX-109378890 | 2D | 93363349 | -0.7961 | | 5.83E-06 | BLUPmrMLM | SUG |
| FD_17ZMD | AX-109378890 | 2D | 93363349 | 0.2936 | | 7.76E-09 | IIIVmrMLM | SIG |
| FD_17ZMD | AX-109854381 | 2D | 115620359 | 1.3198 | | 1.61E-14 | BLUPmrMLM | SIG |
| FD_17ZMD | AX-110058357 | 2D | 607143374 | -0.1104 | 0.4927 | 4.11E-07 | IIIVmrMLM | SUG |
| FD_17ZMD | AX-108732118 | 3A | 136276069 | 0.0439 | 0.8546 | 1.28E-04 | IIIVmrMLM | SUG |
| FD_17ZMD | AX-109936048 | 3A | 162357149 | 0.0428 | -1.1823 | 1.35E-06 | IIIVmrMLM | SUG |
| FD_17ZMD | AX-109620355 | 3A | 475976393 | -0.5248 | | 1.89E-05 | BLUPmrMLM | SUG |
| FD_17ZMD | AX-108738012 | 3A | 523737968 | 0.0567 | -1.7593 | 5.67E-38 | IIIVmrMLM | SIG |
| FD_17ZMD | AX-108941177 | 3A | 705259848 | -2.1453 | | 9.13E-07 | EMMAX | SUG |
| FD_17ZMD | AX-108941177 | 3A | 705259848 | 2.1342 | | 5.37E-07 | GEMMA | SUG |
| FD_17ZMD | AX-110484538 | 3A | 705296840 | -4.9605 | | 4.86E-07 | EMMAX | SUG |
| FD_17ZMD | AX-110484538 | 3A | 705296840 | 4.9389 | | 2.46E-07 | GEMMA | SUG |
| FD_17ZMD | AX-110006232 | 3B | 11631075 | -0.8065 | | 1.49E-10 | BLUPmrMLM | SIG |
| FD_17ZMD | AX-108969017 | 3B | 13442822 | 1.0294 | | 4.32E-12 | BLUPmrMLM | SIG |
| FD_17ZMD | AX-111049518 | 3B | 33759379 | -0.4908 | | 2.83E-06 | BLUPmrMLM | SUG |
| FD_17ZMD | AX-111088141 | 3B | 315427726 | 0.9146 | 2.0645 | 3.15E-62 | IIIVmrMLM | SIG |
| FD_17ZMD | AX-110390549 | 3B | 397298731 | -0.3840 | -0.3681 | 8.88E-13 | IIIVmrMLM | SIG |
| FD_17ZMD | AX-110062183 | 3B | 659281390 | -0.4634 | | 8.17E-07 | BLUPmrMLM | SUG |
| FD_17ZMD | AX-94627268 | 3D | 400693482 | 0.5164 | | 3.72E-06 | BLUPmrMLM | SUG |
| FD_17ZMD | AX-94395861 | 4A | 230998878 | -1.5251 | | 4.90E-10 | BLUPmrMLM | SIG |
| FD_17ZMD | AX-108906441 | 4A | 310078696 | -0.2984 | -3.1176 | 2.41E-13 | IIIVmrMLM | SIG |
| FD_17ZMD | AX-111699414 | 4A | 568530042 | 0.0537 | -0.9844 | 4.09E-10 | IIIVmrMLM | SIG |
| FD_17ZMD | AX-110957600 | 4A | 618314675 | 0.3684 | | 1.96E-07 | BLUPmrMLM | SUG |
| FD_17ZMD | AX-109581501 | 4A | 722135793 | -0.0783 | -1.0856 | 1.65E-07 | IIIVmrMLM | SUG |
| FD_17ZMD | AX-89605275 | 4B | 647667251 | -0.3333 | | 1.63E-05 | BLUPmrMLM | SUG |
| FD_17ZMD | AX-109584296 | 4D | 297099057 | -13.3341 | -12.802 | 0.00E+00 | IIIVmrMLM | SIG |

| | | | | | | | | |
|---|---|---|---|---|---|---|---|---|
| FD_17ZMD | AX-110987476 | 5A | 35638010 | -0.0839 | 1.0972 | 5.15E-09 | IIIVmrMLM | SIG |
| FD_17ZMD | AX-109383929 | 5A | 159802105 | -0.2247 | | 8.80E-06 | IIIVmrMLM | SUG |
| FD_17ZMD | AX-89496688 | 5A | 466614771 | 0.1393 | -2.3176 | 1.59E-10 | IIIVmrMLM | SIG |
| FD_17ZMD | AX-110506335 | 5A | 528114941 | -0.6050 | | 1.57E-08 | BLUPmrMLM | SIG |
| FD_17ZMD | AX-110506335 | 5A | 528114941 | 0.4876 | -0.0888 | 8.42E-16 | IIIVmrMLM | SIG |
| FD_17ZMD | AX-111165329 | 5B | 83784325 | 0.0554 | -1.095 | 4.92E-08 | IIIVmrMLM | SIG |
| FD_17ZMD | AX-108759044 | 5B | 242804477 | -0.3510 | -0.8711 | 1.19E-17 | IIIVmrMLM | SIG |
| FD_17ZMD | AX-109551675 | 5B | 379811358 | 0.0340 | 0.9513 | 2.59E-07 | IIIVmrMLM | SUG |
| FD_17ZMD | AX-111612134 | 5B | 680347003 | 0.4217 | | 1.02E-05 | BLUPmrMLM | SUG |
| FD_17ZMD | AX-111081549 | 5B | 701172230 | 0.5513 | | 1.79E-08 | BLUPmrMLM | SIG |
| FD_17ZMD | AX-94952724 | 5D | 57109549 | -0.4248 | | 1.84E-16 | IIIVmrMLM | SIG |
| FD_17ZMD | AX-108970703 | 5D | 275606471 | -0.6243 | | 5.50E-05 | BLUPmrMLM | SUG |
| FD_17ZMD | AX-110035093 | 5D | 464129291 | 0.9846 | | 1.44E-06 | GEMMA | SUG |
| FD_17ZMD | AX-111175690 | 5D | 464415388 | 1.3256 | | 1.70E-06 | EMMAX | SUG |
| FD_17ZMD | AX-111175690 | 5D | 464415388 | -1.3124 | | 1.44E-06 | GEMMA | SUG |
| FD_17ZMD | AX-110717870 | 5D | 465106394 | -1.0304 | | 7.62E-08 | EMMAX | SIG |
| FD_17ZMD | AX-110717870 | 5D | 465106394 | 1.0427 | | 4.93E-08 | GEMMA | SIG |
| FD_17ZMD | AX-94390434 | 5D | 466230373 | -0.4296 | | 2.80E-06 | BLUPmrMLM | SUG |
| FD_17ZMD | AX-109750213 | 5D | 466694319 | -0.9682 | | 4.37E-07 | EMMAX | SUG |
| FD_17ZMD | AX-109750213 | 5D | 466694319 | 0.9748 | | 3.03E-07 | GEMMA | SUG |
| FD_17ZMD | AX-110958036 | 5D | 469179441 | -0.9174 | | 1.47E-06 | EMMAX | SUG |
| FD_17ZMD | AX-110958036 | 5D | 469179441 | 0.9247 | | 1.08E-06 | GEMMA | SUG |
| FD_17ZMD | AX-109360916 | 5D | 469286340 | 3.4636 | | 3.72E-09 | EMMAX | SIG |
| FD_17ZMD | AX-109360916 | 5D | 469286340 | -3.4488 | | 2.46E-09 | GEMMA | SIG |
| FD_17ZMD | AX-109360916 | 5D | 469286340 | 0.8903 | | 5.30E-57 | IIIVmrMLM | SIG |
| FD_17ZMD | AX-108787427 | 5D | 470769698 | -0.4800 | | 4.79E-08 | BLUPmrMLM | SIG |
| FD_17ZMD | AX-111599209 | 6A | 1732483 | 0.4842 | 0.0639 | 2.01E-17 | IIIVmrMLM | SIG |
| FD_17ZMD | AX-94857662 | 6A | 17910863 | 0.4637 | | 4.32E-07 | BLUPmrMLM | SUG |
| FD_17ZMD | AX-111474814 | 6B | 675563689 | 0.1371 | 1.4298 | 1.85E-10 | IIIVmrMLM | SIG |
| FD_17ZMD | AX-110989869 | 6B | 693693269 | 0.5797 | | 1.31E-09 | BLUPmrMLM | SIG |
| FD_17ZMD | AX-108998615 | 6B | 719675442 | -0.0526 | -0.9475 | 7.55E-14 | IIIVmrMLM | SIG |
| FD_17ZMD | AX-110382511 | 6D | 24026993 | 0.2151 | | 1.25E-04 | BLUPmrMLM | SUG |
| FD_17ZMD | AX-95009409 | 6D | 470044057 | 1.0467 | | 1.65E-10 | BLUPmrMLM | SIG |
| FD_17ZMD | AX-109864970 | 7A | 47458495 | -0.3795 | 0.395 | 1.21E-12 | IIIVmrMLM | SIG |
| FD_17ZMD | AX-111686136 | 7A | 63892737 | -0.0114 | 2.161 | 1.01E-04 | IIIVmrMLM | SUG |
| FD_17ZMD | AX-109815375 | 7A | 353144146 | 0.0904 | -2.7042 | 1.42E-08 | IIIVmrMLM | SIG |
| FD_17ZMD | AX-110191443 | 7A | 645720640 | 0.9276 | | 2.72E-06 | BLUPmrMLM | SUG |
| FD_17ZMD | AX-111018476 | 7A | 668982701 | 0.7026 | | 8.28E-39 | IIIVmrMLM | SIG |
| FD_17ZMD | AX-94524732 | 7A | 670781122 | -0.2416 | | 1.82E-06 | IIIVmrMLM | SUG |
| FD_17ZMD | AX-110674101 | 7B | 41034820 | 0.3518 | | 2.02E-05 | BLUPmrMLM | SUG |
| FD_17ZMD | AX-95200013 | 7B | 59180076 | -0.0684 | -0.8969 | 5.54E-07 | IIIVmrMLM | SUG |
| FD_17ZMD | AX-109924435 | 7B | 127213876 | 0.5680 | 0.219 | 1.25E-23 | IIIVmrMLM | SIG |
| FD_17ZMD | AX-111644834 | 7B | 642828675 | -0.2881 | | 1.46E-05 | BLUPmrMLM | SUG |
| FD_17ZMD | AX-95661646 | 7B | 716958100 | -0.3552 | | 3.88E-12 | IIIVmrMLM | SIG |

| | | | | | | | | |
|---|---|---|---|---|---|---|---|---|
| FD_17ZMD | AX-95199297 | 7D | 3857123 | 0.5562 | | 3.80E-26 | IIIVmrMLM | SIG |
| FD_17ZMD | AX-109870041 | 7D | 581610668 | -0.1900 | 1.711 | 1.31E-08 | IIIVmrMLM | SIG |
| FD_18AY | AX-110180533 | 1A | 393335356 | -2.1459 | | 3.99E-08 | FarmCPU | SIG |
| FD_18AY | AX-111462775 | 1A | 494810364 | 1.8430 | | 2.11E-06 | EMMAX | SUG |
| FD_18AY | AX-111462775 | 1A | 494810364 | -1.8460 | | 3.59E-07 | GEMMA | SUG |
| FD_18AY | AX-111453306 | 1A | 494818292 | -1.6946 | | 1.59E-06 | GEMMA | SUG |
| FD_18AY | AX-109886157 | 1A | 494887518 | -1.7577 | | 8.75E-07 | GEMMA | SUG |
| FD_18AY | AX-111087202 | 1A | 494941701 | -1.6911 | | 1.75E-06 | GEMMA | SUG |
| FD_18AY | AX-109426053 | 1A | 494942486 | -1.7577 | | 8.75E-07 | GEMMA | SUG |
| FD_18AY | AX-111606000 | 1A | 494943039 | -1.7429 | | 8.59E-07 | GEMMA | SUG |
| FD_18AY | AX-111488547 | 1A | 495013458 | -1.7577 | | 8.75E-07 | GEMMA | SUG |
| FD_18AY | AX-109817102 | 1A | 495013803 | -1.7036 | | 1.31E-06 | GEMMA | SUG |
| FD_18AY | AX-109490911 | 1A | 495014197 | 1.8430 | | 2.11E-06 | EMMAX | SUG |
| FD_18AY | AX-109490911 | 1A | 495014197 | -1.8460 | | 3.59E-07 | GEMMA | SUG |
| FD_18AY | AX-111609168 | 1A | 495014994 | 1.8693 | | 1.23E-06 | EMMAX | SUG |
| FD_18AY | AX-111609168 | 1A | 495014994 | -1.8667 | | 2.76E-07 | GEMMA | SUG |
| FD_18AY | AX-109937348 | 1A | 495035706 | -1.6640 | | 2.33E-06 | GEMMA | SUG |
| FD_18AY | AX-110606043 | 1A | 495036799 | -1.7577 | | 8.75E-07 | GEMMA | SUG |
| FD_18AY | AX-111550437 | 1A | 495037353 | 1.8430 | | 2.11E-06 | EMMAX | SUG |
| FD_18AY | AX-111550437 | 1A | 495037353 | -1.8460 | | 3.59E-07 | GEMMA | SUG |
| FD_18AY | AX-111002026 | 1A | 495037475 | -1.7577 | | 8.75E-07 | GEMMA | SUG |
| FD_18AY | AX-108796834 | 1A | 495037699 | 1.8430 | | 2.11E-06 | EMMAX | SUG |
| FD_18AY | AX-108796834 | 1A | 495037699 | -1.8460 | | 3.59E-07 | GEMMA | SUG |
| FD_18AY | AX-111767357 | 1A | 495125499 | -1.7451 | | 8.72E-07 | GEMMA | SUG |
| FD_18AY | AX-110714062 | 1A | 495126545 | -1.6911 | | 1.75E-06 | GEMMA | SUG |
| FD_18AY | AX-110995371 | 1A | 495127554 | -1.6815 | | 1.08E-06 | GEMMA | SUG |
| FD_18AY | AX-110997823 | 1A | 495133578 | -1.7108 | | 1.22E-06 | GEMMA | SUG |
| FD_18AY | AX-108926906 | 1A | 495181892 | -1.7693 | | 1.27E-06 | GEMMA | SUG |
| FD_18AY | AX-109448112 | 1A | 495182069 | -1.7577 | | 8.75E-07 | GEMMA | SUG |
| FD_18AY | AX-111555272 | 1A | 495189010 | -1.4707 | | 1.17E-06 | GEMMA | SUG |
| FD_18AY | AX-94652392 | 1A | 517378589 | 0.3891 | | 6.97E-05 | BLUPmrMLM | SUG |
| FD_18AY | AX-94652392 | 1A | 517378589 | -0.4452 | -0.1841 | 4.88E-12 | IIIVmrMLM | SIG |
| FD_18AY | AX-94381634 | 1A | 544172916 | 0.0592 | -0.4989 | 5.69E-06 | IIIVmrMLM | SUG |
| FD_18AY | AX-110013315 | 1A | 560884406 | 0.3316 | | 4.34E-09 | IIIVmrMLM | SIG |
| FD_18AY | AX-108780402 | 1A | 564570992 | -1.6926 | | 1.37E-07 | FarmCPU | SUG |
| FD_18AY | AX-94684170 | 1B | 597537011 | -2.4495 | | 1.00E-07 | FarmCPU | SIG |
| FD_18AY | AX-109395485 | 2A | 17557922 | 0.3640 | | 9.32E-05 | BLUPmrMLM | SUG |
| FD_18AY | AX-109543143 | 2A | 17596321 | -0.0139 | -1.1765 | 1.21E-26 | IIIVmrMLM | SIG |
| FD_18AY | AX-109906471 | 2B | 46592661 | 0.0281 | -2.3484 | 1.31E-07 | IIIVmrMLM | SUG |
| FD_18AY | AX-94978498 | 2B | 92996342 | -0.3185 | | 1.65E-08 | IIIVmrMLM | SIG |
| FD_18AY | AX-94697254 | 2D | 33710135 | -1.9931 | | 7.16E-07 | FarmCPU | SUG |
| FD_18AY | AX-109928221 | 2D | 81438313 | 0.0182 | -3.1115 | 2.69E-07 | IIIVmrMLM | SUG |
| FD_18AY | AX-111946160 | 2D | 320241808 | -1.8342 | | 2.22E-06 | FarmCPU | SUG |
| FD_18AY | AX-94442651 | 2D | 507050616 | -2.2548 | | 2.17E-06 | FarmCPU | SUG |

| | | | | | | | | |
|---|---|---|---|---|---|---|---|---|
| FD_18AY | AX-111988974 | 2D | 573477416 | 1.6418 | | 1.97E-06 | FarmCPU | SUG |
| FD_18AY | AX-109489829 | 3A | 56731796 | -0.0061 | -0.6112 | 8.41E-04 | IIIVmrMLM | SUG |
| FD_18AY | AX-110964081 | 3A | 189317195 | 0.5849 | | 7.63E-24 | IIIVmrMLM | SIG |
| FD_18AY | AX-109286245 | 3A | 421119830 | -1.9978 | | 1.68E-07 | FarmCPU | SUG |
| FD_18AY | AX-94522205 | 3A | 638010015 | -1.2916 | | 3.14E-08 | FarmCPU | SIG |
| FD_18AY | AX-95151919 | 3A | 689498082 | -0.4353 | | 8.39E-07 | BLUPmrMLM | SUG |
| FD_18AY | AX-110562555 | 3B | 11533876 | -0.7947 | | 1.22E-07 | BLUPmrMLM | SUG |
| FD_18AY | AX-110709880 | 3B | 78815408 | 0.7167 | | 5.17E-06 | BLUPmrMLM | SUG |
| FD_18AY | AX-110116341 | 3B | 132531497 | 0.0669 | 0.5903 | 3.23E-08 | IIIVmrMLM | SIG |
| FD_18AY | AX-111584157 | 3B | 328721122 | -0.1799 | 1.8575 | 3.75E-10 | IIIVmrMLM | SIG |
| FD_18AY | AX-110390549 | 3B | 397298731 | -0.0401 | -0.7736 | 7.63E-05 | IIIVmrMLM | SUG |
| FD_18AY | AX-108951664 | 3B | 554961692 | -1.0602 | -1.8479 | 1.23E-79 | IIIVmrMLM | SIG |
| FD_18AY | AX-110949850 | 3B | 556386549 | -0.4417 | | 1.46E-05 | BLUPmrMLM | SUG |
| FD_18AY | AX-110104032 | 3B | 618689236 | -0.0503 | 0.9895 | 1.43E-04 | IIIVmrMLM | SUG |
| FD_18AY | AX-111724657 | 3D | 28499634 | -0.5651 | -0.2035 | 7.83E-16 | IIIVmrMLM | SIG |
| FD_18AY | AX-109211753 | 3D | 231390696 | -1.8671 | | 6.07E-07 | FarmCPU | SUG |
| FD_18AY | AX-108904508 | 3D | 307390318 | -0.3616 | | 6.63E-05 | BLUPmrMLM | SUG |
| FD_18AY | AX-108948871 | 3D | 376977635 | -2.1172 | | 8.48E-07 | FarmCPU | SUG |
| FD_18AY | AX-110454130 | 3D | 593222451 | -2.7520 | -3.4981 | 8.29E-225 | IIIVmrMLM | SIG |
| FD_18AY | AX-109932274 | 4A | 60712364 | 0.4662 | 0.2869 | 4.65E-15 | IIIVmrMLM | SIG |
| FD_18AY | AX-108824978 | 4A | 191189140 | -2.5457 | | 1.49E-06 | GEMMA | SUG |
| FD_18AY | AX-109383852 | 4A | 198122747 | -1.2161 | | 3.90E-06 | BLUPmrMLM | SUG |
| FD_18AY | AX-110701651 | 4A | 230241219 | 0.2787 | -3.2711 | 1.51E-09 | IIIVmrMLM | SIG |
| FD_18AY | AX-109846555 | 4A | 237113713 | -3.8423 | | 7.05E-07 | GEMMA | SUG |
| FD_18AY | AX-110727256 | 4A | 237695603 | -3.5551 | | 7.39E-07 | GEMMA | SUG |
| FD_18AY | AX-111665679 | 4A | 299019804 | -2.8389 | | 1.42E-06 | GEMMA | SUG |
| FD_18AY | AX-111681450 | 4A | 349171706 | -1.9948 | | 2.25E-06 | GEMMA | SUG |
| FD_18AY | AX-110608887 | 4A | 361305064 | -2.7930 | | 1.71E-06 | GEMMA | SUG |
| FD_18AY | AX-111045266 | 4A | 381459131 | -3.3501 | | 9.21E-07 | GEMMA | SUG |
| FD_18AY | AX-110065075 | 4A | 396690115 | 3.6838 | | 1.48E-06 | EMMAX | SUG |
| FD_18AY | AX-110065075 | 4A | 396690115 | -3.6888 | | 2.10E-07 | GEMMA | SUG |
| FD_18AY | AX-111572808 | 4A | 404228421 | -2.0147 | | 1.27E-06 | GEMMA | SUG |
| FD_18AY | AX-109608730 | 4A | 413757062 | -2.6765 | | 1.31E-06 | GEMMA | SUG |
| FD_18AY | AX-109286260 | 4A | 503210659 | 1.2474 | | 8.79E-07 | FarmCPU | SUG |
| FD_18AY | AX-111027628 | 4A | 597845056 | 0.4476 | | 1.88E-05 | BLUPmrMLM | SUG |
| FD_18AY | AX-94699427 | 4A | 688095838 | -0.3061 | -0.8653 | 1.92E-19 | IIIVmrMLM | SIG |
| FD_18AY | AX-110620525 | 4B | 612215473 | 0.4200 | 0.033 | 4.02E-11 | IIIVmrMLM | SIG |
| FD_18AY | AX-111970247 | 4D | 32225033 | 0.4540 | 0.1795 | 7.01E-12 | IIIVmrMLM | SIG |
| FD_18AY | AX-111714870 | 4D | 68054117 | -2.5240 | | 4.17E-07 | FarmCPU | SUG |
| FD_18AY | AX-110604883 | 4D | 177489950 | -2.4134 | | 2.38E-07 | FarmCPU | SUG |
| FD_18AY | AX-110345857 | 4D | 236594032 | -1.9781 | | 3.35E-07 | FarmCPU | SUG |
| FD_18AY | AX-109250027 | 4D | 496418052 | -0.4155 | | 2.90E-13 | IIIVmrMLM | SIG |
| FD_18AY | AX-108889346 | 5A | 411690705 | -1.8089 | | 1.62E-06 | FarmCPU | SUG |

| | | | | | | | | |
|---|---|---|---|---|---|---|---|---|
| FD_18AY | AX-109579828 | 5A | 476898675 | 1.3515 | | 1.11E-06 | FarmCPU | SUG |
| FD_18AY | AX-95652893 | 5B | 4862088 | 0.1821 | 1.8144 | 2.81E-11 | IIIVmrMLM | SIG |
| FD_18AY | AX-110435059 | 5B | 15036583 | -0.1224 | -1.7759 | 3.96E-14 | IIIVmrMLM | SIG |
| FD_18AY | AX-110144727 | 5B | 409680012 | 0.0740 | 0.994 | 5.16E-05 | IIIVmrMLM | SUG |
| FD_18AY | AX-109102787 | 5B | 502122832 | -0.0494 | -0.8621 | 1.54E-10 | IIIVmrMLM | SIG |
| FD_18AY | AX-109968215 | 5B | 516100266 | 0.0803 | -1.3434 | 5.46E-10 | IIIVmrMLM | SIG |
| FD_18AY | AX-110483836 | 5B | 545794046 | 0.0288 | 1.5238 | 2.53E-07 | IIIVmrMLM | SUG |
| FD_18AY | AX-110997009 | 5B | 568171719 | -0.4244 | -0.9989 | 1.02E-13 | IIIVmrMLM | SIG |
| FD_18AY | AX-111049187 | 5B | 569827392 | -0.9882 | | 8.65E-07 | EMMAX | SUG |
| FD_18AY | AX-111049187 | 5B | 569827392 | 0.9832 | | 4.65E-07 | GEMMA | SUG |
| FD_18AY | AX-108731931 | 5B | 587477374 | -2.2695 | | 2.20E-06 | GEMMA | SUG |
| FD_18AY | AX-109498841 | 5B | 587477780 | -2.4768 | | 9.91E-07 | GEMMA | SUG |
| FD_18AY | AX-111682790 | 5B | 685427202 | -0.3801 | | 1.68E-05 | BLUPmrMLM | SUG |
| FD_18AY | AX-108987566 | 5D | 164594241 | -2.2313 | | 4.51E-08 | FarmCPU | SIG |
| FD_18AY | AX-111496494 | 5D | 370135591 | 0.5922 | 0.5507 | 3.20E-23 | IIIVmrMLM | SIG |
| FD_18AY | AX-94556621 | 5D | 391737205 | 0.3942 | | 3.96E-12 | IIIVmrMLM | SIG |
| FD_18AY | AX-111486410 | 6A | 3008615 | 0.0913 | -0.8205 | 9.96E-08 | IIIVmrMLM | SIG |
| FD_18AY | AX-95092292 | 6A | 63303203 | 0.0015 | -0.5662 | 3.41E-09 | IIIVmrMLM | SIG |
| FD_18AY | AX-111256802 | 6A | 260660724 | -1.9924 | | 5.53E-07 | FarmCPU | SUG |
| FD_18AY | AX-108742970 | 6B | 27991047 | 0.3874 | | 1.79E-04 | BLUPmrMLM | SUG |
| FD_18AY | AX-109329712 | 6B | 58106508 | -0.3199 | | 1.44E-08 | IIIVmrMLM | SIG |
| FD_18AY | AX-109391962 | 6B | 339268873 | -2.5981 | | 1.25E-07 | FarmCPU | SUG |
| FD_18AY | AX-111123882 | 6D | 272759669 | -2.2889 | | 1.12E-06 | FarmCPU | SUG |
| FD_18AY | AX-109647325 | 6D | 438653466 | 0.3530 | | 4.44E-10 | IIIVmrMLM | SIG |
| FD_18AY | AX-111872610 | 6D | 457851551 | -1.8637 | | 2.05E-06 | FarmCPU | SUG |
| FD_18AY | AX-108806541 | 7A | 49398896 | 0.4941 | | 1.38E-05 | BLUPmrMLM | SUG |
| FD_18AY | AX-108806541 | 7A | 49398896 | -0.3411 | 0.244 | 1.87E-08 | IIIVmrMLM | SIG |
| FD_18AY | AX-109647141 | 7A | 285789901 | 0.2640 | 2.7028 | 8.12E-12 | IIIVmrMLM | SIG |
| FD_18AY | AX-109452241 | 7A | 519187081 | -1.8647 | | 4.25E-07 | FarmCPU | SUG |
| FD_18AY | AX-94842225 | 7A | 641317155 | -2.0600 | | 1.34E-06 | FarmCPU | SUG |
| FD_18AY | AX-111536008 | 7A | 712070970 | 0.0963 | 1.6665 | 1.76E-11 | IIIVmrMLM | SIG |
| FD_18AY | AX-111640910 | 7B | 540073505 | 0.1047 | 2.214 | 1.87E-06 | IIIVmrMLM | SUG |
| FD_18AY | AX-109829922 | 7B | 732596318 | 0.5728 | | 5.26E-23 | IIIVmrMLM | SIG |
| FD_18AY | AX-110835487 | 7D | 426153254 | -2.5209 | | 8.97E-07 | FarmCPU | SUG |
| FD_18AY | AX-94480836 | 7D | 592199944 | 0.1864 | 1.4556 | 1.22E-06 | IIIVmrMLM | SUG |
| FD_18JY | AX-111469353 | 1A | 301373345 | 0.3543 | | 1.19E-04 | BLUPmrMLM | SUG |
| FD_18JY | AX-111102601 | 1A | 539320590 | -0.6804 | | 1.52E-08 | BLUPmrMLM | SIG |
| FD_18JY | AX-94495040 | 1A | 549619420 | 0.5908 | | 2.80E-06 | BLUPmrMLM | SUG |
| FD_18JY | AX-109815012 | 1B | 573051141 | 0.6867 | | 4.41E-09 | BLUPmrMLM | SIG |
| FD_18JY | AX-108829556 | 1B | 581099719 | -0.1989 | 1.1491 | 3.00E-07 | IIIVmrMLM | SUG |
| FD_18JY | AX-94496522 | 1B | 622886614 | 2.3759 | 1.4067 | 1.12E-86 | IIIVmrMLM | SIG |
| FD_18JY | AX-111517692 | 1B | 677570622 | -0.2965 | | 2.53E-05 | BLUPmrMLM | SUG |
| FD_18JY | AX-110967256 | 1B | 677617120 | -0.0214 | -1.0297 | 1.81E-14 | IIIVmrMLM | SIG |
| FD_18JY | AX-111528085 | 1D | 2432202 | 0.2739 | | 8.13E-06 | IIIVmrMLM | SUG |

| | | | | | | | | |
|---|---|---|---|---|---|---|---|---|
| FD_18JY | AX-111713476 | 1D | 41150395 | -0.0523 | 3.6579 | 5.39E-06 | IIIVmrMLM | SUG |
| FD_18JY | AX-110422891 | 2A | 536310677 | -0.5786 | -0.5936 | 5.16E-19 | IIIVmrMLM | SIG |
| FD_18JY | AX-111768401 | 2A | 605590887 | 0.7109 | | 1.76E-08 | BLUPmrMLM | SIG |
| FD_18JY | AX-111768401 | 2A | 605590887 | -0.5374 | 0.0083 | 5.11E-12 | IIIVmrMLM | SIG |
| FD_18JY | AX-109439440 | 2B | 11383490 | -0.3542 | 0.0986 | 1.76E-07 | IIIVmrMLM | SUG |
| FD_18JY | AX-111001399 | 2B | 47273270 | -0.0254 | -1.1114 | 1.21E-06 | IIIVmrMLM | SUG |
| FD_18JY | AX-111087936 | 2B | 777308839 | -0.2622 | | 7.80E-05 | BLUPmrMLM | SUG |
| FD_18JY | AX-111956072 | 2D | 34428838 | 0.6034 | | 1.74E-10 | BLUPmrMLM | SIG |
| FD_18JY | AX-111956072 | 2D | 34428838 | -0.7556 | | 4.44E-07 | EMMAX | SUG |
| FD_18JY | AX-111956072 | 2D | 34428838 | 0.7610 | | 1.60E-07 | GEMMA | SUG |
| FD_18JY | AX-111956072 | 2D | 34428838 | -0.4636 | -1.0555 | 1.82E-13 | IIIVmrMLM | SIG |
| FD_18JY | AX-108887820 | 2D | 43666174 | -0.3599 | | 4.58E-07 | BLUPmrMLM | SUG |
| FD_18JY | AX-109901903 | 2D | 610366462 | 0.2237 | | 7.30E-05 | BLUPmrMLM | SUG |
| FD_18JY | AX-108756128 | 3A | 19988117 | 0.9165 | | 2.31E-06 | FarmCPU | SUG |
| FD_18JY | AX-110938240 | 3A | 269089641 | 0.3428 | 0.2983 | 2.23E-07 | IIIVmrMLM | SUG |
| FD_18JY | AX-111512557 | 3A | 512859054 | -0.4949 | | 1.16E-07 | BLUPmrMLM | SIG |
| FD_18JY | AX-111648969 | 3A | 691218716 | -0.3995 | | 2.10E-05 | BLUPmrMLM | SUG |
| FD_18JY | AX-110491100 | 3A | 705251779 | -1.8934 | | 2.63E-07 | EMMAX | SUG |
| FD_18JY | AX-110491100 | 3A | 705251779 | 1.9003 | | 1.08E-07 | GEMMA | SIG |
| FD_18JY | AX-108941177 | 3A | 705259848 | -1.8193 | | 1.39E-07 | EMMAX | SUG |
| FD_18JY | AX-108941177 | 3A | 705259848 | 1.8193 | | 6.25E-08 | GEMMA | SIG |
| FD_18JY | AX-111695263 | 3A | 705274739 | -1.7860 | | 6.53E-07 | EMMAX | SUG |
| FD_18JY | AX-111695263 | 3A | 705274739 | 1.7888 | | 2.92E-07 | GEMMA | SUG |
| FD_18JY | AX-110976174 | 3A | 705274782 | -3.5204 | | 2.75E-07 | EMMAX | SUG |
| FD_18JY | AX-110976174 | 3A | 705274782 | 3.5197 | | 1.26E-07 | GEMMA | SUG |
| FD_18JY | AX-110484538 | 3A | 705296840 | -4.0393 | | 2.32E-07 | EMMAX | SUG |
| FD_18JY | AX-110484538 | 3A | 705296840 | 4.0361 | | 1.15E-07 | GEMMA | SIG |
| FD_18JY | AX-110484538 | 3A | 705296840 | -1.5497 | | 1.34E-97 | IIIVmrMLM | SIG |
| FD_18JY | AX-111000803 | 3B | 115817070 | -0.2388 | | 1.75E-04 | BLUPmrMLM | SUG |
| FD_18JY | AX-110602814 | 3B | 476265425 | 0.4269 | | 4.87E-08 | BLUPmrMLM | SIG |
| FD_18JY | AX-108800638 | 3B | 495480830 | 0.5927 | | 6.53E-21 | IIIVmrMLM | SIG |
| FD_18JY | AX-109022331 | 3D | 276538349 | -8.7431 | | 1.32E-06 | EMMAX | SUG |
| FD_18JY | AX-109270564 | 3D | 283965882 | -8.7431 | | 1.32E-06 | EMMAX | SUG |
| FD_18JY | AX-94395861 | 4A | 230998878 | -0.8286 | | 3.23E-05 | BLUPmrMLM | SUG |
| FD_18JY | AX-108907684 | 4A | 603659293 | 0.5033 | -0.7216 | 7.09E-15 | IIIVmrMLM | SIG |
| FD_18JY | AX-109816200 | 4A | 649204800 | -0.5815 | | 1.30E-06 | EMMAX | SUG |
| FD_18JY | AX-109816200 | 4A | 649204800 | 0.5838 | | 5.24E-07 | GEMMA | SUG |
| FD_18JY | AX-108812049 | 4A | 650375652 | -0.5764 | | 1.15E-06 | EMMAX | SUG |
| FD_18JY | AX-108812049 | 4A | 650375652 | 0.5758 | | 5.68E-07 | GEMMA | SUG |
| FD_18JY | AX-109817479 | 4A | 657098867 | 0.5100 | | 1.53E-11 | BLUPmrMLM | SIG |
| FD_18JY | AX-109817479 | 4A | 657098867 | -0.5977 | | 6.38E-07 | EMMAX | SUG |
| FD_18JY | AX-109817479 | 4A | 657098867 | 0.5976 | | 2.90E-07 | GEMMA | SUG |
| FD_18JY | AX-109817479 | 4A | 657098867 | -0.3544 | -0.0546 | 8.15E-08 | IIIVmrMLM | SIG |
| FD_18JY | AX-94928833 | 4B | 604204038 | -8.7431 | | 1.32E-06 | EMMAX | SUG |

| | | | | | | | | |
|---|---|---|---|---|---|---|---|---|
| FD_18JY | AX-109246746 | 4D | 302169261 | -0.3120 | | 3.99E-07 | IIIVmrMLM | SUG |
| FD_18JY | AX-110477989 | 5A | 528485397 | -0.6101 | | 7.46E-11 | BLUPmrMLM | SIG |
| FD_18JY | AX-109896503 | 5A | 553097144 | 0.4252 | | 1.79E-05 | BLUPmrMLM | SUG |
| FD_18JY | AX-94464140 | 5B | 69644430 | 5.7140 | | 3.70E-07 | EMMAX | SUG |
| FD_18JY | AX-111544597 | 5B | 429106362 | 0.0645 | 1.2198 | 4.32E-05 | IIIVmrMLM | SUG |
| FD_18JY | AX-110426343 | 5D | 430363560 | -2.5065 | | 9.01E-07 | GEMMA | SUG |
| FD_18JY | AX-110529558 | 6A | 1144693 | 0.0849 | -0.9857 | 2.28E-25 | IIIVmrMLM | SIG |
| FD_18JY | AX-108834300 | 6A | 1805647 | 0.4430 | | 3.40E-06 | BLUPmrMLM | SUG |
| FD_18JY | AX-94425880 | 6B | 685669879 | 5.3594 | | 3.41E-07 | EMMAX | SUG |
| FD_18JY | AX-94506692 | 6B | 716477158 | 0.7277 | | 6.93E-06 | BLUPmrMLM | SUG |
| FD_18JY | AX-109308202 | 7A | 48841017 | -0.3992 | | 1.37E-04 | BLUPmrMLM | SUG |
| FD_18JY | AX-111084178 | 7A | 55338625 | -0.0572 | 0.5035 | 1.45E-07 | IIIVmrMLM | SUG |
| FD_18JY | AX-110053794 | 7B | 125912176 | 0.4462 | | 3.63E-07 | BLUPmrMLM | SUG |
| FD_18JY | AX-111575622 | 7B | 328586876 | 0.4183 | 3.4341 | 1.05E-13 | IIIVmrMLM | SIG |
| FD_18JY | AX-108914113 | 7B | 553640645 | -8.7431 | | 1.32E-06 | EMMAX | SUG |
| FD_18JY | AX-95191225 | 7D | 31435113 | -8.7431 | | 1.32E-06 | EMMAX | SUG |
| FD_18JY | AX-94963724 | 7D | 519966772 | 0.3278 | | 1.05E-04 | BLUPmrMLM | SUG |
| FD_18JY | AX-111034418 | 7D | 619011965 | 0.4071 | | 5.14E-11 | IIIVmrMLM | SIG |
| FD_18XX | AX-110521336 | 1A | 289053265 | 0.3898 | | 1.27E-05 | BLUPmrMLM | SUG |
| FD_18XX | AX-109035993 | 1A | 504787723 | 0.4217 | | 6.17E-05 | BLUPmrMLM | SUG |
| FD_18XX | AX-109035993 | 1A | 504787723 | -0.5850 | 0.0259 | 3.39E-32 | IIIVmrMLM | SIG |
| FD_18XX | AX-108958046 | 1A | 532670133 | -0.6417 | 0.5988 | 5.51E-40 | IIIVmrMLM | SIG |
| FD_18XX | AX-108780402 | 1A | 564570992 | -1.0598 | | 1.12E-08 | FarmCPU | SIG |
| FD_18XX | AX-111627734 | 1B | 62570661 | -0.0473 | -1.2521 | 5.34E-13 | IIIVmrMLM | SIG |
| FD_18XX | AX-109938799 | 1B | 236100245 | 0.2746 | | 8.93E-10 | IIIVmrMLM | SIG |
| FD_18XX | AX-110503854 | 1B | 398790050 | -0.0437 | 0.7269 | 4.78E-10 | IIIVmrMLM | SIG |
| FD_18XX | AX-94778791 | 1B | 407544403 | 8.4893 | | 1.16E-08 | EMMAX | SIG |
| FD_18XX | AX-111022013 | 1B | 549361711 | -0.7100 | | 3.50E-07 | FarmCPU | SUG |
| FD_18XX | AX-109932845 | 1B | 549676006 | -0.0039 | -0.5327 | 2.01E-05 | IIIVmrMLM | SUG |
| FD_18XX | AX-94829936 | 1B | 589489117 | 0.3511 | | 3.41E-07 | BLUPmrMLM | SUG |
| FD_18XX | AX-108803758 | 1B | 595426442 | -0.0076 | -3.0859 | 1.58E-07 | IIIVmrMLM | SUG |
| FD_18XX | AX-110374582 | 1D | 14485920 | 17.3880 | | 1.30E-15 | EMMAX | SIG |
| FD_18XX | AX-111585715 | 1D | 20752378 | -4.7656 | -4.1364 | 0.00E+00 | IIIVmrMLM | SIG |
| FD_18XX | AX-109523909 | 1D | 34626632 | 0.3933 | | 6.28E-06 | BLUPmrMLM | SUG |
| FD_18XX | AX-110299313 | 1D | 90823366 | 17.3880 | | 1.30E-15 | EMMAX | SIG |
| FD_18XX | AX-110302439 | 1D | 230920818 | 12.5390 | | 1.21E-10 | EMMAX | SIG |
| FD_18XX | AX-94498489 | 1D | 447861813 | 17.3880 | | 1.30E-15 | EMMAX | SIG |
| FD_18XX | AX-111035423 | 2A | 136058841 | 0.1253 | -3.0982 | 2.93E-20 | IIIVmrMLM | SIG |
| FD_18XX | AX-94991562 | 2A | 712580484 | -0.5101 | | 6.10E-05 | BLUPmrMLM | SUG |
| FD_18XX | AX-108753274 | 2A | 750145504 | -0.0156 | -1.4489 | 3.97E-07 | IIIVmrMLM | SUG |
| FD_18XX | AX-94478206 | 2B | 19415103 | -0.4775 | | 9.07E-07 | BLUPmrMLM | SUG |
| FD_18XX | AX-94652369 | 2B | 106129285 | -0.3248 | -1.9595 | 1.58E-17 | IIIVmrMLM | SIG |
| FD_18XX | AX-109558760 | 2B | 193395342 | -0.2965 | | 3.85E-06 | BLUPmrMLM | SUG |
| FD_18XX | AX-110528518 | 2B | 211009942 | 17.3880 | | 1.30E-15 | EMMAX | SIG |

| | | | | | | | | |
|---|---|---|---|---|---|---|---|---|
| FD_18XX | AX-95069032 | 2B | 488912865 | 5.4078 | | 1.31E-06 | EMMAX | SUG |
| FD_18XX | AX-110971125 | 2B | 632622394 | 3.6055 | | 1.57E-08 | EMMAX | SIG |
| FD_18XX | AX-109951916 | 2B | 765280116 | 0.0683 | 1.51 | 9.86E-06 | IIIVmrMLM | SUG |
| FD_18XX | AX-110145954 | 2B | 769393040 | -0.4521 | | 3.29E-06 | BLUPmrMLM | SUG |
| FD_18XX | AX-108823023 | 2B | 773357574 | 0.1759 | 3.4969 | 1.38E-14 | IIIVmrMLM | SIG |
| FD_18XX | AX-94976683 | 2D | 569949805 | 0.3707 | -0.0777 | 8.98E-12 | IIIVmrMLM | SIG |
| FD_18XX | AX-108996295 | 3A | 562102052 | 0.3056 | | 6.65E-05 | BLUPmrMLM | SUG |
| FD_18XX | AX-94922900 | 3A | 620755626 | 12.5390 | | 1.21E-10 | EMMAX | SIG |
| FD_18XX | AX-109477508 | 3A | 637900571 | -1.4852 | | 3.95E-07 | FarmCPU | SUG |
| FD_18XX | AX-108977417 | 3A | 652023586 | 0.2977 | 0.2087 | 4.09E-10 | IIIVmrMLM | SIG |
| FD_18XX | AX-108953154 | 3A | 652577528 | -0.4046 | | 2.11E-07 | BLUPmrMLM | SUG |
| FD_18XX | AX-109333590 | 3B | 140848912 | -0.0432 | -1.1809 | 1.24E-07 | IIIVmrMLM | SUG |
| FD_18XX | AX-108791633 | 3B | 154208013 | 3.4119 | | 1.25E-06 | EMMAX | SUG |
| FD_18XX | AX-108791633 | 3B | 154208013 | -3.3906 | | 6.31E-07 | GEMMA | SUG |
| FD_18XX | AX-94471116 | 3B | 203556820 | 17.3880 | | 1.30E-15 | EMMAX | SIG |
| FD_18XX | AX-94607525 | 3B | 262853282 | 6.8790 | | 8.98E-08 | EMMAX | SIG |
| FD_18XX | AX-110427131 | 3B | 469648875 | -0.6591 | | 7.29E-12 | BLUPmrMLM | SIG |
| FD_18XX | AX-110427131 | 3B | 469648875 | 0.8407 | | 1.17E-06 | EMMAX | SUG |
| FD_18XX | AX-110427131 | 3B | 469648875 | -0.8403 | | 9.20E-07 | GEMMA | SUG |
| FD_18XX | AX-110427131 | 3B | 469648875 | 0.2984 | -0.0484 | 1.24E-08 | IIIVmrMLM | SIG |
| FD_18XX | AX-109821150 | 3B | 583382718 | 3.4439 | | 3.05E-08 | EMMAX | SIG |
| FD_18XX | AX-109821150 | 3B | 583382718 | -3.4466 | | 1.60E-08 | GEMMA | SIG |
| FD_18XX | AX-111489277 | 3B | 625336740 | 0.3373 | | 8.13E-14 | IIIVmrMLM | SIG |
| FD_18XX | AX-109357603 | 3B | 667162788 | 2.2225 | | 3.70E-07 | EMMAX | SUG |
| FD_18XX | AX-109357603 | 3B | 667162788 | -2.2335 | | 3.26E-07 | GEMMA | SUG |
| FD_18XX | AX-108960339 | 3B | 792963927 | 3.3271 | | 1.55E-06 | EMMAX | SUG |
| FD_18XX | AX-108960339 | 3B | 792963927 | -3.3529 | | 6.04E-07 | GEMMA | SUG |
| FD_18XX | AX-109415956 | 3B | 810332933 | 2.7398 | | 1.06E-06 | EMMAX | SUG |
| FD_18XX | AX-109415956 | 3B | 810332933 | -2.6899 | | 3.58E-07 | GEMMA | SUG |
| FD_18XX | AX-110416128 | 3B | 810334958 | 5.4796 | | 1.06E-06 | EMMAX | SUG |
| FD_18XX | AX-108845796 | 3B | 810335131 | 5.4796 | | 1.06E-06 | EMMAX | SUG |
| FD_18XX | AX-108920352 | 3B | 810358775 | 5.4796 | | 1.06E-06 | EMMAX | SUG |
| FD_18XX | AX-109376714 | 3B | 810448196 | 2.7398 | | 1.06E-06 | EMMAX | SUG |
| FD_18XX | AX-109376714 | 3B | 810448196 | -2.6899 | | 3.58E-07 | GEMMA | SUG |
| FD_18XX | AX-108783932 | 3B | 810528102 | 5.4796 | | 1.06E-06 | EMMAX | SUG |
| FD_18XX | AX-111565872 | 3B | 810528155 | 2.7398 | | 1.06E-06 | EMMAX | SUG |
| FD_18XX | AX-111565872 | 3B | 810528155 | -2.6899 | | 3.58E-07 | GEMMA | SUG |
| FD_18XX | AX-111196578 | 3B | 810528242 | 5.0777 | | 1.18E-06 | EMMAX | SUG |
| FD_18XX | AX-108782991 | 3B | 810544591 | 5.4796 | | 1.06E-06 | EMMAX | SUG |
| FD_18XX | AX-111131199 | 3B | 810564183 | 5.4796 | | 1.06E-06 | EMMAX | SUG |
| FD_18XX | AX-109017734 | 3B | 810565289 | 5.4796 | | 1.06E-06 | EMMAX | SUG |
| FD_18XX | AX-110060618 | 3B | 810622955 | 5.4796 | | 1.06E-06 | EMMAX | SUG |
| FD_18XX | AX-108757746 | 3B | 810625226 | 5.4796 | | 1.06E-06 | EMMAX | SUG |
| FD_18XX | AX-111657522 | 3B | 810783132 | 5.4796 | | 1.06E-06 | EMMAX | SUG |

| | | | | | | | | |
|---|---|---|---|---|---|---|---|---|
| FD_18XX | AX-108903892 | 3B | 810902194 | 5.4796 | | 1.06E-06 | EMMAX | SUG |
| FD_18XX | AX-108844678 | 3B | 820863427 | -0.0377 | -1.9635 | 2.10E-06 | IIIVmrMLM | SUG |
| FD_18XX | AX-108727595 | 3D | 595882666 | 2.2037 | | 4.59E-07 | FarmCPU | SUG |
| FD_18XX | AX-95188288 | 4A | 21628352 | 5.9751 | | 1.47E-06 | EMMAX | SUG |
| FD_18XX | AX-110458146 | 4A | 38722201 | 0.5688 | | 8.33E-34 | IIIVmrMLM | SIG |
| FD_18XX | AX-110185660 | 4A | 117173937 | 0.8090 | | 1.59E-09 | BLUPmrMLM | SIG |
| FD_18XX | AX-109293865 | 4A | 455296478 | -0.0965 | -2.1009 | 5.00E-11 | IIIVmrMLM | SIG |
| FD_18XX | AX-110163554 | 4A | 710270913 | 0.0195 | -4.6494 | 1.88E-08 | IIIVmrMLM | SIG |
| FD_18XX | AX-109324407 | 4B | 132721966 | 0.0939 | 0.9495 | 6.92E-08 | IIIVmrMLM | SIG |
| FD_18XX | AX-94415914 | 4B | 440328146 | 8.9867 | | 2.73E-07 | EMMAX | SUG |
| FD_18XX | AX-108973407 | 4B | 495635342 | -0.1231 | -0.9628 | 7.38E-13 | IIIVmrMLM | SIG |
| FD_18XX | AX-111136038 | 4B | 496286763 | 0.3482 | | 2.99E-05 | BLUPmrMLM | SUG |
| FD_18XX | AX-110853069 | 4D | 26219559 | -0.8413 | | 1.34E-06 | BLUPmrMLM | SUG |
| FD_18XX | AX-109761837 | 4D | 248705348 | 6.3733 | | 1.27E-06 | EMMAX | SUG |
| FD_18XX | AX-94405671 | 4D | 293383599 | 17.3880 | | 1.30E-15 | EMMAX | SIG |
| FD_18XX | AX-110973123 | 5A | 447951616 | 0.0388 | -0.9557 | 2.38E-08 | IIIVmrMLM | SIG |
| FD_18XX | AX-111689312 | 5A | 547056411 | -0.4877 | 0.4693 | 6.95E-25 | IIIVmrMLM | SIG |
| FD_18XX | AX-109394823 | 5A | 570133695 | -0.3476 | | 5.37E-07 | BLUPmrMLM | SUG |
| FD_18XX | AX-94430561 | 5A | 596013453 | 4.1292 | | 2.02E-07 | EMMAX | SUG |
| FD_18XX | AX-94430561 | 5A | 596013453 | -4.2055 | | 2.49E-08 | GEMMA | SIG |
| FD_18XX | AX-94613594 | 5A | 596013552 | 4.3836 | | 5.30E-08 | EMMAX | SIG |
| FD_18XX | AX-94613594 | 5A | 596013552 | -4.5366 | | 3.40E-09 | GEMMA | SIG |
| FD_18XX | AX-94987862 | 5A | 596369178 | 4.0387 | | 5.11E-08 | EMMAX | SIG |
| FD_18XX | AX-94987862 | 5A | 596369178 | -4.1553 | | 4.14E-09 | GEMMA | SIG |
| FD_18XX | AX-110939576 | 5B | 32811877 | -0.2796 | | 6.65E-05 | BLUPmrMLM | SUG |
| FD_18XX | AX-109821989 | 5B | 565424439 | -3.4822 | | 1.86E-06 | GEMMA | SUG |
| FD_18XX | AX-109821989 | 5B | 565424439 | 0.2241 | -2.6022 | 9.98E-21 | IIIVmrMLM | SIG |
| FD_18XX | AX-111528735 | 5B | 574063657 | -1.5708 | | 1.53E-06 | GEMMA | SUG |
| FD_18XX | AX-108987083 | 5B | 575972456 | 3.2925 | | 3.10E-07 | EMMAX | SUG |
| FD_18XX | AX-108987083 | 5B | 575972456 | -3.2594 | | 1.06E-07 | GEMMA | SIG |
| FD_18XX | AX-110595998 | 5B | 576142349 | -1.6885 | | 1.04E-12 | BLUPmrMLM | SIG |
| FD_18XX | AX-110595998 | 5B | 576142349 | 2.3036 | | 1.72E-10 | EMMAX | SIG |
| FD_18XX | AX-110595998 | 5B | 576142349 | -2.2987 | | 1.53E-11 | GEMMA | SIG |
| FD_18XX | AX-110595998 | 5B | 576142349 | 1.0632 | 2.0782 | 1.53E-89 | IIIVmrMLM | SIG |
| FD_18XX | AX-109978928 | 5B | 578348881 | 1.8019 | | 2.98E-07 | EMMAX | SUG |
| FD_18XX | AX-109978928 | 5B | 578348881 | -1.8574 | | 3.12E-08 | GEMMA | SIG |
| FD_18XX | AX-109921216 | 5B | 582423311 | 2.4647 | | 3.26E-09 | EMMAX | SIG |
| FD_18XX | AX-109921216 | 5B | 582423311 | -2.4454 | | 1.41E-09 | GEMMA | SIG |
| FD_18XX | AX-108931712 | 5B | 583475063 | 3.6308 | | 1.65E-06 | EMMAX | SUG |
| FD_18XX | AX-108931712 | 5B | 583475063 | -3.5874 | | 5.05E-07 | GEMMA | SUG |
| FD_18XX | AX-108946482 | 5B | 583616090 | 4.1292 | | 2.02E-07 | EMMAX | SUG |
| FD_18XX | AX-108946482 | 5B | 583616090 | -4.2055 | | 2.49E-08 | GEMMA | SIG |
| FD_18XX | AX-110627615 | 5B | 583714194 | 4.1292 | | 2.02E-07 | EMMAX | SUG |
| FD_18XX | AX-110627615 | 5B | 583714194 | -4.2055 | | 2.49E-08 | GEMMA | SIG |

| | | | | | | | |
|---|---|---|---|---|---|---|---|
| FD_18XX | AX-111742138 | 5B | 583911580 | 4.1292 | 2.02E-07 | EMMAX | SUG |
| FD_18XX | AX-111742138 | 5B | 583911580 | -4.2055 | 2.49E-08 | GEMMA | SIG |
| FD_18XX | AX-94973664 | 5B | 583913905 | 4.1292 | 2.02E-07 | EMMAX | SUG |
| FD_18XX | AX-94973664 | 5B | 583913905 | -4.2055 | 2.49E-08 | GEMMA | SIG |
| FD_18XX | AX-110447642 | 5B | 584068823 | 2.0646 | 2.02E-07 | EMMAX | SUG |
| FD_18XX | AX-110447642 | 5B | 584068823 | -2.1028 | 2.49E-08 | GEMMA | SIG |
| FD_18XX | AX-110402121 | 5B | 584068898 | 2.0191 | 2.46E-07 | EMMAX | SUG |
| FD_18XX | AX-110402121 | 5B | 584068898 | -2.0568 | 3.20E-08 | GEMMA | SIG |
| FD_18XX | AX-86180204 | 5B | 584069243 | 3.7178 | 2.35E-07 | EMMAX | SUG |
| FD_18XX | AX-86180204 | 5B | 584069243 | -3.7525 | 4.77E-08 | GEMMA | SIG |
| FD_18XX | AX-94457982 | 5B | 584070960 | 4.1292 | 2.02E-07 | EMMAX | SUG |
| FD_18XX | AX-94457982 | 5B | 584070960 | -4.2055 | 2.49E-08 | GEMMA | SIG |
| FD_18XX | AX-110740302 | 5B | 584108556 | 4.1292 | 2.02E-07 | EMMAX | SUG |
| FD_18XX | AX-110740302 | 5B | 584108556 | -4.2055 | 2.49E-08 | GEMMA | SIG |
| FD_18XX | AX-111675975 | 5B | 584109393 | 2.0646 | 2.02E-07 | EMMAX | SUG |
| FD_18XX | AX-111675975 | 5B | 584109393 | -2.1028 | 2.49E-08 | GEMMA | SIG |
| FD_18XX | AX-111019260 | 5B | 584110237 | 4.1292 | 2.02E-07 | EMMAX | SUG |
| FD_18XX | AX-111019260 | 5B | 584110237 | -4.2055 | 2.49E-08 | GEMMA | SIG |
| FD_18XX | AX-109879550 | 5B | 584110814 | 4.1292 | 2.02E-07 | EMMAX | SUG |
| FD_18XX | AX-109879550 | 5B | 584110814 | -4.2055 | 2.49E-08 | GEMMA | SIG |
| FD_18XX | AX-109529914 | 5B | 584111028 | 4.1292 | 2.02E-07 | EMMAX | SUG |
| FD_18XX | AX-109529914 | 5B | 584111028 | -4.2055 | 2.49E-08 | GEMMA | SIG |
| FD_18XX | AX-111196036 | 5B | 584142374 | 3.6308 | 1.65E-06 | EMMAX | SUG |
| FD_18XX | AX-111196036 | 5B | 584142374 | -3.5874 | 5.05E-07 | GEMMA | SUG |
| FD_18XX | AX-111542649 | 5B | 584582150 | 4.1292 | 2.02E-07 | EMMAX | SUG |
| FD_18XX | AX-111542649 | 5B | 584582150 | -4.2055 | 2.49E-08 | GEMMA | SIG |
| FD_18XX | AX-108945007 | 5B | 584616343 | 4.1292 | 2.02E-07 | EMMAX | SUG |
| FD_18XX | AX-108945007 | 5B | 584616343 | -4.2055 | 2.49E-08 | GEMMA | SIG |
| FD_18XX | AX-109907290 | 5B | 584639041 | 4.1292 | 2.02E-07 | EMMAX | SUG |
| FD_18XX | AX-109907290 | 5B | 584639041 | -4.2055 | 2.49E-08 | GEMMA | SIG |
| FD_18XX | AX-108916036 | 5B | 584644380 | 3.0803 | 2.92E-10 | EMMAX | SIG |
| FD_18XX | AX-108916036 | 5B | 584644380 | -3.0605 | 7.14E-11 | GEMMA | SIG |
| FD_18XX | AX-109373963 | 5B | 584796270 | 4.1292 | 2.02E-07 | EMMAX | SUG |
| FD_18XX | AX-109373963 | 5B | 584796270 | -4.2055 | 2.49E-08 | GEMMA | SIG |
| FD_18XX | AX-109443147 | 5B | 584796550 | 4.1292 | 2.02E-07 | EMMAX | SUG |
| FD_18XX | AX-109443147 | 5B | 584796550 | -4.2055 | 2.49E-08 | GEMMA | SIG |
| FD_18XX | AX-108833623 | 5B | 584796716 | 3.6308 | 1.65E-06 | EMMAX | SUG |
| FD_18XX | AX-108833623 | 5B | 584796716 | -3.5874 | 5.05E-07 | GEMMA | SUG |
| FD_18XX | AX-110904949 | 5B | 584825940 | -3.3751 | 9.81E-07 | GEMMA | SUG |
| FD_18XX | AX-109005670 | 5B | 584855079 | -3.5346 | 5.84E-07 | GEMMA | SUG |
| FD_18XX | AX-94504636 | 5B | 584884520 | 3.8585 | 4.14E-07 | EMMAX | SUG |
| FD_18XX | AX-94504636 | 5B | 584884520 | -3.9003 | 7.21E-08 | GEMMA | SIG |
| FD_18XX | AX-109468686 | 5B | 584894906 | 2.0708 | 1.88E-07 | EMMAX | SUG |
| FD_18XX | AX-109468686 | 5B | 584894906 | -2.0750 | 3.58E-08 | GEMMA | SIG |

| | | | | | | | | |
|---|---|---|---|---|---|---|---|---|
| FD_18XX | AX-111458601 | 5B | 585025969 | 3.8585 | | 4.14E-07 | EMMAX | SUG |
| FD_18XX | AX-111458601 | 5B | 585025969 | -3.9003 | | 7.21E-08 | GEMMA | SIG |
| FD_18XX | AX-109008501 | 5B | 587803667 | 2.6998 | | 1.21E-08 | EMMAX | SIG |
| FD_18XX | AX-109008501 | 5B | 587803667 | -2.7143 | | 1.94E-09 | GEMMA | SIG |
| FD_18XX | AX-109368028 | 5B | 616441050 | -0.2967 | -0.3291 | 3.14E-10 | IIIVmrMLM | SIG |
| FD_18XX | AX-111662794 | 5B | 663821905 | 0.0786 | -0.7596 | 1.04E-05 | IIIVmrMLM | SUG |
| FD_18XX | AX-108928208 | 5D | 40773135 | 17.3880 | | 1.30E-15 | EMMAX | SIG |
| FD_18XX | AX-94785649 | 5D | 87185087 | 9.5322 | | 1.31E-11 | EMMAX | SIG |
| FD_18XX | AX-111913736 | 5D | 134637128 | 8.7591 | | 1.59E-11 | EMMAX | SIG |
| FD_18XX | AX-111045825 | 5D | 159257059 | 11.7230 | | 1.02E-12 | EMMAX | SIG |
| FD_18XX | AX-111857618 | 5D | 164294326 | 16.9090 | | 2.73E-15 | EMMAX | SIG |
| FD_18XX | AX-111208523 | 5D | 259235163 | 9.4776 | | 2.68E-08 | EMMAX | SIG |
| FD_18XX | AX-111754571 | 5D | 266755425 | 0.3599 | | 1.95E-15 | IIIVmrMLM | SIG |
| FD_18XX | AX-110459577 | 5D | 344875146 | 8.4665 | | 5.23E-09 | EMMAX | SIG |
| FD_18XX | AX-109511204 | 5D | 365700652 | 7.7266 | | 4.20E-07 | EMMAX | SUG |
| FD_18XX | AX-109762451 | 5D | 405638329 | 17.3880 | | 1.30E-15 | EMMAX | SIG |
| FD_18XX | AX-111497172 | 5D | 454427605 | -0.2272 | -0.0055 | 3.62E-06 | IIIVmrMLM | SUG |
| FD_18XX | AX-95112546 | 5D | 464092775 | 7.9351 | | 4.69E-08 | EMMAX | SIG |
| FD_18XX | AX-109360916 | 5D | 469286340 | 3.0734 | | 1.79E-09 | EMMAX | SIG |
| FD_18XX | AX-109360916 | 5D | 469286340 | -3.1094 | | 1.22E-09 | GEMMA | SIG |
| FD_18XX | AX-109360916 | 5D | 469286340 | 0.9626 | | 1.95E-78 | IIIVmrMLM | SIG |
| FD_18XX | AX-111177872 | 5D | 476671897 | 3.6308 | | 1.65E-06 | EMMAX | SUG |
| FD_18XX | AX-111177872 | 5D | 476671897 | -3.5874 | | 5.05E-07 | GEMMA | SUG |
| FD_18XX | AX-94486090 | 5D | 487747659 | 5.6856 | | 1.60E-06 | EMMAX | SUG |
| FD_18XX | AX-94609037 | 5D | 493834097 | 0.5285 | | 7.56E-30 | IIIVmrMLM | SIG |
| FD_18XX | AX-94951925 | 5D | 543803852 | -0.3076 | -0.6615 | 1.36E-19 | IIIVmrMLM | SIG |
| FD_18XX | AX-110954471 | 6A | 2950843 | 0.1047 | -0.5672 | 2.43E-09 | IIIVmrMLM | SIG |
| FD_18XX | AX-95215294 | 6A | 17533994 | 0.2694 | | 5.79E-05 | BLUPmrMLM | SUG |
| FD_18XX | AX-108974037 | 6A | 100258422 | 4.4452 | | 4.51E-07 | EMMAX | SUG |
| FD_18XX | AX-108974037 | 6A | 100258422 | -4.5018 | | 3.21E-07 | GEMMA | SUG |
| FD_18XX | AX-110565182 | 6A | 100282138 | 4.4452 | | 4.51E-07 | EMMAX | SUG |
| FD_18XX | AX-110565182 | 6A | 100282138 | -4.5018 | | 3.21E-07 | GEMMA | SUG |
| FD_18XX | AX-108793857 | 6A | 100358812 | 4.4452 | | 4.51E-07 | EMMAX | SUG |
| FD_18XX | AX-108793857 | 6A | 100358812 | -4.5018 | | 3.21E-07 | GEMMA | SUG |
| FD_18XX | AX-111477610 | 6A | 100371870 | -2.0821 | | 1.71E-06 | GEMMA | SUG |
| FD_18XX | AX-94529047 | 6A | 107799490 | -2.2654 | | 2.23E-06 | GEMMA | SUG |
| FD_18XX | AX-109530167 | 6A | 117072369 | 2.4764 | | 3.19E-07 | EMMAX | SUG |
| FD_18XX | AX-109530167 | 6A | 117072369 | -2.4965 | | 2.18E-07 | GEMMA | SUG |
| FD_18XX | AX-111488753 | 6A | 125881807 | -2.2654 | | 2.23E-06 | GEMMA | SUG |
| FD_18XX | AX-109879343 | 6A | 498830663 | -0.3785 | | 3.60E-06 | BLUPmrMLM | SUG |
| FD_18XX | AX-111454028 | 6A | 517491134 | 3.7143 | | 3.47E-07 | EMMAX | SUG |
| FD_18XX | AX-111476435 | 6B | 75268723 | 12.5390 | | 1.21E-10 | EMMAX | SIG |
| FD_18XX | AX-110384791 | 6B | 102066396 | -0.3296 | | 2.05E-05 | BLUPmrMLM | SUG |
| FD_18XX | AX-110997969 | 6B | 106540703 | 0.3364 | | 9.41E-14 | IIIVmrMLM | SIG |

| | | | | | | | | |
|---|---|---|---|---|---|---|---|---|
| FD_18XX | AX-110360699 | 6B | 201254117 | -0.5674 | | 7.98E-08 | BLUPmrMLM | SIG |
| FD_18XX | AX-108954852 | 6B | 682326705 | 0.3550 | | 5.17E-09 | BLUPmrMLM | SIG |
| FD_18XX | AX-94949358 | 6B | 716259413 | -0.2681 | | 5.81E-05 | BLUPmrMLM | SUG |
| FD_18XX | AX-111482552 | 6D | 34505874 | 6.9622 | | 5.41E-08 | EMMAX | SIG |
| FD_18XX | AX-94413274 | 6D | 292064315 | 8.1433 | | 1.41E-07 | EMMAX | SUG |
| FD_18XX | AX-111339758 | 6D | 412586914 | -0.2647 | | 9.00E-06 | BLUPmrMLM | SUG |
| FD_18XX | AX-108838582 | 6D | 451803002 | 17.3880 | | 1.30E-15 | EMMAX | SIG |
| FD_18XX | AX-108896352 | 6D | 465742631 | -0.3801 | 0.0138 | 1.53E-10 | IIIVmrMLM | SIG |
| FD_18XX | AX-109912510 | 7A | 48483355 | -0.4506 | | 1.19E-07 | BLUPmrMLM | SUG |
| FD_18XX | AX-110128567 | 7A | 225701352 | -0.1292 | 0.6453 | 3.72E-08 | IIIVmrMLM | SIG |
| FD_18XX | AX-94442209 | 7A | 430242647 | 17.3880 | | 1.30E-15 | EMMAX | SIG |
| FD_18XX | AX-110907641 | 7A | 548334437 | 7.4953 | | 8.02E-08 | EMMAX | SIG |
| FD_18XX | AX-109850764 | 7A | 644610427 | 0.0448 | -2.7958 | 1.38E-10 | IIIVmrMLM | SIG |
| FD_18XX | AX-109819108 | 7A | 652230718 | -0.5610 | | 1.32E-13 | BLUPmrMLM | SIG |
| FD_18XX | AX-109819108 | 7A | 652230718 | 0.3034 | -0.1793 | 1.73E-10 | IIIVmrMLM | SIG |
| FD_18XX | AX-108866381 | 7A | 691831775 | -0.0810 | -0.6202 | 6.61E-12 | IIIVmrMLM | SIG |
| FD_18XX | AX-109438992 | 7B | 12196726 | -0.4126 | | 6.21E-05 | BLUPmrMLM | SUG |
| FD_18XX | AX-108786109 | 7B | 46682947 | 0.0015 | 2.0878 | 1.78E-08 | IIIVmrMLM | SIG |
| FD_18XX | AX-94571213 | 7B | 126556450 | 17.3880 | | 1.30E-15 | EMMAX | SIG |
| FD_18XX | AX-95160924 | 7B | 601117142 | -0.6781 | | 1.62E-07 | BLUPmrMLM | SUG |
| FD_18XX | AX-95149392 | 7B | 713217152 | 17.3880 | | 1.30E-15 | EMMAX | SIG |
| FD_18XX | AX-111000473 | 7D | 138439894 | 8.7207 | | 1.72E-09 | EMMAX | SIG |
| FD_18XX | AX-94820538 | 7D | 167228862 | 12.9450 | | 3.36E-11 | EMMAX | SIG |
| FD_18XX | AX-111590438 | 7D | 205149703 | 0.2277 | | 3.33E-07 | IIIVmrMLM | SUG |
| FD_18XX | AX-109642382 | 7D | 355887139 | 8.6388 | | 7.66E-08 | EMMAX | SIG |
| FD_18XX | AX-111726828 | 7D | 409610208 | 12.5390 | | 1.21E-10 | EMMAX | SIG |
| FD_18XX | AX-111577993 | 7D | 472285202 | 0.4811 | | 6.90E-05 | BLUPmrMLM | SUG |
| FD_18XX | AX-110923250 | 7D | 566899316 | 1.4130 | | 3.50E-08 | FarmCPU | SIG |
| FD_19XX | AX-109844324 | 1A | 5036399 | 0.5974 | | 4.79E-06 | BLUPmrMLM | SUG |
| FD_19XX | AX-109504656 | 1A | 495122808 | -0.8928 | | 3.64E-06 | BLUPmrMLM | SUG |
| FD_19XX | AX-109276285 | 1A | 507078569 | 1.1126 | | 3.99E-07 | BLUPmrMLM | SUG |
| FD_19XX | AX-94882326 | 1A | 518972289 | -0.3290 | | 8.58E-09 | IIIVmrMLM | SIG |
| FD_19XX | AX-109276138 | 1A | 532808340 | 1.0657 | 2.0871 | 1.64E-61 | IIIVmrMLM | SIG |
| FD_19XX | AX-111115691 | 1A | 532935799 | 3.3589 | | 1.40E-08 | EMMAX | SIG |
| FD_19XX | AX-111115691 | 1A | 532935799 | -3.4002 | | 1.10E-08 | GEMMA | SIG |
| FD_19XX | AX-111602618 | 1A | 533151001 | -0.0073 | 3.8418 | 3.55E-14 | IIIVmrMLM | SIG |
| FD_19XX | AX-110385762 | 1A | 539598829 | -0.0409 | -2.5485 | 1.13E-07 | IIIVmrMLM | SIG |
| FD_19XX | AX-109283005 | 1B | 384239973 | -0.2004 | 1.2292 | 1.20E-11 | IIIVmrMLM | SIG |
| FD_19XX | AX-109043016 | 1B | 538461468 | 0.5728 | | 1.88E-22 | IIIVmrMLM | SIG |
| FD_19XX | AX-111147652 | 1B | 662213025 | 6.0411 | | 5.37E-07 | EMMAX | SUG |
| FD_19XX | AX-111147652 | 1B | 662213025 | -6.0077 | | 2.28E-07 | GEMMA | SUG |
| FD_19XX | AX-109586144 | 1B | 662489030 | 0.8927 | -0.3967 | 1.28E-44 | IIIVmrMLM | SIG |
| FD_19XX | AX-109832121 | 1B | 662940826 | -4.8174 | | 1.40E-06 | GEMMA | SUG |
| FD_19XX | AX-109917800 | 1D | 10157555 | 0.2947 | -2.9034 | 1.60E-17 | IIIVmrMLM | SIG |

| | | | | | | | | |
|---|---|---|---|---|---|---|---|---|
| FD_19XX | AX-111340373 | 1D | 284308114 | -0.4935 | -12.3509 | 4.76E-45 | IIIVmrMLM | SIG |
| FD_19XX | AX-111012152 | 1D | 345728083 | 2.4093 | | 1.18E-06 | EMMAX | SUG |
| FD_19XX | AX-111012152 | 1D | 345728083 | -2.4608 | | 1.01E-06 | GEMMA | SUG |
| FD_19XX | AX-94464776 | 1D | 419061486 | -0.3583 | -2.3798 | 4.80E-17 | IIIVmrMLM | SIG |
| FD_19XX | AX-112288504 | 2A | 77528386 | 0.4374 | | 2.50E-05 | BLUPmrMLM | SUG |
| FD_19XX | AX-110951755 | 2A | 114862888 | 0.5119 | | 7.44E-05 | BLUPmrMLM | SUG |
| FD_19XX | AX-94464444 | 2A | 293053413 | 0.2629 | 4.2058 | 1.02E-27 | IIIVmrMLM | SIG |
| FD_19XX | AX-95191599 | 2A | 319029074 | 0.7099 | | 9.41E-05 | BLUPmrMLM | SUG |
| FD_19XX | AX-111106303 | 2B | 13429106 | 0.6088 | | 1.17E-05 | BLUPmrMLM | SUG |
| FD_19XX | AX-109360074 | 2B | 706565459 | -1.2567 | | 5.80E-10 | BLUPmrMLM | SIG |
| FD_19XX | AX-110523484 | 2B | 725104222 | -0.3963 | | 3.73E-05 | BLUPmrMLM | SUG |
| FD_19XX | AX-111554268 | 2B | 789434794 | 0.1462 | -2.9505 | 3.17E-08 | IIIVmrMLM | SIG |
| FD_19XX | AX-110606479 | 2B | 800257921 | 0.2160 | 1.1163 | 1.20E-11 | IIIVmrMLM | SIG |
| FD_19XX | AX-94689584 | 3A | 10592927 | -0.2204 | -2.8569 | 8.76E-16 | IIIVmrMLM | SIG |
| FD_19XX | AX-111562115 | 3A | 278156480 | 0.0352 | -2.7487 | 1.42E-09 | IIIVmrMLM | SIG |
| FD_19XX | AX-110370081 | 3A | 427893261 | -0.2050 | -4.6479 | 5.98E-11 | IIIVmrMLM | SIG |
| FD_19XX | AX-109991086 | 3A | 547376579 | 0.1636 | -4.7885 | 1.96E-10 | IIIVmrMLM | SIG |
| FD_19XX | AX-111161673 | 3A | 557419004 | -0.0804 | -1.7698 | 3.38E-10 | IIIVmrMLM | SIG |
| FD_19XX | AX-111579209 | 3A | 724038650 | -0.6731 | | 1.87E-07 | BLUPmrMLM | SUG |
| FD_19XX | AX-108793968 | 3B | 25312665 | -0.8393 | | 9.04E-06 | BLUPmrMLM | SUG |
| FD_19XX | AX-111129787 | 3B | 140859817 | -0.2107 | -0.7671 | 7.91E-16 | IIIVmrMLM | SIG |
| FD_19XX | AX-109508415 | 3B | 172700618 | 0.3199 | -6.411 | 6.30E-25 | IIIVmrMLM | SIG |
| FD_19XX | AX-111833531 | 3B | 332237107 | -2.7157 | | 5.19E-07 | FarmCPU | SUG |
| FD_19XX | AX-110736238 | 3B | 380394917 | 0.0873 | -1.2419 | 3.76E-12 | IIIVmrMLM | SIG |
| FD_19XX | AX-111474822 | 3B | 419233398 | -6.0188 | | 1.45E-06 | FarmCPU | SUG |
| FD_19XX | AX-111057460 | 3B | 759231400 | -0.4248 | | 1.30E-05 | BLUPmrMLM | SUG |
| FD_19XX | AX-111453739 | 3B | 759265358 | 0.3727 | -0.2186 | 1.08E-09 | IIIVmrMLM | SIG |
| FD_19XX | AX-94391826 | 3D | 4066279 | 0.1976 | -0.9864 | 8.21E-15 | IIIVmrMLM | SIG |
| FD_19XX | AX-108738283 | 3D | 479119536 | -0.1819 | -4.6936 | 5.94E-13 | IIIVmrMLM | SIG |
| FD_19XX | AX-110472229 | 4A | 598587967 | 0.0385 | -2.5622 | 9.87E-08 | IIIVmrMLM | SIG |
| FD_19XX | AX-110077550 | 4A | 610038559 | 2.0446 | | 8.86E-07 | EMMAX | SUG |
| FD_19XX | AX-110077550 | 4A | 610038559 | -2.0339 | | 4.42E-07 | GEMMA | SUG |
| FD_19XX | AX-111105228 | 4A | 610609020 | 2.7913 | | 6.87E-07 | EMMAX | SUG |
| FD_19XX | AX-111105228 | 4A | 610609020 | -2.7394 | | 4.84E-07 | GEMMA | SUG |
| FD_19XX | AX-109846294 | 4A | 610629973 | 3.5959 | | 1.97E-06 | EMMAX | SUG |
| FD_19XX | AX-109846294 | 4A | 610629973 | -3.5191 | | 1.47E-06 | GEMMA | SUG |
| FD_19XX | AX-86183809 | 4A | 612060283 | 4.6491 | | 1.08E-07 | EMMAX | SIG |
| FD_19XX | AX-86183809 | 4A | 612060283 | -4.5712 | | 1.17E-07 | GEMMA | SUG |
| FD_19XX | AX-110730034 | 4A | 612088918 | 1.8992 | | 6.77E-07 | EMMAX | SUG |
| FD_19XX | AX-110730034 | 4A | 612088918 | -1.8830 | | 4.47E-07 | GEMMA | SUG |
| FD_19XX | AX-89491834 | 4A | 612091001 | -1.7550 | | 1.90E-06 | GEMMA | SUG |
| FD_19XX | AX-111033323 | 4A | 612437711 | -3.6015 | | 1.97E-06 | GEMMA | SUG |
| FD_19XX | AX-109043286 | 4A | 612484737 | -3.6015 | | 1.97E-06 | GEMMA | SUG |
| FD_19XX | AX-111483645 | 4A | 612492000 | 1.8713 | | 1.91E-06 | EMMAX | SUG |

| | | | | | | | | |
|---|---|---|---|---|---|---|---|---|
| FD_19XX | AX-111483645 | 4A | 612492000 | -1.8771 | | 8.23E-07 | GEMMA | SUG |
| FD_19XX | AX-111149628 | 4A | 612493042 | 1.8713 | | 1.91E-06 | EMMAX | SUG |
| FD_19XX | AX-111149628 | 4A | 612493042 | -1.8771 | | 8.23E-07 | GEMMA | SUG |
| FD_19XX | AX-110371776 | 4A | 612493121 | 1.8713 | | 1.91E-06 | EMMAX | SUG |
| FD_19XX | AX-110371776 | 4A | 612493121 | -1.8771 | | 8.23E-07 | GEMMA | SUG |
| FD_19XX | AX-111768266 | 4A | 612494779 | -3.6015 | | 1.97E-06 | GEMMA | SUG |
| FD_19XX | AX-110926040 | 4A | 612517494 | 1.8713 | | 1.91E-06 | EMMAX | SUG |
| FD_19XX | AX-110926040 | 4A | 612517494 | -1.8771 | | 8.23E-07 | GEMMA | SUG |
| FD_19XX | AX-109589014 | 4A | 612639170 | -3.6015 | | 1.97E-06 | GEMMA | SUG |
| FD_19XX | AX-108861327 | 4A | 612642486 | -3.6015 | | 1.97E-06 | GEMMA | SUG |
| FD_19XX | AX-111493275 | 4A | 612662027 | -3.6015 | | 1.97E-06 | GEMMA | SUG |
| FD_19XX | AX-111705164 | 4A | 612667681 | -1.5101 | | 3.26E-11 | BLUPmrMLM | SIG |
| FD_19XX | AX-111705164 | 4A | 612667681 | 1.9639 | | 5.32E-07 | EMMAX | SUG |
| FD_19XX | AX-111705164 | 4A | 612667681 | -1.9700 | | 2.10E-07 | GEMMA | SUG |
| FD_19XX | AX-110363512 | 4A | 612680091 | 1.8799 | | 1.58E-06 | EMMAX | SUG |
| FD_19XX | AX-110363512 | 4A | 612680091 | -1.8810 | | 7.28E-07 | GEMMA | SUG |
| FD_19XX | AX-111581594 | 4A | 612973608 | 1.9324 | | 1.10E-06 | EMMAX | SUG |
| FD_19XX | AX-111581594 | 4A | 612973608 | -1.9382 | | 4.60E-07 | GEMMA | SUG |
| FD_19XX | AX-110432011 | 4A | 613078092 | 3.8418 | | 8.22E-07 | EMMAX | SUG |
| FD_19XX | AX-110432011 | 4A | 613078092 | -3.8480 | | 3.65E-07 | GEMMA | SUG |
| FD_19XX | AX-109971768 | 4A | 613118940 | 3.8418 | | 8.22E-07 | EMMAX | SUG |
| FD_19XX | AX-109971768 | 4A | 613118940 | -3.8480 | | 3.65E-07 | GEMMA | SUG |
| FD_19XX | AX-111457878 | 4A | 613170537 | 1.8705 | | 1.52E-06 | EMMAX | SUG |
| FD_19XX | AX-111457878 | 4A | 613170537 | -1.8710 | | 7.08E-07 | GEMMA | SUG |
| FD_19XX | AX-109281612 | 4A | 613337260 | 4.1439 | | 3.76E-08 | EMMAX | SIG |
| FD_19XX | AX-109281612 | 4A | 613337260 | -4.1179 | | 1.99E-08 | GEMMA | SIG |
| FD_19XX | AX-110407094 | 4A | 613365713 | 3.4807 | | 6.04E-08 | EMMAX | SIG |
| FD_19XX | AX-110407094 | 4A | 613365713 | -3.4109 | | 6.27E-08 | GEMMA | SIG |
| FD_19XX | AX-95182838 | 4A | 613367700 | 3.2544 | | 9.23E-08 | EMMAX | SIG |
| FD_19XX | AX-95182838 | 4A | 613367700 | -3.1911 | | 9.71E-08 | GEMMA | SIG |
| FD_19XX | AX-111638733 | 4A | 613368868 | 2.0447 | | 8.90E-08 | EMMAX | SIG |
| FD_19XX | AX-111638733 | 4A | 613368868 | -2.0304 | | 4.82E-08 | GEMMA | SIG |
| FD_19XX | AX-110501116 | 4A | 613368960 | 2.0447 | | 8.90E-08 | EMMAX | SIG |
| FD_19XX | AX-110501116 | 4A | 613368960 | -2.0304 | | 4.82E-08 | GEMMA | SIG |
| FD_19XX | AX-109376087 | 4A | 613369207 | 3.4807 | | 6.04E-08 | EMMAX | SIG |
| FD_19XX | AX-109376087 | 4A | 613369207 | -3.4109 | | 6.27E-08 | GEMMA | SIG |
| FD_19XX | AX-109456475 | 4A | 613404437 | 2.0447 | | 8.90E-08 | EMMAX | SIG |
| FD_19XX | AX-109456475 | 4A | 613404437 | -2.0304 | | 4.82E-08 | GEMMA | SIG |
| FD_19XX | AX-109424652 | 4A | 615937289 | -0.1059 | -2.7483 | 1.20E-07 | IIIVmrMLM | SUG |
| FD_19XX | AX-111630051 | 4A | 617355325 | 0.4940 | | 6.14E-07 | BLUPmrMLM | SUG |
| FD_19XX | AX-108795868 | 4A | 673679914 | -0.1469 | -0.9042 | 4.28E-14 | IIIVmrMLM | SIG |
| FD_19XX | AX-111587367 | 4A | 706726644 | 0.1214 | -8.5137 | 3.14E-32 | IIIVmrMLM | SIG |
| FD_19XX | AX-94729361 | 4A | 708650486 | -1.3123 | | 1.39E-05 | BLUPmrMLM | SUG |
| FD_19XX | AX-94729361 | 4A | 708650486 | 3.2597 | | 8.19E-09 | EMMAX | SIG |

| | | | | | | | | |
|---|---|---|---|---|---|---|---|---|
| FD_19XX | AX-94729361 | 4A | 708650486 | -3.2580 | | 4.81E-09 | GEMMA | SIG |
| FD_19XX | AX-110383634 | 4B | 28716715 | -0.9680 | | 4.36E-06 | BLUPmrMLM | SUG |
| FD_19XX | AX-108764506 | 4B | 95130098 | 1.3922 | | 2.19E-09 | BLUPmrMLM | SIG |
| FD_19XX | AX-108764506 | 4B | 95130098 | -2.7334 | | 4.95E-07 | EMMAX | SUG |
| FD_19XX | AX-108764506 | 4B | 95130098 | 2.9391 | | 2.59E-07 | GEMMA | SUG |
| FD_19XX | AX-111097142 | 4B | 608663022 | 1.3498 | | 2.18E-11 | BLUPmrMLM | SIG |
| FD_19XX | AX-110994384 | 4B | 626459030 | 0.5089 | | 1.58E-05 | BLUPmrMLM | SUG |
| FD_19XX | AX-108852281 | 4B | 647668190 | 0.3331 | -4.0965 | 1.16E-20 | IIIVmrMLM | SIG |
| FD_19XX | AX-108939109 | 4D | 482135480 | -0.5652 | -0.0098 | 1.07E-20 | IIIVmrMLM | SIG |
| FD_19XX | AX-111558091 | 5A | 7170601 | -4.4688 | | 9.28E-08 | FarmCPU | SIG |
| FD_19XX | AX-109333173 | 5A | 36015682 | 0.8356 | | 6.96E-06 | BLUPmrMLM | SUG |
| FD_19XX | AX-109520332 | 5A | 247573224 | -0.1298 | -4.3607 | 9.70E-21 | IIIVmrMLM | SIG |
| FD_19XX | AX-109423646 | 5A | 389162899 | 0.0475 | -0.9473 | 4.55E-07 | IIIVmrMLM | SUG |
| FD_19XX | AX-111469175 | 5A | 454763388 | 1.0928 | | 3.66E-08 | BLUPmrMLM | SIG |
| FD_19XX | AX-110169212 | 5A | 455055483 | -0.0775 | -0.8977 | 1.39E-07 | IIIVmrMLM | SUG |
| FD_19XX | AX-110360199 | 5A | 455550448 | 1.4127 | | 1.18E-06 | GEMMA | SUG |
| FD_19XX | AX-109885414 | 5A | 455859986 | 1.3825 | | 2.10E-06 | GEMMA | SUG |
| FD_19XX | AX-111614571 | 5A | 455892282 | 1.3825 | | 2.10E-06 | GEMMA | SUG |
| FD_19XX | AX-94536406 | 5A | 455998344 | -2.8133 | | 1.46E-06 | EMMAX | SUG |
| FD_19XX | AX-94536406 | 5A | 455998344 | 2.8320 | | 5.45E-07 | GEMMA | SUG |
| FD_19XX | AX-94456504 | 5A | 455998430 | 1.3825 | | 2.10E-06 | GEMMA | SUG |
| FD_19XX | AX-110520341 | 5A | 456025515 | -1.4435 | | 1.94E-06 | EMMAX | SUG |
| FD_19XX | AX-110520341 | 5A | 456025515 | 1.4551 | | 7.34E-07 | GEMMA | SUG |
| FD_19XX | AX-108729319 | 5A | 456067035 | 1.3825 | | 2.10E-06 | GEMMA | SUG |
| FD_19XX | AX-109894323 | 5A | 456080840 | 2.7390 | | 1.97E-06 | GEMMA | SUG |
| FD_19XX | AX-94455614 | 5A | 456608136 | 2.7748 | | 1.74E-06 | GEMMA | SUG |
| FD_19XX | AX-109823787 | 5A | 510081595 | -2.2014 | | 1.92E-06 | GEMMA | SUG |
| FD_19XX | AX-109837859 | 5A | 511405176 | -1.0558 | | 4.97E-06 | BLUPmrMLM | SUG |
| FD_19XX | AX-109837859 | 5A | 511405176 | 2.5379 | | 2.91E-07 | EMMAX | SUG |
| FD_19XX | AX-109837859 | 5A | 511405176 | -2.5329 | | 1.77E-07 | GEMMA | SUG |
| FD_19XX | AX-111641807 | 5A | 538948758 | -0.0611 | -2.79 | 1.04E-11 | IIIVmrMLM | SIG |
| FD_19XX | AX-111235500 | 5A | 556150580 | 0.7457 | | 8.34E-06 | BLUPmrMLM | SUG |
| FD_19XX | AX-109503011 | 5A | 590599714 | 4.6748 | | 1.79E-06 | EMMAX | SUG |
| FD_19XX | AX-109503011 | 5A | 590599714 | -4.6236 | | 1.29E-06 | GEMMA | SUG |
| FD_19XX | AX-111747299 | 5A | 590680035 | 5.0238 | | 1.29E-06 | EMMAX | SUG |
| FD_19XX | AX-111747299 | 5A | 590680035 | -4.9596 | | 1.02E-06 | GEMMA | SUG |
| FD_19XX | AX-110593755 | 5A | 590848863 | 5.0238 | | 1.29E-06 | EMMAX | SUG |
| FD_19XX | AX-110593755 | 5A | 590848863 | -4.9596 | | 1.02E-06 | GEMMA | SUG |
| FD_19XX | AX-109497126 | 5A | 590996771 | 5.1965 | | 5.80E-07 | EMMAX | SUG |
| FD_19XX | AX-109497126 | 5A | 590996771 | -5.1305 | | 4.41E-07 | GEMMA | SUG |
| FD_19XX | AX-111057571 | 5A | 591004477 | 2.7429 | | 2.13E-07 | EMMAX | SUG |
| FD_19XX | AX-111057571 | 5A | 591004477 | -2.7211 | | 1.44E-07 | GEMMA | SUG |
| FD_19XX | AX-95659239 | 5A | 591091029 | 5.0399 | | 4.69E-07 | EMMAX | SUG |
| FD_19XX | AX-95659239 | 5A | 591091029 | -4.9986 | | 3.09E-07 | GEMMA | SUG |

| | | | | | | | | |
|---|---|---|---|---|---|---|---|---|
| FD_19XX | AX-108893364 | 5A | 591097621 | 5.0399 | | 4.69E-07 | EMMAX | SUG |
| FD_19XX | AX-108893364 | 5A | 591097621 | -4.9986 | | 3.09E-07 | GEMMA | SUG |
| FD_19XX | AX-110435701 | 5A | 591126755 | 2.4647 | | 9.54E-07 | EMMAX | SUG |
| FD_19XX | AX-110435701 | 5A | 591126755 | -2.4441 | | 6.29E-07 | GEMMA | SUG |
| FD_19XX | AX-111541150 | 5A | 670137320 | -0.3696 | -2.5627 | 1.58E-26 | IIIVmrMLM | SIG |
| FD_19XX | AX-111598398 | 5B | 346306475 | 0.7183 | | 6.24E-10 | BLUPmrMLM | SIG |
| FD_19XX | AX-89323998 | 5B | 582582356 | 0.5428 | | 3.73E-06 | BLUPmrMLM | SUG |
| FD_19XX | AX-94553423 | 5B | 583913934 | -0.6550 | | 3.20E-28 | IIIVmrMLM | SIG |
| FD_19XX | AX-110428229 | 5B | 662031427 | -0.1474 | -4.5359 | 3.19E-18 | IIIVmrMLM | SIG |
| FD_19XX | AX-108951199 | 5B | 665188926 | 0.5740 | | 3.52E-06 | BLUPmrMLM | SUG |
| FD_19XX | AX-95629413 | 5B | 704725351 | 4.0695 | | 4.32E-07 | EMMAX | SUG |
| FD_19XX | AX-95629413 | 5B | 704725351 | -4.0535 | | 2.05E-07 | GEMMA | SUG |
| FD_19XX | AX-95629413 | 5B | 704725351 | 1.0868 | | 2.98E-64 | IIIVmrMLM | SIG |
| FD_19XX | AX-111688062 | 5B | 711711042 | 0.3566 | | 7.14E-05 | BLUPmrMLM | SUG |
| FD_19XX | AX-109360916 | 5D | 469286340 | 0.7942 | | 5.64E-39 | IIIVmrMLM | SIG |
| FD_19XX | AX-110093799 | 5D | 470390566 | 4.9441 | | 5.21E-08 | EMMAX | SIG |
| FD_19XX | AX-110093799 | 5D | 470390566 | -4.8850 | | 4.05E-08 | GEMMA | SIG |
| FD_19XX | AX-108831548 | 6A | 614449 | 0.2457 | -2.6497 | 3.77E-14 | IIIVmrMLM | SIG |
| FD_19XX | AX-108927103 | 6A | 46692482 | 0.2466 | 2.0519 | 1.13E-13 | IIIVmrMLM | SIG |
| FD_19XX | AX-110530579 | 6B | 29770998 | 0.4476 | | 4.46E-07 | BLUPmrMLM | SUG |
| FD_19XX | AX-111013117 | 6B | 117677111 | 0.8440 | | 3.76E-07 | BLUPmrMLM | SUG |
| FD_19XX | AX-110559773 | 6B | 137540357 | 2.7207 | | 1.63E-06 | EMMAX | SUG |
| FD_19XX | AX-110559773 | 6B | 137540357 | -2.7484 | | 5.34E-07 | GEMMA | SUG |
| FD_19XX | AX-110935567 | 6B | 141098799 | 0.3610 | | 1.54E-04 | BLUPmrMLM | SUG |
| FD_19XX | AX-110914313 | 6B | 156531878 | 0.3426 | -0.5197 | 1.59E-11 | IIIVmrMLM | SIG |
| FD_19XX | AX-94602984 | 6B | 462170032 | -0.1725 | -1.5873 | 1.87E-17 | IIIVmrMLM | SIG |
| FD_19XX | AX-94751075 | 6D | 60796504 | -0.7640 | | 4.41E-05 | BLUPmrMLM | SUG |
| FD_19XX | AX-94944360 | 6D | 465742743 | 0.5402 | | 6.01E-07 | BLUPmrMLM | SUG |
| FD_19XX | AX-95174425 | 6D | 469913454 | 0.7365 | | 7.62E-06 | BLUPmrMLM | SUG |
| FD_19XX | AX-111776821 | 7A | 451838535 | -0.2001 | -0.896 | 9.29E-08 | IIIVmrMLM | SIG |
| FD_19XX | AX-108795619 | 7A | 635052697 | -1.0684 | | 2.80E-05 | BLUPmrMLM | SUG |
| FD_19XX | AX-110433605 | 7A | 701367536 | 0.0460 | -0.9943 | 6.72E-08 | IIIVmrMLM | SIG |
| FD_19XX | AX-109597556 | 7B | 45874468 | 0.8648 | | 9.72E-07 | BLUPmrMLM | SUG |
| FD_19XX | AX-110626661 | 7B | 154812167 | -0.5846 | -0.8769 | 2.73E-24 | IIIVmrMLM | SIG |
| FD_19XX | AX-109473119 | 7B | 206007620 | -0.8339 | | 3.63E-06 | BLUPmrMLM | SUG |
| FD_19XX | AX-110390529 | 7B | 243278368 | 0.8862 | 0.5359 | 1.43E-42 | IIIVmrMLM | SIG |
| FD_19XX | AX-109490684 | 7B | 564023603 | -0.0353 | -1.374 | 5.26E-07 | IIIVmrMLM | SUG |
| FD_19XX | AX-111731206 | 7B | 611971167 | 0.6559 | 0.0082 | 2.14E-24 | IIIVmrMLM | SIG |
| FD_19XX | AX-110915122 | 7B | 628349874 | 0.1393 | -8.9386 | 9.80E-20 | IIIVmrMLM | SIG |
| FD_19XX | AX-110440910 | 7B | 739987627 | -0.8030 | | 2.23E-07 | BLUPmrMLM | SUG |
| FD_19XX | AX-110371659 | 7D | 6017853 | 4.5043 | | 1.12E-06 | EMMAX | SUG |
| FD_19XX | AX-110371659 | 7D | 6017853 | -4.5273 | | 1.15E-06 | GEMMA | SUG |
| FD_19XX | AX-110932523 | 7D | 138440518 | 10.9700 | | 3.93E-07 | EMMAX | SUG |
| FD_19XX | AX-111421702 | 7D | 553416685 | 0.1076 | -1.1732 | 1.09E-05 | IIIVmrMLM | SUG |

| | | | | | | | | |
|---|---|---|---|---|---|---|---|---|
| FD_19XX | AX-111525694 | 7D | 617115308 | 0.4131 | -0.971 | 1.43E-13 | IIIVmrMLM | SIG |
| FD_BLUP | AX-109035993 | 1A | 504787723 | -0.2800 | -0.3215 | 4.58E-24 | IIIVmrMLM | SIG |
| FD_BLUP | AX-109388229 | 1A | 557956782 | -0.2670 | | 7.89E-06 | BLUPmrMLM | SUG |
| FD_BLUP | AX-108780402 | 1A | 564570992 | -1.0812 | | 1.17E-06 | FarmCPU | SUG |
| FD_BLUP | AX-109332672 | 1A | 565292715 | 0.0754 | 0.6265 | 1.14E-11 | IIIVmrMLM | SIG |
| FD_BLUP | AX-109983107 | 1B | 86774625 | 0.4713 | | 1.96E-06 | BLUPmrMLM | SUG |
| FD_BLUP | AX-109634576 | 1B | 169711402 | 1.1578 | | 1.91E-06 | EMMAX | SUG |
| FD_BLUP | AX-109634576 | 1B | 169711402 | -1.1488 | | 1.64E-06 | GEMMA | SUG |
| FD_BLUP | AX-109634576 | 1B | 169711402 | -0.0156 | -0.6236 | 1.39E-21 | IIIVmrMLM | SIG |
| FD_BLUP | AX-109283005 | 1B | 384239973 | 0.0025 | 0.4688 | 1.65E-06 | IIIVmrMLM | SUG |
| FD_BLUP | AX-94411493 | 1B | 574463899 | -0.2428 | | 2.37E-19 | IIIVmrMLM | SIG |
| FD_BLUP | AX-94496522 | 1B | 622886614 | 1.0356 | 0.1184 | 7.96E-14 | IIIVmrMLM | SIG |
| FD_BLUP | AX-110967256 | 1B | 677617120 | -0.0437 | -0.3795 | 1.81E-11 | IIIVmrMLM | SIG |
| FD_BLUP | AX-110558601 | 1B | 687418455 | -0.2691 | | 1.76E-06 | BLUPmrMLM | SUG |
| FD_BLUP | AX-111194978 | 1B | 687530101 | 0.1571 | 0.1791 | 2.15E-08 | IIIVmrMLM | SIG |
| FD_BLUP | AX-111680033 | 2A | 162275567 | -0.2160 | | 8.12E-16 | IIIVmrMLM | SIG |
| FD_BLUP | AX-109924835 | 2A | 535801035 | 0.0585 | -0.4785 | 6.94E-17 | IIIVmrMLM | SIG |
| FD_BLUP | AX-94496472 | 2A | 729199454 | -0.2863 | | 7.47E-06 | BLUPmrMLM | SUG |
| FD_BLUP | AX-111001399 | 2B | 47273270 | -0.0784 | -0.3391 | 1.41E-05 | IIIVmrMLM | SUG |
| FD_BLUP | AX-94664524 | 2B | 117624330 | 0.2305 | 0.4438 | 1.64E-17 | IIIVmrMLM | SIG |
| FD_BLUP | AX-110046269 | 2B | 251778087 | -2.5029 | | 2.16E-07 | FarmCPU | SUG |
| FD_BLUP | AX-94491543 | 2B | 730998028 | 0.4527 | | 8.72E-07 | BLUPmrMLM | SUG |
| FD_BLUP | AX-110580387 | 2B | 760929932 | 0.3466 | | 2.87E-07 | BLUPmrMLM | SUG |
| FD_BLUP | AX-94697254 | 2D | 33710135 | -1.3352 | | 1.26E-06 | FarmCPU | SUG |
| FD_BLUP | AX-111080522 | 2D | 406260110 | 0.2196 | | 2.84E-16 | IIIVmrMLM | SIG |
| FD_BLUP | AX-109981832 | 2D | 487626614 | 0.3168 | | 1.02E-08 | BLUPmrMLM | SIG |
| FD_BLUP | AX-108756128 | 3A | 19988117 | 0.8519 | | 9.85E-07 | FarmCPU | SUG |
| FD_BLUP | AX-110719914 | 3A | 25559909 | 0.1359 | -1.782 | 1.51E-15 | IIIVmrMLM | SIG |
| FD_BLUP | AX-108841533 | 3A | 36476031 | 0.3022 | | 3.82E-07 | BLUPmrMLM | SUG |
| FD_BLUP | AX-109478207 | 3A | 395575072 | -0.1557 | -0.7726 | 5.32E-11 | IIIVmrMLM | SIG |
| FD_BLUP | AX-95204178 | 3A | 479782760 | 0.3320 | | 5.45E-08 | BLUPmrMLM | SIG |
| FD_BLUP | AX-108996295 | 3A | 562102052 | 0.1349 | 0.6764 | 4.71E-19 | IIIVmrMLM | SIG |
| FD_BLUP | AX-94522205 | 3A | 638010015 | -0.8025 | | 5.10E-07 | FarmCPU | SUG |
| FD_BLUP | AX-110401245 | 3A | 650425341 | 0.4116 | -1.1859 | 7.58E-48 | IIIVmrMLM | SIG |
| FD_BLUP | AX-111634763 | 3A | 654584208 | 0.2596 | | 1.99E-06 | BLUPmrMLM | SUG |
| FD_BLUP | AX-110491100 | 3A | 705251779 | -1.7134 | | 7.02E-10 | EMMAX | SIG |
| FD_BLUP | AX-110491100 | 3A | 705251779 | 1.7084 | | 2.83E-10 | GEMMA | SIG |
| FD_BLUP | AX-108941177 | 3A | 705259848 | 1.1620 | | 5.38E-12 | BLUPmrMLM | SIG |
| FD_BLUP | AX-108941177 | 3A | 705259848 | -1.6538 | | 2.97E-10 | EMMAX | SIG |
| FD_BLUP | AX-108941177 | 3A | 705259848 | 1.6476 | | 1.13E-10 | GEMMA | SIG |
| FD_BLUP | AX-111695263 | 3A | 705274739 | -1.6412 | | 1.39E-09 | EMMAX | SIG |
| FD_BLUP | AX-111695263 | 3A | 705274739 | 1.6426 | | 4.73E-10 | GEMMA | SIG |
| FD_BLUP | AX-110976174 | 3A | 705274782 | -3.1839 | | 1.09E-09 | EMMAX | SIG |
| FD_BLUP | AX-110976174 | 3A | 705274782 | 3.1570 | | 5.61E-10 | GEMMA | SIG |

| | | | | | | | | |
|---|---|---|---|---|---|---|---|---|
| FD_BLUP | AX-110484538 | 3A | 705296840 | -3.6973 | | 3.56E-10 | EMMAX | SIG |
| FD_BLUP | AX-110484538 | 3A | 705296840 | 3.6699 | | 1.53E-10 | GEMMA | SIG |
| FD_BLUP | AX-110526487 | 3A | 705298427 | 0.8777 | | 2.14E-06 | GEMMA | SUG |
| FD_BLUP | AX-110959604 | 3A | 743362286 | 0.0257 | 0.4356 | 5.76E-13 | IIIVmrMLM | SIG |
| FD_BLUP | AX-111197043 | 3B | 9088741 | 0.4173 | | 1.33E-06 | BLUPmrMLM | SUG |
| FD_BLUP | AX-109307433 | 3B | 61655711 | -0.2899 | | 1.31E-06 | BLUPmrMLM | SUG |
| FD_BLUP | AX-109850027 | 3B | 62742477 | 0.2717 | -0.0872 | 1.87E-16 | IIIVmrMLM | SIG |
| FD_BLUP | AX-111833531 | 3B | 332237107 | -1.5838 | | 2.99E-07 | FarmCPU | SUG |
| FD_BLUP | AX-109529747 | 3B | 466300144 | -0.7029 | -1.9018 | 8.39E-110 | IIIVmrMLM | SIG |
| FD_BLUP | AX-110427131 | 3B | 469648875 | -0.4273 | | 3.12E-10 | BLUPmrMLM | SIG |
| FD_BLUP | AX-110427131 | 3B | 469648875 | -0.4912 | 0.6946 | 3.72E-70 | IIIVmrMLM | SIG |
| FD_BLUP | AX-108771406 | 3B | 482774793 | -0.0097 | 0.9139 | 1.59E-09 | IIIVmrMLM | SIG |
| FD_BLUP | AX-108804485 | 3B | 749983803 | -0.6061 | -0.2397 | 4.49E-73 | IIIVmrMLM | SIG |
| FD_BLUP | AX-109514530 | 3D | 62562233 | 0.0106 | -1.2162 | 5.24E-06 | IIIVmrMLM | SUG |
| FD_BLUP | AX-111026826 | 3D | 560126633 | 0.4489 | | 5.11E-08 | BLUPmrMLM | SIG |
| FD_BLUP | AX-109576258 | 4A | 6642195 | -0.2962 | 0.2055 | 5.94E-26 | IIIVmrMLM | SIG |
| FD_BLUP | AX-108946086 | 4A | 43236449 | 0.4358 | | 1.23E-10 | BLUPmrMLM | SIG |
| FD_BLUP | AX-111256892 | 4A | 342811372 | -0.4355 | -2.4232 | 7.89E-66 | IIIVmrMLM | SIG |
| FD_BLUP | AX-108755559 | 4A | 455270539 | 0.0696 | -0.7819 | 9.99E-08 | IIIVmrMLM | SIG |
| FD_BLUP | AX-109424652 | 4A | 615937289 | 0.0172 | -1.9104 | 2.11E-13 | IIIVmrMLM | SIG |
| FD_BLUP | AX-109816200 | 4A | 649204800 | -0.2224 | 0.6755 | 3.57E-16 | IIIVmrMLM | SIG |
| FD_BLUP | AX-89438414 | 4B | 598261950 | 0.1924 | 0.2806 | 2.56E-12 | IIIVmrMLM | SIG |
| FD_BLUP | AX-110905638 | 4B | 649656041 | 0.2013 | -0.6282 | 1.54E-14 | IIIVmrMLM | SIG |
| FD_BLUP | AX-109466196 | 5A | 528379998 | 0.2600 | 1.2169 | 3.57E-23 | IIIVmrMLM | SIG |
| FD_BLUP | AX-111641807 | 5A | 538948758 | 0.1660 | -1.159 | 2.84E-17 | IIIVmrMLM | SIG |
| FD_BLUP | AX-110513698 | 5A | 549378654 | 0.3433 | | 1.42E-06 | BLUPmrMLM | SUG |
| FD_BLUP | AX-111235500 | 5A | 556150580 | 0.5896 | | 2.05E-13 | BLUPmrMLM | SIG |
| FD_BLUP | AX-111235500 | 5A | 556150580 | -0.2324 | -0.2944 | 3.25E-17 | IIIVmrMLM | SIG |
| FD_BLUP | AX-108836029 | 5A | 559135112 | -0.0584 | -1.2437 | 5.44E-08 | IIIVmrMLM | SIG |
| FD_BLUP | AX-111022811 | 5A | 595493895 | 0.3802 | | 6.79E-07 | BLUPmrMLM | SUG |
| FD_BLUP | AX-94464140 | 5B | 69644430 | 3.9965 | | 9.62E-07 | EMMAX | SUG |
| FD_BLUP | AX-111657935 | 5B | 382332802 | -0.1448 | -0.2976 | 5.05E-09 | IIIVmrMLM | SIG |
| FD_BLUP | AX-108831460 | 5B | 527056847 | 0.0887 | -0.3947 | 2.54E-13 | IIIVmrMLM | SIG |
| FD_BLUP | AX-110538089 | 5B | 539858157 | 0.1699 | -0.2641 | 1.27E-10 | IIIVmrMLM | SIG |
| FD_BLUP | AX-109821989 | 5B | 565424439 | 0.2476 | -1.443 | 2.66E-30 | IIIVmrMLM | SIG |
| FD_BLUP | AX-109276437 | 5B | 653704397 | -0.1536 | -0.4161 | 1.35E-19 | IIIVmrMLM | SIG |
| FD_BLUP | AX-94509671 | 5D | 331402475 | -0.3549 | | 1.80E-05 | BLUPmrMLM | SUG |
| FD_BLUP | AX-110035093 | 5D | 464129291 | -0.2015 | -0.2007 | 4.57E-13 | IIIVmrMLM | SIG |
| FD_BLUP | AX-111175690 | 5D | 464415388 | 0.8397 | | 7.72E-07 | EMMAX | SUG |
| FD_BLUP | AX-111175690 | 5D | 464415388 | -0.8389 | | 5.52E-07 | GEMMA | SUG |
| FD_BLUP | AX-110958036 | 5D | 469179441 | 0.2331 | | 2.22E-05 | BLUPmrMLM | SUG |
| FD_BLUP | AX-109360916 | 5D | 469286340 | 2.1977 | | 1.08E-09 | EMMAX | SIG |
| FD_BLUP | AX-109360916 | 5D | 469286340 | -2.1988 | | 5.88E-10 | GEMMA | SIG |

| | | | | | | | | |
|---|---|---|---|---|---|---|---|---|
| FD_BLUP | AX-109360916 | 5D | 469286340 | 0.6981 | | 3.57E-105 | IIIVmrMLM | SIG |
| FD_BLUP | AX-89336793 | 5D | 489775906 | -1.1293 | | 2.10E-06 | GEMMA | SUG |
| FD_BLUP | AX-111617101 | 6A | 2413580 | 0.1811 | | 2.88E-05 | BLUPmrMLM | SUG |
| FD_BLUP | AX-110980257 | 6A | 254597028 | -0.1821 | -0.0831 | 2.87E-09 | IIIVmrMLM | SIG |
| FD_BLUP | AX-109384863 | 6A | 611924505 | -0.1703 | -0.4611 | 7.60E-14 | IIIVmrMLM | SIG |
| FD_BLUP | AX-110981223 | 6A | 614369572 | -0.0759 | -0.2221 | 1.06E-06 | IIIVmrMLM | SUG |
| FD_BLUP | AX-109523386 | 6B | 54662589 | -0.3270 | | 6.70E-06 | BLUPmrMLM | SUG |
| FD_BLUP | AX-110914313 | 6B | 156531878 | 0.2356 | -0.1928 | 8.92E-16 | IIIVmrMLM | SIG |
| FD_BLUP | AX-109391962 | 6B | 339268873 | -1.6813 | | 6.27E-07 | FarmCPU | SUG |
| FD_BLUP | AX-110952052 | 6D | 7030785 | -0.1721 | -0.1666 | 7.96E-10 | IIIVmrMLM | SIG |
| FD_BLUP | AX-95074292 | 6D | 404222344 | -0.5015 | | 7.00E-07 | BLUPmrMLM | SUG |
| FD_BLUP | AX-111872610 | 6D | 457851551 | -1.2724 | | 2.28E-06 | FarmCPU | SUG |
| FD_BLUP | AX-94612915 | 6D | 464704396 | -0.3518 | | 5.53E-06 | BLUPmrMLM | SUG |
| FD_BLUP | AX-109985214 | 6D | 465508851 | -0.3248 | 0.0728 | 8.75E-20 | IIIVmrMLM | SIG |
| FD_BLUP | AX-109864970 | 7A | 47458495 | -0.2505 | 0.2003 | 4.72E-19 | IIIVmrMLM | SIG |
| FD_BLUP | AX-109308202 | 7A | 48841017 | -0.2866 | | 2.22E-05 | BLUPmrMLM | SUG |
| FD_BLUP | AX-111004071 | 7A | 151956510 | -1.2018 | | 1.74E-06 | FarmCPU | SUG |
| FD_BLUP | AX-111453256 | 7A | 163614277 | -0.1283 | 1.114 | 3.06E-11 | IIIVmrMLM | SIG |
| FD_BLUP | AX-111776821 | 7A | 451838535 | -0.0320 | -0.4469 | 4.46E-06 | IIIVmrMLM | SUG |
| FD_BLUP | AX-109473440 | 7A | 659838893 | 0.1198 | -1.3806 | 1.12E-11 | IIIVmrMLM | SIG |
| FD_BLUP | AX-111018476 | 7A | 668982701 | 0.3838 | | 3.66E-42 | IIIVmrMLM | SIG |
| FD_BLUP | AX-109869065 | 7B | 49175029 | 0.0375 | -0.4979 | 1.39E-08 | IIIVmrMLM | SIG |
| FD_BLUP | AX-111494449 | 7B | 637962621 | -0.3256 | | 6.60E-08 | BLUPmrMLM | SIG |
| FD_BLUP | AX-110995137 | 7B | 740045196 | -0.2661 | | 8.33E-06 | BLUPmrMLM | SUG |
| FD_BLUP | AX-110080714 | 7D | 40595686 | -0.2269 | -0.1287 | 9.16E-15 | IIIVmrMLM | SIG |
| FD_BLUP | AX-110576947 | 7D | 65436205 | 0.7781 | | 1.01E-06 | EMMAX | SUG |
| FD_BLUP | AX-110576947 | 7D | 65436205 | -0.7755 | | 6.19E-07 | GEMMA | SUG |
| FD_BLUP | AX-111061288 | 7D | 66537922 | -0.2011 | 0.009 | 7.54E-12 | IIIVmrMLM | SIG |
| FD_BLUP | AX-109862696 | 7D | 223131934 | -0.3346 | 0.1531 | 1.32E-31 | IIIVmrMLM | SIG |
| FD_BLUP | AX-110875275 | 7D | 257526220 | 0.2212 | | 4.29E-05 | BLUPmrMLM | SUG |
| FD_BLUP | AX-109870041 | 7D | 581610668 | 0.0083 | 1.0621 | 4.06E-08 | IIIVmrMLM | SIG |

FD: flowering date.

**Supplementary Table S21**. All QTNs identified as associated with heading date in wheat.

| Trait | Marker | Chr | Position | Add.effect | Dom.effect | P-value | Method | Sig. |
|---|---|---|---|---|---|---|---|---|
| HD_17AY | AX-111449556 | 1A | 461994223 | 0.1388 | 1.3022 | 1.97E-17 | IIIVmrMLM | SIG |
| HD_17AY | AX-111053694 | 1A | 500990019 | -0.3552 | | 1.22E-05 | BLUPmrMLM | SUG |
| HD_17AY | AX-94849470 | 1A | 551308340 | -0.2714 | | 6.14E-05 | BLUPmrMLM | SUG |
| HD_17AY | AX-111700863 | 1A | 557968013 | -0.3528 | | 2.22E-06 | BLUPmrMLM | SUG |
| HD_17AY | AX-109360506 | 1A | 586279275 | 0.0164 | -2.2495 | 2.27E-12 | IIIVmrMLM | SIG |
| HD_17AY | AX-94447402 | 1B | 637825918 | -0.1672 | -0.4106 | 2.64E-05 | IIIVmrMLM | SUG |
| HD_17AY | AX-108750878 | 1D | 14147616 | -0.0453 | 1.2518 | 1.08E-29 | IIIVmrMLM | SIG |
| HD_17AY | AX-108898996 | 1D | 454766541 | 0.0833 | 1.1612 | 4.52E-09 | IIIVmrMLM | SIG |
| HD_17AY | AX-111810888 | 2A | 40103967 | 0.1949 | | 1.70E-07 | IIIVmrMLM | SUG |
| HD_17AY | AX-110731402 | 2A | 308034792 | -0.1732 | -0.92 | 1.85E-28 | IIIVmrMLM | SIG |
| HD_17AY | AX-108912390 | 2A | 428552045 | 0.1541 | 0.4432 | 3.25E-08 | IIIVmrMLM | SIG |
| HD_17AY | AX-109288544 | 2B | 150899516 | 0.2415 | | 1.02E-04 | BLUPmrMLM | SUG |
| HD_17AY | AX-108864407 | 2B | 242535487 | 0.1806 | 0.7385 | 1.71E-13 | IIIVmrMLM | SIG |
| HD_17AY | AX-110993119 | 2B | 290868022 | 0.0716 | 0.8408 | 4.75E-20 | IIIVmrMLM | SIG |
| HD_17AY | AX-108745839 | 2B | 448783756 | 0.0887 | -2.1157 | 2.87E-13 | IIIVmrMLM | SIG |
| HD_17AY | AX-111537086 | 2B | 601183765 | 0.1862 | | 5.70E-07 | IIIVmrMLM | SUG |
| HD_17AY | AX-111468726 | 2B | 657791701 | -0.5063 | | 2.55E-08 | BLUPmrMLM | SIG |
| HD_17AY | AX-94932983 | 2B | 658996872 | 0.2587 | | 5.90E-12 | IIIVmrMLM | SIG |
| HD_17AY | AX-111585974 | 2B | 765900766 | -0.3786 | -0.0253 | 9.75E-11 | IIIVmrMLM | SIG |
| HD_17AY | AX-109415150 | 2B | 794428137 | 0.1248 | -0.483 | 3.16E-05 | IIIVmrMLM | SUG |
| HD_17AY | AX-110922927 | 2D | 405252279 | 0.3835 | -0.7954 | 4.11E-23 | IIIVmrMLM | SIG |
| HD_17AY | AX-108976454 | 2D | 408537887 | -0.4301 | | 5.08E-07 | BLUPmrMLM | SUG |
| HD_17AY | AX-95240693 | 3A | 10302316 | 0.0263 | -0.5194 | 1.08E-11 | IIIVmrMLM | SIG |
| HD_17AY | AX-111659712 | 3A | 134672225 | -0.3001 | | 1.30E-04 | BLUPmrMLM | SUG |
| HD_17AY | AX-110381099 | 3A | 428570654 | 0.7274 | | 1.53E-07 | BLUPmrMLM | SUG |
| HD_17AY | AX-110030954 | 3A | 540667176 | -0.3374 | | 5.73E-05 | BLUPmrMLM | SUG |
| HD_17AY | AX-111470092 | 3A | 604840137 | 0.3782 | -0.1025 | 8.64E-19 | IIIVmrMLM | SIG |
| HD_17AY | AX-110406923 | 3A | 652775501 | -0.2626 | | 1.44E-04 | BLUPmrMLM | SUG |
| HD_17AY | AX-109864277 | 3A | 664732775 | 0.0335 | -0.6843 | 2.21E-06 | IIIVmrMLM | SUG |
| HD_17AY | AX-108777982 | 3A | 684756427 | 0.2897 | -0.2264 | 6.19E-13 | IIIVmrMLM | SIG |
| HD_17AY | AX-110491100 | 3A | 705251779 | -1.9527 | | 7.33E-08 | EMMAX | SIG |
| HD_17AY | AX-110491100 | 3A | 705251779 | 1.9636 | | 1.65E-08 | GEMMA | SIG |
| HD_17AY | AX-108941177 | 3A | 705259848 | -1.8442 | | 6.44E-08 | EMMAX | SIG |
| HD_17AY | AX-108941177 | 3A | 705259848 | 1.8438 | | 1.70E-08 | GEMMA | SIG |
| HD_17AY | AX-111695263 | 3A | 705274739 | -1.8973 | | 8.74E-08 | EMMAX | SIG |
| HD_17AY | AX-111695263 | 3A | 705274739 | 1.9056 | | 2.05E-08 | GEMMA | SIG |
| HD_17AY | AX-110976174 | 3A | 705274782 | -3.5966 | | 1.02E-07 | EMMAX | SIG |
| HD_17AY | AX-110976174 | 3A | 705274782 | 3.5945 | | 2.92E-08 | GEMMA | SIG |
| HD_17AY | AX-110484538 | 3A | 705296840 | -4.1005 | | 8.31E-08 | EMMAX | SIG |
| HD_17AY | AX-110484538 | 3A | 705296840 | 4.0952 | | 2.28E-08 | GEMMA | SIG |
| HD_17AY | AX-110390871 | 3B | 7252439 | 0.5207 | | 2.04E-07 | BLUPmrMLM | SUG |

| | | | | | | | | |
|---|---|---|---|---|---|---|---|---|
| HD_17AY | AX-110390871 | 3B | 7252439 | -0.7500 | | 3.17E-07 | EMMAX | SUG |
| HD_17AY | AX-110390871 | 3B | 7252439 | 0.7503 | | 1.78E-07 | GEMMA | SUG |
| HD_17AY | AX-111081536 | 3B | 8806517 | -0.2787 | 0.2713 | 1.69E-12 | IIIVmrMLM | SIG |
| HD_17AY | AX-110989677 | 3B | 61104501 | -0.3170 | -0.3661 | 1.37E-16 | IIIVmrMLM | SIG |
| HD_17AY | AX-108818769 | 3B | 61657123 | 0.3000 | -0.979 | 3.31E-28 | IIIVmrMLM | SIG |
| HD_17AY | AX-109967710 | 3B | 469873446 | 0.4528 | | 1.00E-06 | BLUPmrMLM | SUG |
| HD_17AY | AX-109967710 | 3B | 469873446 | -0.3646 | 0.0439 | 2.52E-18 | IIIVmrMLM | SIG |
| HD_17AY | AX-111543008 | 3B | 727794913 | 0.3183 | 0.5457 | 6.37E-17 | IIIVmrMLM | SIG |
| HD_17AY | AX-108973246 | 3B | 779641300 | -0.1001 | 1.0396 | 1.44E-11 | IIIVmrMLM | SIG |
| HD_17AY | AX-95108623 | 3D | 7028397 | -0.4805 | | 2.93E-05 | BLUPmrMLM | SUG |
| HD_17AY | AX-95209443 | 3D | 45938070 | -0.2143 | | 9.66E-09 | IIIVmrMLM | SIG |
| HD_17AY | AX-94436939 | 3D | 461019453 | 0.4232 | | 9.41E-28 | IIIVmrMLM | SIG |
| HD_17AY | AX-109519108 | 4A | 86992797 | -1.0204 | | 2.18E-06 | FarmCPU | SUG |
| HD_17AY | AX-109033918 | 4A | 388253996 | 0.3311 | | 4.84E-05 | BLUPmrMLM | SUG |
| HD_17AY | AX-111190872 | 4A | 540416431 | -0.3908 | | 3.24E-24 | IIIVmrMLM | SIG |
| HD_17AY | AX-111048595 | 4B | 37529724 | 0.4009 | | 8.21E-05 | BLUPmrMLM | SUG |
| HD_17AY | AX-108838330 | 4B | 465981268 | 0.0724 | -0.7154 | 9.74E-09 | IIIVmrMLM | SIG |
| HD_17AY | AX-95158490 | 4B | 592986558 | -0.7503 | | 5.47E-10 | BLUPmrMLM | SIG |
| HD_17AY | AX-94433890 | 4D | 142766146 | 0.2109 | | 1.64E-08 | IIIVmrMLM | SIG |
| HD_17AY | AX-110599156 | 5A | 82876910 | -0.3411 | | 2.19E-05 | BLUPmrMLM | SUG |
| HD_17AY | AX-110169212 | 5A | 455055483 | 0.0374 | -1.0899 | 6.75E-22 | IIIVmrMLM | SIG |
| HD_17AY | AX-111689418 | 5A | 540026591 | -0.0799 | 2.1397 | 1.18E-17 | IIIVmrMLM | SIG |
| HD_17AY | AX-110033906 | 5B | 32962770 | 0.2676 | -0.0725 | 2.34E-11 | IIIVmrMLM | SIG |
| HD_17AY | AX-89465333 | 5B | 41285358 | 0.0593 | 0.8315 | 8.28E-13 | IIIVmrMLM | SIG |
| HD_17AY | AX-111682490 | 5B | 59932775 | 1.1438 | | 1.15E-06 | FarmCPU | SUG |
| HD_17AY | AX-108822938 | 5B | 66339657 | 1.2996 | | 1.87E-06 | FarmCPU | SUG |
| HD_17AY | AX-94638868 | 5B | 103569861 | 0.3271 | | 7.69E-18 | IIIVmrMLM | SIG |
| HD_17AY | AX-111556536 | 5B | 529880097 | 0.1759 | -1.1797 | 2.16E-12 | IIIVmrMLM | SIG |
| HD_17AY | AX-111161208 | 5B | 529989075 | -0.1449 | 4.4739 | 3.90E-09 | IIIVmrMLM | SIG |
| HD_17AY | AX-108938523 | 5B | 536846649 | -0.4028 | 0.102 | 2.06E-12 | IIIVmrMLM | SIG |
| HD_17AY | AX-109308790 | 5B | 596995779 | 0.0014 | -1.3708 | 3.71E-05 | IIIVmrMLM | SUG |
| HD_17AY | AX-94582317 | 5D | 29585874 | -0.0869 | 1.0344 | 1.30E-10 | IIIVmrMLM | SIG |
| HD_17AY | AX-95091790 | 5D | 562341122 | -0.3249 | | 7.62E-05 | BLUPmrMLM | SUG |
| HD_17AY | AX-109526267 | 6A | 22813165 | 0.0782 | 1.0317 | 2.99E-17 | IIIVmrMLM | SIG |
| HD_17AY | AX-94497617 | 6B | 32328193 | -0.3898 | | 5.11E-07 | BLUPmrMLM | SUG |
| HD_17AY | AX-110360699 | 6B | 201254117 | -0.5923 | | 9.94E-08 | BLUPmrMLM | SIG |
| HD_17AY | AX-108923678 | 6B | 459374936 | -0.4511 | | 4.22E-07 | BLUPmrMLM | SUG |
| HD_17AY | AX-108950523 | 6B | 509915792 | 0.0436 | 0.7211 | 1.99E-06 | IIIVmrMLM | SUG |
| HD_17AY | AX-108779531 | 6B | 686383283 | -0.4033 | | 5.14E-05 | BLUPmrMLM | SUG |
| HD_17AY | AX-108901786 | 6B | 712475275 | -0.0192 | -1.2164 | 1.51E-09 | IIIVmrMLM | SIG |
| HD_17AY | AX-110390209 | 6D | 459060764 | -0.0982 | 1.6603 | 1.30E-08 | IIIVmrMLM | SIG |
| HD_17AY | AX-109833655 | 7A | 10110972 | 0.5540 | | 1.31E-07 | BLUPmrMLM | SUG |
| HD_17AY | AX-110977044 | 7A | 12725608 | -0.2695 | -0.9503 | 1.17E-12 | IIIVmrMLM | SIG |
| HD_17AY | AX-108940753 | 7A | 541337103 | -0.0276 | 0.5122 | 1.96E-07 | IIIVmrMLM | SUG |

| | | | | | | | | |
|---|---|---|---|---|---|---|---|---|
| HD_17AY | AX-111184903 | 7A | 652293442 | 0.2696 | -0.2178 | 8.34E-12 | IIIVmrMLM | SIG |
| HD_17AY | AX-109539867 | 7B | 52614647 | -0.0081 | 1.0358 | 6.59E-11 | IIIVmrMLM | SIG |
| HD_17AY | AX-110395012 | 7B | 66499791 | -0.0010 | -0.4633 | 1.10E-07 | IIIVmrMLM | SIG |
| HD_17AY | AX-89516048 | 7B | 145075160 | -0.3144 | 0.7176 | 3.87E-16 | IIIVmrMLM | SIG |
| HD_17AY | AX-111184541 | 7D | 66833728 | -1.0196 | | 1.28E-06 | GEMMA | SUG |
| HD_17AY | AX-111843581 | 7D | 67448053 | -1.0050 | | 2.19E-06 | GEMMA | SUG |
| HD_17AY | AX-109189716 | 7D | 202544957 | -0.2853 | -0.1878 | 5.69E-13 | IIIVmrMLM | SIG |
| HD_17AY | AX-109147840 | 7D | 529898254 | -0.1722 | -0.5931 | 2.38E-19 | IIIVmrMLM | SIG |
| HD_17AY | AX-110381734 | 7D | 556313472 | 0.2606 | | 2.29E-05 | BLUPmrMLM | SUG |
| HD_17ZMD | AX-109838328 | 1A | 5495926 | -0.5634 | -0.103 | 6.98E-21 | IIIVmrMLM | SIG |
| HD_17ZMD | AX-110072329 | 1A | 420379515 | 0.1401 | 3.0555 | 1.68E-10 | IIIVmrMLM | SIG |
| HD_17ZMD | AX-110933445 | 1A | 507745252 | -0.5618 | | 3.44E-05 | BLUPmrMLM | SUG |
| HD_17ZMD | AX-109414027 | 1A | 558500274 | 0.2177 | -1.1623 | 8.99E-15 | IIIVmrMLM | SIG |
| HD_17ZMD | AX-110079714 | 1B | 11185541 | 0.0922 | -1.24 | 1.09E-32 | IIIVmrMLM | SIG |
| HD_17ZMD | AX-109011761 | 1B | 310367970 | -0.0423 | -4.3822 | 1.37E-09 | IIIVmrMLM | SIG |
| HD_17ZMD | AX-109424410 | 1B | 420585198 | -0.5695 | 0.2534 | 5.90E-19 | IIIVmrMLM | SIG |
| HD_17ZMD | AX-111705530 | 1B | 465975682 | 0.5723 | | 4.73E-06 | BLUPmrMLM | SUG |
| HD_17ZMD | AX-111705530 | 1B | 465975682 | -0.4437 | -0.0067 | 1.99E-11 | IIIVmrMLM | SIG |
| HD_17ZMD | AX-110496206 | 1B | 566405225 | -0.3095 | 0.086 | 3.43E-07 | IIIVmrMLM | SUG |
| HD_17ZMD | AX-109343976 | 1B | 662796945 | -0.6993 | | 5.01E-08 | BLUPmrMLM | SIG |
| HD_17ZMD | AX-109343976 | 1B | 662796945 | -0.5208 | -0.1176 | 5.22E-11 | IIIVmrMLM | SIG |
| HD_17ZMD | AX-111581326 | 1D | 440661502 | 0.5544 | | 9.79E-22 | IIIVmrMLM | SIG |
| HD_17ZMD | AX-109239005 | 1D | 465588213 | -0.6113 | | 1.07E-25 | IIIVmrMLM | SIG |
| HD_17ZMD | AX-109142090 | 1D | 476910568 | -0.9505 | | 1.07E-09 | BLUPmrMLM | SIG |
| HD_17ZMD | AX-110062614 | 2A | 252390785 | -0.1751 | 3.0609 | 3.09E-11 | IIIVmrMLM | SIG |
| HD_17ZMD | AX-111728180 | 2B | 407912921 | -0.0832 | 2.5272 | 8.85E-08 | IIIVmrMLM | SIG |
| HD_17ZMD | AX-108881724 | 2B | 580862345 | -0.1887 | -0.7698 | 3.99E-06 | IIIVmrMLM | SUG |
| HD_17ZMD | AX-110978145 | 2B | 697974662 | 0.7513 | | 6.72E-10 | BLUPmrMLM | SIG |
| HD_17ZMD | AX-109328386 | 2B | 748722837 | -0.4836 | -1.8839 | 1.43E-30 | IIIVmrMLM | SIG |
| HD_17ZMD | AX-111956072 | 2D | 34428838 | -0.3716 | -0.7894 | 1.37E-10 | IIIVmrMLM | SIG |
| HD_17ZMD | AX-108841533 | 3A | 36476031 | -0.7514 | -0.0367 | 1.13E-23 | IIIVmrMLM | SIG |
| HD_17ZMD | AX-109936048 | 3A | 162357149 | 0.3085 | -1.3251 | 1.62E-12 | IIIVmrMLM | SIG |
| HD_17ZMD | AX-109838601 | 3A | 500970769 | -0.6614 | | 2.27E-06 | BLUPmrMLM | SUG |
| HD_17ZMD | AX-109099109 | 3A | 525564414 | -0.4173 | -1.4131 | 2.03E-28 | IIIVmrMLM | SIG |
| HD_17ZMD | AX-111634763 | 3A | 654584208 | 0.3974 | | 7.45E-05 | BLUPmrMLM | SUG |
| HD_17ZMD | AX-110491100 | 3A | 705251779 | 2.6755 | | 3.85E-08 | GEMMA | SIG |
| HD_17ZMD | AX-108941177 | 3A | 705259848 | 2.5669 | | 2.09E-08 | GEMMA | SIG |
| HD_17ZMD | AX-111695263 | 3A | 705274739 | 2.3824 | | 4.90E-07 | GEMMA | SUG |
| HD_17ZMD | AX-110976174 | 3A | 705274782 | 4.8379 | | 1.03E-07 | GEMMA | SIG |
| HD_17ZMD | AX-110484538 | 3A | 705296840 | 6.3330 | | 7.79E-10 | GEMMA | SIG |
| HD_17ZMD | AX-94718872 | 3A | 709375804 | -0.2505 | | 8.06E-06 | IIIVmrMLM | SUG |
| HD_17ZMD | AX-94654843 | 3B | 459445445 | -1.2152 | | 1.19E-05 | BLUPmrMLM | SUG |
| HD_17ZMD | AX-110208552 | 3D | 7087589 | -0.6834 | | 4.54E-31 | IIIVmrMLM | SIG |
| HD_17ZMD | AX-110331371 | 3D | 281100831 | -0.4531 | | 2.25E-15 | IIIVmrMLM | SIG |

| | | | | | | | | |
|---|---|---|---|---|---|---|---|---|
| HD_17ZMD | AX-94416014 | 3D | 507042732 | -1.6084 | -1.0329 | 1.61E-98 | IIIVmrMLM | SIG |
| HD_17ZMD | AX-109270750 | 4A | 29026205 | -0.4090 | | 1.54E-04 | BLUPmrMLM | SUG |
| HD_17ZMD | AX-110667425 | 4A | 76899363 | 0.5179 | | 2.48E-19 | IIIVmrMLM | SIG |
| HD_17ZMD | AX-110095653 | 4A | 617934669 | -0.7892 | | 2.14E-07 | BLUPmrMLM | SUG |
| HD_17ZMD | AX-108795868 | 4A | 673679914 | 0.1013 | -0.7169 | 5.97E-09 | IIIVmrMLM | SIG |
| HD_17ZMD | AX-110492242 | 4A | 717307287 | 0.0767 | 0.7513 | 4.77E-14 | IIIVmrMLM | SIG |
| HD_17ZMD | AX-108879237 | 4A | 728354289 | -0.0565 | 1.5526 | 1.12E-05 | IIIVmrMLM | SUG |
| HD_17ZMD | AX-89414036 | 4B | 13419503 | -0.5179 | | 2.58E-06 | BLUPmrMLM | SUG |
| HD_17ZMD | AX-110040941 | 4B | 496161724 | -0.0057 | -0.4851 | 2.57E-08 | IIIVmrMLM | SIG |
| HD_17ZMD | AX-108805422 | 4B | 637580204 | -0.5640 | | 3.73E-06 | BLUPmrMLM | SUG |
| HD_17ZMD | AX-109491309 | 4D | 279848226 | 0.1725 | 1.9873 | 2.69E-10 | IIIVmrMLM | SIG |
| HD_17ZMD | AX-109208291 | 4D | 471559831 | -0.2933 | | 2.56E-05 | BLUPmrMLM | SUG |
| HD_17ZMD | AX-110021183 | 4D | 509176360 | -0.1617 | -1.4146 | 1.81E-09 | IIIVmrMLM | SIG |
| HD_17ZMD | AX-108762405 | 5A | 330504308 | 0.3675 | | 1.83E-04 | BLUPmrMLM | SUG |
| HD_17ZMD | AX-111589398 | 5A | 698638188 | 0.8180 | | 2.27E-07 | BLUPmrMLM | SUG |
| HD_17ZMD | AX-110957570 | 5B | 516094952 | -0.6951 | | 6.99E-06 | BLUPmrMLM | SUG |
| HD_17ZMD | AX-110957570 | 5B | 516094952 | 0.6330 | -0.2205 | 1.74E-23 | IIIVmrMLM | SIG |
| HD_17ZMD | AX-111103826 | 5B | 638671923 | 0.0941 | 0.954 | 2.27E-20 | IIIVmrMLM | SIG |
| HD_17ZMD | AX-89752714 | 5B | 669489097 | 0.3411 | | 4.21E-05 | BLUPmrMLM | SUG |
| HD_17ZMD | AX-95008036 | 5B | 691860601 | -0.4083 | | 4.19E-05 | BLUPmrMLM | SUG |
| HD_17ZMD | AX-110044748 | 5D | 464108412 | -2.8135 | | 9.15E-07 | GEMMA | SUG |
| HD_17ZMD | AX-110035093 | 5D | 464129291 | 0.7259 | | 6.84E-07 | BLUPmrMLM | SUG |
| HD_17ZMD | AX-111175690 | 5D | 464415388 | -1.5090 | | 2.40E-07 | GEMMA | SUG |
| HD_17ZMD | AX-109360916 | 5D | 469286340 | -3.6903 | | 3.27E-09 | GEMMA | SIG |
| HD_17ZMD | AX-109360916 | 5D | 469286340 | 0.5862 | | 6.45E-24 | IIIVmrMLM | SIG |
| HD_17ZMD | AX-111660163 | 5D | 475817480 | 0.3543 | | 5.44E-05 | BLUPmrMLM | SUG |
| HD_17ZMD | AX-109458488 | 6A | 17550625 | 0.4541 | -0.4264 | 7.87E-14 | IIIVmrMLM | SIG |
| HD_17ZMD | AX-110932982 | 6A | 53073622 | 0.8645 | | 6.01E-46 | IIIVmrMLM | SIG |
| HD_17ZMD | AX-110406728 | 6A | 558931174 | -0.1463 | -1.0587 | 1.47E-09 | IIIVmrMLM | SIG |
| HD_17ZMD | AX-109330846 | 6B | 32946759 | 0.4610 | | 1.70E-06 | BLUPmrMLM | SUG |
| HD_17ZMD | AX-109483442 | 6B | 54219152 | -0.4445 | | 5.18E-05 | BLUPmrMLM | SUG |
| HD_17ZMD | AX-109312707 | 6B | 55643674 | 0.0064 | -1.9368 | 4.97E-08 | IIIVmrMLM | SIG |
| HD_17ZMD | AX-111628700 | 6B | 151148378 | -0.2254 | 0.7197 | 6.86E-07 | IIIVmrMLM | SUG |
| HD_17ZMD | AX-111485241 | 6B | 193643528 | -0.0237 | 2.364 | 8.28E-08 | IIIVmrMLM | SIG |
| HD_17ZMD | AX-109874207 | 6B | 377511822 | 0.2085 | -1.0344 | 7.30E-12 | IIIVmrMLM | SIG |
| HD_17ZMD | AX-89761401 | 6B | 713498425 | -0.0763 | -1.1053 | 1.49E-11 | IIIVmrMLM | SIG |
| HD_17ZMD | AX-94612915 | 6D | 464704396 | -0.4861 | | 7.34E-05 | BLUPmrMLM | SUG |
| HD_17ZMD | AX-111068393 | 6D | 464943134 | 0.9454 | 1.6987 | 2.07E-78 | IIIVmrMLM | SIG |
| HD_17ZMD | AX-109414162 | 6D | 470046015 | 0.3785 | 1.8175 | 1.78E-15 | IIIVmrMLM | SIG |
| HD_17ZMD | AX-89372694 | 6D | 471905419 | -0.0268 | -1.5219 | 1.22E-06 | IIIVmrMLM | SUG |
| HD_17ZMD | AX-109864970 | 7A | 47458495 | -0.4677 | 0.5431 | 1.98E-15 | IIIVmrMLM | SIG |
| HD_17ZMD | AX-111026347 | 7A | 75211416 | 0.5679 | | 7.38E-06 | BLUPmrMLM | SUG |
| HD_17ZMD | AX-111490522 | 7A | 707103478 | -2.5297 | | 2.21E-06 | FarmCPU | SUG |
| HD_17ZMD | AX-109899262 | 7A | 714598773 | 0.3374 | 0.785 | 1.60E-09 | IIIVmrMLM | SIG |

| | | | | | | | | |
|---|---|---|---|---|---|---|---|---|
| HD_17ZMD | AX-110396753 | 7B | 43207274 | 0.3180 | | 2.42E-05 | BLUPmrMLM | SUG |
| HD_17ZMD | AX-110378224 | 7B | 135538546 | 0.5762 | | 9.20E-06 | BLUPmrMLM | SUG |
| HD_17ZMD | AX-109975151 | 7B | 603653958 | -0.4054 | | 9.45E-05 | BLUPmrMLM | SUG |
| HD_17ZMD | AX-109962190 | 7B | 664039785 | 0.0167 | -0.9408 | 3.72E-13 | IIIVmrMLM | SIG |
| HD_17ZMD | AX-108822678 | 7D | 75079850 | 0.2654 | 4.0074 | 1.24E-12 | IIIVmrMLM | SIG |
| HD_17ZMD | AX-111000058 | 7D | 529064922 | 0.6168 | | 1.34E-04 | BLUPmrMLM | SUG |
| HD_18AY | AX-111587939 | 1A | 308996879 | 0.4259 | 0.6281 | 5.58E-18 | IIIVmrMLM | SIG |
| HD_18AY | AX-110186328 | 1A | 332380980 | -1.5971 | | 1.29E-06 | GEMMA | SUG |
| HD_18AY | AX-111454083 | 1A | 335612263 | -0.6530 | | 2.03E-06 | BLUPmrMLM | SUG |
| HD_18AY | AX-108988200 | 1A | 355334713 | -1.6771 | | 9.44E-07 | GEMMA | SUG |
| HD_18AY | AX-111668333 | 1A | 388937021 | -1.7448 | | 1.87E-06 | GEMMA | SUG |
| HD_18AY | AX-111462775 | 1A | 494810364 | 1.6851 | | 1.74E-06 | EMMAX | SUG |
| HD_18AY | AX-111462775 | 1A | 494810364 | -1.6849 | | 3.77E-07 | GEMMA | SUG |
| HD_18AY | AX-111453306 | 1A | 494818292 | -1.5244 | | 2.30E-06 | GEMMA | SUG |
| HD_18AY | AX-109886157 | 1A | 494887518 | -1.5952 | | 1.06E-06 | GEMMA | SUG |
| HD_18AY | AX-109426053 | 1A | 494942486 | -1.5952 | | 1.06E-06 | GEMMA | SUG |
| HD_18AY | AX-111488547 | 1A | 495013458 | -1.5952 | | 1.06E-06 | GEMMA | SUG |
| HD_18AY | AX-109817102 | 1A | 495013803 | -1.5226 | | 2.27E-06 | GEMMA | SUG |
| HD_18AY | AX-109490911 | 1A | 495014197 | 1.6851 | | 1.74E-06 | EMMAX | SUG |
| HD_18AY | AX-109490911 | 1A | 495014197 | -1.6849 | | 3.77E-07 | GEMMA | SUG |
| HD_18AY | AX-111609168 | 1A | 495014994 | -1.6403 | | 7.12E-07 | GEMMA | SUG |
| HD_18AY | AX-110606043 | 1A | 495036799 | -1.5952 | | 1.06E-06 | GEMMA | SUG |
| HD_18AY | AX-111550437 | 1A | 495037353 | 1.6851 | | 1.74E-06 | EMMAX | SUG |
| HD_18AY | AX-111550437 | 1A | 495037353 | -1.6849 | | 3.77E-07 | GEMMA | SUG |
| HD_18AY | AX-111002026 | 1A | 495037475 | -1.5952 | | 1.06E-06 | GEMMA | SUG |
| HD_18AY | AX-108796834 | 1A | 495037699 | 1.6851 | | 1.74E-06 | EMMAX | SUG |
| HD_18AY | AX-108796834 | 1A | 495037699 | -1.6849 | | 3.77E-07 | GEMMA | SUG |
| HD_18AY | AX-111767357 | 1A | 495125499 | 1.6348 | | 2.16E-06 | EMMAX | SUG |
| HD_18AY | AX-111767357 | 1A | 495125499 | -1.6293 | | 5.15E-07 | GEMMA | SUG |
| HD_18AY | AX-110995371 | 1A | 495127554 | -1.4917 | | 2.13E-06 | GEMMA | SUG |
| HD_18AY | AX-108926906 | 1A | 495181892 | -1.6458 | | 8.34E-07 | GEMMA | SUG |
| HD_18AY | AX-109448112 | 1A | 495182069 | -1.5952 | | 1.06E-06 | GEMMA | SUG |
| HD_18AY | AX-110667522 | 1A | 495183206 | -1.5558 | | 2.28E-06 | GEMMA | SUG |
| HD_18AY | AX-111555272 | 1A | 495189010 | -1.3640 | | 7.79E-07 | GEMMA | SUG |
| HD_18AY | AX-94652392 | 1A | 517378589 | 0.3593 | | 1.07E-05 | BLUPmrMLM | SUG |
| HD_18AY | AX-111096655 | 1B | 324586060 | -0.4818 | | 3.11E-05 | BLUPmrMLM | SUG |
| HD_18AY | AX-111654753 | 1B | 563555962 | 0.1716 | 1.1495 | 1.05E-12 | IIIVmrMLM | SIG |
| HD_18AY | AX-94425571 | 2A | 778442411 | 0.4169 | | 3.44E-05 | BLUPmrMLM | SUG |
| HD_18AY | AX-95001020 | 2B | 22783709 | -1.8210 | | 2.21E-06 | GEMMA | SUG |
| HD_18AY | AX-109924683 | 2B | 70735536 | -0.4553 | | 1.92E-06 | BLUPmrMLM | SUG |
| HD_18AY | AX-108913257 | 2B | 434968097 | -0.0710 | 1.1372 | 2.07E-12 | IIIVmrMLM | SIG |
| HD_18AY | AX-111047349 | 2B | 643224024 | -0.0265 | -0.7948 | 2.17E-05 | IIIVmrMLM | SUG |
| HD_18AY | AX-95249141 | 2B | 776410822 | 0.0676 | 0.6445 | 5.32E-18 | IIIVmrMLM | SIG |
| HD_18AY | AX-111049506 | 2D | 15384022 | -0.1701 | -0.8954 | 3.09E-22 | IIIVmrMLM | SIG |

| | | | | | | | | |
|---|---|---|---|---|---|---|---|---|
| HD_18AY | AX-108781190 | 3A | 1622310 | -0.1395 | 1.9148 | 4.26E-10 | IIIVmrMLM | SIG |
| HD_18AY | AX-109493197 | 3A | 13042087 | -0.1278 | 0.9238 | 3.21E-09 | IIIVmrMLM | SIG |
| HD_18AY | AX-110447068 | 3A | 13900133 | -0.3951 | | 2.67E-05 | BLUPmrMLM | SUG |
| HD_18AY | AX-110964081 | 3A | 189317195 | -0.9050 | | 2.80E-07 | BLUPmrMLM | SUG |
| HD_18AY | AX-110964081 | 3A | 189317195 | 0.3062 | | 3.93E-10 | IIIVmrMLM | SIG |
| HD_18AY | AX-110365195 | 3B | 123570896 | 0.0356 | 0.8101 | 1.88E-12 | IIIVmrMLM | SIG |
| HD_18AY | AX-110123193 | 3B | 412602158 | -0.0426 | -1.2096 | 1.02E-11 | IIIVmrMLM | SIG |
| HD_18AY | AX-110449389 | 3B | 508303956 | -0.0199 | -1.0935 | 8.03E-09 | IIIVmrMLM | SIG |
| HD_18AY | AX-110147842 | 3B | 685806815 | 0.1409 | 1.3433 | 2.97E-11 | IIIVmrMLM | SIG |
| HD_18AY | AX-110055565 | 3B | 757886685 | -3.4092 | | 5.24E-07 | FarmCPU | SUG |
| HD_18AY | AX-110601323 | 3B | 760960881 | -0.1140 | -1.2308 | 3.08E-06 | IIIVmrMLM | SUG |
| HD_18AY | AX-109304549 | 4A | 29062186 | 0.4653 | 0.2147 | 1.70E-19 | IIIVmrMLM | SIG |
| HD_18AY | AX-109107599 | 4A | 75884266 | 0.0799 | 1.9978 | 9.97E-07 | IIIVmrMLM | SUG |
| HD_18AY | AX-111036297 | 4A | 174713402 | -1.9701 | | 1.82E-06 | GEMMA | SUG |
| HD_18AY | AX-109872804 | 4A | 174817775 | -2.4335 | | 2.82E-07 | GEMMA | SUG |
| HD_18AY | AX-109895285 | 4A | 175005664 | -1.9701 | | 1.82E-06 | GEMMA | SUG |
| HD_18AY | AX-109505745 | 4A | 175121280 | -1.9701 | | 1.82E-06 | GEMMA | SUG |
| HD_18AY | AX-95117398 | 4A | 176387079 | -2.0233 | | 3.17E-07 | GEMMA | SUG |
| HD_18AY | AX-94622840 | 4A | 176387090 | -2.6398 | | 1.80E-06 | GEMMA | SUG |
| HD_18AY | AX-108919346 | 4A | 177637530 | -2.0233 | | 3.17E-07 | GEMMA | SUG |
| HD_18AY | AX-110699780 | 4A | 178490385 | -3.5038 | | 8.12E-07 | GEMMA | SUG |
| HD_18AY | AX-111229340 | 4A | 178746629 | -2.0233 | | 3.17E-07 | GEMMA | SUG |
| HD_18AY | AX-110385657 | 4A | 179474569 | 2.1851 | | 1.62E-06 | EMMAX | SUG |
| HD_18AY | AX-110385657 | 4A | 179474569 | -2.2006 | | 2.04E-07 | GEMMA | SUG |
| HD_18AY | AX-109066809 | 4A | 179941606 | 2.1851 | | 1.62E-06 | EMMAX | SUG |
| HD_18AY | AX-109066809 | 4A | 179941606 | -2.2006 | | 2.04E-07 | GEMMA | SUG |
| HD_18AY | AX-94859293 | 4A | 180405938 | -2.0233 | | 3.17E-07 | GEMMA | SUG |
| HD_18AY | AX-109832498 | 4A | 182917881 | -2.0233 | | 3.17E-07 | GEMMA | SUG |
| HD_18AY | AX-109947779 | 4A | 183286010 | -2.0233 | | 3.17E-07 | GEMMA | SUG |
| HD_18AY | AX-108733392 | 4A | 183976918 | -2.0233 | | 3.17E-07 | GEMMA | SUG |
| HD_18AY | AX-110034111 | 4A | 184801424 | -2.0233 | | 3.17E-07 | GEMMA | SUG |
| HD_18AY | AX-109453226 | 4A | 189934998 | -2.0233 | | 3.17E-07 | GEMMA | SUG |
| HD_18AY | AX-95160821 | 4A | 190189153 | -2.0233 | | 3.17E-07 | GEMMA | SUG |
| HD_18AY | AX-110992403 | 4A | 191051248 | -2.0233 | | 3.17E-07 | GEMMA | SUG |
| HD_18AY | AX-94502524 | 4A | 191131328 | -2.0233 | | 3.17E-07 | GEMMA | SUG |
| HD_18AY | AX-108824978 | 4A | 191189140 | 2.5182 | | 1.61E-06 | EMMAX | SUG |
| HD_18AY | AX-108824978 | 4A | 191189140 | -2.5432 | | 1.68E-07 | GEMMA | SUG |
| HD_18AY | AX-110482544 | 4A | 191586585 | 2.1234 | | 1.70E-06 | EMMAX | SUG |
| HD_18AY | AX-110482544 | 4A | 191586585 | -2.1521 | | 1.81E-07 | GEMMA | SUG |
| HD_18AY | AX-111698357 | 4A | 192774566 | -2.0233 | | 3.17E-07 | GEMMA | SUG |
| HD_18AY | AX-108945836 | 4A | 193097707 | -2.0233 | | 3.17E-07 | GEMMA | SUG |
| HD_18AY | AX-108958423 | 4A | 193626324 | -2.0233 | | 3.17E-07 | GEMMA | SUG |
| HD_18AY | AX-109955137 | 4A | 195392733 | -2.0233 | | 3.17E-07 | GEMMA | SUG |
| HD_18AY | AX-111041723 | 4A | 196730337 | -2.0233 | | 3.17E-07 | GEMMA | SUG |

| | | | | | | | | |
|---|---|---|---|---|---|---|---|---|
| HD_18AY | AX-109045665 | 4A | 196759730 | -2.0233 | | 3.17E-07 | GEMMA | SUG |
| HD_18AY | AX-111161699 | 4A | 198183921 | -2.0233 | | 3.17E-07 | GEMMA | SUG |
| HD_18AY | AX-94957105 | 4A | 199408800 | -2.0233 | | 3.17E-07 | GEMMA | SUG |
| HD_18AY | AX-108853783 | 4A | 199877646 | -1.9701 | | 1.82E-06 | GEMMA | SUG |
| HD_18AY | AX-110052595 | 4A | 200393342 | -1.9701 | | 1.82E-06 | GEMMA | SUG |
| HD_18AY | AX-111606547 | 4A | 202107240 | -2.0233 | | 3.17E-07 | GEMMA | SUG |
| HD_18AY | AX-109355314 | 4A | 202580524 | -2.0233 | | 3.17E-07 | GEMMA | SUG |
| HD_18AY | AX-111256735 | 4A | 203836155 | -2.1126 | | 1.05E-06 | GEMMA | SUG |
| HD_18AY | AX-109049354 | 4A | 203933114 | -2.0233 | | 3.17E-07 | GEMMA | SUG |
| HD_18AY | AX-109942778 | 4A | 204509238 | -2.0233 | | 3.17E-07 | GEMMA | SUG |
| HD_18AY | AX-110945683 | 4A | 204580763 | -2.5725 | | 4.12E-07 | GEMMA | SUG |
| HD_18AY | AX-110711093 | 4A | 204665120 | -2.0233 | | 3.17E-07 | GEMMA | SUG |
| HD_18AY | AX-110683722 | 4A | 206237731 | -1.9701 | | 1.82E-06 | GEMMA | SUG |
| HD_18AY | AX-111530201 | 4A | 207290085 | -2.0233 | | 3.17E-07 | GEMMA | SUG |
| HD_18AY | AX-111115180 | 4A | 207365022 | -2.0233 | | 3.17E-07 | GEMMA | SUG |
| HD_18AY | AX-110925613 | 4A | 209932966 | -2.0233 | | 3.17E-07 | GEMMA | SUG |
| HD_18AY | AX-110025666 | 4A | 210999660 | -1.9701 | | 1.82E-06 | GEMMA | SUG |
| HD_18AY | AX-111769097 | 4A | 211708279 | -2.0233 | | 3.17E-07 | GEMMA | SUG |
| HD_18AY | AX-110699681 | 4A | 214131106 | -2.0233 | | 3.17E-07 | GEMMA | SUG |
| HD_18AY | AX-110977715 | 4A | 214196913 | -2.0233 | | 3.17E-07 | GEMMA | SUG |
| HD_18AY | AX-109889920 | 4A | 214736951 | -2.0233 | | 3.17E-07 | GEMMA | SUG |
| HD_18AY | AX-95233275 | 4A | 215223886 | 4.0724 | | 3.42E-07 | GEMMA | SUG |
| HD_18AY | AX-110583534 | 4A | 216851463 | -2.0233 | | 3.17E-07 | GEMMA | SUG |
| HD_18AY | AX-111497019 | 4A | 219409174 | -2.0233 | | 3.17E-07 | GEMMA | SUG |
| HD_18AY | AX-108827229 | 4A | 219799342 | -2.0233 | | 3.17E-07 | GEMMA | SUG |
| HD_18AY | AX-94550347 | 4A | 220739506 | -2.0233 | | 3.17E-07 | GEMMA | SUG |
| HD_18AY | AX-108978175 | 4A | 221043861 | -2.0233 | | 3.17E-07 | GEMMA | SUG |
| HD_18AY | AX-110681429 | 4A | 221491260 | -2.0233 | | 3.17E-07 | GEMMA | SUG |
| HD_18AY | AX-111133928 | 4A | 221778743 | -2.0233 | | 3.17E-07 | GEMMA | SUG |
| HD_18AY | AX-110435896 | 4A | 222105529 | -2.1311 | | 6.44E-07 | GEMMA | SUG |
| HD_18AY | AX-111011457 | 4A | 222105794 | -1.9701 | | 1.82E-06 | GEMMA | SUG |
| HD_18AY | AX-109363658 | 4A | 225529259 | -2.0233 | | 3.17E-07 | GEMMA | SUG |
| HD_18AY | AX-111670018 | 4A | 226044207 | -2.0233 | | 3.17E-07 | GEMMA | SUG |
| HD_18AY | AX-110463728 | 4A | 226797375 | -1.7581 | | 6.47E-07 | GEMMA | SUG |
| HD_18AY | AX-108937128 | 4A | 227361546 | -2.1126 | | 1.05E-06 | GEMMA | SUG |
| HD_18AY | AX-109591336 | 4A | 227856562 | 2.2213 | | 3.20E-07 | EMMAX | SUG |
| HD_18AY | AX-109591336 | 4A | 227856562 | -2.2325 | | 3.44E-08 | GEMMA | SIG |
| HD_18AY | AX-109960939 | 4A | 228282559 | -2.0233 | | 3.17E-07 | GEMMA | SUG |
| HD_18AY | AX-109501519 | 4A | 229478509 | -2.0233 | | 3.17E-07 | GEMMA | SUG |
| HD_18AY | AX-109638456 | 4A | 231309753 | -2.0233 | | 3.17E-07 | GEMMA | SUG |
| HD_18AY | AX-110188041 | 4A | 231836914 | -1.9701 | | 1.82E-06 | GEMMA | SUG |
| HD_18AY | AX-110176657 | 4A | 232213093 | 2.1851 | | 1.62E-06 | EMMAX | SUG |
| HD_18AY | AX-110176657 | 4A | 232213093 | -2.2006 | | 2.04E-07 | GEMMA | SUG |
| HD_18AY | AX-111808836 | 4A | 232381447 | -2.0233 | | 3.17E-07 | GEMMA | SUG |

| | | | | | | | |
|---|---|---|---|---|---|---|---|
| HD_18AY | AX-109933740 | 4A | 232523882 | -2.3689 | 1.40E-06 | GEMMA | SUG |
| HD_18AY | AX-109350236 | 4A | 233866634 | -2.3042 | 3.11E-07 | GEMMA | SUG |
| HD_18AY | AX-109846101 | 4A | 234888406 | -2.0233 | 3.17E-07 | GEMMA | SUG |
| HD_18AY | AX-111496118 | 4A | 234925642 | -2.0233 | 3.17E-07 | GEMMA | SUG |
| HD_18AY | AX-109319578 | 4A | 234983636 | -2.0233 | 3.17E-07 | GEMMA | SUG |
| HD_18AY | AX-109838175 | 4A | 235052820 | -1.9701 | 1.82E-06 | GEMMA | SUG |
| HD_18AY | AX-111154785 | 4A | 235836721 | -2.0233 | 3.17E-07 | GEMMA | SUG |
| HD_18AY | AX-108892689 | 4A | 236087580 | -1.9701 | 1.82E-06 | GEMMA | SUG |
| HD_18AY | AX-95244223 | 4A | 236130151 | 2.8026 | 2.79E-07 | BLUPmrMLM | SUG |
| HD_18AY | AX-95244223 | 4A | 236130151 | 3.7485 | 4.20E-07 | GEMMA | SUG |
| HD_18AY | AX-110365001 | 4A | 236145472 | -2.1311 | 6.44E-07 | GEMMA | SUG |
| HD_18AY | AX-109467417 | 4A | 237103748 | -2.3482 | 2.86E-07 | GEMMA | SUG |
| HD_18AY | AX-109846555 | 4A | 237113713 | 3.5261 | 2.29E-06 | EMMAX | SUG |
| HD_18AY | AX-109846555 | 4A | 237113713 | -3.5896 | 3.08E-07 | GEMMA | SUG |
| HD_18AY | AX-111268830 | 4A | 237154214 | -1.9964 | 1.81E-06 | GEMMA | SUG |
| HD_18AY | AX-108858024 | 4A | 237687734 | -2.0233 | 3.17E-07 | GEMMA | SUG |
| HD_18AY | AX-110727256 | 4A | 237695603 | 3.4926 | 5.72E-07 | EMMAX | SUG |
| HD_18AY | AX-110727256 | 4A | 237695603 | -3.5403 | 6.72E-08 | GEMMA | SIG |
| HD_18AY | AX-111590554 | 4A | 238193130 | -1.9108 | 6.42E-07 | GEMMA | SUG |
| HD_18AY | AX-110166384 | 4A | 238245029 | -2.0233 | 3.17E-07 | GEMMA | SUG |
| HD_18AY | AX-109357719 | 4A | 238292732 | -2.0233 | 3.17E-07 | GEMMA | SUG |
| HD_18AY | AX-109085773 | 4A | 239945683 | -3.4533 | 3.08E-07 | GEMMA | SUG |
| HD_18AY | AX-109431068 | 4A | 241305627 | -2.0233 | 3.17E-07 | GEMMA | SUG |
| HD_18AY | AX-109621060 | 4A | 241792508 | -2.0233 | 3.17E-07 | GEMMA | SUG |
| HD_18AY | AX-94507064 | 4A | 242877373 | -2.0233 | 3.17E-07 | GEMMA | SUG |
| HD_18AY | AX-109519712 | 4A | 243824586 | -1.9964 | 1.81E-06 | GEMMA | SUG |
| HD_18AY | AX-111625147 | 4A | 244380108 | -2.0233 | 3.17E-07 | GEMMA | SUG |
| HD_18AY | AX-109034391 | 4A | 245959001 | -2.0233 | 3.17E-07 | GEMMA | SUG |
| HD_18AY | AX-108919499 | 4A | 246025438 | -2.0233 | 3.17E-07 | GEMMA | SUG |
| HD_18AY | AX-110136683 | 4A | 246214232 | -2.0233 | 3.17E-07 | GEMMA | SUG |
| HD_18AY | AX-109638506 | 4A | 246221927 | -2.2317 | 1.56E-06 | GEMMA | SUG |
| HD_18AY | AX-110418569 | 4A | 246720295 | -1.9701 | 1.82E-06 | GEMMA | SUG |
| HD_18AY | AX-109515202 | 4A | 247601274 | -2.0233 | 3.17E-07 | GEMMA | SUG |
| HD_18AY | AX-111514566 | 4A | 248887725 | -2.0233 | 3.17E-07 | GEMMA | SUG |
| HD_18AY | AX-111150736 | 4A | 250043729 | -2.0233 | 3.17E-07 | GEMMA | SUG |
| HD_18AY | AX-111012469 | 4A | 251732649 | -2.0233 | 3.17E-07 | GEMMA | SUG |
| HD_18AY | AX-111545105 | 4A | 251862942 | -2.0233 | 3.17E-07 | GEMMA | SUG |
| HD_18AY | AX-110453968 | 4A | 252123154 | -2.0233 | 3.17E-07 | GEMMA | SUG |
| HD_18AY | AX-110703922 | 4A | 253017373 | -2.0233 | 3.17E-07 | GEMMA | SUG |
| HD_18AY | AX-109496160 | 4A | 254470697 | -2.0447 | 3.20E-07 | GEMMA | SUG |
| HD_18AY | AX-108861243 | 4A | 255163375 | -2.0233 | 3.17E-07 | GEMMA | SUG |
| HD_18AY | AX-111208327 | 4A | 255337212 | -2.0233 | 3.17E-07 | GEMMA | SUG |
| HD_18AY | AX-108845537 | 4A | 257351928 | -2.1126 | 1.05E-06 | GEMMA | SUG |
| HD_18AY | AX-109446132 | 4A | 257352494 | -2.1126 | 1.05E-06 | GEMMA | SUG |

| | | | | | | | |
|---|---|---|---|---|---|---|---|
| HD_18AY | AX-109078019 | 4A | 258632382 | -1.8889 | 7.00E-07 | GEMMA | SUG |
| HD_18AY | AX-110679344 | 4A | 260573692 | -1.9306 | 6.08E-07 | GEMMA | SUG |
| HD_18AY | AX-109855587 | 4A | 260611119 | -2.0233 | 3.17E-07 | GEMMA | SUG |
| HD_18AY | AX-111682476 | 4A | 260724062 | -2.0233 | 3.17E-07 | GEMMA | SUG |
| HD_18AY | AX-109964578 | 4A | 261107021 | -2.0233 | 3.17E-07 | GEMMA | SUG |
| HD_18AY | AX-110601497 | 4A | 261528420 | -3.0402 | 1.14E-06 | GEMMA | SUG |
| HD_18AY | AX-109998941 | 4A | 261553137 | -3.0402 | 1.14E-06 | GEMMA | SUG |
| HD_18AY | AX-109008762 | 4A | 261919735 | -2.0233 | 3.17E-07 | GEMMA | SUG |
| HD_18AY | AX-110506031 | 4A | 262333845 | -2.0233 | 3.17E-07 | GEMMA | SUG |
| HD_18AY | AX-110613121 | 4A | 262480899 | 1.9461 | 1.49E-06 | EMMAX | SUG |
| HD_18AY | AX-110613121 | 4A | 262480899 | -1.9778 | 1.14E-07 | GEMMA | SIG |
| HD_18AY | AX-110139570 | 4A | 262633186 | -1.9701 | 1.82E-06 | GEMMA | SUG |
| HD_18AY | AX-110063999 | 4A | 262730149 | -2.0233 | 3.17E-07 | GEMMA | SUG |
| HD_18AY | AX-108801245 | 4A | 262731738 | -2.0233 | 3.17E-07 | GEMMA | SUG |
| HD_18AY | AX-111053161 | 4A | 262810740 | -2.0233 | 3.17E-07 | GEMMA | SUG |
| HD_18AY | AX-109082147 | 4A | 263018072 | -2.0447 | 3.20E-07 | GEMMA | SUG |
| HD_18AY | AX-111565544 | 4A | 263192525 | -2.0233 | 3.17E-07 | GEMMA | SUG |
| HD_18AY | AX-110566534 | 4A | 263466377 | -2.0233 | 3.17E-07 | GEMMA | SUG |
| HD_18AY | AX-108798054 | 4A | 263501794 | -1.9701 | 1.82E-06 | GEMMA | SUG |
| HD_18AY | AX-94477308 | 4A | 277990869 | -2.1311 | 6.44E-07 | GEMMA | SUG |
| HD_18AY | AX-110495470 | 4A | 295000074 | -2.0233 | 3.17E-07 | GEMMA | SUG |
| HD_18AY | AX-95219351 | 4A | 296862915 | -2.0233 | 3.17E-07 | GEMMA | SUG |
| HD_18AY | AX-111629781 | 4A | 297202240 | -3.5038 | 8.12E-07 | GEMMA | SUG |
| HD_18AY | AX-109449887 | 4A | 297257813 | -2.0233 | 3.17E-07 | GEMMA | SUG |
| HD_18AY | AX-108733323 | 4A | 297261385 | -2.0233 | 3.17E-07 | GEMMA | SUG |
| HD_18AY | AX-109501673 | 4A | 297970326 | -2.0233 | 3.17E-07 | GEMMA | SUG |
| HD_18AY | AX-109624312 | 4A | 298166299 | -3.4532 | 3.14E-07 | GEMMA | SUG |
| HD_18AY | AX-109090565 | 4A | 298352750 | -1.9701 | 1.82E-06 | GEMMA | SUG |
| HD_18AY | AX-108860621 | 4A | 298550082 | -1.9701 | 1.82E-06 | GEMMA | SUG |
| HD_18AY | AX-109642318 | 4A | 298801739 | -2.0233 | 3.17E-07 | GEMMA | SUG |
| HD_18AY | AX-111665679 | 4A | 299019804 | -2.7173 | 6.01E-07 | GEMMA | SUG |
| HD_18AY | AX-108874744 | 4A | 299564783 | -2.0233 | 3.17E-07 | GEMMA | SUG |
| HD_18AY | AX-111552703 | 4A | 300466283 | -2.0233 | 3.17E-07 | GEMMA | SUG |
| HD_18AY | AX-111016394 | 4A | 300956481 | -2.0233 | 3.17E-07 | GEMMA | SUG |
| HD_18AY | AX-110134384 | 4A | 302187470 | -1.9051 | 8.35E-07 | GEMMA | SUG |
| HD_18AY | AX-108892346 | 4A | 302528355 | -2.0233 | 3.17E-07 | GEMMA | SUG |
| HD_18AY | AX-110470114 | 4A | 302909715 | -2.0233 | 3.17E-07 | GEMMA | SUG |
| HD_18AY | AX-111656906 | 4A | 303284456 | 2.0163 | 2.07E-06 | EMMAX | SUG |
| HD_18AY | AX-111656906 | 4A | 303284456 | -2.0453 | 1.65E-07 | GEMMA | SUG |
| HD_18AY | AX-110918601 | 4A | 303420004 | -1.9957 | 9.60E-07 | GEMMA | SUG |
| HD_18AY | AX-110962439 | 4A | 303776128 | -1.9957 | 9.60E-07 | GEMMA | SUG |
| HD_18AY | AX-111563778 | 4A | 303813393 | -1.9957 | 9.60E-07 | GEMMA | SUG |
| HD_18AY | AX-109865197 | 4A | 305321313 | -1.9701 | 1.82E-06 | GEMMA | SUG |
| HD_18AY | AX-110704572 | 4A | 306425580 | -2.0233 | 3.17E-07 | GEMMA | SUG |

| | | | | | | | |
|---|---|---|---|---|---|---|---|
| HD_18AY | AX-108897689 | 4A | 306918700 | -2.0233 | 3.17E-07 | GEMMA | SUG |
| HD_18AY | AX-110166551 | 4A | 309010201 | -2.0233 | 3.17E-07 | GEMMA | SUG |
| HD_18AY | AX-111263014 | 4A | 309018917 | -2.0233 | 3.17E-07 | GEMMA | SUG |
| HD_18AY | AX-109431295 | 4A | 309084001 | -2.0233 | 3.17E-07 | GEMMA | SUG |
| HD_18AY | AX-111059298 | 4A | 309201953 | -2.0233 | 3.17E-07 | GEMMA | SUG |
| HD_18AY | AX-108906737 | 4A | 309201984 | -2.0233 | 3.17E-07 | GEMMA | SUG |
| HD_18AY | AX-109981270 | 4A | 309388007 | -1.9921 | 3.49E-07 | GEMMA | SUG |
| HD_18AY | AX-111269759 | 4A | 309542418 | -1.9701 | 1.82E-06 | GEMMA | SUG |
| HD_18AY | AX-111672800 | 4A | 309806911 | -2.2447 | 4.12E-07 | GEMMA | SUG |
| HD_18AY | AX-109635876 | 4A | 310387847 | -3.1193 | 6.38E-07 | GEMMA | SUG |
| HD_18AY | AX-111501964 | 4A | 312675275 | -2.0233 | 3.17E-07 | GEMMA | SUG |
| HD_18AY | AX-110943747 | 4A | 312722683 | -2.0233 | 3.17E-07 | GEMMA | SUG |
| HD_18AY | AX-110695947 | 4A | 314240348 | -2.0846 | 3.72E-07 | GEMMA | SUG |
| HD_18AY | AX-109063836 | 4A | 314707804 | -2.0233 | 3.17E-07 | GEMMA | SUG |
| HD_18AY | AX-109078940 | 4A | 321565249 | -1.9701 | 1.82E-06 | GEMMA | SUG |
| HD_18AY | AX-109606131 | 4A | 322238693 | -2.0233 | 3.17E-07 | GEMMA | SUG |
| HD_18AY | AX-109024444 | 4A | 322881780 | -2.0233 | 3.17E-07 | GEMMA | SUG |
| HD_18AY | AX-111544686 | 4A | 322883021 | -1.9964 | 1.81E-06 | GEMMA | SUG |
| HD_18AY | AX-109046234 | 4A | 323173734 | -2.0233 | 3.17E-07 | GEMMA | SUG |
| HD_18AY | AX-111702369 | 4A | 324631545 | -2.0233 | 3.17E-07 | GEMMA | SUG |
| HD_18AY | AX-110421400 | 4A | 324639269 | -2.0233 | 3.17E-07 | GEMMA | SUG |
| HD_18AY | AX-109303419 | 4A | 326168995 | -2.0233 | 3.17E-07 | GEMMA | SUG |
| HD_18AY | AX-111529741 | 4A | 327087786 | -2.0233 | 3.17E-07 | GEMMA | SUG |
| HD_18AY | AX-108760287 | 4A | 327280819 | -1.9964 | 1.81E-06 | GEMMA | SUG |
| HD_18AY | AX-110921824 | 4A | 327942869 | -1.7155 | 5.93E-07 | GEMMA | SUG |
| HD_18AY | AX-109955677 | 4A | 343199577 | -2.0233 | 3.17E-07 | GEMMA | SUG |
| HD_18AY | AX-111804447 | 4A | 343340890 | -2.0233 | 3.17E-07 | GEMMA | SUG |
| HD_18AY | AX-110973932 | 4A | 344035688 | -1.9701 | 1.82E-06 | GEMMA | SUG |
| HD_18AY | AX-109943224 | 4A | 344389722 | -2.0233 | 3.17E-07 | GEMMA | SUG |
| HD_18AY | AX-110155922 | 4A | 344657840 | -2.0233 | 3.17E-07 | GEMMA | SUG |
| HD_18AY | AX-111559555 | 4A | 346137584 | -2.0233 | 3.17E-07 | GEMMA | SUG |
| HD_18AY | AX-110012070 | 4A | 346932242 | -2.0233 | 3.17E-07 | GEMMA | SUG |
| HD_18AY | AX-108909395 | 4A | 347266134 | -2.0377 | 2.14E-06 | GEMMA | SUG |
| HD_18AY | AX-108769971 | 4A | 347426354 | -1.9701 | 1.82E-06 | GEMMA | SUG |
| HD_18AY | AX-108891522 | 4A | 347884168 | -2.0233 | 3.17E-07 | GEMMA | SUG |
| HD_18AY | AX-110608805 | 4A | 348610204 | -2.0233 | 3.17E-07 | GEMMA | SUG |
| HD_18AY | AX-111681450 | 4A | 349171706 | 2.0389 | 1.39E-06 | EMMAX | SUG |
| HD_18AY | AX-111681450 | 4A | 349171706 | -2.0701 | 9.83E-08 | GEMMA | SIG |
| HD_18AY | AX-111108057 | 4A | 349299114 | -2.0233 | 3.17E-07 | GEMMA | SUG |
| HD_18AY | AX-110177447 | 4A | 349303567 | -2.0233 | 3.17E-07 | GEMMA | SUG |
| HD_18AY | AX-108940071 | 4A | 349855742 | -2.0233 | 3.17E-07 | GEMMA | SUG |
| HD_18AY | AX-110915668 | 4A | 349931328 | -2.0233 | 3.17E-07 | GEMMA | SUG |
| HD_18AY | AX-110575029 | 4A | 349974022 | -1.9701 | 1.82E-06 | GEMMA | SUG |
| HD_18AY | AX-109283082 | 4A | 350119133 | -2.0233 | 3.17E-07 | GEMMA | SUG |

| | | | | | | | |
|---|---|---|---|---|---|---|---|
| HD_18AY | AX-110187659 | 4A | 350347493 | -2.0233 | 3.17E-07 | GEMMA | SUG |
| HD_18AY | AX-111563737 | 4A | 350731597 | -2.0233 | 3.17E-07 | GEMMA | SUG |
| HD_18AY | AX-111262946 | 4A | 351093736 | -2.0233 | 3.17E-07 | GEMMA | SUG |
| HD_18AY | AX-111792757 | 4A | 351093821 | -2.0447 | 3.20E-07 | GEMMA | SUG |
| HD_18AY | AX-111592274 | 4A | 351793378 | -2.0233 | 3.17E-07 | GEMMA | SUG |
| HD_18AY | AX-109008181 | 4A | 352485867 | -2.0233 | 3.17E-07 | GEMMA | SUG |
| HD_18AY | AX-110407556 | 4A | 352513820 | -2.0233 | 3.17E-07 | GEMMA | SUG |
| HD_18AY | AX-108927323 | 4A | 352514126 | -2.0233 | 3.17E-07 | GEMMA | SUG |
| HD_18AY | AX-109877185 | 4A | 353466969 | -2.0233 | 3.17E-07 | GEMMA | SUG |
| HD_18AY | AX-111731859 | 4A | 353873985 | -2.0233 | 3.17E-07 | GEMMA | SUG |
| HD_18AY | AX-109942758 | 4A | 357436291 | -2.0233 | 3.17E-07 | GEMMA | SUG |
| HD_18AY | AX-109829567 | 4A | 358123548 | -2.0233 | 3.17E-07 | GEMMA | SUG |
| HD_18AY | AX-109913904 | 4A | 358982866 | -2.0233 | 3.17E-07 | GEMMA | SUG |
| HD_18AY | AX-109033949 | 4A | 360635039 | -1.8940 | 1.25E-06 | GEMMA | SUG |
| HD_18AY | AX-110608887 | 4A | 361305064 | -2.6672 | 7.62E-07 | GEMMA | SUG |
| HD_18AY | AX-111619720 | 4A | 361541057 | -2.0233 | 3.17E-07 | GEMMA | SUG |
| HD_18AY | AX-109514447 | 4A | 362059730 | -2.0233 | 3.17E-07 | GEMMA | SUG |
| HD_18AY | AX-109400620 | 4A | 363329924 | -2.0233 | 3.17E-07 | GEMMA | SUG |
| HD_18AY | AX-109385715 | 4A | 365203816 | -2.0233 | 3.17E-07 | GEMMA | SUG |
| HD_18AY | AX-94544093 | 4A | 365650024 | 2.1345 | 1.66E-06 | EMMAX | SUG |
| HD_18AY | AX-94544093 | 4A | 365650024 | -2.1701 | 1.09E-07 | GEMMA | SIG |
| HD_18AY | AX-108864812 | 4A | 366643698 | -2.0233 | 3.17E-07 | GEMMA | SUG |
| HD_18AY | AX-110996449 | 4A | 367347533 | -1.9701 | 1.82E-06 | GEMMA | SUG |
| HD_18AY | AX-110967044 | 4A | 368566235 | -2.0233 | 3.17E-07 | GEMMA | SUG |
| HD_18AY | AX-111632994 | 4A | 368931761 | -2.0233 | 3.17E-07 | GEMMA | SUG |
| HD_18AY | AX-110170422 | 4A | 370926619 | -1.9147 | 1.73E-06 | GEMMA | SUG |
| HD_18AY | AX-110527185 | 4A | 371053253 | -2.0233 | 3.17E-07 | GEMMA | SUG |
| HD_18AY | AX-110413368 | 4A | 372105907 | -2.0233 | 3.17E-07 | GEMMA | SUG |
| HD_18AY | AX-110152356 | 4A | 375908026 | 2.2213 | 3.20E-07 | EMMAX | SUG |
| HD_18AY | AX-110152356 | 4A | 375908026 | -2.2325 | 3.44E-08 | GEMMA | SIG |
| HD_18AY | AX-110422521 | 4A | 376360887 | -2.0233 | 3.17E-07 | GEMMA | SUG |
| HD_18AY | AX-110492962 | 4A | 376758845 | -2.0233 | 3.17E-07 | GEMMA | SUG |
| HD_18AY | AX-109373474 | 4A | 377259904 | -2.0447 | 3.20E-07 | GEMMA | SUG |
| HD_18AY | AX-109454861 | 4A | 378292942 | -1.9701 | 1.82E-06 | GEMMA | SUG |
| HD_18AY | AX-109529885 | 4A | 379462816 | -2.0233 | 3.17E-07 | GEMMA | SUG |
| HD_18AY | AX-108762665 | 4A | 379714219 | -2.3674 | 9.30E-07 | GEMMA | SUG |
| HD_18AY | AX-110546053 | 4A | 380139725 | -2.0233 | 3.17E-07 | GEMMA | SUG |
| HD_18AY | AX-111678616 | 4A | 380592323 | -1.9701 | 1.82E-06 | GEMMA | SUG |
| HD_18AY | AX-110679455 | 4A | 380592843 | -2.0233 | 3.17E-07 | GEMMA | SUG |
| HD_18AY | AX-110061118 | 4A | 380594177 | -2.0233 | 3.17E-07 | GEMMA | SUG |
| HD_18AY | AX-109635573 | 4A | 381268405 | -2.0233 | 3.17E-07 | GEMMA | SUG |
| HD_18AY | AX-111045266 | 4A | 381459131 | 3.3313 | 8.12E-07 | EMMAX | SUG |
| HD_18AY | AX-111045266 | 4A | 381459131 | -3.3730 | 7.57E-08 | GEMMA | SIG |
| HD_18AY | AX-111668662 | 4A | 383186144 | -2.0233 | 3.17E-07 | GEMMA | SUG |

| | | | | | | | |
|---|---|---|---|---|---|---|---|
| HD_18AY | AX-108848178 | 4A | 383541570 | -1.9701 | 1.82E-06 | GEMMA | SUG |
| HD_18AY | AX-111002437 | 4A | 384206256 | -2.1816 | 3.43E-07 | GEMMA | SUG |
| HD_18AY | AX-109994432 | 4A | 384381830 | -1.9964 | 1.81E-06 | GEMMA | SUG |
| HD_18AY | AX-110433402 | 4A | 388755277 | -2.0233 | 3.17E-07 | GEMMA | SUG |
| HD_18AY | AX-108918290 | 4A | 388881263 | -1.9701 | 1.82E-06 | GEMMA | SUG |
| HD_18AY | AX-109007570 | 4A | 389231572 | -2.0233 | 3.17E-07 | GEMMA | SUG |
| HD_18AY | AX-108852119 | 4A | 390455289 | -2.0233 | 3.17E-07 | GEMMA | SUG |
| HD_18AY | AX-108820793 | 4A | 390856898 | -2.2317 | 1.56E-06 | GEMMA | SUG |
| HD_18AY | AX-108770359 | 4A | 393594047 | -2.0233 | 3.17E-07 | GEMMA | SUG |
| HD_18AY | AX-109605630 | 4A | 393907277 | -2.0233 | 3.17E-07 | GEMMA | SUG |
| HD_18AY | AX-110566563 | 4A | 394181847 | -2.0233 | 3.17E-07 | GEMMA | SUG |
| HD_18AY | AX-111157719 | 4A | 395278476 | -1.9701 | 1.82E-06 | GEMMA | SUG |
| HD_18AY | AX-110488599 | 4A | 396066205 | -2.0233 | 3.17E-07 | GEMMA | SUG |
| HD_18AY | AX-109343087 | 4A | 396112866 | -1.9701 | 1.82E-06 | GEMMA | SUG |
| HD_18AY | AX-110065075 | 4A | 396690115 | 3.5444 | 4.00E-07 | EMMAX | SUG |
| HD_18AY | AX-110065075 | 4A | 396690115 | -3.5723 | 4.18E-08 | GEMMA | SIG |
| HD_18AY | AX-110947034 | 4A | 397370118 | -2.0233 | 3.17E-07 | GEMMA | SUG |
| HD_18AY | AX-108927015 | 4A | 397898825 | -2.0233 | 3.17E-07 | GEMMA | SUG |
| HD_18AY | AX-109909525 | 4A | 397960287 | -2.0233 | 3.17E-07 | GEMMA | SUG |
| HD_18AY | AX-111467951 | 4A | 398463372 | -2.1311 | 6.44E-07 | GEMMA | SUG |
| HD_18AY | AX-110124990 | 4A | 398747845 | -2.0233 | 3.17E-07 | GEMMA | SUG |
| HD_18AY | AX-111508119 | 4A | 398871722 | -2.0233 | 3.17E-07 | GEMMA | SUG |
| HD_18AY | AX-111115246 | 4A | 398872758 | -2.0233 | 3.17E-07 | GEMMA | SUG |
| HD_18AY | AX-108741917 | 4A | 398988590 | -2.1126 | 1.05E-06 | GEMMA | SUG |
| HD_18AY | AX-109422364 | 4A | 399223501 | -2.0233 | 3.17E-07 | GEMMA | SUG |
| HD_18AY | AX-110587927 | 4A | 401607667 | -2.0233 | 3.17E-07 | GEMMA | SUG |
| HD_18AY | AX-109638869 | 4A | 404148515 | -2.1311 | 6.44E-07 | GEMMA | SUG |
| HD_18AY | AX-111572808 | 4A | 404228421 | 2.0701 | 6.12E-07 | EMMAX | SUG |
| HD_18AY | AX-111572808 | 4A | 404228421 | -2.0978 | 4.27E-08 | GEMMA | SIG |
| HD_18AY | AX-111703243 | 4A | 405493041 | -2.0233 | 3.17E-07 | GEMMA | SUG |
| HD_18AY | AX-111003428 | 4A | 405757048 | -1.9848 | 3.66E-07 | GEMMA | SUG |
| HD_18AY | AX-108856312 | 4A | 409600371 | -1.9663 | 4.42E-07 | GEMMA | SUG |
| HD_18AY | AX-109069284 | 4A | 409715641 | -2.0233 | 3.17E-07 | GEMMA | SUG |
| HD_18AY | AX-110020835 | 4A | 409744478 | -1.9701 | 1.82E-06 | GEMMA | SUG |
| HD_18AY | AX-111225441 | 4A | 409857891 | -2.0233 | 3.17E-07 | GEMMA | SUG |
| HD_18AY | AX-111491783 | 4A | 410392203 | -2.0233 | 3.17E-07 | GEMMA | SUG |
| HD_18AY | AX-108887564 | 4A | 410396554 | -2.0233 | 3.17E-07 | GEMMA | SUG |
| HD_18AY | AX-111731411 | 4A | 410397413 | -2.0233 | 3.17E-07 | GEMMA | SUG |
| HD_18AY | AX-109938548 | 4A | 410539662 | -2.0233 | 3.17E-07 | GEMMA | SUG |
| HD_18AY | AX-109955380 | 4A | 410719058 | -1.9701 | 1.82E-06 | GEMMA | SUG |
| HD_18AY | AX-111269815 | 4A | 411555312 | -2.0233 | 3.17E-07 | GEMMA | SUG |
| HD_18AY | AX-109439820 | 4A | 412138385 | -2.0233 | 3.17E-07 | GEMMA | SUG |
| HD_18AY | AX-111093231 | 4A | 412919825 | -2.0233 | 3.17E-07 | GEMMA | SUG |
| HD_18AY | AX-109608730 | 4A | 413757062 | 2.8229 | 2.44E-07 | EMMAX | SUG |

| | | | | | | | | |
|---|---|---|---|---|---|---|---|---|
| HD_18AY | AX-109608730 | 4A | 413757062 | -2.8396 | | 2.40E-08 | GEMMA | SIG |
| HD_18AY | AX-111785166 | 4A | 413758721 | -1.9484 | | 2.03E-07 | GEMMA | SUG |
| HD_18AY | AX-110175912 | 4A | 414083605 | -2.0233 | | 3.17E-07 | GEMMA | SUG |
| HD_18AY | AX-110695426 | 4A | 414096110 | -2.0233 | | 3.17E-07 | GEMMA | SUG |
| HD_18AY | AX-110682029 | 4A | 414867539 | -1.8899 | | 1.14E-06 | GEMMA | SUG |
| HD_18AY | AX-111559596 | 4A | 415380598 | -2.0233 | | 3.17E-07 | GEMMA | SUG |
| HD_18AY | AX-108843640 | 4A | 415470100 | -2.0233 | | 3.17E-07 | GEMMA | SUG |
| HD_18AY | AX-109379729 | 4A | 415622605 | -2.0233 | | 3.17E-07 | GEMMA | SUG |
| HD_18AY | AX-95185102 | 4A | 415683860 | -2.0233 | | 3.17E-07 | GEMMA | SUG |
| HD_18AY | AX-109918424 | 4A | 417150345 | -2.0233 | | 3.17E-07 | GEMMA | SUG |
| HD_18AY | AX-110711044 | 4A | 418762817 | -2.0233 | | 3.17E-07 | GEMMA | SUG |
| HD_18AY | AX-109080721 | 4A | 422865265 | -2.0233 | | 3.17E-07 | GEMMA | SUG |
| HD_18AY | AX-108839433 | 4A | 425663056 | 2.6410 | | 5.63E-07 | EMMAX | SUG |
| HD_18AY | AX-108839433 | 4A | 425663056 | -2.6679 | | 6.64E-08 | GEMMA | SIG |
| HD_18AY | AX-111075384 | 4A | 425837222 | -2.0233 | | 3.17E-07 | GEMMA | SUG |
| HD_18AY | AX-111511015 | 4A | 426559569 | -3.6340 | | 3.26E-07 | GEMMA | SUG |
| HD_18AY | AX-111577223 | 4A | 427078459 | -2.0233 | | 3.17E-07 | GEMMA | SUG |
| HD_18AY | AX-95204648 | 4A | 427882980 | -1.9147 | | 1.73E-06 | GEMMA | SUG |
| HD_18AY | AX-111486490 | 4A | 428091895 | -1.9369 | | 5.71E-07 | GEMMA | SUG |
| HD_18AY | AX-110964486 | 4A | 429567889 | -2.0233 | | 3.17E-07 | GEMMA | SUG |
| HD_18AY | AX-110444220 | 4A | 429807122 | -2.0233 | | 3.17E-07 | GEMMA | SUG |
| HD_18AY | AX-108778808 | 4A | 431576669 | -2.0233 | | 3.17E-07 | GEMMA | SUG |
| HD_18AY | AX-110499747 | 4A | 433180618 | -2.1110 | | 2.29E-06 | GEMMA | SUG |
| HD_18AY | AX-111450730 | 4A | 433916310 | -2.0233 | | 3.17E-07 | GEMMA | SUG |
| HD_18AY | AX-108778812 | 4A | 434034619 | -2.0233 | | 3.17E-07 | GEMMA | SUG |
| HD_18AY | AX-110584903 | 4A | 435827430 | -2.0079 | | 6.59E-07 | GEMMA | SUG |
| HD_18AY | AX-110026838 | 4A | 438319693 | -2.0233 | | 3.17E-07 | GEMMA | SUG |
| HD_18AY | AX-111033575 | 4A | 438400938 | -2.0233 | | 3.17E-07 | GEMMA | SUG |
| HD_18AY | AX-111509680 | 4A | 439610889 | -1.9701 | | 1.82E-06 | GEMMA | SUG |
| HD_18AY | AX-111788263 | 4A | 440179446 | -2.0233 | | 3.17E-07 | GEMMA | SUG |
| HD_18AY | AX-111763604 | 4A | 440740721 | -2.0233 | | 3.17E-07 | GEMMA | SUG |
| HD_18AY | AX-110998443 | 4A | 441948741 | -2.0233 | | 3.17E-07 | GEMMA | SUG |
| HD_18AY | AX-110047333 | 4A | 443806606 | -2.0233 | | 3.17E-07 | GEMMA | SUG |
| HD_18AY | AX-111634630 | 4A | 444474180 | -2.0233 | | 3.17E-07 | GEMMA | SUG |
| HD_18AY | AX-110031134 | 4A | 446743459 | 2.1021 | | 6.39E-07 | EMMAX | SUG |
| HD_18AY | AX-110031134 | 4A | 446743459 | -2.1208 | | 5.38E-08 | GEMMA | SIG |
| HD_18AY | AX-94542052 | 4A | 448254696 | -1.9701 | | 1.82E-06 | GEMMA | SUG |
| HD_18AY | AX-110004800 | 4A | 448504214 | 0.3886 | -3.131 | 1.00E-20 | IIIVmrMLM | SIG |
| HD_18AY | AX-109441116 | 4A | 448753319 | -2.0233 | | 3.17E-07 | GEMMA | SUG |
| HD_18AY | AX-109363581 | 4A | 449024375 | -1.9306 | | 4.18E-07 | GEMMA | SUG |
| HD_18AY | AX-110906601 | 4A | 449116604 | -1.9701 | | 1.82E-06 | GEMMA | SUG |
| HD_18AY | AX-111027628 | 4A | 597845056 | 0.3574 | | 3.71E-05 | BLUPmrMLM | SUG |
| HD_18AY | AX-108962195 | 4A | 601009409 | 0.4265 | 0.0407 | 4.19E-11 | IIIVmrMLM | SIG |
| HD_18AY | AX-89552384 | 4A | 616986956 | -0.0036 | 1.5367 | 4.91E-05 | IIIVmrMLM | SUG |

| | | | | | | | | |
|---|---|---|---|---|---|---|---|---|
| HD_18AY | AX-95111537 | 4A | 732004989 | 0.3692 | | 6.87E-14 | IIIVmrMLM | SIG |
| HD_18AY | AX-94482010 | 4B | 211981763 | -2.0233 | | 3.17E-07 | GEMMA | SUG |
| HD_18AY | AX-109010485 | 4B | 362147497 | -2.0431 | | 2.22E-06 | GEMMA | SUG |
| HD_18AY | AX-110620525 | 4B | 612215473 | 0.3902 | 0.1337 | 7.78E-13 | IIIVmrMLM | SIG |
| HD_18AY | AX-110394481 | 4D | 27392522 | 0.5359 | -2.2838 | 7.92E-26 | IIIVmrMLM | SIG |
| HD_18AY | AX-111970247 | 4D | 32225033 | -0.3862 | | 3.97E-06 | BLUPmrMLM | SUG |
| HD_18AY | AX-94920138 | 4D | 271209029 | -2.8176 | | 1.50E-06 | GEMMA | SUG |
| HD_18AY | AX-109974153 | 5A | 324460015 | 0.1274 | -3.6808 | 2.95E-10 | IIIVmrMLM | SIG |
| HD_18AY | AX-110435059 | 5B | 15036583 | -0.1473 | -1.52 | 1.01E-14 | IIIVmrMLM | SIG |
| HD_18AY | AX-111146075 | 5B | 381383662 | 0.0425 | 2.2145 | 4.34E-06 | IIIVmrMLM | SUG |
| HD_18AY | AX-110666622 | 5B | 477885647 | -0.5771 | | 1.30E-06 | BLUPmrMLM | SUG |
| HD_18AY | AX-111606848 | 5B | 487218162 | 0.5259 | 0.2021 | 1.93E-19 | IIIVmrMLM | SIG |
| HD_18AY | AX-109102787 | 5B | 502122832 | -0.7743 | | 3.75E-06 | BLUPmrMLM | SUG |
| HD_18AY | AX-108787864 | 5B | 508062299 | -0.1664 | -0.731 | 1.23E-08 | IIIVmrMLM | SIG |
| HD_18AY | AX-110999541 | 5B | 524224803 | 0.4785 | -2.8039 | 1.43E-25 | IIIVmrMLM | SIG |
| HD_18AY | AX-110919389 | 5B | 547313539 | -0.2763 | 0.1719 | 1.62E-07 | IIIVmrMLM | SUG |
| HD_18AY | AX-111049187 | 5B | 569827392 | 0.3991 | | 8.49E-05 | BLUPmrMLM | SUG |
| HD_18AY | AX-109498841 | 5B | 587477780 | -2.1969 | | 2.28E-06 | GEMMA | SUG |
| HD_18AY | AX-109216165 | 5D | 250785260 | 0.4179 | | 3.56E-17 | IIIVmrMLM | SIG |
| HD_18AY | AX-95092292 | 6A | 63303203 | -0.4164 | | 1.89E-05 | BLUPmrMLM | SUG |
| HD_18AY | AX-109908385 | 6A | 610673853 | -0.0511 | -1.3471 | 6.91E-10 | IIIVmrMLM | SIG |
| HD_18AY | AX-109329712 | 6B | 58106508 | -0.2805 | | 9.12E-09 | IIIVmrMLM | SIG |
| HD_18AY | AX-110062747 | 7A | 36164187 | -0.4979 | | 3.71E-08 | BLUPmrMLM | SIG |
| HD_18AY | AX-109868635 | 7A | 36485440 | 0.5939 | -0.0513 | 5.05E-30 | IIIVmrMLM | SIG |
| HD_18AY | AX-94584749 | 7B | 23908708 | -3.3891 | | 2.04E-06 | GEMMA | SUG |
| HD_18AY | AX-110393117 | 7B | 80591805 | -0.1245 | -1.0099 | 1.18E-05 | IIIVmrMLM | SUG |
| HD_18AY | AX-111200188 | 7B | 725964729 | -0.2959 | | 9.04E-06 | BLUPmrMLM | SUG |
| HD_18AY | AX-109829922 | 7B | 732596318 | 0.4411 | | 7.60E-19 | IIIVmrMLM | SIG |
| HD_18AY | AX-110798092 | 7D | 156962271 | -0.0342 | -1.2006 | 1.31E-08 | IIIVmrMLM | SIG |
| HD_18AY | AX-110448713 | 7D | 313395287 | -0.1124 | -3.0453 | 8.20E-09 | IIIVmrMLM | SIG |
| HD_18AY | AX-108928296 | 7D | 448853259 | 0.1532 | -0.6119 | 5.64E-11 | IIIVmrMLM | SIG |
| HD_18AY | AX-110121781 | 7D | 526776624 | 0.0532 | 3.3517 | 9.61E-08 | IIIVmrMLM | SIG |
| HD_18JY | AX-110988612 | 1A | 537530548 | -0.5845 | | 1.09E-05 | BLUPmrMLM | SUG |
| HD_18JY | AX-109870442 | 1A | 543689983 | -0.3627 | | 1.23E-06 | BLUPmrMLM | SUG |
| HD_18JY | AX-111136687 | 1A | 554211078 | -0.5182 | | 5.05E-09 | BLUPmrMLM | SIG |
| HD_18JY | AX-110185658 | 1B | 549031437 | -1.0306 | | 8.21E-07 | FarmCPU | SUG |
| HD_18JY | AX-111022013 | 1B | 549361711 | -1.0462 | | 5.58E-07 | FarmCPU | SUG |
| HD_18JY | AX-95117433 | 1B | 572293440 | 0.5491 | | 6.62E-10 | BLUPmrMLM | SIG |
| HD_18JY | AX-109946093 | 1D | 156055441 | 0.5551 | | 1.48E-06 | BLUPmrMLM | SUG |
| HD_18JY | AX-94570276 | 1D | 338942538 | 0.4943 | | 1.66E-06 | BLUPmrMLM | SUG |
| HD_18JY | AX-108825757 | 2A | 541883453 | -0.4224 | | 7.12E-06 | BLUPmrMLM | SUG |
| HD_18JY | AX-111768401 | 2A | 605590887 | 0.8282 | | 4.17E-10 | BLUPmrMLM | SIG |
| HD_18JY | AX-111116744 | 2A | 656942335 | -0.8829 | | 6.65E-08 | BLUPmrMLM | SIG |
| HD_18JY | AX-108921222 | 2B | 58722811 | -0.3042 | | 2.66E-05 | BLUPmrMLM | SUG |

| | | | | | | | | |
|---|---|---|---|---|---|---|---|---|
| HD_18JY | AX-109944761 | 2B | 290009644 | -0.6424 | | 4.80E-07 | BLUPmrMLM | SUG |
| HD_18JY | AX-111956072 | 2D | 34428838 | 0.7513 | | 1.40E-06 | GEMMA | SUG |
| HD_18JY | AX-94552380 | 2D | 406149072 | -4.1315 | | 1.21E-06 | EMMAX | SUG |
| HD_18JY | AX-94738789 | 3A | 13413518 | -0.5985 | | 4.17E-06 | BLUPmrMLM | SUG |
| HD_18JY | AX-110048135 | 3A | 22733613 | 0.3118 | | 1.85E-04 | BLUPmrMLM | SUG |
| HD_18JY | AX-95204178 | 3A | 479782760 | 0.4283 | | 2.72E-05 | BLUPmrMLM | SUG |
| HD_18JY | AX-111648969 | 3A | 691218716 | -0.5049 | | 2.32E-07 | BLUPmrMLM | SUG |
| HD_18JY | AX-110491100 | 3A | 705251779 | -2.0106 | | 3.91E-07 | EMMAX | SUG |
| HD_18JY | AX-110491100 | 3A | 705251779 | 2.0157 | | 1.76E-07 | GEMMA | SUG |
| HD_18JY | AX-108941177 | 3A | 705259848 | -2.0025 | | 7.58E-08 | EMMAX | SIG |
| HD_18JY | AX-108941177 | 3A | 705259848 | 2.0040 | | 3.43E-08 | GEMMA | SIG |
| HD_18JY | AX-111695263 | 3A | 705274739 | -1.8976 | | 9.30E-07 | EMMAX | SUG |
| HD_18JY | AX-111695263 | 3A | 705274739 | 1.9041 | | 4.01E-07 | GEMMA | SUG |
| HD_18JY | AX-110976174 | 3A | 705274782 | -3.8425 | | 2.01E-07 | EMMAX | SUG |
| HD_18JY | AX-110976174 | 3A | 705274782 | 3.8459 | | 9.47E-08 | GEMMA | SIG |
| HD_18JY | AX-110976174 | 3A | 705274782 | -1.8910 | | 1.32E-76 | IIIVmrMLM | SIG |
| HD_18JY | AX-110484538 | 3A | 705296840 | -4.2669 | | 4.00E-07 | EMMAX | SUG |
| HD_18JY | AX-110484538 | 3A | 705296840 | 4.2486 | | 2.26E-07 | GEMMA | SUG |
| HD_18JY | AX-110641031 | 3B | 7601649 | -0.6328 | | 1.79E-06 | EMMAX | SUG |
| HD_18JY | AX-110641031 | 3B | 7601649 | 0.6332 | | 1.45E-06 | GEMMA | SUG |
| HD_18JY | AX-110364271 | 3B | 7610526 | 0.6122 | | 1.65E-06 | EMMAX | SUG |
| HD_18JY | AX-110364271 | 3B | 7610526 | -0.6143 | | 1.32E-06 | GEMMA | SUG |
| HD_18JY | AX-108865439 | 3B | 7614235 | 0.6271 | | 1.51E-06 | EMMAX | SUG |
| HD_18JY | AX-108865439 | 3B | 7614235 | -0.6284 | | 1.17E-06 | GEMMA | SUG |
| HD_18JY | AX-111612892 | 3B | 7615861 | 0.4681 | | 1.13E-08 | BLUPmrMLM | SIG |
| HD_18JY | AX-111612892 | 3B | 7615861 | -0.6495 | | 1.04E-06 | EMMAX | SUG |
| HD_18JY | AX-111612892 | 3B | 7615861 | 0.6507 | | 8.31E-07 | GEMMA | SUG |
| HD_18JY | AX-110585859 | 3B | 39795435 | 0.3014 | | 1.37E-04 | BLUPmrMLM | SUG |
| HD_18JY | AX-109022331 | 3D | 276538349 | -9.2748 | | 6.81E-07 | EMMAX | SUG |
| HD_18JY | AX-109270564 | 3D | 283965882 | -9.2748 | | 6.81E-07 | EMMAX | SUG |
| HD_18JY | AX-110471609 | 3D | 604263446 | -0.4391 | | 2.25E-07 | BLUPmrMLM | SUG |
| HD_18JY | AX-109849572 | 4A | 465273521 | 0.8652 | 1.1657 | 2.42E-20 | IIIVmrMLM | SIG |
| HD_18JY | AX-109816200 | 4A | 649204800 | -0.6187 | | 1.77E-06 | EMMAX | SUG |
| HD_18JY | AX-109816200 | 4A | 649204800 | 0.6162 | | 9.86E-07 | GEMMA | SUG |
| HD_18JY | AX-109817479 | 4A | 657098867 | -0.6138 | | 2.09E-06 | EMMAX | SUG |
| HD_18JY | AX-109817479 | 4A | 657098867 | 0.6096 | | 1.32E-06 | GEMMA | SUG |
| HD_18JY | AX-110468849 | 4A | 668659171 | -1.0797 | -4.2981 | 7.61E-43 | IIIVmrMLM | SIG |
| HD_18JY | AX-94928833 | 4B | 604204038 | -9.2748 | | 6.81E-07 | EMMAX | SUG |
| HD_18JY | AX-110048398 | 5A | 529719353 | -0.4542 | | 4.89E-06 | BLUPmrMLM | SUG |
| HD_18JY | AX-109271377 | 5A | 702235368 | 0.2955 | | 1.68E-04 | BLUPmrMLM | SUG |
| HD_18JY | AX-109522213 | 5B | 32626571 | 0.4900 | | 2.90E-07 | BLUPmrMLM | SUG |
| HD_18JY | AX-110966766 | 5B | 492001241 | 0.3639 | | 1.05E-04 | BLUPmrMLM | SUG |
| HD_18JY | AX-109933949 | 5B | 526667566 | 0.8420 | | 2.46E-07 | BLUPmrMLM | SUG |
| HD_18JY | AX-108755254 | 6B | 460094100 | -0.4834 | | 1.02E-08 | BLUPmrMLM | SIG |

| | | | | | | | | |
|---|---|---|---|---|---|---|---|---|
| HD_18JY | AX-94425880 | 6B | 685669879 | 5.4583 | | 1.69E-06 | EMMAX | SUG |
| HD_18JY | AX-86178536 | 6B | 716211158 | 0.3605 | | 2.16E-07 | BLUPmrMLM | SUG |
| HD_18JY | AX-110898479 | 6D | 78533136 | -0.6593 | | 2.38E-07 | BLUPmrMLM | SUG |
| HD_18JY | AX-110021490 | 7A | 47697933 | -0.7321 | | 5.33E-07 | EMMAX | SUG |
| HD_18JY | AX-110021490 | 7A | 47697933 | 0.7307 | | 3.30E-07 | GEMMA | SUG |
| HD_18JY | AX-111183447 | 7A | 47698107 | -0.7315 | | 5.78E-07 | EMMAX | SUG |
| HD_18JY | AX-111183447 | 7A | 47698107 | 0.7300 | | 3.61E-07 | GEMMA | SUG |
| HD_18JY | AX-111524711 | 7A | 48562501 | -1.4228 | | 1.31E-06 | GEMMA | SUG |
| HD_18JY | AX-108921785 | 7A | 48624311 | -1.3948 | | 1.19E-06 | GEMMA | SUG |
| HD_18JY | AX-109296563 | 7A | 444924900 | -0.4485 | | 1.13E-06 | BLUPmrMLM | SUG |
| HD_18JY | AX-108914113 | 7B | 553640645 | -9.2748 | | 6.81E-07 | EMMAX | SUG |
| HD_18JY | AX-95191225 | 7D | 31435113 | -9.2748 | | 6.81E-07 | EMMAX | SUG |
| HD_18JY | AX-109767598 | 7D | 476504044 | 1.1517 | -4.2511 | 7.30E-39 | IIIVmrMLM | SIG |
| HD_18XX | AX-111660890 | 1A | 6417405 | 0.0234 | 0.7425 | 2.49E-10 | IIIVmrMLM | SIG |
| HD_18XX | AX-109628437 | 1A | 482221054 | 0.1099 | 0.5609 | 4.41E-06 | IIIVmrMLM | SUG |
| HD_18XX | AX-109276285 | 1A | 507078569 | 0.0385 | -1.6654 | 1.42E-04 | IIIVmrMLM | SUG |
| HD_18XX | AX-108958046 | 1A | 532670133 | 0.7188 | 3.4402 | 1.38E-60 | IIIVmrMLM | SIG |
| HD_18XX | AX-109928662 | 1B | 63083627 | -0.1394 | -0.7978 | 1.45E-04 | IIIVmrMLM | SUG |
| HD_18XX | AX-108949682 | 1B | 66755144 | 0.1964 | -0.8009 | 1.31E-12 | IIIVmrMLM | SIG |
| HD_18XX | AX-109634576 | 1B | 169711402 | -1.0598 | | 1.96E-06 | BLUPmrMLM | SUG |
| HD_18XX | AX-111193379 | 1D | 10985400 | 0.6857 | -0.0364 | 4.03E-34 | IIIVmrMLM | SIG |
| HD_18XX | AX-111913284 | 1D | 21640736 | 0.5399 | | 2.34E-09 | BLUPmrMLM | SIG |
| HD_18XX | AX-109857857 | 1D | 488348964 | -0.3833 | | 1.21E-05 | BLUPmrMLM | SUG |
| HD_18XX | AX-111538390 | 2A | 59637740 | -0.4359 | | 3.03E-05 | BLUPmrMLM | SUG |
| HD_18XX | AX-109575515 | 2A | 65395975 | 0.3642 | | 6.65E-05 | BLUPmrMLM | SUG |
| HD_18XX | AX-110000162 | 2A | 524815534 | 0.4039 | -0.1949 | 6.05E-13 | IIIVmrMLM | SIG |
| HD_18XX | AX-111116744 | 2A | 656942335 | 0.0823 | -0.6068 | 1.10E-05 | IIIVmrMLM | SUG |
| HD_18XX | AX-110948179 | 2A | 738974772 | -0.2349 | 0.8745 | 1.39E-09 | IIIVmrMLM | SIG |
| HD_18XX | AX-111589712 | 2B | 22781589 | -0.5458 | | 1.90E-05 | BLUPmrMLM | SUG |
| HD_18XX | AX-109861243 | 2B | 129214745 | 0.0562 | -1.876 | 2.68E-04 | IIIVmrMLM | SUG |
| HD_18XX | AX-109887392 | 2B | 291603441 | -6.1533 | | 3.80E-08 | FarmCPU | SIG |
| HD_18XX | AX-110511037 | 2B | 444179967 | 0.5856 | | 2.95E-05 | BLUPmrMLM | SUG |
| HD_18XX | AX-111607493 | 2B | 699289728 | -0.0694 | -1.5153 | 4.07E-05 | IIIVmrMLM | SUG |
| HD_18XX | AX-110398503 | 2B | 787954586 | 2.7192 | | 4.65E-07 | EMMAX | SUG |
| HD_18XX | AX-110398503 | 2B | 787954586 | -2.7389 | | 2.52E-07 | GEMMA | SUG |
| HD_18XX | AX-108904634 | 2B | 796344276 | 0.0422 | -1.0928 | 4.60E-09 | IIIVmrMLM | SIG |
| HD_18XX | AX-110886525 | 2D | 59334830 | -12.9110 | | 1.53E-06 | EMMAX | SUG |
| HD_18XX | AX-94976683 | 2D | 569949805 | 0.1283 | -0.5403 | 7.04E-08 | IIIVmrMLM | SIG |
| HD_18XX | AX-94686674 | 2D | 600161540 | 0.2775 | | 6.59E-05 | BLUPmrMLM | SUG |
| HD_18XX | AX-111680059 | 3A | 20797121 | -0.1025 | 1.9406 | 1.28E-04 | IIIVmrMLM | SUG |
| HD_18XX | AX-110949095 | 3A | 651640332 | 0.1905 | -0.9862 | 6.80E-11 | IIIVmrMLM | SIG |
| HD_18XX | AX-108888346 | 3A | 652775915 | -0.4171 | | 3.70E-05 | BLUPmrMLM | SUG |
| HD_18XX | AX-110491100 | 3A | 705251779 | -3.1001 | | 9.17E-12 | EMMAX | SIG |
| HD_18XX | AX-110491100 | 3A | 705251779 | 3.0844 | | 3.67E-12 | GEMMA | SIG |

| | | | | | | | | |
|---|---|---|---|---|---|---|---|---|
| HD_18XX | AX-108941177 | 3A | 705259848 | -2.7909 | | 8.72E-11 | EMMAX | SIG |
| HD_18XX | AX-108941177 | 3A | 705259848 | 2.7698 | | 4.17E-11 | GEMMA | SIG |
| HD_18XX | AX-111695263 | 3A | 705274739 | -2.9109 | | 4.28E-11 | EMMAX | SIG |
| HD_18XX | AX-111695263 | 3A | 705274739 | 2.9038 | | 1.60E-11 | GEMMA | SIG |
| HD_18XX | AX-110976174 | 3A | 705274782 | -5.4649 | | 1.76E-10 | EMMAX | SIG |
| HD_18XX | AX-110976174 | 3A | 705274782 | 5.3971 | | 1.13E-10 | GEMMA | SIG |
| HD_18XX | AX-110484538 | 3A | 705296840 | -7.0380 | | 3.93E-13 | EMMAX | SIG |
| HD_18XX | AX-110484538 | 3A | 705296840 | 6.9876 | | 1.25E-13 | GEMMA | SIG |
| HD_18XX | AX-110526487 | 3A | 705298427 | -1.5053 | | 6.30E-07 | EMMAX | SUG |
| HD_18XX | AX-110526487 | 3A | 705298427 | 1.5027 | | 3.79E-07 | GEMMA | SUG |
| HD_18XX | AX-109510662 | 3A | 705311544 | -1.3821 | | 5.36E-07 | EMMAX | SUG |
| HD_18XX | AX-109510662 | 3A | 705311544 | 1.3678 | | 4.44E-07 | GEMMA | SUG |
| HD_18XX | AX-108929886 | 3B | 326980 | -0.2809 | -1.098 | 1.24E-10 | IIIVmrMLM | SIG |
| HD_18XX | AX-111129787 | 3B | 140859817 | -0.2985 | -0.7363 | 6.13E-20 | IIIVmrMLM | SIG |
| HD_18XX | AX-109396226 | 3B | 212216397 | -0.0239 | -1.8081 | 1.07E-12 | IIIVmrMLM | SIG |
| HD_18XX | AX-94658057 | 3B | 386595171 | 0.3214 | | 1.14E-04 | BLUPmrMLM | SUG |
| HD_18XX | AX-108881562 | 3B | 463233881 | -0.3315 | | 9.89E-05 | BLUPmrMLM | SUG |
| HD_18XX | AX-110450538 | 3B | 492567942 | -12.9110 | | 1.53E-06 | EMMAX | SUG |
| HD_18XX | AX-111489277 | 3B | 625336740 | 0.4552 | | 4.74E-18 | IIIVmrMLM | SIG |
| HD_18XX | AX-110914777 | 3B | 635196909 | -2.2302 | | 1.00E-06 | GEMMA | SUG |
| HD_18XX | AX-108918805 | 3B | 655638189 | 3.9895 | | 2.48E-07 | EMMAX | SUG |
| HD_18XX | AX-108918805 | 3B | 655638189 | -4.0341 | | 1.33E-07 | GEMMA | SUG |
| HD_18XX | AX-109415956 | 3B | 810332933 | 3.3757 | | 1.92E-07 | EMMAX | SUG |
| HD_18XX | AX-109415956 | 3B | 810332933 | -3.2433 | | 2.34E-07 | GEMMA | SUG |
| HD_18XX | AX-110416128 | 3B | 810334958 | 6.7513 | | 1.92E-07 | EMMAX | SUG |
| HD_18XX | AX-108845796 | 3B | 810335131 | 6.7513 | | 1.92E-07 | EMMAX | SUG |
| HD_18XX | AX-108920352 | 3B | 810358775 | 6.7513 | | 1.92E-07 | EMMAX | SUG |
| HD_18XX | AX-109376714 | 3B | 810448196 | 3.3757 | | 1.92E-07 | EMMAX | SUG |
| HD_18XX | AX-109376714 | 3B | 810448196 | -3.2433 | | 2.34E-07 | GEMMA | SUG |
| HD_18XX | AX-108783932 | 3B | 810528102 | 6.7513 | | 1.92E-07 | EMMAX | SUG |
| HD_18XX | AX-111565872 | 3B | 810528155 | 3.3757 | | 1.92E-07 | EMMAX | SUG |
| HD_18XX | AX-111565872 | 3B | 810528155 | -3.2433 | | 2.34E-07 | GEMMA | SUG |
| HD_18XX | AX-110080744 | 3B | 810528192 | 5.0898 | | 1.75E-06 | EMMAX | SUG |
| HD_18XX | AX-111196578 | 3B | 810528242 | 6.2904 | | 1.87E-07 | EMMAX | SUG |
| HD_18XX | AX-108782991 | 3B | 810544591 | 6.7513 | | 1.92E-07 | EMMAX | SUG |
| HD_18XX | AX-111131199 | 3B | 810564183 | 6.7513 | | 1.92E-07 | EMMAX | SUG |
| HD_18XX | AX-109017734 | 3B | 810565289 | 6.7513 | | 1.92E-07 | EMMAX | SUG |
| HD_18XX | AX-110060618 | 3B | 810622955 | 6.7513 | | 1.92E-07 | EMMAX | SUG |
| HD_18XX | AX-108757746 | 3B | 810625226 | 6.7513 | | 1.92E-07 | EMMAX | SUG |
| HD_18XX | AX-111657522 | 3B | 810783132 | 6.7513 | | 1.92E-07 | EMMAX | SUG |
| HD_18XX | AX-108903892 | 3B | 810902194 | 6.7513 | | 1.92E-07 | EMMAX | SUG |
| HD_18XX | AX-111193358 | 3D | 51785648 | 5.3040 | | 1.84E-06 | EMMAX | SUG |
| HD_18XX | AX-111026826 | 3D | 560126633 | 0.6681 | | 2.64E-07 | BLUPmrMLM | SUG |
| HD_18XX | AX-109188381 | 3D | 604693879 | -0.3640 | | 5.48E-05 | BLUPmrMLM | SUG |

| | | | | | | | | |
|---|---|---|---|---|---|---|---|---|
| HD_18XX | AX-94385414 | 4A | 26465663 | -12.9110 | | 1.53E-06 | EMMAX | SUG |
| HD_18XX | AX-111256892 | 4A | 342811372 | 0.1285 | -3.1385 | 2.58E-12 | IIIVmrMLM | SIG |
| HD_18XX | AX-109855028 | 4A | 452086547 | -0.5038 | | 1.56E-06 | BLUPmrMLM | SUG |
| HD_18XX | AX-109816200 | 4A | 649204800 | 0.3314 | | 3.88E-06 | BLUPmrMLM | SUG |
| HD_18XX | AX-109816200 | 4A | 649204800 | -0.2820 | 0.6316 | 2.45E-07 | IIIVmrMLM | SUG |
| HD_18XX | AX-110984523 | 4B | 76044642 | 4.6915 | | 1.55E-06 | EMMAX | SUG |
| HD_18XX | AX-110461577 | 4B | 560942256 | 0.4852 | | 3.06E-06 | BLUPmrMLM | SUG |
| HD_18XX | AX-111643602 | 4D | 309989236 | -12.9110 | | 1.53E-06 | EMMAX | SUG |
| HD_18XX | AX-109522149 | 5A | 417052887 | 0.0065 | -4.3942 | 6.80E-06 | IIIVmrMLM | SUG |
| HD_18XX | AX-95631615 | 5A | 512234990 | -0.2810 | | 8.15E-06 | BLUPmrMLM | SUG |
| HD_18XX | AX-111689312 | 5A | 547056411 | -1.4001 | | 1.90E-06 | EMMAX | SUG |
| HD_18XX | AX-111689312 | 5A | 547056411 | 1.4074 | | 9.90E-07 | GEMMA | SUG |
| HD_18XX | AX-111689312 | 5A | 547056411 | -0.7231 | 0.5875 | 1.90E-38 | IIIVmrMLM | SIG |
| HD_18XX | AX-94987862 | 5A | 596369178 | 4.0590 | | 1.31E-06 | EMMAX | SUG |
| HD_18XX | AX-94987862 | 5A | 596369178 | -4.0510 | | 8.37E-07 | GEMMA | SUG |
| HD_18XX | AX-108868375 | 5B | 319333756 | -0.3774 | | 5.25E-06 | BLUPmrMLM | SUG |
| HD_18XX | AX-109835655 | 5B | 513608526 | -0.4395 | | 2.31E-05 | BLUPmrMLM | SUG |
| HD_18XX | AX-109821989 | 5B | 565424439 | 0.4095 | -1.7822 | 8.62E-19 | IIIVmrMLM | SIG |
| HD_18XX | AX-111528735 | 5B | 574063657 | 2.7583 | | 4.20E-12 | EMMAX | SIG |
| HD_18XX | AX-111528735 | 5B | 574063657 | -2.7097 | | 3.46E-12 | GEMMA | SIG |
| HD_18XX | AX-108987083 | 5B | 575972456 | 3.7683 | | 3.80E-07 | EMMAX | SUG |
| HD_18XX | AX-108987083 | 5B | 575972456 | -3.7148 | | 3.01E-07 | GEMMA | SUG |
| HD_18XX | AX-110595998 | 5B | 576142349 | -2.3099 | | 1.57E-18 | BLUPmrMLM | SIG |
| HD_18XX | AX-110595998 | 5B | 576142349 | 2.6707 | | 1.27E-10 | EMMAX | SIG |
| HD_18XX | AX-110595998 | 5B | 576142349 | -2.6358 | | 7.14E-11 | GEMMA | SIG |
| HD_18XX | AX-110595998 | 5B | 576142349 | 1.6704 | 4.3773 | ####### | IIIVmrMLM | SIG |
| HD_18XX | AX-109547794 | 5B | 585025719 | -0.9723 | -1.5849 | 1.17E-68 | IIIVmrMLM | SIG |
| HD_18XX | AX-109008501 | 5B | 587803667 | 2.6814 | | 9.25E-07 | EMMAX | SUG |
| HD_18XX | AX-109008501 | 5B | 587803667 | -2.6719 | | 6.48E-07 | GEMMA | SUG |
| HD_18XX | AX-109396197 | 5B | 692713178 | 0.3895 | | 4.13E-05 | BLUPmrMLM | SUG |
| HD_18XX | AX-111386686 | 5D | 95927904 | -1.3998 | | 3.80E-07 | FarmCPU | SUG |
| HD_18XX | AX-110371497 | 5D | 453016646 | 1.2226 | | 1.70E-06 | EMMAX | SUG |
| HD_18XX | AX-110371497 | 5D | 453016646 | -1.2398 | | 8.58E-07 | GEMMA | SUG |
| HD_18XX | AX-111216464 | 5D | 453827161 | 1.2226 | | 1.70E-06 | EMMAX | SUG |
| HD_18XX | AX-111216464 | 5D | 453827161 | -1.2398 | | 8.58E-07 | GEMMA | SUG |
| HD_18XX | AX-110546225 | 5D | 454111662 | 1.2086 | | 1.78E-06 | EMMAX | SUG |
| HD_18XX | AX-110546225 | 5D | 454111662 | -1.2254 | | 9.13E-07 | GEMMA | SUG |
| HD_18XX | AX-111497172 | 5D | 454427605 | -0.3344 | -0.7333 | 6.02E-11 | IIIVmrMLM | SIG |
| HD_18XX | AX-111515523 | 5D | 456032452 | 1.1932 | | 2.06E-06 | EMMAX | SUG |
| HD_18XX | AX-111515523 | 5D | 456032452 | -1.2006 | | 1.12E-06 | GEMMA | SUG |
| HD_18XX | AX-110783573 | 5D | 459430740 | 2.3722 | | 1.97E-06 | EMMAX | SUG |
| HD_18XX | AX-110783573 | 5D | 459430740 | -2.3843 | | 1.10E-06 | GEMMA | SUG |
| HD_18XX | AX-111917011 | 5D | 462565152 | 1.3856 | | 7.60E-08 | EMMAX | SIG |
| HD_18XX | AX-111917011 | 5D | 462565152 | -1.3863 | | 4.17E-08 | GEMMA | SIG |

| | | | | | | | | |
|---|---|---|---|---|---|---|---|---|
| HD_18XX | AX-111234923 | 5D | 463118750 | -1.7561 | | 7.18E-27 | BLUPmrMLM | SIG |
| HD_18XX | AX-111234923 | 5D | 463118750 | 1.4099 | | 2.61E-08 | EMMAX | SIG |
| HD_18XX | AX-111234923 | 5D | 463118750 | -1.4115 | | 1.35E-08 | GEMMA | SIG |
| HD_18XX | AX-110044748 | 5D | 464108412 | 2.9396 | | 2.59E-08 | EMMAX | SIG |
| HD_18XX | AX-110044748 | 5D | 464108412 | -2.9386 | | 1.47E-08 | GEMMA | SIG |
| HD_18XX | AX-111175690 | 5D | 464415388 | 1.7092 | | 3.34E-10 | EMMAX | SIG |
| HD_18XX | AX-111175690 | 5D | 464415388 | -1.7079 | | 1.85E-10 | GEMMA | SIG |
| HD_18XX | AX-109360916 | 5D | 469286340 | 4.6652 | | 3.21E-16 | EMMAX | SIG |
| HD_18XX | AX-109360916 | 5D | 469286340 | -4.6587 | | 1.05E-16 | GEMMA | SIG |
| HD_18XX | AX-109360916 | 5D | 469286340 | 1.6512 | | ####### | IIIVmrMLM | SIG |
| HD_18XX | AX-109838967 | 6A | 24125213 | -0.1858 | 0.7267 | 1.78E-07 | IIIVmrMLM | SUG |
| HD_18XX | AX-94523368 | 6A | 584913052 | -0.4855 | | 2.09E-05 | BLUPmrMLM | SUG |
| HD_18XX | AX-94866773 | 6A | 611661195 | -0.1380 | -1.6043 | 5.72E-11 | IIIVmrMLM | SIG |
| HD_18XX | AX-95153605 | 6B | 31516855 | -0.4265 | | 3.90E-16 | IIIVmrMLM | SIG |
| HD_18XX | AX-111549096 | 6B | 708067991 | -0.3431 | -0.411 | 1.97E-10 | IIIVmrMLM | SIG |
| HD_18XX | AX-94801617 | 6D | 454588858 | 0.8675 | | 2.75E-05 | BLUPmrMLM | SUG |
| HD_18XX | AX-111075771 | 7A | 167267363 | 2.7872 | | 1.01E-07 | FarmCPU | SIG |
| HD_18XX | AX-109925674 | 7A | 211414392 | -0.0489 | -4.0232 | 2.23E-07 | IIIVmrMLM | SUG |
| HD_18XX | AX-111776821 | 7A | 451838535 | -0.1414 | -0.9759 | 1.61E-08 | IIIVmrMLM | SIG |
| HD_18XX | AX-110179029 | 7A | 550612870 | 0.4982 | | 8.52E-05 | BLUPmrMLM | SUG |
| HD_18XX | AX-109404223 | 7B | 329788042 | 0.0533 | 1.4655 | 1.49E-05 | IIIVmrMLM | SUG |
| HD_18XX | AX-111089471 | 7B | 638510858 | -0.3709 | | 7.15E-05 | BLUPmrMLM | SUG |
| HD_18XX | AX-109962190 | 7B | 664039785 | -0.0714 | -0.7155 | 8.07E-10 | IIIVmrMLM | SIG |
| HD_18XX | AX-110020643 | 7B | 678854267 | -0.3060 | | 2.64E-05 | BLUPmrMLM | SUG |
| HD_18XX | AX-89423300 | 7D | 19396105 | -0.3751 | | 1.39E-04 | BLUPmrMLM | SUG |
| HD_18XX | AX-110576947 | 7D | 65436205 | -1.2829 | | 1.15E-16 | BLUPmrMLM | SIG |
| HD_18XX | AX-110826147 | 7D | 65503559 | 1.0563 | 0.3899 | 2.47E-71 | IIIVmrMLM | SIG |
| HD_18XX | AX-94587603 | 7D | 102900636 | 0.2293 | -1.1633 | 4.15E-09 | IIIVmrMLM | SIG |
| HD_19XX | AX-109856872 | 1A | 4062943 | 0.4566 | | 6.34E-05 | BLUPmrMLM | SUG |
| HD_19XX | AX-108863576 | 1A | 156514000 | 0.4912 | | 9.69E-05 | BLUPmrMLM | SUG |
| HD_19XX | AX-110534762 | 1A | 326338261 | -0.0130 | -1.9968 | 9.70E-08 | IIIVmrMLM | SIG |
| HD_19XX | AX-111483492 | 1A | 465427783 | 0.0711 | 3.0858 | 6.37E-09 | IIIVmrMLM | SIG |
| HD_19XX | AX-109980625 | 1B | 409304929 | -0.4326 | -1.5021 | 1.04E-12 | IIIVmrMLM | SIG |
| HD_19XX | AX-111026585 | 1B | 547581461 | 0.1556 | -3.196 | 4.49E-08 | IIIVmrMLM | SIG |
| HD_19XX | AX-109868897 | 1D | 15876375 | 0.1588 | -1.5512 | 7.80E-10 | IIIVmrMLM | SIG |
| HD_19XX | AX-110541159 | 1D | 16237393 | -0.8493 | | 1.60E-07 | BLUPmrMLM | SUG |
| HD_19XX | AX-109496774 | 2A | 184915446 | 0.1561 | 1.5903 | 5.58E-08 | IIIVmrMLM | SIG |
| HD_19XX | AX-109831401 | 2A | 292129504 | -0.0832 | 2.0054 | 2.71E-06 | IIIVmrMLM | SUG |
| HD_19XX | AX-110492350 | 2A | 755054872 | 0.7757 | 0.1751 | 2.47E-22 | IIIVmrMLM | SIG |
| HD_19XX | AX-110907953 | 2B | 19137033 | 0.9432 | | 2.93E-06 | BLUPmrMLM | SUG |
| HD_19XX | AX-112288145 | 2B | 121367425 | 0.7393 | | 1.40E-06 | BLUPmrMLM | SUG |
| HD_19XX | AX-110515578 | 2B | 275364099 | 3.3741 | | 8.00E-07 | EMMAX | SUG |
| HD_19XX | AX-111535697 | 2B | 302148498 | 6.7483 | | 8.00E-07 | EMMAX | SUG |
| HD_19XX | AX-108802600 | 2B | 597011824 | -0.1019 | -1.8519 | 6.01E-06 | IIIVmrMLM | SUG |

| | | | | | | | | |
|---|---|---|---|---|---|---|---|---|
| HD_19XX | AX-111271294 | 2B | 767155693 | -0.3127 | -2.8615 | 1.49E-09 | IIIVmrMLM | SIG |
| HD_19XX | AX-110928030 | 2B | 779303114 | -1.3166 | | 4.52E-07 | FarmCPU | SUG |
| HD_19XX | AX-110045144 | 2B | 779303442 | -1.3309 | | 4.21E-07 | FarmCPU | SUG |
| HD_19XX | AX-110048628 | 2B | 779855175 | -0.0313 | -6.3317 | 7.01E-04 | IIIVmrMLM | SUG |
| HD_19XX | AX-110615313 | 2D | 616576649 | -0.9324 | | 3.66E-07 | BLUPmrMLM | SUG |
| HD_19XX | AX-111695263 | 3A | 705274739 | 2.3571 | | 1.07E-06 | GEMMA | SUG |
| HD_19XX | AX-94598344 | 3B | 59032971 | 0.3594 | | 1.30E-07 | IIIVmrMLM | SUG |
| HD_19XX | AX-94616860 | 3B | 222192671 | 0.6098 | 3.1543 | 4.45E-30 | IIIVmrMLM | SIG |
| HD_19XX | AX-109823264 | 3B | 651679343 | -0.5434 | | 3.57E-15 | IIIVmrMLM | SIG |
| HD_19XX | AX-108781140 | 4A | 5814680 | 0.5390 | | 6.35E-06 | BLUPmrMLM | SUG |
| HD_19XX | AX-109519857 | 4A | 47342278 | -0.1255 | -3.6392 | 1.57E-07 | IIIVmrMLM | SUG |
| HD_19XX | AX-110381551 | 4A | 602894161 | -0.6122 | -0.9565 | 8.46E-18 | IIIVmrMLM | SIG |
| HD_19XX | AX-109817340 | 4A | 717300838 | 0.5004 | | 3.15E-06 | BLUPmrMLM | SUG |
| HD_19XX | AX-110021771 | 4B | 1248675 | 0.1818 | -3.8708 | 5.66E-08 | IIIVmrMLM | SIG |
| HD_19XX | AX-109844329 | 4B | 422778745 | -0.2847 | 1.4558 | 5.11E-07 | IIIVmrMLM | SUG |
| HD_19XX | AX-110320914 | 4D | 44529297 | 1.0285 | | 8.22E-07 | FarmCPU | SUG |
| HD_19XX | AX-111963865 | 4D | 45927347 | 1.0286 | | 1.62E-07 | FarmCPU | SUG |
| HD_19XX | AX-109580054 | 5A | 35812034 | -0.5518 | 0.2775 | 1.32E-13 | IIIVmrMLM | SIG |
| HD_19XX | AX-109830637 | 5A | 504285711 | -0.1160 | 1.1237 | 1.81E-07 | IIIVmrMLM | SUG |
| HD_19XX | AX-110470925 | 5A | 547357165 | -0.0375 | 2.93 | 2.88E-09 | IIIVmrMLM | SIG |
| HD_19XX | AX-108836029 | 5A | 559135112 | 0.3502 | -2.1614 | 3.50E-09 | IIIVmrMLM | SIG |
| HD_19XX | AX-94487274 | 5A | 573755065 | -0.3422 | | 4.86E-07 | IIIVmrMLM | SUG |
| HD_19XX | AX-111249882 | 5A | 623379521 | -0.5357 | | 1.14E-05 | BLUPmrMLM | SUG |
| HD_19XX | AX-110926985 | 5B | 11485073 | 0.0252 | -1.4013 | 4.27E-13 | IIIVmrMLM | SIG |
| HD_19XX | AX-111127637 | 5B | 242223397 | -0.0668 | -1.7239 | 7.81E-05 | IIIVmrMLM | SUG |
| HD_19XX | AX-110972217 | 5B | 458727558 | 0.2823 | -6.3317 | 1.65E-07 | IIIVmrMLM | SUG |
| HD_19XX | AX-108960709 | 5D | 162761169 | -0.2192 | -5.454 | 4.39E-16 | IIIVmrMLM | SIG |
| HD_19XX | AX-109188063 | 5D | 461108082 | -0.3568 | | 6.91E-05 | BLUPmrMLM | SUG |
| HD_19XX | AX-111689944 | 5D | 466287415 | 0.7675 | -9.4787 | 5.67E-32 | IIIVmrMLM | SIG |
| HD_19XX | AX-109360916 | 5D | 469286340 | 1.0338 | | 2.71E-45 | IIIVmrMLM | SIG |
| HD_19XX | AX-110696524 | 6A | 345209734 | -0.1739 | 1.9175 | 5.72E-05 | IIIVmrMLM | SUG |
| HD_19XX | AX-95019685 | 6A | 596522253 | -0.3984 | 1.112 | 5.49E-12 | IIIVmrMLM | SIG |
| HD_19XX | AX-111535116 | 6B | 84670424 | -0.1999 | -5.7558 | 1.73E-12 | IIIVmrMLM | SIG |
| HD_19XX | AX-94688060 | 6B | 247002932 | 0.0476 | 1.129 | 2.28E-15 | IIIVmrMLM | SIG |
| HD_19XX | AX-89669056 | 6B | 573572832 | -0.3835 | -1.37 | 5.59E-08 | IIIVmrMLM | SIG |
| HD_19XX | AX-109246594 | 6D | 259031572 | -0.6102 | | 1.47E-05 | BLUPmrMLM | SUG |
| HD_19XX | AX-109295112 | 7A | 35691569 | 0.0503 | -2.5226 | 3.20E-06 | IIIVmrMLM | SUG |
| HD_19XX | AX-109475180 | 7A | 232455842 | 0.6497 | | 2.42E-05 | BLUPmrMLM | SUG |
| HD_19XX | AX-109437286 | 7A | 496821001 | 0.1412 | -1.355 | 1.12E-06 | IIIVmrMLM | SUG |
| HD_19XX | AX-108848825 | 7A | 537663959 | 0.9493 | | 2.81E-07 | BLUPmrMLM | SUG |
| HD_19XX | AX-94398361 | 7A | 617684953 | -0.8429 | | 5.97E-07 | BLUPmrMLM | SUG |
| HD_19XX | AX-111063800 | 7A | 681607244 | 0.0566 | 0.9447 | 3.67E-10 | IIIVmrMLM | SIG |
| HD_19XX | AX-111074941 | 7A | 733371574 | -0.3202 | -3.7406 | 1.09E-09 | IIIVmrMLM | SIG |
| HD_19XX | AX-94946714 | 7B | 19120874 | -0.4182 | | 1.37E-04 | BLUPmrMLM | SUG |

| | | | | | | | | |
|---|---|---|---|---|---|---|---|---|
| HD_19XX | AX-110559866 | 7B | 227591695 | -0.0486 | -1.8994 | 2.25E-06 | IIIVmrMLM | SUG |
| HD_19XX | AX-108892290 | 7B | 552033864 | 0.0227 | -2.6839 | 1.35E-11 | IIIVmrMLM | SIG |
| HD_19XX | AX-108891067 | 7B | 628845066 | -0.1574 | 0.8856 | 4.07E-05 | IIIVmrMLM | SUG |
| HD_19XX | AX-111061288 | 7D | 66537922 | 0.8583 | | 2.25E-10 | BLUPmrMLM | SIG |
| HD_19XX | AX-111061288 | 7D | 66537922 | -0.7998 | 0.7019 | 9.17E-28 | IIIVmrMLM | SIG |
| HD_19XX | AX-109910604 | 7D | 347866686 | 2.8919 | 1.7554 | ####### | IIIVmrMLM | SIG |
| HD_19XX | AX-110152804 | 7D | 602042928 | -0.7129 | | 3.11E-05 | BLUPmrMLM | SUG |
| HD_BLUP | AX-110521336 | 1A | 289053265 | 0.2644 | | 7.92E-05 | BLUPmrMLM | SUG |
| HD_BLUP | AX-109388229 | 1A | 557956782 | -0.3037 | | 4.83E-07 | BLUPmrMLM | SUG |
| HD_BLUP | AX-108780402 | 1A | 564570992 | -1.0746 | | 2.13E-06 | FarmCPU | SUG |
| HD_BLUP | AX-109332672 | 1A | 565292715 | 0.1241 | 0.7221 | 1.08E-11 | IIIVmrMLM | SIG |
| HD_BLUP | AX-111678027 | 1B | 4250649 | 0.2273 | | 5.33E-05 | BLUPmrMLM | SUG |
| HD_BLUP | AX-109634576 | 1B | 169711402 | 0.2128 | -0.3032 | 1.72E-11 | IIIVmrMLM | SIG |
| HD_BLUP | AX-95105311 | 1B | 453524339 | 0.3043 | 0.3198 | 8.67E-19 | IIIVmrMLM | SIG |
| HD_BLUP | AX-108738407 | 1B | 676245447 | -0.2449 | | 1.14E-07 | BLUPmrMLM | SIG |
| HD_BLUP | AX-110967256 | 1B | 677617120 | -0.0695 | -0.5653 | 3.78E-16 | IIIVmrMLM | SIG |
| HD_BLUP | AX-108834310 | 1B | 687418381 | -0.2050 | | 3.49E-05 | BLUPmrMLM | SUG |
| HD_BLUP | AX-109924835 | 2A | 535801035 | 0.0069 | -0.4945 | 1.08E-11 | IIIVmrMLM | SIG |
| HD_BLUP | AX-94991562 | 2A | 712580484 | -0.4873 | | 3.71E-07 | BLUPmrMLM | SUG |
| HD_BLUP | AX-110546696 | 2B | 53000344 | -0.0288 | -1.0088 | 2.74E-06 | IIIVmrMLM | SUG |
| HD_BLUP | AX-111062205 | 2B | 93113255 | -0.3454 | | 1.34E-06 | BLUPmrMLM | SUG |
| HD_BLUP | AX-94652369 | 2B | 106129285 | -0.0257 | -1.5537 | 5.16E-08 | IIIVmrMLM | SIG |
| HD_BLUP | AX-111550170 | 2B | 118475520 | -0.1445 | -0.2582 | 7.34E-08 | IIIVmrMLM | SIG |
| HD_BLUP | AX-111741573 | 2B | 119241898 | 0.1526 | -0.5565 | 6.91E-15 | IIIVmrMLM | SIG |
| HD_BLUP | AX-109975037 | 2B | 172179290 | 0.3043 | | 4.19E-10 | BLUPmrMLM | SIG |
| HD_BLUP | AX-109975037 | 2B | 172179290 | -0.2427 | 0.0725 | 1.08E-11 | IIIVmrMLM | SIG |
| HD_BLUP | AX-110046269 | 2B | 251778087 | -2.5034 | | 3.58E-07 | FarmCPU | SUG |
| HD_BLUP | AX-111074147 | 2B | 404761193 | 0.1309 | -1.0859 | 2.11E-08 | IIIVmrMLM | SIG |
| HD_BLUP | AX-109856958 | 2B | 575869371 | 0.1901 | | 6.88E-05 | BLUPmrMLM | SUG |
| HD_BLUP | AX-94796377 | 2B | 731536579 | 0.3861 | | 2.07E-08 | BLUPmrMLM | SIG |
| HD_BLUP | AX-109836946 | 2D | 32795318 | -0.2615 | 0.0988 | 5.97E-14 | IIIVmrMLM | SIG |
| HD_BLUP | AX-94697254 | 2D | 33710135 | -1.3335 | | 2.03E-06 | FarmCPU | SUG |
| HD_BLUP | AX-110922927 | 2D | 405252279 | 0.1667 | 0.3658 | 1.21E-06 | IIIVmrMLM | SUG |
| HD_BLUP | AX-94502744 | 2D | 460205143 | -0.3861 | | 3.17E-05 | BLUPmrMLM | SUG |
| HD_BLUP | AX-111071407 | 3A | 32177342 | -0.1739 | -0.723 | 4.73E-19 | IIIVmrMLM | SIG |
| HD_BLUP | AX-109860768 | 3A | 652664372 | 0.1888 | 0.3205 | 9.34E-09 | IIIVmrMLM | SIG |
| HD_BLUP | AX-108888346 | 3A | 652775915 | -0.3093 | | 2.36E-06 | BLUPmrMLM | SUG |
| HD_BLUP | AX-110491100 | 3A | 705251779 | -1.9554 | | 2.02E-12 | EMMAX | SIG |
| HD_BLUP | AX-110491100 | 3A | 705251779 | 1.9525 | | 5.34E-13 | GEMMA | SIG |
| HD_BLUP | AX-108941177 | 3A | 705259848 | -1.8008 | | 7.63E-12 | EMMAX | SIG |
| HD_BLUP | AX-108941177 | 3A | 705259848 | 1.7940 | | 2.20E-12 | GEMMA | SIG |
| HD_BLUP | AX-111695263 | 3A | 705274739 | -1.8302 | | 1.57E-11 | EMMAX | SIG |
| HD_BLUP | AX-111695263 | 3A | 705274739 | 1.8357 | | 3.41E-12 | GEMMA | SIG |
| HD_BLUP | AX-110976174 | 3A | 705274782 | -3.5583 | | 1.02E-11 | EMMAX | SIG |

| | | | | | | | | |
|---|---|---|---|---|---|---|---|---|
| HD_BLUP | AX-110976174 | 3A | 705274782 | 3.5368 | | 3.66E-12 | GEMMA | SIG |
| HD_BLUP | AX-110484538 | 3A | 705296840 | -4.1657 | | 1.64E-12 | EMMAX | SIG |
| HD_BLUP | AX-110484538 | 3A | 705296840 | 4.1388 | | 4.71E-13 | GEMMA | SIG |
| HD_BLUP | AX-110526487 | 3A | 705298427 | -1.0107 | | 1.06E-07 | EMMAX | SIG |
| HD_BLUP | AX-110526487 | 3A | 705298427 | 1.0124 | | 5.13E-08 | GEMMA | SIG |
| HD_BLUP | AX-109510662 | 3A | 705311544 | -0.8746 | | 5.86E-07 | EMMAX | SUG |
| HD_BLUP | AX-109510662 | 3A | 705311544 | 0.8715 | | 4.11E-07 | GEMMA | SUG |
| HD_BLUP | AX-110390871 | 3B | 7252439 | 0.2046 | | 5.45E-05 | BLUPmrMLM | SUG |
| HD_BLUP | AX-95150475 | 3B | 378207527 | 0.2300 | | 2.80E-12 | IIIVmrMLM | SIG |
| HD_BLUP | AX-108862946 | 3B | 669390859 | 0.1759 | | 1.80E-05 | BLUPmrMLM | SUG |
| HD_BLUP | AX-111173784 | 3B | 736308550 | 0.2251 | | 7.72E-12 | IIIVmrMLM | SIG |
| HD_BLUP | AX-111546966 | 3B | 792721557 | 0.2990 | 0.3289 | 3.32E-18 | IIIVmrMLM | SIG |
| HD_BLUP | AX-95253038 | 3B | 793295044 | -0.1212 | -0.6417 | 2.09E-16 | IIIVmrMLM | SIG |
| HD_BLUP | AX-95209443 | 3D | 45938070 | -0.2139 | | 7.37E-11 | IIIVmrMLM | SIG |
| HD_BLUP | AX-111737155 | 3D | 608925693 | -0.0960 | 0.8337 | 1.08E-06 | IIIVmrMLM | SUG |
| HD_BLUP | AX-109420906 | 4A | 115634351 | -1.5516 | | 1.59E-06 | FarmCPU | SUG |
| HD_BLUP | AX-111256892 | 4A | 342811372 | -0.0459 | -3.0356 | 5.68E-24 | IIIVmrMLM | SIG |
| HD_BLUP | AX-111458403 | 4A | 620776679 | -0.2317 | -0.0603 | 2.57E-11 | IIIVmrMLM | SIG |
| HD_BLUP | AX-109817479 | 4A | 657098867 | -0.3081 | -0.0786 | 4.97E-19 | IIIVmrMLM | SIG |
| HD_BLUP | AX-95245458 | 4B | 74277174 | -1.9929 | | 2.09E-06 | GEMMA | SUG |
| HD_BLUP | AX-89438414 | 4B | 598261950 | -0.2214 | | 2.47E-05 | BLUPmrMLM | SUG |
| HD_BLUP | AX-110968968 | 4B | 637467667 | 0.1823 | -0.8657 | 9.13E-10 | IIIVmrMLM | SIG |
| HD_BLUP | AX-110604883 | 4D | 177489950 | -1.5516 | | 2.07E-06 | FarmCPU | SUG |
| HD_BLUP | AX-109903760 | 5A | 30056842 | 1.1313 | | 1.33E-06 | EMMAX | SUG |
| HD_BLUP | AX-109903760 | 5A | 30056842 | -1.1231 | | 1.34E-06 | GEMMA | SUG |
| HD_BLUP | AX-109408359 | 5A | 512124032 | 0.2300 | 0.5039 | 2.92E-13 | IIIVmrMLM | SIG |
| HD_BLUP | AX-111235500 | 5A | 556150580 | 0.4463 | | 1.89E-08 | BLUPmrMLM | SIG |
| HD_BLUP | AX-108836029 | 5A | 559135112 | 0.2998 | -0.8227 | 1.19E-19 | IIIVmrMLM | SIG |
| HD_BLUP | AX-111531227 | 5B | 38134179 | 0.1307 | 0.6479 | 7.14E-07 | IIIVmrMLM | SUG |
| HD_BLUP | AX-111455650 | 5B | 460217332 | 0.2421 | | 4.85E-05 | BLUPmrMLM | SUG |
| HD_BLUP | AX-109394866 | 5B | 513849166 | -0.5632 | | 3.81E-13 | BLUPmrMLM | SIG |
| HD_BLUP | AX-108831460 | 5B | 527056847 | 0.2701 | -0.4468 | 2.04E-20 | IIIVmrMLM | SIG |
| HD_BLUP | AX-110538089 | 5B | 539858157 | 0.3403 | 0.0126 | 3.22E-21 | IIIVmrMLM | SIG |
| HD_BLUP | AX-109821989 | 5B | 565424439 | -0.1880 | -0.8842 | 5.58E-11 | IIIVmrMLM | SIG |
| HD_BLUP | AX-109503531 | 5B | 692425935 | -0.2760 | | 1.66E-07 | BLUPmrMLM | SUG |
| HD_BLUP | AX-109503531 | 5B | 692425935 | 0.2170 | 0.0014 | 2.05E-09 | IIIVmrMLM | SIG |
| HD_BLUP | AX-108983843 | 5B | 712898248 | 0.0626 | -1.3013 | 4.15E-07 | IIIVmrMLM | SUG |
| HD_BLUP | AX-94727027 | 5D | 60249666 | 0.0847 | 0.4551 | 1.21E-10 | IIIVmrMLM | SIG |
| HD_BLUP | AX-110554978 | 5D | 166555781 | -0.1125 | -1.7626 | 2.29E-09 | IIIVmrMLM | SIG |
| HD_BLUP | AX-110044748 | 5D | 464108412 | 1.5822 | | 2.14E-06 | EMMAX | SUG |
| HD_BLUP | AX-110044748 | 5D | 464108412 | -1.5727 | | 1.58E-06 | GEMMA | SUG |
| HD_BLUP | AX-111175690 | 5D | 464415388 | 0.9372 | | 3.85E-08 | EMMAX | SIG |
| HD_BLUP | AX-111175690 | 5D | 464415388 | -0.9279 | | 3.34E-08 | GEMMA | SIG |
| HD_BLUP | AX-110958036 | 5D | 469179441 | 0.4226 | | 7.81E-11 | BLUPmrMLM | SIG |

| | | | | | | | | |
|---|---|---|---|---|---|---|---|---|
| HD_BLUP | AX-109360916 | 5D | 469286340 | 2.3981 | | 3.28E-11 | EMMAX | SIG |
| HD_BLUP | AX-109360916 | 5D | 469286340 | -2.3935 | | 1.85E-11 | GEMMA | SIG |
| HD_BLUP | AX-109360916 | 5D | 469286340 | 0.7925 | | 1.93E-93 | IIIVmrMLM | SIG |
| HD_BLUP | AX-89336793 | 5D | 489775906 | 0.5708 | 1.1562 | 4.14E-56 | IIIVmrMLM | SIG |
| HD_BLUP | AX-108786115 | 6A | 805880 | -0.3417 | | 1.72E-06 | BLUPmrMLM | SUG |
| HD_BLUP | AX-94523368 | 6A | 584913052 | -0.3717 | | 3.89E-06 | BLUPmrMLM | SUG |
| HD_BLUP | AX-109969056 | 6B | 135264060 | 0.2496 | | 4.15E-14 | IIIVmrMLM | SIG |
| HD_BLUP | AX-111507306 | 6B | 320638540 | -0.3386 | | 4.53E-07 | BLUPmrMLM | SUG |
| HD_BLUP | AX-109391962 | 6B | 339268873 | -1.7124 | | 6.27E-07 | FarmCPU | SUG |
| HD_BLUP | AX-110076917 | 6B | 454024791 | 0.2512 | | 1.80E-06 | BLUPmrMLM | SUG |
| HD_BLUP | AX-111159598 | 6D | 468835389 | -0.6094 | | 5.77E-10 | BLUPmrMLM | SIG |
| HD_BLUP | AX-111004071 | 7A | 151956510 | -1.2364 | | 1.36E-06 | FarmCPU | SUG |
| HD_BLUP | AX-109430816 | 7A | 458990531 | 0.0601 | 0.5058 | 2.42E-08 | IIIVmrMLM | SIG |
| HD_BLUP | AX-109819108 | 7A | 652230718 | 0.1995 | -0.0794 | 1.20E-08 | IIIVmrMLM | SIG |
| HD_BLUP | AX-109520645 | 7A | 652603840 | -0.1605 | 0.0745 | 9.32E-06 | IIIVmrMLM | SUG |
| HD_BLUP | AX-109473440 | 7A | 659838893 | 0.1334 | -1.7021 | 7.26E-11 | IIIVmrMLM | SIG |
| HD_BLUP | AX-108895338 | 7B | 640567795 | 0.0289 | -0.8261 | 1.53E-06 | IIIVmrMLM | SUG |
| HD_BLUP | AX-109315768 | 7B | 649097785 | -0.2605 | | 3.35E-07 | BLUPmrMLM | SUG |
| HD_BLUP | AX-110984479 | 7B | 705251781 | -0.0279 | 0.7898 | 4.60E-07 | IIIVmrMLM | SUG |
| HD_BLUP | AX-110576947 | 7D | 65436205 | -0.7728 | | 1.39E-14 | BLUPmrMLM | SIG |
| HD_BLUP | AX-110576947 | 7D | 65436205 | 0.7936 | | 7.08E-07 | EMMAX | SUG |
| HD_BLUP | AX-110576947 | 7D | 65436205 | -0.7927 | | 3.98E-07 | GEMMA | SUG |
| HD_BLUP | AX-110576947 | 7D | 65436205 | 0.5561 | 0.2033 | 4.55E-52 | IIIVmrMLM | SIG |
| HD_BLUP | AX-110875275 | 7D | 257526220 | -0.3699 | -0.2286 | 2.41E-26 | IIIVmrMLM | SIG |

HD: heading date.

**Supplementary Table S22.** All the known genes for two growth period-related wheat traits in genome-wide association studies and their pieces of evidence

| Trait | Genome-wide association studies | | | | | | | Gene | Symbol | Reference |
|---|---|---|---|---|---|---|---|---|---|---|
| | Environment | marker | Chr | Position | add/dom | P-value | Method | | | |
| FD | FD_18JY | AX-108829556 | 1B | 581099719 | -0.20 /1.15 | 3.00E-07 | IIIVmrMLM | TraesCS1B01G351100 | *TaFT3-B1* | Zikhali et al.,2017 |
| FD | FD_18JY | AX-111713476 | 1D | 41150395 | -0.05 /3.66 | 5.39E-06 | IIIVmrMLM | TraesCS1D01G059200 | *TaTOE-D1* | Zikhali et al.,2017 |
| FD | FD_18XX | AX-108753274 | 2A | 750145504 | -0.02 /-1.45 | 3.97E-07 | IIIVmrMLM | TraesCS2A01G536600 | *TaFT8-1* | Pshenichnikova et al.,2021 |
| FD | FD_18XX | AX-108753274 | 2A | 750145504 | -0.02 /-1.45 | 3.97E-07 | IIIVmrMLM | TraesCS2A01G536700 | *TaFT8-4* | Pshenichnikova et al.,2021 |
| FD | FD_18XX | AX-108753274 | 2A | 750145504 | -0.02 /-1.45 | 3.97E-07 | IIIVmrMLM | TraesCS2A01G536900 | *TaFT8-3* | Pshenichnikova et al.,2021 |
| HD | HD_17ZMD | AX-89414036 | 4B | 13419503 | -0.52 /-- | 2.58E-06 | BLUPmrMLM | TraesCS4B01G020300 | *TaGRP2* | Xiao et al.,2014 |
| HD | HD_18XX | AX-111528735 | 5B | 574063657 | -1.57 /-- | 4.18E-06 | GEMMA | TraesCS5B01G396200 | *PhyC* | Mizuno et al.,2014 |
| HD | HD_18XX | AX-111528735 | 5B | 574063657 | 2.76 /-- | 4.20E-12 | EMMAX | TraesCS5B01G396200 | *PhyC* | Mizuno et al.,2014 |
| HD | HD_17ZMD | AX-111175690 | 5D | 464415388 | -1.51 /-- | 4.75E-07 | GEMMA | TraesCS5D01G401000 | *PhyC* | Mizuno et al.,2014 |
| HD | HD_19XX | AX-111689944 | 5D | 466287415 | 0.77 /-9.48 | 5.67E-32 | IIIVmrMLM | TraesCS5D01G401000 | *PhyC* | Mizuno et al.,2014 |
| HD | HD_BLUP | AX-111175690 | 5D | 464415388 | -0.93 /-- | 7.28E-08 | GEMMA | TraesCS5D01G401000 | *PhyC* | Mizuno et al.,2014 |
| HD | HD_18XX | AX-111175690 | 5D | 464415388 | -1.71 /-- | 7.21E-10 | GEMMA | TraesCS5D01G401000 | *PhyC* | Mizuno et al.,2014 |
| HD | HD_BLUP | AX-111175690 | 5D | 464415388 | 0.94 /-- | 3.85E-08 | EMMAX | TraesCS5D01G401000 | *PhyC* | Mizuno et al.,2014 |
| HD | HD_18XX | AX-111175690 | 5D | 464415388 | 1.71 /-- | 3.34E-10 | EMMAX | TraesCS5D01G401000 | *PhyC* | Mizuno et al.,2014 |
| HD | HD_BLUP | AX-110576947 | 7D | 65436205 | 0.79 /-- | 7.08E-07 | EMMAX | TraesCS7D01G107500 | *TaGW2L-7D* | Zhang et al.,2021 |
| HD | HD_19XX | AX-111061288 | 7D | 66537922 | 0.86 /-- | 2.25E-10 | BLUPmrMLM | TraesCS7D01G107500 | *TaGW2L-7D* | Zhang et al.,2021 |
| HD | HD_18XX | AX-110826147 | 7D | 65503559 | 1.06 /0.39 | 2.47E-71 | IIIVmrMLM | TraesCS7D01G107500 | *TaGW2L-7D* | Zhang et al.,2021 |
| HD | HD_BLUP | AX-110576947 | 7D | 65436205 | -0.79 /-- | 7.84E-07 | GEMMA | TraesCS7D01G107500 | *TaGW2L-7D* | Zhang et al.,2021 |
| HD | HD_BLUP | AX-110576947 | 7D | 65436205 | 0.56 /0.20 | 4.55E-52 | IIIVmrMLM | TraesCS7D01G107500 | *TaGW2L-7D* | Zhang et al.,2021 |
| HD | HD_19XX | AX-111061288 | 7D | 66537922 | -1.142857143 | 9.17E-28 | IIIVmrMLM | TraesCS7D01G107500 | *TaGW2L-7D* | Zhang et al.,2021 |
| HD | HD_18XX | AX-110576947 | 7D | 65436205 | -1.28 /-- | 1.15E-16 | BLUPmrMLM | TraesCS7D01G107500 | *TaGW2L-7D* | Zhang et al.,2021 |
| HD | HD_BLUP | AX-110576947 | 7D | 65436205 | -0.77 /-- | 1.39E-14 | BLUPmrMLM | TraesCS7D01G107500 | *TaGW2L-7D* | Zhang et al.,2021 |

HD: heading date, FD: flowering date.

**Supplementary Table S23**. Candidate genes for two growth period-related traits in genome-wide association studies and their pieces of evidence

| Trait | Genome-wide association studies | | | | | | | Differential expression analysis | | | Homologous genes/Genes reported in Rice/wheat | GO annotation analysis | | P-value in haplotype analysis | Reference |
|---|---|---|---|---|---|---|---|---|---|---|---|---|---|---|---|
| | Environment | Marker | Chr | Position | Add/Dom | P-value | Method | Gene | $Log_2$\|FC\| | P. adj | | ID | Annotation | | |
| HD | HD_18AY | AX-108796834 | 1A | 495037699 | 1.6851/-- | | EMMAX | TraesCS1A01G301400 | 1.97 [HD] | 1.81E-17 [HD] | *OsSTN8* | | | 5.82E-05 | Nath et al.,2013 |
| HD | HD_18AY | AX-108796834 | 1A | 495037699 | -1.68494/-- | 3.77E-07 | GEMMA | TraesCS1A01G301400 | 1.97 [HD] | 1.81E-17 [HD] | *OsSTN8* | | | 5.82E-05 | Nath et al.,2013 |
| HD | HD_18AY | AX-108926906 | 1A | 495181892 | -1.645836/-- | 8.34E-07 | GEMMA | TraesCS1A01G301400 | 1.97 [HD] | 1.81E-17 [HD] | *OsSTN8* | | | 5.82E-05 | Nath et al.,2013 |
| HD | HD_18AY | AX-109426053 | 1A | 494942486 | -1.595152/-- | 1.06E-06 | GEMMA | TraesCS1A01G301400 | 1.97 [HD] | 1.81E-17 [HD] | *OsSTN8* | | | 5.82E-05 | Nath et al.,2013 |
| HD | HD_18AY | AX-109448112 | 1A | 495182069 | -1.595152/-- | 1.06E-06 | GEMMA | TraesCS1A01G301400 | 1.97 [HD] | 1.81E-17 [HD] | *OsSTN8* | | | 5.82E-05 | Nath et al.,2013 |
| HD | HD_18AY | AX-109490911 | 1A | 495014197 | 1.6851/-- | 1.74E-06 | EMMAX | TraesCS1A01G301400 | 1.97 [HD] | 1.81E-17 [HD] | *OsSTN8* | | | 5.82E-05 | Nath et al.,2013 |
| HD | HD_18AY | AX-109490911 | 1A | 495014197 | -1.68494/-- | 3.77E-07 | GEMMA | TraesCS1A01G301400 | 1.97 [HD] | 1.81E-17 [HD] | *OsSTN8* | | | 5.82E-05 | Nath et al.,2013 |
| HD | HD_18AY | AX-109817102 | 1A | 495013803 | -1.52259/-- | 2.27E-06 | GEMMA | TraesCS1A01G301400 | 1.97 [HD] | 1.81E-17 [HD] | *OsSTN8* | | | 5.82E-05 | Nath et al.,2013 |
| HD | HD_18AY | AX-109886157 | 1A | 494887518 | -1.595152/-- | 1.06E-06 | GEMMA | TraesCS1A01G301400 | 1.97 [HD] | 1.81E-17 [HD] | *OsSTN8* | | | 5.82E-05 | Nath et al.,2013 |
| HD | HD_18AY | AX-110606043 | 1A | 495036799 | -1.595152/-- | 1.06E-06 | GEMMA | TraesCS1A01G301400 | 1.97 [HD] | 1.81E-17 [HD] | *OsSTN8* | | | 5.82E-05 | Nath et al.,2013 |
| HD | HD_18AY | AX-110667522 | 1A | 495183206 | -1.555753/-- | 2.28E-06 | GEMMA | TraesCS1A01G301400 | 1.97 [HD] | 1.81E-17 [HD] | *OsSTN8* | | | 5.82E-05 | Nath et al.,2013 |
| HD | HD_18AY | AX-110995371 | 1A | 495127554 | -1.491663/-- | 2.13E-06 | GEMMA | TraesCS1A01G301400 | 1.97 [HD] | 1.81E-17 [HD] | *OsSTN8* | | | 5.82E-05 | Nath et al.,2013 |
| HD | HD_18AY | AX-111002026 | 1A | 495037475 | -1.595152/-- | 1.06E-06 | GEMMA | TraesCS1A01G301400 | 1.97 [HD] | 1.81E-17 [HD] | *OsSTN8* | | | 5.82E-05 | Nath et al.,2013 |
| HD | HD_18AY | AX-111453306 | 1A | 494818292 | -1.524407/-- | 2.30E-06 | GEMMA | TraesCS1A01G301400 | 1.97 [HD] | 1.81E-17 [HD] | *OsSTN8* | | | 5.82E-05 | Nath et al.,2013 |
| HD | HD_18AY | AX-111462775 | 1A | 494810364 | 1.6851/-- | 1.74E-06 | EMMAX | TraesCS1A01G301400 | 1.97 [HD] | 1.81E-17 [HD] | *OsSTN8* | | | 5.82E-05 | Nath et al.,2013 |
| HD | HD_18AY | AX-111462775 | 1A | 494810364 | -1.68494/-- | 3.77E-07 | GEMMA | TraesCS1A01G301400 | 1.97 [HD] | 1.81E-17 [HD] | *OsSTN8* | | | 5.82E-05 | Nath et al.,2013 |
| HD | HD_18AY | AX-111488547 | 1A | 495013458 | -1.595152/-- | 1.06E-06 | GEMMA | TraesCS1A01G301400 | 1.97 [HD] | 1.81E-17 [HD] | *OsSTN8* | | | 5.82E-05 | Nath et al.,2013 |
| HD | HD_18AY | AX-111550437 | 1A | 495037353 | 1.6851/-- | 1.74E-06 | EMMAX | TraesCS1A01G301400 | 1.97 [HD] | 1.81E-17 [HD] | *OsSTN8* | | | 5.82E-05 | Nath et al.,2013 |
| HD | HD_18AY | AX-111550437 | 1A | 495037353 | -1.68494/-- | 3.77E-07 | GEMMA | TraesCS1A01G301400 | 1.97 [HD] | 1.81E-17 [HD] | *OsSTN8* | | | 5.82E-05 | Nath et al.,2013 |
| HD | HD_18AY | AX-111555272 | 1A | 495189010 | -1.363981/-- | 7.79E-07 | GEMMA | TraesCS1A01G301400 | 1.97 [HD] | 1.81E-17 [HD] | *OsSTN8* | | | 5.82E-05 | Nath et al.,2013 |

| | | | | | | | | | | | | | | | |
|---|---|---|---|---|---|---|---|---|---|---|---|---|---|---|---|
| HD | HD_18AY | AX-111609168 | 1A | 495014994 | -1.640345/-- | 7.12E-07 | GEMMA | TraesCS1A01G301400 | 1.97 $^{HD}$ | 1.81E-17 $^{HD}$ | *OsSTN8* | | | 5.82E-05 | Nath et al.,2013 |
| HD | HD_18AY | AX-111767357 | 1A | 495125499 | 1.6348/-- | 2.16E-06 | EMMAX | TraesCS1A01G301400 | 1.97 $^{HD}$ | 1.81E-17 $^{HD}$ | *OsSTN8* | | | 5.82E-05 | Nath et al.,2013 |
| HD | HD_18AY | AX-111767357 | 1A | 495125499 | -1.629327/-- | 5.15E-07 | GEMMA | TraesCS1A01G301400 | 1.97 $^{HD}$ | 1.81E-17 $^{HD}$ | *OsSTN8* | | | 5.82E-05 | Nath et al.,2013 |
| HD | HD_17AY | AX-109360506 | 1A | 586279275 | 0.0164/-2.2495 | 2.27E-12 | IIIVmrMLM | TraesCS1A01G435000 | 7.71 $^{HD}$ | 2.29E-09 $^{HD}$ | *STH1* | | | 1.18E-11 | Xiang et al.,2022 |
| HD | HD_18XX | AX-109857857 | 1D | 488348964 | -0.3833/-- | 1.21E-05 | BLUPmrMLM | TraesCS1D01G444100 | 5.40 $^{HD}$ | 1.14E-02 $^{HD}$ | *STH1* | | | 3.73E-19 | Xiang et al.,2022 |
| HD | HD_17AY | AX-110390871 | 3B | 7252439 | -0.75004/-- | 3.17E-07 | EMMAX | TraesCS3B01G017400 | 1.11 $^{HD}$ | 5.30E-06 $^{HD}$ | | GO:0048574 | long-day photoperiodism, flowering | 7.43E-24 | |
| HD | HD_17AY | AX-110390871 | 3B | 7252439 | -0.75004/-- | 3.17E-07 | EMMAX | TraesCS3B01G017400 | 1.11 $^{HD}$ | 5.30E-06 $^{HD}$ | | GO:0048571 | long-day photoperiodism | 7.43E-24 | |
| HD | HD_BLUP | AX-110390871 | 3B | 7252439 | 0.2046/-- | 5.45E-05 | BLUPmrMLM | TraesCS3B01G017400 | 1.11 $^{HD}$ | 5.30E-06 $^{HD}$ | | GO:0048574 | long-day photoperiodism, flowering | 2.71E-25 | |
| HD | HD_17AY | AX-110390871 | 3B | 7252439 | 0.5207/-- | 2.04E-07 | BLUPmrMLM | TraesCS3B01G017400 | 1.11 $^{HD}$ | 5.30E-06 $^{HD}$ | | GO:0048574 | long-day photoperiodism, flowering | 7.43E-24 | |
| HD | HD_17AY | AX-110390871 | 3B | 7252439 | -0.75004/-- | 3.17E-07 | EMMAX | TraesCS3B01G017400 | 1.11 $^{HD}$ | 5.30E-06 $^{HD}$ | | GO:0048573 | photoperiodism, flowering | 7.43E-24 | |
| HD | HD_BLUP | AX-110390871 | 3B | 7252439 | 0.2046/-- | 5.45E-05 | BLUPmrMLM | TraesCS3B01G017400 | 1.11 $^{HD}$ | 5.30E-06 $^{HD}$ | | GO:0048573 | photoperiodism, flowering | 2.71E-25 | |
| HD | HD_17AY | AX-110390871 | 3B | 7252439 | 0.5207/-- | 2.04E-07 | BLUPmrMLM | TraesCS3B01G017400 | 1.11 $^{HD}$ | 5.30E-06 $^{HD}$ | | GO:0009648 | photoperiodism | 7.43E-24 | |
| HD | HD_17AY | AX-110390871 | 3B | 7252439 | 0.5207/-- | 2.04E-07 | BLUPmrMLM | TraesCS3B01G017400 | 1.11 $^{HD}$ | 5.30E-06 $^{HD}$ | | GO:0048571 | long-day photoperiodism | 7.43E-24 | |
| HD | HD_BLUP | AX-110390871 | 3B | 7252439 | 0.2046/-- | 5.45E-05 | BLUPmrMLM | TraesCS3B01G017400 | 1.11 $^{HD}$ | 5.30E-06 $^{HD}$ | | GO:0048571 | long-day photoperiodism | 2.71E-25 | |
| HD | HD_17AY | AX-110390871 | 3B | 7252439 | 0.5207/-- | 2.04E-07 | BLUPmrMLM | TraesCS3B01G017400 | 1.11 $^{HD}$ | 5.30E-06 $^{HD}$ | | GO:0048573 | photoperiodism, flowering | 7.43E-24 | |
| HD | HD_18JY | AX-111612892 | 3B | 7615861 | -0.64947/-- | 1.04E-06 | EMMAX | TraesCS3B01G017400 | 1.11 $^{HD}$ | 5.30E-06 $^{HD}$ | | GO:0048571 | long-day photoperiodism | 1.10E-21 | |
| HD | HD_18JY | AX-111612892 | 3B | 7615861 | 0.6507201/-- | 8.31E-07 | GEMMA | TraesCS3B01G017400 | 1.11 $^{HD}$ | 5.30E-06 $^{HD}$ | | GO:0048573 | photoperiodism, flowering | 1.10E-21 | |
| HD | HD_18JY | AX-111612892 | 3B | 7615861 | -0.64947/-- | 1.04E-06 | EMMAX | TraesCS3B01G017400 | 1.11 $^{HD}$ | 5.30E-06 $^{HD}$ | | GO:0048573 | photoperiodism, flowering | 1.10E-21 | |
| HD | HD_18JY | AX-111612892 | 3B | 7615861 | -0.64947/-- | 1.04E-06 | EMMAX | TraesCS3B01G017400 | 1.11 $^{HD}$ | 5.30E-06 $^{HD}$ | | GO:0009648 | photoperiodism | 1.10E-21 | |
| HD | HD_17AY | AX-110390871 | 3B | 7252439 | 0.7502504/-- | 1.78E-07 | GEMMA | TraesCS3B01G017400 | 1.11 $^{HD}$ | 5.30E-06 $^{HD}$ | | GO:0048573 | photoperiodism, flowering | 7.43E-24 | |
| HD | HD_17AY | AX-110390871 | 3B | 7252439 | 0.7502504/-- | 1.78E-07 | GEMMA | TraesCS3B01G017400 | 1.11 $^{HD}$ | 5.30E-06 $^{HD}$ | | GO:0009648 | photoperiodism | 7.43E-24 | |
| HD | HD_17AY | AX-110390871 | 3B | 7252439 | -0.75004/-- | 3.17E-07 | EMMAX | TraesCS3B01G017400 | 1.11 $^{HD}$ | 5.30E-06 $^{HD}$ | | GO:0009648 | photoperiodism | 7.43E-24 | |
| HD | HD_17AY | AX-110390871 | 3B | 7252439 | 0.7502504/-- | 1.78E-07 | GEMMA | TraesCS3B01G017400 | 1.11 $^{HD}$ | 5.30E-06 $^{HD}$ | | GO:0048574 | long-day photoperiodism, flowering | 7.43E-24 | |
| HD | HD_17AY | AX-111081536 | 3B | 8806517 | -1.02728 | 1.69E-12 | IIIVmrMLM | TraesCS3B01G017400 | 1.11 $^{HD}$ | 5.30E-06 $^{HD}$ | | GO:0048573 | photoperiodism, flowering | 7.43E-24 | |

| | | | | | | | | | | | | | |
|---|---|---|---|---|---|---|---|---|---|---|---|---|---|
| HD | HD_17AY | AX-111081536 | 3B | 8806517 | -1.02728 | 1.69E-12 | IIIVmrMLM | TraesCS3B01G017400 | 1.11 HD | 5.30E-06 HD | GO:0048571 | long-day photoperiodism | 7.43E-24 |
| HD | HD_17AY | AX-111081536 | 3B | 8806517 | -1.02728 | 1.69E-12 | IIIVmrMLM | TraesCS3B01G017400 | 1.11 HD | 5.30E-06 HD | GO:0009648 | photoperiodism | 7.43E-24 |
| HD | HD_18JY | AX-108865439 | 3B | 7614235 | 0.62705/-- | 1.51E-06 | EMMAX | TraesCS3B01G017400 | 1.11 HD | 5.30E-06 HD | GO:0048573 | photoperiodism, flowering | 1.10E-21 |
| HD | HD_18JY | AX-108865439 | 3B | 7614235 | 0.62705/-- | 1.51E-06 | EMMAX | TraesCS3B01G017400 | 1.11 HD | 5.30E-06 HD | GO:0009648 | photoperiodism | 1.10E-21 |
| HD | HD_18JY | AX-108865439 | 3B | 7614235 | -0.6283741/-- | 1.17E-06 | GEMMA | TraesCS3B01G017400 | 1.11 HD | 5.30E-06 HD | GO:0048574 | long-day photoperiodism, flowering | 1.10E-21 |
| HD | HD_18JY | AX-108865439 | 3B | 7614235 | -0.6283741/-- | 1.17E-06 | GEMMA | TraesCS3B01G017400 | 1.11 HD | 5.30E-06 HD | GO:0009648 | photoperiodism | 1.10E-21 |
| HD | HD_18JY | AX-108865439 | 3B | 7614235 | -0.6283741/-- | 1.17E-06 | GEMMA | TraesCS3B01G017400 | 1.11 HD | 5.30E-06 HD | GO:0048571 | long-day photoperiodism | 1.10E-21 |
| HD | HD_18JY | AX-108865439 | 3B | 7614235 | -0.6283741/-- | 1.17E-06 | GEMMA | TraesCS3B01G017400 | 1.11 HD | 5.30E-06 HD | GO:0048573 | photoperiodism, flowering | 1.10E-21 |
| HD | HD_18JY | AX-110364271 | 3B | 7610526 | 0.61218/-- | 1.65E-06 | EMMAX | TraesCS3B01G017400 | 1.11 HD | 5.30E-06 HD | GO:0048573 | photoperiodism, flowering | 1.10E-21 |
| HD | HD_18JY | AX-110364271 | 3B | 7610526 | 0.61218/-- | 1.65E-06 | EMMAX | TraesCS3B01G017400 | 1.11 HD | 5.30E-06 HD | GO:0048574 | long-day photoperiodism, flowering | 1.10E-21 |
| HD | HD_17AY | AX-110390871 | 3B | 7252439 | 0.7502504/-- | 1.78E-07 | GEMMA | TraesCS3B01G017400 | 1.11 HD | 5.30E-06 HD | GO:0048571 | long-day photoperiodism | 7.43E-24 |
| HD | HD_17AY | AX-111081536 | 3B | 8806517 | -1.02728 | 1.69E-12 | IIIVmrMLM | TraesCS3B01G017400 | 1.11 HD | 5.30E-06 HD | GO:0048574 | long-day photoperiodism, flowering | 7.43E-24 |
| HD | HD_18JY | AX-110364271 | 3B | 7610526 | -0.6143386/-- | 1.32E-06 | GEMMA | TraesCS3B01G017400 | 1.11 HD | 5.30E-06 HD | GO:0048571 | long-day photoperiodism | 1.10E-21 |
| HD | HD_18JY | AX-110364271 | 3B | 7610526 | -0.6143386/-- | 1.32E-06 | GEMMA | TraesCS3B01G017400 | 1.11 HD | 5.30E-06 HD | GO:0048573 | photoperiodism, flowering | 1.10E-21 |
| HD | HD_18JY | AX-110364271 | 3B | 7610526 | -0.6143386/-- | 1.32E-06 | GEMMA | TraesCS3B01G017400 | 1.11 HD | 5.30E-06 HD | GO:0048574 | long-day photoperiodism, flowering | 1.10E-21 |
| HD | HD_18JY | AX-110364271 | 3B | 7610526 | -0.6143386/-- | 1.32E-06 | GEMMA | TraesCS3B01G017400 | 1.11 HD | 5.30E-06 HD | GO:0009648 | photoperiodism | 1.10E-21 |
| HD | HD_18JY | AX-110641031 | 3B | 7601649 | -0.63279/-- | 1.79E-06 | EMMAX | TraesCS3B01G017400 | 1.11 HD | 5.30E-06 HD | GO:0048574 | long-day photoperiodism, flowering | 1.10E-21 |
| HD | HD_18JY | AX-110641031 | 3B | 7601649 | -0.63279/-- | 1.79E-06 | EMMAX | TraesCS3B01G017400 | 1.11 HD | 5.30E-06 HD | GO:0048573 | photoperiodism, flowering | 1.10E-21 |
| HD | HD_18JY | AX-110641031 | 3B | 7601649 | -0.63279/-- | 1.79E-06 | EMMAX | TraesCS3B01G017400 | 1.11 HD | 5.30E-06 HD | GO:0048571 | long-day photoperiodism | 1.10E-21 |
| HD | HD_18JY | AX-110641031 | 3B | 7601649 | -0.63279/-- | 1.79E-06 | EMMAX | TraesCS3B01G017400 | 1.11 HD | 5.30E-06 HD | GO:0009648 | photoperiodism | 1.10E-21 |
| HD | HD_18JY | AX-110641031 | 3B | 7601649 | 0.6332392/-- | 1.45E-06 | GEMMA | TraesCS3B01G017400 | 1.11 HD | 5.30E-06 HD | GO:0009648 | photoperiodism | 1.10E-21 |
| HD | HD_18JY | AX-110641031 | 3B | 7601649 | 0.6332392/-- | 1.45E-06 | GEMMA | TraesCS3B01G017400 | 1.11 HD | 5.30E-06 HD | GO:0048574 | long-day photoperiodism, flowering | 1.10E-21 |
| HD | HD_18JY | AX-110641031 | 3B | 7601649 | 0.6332392/-- | 1.45E-06 | GEMMA | TraesCS3B01G017400 | 1.11 HD | 5.30E-06 HD | GO:0048573 | photoperiodism, flowering | 1.10E-21 |
| HD | HD_18JY | AX-110364271 | 3B | 7610526 | 0.61218/-- | 1.65E-06 | EMMAX | TraesCS3B01G017400 | 1.11 HD | 5.30E-06 HD | GO:0048571 | long-day photoperiodism | 1.10E-21 |

| | | | | | | | | | | | | | |
|---|---|---|---|---|---|---|---|---|---|---|---|---|---|
| HD | HD_18JY | AX-110364271 | 3B | 7610526 | 0.61218/-- | 1.65E-06 | EMMAX | TraesCS3B01G017400 | 1.11 $^{HD}$ | 5.30E-06 $^{HD}$ | GO:0009648 | photoperiodism | 1.10E-21 |
| HD | HD_18JY | AX-111612892 | 3B | 7615861 | 0.4681/-- | 1.13E-08 | BLUPmrMLM | TraesCS3B01G017400 | 1.11 $^{HD}$ | 5.30E-06 $^{HD}$ | GO:0048573 | photoperiodism, flowering | 1.10E-21 |
| HD | HD_18JY | AX-111612892 | 3B | 7615861 | 0.4681/-- | 1.13E-08 | BLUPmrMLM | TraesCS3B01G017400 | 1.11 $^{HD}$ | 5.30E-06 $^{HD}$ | GO:0009648 | photoperiodism | 1.10E-21 |
| HD | HD_18JY | AX-111612892 | 3B | 7615861 | 0.4681/-- | 1.13E-08 | BLUPmrMLM | TraesCS3B01G017400 | 1.11 $^{HD}$ | 5.30E-06 $^{HD}$ | GO:0048571 | long-day photoperiodism | 1.10E-21 |
| HD | HD_18JY | AX-111612892 | 3B | 7615861 | -0.64947/-- | 1.04E-06 | EMMAX | TraesCS3B01G017400 | 1.11 $^{HD}$ | 5.30E-06 $^{HD}$ | GO:0048574 | long-day photoperiodism, flowering | 1.10E-21 |
| HD | HD_18JY | AX-111612892 | 3B | 7615861 | 0.6507201/-- | 8.31E-07 | GEMMA | TraesCS3B01G017400 | 1.11 $^{HD}$ | 5.30E-06 $^{HD}$ | GO:0048574 | long-day photoperiodism, flowering | 1.10E-21 |
| HD | HD_18JY | AX-111612892 | 3B | 7615861 | 0.6507201/-- | 8.31E-07 | GEMMA | TraesCS3B01G017400 | 1.11 $^{HD}$ | 5.30E-06 $^{HD}$ | GO:0048571 | long-day photoperiodism | 1.10E-21 |
| HD | HD_18JY | AX-111612892 | 3B | 7615861 | 0.6507201/-- | 8.31E-07 | GEMMA | TraesCS3B01G017400 | 1.11 $^{HD}$ | 5.30E-06 $^{HD}$ | GO:0009648 | photoperiodism | 1.10E-21 |
| HD | HD_18JY | AX-110641031 | 3B | 7601649 | 0.6332392/-- | 1.45E-06 | GEMMA | TraesCS3B01G017400 | 1.11 $^{HD}$ | 5.30E-06 $^{HD}$ | GO:0048571 | long-day photoperiodism | 1.10E-21 |
| HD | HD_18JY | AX-111612892 | 3B | 7615861 | 0.4681/-- | 1.13E-08 | BLUPmrMLM | TraesCS3B01G017400 | 1.11 $^{HD}$ | 5.30E-06 $^{HD}$ | GO:0048574 | long-day photoperiodism, flowering | 1.10E-21 |
| HD | HD_BLUP | AX-110390871 | 3B | 7252439 | 0.2046/-- | 5.45E-05 | BLUPmrMLM | TraesCS3B01G017400 | 1.11 $^{HD}$ | 5.30E-06 $^{HD}$ | GO:0009648 | photoperiodism | 2.71E-25 |
| HD | HD_18JY | AX-108865439 | 3B | 7614235 | 0.62705/-- | 1.51E-06 | EMMAX | TraesCS3B01G017400 | 1.11 $^{HD}$ | 5.30E-06 $^{HD}$ | GO:0048574 | long-day photoperiodism, flowering | 1.10E-21 |
| HD | HD_18JY | AX-108865439 | 3B | 7614235 | 0.62705/-- | 1.51E-06 | EMMAX | TraesCS3B01G017400 | 1.11 $^{HD}$ | 5.30E-06 $^{HD}$ | GO:0048571 | long-day photoperiodism | 1.10E-21 |
| HD | HD_18AY | AX-110601323 | 3B | 760960881 | 0.092623 | 3.08E-06 | IIIVmrMLM | TraesCS3B01G517100 | -1.75 $^{HD}$ | 1.99E-20 $^{HD}$ | GO:0009648 | photoperiodism | 1.18E-55 |
| HD | HD_18AY | AX-110601323 | 3B | 760960881 | 0.092623 | 3.08E-06 | IIIVmrMLM | TraesCS3B01G517100 | -1.75 $^{HD}$ | 1.99E-20 $^{HD}$ | GO:0048573 | photoperiodism, flowering | 1.18E-55 |
| HD | HD_18AY | AX-110601323 | 3B | 760960881 | 0.092623 | 3.08E-06 | IIIVmrMLM | TraesCS3B01G517100 | -1.75 $^{HD}$ | 1.99E-20 $^{HD}$ | GO:2000028 | regulation of photoperiodism, flowering | 1.18E-55 |
| HD | HD_BLUP | AX-109503531 | 5B | 692425935 | -0.276/-- | 1.66E-07 | BLUPmrMLM | TraesCS5B01G538100 | 3.27 $^{HD}$ | 4.93E-03 $^{HD}$ | GO:0009648 | photoperiodism | 1.10E-32 |
| HD | HD_17ZMD | AX-95008036 | 5B | 691860601 | -0.4083/-- | 4.19E-05 | BLUPmrMLM | TraesCS5B01G538100 | 3.27 $^{HD}$ | 4.93E-03 $^{HD}$ | GO:0048573 | photoperiodism, flowering | 4.15E-22 |
| HD | HD_BLUP | AX-109503531 | 5B | 692425935 | 0.217/0.0014 | 2.05E-09 | IIIVmrMLM | TraesCS5B01G538100 | 3.27 $^{HD}$ | 4.93E-03 $^{HD}$ | GO:0048573 | photoperiodism, flowering | 1.10E-32 |
| HD | HD_BLUP | AX-109503531 | 5B | 692425935 | -0.276/-- | 1.66E-07 | BLUPmrMLM | TraesCS5B01G538100 | 3.27 $^{HD}$ | 4.93E-03 $^{HD}$ | GO:0048573 | photoperiodism, flowering | 1.10E-32 |
| HD | HD_18XX | AX-109396197 | 5B | 692713178 | 0.3895/-- | 4.13E-05 | BLUPmrMLM | TraesCS5B01G538100 | 3.27 $^{HD}$ | 4.93E-03 $^{HD}$ | GO:0048573 | photoperiodism, flowering | 1.81E-27 |
| HD | HD_18XX | AX-109396197 | 5B | 692713178 | 0.3895/-- | 4.13E-05 | BLUPmrMLM | TraesCS5B01G538100 | 3.27 $^{HD}$ | 4.93E-03 $^{HD}$ | GO:0009648 | photoperiodism | 1.81E-27 |
| HD | HD_BLUP | AX-109503531 | 5B | 692425935 | 0.217/0.0014 | 2.05E-09 | IIIVmrMLM | TraesCS5B01G538100 | 3.27 $^{HD}$ | 4.93E-03 $^{HD}$ | GO:0009648 | photoperiodism | 1.10E-32 |
| HD | HD_17ZMD | AX-95008036 | 5B | 691860601 | -0.4083/-- | 4.19E-05 | BLUPmrMLM | TraesCS5B01G538100 | 3.27 $^{HD}$ | 4.93E-03 $^{HD}$ | GO:0009648 | photoperiodism | 4.15E-22 |

| | | | | | | | | | | | | | | | |
|---|---|---|---|---|---|---|---|---|---|---|---|---|---|---|---|
| HD | HD_BLUP | AX-110958036 | 5D | 469179441 | 0.4226/-- | 7.81E-11 | BLUPmrMLM | TraesCS5D01G401500 | -1.22 [HD] | 5.84E-03 [HD] | *OsMADS14* | | | 8.40E-06 | Kobayashi et al.,2012 |
| HD | HD_19XX | AX-111689944 | 5D | 466287415 | 0.7675/-9.4787 | 5.67E-32 | IIIVmrMLM | TraesCS5D01G401500 | -1.22 [HD] | 5.84E-03 [HD] | *OsMADS14* | | | 3.14E-03 | Kobayashi et al.,2012 |
| HD | HD_BLUP | AX-109360916 | 5D | 469286340 | 2.3981/-- | 3.28E-11 | EMMAX | TraesCS5D01G403200 | -1.38 [HD] | 3.24E-05 [HD] | | GO:0010048 | vernalization response | 1.20E-05 | |
| HD | HD_BLUP | AX-109360916 | 5D | 469286340 | -2.393489/-- | 1.85E-11 | GEMMA | TraesCS5D01G403200 | -1.38 [HD] | 3.24E-05 [HD] | | GO:0010048 | vernalization response | 1.20E-05 | |
| HD | HD_18XX | AX-109360916 | 5D | 469286340 | 1.6512/-- | ###### | IIIVmrMLM | TraesCS5D01G403200 | -1.38 [HD] | 3.24E-05 [HD] | | GO:0010048 | vernalization response | 8.31E-03 | |
| HD | HD_18XX | AX-109360916 | 5D | 469286340 | 4.6652/-- | 3.21E-16 | EMMAX | TraesCS5D01G403200 | -1.38 [HD] | 3.24E-05 [HD] | | GO:0010048 | vernalization response | 8.31E-03 | |
| HD | HD_BLUP | AX-109360916 | 5D | 469286340 | 0.7925/-- | 1.93E-93 | IIIVmrMLM | TraesCS5D01G403200 | -1.38 [HD] | 3.24E-05 [HD] | | GO:0010048 | vernalization response | 1.20E-05 | |
| HD | HD_17ZMD | AX-109360916 | 5D | 469286340 | 0.5862/-- | 6.45E-24 | IIIVmrMLM | TraesCS5D01G403200 | -1.38 [HD] | 3.24E-05 [HD] | | GO:0010048 | vernalization response | 1.62E-05 | |
| HD | HD_18XX | AX-109360916 | 5D | 469286340 | -4.65868/-- | 1.05E-16 | GEMMA | TraesCS5D01G403200 | -1.38 [HD] | 3.24E-05 [HD] | | GO:0010048 | vernalization response | 8.31E-03 | |
| HD | HD_17ZMD | AX-109360916 | 5D | 469286340 | -3.690295/-- | 3.27E-09 | GEMMA | TraesCS5D01G403200 | -1.38 [HD] | 3.24E-05 [HD] | | GO:0010048 | vernalization response | 1.62E-05 | |
| HD | HD_BLUP | AX-110958036 | 5D | 469179441 | 0.4226/-- | 7.81E-11 | BLUPmrMLM | TraesCS5D01G403200 | -1.38 [HD] | 3.24E-05 [HD] | | GO:0010048 | vernalization response | 1.20E-05 | |
| HD | HD_19XX | AX-109360916 | 5D | 469286340 | 1.0338/-- | 2.71E-45 | IIIVmrMLM | TraesCS5D01G403200 | -1.38 [HD] | 3.24E-05 [HD] | | GO:0010048 | vernalization response | 9.68E-04 | |
| HD | HD_17AY | AX-94497617 | 6B | 32328193 | -0.3898/-- | 5.11E-07 | BLUPmrMLM | TraesCS6B01G054100 | 5.08 [HD] | 1.15E-02 [HD] | | GO:0009648 | photoperiodism | 2.68E-11 | |
| HD | HD_17ZMD | AX-109330846 | 6B | 32946759 | 0.461/-- | 1.70E-06 | BLUPmrMLM | TraesCS6B01G054100 | 5.08 [HD] | 1.15E-02 [HD] | | GO:0009648 | photoperiodism | 8.85E-14 | |
| HD | HD_17ZMD | AX-109330846 | 6B | 32946759 | 0.461/-- | 1.70E-06 | BLUPmrMLM | TraesCS6B01G054100 | 5.08 [HD] | 1.15E-02 [HD] | | GO:0048573 | photoperiodism, flowering | 8.85E-14 | |
| HD | HD_17AY | AX-94497617 | 6B | 32328193 | -0.3898/-- | 5.11E-07 | BLUPmrMLM | TraesCS6B01G054100 | 5.08 [HD] | 1.15E-02 [HD] | | GO:0048573 | photoperiodism, flowering | 2.68E-11 | |
| FD | FD_18JY | AX-94495040 | 1A | 549619420 | 0.5908/-- | 2.80E-06 | BLUPmrMLM | TraesCS1A01G372200 | -4.63 [FD1]; -3.18 [FD2] | 5.42E-03 [FD1]; 8.95E-03 [FD2] | | GO:0048573 | photoperiodism, flowering | 1.75E-24 | |
| FD | FD_18JY | AX-94495040 | 1A | 549619420 | 0.5908/-- | 2.80E-06 | BLUPmrMLM | TraesCS1A01G372200 | -4.63 [FD1]; -3.18 [FD2] | 5.42E-03 [FD1]; 8.95E-03 [FD2] | | GO:0009648 | photoperiodism | 1.75E-24 | |
| FD | FD_18JY | AX-94495040 | 1A | 549619420 | 0.5908/-- | 2.80E-06 | BLUPmrMLM | TraesCS1A01G372300 | -5.22 [FD1] | 1.34E-06 [FD1] | | GO:0048573 | photoperiodism, flowering | 3.74E-37 | |
| FD | FD_18JY | AX-94495040 | 1A | 549619420 | 0.5908/-- | 2.80E-06 | BLUPmrMLM | TraesCS1A01G372300 | -5.22 [FD1] | 1.34E-06 [FD1] | | GO:0009648 | photoperiodism | 3.74E-37 | |
| FD | FD_17AY | AX-108898996 | 1D | 454766541 | -0.0082 | 1.02E-11 | IIIVmrMLM | TraesCS1D01G378800 | -6.40 [FD1]; -7.93 [FD2] | 3.56E-08 [FD1]; 4.69E-08 [FD2] | | GO:0009648 | photoperiodism | 1.40E-05 | |
| FD | FD_17AY | AX-108898996 | 1D | 454766541 | -0.0082 | 1.02E-11 | IIIVmrMLM | TraesCS1D01G378800 | -6.40 [FD1]; -7.93 [FD2] | 3.56E-08 [FD1]; 4.69E-08 [FD2] | | GO:0048573 | photoperiodism, flowering | 1.40E-05 | |
| FD | FD_17AY | AX-108898996 | 1D | 454766541 | -0.0082 | 1.02E-11 | IIIVmrMLM | TraesCS1D01G378900 | -6.68 [FD2]; -1.85 [FD1] | 1.04E-09 [FD2]; 8.10E-03 [FD1] | | GO:0009648 | photoperiodism | 1.07E-04 | |
| FD | FD_17AY | AX-108898996 | 1D | 454766541 | -0.0082 | 1.02E-11 | IIIVmrMLM | TraesCS1D01G378900 | -6.68 [FD2]; -1.85 [FD1] | 1.04E-09 [FD2]; 8.10E-03 [FD1] | | GO:0048573 | photoperiodism, flowering | 1.07E-04 | |

| | | | | | | | | | | | | | | | |
|---|---|---|---|---|---|---|---|---|---|---|---|---|---|---|---|
| FD | FD_17ZMD | AX-110374023 | 2A | 71438950 | 0.4007/-- | 2.37E-05 | BLUPmrMLM | TraesCS2A01G119600 | 7.60 $^{FD1}$; 7.60 $^{FD2}$ | 4.32E-47 $^{FD1}$; 4.32E-47 $^{FD2}$ | *OsCOL13* | | | 2.03E-33 | Sheng et al.,2016 |
| FD | FD_19XX | AX-112288504 | 2A | 77528386 | 0.4374/-- | 2.50E-05 | BLUPmrMLM | TraesCS2A01G132300 | -10.30 $^{FD1}$; -10.68 $^{FD2}$ | 1.12E-50 $^{FD1}$; 3.28E-23 $^{FD2}$ | *OsFTL12* | | | 1.79E-03 | Zhang et al., 2023 |
| FD | FD_18XX | AX-111035423 | 2A | 136058841 | 0.1253/-3.0982 | 2.93E-20 | IIIVmrMLM | TraesCS2A01G179000 | 3.94 $^{FD2}$; 9.05 $^{FD1}$ | 3.60E-02 $^{FD2}$; 8.41E-11 $^{FD1}$ | | GO:2000028 | regulation of photoperiodism, flowering | 1.99E-32 | |
| FD | FD_18XX | AX-111035423 | 2A | 136058841 | 0.1253/-3.0982 | 2.93E-20 | IIIVmrMLM | TraesCS2A01G179200 | 11.21 $^{FD2}$ | 3.75E-17 $^{FD2}$ | | GO:2000028 | regulation of photoperiodism, flowering | 3.35E-32 | |
| FD | FD_BLUP | AX-111680033 | 2A | 162275567 | -0.216/-- | 8.12E-16 | IIIVmrMLM | TraesCS2A01G194500 | 1.13 $^{FD1}$ | 2.35E-02 $^{FD1}$ | *OsLBD37* | | | 4.07E-13 | Li et al.,2017 |
| FD | FD_18XX | AX-108753274 | 2A | 750145504 | 0.010767 | 3.97E-07 | IIIVmrMLM | TraesCS2A01G534200 | -5.01 $^{FD1}$; -5.95 $^{FD2}$ | 6.74E-03 $^{FD1}$; 5.34E-04 $^{FD2}$ | | GO:2000028 | regulation of photoperiodism, flowering | 3.27E-09 | |
| FD | FD_18XX | AX-108753274 | 2A | 750145504 | 0.010767 | 3.97E-07 | IIIVmrMLM | TraesCS2A01G534200 | -5.01 $^{FD1}$; -5.95 $^{FD2}$ | 6.74E-03 $^{FD1}$; 5.34E-04 $^{FD2}$ | | GO:0009648 | photoperiodism | 3.27E-09 | |
| FD | FD_18XX | AX-108753274 | 2A | 750145504 | 0.010767 | 3.97E-07 | IIIVmrMLM | TraesCS2A01G534300 | -6.38 $^{FD2}$; -2.88 $^{FD1}$ | 2.15E-05 $^{FD2}$; 2.21E-03 $^{FD1}$ | | GO:0009648 | photoperiodism | 3.93E-09 | |
| FD | FD_18XX | AX-108753274 | 2A | 750145504 | 0.010767 | 3.97E-07 | IIIVmrMLM | TraesCS2A01G534300 | -6.38 $^{FD2}$; -2.88 $^{FD1}$ | 2.15E-05 $^{FD2}$; 2.21E-03 $^{FD1}$ | | GO:2000028 | regulation of photoperiodism, flowering | 3.93E-09 | |
| FD | FD_18XX | AX-94652369 | 2B | 106129285 | 0.165757 | 1.58E-17 | IIIVmrMLM | TraesCS2B01G140300 | 5.81 $^{FD2}$; 3.69 $^{FD1}$ | 3.64E-09 $^{FD2}$; 7.18E-03 $^{FD1}$ | *OsCOL13* | | | 1.42E-06 | Sheng et al.,2016 |
| FD | FD_BLUP | AX-94697254 | 2D | 33710135 | -1.3352/-- | 1.26E-06 | FarmCPU | TraesCS2D01G082000 | -2.69 $^{FD1}$; -1.38 $^{FD2}$ | 2.40E-36 $^{FD1}$; 7.13E-08 $^{FD2}$ | *OsPLC1* | | | 2.27E-02 | Sheng et al.,2016 |
| FD | FD_17AY | AX-110390871 | 3B | 7252439 | 0.7323/-- | 7.35E-07 | GEMMA | TraesCS3B01G017400 | -5.65 $^{FD1}$; -3.60 $^{FD2}$ | 2.08E-55 $^{FD1}$; 2.15E-44 $^{FD2}$ | | GO:0048571 | long-day photoperiodism | 4.71E-22 | |
| FD | FD_BLUP | AX-111197043 | 3B | 9088741 | 0.4173/-- | 1.33E-06 | BLUPmrMLM | TraesCS3B01G017400 | -5.65 $^{FD1}$; -3.60 $^{FD2}$ | 2.08E-55 $^{FD1}$; 2.15E-44 $^{FD2}$ | | GO:0048571 | long-day photoperiodism | 2.84E-24 | |
| FD | FD_17AY | AX-110390871 | 3B | 7252439 | 0.7323/-- | 7.35E-07 | GEMMA | TraesCS3B01G017400 | -5.65 $^{FD1}$; -3.60 $^{FD2}$ | 2.08E-55 $^{FD1}$; 2.15E-44 $^{FD2}$ | | GO:0048574 | long-day photoperiodism, flowering | 4.71E-22 | |
| FD | FD_17AY | AX-110390871 | 3B | 7252439 | 0.7323/-- | 7.35E-07 | GEMMA | TraesCS3B01G017400 | -5.65 $^{FD1}$; -3.60 $^{FD2}$ | 2.08E-55 $^{FD1}$; 2.15E-44 $^{FD2}$ | | GO:0048573 | photoperiodism, flowering | 4.71E-22 | |
| FD | FD_17AY | AX-110390871 | 3B | 7252439 | 0.7323/-- | 7.35E-07 | GEMMA | TraesCS3B01G017400 | -5.65 $^{FD1}$; -3.60 $^{FD2}$ | 2.08E-55 $^{FD1}$; 2.15E-44 $^{FD2}$ | | GO:0009648 | photoperiodism | 4.71E-22 | |
| FD | FD_17AY | AX-111081536 | 3B | 8806517 | -0.95138 | 1.63E-08 | IIIVmrMLM | TraesCS3B01G017400 | -5.65 $^{FD1}$; -3.60 $^{FD2}$ | 2.08E-55 $^{FD1}$; 2.15E-44 $^{FD2}$ | | GO:0048571 | long-day photoperiodism | 4.71E-22 | |
| FD | FD_17AY | AX-111081536 | 3B | 8806517 | -0.95138 | 1.63E-08 | IIIVmrMLM | TraesCS3B01G017400 | -5.65 $^{FD1}$; -3.60 $^{FD2}$ | 2.08E-55 $^{FD1}$; 2.15E-44 $^{FD2}$ | | GO:0048574 | long-day photoperiodism, flowering | 4.71E-22 | |
| FD | FD_17AY | AX-111081536 | 3B | 8806517 | -0.95138 | 1.63E-08 | IIIVmrMLM | TraesCS3B01G017400 | -5.65 $^{FD1}$; -3.60 $^{FD2}$ | 2.08E-55 $^{FD1}$; 2.15E-44 $^{FD2}$ | | GO:0048573 | photoperiodism, flowering | 4.71E-22 | |
| FD | FD_17AY | AX-111081536 | 3B | 8806517 | -0.95138 | 1.63E-08 | IIIVmrMLM | TraesCS3B01G017400 | -5.65 $^{FD1}$; -3.60 $^{FD2}$ | 2.08E-55 $^{FD1}$; 2.15E-44 $^{FD2}$ | | GO:0009648 | photoperiodism | 4.71E-22 | |
| FD | FD_17AY | AX-110390871 | 3B | 7252439 | 0.5327/-- | 4.63E-08 | BLUPmrMLM | TraesCS3B01G017400 | -5.65 $^{FD1}$; -3.60 $^{FD2}$ | 2.08E-55 $^{FD1}$; 2.15E-44 $^{FD2}$ | | GO:0048573 | photoperiodism, flowering | 4.71E-22 | |
| FD | FD_17AY | AX-110390871 | 3B | 7252439 | 0.5327/-- | 4.63E-08 | BLUPmrMLM | TraesCS3B01G017400 | -5.65 $^{FD1}$; -3.60 $^{FD2}$ | 2.08E-55 $^{FD1}$; 2.15E-44 $^{FD2}$ | | GO:0048574 | long-day photoperiodism, flowering | 4.71E-22 | |
| FD | FD_BLUP | AX-111197043 | 3B | 9088741 | 0.4173/-- | 1.33E-06 | BLUPmrMLM | TraesCS3B01G017400 | -5.65 $^{FD1}$; -3.60 $^{FD2}$ | 2.08E-55 $^{FD1}$; 2.15E-44 $^{FD2}$ | | GO:0048573 | photoperiodism, flowering | 2.84E-24 | |
| FD | FD_BLUP | AX-111197043 | 3B | 9088741 | 0.4173/-- | 1.33E-06 | BLUPmrMLM | TraesCS3B01G017400 | -5.65 $^{FD1}$; -3.60 $^{FD2}$ | 2.08E-55 $^{FD1}$; 2.15E-44 $^{FD2}$ | | GO:0048574 | long-day photoperiodism, flowering | 2.84E-24 | |

| | | | | | | | | | | | | | | | |
|---|---|---|---|---|---|---|---|---|---|---|---|---|---|---|---|
| FD | FD_17AY | AX-110390871 | 3B | 7252439 | 0.5327/-- | 4.63E-08 | BLUPmrMLM | TraesCS3B01G017400 | -5.65 [FD1]; -3.60 [FD2] | 2.08E-55 [FD1]; 2.15E-44 [FD2] | | GO:0009648 | photoperiodism | 4.71E-22 | |
| FD | FD_BLUP | AX-111197043 | 3B | 9088741 | 0.4173/-- | 1.33E-06 | BLUPmrMLM | TraesCS3B01G017400 | -5.65 [FD1]; -3.60 [FD2] | 2.08E-55 [FD1]; 2.15E-44 [FD2] | | GO:0009648 | photoperiodism | 2.84E-24 | |
| FD | FD_17AY | AX-110390871 | 3B | 7252439 | 0.5327/-- | 4.63E-08 | BLUPmrMLM | TraesCS3B01G017400 | -5.65 [FD1]; -3.60 [FD2] | 2.08E-55 [FD1]; 2.15E-44 [FD2] | | GO:0048571 | long-day photoperiodism | 4.71E-22 | |
| FD | FD_18XX | AX-109357603 | 3B | 667162788 | 2.2225/-- | 3.70E-07 | EMMAX | TraesCS3B01G428100 | -3.96 [FD1]; 1.41 [FD2] | 7.11E-08 [FD1]; 6.67E-05 [FD2] | | GO:2000028 | regulation of photoperiodism, flowering | 1.37E-08 | |
| FD | FD_18XX | AX-109357603 | 3B | 667162788 | -2.2335/-- | 3.26E-07 | GEMMA | TraesCS3B01G428100 | -3.96 [FD1]; 1.41 [FD2] | 7.11E-08 [FD1]; 6.67E-05 [FD2] | | GO:2000028 | regulation of photoperiodism, flowering | 1.37E-08 | |
| FD | FD_19XX | AX-111453739 | 3B | 759265358 | 0.3727/-0.2186 | 1.08E-09 | IIIVmrMLM | TraesCS3B01G517100 | 3.25 [FD2]; 6.35 [FD1] | 1.88E-04 [FD2]; 9.56E-40 [FD1] | | GO:2000028 | regulation of photoperiodism, flowering | 7.29E-16 | |
| FD | FD_19XX | AX-111453739 | 3B | 759265358 | 0.3727/-0.2186 | 1.08E-09 | IIIVmrMLM | TraesCS3B01G517100 | 3.25 [FD2]; 6.35 [FD1] | 1.88E-04 [FD2]; 9.56E-40 [FD1] | | GO:0009648 | photoperiodism | 7.29E-16 | |
| FD | FD_19XX | AX-111057460 | 3B | 759231400 | -0.4248/-- | 1.30E-05 | BLUPmrMLM | TraesCS3B01G517100 | 3.25 [FD2]; 6.35 [FD1] | 1.88E-04 [FD2]; 9.56E-40 [FD1] | | GO:0009648 | photoperiodism | 7.29E-16 | |
| FD | FD_19XX | AX-111453739 | 3B | 759265358 | 0.3727/-0.2186 | 1.08E-09 | IIIVmrMLM | TraesCS3B01G517100 | 3.25 [FD2]; 6.35 [FD1] | 1.88E-04 [FD2]; 9.56E-40 [FD1] | | GO:0048573 | photoperiodism, flowering | 7.29E-16 | |
| FD | FD_19XX | AX-111057460 | 3B | 759231400 | -0.4248/-- | 1.30E-05 | BLUPmrMLM | TraesCS3B01G517100 | 3.25 [FD2]; 6.35 [FD1] | 1.88E-04 [FD2]; 9.56E-40 [FD1] | | GO:0048573 | photoperiodism, flowering | 7.29E-16 | |
| FD | FD_19XX | AX-111057460 | 3B | 759231400 | -0.4248/-- | 1.30E-05 | BLUPmrMLM | TraesCS3B01G517100 | 3.25 [FD2]; 6.35 [FD1] | 1.88E-04 [FD2]; 9.56E-40 [FD1] | | GO:2000028 | regulation of photoperiodism, flowering | 7.29E-16 | |
| FD | FD_18XX | AX-110458146 | 4A | 38722201 | 0.5688/-- | 8.33E-34 | IIIVmrMLM | TraesCS4A01G044800 | 2.26 [FD2]; 1.09 [FD1] | 1.65E-13 [FD2]; 7.71E-03 [FD1] | *OsTPR075* | | | 6.79E-30 | Zhang et al.,2022 |
| FD | FD_18XX | AX-110185660 | 4A | 117173937 | 0.809/-- | 1.59E-09 | BLUPmrMLM | TraesCS4A01G105300 | 2.57 [FD2] | 1.63E-32 [FD2] | *DTH7* | | | 3.78E-17 | Gao et al.,2014 |
| FD | FD_18XX | AX-110185660 | 4A | 117173937 | 0.809/-- | 1.59E-09 | BLUPmrMLM | TraesCS4A01G105300 | 2.57 [FD2] | 1.63E-32 [FD2] | *OsCCT11* | | | 3.78E-17 | Zhang et al. 2015 |
| FD | FD_19XX | AX-108861327 | 4A | 612642486 | -3.6015/-- | 1.97E-06 | GEMMA | TraesCS4A01G324900 | 7.32 [FD2]; 8.56 [FD1] | 5.08E-07 [FD2]; 2.47E-10 [FD1] | | GO:0048573 | photoperiodism, flowering | 6.30E-06 | |
| FD | FD_19XX | AX-108861327 | 4A | 612642486 | -3.6015/-- | 1.97E-06 | GEMMA | TraesCS4A01G324900 | 7.32 [FD2]; 8.56 [FD1] | 5.08E-07 [FD2]; 2.47E-10 [FD1] | | GO:0009648 | photoperiodism | 6.30E-06 | |
| FD | FD_19XX | AX-110371776 | 4A | 612493121 | 1.8713/-- | 1.91E-06 | EMMAX | TraesCS4A01G324900 | 7.32 [FD2]; 8.56 [FD1] | 5.08E-07 [FD2]; 2.47E-10 [FD1] | | GO:0009648 | photoperiodism | 6.30E-06 | |
| FD | FD_19XX | AX-109043286 | 4A | 612484737 | -3.6015/-- | 1.97E-06 | GEMMA | TraesCS4A01G324900 | 7.32 [FD2]; 8.56 [FD1] | 5.08E-07 [FD2]; 2.47E-10 [FD1] | | GO:0048573 | photoperiodism, flowering | 6.30E-06 | |
| FD | FD_19XX | AX-110371776 | 4A | 612493121 | 1.8713/-- | 1.91E-06 | EMMAX | TraesCS4A01G324900 | 7.32 [FD2]; 8.56 [FD1] | 5.08E-07 [FD2]; 2.47E-10 [FD1] | | GO:0048573 | photoperiodism, flowering | 6.30E-06 | |
| FD | FD_19XX | AX-109043286 | 4A | 612484737 | -3.6015/-- | 1.97E-06 | GEMMA | TraesCS4A01G324900 | 7.32 [FD2]; 8.56 [FD1] | 5.08E-07 [FD2]; 2.47E-10 [FD1] | | GO:0009648 | photoperiodism | 6.30E-06 | |
| FD | FD_19XX | AX-109281612 | 4A | 613337260 | 4.1439/-- | 3.76E-08 | EMMAX | TraesCS4A01G324900 | 7.32 [FD2]; 8.56 [FD1] | 5.08E-07 [FD2]; 2.47E-10 [FD1] | | GO:0048573 | photoperiodism, flowering | 6.30E-06 | |
| FD | FD_19XX | AX-109281612 | 4A | 613337260 | 4.1439/-- | 3.76E-08 | EMMAX | TraesCS4A01G324900 | 7.32 [FD2]; 8.56 [FD1] | 5.08E-07 [FD2]; 2.47E-10 [FD1] | | GO:0009648 | photoperiodism | 6.30E-06 | |
| FD | FD_19XX | AX-109281612 | 4A | 613337260 | -4.1179/-- | 1.99E-08 | GEMMA | TraesCS4A01G324900 | 7.32 [FD2]; 8.56 [FD1] | 5.08E-07 [FD2]; 2.47E-10 [FD1] | | GO:0048573 | photoperiodism, flowering | 6.30E-06 | |
| FD | FD_19XX | AX-109281612 | 4A | 613337260 | -4.1179/-- | 1.99E-08 | GEMMA | TraesCS4A01G324900 | 7.32 [FD2]; 8.56 [FD1] | 5.08E-07 [FD2]; 2.47E-10 [FD1] | | GO:0009648 | photoperiodism | 6.30E-06 | |

| | | | | | | | | | | | | | |
|---|---|---|---|---|---|---|---|---|---|---|---|---|---|
| FD | FD_19XX | AX-109376087 | 4A | 613369207 | 3.4807/-- | 6.04E-08 | EMMAX | TraesCS4A01G324900 | 7.32 [FD2]; 8.56 [FD1] | 5.08E-07 [FD2]; 2.47E-10 [FD1] | GO:0048573 | photoperiodism, flowering | 6.30E-06 |
| FD | FD_19XX | AX-109376087 | 4A | 613369207 | 3.4807/-- | 6.04E-08 | EMMAX | TraesCS4A01G324900 | 7.32 [FD2]; 8.56 [FD1] | 5.08E-07 [FD2]; 2.47E-10 [FD1] | GO:0009648 | photoperiodism | 6.30E-06 |
| FD | FD_19XX | AX-109376087 | 4A | 613369207 | -3.4109/-- | 6.27E-08 | GEMMA | TraesCS4A01G324900 | 7.32 [FD2]; 8.56 [FD1] | 5.08E-07 [FD2]; 2.47E-10 [FD1] | GO:0048573 | photoperiodism, flowering | 6.30E-06 |
| FD | FD_19XX | AX-109376087 | 4A | 613369207 | -3.4109/-- | 6.27E-08 | GEMMA | TraesCS4A01G324900 | 7.32 [FD2]; 8.56 [FD1] | 5.08E-07 [FD2]; 2.47E-10 [FD1] | GO:0009648 | photoperiodism | 6.30E-06 |
| FD | FD_19XX | AX-109456475 | 4A | 613404437 | 2.0447/-- | 8.90E-08 | EMMAX | TraesCS4A01G324900 | 7.32 [FD2]; 8.56 [FD1] | 5.08E-07 [FD2]; 2.47E-10 [FD1] | GO:0048573 | photoperiodism, flowering | 6.30E-06 |
| FD | FD_19XX | AX-109456475 | 4A | 613404437 | 2.0447/-- | 8.90E-08 | EMMAX | TraesCS4A01G324900 | 7.32 [FD2]; 8.56 [FD1] | 5.08E-07 [FD2]; 2.47E-10 [FD1] | GO:0009648 | photoperiodism | 6.30E-06 |
| FD | FD_19XX | AX-109456475 | 4A | 613404437 | -2.0304/-- | 4.82E-08 | GEMMA | TraesCS4A01G324900 | 7.32 [FD2]; 8.56 [FD1] | 5.08E-07 [FD2]; 2.47E-10 [FD1] | GO:0048573 | photoperiodism, flowering | 6.30E-06 |
| FD | FD_19XX | AX-109456475 | 4A | 613404437 | -2.0304/-- | 4.82E-08 | GEMMA | TraesCS4A01G324900 | 7.32 [FD2]; 8.56 [FD1] | 5.08E-07 [FD2]; 2.47E-10 [FD1] | GO:0009648 | photoperiodism | 6.30E-06 |
| FD | FD_19XX | AX-110501116 | 4A | 613368960 | -2.0304/-- | 4.82E-08 | GEMMA | TraesCS4A01G324900 | 7.32 [FD2]; 8.56 [FD1] | 5.08E-07 [FD2]; 2.47E-10 [FD1] | GO:0009648 | photoperiodism | 6.30E-06 |
| FD | FD_19XX | AX-110501116 | 4A | 613368960 | -2.0304/-- | 4.82E-08 | GEMMA | TraesCS4A01G324900 | 7.32 [FD2]; 8.56 [FD1] | 5.08E-07 [FD2]; 2.47E-10 [FD1] | GO:0048573 | photoperiodism, flowering | 6.30E-06 |
| FD | FD_19XX | AX-109589014 | 4A | 612639170 | -3.6015/-- | 1.97E-06 | GEMMA | TraesCS4A01G324900 | 7.32 [FD2]; 8.56 [FD1] | 5.08E-07 [FD2]; 2.47E-10 [FD1] | GO:0009648 | photoperiodism | 6.30E-06 |
| FD | FD_19XX | AX-109589014 | 4A | 612639170 | -3.6015/-- | 1.97E-06 | GEMMA | TraesCS4A01G324900 | 7.32 [FD2]; 8.56 [FD1] | 5.08E-07 [FD2]; 2.47E-10 [FD1] | GO:0048573 | photoperiodism, flowering | 6.30E-06 |
| FD | FD_19XX | AX-109846294 | 4A | 610629973 | 3.5959/-- | 1.97E-06 | EMMAX | TraesCS4A01G324900 | 7.32 [FD2]; 8.56 [FD1] | 5.08E-07 [FD2]; 2.47E-10 [FD1] | GO:0048573 | photoperiodism, flowering | 6.30E-06 |
| FD | FD_19XX | AX-109846294 | 4A | 610629973 | 3.5959/-- | 1.97E-06 | EMMAX | TraesCS4A01G324900 | 7.32 [FD2]; 8.56 [FD1] | 5.08E-07 [FD2]; 2.47E-10 [FD1] | GO:0009648 | photoperiodism | 6.30E-06 |
| FD | FD_19XX | AX-109846294 | 4A | 610629973 | -3.5191/-- | 1.47E-06 | GEMMA | TraesCS4A01G324900 | 7.32 [FD2]; 8.56 [FD1] | 5.08E-07 [FD2]; 2.47E-10 [FD1] | GO:0009648 | photoperiodism | 6.30E-06 |
| FD | FD_19XX | AX-109846294 | 4A | 610629973 | -3.5191/-- | 1.47E-06 | GEMMA | TraesCS4A01G324900 | 7.32 [FD2]; 8.56 [FD1] | 5.08E-07 [FD2]; 2.47E-10 [FD1] | GO:0048573 | photoperiodism, flowering | 6.30E-06 |
| FD | FD_19XX | AX-109971768 | 4A | 613118940 | 3.8418/-- | 8.22E-07 | EMMAX | TraesCS4A01G324900 | 7.32 [FD2]; 8.56 [FD1] | 5.08E-07 [FD2]; 2.47E-10 [FD1] | GO:0009648 | photoperiodism | 6.30E-06 |
| FD | FD_19XX | AX-109971768 | 4A | 613118940 | 3.8418/-- | 8.22E-07 | EMMAX | TraesCS4A01G324900 | 7.32 [FD2]; 8.56 [FD1] | 5.08E-07 [FD2]; 2.47E-10 [FD1] | GO:0048573 | photoperiodism, flowering | 6.30E-06 |
| FD | FD_19XX | AX-109971768 | 4A | 613118940 | -3.8480/-- | 3.65E-07 | GEMMA | TraesCS4A01G324900 | 7.32 [FD2]; 8.56 [FD1] | 5.08E-07 [FD2]; 2.47E-10 [FD1] | GO:0048573 | photoperiodism, flowering | 6.30E-06 |
| FD | FD_19XX | AX-109971768 | 4A | 613118940 | -3.8480/-- | 3.65E-07 | GEMMA | TraesCS4A01G324900 | 7.32 [FD2]; 8.56 [FD1] | 5.08E-07 [FD2]; 2.47E-10 [FD1] | GO:0009648 | photoperiodism | 6.30E-06 |
| FD | FD_19XX | AX-110363512 | 4A | 612680091 | 1.8799/-- | 1.58E-06 | EMMAX | TraesCS4A01G324900 | 7.32 [FD2]; 8.56 [FD1] | 5.08E-07 [FD2]; 2.47E-10 [FD1] | GO:0048573 | photoperiodism, flowering | 6.30E-06 |
| FD | FD_19XX | AX-110363512 | 4A | 612680091 | 1.8799/-- | 1.58E-06 | EMMAX | TraesCS4A01G324900 | 7.32 [FD2]; 8.56 [FD1] | 5.08E-07 [FD2]; 2.47E-10 [FD1] | GO:0009648 | photoperiodism | 6.30E-06 |
| FD | FD_19XX | AX-110363512 | 4A | 612680091 | -1.8810/-- | 7.28E-07 | GEMMA | TraesCS4A01G324900 | 7.32 [FD2]; 8.56 [FD1] | 5.08E-07 [FD2]; 2.47E-10 [FD1] | GO:0048573 | photoperiodism, flowering | 6.30E-06 |
| FD | FD_19XX | AX-110363512 | 4A | 612680091 | -1.8810/-- | 7.28E-07 | GEMMA | TraesCS4A01G324900 | 7.32 [FD2]; 8.56 [FD1] | 5.08E-07 [FD2]; 2.47E-10 [FD1] | GO:0009648 | photoperiodism | 6.30E-06 |

| | | | | | | | | | | | | | |
|---|---|---|---|---|---|---|---|---|---|---|---|---|---|
| FD | FD_19XX | AX-111705164 | 4A | 612667681 | -1.5101/-- | 3.26E-11 | BLUPmrMLM | TraesCS4A01G324900 | 7.32 [FD2]; 8.56 [FD1] | 5.08E-07 [FD2]; 2.47E-10 [FD1] | GO:0048573 | photoperiodism, flowering | 6.30E-06 |
| FD | FD_19XX | AX-110371776 | 4A | 612493121 | -1.8771/-- | 8.23E-07 | GEMMA | TraesCS4A01G324900 | 7.32 [FD2]; 8.56 [FD1] | 5.08E-07 [FD2]; 2.47E-10 [FD1] | GO:0048573 | photoperiodism, flowering | 6.30E-06 |
| FD | FD_19XX | AX-110371776 | 4A | 612493121 | -1.8771/-- | 8.23E-07 | GEMMA | TraesCS4A01G324900 | 7.32 [FD2]; 8.56 [FD1] | 5.08E-07 [FD2]; 2.47E-10 [FD1] | GO:0009648 | photoperiodism | 6.30E-06 |
| FD | FD_19XX | AX-110407094 | 4A | 613365713 | 3.4807/-- | 6.04E-08 | EMMAX | TraesCS4A01G324900 | 7.32 [FD2]; 8.56 [FD1] | 5.08E-07 [FD2]; 2.47E-10 [FD1] | GO:0009648 | photoperiodism | 6.30E-06 |
| FD | FD_19XX | AX-110407094 | 4A | 613365713 | 3.4807/-- | 6.04E-08 | EMMAX | TraesCS4A01G324900 | 7.32 [FD2]; 8.56 [FD1] | 5.08E-07 [FD2]; 2.47E-10 [FD1] | GO:0048573 | photoperiodism, flowering | 6.30E-06 |
| FD | FD_19XX | AX-110407094 | 4A | 613365713 | -3.4109/-- | 6.27E-08 | GEMMA | TraesCS4A01G324900 | 7.32 [FD2]; 8.56 [FD1] | 5.08E-07 [FD2]; 2.47E-10 [FD1] | GO:0048573 | photoperiodism, flowering | 6.30E-06 |
| FD | FD_19XX | AX-110407094 | 4A | 613365713 | -3.4109/-- | 6.27E-08 | GEMMA | TraesCS4A01G324900 | 7.32 [FD2]; 8.56 [FD1] | 5.08E-07 [FD2]; 2.47E-10 [FD1] | GO:0009648 | photoperiodism | 6.30E-06 |
| FD | FD_19XX | AX-110432011 | 4A | 613078092 | 3.8418/-- | 8.22E-07 | EMMAX | TraesCS4A01G324900 | 7.32 [FD2]; 8.56 [FD1] | 5.08E-07 [FD2]; 2.47E-10 [FD1] | GO:0048573 | photoperiodism, flowering | 6.30E-06 |
| FD | FD_19XX | AX-110432011 | 4A | 613078092 | 3.8418/-- | 8.22E-07 | EMMAX | TraesCS4A01G324900 | 7.32 [FD2]; 8.56 [FD1] | 5.08E-07 [FD2]; 2.47E-10 [FD1] | GO:0009648 | photoperiodism | 6.30E-06 |
| FD | FD_19XX | AX-110432011 | 4A | 613078092 | -3.8480/-- | 3.65E-07 | GEMMA | TraesCS4A01G324900 | 7.32 [FD2]; 8.56 [FD1] | 5.08E-07 [FD2]; 2.47E-10 [FD1] | GO:0048573 | photoperiodism, flowering | 6.30E-06 |
| FD | FD_19XX | AX-110432011 | 4A | 613078092 | -3.8480/-- | 3.65E-07 | GEMMA | TraesCS4A01G324900 | 7.32 [FD2]; 8.56 [FD1] | 5.08E-07 [FD2]; 2.47E-10 [FD1] | GO:0009648 | photoperiodism | 6.30E-06 |
| FD | FD_19XX | AX-86183809 | 4A | 612060283 | -4.5712/-- | 1.17E-07 | GEMMA | TraesCS4A01G324900 | 7.32 [FD2]; 8.56 [FD1] | 5.08E-07 [FD2]; 2.47E-10 [FD1] | GO:0048573 | photoperiodism, flowering | 6.30E-06 |
| FD | FD_19XX | AX-86183809 | 4A | 612060283 | -4.5712/-- | 1.17E-07 | GEMMA | TraesCS4A01G324900 | 7.32 [FD2]; 8.56 [FD1] | 5.08E-07 [FD2]; 2.47E-10 [FD1] | GO:0009648 | photoperiodism | 6.30E-06 |
| FD | FD_19XX | AX-110501116 | 4A | 613368960 | 2.0447/-- | 8.90E-08 | EMMAX | TraesCS4A01G324900 | 7.32 [FD2]; 8.56 [FD1] | 5.08E-07 [FD2]; 2.47E-10 [FD1] | GO:0009648 | photoperiodism | 6.30E-06 |
| FD | FD_19XX | AX-110501116 | 4A | 613368960 | 2.0447/-- | 8.90E-08 | EMMAX | TraesCS4A01G324900 | 7.32 [FD2]; 8.56 [FD1] | 5.08E-07 [FD2]; 2.47E-10 [FD1] | GO:0048573 | photoperiodism, flowering | 6.30E-06 |
| FD | FD_19XX | AX-111457878 | 4A | 613170537 | -1.8710/-- | 7.08E-07 | GEMMA | TraesCS4A01G324900 | 7.32 [FD2]; 8.56 [FD1] | 5.08E-07 [FD2]; 2.47E-10 [FD1] | GO:0009648 | photoperiodism | 6.30E-06 |
| FD | FD_19XX | AX-111457878 | 4A | 613170537 | -1.8710/-- | 7.08E-07 | GEMMA | TraesCS4A01G324900 | 7.32 [FD2]; 8.56 [FD1] | 5.08E-07 [FD2]; 2.47E-10 [FD1] | GO:0048573 | photoperiodism, flowering | 6.30E-06 |
| FD | FD_19XX | AX-111483645 | 4A | 612492000 | 1.8713/-- | 1.91E-06 | EMMAX | TraesCS4A01G324900 | 7.32 [FD2]; 8.56 [FD1] | 5.08E-07 [FD2]; 2.47E-10 [FD1] | GO:0048573 | photoperiodism, flowering | 6.30E-06 |
| FD | FD_19XX | AX-110730034 | 4A | 612088918 | 1.8992/-- | 6.77E-07 | EMMAX | TraesCS4A01G324900 | 7.32 [FD2]; 8.56 [FD1] | 5.08E-07 [FD2]; 2.47E-10 [FD1] | GO:0048573 | photoperiodism, flowering | 6.30E-06 |
| FD | FD_19XX | AX-110730034 | 4A | 612088918 | 1.8992/-- | 6.77E-07 | EMMAX | TraesCS4A01G324900 | 7.32 [FD2]; 8.56 [FD1] | 5.08E-07 [FD2]; 2.47E-10 [FD1] | GO:0009648 | photoperiodism | 6.30E-06 |
| FD | FD_19XX | AX-110730034 | 4A | 612088918 | -1.8830/-- | 4.47E-07 | GEMMA | TraesCS4A01G324900 | 7.32 [FD2]; 8.56 [FD1] | 5.08E-07 [FD2]; 2.47E-10 [FD1] | GO:0048573 | photoperiodism, flowering | 6.30E-06 |
| FD | FD_19XX | AX-110730034 | 4A | 612088918 | -1.8830/-- | 4.47E-07 | GEMMA | TraesCS4A01G324900 | 7.32 [FD2]; 8.56 [FD1] | 5.08E-07 [FD2]; 2.47E-10 [FD1] | GO:0009648 | photoperiodism | 6.30E-06 |
| FD | FD_19XX | AX-110926040 | 4A | 612517494 | 1.8713/-- | 1.91E-06 | EMMAX | TraesCS4A01G324900 | 7.32 [FD2]; 8.56 [FD1] | 5.08E-07 [FD2]; 2.47E-10 [FD1] | GO:0048573 | photoperiodism, flowering | 6.30E-06 |
| FD | FD_19XX | AX-110926040 | 4A | 612517494 | 1.8713/-- | 1.91E-06 | EMMAX | TraesCS4A01G324900 | 7.32 [FD2]; 8.56 [FD1] | 5.08E-07 [FD2]; 2.47E-10 [FD1] | GO:0009648 | photoperiodism | 6.30E-06 |

| | | | | | | | | | | | | | |
|---|---|---|---|---|---|---|---|---|---|---|---|---|---|
| FD | FD_19XX | AX-110926040 | 4A | 612517494 | -1.8771/-- | 8.23E-07 | GEMMA | TraesCS4A01G324900 | 7.32 $^{FD2}$; 8.56 $^{FD1}$ | 5.08E-07 $^{FD2}$; 2.47E-10 $^{FD1}$ | GO:0048573 | photoperiodism, flowering | 6.30E-06 |
| FD | FD_19XX | AX-110926040 | 4A | 612517494 | -1.8771/-- | 8.23E-07 | GEMMA | TraesCS4A01G324900 | 7.32 $^{FD2}$; 8.56 $^{FD1}$ | 5.08E-07 $^{FD2}$; 2.47E-10 $^{FD1}$ | GO:0009648 | photoperiodism | 6.30E-06 |
| FD | FD_19XX | AX-111033323 | 4A | 612437711 | -3.6015/-- | 1.97E-06 | GEMMA | TraesCS4A01G324900 | 7.32 $^{FD2}$; 8.56 $^{FD1}$ | 5.08E-07 $^{FD2}$; 2.47E-10 $^{FD1}$ | GO:0048573 | photoperiodism, flowering | 6.30E-06 |
| FD | FD_19XX | AX-111033323 | 4A | 612437711 | -3.6015/-- | 1.97E-06 | GEMMA | TraesCS4A01G324900 | 7.32 $^{FD2}$; 8.56 $^{FD1}$ | 5.08E-07 $^{FD2}$; 2.47E-10 $^{FD1}$ | GO:0009648 | photoperiodism | 6.30E-06 |
| FD | FD_19XX | AX-111638733 | 4A | 613368868 | 2.0447/-- | 8.90E-08 | EMMAX | TraesCS4A01G324900 | 7.32 $^{FD2}$; 8.56 $^{FD1}$ | 5.08E-07 $^{FD2}$; 2.47E-10 $^{FD1}$ | GO:0048573 | photoperiodism, flowering | 6.30E-06 |
| FD | FD_19XX | AX-111638733 | 4A | 613368868 | 2.0447/-- | 8.90E-08 | EMMAX | TraesCS4A01G324900 | 7.32 $^{FD2}$; 8.56 $^{FD1}$ | 5.08E-07 $^{FD2}$; 2.47E-10 $^{FD1}$ | GO:0009648 | photoperiodism | 6.30E-06 |
| FD | FD_19XX | AX-111638733 | 4A | 613368868 | -2.0304/-- | 4.82E-08 | GEMMA | TraesCS4A01G324900 | 7.32 $^{FD2}$; 8.56 $^{FD1}$ | 5.08E-07 $^{FD2}$; 2.47E-10 $^{FD1}$ | GO:0048573 | photoperiodism, flowering | 6.30E-06 |
| FD | FD_19XX | AX-111638733 | 4A | 613368868 | -2.0304/-- | 4.82E-08 | GEMMA | TraesCS4A01G324900 | 7.32 $^{FD2}$; 8.56 $^{FD1}$ | 5.08E-07 $^{FD2}$; 2.47E-10 $^{FD1}$ | GO:0009648 | photoperiodism | 6.30E-06 |
| FD | FD_19XX | AX-86183809 | 4A | 612060283 | 4.6491/-- | 1.08E-07 | EMMAX | TraesCS4A01G324900 | 7.32 $^{FD2}$; 8.56 $^{FD1}$ | 5.08E-07 $^{FD2}$; 2.47E-10 $^{FD1}$ | GO:0009648 | photoperiodism | 6.30E-06 |
| FD | FD_19XX | AX-111705164 | 4A | 612667681 | -1.5101/-- | 3.26E-11 | BLUPmrMLM | TraesCS4A01G324900 | 7.32 $^{FD2}$; 8.56 $^{FD1}$ | 5.08E-07 $^{FD2}$; 2.47E-10 $^{FD1}$ | GO:0009648 | photoperiodism | 6.30E-06 |
| FD | FD_19XX | AX-111105228 | 4A | 610609020 | 2.7913/-- | 6.87E-07 | EMMAX | TraesCS4A01G324900 | 7.32 $^{FD2}$; 8.56 $^{FD1}$ | 5.08E-07 $^{FD2}$; 2.47E-10 $^{FD1}$ | GO:0009648 | photoperiodism | 6.30E-06 |
| FD | FD_19XX | AX-111105228 | 4A | 610609020 | 2.7913/-- | 6.87E-07 | EMMAX | TraesCS4A01G324900 | 7.32 $^{FD2}$; 8.56 $^{FD1}$ | 5.08E-07 $^{FD2}$; 2.47E-10 $^{FD1}$ | GO:0048573 | photoperiodism, flowering | 6.30E-06 |
| FD | FD_19XX | AX-111105228 | 4A | 610609020 | -2.7394/-- | 4.84E-07 | GEMMA | TraesCS4A01G324900 | 7.32 $^{FD2}$; 8.56 $^{FD1}$ | 5.08E-07 $^{FD2}$; 2.47E-10 $^{FD1}$ | GO:0048573 | photoperiodism, flowering | 6.30E-06 |
| FD | FD_19XX | AX-111105228 | 4A | 610609020 | -2.7394/-- | 4.84E-07 | GEMMA | TraesCS4A01G324900 | 7.32 $^{FD2}$; 8.56 $^{FD1}$ | 5.08E-07 $^{FD2}$; 2.47E-10 $^{FD1}$ | GO:0009648 | photoperiodism | 6.30E-06 |
| FD | FD_19XX | AX-111149628 | 4A | 612493042 | 1.8713/-- | 1.91E-06 | EMMAX | TraesCS4A01G324900 | 7.32 $^{FD2}$; 8.56 $^{FD1}$ | 5.08E-07 $^{FD2}$; 2.47E-10 $^{FD1}$ | GO:0048573 | photoperiodism, flowering | 6.30E-06 |
| FD | FD_19XX | AX-111149628 | 4A | 612493042 | 1.8713/-- | 1.91E-06 | EMMAX | TraesCS4A01G324900 | 7.32 $^{FD2}$; 8.56 $^{FD1}$ | 5.08E-07 $^{FD2}$; 2.47E-10 $^{FD1}$ | GO:0009648 | photoperiodism | 6.30E-06 |
| FD | FD_19XX | AX-111149628 | 4A | 612493042 | -1.8771/-- | 8.23E-07 | GEMMA | TraesCS4A01G324900 | 7.32 $^{FD2}$; 8.56 $^{FD1}$ | 5.08E-07 $^{FD2}$; 2.47E-10 $^{FD1}$ | GO:0048573 | photoperiodism, flowering | 6.30E-06 |
| FD | FD_19XX | AX-111149628 | 4A | 612493042 | -1.8771/-- | 8.23E-07 | GEMMA | TraesCS4A01G324900 | 7.32 $^{FD2}$; 8.56 $^{FD1}$ | 5.08E-07 $^{FD2}$; 2.47E-10 $^{FD1}$ | GO:0009648 | photoperiodism | 6.30E-06 |
| FD | FD_19XX | AX-86183809 | 4A | 612060283 | 4.6491/-- | 1.08E-07 | EMMAX | TraesCS4A01G324900 | 7.32 $^{FD2}$; 8.56 $^{FD1}$ | 5.08E-07 $^{FD2}$; 2.47E-10 $^{FD1}$ | GO:0048573 | photoperiodism, flowering | 6.30E-06 |
| FD | FD_19XX | AX-111493275 | 4A | 612662027 | -3.6015/-- | 1.97E-06 | GEMMA | TraesCS4A01G324900 | 7.32 $^{FD2}$; 8.56 $^{FD1}$ | 5.08E-07 $^{FD2}$; 2.47E-10 $^{FD1}$ | GO:0048573 | photoperiodism, flowering | 6.30E-06 |
| FD | FD_19XX | AX-111493275 | 4A | 612662027 | -3.6015/-- | 1.97E-06 | GEMMA | TraesCS4A01G324900 | 7.32 $^{FD2}$; 8.56 $^{FD1}$ | 5.08E-07 $^{FD2}$; 2.47E-10 $^{FD1}$ | GO:0009648 | photoperiodism | 6.30E-06 |
| FD | FD_19XX | AX-111457878 | 4A | 613170537 | 1.8705/-- | 1.52E-06 | EMMAX | TraesCS4A01G324900 | 7.32 $^{FD2}$; 8.56 $^{FD1}$ | 5.08E-07 $^{FD2}$; 2.47E-10 $^{FD1}$ | GO:0048573 | photoperiodism, flowering | 6.30E-06 |
| FD | FD_19XX | AX-111457878 | 4A | 613170537 | 1.8705/-- | 1.52E-06 | EMMAX | TraesCS4A01G324900 | 7.32 $^{FD2}$; 8.56 $^{FD1}$ | 5.08E-07 $^{FD2}$; 2.47E-10 $^{FD1}$ | GO:0009648 | photoperiodism | 6.30E-06 |
| FD | FD_19XX | AX-95182838 | 4A | 613367700 | 3.2544/-- | 9.23E-08 | EMMAX | TraesCS4A01G324900 | 7.32 $^{FD2}$; 8.56 $^{FD1}$ | 5.08E-07 $^{FD2}$; 2.47E-10 $^{FD1}$ | GO:0048573 | photoperiodism, flowering | 6.30E-06 |

| | | | | | | | | | | | | | | | |
|---|---|---|---|---|---|---|---|---|---|---|---|---|---|---|---|
| FD | FD_19XX | AX-95182838 | 4A | 613367700 | 3.2544/-- | 9.23E-08 | EMMAX | TraesCS4A01G324900 | 7.32 [FD2]; 8.56 [FD1] | 5.08E-07 [FD2]; 2.47E-10 [FD1] | | GO:0009648 | photoperiodism | 6.30E-06 | |
| FD | FD_19XX | AX-111483645 | 4A | 612492000 | 1.8713/-- | 1.91E-06 | EMMAX | TraesCS4A01G324900 | 7.32 [FD2]; 8.56 [FD1] | 5.08E-07 [FD2]; 2.47E-10 [FD1] | | GO:0009648 | photoperiodism | 6.30E-06 | |
| FD | FD_19XX | AX-111483645 | 4A | 612492000 | -1.8771/-- | 8.23E-07 | GEMMA | TraesCS4A01G324900 | 7.32 [FD2]; 8.56 [FD1] | 5.08E-07 [FD2]; 2.47E-10 [FD1] | | GO:0048573 | photoperiodism, flowering | 6.30E-06 | |
| FD | FD_19XX | AX-111483645 | 4A | 612492000 | -1.8771/-- | 8.23E-07 | GEMMA | TraesCS4A01G324900 | 7.32 [FD2]; 8.56 [FD1] | 5.08E-07 [FD2]; 2.47E-10 [FD1] | | GO:0009648 | photoperiodism | 6.30E-06 | |
| FD | FD_19XX | AX-111581594 | 4A | 612973608 | 1.9324/-- | 1.10E-06 | EMMAX | TraesCS4A01G324900 | 7.32 [FD2]; 8.56 [FD1] | 5.08E-07 [FD2]; 2.47E-10 [FD1] | | GO:0009648 | photoperiodism | 6.30E-06 | |
| FD | FD_19XX | AX-111581594 | 4A | 612973608 | 1.9324/-- | 1.10E-06 | EMMAX | TraesCS4A01G324900 | 7.32 [FD2]; 8.56 [FD1] | 5.08E-07 [FD2]; 2.47E-10 [FD1] | | GO:0048573 | photoperiodism, flowering | 6.30E-06 | |
| FD | FD_19XX | AX-111768266 | 4A | 612494779 | -3.6015/-- | 1.97E-06 | GEMMA | TraesCS4A01G324900 | 7.32 [FD2]; 8.56 [FD1] | 5.08E-07 [FD2]; 2.47E-10 [FD1] | | GO:0048573 | photoperiodism, flowering | 6.30E-06 | |
| FD | FD_19XX | AX-111581594 | 4A | 612973608 | -1.9382/-- | 4.60E-07 | GEMMA | TraesCS4A01G324900 | 7.32 [FD2]; 8.56 [FD1] | 5.08E-07 [FD2]; 2.47E-10 [FD1] | | GO:0048573 | photoperiodism, flowering | 6.30E-06 | |
| FD | FD_19XX | AX-111581594 | 4A | 612973608 | -1.9382/-- | 4.60E-07 | GEMMA | TraesCS4A01G324900 | 7.32 [FD2]; 8.56 [FD1] | 5.08E-07 [FD2]; 2.47E-10 [FD1] | | GO:0009648 | photoperiodism | 6.30E-06 | |
| FD | FD_19XX | AX-111705164 | 4A | 612667681 | 1.9639/-- | 5.32E-07 | EMMAX | TraesCS4A01G324900 | 7.32 [FD2]; 8.56 [FD1] | 5.08E-07 [FD2]; 2.47E-10 [FD1] | | GO:0048573 | photoperiodism, flowering | 6.30E-06 | |
| FD | FD_19XX | AX-111705164 | 4A | 612667681 | 1.9639/-- | 5.32E-07 | EMMAX | TraesCS4A01G324900 | 7.32 [FD2]; 8.56 [FD1] | 5.08E-07 [FD2]; 2.47E-10 [FD1] | | GO:0009648 | photoperiodism | 6.30E-06 | |
| FD | FD_19XX | AX-111768266 | 4A | 612494779 | -3.6015/-- | 1.97E-06 | GEMMA | TraesCS4A01G324900 | 7.32 [FD2]; 8.56 [FD1] | 5.08E-07 [FD2]; 2.47E-10 [FD1] | | GO:0009648 | photoperiodism | 6.30E-06 | |
| FD | FD_19XX | AX-89491834 | 4A | 612091001 | -1.7550/-- | 1.90E-06 | GEMMA | TraesCS4A01G324900 | 7.32 [FD2]; 8.56 [FD1] | 5.08E-07 [FD2]; 2.47E-10 [FD1] | | GO:0009648 | photoperiodism | 6.30E-06 | |
| FD | FD_19XX | AX-95182838 | 4A | 613367700 | -3.1911/-- | 9.71E-08 | GEMMA | TraesCS4A01G324900 | 7.32 [FD2]; 8.56 [FD1] | 5.08E-07 [FD2]; 2.47E-10 [FD1] | | GO:0048573 | photoperiodism, flowering | 6.30E-06 | |
| FD | FD_19XX | AX-111705164 | 4A | 612667681 | -1.9700/-- | 2.10E-07 | GEMMA | TraesCS4A01G324900 | 7.32 [FD2]; 8.56 [FD1] | 5.08E-07 [FD2]; 2.47E-10 [FD1] | | GO:0048573 | photoperiodism, flowering | 6.30E-06 | |
| FD | FD_19XX | AX-111705164 | 4A | 612667681 | -1.9700/-- | 2.10E-07 | GEMMA | TraesCS4A01G324900 | 7.32 [FD2]; 8.56 [FD1] | 5.08E-07 [FD2]; 2.47E-10 [FD1] | | GO:0009648 | photoperiodism | 6.30E-06 | |
| FD | FD_19XX | AX-89491834 | 4A | 612091001 | -1.7550/-- | 1.90E-06 | GEMMA | TraesCS4A01G324900 | 7.32 [FD2]; 8.56 [FD1] | 5.08E-07 [FD2]; 2.47E-10 [FD1] | | GO:0048573 | photoperiodism, flowering | 6.30E-06 | |
| FD | FD_19XX | AX-95182838 | 4A | 613367700 | -3.1911/-- | 9.71E-08 | GEMMA | TraesCS4A01G324900 | 7.32 [FD2]; 8.56 [FD1] | 5.08E-07 [FD2]; 2.47E-10 [FD1] | | GO:0009648 | photoperiodism | 6.30E-06 | |
| FD | FD_BLUP | AX-109816200 | 4A | 649204800 | -0.3292 | 3.57E-16 | IIIVmrMLM | TraesCS4A01G375100 | 1.24 [FD2]; 1.24 [FD1] | 4.47E-05 [FD2]; 4.47E-05 [FD1] | | GO:2000028 | regulation of photoperiodism, flowering | 3.93E-07 | |
| FD | FD_18JY | AX-109816200 | 4A | 649204800 | -0.58148/-- | 1.30E-06 | EMMAX | TraesCS4A01G375100 | 1.24 [FD2]; 1.24 [FD1] | 4.47E-05 [FD2]; 4.47E-05 [FD1] | | GO:2000028 | regulation of photoperiodism, flowering | 6.57E-06 | |
| FD | FD_18JY | AX-108812049 | 4A | 650375652 | -0.57644/-- | 1.15E-06 | EMMAX | TraesCS4A01G375100 | 1.24 [FD2]; 1.24 [FD1] | 4.47E-05 [FD2]; 4.47E-05 [FD1] | | GO:2000028 | regulation of photoperiodism, flowering | 6.57E-06 | |
| FD | FD_18JY | AX-108812049 | 4A | 650375652 | 0.57583/-- | 5.68E-07 | GEMMA | TraesCS4A01G375100 | 1.24 [FD2]; 1.24 [FD1] | 4.47E-05 [FD2]; 4.47E-05 [FD1] | | GO:2000028 | regulation of photoperiodism, flowering | 6.57E-06 | |
| FD | FD_18JY | AX-109816200 | 4A | 649204800 | 0.58382/-- | 5.24E-07 | GEMMA | TraesCS4A01G375100 | 1.24 [FD2]; 1.24 [FD1] | 4.47E-05 [FD2]; 4.47E-05 [FD1] | | GO:2000028 | regulation of photoperiodism, flowering | 6.57E-06 | |
| FD | FD_19XX | AX-111097142 | 4B | 608663022 | 1.3498/-- | 2.18E-11 | BLUPmrMLM | TraesCS4B01G317400 | 1.24 [FD1] | 8.02E-09 [FD1] | *Se14* | GO:0048579 | negative regulation of long-day photoperiodism, flowering | 2.79E-02 | Yokoo et al.,2014 |

| | | | | | | | | | | | | | | | |
|---|---|---|---|---|---|---|---|---|---|---|---|---|---|---|---|
| FD | FD_19XX | AX-111097142 | 4B | 608663022 | 1.3498/-- | 2.18E-11 | BLUPmrMLM | TraesCS4B01G317400 | 1.24 FD1 | 8.02E-09 FD1 | *Se14* | GO:0048587 | regulation of short-day photoperiodism, flowering | 2.79E-02 | Yokoo et al.,2014 |
| FD | FD_19XX | AX-111097142 | 4B | 608663022 | 1.3498/-- | 2.18E-11 | BLUPmrMLM | TraesCS4B01G317400 | 1.24 FD1 | 8.02E-09 FD1 | *Se14* | GO:0048577 | negative regulation of short-day photoperiodism, flowering | 2.79E-02 | Yokoo et al.,2014 |
| FD | FD_19XX | AX-111097142 | 4B | 608663022 | 1.3498/-- | 2.18E-11 | BLUPmrMLM | TraesCS4B01G317400 | 1.24 FD1 | 8.02E-09 FD1 | *Se14* | GO:0048586 | regulation of long-day photoperiodism, flowering | 2.79E-02 | Yokoo et al.,2014 |
| FD | FD_19XX | AX-111097142 | 4B | 608663022 | 1.3498/-- | 2.18E-11 | BLUPmrMLM | TraesCS4B01G317400 | 1.24 FD1 | 8.02E-09 FD1 | *Se14* | GO:2000028 | regulation of photoperiodism, flowering | 2.79E-02 | Yokoo et al.,2014 |
| FD | FD_19XX | AX-108893364 | 5A | 591097621 | -4.9986/-- | 3.09E-07 | GEMMA | TraesCS5A01G393400 | -2.38 FD2 | 3.58E-04 FD2 | | GO:0010048 | vernalization response | 1.27E-05 | |
| FD | FD_19XX | AX-108893364 | 5A | 591097621 | 5.0399/-- | 4.69E-07 | EMMAX | TraesCS5A01G393400 | -2.38 FD2 | 3.58E-04 FD2 | | GO:0010048 | vernalization response | 1.27E-05 | |
| FD | FD_19XX | AX-111057571 | 5A | 591004477 | -2.7211/-- | 1.44E-07 | GEMMA | TraesCS5A01G393400 | -2.38 FD2 | 3.58E-04 FD2 | | GO:0010048 | vernalization response | 1.27E-05 | |
| FD | FD_19XX | AX-95659239 | 5A | 591091029 | 5.0399/-- | 4.69E-07 | EMMAX | TraesCS5A01G393400 | -2.38 FD2 | 3.58E-04 FD2 | | GO:0010048 | vernalization response | 1.27E-05 | |
| FD | FD_19XX | AX-95659239 | 5A | 591091029 | -4.9986/-- | 3.09E-07 | GEMMA | TraesCS5A01G393400 | -2.38 FD2 | 3.58E-04 FD2 | | GO:0010048 | vernalization response | 1.27E-05 | |
| FD | FD_19XX | AX-109503011 | 5A | 590599714 | -4.6236/-- | 1.29E-06 | GEMMA | TraesCS5A01G393400 | -2.38 FD2 | 3.58E-04 FD2 | | GO:0010048 | vernalization response | 1.27E-05 | |
| FD | FD_19XX | AX-111057571 | 5A | 591004477 | 2.7429/-- | 2.13E-07 | EMMAX | TraesCS5A01G393400 | -2.38 FD2 | 3.58E-04 FD2 | | GO:0010048 | vernalization response | 1.27E-05 | |
| FD | FD_19XX | AX-111747299 | 5A | 590680035 | -4.9596/-- | 1.02E-06 | GEMMA | TraesCS5A01G393400 | -2.38 FD2 | 3.58E-04 FD2 | | GO:0010048 | vernalization response | 1.27E-05 | |
| FD | FD_19XX | AX-110593755 | 5A | 590848863 | -4.9596/-- | 1.02E-06 | GEMMA | TraesCS5A01G393400 | -2.38 FD2 | 3.58E-04 FD2 | | GO:0010048 | vernalization response | 1.27E-05 | |
| FD | FD_19XX | AX-110435701 | 5A | 591126755 | 2.4647/-- | 9.54E-07 | EMMAX | TraesCS5A01G393400 | -2.38 FD2 | 3.58E-04 FD2 | | GO:0010048 | vernalization response | 1.27E-05 | |
| FD | FD_19XX | AX-110435701 | 5A | 591126755 | -2.4441/-- | 6.29E-07 | GEMMA | TraesCS5A01G393400 | -2.38 FD2 | 3.58E-04 FD2 | | GO:0010048 | vernalization response | 1.27E-05 | |
| FD | FD_19XX | AX-109497126 | 5A | 590996771 | 5.1965/-- | 5.80E-07 | EMMAX | TraesCS5A01G393400 | -2.38 FD2 | 3.58E-04 FD2 | | GO:0010048 | vernalization response | 1.27E-05 | |
| FD | FD_19XX | AX-109497126 | 5A | 590996771 | -5.1305/-- | 4.41E-07 | GEMMA | TraesCS5A01G393400 | -2.38 FD2 | 3.58E-04 FD2 | | GO:0010048 | vernalization response | 1.27E-05 | |
| FD | FD_19XX | AX-109503011 | 5A | 590599714 | 4.6748/-- | 1.79E-06 | EMMAX | TraesCS5A01G393400 | -2.38 FD2 | 3.58E-04 FD2 | | GO:0010048 | vernalization response | 1.27E-05 | |
| FD | FD_19XX | AX-111747299 | 5A | 590680035 | 5.0238/-- | 1.29E-06 | EMMAX | TraesCS5A01G393400 | -2.38 FD2 | 3.58E-04 FD2 | | GO:0010048 | vernalization response | 1.27E-05 | |
| FD | FD_19XX | AX-110593755 | 5A | 590848863 | 5.0238/-- | 1.29E-06 | EMMAX | TraesCS5A01G393400 | -2.38 FD2 | 3.58E-04 FD2 | | GO:0010048 | vernalization response | 1.27E-05 | |
| FD | FD_BLUP | AX-110538089 | 5B | 539858157 | 0.1699/-0.2641 | 1.27E-10 | IIIVmrMLM | TraesCS5B01G358500 | -1.68 FD2 | 3.13E-10 FD2 | | GO:0009648 | photoperiodism | 2.37E-20 | |
| FD | FD_BLUP | AX-110538089 | 5B | 539858157 | 0.1699/-0.2641 | 1.27E-10 | IIIVmrMLM | TraesCS5B01G358500 | -1.68 FD2 | 3.13E-10 FD2 | | GO:0048574 | long-day photoperiodism, flowering | 2.37E-20 | |
| FD | FD_BLUP | AX-110538089 | 5B | 539858157 | 0.1699/-0.2641 | 1.27E-10 | IIIVmrMLM | TraesCS5B01G358500 | -1.68 FD2 | 3.13E-10 FD2 | | GO:0048571 | long-day photoperiodism | 2.37E-20 | |
| FD | FD_BLUP | AX-110538089 | 5B | 539858157 | 0.1699/-0.2641 | 1.27E-10 | IIIVmrMLM | TraesCS5B01G358500 | -1.68 FD2 | 3.13E-10 FD2 | | GO:0048573 | photoperiodism, flowering | 2.37E-20 | |

| | | | | | | | | | | | | | | | |
|---|---|---|---|---|---|---|---|---|---|---|---|---|---|---|---|
| FD | FD_18XX | AX-111754571 | 5D | 266755425 | 0.3599/-- | 1.95E-15 | IIIVmrMLM | TraesCS5D01G170100 | 1.36 [FD1]; 1.36 [FD2] | 1.90E-12 [FD1]; 1.90E-12 [FD2] | *OsTrx1* | | | 6.14E-35 | Xu et al.,2022 |
| FD | FD_18JY | AX-110426343 | 5D | 430363560 | -2.5065/-- | 9.01E-07 | GEMMA | TraesCS5D01G341000 | 5.43 [FD2] | 7.42E-04 [FD2] | *DHD4* | | | 1.26E-13 | Cai et al.,2021 |
| FD | FD_18JY | AX-110426343 | 5D | 430363560 | -2.5065/-- | 9.01E-07 | GEMMA | TraesCS5D01G341000 | 5.43 [FD2] | 7.42E-04 [FD2] | *OsCCT19* | | | 1.26E-13 | Zhang et al. 2015 |
| FD | FD_BLUP | AX-109360916 | 5D | 469286340 | 2.1977/-- | 1.08E-09 | EMMAX | TraesCS5D01G403200 | -1.26 [FD2] | 2.36E-03 [FD2] | | GO:0010048 | vernalization response | 2.00E-05 | |
| FD | FD_BLUP | AX-109360916 | 5D | 469286340 | -2.1988/-- | 5.88E-10 | GEMMA | TraesCS5D01G403200 | -1.26 [FD2] | 2.36E-03 [FD2] | | GO:0010048 | vernalization response | 2.00E-05 | |
| FD | FD_BLUP | AX-109360916 | 5D | 469286340 | 0.6981/-- | ###### | IIIVmrMLM | TraesCS5D01G403200 | -1.26 [FD2] | 2.36E-03 [FD2] | | GO:0010048 | vernalization response | 2.00E-05 | |
| FD | FD_17ZMD | AX-109360916 | 5D | 469286340 | 3.4636/-- | 3.72E-09 | EMMAX | TraesCS5D01G403200 | -1.26 [FD2] | 2.36E-03 [FD2] | | GO:0010048 | vernalization response | 3.99E-08 | |
| FD | FD_18XX | AX-109360916 | 5D | 469286340 | 3.0734/-- | 1.79E-09 | EMMAX | TraesCS5D01G403200 | -1.26 [FD2] | 2.36E-03 [FD2] | | GO:0010048 | vernalization response | 1.66E-02 | |
| FD | FD_18XX | AX-109360916 | 5D | 469286340 | -3.1094/-- | 1.22E-09 | GEMMA | TraesCS5D01G403200 | -1.26 [FD2] | 2.36E-03 [FD2] | | GO:0010048 | vernalization response | 1.66E-02 | |
| FD | FD_18XX | AX-109360916 | 5D | 469286340 | 0.9626/-- | 1.95E-78 | IIIVmrMLM | TraesCS5D01G403200 | -1.26 [FD2] | 2.36E-03 [FD2] | | GO:0010048 | vernalization response | 1.66E-02 | |
| FD | FD_17ZMD | AX-110958036 | 5D | 469179441 | 0.9247/-- | 1.08E-06 | GEMMA | TraesCS5D01G403200 | -1.26 [FD2] | 2.36E-03 [FD2] | | GO:0010048 | vernalization response | 3.99E-08 | |
| FD | FD_17ZMD | AX-109360916 | 5D | 469286340 | -3.4488/-- | 2.46E-09 | GEMMA | TraesCS5D01G403200 | -1.26 [FD2] | 2.36E-03 [FD2] | | GO:0010048 | vernalization response | 3.99E-08 | |
| FD | FD_BLUP | AX-110958036 | 5D | 469179441 | 0.2331/-- | 2.22E-05 | BLUPmrMLM | TraesCS5D01G403200 | -1.26 [FD2] | 2.36E-03 [FD2] | | GO:0010048 | vernalization response | 2.00E-05 | |
| FD | FD_17ZMD | AX-108787427 | 5D | 470769698 | -0.48/-- | 4.79E-08 | BLUPmrMLM | TraesCS5D01G403200 | -1.26 [FD2] | 2.36E-03 [FD2] | | GO:0010048 | vernalization response | 3.99E-08 | |
| FD | FD_17ZMD | AX-109360916 | 5D | 469286340 | 0.8903/-- | 5.30E-57 | IIIVmrMLM | TraesCS5D01G403200 | -1.26 [FD2] | 2.36E-03 [FD2] | | GO:0010048 | vernalization response | 3.99E-08 | |
| FD | FD_17ZMD | AX-110958036 | 5D | 469179441 | -0.9174/-- | 1.47E-06 | EMMAX | TraesCS5D01G403200 | -1.26 [FD2] | 2.36E-03 [FD2] | | GO:0010048 | vernalization response | 3.99E-08 | |
| FD | FD_17AY | AX-109068845 | 7A | 117272978 | 0.1902/2.7686 | 1.90E-09 | IIIVmrMLM | TraesCS7A01G162200 | -2.83 [FD2] | 7.29E-15 [FD2] | *OsHAL3* | | | 1.41E-05 | Su et al.,2016 |
| FD | FD_18AY | AX-111640910 | 7B | 540073505 | 0.1047/2.214 | 1.87E-06 | IIIVmrMLM | TraesCS7B01G302900 | -4.86 [FD2] | 1.88E-04 [FD2] | *OsFTIP1* | | | 4.54E-43 | Chen et al.,2021 |
| FD | FD_17ZMD | AX-95199297 | 7D | 3857123 | 0.5562/-- | 3.80E-26 | IIIVmrMLM | TraesCS7D01G003500 | 5.71 [FD2] | 3.61E-03 [FD2] | *DHD4* | | | 1.61E-04 | Cai et al.,2021 |
| FD | FD_17ZMD | AX-95199297 | 7D | 3857123 | 0.5562/-- | 3.80E-26 | IIIVmrMLM | TraesCS7D01G003500 | 5.71 [FD2] | 3.61E-03 [FD2] | *OsCCT19* | | | 1.61E-04 | Zhang et al. 2015 |
| FD | FD_18JY | AX-95191225 | 7D | 31435113 | -8.7431/-- | 1.32E-06 | EMMAX | TraesCS7D01G061300 | -3.77 [FD2] | 1.55E-06 [FD2] | *OsGBP1* | | | 4.49E-21 | Gong et al.,2018 |
| FD | FD_18XX | AX-111000473 | 7D | 138439894 | 8.7207/-- | 1.72E-09 | EMMAX | TraesCS7D01G183600 | 3.21 [FD1] | 2.23E-34 [FD1] | | GO:2000028 | regulation of photoperiodism, flowering | 8.71E-18 | |
| FD | FD_19XX | AX-110932523 | 7D | 138440518 | 10.97/-- | 3.93E-07 | EMMAX | TraesCS7D01G183600 | 3.21 [FD1] | 2.23E-34 [FD1] | | GO:2000028 | regulation of photoperiodism, flowering | 3.05E-11 | |

**Supplementary Table S24.** The genomic prediction accuracy of thirteen wheat traits using all the significant/suggested QTLs and rrBLUP with fivefold cross-validation

| Trait | Replication | | | | | | | | | | | | | | | Average |
|---|---|---|---|---|---|---|---|---|---|---|---|---|---|---|---|---|
| | 1 | 2 | 3 | 4 | 5 | 6 | 7 | 8 | 9 | 10 | 11 | 12 | 13 | 14 | 15 | |
| PH | 0.9881 | 0.9878 | 0.9880 | 0.9877 | 0.9890 | 0.9867 | 0.9890 | 0.9879 | 0.9875 | 0.9877 | 0.9887 | 0.9885 | 0.9887 | 0.9888 | 0.9881 | 0.9881 |
| SL | 0.8144 | 0.8204 | 0.8194 | 0.8172 | 0.8284 | 0.8192 | 0.8195 | 0.8217 | 0.8098 | 0.8277 | 0.8210 | 0.8318 | 0.8155 | 0.8189 | 0.8134 | 0.8199 |
| IHR | 0.6161 | 0.6245 | 0.6201 | 0.6181 | 0.6094 | 0.6414 | 0.6178 | 0.6172 | 0.6041 | 0.6120 | 0.6175 | 0.6205 | 0.6299 | 0.6290 | 0.6413 | 0.6213 |
| TGW | 0.9484 | 0.9466 | 0.9468 | 0.9482 | 0.9462 | 0.9500 | 0.9493 | 0.9495 | 0.9500 | 0.9520 | 0.9469 | 0.9474 | 0.9468 | 0.9484 | 0.9494 | 0.9484 |
| SNS | 0.7292 | 0.7209 | 0.7307 | 0.7373 | 0.7314 | 0.7318 | 0.7409 | 0.7408 | 0.7342 | 0.7337 | 0.7407 | 0.7260 | 0.7284 | 0.7286 | 0.7431 | 0.7332 |
| GL | 0.8148 | 0.8090 | 0.8061 | 0.8077 | 0.8084 | 0.8102 | 0.8084 | 0.8137 | 0.8083 | 0.7992 | 0.8029 | 0.8128 | 0.8178 | 0.8130 | 0.8097 | 0.8095 |
| GW | 0.7948 | 0.7918 | 0.7872 | 0.7956 | 0.7883 | 0.7913 | 0.7887 | 0.7957 | 0.7980 | 0.7816 | 0.7893 | 0.7896 | 0.7897 | 0.7937 | 0.7955 | 0.7914 |
| GWLR | 0.7364 | 0.7377 | 0.7333 | 0.7417 | 0.7238 | 0.7193 | 0.7293 | 0.7397 | 0.7413 | 0.7355 | 0.7373 | 0.7290 | 0.7329 | 0.7269 | 0.7428 | 0.7338 |
| V | 0.8218 | 0.8253 | 0.8330 | 0.8233 | 0.8320 | 0.8292 | 0.8279 | 0.8322 | 0.8325 | 0.8276 | 0.8291 | 0.8300 | 0.8267 | 0.8306 | 0.8198 | 0.8281 |
| T | 0.4165 | 0.3983 | 0.4007 | 0.4153 | 0.4322 | 0.4175 | 0.4128 | 0.4295 | 0.4131 | 0.4246 | 0.4039 | 0.4108 | 0.3977 | 0.4240 | 0.3948 | 0.4128 |
| Vmax | 0.8362 | 0.8325 | 0.8426 | 0.8451 | 0.8322 | 0.8452 | 0.8335 | 0.8406 | 0.8382 | 0.8301 | 0.8402 | 0.8388 | 0.8382 | 0.8381 | 0.8340 | 0.8377 |
| HD | 0.9366 | 0.9359 | 0.9396 | 0.9357 | 0.9341 | 0.9369 | 0.9387 | 0.9349 | 0.9401 | 0.9384 | 0.9384 | 0.9351 | 0.9370 | 0.9388 | 0.9371 | 0.9372 |
| FD | 0.9342 | 0.9294 | 0.9314 | 0.9338 | 0.9278 | 0.9285 | 0.9315 | 0.9327 | 0.9296 | 0.9325 | 0.9328 | 0.9274 | 0.9327 | 0.9340 | 0.9341 | 0.9315 |

PH: plant height, SL: spike length, PPR: the ratio of peduncle length to plant height, SNS: spikelet number per spike, TGW: thousand grain weight, GL: grain length, GW: grain width, GWLR: grain width/length ratio, Vmax: maximum filling rate, V: average grain filling rate, T: active filling period, HD: heading date, FD: flowering date.

**Supplementary Table S25.** Frequencies and genetic values of superior haplotypes for eighty genes associated with the trait changes between landraces and cultivars in different time periods

| Trait type | Trait | Gene | Value type | LR | 50s | 60s-70s | 80s-90s | 00s- | r | P-value |
|---|---|---|---|---|---|---|---|---|---|---|
| Morphologial traits | Plant height (cm) | | **Trait average** | **135.77** | **127.90** | **96.30** | **79.75** | **73.28** | | |
| | | TraesCS1A01G218800 | Frequency | 0.73 | 0.68 | 0.75 | 0.89 | 0.91 | -0.9035 | 3.55E-02 |
| | | | Genetic value | 135.38 | 127.91 | 100.26 | 79.86 | 73.04 | | |
| | | TraesCS1A01G263600 | Frequency | 0.10 | 0.54 | 0.82 | 0.95 | 0.98 | -0.9356 | 1.94E-02 |
| | | | Genetic value | 133.27 | 122.55 | 94.05 | 79.58 | 73.18 | | |
| | | TraesCS2A01G545400 | Frequency | 0.91 | 0.89 | 0.96 | 1.00 | 0.99 | -0.9708 | 5.97E-03 |
| | | | Genetic value | 135.54 | 127.19 | 95.12 | 79.80 | 73.20 | | |
| | | TraesCS2A01G547800 | Frequency | 0.37 | 0.62 | 0.86 | 0.86 | 0.81 | -0.8821 | 4.77E-02 |
| | | | Genetic value | 136.44 | 125.13 | 92.50 | 80.30 | 72.49 | | |
| | | TraesCS2B01G575900 | Frequency | 0.01 | 0.00 | 0.04 | 0.09 | 0.07 | -0.9394 | 1.78E-02 |
| | | | Genetic value | 137.29 | -- | 74.20 | 77.95 | 69.16 | | |
| | | TraesCS4A01G005600 | Frequency | 0.00 | 0.00 | 0.04 | 0.09 | 0.07 | -0.9346 | 1.99E-02 |
| | | | Genetic value | -- | -- | 76.17 | 79.01 | 72.75 | | |
| | | TraesCS4A01G016600 | Frequency | 0.21 | 0.57 | 0.86 | 0.95 | 0.94 | -0.9361 | 1.92E-02 |
| | | | Genetic value | 135.28 | 121.62 | 96.15 | 79.43 | 73.24 | | |
| | | TraesCS5A01G558600 | Frequency | 0.01 | 0.19 | 0.14 | 0.62 | 0.62 | -0.8796 | 4.92E-02 |
| | | | Genetic value | 131.20 | 109.27 | 97.66 | 78.59 | 73.54 | | |
| | | TraesCS5B01G232100 | Frequency | 0.01 | 0.41 | 0.64 | 0.62 | 0.76 | -0.8981 | 3.84E-02 |
| | | | Genetic value | 138.23 | 120.21 | 93.41 | 79.33 | 72.77 | | |
| | | TraesCS5B01G236500 | Frequency | 0.03 | 0.43 | 0.75 | 0.75 | 0.84 | -0.9204 | 2.66E-02 |
| | | | Genetic value | 133.95 | 120.70 | 94.17 | 79.91 | 73.13 | | |
| | | TraesCS7A01G362300 | Frequency | 0.06 | 0.51 | 0.68 | 0.83 | 0.94 | -0.9251 | 2.43E-02 |
| | | | Genetic value | 134.63 | 120.37 | 97.83 | 79.66 | 73.35 | | |
| | Ratio of peduncle length to plant height | | **Trait average** | **0.30** | **0.32** | **0.36** | **0.37** | **0.34** | | |
| | | TraesCS2A01G547800 | Frequency | 0.37 | 0.62 | 0.86 | 0.86 | 0.81 | 0.9408 | 1.71E-02 |
| | | | Genetic value | 0.31 | 0.33 | 0.36 | 0.37 | 0.34 | | |
| | | TraesCS2A01G552400 | Frequency | 0.01 | 0.08 | 0.14 | 0.11 | 0.07 | 0.8813 | 4.82E-02 |
| | | | Genetic value | 0.37 | 0.33 | 0.34 | 0.37 | 0.35 | | |
| | | TraesCS2B01G010100 | Frequency | 0.04 | 0.38 | 0.50 | 0.60 | 0.35 | 0.9435 | 1.60E-02 |
| | | | Genetic value | 0.33 | 0.34 | 0.35 | 0.36 | 0.33 | | |
| | | TraesCS4A01G016600 | Frequency | 0.21 | 0.57 | 0.86 | 0.95 | 0.94 | 0.9023 | 3.61E-02 |
| | | | Genetic value | 0.30 | 0.33 | 0.36 | 0.37 | 0.34 | | |
| | | TraesCS4B01G263400 | Frequency | 0.01 | 0.35 | 0.64 | 0.48 | 0.30 | 0.8855 | 4.57E-02 |
| | | | Genetic value | 0.33 | 0.34 | 0.36 | 0.37 | 0.35 | | |
| | | TraesCS5B01G377500 | Frequency | 0.03 | 0.38 | 0.57 | 0.60 | 0.30 | 0.9395 | 1.77E-02 |
| | | | Genetic value | 0.32 | 0.33 | 0.36 | 0.37 | 0.33 | | |
| | | | **Trait average** | **9.00** | **9.02** | **8.64** | **8.62** | **8.57** | | |
| | | TraesCS1A01G334200 | Frequency | 0.19 | 0.38 | 0.57 | 0.74 | 0.84 | -0.9193 | 2.72E-02 |

| | | | | | | | | | | |
|---|---|---|---|---|---|---|---|---|---|---|
| | | | Genetic value | 8.87 | 8.98 | 8.52 | 8.55 | 8.56 | | |
| | | TraesCS1B01G005300 | Frequency | 0.90 | 0.89 | 0.96 | 0.97 | 0.95 | -0.9480 | 1.41E-02 |
| | Spike length (cm) | | Genetic value | 8.95 | 9.06 | 8.65 | 8.64 | 8.58 | | |
| | | TraesCS4A01G016600 | Frequency | 0.21 | 0.57 | 0.86 | 0.95 | 0.94 | -0.8991 | 3.79E-02 |
| | | | Genetic value | 9.07 | 9.23 | 8.61 | 8.60 | 8.55 | | |
| | | TraesCS5D01G435800 | Frequency | 0.33 | 0.46 | 0.82 | 0.68 | 0.75 | -0.9214 | 2.62E-02 |
| | | | Genetic value | 8.82 | 9.02 | 8.55 | 8.70 | 8.51 | | |
| Yield related traits | Grain length (mm) | | **Trait average** | **6.06** | **6.38** | **6.47** | **6.65** | **6.62** | | |
| | | TraesCS2A01G010600 | Frequency | 0.60 | 0.76 | 0.96 | 0.97 | 0.99 | 0.9449 | 1.54E-02 |
| | | | Genetic value | 6.05 | 6.45 | 6.47 | 6.65 | 6.63 | | |
| | | TraesCS4B01G047400 | Frequency | 0.68 | 0.86 | 1.00 | 0.97 | 0.99 | 0.9285 | 2.27E-02 |
| | | | Genetic value | 6.03 | 6.43 | 6.47 | 6.66 | 6.63 | | |
| | | TraesCS6A01G382400 | Frequency | 0.31 | 0.65 | 0.96 | 0.94 | 0.96 | 0.9426 | 1.64E-02 |
| | | | Genetic value | 6.15 | 6.48 | 6.47 | 6.67 | 6.63 | | |
| | | TraesCS6A01G382500 | Frequency | 0.31 | 0.65 | 0.96 | 0.94 | 0.96 | 0.9426 | 1.64E-02 |
| | | | Genetic value | 6.15 | 6.48 | 6.47 | 6.67 | 6.63 | | |
| | Grain width (mm) | | **Trait average** | **3.04** | **3.13** | **3.20** | **3.31** | **3.36** | | |
| | | TraesCS1A01G247100 | Frequency | 0.16 | 0.43 | 0.82 | 0.82 | 0.84 | 0.9099 | 3.20E-02 |
| | | | Genetic value | 3.08 | 3.16 | 3.20 | 3.31 | 3.35 | | |
| | | TraesCS1A01G329900 | Frequency | 0.07 | 0.57 | 0.89 | 0.98 | 0.99 | 0.9194 | 2.71E-02 |
| | | | Genetic value | 3.07 | 3.17 | 3.20 | 3.31 | 3.36 | | |
| | | TraesCS3A01G311100 | Frequency | 0.02 | 0.51 | 0.79 | 0.92 | 0.96 | 0.9407 | 1.72E-02 |
| | | | Genetic value | 3.14 | 3.18 | 3.22 | 3.31 | 3.37 | | |
| | | TraesCS6A01G382400 | Frequency | 0.31 | 0.65 | 0.96 | 0.94 | 0.96 | 0.8821 | 4.77E-02 |
| | | | Genetic value | 3.05 | 3.16 | 3.19 | 3.32 | 3.37 | | |
| | | TraesCS6D01G012100 | Frequency | 0.55 | 0.65 | 0.75 | 0.80 | 0.86 | 0.9865 | 1.88E-03 |
| | | | Genetic value | 3.04 | 3.16 | 3.20 | 3.30 | 3.37 | | |
| | | TraesCS7B01G160000 | Frequency | 0.31 | 0.68 | 0.82 | 0.88 | 0.87 | 0.8927 | 4.15E-02 |
| | | | Genetic value | 3.06 | 3.15 | 3.21 | 3.32 | 3.36 | | |
| | Thousand grain weight (g) | | **Trait average** | **31.37** | **34.72** | **38.20** | **40.83** | **42.22** | | |
| | | TraesCS1D01G059200 | Frequency | 0.58 | 0.73 | 0.79 | 0.98 | 0.99 | 0.9776 | 4.01E-03 |
| | | | Genetic value | 31.26 | 35.79 | 38.86 | 40.91 | 42.24 | | |
| | | TraesCS2A01G547800 | Frequency | 0.37 | 0.62 | 0.86 | 0.86 | 0.81 | 0.9119 | 3.10E-02 |
| | | | Genetic value | 32.35 | 35.63 | 38.30 | 40.95 | 42.18 | | |
| | | TraesCS2B01G012500 | Frequency | 0.03 | 0.35 | 0.71 | 0.66 | 0.67 | 0.9197 | 2.70E-02 |
| | | | Genetic value | 35.63 | 37.93 | 38.73 | 40.91 | 41.99 | | |
| | | TraesCS3A01G311000 | Frequency | 0.02 | 0.49 | 0.79 | 0.92 | 0.95 | 0.9725 | 5.44E-03 |
| | | | Genetic value | 33.73 | 36.64 | 38.24 | 40.78 | 42.32 | | |
| | | TraesCS3A01G311100 | Frequency | 0.02 | 0.51 | 0.79 | 0.92 | 0.96 | 0.9690 | 6.52E-03 |
| | | | Genetic value | 33.73 | 36.52 | 38.24 | 40.78 | 42.31 | | |
| | | TraesCS3B01G252000 | Frequency | 0.07 | 0.51 | 0.68 | 0.78 | 0.83 | 0.9574 | 1.05E-02 |
| | | | Genetic value | 34.19 | 37.06 | 38.98 | 40.94 | 42.51 | | |
| | | TraesCS5B01G292100 | Frequency | 0.15 | 0.38 | 0.54 | 0.80 | 0.99 | 0.9837 | 2.49E-03 |

| | | | | | | | | | | |
|---|---|---|---|---|---|---|---|---|---|---|
| | | | Genetic value | 31.64 | 37.42 | 38.51 | 40.90 | 42.22 | | |
| | | TraesCS6A01G382500 | Frequency | 0.31 | 0.65 | 0.96 | 0.94 | 0.96 | 0.9300 | 2.20E-02 |
| | | | Genetic value | 31.59 | 35.35 | 38.12 | 41.09 | 42.32 | | |
| | | TraesCS6D01G012100 | Frequency | 0.57 | 0.65 | 0.79 | 0.83 | 0.87 | 0.9928 | 7.28E-04 |
| | | | Genetic value | 31.53 | 35.98 | 38.25 | 40.89 | 42.58 | | |
| | | TraesCS7D01G309400 | Frequency | 0.76 | 0.84 | 1.00 | 1.00 | 0.99 | 0.9300 | 2.20E-02 |
| | | | Genetic value | 31.43 | 34.76 | 38.19 | 40.81 | 42.21 | | |
| | | TraesCS7D01G525100 | Frequency | 0.83 | 0.89 | 1.00 | 0.98 | 0.97 | 0.8885 | 4.39E-02 |
| | | | Genetic value | 31.27 | 34.66 | 38.19 | 40.80 | 42.23 | | |
| | Spikelet number per spike | | **Trait average** | **17.92** | **17.52** | **16.93** | **17.28** | **17.71** | | |
| | | TraesCS7B01G271200 | Frequency | 0.97 | 0.78 | 0.36 | 0.72 | 0.76 | 0.9343 | 2.00E-02 |
| | | | Genetic value | 17.93 | 17.88 | 17.04 | 17.30 | 17.81 | | |
| Grain filling related traits | Active filling period (g/DAP) | | **Trait average** | **25.80** | **26.17** | **28.30** | **28.45** | **27.52** | | |
| | | TraesCS2B01G578500 | Frequency | 0.41 | 0.73 | 0.93 | 0.98 | 0.96 | 0.8878 | 4.44E-02 |
| | | | Genetic value | 25.82 | 26.47 | 28.45 | 28.44 | 27.59 | | |
| | | TraesCS7D01G159800 | Frequency | 0.03 | 0.41 | 0.64 | 0.63 | 0.47 | 0.8945 | 4.05E-02 |
| | | | Genetic value | 25.78 | 27.27 | 28.56 | 28.63 | 28.14 | | |
| | | TraesCS7D01G512100 | Frequency | 0.30 | 0.59 | 0.89 | 0.95 | 0.93 | 0.9197 | 2.70E-02 |
| | | | Genetic value | 25.93 | 26.50 | 28.35 | 28.53 | 27.54 | | |
| | Average grain filling rate (g/DAP) | | **Trait average** | **1.28** | **1.38** | **1.44** | **1.65** | **1.70** | | |
| | | TraesCS1A01G214700 | Frequency | 0.05 | 0.30 | 0.43 | 0.62 | 0.70 | 0.9748 | 4.79E-03 |
| | | | Genetic value | 1.42 | 1.43 | 1.37 | 1.64 | 1.70 | | |
| | | TraesCS1B01G066900 | Frequency | 0.02 | 0.08 | 0.07 | 0.40 | 0.52 | 0.9686 | 6.66E-03 |
| | | | Genetic value | 1.29 | 1.40 | 1.33 | 1.64 | 1.74 | | |
| | | TraesCS2A01G293400 | Frequency | 0.04 | 0.35 | 0.61 | 0.62 | 0.84 | 0.9044 | 3.50E-02 |
| | | | Genetic value | 1.26 | 1.40 | 1.46 | 1.66 | 1.70 | | |
| | | TraesCS2B01G202300 | Frequency | 0.03 | 0.30 | 0.21 | 0.51 | 0.50 | 0.9381 | 1.83E-02 |
| | | | Genetic value | 1.47 | 1.41 | 1.32 | 1.68 | 1.71 | | |
| | | TraesCS3B01G469000 | Frequency | 0.01 | 0.14 | 0.14 | 0.26 | 0.25 | 0.9600 | 9.54E-03 |
| | | | Genetic value | 1.41 | 1.45 | 1.40 | 1.65 | 1.69 | | |
| | | TraesCS3B01G560400 | Frequency | 0.05 | 0.05 | 0.00 | 0.25 | 0.39 | 0.8839 | 4.66E-02 |
| | | | Genetic value | 1.32 | 1.49 | -- | 1.68 | 1.71 | | |
| | | TraesCS3B01G561000 | Frequency | 0.06 | 0.05 | 0.00 | 0.26 | 0.40 | 0.8876 | 4.44E-02 |
| | | | Genetic value | 1.32 | 1.49 | -- | 1.67 | 1.70 | | |
| | | TraesCS4A01G016600 | Frequency | 0.21 | 0.57 | 0.86 | 0.95 | 0.94 | 0.8822 | 4.76E-02 |
| | | | Genetic value | 1.28 | 1.45 | 1.47 | 1.65 | 1.70 | | |
| | | TraesCS5B01G000100 | Frequency | 0.03 | 0.14 | 0.36 | 0.48 | 0.49 | 0.9476 | 1.43E-02 |
| | | | Genetic value | 1.42 | 1.46 | 1.48 | 1.67 | 1.69 | | |
| | | TraesCS6D01G403800 | Frequency | 0.02 | 0.22 | 0.43 | 0.52 | 0.47 | 0.8891 | 4.36E-02 |
| | | | Genetic value | 1.33 | 1.36 | 1.43 | 1.66 | 1.69 | | |
| | | TraesCS7A01G549300 | Frequency | 0.00 | 0.00 | 0.04 | 0.09 | 0.10 | 0.9731 | 5.26E-03 |
| | | | Genetic value | 1.38 | -- | 1.33 | 1.62 | 1.66 | | |
| | | | **Trait average** | **1.93** | **2.08** | **2.20** | **2.49** | **2.61** | | |

| | | | | | | | | | | |
|---|---|---|---|---|---|---|---|---|---|---|
| | Maximum filling rate (g/DAP) | TraesCS1A01G033500 | Frequency | 0.00 | 0.11 | 0.07 | 0.48 | 0.54 | 0.9635 | 8.32E-03 |
| | | | Genetic value | -- | 2.27 | 2.15 | 2.52 | 2.65 | | |
| | | TraesCS1A01G033600 | Frequency | 0.00 | 0.00 | 0.04 | 0.14 | 0.20 | 0.9700 | 6.20E-03 |
| | | | Genetic value | -- | -- | 2.39 | 2.51 | 2.66 | | |
| | | TraesCS1A01G219400 | Frequency | 0.01 | 0.14 | 0.32 | 0.57 | 0.59 | 0.9865 | 1.88E-03 |
| | | | Genetic value | 2.39 | 2.15 | 2.26 | 2.50 | 2.62 | | |
| | | TraesCS1B01G066900 | Frequency | 0.02 | 0.08 | 0.07 | 0.40 | 0.52 | 0.9647 | 7.93E-03 |
| | | | Genetic value | 1.92 | 2.18 | 2.15 | 2.48 | 2.66 | | |
| | | TraesCS2A01G293400 | Frequency | 0.04 | 0.35 | 0.61 | 0.62 | 0.84 | 0.9191 | 2.73E-02 |
| | | | Genetic value | 1.89 | 2.14 | 2.20 | 2.50 | 2.61 | | |
| | | TraesCS2B01G202300 | Frequency | 0.03 | 0.30 | 0.21 | 0.51 | 0.50 | 0.9279 | 2.30E-02 |
| | | | Genetic value | 2.26 | 2.15 | 2.02 | 2.54 | 2.61 | | |
| | | TraesCS3B01G469000 | Frequency | 0.01 | 0.14 | 0.14 | 0.26 | 0.25 | 0.9538 | 1.18E-02 |
| | | | Genetic value | 2.16 | 2.20 | 2.24 | 2.47 | 2.60 | | |
| | | TraesCS4A01G016600 | Frequency | 0.21 | 0.57 | 0.86 | 0.95 | 0.94 | 0.8873 | 4.46E-02 |
| | | | Genetic value | 1.93 | 2.19 | 2.24 | 2.48 | 2.60 | | |
| | | TraesCS4B01G390500 | Frequency | 0.10 | 0.41 | 0.39 | 0.82 | 0.77 | 0.9602 | 9.48E-03 |
| | | | Genetic value | 2.01 | 2.04 | 2.19 | 2.50 | 2.62 | | |
| | | TraesCS5B01G400000 | Frequency | 0.10 | 0.54 | 0.86 | 0.97 | 0.98 | 0.8911 | 4.24E-02 |
| | | | Genetic value | 1.97 | 2.18 | 2.19 | 2.48 | 2.61 | | |
| | | TraesCS6D01G403800 | Frequency | 0.02 | 0.14 | 0.32 | 0.48 | 0.43 | 0.9413 | 1.69E-02 |
| | | | Genetic value | 1.99 | 2.17 | 2.25 | 2.51 | 2.59 | | |
| | | TraesCS7B01G439900 | Frequency | 0.06 | 0.22 | 0.32 | 0.51 | 0.55 | 0.9894 | 1.30E-03 |
| | | | Genetic value | 2.07 | 2.25 | 2.25 | 2.50 | 2.64 | | |
| Growth period traits | Flowering date (g) | | **Trait average** | **195.74** | **194.49** | **192.26** | **189.95** | **190.95** | | |
| | | TraesCS1A01G372200 | Frequency | 0.19 | 0.49 | 0.82 | 0.88 | 0.88 | -0.9486 | 1.39E-02 |
| | | | Genetic value | 196.16 | 194.23 | 192.57 | 189.97 | 190.93 | | |
| | | TraesCS1D01G059200 | Frequency | 0.58 | 0.73 | 0.79 | 0.98 | 0.99 | -0.9617 | 8.95E-03 |
| | | | Genetic value | 195.74 | 194.24 | 192.02 | 189.92 | 190.98 | | |
| | | TraesCS2A01G132300 | Frequency | 0.27 | 0.49 | 0.71 | 0.74 | 0.73 | -0.9377 | 1.85E-02 |
| | | | Genetic value | 196.36 | 194.54 | 192.71 | 190.03 | 191.02 | | |
| | | TraesCS4A01G105300 | Frequency | 0.03 | 0.35 | 0.46 | 0.58 | 0.75 | -0.9023 | 3.61E-02 |
| | | | Genetic value | 197.43 | 195.21 | 192.18 | 189.65 | 191.09 | | |
| | | TraesCS4B01G317400 | Frequency | 0.01 | 0.08 | 0.21 | 0.26 | 0.32 | -0.9510 | 1.29E-02 |
| | | | Genetic value | 195.67 | 196.72 | 192.25 | 191.48 | 191.05 | | |
| | | TraesCS5D01G341000 | Frequency | 0.53 | 0.78 | 1.00 | 0.97 | 1.00 | -0.8900 | 4.30E-02 |
| | | | Genetic value | 195.75 | 194.28 | 192.26 | 189.97 | 190.96 | | |
| | | TraesCS7A01G162200 | Frequency | 0.89 | 0.92 | 0.96 | 1.00 | 1.00 | -0.9891 | 1.37E-03 |
| | | | Genetic value | 195.71 | 194.62 | 192.15 | 189.93 | 190.96 | | |
| | | TraesCS7D01G061300 | Frequency | 0.44 | 0.70 | 0.96 | 0.97 | 0.95 | -0.9193 | 2.72E-02 |
| | | | Genetic value | 195.67 | 194.43 | 192.26 | 189.88 | 191.03 | | |
| | | TraesCS7D01G183600 | Frequency | 0.01 | 0.24 | 0.36 | 0.55 | 0.44 | -0.9728 | 5.35E-03 |
| | | | Genetic value | 194.27 | 193.44 | 192.02 | 189.72 | 190.77 | | |

| | | | | | | | | | | |
|---|---|---|---|---|---|---|---|---|---|---|
| | Heading date (g) | | **Trait average** | **190.00** | **188.56** | **186.49** | **184.11** | **184.89** | | |
| | | TraesCS6B01G054100 | Frequency | 0.18 | 0.24 | 0.61 | 0.68 | 0.52 | -0.9230 | 2.54E-02 |
| | | | Genetic value | 190.05 | 187.48 | 186.05 | 183.97 | 184.78 | | |
| | | TraesCS7D01G107500 | Frequency | 0.85 | 0.86 | 0.93 | 0.98 | 0.92 | -0.9339 | 2.02E-02 |
| | | | Genetic value | 190.10 | 188.79 | 186.33 | 184.11 | 184.84 | | |

Genetic value: the average trait value of individuals with the superior haplotype during the corresponding period.

**Supplementary Table S26**. Frequencies and genetic values of superior genotypes for 50 loci associated with the trait changes between landraces and cultivars in different periods

| Trait type | Trait | SNP | Superior genotype | Value type | LR | 50s | 60s-70s | 80s-90s | 00s- | r | P-value |
|---|---|---|---|---|---|---|---|---|---|---|---|
| Morphologial traits | Plant height (cm) | | | **Trait average** | **135.77** | **127.90** | **96.30** | **79.75** | **73.28** | | |
| | | AX-94728034 | 1 | Frequency | 0.14 | 0.16 | 0.29 | 0.58 | 0.81 | -0.9215 | 2.61E-02 |
| | | AX-110018488 | 2 | Frequency | 0.01 | 0.16 | 0.36 | 0.88 | 0.99 | -0.9598 | 9.60E-03 |
| | | AX-109434530 | 1 | Frequency | 0.00 | 0.00 | 0.14 | 0.17 | 0.12 | -0.9069 | 3.36E-02 |
| | | AX-109355416 | 1 | Frequency | 0.02 | 0.41 | 0.79 | 0.97 | 1.00 | -0.9680 | 6.83E-03 |
| | | AX-95240191 | 0 | Frequency | 0.19 | 0.57 | 0.82 | 0.97 | 0.92 | -0.9292 | 2.24E-02 |
| | | AX-110814077 | 1 | Frequency | 0.02 | 0.24 | 0.75 | 0.71 | 0.80 | -0.9518 | 1.26E-02 |
| | | AX-109325020 | 2 | Frequency | 0.02 | 0.16 | 0.43 | 0.86 | 0.99 | -0.9780 | 3.91E-03 |
| | | AX-108738714 | 0 | Frequency | 0.99 | 0.89 | 0.75 | 0.49 | 0.39 | 0.9717 | 5.68E-03 |
| | Spike length (cm) | | | **Trait average** | **9.00** | **9.02** | **8.64** | **8.62** | **8.57** | | |
| | | AX-110386923 | 0 | Frequency | 0.96 | 0.95 | 1.00 | 1.00 | 1.00 | -0.9863 | 1.92E-03 |
| | Ratio of peduncle length to plant height | | | **Trait average** | **0.30** | **0.32** | **0.36** | **0.37** | **0.34** | | |
| | | AX-109838169 | 1 | Frequency | 0.01 | 0.11 | 0.36 | 0.34 | 0.17 | 0.9792 | 3.59E-03 |
| | | AX-110594254 | 1 | Frequency | 0.01 | 0.22 | 0.50 | 0.37 | 0.17 | 0.8903 | 4.29E-02 |
| | | AX-108894246 | 2 | Frequency | 0.05 | 0.51 | 0.82 | 0.78 | 0.48 | 0.9458 | 1.50E-02 |
| | | AX-108925664 | 2 | Frequency | 0.14 | 0.43 | 0.64 | 0.75 | 0.76 | 0.8825 | 4.75E-02 |
| | | AX-110362233 | 2 | Frequency | 0.04 | 0.30 | 0.36 | 0.48 | 0.30 | 0.9317 | 2.12E-02 |
| | | AX-110461119 | 2 | Frequency | 0.05 | 0.27 | 0.57 | 0.58 | 0.38 | 0.9929 | 7.20E-04 |
| | | AX-110374260 | 0 | Frequency | 0.38 | 0.70 | 0.93 | 0.97 | 0.99 | 0.8847 | 4.62E-02 |
| Yield related traits | Thousand grain weight (g) | | | **Trait average** | **31.37** | **34.72** | **38.20** | **40.83** | **42.22** | | |
| | | AX-95082229 | 1 | Frequency | 0.01 | 0.27 | 0.39 | 0.48 | 0.65 | 0.9794 | 3.54E-03 |
| | | AX-110675322 | 0 | Frequency | 0.10 | 0.43 | 0.43 | 0.54 | 0.63 | 0.9313 | 2.14E-02 |
| | | AX-110990714 | 0 | Frequency | 0.05 | 0.54 | 0.61 | 0.72 | 0.81 | 0.9368 | 1.89E-02 |
| | | AX-108739437 | 2 | Frequency | 0.22 | 0.32 | 0.43 | 0.63 | 0.71 | 0.9787 | 3.72E-03 |
| | | AX-109423631 | 0 | Frequency | 0.03 | 0.41 | 0.89 | 0.94 | 0.92 | 0.9532 | 1.21E-02 |
| | | AX-111077704 | 2 | Frequency | 0.04 | 0.38 | 0.39 | 0.69 | 0.74 | 0.9642 | 8.09E-03 |
| | | AX-108817049 | 0 | Frequency | 0.25 | 0.51 | 0.68 | 0.78 | 0.79 | 0.9791 | 3.63E-03 |
| | | AX-108957255 | 2 | Frequency | 0.01 | 0.41 | 0.57 | 0.66 | 0.73 | 0.9668 | 7.24E-03 |
| | Spikelet number per spike | | | **Trait average** | **17.92** | **17.52** | **16.93** | **17.28** | **17.71** | | |
| | | AX-94465953 | 0 | Frequency | 0.97 | 0.84 | 0.54 | 0.60 | 0.76 | 0.9278 | 2.30E-02 |
| | | AX-110457590 | 0 | Frequency | 0.99 | 0.84 | 0.50 | 0.66 | 0.73 | 0.9139 | 2.99E-02 |
| | Grain length (mm) | | | **Trait average** | **6.06** | **6.38** | **6.47** | **6.65** | **6.62** | | |
| | | AX-111570261 | 2 | Frequency | 0.07 | 0.59 | 0.82 | 0.92 | 0.93 | 0.9821 | 2.88E-03 |
| | Grain width (mm) | | | **Trait average** | **3.04** | **3.13** | **3.20** | **3.31** | **3.36** | | |
| | | AX-111460358 | 0 | Frequency | 0.94 | 0.95 | 1.00 | 1.00 | 1.00 | 0.8798 | 4.91E-02 |
| | | AX-110922076 | 1 | Frequency | 0.01 | 0.11 | 0.07 | 0.23 | 0.36 | 0.9238 | 2.50E-02 |

| | | | | | | | | | | | |
|---|---|---|---|---|---|---|---|---|---|---|---|
| | | AX-95242102 | 2 | Frequency | 0.00 | 0.08 | 0.07 | 0.43 | 0.54 | 0.9355 | 1.95E-02 |
| | | AX-111055136 | 2 | Frequency | 0.02 | 0.32 | 0.79 | 0.83 | 0.94 | 0.9492 | 1.36E-02 |
| Grain filling related traits | Average grain filling rate (g/DAP) | | | **Trait average** | **1.28** | **1.38** | **1.44** | **1.65** | **1.70** | | |
| | | AX-110996750 | 1 | Frequency | 0.06 | 0.11 | 0.14 | 0.43 | 0.64 | 0.9598 | 9.62E-03 |
| | | AX-111101313 | 2 | Frequency | 0.00 | 0.00 | 0.00 | 0.22 | 0.22 | 0.9426 | 1.64E-02 |
| | | AX-111559203 | 2 | Frequency | 0.10 | 0.16 | 0.14 | 0.45 | 0.42 | 0.9593 | 9.79E-03 |
| | | AX-94971846 | 1 | Frequency | 0.09 | 0.27 | 0.39 | 0.48 | 0.49 | 0.9261 | 2.39E-02 |
| | | AX-109100815 | 1 | Frequency | 0.03 | 0.08 | 0.25 | 0.38 | 0.30 | 0.9085 | 3.27E-02 |
| | | AX-110961703 | 2 | Frequency | 0.02 | 0.24 | 0.57 | 0.62 | 0.80 | 0.9255 | 2.41E-02 |
| | Active filling period (g/DAP) | | | **Trait average** | **25.80** | **26.17** | **28.30** | **28.45** | **27.52** | | |
| | | AX-110942379 | 0 | Frequency | 0.82 | 0.89 | 1.00 | 1.00 | 0.99 | 0.9548 | 1.15E-02 |
| | | AX-111536488 | 0 | Frequency | 0.13 | 0.57 | 0.86 | 1.00 | 1.00 | 0.8785 | 4.99E-02 |
| | | AX-109346027 | 2 | Frequency | 0.04 | 0.46 | 0.96 | 0.82 | 0.89 | 0.9029 | 3.58E-02 |
| | Maximum filling rate (g/DAP) | | | **Trait average** | **1.93** | **2.08** | **2.20** | **2.49** | **2.61** | | |
| | | AX-111101313 | 2 | Frequency | 0.00 | 0.00 | 0.00 | 0.22 | 0.22 | 0.9310 | 2.15E-02 |
| | | AX-110961703 | 2 | Frequency | 0.02 | 0.24 | 0.57 | 0.62 | 0.80 | 0.9386 | 1.81E-02 |
| | | AX-111065061 | 0 | Frequency | 0.42 | 0.51 | 0.54 | 0.68 | 0.81 | 0.9832 | 2.61E-03 |
| | | AX-109338715 | 0 | Frequency | 0.78 | 0.73 | 0.75 | 0.91 | 0.98 | 0.8922 | 4.18E-02 |
| Growth period traits | Heading date (d) | | | **Trait average** | **190.00** | **188.56** | **186.49** | **184.11** | **184.89** | | |
| | | AX-110932982 | 1 | Frequency | 0.01 | 0.03 | 0.04 | 0.15 | 0.19 | -0.8801 | 4.89E-02 |
| | | AX-109767598 | 2 | Frequency | 0.02 | 0.46 | 0.89 | 0.97 | 0.99 | -0.9482 | 1.40E-02 |
| | | AX-110079714 | 1 | Frequency | 0.10 | 0.30 | 0.39 | 0.66 | 0.79 | -0.9461 | 1.49E-02 |
| | Flowering date (d) | | | **Trait average** | **195.74** | **194.49** | **192.26** | **189.95** | **190.95** | | |
| | | AX-109584296 | 0 | Frequency | 0.61 | 0.68 | 0.68 | 0.82 | 0.86 | -0.8903 | 4.29E-02 |
| | | AX-111088141 | 2 | Frequency | 0.08 | 0.49 | 0.64 | 0.74 | 0.73 | -0.9154 | 2.92E-02 |
| | | AX-110390529 | 2 | Frequency | 0.01 | 0.19 | 0.39 | 0.49 | 0.63 | -0.9371 | 1.87E-02 |

**Supplementary Table S27**. Gene superior haplotypes in 30 varieties with the top 5% SNS, V and Vmax phenotypes

| Trait | Accession | Count | Gene |
|---|---|---|---|
| SNS | HXM181 | 1 | TraesCS7B01G271200 |
| SNS | HXM286 | 1 | TraesCS7B01G271200 |
| SNS | HXZ171 | 1 | TraesCS7B01G271200 |
| SNS | HXM314 | 1 | TraesCS7B01G271200 |
| SNS | HXM280 | 1 | TraesCS7B01G271200 |
| SNS | HXZ168 | 1 | TraesCS7B01G271200 |
| SNS | HXM523 | 0 | |
| SNS | HXM220 | 1 | TraesCS7B01G271200 |
| SNS | HXZ362 | 1 | TraesCS7B01G271200 |
| SNS | HXZ263 | 1 | TraesCS7B01G271200 |
| SNS | HXM369 | 1 | TraesCS7B01G271200 |
| SNS | HXZ167 | 1 | TraesCS7B01G271200 |
| SNS | HXM294 | 1 | TraesCS7B01G271200 |
| SNS | HXM483 | 1 | TraesCS7B01G271200 |
| SNS | HXM130 | 1 | TraesCS7B01G271200 |
| SNS | HXM382 | 1 | TraesCS7B01G271200 |
| SNS | HXM277 | 1 | TraesCS7B01G271200 |
| SNS | HXM500 | 0 | |
| SNS | HXM570 | 1 | TraesCS7B01G271200 |
| SNS | HXZ457 | 0 | |
| SNS | HXM635 | 1 | TraesCS7B01G271200 |
| SNS | HXM199 | 1 | TraesCS7B01G271200 |
| SNS | HXM285 | 1 | TraesCS7B01G271200 |
| SNS | HXZ361 | 1 | TraesCS7B01G271200 |
| SNS | HXZ165 | 1 | TraesCS7B01G271200 |
| SNS | HXM36 | 1 | TraesCS7B01G271200 |
| SNS | HXM628 | 1 | TraesCS7B01G271200 |
| SNS | HXM334 | 1 | TraesCS7B01G271200 |
| SNS | HXM398 | 1 | TraesCS7B01G271200 |
| SNS | HXZ363 | 1 | TraesCS7B01G271200 |
| V | HXM569 | 7 | TraesCS1A01G214700/TraesCS1B01G066900/TraesCS2A01G293400/TraesCS2B01G202300/TraesCS4A01G016600/TraesCS5B01G000100/TraesCS6D01G403800 |
| V | HXM571 | 6 | TraesCS1A01G214700/TraesCS1B01G066900/TraesCS2A01G293400/TraesCS2B01G202300/TraesCS4A01G016600/TraesCS5B01G000100 |
| V | HXM523 | 9 | TraesCS1A01G214700/TraesCS1B01G066900/TraesCS2B01G202300/TraesCS3B01G469000/TraesCS3B01G560400/TraesCS3B01G561000/TraesCS4A01G016600/TraesCS5B01G000100/TraesCS6D01G403800 |
| V | HXM593 | 4 | TraesCS1B01G066900/TraesCS2A01G293400/TraesCS3B01G469000/TraesCS4A01G016600 |

| | | | |
|---|---|---|---|
| V | HXM619 | 6 | TraesCS1B01G066900/TraesCS2A01G293400/TraesCS2B01G202300/TraesCS4A01G016600/TraesCS5B01G000100/TraesCS6D01G403800 |
| V | HXM592 | 5 | TraesCS1A01G214700/TraesCS1B01G066900/TraesCS3B01G560400/TraesCS3B01G561000/TraesCS4A01G016600 |
| V | HXM445 | 6 | TraesCS2A01G293400/TraesCS2B01G202300/TraesCS3B01G469000/TraesCS4A01G016600/TraesCS5B01G000100/TraesCS6D01G403800 |
| V | HXZ556 | 7 | TraesCS1A01G214700/TraesCS1B01G066900/TraesCS2A01G293400/TraesCS3B01G560400/TraesCS3B01G561000/TraesCS4A01G016600/TraesCS5B01G000100 |
| V | HXM465 | 7 | TraesCS1A01G214700/TraesCS2A01G293400/TraesCS2B01G202300/TraesCS3B01G560400/TraesCS3B01G561000/TraesCS4A01G016600/TraesCS5B01G000100 |
| V | HXM535 | 6 | TraesCS1A01G214700/TraesCS1B01G066900/TraesCS2B01G202300/TraesCS3B01G560400/TraesCS3B01G561000/TraesCS4A01G016600 |
| V | HXZ458 | 6 | TraesCS1A01G214700/TraesCS1B01G066900/TraesCS2A01G293400/TraesCS2B01G202300/TraesCS4A01G016600/TraesCS6D01G403800 |
| V | HXM588 | 5 | TraesCS1A01G214700/TraesCS2A01G293400/TraesCS2B01G202300/TraesCS4A01G016600/TraesCS5B01G000100 |
| V | HXZ464 | 2 | TraesCS4A01G016600/TraesCS6D01G403800 |
| V | HXM594 | 3 | TraesCS1B01G066900/TraesCS2A01G293400/TraesCS4A01G016600 |
| V | HXM473 | 8 | TraesCS1A01G214700/TraesCS2A01G293400/TraesCS2B01G202300/TraesCS3B01G560400/TraesCS3B01G561000/TraesCS4A01G016600/TraesCS5B01G000100/TraesCS6D01G403800 |
| V | HXM500 | 10 | TraesCS1A01G214700/TraesCS1B01G066900/TraesCS2A01G293400/TraesCS2B01G202300/TraesCS3B01G469000/TraesCS3B01G560400/TraesCS3B01G561000/TraesCS4A01G016600/TraesCS5B01G000100/TraesCS6D01G403800 |
| V | HXM541 | 8 | TraesCS1B01G066900/TraesCS2A01G293400/TraesCS2B01G202300/TraesCS3B01G560400/TraesCS3B01G561000/TraesCS4A01G016600/TraesCS5B01G000100/TraesCS6D01G403800 |
| V | HXM542 | 4 | TraesCS1A01G214700/TraesCS1B01G066900/TraesCS2A01G293400/TraesCS4A01G016600 |
| V | HXM618 | 8 | TraesCS1A01G214700/TraesCS1B01G066900/TraesCS2A01G293400/TraesCS2B01G202300/TraesCS3B01G469000/TraesCS3B01G560400/TraesCS3B01G561000/TraesCS4A01G016600 |
| V | HXM526 | 4 | TraesCS1A01G214700/TraesCS2B01G202300/TraesCS4A01G016600/TraesCS5B01G000100 |
| V | HXM521 | 5 | TraesCS1A01G214700/TraesCS2A01G293400/TraesCS2B01G202300/TraesCS3B01G469000/TraesCS4A01G016600 |
| V | HXM609 | 9 | TraesCS1B01G066900/TraesCS2A01G293400/TraesCS2B01G202300/TraesCS3B01G469000/TraesCS3B01G560400/TraesCS3B01G561000/TraesCS4A01G016600/TraesCS5B01G000100/TraesCS6D01G403800 |
| V | HXM610 | 5 | TraesCS1A01G214700/TraesCS2A01G293400/TraesCS3B01G469000/TraesCS4A01G016600/TraesCS6D01G403800 |
| V | HXM421 | 2 | TraesCS1A01G214700/TraesCS4A01G016600 |
| V | HXM527 | 4 | TraesCS1A01G214700/TraesCS2A01G293400/TraesCS4A01G016600/TraesCS5B01G000100 |

| | | | |
|---|---|---|---|
| V | HXM471 | 4 | TraesCS1A01G214700/TraesCS2B01G202300/TraesCS4A01G016600/TraesCS5B01G000100 |
| V | HXM438 | 3 | TraesCS1B01G066900/TraesCS2A01G293400/TraesCS2B01G202300 |
| V | HXM475 | 6 | TraesCS1A01G214700/TraesCS1B01G066900/TraesCS2A01G293400/TraesCS3B01G469000/TraesCS4A01G016600/TraesCS6D01G403800 |
| V | HXM499 | 6 | TraesCS1A01G214700/TraesCS1B01G066900/TraesCS2A01G293400/TraesCS3B01G560400/TraesCS3B01G561000/TraesCS4A01G016600 |
| V | HXM515 | 8 | TraesCS1A01G214700/TraesCS2A01G293400/TraesCS2B01G202300/TraesCS3B01G560400/TraesCS3B01G561000/TraesCS4A01G016600/TraesCS5B01G000100/TraesCS6D01G403800 |
| Vmax | HXM571 | 9 | TraesCS1A01G033500/TraesCS1A01G219400/TraesCS1B01G066900/TraesCS2A01G293400/TraesCS2B01G202300/TraesCS4A01G016600/TraesCS4B01G390500/TraesCS5B01G400000/TraesCS7B01G439900 |
| Vmax | HXM569 | 8 | TraesCS1A01G033500/TraesCS1A01G219400/TraesCS1B01G066900/TraesCS2A01G293400/TraesCS2B01G202300/TraesCS4A01G016600/TraesCS4B01G390500/TraesCS5B01G400000 |
| Vmax | HXM568 | 9 | TraesCS1A01G033500/TraesCS1A01G219400/TraesCS1B01G066900/TraesCS2A01G293400/TraesCS4A01G016600/TraesCS4B01G390500/TraesCS5B01G400000/TraesCS6D01G403800/TraesCS7B01G439900 |
| Vmax | HXM593 | 10 | TraesCS1A01G033500/TraesCS1A01G033600/TraesCS1A01G219400/TraesCS1B01G066900/TraesCS2A01G293400/TraesCS3B01G469000/TraesCS4A01G016600/TraesCS4B01G390500/TraesCS5B01G400000/TraesCS7B01G439900 |
| Vmax | HXM624 | 9 | TraesCS1A01G033500/TraesCS1A01G219400/TraesCS1B01G066900/TraesCS2A01G293400/TraesCS4A01G016600/TraesCS4B01G390500/TraesCS5B01G400000/TraesCS6D01G403800/TraesCS7B01G439900 |
| Vmax | HXZ557 | 6 | TraesCS1A01G219400/TraesCS2A01G293400/TraesCS3B01G469000/TraesCS4A01G016600/TraesCS4B01G390500/TraesCS5B01G400000 |
| Vmax | HXM523 | 9 | TraesCS1A01G033500/TraesCS1A01G219400/TraesCS1B01G066900/TraesCS2B01G202300/TraesCS3B01G469000/TraesCS4A01G016600/TraesCS4B01G390500/TraesCS5B01G400000/TraesCS6D01G403800 |
| Vmax | HXM572 | 9 | TraesCS1A01G033500/TraesCS1A01G219400/TraesCS1B01G066900/TraesCS2A01G293400/TraesCS4A01G016600/TraesCS4B01G390500/TraesCS5B01G400000/TraesCS6D01G403800/TraesCS7B01G439900 |
| Vmax | HXM465 | 7 | TraesCS1A01G033500/TraesCS1A01G219400/TraesCS2A01G293400/TraesCS2B01G202300/TraesCS4A01G016600/TraesCS4B01G390500/TraesCS5B01G400000 |
| Vmax | HXM609 | 11 | TraesCS1A01G033500/TraesCS1A01G219400/TraesCS1B01G066900/TraesCS2A01G293400/TraesCS2B01G202300/TraesCS3B01G469000/TraesCS4A01G016600/TraesCS4B01G390500/TraesCS5B01G400000/TraesCS6D01G403800/TraesCS7B01G439900 |
| Vmax | HXM619 | 9 | TraesCS1A01G033500/TraesCS1B01G066900/TraesCS2A01G293400/TraesCS2B01G202300/TraesCS4A01G016600/TraesCS4B01G390500/TraesCS5B01G400000/TraesCS6D01G403800/TraesCS7B01G439900 |
| Vmax | HXM541 | 10 | TraesCS1A01G033500/TraesCS1A01G033600/TraesCS1B01G066900/TraesCS2A01G293400/TraesCS2B01G202300/TraesCS4A01G016600/TraesCS4B01G390500/TraesCS5B01G400000/TraesCS6D01G403800/TraesCS7B01G439900 |

| | | | |
|---|---|---|---|
| Vmax | HXM592 | 8 | TraesCS1A01G033500/TraesCS1A01G033600/TraesCS1A01G219400/TraesCS1B01G066900/TraesCS4A01G016600/TraesCS4B01G390500/TraesCS5B01G400000/TraesCS7B01G439900 |
| Vmax | HXM220 | 0 | |
| Vmax | HXM445 | 9 | TraesCS1A01G033500/TraesCS2A01G293400/TraesCS2B01G202300/TraesCS3B01G469000/TraesCS4A01G016600/TraesCS4B01G390500/TraesCS5B01G400000/TraesCS6D01G403800/TraesCS7B01G439900 |
| Vmax | HXZ556 | 8 | TraesCS1A01G033500/TraesCS1A01G219400/TraesCS1B01G066900/TraesCS2A01G293400/TraesCS4A01G016600/TraesCS4B01G390500/TraesCS5B01G400000/TraesCS7B01G439900 |
| Vmax | HXM535 | 9 | TraesCS1A01G033500/TraesCS1A01G033600/TraesCS1A01G219400/TraesCS1B01G066900/TraesCS2B01G202300/TraesCS4A01G016600/TraesCS4B01G390500/TraesCS5B01G400000/TraesCS7B01G439900 |
| Vmax | HXZ458 | 10 | TraesCS1A01G033500/TraesCS1A01G219400/TraesCS1B01G066900/TraesCS2A01G293400/TraesCS2B01G202300/TraesCS4A01G016600/TraesCS4B01G390500/TraesCS5B01G400000/TraesCS6D01G403800/TraesCS7B01G439900 |
| Vmax | HXM588 | 5 | TraesCS2A01G293400/TraesCS2B01G202300/TraesCS4A01G016600/TraesCS4B01G390500/TraesCS5B01G400000 |
| Vmax | HXM608 | 6 | TraesCS2B01G202300/TraesCS4A01G016600/TraesCS4B01G390500/TraesCS5B01G400000/TraesCS6D01G403800/TraesCS7B01G439900 |
| Vmax | HXZ464 | 6 | TraesCS1A01G219400/TraesCS4A01G016600/TraesCS4B01G390500/TraesCS5B01G400000/TraesCS6D01G403800/TraesCS7B01G439900 |
| Vmax | HXM594 | 8 | TraesCS1A01G033500/TraesCS1A01G033600/TraesCS1B01G066900/TraesCS2A01G293400/TraesCS4A01G016600/TraesCS4B01G390500/TraesCS5B01G400000/TraesCS7B01G439900 |
| Vmax | HXM473 | 7 | TraesCS1A01G219400/TraesCS2A01G293400/TraesCS2B01G202300/TraesCS4A01G016600/TraesCS4B01G390500/TraesCS5B01G400000/TraesCS6D01G403800 |
| Vmax | HXM500 | 10 | TraesCS1A01G033500/TraesCS1A01G219400/TraesCS1B01G066900/TraesCS2A01G293400/TraesCS2B01G202300/TraesCS3B01G469000/TraesCS4A01G016600/TraesCS4B01G390500/TraesCS5B01G400000/TraesCS6D01G403800 |
| Vmax | HXM575 | 6 | TraesCS1A01G033500/TraesCS2A01G293400/TraesCS2B01G202300/TraesCS4A01G016600/TraesCS4B01G390500/TraesCS5B01G400000 |
| Vmax | HXM614 | 5 | TraesCS2A01G293400/TraesCS4A01G016600/TraesCS4B01G390500/TraesCS5B01G400000/TraesCS7B01G439900 |
| Vmax | HXM618 | 10 | TraesCS1A01G033500/TraesCS1A01G033600/TraesCS1B01G066900/TraesCS2A01G293400/TraesCS2B01G202300/TraesCS3B01G469000/TraesCS4A01G016600/TraesCS4B01G390500/TraesCS5B01G400000/TraesCS7B01G439900 |
| Vmax | HXM542 | 8 | TraesCS1A01G033500/TraesCS1A01G219400/TraesCS1B01G066900/TraesCS2A01G293400/TraesCS4A01G016600/TraesCS4B01G390500/TraesCS5B01G400000/TraesCS7B01G439900 |
| Vmax | HXM472 | 5 | TraesCS2B01G202300/TraesCS4A01G016600/TraesCS4B01G390500/TraesCS5B01G400000/TraesCS6D01G403800 |
| Vmax | HXM596 | 7 | TraesCS1A01G219400/TraesCS2A01G293400/TraesCS2B01G202300/TraesCS3B01G469000/TraesCS4A01G016600/TraesCS5B01G400000/TraesCS7B01G439900 |

SNS: spikelet number per spike, Vmax: maximum filling rate, V: average grain filling rate.

**Supplementary Table S28**. Transmission of superior haplotypes of HXM497 (Zhoumai 17) in its varietal pedigree

| Gene | HXM416 (Zhengzhou 761) | HXM423 (Yanshi 4) | HXM444 (Yumai 18) | HXM446 (Yumai 21) | HXM574 (Zhou 8425B) |
|---|---|---|---|---|---|
| TraesCS1A01G033500 | | | | 1 | |
| TraesCS1A01G214700 | 1 | | | | |
| TraesCS1A01G218800 | | 1 | | 1 | 1 |
| TraesCS1A01G219400 | | 1 | | 1 | |
| TraesCS1A01G263600 | 1 | | 1 | 1 | 1 |
| TraesCS1A01G329900 | 1 | 1 | 1 | 1 | 1 |
| TraesCS1A01G334200 | 1 | 1 | | 1 | |
| TraesCS1A01G372200 | 1 | | 1 | 1 | |
| TraesCS1B01G005300 | | | | 1 | 1 |
| TraesCS1B01G066900 | | | | 1 | 1 |
| TraesCS1D01G059200 | | 1 | 1 | 1 | 1 |
| TraesCS2A01G010600 | 1 | 1 | 1 | 1 | 1 |
| TraesCS2A01G132300 | | | | | 1 |
| TraesCS2A01G293400 | | | | | 1 |
| TraesCS2A01G545400 | 1 | 1 | 1 | 1 | 1 |
| TraesCS2A01G547800 | 1 | 1 | 1 | 1 | |
| TraesCS2B01G012500 | | | 1 | | |
| TraesCS2B01G578500 | 1 | 1 | 1 | 1 | |
| TraesCS3A01G311000 | 1 | 1 | 1 | 1 | 1 |
| TraesCS3A01G311100 | 1 | 1 | 1 | 1 | 1 |
| TraesCS3B01G252000 | | 1 | 1 | 1 | 1 |
| TraesCS4A01G005600 | 1 | | | | |
| TraesCS4A01G016600 | 1 | 1 | 1 | 1 | 1 |
| TraesCS4A01G105300 | 1 | 1 | 1 | 1 | 1 |
| TraesCS4B01G047400 | 1 | 1 | 1 | 1 | 1 |
| TraesCS4B01G390500 | | | | 1 | |
| TraesCS5A01G558600 | | | | 1 | 1 |
| TraesCS5B01G000100 | | | | | |
| TraesCS5B01G292100 | 1 | 1 | 1 | 1 | 1 |
| TraesCS5B01G400000 | 1 | 1 | 1 | 1 | 1 |
| TraesCS5D01G341000 | 1 | 1 | 1 | 1 | 1 |
| TraesCS5D01G435800 | 1 | 1 | 1 | 1 | 1 |
| TraesCS6A01G382400 | 1 | 1 | 1 | 1 | 1 |
| TraesCS6A01G382500 | 1 | 1 | 1 | 1 | 1 |
| TraesCS6D01G012100 | 1 | | | 1 | 1 |
| TraesCS6D01G403800 | | | 1 | | 1 |
| TraesCS7A01G162200 | 1 | 1 | 1 | 1 | 1 |
| TraesCS7A01G362300 | | 1 | 1 | 1 | 1 |
| TraesCS7A01G549300 | | | | | 1 |

| | | | | | |
|---|---|---|---|---|---|
| TraesCS7B01G160000 | | 1 | 1 | 1 | |
| TraesCS7B01G271200 | 1 | | 1 | 1 | |
| TraesCS7D01G061300 | 1 | 1 | 1 | 1 | 1 |
| TraesCS7D01G107500 | 1 | 1 | 1 | 1 | 1 |
| TraesCS7D01G159800 | 1 | 1 | 1 | 1 | |
| TraesCS7D01G309400 | 1 | 1 | 1 | 1 | 1 |
| TraesCS7D01G512100 | | 1 | 1 | 1 | 1 |
| TraesCS7D01G525100 | 1 | 1 | 1 | 1 | 1 |

**Supplementary Table S29.** Transmission of superior haplotypes of HXM518 (Zhengmai 7698) in its varietal pedigree

| Gene | HXM446 (Yumai 21) | HXM496 (Zhoumai 16) | HXM516 (Zhengmai 9405) | HXM573 (4B269) | HXM574 (Zhou 8425B) |
|---|---|---|---|---|---|
| TraesCS1A01G033500 | | 1 | | 1 | 1 |
| TraesCS1A01G214700 | | | | 1 | |
| TraesCS1A01G218800 | 1 | 1 | 1 | 1 | 1 |
| TraesCS1A01G219400 | 1 | 1 | 1 | | |
| TraesCS1A01G247100 | | 1 | 1 | 1 | 1 |
| TraesCS1A01G329900 | | | | 1 | |
| TraesCS1A01G334200 | 1 | 1 | | 1 | |
| TraesCS1A01G372200 | 1 | 1 | | 1 | |
| TraesCS1B01G005300 | 1 | 1 | | 1 | 1 |
| TraesCS1B01G066900 | 1 | 1 | | 1 | 1 |
| TraesCS1D01G059200 | 1 | 1 | 1 | 1 | 1 |
| TraesCS2A01G010600 | 1 | 1 | 1 | 1 | 1 |
| TraesCS2A01G293400 | 1 | 1 | | 1 | |
| TraesCS2A01G545400 | 1 | 1 | 1 | 1 | 1 |
| TraesCS2A01G547800 | 1 | 1 | 1 | 1 | |
| TraesCS2B01G012500 | 1 | 1 | | 1 | |
| TraesCS2B01G578500 | 1 | 1 | 1 | 1 | |
| TraesCS3A01G311000 | 1 | 1 | 1 | 1 | 1 |
| TraesCS3A01G311100 | 1 | 1 | 1 | 1 | 1 |
| TraesCS3B01G252000 | 1 | 1 | 1 | 1 | 1 |
| TraesCS3B01G560400 | | 1 | | | 1 |
| TraesCS3B01G561000 | | 1 | | | 1 |
| TraesCS4A01G005600 | | | | | |
| TraesCS4A01G016600 | 1 | 1 | 1 | 1 | 1 |
| TraesCS4A01G105300 | 1 | 1 | | | 1 |
| TraesCS4B01G047400 | 1 | 1 | 1 | 1 | 1 |
| TraesCS4B01G390500 | 1 | 1 | | 1 | |
| TraesCS5A01G558600 | 1 | 1 | 1 | 1 | 1 |
| TraesCS5B01G232100 | | 1 | 1 | | 1 |
| TraesCS5B01G236500 | | 1 | 1 | 1 | 1 |
| TraesCS5B01G292100 | 1 | 1 | 1 | 1 | 1 |
| TraesCS5B01G400000 | 1 | 1 | 1 | 1 | 1 |
| TraesCS5D01G341000 | 1 | 1 | 1 | 1 | 1 |
| TraesCS5D01G435800 | 1 | 1 | 1 | 1 | 1 |
| TraesCS6A01G382400 | 1 | 1 | 1 | 1 | 1 |
| TraesCS6A01G382500 | 1 | 1 | 1 | 1 | 1 |
| TraesCS6B01G054100 | | | | 1 | |
| TraesCS6D01G012100 | 1 | 1 | 1 | 1 | 1 |
| TraesCS7A01G162200 | 1 | 1 | 1 | 1 | 1 |

| | | | | | |
|---|---|---|---|---|---|
| TraesCS7B01G160000 | 1 | 1 | 1 | 1 | |
| TraesCS7B01G271200 | 1 | 1 | | 1 | |
| TraesCS7B01G439900 | | | | | |
| TraesCS7D01G061300 | 1 | 1 | 1 | 1 | 1 |
| TraesCS7D01G309400 | 1 | 1 | 1 | 1 | 1 |
| TraesCS7D01G512100 | 1 | 1 | 1 | 1 | 1 |
| TraesCS7D01G525100 | 1 | 1 | 1 | 1 | 1 |

**Supplementary Table S30.** Transmission of superior haplotypes of HXM523 (Luomai 21) in its varietal pedigree

| Gene | HXM446 (Yumai 21) | HXM487 (Luomai 1) | HXM495 (Zhoumai 13) | HXM574 (Zhou 8425B) |
|---|---|---|---|---|
| TraesCS1A01G033500 | | 1 | | 1 |
| TraesCS1A01G214700 | | 1 | 1 | |
| TraesCS1A01G218800 | 1 | 1 | 1 | 1 |
| TraesCS1A01G219400 | 1 | 1 | 1 | |
| TraesCS1A01G247100 | | 1 | | 1 |
| TraesCS1A01G263600 | 1 | 1 | 1 | 1 |
| TraesCS1A01G329900 | 1 | 1 | 1 | 1 |
| TraesCS1A01G334200 | 1 | | | |
| TraesCS1A01G372200 | 1 | 1 | | |
| TraesCS1B01G005300 | 1 | 1 | 1 | 1 |
| TraesCS1B01G066900 | 1 | 1 | 1 | 1 |
| TraesCS1D01G059200 | 1 | 1 | 1 | 1 |
| TraesCS2A01G010600 | 1 | 1 | 1 | 1 |
| TraesCS2A01G132300 | | | 1 | 1 |
| TraesCS2A01G545400 | 1 | 1 | 1 | 1 |
| TraesCS2B01G202300 | 1 | | 1 | |
| TraesCS2B01G578500 | | | 1 | 1 |
| TraesCS3A01G311000 | 1 | 1 | 1 | 1 |
| TraesCS3A01G311100 | 1 | 1 | 1 | 1 |
| TraesCS3B01G469000 | | | 1 | 1 |
| TraesCS3B01G560400 | | | 1 | 1 |
| TraesCS3B01G561000 | | | 1 | 1 |
| TraesCS4A01G016600 | 1 | 1 | 1 | 1 |
| TraesCS4A01G105300 | 1 | 1 | 1 | 1 |
| TraesCS4B01G047400 | 1 | 1 | 1 | 1 |
| TraesCS4B01G317400 | | | 1 | 1 |
| TraesCS4B01G390500 | 1 | 1 | 1 | |
| TraesCS5B01G000100 | 1 | 1 | | |
| TraesCS5B01G232100 | | 1 | | 1 |
| TraesCS5B01G236500 | | 1 | | 1 |
| TraesCS5B01G292100 | 1 | 1 | 1 | 1 |
| TraesCS5B01G377500 | | 1 | 1 | 1 |
| TraesCS5B01G400000 | 1 | | 1 | 1 |
| TraesCS5D01G341000 | 1 | 1 | 1 | 1 |
| TraesCS6A01G382400 | 1 | 1 | 1 | 1 |
| TraesCS6A01G382500 | 1 | 1 | 1 | 1 |
| TraesCS6D01G012100 | 1 | 1 | 1 | 1 |
| TraesCS6D01G403800 | | | | 1 |

| | | | | |
|---|---|---|---|---|
| TraesCS7A01G162200 | 1 | 1 | 1 | 1 |
| TraesCS7A01G362300 | 1 | 1 | 1 | 1 |
| TraesCS7B01G160000 | 1 | 1 | 1 | |
| TraesCS7D01G061300 | 1 | 1 | 1 | 1 |
| TraesCS7D01G107500 | 1 | 1 | 1 | 1 |
| TraesCS7D01G159800 | 1 | | | |
| TraesCS7D01G309400 | 1 | 1 | 1 | 1 |
| TraesCS7D01G512100 | 1 | 1 | 1 | 1 |

**Supplementary Table S31.** Transmission of superior haplotypes of HXM590 (Xinmai 30) in its varietal pedigree

| Gene | HXM446 (Yumai 21) | HXM496 (Zhoumai 16) | HXM509 (Xinmai 11) | HXM574 (Zhou 8425B) |
|---|---|---|---|---|
| TraesCS1A01G214700 | | | 1 | |
| TraesCS1A01G218800 | 1 | 1 | 1 | 1 |
| TraesCS1A01G219400 | 1 | 1 | | |
| TraesCS1A01G247100 | | 1 | | 1 |
| TraesCS1A01G263600 | 1 | 1 | 1 | 1 |
| TraesCS1A01G329900 | 1 | 1 | 1 | 1 |
| TraesCS1A01G334200 | | | | |
| TraesCS1A01G372200 | 1 | 1 | 1 | |
| TraesCS1B01G005300 | | | | |
| TraesCS1D01G059200 | 1 | 1 | 1 | 1 |
| TraesCS2A01G010600 | 1 | 1 | 1 | 1 |
| TraesCS2A01G132300 | | | 1 | 1 |
| TraesCS2A01G293400 | 1 | 1 | 1 | |
| TraesCS2A01G545400 | 1 | 1 | 1 | 1 |
| TraesCS2A01G547800 | 1 | 1 | 1 | |
| TraesCS2B01G012500 | 1 | 1 | 1 | |
| TraesCS2B01G202300 | | | | 1 |
| TraesCS2B01G578500 | 1 | 1 | 1 | |
| TraesCS3A01G311000 | 1 | 1 | 1 | 1 |
| TraesCS3A01G311100 | 1 | 1 | 1 | 1 |
| TraesCS3B01G252000 | 1 | 1 | 1 | 1 |
| TraesCS4A01G016600 | 1 | 1 | 1 | 1 |
| TraesCS4A01G105300 | 1 | 1 | 1 | 1 |
| TraesCS4B01G047400 | 1 | 1 | 1 | 1 |
| TraesCS4B01G390500 | 1 | 1 | 1 | |
| TraesCS5A01G558600 | 1 | 1 | 1 | 1 |
| TraesCS5B01G000100 | 1 | | | |
| TraesCS5B01G232100 | | | 1 | |
| TraesCS5B01G236500 | | 1 | 1 | 1 |
| TraesCS5B01G292100 | 1 | 1 | 1 | 1 |
| TraesCS5B01G400000 | 1 | 1 | 1 | 1 |
| TraesCS5D01G341000 | 1 | 1 | 1 | 1 |
| TraesCS5D01G435800 | 1 | 1 | 1 | 1 |
| TraesCS6A01G382400 | 1 | 1 | 1 | 1 |
| TraesCS6A01G382500 | 1 | 1 | 1 | 1 |
| TraesCS6B01G054100 | | | 1 | |
| TraesCS6D01G012100 | 1 | 1 | 1 | 1 |
| TraesCS7A01G162200 | 1 | 1 | 1 | 1 |
| TraesCS7A01G362300 | 1 | 1 | 1 | 1 |
| TraesCS7B01G160000 | 1 | 1 | 1 | |
| TraesCS7B01G271200 | 1 | 1 | 1 | |
| TraesCS7B01G439900 | | | 1 | |
| TraesCS7D01G061300 | 1 | 1 | 1 | 1 |
| TraesCS7D01G107500 | 1 | 1 | 1 | 1 |
| TraesCS7D01G159800 | 1 | | 1 | |
| TraesCS7D01G309400 | 1 | 1 | 1 | 1 |
| TraesCS7D01G512100 | | | 1 | |
| TraesCS7D01G525100 | 1 | 1 | 1 | 1 |

**Supplementary Table S32.** The superior haplotype frequencies for 13 wheat traits between landraces and cultivars with different periods

| Trait | Gene | LR | 50s | 60s-70s | 80s-90s | 00s- |
|---|---|---|---|---|---|---|
| PH | TraesCS1A01G218800 | 0.73 | 0.68 | 0.75 | 0.89 | 0.91 |
| PH | TraesCS1A01G263600 | 0.10 | 0.54 | 0.82 | 0.95 | 0.98 |
| PH | TraesCS1B01G462300 | 0.01 | 0.08 | 0.00 | 0.14 | 0.26 |
| PH | TraesCS2A01G453300 | 0.02 | 0.11 | 0.11 | 0.06 | 0.10 |
| PH | TraesCS2A01G545400 | 0.91 | 0.89 | 0.96 | 1.00 | 0.99 |
| PH | TraesCS2A01G547800 | 0.37 | 0.62 | 0.86 | 0.86 | 0.81 |
| PH | TraesCS2A01G552400 | 0.01 | 0.08 | 0.14 | 0.11 | 0.07 |
| PH | TraesCS2B01G575900 | 0.01 | 0.00 | 0.04 | 0.09 | 0.07 |
| PH | TraesCS2B01G578500 | 0.04 | 0.11 | 0.07 | 0.14 | 0.14 |
| PH | TraesCS3B01G039100 | 0.12 | 0.49 | 0.54 | 0.46 | 0.63 |
| PH | TraesCS3D01G540600 | 0.00 | 0.00 | 0.00 | 0.00 | 0.00 |
| PH | TraesCS4A01G005600 | 0.00 | 0.00 | 0.04 | 0.09 | 0.07 |
| PH | TraesCS4A01G016600 | 0.21 | 0.57 | 0.86 | 0.95 | 0.94 |
| PH | TraesCS4A01G455200 | 0.01 | 0.14 | 0.11 | 0.22 | 0.12 |
| PH | TraesCS4A01G460100 | 0.01 | 0.27 | 0.39 | 0.57 | 0.31 |
| PH | TraesCS4D01G301100 | 0.83 | 0.81 | 0.68 | 0.88 | 0.90 |
| PH | TraesCS5A01G330500 | 0.00 | 0.00 | 0.00 | 0.00 | 0.03 |
| PH | TraesCS5A01G543100 | 0.40 | 0.49 | 0.39 | 0.52 | 0.86 |
| PH | TraesCS5A01G558600 | 0.01 | 0.19 | 0.14 | 0.62 | 0.62 |
| PH | TraesCS5B01G232100 | 0.01 | 0.41 | 0.64 | 0.62 | 0.76 |
| PH | TraesCS5B01G236500 | 0.03 | 0.43 | 0.75 | 0.75 | 0.84 |
| PH | TraesCS5B01G523100 | 0.00 | 0.16 | 0.11 | 0.26 | 0.45 |
| PH | TraesCS5D01G097400 | 0.99 | 1.00 | 1.00 | 1.00 | 1.00 |
| PH | TraesCS6B01G178300 | 0.16 | 0.27 | 0.50 | 0.17 | 0.53 |
| PH | TraesCS6B01G293600 | 0.08 | 0.57 | 0.89 | 0.88 | 0.79 |
| PH | TraesCS6D01G173000 | 0.03 | 0.54 | 0.93 | 0.83 | 0.86 |
| PH | TraesCS7A01G362300 | 0.06 | 0.51 | 0.68 | 0.83 | 0.94 |
| PH | TraesCS7A01G424700 | 0.01 | 0.00 | 0.00 | 0.00 | 0.00 |
| SL | TraesCS1A01G320200 | 0.01 | 0.00 | 0.00 | 0.00 | 0.00 |
| SL | TraesCS1A01G334200 | 0.19 | 0.38 | 0.57 | 0.74 | 0.84 |
| SL | TraesCS1A01G334900 | 0.03 | 0.05 | 0.21 | 0.12 | 0.02 |
| SL | TraesCS1B01G005300 | 0.90 | 0.89 | 0.96 | 0.97 | 0.95 |
| SL | TraesCS1B01G133500 | 0.01 | 0.00 | 0.00 | 0.00 | 0.00 |

| | | | | | | |
|---|---|---|---|---|---|---|
| SL | TraesCS1B01G318800 | 0.98 | 0.95 | 1.00 | 1.00 | 0.98 |
| SL | TraesCS2A01G009800 | 0.26 | 0.11 | 0.00 | 0.02 | 0.02 |
| SL | TraesCS2A01G013000 | 0.98 | 1.00 | 1.00 | 1.00 | 1.00 |
| SL | TraesCS2A01G445500 | 0.00 | 0.00 | 0.00 | 0.00 | 0.00 |
| SL | TraesCS2A01G590600 | 0.01 | 0.00 | 0.00 | 0.00 | 0.00 |
| SL | TraesCS2B01G066100 | 0.88 | 0.73 | 0.75 | 0.89 | 0.96 |
| SL | TraesCS2B01G578500 | 0.37 | 0.62 | 0.86 | 0.85 | 0.82 |
| SL | TraesCS3A01G019600 | 0.03 | 0.00 | 0.00 | 0.02 | 0.01 |
| SL | TraesCS3A01G103800 | 0.03 | 0.51 | 0.79 | 0.92 | 0.95 |
| SL | TraesCS3A01G280200 | 0.69 | 0.92 | 1.00 | 0.98 | 0.99 |
| SL | TraesCS3A01G481400 | 0.00 | 0.00 | 0.00 | 0.00 | 0.01 |
| SL | TraesCS4A01G016600 | 0.21 | 0.57 | 0.86 | 0.95 | 0.94 |
| SL | TraesCS5A01G295800 | 0.00 | 0.00 | 0.00 | 0.00 | 0.00 |
| SL | TraesCS5B01G014100 | 0.27 | 0.59 | 0.54 | 0.58 | 0.72 |
| SL | TraesCS5B01G236500 | 0.00 | 0.00 | 0.00 | 0.00 | 0.00 |
| SL | TraesCS5B01G277500 | 0.00 | 0.00 | 0.04 | 0.00 | 0.00 |
| SL | TraesCS5D01G435800 | 0.33 | 0.46 | 0.82 | 0.68 | 0.75 |
| SL | TraesCS6A01G313800 | 0.73 | 0.68 | 0.71 | 0.74 | 0.70 |
| SL | TraesCS7B01G478200 | 0.00 | 0.00 | 0.00 | 0.00 | 0.00 |
| SL | TraesCS7B01G484200 | 0.00 | 0.00 | 0.00 | 0.00 | 0.00 |
| PPR | TraesCS2A01G521500 | 0.00 | 0.00 | 0.04 | 0.00 | 0.00 |
| PPR | TraesCS2A01G547800 | 0.37 | 0.62 | 0.86 | 0.86 | 0.81 |
| PPR | TraesCS2A01G552400 | 0.01 | 0.08 | 0.14 | 0.11 | 0.07 |
| PPR | TraesCS2B01G010100 | 0.04 | 0.38 | 0.50 | 0.60 | 0.35 |
| PPR | TraesCS3A01G535100 | 0.00 | 0.00 | 0.00 | 0.02 | 0.01 |
| PPR | TraesCS4A01G016600 | 0.21 | 0.57 | 0.86 | 0.95 | 0.94 |
| PPR | TraesCS4A01G079300 | 0.76 | 0.89 | 0.93 | 0.98 | 1.00 |
| PPR | TraesCS4B01G263400 | 0.01 | 0.35 | 0.64 | 0.48 | 0.30 |
| PPR | TraesCS5B01G377500 | 0.03 | 0.38 | 0.57 | 0.60 | 0.30 |
| PPR | TraesCS5B01G523100 | 0.07 | 0.57 | 0.68 | 0.91 | 0.91 |
| PPR | TraesCS5D01G097400 | 0.01 | 0.00 | 0.00 | 0.00 | 0.00 |
| PPR | TraesCS6B01G293600 | 0.00 | 0.00 | 0.00 | 0.03 | 0.00 |
| PPR | TraesCS7A01G059000 | 0.07 | 0.16 | 0.29 | 0.23 | 0.12 |
| GL | TraesCS1A01G273900 | 0.01 | 0.00 | 0.00 | 0.00 | 0.00 |
| GL | TraesCS2A01G010600 | 0.60 | 0.76 | 0.96 | 0.97 | 0.99 |
| GL | TraesCS2A01G482400 | 0.00 | 0.00 | 0.00 | 0.00 | 0.01 |
| GL | TraesCS2D01G011200 | 0.01 | 0.00 | 0.00 | 0.00 | 0.00 |

| | | | | | | |
|---|---|---|---|---|---|---|
| GL | TraesCS2D01G016900 | 0.01 | 0.00 | 0.00 | 0.00 | 0.00 |
| GL | TraesCS2D01G082000 | 0.02 | 0.00 | 0.04 | 0.00 | 0.00 |
| GL | TraesCS2D01G482500 | 0.00 | 0.00 | 0.00 | 0.00 | 0.00 |
| GL | TraesCS3B01G560400 | 0.06 | 0.05 | 0.04 | 0.25 | 0.40 |
| GL | TraesCS3B01G561000 | 0.06 | 0.05 | 0.00 | 0.26 | 0.40 |
| GL | TraesCS4A01G266900 | 0.01 | 0.00 | 0.00 | 0.00 | 0.00 |
| GL | TraesCS4A01G444300 | 0.01 | 0.00 | 0.00 | 0.00 | 0.00 |
| GL | TraesCS4B01G047400 | 0.68 | 0.86 | 1.00 | 0.97 | 0.99 |
| GL | TraesCS4B01G321800 | 0.01 | 0.00 | 0.00 | 0.00 | 0.00 |
| GL | TraesCS4B01G360800 | 0.00 | 0.00 | 0.00 | 0.00 | 0.00 |
| GL | TraesCS4D01G318400 | 0.01 | 0.00 | 0.00 | 0.00 | 0.00 |
| GL | TraesCS4D01G325200 | 0.01 | 0.00 | 0.00 | 0.00 | 0.00 |
| GL | TraesCS5A01G377700 | 0.02 | 0.00 | 0.00 | 0.00 | 0.01 |
| GL | TraesCS5B01G307600 | 0.11 | 0.38 | 0.43 | 0.28 | 0.10 |
| GL | TraesCS5B01G375800 | 0.01 | 0.00 | 0.00 | 0.00 | 0.00 |
| GL | TraesCS6A01G009100 | 0.01 | 0.00 | 0.00 | 0.00 | 0.00 |
| GL | TraesCS6A01G382400 | 0.31 | 0.65 | 0.96 | 0.94 | 0.96 |
| GL | TraesCS6A01G382500 | 0.31 | 0.65 | 0.96 | 0.94 | 0.96 |
| GL | TraesCS6A01G382600 | 0.09 | 0.35 | 0.50 | 0.31 | 0.34 |
| GL | TraesCS6D01G378300 | 0.01 | 0.00 | 0.00 | 0.00 | 0.00 |
| GL | TraesCS7B01G018600 | 0.01 | 0.00 | 0.00 | 0.00 | 0.00 |
| GL | TraesCS7D01G039500 | 0.01 | 0.00 | 0.00 | 0.00 | 0.00 |
| GW | TraesCS1A01G247100 | 0.16 | 0.43 | 0.82 | 0.82 | 0.84 |
| GW | TraesCS1A01G329900 | 0.07 | 0.57 | 0.89 | 0.98 | 0.99 |
| GW | TraesCS1B01G155700 | 0.00 | 0.11 | 0.14 | 0.09 | 0.05 |
| GW | TraesCS1B01G462000 | 0.01 | 0.14 | 0.11 | 0.12 | 0.30 |
| GW | TraesCS2A01G464000 | 0.00 | 0.00 | 0.00 | 0.00 | 0.01 |
| GW | TraesCS2B01G485700 | 0.00 | 0.16 | 0.29 | 0.34 | 0.25 |
| GW | TraesCS2D01G505700 | 0.00 | 0.05 | 0.04 | 0.05 | 0.04 |
| GW | TraesCS3A01G311100 | 0.02 | 0.51 | 0.79 | 0.92 | 0.96 |
| GW | TraesCS6A01G382400 | 0.31 | 0.65 | 0.96 | 0.94 | 0.96 |
| GW | TraesCS6B01G015100 | 0.00 | 0.03 | 0.00 | 0.00 | 0.01 |
| GW | TraesCS6B01G130700 | 0.00 | 0.00 | 0.00 | 0.00 | 0.01 |
| GW | TraesCS6B01G186700 | 0.00 | 0.00 | 0.00 | 0.02 | 0.00 |
| GW | TraesCS6D01G009700 | 0.75 | 0.65 | 0.68 | 0.85 | 0.89 |
| GW | TraesCS6D01G012100 | 0.55 | 0.65 | 0.75 | 0.80 | 0.86 |
| GW | TraesCS7A01G043700 | 0.00 | 0.03 | 0.04 | 0.09 | 0.04 |

| | | | | | | |
|---|---|---|---|---|---|---|
| GW | TraesCS7A01G120300 | 0.00 | 0.00 | 0.00 | 0.00 | 0.01 |
| GW | TraesCS7B01G115300 | 0.29 | 0.76 | 0.93 | 0.98 | 0.98 |
| GW | TraesCS7B01G160000 | 0.31 | 0.68 | 0.82 | 0.88 | 0.87 |
| GWLR | TraesCS1A01G353300 | 0.97 | 0.97 | 1.00 | 0.98 | 1.00 |
| GWLR | TraesCS2A01G548100 | 0.00 | 0.00 | 0.00 | 0.00 | 0.00 |
| GWLR | TraesCS2B01G485700 | 0.85 | 0.73 | 0.71 | 0.60 | 0.73 |
| GWLR | TraesCS2D01G011200 | 0.13 | 0.11 | 0.00 | 0.00 | 0.01 |
| GWLR | TraesCS2D01G016900 | 0.99 | 1.00 | 1.00 | 1.00 | 1.00 |
| GWLR | TraesCS3B01G128700 | 0.10 | 0.11 | 0.07 | 0.02 | 0.04 |
| GWLR | TraesCS4A01G074300 | 0.26 | 0.14 | 0.07 | 0.02 | 0.01 |
| GWLR | TraesCS4A01G227900 | 0.96 | 1.00 | 1.00 | 1.00 | 1.00 |
| GWLR | TraesCS4A01G418200 | 0.99 | 0.95 | 1.00 | 0.97 | 0.95 |
| GWLR | TraesCS4D01G350500 | 0.99 | 1.00 | 1.00 | 1.00 | 1.00 |
| GWLR | TraesCS4D01G353800 | 0.99 | 1.00 | 1.00 | 1.00 | 0.99 |
| GWLR | TraesCS5B01G375800 | 0.01 | 0.00 | 0.00 | 0.05 | 0.01 |
| GWLR | TraesCS5D01G387700 | 0.99 | 1.00 | 1.00 | 1.00 | 1.00 |
| GWLR | TraesCS6A01G009100 | 0.00 | 0.00 | 0.00 | 0.03 | 0.00 |
| GWLR | TraesCS6B01G015100 | 0.02 | 0.00 | 0.00 | 0.00 | 0.01 |
| GWLR | TraesCS6B01G327500 | 0.95 | 0.76 | 0.50 | 0.69 | 0.85 |
| GWLR | TraesCS7A01G549100 | 0.00 | 0.00 | 0.00 | 0.00 | 0.01 |
| GWLR | TraesCS7A01G549200 | 0.01 | 0.00 | 0.00 | 0.00 | 0.01 |
| GWLR | TraesCS7A01G549300 | 0.01 | 0.00 | 0.00 | 0.00 | 0.00 |
| GWLR | TraesCS7D01G039500 | 0.99 | 1.00 | 0.96 | 1.00 | 0.99 |
| TGW | TraesCS1A01G050400 | 0.00 | 0.00 | 0.00 | 0.00 | 0.01 |
| TGW | TraesCS1A01G058400 | 0.00 | 0.00 | 0.00 | 0.00 | 0.01 |
| TGW | TraesCS1A01G063200 | 0.00 | 0.00 | 0.00 | 0.03 | 0.15 |
| TGW | TraesCS1A01G329900 | 0.00 | 0.03 | 0.00 | 0.09 | 0.13 |
| TGW | TraesCS1A01G435000 | 0.00 | 0.00 | 0.00 | 0.00 | 0.01 |
| TGW | TraesCS1B01G368500 | 0.00 | 0.00 | 0.21 | 0.09 | 0.05 |
| TGW | TraesCS1B01G462000 | 0.00 | 0.08 | 0.04 | 0.12 | 0.27 |
| TGW | TraesCS1D01G059200 | 0.58 | 0.73 | 0.79 | 0.98 | 0.99 |
| TGW | TraesCS1D01G063500 | 0.01 | 0.00 | 0.00 | 0.00 | 0.00 |
| TGW | TraesCS1D01G063900 | 0.01 | 0.00 | 0.00 | 0.02 | 0.01 |
| TGW | TraesCS2A01G310300 | 0.00 | 0.00 | 0.00 | 0.00 | 0.01 |
| TGW | TraesCS2A01G547800 | 0.37 | 0.62 | 0.86 | 0.86 | 0.81 |
| TGW | TraesCS2A01G548100 | 0.00 | 0.00 | 0.00 | 0.03 | 0.03 |
| TGW | TraesCS2A01G555600 | 0.00 | 0.00 | 0.00 | 0.02 | 0.01 |

| | | | | | | |
|---|---|---|---|---|---|---|
| TGW | TraesCS2B01G010800 | 0.00 | 0.00 | 0.00 | 0.00 | 0.01 |
| TGW | TraesCS2B01G012500 | 0.03 | 0.35 | 0.71 | 0.66 | 0.67 |
| TGW | TraesCS2B01G201900 | 0.98 | 1.00 | 0.93 | 1.00 | 1.00 |
| TGW | TraesCS2B01G202300 | 0.01 | 0.22 | 0.11 | 0.14 | 0.33 |
| TGW | TraesCS2B01G533300 | 0.00 | 0.00 | 0.00 | 0.00 | 0.01 |
| TGW | TraesCS3A01G311000 | 0.02 | 0.49 | 0.79 | 0.92 | 0.95 |
| TGW | TraesCS3A01G311100 | 0.02 | 0.51 | 0.79 | 0.92 | 0.96 |
| TGW | TraesCS3B01G252000 | 0.07 | 0.51 | 0.68 | 0.78 | 0.83 |
| TGW | TraesCS4A01G289300 | 0.92 | 1.00 | 1.00 | 1.00 | 1.00 |
| TGW | TraesCS4A01G444300 | 0.00 | 0.00 | 0.00 | 0.02 | 0.04 |
| TGW | TraesCS4B01G029700 | 0.04 | 0.41 | 0.46 | 0.71 | 0.45 |
| TGW | TraesCS4D01G274200 | 0.01 | 0.00 | 0.00 | 0.00 | 0.00 |
| TGW | TraesCS4D01G350500 | 0.01 | 0.00 | 0.00 | 0.00 | 0.00 |
| TGW | TraesCS4D01G353800 | 0.01 | 0.00 | 0.00 | 0.00 | 0.01 |
| TGW | TraesCS5B01G292100 | 0.15 | 0.38 | 0.54 | 0.80 | 0.99 |
| TGW | TraesCS6A01G194600 | 0.91 | 0.89 | 0.89 | 0.94 | 0.94 |
| TGW | TraesCS6A01G382500 | 0.31 | 0.65 | 0.96 | 0.94 | 0.96 |
| TGW | TraesCS6A01G382600 | 0.09 | 0.35 | 0.50 | 0.31 | 0.34 |
| TGW | TraesCS6B01G015100 | 0.00 | 0.03 | 0.00 | 0.00 | 0.00 |
| TGW | TraesCS6B01G253900 | 0.01 | 0.24 | 0.46 | 0.31 | 0.19 |
| TGW | TraesCS6D01G009700 | 0.01 | 0.00 | 0.00 | 0.00 | 0.00 |
| TGW | TraesCS6D01G012100 | 0.57 | 0.65 | 0.79 | 0.83 | 0.87 |
| TGW | TraesCS7A01G043700 | 0.00 | 0.00 | 0.00 | 0.00 | 0.01 |
| TGW | TraesCS7B01G455000 | 0.02 | 0.32 | 0.25 | 0.28 | 0.21 |
| TGW | TraesCS7D01G309400 | 0.76 | 0.84 | 1.00 | 1.00 | 0.99 |
| TGW | TraesCS7D01G525100 | 0.83 | 0.89 | 1.00 | 0.98 | 0.97 |
| SNS | TraesCS2B01G464900 | 0.07 | 0.00 | 0.00 | 0.00 | 0.01 |
| SNS | TraesCS3A01G022700 | 0.01 | 0.00 | 0.00 | 0.00 | 0.00 |
| SNS | TraesCS3A01G391600 | 0.01 | 0.00 | 0.00 | 0.00 | 0.00 |
| SNS | TraesCS3A01G502900 | 0.02 | 0.00 | 0.00 | 0.00 | 0.00 |
| SNS | TraesCS3B01G428100 | 0.01 | 0.00 | 0.00 | 0.00 | 0.00 |
| SNS | TraesCS4A01G079300 | 0.25 | 0.11 | 0.07 | 0.02 | 0.00 |
| SNS | TraesCS4A01G266900 | 1.00 | 1.00 | 1.00 | 1.00 | 1.00 |
| SNS | TraesCS5B01G469600 | 0.22 | 0.11 | 0.14 | 0.11 | 0.12 |
| SNS | TraesCS7A01G433000 | 0.00 | 0.00 | 0.00 | 0.00 | 0.00 |
| SNS | TraesCS7B01G271200 | 0.97 | 0.78 | 0.36 | 0.72 | 0.76 |
| T | TraesCS1A01G033500 | 0.02 | 0.22 | 0.14 | 0.57 | 0.67 |

| | | | | | | |
|---|---|---|---|---|---|---|
| T | TraesCS2A01G176000 | 0.01 | 0.03 | 0.00 | 0.18 | 0.00 |
| T | TraesCS2B01G578500 | 0.41 | 0.73 | 0.93 | 0.98 | 0.96 |
| T | TraesCS4A01G302000 | 0.00 | 0.00 | 0.00 | 0.00 | 0.00 |
| T | TraesCS7D01G159800 | 0.03 | 0.41 | 0.64 | 0.63 | 0.47 |
| T | TraesCS7D01G512100 | 0.30 | 0.59 | 0.89 | 0.95 | 0.93 |
| V | TraesCS1A01G214700 | 0.05 | 0.30 | 0.43 | 0.62 | 0.70 |
| V | TraesCS1B01G066900 | 0.02 | 0.08 | 0.07 | 0.40 | 0.52 |
| V | TraesCS2A01G293400 | 0.04 | 0.35 | 0.61 | 0.62 | 0.84 |
| V | TraesCS2B01G202300 | 0.03 | 0.30 | 0.21 | 0.51 | 0.50 |
| V | TraesCS3A01G015500 | 0.00 | 0.00 | 0.00 | 0.00 | 0.01 |
| V | TraesCS3B01G469000 | 0.01 | 0.14 | 0.14 | 0.26 | 0.25 |
| V | TraesCS3B01G560400 | 0.05 | 0.05 | 0.00 | 0.25 | 0.39 |
| V | TraesCS3B01G561000 | 0.06 | 0.05 | 0.00 | 0.26 | 0.40 |
| V | TraesCS4A01G016600 | 0.21 | 0.57 | 0.86 | 0.95 | 0.94 |
| V | TraesCS5B01G000100 | 0.03 | 0.14 | 0.36 | 0.48 | 0.49 |
| V | TraesCS6B01G009600 | 0.21 | 0.49 | 0.82 | 0.68 | 0.79 |
| V | TraesCS6D01G403800 | 0.02 | 0.22 | 0.43 | 0.52 | 0.47 |
| V | TraesCS7A01G040900 | 0.00 | 0.00 | 0.00 | 0.00 | 0.01 |
| V | TraesCS7A01G518200 | 0.00 | 0.00 | 0.00 | 0.02 | 0.00 |
| V | TraesCS7A01G518300 | 0.00 | 0.00 | 0.04 | 0.02 | 0.01 |
| V | TraesCS7A01G549100 | 0.04 | 0.41 | 0.64 | 0.49 | 0.58 |
| V | TraesCS7A01G549200 | 0.00 | 0.00 | 0.00 | 0.02 | 0.00 |
| V | TraesCS7A01G549300 | 0.00 | 0.00 | 0.04 | 0.09 | 0.10 |
| V | TraesCS7B01G034600 | 0.00 | 0.00 | 0.00 | 0.03 | 0.19 |
| V | TraesCS7B01G115300 | 0.29 | 0.76 | 0.93 | 0.98 | 0.99 |
| V | TraesCS7B01G439900 | 0.00 | 0.00 | 0.04 | 0.00 | 0.01 |
| V | TraesCS7B01G472500 | 0.00 | 0.00 | 0.00 | 0.02 | 0.00 |
| V | TraesCS7D01G215600 | 0.47 | 0.76 | 1.00 | 1.00 | 0.99 |
| Vmax | TraesCS1A01G033500 | 0.00 | 0.11 | 0.07 | 0.48 | 0.54 |
| Vmax | TraesCS1A01G033600 | 0.00 | 0.00 | 0.04 | 0.14 | 0.20 |
| Vmax | TraesCS1A01G219400 | 0.01 | 0.14 | 0.32 | 0.57 | 0.59 |
| Vmax | TraesCS1A01G247100 | 0.16 | 0.43 | 0.82 | 0.82 | 0.84 |
| Vmax | TraesCS1B01G066900 | 0.02 | 0.08 | 0.07 | 0.40 | 0.52 |
| Vmax | TraesCS1B01G422700 | 0.00 | 0.00 | 0.00 | 0.02 | 0.01 |
| Vmax | TraesCS2A01G293400 | 0.04 | 0.35 | 0.61 | 0.62 | 0.84 |
| Vmax | TraesCS2B01G202300 | 0.03 | 0.30 | 0.21 | 0.51 | 0.50 |
| Vmax | TraesCS3B01G469000 | 0.01 | 0.14 | 0.14 | 0.26 | 0.25 |

| | | | | | | |
|---|---|---|---|---|---|---|
| Vmax | TraesCS4A01G016600 | 0.21 | 0.57 | 0.86 | 0.95 | 0.94 |
| Vmax | TraesCS4A01G093100 | 0.83 | 0.89 | 0.82 | 0.94 | 0.91 |
| Vmax | TraesCS4B01G390500 | 0.10 | 0.41 | 0.39 | 0.82 | 0.77 |
| Vmax | TraesCS5B01G039800 | 0.00 | 0.00 | 0.00 | 0.02 | 0.00 |
| Vmax | TraesCS5B01G400000 | 0.10 | 0.54 | 0.86 | 0.97 | 0.98 |
| Vmax | TraesCS6B01G009600 | 0.21 | 0.49 | 0.82 | 0.68 | 0.79 |
| Vmax | TraesCS6B01G369700 | 0.00 | 0.08 | 0.18 | 0.23 | 0.07 |
| Vmax | TraesCS6D01G403800 | 0.02 | 0.14 | 0.32 | 0.48 | 0.43 |
| Vmax | TraesCS7A01G549100 | 0.00 | 0.00 | 0.00 | 0.00 | 0.01 |
| Vmax | TraesCS7A01G549200 | 0.00 | 0.00 | 0.00 | 0.02 | 0.00 |
| Vmax | TraesCS7A01G549300 | 0.03 | 0.41 | 0.64 | 0.49 | 0.56 |
| Vmax | TraesCS7B01G034600 | 0.00 | 0.00 | 0.00 | 0.03 | 0.19 |
| Vmax | TraesCS7B01G115300 | 0.29 | 0.76 | 0.93 | 0.98 | 0.99 |
| Vmax | TraesCS7B01G439900 | 0.06 | 0.22 | 0.32 | 0.51 | 0.55 |
| Vmax | TraesCS7B01G472500 | 0.00 | 0.00 | 0.00 | 0.02 | 0.00 |
| Vmax | TraesCS7D01G215600 | 0.47 | 0.76 | 1.00 | 1.00 | 0.99 |
| FD | TraesCS1A01G372200 | 0.19 | 0.49 | 0.82 | 0.88 | 0.88 |
| FD | TraesCS1A01G372300 | 0.01 | 0.00 | 0.00 | 0.00 | 0.01 |
| FD | TraesCS1D01G059200 | 0.58 | 0.73 | 0.79 | 0.98 | 0.99 |
| FD | TraesCS1D01G378800 | 0.00 | 0.05 | 0.18 | 0.12 | 0.22 |
| FD | TraesCS1D01G378900 | 0.38 | 0.49 | 0.46 | 0.55 | 0.68 |
| FD | TraesCS2A01G119600 | 0.00 | 0.00 | 0.00 | 0.00 | 0.00 |
| FD | TraesCS2A01G132300 | 0.27 | 0.49 | 0.71 | 0.74 | 0.73 |
| FD | TraesCS2A01G179000 | 0.00 | 0.00 | 0.00 | 0.02 | 0.00 |
| FD | TraesCS2A01G179200 | 0.00 | 0.00 | 0.04 | 0.00 | 0.01 |
| FD | TraesCS2A01G194500 | 0.73 | 0.97 | 1.00 | 1.00 | 1.00 |
| FD | TraesCS2A01G534200 | 0.27 | 0.46 | 0.43 | 0.66 | 0.80 |
| FD | TraesCS2A01G534300 | 0.27 | 0.46 | 0.43 | 0.66 | 0.80 |
| FD | TraesCS2B01G140300 | 0.71 | 0.95 | 0.96 | 1.00 | 1.00 |
| FD | TraesCS2D01G082000 | 0.98 | 1.00 | 0.96 | 1.00 | 1.00 |
| FD | TraesCS3B01G017400 | 0.28 | 0.16 | 0.00 | 0.00 | 0.00 |
| FD | TraesCS3B01G428100 | 0.00 | 0.03 | 0.04 | 0.09 | 0.01 |
| FD | TraesCS3B01G517100 | 0.01 | 0.00 | 0.00 | 0.00 | 0.00 |
| FD | TraesCS4A01G044800 | 0.00 | 0.00 | 0.00 | 0.00 | 0.00 |
| FD | TraesCS4A01G105300 | 0.03 | 0.35 | 0.46 | 0.58 | 0.75 |
| FD | TraesCS4A01G324900 | 0.00 | 0.00 | 0.00 | 0.02 | 0.01 |
| FD | TraesCS4A01G375100 | 0.18 | 0.19 | 0.29 | 0.35 | 0.51 |

| | | | | | | |
|---|---|---|---|---|---|---|
| FD | TraesCS4B01G317400 | 0.01 | 0.08 | 0.21 | 0.26 | 0.32 |
| FD | TraesCS5A01G393400 | 0.00 | 0.00 | 0.04 | 0.02 | 0.00 |
| FD | TraesCS5B01G358500 | 0.20 | 0.30 | 0.36 | 0.60 | 0.83 |
| FD | TraesCS5D01G170100 | 0.10 | 0.68 | 0.86 | 0.89 | 0.92 |
| FD | TraesCS5D01G341000 | 0.53 | 0.78 | 1.00 | 0.97 | 1.00 |
| FD | TraesCS5D01G403200 | 0.99 | 1.00 | 1.00 | 1.00 | 1.00 |
| FD | TraesCS7A01G162200 | 0.89 | 0.92 | 0.96 | 1.00 | 1.00 |
| FD | TraesCS7B01G302900 | 0.04 | 0.32 | 0.61 | 0.54 | 0.73 |
| FD | TraesCS7D01G003500 | 0.94 | 1.00 | 1.00 | 0.98 | 0.99 |
| FD | TraesCS7D01G061300 | 0.44 | 0.70 | 0.96 | 0.97 | 0.95 |
| FD | TraesCS7D01G183600 | 0.01 | 0.24 | 0.36 | 0.55 | 0.44 |
| HD | TraesCS1A01G301400 | 0.00 | 0.08 | 0.11 | 0.06 | 0.07 |
| HD | TraesCS1A01G435000 | 0.01 | 0.00 | 0.00 | 0.00 | 0.00 |
| HD | TraesCS1D01G444100 | 0.01 | 0.00 | 0.00 | 0.02 | 0.00 |
| HD | TraesCS3B01G017400 | 0.28 | 0.16 | 0.00 | 0.00 | 0.00 |
| HD | TraesCS3B01G517100 | 0.01 | 0.00 | 0.00 | 0.00 | 0.00 |
| HD | TraesCS4B01G020300 | 0.05 | 0.35 | 0.39 | 0.31 | 0.79 |
| HD | TraesCS5B01G538100 | 0.00 | 0.00 | 0.00 | 0.00 | 0.00 |
| HD | TraesCS5D01G401500 | 0.99 | 1.00 | 1.00 | 1.00 | 1.00 |
| HD | TraesCS5D01G403200 | 0.99 | 1.00 | 1.00 | 1.00 | 1.00 |
| HD | TraesCS6B01G054100 | 0.18 | 0.24 | 0.61 | 0.68 | 0.52 |
| HD | TraesCS7D01G107500 | 0.85 | 0.86 | 0.93 | 0.98 | 0.92 |

**Supplementary Table S33**. Haplotype frequencies of the genes for important traits in landraces and cultivars with different periods

| Gene | Symbol | LR | 50s | 60s-70s | 80s-90s | 00s- | Trait |
|---|---|---|---|---|---|---|---|
| TraesCS4A01G066200 | *Lr67* | 0.75 | 0.89 | 0.93 | 1.00 | 1.00 | Powdery Mildew |
| TraesCS4D01G243100 | *Lr67* | 0.01 | 0.00 | 0.00 | 0.00 | 0.00 | Powdery Mildew |
| TraesCS4D01G319100 | *TaMlo-D1* | 0.01 | 0.00 | 0.00 | 0.00 | 0.00 | Powdery Mildew |
| TraesCS5A01G494800 | *TaMlo-A1* | 0.42 | 0.41 | 0.25 | 0.38 | 0.53 | Powdery Mildew |
| TraesCS7B01G441700 | *Pm5e* | 0.26 | 0.22 | 0.25 | 0.35 | 0.39 | Powdery Mildew |
| TraesCS7D01G080300 | *Lr34* | 0.07 | 0.41 | 0.57 | 0.66 | 0.59 | Powdery Mildew |
| TraesCS2B01G488000 | *YrSP* | 0.22 | 0.24 | 0.14 | 0.29 | 0.30 | Stripe Rust |
| TraesCS1A01G061500 | *Yr15-1A* | 0.49 | 0.51 | 0.39 | 0.37 | 0.60 | Stripe Rust |
| TraesCS2B01G182800 | *Lr13* | 0.60 | 0.38 | 0.21 | 0.12 | 0.26 | Stripe Rust |
| TraesCS4A01G066200 | *Lr67* | 0.24 | 0.11 | 0.07 | 0.00 | 0.00 | Stripe Rust |
| TraesCS4D01G243100 | *Lr67* | 0.99 | 1.00 | 1.00 | 1.00 | 1.00 | Stripe Rust |
| TraesCS5A01G344100 | *YrU1* | 0.85 | 0.70 | 0.61 | 0.52 | 0.72 | Stripe Rust |
| TraesCS5B01G344200 | *YrU1-5B* | 0.82 | 0.70 | 0.82 | 0.71 | 0.87 | Stripe Rust |
| TraesCS5D01G349800 | *YrU1-5D* | 0.93 | 0.89 | 0.96 | 0.95 | 0.96 | Stripe Rust |
| TraesCS6B01G167900 | *TaPsIPK1-6B* | 0.13 | 0.05 | 0.00 | 0.00 | 0.00 | Stripe Rust |
| TraesCS6B01G185700 | *TaCBF1b* | 0.48 | 0.08 | 0.04 | 0.40 | 0.53 | Stripe Rust |
| TraesCS7D01G080300 | *Lr34* | 0.92 | 0.57 | 0.43 | 0.32 | 0.41 | Stripe Rust |
| TraesCS1A01G007934 | *Glu-A3* | 0.96 | 0.76 | 0.68 | 0.63 | 0.64 | High Protein Content |
| TraesCS1A01G008000 | *Glu-A3* | 0.91 | 0.70 | 0.61 | 0.54 | 0.54 | High Protein Content |
| TraesCS1A01G010905 | *Glu-A3* | 0.95 | 0.78 | 0.50 | 0.26 | 0.32 | High Protein Content |
| TraesCS1A01G317311 | *Glu-1Ax* | 0.97 | 0.62 | 0.25 | 0.31 | 0.51 | High Protein Content |
| TraesCS1B01G329711 | *Glu-1Bx* | 1.00 | 1.00 | 1.00 | 1.00 | 1.00 | High Protein Content |
| TraesCS1B01G329992 | *Glu-1By* | 0.88 | 0.73 | 0.79 | 0.62 | 0.69 | High Protein Content |
| TraesCS1D01G007400 | *Glu-D3* | 0.93 | 0.86 | 0.82 | 0.75 | 0.84 | High Protein Content |
| TraesCS1D01G008600 | *Glu-D3* | 0.13 | 0.11 | 0.11 | 0.02 | 0.01 | High Protein Content |
| TraesCS1D01G009400 | *Glu-D3* | 0.97 | 0.92 | 1.00 | 0.91 | 0.76 | High Protein Content |
| TraesCS1D01G009900 | *Glu-D3* | 0.64 | 0.73 | 0.71 | 0.58 | 0.72 | High Protein Content |
| TraesCS1D01G015100 | *Glu-D3* | 1.00 | 1.00 | 1.00 | 1.00 | 1.00 | High Protein Content |
| TraesCS3A01G077900 | *TaNAC019-3A* | 0.98 | 0.92 | 0.93 | 0.88 | 0.73 | High Protein Content |
| TraesCS1A01G007934 | *Glu-A3* | 0.00 | 0.19 | 0.11 | 0.32 | 0.31 | Low Protein Content |
| TraesCS1A01G008000 | *Glu-A3* | 0.02 | 0.22 | 0.25 | 0.37 | 0.39 | Low Protein Content |
| TraesCS1A01G010905 | *Glu-A3* | 0.05 | 0.19 | 0.50 | 0.72 | 0.68 | Low Protein Content |
| TraesCS1A01G317311 | *Glu-1Ax* | 0.03 | 0.38 | 0.75 | 0.69 | 0.49 | Low Protein Content |
| TraesCS1B01G329992 | *Glu-1By* | 0.01 | 0.08 | 0.04 | 0.20 | 0.20 | Low Protein Content |
| TraesCS1D01G007400 | *Glu-D3* | 0.05 | 0.11 | 0.14 | 0.20 | 0.10 | Low Protein Content |
| TraesCS1D01G009400 | *Glu-D3* | 0.02 | 0.08 | 0.00 | 0.09 | 0.23 | Low Protein Content |
| TraesCS1D01G009900 | *Glu-D3* | 0.32 | 0.24 | 0.25 | 0.38 | 0.27 | Low Protein Content |
| TraesCS3A01G077900 | *TaNAC019-3A* | 0.02 | 0.08 | 0.07 | 0.12 | 0.27 | Low Protein Content |

| | | | | | | | |
|---|---|---|---|---|---|---|---|
| TraesCS1A01G058400 | *TaTOE-A1* | 0.01 | 0.00 | 0.00 | 0.00 | 0.00 | Flowring Date |
| TraesCS1A01G220300 | *WCO-1* | 0.00 | 0.03 | 0.07 | 0.14 | 0.02 | Flowring Date |
| TraesCS1A01G338600 | *TaFT3-A1* | 0.81 | 0.73 | 0.79 | 0.92 | 0.96 | Flowring Date |
| TraesCS3B01G007400 | *TaMFT_3B2* | 0.00 | 0.00 | 0.00 | 0.03 | 0.00 | Flowring Date |
| TraesCS3B01G135400 | *TaGI-3B* | 0.00 | 0.00 | 0.00 | 0.00 | 0.00 | Flowring Date |
| TraesCS3D01G118200 | *TaGI-3D* | 0.99 | 1.00 | 1.00 | 1.00 | 1.00 | Flowring Date |
| TraesCS4B01G346700 | *WSOC1-4B* | 0.00 | 0.00 | 0.00 | 0.02 | 0.00 | Flowring Date |
| TraesCS5A01G515500 | *WSOC1-5A* | 0.00 | 0.00 | 0.00 | 0.02 | 0.00 | Flowring Date |
| TraesCS6A01G091300 | *TaOGT-6A* | 0.00 | 0.00 | 0.00 | 0.02 | 0.00 | Flowring Date |
| TraesCS6B01G119500 | *TaOGT-6B* | 0.08 | 0.49 | 0.79 | 0.72 | 0.87 | Flowring Date |
| TraesCS6B01G319500 | *TaHd1-6B* | 0.27 | 0.59 | 0.89 | 0.89 | 0.61 | Flowring Date |
| TraesCS7D01G176700 | *TaVRT-2* | 0.01 | 0.16 | 0.21 | 0.23 | 0.09 | Flowring Date |
| TraesCS2B01G200800 | *FUL3-2B* | 0.01 | 0.00 | 0.00 | 0.00 | 0.00 | Flowring Date |
| TraesCS7B01G020800 | *SEP1-4-7B* | 0.02 | 0.32 | 0.43 | 0.38 | 0.61 | Flowring Date |
| TraesCS7D01G120500 | *SEP1-4-7D* | 0.99 | 1.00 | 1.00 | 1.00 | 1.00 | Flowring Date |
| TraesCS7A01G423100 | *TaZIM-A1* | 0.00 | 0.00 | 0.00 | 0.02 | 0.00 | Flowring Date |
| TraesCS7D01G416000 | *TaZIM-D1* | 0.99 | 1.00 | 1.00 | 1.00 | 1.00 | Flowring Date |
| TraesCS2B01G202300 | *TaGW7* | 0.01 | 0.22 | 0.11 | 0.14 | 0.33 | Thousand Grain Weight |
| TraesCS2B01G201900 | *TaCYP78A5-2B* | 0.98 | 1.00 | 0.93 | 1.00 | 1.00 | Thousand Grain Weight |
| TraesCS2B01G578500 | *TaARF12-2B* | 0.04 | 0.11 | 0.07 | 0.14 | 0.14 | Thousand Grain Weight |
| TraesCS2D01G548900 | *TaARF12-2D* | 0.00 | 0.00 | 0.00 | 0.02 | 0.00 | Thousand Grain Weight |
| TraesCS2A01G555600 | *TaSTT3b-2A* | 0.00 | 0.00 | 0.00 | 0.02 | 0.01 | Thousand Grain Weight |
| TraesCS2B01G587900 | *TaSTT3b-2B* | 0.00 | 0.00 | 0.00 | 0.00 | 0.01 | Thousand Grain Weight |
| TraesCS5B01G039800 | *TaARF25-5B* | 0.01 | 0.00 | 0.00 | 0.09 | 0.13 | Thousand Grain Weight |
| TraesCS6D01G176900 | *TaGW2-D1* | 0.01 | 0.00 | 0.00 | 0.00 | 0.00 | Thousand Grain Weight |
| TraesCS7D01G416000 | *TaZIM-D1* | 0.01 | 0.00 | 0.00 | 0.00 | 0.00 | Thousand Grain Weight |
| TraesCS6D01G207100 | *TaPRR1-D1* | 0.01 | 0.00 | 0.00 | 0.00 | 0.01 | Thousand Grain Weight |
| TraesCS6D01G168200 | *TaBT1-6D* | 0.01 | 0.00 | 0.00 | 0.02 | 0.07 | Thousand Grain Weight |
| TraesCS6A01G392400 | *KAT-2A* | 0.01 | 0.08 | 0.18 | 0.45 | 0.47 | Thousand Grain Weight |
| TraesCS7A01G423100 | *TaZIM-A1* | 0.03 | 0.19 | 0.43 | 0.31 | 0.33 | Thousand Grain Weight |
| TraesCS2D01G558800 | *TaSTT3b-2D* | 0.19 | 0.49 | 0.75 | 0.85 | 0.52 | Thousand Grain Weight |
| TraesCS3A01G311100 | *CKX2.2.1_3A* | 0.02 | 0.51 | 0.79 | 0.92 | 0.96 | Thousand Grain Weight |
| TraesCS2B01G200800 | *FUL3-2B* | 0.00 | 0.16 | 0.25 | 0.18 | 0.44 | Plant Height |
| TraesCS4A01G105300 | *TaPRR73-4A* | 0.03 | 0.35 | 0.46 | 0.58 | 0.75 | Plant Height |
| TraesCS4B01G198700 | *TaPRR73-4B* | 0.03 | 0.32 | 0.50 | 0.43 | 0.16 | Plant Height |
| TraesCS4A01G005600 | *TaSRL1-4A* | 0.00 | 0.00 | 0.04 | 0.09 | 0.07 | Plant Height |
| TraesCS5A01G543100 | *Rht12* | 0.40 | 0.49 | 0.39 | 0.52 | 0.86 | Plant Height |
| TraesCS5B01G208700 | *TaDEP1-5B* | 0.00 | 0.00 | 0.00 | 0.02 | 0.04 | Plant Height |
| TraesCS2B01G578500 | *TaARF12-2B* | 0.04 | 0.11 | 0.07 | 0.14 | 0.14 | Plant Height |
| TraesCS2D01G548900 | *TaARF12-2D* | 0.13 | 0.54 | 0.93 | 0.97 | 0.95 | Plant Height |
| TraesCS6D01G207100 | *TaPRR1-D1* | 0.02 | 0.19 | 0.18 | 0.08 | 0.07 | Plant Height |

**Supplementary Table S34.** Distribution of the superior haplotypes of 164 genes for future breeding

| Trait | Gene | Superior haplotype | Cultivars with superior haplotypes |
|---|---|---|---|
| FD | TraesCS1A01G058400 | 1\|39 | HXM239, HXM280, HXM635 |
| FD | TraesCS1A01G220300 | 1\|7 | HXM104, HXM399, HXM409, HXM415, HXM429, HXM430, HXM431, HXM433, HXM439, HXM445, HXM446, HXZ459, HXM471, HXM484, HXM525, HXM535 |
| FD | TraesCS1A01G372300 | 1\|6 | HXM280, HXM320, HXM336, HXM483 |
| FD | TraesCS1D01G378800 | 2\|3 | HXM20, HXM381, HXM387, HXM389, HXM393, HXM394, HXM415, HXM422, HXM440, HXM447, HXM448, HXZ452, HXZ459, HXM465, HXM468, HXM473, HXM491, HXM515, HXM516, HXM520, HXM522, HXM533, HXM534, HXM535, HXM538, HXM540, HXM541, HXZ552, HXZ557, HXM561, HXM562, HXM570, HXM586, HXM595, HXM598, HXM599, HXM600, HXM602, HXM607, HXM608, HXM609, HXM610, HXM613, HXM615, HXM617, HXM620 |
| FD | TraesCS2A01G119600 | 1\|21 | HXM333 |
| FD | TraesCS2A01G179000 | 1\|10 | HXM478 |
| FD | TraesCS2A01G179200 | 1\|8 | HXM423, HXM534 |
| FD | TraesCS2B01G200800 | 1\|40 | HXM239, HXM635 |
| FD | TraesCS3B01G007400 | 1\|5 | HXM428, HXZ461 |
| FD | TraesCS3B01G017400 | 1\|19 | HXM22, HXM28, HXM37, HXM55, HXM59, HXZ77, HXZ78, HXZ79, HXM81, HXM83, HXM85, HXM89, HXM93, HXM96, HXM102, HXM105, HXM111, HXM125, HXM127, HXM129, HXM130, HXM135, HXM137, HXM138, HXM139, HXM141, HXM142, HXM144, HXM146, HXM148, HXM151, HXM154, HXZ172, HXZ173, HXM179, HXM181, HXM184, HXM185, HXM188, HXM190, HXM191, HXM192, HXM194, HXM195, HXM196, HXM197, HXM198, HXM199, HXM207, HXM209, HXM221, HXM225, HXM234, HXM239, HXM243, HXM247, HXM249, HXM253, HXM254, HXZ263, HXZ270, HXZ271, HXM275, HXM278, HXM284, HXM285, HXM288, HXM289, HXM292, HXM293, HXM296, HXM299, HXM303, HXM304, HXM307, HXM309, HXM314, HXM319, HXM321, HXM322, HXM324, HXM326, HXM327, HXM329, HXM330, HXM336, HXM337, HXM339, HXM340, HXM342, HXM345, HXM348, HXM349, HXM351, HXM352, HXZ353, HXZ354, HXZ356, HXZ357, HXZ361, HXZ362, HXZ366, HXM369, HXM374, HXM377, HXM635 |
| FD | TraesCS3B01G135400 | 1\|10 | HXM282 |
| FD | TraesCS3B01G428100 | 1\|6 | HXM226, HXM400, HXM401, HXM438, HXM442, HXM443, HXM445, HXZ456, HXM466, HXM533 |
| FD | TraesCS3B01G517100 | 1\|21 | HXM298, HXM300 |
| FD | TraesCS4A01G044800 | 1\|10 | HXM280 |
| FD | TraesCS4A01G324900 | 1\|7 | HXZ463, HXM599 |
| FD | TraesCS4B01G317400 | 1\|2 | HXZ75, HXM84, HXZ163, HXZ171, HXM273, HXM379, HXM390, HXM394, HXM398, HXM405, HXM407, HXM409, HXM419, HXM422, HXM426, HXM427, HXM436, HXM448, HXZ452, HXZ457, HXZ459, HXM465, HXM468, HXM469, HXM471, HXM472, HXM473, HXM479, HXM480, HXM485, HXM491, HXM492, HXM495, HXM498, HXM511, HXM513, HXM515, HXM522, HXM523, HXM533, |

| | | | |
|---|---|---|---|
| | | | HXM536, HXM537, HXM541, HXM543, HXM544, HXZ550, HXZ551, HXZ553, HXZ555, HXZ558, HXZ560, HXM561, HXM562, HXM565, HXM569, HXM574, HXM575, HXM576, HXM577, HXM586, HXM587, HXM591, HXM592, HXM594, HXM595, HXM596, HXM607, HXM608, HXM609, HXM613, HXM615, HXM618, HXM619, HXM622 |
| FD | TraesCS4B01G346700 | 1\|10 | HXZ449 |
| FD | TraesCS5A01G393400 | 2\|11 | HXM423, HXZ449 |
| FD | TraesCS5A01G515500 | 1\|13 | HXM445 |
| FD | TraesCS6A01G091300 | 1\|26 | HXM434 |
| FD | TraesCS7A01G423100 | 1\|14 | HXM443 |
| FD | TraesCS7D01G176700 | 2\|8 | HXM104, HXZ163, HXM226, HXM379, HXM384, HXM390, HXM391, HXM392, HXM395, HXM405, HXM406, HXM408, HXM409, HXM419, HXM425, HXM426, HXM430, HXM434, HXM436, HXM438, HXM445, HXM446, HXZ449, HXZ450, HXZ453, HXZ464, HXM467, HXM471, HXM474, HXM482, HXM498, HXM508, HXM509, HXM517, HXM536, HXZ545, HXZ551, HXM607, HXM608, HXM619, HXM622, HXM637 |
| FD | TraesCS7D01G183600 | 2\|6 | HXM20, HXM30, HXM229, HXM279, HXM311, HXM378, HXM380, HXM382, HXM385, HXM389, HXM396, HXM401, HXM403, HXM404, HXM411, HXM412, HXM413, HXM417, HXM420, HXM421, HXM423, HXM425, HXM427, HXM428, HXM429, HXM431, HXM432, HXM433, HXM435, HXM440, HXM443, HXM444, HXM446, HXZ449, HXZ450, HXZ453, HXZ454, HXZ456, HXZ459, HXZ460, HXZ461, HXZ462, HXZ464, HXM465, HXM467, HXM469, HXM470, HXM471, HXM472, HXM475, HXM476, HXM477, HXM479, HXM480, HXM482, HXM486, HXM487, HXM488, HXM489, HXM490, HXM491, HXM492, HXM499, HXM501, HXM502, HXM503, HXM504, HXM505, HXM506, HXM508, HXM509, HXM510, HXM511, HXM512, HXM517, HXM521, HXM522, HXM524, HXM528, HXM530, HXM532, HXM533, HXM534, HXM541, HXM542, HXM543, HXZ545, HXZ546, HXZ547, HXZ549, HXZ550, HXZ552, HXZ553, HXZ556, HXZ559, HXM562, HXM563, HXM565, HXM566, HXM573, HXM587, HXM588, HXM591, HXM593, HXM596, HXM600, HXM604, HXM608, HXM610, HXM612, HXM613, HXM614, HXM615, HXM619, HXM621, HXM622, HXM637, HXM638 |
| GL | TraesCS1A01G273900 | 1\|2 | HXM232, HXM239, HXM280, HXM635 |
| GL | TraesCS2A01G482400 | 1\|17 | HXM541 |
| GL | TraesCS2D01G011200 | 1\|3 | HXM239, HXZ267, HXM280, HXM635 |
| GL | TraesCS2D01G016900 | 1\|4 | HXM232, HXM239, HXZ267, HXM635 |
| GL | TraesCS2D01G082000 | 1\|2 | HXM109, HXM239, HXZ269, HXM280, HXM343, HXM372, HXM423, HXM635 |
| GL | TraesCS2D01G482500 | 1\|12 | HXM232 |
| GL | TraesCS3B01G560400 | 3\|4 | HXM38, HXM111, HXM125, HXM154, HXM182, HXM209, HXM220, HXM224, HXM232, HXM239, HXM280, HXM292, HXM296, HXM300, HXM307, HXM313, HXM348, HXM349, HXM351, HXZ354, HXZ356, HXZ363, HXM423, HXM429, HXM436, HXM447, HXM448, HXZ450, HXZ452, HXZ455, HXZ462, HXM465, HXM468, HXM473, HXM478, HXM484, HXM485, HXM490, HXM495, HXM496, HXM498, HXM499, HXM500, HXM510, HXM511, HXM513, HXM515, HXM518, HXM519, HXM522, HXM523, HXM529, HXM533, HXM534, HXM535, HXM537, |

| | | | |
|---|---|---|---|
| | | | HXM538, HXM540, HXM541, HXM544, HXZ549, HXZ550, HXZ551, HXZ556, HXZ557, HXZ558, HXZ560, HXM564, HXM565, HXM567, HXM570, HXM572, HXM574, HXM591, HXM592, HXM595, HXM596, HXM597, HXM599, HXM602, HXM603, HXM604, HXM609, HXM611, HXM612, HXM613, HXM614, HXM615, HXM617, HXM618, HXM620, HXM623, HXM634, HXM635 |
| GL | TraesCS3B01G561000 | 2\|4 | HXM38, HXM125, HXM154, HXM182, HXM209, HXM220, HXM224, HXM232, HXM239, HXM280, HXM292, HXM296, HXM300, HXM307, HXM313, HXM348, HXM349, HXM351, HXZ354, HXZ356, HXZ363, HXM429, HXM436, HXM447, HXM448, HXZ450, HXZ452, HXZ455, HXZ462, HXM465, HXM468, HXM473, HXM478, HXM481, HXM484, HXM485, HXM490, HXM495, HXM496, HXM498, HXM499, HXM500, HXM510, HXM511, HXM513, HXM515, HXM518, HXM519, HXM522, HXM523, HXM529, HXM533, HXM534, HXM535, HXM537, HXM538, HXM540, HXM541, HXM544, HXZ549, HXZ550, HXZ551, HXZ556, HXZ557, HXZ558, HXZ560, HXM564, HXM565, HXM567, HXM570, HXM572, HXM574, HXM591, HXM592, HXM595, HXM596, HXM597, HXM599, HXM602, HXM603, HXM604, HXM609, HXM611, HXM612, HXM613, HXM614, HXM615, HXM617, HXM618, HXM620, HXM623, HXM634, HXM635 |
| GL | TraesCS4A01G266900 | 1\|6 | HXZ73, HXM239, HXM635 |
| GL | TraesCS4A01G444300 | 2\|37 | HXM239, HXM635 |
| GL | TraesCS4B01G321800 | 1\|32 | HXM232, HXM239, HXM635 |
| GL | TraesCS4B01G360800 | 1\|39 | HXM232 |
| GL | TraesCS4D01G318400 | 1\|2 | HXM232, HXM239, HXM280, HXM635 |
| GL | TraesCS4D01G325200 | 2\|4 | HXM232, HXM239, HXM280, HXM635 |
| GL | TraesCS5A01G377700 | 2\|9 | HXM21, HXM153, HXM239, HXZ268, HXM370, HXZ551, HXM597, HXM635 |
| GL | TraesCS5B01G307600 | 2\|3 | HXZ9, HXZ14, HXM20, HXM21, HXM30, HXZ69, HXM120, HXM126, HXM133, HXM148, HXZ161, HXZ163, HXM179, HXM185, HXM188, HXM191, HXM192, HXM194, HXM197, HXM199, HXM208, HXM231, HXM232, HXM239, HXM243, HXM249, HXM253, HXZ263, HXZ271, HXM280, HXM286, HXM297, HXM309, HXM318, HXM333, HXM335, HXM344, HXZ364, HXM374, HXM378, HXM381, HXM382, HXM387, HXM389, HXM390, HXM393, HXM394, HXM395, HXM397, HXM398, HXM399, HXM400, HXM403, HXM404, HXM405, HXM408, HXM411, HXM413, HXM415, HXM417, HXM419, HXM426, HXM431, HXM437, HXM438, HXM439, HXM440, HXZ453, HXZ455, HXZ458, HXZ462, HXZ464, HXM466, HXM467, HXM472, HXM475, HXM481, HXM484, HXM493, HXM499, HXM505, HXZ546, HXZ548, HXZ553, HXZ554, HXM569, HXM573, HXM576, HXM590, HXM611, HXM616, HXM629, HXM632, HXM634, HXM635, HXM640 |
| GL | TraesCS5B01G375800 | 1\|22 | HXM232, HXM239, HXM280, HXM635 |
| GL | TraesCS6A01G009100 | 1\|30 | HXM239, HXM635 |
| GL | TraesCS6A01G382600 | 1\|3 | HXZ1, HXZ3, HXM33, HXZ76, HXM117, HXM118, HXM126, HXM127, HXM128, HXM133, HXM139, HXZ163, HXZ167, HXZ169, HXZ172, HXM196, HXM231, HXM232, HXM237, HXM239, HXM254, HXZ261, HXZ265, HXZ268, HXM280, HXM282, HXM296, HXM304, HXM337, HXM344, HXM346, HXM370, HXM371, HXM378, HXM379, HXM382, HXM385, HXM386, HXM387, HXM388, HXM389, HXM390, HXM393, HXM395, HXM396, HXM398, HXM400, HXM401, |

| | | | |
|---|---|---|---|
| | | | HXM402, HXM405, HXM407, HXM408, HXM409, HXM413, HXM418, HXM419, HXM420, HXM423, HXM424, HXM425, HXM427, HXM436, HXM439, HXM442, HXZ449, HXZ450, HXZ451, HXZ453, HXZ456, HXM466, HXM467, HXM468, HXM469, HXM471, HXM476, HXM477, HXM483, HXM486, HXM488, HXM490, HXM493, HXM494, HXM495, HXM496, HXM497, HXM498, HXM506, HXM507, HXM508, HXM510, HXM511, HXM512, HXM514, HXM517, HXM518, HXM520, HXM526, HXM527, HXM532, HXM536, HXM537, HXM540, HXM544, HXZ551, HXZ557, HXZ559, HXM589, HXM590, HXM591, HXM592, HXM594, HXM602, HXM603, HXM609, HXM610, HXM611, HXM617, HXM618, HXM619, HXM620, HXM624, HXM625, HXM626, HXM627, HXM628 |
| GL | TraesCS6D01G378300 | 1\|4 | HXM232, HXM635 |
| GL | TraesCS7B01G018600 | 1\|2 | HXM232, HXM239, HXM280, HXM635 |
| GL | TraesCS7D01G039500 | 2\|5 | HXM239, HXM635 |
| GW | TraesCS1B01G155700 | 1\|3 | HXZ163, HXM392, HXM395, HXM398, HXM406, HXM409, HXM422, HXM424, HXM426, HXM434, HXM436, HXM444, HXZ459, HXM478, HXM501, HXM507, HXM516, HXM532, HXM563, HXM564, HXM621, HXM632 |
| GW | TraesCS1B01G462000 | 11\|24 | HXM21, HXM300, HXM375, HXM378, HXM380, HXM382, HXM386, HXM391, HXM415, HXM419, HXM440, HXM446, HXM447, HXZ450, HXZ452, HXM465, HXM467, HXM473, HXM496, HXM497, HXM498, HXM508, HXM510, HXM513, HXM517, HXM519, HXM522, HXM529, HXM534, HXM535, HXM539, HXM540, HXM541, HXZ549, HXZ550, HXZ551, HXZ555, HXZ560, HXM567, HXM568, HXM570, HXM571, HXM572, HXM589, HXM593, HXM597, HXM600, HXM606, HXM612, HXM614, HXM617, HXM622, HXM624, HXM625, HXM626, HXM627, HXM628, HXM629 |
| GW | TraesCS2A01G464000 | 1\|32 | HXM541 |
| GW | TraesCS2B01G485700 | 2\|5 | HXM104, HXM379, HXM384, HXM386, HXM391, HXM392, HXM394, HXM400, HXM409, HXM410, HXM416, HXM418, HXM420, HXM422, HXM429, HXM430, HXM431, HXM437, HXM439, HXM442, HXM443, HXZ449, HXZ450, HXZ454, HXZ456, HXZ458, HXZ459, HXZ460, HXM466, HXM472, HXM476, HXM480, HXM481, HXM490, HXM494, HXM495, HXM502, HXM516, HXM520, HXM524, HXM530, HXM532, HXM534, HXM536, HXM537, HXM538, HXM541, HXZ548, HXZ550, HXZ554, HXZ557, HXZ558, HXZ560, HXM561, HXM566, HXM569, HXM574, HXM575, HXM576, HXM577, HXM587, HXM588, HXM591, HXM600, HXM603, HXM611, HXM614, HXM615, HXM618, HXM620, HXM640 |
| GW | TraesCS2D01G505700 | 4\|20 | HXM392, HXM399, HXM442, HXZ460, HXM476, HXM494, HXM541, HXZ559, HXM591, HXM600, HXM622, HXM640 |
| GW | TraesCS6B01G015100 | 2\|45 | HXM393, HXM613 |
| GW | TraesCS6B01G130700 | 1\|40 | HXM624 |
| GW | TraesCS6B01G186700 | 1\|16 | HXZ449 |
| GW | TraesCS7A01G043700 | 9\|37 | HXM300, HXM400, HXM423, HXM437, HXM441, HXZ449, HXZ464, HXM471, HXM480, HXM499, HXM521, HXZ548, HXM600, HXM605, HXM611 |
| GW | TraesCS7A01G120300 | 1\|18 | HXM525 |
| GWLR | TraesCS2A01G548100 | 1\|29 | HXZ368 |

| | | | |
|---|---|---|---|
| GWLR | TraesCS2D01G011200 | 1\|3 | HXZ11, HXZ16, HXM25, HXM62, HXZ69, HXZ70, HXZ74, HXM117, HXM118, HXM121, HXM122, HXM124, HXM128, HXM155, HXM156, HXM159, HXM160, HXZ162, HXZ167, HXZ168, HXM178, HXM186, HXM187, HXM193, HXM223, HXM240, HXM244, HXZ260, HXZ264, HXM277, HXM295, HXM297, HXM300, HXM308, HXM314, HXM319, HXM324, HXM325, HXM329, HXM331, HXM333, HXM334, HXM337, HXM341, HXM344, HXZ358, HXZ359, HXZ361, HXZ362, HXZ364, HXM395, HXZ559 |
| GWLR | TraesCS3B01G128700 | 1\|8 | HXZ4, HXZ7, HXZ10, HXM20, HXM32, HXM39, HXM42, HXM50, HXM63, HXZ80, HXM88, HXM97, HXM106, HXM109, HXM114, HXM134, HXM142, HXM148, HXM154, HXM159, HXZ162, HXM192, HXM198, HXM199, HXM201, HXM209, HXM231, HXZ259, HXZ261, HXZ266, HXM275, HXM278, HXM329, HXM343, HXZ358, HXM372, HXM377, HXM383, HXM401, HXM410, HXM421, HXM473, HXM493, HXM533, HXM535, HXM536, HXM538, HXM540 |
| GWLR | TraesCS4A01G074300 | 6\|8 | HXZ3, HXZ9, HXZ13, HXZ16, HXM24, HXM32, HXM34, HXM39, HXM141, HXM148, HXM153, HXM156, HXM157, HXM158, HXM159, HXM160, HXZ161, HXZ162, HXZ167, HXZ169, HXZ170, HXZ171, HXZ172, HXZ173, HXZ174, HXZ176, HXM177, HXM178, HXM179, HXM180, HXM181, HXM182, HXM183, HXM184, HXM185, HXM186, HXM187, HXM188, HXM189, HXM190, HXM191, HXM192, HXM193, HXM194, HXM195, HXM197, HXM198, HXM199, HXM220, HXM223, HXM224, HXM240, HXM243, HXM249, HXM252, HXM253, HXM254, HXM256, HXZ259, HXZ263, HXZ267, HXZ268, HXZ270, HXZ271, HXM275, HXM277, HXM303, HXM304, HXM307, HXM309, HXM314, HXM318, HXM319, HXM324, HXM325, HXM329, HXM334, HXM337, HXM340, HXM342, HXM350, HXM352, HXZ353, HXZ357, HXZ358, HXZ359, HXZ361, HXZ362, HXZ363, HXM369, HXM370, HXM371, HXM374, HXM377, HXM391, HXZ458, HXM561, HXM566, HXM637, HXM638 |
| GWLR | TraesCS5B01G375800 | 5\|22 | HXZ6, HXZ167, HXM432, HXZ452, HXM467, HXM488, HXM532 |
| GWLR | TraesCS6A01G009100 | 1\|30 | HXM444, HXM472 |
| GWLR | TraesCS6B01G015100 | 6\|45 | HXM112, HXM184, HXM198, HXZ261, HXM309, HXM340, HXM348, HXM352, HXM526, HXM613 |
| GWLR | TraesCS7A01G549100 | 1\|15 | HXM512 |
| GWLR | TraesCS7A01G549200 | 4\|18 | HXM109, HXM229, HXZ360, HXZ559 |
| GWLR | TraesCS7A01G549300 | 1\|13 | HXM229, HXM300 |
| HD | TraesCS1A01G301400 | 1\|3 | HXM30, HXM396, HXM403, HXM404, HXM411, HXM417, HXM425, HXM435, HXZ456, HXM483, HXM494, HXM518, HXM526, HXM529, HXM569, HXM573, HXM575, HXM576, HXM577 |
| HD | TraesCS1A01G435000 | 1\|16 | HXM232, HXM280, HXM334 |
| HD | TraesCS1D01G444100 | 1\|6 | HXM21, HXM226, HXM280, HXZ452 |
| HD | TraesCS3B01G017400 | 1\|19 | HXM22, HXM28, HXM37, HXM55, HXM59, HXZ77, HXZ78, HXZ79, HXM81, HXM83, HXM85, HXM89, HXM93, HXM96, HXM102, HXM105, HXM111, HXM125, HXM127, HXM129, HXM130, HXM135, HXM137, HXM138, HXM139, HXM141, HXM142, HXM144, HXM146, HXM148, HXM151, HXM154, HXZ172, HXZ173, HXM179, HXM181, HXM184, HXM185, HXM188, HXM190, HXM191, HXM192, HXM194, HXM195, HXM196, HXM197, HXM198, HXM199, HXM207, |

| | | | |
|---|---|---|---|
| | | | HXM209, HXM221, HXM225, HXM234, HXM239, HXM243, HXM247, HXM249, HXM253, HXM254, HXZ263, HXZ270, HXZ271, HXM275, HXM278, HXM284, HXM285, HXM288, HXM289, HXM292, HXM293, HXM296, HXM299, HXM303, HXM304, HXM307, HXM309, HXM314, HXM319, HXM321, HXM322, HXM324, HXM326, HXM327, HXM329, HXM330, HXM336, HXM337, HXM339, HXM340, HXM342, HXM345, HXM348, HXM349, HXM351, HXM352, HXZ353, HXZ354, HXZ356, HXZ357, HXZ361, HXZ362, HXZ366, HXM369, HXM374, HXM377, HXM635 |
| HD | TraesCS3B01G517100 | 1\|21 | HXM298, HXM300 |
| HD | TraesCS5B01G538100 | 1\|20 | HXM139 |
| PH | TraesCS1B01G462300 | 2\|9 | HXM21, HXM232, HXM239, HXM380, HXM382, HXM391, HXM440, HXM446, HXM447, HXZ450, HXZ452, HXZ457, HXM465, HXM467, HXM473, HXM496, HXM497, HXM498, HXM508, HXM510, HXM513, HXM517, HXM519, HXM522, HXM529, HXM535, HXM540, HXM544, HXZ549, HXZ550, HXZ551, HXZ560, HXM567, HXM568, HXM570, HXM571, HXM572, HXM593, HXM597, HXM606, HXM612, HXM614, HXM617, HXM622, HXM624, HXM625, HXM626, HXM627, HXM628, HXM629, HXM635 |
| PH | TraesCS2A01G453300 | 1\|2 | HXZ6, HXM21, HXM45, HXM227, HXM239, HXM385, HXM388, HXM389, HXM398, HXM402, HXM410, HXM447, HXZ462, HXM482, HXM484, HXM485, HXM502, HXM507, HXM531, HXZ547, HXZ548, HXM563, HXM564, HXM599, HXM601, HXM605, HXM621, HXM632, HXM634, HXM635, HXM639 |
| PH | TraesCS2A01G552400 | 1\|4 | HXM239, HXM388, HXM397, HXM402, HXM412, HXM416, HXM430, HXM432, HXM442, HXM445, HXZ460, HXZ464, HXM476, HXM491, HXM539, HXM541, HXZ551, HXM565, HXM597, HXM607, HXM608, HXM615, HXM632, HXM635, HXM640 |
| PH | TraesCS2B01G200800 | 7\|40 | HXM383, HXM392, HXM395, HXM396, HXM398, HXM413, HXM418, HXM419, HXM420, HXM423, HXM436, HXM447, HXZ455, HXZ460, HXZ461, HXZ463, HXM465, HXM479, HXM486, HXM488, HXM490, HXM491, HXM493, HXM503, HXM511, HXM513, HXM527, HXM530, HXM532, HXM534, HXM536, HXM539, HXM540, HXM541, HXM543, HXM544, HXZ545, HXZ546, HXZ548, HXZ549, HXZ550, HXZ551, HXZ553, HXZ557, HXZ558, HXZ559, HXZ560, HXM564, HXM565, HXM566, HXM569, HXM570, HXM571, HXM573, HXM574, HXM575, HXM577, HXM586, HXM595, HXM596, HXM598, HXM600, HXM602, HXM603, HXM604, HXM605, HXM606, HXM607, HXM609, HXM611, HXM612, HXM613, HXM614, HXM615, HXM616, HXM617, HXM618, HXM619, HXM620, HXM622, HXM623, HXM637, HXM638, HXM640 |
| PH | TraesCS2B01G575900 | 2\|6 | HXM21, HXM34, HXM443, HXZ456, HXM470, HXM474, HXM477, HXM486, HXM491, HXM494, HXM562, HXM565, HXM586, HXM605, HXM607, HXM608, HXM609, HXM628, HXM640 |
| PH | TraesCS2B01G578500 | 1\|3 | HXM38, HXM45, HXM61, HXM62, HXM96, HXM99, HXM111, HXM226, HXM231, HXM233, HXM255, HXZ258, HXM329, HXM335, HXM385, HXM389, HXM398, HXM408, HXM421, HXM435, HXZ451, HXZ454, HXZ462, HXM478, HXM481, HXM485, HXM495, HXM499, HXM500, HXM502, HXM523, HXM527, |

| | | | |
|---|---|---|---|
| | | | HXM531, HXM533, HXM544, HXZ548, HXZ549, HXZ560, HXM563, HXM574, HXM587, HXM596, HXM599, HXM613, HXM618, HXM621, HXM632, HXM639 |
| PH | TraesCS2B01G578500 | 1\|3 | HXM38, HXM45, HXM61, HXM62, HXM96, HXM99, HXM111, HXM226, HXM231, HXM233, HXM255, HXZ258, HXM329, HXM335, HXM385, HXM389, HXM398, HXM408, HXM421, HXM435, HXZ451, HXZ454, HXZ462, HXM478, HXM481, HXM485, HXM495, HXM499, HXM500, HXM502, HXM523, HXM527, HXM531, HXM533, HXM544, HXZ548, HXZ549, HXZ560, HXM563, HXM574, HXM587, HXM596, HXM599, HXM613, HXM618, HXM621, HXM632, HXM639 |
| PH | TraesCS3D01G540600 | 1\|26 | HXM159 |
| PH | TraesCS4A01G005600 | 5\|14 | HXM416, HXM430, HXM434, HXZ449, HXZ459, HXM468, HXM473, HXM488, HXM497, HXM501, HXM508, HXM518, HXM529, HXZ557, HXM587, HXM605 |
| PH | TraesCS4A01G005600 | 5\|14 | HXM416, HXM430, HXM434, HXZ449, HXZ459, HXM468, HXM473, HXM488, HXM497, HXM501, HXM508, HXM518, HXM529, HXZ557, HXM587, HXM605 |
| PH | TraesCS4A01G455200 | 1\|9 | HXM229, HXM311, HXM383, HXM391, HXM398, HXM399, HXM400, HXM401, HXM412, HXM414, HXM428, HXM431, HXM432, HXM433, HXM440, HXM441, HXM445, HXM447, HXZ460, HXM471, HXM480, HXM481, HXM482, HXM484, HXM487, HXM488, HXM530, HXZ546, HXZ556, HXZ559, HXM564, HXM569, HXM573, HXM575, HXM576, HXM577, HXM588, HXM604, HXM605, HXM639 |
| PH | TraesCS4A01G460100 | 8\|44 | HXM95, HXM274, HXM284, HXM338, HXM379, HXM383, HXM384, HXM392, HXM393, HXM394, HXM395, HXM396, HXM401, HXM402, HXM403, HXM404, HXM412, HXM413, HXM414, HXM415, HXM416, HXM417, HXM422, HXM425, HXM428, HXM430, HXM431, HXM432, HXM433, HXM434, HXM435, HXM436, HXM437, HXM439, HXM440, HXM442, HXM443, HXM444, HXM445, HXM447, HXZ449, HXZ450, HXZ451, HXZ456, HXZ459, HXZ460, HXZ464, HXM466, HXM467, HXM471, HXM474, HXM475, HXM476, HXM478, HXM480, HXM481, HXM482, HXM484, HXM488, HXM489, HXM490, HXM499, HXM501, HXM502, HXM504, HXM507, HXM510, HXM514, HXM516, HXM521, HXM524, HXM530, HXM531, HXM534, HXM537, HXM542, HXZ546, HXZ548, HXZ554, HXZ557, HXZ559, HXM563, HXM564, HXM566, HXM569, HXM573, HXM575, HXM576, HXM577, HXM586, HXM588, HXM592, HXM598, HXM599, HXM600, HXM602, HXM604, HXM605, HXM620, HXM621, HXM634, HXM639, HXM640 |
| PH | TraesCS4B01G198700 | 5\|9 | HXM34, HXM104, HXM153, HXZ163, HXM227, HXZ268, HXM311, HXM335, HXM370, HXM379, HXM384, HXM390, HXM392, HXM393, HXM395, HXM397, HXM398, HXM402, HXM403, HXM404, HXM405, HXM406, HXM407, HXM408, HXM413, HXM414, HXM415, HXM416, HXM417, HXM419, HXM420, HXM421, HXM423, HXM424, HXM425, HXM426, HXM428, HXM430, HXM431, HXM436, HXM437, HXM440, HXM441, HXM442, HXM443, HXM444, HXZ449, HXZ451, HXZ455, HXZ457, HXZ459, HXZ460, HXZ461, HXM470, HXM471, HXM472, HXM474, HXM476, HXM477, HXM478, HXM479, HXM486, HXM487, HXM491, HXM494, HXM501, HXM505, HXM507, HXM510, HXM512, HXM514, HXM524, HXM527, HXM539, HXZ545, HXZ548, HXZ551, HXZ553, HXM590, HXM608, HXM611, HXM612, HXM613, HXM616, HXM640 |
| PH | TraesCS5A01G330500 | 3\|6 | HXM526, HXM534, HXM592, HXM600 |
| PH | TraesCS5B01G208700 | 2\|15 | HXM490, HXM511, HXZ546, HXM572, HXM615, HXM621 |

| | | | |
|---|---|---|---|
| PH | TraesCS5B01G523100 | 2\|13 | HXM20, HXM378, HXM379, HXM385, HXM388, HXM393, HXM395, HXM418, HXM420, HXM423, HXM439, HXM440, HXM446, HXZ453, HXZ454, HXZ455, HXZ457, HXZ458, HXZ460, HXZ462, HXM467, HXM474, HXM478, HXM481, HXM482, HXM483, HXM485, HXM487, HXM488, HXM495, HXM496, HXM498, HXM499, HXM500, HXM503, HXM505, HXM507, HXM508, HXM509, HXM510, HXM511, HXM514, HXM515, HXM517, HXM520, HXM523, HXM524, HXM529, HXM532, HXM534, HXM535, HXM538, HXM539, HXM540, HXM544, HXZ545, HXZ547, HXZ548, HXZ550, HXZ551, HXZ552, HXZ556, HXZ558, HXZ560, HXM564, HXM568, HXM570, HXM571, HXM572, HXM574, HXM587, HXM588, HXM589, HXM590, HXM593, HXM594, HXM595, HXM596, HXM599, HXM601, HXM606, HXM610, HXM612, HXM617, HXM618, HXM623, HXM629, HXM634 |
| PH | TraesCS6D01G207100 | 1\|3 | HXM50, HXZ80, HXM84, HXM85, HXM104, HXM286, HXM383, HXM384, HXM392, HXM393, HXM398, HXM403, HXM413, HXM415, HXM419, HXM420, HXM422, HXM426, HXZ452, HXZ459, HXM466, HXM494, HXM504, HXM543, HXM544, HXM590, HXM603, HXM612, HXM616, HXM622, HXM637, HXM639 |
| PH | TraesCS7A01G424700 | 1\|9 | HXM232, HXM239, HXM635 |
| Pm | TraesCS7B01G441700 | 3\|105 | HXZ258, HXZ262, HXZ559 |
| Pm | TraesCS7D01G080300 | 1\|5 | HXM232, HXM239, HXM280, HXM635 |
| PPR | TraesCS2A01G521500 | 1\|19 | HXM423 |
| PPR | TraesCS2A01G552400 | 1\|4 | HXM239, HXM388, HXM397, HXM402, HXM412, HXM416, HXM430, HXM432, HXM442, HXM445, HXZ460, HXZ464, HXM476, HXM491, HXM539, HXM541, HXZ551, HXM565, HXM597, HXM607, HXM608, HXM615, HXM632, HXM635, HXM640 |
| PPR | TraesCS2B01G010100 | 2\|3 | HXZ5, HXM20, HXM34, HXM104, HXM145, HXM153, HXM229, HXM231, HXM232, HXM239, HXZ269, HXM311, HXM378, HXM379, HXM381, HXM384, HXM385, HXM386, HXM387, HXM389, HXM392, HXM393, HXM396, HXM397, HXM399, HXM400, HXM403, HXM404, HXM405, HXM407, HXM410, HXM412, HXM413, HXM414, HXM417, HXM419, HXM420, HXM421, HXM424, HXM425, HXM427, HXM428, HXM431, HXM432, HXM433, HXM435, HXM436, HXM439, HXM440, HXM441, HXM442, HXM443, HXM445, HXZ449, HXZ451, HXZ454, HXZ455, HXZ456, HXZ457, HXZ458, HXZ460, HXZ461, HXZ463, HXZ464, HXM467, HXM470, HXM471, HXM474, HXM475, HXM476, HXM477, HXM479, HXM480, HXM481, HXM484, HXM485, HXM486, HXM487, HXM488, HXM491, HXM492, HXM493, HXM494, HXM502, HXM507, HXM508, HXM509, HXM512, HXM514, HXM521, HXM522, HXM524, HXM527, HXM531, HXM533, HXM535, HXM538, HXM540, HXM542, HXZ547, HXZ548, HXZ551, HXZ555, HXZ559, HXM562, HXM563, HXM566, HXM570, HXM587, HXM589, HXM591, HXM597, HXM601, HXM603, HXM607, HXM608, HXM611, HXM613, HXM617, HXM621, HXM622, HXM623, HXM634, HXM635, HXM636, HXM639, HXM640 |
| PPR | TraesCS3A01G535100 | 1\|23 | HXZ461, HXM599 |
| PPR | TraesCS4B01G263400 | 3\|6 | HXZ75, HXM104, HXZ163, HXM300, HXM311, HXM379, HXM384, HXM385, HXM388, HXM389, HXM390, HXM392, HXM393, HXM394, HXM395, HXM397, HXM402, HXM403, HXM404, HXM405, HXM406, HXM407, HXM408, HXM409, HXM413, HXM415, HXM417, HXM418, HXM419, HXM420, HXM421, HXM422, |

| | | | |
|---|---|---|---|
| | | | HXM423, HXM424, HXM425, HXM426, HXM428, HXM429, HXM431, HXM434, HXM435, HXM436, HXM437, HXM440, HXM443, HXZ449, HXZ450, HXZ451, HXZ455, HXZ456, HXZ457, HXZ459, HXZ460, HXZ461, HXZ462, HXZ464, HXM469, HXM470, HXM471, HXM476, HXM477, HXM479, HXM484, HXM486, HXM487, HXM488, HXM489, HXM490, HXM491, HXM494, HXM504, HXM506, HXM512, HXM516, HXM527, HXM528, HXM529, HXM530, HXM532, HXM534, HXM542, HXM543, HXZ545, HXZ547, HXZ548, HXZ551, HXZ552, HXZ557, HXZ559, HXM566, HXM569, HXM575, HXM576, HXM577, HXM588, HXM599, HXM601, HXM603, HXM604, HXM607, HXM608, HXM610, HXM611, HXM612, HXM620, HXM632, HXM634, HXM637 |
| PPR | TraesCS5B01G377500 | 3\|11 | HXM20, HXM21, HXM24, HXM30, HXM34, HXM153, HXZ163, HXM226, HXM344, HXM371, HXM378, HXM379, HXM382, HXM385, HXM386, HXM389, HXM390, HXM391, HXM393, HXM394, HXM396, HXM398, HXM399, HXM400, HXM401, HXM403, HXM404, HXM405, HXM406, HXM407, HXM408, HXM410, HXM411, HXM413, HXM415, HXM417, HXM419, HXM425, HXM426, HXM427, HXM428, HXM429, HXM430, HXM431, HXM435, HXM441, HXM442, HXM448, HXZ449, HXZ450, HXZ451, HXZ452, HXZ453, HXZ454, HXZ455, HXZ456, HXZ457, HXZ458, HXZ459, HXZ464, HXM465, HXM466, HXM467, HXM471, HXM472, HXM473, HXM476, HXM478, HXM479, HXM480, HXM481, HXM483, HXM485, HXM487, HXM489, HXM490, HXM494, HXM495, HXM499, HXM500, HXM502, HXM512, HXM516, HXM521, HXM523, HXM527, HXM530, HXM533, HXM542, HXZ546, HXZ548, HXZ549, HXZ550, HXZ553, HXZ554, HXZ555, HXZ556, HXZ559, HXZ560, HXM561, HXM563, HXM564, HXM567, HXM569, HXM572, HXM573, HXM574, HXM587, HXM588, HXM596, HXM605, HXM611, HXM613, HXM616, HXM618, HXM621, HXM632, HXM639, HXM640 |
| PPR | TraesCS5D01G097400 | 1\|3 | HXM232, HXM239, HXM280, HXM635 |
| PPR | TraesCS6B01G293600 | 1\|6 | HXM441, HXM443 |
| PPR | TraesCS7A01G059000 | 1\|2 | HXM21, HXM112, HXZ163, HXZ167, HXM181, HXM193, HXM227, HXM232, HXM239, HXZ262, HXM280, HXM285, HXM306, HXM314, HXM320, HXM324, HXM327, HXM329, HXM336, HXM337, HXM338, HXM342, HXZ361, HXZ362, HXM372, HXM379, HXM385, HXM389, HXM390, HXM394, HXM395, HXM398, HXM402, HXM406, HXM407, HXM408, HXM409, HXM419, HXM424, HXM425, HXM426, HXM430, HXM431, HXM433, HXM436, HXM443, HXZ451, HXZ456, HXZ457, HXZ458, HXZ459, HXZ464, HXM467, HXM480, HXM483, HXM488, HXM489, HXM494, HXM527, HXM529, HXM530, HXZ546, HXZ549, HXZ555, HXZ557, HXZ559, HXM588, HXM602, HXM604, HXM620, HXM635 |
| SL | TraesCS1A01G320200 | 1\|11 | HXM21, HXM43 |
| SL | TraesCS1A01G334900 | 2\|5 | HXZ78, HXM93, HXM110, HXZ163, HXM205, HXM244, HXM256, HXZ259, HXM300, HXM317, HXM400, HXM402, HXM406, HXM407, HXM409, HXM411, HXM422, HXM424, HXM426, HXM428, HXM441, HXM442, HXM448, HXZ458, HXZ459, HXZ461, HXM517, HXZ551, HXM599 |
| SL | TraesCS1B01G133500 | 1\|10 | HXM56, HXM298 |
| SL | TraesCS2A01G009800 | 1\|2 | HXZ10, HXM17, HXM18, HXM22, HXM24, HXM25, HXM31, HXM32, HXM35, HXM37, HXM40, HXM41, HXM42, HXM43, HXM44, HXM46, HXM47, HXM48, |

| | | | |
|---|---|---|---|
| | | | HXM49, HXM50, HXM51, HXM53, HXM54, HXM55, HXM56, HXM57, HXM59, HXM60, HXZ74, HXM83, HXM93, HXM94, HXM95, HXM99, HXM100, HXM101, HXM102, HXM105, HXM106, HXM107, HXM109, HXM110, HXM114, HXM115, HXM134, HXM135, HXM137, HXM138, HXM145, HXM146, HXM149, HXM150, HXM151, HXM160, HXZ164, HXZ171, HXZ172, HXZ176, HXM183, HXM184, HXM196, HXM198, HXM204, HXM221, HXM228, HXM229, HXM233, HXM239, HXM245, HXM255, HXZ257, HXZ258, HXZ270, HXM276, HXM278, HXM281, HXM290, HXM291, HXM300, HXM310, HXM312, HXM323, HXM330, HXM332, HXM336, HXM339, HXM340, HXM352, HXZ353, HXZ366, HXM376, HXM383, HXM397, HXZ453, HXM498, HXM540, HXM609, HXM632, HXM635 |
| SL | TraesCS2A01G445500 | 1\|8 | HXZ269 |
| SL | TraesCS2A01G590600 | 1\|21 | HXM111, HXM298, HXM333 |
| SL | TraesCS3A01G019600 | 3\|12 | HXZ9, HXM40, HXM50, HXM228, HXM256, HXZ258, HXZ261, HXZ267, HXM300, HXZ460, HXM600 |
| SL | TraesCS3A01G481400 | 1\|13 | HXM538 |
| SL | TraesCS5A01G295800 | 1\|12 | HXM232 |
| SL | TraesCS5B01G236500 | 1\|16 | HXM227 |
| SL | TraesCS5B01G277500 | 1\|13 | HXM405 |
| SL | TraesCS7B01G478200 | 1\|31 | HXM232 |
| SL | TraesCS7B01G484200 | 1\|23 | HXM123 |
| SNS | TraesCS2B01G464900 | 1\|3 | HXZ15, HXZ16, HXM18, HXM61, HXZ78, HXM112, HXM177, HXM191, HXM192, HXM195, HXM207, HXM224, HXM227, HXM237, HXM249, HXM282, HXM294, HXM300, HXM303, HXM304, HXM316, HXM328, HXM334, HXM338, HXM600 |
| SNS | TraesCS3A01G022700 | 2\|34 | HXM239, HXM280, HXM635 |
| SNS | TraesCS3A01G391600 | 1\|8 | HXM239, HXM280, HXM635 |
| SNS | TraesCS3A01G502900 | 1\|7 | HXM31, HXZ161, HXM229, HXM252, HXM293, HXM333 |
| SNS | TraesCS3B01G428100 | 1\|6 | HXM34, HXM153, HXZ268, HXM370, HXM371 |
| SNS | TraesCS4A01G079300 | 2\|4 | HXZ3, HXZ9, HXZ13, HXZ16, HXM24, HXM39, HXM141, HXM148, HXM156, HXM157, HXM158, HXM159, HXM160, HXZ161, HXZ162, HXZ167, HXZ169, HXZ170, HXZ171, HXZ172, HXZ173, HXZ174, HXZ176, HXM177, HXM178, HXM179, HXM180, HXM181, HXM182, HXM183, HXM184, HXM185, HXM186, HXM187, HXM188, HXM189, HXM190, HXM191, HXM192, HXM193, HXM194, HXM195, HXM197, HXM198, HXM199, HXM220, HXM223, HXM224, HXM232, HXM240, HXM243, HXM249, HXM252, HXM253, HXM254, HXM256, HXZ259, HXZ263, HXZ267, HXZ270, HXZ271, HXM275, HXM277, HXM303, HXM304, HXM307, HXM309, HXM314, HXM319, HXM324, HXM325, HXM326, HXM329, HXM333, HXM334, HXM337, HXM340, HXM342, HXM350, HXM352, HXZ353, HXZ357, HXZ358, HXZ359, HXZ361, HXZ362, HXZ363, HXM369, HXM371, HXM374, HXM377, HXM430, HXM637, HXM638 |
| SNS | TraesCS5B01G469600 | 2\|5 | HXZ10, HXZ12, HXZ13, HXZ14, HXM45, HXM62, HXM63, HXM64, HXZ65, HXZ67, HXZ75, HXZ76, HXZ77, HXZ78, HXZ79, HXM82, HXM84, HXM85, HXM86, HXM87, HXM88, HXM95, HXM112, HXM113, HXM114, HXM119, HXM120, HXM126, HXM127, HXM129, HXM130, HXM131, HXM133, HXM139, |

| | | | |
|---|---|---|---|
| | | | HXM140, HXM143, HXM157, HXZ165, HXZ167, HXZ171, HXM177, HXM178, HXM180, HXM181, HXM182, HXM183, HXM193, HXM196, HXM207, HXM220, HXM224, HXM234, HXM275, HXM277, HXM284, HXM286, HXM291, HXM294, HXM295, HXM297, HXM299, HXM306, HXM307, HXM314, HXM317, HXM320, HXM324, HXM329, HXM333, HXM337, HXM342, HXM345, HXM349, HXM350, HXZ357, HXZ359, HXZ361, HXZ362, HXZ363, HXZ368, HXM373, HXM394, HXM402, HXM409, HXM416, HXM422, HXM433, HXM434, HXM439, HXM444, HXZ452, HXM469, HXM478, HXM484, HXM497, HXM500, HXM501, HXM507, HXM513, HXM514, HXM523, HXM543, HXZ552, HXZ558, HXM566, HXM589, HXM596, HXM606, HXM629 |
| SNS | TraesCS7A01G433000 | 1\|24 | HXM280 |
| T | TraesCS2A01G176000 | 1\|4 | HXM153, HXM227, HXM231, HXM421, HXM425, HXM431, HXM434, HXM435, HXM440, HXM441, HXM443, HXZ450, HXZ452, HXZ453, HXZ454, HXZ463 |
| T | TraesCS4A01G302000 | 1\|19 | HXM194 |
| T | TraesCS7D01G159800 | 1\|3 | HXM20, HXM21, HXM34, HXZ163, HXM226, HXM227, HXM229, HXM279, HXM311, HXM378, HXM379, HXM380, HXM382, HXM383, HXM385, HXM390, HXM391, HXM394, HXM395, HXM396, HXM398, HXM401, HXM402, HXM403, HXM405, HXM406, HXM407, HXM408, HXM409, HXM413, HXM415, HXM416, HXM418, HXM419, HXM420, HXM421, HXM422, HXM423, HXM424, HXM425, HXM426, HXM428, HXM429, HXM431, HXM432, HXM433, HXM434, HXM435, HXM436, HXM438, HXM439, HXM440, HXM442, HXM443, HXM444, HXM445, HXM446, HXZ450, HXZ453, HXZ456, HXZ458, HXZ459, HXZ460, HXZ461, HXZ462, HXM466, HXM467, HXM470, HXM471, HXM472, HXM475, HXM476, HXM477, HXM479, HXM480, HXM482, HXM486, HXM488, HXM489, HXM490, HXM491, HXM493, HXM494, HXM497, HXM499, HXM500, HXM501, HXM502, HXM503, HXM505, HXM508, HXM509, HXM510, HXM512, HXM516, HXM517, HXM520, HXM521, HXM523, HXM524, HXM531, HXM532, HXM533, HXM536, HXM539, HXM541, HXM542, HXZ545, HXZ546, HXZ550, HXZ553, HXZ554, HXZ555, HXZ557, HXZ559, HXZ560, HXM562, HXM563, HXM565, HXM569, HXM573, HXM575, HXM576, HXM577, HXM587, HXM590, HXM596, HXM598, HXM600, HXM602, HXM603, HXM605, HXM608, HXM612, HXM613, HXM614, HXM615, HXM618, HXM619, HXM620, HXM621, HXM629, HXM636, HXM637, HXM638, HXM639, HXM640 |
| TGW | TraesCS1A01G008000 | 8\|16 | HXM30, HXM379, HXM385, HXM387, HXM388, HXM389, HXM391, HXM396, HXM400, HXM411, HXM414, HXM420, HXM421, HXM422, HXM433, HXM435, HXM443, HXM446, HXM448, HXZ451, HXZ454, HXZ455, HXZ457, HXZ459, HXZ462, HXZ463, HXM465, HXM467, HXM468, HXM470, HXM473, HXM475, HXM476, HXM477, HXM484, HXM485, HXM486, HXM491, HXM493, HXM495, HXM497, HXM502, HXM508, HXM510, HXM513, HXM515, HXM522, HXM524, HXM532, HXM533, HXM534, HXM535, HXM536, HXM537, HXM538, HXM540, HXM541, HXM543, HXZ545, HXZ547, HXZ550, HXZ560, HXM561, HXM562, HXM565, HXM570, HXM586, HXM587, HXM592, HXM593, HXM594, HXM597, HXM599, HXM600, HXM601, HXM607, HXM608, HXM611, HXM613, HXM615, HXM617, HXM618, HXM619, HXM622, HXM629, HXM638, HXM640 |

| | | | |
|---|---|---|---|
| TGW | TraesCS1A01G010900 | 8\|15 | HXZ15, HXM41, HXM232, HXM239, HXM379, HXM386, HXM389, HXM391, HXM398, HXM400, HXM410, HXM411, HXM414, HXM420, HXM421, HXM422, HXM423, HXM425, HXM433, HXM437, HXM442, HXM443, HXM445, HXM446, HXM448, HXZ451, HXZ455, HXZ457, HXZ459, HXZ463, HXM465, HXM467, HXM470, HXM473, HXM477, HXM481, HXM484, HXM485, HXM486, HXM491, HXM495, HXM497, HXM508, HXM510, HXM513, HXM515, HXM525, HXM526, HXM527, HXM532, HXM533, HXM534, HXM537, HXM540, HXM541, HXM543, HXZ545, HXZ546, HXZ547, HXZ548, HXZ557, HXZ559, HXM561, HXM565, HXM570, HXM586, HXM592, HXM593, HXM594, HXM597, HXM599, HXM601, HXM602, HXM608, HXM611, HXM615, HXM617, HXM618, HXM620, HXM629, HXM635, HXM638, HXM640 |
| TGW | TraesCS1A01G050400 | 1\|15 | HXM617 |
| TGW | TraesCS1A01G058400 | 1\|39 | HXM521, HXZ560 |
| TGW | TraesCS1A01G063200 | 1\|4 | HXM371, HXZ458, HXM479, HXM489, HXM492, HXM499, HXM503, HXM505, HXM506, HXM508, HXM509, HXM523, HXM528, HXM531, HXM536, HXM542, HXZ555, HXZ558, HXM564, HXM587, HXM591, HXM612, HXM622 |
| TGW | TraesCS1A01G329900 | 2\|7 | HXM396, HXM433, HXM435, HXM443, HXZ454, HXM475, HXM483, HXM493, HXM502, HXM518, HXM567, HXM568, HXM569, HXM571, HXM572, HXM573, HXM575, HXM576, HXM577, HXM624, HXM625, HXM626, HXM627, HXM628 |
| TGW | TraesCS1A01G435000 | 1\|16 | HXM538 |
| TGW | TraesCS1B01G368500 | 1\|3 | HXZ163, HXM394, HXM406, HXM407, HXM409, HXM422, HXM423, HXM426, HXM431, HXM434, HXM444, HXZ459, HXZ463, HXM501, HXM507, HXM510, HXM514, HXM516, HXM589, HXM606 |
| TGW | TraesCS1B01G462000 | 6\|24 | HXM21, HXM380, HXM382, HXM391, HXM415, HXM440, HXM446, HXM447, HXZ450, HXZ452, HXM465, HXM467, HXM473, HXM496, HXM497, HXM498, HXM508, HXM510, HXM513, HXM517, HXM519, HXM522, HXM529, HXM534, HXM535, HXM540, HXZ549, HXZ550, HXZ551, HXZ555, HXZ560, HXM567, HXM568, HXM570, HXM571, HXM572, HXM593, HXM597, HXM606, HXM612, HXM614, HXM617, HXM622, HXM624, HXM625, HXM626, HXM627, HXM628, HXM629 |
| TGW | TraesCS1D01G009400 | 1\|2 | HXM232, HXM239, HXM400, HXM423, HXM442, HXM466, HXM467, HXM525, HXM526, HXM629, HXM635, HXM640 |
| TGW | TraesCS1D01G009900 | 2\|4 | HXM30, HXM98, HXM226, HXM379, HXM389, HXM411, HXM420, HXM422, HXM435, HXM437, HXM445, HXM446, HXM448, HXZ454, HXZ457, HXZ459, HXZ462, HXM465, HXM467, HXM468, HXM473, HXM476, HXM486, HXM495, HXM497, HXM508, HXM510, HXM513, HXM515, HXM522, HXM532, HXM533, HXM534, HXM535, HXM536, HXM537, HXM538, HXM540, HXM543, HXZ545, HXZ550, HXZ560, HXM570, HXM587, HXM592, HXM593, HXM594, HXM597, HXM599, HXM608, HXM611, HXM617, HXM618, HXM619, HXM622 |
| TGW | TraesCS1D01G063500 | 1\|7 | HXM232, HXM239, HXM280, HXM635 |
| TGW | TraesCS1D01G063900 | 1\|3 | HXM111, HXZ161, HXZ262, HXM280, HXM495, HXM521, HXM535 |
| TGW | TraesCS2A01G310300 | 1\|16 | HXM567 |
| TGW | TraesCS2A01G548100 | 3\|29 | HXM466, HXM495, HXZ560, HXM599, HXM613, HXM618 |
| TGW | TraesCS2A01G555600 | 2\|15 | HXM438, HXM535 |

| | | | |
|---|---|---|---|
| TGW | TraesCS2A01G555600 | 2\|15 | HXM438, HXM535 |
| TGW | TraesCS2B01G010800 | 1\|10 | HXM612 |
| TGW | TraesCS2B01G202300 | 1\|7 | HXM239, HXM280, HXM379, HXM392, HXM393, HXM395, HXM396, HXM398, HXM403, HXM413, HXM418, HXM420, HXM432, HXM438, HXM445, HXM447, HXZ457, HXM469, HXM482, HXM486, HXM491, HXM503, HXM511, HXM512, HXM526, HXM530, HXM544, HXZ545, HXZ546, HXZ548, HXZ550, HXZ551, HXZ553, HXZ555, HXZ558, HXZ560, HXM562, HXM564, HXM565, HXM566, HXM569, HXM570, HXM571, HXM573, HXM574, HXM586, HXM590, HXM595, HXM596, HXM597, HXM598, HXM600, HXM603, HXM604, HXM607, HXM608, HXM609, HXM613, HXM615, HXM616, HXM617, HXM618, HXM619, HXM622, HXM623, HXM635, HXM640 |
| TGW | TraesCS2B01G202300 | 1\|7 | HXM239, HXM280, HXM379, HXM392, HXM393, HXM395, HXM396, HXM398, HXM403, HXM413, HXM418, HXM420, HXM432, HXM438, HXM445, HXM447, HXZ457, HXM469, HXM482, HXM486, HXM491, HXM503, HXM511, HXM512, HXM526, HXM530, HXM544, HXZ545, HXZ546, HXZ548, HXZ550, HXZ551, HXZ553, HXZ555, HXZ558, HXZ560, HXM562, HXM564, HXM565, HXM566, HXM569, HXM570, HXM571, HXM573, HXM574, HXM586, HXM590, HXM595, HXM596, HXM597, HXM598, HXM600, HXM603, HXM604, HXM607, HXM608, HXM609, HXM613, HXM615, HXM616, HXM617, HXM618, HXM619, HXM622, HXM623, HXM635, HXM640 |
| TGW | TraesCS2B01G533300 | 1\|11 | HXM531 |
| TGW | TraesCS2B01G578500 | 1\|3 | HXM38, HXM45, HXM61, HXM62, HXM96, HXM99, HXM111, HXM226, HXM231, HXM233, HXM255, HXZ258, HXM329, HXM335, HXM385, HXM389, HXM398, HXM408, HXM421, HXM435, HXZ451, HXZ454, HXZ462, HXM478, HXM481, HXM485, HXM495, HXM499, HXM500, HXM502, HXM523, HXM527, HXM531, HXM533, HXM544, HXZ548, HXZ549, HXZ560, HXM563, HXM574, HXM587, HXM596, HXM599, HXM613, HXM618, HXM621, HXM632, HXM639 |
| TGW | TraesCS2B01G587900 | 1\|8 | HXM535 |
| TGW | TraesCS2D01G548900 | 1\|14 | HXM438 |
| TGW | TraesCS4A01G444300 | 5\|37 | HXM239, HXM486, HXM518, HXM541, HXM592, HXM608, HXM612, HXM617 |
| TGW | TraesCS4B01G029700 | 1\|3 | HXM21, HXZ70, HXZ75, HXM104, HXM117, HXM118, HXM124, HXM128, HXZ163, HXM226, HXM244, HXZ260, HXZ264, HXZ267, HXM311, HXM335, HXM379, HXM383, HXM384, HXM385, HXM391, HXM392, HXM393, HXM394, HXM395, HXM396, HXM397, HXM399, HXM401, HXM403, HXM404, HXM405, HXM406, HXM412, HXM413, HXM414, HXM415, HXM417, HXM419, HXM420, HXM421, HXM423, HXM425, HXM426, HXM427, HXM428, HXM431, HXM432, HXM433, HXM434, HXM435, HXM436, HXM437, HXM439, HXM440, HXM442, HXM443, HXM447, HXM448, HXZ449, HXZ451, HXZ452, HXZ453, HXZ454, HXZ455, HXZ456, HXZ457, HXZ458, HXZ459, HXZ460, HXZ464, HXM465, HXM466, HXM468, HXM470, HXM471, HXM473, HXM474, HXM475, HXM476, HXM477, HXM479, HXM480, HXM481, HXM483, HXM484, HXM486, HXM487, HXM488, HXM489, HXM491, HXM493, HXM494, HXM498, HXM499, HXM502, HXM504, HXM505, HXM511, HXM512, HXM513, HXM515, HXM519, HXM522, HXM524, HXM525, HXM526, HXM527, HXM530, HXM533, HXM539, HXM542, |

| | | | |
|---|---|---|---|
| | | | HXZ545, HXZ546, HXZ547, HXZ548, HXZ550, HXZ551, HXZ555, HXZ557, HXZ560, HXM561, HXM562, HXM566, HXM567, HXM568, HXM571, HXM573, HXM587, HXM588, HXM590, HXM599, HXM601, HXM604, HXM607, HXM608, HXM609, HXM613, HXM616, HXM618, HXM619, HXM620, HXM624, HXM625, HXM626, HXM627, HXM628, HXM636, HXM639, HXM640 |
| TGW | TraesCS4D01G274200 | 1\|3 | HXM232, HXM239, HXM280, HXM635 |
| TGW | TraesCS4D01G350500 | 1\|3 | HXM232, HXM239, HXM280, HXM635 |
| TGW | TraesCS4D01G353800 | 1\|2 | HXM232, HXM239, HXM280, HXZ559, HXM635 |
| TGW | TraesCS5B01G039800 | 3\|8 | HXZ267, HXZ268, HXM338, HXM370, HXM371, HXM447, HXM448, HXZ452, HXM465, HXM468, HXM473, HXM483, HXM491, HXM511, HXM513, HXM515, HXM522, HXM525, HXM526, HXM534, HXM541, HXZ551, HXM562, HXM586, HXM605, HXM607, HXM615, HXM619 |
| TGW | TraesCS6A01G382600 | 1\|3 | HXZ1, HXZ3, HXM33, HXZ76, HXM117, HXM118, HXM126, HXM127, HXM128, HXM133, HXM139, HXZ163, HXZ167, HXZ169, HXZ172, HXM196, HXM231, HXM232, HXM237, HXM239, HXM254, HXZ261, HXZ265, HXZ268, HXM280, HXM282, HXM296, HXM304, HXM337, HXM344, HXM346, HXM370, HXM371, HXM378, HXM379, HXM382, HXM385, HXM386, HXM387, HXM388, HXM389, HXM390, HXM393, HXM395, HXM396, HXM398, HXM400, HXM401, HXM402, HXM405, HXM407, HXM408, HXM409, HXM413, HXM418, HXM419, HXM420, HXM423, HXM424, HXM425, HXM427, HXM436, HXM439, HXM442, HXZ449, HXZ450, HXZ451, HXZ453, HXZ456, HXM466, HXM467, HXM468, HXM469, HXM471, HXM476, HXM477, HXM483, HXM486, HXM488, HXM490, HXM493, HXM494, HXM495, HXM496, HXM497, HXM498, HXM506, HXM507, HXM508, HXM510, HXM511, HXM512, HXM514, HXM517, HXM518, HXM520, HXM526, HXM527, HXM532, HXM536, HXM537, HXM540, HXM544, HXZ551, HXZ557, HXZ559, HXM589, HXM590, HXM591, HXM592, HXM594, HXM602, HXM603, HXM609, HXM610, HXM611, HXM617, HXM618, HXM619, HXM620, HXM624, HXM625, HXM626, HXM627, HXM628 |
| TGW | TraesCS6A01G392400 | 7\|16 | HXM30, HXM232, HXM239, HXM280, HXM300, HXM388, HXM399, HXM402, HXM416, HXM418, HXM423, HXM427, HXM431, HXM433, HXM436, HXM442, HXM445, HXM447, HXM448, HXZ449, HXZ452, HXZ455, HXZ456, HXZ457, HXZ458, HXZ460, HXZ463, HXM465, HXM466, HXM468, HXM469, HXM470, HXM472, HXM473, HXM474, HXM477, HXM479, HXM480, HXM482, HXM484, HXM485, HXM486, HXM487, HXM488, HXM491, HXM492, HXM494, HXM497, HXM500, HXM502, HXM503, HXM504, HXM505, HXM507, HXM508, HXM509, HXM511, HXM512, HXM513, HXM514, HXM515, HXM516, HXM518, HXM522, HXM523, HXM524, HXM525, HXM526, HXM527, HXM533, HXM534, HXM541, HXM543, HXZ547, HXZ553, HXZ558, HXM562, HXM563, HXM564, HXM565, HXM566, HXM568, HXM574, HXM586, HXM587, HXM590, HXM593, HXM595, HXM596, HXM599, HXM601, HXM605, HXM606, HXM608, HXM609, HXM615, HXM616, HXM621, HXM622, HXM624, HXM625, HXM626, HXM627, HXM628, HXM635, HXM640 |
| TGW | TraesCS6B01G015100 | 1\|45 | HXM393 |

| | | | |
|---|---|---|---|
| TGW | TraesCS6B01G253900 | 2\|6 | HXM20, HXZ75, HXM311, HXM344, HXM378, HXM380, HXM381, HXM382, HXM385, HXM386, HXM387, HXM389, HXM393, HXM394, HXM403, HXM405, HXM409, HXM410, HXM413, HXM414, HXM415, HXM419, HXM420, HXM421, HXM422, HXM423, HXM428, HXM429, HXM430, HXM434, HXM436, HXM439, HXM443, HXM444, HXZ450, HXZ459, HXZ460, HXZ463, HXM470, HXM474, HXM477, HXM480, HXM485, HXM486, HXM490, HXM495, HXM497, HXM498, HXM500, HXM501, HXM523, HXM531, HXM532, HXM535, HXM536, HXM537, HXM538, HXM540, HXM543, HXM544, HXZ551, HXM563, HXM570, HXM574, HXM592, HXM593, HXM595, HXM609, HXM611, HXM617, HXM621 |
| TGW | TraesCS6D01G009700 | 1\|4 | HXM239, HXM635 |
| TGW | TraesCS6D01G168200 | 2\|4 | HXM232, HXM239, HXM280, HXZ455, HXM506, HXM516, HXM518, HXM520, HXM528, HXM530, HXZ546, HXM588, HXM604, HXM635 |
| TGW | TraesCS6D01G176900 | 1\|3 | HXM232, HXM239, HXM280, HXM635 |
| TGW | TraesCS6D01G207100 | 1\|3 | HXZ76, HXM232, HXM239, HXM280, HXZ557, HXM635 |
| TGW | TraesCS7A01G043700 | 1\|37 | HXM521 |
| TGW | TraesCS7A01G423100 | 5\|14 | HXM20, HXM226, HXM229, HXM279, HXM280, HXM311, HXM325, HXM343, HXM344, HXM378, HXM380, HXM386, HXM393, HXM394, HXM401, HXM402, HXM403, HXM407, HXM410, HXM413, HXM414, HXM415, HXM416, HXM419, HXM420, HXM421, HXM428, HXM431, HXM437, HXM442, HXM443, HXM444, HXZ458, HXZ459, HXZ460, HXZ461, HXZ462, HXM466, HXM470, HXM471, HXM472, HXM477, HXM479, HXM480, HXM483, HXM486, HXM487, HXM488, HXM489, HXM491, HXM494, HXM496, HXM497, HXM500, HXM501, HXM504, HXM507, HXM510, HXM514, HXM530, HXM532, HXM533, HXM535, HXM537, HXM538, HXM539, HXM541, HXZ551, HXZ553, HXZ555, HXZ556, HXZ559, HXM562, HXM566, HXM567, HXM570, HXM572, HXM574, HXM586, HXM588, HXM589, HXM592, HXM593, HXM595, HXM597, HXM606, HXM608, HXM609, HXM612, HXM615, HXM617, HXM635, HXM638, HXM640 |
| TGW | TraesCS7B01G455000 | 1\|3 | HXZ3, HXM17, HXM30, HXM104, HXM226, HXM230, HXZ267, HXM311, HXM379, HXM384, HXM388, HXM391, HXM392, HXM393, HXM394, HXM395, HXM397, HXM402, HXM403, HXM404, HXM413, HXM415, HXM417, HXM418, HXM420, HXM421, HXM427, HXM428, HXM432, HXM437, HXM440, HXM447, HXZ452, HXZ457, HXZ460, HXZ463, HXZ464, HXM465, HXM468, HXM469, HXM473, HXM474, HXM479, HXM480, HXM487, HXM492, HXM500, HXM504, HXM506, HXM510, HXM512, HXM513, HXM515, HXM522, HXM523, HXM525, HXM526, HXM528, HXM530, HXM531, HXM533, HXM539, HXM543, HXM544, HXZ546, HXZ550, HXZ558, HXM561, HXM588, HXM591, HXM605, HXM610, HXM622 |
| TGW | TraesCS7D01G416000 | 1\|2 | HXM232, HXM239, HXM280, HXM635 |
| V | TraesCS2B01G202300 | 3\|7 | HXM21, HXM34, HXM104, HXM226, HXM227, HXM232, HXM239, HXM280, HXM379, HXM384, HXM391, HXM392, HXM393, HXM395, HXM396, HXM398, HXM400, HXM403, HXM404, HXM411, HXM413, HXM417, HXM418, HXM420, HXM427, HXM432, HXM435, HXM438, HXM440, HXM442, HXM445, HXM446, HXM447, HXM448, HXZ449, HXZ452, HXZ457, HXZ458, HXZ462, HXM465, HXM467, HXM468, HXM469, HXM470, HXM471, HXM472, HXM473, HXM474, |

| | | | |
|---|---|---|---|
| | | | HXM476, HXM477, HXM479, HXM480, HXM481, HXM482, HXM486, HXM489, HXM491, HXM492, HXM493, HXM495, HXM498, HXM500, HXM503, HXM505, HXM508, HXM509, HXM511, HXM512, HXM513, HXM515, HXM521, HXM522, HXM523, HXM525, HXM526, HXM530, HXM531, HXM535, HXM540, HXM541, HXM543, HXM544, HXZ545, HXZ546, HXZ548, HXZ550, HXZ551, HXZ553, HXZ555, HXZ558, HXZ560, HXM562, HXM564, HXM565, HXM566, HXM569, HXM570, HXM571, HXM573, HXM574, HXM575, HXM576, HXM577, HXM586, HXM588, HXM590, HXM595, HXM596, HXM597, HXM598, HXM600, HXM603, HXM604, HXM607, HXM608, HXM609, HXM613, HXM615, HXM616, HXM617, HXM618, HXM619, HXM622, HXM623, HXM635, HXM640 |
| V | TraesCS3A01G015500 | 1\|107 | HXZ556 |
| V | TraesCS3B01G469000 | 1\|3 | HXM20, HXM21, HXM227, HXM335, HXM344, HXM378, HXM382, HXM393, HXM394, HXM399, HXM400, HXM401, HXM415, HXM431, HXM433, HXM439, HXM442, HXM443, HXM445, HXZ452, HXZ456, HXZ457, HXZ463, HXM469, HXM470, HXM475, HXM477, HXM483, HXM484, HXM485, HXM486, HXM489, HXM491, HXM494, HXM495, HXM496, HXM500, HXM502, HXM521, HXM523, HXM524, HXM536, HXM539, HXM543, HXM544, HXZ552, HXZ557, HXZ558, HXZ560, HXM562, HXM563, HXM567, HXM574, HXM587, HXM593, HXM595, HXM596, HXM603, HXM607, HXM609, HXM610, HXM613, HXM618, HXM621, HXM638 |
| V | TraesCS3B01G560400 | 2\|4 | HXM38, HXM125, HXM154, HXM182, HXM209, HXM220, HXM224, HXM232, HXM239, HXM280, HXM292, HXM296, HXM307, HXM313, HXM348, HXM349, HXM351, HXZ354, HXZ356, HXZ363, HXM429, HXM436, HXM447, HXM448, HXZ450, HXZ452, HXZ455, HXZ462, HXM465, HXM468, HXM473, HXM478, HXM484, HXM485, HXM490, HXM495, HXM496, HXM498, HXM499, HXM500, HXM510, HXM511, HXM513, HXM515, HXM518, HXM519, HXM522, HXM523, HXM529, HXM533, HXM534, HXM535, HXM537, HXM538, HXM540, HXM541, HXM544, HXZ549, HXZ550, HXZ551, HXZ556, HXZ557, HXZ558, HXZ560, HXM564, HXM565, HXM567, HXM570, HXM572, HXM574, HXM591, HXM592, HXM595, HXM596, HXM597, HXM602, HXM603, HXM604, HXM609, HXM611, HXM612, HXM613, HXM614, HXM615, HXM617, HXM618, HXM620, HXM623, HXM634, HXM635 |
| V | TraesCS3B01G561000 | 2\|4 | HXM38, HXM125, HXM154, HXM182, HXM209, HXM220, HXM224, HXM232, HXM239, HXM280, HXM292, HXM296, HXM300, HXM307, HXM313, HXM348, HXM349, HXM351, HXZ354, HXZ356, HXZ363, HXM429, HXM436, HXM447, HXM448, HXZ450, HXZ452, HXZ455, HXZ462, HXM465, HXM468, HXM473, HXM478, HXM481, HXM484, HXM485, HXM490, HXM495, HXM496, HXM498, HXM499, HXM500, HXM510, HXM511, HXM513, HXM515, HXM518, HXM519, HXM522, HXM523, HXM529, HXM533, HXM534, HXM535, HXM537, HXM538, HXM540, HXM541, HXM544, HXZ549, HXZ550, HXZ551, HXZ556, HXZ557, HXZ558, HXZ560, HXM564, HXM565, HXM567, HXM570, HXM572, HXM574, HXM591, HXM592, HXM595, HXM596, HXM597, HXM599, HXM602, HXM603, HXM604, HXM609, HXM611, HXM612, HXM613, HXM614, HXM615, HXM617, HXM618, HXM620, HXM623, HXM634, HXM635 |

| | | | |
|---|---|---|---|
| V | TraesCS5B01G000100 | 7\|15 | HXM34, HXZ75, HXZ163, HXM226, HXZ267, HXZ268, HXM335, HXM370, HXM371, HXM379, HXM385, HXM388, HXM390, HXM391, HXM394, HXM395, HXM405, HXM406, HXM408, HXM409, HXM415, HXM418, HXM423, HXM424, HXM427, HXM433, HXM434, HXM436, HXM437, HXM439, HXM440, HXM441, HXM445, HXM446, HXM447, HXM448, HXZ449, HXZ452, HXZ453, HXZ454, HXZ456, HXZ459, HXZ462, HXM465, HXM467, HXM468, HXM469, HXM471, HXM472, HXM473, HXM478, HXM482, HXM483, HXM484, HXM487, HXM491, HXM492, HXM494, HXM497, HXM498, HXM500, HXM504, HXM505, HXM506, HXM508, HXM510, HXM511, HXM513, HXM514, HXM515, HXM519, HXM522, HXM523, HXM525, HXM526, HXM527, HXM528, HXM530, HXM531, HXM533, HXM534, HXM536, HXM541, HXM544, HXZ548, HXZ550, HXZ551, HXZ553, HXZ555, HXZ556, HXM562, HXM564, HXM565, HXM566, HXM567, HXM569, HXM571, HXM573, HXM575, HXM576, HXM577, HXM588, HXM590, HXM595, HXM600, HXM604, HXM605, HXM607, HXM609, HXM615, HXM616, HXM619, HXM622, HXM624, HXM625, HXM626, HXM627, HXM628, HXM634, HXM636, HXM639 |
| V | TraesCS6D01G403800 | 5\|13 | HXM20, HXM21, HXM30, HXZ75, HXM183, HXM279, HXM313, HXM325, HXM344, HXM378, HXM380, HXM381, HXM385, HXM387, HXM389, HXM393, HXM394, HXM396, HXM403, HXM404, HXM409, HXM411, HXM412, HXM413, HXM415, HXM416, HXM417, HXM428, HXM433, HXM434, HXM435, HXM436, HXM442, HXM443, HXM444, HXM445, HXM447, HXM448, HXZ449, HXZ452, HXZ455, HXZ456, HXZ457, HXZ458, HXZ460, HXZ464, HXM466, HXM468, HXM470, HXM472, HXM473, HXM474, HXM475, HXM476, HXM477, HXM478, HXM479, HXM480, HXM482, HXM483, HXM484, HXM485, HXM486, HXM488, HXM489, HXM491, HXM493, HXM497, HXM498, HXM500, HXM501, HXM504, HXM505, HXM507, HXM510, HXM511, HXM512, HXM514, HXM515, HXM523, HXM532, HXM537, HXM538, HXM539, HXM541, HXM543, HXZ547, HXZ551, HXZ552, HXZ553, HXM562, HXM563, HXM564, HXM565, HXM566, HXM567, HXM568, HXM569, HXM572, HXM574, HXM575, HXM576, HXM577, HXM586, HXM589, HXM591, HXM599, HXM601, HXM606, HXM607, HXM608, HXM609, HXM610, HXM611, HXM613, HXM615, HXM619, HXM621, HXM623, HXM624, HXM625, HXM626, HXM627, HXM628, HXM636, HXM640 |
| V | TraesCS7A01G040900 | 1\|59 | HXM521 |
| V | TraesCS7A01G518200 | 1\|92 | HXM473 |
| V | TraesCS7A01G518300 | 1\|6 | HXM423, HXZ450, HXM535 |
| V | TraesCS7A01G549200 | 1\|18 | HXM465 |
| V | TraesCS7A01G549300 | 1\|13 | HXM279, HXM405, HXZ462, HXM470, HXM477, HXM484, HXM485, HXM486, HXM491, HXM497, HXM534, HXZ554, HXM562, HXM565, HXM574, HXM586, HXM587, HXM595, HXM596, HXM613, HXM634 |
| V | TraesCS7B01G034600 | 2\|10 | HXM232, HXM483, HXM495, HXM496, HXM520, HXM521, HXM529, HXM535, HXM536, HXM539, HXZ550, HXZ560, HXM565, HXM567, HXM570, HXM572, HXM574, HXM593, HXM600, HXM606, HXM612, HXM617, HXM623, HXM624, HXM625, HXM626, HXM627, HXM628 |
| V | TraesCS7B01G439900 | 1\|30 | HXM408, HXM600 |

| | | | |
|---|---|---|---|
| V | TraesCS7B01G472500 | 1\|13 | HXM445 |
| Vmax | TraesCS1A01G033600 | 1\|5 | HXM401, HXM433, HXM443, HXM446, HXZ461, HXM467, HXM470, HXM477, HXM484, HXM485, HXM486, HXM491, HXM495, HXM508, HXM517, HXM535, HXM537, HXM538, HXM540, HXM541, HXM543, HXM544, HXZ550, HXZ560, HXM561, HXM570, HXM586, HXM592, HXM593, HXM594, HXM597, HXM607, HXM613, HXM615, HXM617, HXM618, HXM629 |
| Vmax | TraesCS1B01G422700 | 1\|9 | HXM448, HXM597 |
| Vmax | TraesCS2B01G202300 | 3\|7 | HXM21, HXM34, HXM104, HXM226, HXM227, HXM232, HXM239, HXM280, HXM379, HXM384, HXM391, HXM392, HXM393, HXM395, HXM396, HXM398, HXM400, HXM403, HXM404, HXM411, HXM413, HXM417, HXM418, HXM420, HXM427, HXM432, HXM435, HXM438, HXM440, HXM442, HXM445, HXM446, HXM447, HXM448, HXZ449, HXZ452, HXZ457, HXZ458, HXZ462, HXM465, HXM467, HXM468, HXM469, HXM470, HXM471, HXM472, HXM473, HXM474, HXM476, HXM477, HXM479, HXM480, HXM481, HXM482, HXM486, HXM489, HXM491, HXM492, HXM493, HXM495, HXM498, HXM500, HXM503, HXM505, HXM508, HXM509, HXM511, HXM512, HXM513, HXM515, HXM521, HXM522, HXM523, HXM525, HXM526, HXM530, HXM531, HXM535, HXM540, HXM541, HXM543, HXM544, HXZ545, HXZ546, HXZ548, HXZ550, HXZ551, HXZ553, HXZ555, HXZ558, HXZ560, HXM562, HXM564, HXM565, HXM566, HXM569, HXM570, HXM571, HXM573, HXM574, HXM575, HXM576, HXM577, HXM586, HXM588, HXM590, HXM595, HXM596, HXM597, HXM598, HXM600, HXM603, HXM604, HXM607, HXM608, HXM609, HXM613, HXM615, HXM616, HXM617, HXM618, HXM619, HXM622, HXM623, HXM635, HXM640 |
| Vmax | TraesCS3B01G469000 | 1\|3 | HXM20, HXM21, HXM227, HXM335, HXM344, HXM378, HXM382, HXM393, HXM394, HXM399, HXM400, HXM401, HXM415, HXM431, HXM433, HXM439, HXM442, HXM443, HXM445, HXZ452, HXZ456, HXZ457, HXZ463, HXM469, HXM470, HXM475, HXM477, HXM483, HXM484, HXM485, HXM486, HXM489, HXM491, HXM494, HXM495, HXM496, HXM500, HXM502, HXM521, HXM523, HXM524, HXM536, HXM539, HXM543, HXM544, HXZ552, HXZ557, HXZ558, HXZ560, HXM562, HXM563, HXM567, HXM574, HXM587, HXM593, HXM595, HXM596, HXM603, HXM607, HXM609, HXM610, HXM613, HXM618, HXM621, HXM638 |
| Vmax | TraesCS5B01G039800 | 1\|8 | HXM465 |
| Vmax | TraesCS6B01G369700 | 1\|12 | HXM227, HXM390, HXM394, HXM395, HXM399, HXM407, HXM408, HXM409, HXM419, HXM429, HXM433, HXM436, HXM437, HXM439, HXM441, HXM444, HXZ450, HXZ454, HXZ457, HXZ463, HXM469, HXM471, HXM474, HXM482, HXM484, HXM487, HXM490, HXM501, HXZ554, HXZ556, HXM563, HXM611, HXM621 |
| Vmax | TraesCS6D01G403800 | 4\|13 | HXM20, HXM21, HXZ75, HXM183, HXM279, HXM313, HXM325, HXM344, HXM378, HXM380, HXM381, HXM387, HXM393, HXM394, HXM403, HXM409, HXM411, HXM412, HXM413, HXM415, HXM416, HXM428, HXM433, HXM434, HXM436, HXM442, HXM443, HXM444, HXM445, HXM447, HXM448, HXZ449, HXZ452, HXZ455, HXZ456, HXZ457, HXZ458, HXZ460, HXZ464, HXM466, HXM468, HXM470, HXM472, HXM473, HXM475, HXM476, HXM477, HXM478, |

| | | | |
|---|---|---|---|
| | | | HXM479, HXM480, HXM482, HXM485, HXM486, HXM488, HXM489, HXM491, HXM493, HXM497, HXM498, HXM500, HXM501, HXM504, HXM505, HXM507, HXM510, HXM511, HXM512, HXM514, HXM515, HXM523, HXM532, HXM537, HXM538, HXM539, HXM541, HXM543, HXZ547, HXZ551, HXZ552, HXZ553, HXM562, HXM563, HXM564, HXM565, HXM566, HXM567, HXM568, HXM572, HXM574, HXM586, HXM589, HXM591, HXM599, HXM601, HXM606, HXM607, HXM608, HXM609, HXM610, HXM611, HXM613, HXM615, HXM619, HXM621, HXM623, HXM624, HXM625, HXM626, HXM627, HXM628, HXM640 |
| Vmax | TraesCS7A01G549100 | 1\|15 | HXM312, HXM571 |
| Vmax | TraesCS7A01G549200 | 1\|18 | HXM465 |
| Vmax | TraesCS7B01G034600 | 2\|10 | HXM232, HXM483, HXM495, HXM496, HXM520, HXM521, HXM529, HXM535, HXM536, HXM539, HXZ550, HXZ560, HXM565, HXM567, HXM570, HXM572, HXM574, HXM593, HXM600, HXM606, HXM612, HXM617, HXM623, HXM624, HXM625, HXM626, HXM627, HXM628 |
| Vmax | TraesCS7B01G472500 | 1\|13 | HXM445 |
| Yr | TraesCS1A01G061500 | 3\|13 | HXM151, HXM232, HXM239, HXM280, HXZ460, HXM534, HXM635 |
| Yr | TraesCS2B01G182800 | 1\|100 | HXM82 |
| Yr | TraesCS2B01G488000 | 1\|38 | HXZ555 |
| Yr | TraesCS4A01G066200 | 2\|6 | HXZ3, HXZ9, HXZ13, HXZ16, HXM24, HXM39, HXM141, HXM148, HXM156, HXM157, HXM158, HXM159, HXM160, HXZ161, HXZ162, HXZ167, HXZ169, HXZ170, HXZ171, HXZ172, HXZ173, HXZ174, HXZ176, HXM177, HXM178, HXM179, HXM180, HXM181, HXM182, HXM183, HXM184, HXM185, HXM186, HXM187, HXM188, HXM189, HXM190, HXM191, HXM192, HXM193, HXM194, HXM195, HXM197, HXM198, HXM199, HXM220, HXM223, HXM224, HXM240, HXM243, HXM249, HXM252, HXM253, HXM254, HXM256, HXZ259, HXZ263, HXZ267, HXZ270, HXZ271, HXM275, HXM277, HXM303, HXM304, HXM307, HXM309, HXM314, HXM319, HXM324, HXM325, HXM329, HXM333, HXM334, HXM337, HXM340, HXM342, HXM350, HXM352, HXZ353, HXZ357, HXZ358, HXZ359, HXZ361, HXZ362, HXZ363, HXM369, HXM371, HXM374, HXM377, HXM637, HXM638 |
| Yr | TraesCS4D01G243100 | 1\|2 | HXM232, HXM239, HXM280, HXM635 |
| Yr | TraesCS5A01G344100 | 9\|38 | HXM21, HXM239, HXZ360, HXM426, HXM486, HXM493, HXM506, HXM544, HXZ557, HXM634, HXM635 |
| Yr | TraesCS5D01G349800 | 1\|11 | HXM351, HXM428, HXM430, HXM445, HXZ554 |
| Yr | TraesCS6B01G167900 | 2\|3 | HXZ1, HXZ2, HXZ4, HXZ5, HXZ7, HXZ8, HXZ10, HXZ12, HXZ13, HXZ14, HXM17, HXM18, HXM22, HXM23, HXM25, HXM37, HXM38, HXM40, HXZ70, HXM120, HXM132, HXM155, HXM156, HXM158, HXM159, HXM160, HXZ162, HXZ164, HXZ165, HXZ166, HXZ167, HXM186, HXM236, HXM240, HXM244, HXM248, HXM254, HXZ260, HXZ264, HXZ265, HXZ267, HXM275, HXM286, HXM295, HXM341, HXM342 |
| Yr | TraesCS6B01G185700 | 1\|3 | HXM232, HXM239, HXM280, HXM635 |
| Yr | TraesCS7D01G080300 | 1\|5 | HXM232, HXM239, HXM280, HXM635 |

Superior haplotype: The number of superior haplotypes in all the haplotypes.

PH: plant height, SL: spike length, PPR: ratio of peduncle length to plant height, SNS: spikelet number per spike, TGW: thousand grain weight, GL: grain length, GW: grain width, Vmax: maximum filling rate, V: average grain filling rate, T: active filling period, HD: heading date, FD: flowering date, Pm: powdery mildew, Sr: stripe rust.

**Supplementary Table S35.** Four landraces with the most superior haplotypes for future breeding

| Trait type | Trait | Gene | HXM239 | HXM280 | HXM232 | HXM635 |
|---|---|---|---|---|---|---|
| Morphological traits | PH | TraesCS1B01G462300 | 1 | | 1 | 1 |
| | | TraesCS2A01G453300 | 1 | | | 1 |
| | | TraesCS2A01G552400 | 1 | | | 1 |
| | | TraesCS7A01G424700 | 1 | | 1 | 1 |
| | SL | TraesCS2A01G009800 | 1 | | | 1 |
| | | TraesCS5A01G295800 | | | 1 | |
| | | TraesCS7B01G478200 | | | 1 | |
| | PPR | TraesCS2A01G552400 | 1 | | | 1 |
| | | TraesCS2B01G010100 | 1 | | 1 | 1 |
| | | TraesCS5D01G097400 | 1 | 1 | 1 | 1 |
| | | TraesCS7A01G059000 | 1 | 1 | 1 | 1 |
| Yield-related traits | GL | TraesCS1A01G273900 | 1 | 1 | 1 | 1 |
| | | TraesCS2D01G011200 | 1 | 1 | | 1 |
| | | TraesCS2D01G016900 | 1 | | 1 | 1 |
| | | TraesCS2D01G082000 | 1 | 1 | | 1 |
| | | TraesCS2D01G482500 | | | 1 | |
| | | TraesCS3B01G560400 | 1 | 1 | 1 | 1 |
| | | TraesCS3B01G561000 | 1 | 1 | 1 | 1 |
| | | TraesCS4A01G266900 | 1 | | | 1 |
| | | TraesCS4A01G444300 | 1 | | | 1 |
| | | TraesCS4B01G321800 | 1 | | 1 | 1 |
| | | TraesCS4B01G360800 | | | 1 | |
| | | TraesCS4D01G318400 | 1 | 1 | 1 | 1 |
| | | TraesCS4D01G325200 | 1 | 1 | 1 | 1 |
| | | TraesCS5A01G377700 | 1 | | | 1 |
| | | TraesCS5B01G307600 | 1 | 1 | 1 | 1 |
| | | TraesCS5B01G375800 | 1 | 1 | 1 | 1 |
| | | TraesCS6A01G009100 | 1 | | | 1 |
| | | TraesCS6A01G382600 | 1 | 1 | 1 | |
| | | TraesCS6D01G378300 | | | 1 | 1 |
| | | TraesCS7B01G018600 | 1 | 1 | 1 | 1 |
| | | TraesCS7D01G039500 | 1 | | | 1 |
| | TGW | TraesCS1A01G010900 | 1 | | 1 | 1 |
| | | TraesCS1D01G009400 | 1 | | 1 | 1 |
| | | TraesCS1D01G063500 | 1 | 1 | 1 | 1 |
| | | TraesCS1D01G063900 | | 1 | | |
| | | TraesCS2B01G202300 | 1 | 1 | | 1 |
| | | TraesCS2B01G202300 | 1 | 1 | | 1 |
| | | TraesCS4A01G444300 | 1 | | | |
| | | TraesCS4D01G274200 | 1 | 1 | 1 | 1 |

| | | | | | | |
|---|---|---|---|---|---|---|
| | | TraesCS4D01G350500 | 1 | 1 | 1 | 1 |
| | | TraesCS4D01G353800 | 1 | 1 | 1 | 1 |
| | | TraesCS6A01G382600 | 1 | 1 | 1 | |
| | | TraesCS6A01G392400 | 1 | 1 | 1 | 1 |
| | | TraesCS6D01G009700 | 1 | | | 1 |
| | | TraesCS6D01G168200 | 1 | 1 | 1 | 1 |
| | | TraesCS6D01G176900 | 1 | 1 | 1 | 1 |
| | | TraesCS6D01G207100 | 1 | 1 | 1 | 1 |
| | | TraesCS7A01G423100 | | 1 | | 1 |
| | | TraesCS7D01G416000 | 1 | 1 | 1 | 1 |
| | SNS | TraesCS3A01G022700 | 1 | 1 | | 1 |
| | | TraesCS3A01G391600 | 1 | 1 | | 1 |
| | | TraesCS4A01G079300 | | | 1 | |
| | | TraesCS7A01G433000 | | 1 | | |
| Grain filling related traits | V | TraesCS2B01G202300 | 1 | 1 | 1 | 1 |
| | | TraesCS3B01G560400 | 1 | 1 | 1 | 1 |
| | | TraesCS3B01G561000 | 1 | 1 | 1 | 1 |
| | | TraesCS7B01G034600 | | | 1 | |
| | Vmax | TraesCS2B01G202300 | 1 | 1 | 1 | 1 |
| | | TraesCS7B01G034600 | | | 1 | |
| Growth period traits | FD | TraesCS1A01G058400 | 1 | 1 | | 1 |
| | | TraesCS1A01G372300 | | 1 | | |
| | | TraesCS2B01G200800 | 1 | | | 1 |
| | | TraesCS3B01G017400 | 1 | | | 1 |
| | | TraesCS4A01G044800 | | 1 | | |
| | HD | TraesCS1A01G435000 | | 1 | 1 | |
| | | TraesCS1D01G444100 | | 1 | | |
| | | TraesCS3B01G017400 | 1 | | | 1 |
| Resistance | Pm | TraesCS7D01G080300 | 1 | 1 | 1 | 1 |
| | Sr | TraesCS1A01G061500 | 1 | 1 | 1 | 1 |
| | | TraesCS4D01G243100 | 1 | 1 | 1 | 1 |
| | | TraesCS5A01G344100 | 1 | | | 1 |
| | | TraesCS6B01G185700 | 1 | 1 | 1 | 1 |
| | | TraesCS7D01G080300 | 1 | 1 | 1 | 1 |

PH: plant height, SL: spike length, PPR: ratio of peduncle length to plant height, SNS: spikelet number per spike, TGW: thousand grain weight, GL: grain length, Vmax: maximum filling rate, V: average grain filling rate, T: active filling period, HD: heading date, FD: flowering date, Pm: powdery mildew, Sr: stripe rust.

**Supplementary Table S36.** The results for accessible chromatin regions of candidate genes for wheat grain-related traits

| gene | trait | IWGSC_v1.1 | Chr | Start | End | DAP5 | DAP9 | DAP15 | DAP20 |
|---|---|---|---|---|---|---|---|---|---|
| TraesCS1A01G033500 | T | TraesCS1A02G033500 | 1A | 16388445 | 16391082 | 551097 | 34228 | 43 | 83 |
| TraesCS1A01G033600 | T | TraesCS1A02G033600 | 1A | 16407008 | 16410201 | 531978 | 15109 | 252 | 15842 |
| TraesCS2A01G176000 | T | TraesCS2A02G176000 | 2A | 135191755 | 135197161 | 247 | overlap | 3699 | 3743 |
| TraesCS2B01G194200 | T | TraesCS2B02G194200 | 2B | 171030105 | 171036053 | overlap | overlap | 72 | 1860 |
| TraesCS2B01G578500 | T | TraesCS2B02G578500 | 2B | 766557473 | 766568984 | 1022 | 1111 | 755 | 716 |
| TraesCS4A01G302000 | T | TraesCS4A02G302000 | 4A | 598719241 | 598726726 | 1443 | 120 | 3434 | overlap |
| TraesCS4B01G390500 | T | TraesCS4B02G390500 | 4B | 666581148 | 666586473 | overlap | overlap | 101 | 110 |
| TraesCS7D01G159800 | T | TraesCS7D02G159800 | 7D | 109170863 | 109176000 | overlap | overlap | overlap | 801 |
| TraesCS7D01G512100 | T | TraesCS7D02G512100 | 7D | 613867768 | 613870602 | 7 | 82 | 69 | 97 |
| TraesCS1A01G214700 | V | TraesCS1A02G214700 | 1A | 377613069 | 377615762 | overlap | overlap | overlap | 2367 |
| TraesCS2A01G293400 | V | TraesCS2A02G293400 | 2A | 504334128 | 504344701 | overlap | overlap | overlap | overlap |
| TraesCS2B01G202300 | V | TraesCS2B02G202300 | 2B | 182011547 | 182016800 | overlap | 488 | 121 | 21789 |
| TraesCS3A01G015500 | V | TraesCS3A02G015500 | 3A | 10300292 | 10309206 | 76 | 98 | overlap | overlap |
| TraesCS3B01G469000 | V | TraesCS3B02G469000 | 3B | 715879545 | 715885277 | 189 | 96 | overlap | 243 |
| TraesCS3B01G560400 | V | TraesCS3B02G560400 | 3B | 794543281 | 794544515 | 73 | 1757 | 270 | 186500 |
| TraesCS3B01G561000 | V | TraesCS3B02G561000 | 3B | 794839275 | 794840475 | 77 | 33 | 2482 | 2310 |
| TraesCS4A01G016600 | V | TraesCS4A02G016600 | 4A | 11296831 | 11299759 | overlap | 2571 | 1659 | 2789 |
| TraesCS5B01G000100 | V | TraesCS5B02G000100 | 5B | 16235 | 21357 | overlap | overlap | overlap | 39461 |
| TraesCS6B01G009600 | V | TraesCS6B02G009600 | 6B | 6015870 | 6019641 | overlap | overlap | overlap | overlap |
| TraesCS6D01G403800 | V | TraesCS6D02G403800 | 6D | 471917727 | 471921914 | overlap | 6722 | overlap | 6871 |
| TraesCS7A01G518200 | V | TraesCS7A02G518200 | 7A | 701764017 | 701771932 | 222 | 184 | 151 | 81 |
| TraesCS7A01G518300 | V | TraesCS7A02G518300 | 7A | 701803301 | 701806216 | overlap | overlap | overlap | 1086 |
| TraesCS7A01G549100 | V | TraesCS7A02G549100 | 7A | 723259779 | 723265225 | 313 | 252 | overlap | 149 |
| TraesCS7A01G549200 | V | TraesCS7A02G549200 | 7A | 723275296 | 723280279 | 227 | 249 | overlap | 161 |
| TraesCS7A01G549300 | V | TraesCS7A02G549300 | 7A | 723283231 | 723293633 | overlap | overlap | overlap | 82 |
| TraesCS7B01G034600 | V | TraesCS7B02G034600 | 7B | 33906007 | 33915606 | 6186 | 95410 | overlap | 95412 |
| TraesCS7B01G115300 | V | TraesCS7B02G115300 | 7B | 133791961 | 133792855 | 41 | 49 | 48753 | 49602 |
| TraesCS7B01G439900 | V | TraesCS7B02G439900 | 7B | 705251306 | 705254066 | 152 | 113 | 147 | 120 |
| TraesCS7B01G472500 | V | TraesCS7B02G472500 | 7B | 728689640 | 728694839 | 193 | 168 | 235 | 150 |
| TraesCS7D01G215600 | V | TraesCS7D02G215600 | 7D | 175646622 | 175653409 | 405 | 492 | 29760 | 497 |
| TraesCS1A01G033500 | Vmax | TraesCS1A02G033500 | 1A | 16388445 | 16391082 | 551097 | 34228 | 43 | 83 |
| TraesCS1A01G033600 | Vmax | TraesCS1A02G033600 | 1A | 16407008 | 16410201 | 531978 | 15109 | 252 | 15842 |
| TraesCS1A01G219400 | Vmax | TraesCS1A02G219400 | 1A | 388156746 | 388159972 | 80 | 197 | 16 | 166 |
| TraesCS1A01G247100 | Vmax | TraesCS1A02G247100 | 1A | 438365795 | 438368421 | 207 | overlap | overlap | overlap |
| TraesCS2A01G293400 | Vmax | TraesCS2A02G293400 | 2A | 504334128 | 504344701 | overlap | overlap | overlap | overlap |
| TraesCS2B01G202300 | Vmax | TraesCS2B02G202300 | 2B | 182011547 | 182016800 | overlap | 488 | 121 | 21789 |
| TraesCS3B01G469000 | Vmax | TraesCS3B02G469000 | 3B | 715879545 | 715885277 | 189 | 96 | overlap | 243 |
| TraesCS4A01G016600 | Vmax | TraesCS4A02G016600 | 4A | 11296831 | 11299759 | overlap | 2571 | 1659 | 2789 |
| TraesCS4B01G390500 | Vmax | TraesCS4B02G390500 | 4B | 666581148 | 666586473 | overlap | overlap | 101 | 110 |
| TraesCS5B01G039800 | Vmax | TraesCS5B02G039800 | 5B | 44776428 | 44782874 | 2049 | 2050 | 2172 | 2115 |

| | | | | | | | | | |
|---|---|---|---|---|---|---|---|---|---|
| TraesCS5B01G400000 | Vmax | TraesCS5B02G400000 | 5B | 577301899 | 577309490 | 176 | overlap | 41 | 27 |
| TraesCS6B01G009600 | Vmax | TraesCS6B02G009600 | 6B | 6015870 | 6019641 | overlap | overlap | overlap | overlap |
| TraesCS6B01G369700 | Vmax | TraesCS6B02G369700 | 6B | 643259758 | 643265986 | overlap | overlap | overlap | 3216 |
| TraesCS6D01G403800 | Vmax | TraesCS6D02G403800 | 6D | 471917727 | 471921914 | overlap | 6722 | overlap | 6871 |
| TraesCS7A01G549100 | Vmax | TraesCS7A02G549100 | 7A | 723259779 | 723265225 | 313 | 252 | overlap | 149 |
| TraesCS7A01G549200 | Vmax | TraesCS7A02G549200 | 7A | 723275296 | 723280279 | 227 | 249 | overlap | 161 |
| TraesCS7A01G549300 | Vmax | TraesCS7A02G549300 | 7A | 723283231 | 723293633 | overlap | overlap | overlap | 82 |
| TraesCS7B01G034600 | Vmax | TraesCS7B02G034600 | 7B | 33906007 | 33915606 | 6186 | 95410 | overlap | 95412 |
| TraesCS7B01G115300 | Vmax | TraesCS7B02G115300 | 7B | 133791961 | 133792855 | 41 | 49 | 48753 | 49602 |
| TraesCS7B01G439900 | Vmax | TraesCS7B02G439900 | 7B | 705251306 | 705254066 | 152 | 113 | 147 | 120 |
| TraesCS7B01G472500 | Vmax | TraesCS7B02G472500 | 7B | 728689640 | 728694839 | 193 | 168 | 235 | 150 |
| TraesCS7D01G215600 | Vmax | TraesCS7D02G215600 | 7D | 175646622 | 175653409 | 405 | 492 | 29760 | 497 |
| TraesCS2A01G010600 | GL | TraesCS2A02G010600 | 2A | 4083824 | 4089576 | 12290 | overlap | 70 | 77 |
| TraesCS2A01G482400 | GL | TraesCS2A02G482400 | 2A | 718935574 | 718940889 | 1931 | 25335 | 2766 | 1542 |
| TraesCS2D01G011200 | GL | TraesCS2D02G011200 | 2D | 5435218 | 5441031 | overlap | overlap | overlap | 5648 |
| TraesCS2D01G082000 | GL | TraesCS2D02G082000 | 2D | 35259471 | 35263395 | 371 | overlap | 45 | 30 |
| TraesCS2D01G482500 | GL | TraesCS2D02G482500 | 2D | 584799573 | 584804705 | 32788 | 32923 | 34 | 33292 |
| TraesCS3B01G560400 | GL | TraesCS3B02G560400 | 3B | 794543281 | 794544515 | 73 | 1757 | 270 | 186500 |
| TraesCS3B01G561000 | GL | TraesCS3B02G561000 | 3B | 794839275 | 794840475 | 77 | 33 | 2482 | 2310 |
| TraesCS4A01G266900 | GL | TraesCS4A02G266900 | 4A | 579428158 | 579431177 | overlap | 215 | overlap | overlap |
| TraesCS4A01G444300 | GL | TraesCS4A02G444300 | 4A | 712862865 | 712874048 | 6993 | 6787 | 46712 | 46383 |
| TraesCS4B01G047400 | GL | TraesCS4B02G047400 | 4B | 34722272 | 34725256 | overlap | overlap | overlap | 148 |
| TraesCS4B01G321800 | GL | TraesCS4B02G321800 | 4B | 612759007 | 612763642 | 579 | 783 | 327 | 513 |
| TraesCS4B01G360800 | GL | TraesCS4B02G360800 | 4B | 650683204 | 650688747 | overlap | overlap | 89 | overlap |
| TraesCS4D01G318400 | GL | TraesCS4D02G318400 | 4D | 482773327 | 482777407 | overlap | 261 | overlap | 254 |
| TraesCS4D01G325200 | GL | TraesCS4D02G325200 | 4D | 485520708 | 485526115 | overlap | overlap | overlap | overlap |
| TraesCS5A01G377700 | GL | TraesCS5A02G377700 | 5A | 575522382 | 575525543 | 165 | 280 | 130 | 182 |
| TraesCS5B01G307600 | GL | TraesCS5B02G307600 | 5B | 490769556 | 490771476 | overlap | 38931 | overlap | 17738 |
| TraesCS6D01G378300 | GL | TraesCS6D02G378300 | 6D | 460732888 | 460735270 | overlap | overlap | 151 | 195 |
| TraesCS7B01G018600 | GL | TraesCS7B02G018600 | 7B | 15592400 | 15602503 | overlap | overlap | overlap | overlap |
| TraesCS7D01G039500 | GL | TraesCS7D02G039500 | 7D | 20057020 | 20069082 | overlap | overlap | overlap | overlap |
| TraesCS1A01G247100 | GW | TraesCS1A02G247100 | 1A | 438365795 | 438368421 | 207 | overlap | overlap | overlap |
| TraesCS1A01G329900 | GW | TraesCS1A02G329900 | 1A | 518808509 | 518813919 | overlap | overlap | overlap | overlap |
| TraesCS1B01G155700 | GW | TraesCS1B02G155700 | 1B | 263547930 | 263561834 | 100 | 41 | overlap | 260 |
| TraesCS1B01G462000 | GW | TraesCS1B02G462000 | 1B | 674627939 | 674633403 | 65 | 1474 | overlap | overlap |
| TraesCS2A01G464000 | GW | TraesCS2A02G464000 | 2A | 709772645 | 709775780 | 910 | overlap | 712 | 561 |
| TraesCS2D01G505700 | GW | TraesCS2D02G505700 | 2D | 599916155 | 599919148 | 429 | 553 | 6980 | 256 |
| TraesCS6B01G130700 | GW | TraesCS6B02G130700 | 6B | 127321527 | 127326100 | 4888 | 4875 | 4959 | 4872 |
| TraesCS6B01G186700 | GW | TraesCS6B02G186700 | 6B | 212968226 | 212970183 | 2433 | 667 | 270446 | 270557 |
| TraesCS6D01G009700 | GW | TraesCS6D02G009700 | 6D | 3738920 | 3741139 | 4767 | 4854 | 321 | 22890 |
| TraesCS7A01G043700 | GW | TraesCS7A02G043700 | 7A | 20170996 | 20182247 | 98149 | 100697 | 197683 | 832 |
| TraesCS7A01G120300 | GW | TraesCS7A02G120300 | 7A | 77434519 | 77442262 | 16588 | overlap | overlap | 15 |
| TraesCS7B01G115300 | GW | TraesCS7B02G115300 | 7B | 133791961 | 133792855 | 41 | 49 | 48753 | 49602 |
| TraesCS7B01G160000 | GW | TraesCS7B02G160000 | 7B | 218547229 | 218549774 | overlap | overlap | 15389 | 20033 |

| | | | | | | | | | |
|---|---|---|---|---|---|---|---|---|---|
| TraesCS1A01G353300 | GWLR | TraesCS1A02G353300 | 1A | 536958431 | 536965909 | overlap | overlap | overlap | 85095 |
| TraesCS2A01G295400 | GWLR | TraesCS2A02G295400 | 2A | 508030243 | 508034286 | overlap | overlap | overlap | 17777 |
| TraesCS2A01G548100 | GWLR | TraesCS2A02G548100 | 2A | 755927033 | 755931199 | overlap | 18 | overlap | overlap |
| TraesCS2D01G011200 | GWLR | TraesCS2D02G011200 | 2D | 5435218 | 5441031 | overlap | overlap | overlap | 5648 |
| TraesCS3B01G128700 | GWLR | TraesCS3B02G128700 | 3B | 107958655 | 107961101 | 293911 | 325417 | 3142 | 3065 |
| TraesCS4A01G074300 | GWLR | TraesCS4A02G074300 | 4A | 73541616 | 73543573 | 1265 | 1189 | 79970 | 1717 |
| TraesCS4A01G227900 | GWLR | TraesCS4A02G227900 | 4A | 536202316 | 536211275 | 56 | 18 | 439 | 326 |
| TraesCS4A01G418200 | GWLR | TraesCS4A02G418200 | 4A | 688097145 | 688100962 | overlap | overlap | overlap | overlap |
| TraesCS4D01G350500 | GWLR | TraesCS4D02G350500 | 4D | 502708408 | 502712304 | overlap | overlap | overlap | 11797 |
| TraesCS4D01G353800 | GWLR | TraesCS4D02G353800 | 4D | 504087839 | 504093419 | overlap | overlap | overlap | 61 |
| TraesCS5D01G387700 | GWLR | TraesCS5D02G387700 | 5D | 457624283 | 457628075 | 21017 | 20559 | 223 | 37633 |
| TraesCS7A01G549100 | GWLR | TraesCS7A02G549100 | 7A | 723259779 | 723265225 | 313 | 252 | overlap | 149 |
| TraesCS7A01G549200 | GWLR | TraesCS7A02G549200 | 7A | 723275296 | 723280279 | 227 | 249 | overlap | 161 |
| TraesCS7A01G549300 | GWLR | TraesCS7A02G549300 | 7A | 723283231 | 723293633 | overlap | overlap | overlap | 82 |
| TraesCS7D01G039500 | GWLR | TraesCS7D02G039500 | 7D | 20057020 | 20069082 | overlap | overlap | overlap | overlap |
| TraesCS1A01G050400 | TGW | TraesCS1A02G050400 | 1A | 32260043 | 32262299 | 3556 | overlap | 539 | 19754 |
| TraesCS1A01G329900 | TGW | TraesCS1A02G329900 | 1A | 518808509 | 518813919 | overlap | overlap | overlap | overlap |
| TraesCS1A01G435000 | TGW | TraesCS1A02G435000 | 1A | 585625902 | 585628146 | overlap | overlap | overlap | 100 |
| TraesCS1B01G368500 | TGW | TraesCS1B02G368500 | 1B | 598877339 | 598884411 | overlap | overlap | overlap | overlap |
| TraesCS1B01G462000 | TGW | TraesCS1B02G462000 | 1B | 674627939 | 674633403 | 65 | 1474 | overlap | overlap |
| TraesCS1D01G059200 | TGW | TraesCS1D02G059200 | 1D | 39287710 | 39292333 | 4727 | 7685 | 494814 | 7740 |
| TraesCS2A01G310300 | TGW | TraesCS2A02G310300 | 2A | 533893908 | 533915882 | overlap | overlap | overlap | overlap |
| TraesCS2A01G548100 | TGW | TraesCS2A02G548100 | 2A | 755927033 | 755931199 | overlap | 18 | overlap | overlap |
| TraesCS2B01G010800 | TGW | TraesCS2B02G010800 | 2B | 5796815 | 5806855 | 6019 | 5996 | 6170 | 6114 |
| TraesCS2B01G012500 | TGW | TraesCS2B02G012500 | 2B | 6257247 | 6261132 | overlap | overlap | 56 | overlap |
| TraesCS2B01G533300 | TGW | TraesCS2B02G533300 | 2B | 729627351 | 729630308 | 461 | 106 | 1963 | 2320 |
| TraesCS3A01G311000 | TGW | TraesCS3A02G311000 | 3A | 549902478 | 549907314 | overlap | overlap | overlap | overlap |
| TraesCS3B01G252000 | TGW | TraesCS3B02G252000 | 3B | 404826418 | 404833319 | overlap | overlap | overlap | overlap |
| TraesCS4A01G289300 | TGW | TraesCS4A02G289300 | 4A | 593761733 | 593766066 | overlap | 78 | overlap | 38 |
| TraesCS4A01G444300 | TGW | TraesCS4A02G444300 | 4A | 712862865 | 712874048 | 6993 | 6787 | 46712 | 46383 |
| TraesCS4B01G029700 | TGW | TraesCS4B02G029700 | 4B | 21937120 | 21944075 | 1170 | overlap | overlap | 455 |
| TraesCS4D01G350500 | TGW | TraesCS4D02G350500 | 4D | 502708408 | 502712304 | overlap | overlap | overlap | 11797 |
| TraesCS4D01G353800 | TGW | TraesCS4D02G353800 | 4D | 504087839 | 504093419 | overlap | overlap | overlap | 61 |
| TraesCS5B01G292100 | TGW | TraesCS5B02G292100 | 5B | 477609542 | 477611760 | 47194 | 55084 | 55042 | 1874 |
| TraesCS6A01G194600 | TGW | TraesCS6A02G194600 | 6A | 272122365 | 272141924 | overlap | overlap | overlap | overlap |
| TraesCS6D01G009700 | TGW | TraesCS6D02G009700 | 6D | 3738920 | 3741139 | 4767 | 4854 | 321 | 22890 |
| TraesCS7A01G043700 | TGW | TraesCS7A02G043700 | 7A | 20170996 | 20182247 | 98149 | 100697 | 197683 | 832 |
| TraesCS7B01G455000 | TGW | TraesCS7B02G455000 | 7B | 715713483 | 715716700 | overlap | overlap | overlap | overlap |
| TraesCS7D01G309400 | TGW | TraesCS7D02G309400 | 7D | 392744990 | 392749017 | 792 | overlap | 13188 | 8553 |
| TraesCS7D01G525100 | TGW | TraesCS7D02G525100 | 7D | 620989956 | 620992337 | 3267 | 3249 | 3237 | 1023 |
| TraesCS2B01G464900 | SNS | TraesCS2B02G464900 | 2B | 659083257 | 659085942 | 3297 | 2319 | 2628 | 2349 |
| TraesCS3A01G022700 | SNS | TraesCS3A02G022700 | 3A | 12835921 | 12841532 | overlap | overlap | overlap | overlap |
| TraesCS3A01G026900 | SNS | TraesCS3A02G026900 | 3A | 14043984 | 14044938 | 184 | 9 | 72 | 164 |
| TraesCS3A01G391600 | SNS | TraesCS3A02G391600 | 3A | 639479177 | 639483225 | 55126 | 55060 | 729 | 145771 |

| | | | | | | | | | |
|---|---|---|---|---|---|---|---|---|---|
| TraesCS3A01G502900 | SNS | TraesCS3A02G502900 | 3A | 725842411 | 725843058 | 4290 | 480 | 4176 | 599 |
| TraesCS3B01G525300 | SNS | TraesCS3B02G525300 | 3B | 766978875 | 766981579 | 1108 | 1274 | 91 | 2193 |
| TraesCS4A01G079300 | SNS | TraesCS4A02G079300 | 4A | 80856864 | 80857712 | 794 | 1144 | 9692 | 21111 |
| TraesCS4A01G266900 | SNS | TraesCS4A02G266900 | 4A | 579428158 | 579431177 | overlap | 215 | overlap | overlap |
| TraesCS5A01G272400 | SNS | TraesCS5A02G272400 | 5A | 482118145 | 482120100 | overlap | overlap | overlap | overlap |
| TraesCS5B01G469600 | SNS | TraesCS5B02G469600 | 5B | 643260159 | 643261737 | 4220 | 36466 | 42474 | 35458 |
| TraesCS7B01G271200 | SNS | TraesCS7B02G271200 | 7B | 498085563 | 498091364 | overlap | overlap | 89719 | 87151 |

DAP: days after pollination.

**Supplementary Table S37**. GO annotation analysis for the genes with selection signals between landraces and cultivars with different periods

| GO ID | Function | Gene number | P.adj | Scope | Group |
|---|---|---|---|---|---|
| GO:0006468 | protein phosphorylation | 3218 | 2.11E-28 | Whole | All |
| GO:0055114 | oxidation-reduction process | 3109 | 6.50E-123 | Whole | All |
| GO:0006355 | regulation of transcription, DNA-templated | 1424 | 5.48E-65 | Whole | All |
| GO:0006508 | proteolysis | 847 | 1.01E-90 | Whole | All |
| GO:0005975 | carbohydrate metabolic process | 776 | 5.47E-52 | Whole | All |
| GO:0048544 | recognition of pollen | 358 | 4.26E-65 | Whole | All |
| GO:0006629 | lipid metabolic process | 265 | 7.72E-26 | Whole | All |
| GO:0055085 | transmembrane transport | 208 | 7.88E-95 | Whole | All |
| GO:0009733 | response to auxin | 201 | 7.69E-20 | Whole | All |
| GO:0009058 | biosynthetic process | 199 | 3.27E-20 | Whole | All |
| GO:0016567 | protein ubiquitination | 129 | 1.65E-12 | Whole | All |
| GO:0000160 | phosphorelay signal transduction system | 124 | 5.29E-13 | Whole | All |
| GO:0006396 | RNA processing | 105 | 2.44E-12 | Whole | All |
| GO:0009611 | response to wounding | 102 | 5.32E-05 | Whole | All |
| GO:0016579 | protein deubiquitination | 101 | 1.01E-16 | Whole | All |
| GO:0016192 | vesicle-mediated transport | 97 | 1.79E-02 | Whole | All |
| GO:0006139 | nucleobase-containing compound metabolic process | 86 | 4.49E-04 | Whole | All |
| GO:0006096 | glycolytic process | 79 | 1.81E-10 | Whole | All |
| GO:0046488 | phosphatidylinositol metabolic process | 69 | 2.89E-15 | Whole | All |
| GO:0019752 | carboxylic acid metabolic process | 65 | 3.36E-08 | Whole | All |
| GO:0006952 | defense response | 62 | 8.01E-04 | Whole | All |
| GO:0006886 | intracellular protein transport | 61 | 4.03E-05 | Whole | All |
| GO:0042545 | cell wall modification | 46 | 3.12E-02 | Whole | All |
| GO:0031047 | gene silencing by RNA | 45 | 4.35E-03 | Whole | All |
| GO:0005992 | trehalose biosynthetic process | 44 | 6.52E-04 | Whole | All |
| GO:0006364 | rRNA processing | 44 | 1.28E-02 | Whole | All |
| GO:0017148 | negative regulation of translation | 43 | 2.30E-04 | Whole | All |
| GO:0000413 | protein peptidyl-prolyl isomerization | 41 | 1.92E-03 | Whole | All |
| GO:0006457 | protein folding | 40 | 1.00E-02 | Whole | All |
| GO:0006413 | translational initiation | 39 | 8.91E-03 | Whole | All |
| GO:0009415 | response to water | 38 | 4.52E-05 | Whole | All |
| GO:0006807 | nitrogen compound metabolic process | 35 | 9.65E-03 | Whole | All |
| GO:0016998 | cell wall macromolecule catabolic process | 34 | 2.71E-02 | Whole | All |
| GO:0006284 | base-excision repair | 33 | 5.96E-04 | Whole | All |
| GO:0005985 | sucrose metabolic process | 32 | 1.29E-05 | Whole | All |
| GO:0007010 | cytoskeleton organization | 31 | 1.05E-06 | Whole | All |
| GO:0043039 | tRNA aminoacylation | 30 | 3.11E-02 | Whole | All |
| GO:1902600 | proton transmembrane transport | 29 | 4.84E-02 | Whole | All |
| GO:0043631 | RNA polyadenylation | 28 | 8.09E-03 | Whole | All |
| GO:0007275 | multicellular organism development | 26 | 7.44E-03 | Whole | All |
| GO:0050832 | defense response to fungus | 26 | 8.09E-03 | Whole | All |
| GO:0009451 | RNA modification | 26 | 8.09E-03 | Whole | All |
| GO:0006308 | DNA catabolic process | 24 | 4.80E-05 | Whole | All |
| GO:0002100 | tRNA wobble adenosine to inosine editing | 23 | 1.21E-05 | Whole | All |
| GO:0010274 | hydrotropism | 23 | 1.98E-02 | Whole | All |
| GO:0003333 | amino acid transmembrane transport | 22 | 1.14E-02 | Whole | All |
| GO:0015979 | photosynthesis | 21 | 1.45E-19 | Whole | All |
| GO:0046373 | L-arabinose metabolic process | 21 | 9.74E-06 | Whole | All |
| GO:0016226 | iron-sulfur cluster assembly | 21 | 1.79E-02 | Whole | All |
| GO:0009793 | embryo development ending in seed dormancy | 21 | 2.77E-02 | Whole | All |
| GO:0006506 | GPI anchor biosynthetic process | 21 | 3.63E-02 | Whole | All |
| GO:0006164 | purine nucleotide biosynthetic process | 20 | 8.04E-04 | Whole | All |
| GO:0035556 | intracellular signal transduction | 20 | 9.98E-03 | Whole | All |
| GO:0019346 | transsulfuration | 20 | 2.04E-02 | Whole | All |
| GO:0030418 | nicotianamine biosynthetic process | 20 | 2.21E-02 | Whole | All |
| GO:0043248 | proteasome assembly | 19 | 2.60E-04 | Whole | All |
| GO:0006412 | translation | 18 | 1.30E-78 | Whole | All |
| GO:0015937 | coenzyme A biosynthetic process | 18 | 2.58E-04 | Whole | All |
| GO:0006568 | tryptophan metabolic process | 18 | 1.01E-02 | Whole | All |
| GO:0008612 | peptidyl-lysine modification to peptidyl-hypusine | 17 | 1.29E-05 | Whole | All |
| GO:0015977 | carbon fixation | 17 | 6.12E-03 | Whole | All |
| GO:0043666 | regulation of phosphoprotein phosphatase activity | 16 | 3.61E-04 | Whole | All |
| GO:0032259 | methylation | 16 | 4.76E-02 | Whole | All |
| GO:0006221 | pyrimidine nucleotide biosynthetic process | 15 | 2.18E-03 | Whole | All |
| GO:0016573 | histone acetylation | 15 | 1.23E-02 | Whole | All |
| GO:0000724 | double-strand break repair via homologous recombination | 12 | 2.79E-05 | Whole | All |
| GO:0006370 | 7-methylguanosine mRNA capping | 12 | 1.64E-04 | Whole | All |
| GO:0006366 | transcription by RNA polymerase II | 12 | 6.31E-04 | Whole | All |
| GO:0006488 | dolichol-linked oligosaccharide biosynthetic process | 12 | 5.33E-03 | Whole | All |
| GO:0006529 | asparagine biosynthetic process | 12 | 2.48E-02 | Whole | All |
| GO:0071569 | protein ufmylation | 11 | 5.27E-03 | Whole | All |
| GO:0043161 | proteasome-mediated ubiquitin-dependent protein catabolic process | 10 | 1.14E-02 | Whole | All |

| | | | | | |
|---|---|---|---|---|---|
| GO:0008295 | spermidine biosynthetic process | 10 | 1.14E-02 | Whole | All |
| GO:0006000 | fructose metabolic process | 9 | 7.53E-03 | Whole | All |
| GO:0050790 | regulation of catalytic activity | 9 | 1.14E-02 | Whole | All |
| GO:0030638 | polyketide metabolic process | 8 | 1.08E-02 | Whole | All |
| GO:0006888 | endoplasmic reticulum to Golgi vesicle-mediated transport | 7 | 2.13E-04 | Whole | All |
| GO:0000469 | cleavage involved in rRNA processing | 6 | 2.36E-02 | Whole | All |
| GO:0044238 | primary metabolic process | 6 | 3.95E-02 | Whole | All |
| GO:0019684 | photosynthesis, light reaction | 5 | 6.61E-03 | Whole | All |
| GO:0007018 | microtubule-based movement | 4 | 2.19E-10 | Whole | All |
| GO:0045017 | glycerolipid biosynthetic process | 2 | 2.36E-02 | Whole | All |
| GO:0006556 | S-adenosylmethionine biosynthetic process | 2 | 4.63E-02 | Whole | All |
| GO:0042558 | pteridine-containing compound metabolic process | 2 | 4.44E-02 | A | LR ~ Cul |
| GO:0015995 | chlorophyll biosynthetic process | 2 | 4.44E-02 | A | LR ~ Cul |
| GO:0006412 | translation | 3 | 4.33E-09 | A | LR ~ Cul |
| GO:0016480 | negative regulation of transcription by RNA polymerase III | 3 | 6.50E-03 | A | LR ~ Cul |
| GO:0015979 | photosynthesis | 3 | 1.81E-02 | A | LR ~ Cul |
| GO:0071569 | protein ufmylation | 3 | 2.62E-02 | A | LR ~ Cul |
| GO:0018298 | protein-chromophore linkage | 3 | 2.62E-02 | A | LR ~ Cul |
| GO:0006488 | dolichol-linked oligosaccharide biosynthetic process | 3 | 3.69E-02 | A | LR ~ Cul |
| GO:0043085 | positive regulation of catalytic activity | 3 | 3.69E-02 | A | LR ~ Cul |
| GO:0006813 | potassium ion transport | 4 | 2.07E-03 | A | LR ~ Cul |
| GO:0008612 | peptidyl-lysine modification to peptidyl-hypusine | 4 | 4.93E-03 | A | LR ~ Cul |
| GO:0000077 | DNA damage checkpoint | 4 | 1.15E-02 | A | LR ~ Cul |
| GO:0006221 | pyrimidine nucleotide biosynthetic process | 4 | 1.52E-02 | A | LR ~ Cul |
| GO:0006729 | tetrahydrobiopterin biosynthetic process | 5 | 2.71E-04 | A | LR ~ Cul |
| GO:0016226 | iron-sulfur cluster assembly | 5 | 3.24E-02 | A | LR ~ Cul |
| GO:0046373 | L-arabinose metabolic process | 6 | 3.47E-04 | A | LR ~ Cul |
| GO:0006506 | GPI anchor biosynthetic process | 6 | 1.52E-02 | A | LR ~ Cul |
| GO:0006308 | DNA catabolic process | 7 | 3.47E-04 | A | LR ~ Cul |
| GO:0009793 | embryo development ending in seed dormancy | 8 | 5.37E-04 | A | LR ~ Cul |
| GO:0017148 | negative regulation of translation | 8 | 1.54E-02 | A | LR ~ Cul |
| GO:0006364 | rRNA processing | 8 | 4.90E-02 | A | LR ~ Cul |
| GO:0006952 | defense response | 11 | 1.81E-02 | A | LR ~ Cul |
| GO:0000160 | phosphorelay signal transduction system | 14 | 2.62E-02 | A | LR ~ Cul |
| GO:0009058 | biosynthetic process | 18 | 4.78E-02 | A | LR ~ Cul |
| GO:0006629 | lipid metabolic process | 25 | 1.64E-02 | A | LR ~ Cul |
| GO:0055085 | transmembrane transport | 33 | 6.18E-09 | A | LR ~ Cul |
| GO:0009733 | response to auxin | 44 | 1.71E-13 | A | LR ~ Cul |
| GO:0048544 | recognition of pollen | 63 | 1.91E-23 | A | LR ~ Cul |
| GO:0005975 | carbohydrate metabolic process | 66 | 1.71E-02 | A | LR ~ Cul |
| GO:0006508 | proteolysis | 81 | 6.28E-08 | A | LR ~ Cul |
| GO:0006355 | regulation of transcription, DNA-templated | 138 | 2.89E-04 | A | LR ~ Cul |
| GO:0055114 | oxidation-reduction process | 375 | 8.78E-21 | A | LR ~ Cul |
| GO:0006468 | protein phosphorylation | 406 | 1.25E-07 | A | LR ~ Cul |
| GO:0006412 | translation | 1 | 6.57E-07 | B | LR ~ Cul |
| GO:0044238 | primary metabolic process | 2 | 2.93E-02 | B | LR ~ Cul |
| GO:0006303 | double-strand break repair via nonhomologous end joining | 3 | 7.38E-03 | B | LR ~ Cul |
| GO:0006606 | protein import into nucleus | 4 | 1.69E-03 | B | LR ~ Cul |
| GO:0006414 | translational elongation | 4 | 1.60E-02 | B | LR ~ Cul |
| GO:0000105 | histidine biosynthetic process | 4 | 2.04E-02 | B | LR ~ Cul |
| GO:0000724 | double-strand break repair via homologous recombination | 5 | 3.38E-06 | B | LR ~ Cul |
| GO:0045927 | positive regulation of growth | 5 | 4.35E-02 | B | LR ~ Cul |
| GO:0006457 | protein folding | 6 | 3.11E-02 | B | LR ~ Cul |
| GO:0002100 | tRNA wobble adenosine to inosine editing | 7 | 5.66E-06 | B | LR ~ Cul |
| GO:0031047 | gene silencing by RNA | 7 | 1.44E-02 | B | LR ~ Cul |
| GO:0042545 | cell wall modification | 9 | 3.22E-03 | B | LR ~ Cul |
| GO:0055085 | transmembrane transport | 16 | 3.22E-08 | B | LR ~ Cul |
| GO:0009611 | response to wounding | 16 | 1.72E-04 | B | LR ~ Cul |
| GO:0006396 | RNA processing | 17 | 5.21E-07 | B | LR ~ Cul |
| GO:0048544 | recognition of pollen | 24 | 7.37E-06 | B | LR ~ Cul |
| GO:0006508 | proteolysis | 60 | 3.22E-08 | B | LR ~ Cul |
| GO:0005975 | carbohydrate metabolic process | 60 | 4.78E-06 | B | LR ~ Cul |
| GO:0006355 | regulation of transcription, DNA-templated | 118 | 2.40E-09 | B | LR ~ Cul |
| GO:0055114 | oxidation-reduction process | 217 | 7.24E-09 | B | LR ~ Cul |
| GO:0006468 | protein phosphorylation | 226 | 4.88E-02 | B | LR ~ Cul |
| GO:0009052 | pentose-phosphate shunt, non-oxidative branch | 2 | 1.17E-02 | D | LR ~ Cul |
| GO:0010112 | regulation of systemic acquired resistance | 2 | 1.55E-02 | D | LR ~ Cul |
| GO:0016042 | lipid catabolic process | 2 | 2.12E-02 | D | LR ~ Cul |
| GO:0043248 | proteasome assembly | 2 | 2.12E-02 | D | LR ~ Cul |
| GO:0009094 | L-phenylalanine biosynthetic process | 2 | 2.56E-02 | D | LR ~ Cul |
| GO:0008612 | peptidyl-lysine modification to peptidyl-hypusine | 3 | 4.88E-04 | D | LR ~ Cul |
| GO:0006221 | pyrimidine nucleotide biosynthetic process | 3 | 1.31E-03 | D | LR ~ Cul |
| GO:0055085 | transmembrane transport | 3 | 1.95E-03 | D | LR ~ Cul |
| GO:0006308 | DNA catabolic process | 3 | 3.59E-03 | D | LR ~ Cul |
| GO:0007010 | cytoskeleton organization | 3 | 5.57E-03 | D | LR ~ Cul |
| GO:0033014 | tetrapyrrole biosynthetic process | 3 | 9.34E-03 | D | LR ~ Cul |
| GO:0043039 | tRNA aminoacylation | 3 | 2.13E-02 | D | LR ~ Cul |
| GO:0005992 | trehalose biosynthetic process | 3 | 3.37E-02 | D | LR ~ Cul |
| GO:0006096 | glycolytic process | 4 | 1.59E-02 | D | LR ~ Cul |
| GO:0006807 | nitrogen compound metabolic process | 6 | 7.22E-05 | D | LR ~ Cul |

| | | | | | |
|---|---|---|---|---|---|
| GO:0009058 | biosynthetic process | 7 | 1.17E-02 | D | LR ~ Cul |
| GO:0006629 | lipid metabolic process | 7 | 3.31E-02 | D | LR ~ Cul |
| GO:0006508 | proteolysis | 15 | 2.34E-02 | D | LR ~ Cul |
| GO:0048544 | recognition of pollen | 20 | 2.12E-11 | D | LR ~ Cul |
| GO:0005975 | carbohydrate metabolic process | 27 | 6.93E-06 | D | LR ~ Cul |
| GO:0055114 | oxidation-reduction process | 55 | 4.01E-02 | D | LR ~ Cul |
| GO:0006468 | protein phosphorylation | 594 | 2.41E-19 | A | LR ~ 50s |
| GO:0055114 | oxidation-reduction process | 481 | 1.72E-25 | A | LR ~ 50s |
| GO:0006355 | regulation of transcription, DNA-templated | 202 | 1.32E-09 | A | LR ~ 50s |
| GO:0005975 | carbohydrate metabolic process | 99 | 1.71E-05 | A | LR ~ 50s |
| GO:0006508 | proteolysis | 91 | 4.13E-06 | A | LR ~ 50s |
| GO:0048544 | recognition of pollen | 89 | 7.01E-35 | A | LR ~ 50s |
| GO:0055085 | transmembrane transport | 36 | 2.43E-14 | A | LR ~ 50s |
| GO:0006629 | lipid metabolic process | 33 | 3.22E-03 | A | LR ~ 50s |
| GO:0009611 | response to wounding | 24 | 3.37E-04 | A | LR ~ 50s |
| GO:0000160 | phosphorelay signal transduction system | 23 | 9.58E-05 | A | LR ~ 50s |
| GO:0016567 | protein ubiquitination | 21 | 2.53E-03 | A | LR ~ 50s |
| GO:0006952 | defense response | 18 | 1.50E-04 | A | LR ~ 50s |
| GO:0019752 | carboxylic acid metabolic process | 15 | 1.34E-04 | A | LR ~ 50s |
| GO:0016579 | protein deubiquitination | 13 | 6.99E-03 | A | LR ~ 50s |
| GO:0006096 | glycolytic process | 12 | 1.11E-02 | A | LR ~ 50s |
| GO:0006308 | DNA catabolic process | 11 | 8.29E-07 | A | LR ~ 50s |
| GO:0046488 | phosphatidylinositol metabolic process | 10 | 2.34E-03 | A | LR ~ 50s |
| GO:0017148 | negative regulation of translation | 10 | 6.72E-03 | A | LR ~ 50s |
| GO:0005985 | sucrose metabolic process | 9 | 4.57E-04 | A | LR ~ 50s |
| GO:0008612 | peptidyl-lysine modification to peptidyl-hypusine | 8 | 1.79E-06 | A | LR ~ 50s |
| GO:0046373 | L-arabinose metabolic process | 8 | 1.98E-05 | A | LR ~ 50s |
| GO:0009793 | embryo development ending in seed dormancy | 8 | 2.18E-03 | A | LR ~ 50s |
| GO:0002100 | tRNA wobble adenosine to inosine editing | 7 | 4.07E-04 | A | LR ~ 50s |
| GO:0043631 | RNA polyadenylation | 7 | 2.83E-02 | A | LR ~ 50s |
| GO:0009086 | methionine biosynthetic process | 4 | 3.59E-03 | A | LR ~ 50s |
| GO:0006488 | dolichol-linked oligosaccharide biosynthetic process | 4 | 1.33E-02 | A | LR ~ 50s |
| GO:0000077 | DNA damage checkpoint | 4 | 1.99E-02 | A | LR ~ 50s |
| GO:0006412 | translation | 3 | 9.82E-13 | A | LR ~ 50s |
| GO:0016480 | negative regulation of transcription by RNA polymerase III | 3 | 9.31E-03 | A | LR ~ 50s |
| GO:0015979 | photosynthesis | 2 | 2.63E-04 | A | LR ~ 50s |
| GO:0006468 | protein phosphorylation | 424 | 4.11E-05 | B | LR ~ 50s |
| GO:0055114 | oxidation-reduction process | 408 | 2.21E-20 | B | LR ~ 50s |
| GO:0006355 | regulation of transcription, DNA-templated | 212 | 1.82E-17 | B | LR ~ 50s |
| GO:0005975 | carbohydrate metabolic process | 123 | 5.89E-16 | B | LR ~ 50s |
| GO:0006508 | proteolysis | 117 | 5.32E-18 | B | LR ~ 50s |
| GO:0048544 | recognition of pollen | 67 | 1.22E-23 | B | LR ~ 50s |
| GO:0009611 | response to wounding | 34 | 2.63E-10 | B | LR ~ 50s |
| GO:0006629 | lipid metabolic process | 33 | 2.88E-04 | B | LR ~ 50s |
| GO:0055085 | transmembrane transport | 28 | 6.07E-14 | B | LR ~ 50s |
| GO:0006396 | RNA processing | 21 | 4.43E-06 | B | LR ~ 50s |
| GO:0000160 | phosphorelay signal transduction system | 18 | 1.72E-03 | B | LR ~ 50s |
| GO:0006139 | nucleobase-containing compound metabolic process | 17 | 4.88E-03 | B | LR ~ 50s |
| GO:0006749 | glutathione metabolic process | 16 | 6.15E-03 | B | LR ~ 50s |
| GO:0006413 | translational initiation | 12 | 5.00E-04 | B | LR ~ 50s |
| GO:0019752 | carboxylic acid metabolic process | 11 | 4.06E-03 | B | LR ~ 50s |
| GO:0016998 | cell wall macromolecule catabolic process | 9 | 9.32E-03 | B | LR ~ 50s |
| GO:0006568 | tryptophan metabolic process | 8 | 1.38E-04 | B | LR ~ 50s |
| GO:0017148 | negative regulation of translation | 8 | 2.29E-02 | B | LR ~ 50s |
| GO:0002100 | tRNA wobble adenosine to inosine editing | 7 | 1.92E-04 | B | LR ~ 50s |
| GO:0016573 | histone acetylation | 6 | 1.58E-03 | B | LR ~ 50s |
| GO:0005985 | sucrose metabolic process | 6 | 1.46E-02 | B | LR ~ 50s |
| GO:0006366 | transcription by RNA polymerase II | 5 | 3.51E-04 | B | LR ~ 50s |
| GO:0006488 | dolichol-linked oligosaccharide biosynthetic process | 4 | 8.55E-03 | B | LR ~ 50s |
| GO:0015979 | photosynthesis | 4 | 1.76E-02 | B | LR ~ 50s |
| GO:0015937 | coenzyme A biosynthetic process | 4 | 2.09E-02 | B | LR ~ 50s |
| GO:0006370 | 7-methylguanosine mRNA capping | 3 | 1.33E-02 | B | LR ~ 50s |
| GO:0006303 | double-strand break repair via nonhomologous end joining | 3 | 2.73E-02 | B | LR ~ 50s |
| GO:0008612 | peptidyl-lysine modification to peptidyl-hypusine | 3 | 3.20E-02 | B | LR ~ 50s |
| GO:0006412 | translation | 1 | 7.93E-13 | B | LR ~ 50s |
| GO:0006468 | protein phosphorylation | 69 | 1.59E-05 | D | LR ~ 50s |
| GO:0048544 | recognition of pollen | 24 | 9.32E-20 | D | LR ~ 50s |
| GO:0006355 | regulation of transcription, DNA-templated | 21 | 2.42E-02 | D | LR ~ 50s |
| GO:0006508 | proteolysis | 20 | 3.26E-06 | D | LR ~ 50s |
| GO:0006511 | ubiquitin-dependent protein catabolic process | 11 | 1.59E-05 | D | LR ~ 50s |
| GO:0071569 | protein ufmylation | 4 | 3.93E-06 | D | LR ~ 50s |
| GO:0009415 | response to water | 4 | 6.98E-04 | D | LR ~ 50s |
| GO:0033014 | tetrapyrrole biosynthetic process | 3 | 3.39E-03 | D | LR ~ 50s |
| GO:0006298 | mismatch repair | 3 | 2.60E-02 | D | LR ~ 50s |
| GO:0003352 | regulation of cilium movement | 2 | 1.78E-03 | D | LR ~ 50s |
| GO:0043666 | regulation of phosphoprotein phosphatase activity | 2 | 8.73E-03 | D | LR ~ 50s |
| GO:0016042 | lipid catabolic process | 2 | 1.28E-02 | D | LR ~ 50s |
| GO:0006308 | DNA catabolic process | 2 | 1.80E-02 | D | LR ~ 50s |
| GO:0032259 | methylation | 2 | 2.22E-02 | D | LR ~ 50s |
| GO:0005985 | sucrose metabolic process | 2 | 3.26E-02 | D | LR ~ 50s |

| | | | | | |
|---|---|---|---|---|---|
| GO:0006468 | protein phosphorylation | 405 | 2.35E-04 | A | 50s ~ 60s-70s |
| GO:0055114 | oxidation-reduction process | 396 | 1.72E-19 | A | 50s ~ 60s-70s |
| GO:0006355 | regulation of transcription, DNA-templated | 185 | 1.85E-11 | A | 50s ~ 60s-70s |
| GO:0006508 | proteolysis | 113 | 3.18E-17 | A | 50s ~ 60s-70s |
| GO:0005975 | carbohydrate metabolic process | 93 | 5.42E-07 | A | 50s ~ 60s-70s |
| GO:0006629 | lipid metabolic process | 46 | 7.14E-10 | A | 50s ~ 60s-70s |
| GO:0048544 | recognition of pollen | 43 | 3.57E-10 | A | 50s ~ 60s-70s |
| GO:0016579 | protein deubiquitination | 34 | 1.02E-17 | A | 50s ~ 60s-70s |
| GO:0055085 | transmembrane transport | 27 | 1.86E-13 | A | 50s ~ 60s-70s |
| GO:0016192 | vesicle-mediated transport | 24 | 2.57E-04 | A | 50s ~ 60s-70s |
| GO:0009058 | biosynthetic process | 22 | 8.69E-03 | A | 50s ~ 60s-70s |
| GO:0009611 | response to wounding | 18 | 8.32E-03 | A | 50s ~ 60s-70s |
| GO:0019752 | carboxylic acid metabolic process | 13 | 3.14E-04 | A | 50s ~ 60s-70s |
| GO:0046488 | phosphatidylinositol metabolic process | 12 | 4.12E-05 | A | 50s ~ 60s-70s |
| GO:0006952 | defense response | 12 | 1.20E-02 | A | 50s ~ 60s-70s |
| GO:0009415 | response to water | 10 | 3.44E-04 | A | 50s ~ 60s-70s |
| GO:0006096 | glycolytic process | 10 | 2.18E-02 | A | 50s ~ 60s-70s |
| GO:0005985 | sucrose metabolic process | 9 | 1.58E-04 | A | 50s ~ 60s-70s |
| GO:0031047 | gene silencing by RNA | 9 | 2.38E-02 | A | 50s ~ 60s-70s |
| GO:0015937 | coenzyme A biosynthetic process | 7 | 8.48E-05 | A | 50s ~ 60s-70s |
| GO:0043248 | proteasome assembly | 6 | 8.67E-04 | A | 50s ~ 60s-70s |
| GO:0006506 | GPI anchor biosynthetic process | 6 | 1.79E-02 | A | 50s ~ 60s-70s |
| GO:0010274 | hydrotropism | 6 | 2.12E-02 | A | 50s ~ 60s-70s |
| GO:0006284 | base-excision repair | 6 | 3.85E-02 | A | 50s ~ 60s-70s |
| GO:0006221 | pyrimidine nucleotide biosynthetic process | 5 | 2.90E-03 | A | 50s ~ 60s-70s |
| GO:0006886 | intracellular protein transport | 5 | 4.52E-02 | A | 50s ~ 60s-70s |
| GO:0008612 | peptidyl-lysine modification to peptidyl-hypusine | 4 | 5.23E-03 | A | 50s ~ 60s-70s |
| GO:0006488 | dolichol-linked oligosaccharide biosynthetic process | 4 | 8.09E-03 | A | 50s ~ 60s-70s |
| GO:0000077 | DNA damage checkpoint | 4 | 1.17E-02 | A | 50s ~ 60s-70s |
| GO:0046373 | L-arabinose metabolic process | 4 | 1.66E-02 | A | 50s ~ 60s-70s |
| GO:0051260 | protein homooligomerization | 4 | 2.18E-02 | A | 50s ~ 60s-70s |
| GO:0042147 | retrograde transport, endosome to Golgi | 3 | 6.28E-03 | A | 50s ~ 60s-70s |
| GO:0016480 | negative regulation of transcription by RNA polymerase III | 3 | 6.28E-03 | A | 50s ~ 60s-70s |
| GO:0015979 | photosynthesis | 3 | 7.37E-03 | A | 50s ~ 60s-70s |
| GO:0051321 | meiotic cell cycle | 3 | 8.99E-03 | A | 50s ~ 60s-70s |
| GO:0009446 | putrescine biosynthetic process | 3 | 1.22E-02 | A | 50s ~ 60s-70s |
| GO:0006659 | phosphatidylserine biosynthetic process | 3 | 1.66E-02 | A | 50s ~ 60s-70s |
| GO:0050790 | regulation of catalytic activity | 3 | 2.11E-02 | A | 50s ~ 60s-70s |
| GO:0009052 | pentose-phosphate shunt, non-oxidative branch | 3 | 3.49E-02 | A | 50s ~ 60s-70s |
| GO:0015995 | chlorophyll biosynthetic process | 2 | 4.81E-02 | A | 50s ~ 60s-70s |
| GO:0006412 | translation | 1 | 2.13E-12 | A | 50s ~ 60s-70s |
| GO:0006468 | protein phosphorylation | 376 | 5.26E-03 | B | 50s ~ 60s-70s |
| GO:0055114 | oxidation-reduction process | 370 | 2.33E-16 | B | 50s ~ 60s-70s |
| GO:0006355 | regulation of transcription, DNA-templated | 171 | 2.88E-09 | B | 50s ~ 60s-70s |
| GO:0005975 | carbohydrate metabolic process | 124 | 4.54E-18 | B | 50s ~ 60s-70s |
| GO:0006508 | proteolysis | 89 | 2.62E-09 | B | 50s ~ 60s-70s |
| GO:0048544 | recognition of pollen | 38 | 4.84E-08 | B | 50s ~ 60s-70s |
| GO:0009733 | response to auxin | 33 | 5.37E-07 | B | 50s ~ 60s-70s |
| GO:0006351 | transcription, DNA-templated | 32 | 4.28E-03 | B | 50s ~ 60s-70s |
| GO:0006629 | lipid metabolic process | 30 | 8.52E-04 | B | 50s ~ 60s-70s |
| GO:0055085 | transmembrane transport | 28 | 3.69E-12 | B | 50s ~ 60s-70s |
| GO:0016567 | protein ubiquitination | 23 | 1.58E-05 | B | 50s ~ 60s-70s |
| GO:0009058 | biosynthetic process | 23 | 2.63E-03 | B | 50s ~ 60s-70s |
| GO:0006139 | nucleobase-containing compound metabolic process | 16 | 6.18E-03 | B | 50s ~ 60s-70s |
| GO:0006096 | glycolytic process | 13 | 8.20E-04 | B | 50s ~ 60s-70s |
| GO:0016579 | protein deubiquitination | 13 | 1.34E-03 | B | 50s ~ 60s-70s |
| GO:0019752 | carboxylic acid metabolic process | 12 | 8.00E-04 | B | 50s ~ 60s-70s |
| GO:0006396 | RNA processing | 12 | 2.52E-02 | B | 50s ~ 60s-70s |
| GO:0046488 | phosphatidylinositol metabolic process | 11 | 1.23E-04 | B | 50s ~ 60s-70s |
| GO:0006568 | tryptophan metabolic process | 10 | 1.98E-06 | B | 50s ~ 60s-70s |
| GO:0009415 | response to water | 9 | 1.08E-03 | B | 50s ~ 60s-70s |
| GO:0006457 | protein folding | 9 | 1.27E-02 | B | 50s ~ 60s-70s |
| GO:0006364 | rRNA processing | 9 | 2.67E-02 | B | 50s ~ 60s-70s |
| GO:0006352 | DNA-templated transcription, initiation | 9 | 4.84E-02 | B | 50s ~ 60s-70s |
| GO:0043631 | RNA polyadenylation | 6 | 3.19E-02 | B | 50s ~ 60s-70s |
| GO:0017004 | cytochrome complex assembly | 6 | 3.75E-02 | B | 50s ~ 60s-70s |
| GO:0050790 | regulation of catalytic activity | 5 | 2.59E-04 | B | 50s ~ 60s-70s |
| GO:0043666 | regulation of phosphoprotein phosphatase activity | 5 | 1.53E-03 | B | 50s ~ 60s-70s |
| GO:0006164 | purine nucleotide biosynthetic process | 5 | 8.09E-03 | B | 50s ~ 60s-70s |
| GO:0035556 | intracellular signal transduction | 5 | 2.08E-02 | B | 50s ~ 60s-70s |
| GO:0016226 | iron-sulfur cluster assembly | 5 | 3.40E-02 | B | 50s ~ 60s-70s |
| GO:0006366 | transcription by RNA polymerase II | 4 | 2.38E-03 | B | 50s ~ 60s-70s |
| GO:0006488 | dolichol-linked oligosaccharide biosynthetic process | 4 | 6.68E-03 | B | 50s ~ 60s-70s |
| GO:0006606 | protein import into nucleus | 4 | 8.09E-03 | B | 50s ~ 60s-70s |
| GO:0006221 | pyrimidine nucleotide biosynthetic process | 4 | 1.43E-02 | B | 50s ~ 60s-70s |
| GO:0006886 | intracellular protein transport | 4 | 2.14E-02 | B | 50s ~ 60s-70s |
| GO:0016573 | histone acetylation | 4 | 2.76E-02 | B | 50s ~ 60s-70s |
| GO:0006412 | translation | 3 | 9.32E-10 | B | 50s ~ 60s-70s |
| GO:0051301 | cell division | 3 | 1.44E-02 | B | 50s ~ 60s-70s |
| GO:0008612 | peptidyl-lysine modification to peptidyl-hypusine | 3 | 2.73E-02 | B | 50s ~ 60s-70s |

| | | | | | |
|---|---|---|---|---|---|
| GO:0043161 | proteasome-mediated ubiquitin-dependent protein catabolic process | 3 | 2.73E-02 | B | 50s ~ 60s-70s |
| GO:0006188 | IMP biosynthetic process | 2 | 4.56E-02 | B | 50s ~ 60s-70s |
| GO:0042398 | cellular modified amino acid biosynthetic process | 2 | 4.56E-02 | B | 50s ~ 60s-70s |
| GO:0015979 | photosynthesis | 1 | 4.33E-04 | B | 50s ~ 60s-70s |
| GO:0006468 | protein phosphorylation | 359 | 3.33E-05 | D | 50s ~ 60s-70s |
| GO:0055114 | oxidation-reduction process | 331 | 1.73E-15 | D | 50s ~ 60s-70s |
| GO:0006355 | regulation of transcription, DNA-templated | 143 | 8.56E-07 | D | 50s ~ 60s-70s |
| GO:0006508 | proteolysis | 98 | 2.41E-15 | D | 50s ~ 60s-70s |
| GO:0005975 | carbohydrate metabolic process | 90 | 2.77E-09 | D | 50s ~ 60s-70s |
| GO:0048544 | recognition of pollen | 31 | 3.09E-06 | D | 50s ~ 60s-70s |
| GO:0055085 | transmembrane transport | 27 | 1.07E-09 | D | 50s ~ 60s-70s |
| GO:0009058 | biosynthetic process | 24 | 2.15E-04 | D | 50s ~ 60s-70s |
| GO:0016579 | protein deubiquitination | 17 | 1.77E-06 | D | 50s ~ 60s-70s |
| GO:0016567 | protein ubiquitination | 15 | 8.83E-03 | D | 50s ~ 60s-70s |
| GO:0007010 | cytoskeleton organization | 13 | 1.73E-09 | D | 50s ~ 60s-70s |
| GO:0005992 | trehalose biosynthetic process | 12 | 8.16E-05 | D | 50s ~ 60s-70s |
| GO:0006396 | RNA processing | 12 | 1.21E-02 | D | 50s ~ 60s-70s |
| GO:0031047 | gene silencing by RNA | 10 | 3.59E-03 | D | 50s ~ 60s-70s |
| GO:0006370 | 7-methylguanosine mRNA capping | 7 | 2.42E-07 | D | 50s ~ 60s-70s |
| GO:0046488 | phosphatidylinositol metabolic process | 7 | 1.35E-02 | D | 50s ~ 60s-70s |
| GO:0043631 | RNA polyadenylation | 6 | 2.11E-02 | D | 50s ~ 60s-70s |
| GO:0043039 | tRNA aminoacylation | 6 | 4.69E-02 | D | 50s ~ 60s-70s |
| GO:0006412 | translation | 5 | 9.32E-07 | D | 50s ~ 60s-70s |
| GO:0015977 | carbon fixation | 5 | 5.46E-03 | D | 50s ~ 60s-70s |
| GO:0015743 | malate transport | 5 | 9.07E-03 | D | 50s ~ 60s-70s |
| GO:0032259 | methylation | 4 | 4.69E-02 | D | 50s ~ 60s-70s |
| GO:0006366 | transcription by RNA polymerase II | 3 | 1.49E-02 | D | 50s ~ 60s-70s |
| GO:0008612 | peptidyl-lysine modification to peptidyl-hypusine | 3 | 2.11E-02 | D | 50s ~ 60s-70s |
| GO:0009245 | lipid A biosynthetic process | 3 | 2.11E-02 | D | 50s ~ 60s-70s |
| GO:0008295 | spermidine biosynthetic process | 3 | 2.11E-02 | D | 50s ~ 60s-70s |
| GO:0006488 | dolichol-linked oligosaccharide biosynthetic process | 3 | 3.17E-02 | D | 50s ~ 60s-70s |
| GO:0006606 | protein import into nucleus | 3 | 3.72E-02 | D | 50s ~ 60s-70s |
| GO:0015979 | photosynthesis | 2 | 8.06E-03 | D | 50s ~ 60s-70s |
| GO:0042558 | pteridine-containing compound metabolic process | 2 | 4.26E-02 | D | 50s ~ 60s-70s |
| GO:0034477 | U6 snRNA 3-end processing | 2 | 4.26E-02 | D | 50s ~ 60s-70s |
| GO:0055114 | oxidation-reduction process | 579 | 2.55E-44 | A | 60s-70s ~ 80s-90s |
| GO:0006468 | protein phosphorylation | 514 | 7.21E-06 | A | 60s-70s ~ 80s-90s |
| GO:0006355 | regulation of transcription, DNA-templated | 201 | 9.21E-08 | A | 60s-70s ~ 80s-90s |
| GO:0006508 | proteolysis | 137 | 1.85E-19 | A | 60s-70s ~ 80s-90s |
| GO:0005975 | carbohydrate metabolic process | 126 | 3.30E-11 | A | 60s-70s ~ 80s-90s |
| GO:0048544 | recognition of pollen | 83 | 2.04E-29 | A | 60s-70s ~ 80s-90s |
| GO:0055085 | transmembrane transport | 45 | 6.85E-12 | A | 60s-70s ~ 80s-90s |
| GO:0006629 | lipid metabolic process | 41 | 3.69E-05 | A | 60s-70s ~ 80s-90s |
| GO:0009058 | biosynthetic process | 33 | 4.71E-05 | A | 60s-70s ~ 80s-90s |
| GO:0016567 | protein ubiquitination | 30 | 5.91E-07 | A | 60s-70s ~ 80s-90s |
| GO:0006952 | defense response | 23 | 7.47E-07 | A | 60s-70s ~ 80s-90s |
| GO:0000160 | phosphorelay signal transduction system | 22 | 4.77E-04 | A | 60s-70s ~ 80s-90s |
| GO:0016192 | vesicle-mediated transport | 22 | 1.92E-02 | A | 60s-70s ~ 80s-90s |
| GO:0016579 | protein deubiquitination | 18 | 5.87E-05 | A | 60s-70s ~ 80s-90s |
| GO:0019752 | carboxylic acid metabolic process | 11 | 1.56E-02 | A | 60s-70s ~ 80s-90s |
| GO:0017148 | negative regulation of translation | 10 | 1.02E-02 | A | 60s-70s ~ 80s-90s |
| GO:0005992 | trehalose biosynthetic process | 10 | 1.75E-02 | A | 60s-70s ~ 80s-90s |
| GO:0009415 | response to water | 8 | 1.76E-02 | A | 60s-70s ~ 80s-90s |
| GO:0000724 | double-strand break repair via homologous recombination | 7 | 6.16E-07 | A | 60s-70s ~ 80s-90s |
| GO:0009793 | embryo development ending in seed dormancy | 7 | 1.12E-02 | A | 60s-70s ~ 80s-90s |
| GO:0016310 | phosphorylation | 6 | 5.80E-03 | A | 60s-70s ~ 80s-90s |
| GO:0032259 | methylation | 6 | 1.32E-02 | A | 60s-70s ~ 80s-90s |
| GO:0030026 | cellular manganese ion homeostasis | 6 | 2.35E-02 | A | 60s-70s ~ 80s-90s |
| GO:0010112 | regulation of systemic acquired resistance | 4 | 2.42E-02 | A | 60s-70s ~ 80s-90s |
| GO:0006529 | asparagine biosynthetic process | 4 | 3.37E-02 | A | 60s-70s ~ 80s-90s |
| GO:0046907 | intracellular transport | 4 | 3.37E-02 | A | 60s-70s ~ 80s-90s |
| GO:0015937 | coenzyme A biosynthetic process | 4 | 3.84E-02 | A | 60s-70s ~ 80s-90s |
| GO:0016480 | negative regulation of transcription by RNA polymerase III | 3 | 1.12E-02 | A | 60s-70s ~ 80s-90s |
| GO:0042147 | retrograde transport, endosome to Golgi | 3 | 1.12E-02 | A | 60s-70s ~ 80s-90s |
| GO:0051321 | meiotic cell cycle | 3 | 1.67E-02 | A | 60s-70s ~ 80s-90s |
| GO:0030638 | polyketide metabolic process | 3 | 2.26E-02 | A | 60s-70s ~ 80s-90s |
| GO:0006000 | fructose metabolic process | 3 | 2.97E-02 | A | 60s-70s ~ 80s-90s |
| GO:0015979 | photosynthesis | 1 | 2.11E-05 | A | 60s-70s ~ 80s-90s |
| GO:0055114 | oxidation-reduction process | 520 | 3.66E-32 | B | 60s-70s ~ 80s-90s |
| GO:0006468 | protein phosphorylation | 477 | 6.87E-04 | B | 60s-70s ~ 80s-90s |
| GO:0006355 | regulation of transcription, DNA-templated | 273 | 7.65E-27 | B | 60s-70s ~ 80s-90s |
| GO:0005975 | carbohydrate metabolic process | 147 | 3.46E-19 | B | 60s-70s ~ 80s-90s |
| GO:0006508 | proteolysis | 114 | 3.07E-12 | B | 60s-70s ~ 80s-90s |
| GO:0048544 | recognition of pollen | 55 | 1.83E-13 | B | 60s-70s ~ 80s-90s |
| GO:0006629 | lipid metabolic process | 52 | 5.05E-10 | B | 60s-70s ~ 80s-90s |
| GO:0009733 | response to auxin | 47 | 2.45E-11 | B | 60s-70s ~ 80s-90s |
| GO:0055085 | transmembrane transport | 41 | 8.94E-13 | B | 60s-70s ~ 80s-90s |
| GO:0009058 | biosynthetic process | 33 | 4.21E-05 | B | 60s-70s ~ 80s-90s |
| GO:0006396 | RNA processing | 24 | 1.70E-06 | B | 60s-70s ~ 80s-90s |
| GO:0016192 | vesicle-mediated transport | 22 | 1.57E-02 | B | 60s-70s ~ 80s-90s |

| | | | | | |
|---|---|---|---|---|---|
| GO:0016567 | protein ubiquitination | 19 | 9.69E-03 | B | 60s-70s ~ 80s-90s |
| GO:0006096 | glycolytic process | 17 | 8.19E-05 | B | 60s-70s ~ 80s-90s |
| GO:0000160 | phosphorelay signal transduction system | 17 | 1.79E-02 | B | 60s-70s ~ 80s-90s |
| GO:0019752 | carboxylic acid metabolic process | 15 | 1.89E-04 | B | 60s-70s ~ 80s-90s |
| GO:0006364 | rRNA processing | 11 | 1.72E-02 | B | 60s-70s ~ 80s-90s |
| GO:0042545 | cell wall modification | 11 | 3.24E-02 | B | 60s-70s ~ 80s-90s |
| GO:0006298 | mismatch repair | 10 | 4.96E-02 | B | 60s-70s ~ 80s-90s |
| GO:0019346 | transsulfuration | 7 | 5.87E-03 | B | 60s-70s ~ 80s-90s |
| GO:0000272 | polysaccharide catabolic process | 7 | 1.52E-02 | B | 60s-70s ~ 80s-90s |
| GO:0006370 | 7-methylguanosine mRNA capping | 5 | 3.29E-04 | B | 60s-70s ~ 80s-90s |
| GO:0050790 | regulation of catalytic activity | 5 | 7.10E-04 | B | 60s-70s ~ 80s-90s |
| GO:0008295 | spermidine biosynthetic process | 5 | 1.38E-03 | B | 60s-70s ~ 80s-90s |
| GO:0015937 | coenzyme A biosynthetic process | 5 | 7.64E-03 | B | 60s-70s ~ 80s-90s |
| GO:0006886 | intracellular protein transport | 5 | 8.81E-03 | B | 60s-70s ~ 80s-90s |
| GO:0032012 | regulation of ARF protein signal transduction | 5 | 2.15E-02 | B | 60s-70s ~ 80s-90s |
| GO:0006000 | fructose metabolic process | 4 | 3.96E-03 | B | 60s-70s ~ 80s-90s |
| GO:0006606 | protein import into nucleus | 4 | 1.72E-02 | B | 60s-70s ~ 80s-90s |
| GO:0015969 | guanosine tetraphosphate metabolic process | 4 | 2.73E-02 | B | 60s-70s ~ 80s-90s |
| GO:0006221 | pyrimidine nucleotide biosynthetic process | 4 | 2.73E-02 | B | 60s-70s ~ 80s-90s |
| GO:0006071 | glycerol metabolic process | 3 | 2.45E-02 | B | 60s-70s ~ 80s-90s |
| GO:0009411 | response to UV | 3 | 2.45E-02 | B | 60s-70s ~ 80s-90s |
| GO:0006303 | double-strand break repair via nonhomologous end joining | 3 | 3.75E-02 | B | 60s-70s ~ 80s-90s |
| GO:0008612 | peptidyl-lysine modification to peptidyl-hypusine | 3 | 4.59E-02 | B | 60s-70s ~ 80s-90s |
| GO:0015979 | photosynthesis | 1 | 3.22E-05 | B | 60s-70s ~ 80s-90s |
| GO:0007165 | signal transduction | 1 | 2.52E-02 | B | 60s-70s ~ 80s-90s |
| GO:0055114 | oxidation-reduction process | 414 | 4.45E-34 | D | 60s-70s ~ 80s-90s |
| GO:0006468 | protein phosphorylation | 366 | 4.19E-05 | D | 60s-70s ~ 80s-90s |
| GO:0006355 | regulation of transcription, DNA-templated | 148 | 3.29E-07 | D | 60s-70s ~ 80s-90s |
| GO:0006508 | proteolysis | 95 | 1.14E-13 | D | 60s-70s ~ 80s-90s |
| GO:0005975 | carbohydrate metabolic process | 67 | 3.63E-03 | D | 60s-70s ~ 80s-90s |
| GO:0009733 | response to auxin | 46 | 2.04E-15 | D | 60s-70s ~ 80s-90s |
| GO:0009058 | biosynthetic process | 32 | 8.99E-08 | D | 60s-70s ~ 80s-90s |
| GO:0048544 | recognition of pollen | 32 | 2.07E-06 | D | 60s-70s ~ 80s-90s |
| GO:0006629 | lipid metabolic process | 24 | 1.61E-02 | D | 60s-70s ~ 80s-90s |
| GO:0000160 | phosphorelay signal transduction system | 22 | 2.64E-06 | D | 60s-70s ~ 80s-90s |
| GO:0055085 | transmembrane transport | 21 | 2.64E-13 | D | 60s-70s ~ 80s-90s |
| GO:0016567 | protein ubiquitination | 20 | 9.64E-05 | D | 60s-70s ~ 80s-90s |
| GO:0009611 | response to wounding | 14 | 3.92E-02 | D | 60s-70s ~ 80s-90s |
| GO:0006396 | RNA processing | 12 | 1.33E-02 | D | 60s-70s ~ 80s-90s |
| GO:0006096 | glycolytic process | 11 | 3.51E-03 | D | 60s-70s ~ 80s-90s |
| GO:0007010 | cytoskeleton organization | 9 | 1.10E-05 | D | 60s-70s ~ 80s-90s |
| GO:0009415 | response to water | 9 | 5.50E-04 | D | 60s-70s ~ 80s-90s |
| GO:0005992 | trehalose biosynthetic process | 9 | 5.41E-03 | D | 60s-70s ~ 80s-90s |
| GO:0016998 | cell wall macromolecule catabolic process | 7 | 3.24E-02 | D | 60s-70s ~ 80s-90s |
| GO:0005985 | sucrose metabolic process | 6 | 7.33E-03 | D | 60s-70s ~ 80s-90s |
| GO:0006783 | heme biosynthetic process | 5 | 1.05E-04 | D | 60s-70s ~ 80s-90s |
| GO:0015937 | coenzyme A biosynthetic process | 5 | 1.98E-03 | D | 60s-70s ~ 80s-90s |
| GO:0009094 | L-phenylalanine biosynthetic process | 5 | 4.89E-03 | D | 60s-70s ~ 80s-90s |
| GO:0006164 | purine nucleotide biosynthetic process | 4 | 2.60E-02 | D | 60s-70s ~ 80s-90s |
| GO:0032508 | DNA duplex unwinding | 4 | 3.92E-02 | D | 60s-70s ~ 80s-90s |
| GO:0006412 | translation | 3 | 1.24E-08 | D | 60s-70s ~ 80s-90s |
| GO:0006370 | 7-methylguanosine mRNA capping | 3 | 8.49E-03 | D | 60s-70s ~ 80s-90s |
| GO:0006606 | protein import into nucleus | 3 | 3.64E-02 | D | 60s-70s ~ 80s-90s |
| GO:0006420 | arginyl-tRNA aminoacylation | 3 | 3.64E-02 | D | 60s-70s ~ 80s-90s |
| GO:0006417 | regulation of translation | 2 | 1.97E-02 | D | 60s-70s ~ 80s-90s |
| GO:0034477 | U6 snRNA 3-end processing | 2 | 3.92E-02 | D | 60s-70s ~ 80s-90s |
| GO:0000469 | cleavage involved in rRNA processing | 2 | 3.92E-02 | D | 60s-70s ~ 80s-90s |
| GO:0000723 | telomere maintenance | 2 | 3.92E-02 | D | 60s-70s ~ 80s-90s |
| GO:0006468 | protein phosphorylation | 395 | 1.16E-03 | A | 80s-90s ~ 00s- |
| GO:0055114 | oxidation-reduction process | 389 | 2.19E-18 | A | 80s-90s ~ 00s- |
| GO:0006355 | regulation of transcription, DNA-templated | 182 | 7.73E-11 | A | 80s-90s ~ 00s- |
| GO:0006508 | proteolysis | 117 | 9.88E-19 | A | 80s-90s ~ 00s- |
| GO:0005975 | carbohydrate metabolic process | 100 | 3.60E-09 | A | 80s-90s ~ 00s- |
| GO:0006629 | lipid metabolic process | 44 | 5.05E-09 | A | 80s-90s ~ 00s- |
| GO:0009058 | biosynthetic process | 39 | 5.99E-10 | A | 80s-90s ~ 00s- |
| GO:0048544 | recognition of pollen | 36 | 5.50E-07 | A | 80s-90s ~ 00s- |
| GO:0055085 | transmembrane transport | 24 | 4.86E-15 | A | 80s-90s ~ 00s- |
| GO:0016567 | protein ubiquitination | 22 | 7.13E-05 | A | 80s-90s ~ 00s- |
| GO:0009733 | response to auxin | 22 | 1.00E-02 | A | 80s-90s ~ 00s- |
| GO:0000160 | phosphorelay signal transduction system | 19 | 5.06E-04 | A | 80s-90s ~ 00s- |
| GO:0016579 | protein deubiquitination | 18 | 3.46E-06 | A | 80s-90s ~ 00s- |
| GO:0046488 | phosphatidylinositol metabolic process | 16 | 3.20E-08 | A | 80s-90s ~ 00s- |
| GO:0006096 | glycolytic process | 15 | 9.75E-05 | A | 80s-90s ~ 00s- |
| GO:0007165 | signal transduction | 15 | 2.91E-02 | A | 80s-90s ~ 00s- |
| GO:0006396 | RNA processing | 13 | 1.43E-02 | A | 80s-90s ~ 00s- |
| GO:0006284 | base-excision repair | 11 | 4.65E-05 | A | 80s-90s ~ 00s- |
| GO:0007034 | vacuolar transport | 8 | 1.04E-03 | A | 80s-90s ~ 00s- |
| GO:0006457 | protein folding | 8 | 3.86E-02 | A | 80s-90s ~ 00s- |
| GO:0000724 | double-strand break repair via homologous recombination | 7 | 1.30E-07 | A | 80s-90s ~ 00s- |
| GO:0050832 | defense response to fungus | 7 | 8.18E-03 | A | 80s-90s ~ 00s- |

| | | | | | |
|---|---|---|---|---|---|
| GO:0006367 | transcription initiation from RNA polymerase II promoter | 7 | 3.21E-02 | A | 80s-90s ~ 00s- |
| GO:0006729 | tetrahydrobiopterin biosynthetic process | 6 | 2.71E-05 | A | 80s-90s ~ 00s- |
| GO:0006164 | purine nucleotide biosynthetic process | 6 | 1.92E-03 | A | 80s-90s ~ 00s- |
| GO:0009116 | nucleoside metabolic process | 6 | 3.21E-02 | A | 80s-90s ~ 00s- |
| GO:0043631 | RNA polyadenylation | 6 | 3.79E-02 | A | 80s-90s ~ 00s- |
| GO:0006412 | translation | 5 | 1.83E-08 | A | 80s-90s ~ 00s- |
| GO:0007264 | small GTPase mediated signal transduction | 5 | 6.05E-03 | A | 80s-90s ~ 00s- |
| GO:0002100 | tRNA wobble adenosine to inosine editing | 5 | 6.05E-03 | A | 80s-90s ~ 00s- |
| GO:0006415 | translational termination | 5 | 3.24E-02 | A | 80s-90s ~ 00s- |
| GO:0003333 | amino acid transmembrane transport | 5 | 4.12E-02 | A | 80s-90s ~ 00s- |
| GO:0005985 | sucrose metabolic process | 5 | 4.34E-02 | A | 80s-90s ~ 00s- |
| GO:0008612 | peptidyl-lysine modification to peptidyl-hypusine | 4 | 4.92E-03 | A | 80s-90s ~ 00s- |
| GO:0006886 | intracellular protein transport | 4 | 2.46E-02 | A | 80s-90s ~ 00s- |
| GO:0032012 | regulation of ARF protein signal transduction | 4 | 4.34E-02 | A | 80s-90s ~ 00s- |
| GO:0006308 | DNA catabolic process | 4 | 4.78E-02 | A | 80s-90s ~ 00s- |
| GO:0015979 | photosynthesis | 3 | 9.81E-03 | A | 80s-90s ~ 00s- |
| GO:0006741 | NADP biosynthetic process | 3 | 4.33E-02 | A | 80s-90s ~ 00s- |
| GO:0006468 | protein phosphorylation | 388 | 8.45E-03 | B | 80s-90s ~ 00s- |
| GO:0055114 | oxidation-reduction process | 358 | 4.79E-12 | B | 80s-90s ~ 00s- |
| GO:0006355 | regulation of transcription, DNA-templated | 206 | 1.83E-16 | B | 80s-90s ~ 00s- |
| GO:0005975 | carbohydrate metabolic process | 127 | 7.82E-18 | B | 80s-90s ~ 00s- |
| GO:0006508 | proteolysis | 103 | 4.34E-13 | B | 80s-90s ~ 00s- |
| GO:0048544 | recognition of pollen | 46 | 1.14E-11 | B | 80s-90s ~ 00s- |
| GO:0006629 | lipid metabolic process | 35 | 5.43E-05 | B | 80s-90s ~ 00s- |
| GO:0055085 | transmembrane transport | 30 | 1.60E-12 | B | 80s-90s ~ 00s- |
| GO:0006351 | transcription, DNA-templated | 30 | 2.33E-02 | B | 80s-90s ~ 00s- |
| GO:0016567 | protein ubiquitination | 16 | 1.91E-02 | B | 80s-90s ~ 00s- |
| GO:0006281 | DNA repair | 14 | 1.78E-02 | B | 80s-90s ~ 00s- |
| GO:0019752 | carboxylic acid metabolic process | 12 | 1.28E-03 | B | 80s-90s ~ 00s- |
| GO:0046488 | phosphatidylinositol metabolic process | 10 | 8.24E-04 | B | 80s-90s ~ 00s- |
| GO:0015031 | protein transport | 10 | 2.78E-02 | B | 80s-90s ~ 00s- |
| GO:0005992 | trehalose biosynthetic process | 8 | 3.43E-02 | B | 80s-90s ~ 00s- |
| GO:0045927 | positive regulation of growth | 7 | 3.85E-02 | B | 80s-90s ~ 00s- |
| GO:0050832 | defense response to fungus | 6 | 2.94E-02 | B | 80s-90s ~ 00s- |
| GO:0000724 | double-strand break repair via homologous recombination | 5 | 5.49E-05 | B | 80s-90s ~ 00s- |
| GO:0006370 | 7-methylguanosine mRNA capping | 5 | 1.58E-04 | B | 80s-90s ~ 00s- |
| GO:0046373 | L-arabinose metabolic process | 5 | 3.28E-03 | B | 80s-90s ~ 00s- |
| GO:0030833 | regulation of actin filament polymerization | 5 | 5.03E-03 | B | 80s-90s ~ 00s- |
| GO:0032508 | DNA duplex unwinding | 5 | 1.79E-02 | B | 80s-90s ~ 00s- |
| GO:0006414 | translational elongation | 5 | 2.19E-02 | B | 80s-90s ~ 00s- |
| GO:0006415 | translational termination | 5 | 3.43E-02 | B | 80s-90s ~ 00s- |
| GO:0003333 | amino acid transmembrane transport | 5 | 4.34E-02 | B | 80s-90s ~ 00s- |
| GO:0006606 | protein import into nucleus | 4 | 1.07E-02 | B | 80s-90s ~ 00s- |
| GO:0010142 | farnesyl diphosphate biosynthetic process, mevalonate pathway | 3 | 1.79E-02 | B | 80s-90s ~ 00s- |
| GO:0015979 | photosynthesis | 2 | 1.97E-03 | B | 80s-90s ~ 00s- |
| GO:0006412 | translation | 1 | 1.19E-12 | B | 80s-90s ~ 00s- |
| GO:0055114 | oxidation-reduction process | 282 | 2.08E-17 | D | 80s-90s ~ 00s- |
| GO:0006468 | protein phosphorylation | 262 | 1.56E-02 | D | 80s-90s ~ 00s- |
| GO:0006355 | regulation of transcription, DNA-templated | 123 | 9.43E-08 | D | 80s-90s ~ 00s- |
| GO:0006508 | proteolysis | 93 | 2.32E-19 | D | 80s-90s ~ 00s- |
| GO:0005975 | carbohydrate metabolic process | 76 | 3.51E-09 | D | 80s-90s ~ 00s- |
| GO:0048544 | recognition of pollen | 27 | 2.98E-06 | D | 80s-90s ~ 00s- |
| GO:0006351 | transcription, DNA-templated | 24 | 5.99E-03 | D | 80s-90s ~ 00s- |
| GO:0006629 | lipid metabolic process | 23 | 1.58E-03 | D | 80s-90s ~ 00s- |
| GO:0009058 | biosynthetic process | 16 | 1.78E-02 | D | 80s-90s ~ 00s- |
| GO:0016567 | protein ubiquitination | 13 | 8.45E-03 | D | 80s-90s ~ 00s- |
| GO:0055085 | transmembrane transport | 12 | 2.24E-12 | D | 80s-90s ~ 00s- |
| GO:0046488 | phosphatidylinositol metabolic process | 10 | 3.36E-05 | D | 80s-90s ~ 00s- |
| GO:0042545 | cell wall modification | 9 | 8.45E-03 | D | 80s-90s ~ 00s- |
| GO:0016579 | protein deubiquitination | 9 | 1.01E-02 | D | 80s-90s ~ 00s- |
| GO:0009793 | embryo development ending in seed dormancy | 6 | 2.57E-03 | D | 80s-90s ~ 00s- |
| GO:0002100 | tRNA wobble adenosine to inosine editing | 5 | 1.43E-03 | D | 80s-90s ~ 00s- |
| GO:0009451 | RNA modification | 5 | 2.51E-02 | D | 80s-90s ~ 00s- |
| GO:0006284 | base-excision repair | 5 | 3.42E-02 | D | 80s-90s ~ 00s- |
| GO:0046373 | L-arabinose metabolic process | 4 | 5.58E-03 | D | 80s-90s ~ 00s- |
| GO:0007010 | cytoskeleton organization | 4 | 3.02E-02 | D | 80s-90s ~ 00s- |
| GO:0019346 | transsulfuration | 4 | 3.84E-02 | D | 80s-90s ~ 00s- |
| GO:0006415 | translational termination | 4 | 3.84E-02 | D | 80s-90s ~ 00s- |
| GO:0006606 | protein import into nucleus | 3 | 2.45E-02 | D | 80s-90s ~ 00s- |
| GO:0006412 | translation | 2 | 9.31E-07 | D | 80s-90s ~ 00s- |
| GO:0042558 | pteridine-containing compound metabolic process | 2 | 3.08E-02 | D | 80s-90s ~ 00s- |
| GO:0007094 | mitotic spindle assembly checkpoint | 2 | 3.08E-02 | D | 80s-90s ~ 00s- |
| GO:0034477 | U6 snRNA 3-end processing | 2 | 3.08E-02 | D | 80s-90s ~ 00s- |
| GO:0015979 | photosynthesis | 2 | 4.65E-02 | D | 80s-90s ~ 00s- |
